# Apical Localization of RNA Polymerases Modulate Transcription Dynamics and Supercoiling Domains Revealed by Cryo-ET

**DOI:** 10.64898/2026.03.25.714350

**Authors:** Meng Zhang, Cristhian Cañari-Chumpitaz, Jianfang Liu, Bibiana Onoa, Sinead de Cleir, Enze Cheng, Katherinne I. Requejo, Carlos Bustamante

## Abstract

Protein interactions with canonical B-form DNA are well-characterized, yet the effect on these interactions of torsionally constrained DNA—ubiquitous in cells—remains underexplored. Using cryo-electron tomography (cryo-ET), we 3D-reconstructed entire negatively supercoiled DNA substrates bound to active RNA polymerase (RNAP), revealing diverse DNA supercoiling conformations and their interplay with transcription. RNAP preferentially localizes at plectoneme apices in a swiveled, pause-prone state. RNAP, along with other DNA-melting proteins like dCas9, can act as torsional roadblocks that segregate “twin-supercoiling domains” during active transcription, independent of external DNA/RNAP tethering. Co-transcribing RNAPs further intensify this domain separation: tandem oriented RNAPs relieve negative supercoiling more effectively than opposing ones, promote greater RNAP accumulation and enhanced elongation, both in vitro and in vivo. Topoisomerase I relieves torsional stress and facilitates RNAP escape from apical stalls, thereby supporting apical transcription regulation. Together, these findings support a load-and-release mechanism at plectoneme apices that may underlie supercoiling-dependent transcriptional bursting.

## INTRODUCTION

The influence of DNA supercoiling on both prokaryotic and eukaryotic DNA transactions^1^—including transcription, replication, and chromatin segregation,^2,3^ underscores the role played by deviations from canonical B-form DNA in cellular metabolism. In particular, prokaryotic genomes, inherently accumulate negative torsional stress with an average supercoiling density of ∼ −0.06. Accordingly, the interplay between DNA supercoiling and the activities of DNA-binding proteins, such as the collective transcriptional behavior of RNA polymerases (RNAPs) on supercoiled templates, or the torsional regulation by DNA gyrase and topoisomerase, remain an area of active research.^4–8^

The need to visualize DNA supercoiling dynamics has spurred the development of diverse biophysical approaches. In vitro single-molecule fluorescence microscopy enables real-time tracking of DNA plectoneme formation.^9^ Atomic Force Microscopy (AFM) has further allowed the detection of kinks generated in supercoiled DNA.^10^ However, these approaches typically require partial confinement or surface immobilization of DNA. Cryo-electron tomography (cryo-ET) has revisited this topic using small minicircles in a more native state,^11–13^ but the ∼300 bp minicircles were not of sufficient length to capture the complexity of the plectoneme structures.^14^ In structural studies, DNA transcription has primarily focused on elucidating RNA polymerase (RNAP) states and cofactor-mediated regulation on linear templates,^15–19^ while the supercoiling aspect of the template is often overlooked. The “twin-supercoiling domain model”,^20^ that describes overwinding (positive supercoils) ahead of the RNAP and transcription-induced DNA unwinding (negative supercoils) at its wake, still lacks detailed 3D structural characterization despite strong biochemical support.^21–24^ Consequently, questions persist such as: What are the structures naturally adopted by supercoiled DNA in 3D? What is the spatial relationship between a transcribing RNAP and its supercoiled substrate? And how do supercoiling and transcription mutually affect each other?

In this cryo-ET study, we achieved direct 3D visualization of individual plasmid particles (∼2 kbp) under their naturally occurring state of negative supercoiling, allowing precise quantification of their 3D conformational dynamics. Using this plasmid, we stalled and initiated *E.coli* RNAP to systematically investigate the mutual influence between supercoiling and transcription. We determined the orientation of RNAPs on their templates and discovered that their preferred apical binding on negatively supercoiled DNA plectonemes persists during active transcription. RNAPs were found swiveled in this apical configuration, functionally facilitating initiation but hindering elongation—a previously unrecognized mode. Interestingly, we discover that this apical binding also applies to dCas9, turning it into a “soft” torsional barrier that hinders free DNA rotation. The simultaneous apical binding of dCas9 and RNAP on opposite apices of a plasmid leads to the first structural visualization of supercoiling-organized transcription domains, inducing non-plectonemic DNA loops and promoting multi-RNAP slow co-transcriptional events. This novel apical-torsional regulatory mechanism provides a direct structural basis for understanding and experimentally probing how co-transcribing RNAPs impose mutual torsional constraints under tandem and opposing gene contexts, yielding contrasting in vivo transcriptional outcomes. Additionally, introducing *E.coli* Topoisomerase I (TopI) in the presence of RNAP only partially releases the torsion within the plasmid, which, however, is sufficient to disrupt RNAP’s apical positioning and increase transcription elongation activity. Therefore, we propose an ‘on-off’ switch model of apical constraint that plays a critical role in linking torsional stress to transcription regulation.

## RESULTS

### Cryo-ET per-particle analysis enables full 3D DNA tracing of negatively supercoiled plasmids

To explore RNAP’s interactions with its naturally occurring substrate, we first examined the structure and dynamics of a ∼2 kbp modified pUC19-T7A1U circular plasmid (**Figure 1A)** isolated from early stationary-phase *E. coli* cells. 2D gel analysis revealed that the two DNA strands of the plasmid loop over each other 15 times fewer (ΔLk= –15) compared to their relaxed state (Lk_0_ =186), presenting a physiologically relevant supercoiling density (σ=ΔLk/Lk_0_) of –0.08 (**Figure 1B**). The size of pUC19 permits multiple plectoneme formation, as observed by AFM, leading to significant conformational deviation among ΔLk variants. (**Figure 1C**). To capture the native topological diversity of DNA free from surface constraints, we leveraged recent advances in cryo-electron tomography (cryo-ET)^25–27^ to resolve the 3D conformation of large plasmid molecules.

**Figure 1:**
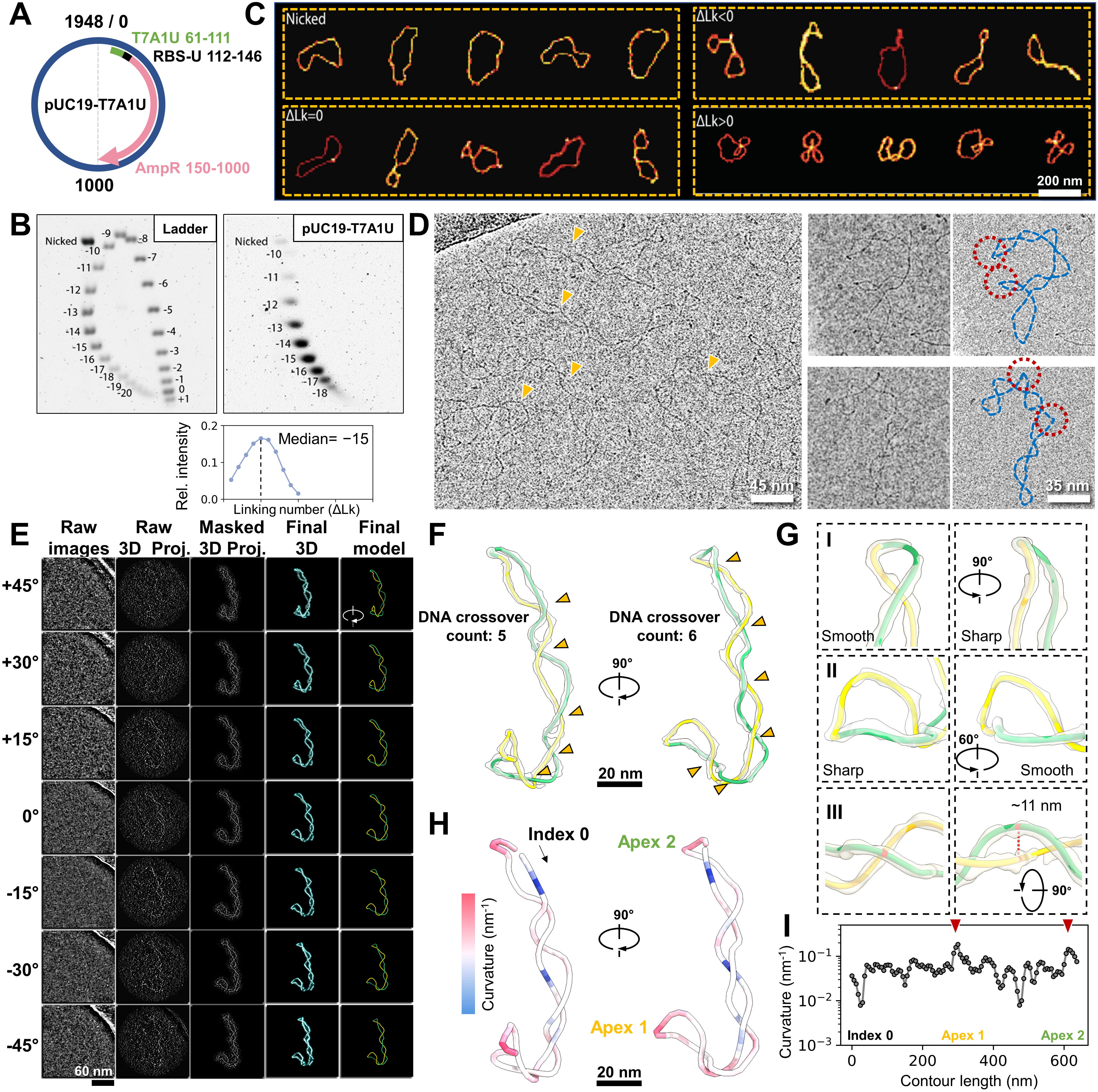
3D reconstruction of an individual negative-supercoiled (-sc) DNA plasmid. **(A)** pUC19 plasmid construct features a T7A1 promoter (green), a U-less stalling site (black), and a transcriptional region (pink). **(B)** 2D gel electrophoresis of the -sc pUC19-T7A1U plasmid and ΔLk quantification. **(C)** AFM images of pUC19 plasmids showing ΔLk variants. **(D)** Cryo-EM images of-sc plasmid (orange arrowhead) highlighting sharp DNA kinks (red circle). **(E)** Cryo-ET per-particle 3D reconstruction steps of a representative -sc plasmid. **(F)** Zoom-in view of the final map and model from E, showing two intertwined DNA segments (yellow and green) with the DNA crossing marked by arrowheads in the view. **(G)** Different 3D views influence 2D assessments of DNA curvature (panels I, II) and spacing (panel III). **(H)** Color-coded map of the plasmid in F, with high-curvature regions in red and low-curvature regions in blue. (I) Tracing of the DNA curvature in H along the circular DNA plasmid.

We initiated cryo-EM imaging under low-salt conditions (5 mM KCl) to capture loosely coiled plasmids,^14^ revealing predominantly long plectonemes (**Figure 1D, orange arrows**) and occasional sharp kinks (**Figure 1D, red circles**) resembling those seen by high-resolution AFM.^10^ Following a deep learning-based segmentation,^28^ local “per-particle” 3D reconstruction^26^, and IsoNet missing wedge correction^29^ (**Video S1 and Supplementary Data 1**), we were able to obtain the non-averaged 3D maps of individual plasmids (**Figure 1E**) at resolution of ∼30-40 Å (**Supplementary Data, Particle gallery #006**). Using a DNA detection algorithm with manual curation (**Supplementary Data 2**), we traced full 3D plasmid contours, enabling precise supercoiling quantification. For the example in **Figure 1E**, spatial quantification^30^ of right-handed crossings in 3D gave a writhe (Wr) of −6.2. Accurate structural analysis is difficult to achieve from 2D projections alone due to projection artifacts—e.g., DNA crossing number varies (**Figure 1F, orange arrows**), smooth curves may appear as sharp kinks, and true bends can be obscured in certain views. (**Figure 1G, panels I-II)**. To overcome these limitations, we measured 3D curvature (reciprocal of osculating sphere radius; **Figure 1H**) and inter-helix spacing in plectonemes (**Figure 1G, panel III**). The curvature distribution showed two peaks (**Figure 1I, red arrows**), corresponding to plectoneme apices.

We then compared plasmids under low salt (5□mM K^⁺^; **Figure S1A**) and high salt (40□mM K^⁺^, 5□mM Mg^²⁺^, near-physiological ionic strength; **Figure S1B**) conditions, (see full dataset in **Supplementary Data Particle Gallery**). Under higher salt, plasmids exhibited increased elongation, indicated by a larger radius of gyration (Rg), and greater DNA intertwining with mean Wr shifting from −5.7 to −10.2; *P* <0.0001 (**Figure S1C-D**). This result aligns with theoretical predictions that electrostatic screening favors DNA crossing,^14^ promoting the conversion of negative supercoiling into writhe at the expense of twist (ΔTw). As a result, plectoneme width decreased with high salt from 13.2 to 8.4 nm; *P* <0.001 (**Figure S1E**), accompanied by increased mean curvature, particularly at distal apices (from 0.19 to 0.23 nm⁻¹; *P* <0.001, **Figure S1F-G**). Given that some proteins recognize highly curved DNA region via indirect readout^31^, we assessed apex number distributions under both conditions and observed comparable results: ∼60% of plasmids remaining unbranched (two apices; **Figure S1H**). Clearly, generation of new apices is energetically unfavorable. These analyses provide a structural framework for understanding supercoiling dynamics (see below).

### RNAP apical binding favors promoter escape and affects the configuration of the negatively supercoiled template

An early study in the 1990s revealed unexpected apical localization of RNAP on supercoiled DNA,^32^ a finding recently revisited by single-molecule experiments.^9^ To gain structural insights, we incubated RNAP with pUC19-T7A1U plasmids (bearing a single T7A1-driven gene; **Figure 2A, top circle**) at a 3:1 ratio under high-salt conditions and obtained stalled transcription elongation complexes (sTECs) via UTP starvation, halting RNAP 19 nt downstream of the transcription start site (TSS). 3D reconstruction revealed that ∼90% of RNAPs localized at plectoneme apices (**Figure 2A, lower panels**). Sub-tomogram averaging (STA) resolved RNAP orientation at the apex.^28^ Averaging all RNAP particles (bound and unbound) yielded a 14 Å map with blurred DNA entry/exit sites (**Figure S2A-B, top**), whereas averaging only plasmid-bound RNAPs (17 Å resolution) revealed a DNA density protrusion indicating the downstream direction (**Figure S2A, bottom, black arrow**). This feature is absent in RNAP structures resolved with linear DNA templates (4YLN, 6CA0, 6JBQ, 6N60,^33–36^ which show only upstream DNA bound to the σ factor (**Figure S2C**). Mapping STA-determined RNAP orientations onto their respective plasmid models (**Figure 2A, top right**) recovered DNA information otherwise averaged out. Superimposing all RNAPs at apical regions highlighted DNA’s conformational flexibility (**Figure 2B**), while superimposing DNA at the same region revealed relatively restricted RNAP apical localization (**Figure 2C**).

**Figure 2:**
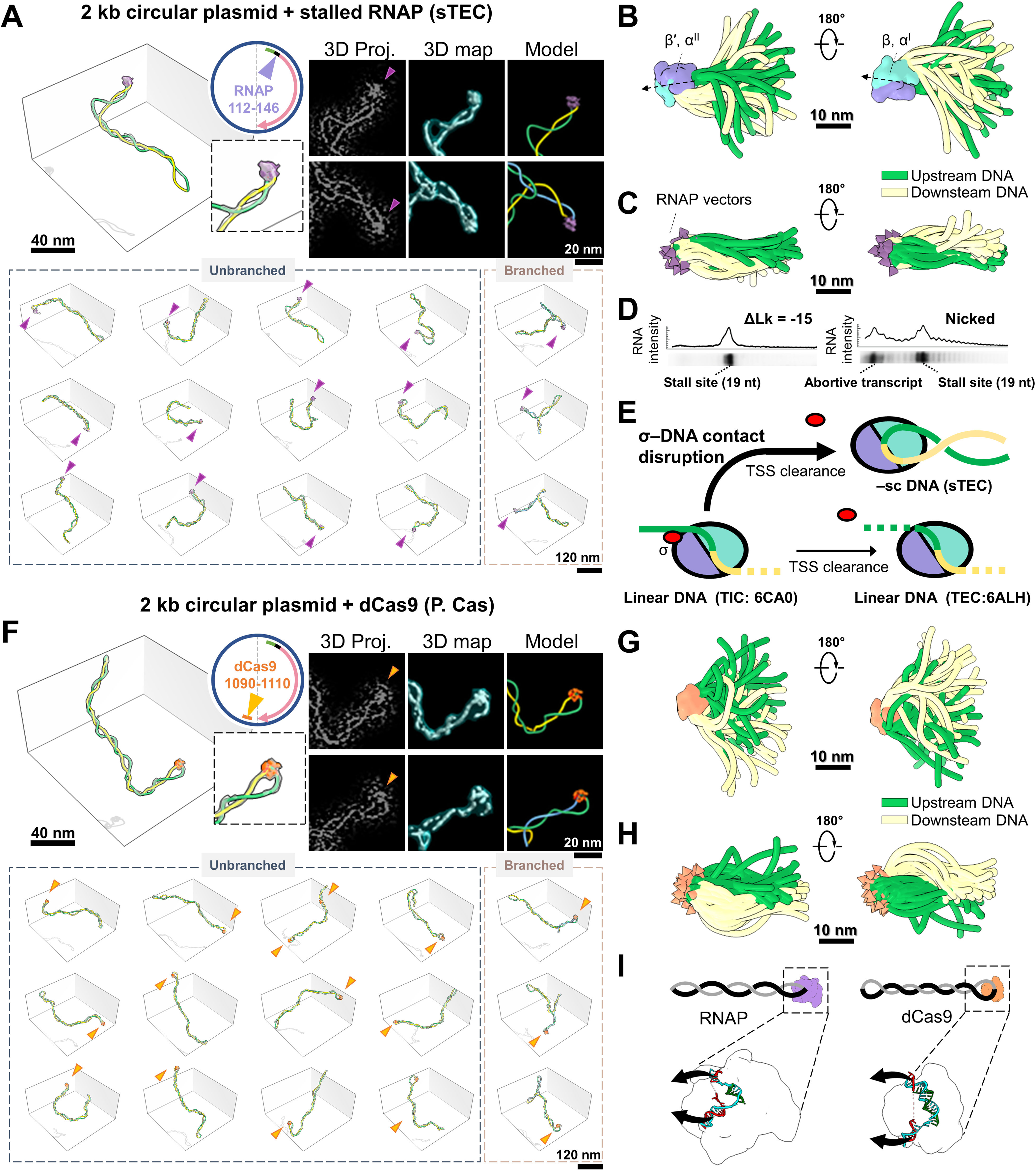
Conformational dynamics and regulatory effect of apically bound proteins on DNA supercoiling. **(A)** Cryo-ET (3D maps and models) of the -sc plasmids bound with stalled RNAP (sTEC), with plasmid construct design in circle and plectoneme apex modeling steps on top right. RNAPs in sTECs are marked by purple arrowheads in the bottom particle collection **(B)** The superimposition of all bound RNAPs reveals apical DNA dynamics. The dashed vector indicates RNAP orientation. **(C)** The superimposition of all apical DNA segments shows the RNAP orientation dynamics (purple arrowheads). **(D)** Electrophoresis gel shows reduced abortive transcript levels on -sc DNA compared to nicked DNA. **(E)** Schematic highlighting DNA geometry conversion in TEC on -sc DNA (top), and in TIC (bottom left) and TEC (bottom right) on linear DNA templates. **(F)** Cryo-ET of dCas9 (orange) bound to -sc plasmids. **(G and H)** Superimposition of all bound dCas9s and all apical DNA segments, respectively. **(I)** Schematic highlights dCas9’s smaller pocket curvature compared to that of RNAP, reshaping bound plasmids with enlarged distal ends and tightly intertwined body.

Negative DNA supercoiling is believed to facilitate transcription initiation by enhancing the kinetics of open complex formation through DNA unwinding.^37,38^ However, its impact on other initiation events, such as promoter search and escape, remains unclear. Interestingly, bulk transcription assays on nicked and negatively supercoiled (–sc) templates revealed more efficient promoter escape on –sc DNA, evidenced by a reduced rate of abortive initiation before RNAP reached the 19 nt stall site (**Figure 2D**). Structural superposition of the apically stalled TECs (sTEC) in –sc plasmids onto transcription initiation complexes (TICs) assembled on linear DNA (such as 6CA0) revealed a significant deviation in the consensus upstream DNA trajectory—approximately 170° in bending angle in the former (**Figure S2C**). This substantial reorientation of the upstream DNA in –sc complexes would disrupt most of DNA-σ-factor interaction, if present (**Figure 2E**). As weakening of DNA-σ contacts is a necessary step for RNAP release from the promoter and transition into elongation^39^, this apical configuration on –sc DNA provides a possible explanation for the increased promotor clearance observed in the bulk assay (**Figure 2E**).

Having characterized RNAP’s apical binding, we asked whether this feature is unique to RNAP or shared by other DNA-binding proteins. With the same imaging setup, we replaced RNAP with catalytically inactive dCas9 guided by an sgRNA targeting the plasmid’s transcription termination site (TTS) (**Figure 2F, top circle**). We chose dCas9 for its RNAP-like DNA binding (i.e. forming a bubble via DNA-RNA hybrid) and its visibility (∼10 nm) in cryo-ET. Notably, dCas9 also exhibited apical binding on –sc plasmids (**Figure 2F, bottom**) with STA analysis revealing the consensus entry-exit DNA orientation **(Figure S2D, arrows**; 17Å resolution, **Figure S2E**). While both proteins share apical binding, quantitative 3D analysis enables the identification of structural differences between dCas9- and RNAP-plasmid complexes (P.Cas vs. sTEC), with the former displaying more elongated, tightly wound plectonemes, as indicated by increased Rg and a more negative Wr (−9.5 to −11.8; *P* <0.0001, **Figure S2F, top**). Enlarged distal loops were also observed at dCas9-bound apices, showing reduced curvature (from 0.25 to 0.22 nm^-1^; *P* <0.05) and increased DNA spacing (from 7.7 to 10.3 nm; *P* <0.05) compared to RNAP (**Figure S2F, bottom**). Mapping dCas9 orientation highlighted these distinctions (**Figure 2G-H**).

Why do two DNA-binding proteins with similar mechanisms reshape plectonemes differently? Structural comparison of RNAP^40^ and dCas9^41^ binding pockets revealed that, despite its larger size (15 vs. 10 nm), RNAP features a more acute binding pocket with a sharper curvature than dCas9 (**Figure 2I**), favoring DNA melting over negative writhe. Coarse-grained molecular dynamics (CGMD) simulations using oxDNA^42^ further support this conclusion, showing that tighter apex bending with strand separations of 4.8 and 6.4 nm (**Figure S3A**) yield sharper apices and reduced negative writhe compared to looser apex bending (8 nm separation) (**Figure S3B, left and middle**). A positive correlation (r =0.51) was observed between apex curvature and writhe number, based on data pooled from all trajectories (**Figure S3B, right**). Taken together, these findings suggest that apex localization is not unique to RNAP, but reflects a general structural adaptation of supercoiled DNA to the geometry of bound proteins that minimizes the energy of the overall configuration. Significantly, we found that DNA sequence at the promoter and dCas9 binding site played a minor effect on proteins apical localization^43^ (**Figure S3C**).

### Transcription induces branched supercoiling while preserving RNAP’s apical localization

RNAP’s translocation along the DNA helix necessitates rotational movement by either the DNA or RNAP to keep the DNA template in register within the RNAP active site. Simultaneous rotational constraints on both RNAP and DNA, such as RNAP tethering to cellular structures^44,45^ and DNA confinement within dense genome architecture,^46–48^ can lead to the formation of twin supercoiling domains.^20^ However, such external constraints might not always be present in various cellular contexts. It is therefore intriguing to investigate whether intrinsic constraints in TECs resulting from RNAP’s apical localization and a negative supercoiling background^48–50^ can sustain transcription and contribute to topological-twin-domain delineation.

To investigate the above scenario, we added the full set of NTPs (100 μM each) to the stalled TECs (sTECs) and imaged the sample 10 minutes after resuming transcription. Remarkably, the 3D reconstruction reveals the continued apical positioning of RNAP in TECs (**Figure 3A-B, purple arrows**), with nascent RNA density intermittently observed around RNAP in both 3D maps (**Figure 3A, right**) and 2D slices (**Figure 3C top, red contours**), but absent in sTECs (**Figure 3C bottom**). However, the limited resolution of cryo-ET, along with conformational heterogeneity of RNA^51^ prevented the tracing of complete RNA transcripts, permitting only the mapping of the RNA protrusion direction relative to RNAP (**Figure S3D, arrows**). Quantitative analysis revealed that, during active transcription, plasmids predominantly adopt a multi-branched configuration for the first time (>78% of structures, **Figure 3D**), resulting in a population with reduced spatial extension and smaller Rg (**Figure S3E, top, red arrow**). Additionally, we observed occasional emergence of blunt apices in TECs (hollow purple arrows in both **Figure 3A and Figure S3E**), a feature rarely seen in sTECs (**Figure 2A**). While the overall plasmid writhe was similar between TECs and sTECs (−9.5 vs. −9.8) (**Figure S3E, top**), segmentation of individual branches (**Figure S3F, I-III**) followed by analysis of writhe density (branch Wr/branch length) showed that TECs exhibit a broader distribution with a slightly increases (−0.13 vs. −0.15 nm⁻¹; *P* <0.05; **Figure S3F, bottom**). These observations suggest that intrinsic −sc constraints, coupled with active DNA translocation relative to RNAP, are sufficient to generate a mild degree of torsional imbalance in the plasmid, promoting the formation of a new supercoiling domains (plectoneme branching). Branching appears to preferentially occur in the region of the plasmid proximal to RNAP (**Figure S3G**), consistent with its role as the source of torsional stress.

**Figure 3:**
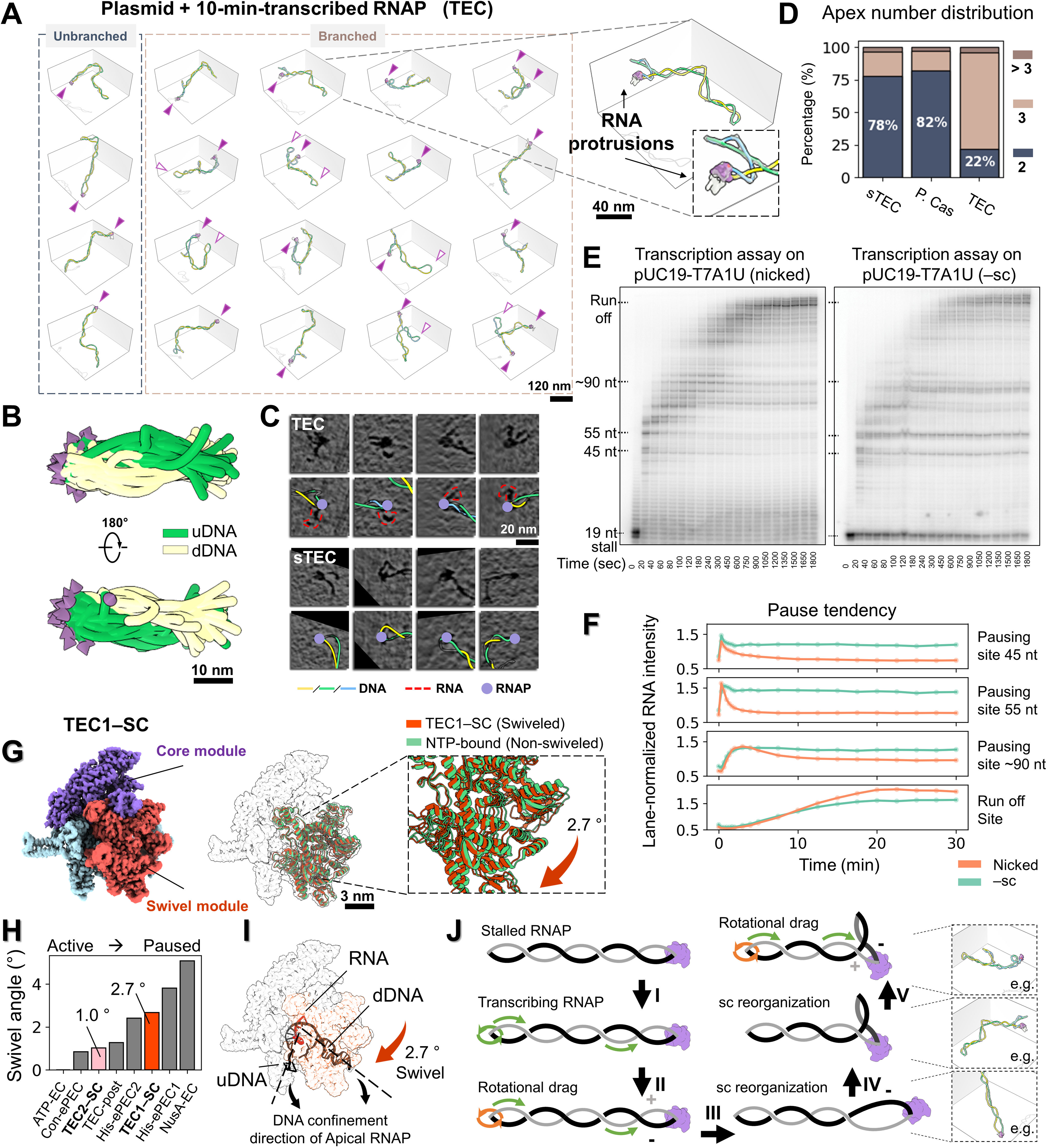
Interplay between apically bound RNAP and -sc DNA during non-equilibrium transcription. **(A)** Cryo-ET of TEC on -sc templates after 10 minutes of transcription, with RNAP and large DNA loops indicated by solid and hollow purple arrowheads, respectively. **(B)** Alignment and superimposition of all apical DNA segments. **(C)** Z- dimensional slices (10 nm-thickness) of sub-tomograms comparing transcribed (top) and stalled (bottom) RNAPs, overlaid with models. **(D)** Plasmid apex number distribution; N=84. **(E-F)** Single-round *in vitro* transcription of RNAP transcription on nicked and -sc templates over time, with quantification of their pause release kinetics, respectively. **(G)** Single-particle cryo-EM reconstruction of the major RNAP class, TEC1-SC on -sc DNA, overlaid with the non-swiveled 6RH3 model. **(H)** Comparison of TEC1-SC swivel angle to reported RNAP structures from active transcription to various paused states. (I) TEC1-SC active-site DNA orientation and subunit swivel direction. **(J)** Schematic illustration of transcription-induced new plectoneme formation on -sc plasmid. DNA translational and rotational motion is indicated by green arrows, with orange color denoting drags.

DNA torsional stress resulting from transcription of torsionally constrained templates is known to impede transcription elongation, causing RNAP stalling and increased backtracking as shown in single-molecule assays.^21^ Given the persistent apical localization of RNAP observed in our experiments during active transcription elongation, we wondered if transcription rate was also affected in these conditions. Using radiolabeled α-^32^P-ATP, single-round in vitro transcription assays on –sc DNA displayed significantly slower elongation kinetics (**Figure 3E**), as well as increased pause frequency and duration (**Figure 3F**), than those performed on nicked DNA. We note that, in our bulk transcription assay, the topological constraint arises from the natural steady-state degree of negative supercoiling of the plasmids extracted from the cell. The slower elongation kinetics observed supports the notion that the apical localization of RNAP hinders its rotation around DNA, and that the inefficient rotation of the larger DNA molecule becomes rate-limiting to transcription.^32^

Alternatively, the topological stress under which the apical RNAP operates could induce a conformational change in the enzyme which could, in turn, affect its dynamics. To test this hypothesis, we performed single-particle analysis (SPA) of TECs on –sc templates. 3D classification yielded three classes, of which Class 1 and Class 2 (TEC1–SC and TEC2–SC), comprise 34% and 40% of particles, respectively (**Figure S4A-C**) attaining in both cases 2.9 Å resolution (**Figure S4D-E**) and clearly showing melted DNA and RNA transcripts at the active site. Minor Class 3 (26%) exhibited faint DNA density and was excluded from further analysis. Close inspection revealed that TEC1–SC and TEC2–SC are both in a post-translocated state, evidenced by an unfolded trigger helix and absence of the incoming NTP (**Figure S4F**). Using the NTP-bound RNAP structure (PDB: 6RH3) as a reference, we measured the swivel angle following Kang et al.^52^ and found that TEC1–SC exhibits significant swiveling (∼2.7°; **Figure 3G**), comparable to the swivel state displayed by the His-elemental paused elongation complex (His-ePEC, **Figure 3H**), that results from a pause stabilized by an RNA hairpin formation. Interestingly, we notice that the TEC1–SC DNA orientation (**Figure 3I, dashed line**) and the DNA confinement geometry in apically located RNAP (**Figure 3I, black arrow**) align with the orientation of the swivel module (**Figure 3I, red arrow**). Because the downstream DNA contacts the swivel module, these observations suggest that bending of the constrained apical DNA drives the swivel conformation, thereby slowing transcription elongation.

Building on these findings, we propose a model for how transcription takes place in intrinsically –sc DNA (**Figure 3J**): The persistent apical positioning of RNAP requires DNA to rotate during transcription as it is thread through the enzyme (**Figure 3J, I**). However, the propagation of DNA rotation throughout the entire plasmid structure is slow,^53,54^ so that transcription induces transient (+) torsional stress ahead of the enzyme and (−) torsional stress at its wake (**Figure 3J, II**). In topologically constrained DNA, the (+) torsional stress is rapidly absorbed by the –sc background, and therefore, the cancellation of (+) and (−) supercoils through diffusion across the plasmid does not occur (**Figure 3J, III**). Meanwhile, since high salt conditions favor partitioning of ΔLk into writhe over twist (**Figure S1D**), the extra (−) torsional stress generated at the wake of the polymerase promotes the formation of a new negative plectoneme branch.

### Dual apical protein binding promotes the formation of topological domains

The results above highlighted RNAP’s role in generating torsional stress during non-equilibrium transcription of supercoiled DNA. Torsional stress during transcription can also accumulate due to the presence of torsional roadblocks, such as DNA-binding proteins and transcription factors,^55,56^ leading to the formation of topological domains that can affect RNAP molecules distant kilobases away.^57^ Nonetheless, the mechanism through which small DNA-binding proteins impose rotational constraints and favor the generation of topological domains remains elusive.

To gain insight into this process, we employed dCas9 again but in this case for its capability of delaying DNA rotation, as reported in magnetic tweezers assays.^58^ To confirm that dCas9 can serve as a programmable torsional roadblock, we performed DNA relaxation experiments using topoisomerase I (TopI) in the absence and presence of dCas9 bound to the transcriptional termination site (TTS) (**Figure 4A, top circle**). TopI relaxes torsion in –sc DNA by rotating the helix until equilibrium (ΔLk□=□0) is achieved. Our results demonstrate that dCas9 delays the relaxation kinetics of pUC19 plasmids, and topoisomers with ΔLk from −3 to −1 still persist after a 30-minute reaction (**Figure S5A**). We repeated the experiment using human Topoisomerase IB (hTopIB), which relaxes supercoiling via a direct DNA rotation mechanism rather than strand passage (as in TopI), and similarly observed that dCas9 delays DNA relaxation (**Figure S5B**). Because dCas9 primarily delays, rather than halts, –sc relaxation, we term it a “soft” torsional block. To understand this mechanism, we performed CGMD simulations to investigate how a –sc plasmid relaxes after nicking, with or without an apical constraint. Results reveal that imposing an apical constraint on one of the two apices slows supercoiling torsional relaxation dynamics when the nick is positioned between the apices (**Video S2 and Figure S5C, Model II-III vs. I**). However, when the nick is placed directly on the apex, relaxation becomes fastest, even with the opposite apex constrained (**Figure S5C, Model IV**). Based on the observed relaxation speeds (IV > I > II and III), and given that model IV is the only scenario where torsional propagation encounters no apices (with the nick created at the very end), it suggests that the apex, particularly the confined apex (mimicking the effect of dCas9 binding), hinders free DNA rotation.

**Figure 4:**
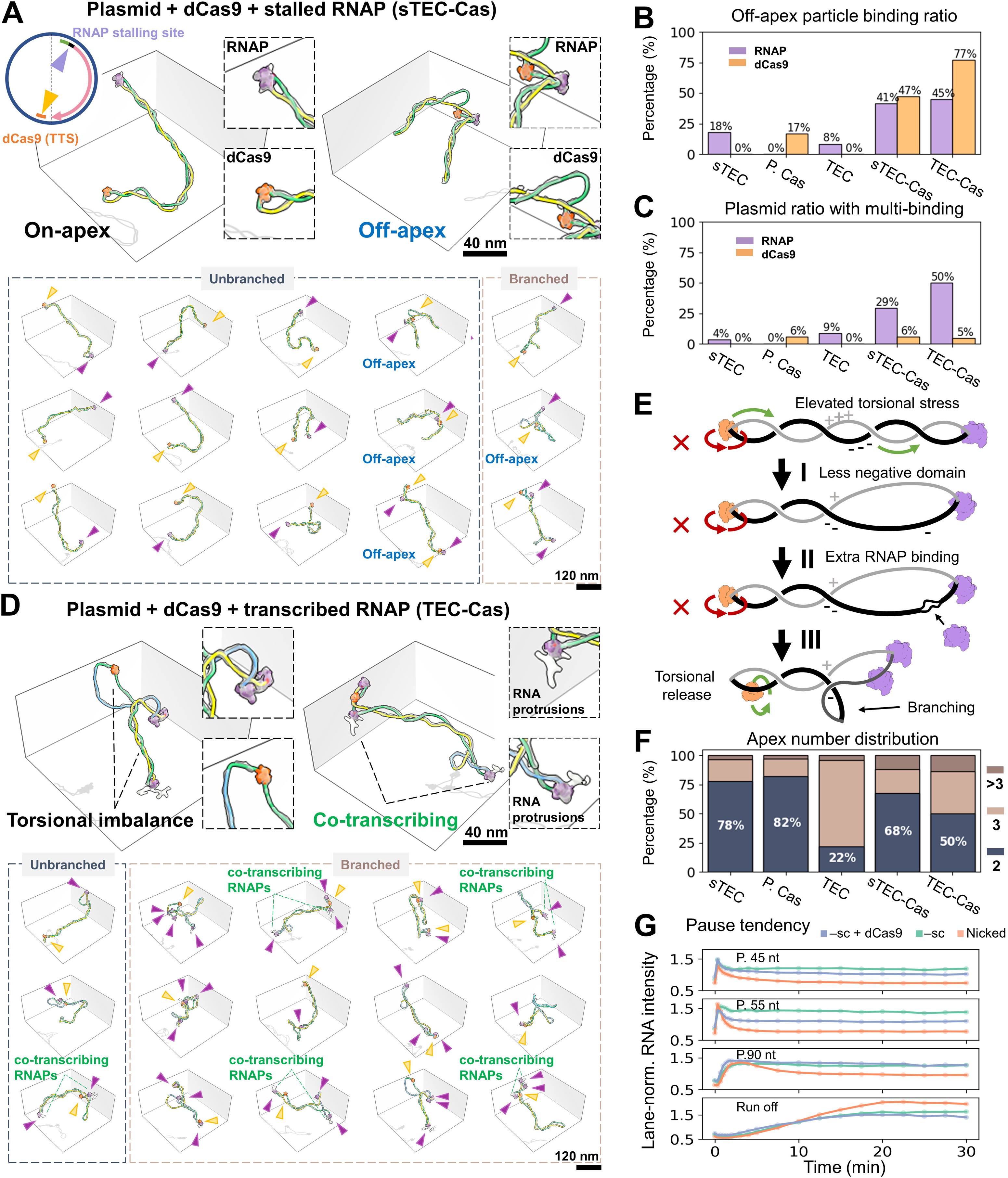
Plectoneme apical confinement promotes topological domains and multi-RNAP transcription. **(A)** Cryo-ET of -sc plasmids simultaneously bound by stalled RNAP (purple) and dCas9 (orange). Representative particles show RNAP and dCas9 co-localized at the apex (top left) or off-apex (top right), both consistent with the opposing-position construct design (circular plot). **(B)** Off-apex particle ratios for RNAP and dCas9 across conditions; N=301 **(C)** Ratio of plasmids bound by multiple RNAPs or dCas9; N=324. **(D)** Cryo-ET of -sc plasmid resuming transcription for 10 minutes in the presence of torsional roadblock dCas9 **(E)** Schematics illustrating RNAP transcription-induced topological domains and supercoiling rearrangement in the presence of dCas9. **(F)** Quantification of plasmids’ apex number; N=162. **(G)** Assessment of RNAP pause release in the presence of dCas9 during transcription via electrophoresis. The RNA band intensities were quantified at three strong pausing sites and run-off site.

To experimentally test the role of torsional roadblocks on transcription, we first initiated transcription on pUC19 plasmids with dCas9 bound at the TTS (**Figure 4A, top circle**). Imaging of stalled TECs (sTEC-Cas) harboring a ∼20 nt transcript revealed that RNAP and dCas9 predominantly localize to opposite plectoneme ends (**Figure 4A top left, and bottom columns 1-3**). We note that on unbranched plasmids, their apical localization is mutually reinforcing, as both favor and induce bending at their target sites (**Figure 2A,F**). Even under this mild transcription and stalling scenario, sTEC-Cas exhibited noticeably less negative writhe domains compared to the dCas9-bound-only plasmid (P.Cas), resembling instead those of active TECs (**Figure S5D, red arrow**). Notably, mild transcription conditions also led to a significant increase in non-apical binding cases (**Figure 4A, blue label** and **Figure S5E**) for both RNAP and dCas9 (from 18% to 41% and from 17% to 47%, respectively, **Figure 4B**). Given that the increased off-apex dCas9 ratio was not due to additional dCas9 binding events (both 6% for P.Cas and sTEC-Cas; **Figure 4C**), it suggests that transcription-induced accumulation of (+) torsional stress between the front of RNAP and dCas9, and (–) stress between dCas9 and the back of RNAP, promotes partial escape of the dCas9 from the apices as the system seeks to minimize its energy.

### Torsional block enhances cooperative RNAP transcription

As shown above, mild (+) and (–) torsional stresses differentially accumulate in each half of the plasmid, delineated by RNAP and dCas9, resulting in more frequent off-apical dCas9 events, suggesting the “soft” nature of dCas9 when acting as a torsional block. Accordingly, we anticipated increased off-apex localization of dCas9 after 10 minutes of active RNAP transcription with the full set of NTPs at 100 μM (TEC-Cas; **Figure 4D**). Indeed, the off-apex ratios of both RNAP and dCas9 increased, with dCas9 reaching 77% (**Figure 4B**). Given a −sc background (ΔLk = −15), the “twin-domain” model should result in the DNA segment in front of the RNAP to be less –sc and the domain at its back more so. Consistent with this expectation, we observed the coexistence of large, non-plectonemic DNA loops and tightly wound DNA regions within the same TEC-Cas particle (**Figure 4D, top left**). Quantitative analysis confirmed that, although overall plasmid morphology remained similar (**Figure S6A, top**), TEC-Cas exhibited a broader writhe density distribution across plectonemic branches, with two peaks above and one slightly below the mean of the plasmid-only control (**Figure S6B, red arrows**). This shift toward less negative writhe was accompanied by blunter apices, characterized by reduced apex curvature (from 2.5 to 2.3 nm^-1^; *P* < 0.05) and increased apical-DNA spacing (from 8.5 to 11.1 nm; *P* < 0.001, **Figure S6A, bottom**). Notably, less-negative domains were more prominent, which seems contrary to the expected symmetry of (+) and (−) torsion generated by RNAP (**Figure 4E, I**). This asymmetry can be rationalized by the marked increase in multi-RNAP binding (from 29% to 50%; **Figure 4C**) observed in TEC-Cas. Accordingly, as each bound RNAP unwinds ∼10 bp of DNA, they absorb excess negative supercoiling (**Figure 4E, II**). Moreover, the increase in the proportion of multi-branch form plasmids in TEC-Cas (from 32% to 50%, **Figure 4F**), providing an additional mechanism to buffer excess negative supercoiling (**Figure 4F, III**).

Since structural studies provide only static snapshots of domain separation, we employed CGMD to better visualize how torsional domains dynamically emerge as a consequence of apical constraint in the absence of external tethering. We treated the domains as harmonic (linear torsional springs)^59^ to mimic the (+) and (−) torsion generated by RNAP. The simulations compare a free versus constrained apex in managing torsional stress (**Video S3**). Notably, the constrained apex successfully recapitulated the formation of large loops and branching structures observed experimentally.

Leveraging dCas9’s gRNA-directed DNA binding as a fiducial marker, we tentatively mapped RNAP positions and orientations on the plasmid construct (**Figure S6C**). Some RNAPs were found outside the transcriptional region or oriented upstream of the TSS (brown and blue arrows), which we interpret as non-specific binding enhanced by transcription-induced −sc domain. In contrast, most RNAPs were located within the transcriptional region and oriented downstream (pink arrows), with some plasmids displaying multiple transcribing RNAPs. The direct visualization of two RNAPs on the same template, both exhibiting nascent RNA protrusions (**Figure 4D, green labels**), supports this interpretation. These findings suggest that dCas9-induced topological imbalance in TECs promotes cooperative RNAP transcription, potentially enhancing transcriptional kinetics. To corroborate this hypothesis and link structure to function, we performed a bulk transcription assay in the presence of dCas9 (**Figure S6D**). The results show that while pause frequency remains unchanged, pause escape is faster in the TEC-Cas—particularly at the 45 nt and 55 nt pause sites (**Figure 4G, blue vs. green curve**). This enhanced pause escape suggests that the accumulation of (+) torsion ahead of RNAP, induced by the dCas9 torsional block, relieves the enzyme’s apical constraint on a –sc template, thereby facilitating elongation—though not to the level observed with nicked templates (**Figure 4G, orange curve**). Together, these results suggest that, in the context of −sc, twin-domain formation facilitates both RNAP initiation and elongation, with a stronger effect on the former.

### TopI removes the apical constraint of RNAP facilitating DNA transcription

Although the presence of a torsional roadblock like dCas9 promotes transcription by multiple RNAPs, it could not fully replicate the elongation dynamics seen on nicked circular plasmids. Accordingly, we sought to determine if TopI^8,60,61^ (**Figure 5A**) could alleviate the apical constraints on RNAP by relaxing the DNA plectoneme and thereby relieve RNAP from its pauses.

**Figure 5:**
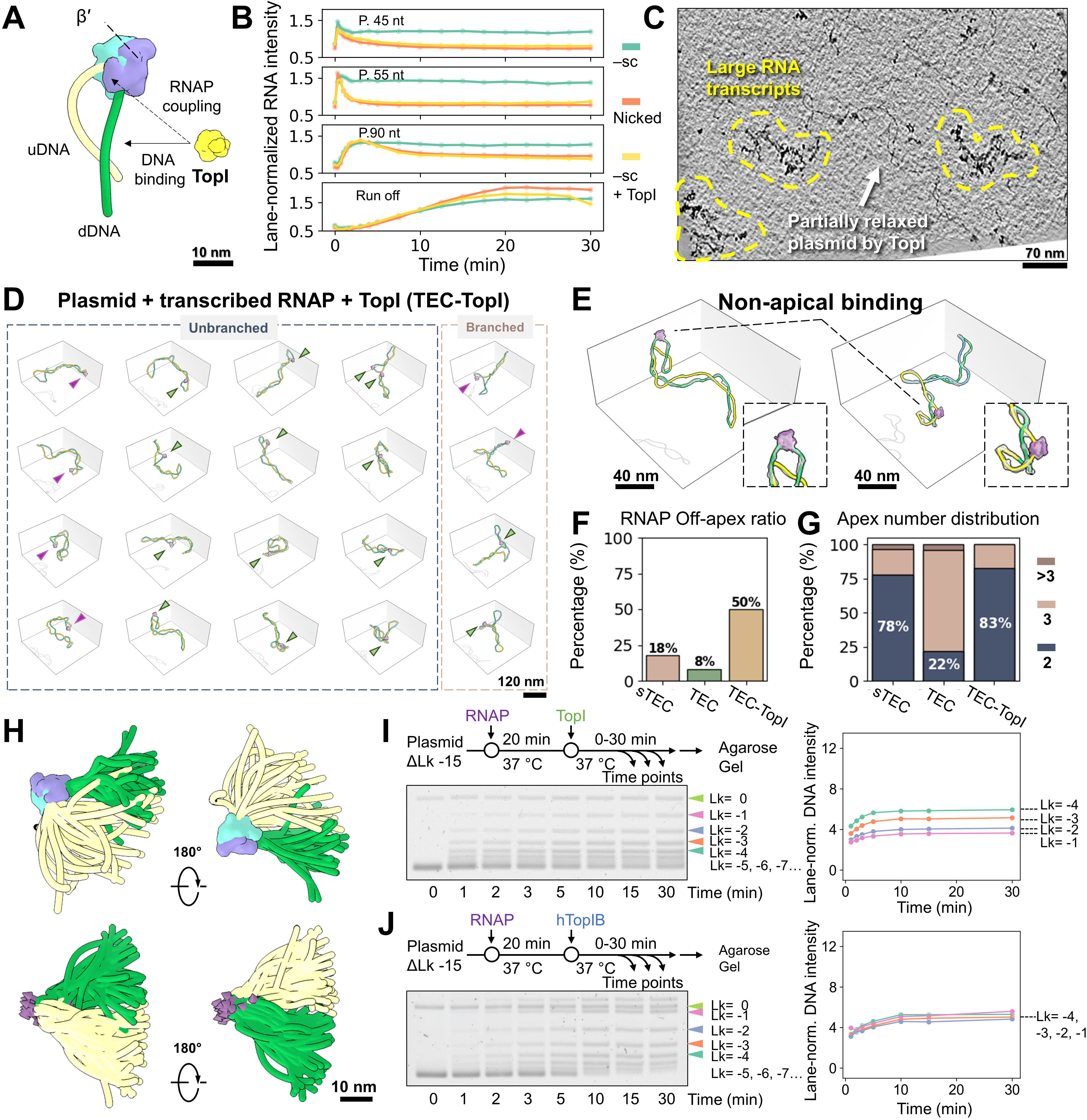
Topi releases RNAP from apical constraint during transcription. **(A)** Schematic showing Topi bound to DNA or coupled with RNAP. **(B)** Assessment of RNAP pause release in the presence of Topi from single-round *in vitro* transcription assay. **(C)** Cryo-ET z-dimensional slice (50 nm-thickness) of sample after 10 minutes of transcription in the presence of Topi. **(D)** Cryo-ET of the plasmid particles in the presence of RNAP and Topi after 10 minutes of of transcription (TEC-Topl). Apical and non-apical RNAP binding are indicated by purple and green arrowheads, respectively. **(E)** Representative TEC-Topl complexes showing RNAPs escape from the apices. **(F-G)** Quantification of the non-apical RNAP ratio (N=90) and plasmid’s apex number (N=85), respectively. **(H)** Superimposition of all bound RNAPs (top panel) and all apical DNA segments (bottom panel). **(I-J)** DNA supercoiling relaxation assays mediated by Topi and hToplB in the presence of RNAP, respectively, with ΔLk quantification shown on the right.

Bulk transcription assays on –sc plasmid with TopI confirmed markedly improved transcription (**Figure 5B, yellow vs. green, and Figure S6E**), exhibiting kinetics similar to those of nicked templates without torsional stress (**Figure 5B, orange**). Based on these results, we conducted a cryo-ET imaging study of RNAP transcription in the presence of TopI (RNAP:TopI:plasmid=3:1:1). 10 minutes after the re-start of the sTEC, TEC-TopI sample revealed the presence of large RNA transcripts (**Figure 5C**). Moreover, TopI activity alone is sufficient to displace 50% of RNAPs from their apical localization, even in the absence of a torsional block (**Figure 5D, green arrows, and Figure 5E**), yielding the highest off-apex ratio across all tested conditions (**Figure 5F**), and without inducing plasmid branching (**Figure 5G**). Quantitative analysis showed that blunt apices predominated in TopI-relaxed plasmids, in contrast to unrelaxed ones, with a marked reduction in curvature (from 0.25 to 0.18 nm⁻¹; *P* <0.0001) and an increase in DNA spacing (from 7.3 to 15.2 nm; *P* <0.0001). (**Figure S6F**). We attribute this prominent apex structural change to the relaxation of the plectoneme (mean Wr =−6.4; **Figure S6F**). Analysis of RNAP orientations in TEC-TopI particles reveals highly dynamic entry–exit DNA (**Figure 5H**), in contrast to the apically constrained configuration observed in TECs (**Figure 3B**). These results suggest that not only torsional constraint, but also a high degree of −sc, is required for RNAP apical localization and the longer pauses observed during single-round transcription.

Interestingly, while TopI facilitates RNAP transcription, RNAP also modulates TopI activity. Previous bulk assays showed that TopI alone (1:1 with plasmid) rapidly relaxed supercoils (ΔLk from –15 to –1) within minutes (**Figure S5A**). However, the addition of RNAP (plasmid:RNAP:TopI = 1:3:1) notably slowed relaxation kinetics, with topoisomer bands clustering at ΔLk < –4 even after 30 minutes (**Figure 5I**). 2D gel further resolved these bands, revealing accumulation of topoisomers at ΔLk = −8 (**Figure S7A**). Given the reported coupling between RNAP and TopI^8^ (**Figure 5A, dashed arrow**), which could potentially impair TopI function, we performed a control experiment with excess TopI (plasmid:RNAP:TopI = 1:3:9; **Figure S7B**). The relaxation kinetics were similar to those observed at a 1:3:1 ratio, indicating that the slower relaxation is not due to TopI inactivation upon interaction with RNAP, but rather reflects RNAP’s torsional block function—similar to the dCas9-mediated TopI downregulation (**Figure S5A**). Moreover, hTopIB, known not to interact with RNAP, showed a similarly slowed relaxation pattern on RNAP-bound plasmids, further supporting the above interpretation (**Figure 5J**). Interestingly, the torsional block effect of RNAP appears to be more pronounced than that of dCas9 (dCas9-topoisomers clustered at ΔLk = −1 to −3), likely due to its larger size and its ability to induce sharper kinks when it melts the DNA duplex. Indeed, supporting the requirement of DNA melting for DNA-binding protein to act as a roadblock, use of RNAP core—which binds DNA but cannot form stable transcription bubbles—resulted in faster relaxation than the holo-enzyme and ended in fully relaxed plasmids. (**Figure S7C**). Consistently, dead EcoRI mutant (dEcoRI), which bends but does not melt DNA, failed to act as a torsional barrier and permitted rapid, full relaxation (**Figure S7D**). These results suggest that both DNA bending and duplex melting are required for a protein to function effectively as a torsional block.

### Tandem promoters facilitate supercoiling domain formation and favors transcription over opposing promoters

We learned from the above experiments that the restricted rotational freedom of RNAP at apices confers dual functions: (1) generation of torsional strain during transcription, a prerequisite for the twin-domain model, and (2) serve as a torsional roadblock for a second transcribing RNAP as is also observed with dCas9. This dual function parallels the cellular scenario where multiple RNAPs co-transcribe the same genomic DNA, enabling transcriptional coupling between neighboring genes through the formation of topological domains.^62^ To gain insight into this system, we designed dual-promoter circular pUC19 plasmids (2.8 kb) containing two identical T7A1 promoters, each with a U-less stalling sequence, driving transcription of AmpR and mEGFP, either in opposite (topologically mixed convergent and divergent; **Figure 6A, left**) or tandem (i.e., same-direction; **Figure 6B, left**) orientations. After UTP starvation–induced stalling, TECs were imaged after 10 minutes of transcription. To facilitate visualization of these more complex structures, we projected plasmids onto a circular layout (**Figure 6A-B, middle**), with the rim color-coded to match the 3D molecular model (**Figure 6A-B, right**). Arrowheads indicate the transcription direction of bound RNAPs, and circles mark apical sites. Inner circular sectors represent individual plectonemes (e.g., I, II, and III), color-coded by its writhe density (blue–pink–red, −0.3 to 0.1) (**Figure 6A, middle**).

**Figure 6:**
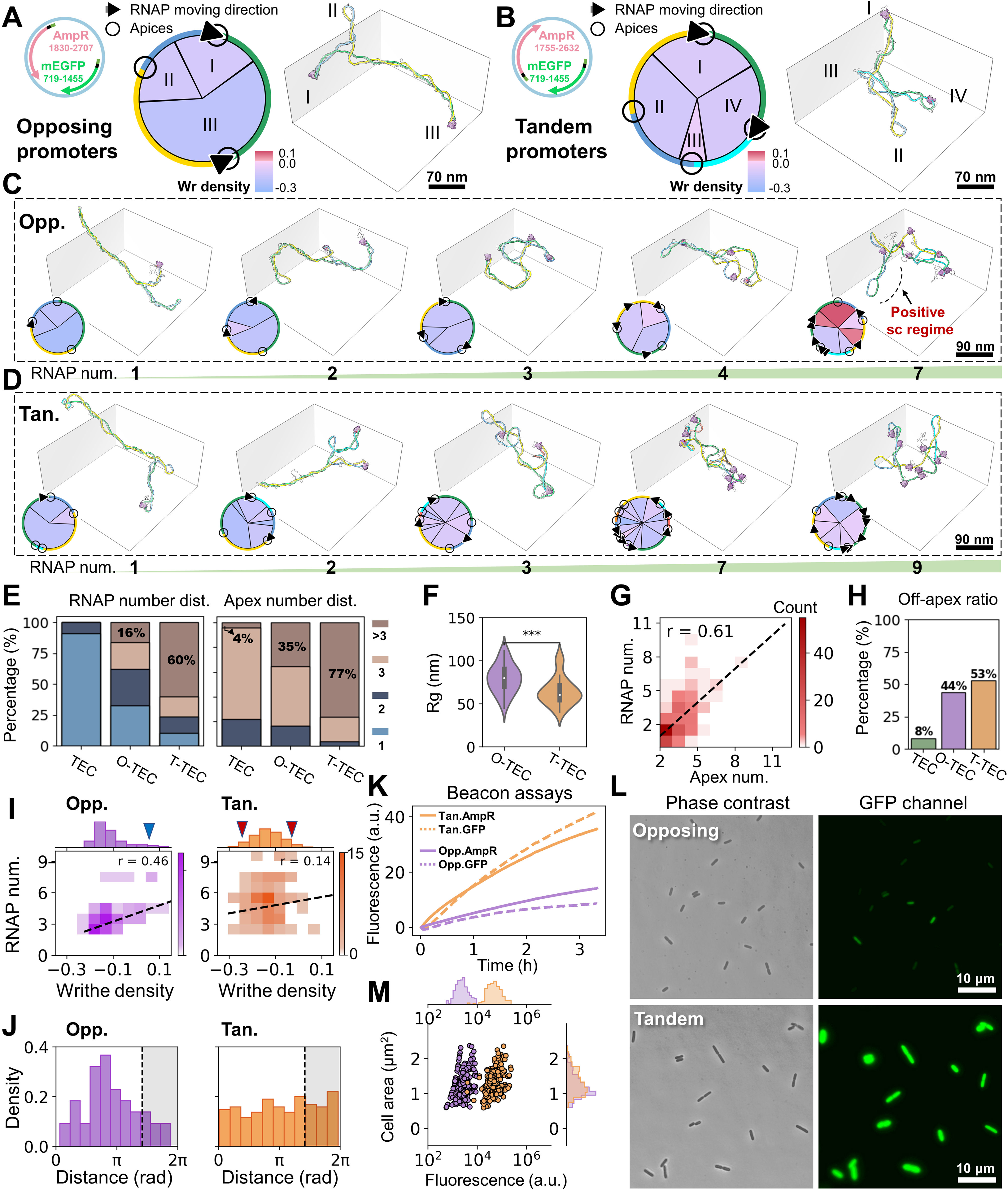
Tandem promoters enhance topological domain formation and transcription compared to opposing promoters. **(A)** Opposing dual-T7A1U promoter construct (left) driving transcription of AmpR and mEGFP. Circular plasmid layout (middle) with rim color-matched to the 3D model (right). Arrowheads indicate RNAP transcription direction; circles mark apical sites. Inner sectors represent individual plectonemes (l-III), color-coded by writhe density (blue-pink-red). **(B)** Tandem promoter construct, identical to A, but with AmpR flipped. **(C-D)** Representative TECs from opposing and tandem constructs, ordered by RNAP count. **(E)** Plasmid distribution by bound RNAP number and apex count; N=90. **(F-H)** Quantification of plasmid Rg (N=67), correlation between RNAP number and apex count (N=206), and ratio of RNAP located off-apex; N=232. (I) Plectoneme writhe density distribution vs. bound RNAP number; N=230. **(J)** Distance distribution of opposing RNAPs (left) and tandem RNAPs (right) on the circular plasmid layout. Shaded regions indicate less likely RNAP separations, attributed to the non-specifically bound RNAP population; N=212. **(K)** Quantification of *in vitro* gene expression by molecular beacon assay. **(L-M)** Imaging and quantification of in vivo GFP gene expression in *E. coli* MG1655 carrying either plasmid of tandem or opposing constructs; N=529.

Significantly, as RNAP loading increased, plasmids underwent major reconfiguration from elongated plectonemes to expanded, globular structures (**Figure 6C, opposing, and Figure 6D, tandem**). Under identical reaction conditions, we observed a higher number of RNAPs in tandem TECs (T-TECs) compared to opposing TECs (O-TECs). T-TEC particles exhibited up to 10 bound RNAPs and 9 apices, with 60% of plasmids bound by more than 3 RNAPs and 77% forming more than 3 branches (**Figure 6E**). In contrast, multi-RNAP loading and multi-apices were down to 16% and 35% in O-TECs, and dropped further to 0% and 4%, respectively in single-promoter TECs, respectively. As a result, the highly branched T-TECs exhibited reduced plectonemic extension, with a significantly smaller Rg compared to O-TECs (64 nm vs. 79 nm; *P* <0.001, **Figure 6F**). Statistical analysis confirmed a positive correlation between RNAP occupancy and plasmid branching (*r* = 0.61, **Figure 6G**), suggesting that RNAP number potentially reflects the generation of domain separation. Notably, the higher number of RNAPs was not necessarily localized at apices. In both O-TECs and T-TECs, increased multi-RNAP binding led to elevated off-apex RNAP localization, reaching 44% for O-TECs and 54% for T-TECs (**Figure 6H**), in contrast to off-apex localization of single TECs (8%), but comparable to levels observed in TEC-Cas and TEC-TopI (45% and 50%, respectively).

The overall phenomena of multi-RNAP binding and multi-branching are not fundamentally distinct from those observed in TEC-Cas, but rather represent more extreme cases, especially in T-TECs. This result supports the continued validity of the TEC-Cas model (**Figure 4E**), with the RNAP version of “dCas9” reinforcing domain separation through its dual functions. In this context, we compare O-TECs and T-TECs in their supercoiling domain formation. Visually, as more RNAPs were loaded, a shift toward non-plectonemic DNA loops (lower negative writhe density; pink circle segments, **Figure S7E**) was more frequently observed in T-TECs than in O-TECs. This trend was further confirmed as increased number of RNAPs led to a broader writhe density distribution of T-TECs, displaying enriched hypo and hyper –sc domains compared to O-TECs (**Figure 6I, red arrows**). In contrast, a subset of O-TECs exhibited plectoneme writhe densities extending into the far-positive regime (**Figure 6I, blue arrow**). Examination of these particles revealed, for the first time, TECs with long positive plectonemes (**Figure 6C and S7E, red label**), supporting that opposing RNAPs, transcribing toward each other mutually act as torsional blocks that accumulate positive supercoiling. One T-TEC also showed short positive supercoiling (**Figure S7E**), but likely due to DNA entanglement between plectonemes. Nevertheless, this observation also suggests the possibility of mutual torsional interference between tandem RNAPs due to differential translocation speeds, consistent with the on–off apex model observed in TEC-Cas particles.

Interestingly, we observed a higher correlation between the writhe density and RNAP number in the O-TECs than T-TECs (r =0.46 vs. r =0,14, **Figure 6I**). We interpret it as the accumulation of torsional stress in the opposing geometry leading to higher changes in writhe compared to the cancellation of torsional stress when RNAP transcribes in tandem (i.e., further multi-RNAP loading has limited impact on torsional accumulation). Furthermore, by identifying all possible RNAP pairs on the plasmid and measuring their angular separation in polar plots, we found that opposing RNAPs were more likely to be separated by 2.7 radians (∼1.2 kbp) (**Figure 6J**), in contrast to the uniform distribution observed for tandem RNAPs—further supporting a model of transcriptional hindrance in opposing configurations.

To confirm the functional consequences of the observed structural differences between tandem and opposing gene arrangements, we measured gene expression both in vitro and in vivo. RNA molecular beacon assays^63^ revealed comparable in vitro transcript levels of AmpR and mEGFP within the same construct (**Figure 6K**), as expected since both genes are driven by identical promoters (T7A1). However, higher RNA levels were observed with the tandem arrangement compared to the opposing one for both genes (**Figure 6K**). We also compared in vivo gene expression of mEGFP by measuring fluorescence emission in live *E. coli* cells harboring either construct, grown in M9GluCAAT medium. Consistent with the in vitro results, tandem constructs showed nearly a ten-fold higher fluorescence than their opposing counterparts (**Figure 6L-M**). These results are highly consistent with our cryo-ET observations: the tandem gene arrangement promotes increased transcriptional activity, including higher RNAP loading, increased numbers of off-apex RNAPs less prone to pausing, and lower accumulated torsional stress due to the cancellation of neighboring RNAPs.

## DISCUSSION

We demonstrate that cryo-ET, combined with modern image processing, enables detailed 3D visualization of highly dynamic DNA/protein complexes previously underexplored. By reconstructing the full DNA backbone of plasmids extracted from cells and bearing natural −sc densities, we precisely quantify plectoneme conformational dynamics. This marks a major advance in understanding DNA supercoiling, previously limited to surface-level AFM,^10,64^ negative-stain electron microscopy,^65^ or cryo-ET imaging of ∼300 bp minicircles.^12^ By directly imaging individual ∼2-3 kb plasmids, we quantify writhe and show that under near-physiological ionic strength (40 mM KCl, 5 mM MgCl₂), torsional stress is stored primarily as writhe (Wr ≈ −10 for ΔLk ∼ −15; **Figure S1B**, **S1D**). We find that lowering the ionic strength reduces Wr to −5, increases DNA strand spacing of plectonemic DNA and lowers apex curvature. Previously, such 3D conformational changes were accessible only through coarse-grained Brownian dynamics simulations.^66,67^ Our findings provide a basis for refining theoretical models to better reflect DNA’s typical topological state in cells.

Our full DNA tracing in 3D reveals that DNA-melting proteins like RNAP and dCas9 preferentially localize at plectoneme apices when bound to –sc DNA (**Figures 2A, 2F**). We find that this apical localization, first noted by ten Heggeler-Bordier et al.^32^ and revisited by Janissen et al.,^9^ is sequence-independent and persists as RNAP translocates along DNA (**Figure 3A**). Direct observation of RNA transcripts shows that this geometry helps segregate nascent RNA from the DNA body (**Figure 3A-B**), potentially aiding co-transcriptional events such as RNA folding, RNA processing, ribosome binding, while preventing RNA-DNA hybrid formation. Notably, RNAP localizes apically only at σ ≈ −0.08 (ΔLk = −15); relaxation by TopI to σ ≈ −0.04 (ΔLk = −8) reduces this localization (**Figure 5F**), suggesting a possible role of −sc on gene expression.

Beyond stabilizing open complex formation,^37^ we show that −sc promotes transcription by facilitating promoter escape and reducing abortive initiation (**Figure 2D**). The ∼170° bending induced on the upstream DNA in the apically positioned TECs disrupts the majority of σ-factor:DNA contacts, a requirement for promoter clearance, offering an additional layer of regulation and helping rationalize the supercoiling sensitivity of certain bacterial promoters.^68^ Negative supercoiling also enhances RNAP binding following the formation of hyper-negative domains (**Figure S6B**), likely accelerating promoter search and triggering transcriptional cascades.

However, supercoiling can also impede transcription by promoting RNAP pausing (**Figure 3F**), in line with single-molecule studies.^21^ In contrast to those studies, which used surface fixed DNA, our work shows that under no additional torsional barrier other than the inherit-torsional stress of plectonemic DNA, the apical localization of RNAP can induce a swiveled state in RNAP with increased propensity to pausing (**Figure 3G**). While nascent RNA structures, transcription factors, and DNA sequence are known to promote pausing by stabilizing the swiveled state, our results reveal that DNA topology, through apical localization, can similarly regulate this conformational transition. This supercoiling-induced pausing is reversed by TopI (**Figure 5B**) and also by the action of co-transcribing RNAPs in tandem, which reduces apical localization (**Figure 6H**), restores elongation, and promote a more “fluid” transcriptional mode.^24^

The twin-supercoiled domain model has long explained transcription-induced supercoiling in prokaryotes and eukaryotes,^5,24,69,70^ but direct visualization of the resulting supercoils (**Figure 7A, left**) has remained elusive. Early models suggested that long RNA transcripts or anchoring to cellular structures were required to prevent RNAP rotation and allow torsional stress to build. Here, we show direct structural evidence that rotationally constrained, apically localized RNAP—with even short (∼20 nt) transcripts—can induce new plectoneme formation. This suggests long transcripts are not necessary to constrain RNAP, aligning with recent RNase-treatment studies.^9^ A single apically positioned RNAP can generate plectoneme branching, as observed in our experiments and predicted by other simulation work,^71^ though the latter required unrealistically high elongation rates and forces. Incorporating our 3D model of the apical RNAP configuration may help reconcile this discrepancy. Under physiological conditions with initial negative supercoiling, positive supercoil formation was rare, contrary to textbook models. Instead, we observed writhe density changes upstream and downstream of RNAP (**Figure 7A, right**), especially when torsional barriers such as neighboring RNAPs were present. This role of RNAP and DNA-melting proteins like dCas9 may extend to similar enzymes in both domains of life. DNA-bending proteins like dEcoRI did not show this effect, but other factors such as nucleoid-associated proteins^72^ or nucleosomes—which store ∼1.7 DNA turns—may also act as torsional barriers.^71,73^ Together, these observations define a revised twin-supercoil model that updates a canonical textbook framework under physiologically relevant, tether-free negative supercoiling.

**Figure 7.**
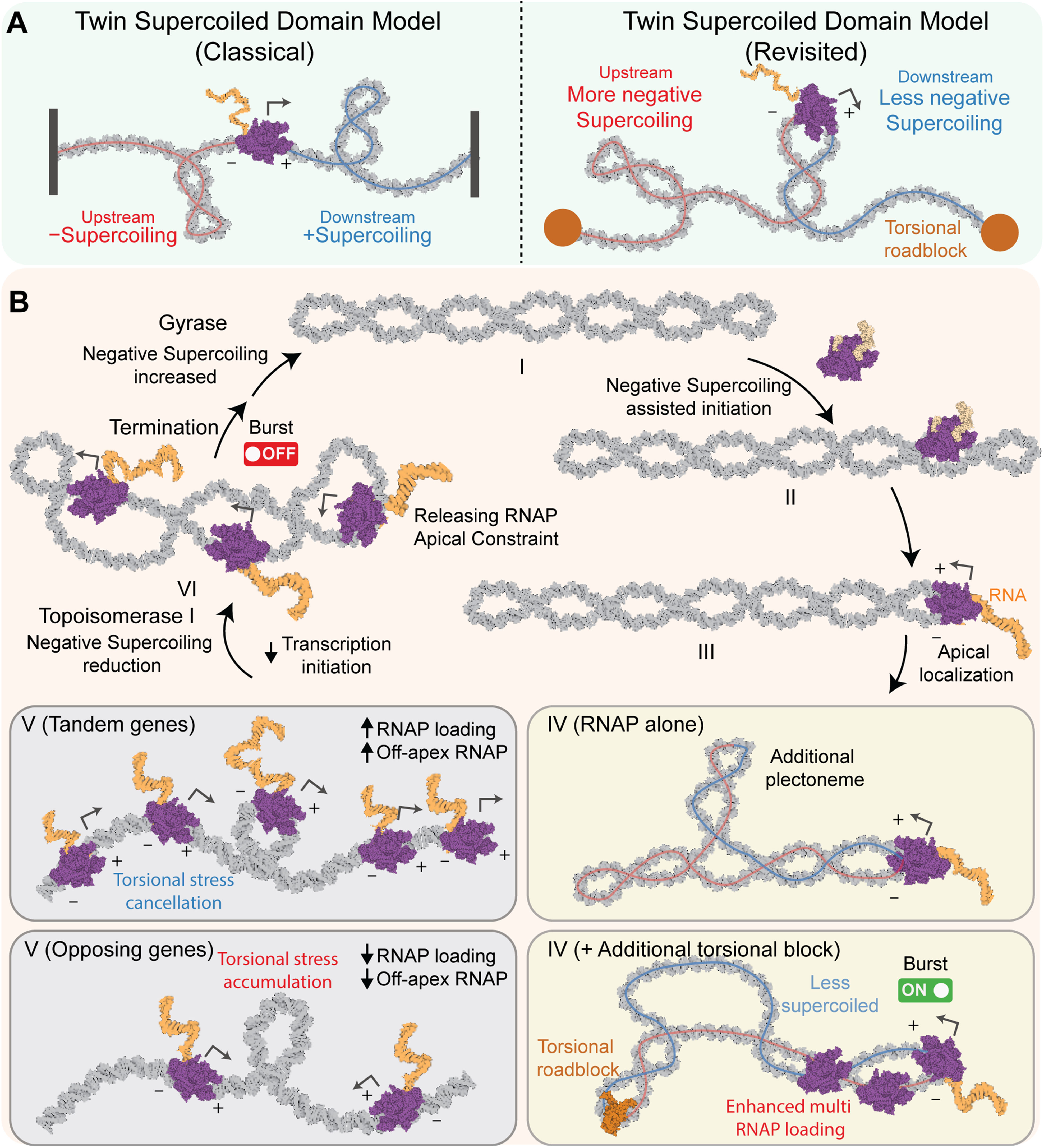
Revisited twin-supercoiled domain model and its connection to transcriptional bursting. **(A)** Comparison of classical twin-supercoiled domain and a revisited model considering the apical localization of RNAP in -sc DNA. **(B)** Summary model connecting transcriptional induced torsional stress and transcriptional bursting in -sc DNA. (See Discussion for further description of the model).

Negative supercoiling promotes RNAP initiation but hinders elongation, supporting a pulse-like transcription model reminiscent of in vivo “transcriptional bursting”.^74,75^ Single-molecule studies have linked supercoiling to this behavior.^22^ Based on our findings, we propose an updated model for −sc DNA (**Figure 7B, I–III**). Initially (**Figure 7B, I–III**), physiological DNA stores negative torsion that facilitates transcription initiation, promoter escape, and apical RNAP localization. In the absence of additional torsional roadblocks besides an apically located RNAP, transcription-induced supercoiling may lead to the formation of additional plectonemes (**Figure 7B, IV, RNAP alone**). In contrast, additional torsional blocks segment topological domains—reducing the negative writhe in front of the RNAP while increasing it behind, leading to enhanced RNAP binding and transcriptional burst initiation (**Figure 7B, IV +additional torsional block**). When RNAPs bind in tandem, they can cancel each other’s torsional stress, promoting transcriptional synergy (**Figure 7B, V, Tandem**); however, opposing RNAPs may stall due to accumulated stress (**Figure 7B, V, Opposing**). To complete transcription, topoisomerase relieves −sc, displacing RNAPs from the apex and facilitating elongation (**Figure 7B, VI**). The requirement for topoisomerase in active transcription is supported by the direct interaction between *E. coli* TopI and RNAP^8,60^ and by the observed recruitment of topoisomerase at transcription start sites.^61^ The resulting relief of negative torsion reduces the likelihood of new initiation, effectively acting as an OFF switch. The cycle can restart via ATP-dependent DNA gyrase activity, which restores negative supercoiling.

## RESOURCE AVAILABILITY

### Lead contact

Further information and requests for resources and reagents should be directed to and will be fulfilled by the Lead Contact, Carlos Bustamante.

### Materials availability

All unique reagents generated in this study are available from the lead contact with a completed Materials Transfer Agreement.

## DATA AVAILABILITY

The cryo-ET maps of ∼2 kb negatively supercoiled pUC19-T7A1U plasmids under various conditions were montaged and deposited in the Electron Microscopy Data Bank (EMDB). These include maps of plasmids under low salt conditions (EMD-47843), high salt conditions without additional proteins (EMD-47844), with stalled RNA polymerase (EMD-47847), with dCas9 (EMD-47849), with RNA polymerase during active transcription (EMD-47850), with both stalled RNA polymerase and dCas9 (EMD-47851), with transcribed RNA polymerase and dCas9 (EMD-47853), and with transcribed RNA polymerase and Topoisomerase I (EMD-47855). The cryo-ET maps of ∼2.8 kb negatively supercoiled pUC19-T7A1U plasmids containing opposing promoters and tandem promoters were montaged and deposited in the EMDB under accession numbers EMD-71622 and EMD-71618, respectively. The single-particle analysis maps of TEC1–SC and TEC2–SC were deposited in the EMDB under accession numbers EMD-71675 and EMD-71676, with their corresponding atomic models available in the PDB under accession codes 9PIP and 9PIQ, respectively. The individual particle 3D reconstructed maps, models, original gel images, fluorescence images, the script calculate the TECs swivel angle, and data used to generate the statistical analysis presented in the figures have been deposited on GitHub Zenodo at (https://doi.org/10.5281/zenodo.16423113).

## ACKNOWLEDGMENTS

We thank Dr. Liang Meng Wee for the expression and purification of dCas9. We thank Rodrigo Fregoso, Matthew Lo, Allison Cordova and Jimena Pavlovitch for assistance in the early stages of the project. We also thank Dr. Gang Ren for technical support and valuable feedback, and Professor Zev Bryant and Dr. Noor Sayyad for their insightful comments on the work. Data were collected at the Cal-Cryo facility, Berkeley QB3 Institute. The work at the molecular foundry, LBNL, was supported by the Office of Science, Office of Basic Energy Sciences of the United States Department of Energy (contract no. DE-AC02-05CH11231). This research was supported by grants from the US National Institutes of Health R01GM032543 (C.B). C.B. is Howard Hughes Medical Institute investigator.

## AUTHOR CONTRIBUTIONS

M.Z., C.C.-C., and C.B. conceived the study and designed the research. C.C.-C. prepared the oligonucleotide and plasmid. C.C.-C. performed the bulk transcription assay. C.C.-C., S.D.C., and K.I.R. performed the relaxation assays. C.C.-C. and K.I.R. performed in vitro and in vivo measurements of gene expression for the opposing and tandem constructs. B.O. conducted the AFM imaging. M.Z. prepared the EM specimens, and M.Z. and J.L. collected the EM data. M.Z. managed the 3D reconstruction workflow. M.Z., S.D.C., and E.C. contributed to the modeling. M.Z. performed single particle cryo-EM analysis. M.Z. interpreted the data and prepared the figures. J.F and M.Z. performed the molecular dynamic simulation. M.Z. and C.C.-C. wrote the original draft, with revisions by B.O., J.L., S.D.C., K.I.R., E.C. and C.B. supervised the work.

## MATERIALS AND METHODS

### Oligonucleotides and RNA

All DNA oligonucleotides listed below were obtained from Integrated DNA Technologies (IDT). Except for PCR primers, all oligonucleotides were purified in-house using denaturing urea-polyacrylamide gels (PAGE) prepared from SequaGel UreaGel 29:1 Concentrate (National Diagnostics).

### Preparation of single guide RNAs (sgRNAs)

sgRNA specific oligonucleotides and scaffold oligonucleotides (CBD03 and CBD04) were ordered from IDT and purified in-house by polyacrylamide-gel electrophoresis (PAGE). In a 30 μL reaction, oligonucleotides (20 μL, 50 μM) were annealed and extended by T4 DNA polymerase in CutSmart 1X buffer to produce the dsDNA template. After heat inactivation, 10 μL of this reaction (∼33.3 μM dsDNA template) were used for a 20 µL in vitro transcription reaction using HiScribe T7 In vitro Transcription Kit (New England Biolabs). In vitro transcription reactions were performed at 37°C for 6 hours, after which Dnase I was added to degrade the DNA template. sgRNAs were purified by urea PAGE. Bands corresponding to the correct RNA size were cut out as gel slices, eluted overnight at room temperature in 2X PK buffer (200 mM Tris-HCl, pH 7.5, 25 mM EDTA, pH 8.0, 300 mM NaCl and 2% SDS (w/v)), phenol-chloroform extracted and precipitated with 2X volume of 200-proof 100% ethanol (Koptec). Then, samples were air dried and suspended in UltraPure DNase/RNase-free distilled water (Invitrogen, Thermo Fisher Scientific).

### Cloning of Single promoter plasmid pUC19-T7A1U, pUC19-T7A1-AmpR-mEGFP-Tandem and pUC19-T7A1-AmpR-mEGFP-Opposing constructs

Single promoter plasmid pUC19-T7A1-U was generated using around-the-horn PCR from template plasmid pUC19 using primers (CBD01/CBD02) with extensions containing the T7A1 promoter sequence as well mutations in the initially transcribed region to form a U-less cassette. These primers also contained EcoRI recognition sites for subsequent digestion and circularization by ligation. PCR products were digested with EcoRI and DpnI and were further purified by agarose-gel extraction. Purified PCR products were ligated using T4 DNA Ligase and transformed into DH5α cells. The tandem construct (pUC19-T7A1-AmpR-mEGFP-Tandem) was generated using Gibson Assembly of three PCR fragment assembly reaction: fragment 1 (PCR with template pUC19-T7A1-U using primers CBD05/06, fragment (PCR with template pUC19-T7A1-U using primers CBD07/08), and fragment 3 (PCR from plasmid containing mEGFP using primers CBD09/10). Each PCR product was purified by gel-extraction. Gibson Assembly was carried using NEB Gibson Assembly Master Mix (NEB, #E2611L) following manufacturer instructions and then assembly reaction transformed into DH5α cells. The opposing construct (pUC19-T7A1-AmpR-mEGFP-Opposing) was obtained using Golden Gate Assembly from two PCR fragments: fragment 1 (PCR with template pUC19-T7A1-AmpR-mEGFP-Tandem using primers CBD11/12) and fragment 2 (PCR with template pUC19-T7A1-U using primers CBD13/14). The PCR fragments were treated with BsaI (NEB) and BsmbI (NEB), respectively, and then ligated using T4 DNA ligase (NEB). Given the instability of this plasmid in DH5α cells, NEB^®^ Stable Competent *E. coli* cells were used for transformation instead. In all cases, plasmids from positive clones were extracted using QIAGEN miniprep kit and their sequences were confirmed by DNA sequencing.

For large scale production of plasmids, plasmids were retransformed into DH5α or NEB Stable competent cells, accordingly. Cells were grown in 1 L of LB media and extracted using QIAGEN Maxiprep kit following manufacturer instructions. For bulk biochemical as well as cryo-ET structural studies, supercoiled plasmid pUC19-T7A1-U was further purified using low-melting agarose (SeaPlaque Agarose, Lonza) from 1% agarose gel (1X TAE with 1X SYBR Safe DNA Staining). Under these electrophoretic conditions nicked DNA was well separated from supercoiled DNA. The band corresponding to supercoiled DNA was excised and extracted using β-Agarase (New England Biolabs), followed by phenol-chloroform extraction and ethanol precipitation. To desalt and concentrate the samples, the purified plasmids were further purified with Monarch PCR & DNA Cleanup Kit (New England Biolabs) using manufacturer instructions. The isolated purified plasmids were stored at −20 °C until further use.

### Generation of different DNA topologies

To generate different DNA topologies (nicked, ΔLk=0, ΔLk < 0, and ΔLk>0) for AFM, pUC19-T7A1-U plasmids were treated with different enzymes to obtain the desired topology and they were further purified by agarose gel extraction using the β-Agarase described above. To generate nicked topology, plasmids (20 µg) were incubated in a 100 µL reaction with nicking endonuclease Nt.BspQI (20 units) in NEB 3.1 Buffer for 6 hours at 50 °C. For ΔLk = 0 topology, nicked plasmids (20 µg) were re-ligated by incubation in a 50 µL mixture with T4 DNA ligase (800 units) in T4 DNA ligase 1X Buffer at room temperature. For ΔLk < 0 topology, we used isolated plasmids with their native supercoiling degree as extracted from the stationary phase cells. For ΔLk > 0 topology, plasmids (600 fmol) were incubated in a 20 μL reaction with reverse gyrase TopR2 (0.1 μM), and ATP (1 mM) in Reverse Gyrase 1X Buffer (50 mM Tris–HCl pH 8.0, 20 mM MgCl_2_, 100 mM NaCl, 0.5 mM DTT, and 0.5 mM EDTA) for 1 hour at 75 °C. Once plasmid topologies were generated, plasmids were purified using the Monarch PCR DNA Clean-up Kit.

### Expression and purification of *Streptococcus pyogenes* catalytically inactive dCas9 and nickase Cas9 D10A (SpCas9 D10A)

Purification of SpCas9 D10A was performed using a reported protocol.^76^ Briefly, plasmid MJ825 (Addgene, #39315) or MJ841 (Addgene, #39318) was transformed into BL21(DE3) cells. One liter of Terrific Broth culture containing 50 µg/mL kanamycin was grown at 37°C. Upon reaching OD 0.6, cells were induced with IPTG to a final concentration of 0.5 mM at 20°C and were grown overnight (∼16 hours). Cells were harvested by spinning at 5000 rpm for 10 minutes and the cell pellet was stored at −80 °C. Cells were resuspended in 50 mL Lysis Buffer (50 mM Tris-HCl pH 7.5, NaCl 500 mM, 5% (v/v) glycerol, 1 mM DTT, supplemented with 4 tablets of mini-EDTA free protease inhibitor) and lysed using a sonicator. The lysate was centrifuged for 30 minutes at 4°C at 20,000 g. The clarified supernatant was passed through a HisTrap 5 mL column, washed with buffer NiA (50 mM Tris-HCl pH 7.5, NaCl 500 mM, 5% (v/v) glycerol, 1 mM DTT, 10 mM imidazole) and eluted on a linear gradient with buffer NiB (50 mM Tris-HCl pH 7.5, NaCl 500 mM, 5% (v/v) glycerol, 1 mM DTT, 300 mM imidazole). Positive fractions containing SpCas9 D10A (verified by SDS-PAGE) were combined, and 2 mL of TEV protease were added and dialyzed overnight on Dialysis buffer (50 mM Tris-HCl pH 7.5, NaCl 500 mM, 5% (v/v) glycerol, 1 mM DTT). The samples were further purified on HiTrap Heparin column using a linear gradient of KCl from 200 mM to 1000 mM (50 mM Tris-HCl pH 7.5, 5% (v/v) glycerol, 1 mM DTT), followed by size-exclusion chromatography on a Superdex S300 column using Gel Filtration buffer (20 mM Tris-HCl pH 7.5, KCl 200 mM, 5% (v/v) glycerol, 1 mM DTT).

### Expression and purification of *S. solfataricus* reverse gyrase (TopR2)

The gene fragment expressing the reverse gyrase (TopR2) was obtained by PCR from *Sulfolobus solfataricus* genomic DNA (ATCC, 35092D-5). This fragment was inserted into an expression plasmid with His6 N-terminal tag with a TEV protease site (2Bc-T, Addgene #37236) using ligation independent cloning. Protein expression was carried with this plasmid (2Bc-T-TopR2) following a reported protocol.^77^ Briefly, plasmid 2Bc-T-TopR2 was transformed into Rosetta (DE3) cells. One liter of 2xYT culture containing 100 μg/mL ampicillin, 34 μg/mL chloramphenicol, and 1% glucose was grown at 37°C. Upon reaching OD 0.5, cells were induced with IPTG to a final concentration of 1 mM at 37°C and were grown for 6 hours. Cells were harvested by spinning at 5000 rpm for 10 min and the cell pellet was stored at −80°C. The pellets of induced cells were resuspended in Lysis buffer (40 mM Tris–HCl pH 8.0, 100 mM NaCl, 1 mM DTT, 0.1 mM EDTA). After sonication on ice, the sample was centrifuged at 30,000 g for 30 minutes. The resulting supernatant underwent heat treatment at 75°C for 13 minutes and was clarified by centrifugation at 30,000 g for 20 minutes. Polyethylenimine was then added to achieve a final concentration of 0.3%, and the mixture was stirred for 1 hour before being centrifuged at 40,000 g for 30 minutes. The supernatant was adjusted to 70% (NH_4_)_2_SO_4_ saturation, stirred for 30 minutes at 4°C, and centrifuged at 20,000 g for 30 minutes. The proteins were dissolved in buffer B (Lysis Buffer with 200 mM NaCl and 10% (v/v) ethylene glycol). After dialysis against buffer B, the proteins were loaded onto a 5 mL HiTrap Heparin HP column pre-equilibrated with buffer B using an FPLC AKTA system (GE Healthcare). The column was washed with 50 mL of buffer B, and bound proteins were eluted with a NaCl gradient ranging from 0.2 to 1.2 M NaCl in buffer B.

### SpCas9 D10A nicking assay

To test the activity of the sgRNA and nickase dCas9, a nicking assay was developed to monitor activity by electrophoretic mobility of DNA in agarose gel. sgRNA (90 fmol) was incubated at room temperature for 5 min with SpCas9 D10A (90 fmol) in Cas9 Reaction Buffer (20 mM HEPES pH 6.5, 100 mM NaCl, 5 mM MgCl_2_, 0.1 mM EDTA). Supercoiled plasmid (9 fmol) was added, and the reaction was kept at 37°C for one hour. The samples were treated with Proteinase K and then loaded into an agarose gel (1% agarose, 1X TAE 2.5 μg/mL chloroquine).

### DNA topoisomerase relaxation assay

The DNA relaxation activity of *E. coli* DNA topoisomerase I on supercoiled pUC19-T7A1U was tested under different reaction conditions. In all reaction conditions, a total reaction volume of ∼40 μL containing ∼200 ng (∼ 0.157 pmol) of negatively supercoiled plasmid was used and relaxation was carried out in TB40 buffer (20 mM Tris pH 8.0, 40 mM KCl, 5 mM MgCl_2_, 0.02 mg/mL BSA, and 1 mM DTT). An incubation at 37°C for 20 minutes was done for all conditions prior to the addition of *E. coli* Topoisomerase I (New England Biolabs, M0301S, 0.720 pmol). (1) For Topoisomerase I only condition, topoisomerase was added after an incubation of 37°C for 20 minutes. (2) To test the effect of a stalled transcription complex, *E.coli* RNA Polymerase Holoenzyme (New England Biolabs, M0551S, 1.700 pmol) was supplemented to the previous reaction condition with 10 μM of ATP, GTP, CTP (Thermo Scientific, R0481) and 50 μM of GpA dinucleotide (TriLink Biotechnologies) and incubated during 20 min at 37°C. (3) To test for the effect of a dCas9: sgRNA complex, a sgRNA:dCas9 complex at a concentration of 1.55 μM was first formed by incubating 15.5 pmoles of sgRNA and 15.5 pmoles of dCas9 in TB40 buffer (10 μL) at 25°C for 10 minutes. Afterward, the sgRNA: dCas9 complex was incubated with the supercoiled plasmids for 20 minutes at 37 °C. (4) To test the effect of *E.coli* RNAP without DNA-melting, *E.coli* RNAP core (NEB, 1.7 pmol) was supplemented to the plasmid-only condition and incubated at 37°C incubation prior to the addition of topoisomerase. (5) To test the effect DNA binding protein that does not melt DNA, dEcoRI (EcoRI Q111, 2 pmol, a gift from ModrichLab) was supplemented to the plasmid-only condition and incubated at the 37°C incubation prior to the addition of topoisomerase.

In each assay, after 20 minutes of incubation, *E. coli* Topoisomerase I (NEB, 0.72 pmol) or human topoisomerase (hTop1B, 4U,Topogen) was added to a total reaction volume of 40 μL and time-points were taken at 0, 1, 2, 3, 5, 10, 15, and 30 minutes and quenched in a 100 μL of a Stop Buffer containing 8 U Proteinase K (NEB), 4 μL Glycogen (5 mg/mL, Invitrogen), and 86 μL of 2X PK buffer (Tris 200 mM pH 7.5, EDTA 25 mM, NaCl 300 mM, SDS 2%). The timepoints were extracted with phenol (pH > 7.5) and phenol/chloroform/isoamyl alcohol (25:24:1 Mixture, pH 6.7/8.0, FisherBioReagents™) and then precipitated with 70% ethanol and redissolved in EB buffer. The time points were electrophoresed using a 1.5% (w/v) agarose gel that was run at 80V for 3 h with 1X TAE buffer (40 mM Tris-acetate, pH 8.1, 2 mM EDTA) in ice and post-stained in a solution of SYBR Safe in water before being visualized using Typhoon imager.

### 2D agarose gel electrophoresis

#### DNA topoisomer ladder generation

Plasmid pUC19-T7A1U was nicked with Nt.BspQI (New England Biolabs). Nicked pUC19-T7A1U plasmids (400 ng) were religated in 1X T4 DNA ligase buffer with increasing amounts of ethidium bromide (0, 40, 80, 120, 160, 200, 400, and 800 ng) using 100 U of T4 DNA ligase (New England Biolabs) in 40 μL reactions. The reaction proceeded overnight at room temperature and was quenched with Proteinase K (NEB), followed by extraction using 2-butanol, phenol-chloroform extraction, and ethanol precipitation. A mixture of the different ligation reactions was made to generate the 2D topoisomer ladder.

#### Separation of DNA topoisomers in 2D agarose gel

A large 2% agarose slab was cast in 1X TAE containing 2.5 μg/mL chloroquine. 100 ng of plasmid was loaded on a narrow lane in the 2D gel and electrophoresis was carried out with the following conditions: 1) For the first dimension: 40V (22 hours) in 1X TAE 2.5 μg/mL chloroquine and 2) For the second dimension: 60V (8 hours) in 1X TAE 25 μg/mL chloroquine. The gel was then stained using SYBR Safe before imaging on Typhoon FLA 9500.

### *In vitro* Transcription assay

To test the effect of DNA supercoiling, soft-torsional block by dCas9 and Topoisomerase I in single-round transcription elongation, we performed experiments using radiolabeled RNA. Stalled elongation complexes labeled with radiolabeled nascent RNA were formed by assembling *E.coli* RNA polymerase Holoenzyme (10 pmol, New England Biolabs) in pUC19-T7A1U (1 pmol) TB40 buffer (20 mM Tris pH 8.0, 40 mM KCl, 5 mM MgCl_2_, 0.02 mg/mL BSA, and 1 mM DTT) in the presence of 10 μM of ATP, GTP, CTP (Thermo Scientific, R0481) and 50 μM of GpA dinucleotide (TriLink Biotechnologies) and α-P32-ATP (Perkin Elmer). The reactions were incubated for 20 min at 37 °C and then placed on ice. The excess of radiolabeled nucleotides was removed using a size-exclusion column Microspin G-25 (GE Healthcare). The stalled complexes were reinitiated by adding NTPs at 10 μM and reaction allowed to proceed at room temperature. Time points were quenched using 2X RNA formamide loading dye and taken at 0, 20, 40, 60, 80, 100, 120, 180, 240, 300, 450, 600, 750, 900, 1050, 1200, 1350, 1500, 1650, 1800 s. Quenched samples were ran in 10% denaturing urea gel 19:1 acrylamide/bisacrylamide, the resolved RNA were imaged on Typhoon FLA 9500 and the bands were quantified with ImageQuant TL 8.2.

The reactions were performed using nicked and supercoiled DNA templates of single promoter pUC19-T7A1U. For conditions where dCas9 was used, dCas9:sgRNA targeting TTS (CBR01) were incubated at 10 min after addition of *E. coli* RNAP Holoenzyme. When Topoisomerase I was used, *E. coli* Topoisomerase I was added 10 min after addition of *E coli* RNAP and allowed to relax DNA before isolating stalled elongation complexes. The abortive initiation measurements in **Figure 2D** were based on autoradiographed RNA intensities at time point 0 for both nicked and –sc pUC19-T7A1U templates. As can be seen from these experiments, more efficient loading and stalling of RNAP was observed on −sc DNA templates than nicked after removal of the excess radiolabeled nucleotides (i.e.: higher absolute amount was observed at 19 nt stalled site for the −sc condition than the nicked condition). To account for this, gel intensities were scaled accordingly in **Figure 3E**, and lane-normalization was used for quantification accordingly (See below in Gel quantification).

### Gel quantification

The gel images were initially adjusted using the GIMP cage transformation function to correct bending along the horizontal direction, facilitating subsequent auto-processing steps. The quantification of gel intensity was facilitated by using the Python script gel_lane_finder script,^78^ which enable the automated annotation of gel lanes. The two-dimensional band was averaged in the horizontal direction, resulting in a one-dimensional array representing the pixel-wise intensity along the annotated gel lane. To standardize the intensity values, which may be influenced by variations in sample loading for each lane, the sum of pixel intensity in the one-dimensional array within each lane was employed to normalize the intensity at each pixel point. Specific representative pausing positions were traced across the temporal evolution of the transcription. For supercoiling relaxation experiments, the presence of a small fraction of pre-existing nicked species in the negatively supercoiled DNA template (i.e., the top band at time zero, ΔLk = 0) may affect the interpretation of the results. Therefore, the ΔLk = 0 lane was excluded from the analysis.

### Real-time transcription assays using a molecular beacon

To measure RNA synthesis in real time for both AmpR and mEGFP gene from the opposing and tandem constructs, we adapted a fluorescence-based methodology using molecular beacons that possess a 2′-*O*-methylribonucleotide backbone.^63^ Molecular beacons CBMB01 and CBMB02 targeting 3’ RNA ends of AmpR and mEGFP transcripts were used in separate reactions. Briefly, each reaction was performed in TB40 buffer (20 mM Tris pH 8.0, 40 mM KCl, 5 mM MgCl_2_, 0.02 mg/mL BSA, and 1 mM DTT) in the presence of 1 mM of ATP, GTP, CTP, UTP (Thermo Scientific, R0481), 50 μM of GpA dinucleotide (TriLink Biotechnologies), 8 nM of plasmid, 85 nM of *E.coli* RNAP Holoenzyme (NEB), and 500 nM of molecular beacon. To ensure that all of the transcription reactions started simultaneously, the reaction mixtures were prepared on an ice-cold metal block. Then, all transcription reactions were incubated simultaneously in an iQ5 Thermal Cycler for 2 hour at 37°C. During the reaction, the fluorescence intensity FAM of the molecular beacon was recorded every 20 s.

### Measurement of GFP fluorescence emission in *E. coli* MG1655 with tandem and opposing constructs

The following experiments were performed using similar experimental conditions as reported by Kim et al.^24^ Briefly, plasmids for the tandem and opposing constructs (pUC19-T7A1-AmpR-mEGFP-Tandem and pUC19-T7A1-AmpR-mEGFP-Tandem) were transformed into *E.coli* MG1655 cells using standard electroporation protocols. Positive clones were selected by their growth on an ampicillin antibiotic plate and by their fluorescence emission in a trans-illuminator under blue light. Three colonies from each construct were selected, mixed and a ∼ 5 mL inoculate grown in M9GluCAAT media containing M9 minimal medium (6 g/L Na_2_HPO_4_, 3 g/L KH_2_PO_4_, 0.5 g/L NaCl, 1 g/L NH_4_Cl, 2 mM MgSO_4_, 0.1 mM CaCl_2_) supplemented with 0.2% glucose, 0.1% casamino acids (Difco Laboratories),1 mg/L thiamine and ampicillin 50 μg/mL at 37 °C. The overnight inoculate was then diluted 10,000 fold and then grown to exponential phase in M9GluCAAT with ampicillin 50 μg/mL to an optical density OD_600_=0.2. Then, ∼0.5 μL of these cells were spotted into an agarose pad made with 1% agar and M9GluCAAT covered with a no. 1.5 coverslip, then imaged in the microscope at room temperature. Imaging by phase contrast and fluorescence microscopy was performed on a Nikon Eclipse Ti microscope equipped with either a 1.40 NA phase-contrast oil objective, a Hamamatsu Orca-Flash4.0 V2 CMOS camera, a Sola Light Engine (Lumencor) and pE-4000 (CoolLED) light source. For GFP excitation/emission the ET470/40x Chroma dichroic was used. The microscopes were controlled by the Nikon Elements software.

### Atomic Force Microscopy

#### Slow scan Atomic Force Microscopy (AFM)

DNA samples of different topologies of pUC19-T7A1U (1-2 nM) were diluted in 10 mM HEPES pH 7.0 and 5 mM MgCl_2_ prior to sample deposition. Two microliters of DNA were deposited on freshly cleaved mica and incubated at room temperature for 2 or 3 minutes. Mica was rinsed with 50 μL of water five times and dried under N_2_ gas flow. AFM measurements were performed with a Multimode AFM Nanoscope 8 (Bruker Co.). The samples were imaged in tapping mode; the silicon cantilevers (Nanosensors) were excited at their resonance frequency (280–350 kHz) with free amplitudes of 2-10 nm. The image amplitude (set point As) and free amplitude (A0) ratio (As/A0) was kept at 0.8. All samples were imaged at room temperature in air, at a relative humidity of 30%. Raw static AFM images acquired in air were flattened and leveled using Gwyddion 2.5. A mask of the entire DNA molecule using thresholding was obtained and only pixels corresponding to DNA regions shown in **Figure 1C**.

### Cryo-ET sample preparation for transcriptional modulation of DNA supercoiling

#### Preparation of circular plasmid under low salt and high salt conditions

For the preliminary investigation into the conformational dynamics of DNA supercoiling in response to varying ionic strength, the above extracted and purified pUC19-T7A1U plasmid was diluted to a concentration of 100 nM under two distinct buffer conditions: low salt and high salt buffer (20 mM Tris-Cl, 0.5 mM DTT, and 2% Trehalose; 20 mM Tris-Cl, 40 mM KCl, 5 mM MgCl_2_, 0.5 mM DTT, and 2% Trehalose, respectively) preceding the application onto the EM grids.

#### Formation of RNAP-pUC19-T7A1U stalled complex

To investigate the localization of RNAP on the plasmid, a 10 μL incubation reaction was prepared to induce RNAP stalling 19 nucleotides downstream from the TSS of the T7A1 promoter. This was achieved by employing UTP starvation in a system composed of 60 nM pUC19-T7A1U plasmid, 180 nM RNAP (3:1 ratio to the plasmid), 50 μM GpA dinucleotide primer (facilitating RNAP initiation), and 10 μM rNTP mix (excluding UTP). The reaction buffer consisted of 20 mM Tris-Cl, 40 mM KCl, 5 mM MgCl_2_, 0.5 mM DTT, and 2% Trehalose. This reaction was then allowed to incubate for 20 minutes at 37°C to create the RNAP stall complex before application onto the TEM grids.

#### Formation of dCas9-pUC19-T7A1U complex

Following the same incubation protocol as described above, the RNAP in the system was replaced by dCas9, an alternative DNA-binding protein. To do so, the dCas9 protein and its guiding RNA (sgRNA) were first assembled in a separate vial at a 1:1 ratio, reaching a concentration of 500 nM. This assembly was achieved by incubating the dCas9-sgRNA mixture in a buffer containing 20 mM Tris-Cl, 40 mM KCl, 5 mM MgCl_2_, and 0.5 mM DTT for 10 minutes at room temperature. Subsequently, the incubated dCas9-sgRNA complex was introduced into the plasmid, yielding a system containing 60 nM pUC19-T7A1U, 120 nM dCas9-sgRNA (2:1 ratio to the plasmid), a GpA dinucleotide primer (50 μM) for initiation, and an rNTP mix (10 μM, excluding UTP). The reaction buffer composition remained consistent with 20 mM Tris-Cl, 40 mM KCl, 5 mM MgCl_2_, 0.5 mM DTT, and 2% Trehalose. This reaction mixture was further incubated for 20 minutes at 37°C before being applied to the TEM grids for further analysis.

#### Formation of transcribed (none-equilibrium) RNAP-pUC19-T7A1U complex

To study the activation of RNAP transcription on the negatively supercoiled plasmid, the RNAP stalled complex was first prepared as described above. At the end of its 20-minute incubation at 37°C, rNTP (including UTP) was added to the solution, adjusting the concentration of each nucleotide to reach 100 μM. The transcription reaction continued for 10 minutes at room temperature before being transferred onto ice prior to application onto the TEM grids. For single-particle analysis, the transcription elongation complex concentration (both RNAP and plasmid) was scaled up 4-fold without changing the buffer conditions.

#### Formation of stalled and transcribed RNAP-dCas9-pUC19-T7A1U complexes

To study active RNAP transcription on the negatively supercoiled plasmid in the presence of dCas9 torsional block, the previous dCas9-pUC19-T7A1U complex was first prepared. Subsequently, RNAP was introduced into the incubated solution at a 3:1:1 ratio to dCas9 and plasmid, with concentrations of 180 nM, 60 nM, and 60 nM, respectively, maintained under the same physiological salt condition. In this sequence, the addition of rNTP (10 μM final concentration, excluding UTP) allowed the accumulation of mild torsion as RNAP initiated transcription from the TSS to the U-less stalling site, spanning 19 nucleotides. To induce additional torsional stress in the system, rNTP (including UTP) was added to the stalled RNAP-dCas9-pUC19-T7A1U complex at 100 μM final concentration. The sample was subjected to a 10 minutes incubation at room temperature for DNA transcription before being transferred to the EM grids.

#### Formation of transcribed RNAP-topI-pUC19-T7A1U complexes

To relieve RNAP from its apical constraint, topoisomerase I was introduced into the transcription system. The stalled RNAP-pUC19-T7A1U complex was initially prepared following the same established protocol. After a 20-minute incubation at 37°C, topoisomerase I was added to the solution at a ratio of 3:1:1 with respect to RNAP and plasmid at concentrations of 180 nM, 60 nM, and 60 nM, respectively, under the same physiological salt condition. Subsequently, rNTP (including UTP) was added to the solution mixture, reaching a final concentration of 100 μM, and the transcription persisted for 10 minutes at room temperature before application onto the EM grids.

#### Formation of transcribed RNAP-pUC19-T7A1U dual-promoter complexes

To study RNAP transcription activation on negatively supercoiled plasmids containing dual promoters (i.e., opposing and tandem constructs), RNAP-stalled complexes were prepared using the same procedure as described above for the single-promoter pUC19-T7A1U template, with the exception that the RNAP concentration was doubled to 360□nM (corresponding to a 6:1 molar ratio of RNAP to plasmid) to accommodate dual-promoter initiation. Following a 20-minute incubation at 37□°C, rNTPs (including UTP) were added to a final concentration of 100□μM for each nucleotide. The transcription reaction was allowed to proceed for 10 minutes at room temperature and then placed on ice prior to application onto TEM grids.

### TEM specimen preparation

3 μL of a sample after the above incubation was deposited onto a glow-discharged (PELCO easiGlow™ Glow Discharge Cleaning System) 200 mesh Quantifoil gold grid (hole size ranging from 1 micron to 2 microns, Electron Microscopy Sciences) for 30 seconds. Following a 20-second on-grid incubation, the grid was plunge-frozen in liquid ethane at ∼95% humidity and 15°C using a Leica EM GP rapid-plunging device (Leica, Buffalo Grove, IL, USA) after controlled blotting with filter paper (3-5 s). The resulting flash-frozen grids were then transferred into liquid nitrogen for storage.

### TEM data acquisition

The Cryo-EM data were acquired using a Titan Krios (FEI) transmission electron microscope operating at 300 kV high tension and equipped with a Gatan energy filter. Imaging was performed with a Gatan K3 Summit direct electron detection camera, employing a magnification of x53,000 (where each pixel of the micrographs corresponds to 1.67 Å in specimens) in super-resolution and correlated double sampling (CDS) mode. Tilt series of the samples were captured from -51° to +51° at 3° step or -55° to +55° at 5° step increments using a dose symmetry scheme. Data collection was automated using SerialEM software^79^ to track the specimen and maintain a defocus of ∼2.5 μm. The total dose for the tilt image series ranged from ∼110-150 e⁻/Å^2^. At each tilt angle, a total of 8 frames were collected, with an exposure time of 0.25 s per frame. Single-particle movie data were collected using a Gatan K3 direct electron detector at a nominal magnification of ×81,000, corresponding to a pixel size of 1.05 Å. Each exposure lasted 7.39 seconds and was recorded over 50 frames, with a total electron dose of 50 e⁻/Å². Micrographs were acquired in non-CDS and super-resolution modes, with a defocus range of 0.6–1.6 μm, using SerialEM.

### Cryo-ET tilt series processing and 3D reconstruction

The beam-induced motion of cryo-EM frames was corrected using MotionCor2.^80^ To improve the contrast of low-dose cryo-ET tilt series images, a deep learning-based denoising method (NOISE2NOISE^81^) was implemented, involving the division of motion-corrected movie frames into even and odd halves for network training and subsequent particle resolution estimation (**Supplementary Data 2, I**). Predictions for even and odd frames for a single tilt were averaged together. The defocus value of the cryo-ET tilt series was determined using GCTF.^82^ A carbon area perpendicular to the tilt axis was included during data collection to assist in contrast transfer function (CTF) detection and later tilt series alignment. The tilt series were initially aligned using IMOD with patch tracking function^83^ and subsequently imported into e2tomo software^28^ for 3D reconstruction (**Supplementary Data 2, II**). Approximately 100 subtomogram patches containing DNA features were cropped for manual annotation and subsequently submitted for training. This convolutional neural network (CNN) was then applied across all tomograms for DNA identification (**Supplementary Data 2, III**).

### Individual particle 3D reconstruction refinement and modeling

After the deep learning-based DNA annotation, individual plasmid particles were manually labeled in the tomogram (**Supplementary Data 1, III, right**). In brief, the DNA-annotated tomograms were binned by 4 (6.68 Apix) and divided into isolated small surface pieces within Chimera.^84^ Density segments from a single plasmid particle were then manually selected and grouped, followed by low-pass filtering to a 6 nm resolution to serve as the particle-shape mask. The criteria for selecting plasmid particles from the annotated tomogram included: 1) the particle should display supercoiling, manifesting as a helical structure (cyan color) without nicking (red color) (**Supplementary Data 1, III, right**), 2) plasmid particles should be isolated and not significantly entangled with others; and 3) particles not partially attached to the carbon area. The center of mass of the particle shape mask was calculated (**Supplementary Data 1, IV, left**), and this information was used to crop 1400x1400 pixel local tilt series within the raw large micrograph before denoising. This cropped tilt series, with a plasmid particle in the center, was binned by 7 (a larger crop area and bin=8 were used for the later opposing and tandem construct plasmid particles for their larger size) and then submitted for local 3D reconstruct with EMAN2.^28^ To minimize artifacts caused by the limited tilt angle range, the 3D reconstructed maps were missing-wedge compensated using IsoNet software^29^ (**Supplementary Data 1, V**), employing the previously created low-resolution mask. The output map was then low-pass filtered to 2 nm, serving as the final map for the modeling.

Plasmid modeling was facilitated through the following steps: from the 3D reconstructed maps of the plasmid particle, a spine line representing the particle’s super-helical axis was initially computed (**Supplementary Data 2, first row, left**). This involved applying a strong Gaussian kernel to the map, followed by skeletonization using the ‘lee’ method from the scikit-image package. Along the determined spine line, a sampling cylinder with dimensions of 60 nm diameter and 15 nm height was generated, utilizing a 5 nm step. The DNA density within the cylinder was divided into two regions, and the weight centers for each region were recorded (**Supplementary Data 2, first row, right**). Subsequently, the tracing centers were interconnected based on rotation and distance to the previous centers, with a subsequent round of manual correction undertaken to address misconnection cases, particularly in low-quality areas of the EM map (**Supplementary Data 2, second row, left**). The resulting threaded points underwent final processing through smoothing and interpolation with a 2 nm spatial interval, yielding the final model for the 3D map of the supercoiled DNA (**Supplementary Data 2, second row, right**).

### Evaluation of the cryo-ET 3D reconstruction resolution

The resolution for the individual particle reconstructions were estimated by two methods. 1) Data-to-Data based analysis: the Fourier Shell Correlation (FSC) was calculated between two independently reconstructed 3D maps, in which each map was based on one-half of the tilt-series (split by even and odd frames for each tilt). The frequencies at which the FSC curve first falls to values of 0.143 were used to represent the reconstruction resolution. 2) Data-to-Model based analysis: the FSC curve between the final 3D reconstruction and the density map converted from the corresponding fitting model was calculated. The frequencies at which the FSC curve fell below 0.5 was used to estimate the resolution. The density map of the fitting model was generated by pdb2mrc in EMAN software.^85^

### Sub-tomogram averaging of RNAP and dCas9

The sub-tomogram averaging of RNAP and dCas9 particles was carried out utilizing the EMAN2 e2tomo. Briefly, the CTF-corrected and imod-aligned raw tilt series were imported into the software, followed by e2tomo 3D reconstruction, yielding a 4×-binned tomogram (equivalent to 6.68 Å with an unbinned pixel size of 1.67 Å). Subsequently, approximately 100 particles for each protein type were manually selected, and ab-initio maps were reconstructed for the subsequent round of reference-based boxing. Employing a template matching threshold value (vthr) of 7.5, a total of 1781 and 875 particles were extracted for RNAP and dCas9, respectively. Particle cropping, alignment, and averaging procedures were executed using the e2spt_refine_new.py script. No symmetry was applied during particle alignment. Following 8 rounds of alignment (iters=p,p,p,t,p,p,t,r), resolutions of 14.1 Å and 17.6 Å were determined for RNAP and dCas9, respectively (0.143 cut-off), employing Fourier Shell Correlation (FSC) with odd and even particles from the masked, final average. By mapping the orientation-determined particles back to their tomogram using e2spt_mapptclstotomo.py, DNA-bound particles were selectively chosen if their center of mass was within a distance of <1 nm to any points of the plasmid model. A subset of 232 RNAP and 116 dCas9 particles was then subjected to another round refinement (iters=p,p,p,t,p), resulting in resolutions of 17.7 Å and 18.9 Å, respectively, evaluated under the same standards.

### Single-particle analysis of TECs on negatively supercoiled DNA template

Single-particle 3D reconstruction was performed using cryoSPARC.^86^ The detailed data processing workflow for TECs on negatively supercoiled DNA templates is shown in **Figure S4A**. A total of 10,476 micrographs underwent patch motion correction, patch CTF estimation, and particle picking, yielding 2,063,152 particles. Initial 2D classification and ab initio reconstruction revealed moderate orientation bias of RNAP particles. To preserve rare views, particle cleanup was performed via three rounds of heterogeneous refinement against 6 models—5 representing noisy junk references and one low-pass-filtered good model from the initial 3D reconstruction—instead of standard 2D classification. This cleanup yielded 611,747 particles (binned 3, in a 108-pixel box), which were used for homogeneous refinement, achieving a 6.5 Å reconstruction. Subsequent 3D classification at 15 Å resolution separated two major classes: TIC (186,816 particles, with σ-factor bound) and TEC (424,931 particles, without σ). The TEC particles were re-boxed (unbinned, in a 324-pixel box) and subjected to ab initio reconstruction, non-uniform refinement, and orientation rebalancing using cryoSPARC’s default settings. Further 3D classification at 10 Å resolution of TEC particles yielded three classes: Class I (TEC1–SC) and Class II (TEC2–SC), both showing DNA in the RNAP active site and a downstream DNA density protrusion, and Class III, which showed only faint DNA density and was not analyzed further. Non-uniform and local refinement of TEC1–SC (46,743 particles) and TEC2–SC (54,869 particles) yielded final structures at 2.9 Å resolution. Maps were post-processed with DeepEMhancer.^87^ Initial models were built by fitting domains from known TEC structures (e.g., PDB: 6ALH) into the density maps using ChimeraX.^88^ Models were iteratively refined using phenix.real_space_refine,^89^ manually adjusted in ISOLDE,^90^ and validated with MolProbity^91^ (**Supplementary Data 4**). The swivel angle calculation for TEC1–SC and TEC2–SC followed the definition of the core and swivel modules as described by Kang et al.,^52^ and using PDB structure 6RH3 as the reference. Additional structures (1:6RH3, 2:8EG8,4:6ALH, 5:8EHI, 7:8EHF, and 8:7PYK) were also included in the measurement as control. Measurements were performed in ChimeraX, and the ChimeraX script is available in the Zenodo repository.

### Plectoneme apical site sequence-based prediction

Prediction of DNA plectoneme formation loci along the pUC19-T7A1U sequence was performed using a computational model developed by Kim et al.^43^ To account for the circular topology of the plasmid while matching the required linear DNA input format, the pUC19-T7A1U sequence was duplicated three times and joined head-to-tail to simulate continuity; only the central region corresponding to the original plasmid was plotted to eliminate edge artifacts. To provide a reference for interpreting the plectoneme formation scores, a control sequence consisting of polyA–Widom 601–polyA was used. This construct includes the well-characterized Widom 601 nucleosome positioning element, flanked by polyadenine tracts, and is known for its high intrinsic bendability, which provides a comparative baseline for sequence-driven plectoneme localization.

### Quantification of supercoiled DNA plasmid morphology

The quantification of the supercoiled DNA plasmid’s morphology involved measurements of radius of gyration, global writhe number, and plectoneme branch writhe density. To quantify these characteristics, the eigenvalues of the gyration tensor were calculated and designated as r1, r2, and r3, with the relationship r1 > r2 > r3. The radius of gyration (R) was determined by the formula R = r1^2^ + r2^2^ + r3^2^. The calculation of the writhe number was facilitated by the E-CAM polymer_data_analysis module, which implements a double integral computation method proposed by Klenin & Langowski.^92^

### Determination of the curvature and apexes along the supercoiled DNA

To determine the curvature of the plasmid along its DNA trajectory, a loop iterated from index 1 to N-1 for each point on the DNA plasmid model. The curvature (κ) at each point (i) was computed with its two neighboring points (i-1 and i+1) using the formula κ = 1/R = 4S/fgh,^93^ where R is the radius of the circle circumscribing a triangle formed by the three points. Here, S represents the area of the triangle, and f, g, and h are the side lengths opposite the vertices of the triangle. The curvature at each point along the DNA trajectory was color-coded using a gradient, with points exhibiting the largest curvature values near the distal ends of each plectoneme branch utilized as the apexes of the plasmid.

### Calculation of the distance between DNA along the plasmid’s helical axis

To calculate the distance between DNA along the supercoiling helical axis, the points on the DNA plasmid model were sequentially sorted and segmented into different color groups using the apical points on the plasmid (**Figure S1E**). For each point (i) in a group, its nearest neighboring point in another color group was determined, and the center of mass as well as the distance between the two points were calculated. This calculation was repeated from point 1 to N for each segment, resulting in a list of centers and distances that were used to represent writhe helix axis and the distances between the DNA along the supercoiling helix (**Figure S3F, II**). Given the difficulty of defining the start and end site for a circular plasmid, the apexes were utilized as the zero index to align the calculated distance list.

### Determination of the local writhe density of the plasmid

Utilizing the previously calculated writhe helical axis trace of the model (**Figure S3F, II**), the branches of the plasmid can be determined and split at the junction (**Figure S3F, I**). Each helical axis branch was color-coded, and these color codes were used to segment the points on the plasmid model based on their vicinity (**Figure S3F, III**). These branch segmentations were treated as smaller closed plasmids and subjected to writhe number calculation using the previously described method.^92^ In the statistical analysis, to account for the greater contribution of longer plectonemes to writhe density, long DNA plectonemes were segmented into 100 nm units. Plectonemes shorter than 100 nm were left unsegmented and included as individual units.

### Construction of superimposed models and mapping RNAP position on plasmid

To align all models from the same experimental condition, orientation-determined particles within the same group were back-mapped to their respective tomograms. Concurrently, the plasmid model was also back-mapped to the tomogram, so that it allows the determination of the relative orientation between the plasmid and plasmid-bound particles (RNAP and dCas9). Given the current resolution limitations of cryo-ET in resolving DNA sequence information, the position of dCas9 was utilized as a marker to infer the TSS, assuming specificity in dCas9 binding as per the construct design. The entire length of the plasmid trace was normalized to 1950 base pairs, with the point nearest to the dCas9 particle designated as 1090 bp. As none of the particles are symmetrical, the upstream and downstream positions of the bound RNAP were determined relative to the plasmid.

### Coarse-grained simulation of DNA supercoiling relaxation

Coarse-grained molecular dynamics (CGMD) simulations were conducted using the GPU-accelerated, sequence-dependent oxDNA2 model.^42,94^ The simulation environment was set up under the canonical NVT ensemble (constant number of particles, volume, and temperature). To enhance computational efficiency, the 525-bp minicircle DNA system were employed. The in-silico plasmid model was generated using TacoxDNA,^95^ with a DNA twist deficit of -5 and a writhe of 0, yielding a supercoiling density of -0.1. The circular DNA plasmid was energy minimized and relaxed using parameters tailored from oxDNA example simulations (at dna.physics.ox.ac.uk). These parameters included the following (thermostat = john, interaction_type = DNA2, newtonian_steps = 103, diff_coeff = 2.5, salt_concentration = 0.5, T = 300K, dt = 0.005, verlet_skin = 0.05, rcut = 2.0).

To study the impact of varying levels of plectoneme apex constraint on DNA supercoiling structural dynamics, simulations were initiated by introducing a small bulge on the minicircle DNA (**Figure S3A, left**), which facilitated apex formation at the target site during relaxation. The DNA base pairs at the two distal ends of the bulge were subjected to different levels of constraint, implemented as a mutual force trap between the two red sites (**Figure S3A, dashed line box**), with spring stiffness set to 1 and spring equilibrium distances (r0) set to 8.0 nm, 6.4 nm (80%), and 4.8 nm (60%). A fully unconstrained condition was also included. For each condition, the plasmid was first energy minimized for 2 × 10^6^ steps, followed by relaxation for 3 × 10^7^ steps (equivalent to 300 μs), with a frame sampling interval of 5 × 10^4^ steps, yielding 600 frames. The final 500 frame structures after plasmid relaxation and apices formation were used to compute the global writhe distribution and plectoneme apex curvature statistics.

To study the effect of plectoneme apex constraint on DNA supercoiling torsional relaxation, simulations were initiated by relaxing the minicircle for 3x10^6^ steps (30 μs) until the formation of plectonemes and the appearance of apices. The final frame structure was edited to create a nicked plasmid by introducing a single-strand break at base pair index 940 for models I and II, 828 for model III, and 564 for model IV (**Figure S5C**). This nicked configuration was designed to mimic the activity of Topoisomerase I (TopI). Four DNA supercoiling torsional relaxation simulations were conducted: For model I, the nicked plasmid was allowed to relax without any external constraints for 3x10^7^ steps (equivalent to 300 μs). For the model II, III, and IV, a mutual force trap was applied to constrain the sharp apex region, covering approximately 20 base pairs. These traps, functioning as springs, were formed between base pairs at indices 798, 251 on one side and 817 and 232 on the other. The spring stiffness was set to 1, and r0 was defined as the initial distance between the mass centers of each pair of base pairs in the initial structure divided by a factor of 2. The second batch of simulations were also run 3x10^7^ steps. All simulations were repeated two times using the same relaxation parameters mentioned above. Movies were generated using the last run of each simulation at a rate of 1x10^5^ steps per frame.

The equilibrated final frame structure prior to the nick was also utilized to simulate the formation of DNA twin domains. In brief, two rotating harmonic traps were applied to the base pairs at indices 107 and 942, as well as indices 366 and 683, positioned within the intermediate region of the plectoneme. This approach segments plectoneme two sections: the first section contains an apex with constraints, while the second section features a free apex. This setup allows for a comparison of the effects of (+) torsional propagation on one strand, (-) torsional propagation on the other, and their mutual cancellation when the rotating trap is applied. The center of rotation was defined as the center of mass of the selected base pair, with the rotation axis determined as the unit vector extending from the center of the base pair to the center of mass of another base pair located 10 base pairs upstream. The stiffness of the trap was set to 10 in oxDNA, with a rotation rate of 1x10^5^. The confined apex was configured as described earlier. The simulation was run for 3x10^6^ steps and with one repeat. Movies were created with frames captured every 1x10^4^ steps.

## SUPPLEMENTAL INFORMATION

Video S1. 3D reconstruction workflow of a representative negatively supercoiled 2 kbp plasmid particle

Video S2. Coarse-grained MD simulation of negatively supercoiled plasmid relaxation with and without apical constraint

Video S3. Coarse-grained MD simulation of negative DNA supercoiling in managing positive and negative torsion with and without apical constraints

## KEY RESOURCE TABLE

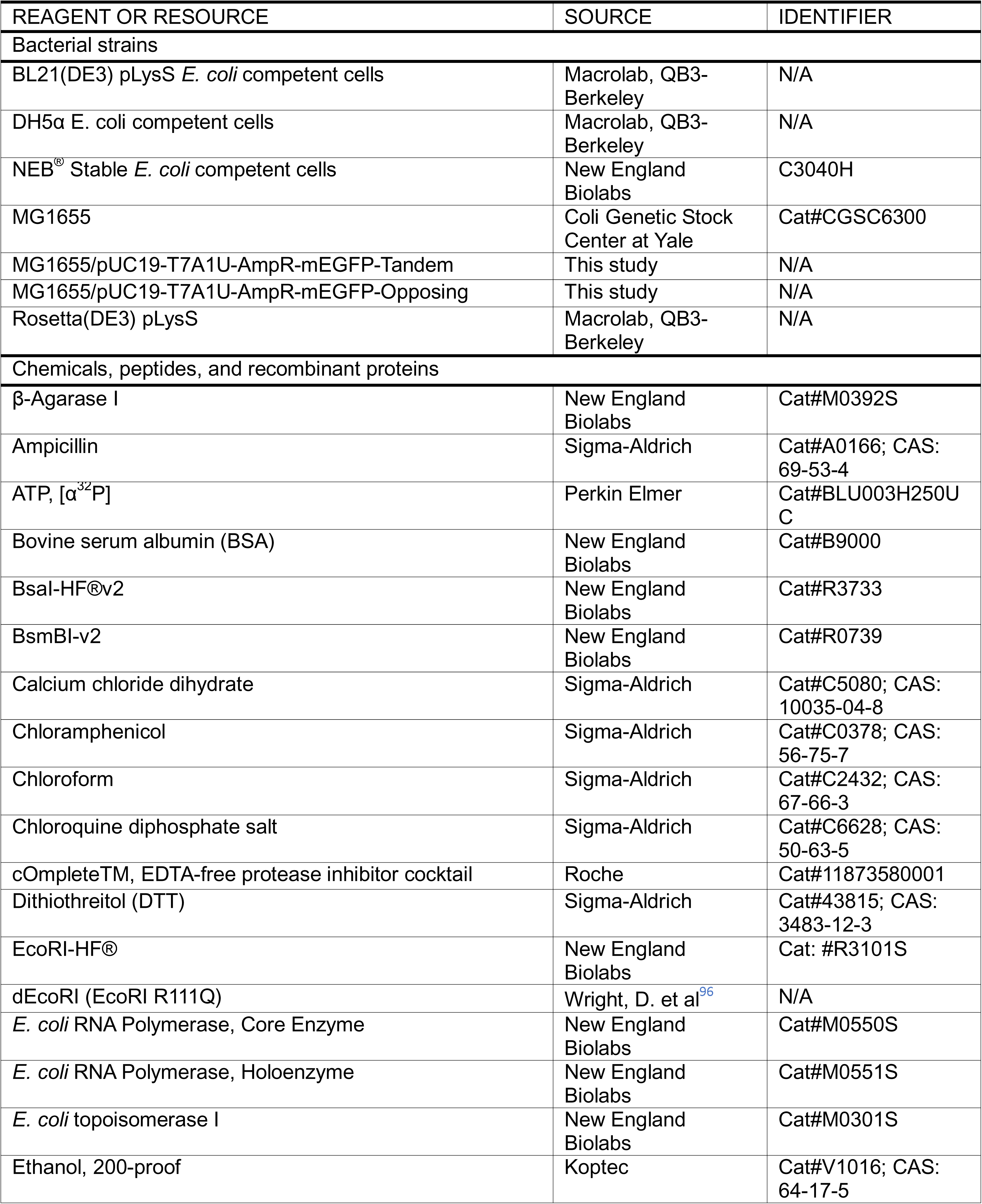
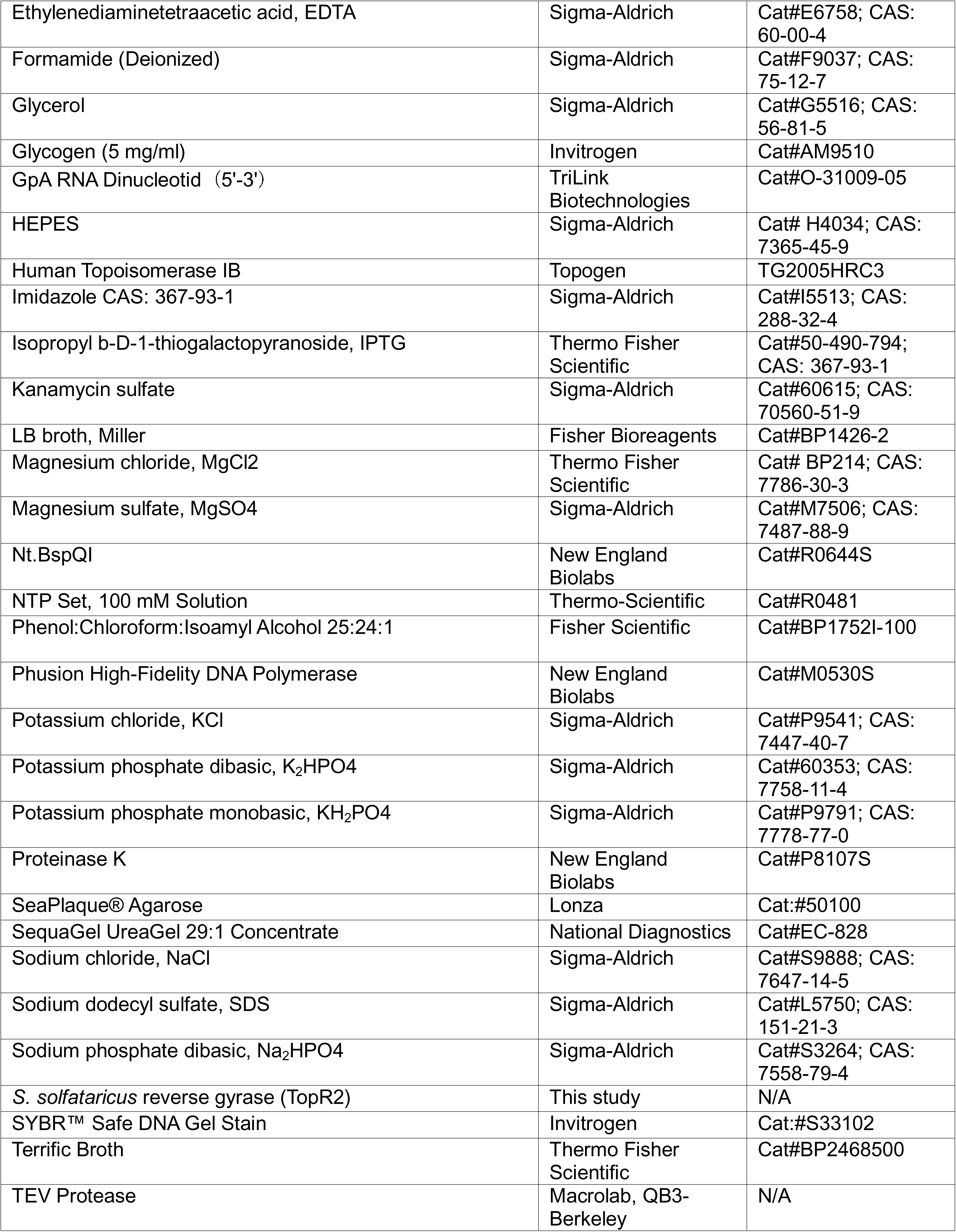
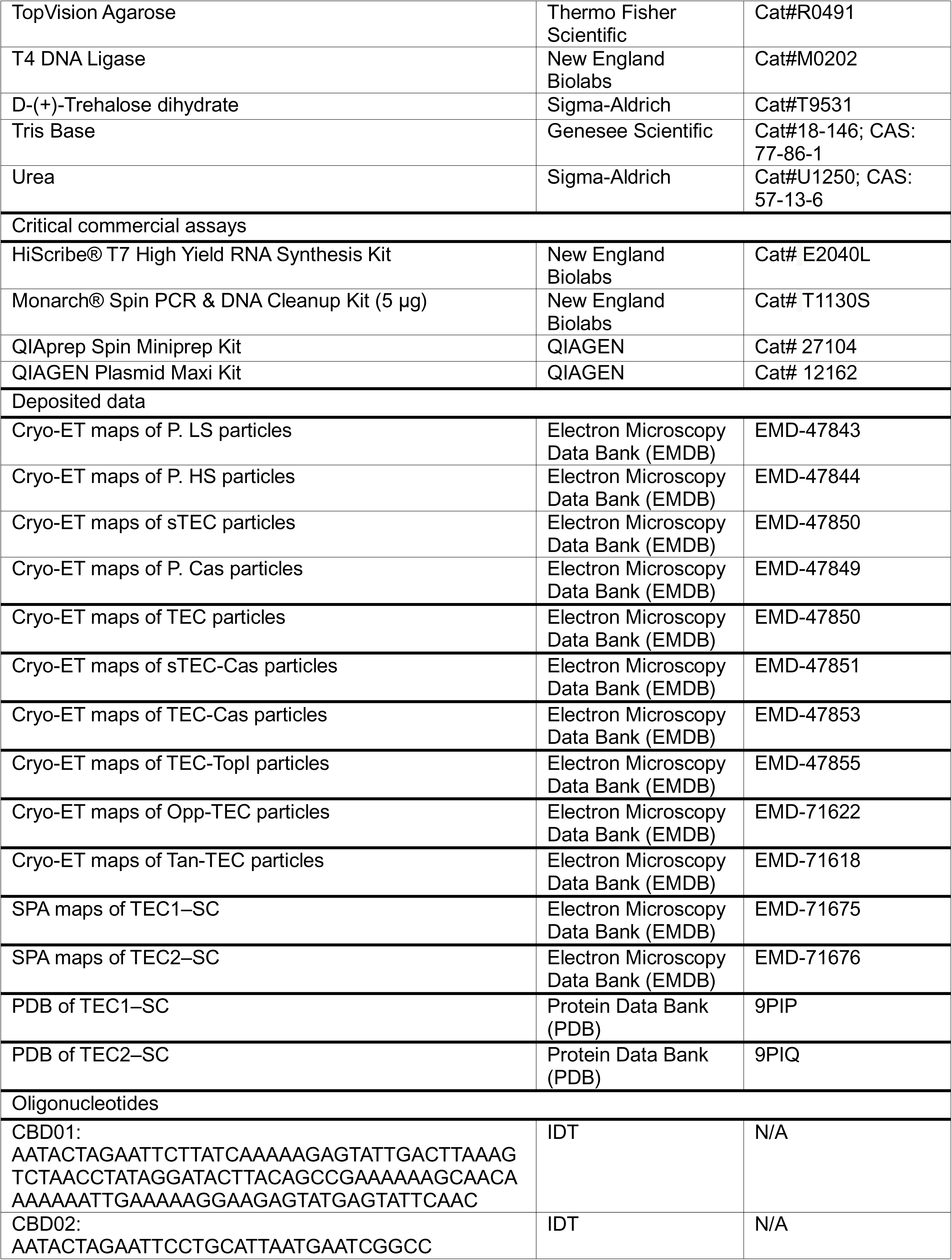
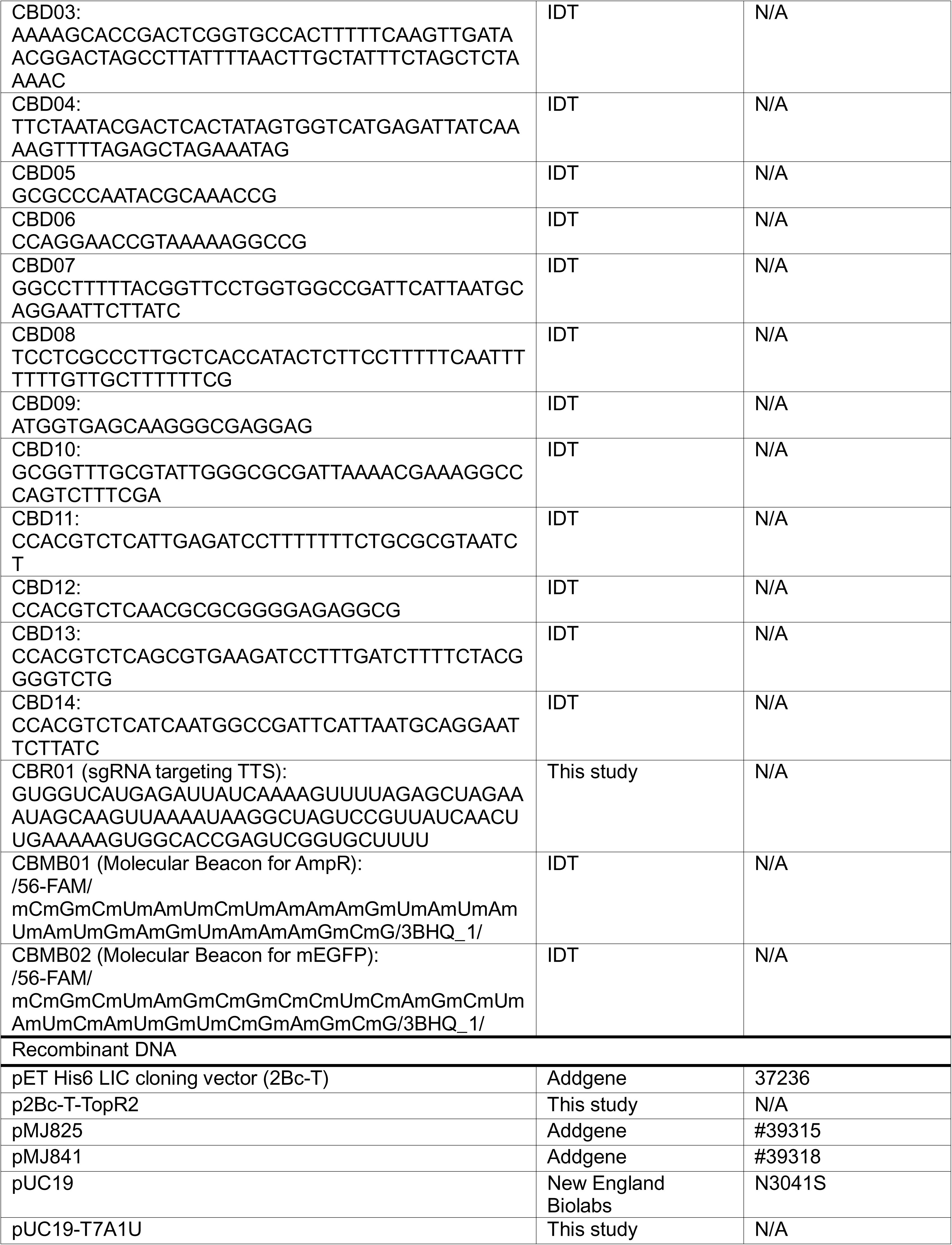
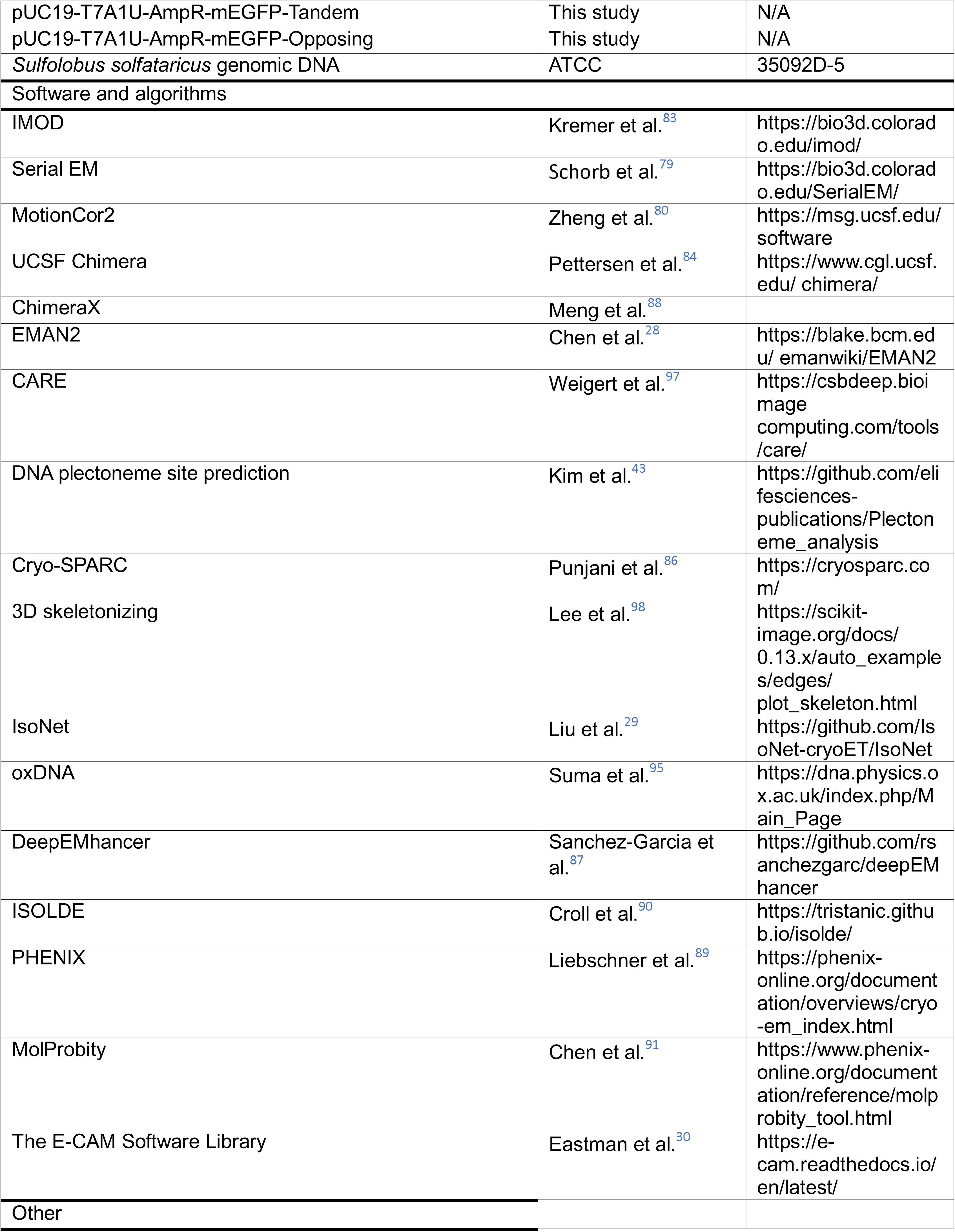
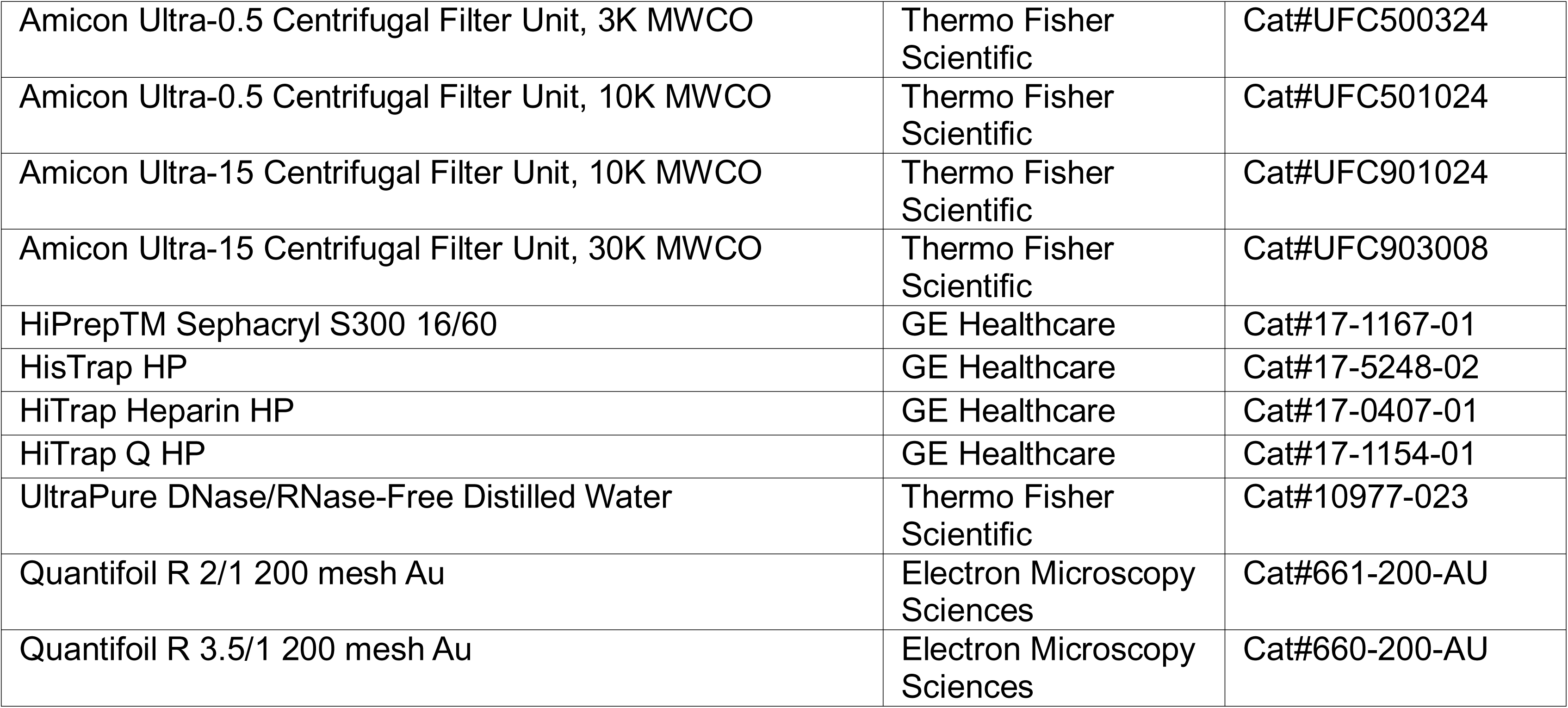

---

**Figure S1:**
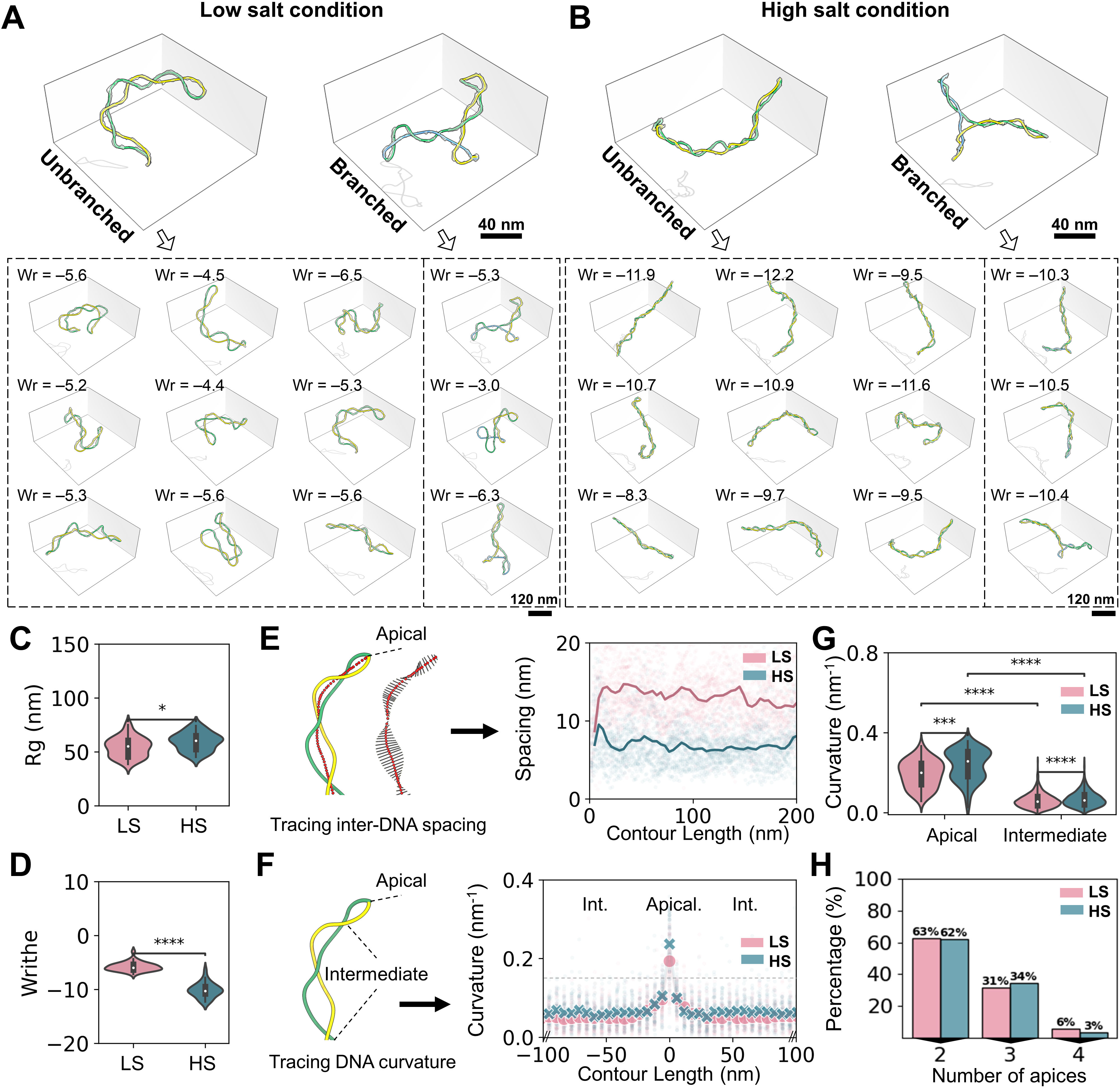
Structural analysis of-sc plasmid under low salt and high salt conditions, related to Figure 1. **(A-B)** Representative cryo-ET map and model of the -sc plasmid (ΔLk centered on -15) under low salt (LS) and high salt (HS) conditions, respectively. Two exemplary particles (unbranched and branch) are displayed on the top panel, with their collection presented on the bottom, featuring writhe numbers. **(C-D)** Statistical analysis of plasmid radius of gyration (Rg) and writhe number using violin plots, respectively; N=64. **(E)** Schematic of inter-DNA spacing quantification (black bars) along the plectoneme axis (red dots). The inter-DNA spacing of the plectoneme was plotted from the plasmid apex toward the intermediate region; N=158 **(F)** Schematic illustrating the apical and intermediate regions of a plectoneme, accompanied by DNA curvature quantification. **(G)** Statistical analysis of DNA curvature near the apical and intermediate regions of the plectoneme under LS and HS conditions; N=158 **(H)** Distribution of the number of apices of the plasmid in LS and HS conditions; N=64. Statistics are calculated using a Mann-Whitney test, where *p < 0.05, **p < 0.01, ***p < 0.001, ****p < 0.0001; and ns, not significant.

**Figure S2:**
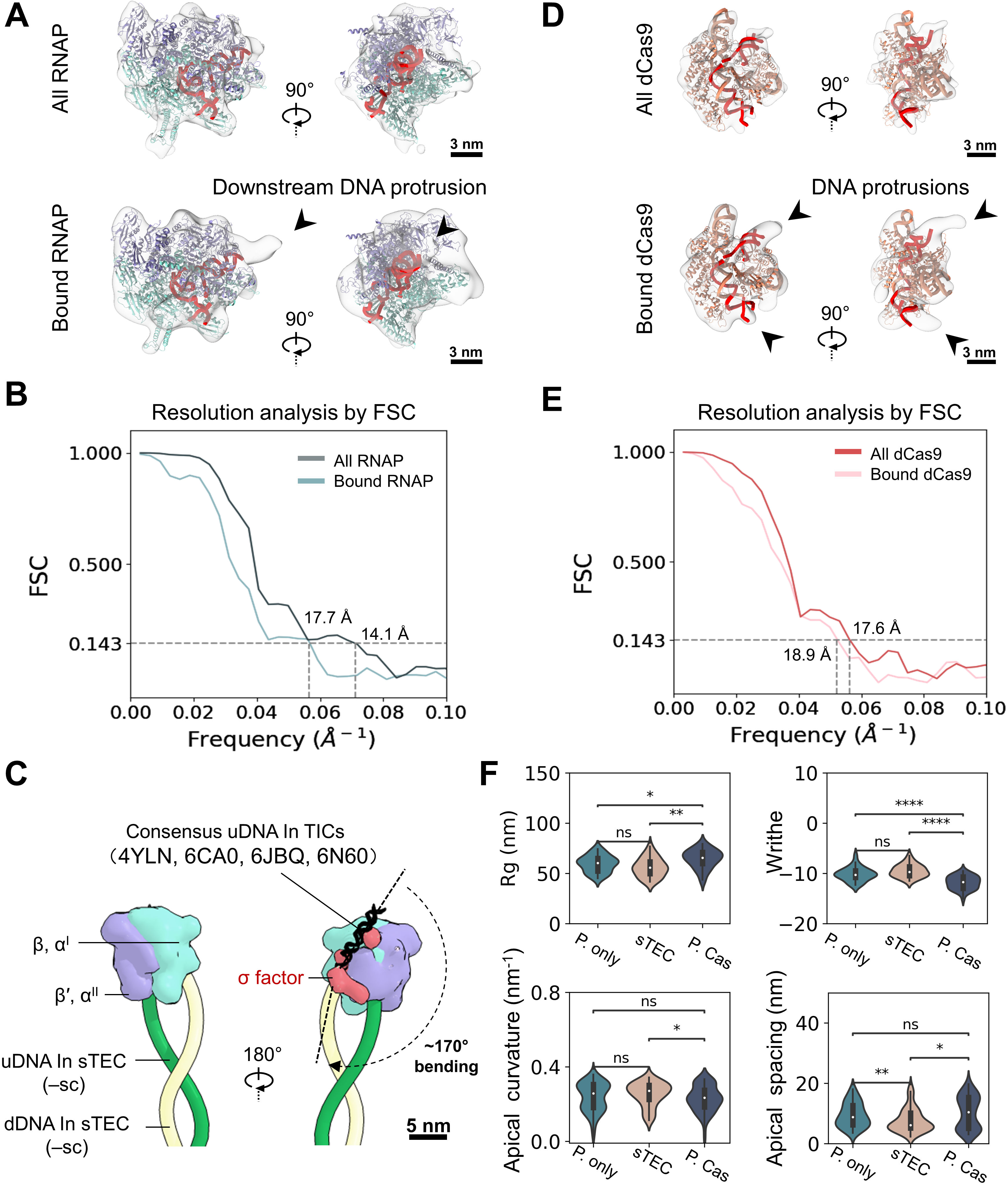
Sub-tomogram averaging analysis of apically bound RNAP and dCas9, related to Figure 2. **(A-B)** Two orthogonal views of the sub-tomogram averaged RNAP map docked with PDB structure 6ALH. Averaging all RNAPs selected from the tomogram (N=1781) yielded a 14.1 Å resolution map (top), while using only plasmid-bound RNAPs after inspection (N=232) resulted in a 17.7 Å resolution map (bottom). **(C)** Superimposition of consensus TIC upstream DNA (black helix) and the TIC o-factor (red) from PDBs 4YLN, 6CA0, 6JBQ, and 6N60 onto the apically stalled TEC on -sc DNA reveals a -170° upstream DNA bend. **(D-E)** Sub-tomogram averaged dCas9 map docked with PDB structure 6O0X. Averaging all dCas9 particles selected from the tomogram (N=875) yielded a 17.6 Å resolution map (top), while the plasmid-bound subset (N=116) achieved a resolution of 18.9 Å (bottom). Resolutions were estimated by measuring the Fourier shell correlation (FSC) between two independently determined half-maps at 0.143. **(F)** Statistical analysis of global plasmid Rg and writhe number (N=90), along with local DNA curvature and inter-DNA spacing at apical sites (N=207), for plasmid only, plasmid with stalled RNAP, and plasmid with dCas9, respectively. Statistics are calculated using a Mann-Whitney test, where *p < 0.05, **p < 0.01, ***p < 0.001, ****p < 0.0001; and ns, not significant.

**Figure S3:**
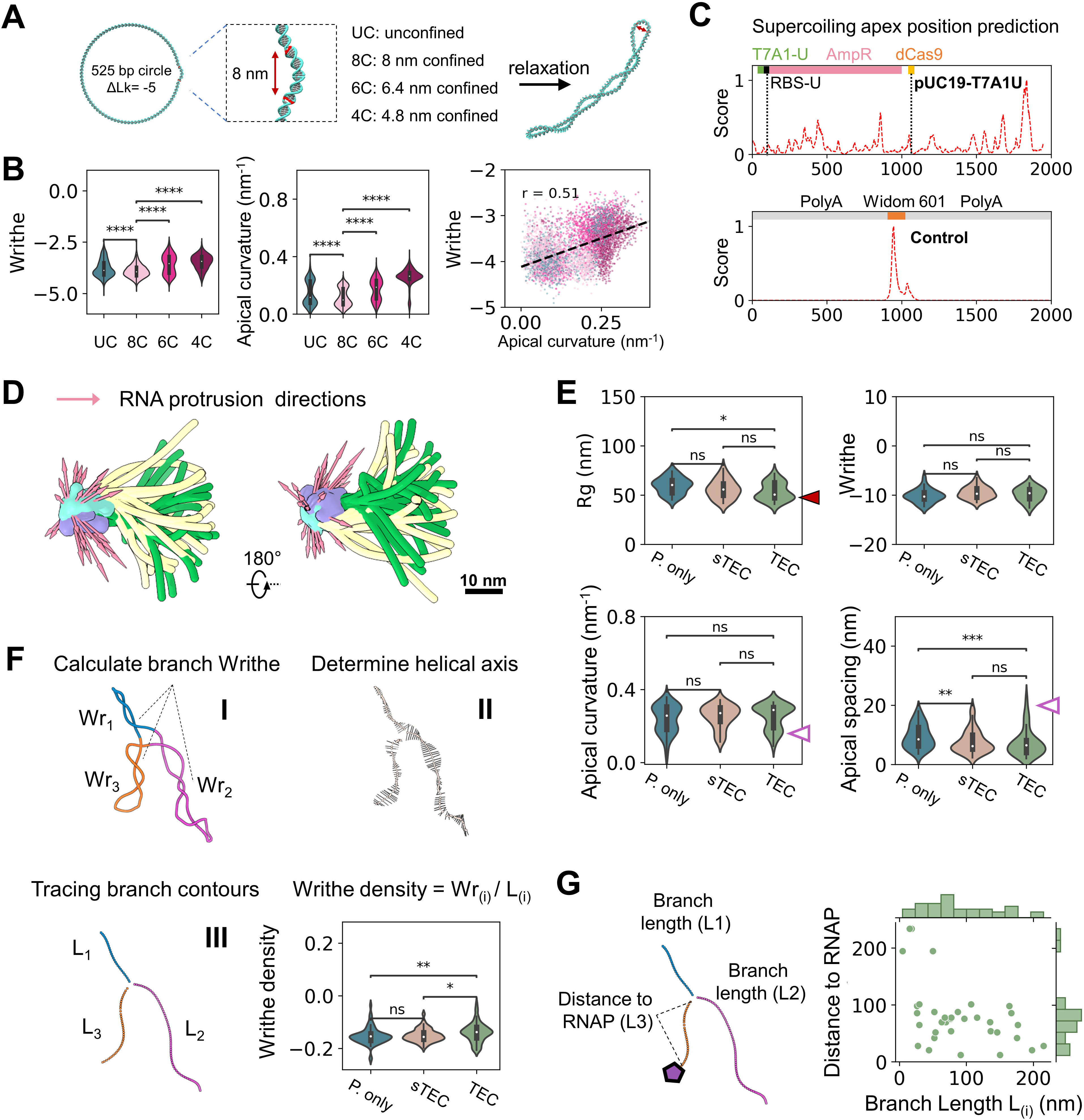
Quantification of plasmid geometry upon RNAP/dCas9 binding, related to Figure 3. **(A)** Generation of -sc DNA minicircles with a target site subjected to varying levels of constraint, defined by spring distances of 8, 6.4, and 4.8 nm, to modulate DNA relaxation. **(B)** Quantification of relaxed plasmid writhe, apical DNA curvature, and their correlation following MD simulation (sampled from frame 100 onward after relaxation; three replicates). **(C)** Predicted DNA plectoneme formation loci along the pUC19-T7A1U sequence. A control was generated using a polyA-Widom601-polyA sequence. **(D)** Overlay of apically bound RNAPs in TECs with RNA protrusion directions indicated by pink arrows. **(E)** Statistical analysis of global plasmid Rg and writhe number (N=79), along with local DNA curvature and inter-DNA spacing at apical sites (N=192) for sTECs and TECs. **(F)** Plectoneme branches were segmented (I), and their superhelical axes (II) and contour lengths (III) were determined. Writhe density was calculated as writhe number divided by segment contour length. Bottom right: writhe density distribution; N=187. **(G)** Plot of plectoneme branch length versus distance from the branch to the apical RNAP; N=38. Statistics are calculated using a Mann-Whitney test, where *p < 0.05, **p < 0.01, ***p < 0.001, ****p < 0.0001; and ns, not significant.

**Figure S4.**
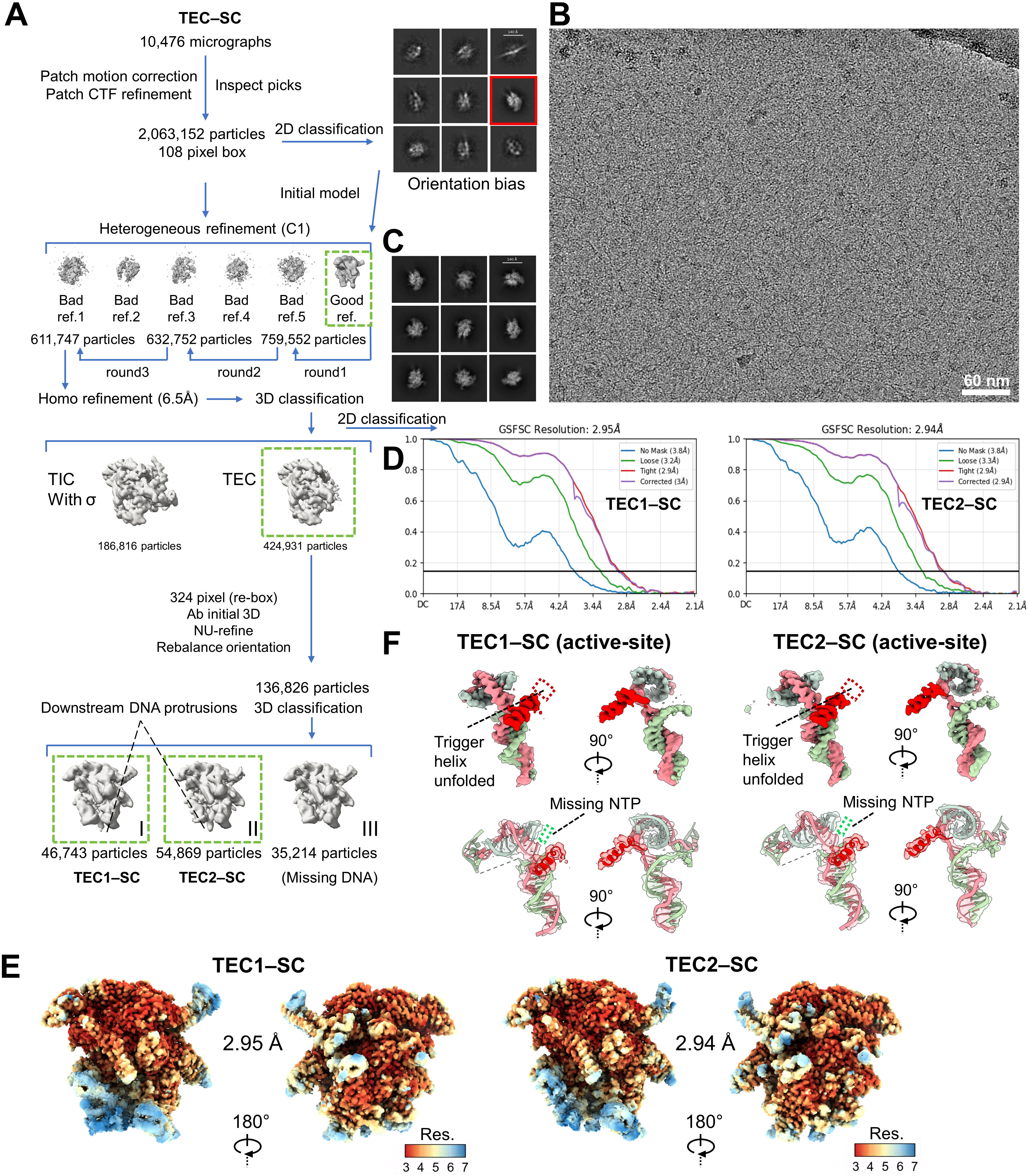
Cryo-EM analysis of the transcription elongation complex (TEC) on a -sc plasmid template, related to Figure 3. **(A)** Workflow of single-particle analysis and 3D reconstruction of TECs. Three final maps were obtained: the first (TEC1-SC, global resolution of 2.95 Å from 46,743 particles) and the second (TEC2-SC, global resolution of 2.94 Å from 54,869 particles) show particles with downstream DNA protrusion. The third class was not further analyzed due to the absence of DNA density, indicating likely unbound RNAPs. **(B)** Representative reference-free 2D class averages of TEC particles, displaying various TECs orientations. **(C)** Representative cryo-EM micrograph of TEC particles assembled on negatively supercoiled pUC19-T7A1U plasmid in vitreous ice. **(D)** Fourier shell correlation (FSC) curves between independently reconstructed even-odd maps. Resolution was estimated at the 0.143 FSC cutoff. **(E)** Cryo-EM density maps of TEC1-SC (left) and TEC2-SC (right), color-coded by local resolution, ranging from 3 Å (red) to yellow to 7 Å (blue). **(F)** Examination of RNAP active sites shows that both TEC1-SC and TEC2-SC are in a post-translocated state, with the trigger helix unfolded and the NTP absent.

**Figure S5:**
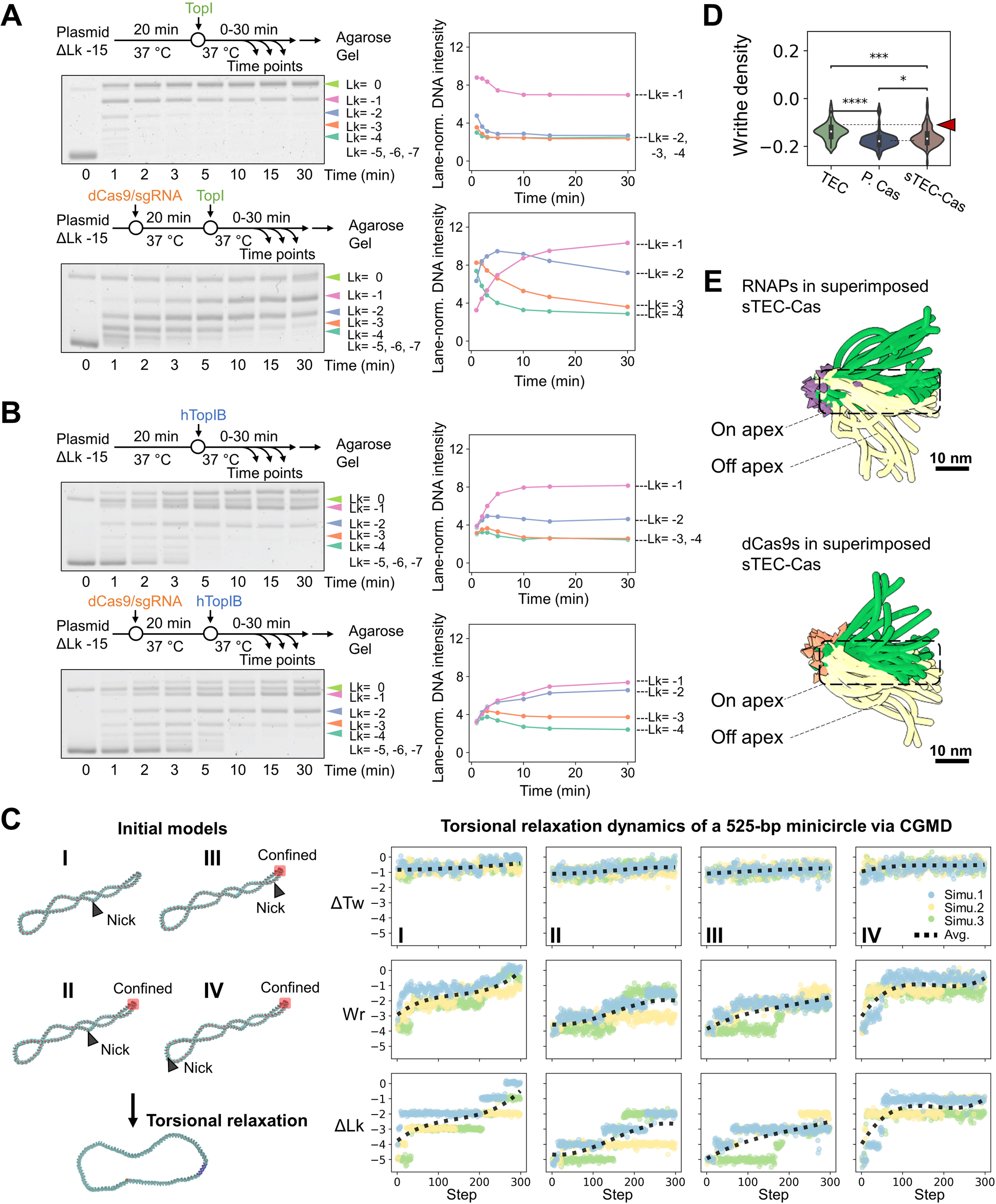
Apical binding of dCas9 hinders DNA rotation, related to Figure 4. **(A)** DNA supercoiling relaxation assay in the absence (top) and presence (bottom) of a dCas9. Right panel: quantification of ΔLk changes over time following Topi addition. **(B)** Control experiment of A, with Topi replaced by hToplB for DNA supercoiling relaxation. **(C)** Coarse-grained MD simulations of 525-bp -sc (ΔLk = -5) minicircle plasmid relaxation dynamics, initiated with different nick locations. Simulations were conducted without apical confinement (Model I) and with apical restriction (Models II—IV). Topological parameters (ΔTw, Wr, and ΔLk) are shown in rows 1-3 for each model. Each simulation was performed in triplicate, with averages indicated by black dashed lines. **(D)** Statistical analysis of plectoneme branch writhe density distribution; N=226. **(E)** Superimposition of apical DNA segments illustrating the dynamics of stalled RNAP and dCas9 when co-present on the plasmid.

**Figure S6:**
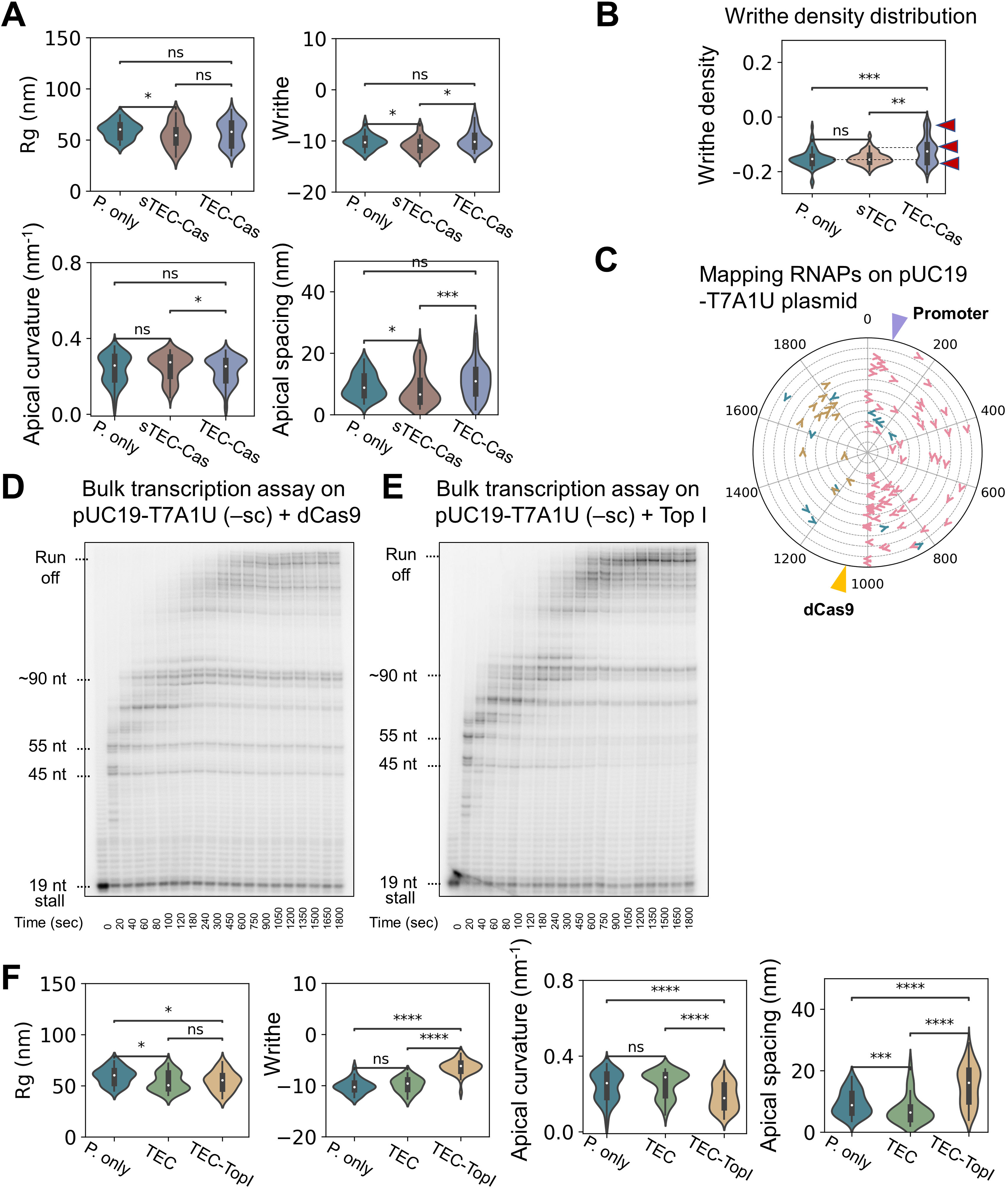
Quantification of plasmid morphology with co-presence of RNAP and dCas9, and of RNAP and Topi, related to Figures 4 and 5. **(A)** Statistical analysis of global plasmid Rg and writhe number (1X1=107), along with local DNA curvature and inter-DNA spacing at apical sites (N=270) for sTEC-Cas and TEC-Cas particles. **(B)** Statistical comparison of the plectoneme writhe density distribution; N=198. **(C)** Mapping all bound RNAPs on pUC19-T7A1U templates using dCas9 as fiducial. For visualization, different plasmids and their associated RNAPs are displayed at distinct radii in the polar plot. RNAP on the transcriptional region, oriented downstream, is marked in pink; RNAP on the non-transcriptional region or pointing upstream, is colored in tan and blue, respectively; N=89 **(D-E)** Electrophoresis assay of RNAP transcription on -sc pUC19-T7A1U templates in the presence of dCas9 and in the presence of Topi over time, respectively. **(F)** Same statistical measurements as in (A), performed on TEC-Topl particles (90 particles and 212 apices total). All statistics are calculated using a Mann-Whitney test, where *p < 0.05, **p < 0.01, ***p < 0.001, ****p < 0.0001; and ns, not significant.

**Figure S7:**
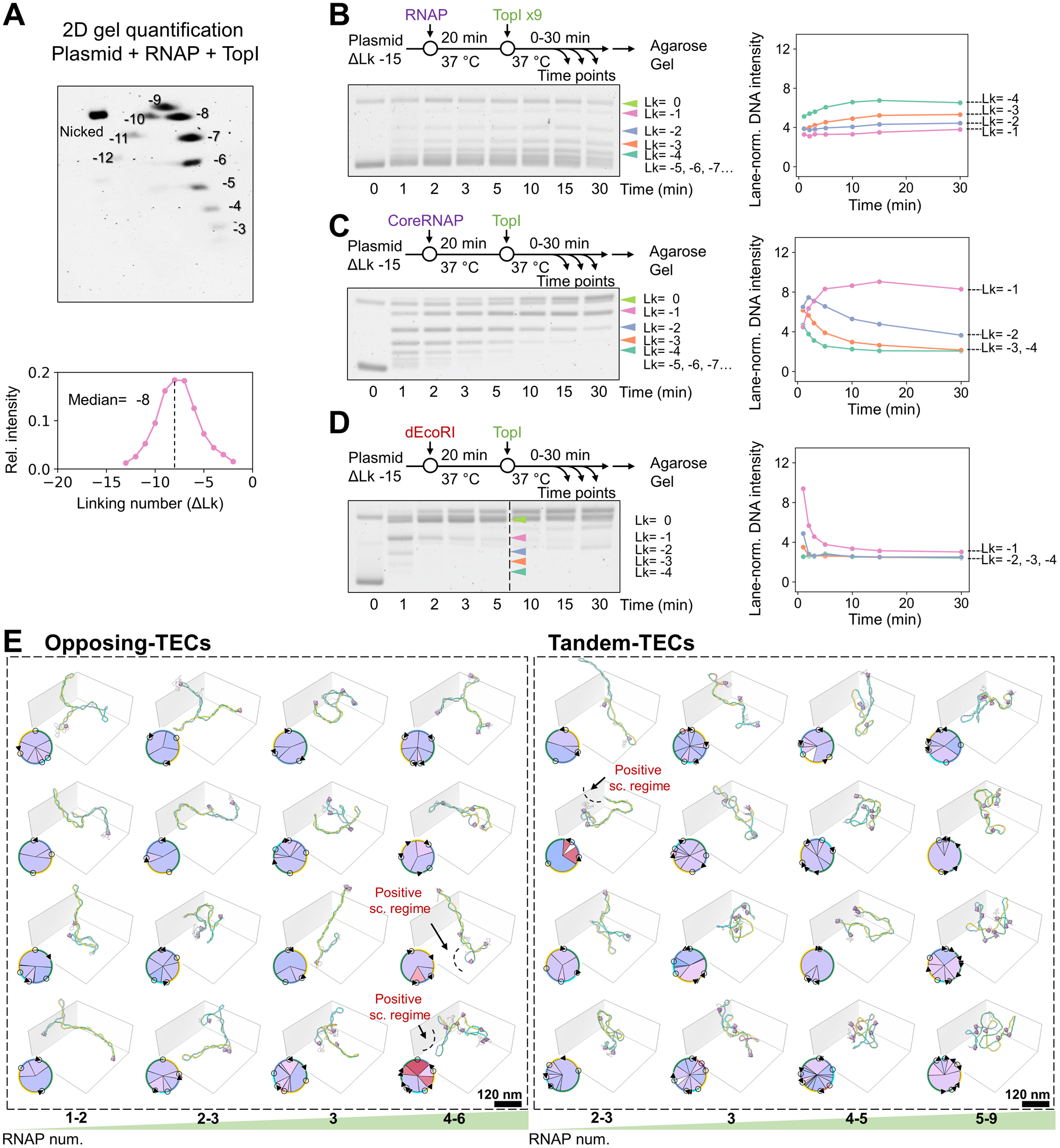
Top I and RNAP coupling slows supercoiling relaxation, and particle collections of dual-promoter TECs, related to Figures 5 and 6. **(A)** 2D electrophoresis gel of-sc pUC19-T7A1U plasmid in the presence of RNAP and Topi, with quantification of the dominant ΔLk value. **(B)** 1D gel-based DNA supercoiling relaxation assay of plasmid in the presence of RNAP and a large excess of Topi. **(C)** DNA supercoiling relaxation assay of plasmid in the presence of RNAP core (lacking σ and unable to form stable transcription bubbles). **(D)** DNA supercoiling relaxation assay of plasmid in the presence of dEcoRI. 1D gel-based linking number quantification is shown in the right panel for B-D. **(E)** Particle collections of opposing-promoter TECs and tandem-promoter TECs, ordered by the number of bound RNAPs. Each dual-promoter TEC is also shown with its corresponding circular plasmid layout, annotated in red to indicate plectonemes extending into the positive supercoiling regime.

## References

1. Baranello, L., Levens, D., Gupta, A., and Kouzine, F. (2012). The importance of being supercoiled: How DNA mechanics regulate dynamic processes. Bba-Gene Regul Mech 1819, 632–638. 10.1016/j.bbagrm.2011.12.007.

2. Gagua, A.V., Belintsev, B.N., and Lyubchenko Yu, L. (1981). Effect of base-pair stability on the melting of superhelical DNA. Nature 294, 662–663. 10.1038/294662a0.

3. Yan, Y., Leng, F., Finzi, L., and Dunlap, D. (2018). Protein-mediated looping of DNA under tension requires supercoiling. Nucleic acids research 46, 2370–2379. 10.1093/nar/gky021.

4. Tripathi, S., Brahmachari, S., Onuchic, J.N., and Levine, H. (2022). DNA supercoiling-mediated collective behavior of co-transcribing RNA polymerases. Nucleic acids research 50, 1269–1279. 10.1093/nar/gkab1252.

5. Patel, H.P., Coppola, S., Pomp, W., Aiello, U., Brouwer, I., Libri, D., and Lenstra, T.L. (2023). DNA supercoiling restricts the transcriptional bursting of neighboring eukaryotic genes. Molecular cell 83, 1573–1587 e1578. 10.1016/j.molcel.2023.04.015.

6. Fu, Z., Guo, M.S., Zhou, W., and Xiao, J. (2024). Differential roles of positive and negative supercoiling in organizing the E. coli genome. Nucleic acids research 52, 724–737. 10.1093/nar/gkad1139.

7. Vayssieres, M., Marechal, N., Yun, L., Lopez Duran, B., Murugasamy, N.K., Fogg, J.M., Zechiedrich, L., Nadal, M., and Lamour, V. (2024). Structural basis of DNA crossover capture by Escherichia coli DNA gyrase. Science 384, 227–232. 10.1126/science.adl5899.

8. Vidmar, V., Borde, C., Bruno, L., Takacs, M., Batisse, C., Saint-Andre, C., Zhu, C., Espeli, O., Lamour, V., and Weixlbaumer, A. (2024). DNA topoisomerase I acts as supercoiling sensor for transcription elongation in E. coli. bioRxiv.

9. Janissen, R., Barth, R., Polinder, M., van der Torre, J., and Dekker, C. (2024). Single-molecule visualization of twin-supercoiled domains generated during transcription. Nucleic acids research 52, 1677–1687. 10.1093/nar/gkad1181.

10. Pyne, A.L.B., Noy, A., Main, K.H.S., Velasco-Berrelleza, V., Piperakis, M.M., Mitchenall, L.A., Cugliandolo, F.M., Beton, J.G., Stevenson, C.E.M., Hoogenboom, B.W., et al. (2021). Base-pair resolution analysis of the effect of supercoiling on DNA flexibility and major groove recognition by triplex-forming oligonucleotides. Nature communications 12, 1053. 10.1038/s41467-021-21243-y.

11. Demurtas, D., Amzallag, A., Rawdon, E.J., Maddocks, J.H., Dubochet, J., and Stasiak, A. (2009). Bending modes of DNA directly addressed by cryo-electron microscopy of DNA minicircles. Nucleic acids research 37, 2882–2893. 10.1093/nar/gkp137.

12. Irobalieva, R.N., Fogg, J.M., Catanese, D.J., Jr., Sutthibutpong, T., Chen, M., Barker, A.K., Ludtke, S.J., Harris, S.A., Schmid, M.F., Chiu, W., and Zechiedrich, L. (2015). Structural diversity of supercoiled DNA. Nature communications 6, 8440. 10.1038/ncomms9440.

13. Amzallag, A., Vaillant, C., Jacob, M., Unser, M., Bednar, J., Kahn, J.D., Dubochet, J., Stasiak, A., and Maddocks, J.H. (2006). 3D reconstruction and comparison of shapes of DNA minicircles observed by cryo-electron microscopy. Nucleic acids research 34, e125. 10.1093/nar/gkl675.

14. Vologodskii, A.V., and Cozzarelli, N.R. (1994). Conformational and Thermodynamic Properties of Supercoiled DNA. Annu Rev Bioph Biom 23, 609–643. DOI 10.1146/annurev.bb.23.060194.003141.

15. Abdella, R., Talyzina, A., Chen, S., Inouye, C.J., Tjian, R., and He, Y. (2021). Structure of the human Mediator-bound transcription preinitiation complex. Science 372, 52–56. 10.1126/science.abg3074.

16. Aibara, S., Schilbach, S., and Cramer, P. (2021). Structures of mammalian RNA polymerase II pre-initiation complexes. Nature 594, 124–128. 10.1038/s41586-021-03554-8.

17. Chen, X.Z., Qi, Y.L., Wu, Z.H., Wang, X.X., Li, J.B., Zhao, D., Hou, H.F., Li, Y., Yu, Z.S., Liu, W.D., et al. (2021). Structural insights into preinitiation complex assembly on core promoters. Science 372, 480–+. ARTN eaba8490 10.1126/science.aba8490.

18. Kettenberger, H., Armache, K.J., and Cramer, P. (2004). Complete RNA polymerase II elongation complex structure and its interactions with NTP and TFIIS. Molecular cell 16, 955–965. DOI 10.1016/j.molcel.2004.11.040.

19. Gnatt, A.L., Cramer, P., Fu, J.H., Bushnell, D.A., and Kornberg, R.D. (2001). Structural basis of transcription:: An RNA polymerase II elongation complex at 3.3 Å resolution. Science 292, 1876–1882. 10.1126/science.1059495.

20. Liu, L.F., and Wang, J.C. (1987). Supercoiling of the DNA-Template during Transcription. Proceedings of the National Academy of Sciences of the United States of America 84, 7024–7027. DOI 10.1073/pnas.84.20.7024.

21. Ma, J., Bai, L., and Wang, M.D. (2013). Transcription Under Torsion. Science 340, 1580–1583. 10.1126/science.1235441.

22. Chong, S.S., Chen, C.Y., Ge, H., and Xie, X.S. (2014). Mechanism of Transcriptional Bursting in Bacteria. Cell 158, 314–326. 10.1016/j.cell.2014.05.038.

23. Leng, F.F., Amado, L., and McMacken, R. (2004). Coupling DNA supercoiling to transcription in defined protein systems. Journal of Biological Chemistry 279, 47564–47571. 10.1074/jbc.M403798200.

24. Kim, S., Beltran, B., Irnov, I., and Jacobs-Wagner, C. (2019). Long-Distance Cooperative and Antagonistic RNA Polymerase Dynamics via DNA Supercoiling. Cell 179, 106–119 e116. 10.1016/j.cell.2019.08.033.

25. Jentink, N., Purnell, C., Kable, B., Swulius, M.T., and Grigoryev, S.A. (2023). Cryoelectron tomography reveals the multiplex anatomy of condensed native chromatin and its unfolding by histone citrullination. Molecular cell 83, 3236–3252 e3237. 10.1016/j.molcel.2023.08.017.

26. Zhang, M., Díaz-Celis, C., Liu, J., Tao, J., Ashby, P.D., Bustamante, C., and Ren, G. (2024). Angle between DNA linker and nucleosome core particle regulates array compaction revealed by individual-particle cryo-electron tomography. Nature communications 15, 4395.

27. Beel, A.J., Azubel, M., Mattei, P.J., and Kornberg, R.D. (2021). Structure of mitotic chromosomes. Molecular cell 81, 4369–4376 e4363. 10.1016/j.molcel.2021.08.020.

28. Chen, M., Bell, J.M., Shi, X., Sun, S.Y., Wang, Z., and Ludtke, S.J. (2019). A complete data processing workflow for cryo-ET and subtomogram averaging. Nature methods 16, 1161–1168. 10.1038/s41592-019-0591-8.

29. Liu, Y.T., Zhang, H., Wang, H., Tao, C.L., Bi, G.Q., and Zhou, Z.H. (2022). Isotropic reconstruction for electron tomography with deep learning. Nature communications 13, 6482. 10.1038/s41467-022-33957-8.

30. Eastman, P., Swails, J., Chodera, J.D., McGibbon, R.T., Zhao, Y., Beauchamp, K.A., Wang, L.P., Simmonett, A.C., Harrigan, M.P., Stern, C.D., et al. (2017). OpenMM 7: Rapid development of high performance algorithms for molecular dynamics. PLoS Comput Biol 13, e1005659. 10.1371/journal.pcbi.1005659.

31. Samee, M.A.H., Bruneau, B.G., and Pollard, K.S. (2019). A De Novo Shape Motif Discovery Algorithm Reveals Preferences of Transcription Factors for DNA Shape Beyond Sequence Motifs. Cell Syst 8, 27–42 e26. 10.1016/j.cels.2018.12.001.

32. ten Heggeler-Bordier, B., Wahli, W., Adrian, M., Stasiak, A., and Dubochet, J. (1992). The apical localization of transcribing RNA polymerases on supercoiled DNA prevents their rotation around the template. The EMBO journal 11, 667–672.

33. Zuo, Y., and Steitz, T.A. (2015). Crystal structures of the E. coli transcription initiation complexes with a complete bubble. Molecular cell 58, 534–540. 10.1016/j.molcel.2015.03.010.

34. Narayanan, A., Vago, F.S., Li, K., Qayyum, M.Z., Yernool, D., Jiang, W., and Murakami, K.S. (2018). Cryo-EM structure of Escherichia coli sigma(70) RNA polymerase and promoter DNA complex revealed a role of sigma non-conserved region during the open complex formation. The Journal of biological chemistry 293, 7367–7375. 10.1074/jbc.RA118.002161.

35. Fang, C., Li, L., Shen, L., Shi, J., Wang, S., Feng, Y., and Zhang, Y. (2019). Structures and mechanism of transcription initiation by bacterial ECF factors. Nucleic acids research 47, 7094–7104. 10.1093/nar/gkz470.

36. Braffman, N.R., Piscotta, F.J., Hauver, J., Campbell, E.A., Link, A.J., and Darst, S.A. (2019). Structural mechanism of transcription inhibition by lasso peptides microcin J25 and capistruin. Proceedings of the National Academy of Sciences of the United States of America 116, 1273–1278. 10.1073/pnas.1817352116.

37. Wood, D.C., and Lebowitz, J. (1984). Effect of supercoiling on the abortive initiation kinetics of the RNA-I promoter of ColE1 plasmid DNA. The Journal of biological chemistry 259, 11184–11187.

38. Ueshima, R., Fujita, N., and Ishihama, A. (1989). DNA supercoiling and temperature shift affect the promoter activity of the Escherichia coli rpoH gene encoding the heat-shock sigma subunit of RNA polymerase. Mol Gen Genet 215, 185–189. 10.1007/BF00339716.

39. Henderson, K.L., Felth, L.C., Molzahn, C.M., Shkel, I., Wang, S., Chhabra, M., Ruff, E.F., Bieter, L., Kraft, J.E., and Record, M.T., Jr. (2017). Mechanism of transcription initiation and promoter escape by E. coli RNA polymerase. Proceedings of the National Academy of Sciences of the United States of America 114, E3032–E3040. 10.1073/pnas.1618675114.

40. Finn, R.D., Orlova, E.V., Gowen, B., Buck, M., and van Heel, M. (2000). Escherichia coli RNA polymerase core and holoenzyme structures. The EMBO journal 19, 6833–6844. 10.1093/emboj/19.24.6833.

41. Josephs, E.A., Kocak, D.D., Fitzgibbon, C.J., McMenemy, J., Gersbach, C.A., and Marszalek, P.E. (2015). Structure and specificity of the RNA-guided endonuclease Cas9 during DNA interrogation, target binding and cleavage. Nucleic acids research 43, 8924–8941. 10.1093/nar/gkv892.

42. Sulc, P., Romano, F., Ouldridge, T.E., Rovigatti, L., Doye, J.P., and Louis, A.A. (2012). Sequence-dependent thermodynamics of a coarse-grained DNA model. The Journal of chemical physics 137, 135101. 10.1063/1.4754132.

43. Kim, S.H., Ganji, M., Kim, E., van der Torre, J., Abbondanzieri, E., and Dekker, C. (2018). DNA sequence encodes the position of DNA supercoils. eLife 7. 10.7554/eLife.36557.

44. Brogna, S., Sato, T.A., and Rosbash, M. (2002). Ribosome components are associated with sites of transcription. Molecular cell 10, 93–104. Doi 10.1016/S1097-2765(02)00565-8.

45. Liu, Y., Li, P.Y., Fan, L., and Wu, M.H. (2018). The nuclear transportation routes of membrane-bound transcription factors. Cell Commun Signal 16. ARTN 12 10.1186/s12964-018-0224-3.

46. Gilbert, N., and Allan, J. (2014). Supercoiling in DNA and chromatin. Current Opinion in Genetics & Development 25, 15–21. 10.1016/j.gde.2013.10.013.

47. Worcel, A., Strogatz, S., and Riley, D. (1981). Structure of Chromatin and the Linking Number of DNA. P Natl Acad Sci-Biol 78, 1461–1465. DOI 10.1073/pnas.78.3.1461.

48. Luijsterburg, M.S., White, M.F., van Driel, R., and Dame, R.T. (2008). The Major Architects of Chromatin: Architectural Proteins in Bacteria, Archaea and Eukaryotes. Crit Rev Biochem Mol 43, 393–418. 10.1080/10409230802528488.

49. Matsumoto, K., and Hirose, S. (2004). Visualization of unconstrained negative supercoils of DNA on polytene chromosomes of Drosophila. Journal of cell science 117, 3797–3805. 10.1242/jcs.01225.

50. Ljungman, M., and Hanawalt, P.C. (1995). Presence of negative torsional tension in the promoter region of the transcriptionally poised dihydrofolate reductase gene in vivo. Nucleic acids research 23, 1782–1789. 10.1093/nar/23.10.1782.

51. Ding, J.Y., Lee, Y.T., Bhandari, Y., Schwieters, C.D., Fan, L.X., Yu, P., Tarosov, S.G., Stagno, J.R., Ma, B.Y., Nussinov, R., et al. (2023). Visualizing RNA conformational and architectural heterogeneity in solution. Nature communications 14. 10.1038/s41467-023-36184-x.

52. Kang, J.Y., Mishanina, T.V., Bao, Y., Chen, J., Llewellyn, E., Liu, J., Darst, S.A., and Landick, R. (2023). An ensemble of interconverting conformations of the elemental paused transcription complex creates regulatory options. Proceedings of the National Academy of Sciences of the United States of America 120, e2215945120. 10.1073/pnas.2215945120.

53. Thomen, P., Bockelmann, U., and Heslot, F. (2002). Rotational drag on DNA: a single molecule experiment. Physical review letters 88, 248102. 10.1103/PhysRevLett.88.248102.

54. Nelson, P. (1999). Transport of torsional stress in DNA. Proceedings of the National Academy of Sciences of the United States of America 96, 14342–14347. 10.1073/pnas.96.25.14342.

55. Chatterjee, P., Goldenfeld, N., and Kim, S. (2021). DNA Supercoiling Drives a Transition between Collective Modes of Gene Synthesis. Physical review letters 127. ARTN 218101 10.1103/PhysRevLett.127.218101.

56. Horberg, J., and Reymer, A. (2020). Specifically bound BZIP transcription factors modulate DNA supercoiling transitions. Scientific reports 10, 18795. 10.1038/s41598-020-75711-4.

57. Kouzine, F., Gupta, A., Baranello, L., Wojtowicz, D., Ben-Aissa, K., Liu, J.H., Przytycka, T.M., and Levens, D. (2013). Transcription-dependent dynamic supercoiling is a short-range genomic force. Nature structural & molecular biology 20, 396–403. 10.1038/nsmb.2517.

58. Aldag, P., Welzel, F., Jakob, L., Schmidbauer, A., Rutkauskas, M., Fettes, F., Grohmann, D., and Seidel, R. (2021). Probing the stability of the SpCas9-DNA complex after cleavage. Nucleic acids research 49, 12411–12421. 10.1093/nar/gkab1072.

59. Bryant, Z., Stone, M.D., Gore, J., Smith, S.B., Cozzarelli, N.R., and Bustamante, C. (2003). Structural transitions and elasticity from torque measurements on DNA. Nature 424, 338–341. 10.1038/nature01810.

60. Cheng, B.K., Zhu, C.X., Ji, C.L., Ahumada, A., and Tse-Dinh, Y.C. (2003). Direct interaction between RNA polymerase and the zinc ribbon domains of DNA topoisomerase I. Journal of Biological Chemistry 278, 30705–30710. 10.1074/jbc.M303403200.

61. Sutormin, D., Galivondzhyan, A., Musharova, O., Travin, D., Rusanova, A., Obraztsova, K., Borukhov, S., and Severinov, K. (2022). Interaction between transcribing RNA polymerase and topoisomerase I prevents R-loop formation in E. coli. Nature communications 13, 4524. 10.1038/s41467-022-32106-5.

62. Bignaud, A., Cockram, C., Borde, C., Groseille, J., Allemand, E., Thierry, A., Marbouty, M., Mozziconacci, J., Espéli, O., and Koszul, R. (2024). Transcription-induced domains form the elementary constraining building blocks of bacterial chromosomes. Nature structural & molecular biology 31. 10.1038/s41594-023-01178-2.

63. Marras, S.A., Gold, B., Kramer, F.R., Smith, I., and Tyagi, S. (2004). Real-time measurement of in vitro transcription. Nucleic acids research 32, e72. 10.1093/nar/gnh068.

64. Lyubchenko, Y.L., and Shlyakhtenko, L.S. (1997). Visualization of supercoiled DNA with atomic force microscopy in situ. Proceedings of the National Academy of Sciences of the United States of America 94, 496–501. 10.1073/pnas.94.2.496.

65. Boles, T.C., White, J.H., and Cozzarelli, N.R. (1990). Structure of plectonemically supercoiled DNA. Journal of molecular biology 213, 931–951. 10.1016/S0022-2836(05)80272-4.

66. Arman, F., Helmut, S., and Reza, E.M. (2015). Molecular Dynamics Simulation of Supercoiled DNA Rings.

67. Huang, J., Schlick, T., and Vologodskii, A. (2001). Dynamics of site juxtaposition in supercoiled DNA. Proceedings of the National Academy of Sciences of the United States of America 98, 968–973. 10.1073/pnas.98.3.968.

68. Forquet, R., Nasser, W., Reverchon, S., and Meyer, S. (2022). Quantitative contribution of the spacer length in the supercoiling-sensitivity of bacterial promoters. Nucleic acids research 50, 7287–7297. 10.1093/nar/gkac579.

69. Bentin, T., and Nielsen, P.E. (2002). In vitro transcription of a torsionally constrained template. Nucleic acids research 30, 803–809. 10.1093/nar/30.3.803.

70. Teves, S.S., and Henikoff, S. (2014). Transcription-generated torsional stress destabilizes nucleosomes. Nature structural & molecular biology 21, 88–94. 10.1038/nsmb.2723.

71. Fosado, Y.A.G., Michieletto, D., Brackley, C.A., and Marenduzzo, D. (2021). Nonequilibrium dynamics and action at a distance in transcriptionally driven DNA supercoiling. Proceedings of the National Academy of Sciences of the United States of America 118. ARTN e1905215118 10.1073/pnas.1905215118.

72. Leng, F.F., and McMacken, R. (2002). Potent stimulation of transcription-coupled DNA supercoiling by sequence-specific DNA-binding proteins. Proceedings of the National Academy of Sciences of the United States of America 99, 9139–9144. 10.1073/pnas.142002099.

73. Brahmachari, S., Tripathi, S., Onuchic, J.N., and Levine, H. (2024). Nucleosomes play a dual role in regulating transcription dynamics. Proceedings of the National Academy of Sciences of the United States of America 121, e2319772121. 10.1073/pnas.2319772121.

74. Golding, I., Paulsson, J., Zawilski, S.M., and Cox, E.C. (2005). Real-time kinetics of gene activity in individual bacteria. Cell 123, 1025–1036. 10.1016/j.cell.2005.09.031.

75. So, L.H., Ghosh, A., Zong, C., Sepulveda, L.A., Segev, R., and Golding, I. (2011). General properties of transcriptional time series in Escherichia coli. Nature genetics 43, 554–560. 10.1038/ng.821.

76. Jinek, M., Chylinski, K., Fonfara, I., Hauer, M., Doudna, J.A., and Charpentier, E. (2012). A programmable dual-RNA-guided DNA endonuclease in adaptive bacterial immunity. Science 337, 816–821. 10.1126/science.1225829.

77. Bizard, A., Garnier, F., and Nadal, M. (2011). TopR2, the second reverse gyrase of Sulfolobus solfataricus, exhibits unusual properties. Journal of molecular biology 408, 839–849. 10.1016/j.jmb.2011.03.030.

78. Aquilina, M., and Dunn, K.E. (2023). Multiplexed Label-Free Biomarker Detection by Targeted Disassembly of Variable-Length DNA Payload Chains. Anal Sens 3. ARTN e202200082 10.1002/anse.202200082.

79. Schorb, M., Haberbosch, I., Hagen, W.J.H., Schwab, Y., and Mastronarde, D.N. (2019). Software tools for automated transmission electron microscopy. Nat Methods 16, 471–477. 10.1038/s41592-019-0396-9.

80. Zheng, S.Q., Palovcak, E., Armache, J.P., Verba, K.A., Cheng, Y., and Agard, D.A. (2017). MotionCor2: anisotropic correction of beam-induced motion for improved cryo-electron microscopy. Nat Methods 14, 331–332. 10.1038/nmeth.4193.

81. Buchholz, T.O., Krull, A., Shahidi, R., Pigino, G., Jekely, G., and Jug, F. (2019). Content-aware image restoration for electron microscopy. Methods Cell Biol 152, 277–289. 10.1016/bs.mcb.2019.05.001.

82. Zhang, K. (2016). Gctf: Real-time CTF determination and correction. J Struct Biol 193, 1–12. 10.1016/j.jsb.2015.11.003.

83. Kremer, J.R., Mastronarde, D.N., and McIntosh, J.R. (1996). Computer visualization of three-dimensional image data using IMOD. J Struct Biol 116, 71–76. 10.1006/jsbi.1996.0013.

84. Pettersen, E.F., Goddard, T.D., Huang, C.C., Couch, G.S., Greenblatt, D.M., Meng, E.C., and Ferrin, T.E. (2004). UCSF Chimera--a visualization system for exploratory research and analysis. J Comput Chem 25, 1605–1612. 10.1002/jcc.20084.

85. Ludtke, S.J., Baldwin, P.R., and Chiu, W. (1999). EMAN: semiautomated software for high-resolution single-particle reconstructions. J Struct Biol 128, 82–97. 10.1006/jsbi.1999.4174.

86. Punjani, A., Rubinstein, J.L., Fleet, D.J., and Brubaker, M.A. (2017). cryoSPARC: algorithms for rapid unsupervised cryo-EM structure determination. Nat Methods 14, 290–296. 10.1038/nmeth.4169.

87. Sanchez-Garcia, R., Gomez-Blanco, J., Cuervo, A., Carazo, J.M., Sorzano, C.O.S., and Vargas, J. (2021). DeepEMhancer: a deep learning solution for cryo-EM volume post-processing. Commun Biol 4, 874. 10.1038/s42003-021-02399-1.

88. Meng, E.C., Goddard, T.D., Pettersen, E.F., Couch, G.S., Pearson, Z.J., Morris, J.H., and Ferrin, T.E. (2023). UCSF ChimeraX: Tools for structure building and analysis. Protein Sci 32, e4792. 10.1002/pro.4792.

89. Liebschner, D., Afonine, P.V., Baker, M.L., Bunkoczi, G., Chen, V.B., Croll, T.I., Hintze, B., Hung, L.W., Jain, S., McCoy, A.J., et al. (2019). Macromolecular structure determination using X-rays, neutrons and electrons: recent developments in Phenix. Acta Crystallogr D Struct Biol 75, 861–877. 10.1107/S2059798319011471.

90. Croll, T.I. (2018). ISOLDE: a physically realistic environment for model building into low-resolution electron-density maps. Acta Crystallogr D Struct Biol 74, 519–530. 10.1107/S2059798318002425.

91. Chen, V.B., Arendall, W.B., 3rd, Headd, J.J., Keedy, D.A., Immormino, R.M., Kapral, G.J., Murray, L.W., Richardson, J.S., and Richardson, D.C. (2010). MolProbity: all-atom structure validation for macromolecular crystallography. Acta crystallographica. Section D, Biological crystallography 66, 12–21. 10.1107/S0907444909042073.

92. Klenin, K., and Langowski, J. (2000). Computation of writhe in modeling of supercoiled DNA. Biopolymers 54, 307–317. 10.1002/1097-0282(20001015)54:5<307::AID-BIP20>3.0.CO;2-Y.

93. Johnson, R.A. (1929). Modern geometry: an elementary treatise on the geometry of the triangle and the circle (Houghton, Mifflin Company).

94. Burns, J.R., Seifert, A., Fertig, N., and Howorka, S. (2016). A biomimetic DNA-based channel for the ligand-controlled transport of charged molecular cargo across a biological membrane. Nat Nanotechnol 11, 152–156.

95. Suma, A., Poppleton, E., Matthies, M., Sulc, P., Romano, F., Louis, A.A., Doye, J.P.K., Micheletti, C., and Rovigatti, L. (2019). TacoxDNA: A user-friendly web server for simulations of complex DNA structures, from single strands to origami. Journal of computational chemistry 40, 2586–2595. 10.1002/jcc.26029.

96. Wright, D.J., King, K., and Modrich, P. (1989). The Negative Charge of Glu-111 Is Required to Activate the Cleavage Center of Ecori Endonuclease. J Biol Chem 264, 11816–11821.

97. Weigert, M., Schmidt, U., Boothe, T., Muller, A., Dibrov, A., Jain, A., Wilhelm, B., Schmidt, D., Broaddus, C., Culley, S., et al. (2018). Content-aware image restoration: pushing the limits of fluorescence microscopy. Nat Methods 15, 1090–1097. 10.1038/s41592-018-0216-7.

98. Lee, T.C., Kashyap, R.L., and Chu, C.N. (1994). Building Skeleton Models Via 3-D Medial Surface Axis Thinning Algorithms. Cvgip-Graph Model Im 56, 462–478. DOI 10.1006/cgip.1994.1042.

