## Supplementary Data for "Apical Localization of RNA Polymerases Modulate Transcription Dynamics and Supercoiling Domains Revealed by Cryo-ET"

[1] California Institute for Quantitative Biosciences, University of California, Berkeley, USA
[2] The Molecular Foundry, Lawrence Berkeley National Laboratory, Berkeley, USA
[3] Department of Chemistry, University of California Berkeley, Berkeley, CA, USA
[4] Department of Biology, Stanford University, Stanford, CA USA
[5] Innovative Genomics Institute, University of California Berkeley, Berkeley, CA, USA
[6] Department of Molecular and Cell Biology, University of California, Berkeley, USA
[7] Howard Hughes Medical Institute, University of California, Berkeley, USA
[8] Department of Physics, University of California, Berkeley, USA
[9] Department of Chemistry, University of California, Berkeley, USA
[10] Molecular Biophysics and Integrative Bioimaging Division, Lawrence Berkeley National Laboratory, USA
[11] Kavli Energy Nanoscience Institute, University of California, Berkeley, USA
[12]Jason Choy laboratory of single molecule biophysics, University of California, Berkeley, USA
* These authors contributed equally

SD1: Cryo-ET 3D reconstruction workflow .......... 2
SD2: Tracing of DNA within the cryo-ET map for plasmid particle modeling .......... 3
SD3: Cryo-EM single-particle data collection and refinement statistics .......... 4
SD4: Cryo-ET per-particle data collection and evaluation table .......... 5
Particle gallery of plasmid in low salt condition (P. LS) .......... 9
Particle gallery of plasmid in high salt condition (P. HS) .......... 26
Particle gallery of stalled transcription elongate complexes (sTEC) .......... 41
Particle gallery of plasmid particles in the presence of dCas9 (P. Cas) .......... 55
Particle gallery of transcription elongate complexes (TEC) .......... 71
Particle gallery of sTECs in the presence of dCas9 (sTEC-Cas9) .......... 83
Particle gallery of TECs in the presence of dCas9 (TEC-Cas9) .......... 100
Particle gallery of TECs in the presence of TopI (TEC-TopI) .......... 121
Particle gallery of TECs containing two opposing promoters (Opp-TEC) .......... 141
Particle gallery of TECs containing two tandem promoters (Tan-TEC) .......... 159

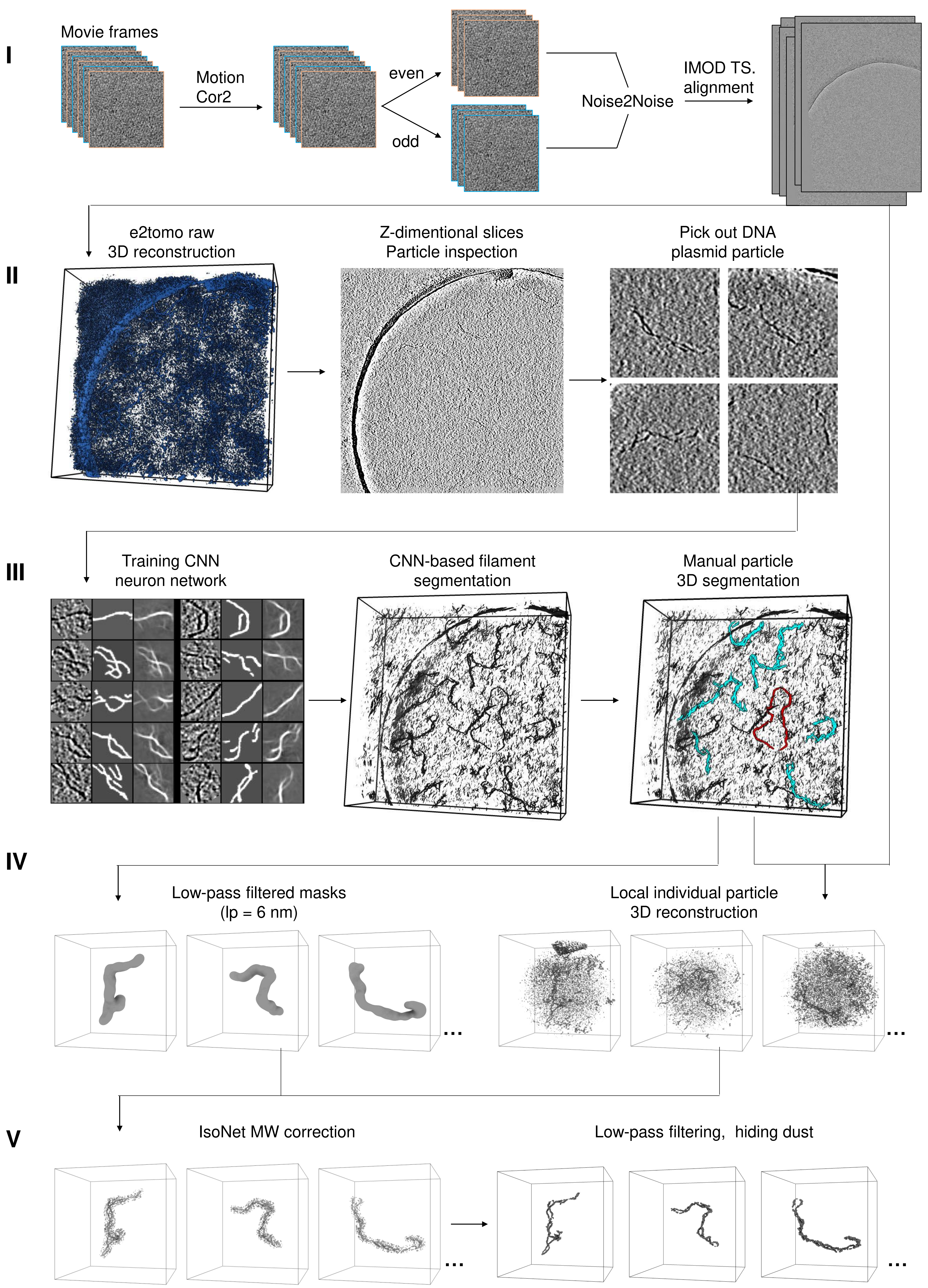

**Supplementary Data 1: Cryo-ET 3D reconstruction workflow, related to Figure 1.** Tomographic 3D reconstruction workflow of the plasmid particle from the tilt series. I: The motion-corrected tilt series were split into even and odd sets and subjected to deep learning-based image denoising (N2N method). II: The denoised tilt series were initially patch tracking aligned in imod and submitted for full tomogram reconstruction in e2tomo. III: DNA segments selected from the large 3D map were used to train a CNN network for annotating all DNA elements in the tomogram. IV: A mask of the target particle was created to calculate its center of mass, which was then used to extract raw tilts from the initial low-contrast micrograph for localized 3D reconstruction of individual particles. V: The map was processed with IsoNet for missing wedge correction, followed by low-pass filtering to 2 nm and dust removal, resulting in final map for modeling.

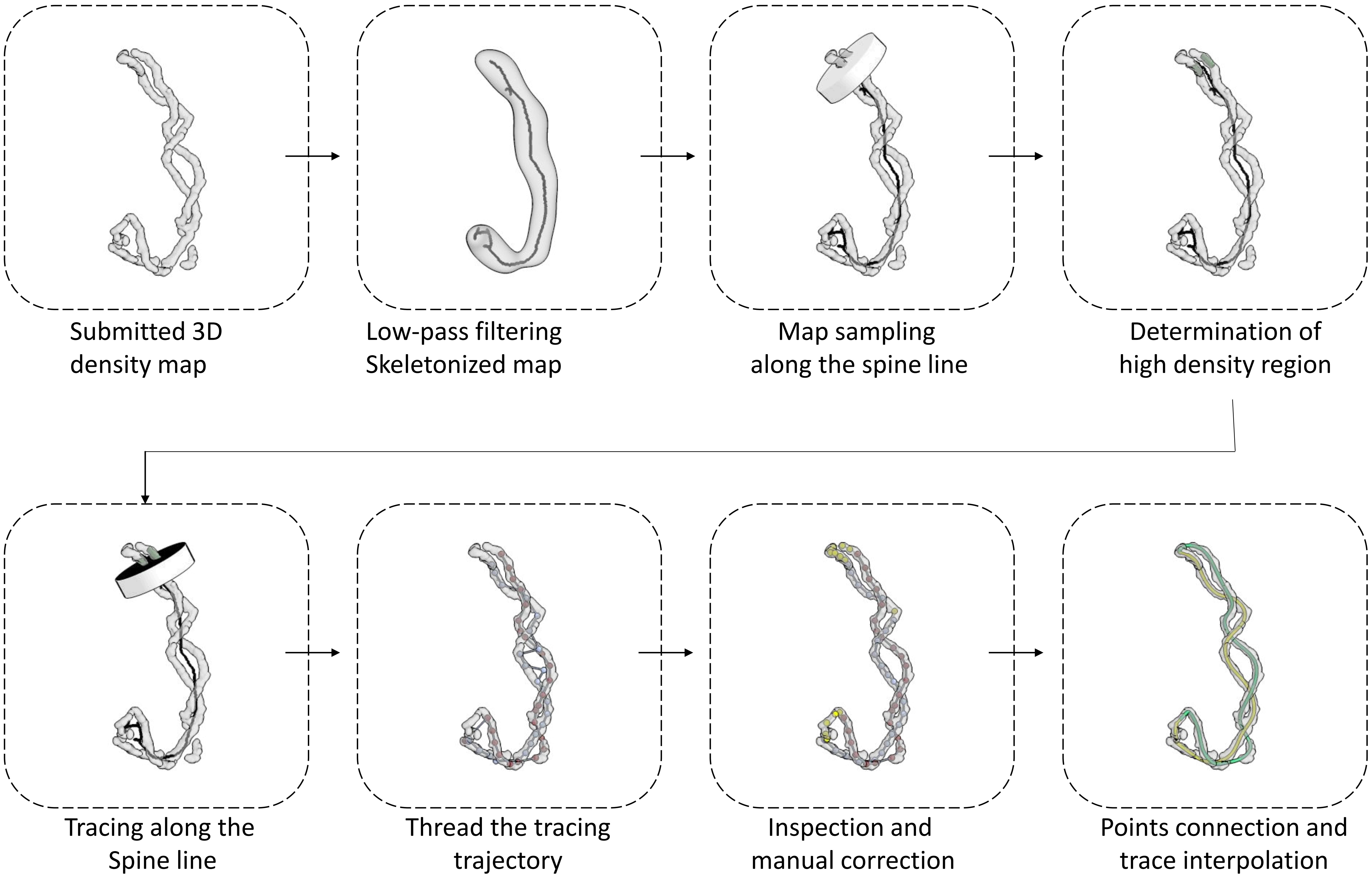

**Supplementary Data 2 : Tracing of DNA within the cryo-ET map for plasmid particle modeling, related to Figure 1.** DNA model fitting workflow, see also materials and methods.

| | TEC1-SC. | TEC2-SC. |
|---|---|---|
| EMDB: Map | EMD-71675 | EMD-71676 |
| PDB: Model | 9PIP | 9PIQ |
| Microscope | Thermo Fisher Krios G3i | Thermo Fisher Krios G3i |
| Detector | Gatan K3 | Gatan K3 |
| Voltage (kV) | 300 | 300 |
| Magnification | 81,000 | 81,000 |
| Electron exposure($e^-/Å^2$) | 50 | 50 |
| Defocus range (μm) | −0.6 to −1.6 | −0.6 to −1.6 |
| Pixel size (Å) | 1.05 | 1.05 |
| Images | 10476 | 10476 |
| Initial particles | 2,063,152 | 2,063,152 |
| Final particles | 46,743 | 54,869 |
| Symmetry | C1 | C1 |
| Box size (pix) | 324 | 324 |
| Resolution (Å) (FSC = 0.143) | 2.95 | 2.94 |
| Map sharpening B factor ($Å^2$) | -- | -- |
| Initial model used (PDB code) | 6ALH | 6ALH |
| Model resolution (Å) | 4.4 | 4.4 |
| FSC threshold | 0.5 | 0.5 |
| Model composition | | |
| Non-hydrogen atoms | 26008 | 26008 |
| Protein residues | 3230 | 3230 |
| Ligands | -- | -- |
| B factors ($Å^2$) | | |
| Protein | 50.7 | 49.0 |
| Ligand | -- | -- |
| R.m.s. deviations | | |
| Bond lengths(Å) | 0.0064 | 0.006 |
| Bond angles (°) | 1.17 | 1.13 |
| Validation | | |
| MolProbity score | 1.64 | 1.71 |
| Clashscore | 5.45 | 5.73 |
| Poor rotamers(%) | 0.04 | 0.00 |
| Ramachandran plot | | |
| Favored (%) | 94.95 | 94.12 |
| Allowed (%) | 5.05 | 5.88 |
| Disallowed (%) | 0.00 | 0.00 |

**Supplementary Data 3. Cryo-EM single-particle data collection and refinement statistics**

### Supplementary Data 4. Cryo-ET per-particle data collection and evaluation

| Par. id | Name abbr. | Bound RNAP | Bound dCas9 | Salt condition | Tilt range | Tilt step | TEM /Cam. | Mag. | Apix | Total dose | EMDB map ID | Montage position | Resolution (FSC=0.143) | Resolution (FSC=0.5) |
|---|---|---|---|---|---|---|---|---|---|---|---|---|---|---|
| # | | # | # | | (°) | (°) | | | (Å²) | (e-/Å²) | # | (row, col) | Even vs. Odd (Å) | Map vs. Model (Å) |
| 1 | P. LS | 0 | 0 | 5 mM K+ | -51 to 51 | 3 | G3/K3 | 53K | 1.67 | 183 | EMD-47843 | 1,1 | 33 | 38.7 |
| 2 | P. LS | 0 | 0 | 5 mM K+ | -51 to 51 | 3 | G3/K3 | 53K | 1.67 | 183 | EMD-47843 | 1,2 | 30 | 36 |
| 3 | P. LS | 0 | 0 | 5 mM K+ | -51 to 51 | 3 | G3/K3 | 53K | 1.67 | 183 | EMD-47843 | 1,3 | 34.1 | 38.6 |
| 4 | P. LS | 0 | 0 | 5 mM K+ | -51 to 51 | 3 | G3/K3 | 53K | 1.67 | 183 | EMD-47843 | 1,4 | 28.7 | 40.9 |
| 5 | P. LS | 0 | 0 | 5 mM K+ | -51 to 51 | 3 | G3/K3 | 53K | 1.67 | 183 | EMD-47843 | 1,5 | 28.2 | 40 |
| 6 | P. LS | 0 | 0 | 5 mM K+ | -51 to 51 | 3 | G3/K3 | 53K | 1.67 | 183 | EMD-47843 | 1,6 | 29.7 | 38.9 |
| 7 | P. LS | 0 | 0 | 5 mM K+ | -51 to 51 | 3 | G3/K3 | 53K | 1.67 | 183 | EMD-47843 | 1,7 | 29.3 | 41.5 |
| 8 | P. LS | 0 | 0 | 5 mM K+ | -51 to 51 | 3 | G3/K3 | 53K | 1.67 | 183 | EMD-47843 | 2,1 | 31.2 | 32.1 |
| 9 | P. LS | 0 | 0 | 5 mM K+ | -51 to 51 | 3 | G3/K3 | 53K | 1.67 | 183 | EMD-47843 | 2,2 | 30.8 | 38.5 |
| 10 | P. LS | 0 | 0 | 5 mM K+ | -51 to 51 | 3 | G3/K3 | 53K | 1.67 | 183 | EMD-47843 | 2,3 | 31 | 42.9 |
| 11 | P. LS | 0 | 0 | 5 mM K+ | -51 to 51 | 3 | G3/K3 | 53K | 1.67 | 183 | EMD-47843 | 2,4 | 29.2 | 39.6 |
| 12 | P. LS | 0 | 0 | 5 mM K+ | -51 to 51 | 3 | G3/K3 | 53K | 1.67 | 183 | EMD-47843 | 2,5 | 30.4 | 37.6 |
| 13 | P. LS | 0 | 0 | 5 mM K+ | -51 to 51 | 3 | G3/K3 | 53K | 1.67 | 183 | EMD-47843 | 2,6 | 31.4 | 32.4 |
| 14 | P. LS | 0 | 0 | 5 mM K+ | -51 to 51 | 3 | G3/K3 | 53K | 1.67 | 183 | EMD-47843 | 2,7 | 29.3 | 38.4 |
| 15 | P. LS | 0 | 0 | 5 mM K+ | -51 to 51 | 3 | G3/K3 | 53K | 1.67 | 183 | EMD-47843 | 3,1 | 30.6 | 42.7 |
| 16 | P. LS | 0 | 0 | 5 mM K+ | -51 to 51 | 3 | G3/K3 | 53K | 1.67 | 183 | EMD-47843 | 3,2 | 31.3 | 47.6 |
| 17 | P. LS | 0 | 0 | 5 mM K+ | -51 to 51 | 3 | G3/K3 | 53K | 1.67 | 183 | EMD-47843 | 3,3 | 29.2 | 39.8 |
| 18 | P. LS | 0 | 0 | 5 mM K+ | -51 to 51 | 3 | G3/K3 | 53K | 1.67 | 183 | EMD-47843 | 3,4 | 27.9 | 35 |
| 19 | P. LS | 0 | 0 | 5 mM K+ | -51 to 51 | 3 | G3/K3 | 53K | 1.67 | 183 | EMD-47843 | 3,5 | 25.7 | 35.3 |
| 20 | P. LS | 0 | 0 | 5 mM K+ | -51 to 51 | 3 | G3/K3 | 53K | 1.67 | 183 | EMD-47843 | 3,6 | 24.7 | 33.6 |
| 21 | P. LS | 0 | 0 | 5 mM K+ | -51 to 51 | 3 | G3/K3 | 53K | 1.67 | 183 | EMD-47843 | 3,7 | 31.4 | 39.3 |
| 22 | P. LS | 0 | 0 | 5 mM K+ | -51 to 51 | 3 | G3/K3 | 53K | 1.67 | 183 | EMD-47843 | 4,1 | 29.8 | 38.9 |
| 23 | P. LS | 0 | 0 | 5 mM K+ | -51 to 51 | 3 | G3/K3 | 53K | 1.67 | 183 | EMD-47843 | 4,2 | 31.2 | 40 |
| 24 | P. LS | 0 | 0 | 5 mM K+ | -51 to 51 | 3 | G3/K3 | 53K | 1.67 | 183 | EMD-47843 | 4,3 | 31.9 | 43.6 |
| 25 | P. LS | 0 | 0 | 5 mM K+ | -51 to 51 | 3 | G3/K3 | 53K | 1.67 | 183 | EMD-47843 | 4,4 | 28.2 | 39.1 |
| 26 | P. LS | 0 | 0 | 5 mM K+ | -51 to 51 | 3 | G3/K3 | 53K | 1.67 | 183 | EMD-47843 | 4,5 | 31.2 | 33 |
| 27 | P. LS | 0 | 0 | 5 mM K+ | -51 to 51 | 3 | G3/K3 | 53K | 1.67 | 183 | EMD-47843 | 4,6 | 27.9 | 37 |
| 28 | P. LS | 0 | 0 | 5 mM K+ | -51 to 51 | 3 | G3/K3 | 53K | 1.67 | 183 | EMD-47843 | 4,7 | 29.9 | 38 |
| 29 | P. LS | 0 | 0 | 5 mM K+ | -51 to 51 | 3 | G3/K3 | 53K | 1.67 | 183 | EMD-47843 | 5,1 | 29 | 33.4 |
| 30 | P. LS | 0 | 0 | 5 mM K+ | -51 to 51 | 3 | G3/K3 | 53K | 1.67 | 183 | EMD-47843 | 5,2 | 29 | 36.2 |
| 31 | P. LS | 0 | 0 | 5 mM K+ | -51 to 51 | 3 | G3/K3 | 53K | 1.67 | 183 | EMD-47843 | 5,3 | 33.6 | 38.2 |
| 32 | P. LS | 0 | 0 | 5 mM K+ | -51 to 51 | 3 | G3/K3 | 53K | 1.67 | 183 | EMD-47843 | 5,4 | 27.1 | 32.9 |
| 33 | P. LS | 0 | 0 | 5 mM K+ | -51 to 51 | 3 | G3/K3 | 53K | 1.67 | 183 | EMD-47843 | 5,5 | 24.7 | 35.7 |
| 34 | P. LS | 0 | 0 | 5 mM K+ | -51 to 51 | 3 | G3/K3 | 53K | 1.67 | 183 | EMD-47843 | 5,6 | 28.1 | 34.6 |
| 35 | P. LS | 0 | 0 | 5 mM K+ | -51 to 51 | 3 | G3/K3 | 53K | 1.67 | 183 | EMD-47843 | 5,7 | 24.6 | 37.1 |
| 36 | P. HS | 0 | 0 | 40 mM K+, 5 mM Mg2+ | -51 to 51 | 3 | G3/K3 | 53K | 1.67 | 183 | EMD-47844 | 1,1 | 27.6 | 34.2 |
| 37 | P. HS | 0 | 0 | 40 mM K+, 5 mM Mg2+ | -51 to 51 | 3 | G3/K3 | 53K | 1.67 | 183 | EMD-47844 | 1,2 | 26.9 | 38.1 |
| 38 | P. HS | 0 | 0 | 40 mM K+, 5 mM Mg2+ | -51 to 51 | 3 | G3/K3 | 53K | 1.67 | 183 | EMD-47844 | 1,3 | 27.8 | 36 |
| 39 | P. HS | 0 | 0 | 40 mM K+, 5 mM Mg2+ | -51 to 51 | 3 | G3/K3 | 53K | 1.67 | 183 | EMD-47844 | 1,4 | 27.3 | 34.1 |
| 40 | P. HS | 0 | 0 | 40 mM K+, 5 mM Mg2+ | -51 to 51 | 3 | G3/K3 | 53K | 1.67 | 183 | EMD-47844 | 1,5 | 29.5 | 36.6 |
| 41 | P. HS | 0 | 0 | 40 mM K+, 5 mM Mg2+ | -51 to 51 | 3 | G3/K3 | 53K | 1.67 | 183 | EMD-47844 | 1,6 | 27 | 32.8 |
| 42 | P. HS | 0 | 0 | 40 mM K+, 5 mM Mg2+ | -51 to 51 | 3 | G3/K3 | 53K | 1.67 | 183 | EMD-47844 | 2,1 | 26.3 | 38 |
| 43 | P. HS | 0 | 0 | 40 mM K+, 5 mM Mg2+ | -51 to 51 | 3 | G3/K3 | 53K | 1.67 | 183 | EMD-47844 | 2,2 | 29 | 37.2 |
| 44 | P. HS | 0 | 0 | 40 mM K+, 5 mM Mg2+ | -51 to 51 | 3 | G3/K3 | 53K | 1.67 | 183 | EMD-47844 | 2,3 | 26.3 | 42.8 |
| 45 | P. HS | 0 | 0 | 40 mM K+, 5 mM Mg2+ | -51 to 51 | 3 | G3/K3 | 53K | 1.67 | 183 | EMD-47844 | 2,4 | 28.9 | 31.5 |
| 46 | P. HS | 0 | 0 | 40 mM K+, 5 mM Mg2+ | -51 to 51 | 3 | G3/K3 | 53K | 1.67 | 183 | EMD-47844 | 2,5 | 27.8 | 35.1 |
| 47 | P. HS | 0 | 0 | 40 mM K+, 5 mM Mg2+ | -51 to 51 | 3 | G3/K3 | 53K | 1.67 | 183 | EMD-47844 | 2,6 | 26.3 | 36.5 |
| 48 | P. HS | 0 | 0 | 40 mM K+, 5 mM Mg2+ | -51 to 51 | 3 | G3/K3 | 53K | 1.67 | 183 | EMD-47844 | 3,1 | 28.1 | 32.3 |
| 49 | P. HS | 0 | 0 | 40 mM K+, 5 mM Mg2+ | -51 to 51 | 3 | G3/K3 | 53K | 1.67 | 183 | EMD-47844 | 3,2 | 27.2 | 32.1 |
| 50 | P. HS | 0 | 0 | 40 mM K+, 5 mM Mg2+ | -51 to 51 | 3 | G3/K3 | 53K | 1.67 | 183 | EMD-47844 | 3,3 | 29.7 | 35.6 |
| 51 | P. HS | 0 | 0 | 40 mM K+, 5 mM Mg2+ | -51 to 51 | 3 | G3/K3 | 53K | 1.67 | 183 | EMD-47844 | 3,4 | 27.4 | 36.9 |
| 52 | P. HS | 0 | 0 | 40 mM K+, 5 mM Mg2+ | -51 to 51 | 3 | G3/K3 | 53K | 1.67 | 183 | EMD-47844 | 3,5 | 27.3 | 34.6 |
| 53 | P. HS | 0 | 0 | 40 mM K+, 5 mM Mg2+ | -51 to 51 | 3 | G3/K3 | 53K | 1.67 | 183 | EMD-47844 | 3,6 | 29.6 | 41.2 |
| 54 | P. HS | 0 | 0 | 40 mM K+, 5 mM Mg2+ | -51 to 51 | 3 | G3/K3 | 53K | 1.67 | 183 | EMD-47844 | 4,1 | 27.3 | 31.7 |
| 55 | P. HS | 0 | 0 | 40 mM K+, 5 mM Mg2+ | -51 to 51 | 3 | G3/K3 | 53K | 1.67 | 183 | EMD-47844 | 4,2 | 26.7 | 39.2 |
| 56 | P. HS | 0 | 0 | 40 mM K+, 5 mM Mg2+ | -51 to 51 | 3 | G3/K3 | 53K | 1.67 | 183 | EMD-47844 | 4,3 | 26.7 | 42 |
| 57 | P. HS | 0 | 0 | 40 mM K+, 5 mM Mg2+ | -51 to 51 | 3 | G3/K3 | 53K | 1.67 | 183 | EMD-47844 | 4,4 | 29.7 | 45.6 |
| 58 | P. HS | 0 | 0 | 40 mM K+, 5 mM Mg2+ | -51 to 51 | 3 | G3/K3 | 53K | 1.67 | 183 | EMD-47844 | 4,5 | 26.9 | 42.4 |
| 59 | P. HS | 0 | 0 | 40 mM K+, 5 mM Mg2+ | -51 to 51 | 3 | G3/K3 | 53K | 1.67 | 183 | EMD-47844 | 4,6 | 27 | 38.9 |
| 60 | P. HS | 0 | 0 | 40 mM K+, 5 mM Mg2+ | -51 to 51 | 3 | G3/K3 | 53K | 1.67 | 183 | EMD-47844 | 5,1 | 28.8 | 41.1 |
| 61 | P. HS | 0 | 0 | 40 mM K+, 5 mM Mg2+ | -51 to 51 | 3 | G3/K3 | 53K | 1.67 | 183 | EMD-47844 | 5,2 | 27.4 | 42.1 |
| 62 | P. HS | 0 | 0 | 40 mM K+, 5 mM Mg2+ | -51 to 51 | 3 | G3/K3 | 53K | 1.67 | 183 | EMD-47844 | 5,3 | 28.1 | 35.4 |
| 63 | P. HS | 0 | 0 | 40 mM K+, 5 mM Mg2+ | -51 to 51 | 3 | G3/K3 | 53K | 1.67 | 183 | EMD-47844 | 5,4 | 26.7 | 53.5 |
| 64 | P. HS | 0 | 0 | 40 mM K+, 5 mM Mg2+ | -51 to 51 | 3 | G3/K3 | 53K | 1.67 | 183 | EMD-47844 | 5,5 | 28.2 | 47.7 |
| 65 | sTEC | 1 | 0 | 40 mM K+, 5 mM Mg2+ | -51 to 51 | 3 | G3/K3 | 53K | 1.67 | 183 | EMD-47847 | 1,1 | 27 | 44.7 |
| 66 | sTEC | 1 | 0 | 40 mM K+, 5 mM Mg2+ | -51 to 51 | 3 | G3/K3 | 53K | 1.67 | 183 | EMD-47847 | 1,2 | 26.9 | 35.6 |
| 67 | sTEC | 1 | 0 | 40 mM K+, 5 mM Mg2+ | -51 to 51 | 3 | G3/K3 | 53K | 1.67 | 183 | EMD-47847 | 1,3 | 28.7 | 45.3 |
| 68 | sTEC | 1 | 0 | 40 mM K+, 5 mM Mg2+ | -51 to 51 | 3 | G3/K3 | 53K | 1.67 | 183 | EMD-47847 | 1,4 | 28.3 | 48.3 |
| 69 | sTEC | 1 | 0 | 40 mM K+, 5 mM Mg2+ | -51 to 51 | 3 | G3/K3 | 53K | 1.67 | 183 | EMD-47847 | 1,5 | 25.9 | 51.5 |
| 70 | sTEC | 1 | 0 | 40 mM K+, 5 mM Mg2+ | -51 to 51 | 3 | G3/K3 | 53K | 1.67 | 183 | EMD-47847 | 1,6 | 28.4 | 44.8 |
| 71 | sTEC | 1 | 0 | 40 mM K+, 5 mM Mg2+ | -51 to 51 | 3 | G3/K3 | 53K | 1.67 | 183 | EMD-47847 | 2,1 | 28.2 | 42.8 |
| 72 | sTEC | 1 | 0 | 40 mM K+, 5 mM Mg2+ | -51 to 51 | 3 | G3/K3 | 53K | 1.67 | 183 | EMD-47847 | 2,2 | 27.3 | 40.1 |
| 73 | sTEC | 2 | 0 | 40 mM K+, 5 mM Mg2+ | -51 to 51 | 3 | G3/K3 | 53K | 1.67 | 183 | EMD-47847 | 2,3 | 25.3 | 41 |
| 74 | sTEC | 1 | 0 | 40 mM K+, 5 mM Mg2+ | -51 to 51 | 3 | G3/K3 | 53K | 1.67 | 183 | EMD-47847 | 2,4 | 28.9 | 53.2 |
| 75 | sTEC | 1 | 0 | 40 mM K+, 5 mM Mg2+ | -51 to 51 | 3 | G3/K3 | 53K | 1.67 | 183 | EMD-47847 | 2,5 | 28 | 45.3 |
| 76 | sTEC | 1 | 0 | 40 mM K+, 5 mM Mg2+ | -51 to 51 | 3 | G3/K3 | 53K | 1.67 | 183 | EMD-47847 | 2,6 | 27.4 | 35.6 |
| 77 | sTEC | 1 | 0 | 40 mM K+, 5 mM Mg2+ | -51 to 51 | 3 | G3/K3 | 53K | 1.67 | 183 | EMD-47847 | 3,1 | 27.2 | 45.3 |
| 78 | sTEC | 1 | 0 | 40 mM K+, 5 mM Mg2+ | -51 to 51 | 3 | G3/K3 | 53K | 1.67 | 183 | EMD-47847 | 3,2 | 30.7 | 46.3 |
| 79 | sTEC | 1 | 0 | 40 mM K+, 5 mM Mg2+ | -51 to 51 | 3 | G3/K3 | 53K | 1.67 | 183 | EMD-47847 | 3,3 | 27.7 | 41.2 |
| 80 | sTEC | 1 | 0 | 40 mM K+, 5 mM Mg2+ | -51 to 51 | 3 | G3/K3 | 53K | 1.67 | 183 | EMD-47847 | 3,4 | 28.7 | 41.8 |
| 81 | sTEC | 1 | 0 | 40 mM K+, 5 mM Mg2+ | -51 to 51 | 3 | G3/K3 | 53K | 1.67 | 183 | EMD-47847 | 3,5 | 29.1 | 54.3 |
| 82 | sTEC | 1 | 0 | 40 mM K+, 5 mM Mg2+ | -51 to 51 | 3 | G3/K3 | 53K | 1.67 | 183 | EMD-47847 | 3,6 | 27 | 41 |
| 83 | sTEC | 1 | 0 | 40 mM K+, 5 mM Mg2+ | -51 to 51 | 3 | G3/K3 | 53K | 1.67 | 183 | EMD-47847 | 4,1 | 30.2 | 46.8 |

| Par. id | Name abbr. | Bound RNAP | Bound dCas9 | Salt condition | Tilt range | Tilt step | TEM /Cam. | Mag. | Apix | Total dose | EMDB map ID | Montage position | Resolution (FSC=0.143) | Resolution (FSC=0.5) |
|---|---|---|---|---|---|---|---|---|---|---|---|---|---|---|
| # | | # | # | | (°) | (°) | | | (Å²) | (e-/Å²) | # | (row, col) | Even vs. Odd (Å) | Map vs. Model (Å) |
| 84 | sTEC | 1 | 0 | 40 mM K+, 5 mM Mg2+ | -51 to 51 | 3 | G3/K3 | 53K | 1.67 | 183 | EMD-47847 | 4,2 | 29.5 | 37.8 |
| 85 | sTEC | 1 | 0 | 40 mM K+, 5 mM Mg2+ | -51 to 51 | 3 | G3/K3 | 53K | 1.67 | 183 | EMD-47847 | 4,3 | 30.9 | 38.5 |
| 86 | sTEC | 1 | 0 | 40 mM K+, 5 mM Mg2+ | -51 to 51 | 3 | G3/K3 | 53K | 1.67 | 183 | EMD-47847 | 4,4 | 26.6 | 45.5 |
| 87 | sTEC | 1 | 0 | 40 mM K+, 5 mM Mg2+ | -51 to 51 | 3 | G3/K3 | 53K | 1.67 | 183 | EMD-47847 | 4,5 | 26.5 | 48.3 |
| 88 | sTEC | 1 | 0 | 40 mM K+, 5 mM Mg2+ | -51 to 51 | 3 | G3/K3 | 53K | 1.67 | 183 | EMD-47847 | 4,6 | 25.6 | 43 |
| 89 | sTEC | 1 | 0 | 40 mM K+, 5 mM Mg2+ | -51 to 51 | 3 | G3/K3 | 53K | 1.67 | 183 | EMD-47847 | 5,1 | 28.1 | 59.7 |
| 90 | sTEC | 1 | 0 | 40 mM K+, 5 mM Mg2+ | -51 to 51 | 3 | G3/K3 | 53K | 1.67 | 183 | EMD-47847 | 5,2 | 28.8 | 43.8 |
| 91 | sTEC | 1 | 0 | 40 mM K+, 5 mM Mg2+ | -51 to 51 | 3 | G3/K3 | 53K | 1.67 | 183 | EMD-47847 | 5,3 | 29.8 | 37.7 |
| 92 | P. Cas | 0 | 1 | 40 mM K+, 5 mM Mg2+ | -51 to 51 | 3 | G3/K3 | 53K | 1.67 | 183 | EMD-47849 | 1,1 | 27.1 | 47.2 |
| 93 | P. Cas | 0 | 1 | 40 mM K+, 5 mM Mg2+ | -51 to 51 | 3 | G3/K3 | 53K | 1.67 | 183 | EMD-47849 | 1,2 | 26.1 | 39.6 |
| 94 | P. Cas | 0 | 1 | 40 mM K+, 5 mM Mg2+ | -51 to 51 | 3 | G3/K3 | 53K | 1.67 | 183 | EMD-47849 | 1,3 | 25.7 | 42.7 |
| 95 | P. Cas | 0 | 1 | 40 mM K+, 5 mM Mg2+ | -51 to 51 | 3 | G3/K3 | 53K | 1.67 | 183 | EMD-47849 | 1,4 | 27.3 | 36.7 |
| 96 | P. Cas | 0 | 2 | 40 mM K+, 5 mM Mg2+ | -51 to 51 | 3 | G3/K3 | 53K | 1.67 | 183 | EMD-47849 | 1,5 | 28.1 | 31.7 |
| 97 | P. Cas | 0 | 1 | 40 mM K+, 5 mM Mg2+ | -51 to 51 | 3 | G3/K3 | 53K | 1.67 | 183 | EMD-47849 | 1,6 | 25.5 | 33.1 |
| 98 | P. Cas | 0 | 1 | 40 mM K+, 5 mM Mg2+ | -51 to 51 | 3 | G3/K3 | 53K | 1.67 | 183 | EMD-47849 | 1,7 | 26.3 | 40.5 |
| 99 | P. Cas | 0 | 1 | 40 mM K+, 5 mM Mg2+ | -51 to 51 | 3 | G3/K3 | 53K | 1.67 | 183 | EMD-47849 | 2,1 | 28.5 | 30.3 |
| 100 | P. Cas | 0 | 1 | 40 mM K+, 5 mM Mg2+ | -51 to 51 | 3 | G3/K3 | 53K | 1.67 | 183 | EMD-47849 | 2,2 | 26.6 | 41.4 |
| 101 | P. Cas | 0 | 1 | 40 mM K+, 5 mM Mg2+ | -51 to 51 | 3 | G3/K3 | 53K | 1.67 | 183 | EMD-47849 | 2,3 | 28 | 32.8 |
| 102 | P. Cas | 0 | 1 | 40 mM K+, 5 mM Mg2+ | -51 to 51 | 3 | G3/K3 | 53K | 1.67 | 183 | EMD-47849 | 2,4 | 27.2 | 31.6 |
| 103 | P. Cas | 0 | 1 | 40 mM K+, 5 mM Mg2+ | -51 to 51 | 3 | G3/K3 | 53K | 1.67 | 183 | EMD-47849 | 2,5 | 28.1 | 30.8 |
| 104 | P. Cas | 0 | 1 | 40 mM K+, 5 mM Mg2+ | -51 to 51 | 3 | G3/K3 | 53K | 1.67 | 183 | EMD-47849 | 2,6 | 27.9 | 32.9 |
| 105 | P. Cas | 0 | 1 | 40 mM K+, 5 mM Mg2+ | -51 to 51 | 3 | G3/K3 | 53K | 1.67 | 183 | EMD-47849 | 2,7 | 28.6 | 34.5 |
| 106 | P. Cas | 0 | 1 | 40 mM K+, 5 mM Mg2+ | -51 to 51 | 3 | G3/K3 | 53K | 1.67 | 183 | EMD-47849 | 3,1 | 26 | 32.1 |
| 107 | P. Cas | 0 | 2 | 40 mM K+, 5 mM Mg2+ | -51 to 51 | 3 | G3/K3 | 53K | 1.67 | 183 | EMD-47849 | 3,2 | 26.2 | 36.2 |
| 108 | P. Cas | 0 | 1 | 40 mM K+, 5 mM Mg2+ | -51 to 51 | 3 | G3/K3 | 53K | 1.67 | 183 | EMD-47849 | 3,3 | 27.1 | 31 |
| 109 | P. Cas | 0 | 1 | 40 mM K+, 5 mM Mg2+ | -51 to 51 | 3 | G3/K3 | 53K | 1.67 | 183 | EMD-47849 | 3,4 | 27.3 | 30.6 |
| 110 | P. Cas | 0 | 1 | 40 mM K+, 5 mM Mg2+ | -51 to 51 | 3 | G3/K3 | 53K | 1.67 | 183 | EMD-47849 | 3,5 | 27.7 | 32.6 |
| 111 | P. Cas | 0 | 1 | 40 mM K+, 5 mM Mg2+ | -51 to 51 | 3 | G3/K3 | 53K | 1.67 | 183 | EMD-47849 | 3,6 | 27.8 | 35.2 |
| 112 | P. Cas | 0 | 1 | 40 mM K+, 5 mM Mg2+ | -51 to 51 | 3 | G3/K3 | 53K | 1.67 | 183 | EMD-47849 | 3,7 | 28.2 | 30.9 |
| 113 | P. Cas | 0 | 1 | 40 mM K+, 5 mM Mg2+ | -51 to 51 | 3 | G3/K3 | 53K | 1.67 | 183 | EMD-47849 | 4,1 | 28.4 | 31.7 |
| 114 | P. Cas | 0 | 1 | 40 mM K+, 5 mM Mg2+ | -51 to 51 | 3 | G3/K3 | 53K | 1.67 | 183 | EMD-47849 | 4,2 | 28.3 | 30.2 |
| 115 | P. Cas | 0 | 1 | 40 mM K+, 5 mM Mg2+ | -51 to 51 | 3 | G3/K3 | 53K | 1.67 | 183 | EMD-47849 | 4,3 | 27.7 | 29 |
| 116 | P. Cas | 0 | 1 | 40 mM K+, 5 mM Mg2+ | -51 to 51 | 3 | G3/K3 | 53K | 1.67 | 183 | EMD-47849 | 4,4 | 27.8 | 28.8 |
| 117 | P. Cas | 0 | 1 | 40 mM K+, 5 mM Mg2+ | -51 to 51 | 3 | G3/K3 | 53K | 1.67 | 183 | EMD-47849 | 4,5 | 26.6 | 31.6 |
| 118 | P. Cas | 0 | 1 | 40 mM K+, 5 mM Mg2+ | -51 to 51 | 3 | G3/K3 | 53K | 1.67 | 183 | EMD-47849 | 4,6 | 26.9 | 30.9 |
| 119 | P. Cas | 0 | 1 | 40 mM K+, 5 mM Mg2+ | -51 to 51 | 3 | G3/K3 | 53K | 1.67 | 183 | EMD-47849 | 4,7 | 26.7 | 33 |
| 120 | P. Cas | 0 | 1 | 40 mM K+, 5 mM Mg2+ | -51 to 51 | 3 | G3/K3 | 53K | 1.67 | 183 | EMD-47849 | 5,1 | 28.8 | 30.8 |
| 121 | P. Cas | 0 | 1 | 40 mM K+, 5 mM Mg2+ | -51 to 51 | 3 | G3/K3 | 53K | 1.67 | 183 | EMD-47849 | 5,2 | 26.2 | 36.1 |
| 122 | P. Cas | 0 | 1 | 40 mM K+, 5 mM Mg2+ | -51 to 51 | 3 | G3/K3 | 53K | 1.67 | 183 | EMD-47849 | 5,3 | 25.8 | 34.6 |
| 123 | P. Cas | 0 | 1 | 40 mM K+, 5 mM Mg2+ | -51 to 51 | 3 | G3/K3 | 53K | 1.67 | 183 | EMD-47849 | 5,4 | 27.1 | 33 |
| 124 | P. Cas | 0 | 1 | 40 mM K+, 5 mM Mg2+ | -51 to 51 | 3 | G3/K3 | 53K | 1.67 | 183 | EMD-47849 | 5,5 | 26.2 | 32.4 |
| 125 | P. Cas | 0 | 1 | 40 mM K+, 5 mM Mg2+ | -51 to 51 | 3 | G3/K3 | 53K | 1.67 | 183 | EMD-47849 | 5,6 | 25.7 | 30.9 |
| 126 | TEC | 1 | 0 | 40 mM K+, 5 mM Mg2+ | -51 to 51 | 3 | G3/K3 | 53K | 1.67 | 183 | EMD-47850 | 1,1 | 27.7 | 37.2 |
| 127 | TEC | 1 | 0 | 40 mM K+, 5 mM Mg2+ | -51 to 51 | 3 | G3/K3 | 53K | 1.67 | 183 | EMD-47850 | 1,2 | 28.6 | 33.4 |
| 128 | TEC | 1 | 0 | 40 mM K+, 5 mM Mg2+ | -51 to 51 | 3 | G3/K3 | 53K | 1.67 | 183 | EMD-47850 | 1,3 | 28.3 | 34.6 |
| 129 | TEC | 1 | 0 | 40 mM K+, 5 mM Mg2+ | -51 to 51 | 3 | G3/K3 | 53K | 1.67 | 183 | EMD-47850 | 1,4 | 28.1 | 40.9 |
| 130 | TEC | 1 | 0 | 40 mM K+, 5 mM Mg2+ | -51 to 51 | 3 | G3/K3 | 53K | 1.67 | 183 | EMD-47850 | 1,5 | 27.5 | 44.3 |
| 131 | TEC | 1 | 0 | 40 mM K+, 5 mM Mg2+ | -51 to 51 | 3 | G3/K3 | 53K | 1.67 | 183 | EMD-47850 | 2,1 | 25.5 | 32.9 |
| 132 | TEC | 1 | 0 | 40 mM K+, 5 mM Mg2+ | -51 to 51 | 3 | G3/K3 | 53K | 1.67 | 183 | EMD-47850 | 2,2 | 26.9 | 34.6 |
| 133 | TEC | 1 | 0 | 40 mM K+, 5 mM Mg2+ | -51 to 51 | 3 | G3/K3 | 53K | 1.67 | 183 | EMD-47850 | 2,3 | 27.7 | 35.1 |
| 134 | TEC | 1 | 0 | 40 mM K+, 5 mM Mg2+ | -51 to 51 | 3 | G3/K3 | 53K | 1.67 | 183 | EMD-47850 | 2,4 | 27.9 | 34.7 |
| 135 | TEC | 2 | 0 | 40 mM K+, 5 mM Mg2+ | -51 to 51 | 3 | G3/K3 | 53K | 1.67 | 183 | EMD-47850 | 2,5 | 27.7 | 35.7 |
| 136 | TEC | 1 | 0 | 40 mM K+, 5 mM Mg2+ | -51 to 51 | 3 | G3/K3 | 53K | 1.67 | 183 | EMD-47850 | 3,1 | 26.5 | 32 |
| 137 | TEC | 1 | 0 | 40 mM K+, 5 mM Mg2+ | -51 to 51 | 3 | G3/K3 | 53K | 1.67 | 183 | EMD-47850 | 3,2 | 28.7 | 31.4 |
| 138 | TEC | 1 | 0 | 40 mM K+, 5 mM Mg2+ | -51 to 51 | 3 | G3/K3 | 53K | 1.67 | 183 | EMD-47850 | 3,3 | 27.3 | 33 |
| 139 | TEC | 1 | 0 | 40 mM K+, 5 mM Mg2+ | -51 to 51 | 3 | G3/K3 | 53K | 1.67 | 183 | EMD-47850 | 3,4 | 28.5 | 34.5 |
| 140 | TEC | 1 | 0 | 40 mM K+, 5 mM Mg2+ | -51 to 51 | 3 | G3/K3 | 53K | 1.67 | 183 | EMD-47850 | 3,5 | 26.9 | 33.3 |
| 141 | TEC | 1 | 0 | 40 mM K+, 5 mM Mg2+ | -51 to 51 | 3 | G3/K3 | 53K | 1.67 | 183 | EMD-47850 | 4,1 | 26.5 | 33.8 |
| 142 | TEC | 1 | 0 | 40 mM K+, 5 mM Mg2+ | -55 to 55 | 5 | G3/K3 | 53K | 1.67 | 120 | EMD-47850 | 4,2 | 27.4 | 35.6 |
| 143 | TEC | 1 | 0 | 40 mM K+, 5 mM Mg2+ | -51 to 51 | 3 | G3/K3 | 53K | 1.67 | 183 | EMD-47850 | 4,3 | 28 | 37.8 |
| 144 | TEC | 0 | 0 | 40 mM K+, 5 mM Mg2+ | -51 to 51 | 3 | G3/K3 | 53K | 1.67 | 183 | EMD-47850 | 4,4 | 27.9 | 36.4 |
| 145 | TEC | 2 | 0 | 40 mM K+, 5 mM Mg2+ | -51 to 51 | 3 | G3/K3 | 53K | 1.67 | 183 | EMD-47850 | 4,5 | 26.3 | 33.3 |
| 146 | TEC | 1 | 0 | 40 mM K+, 5 mM Mg2+ | -51 to 51 | 3 | G3/K3 | 53K | 1.67 | 183 | EMD-47850 | 5,1 | 26.5 | 34 |
| 147 | TEC | 1 | 0 | 40 mM K+, 5 mM Mg2+ | -51 to 51 | 3 | G3/K3 | 53K | 1.67 | 183 | EMD-47850 | 5,2 | 27.8 | 35 |
| 148 | TEC | 1 | 0 | 40 mM K+, 5 mM Mg2+ | -51 to 51 | 3 | G3/K3 | 53K | 1.67 | 183 | EMD-47850 | 5,3 | 29.3 | 33.3 |
| 149 | sTEC-Cas | 3 | 1 | 40 mM K+, 5 mM Mg2+ | -55 to 55 | 5 | G3/K3 | 53K | 1.67 | 120 | EMD-47851 | 1,1 | 26.8 | 45.5 |
| 150 | sTEC-Cas | 2 | 1 | 40 mM K+, 5 mM Mg2+ | -55 to 55 | 5 | G3/K3 | 53K | 1.67 | 120 | EMD-47851 | 1,2 | 27.8 | 41.2 |
| 151 | sTEC-Cas | 1 | 1 | 40 mM K+, 5 mM Mg2+ | -55 to 35 | 5 | G3/K3 | 53K | 1.67 | 99 | EMD-47851 | 1,3 | 25.4 | 45.4 |
| 152 | sTEC-Cas | 1 | 1 | 40 mM K+, 5 mM Mg2+ | -55 to 35 | 5 | G3/K3 | 53K | 1.67 | 99 | EMD-47851 | 1,4 | 26.5 | 39.8 |
| 153 | sTEC-Cas | 1 | 1 | 40 mM K+, 5 mM Mg2+ | -55 to 35 | 5 | G3/K3 | 53K | 1.67 | 99 | EMD-47851 | 1,5 | 25.7 | 45.7 |
| 154 | sTEC-Cas | 1 | 1 | 40 mM K+, 5 mM Mg2+ | -55 to 35 | 5 | G3/K3 | 53K | 1.67 | 99 | EMD-47851 | 1,6 | 26.3 | 43.5 |
| 155 | sTEC-Cas | 3 | 1 | 40 mM K+, 5 mM Mg2+ | -55 to 55 | 5 | G3/K3 | 53K | 1.67 | 120 | EMD-47851 | 1,7 | 26.1 | 58.8 |
| 156 | sTEC-Cas | 3 | 2 | 40 mM K+, 5 mM Mg2+ | -55 to 55 | 5 | G3/K3 | 53K | 1.67 | 120 | EMD-47851 | 2,1 | 25.8 | 57.4 |
| 157 | sTEC-Cas | 1 | 1 | 40 mM K+, 5 mM Mg2+ | -55 to 35 | 5 | G3/K3 | 53K | 1.67 | 99 | EMD-47851 | 2,2 | 24 | 39.7 |
| 158 | sTEC-Cas | 1 | 1 | 40 mM K+, 5 mM Mg2+ | -55 to 55 | 5 | G3/K3 | 53K | 1.67 | 120 | EMD-47851 | 2,3 | 27.4 | 46.3 |
| 159 | sTEC-Cas | 3 | 0 | 40 mM K+, 5 mM Mg2+ | -55 to 55 | 5 | G3/K3 | 53K | 1.67 | 120 | EMD-47851 | 2,4 | 28 | 54.3 |
| 160 | sTEC-Cas | 1 | 1 | 40 mM K+, 5 mM Mg2+ | -55 to 55 | 5 | G3/K3 | 53K | 1.67 | 120 | EMD-47851 | 2,5 | 26.7 | 46.6 |
| 161 | sTEC-Cas | 1 | 1 | 40 mM K+, 5 mM Mg2+ | -55 to 55 | 5 | G3/K3 | 53K | 1.67 | 120 | EMD-47851 | 2,6 | 26.6 | 40.5 |
| 162 | sTEC-Cas | 0 | 1 | 40 mM K+, 5 mM Mg2+ | -55 to 55 | 5 | G3/K3 | 53K | 1.67 | 120 | EMD-47851 | 2,7 | 26.6 | 34 |
| 163 | sTEC-Cas | 2 | 1 | 40 mM K+, 5 mM Mg2+ | -55 to 55 | 5 | G3/K3 | 53K | 1.67 | 120 | EMD-47851 | 3,1 | 27.9 | 48.3 |
| 164 | sTEC-Cas | 1 | 1 | 40 mM K+, 5 mM Mg2+ | -55 to 55 | 5 | G3/K3 | 53K | 1.67 | 120 | EMD-47851 | 3,2 | 26.9 | 38.4 |
| 165 | sTEC-Cas | 1 | 1 | 40 mM K+, 5 mM Mg2+ | -55 to 55 | 5 | G3/K3 | 53K | 1.67 | 120 | EMD-47851 | 3,3 | 26.5 | 47.6 |
| 166 | sTEC-Cas | 1 | 1 | 40 mM K+, 5 mM Mg2+ | -55 to 45 | 5 | G3/K3 | 53K | 1.67 | 109 | EMD-47851 | 3,4 | 27.4 | 34.5 |

| Par. id | Name abbr. | Bound RNAP | Bound dCas9 | Salt condition | Tilt range | Tilt step | TEM /Cam. | Mag. | Apix | Total dose | EMDB map ID | Montage position | Resolution (FSC=0.143) | Resolution (FSC=0.5) |
|---|---|---|---|---|---|---|---|---|---|---|---|---|---|---|
| # | | # | # | | (°) | (°) | | | (Å²) | (e-/Å²) | # | (row, col) | Even vs. Odd (Å) | Map vs. Model (Å) |
| 167 | sTEC-Cas | 1 | 1 | 40 mM K+, 5 mM Mg2+ | -55 to 55 | 5 | G3/K3 | 53K | 1.67 | 120 | EMD-47851 | 3,5 | 25.2 | 44.5 |
| 168 | sTEC-Cas | 3 | 1 | 40 mM K+, 5 mM Mg2+ | -55 to 45 | 5 | G3/K3 | 53K | 1.67 | 109 | EMD-47851 | 3,6 | 24.9 | 43.7 |
| 169 | sTEC-Cas | 2 | 1 | 40 mM K+, 5 mM Mg2+ | -55 to 45 | 5 | G3/K3 | 53K | 1.67 | 109 | EMD-47851 | 3,7 | 28.1 | 43.2 |
| 170 | sTEC-Cas | 1 | 1 | 40 mM K+, 5 mM Mg2+ | -55 to 40 | 5 | G3/K3 | 53K | 1.67 | 104 | EMD-47851 | 4,1 | 26.8 | 34.6 |
| 171 | sTEC-Cas | 1 | 1 | 40 mM K+, 5 mM Mg2+ | -55 to 40 | 5 | G3/K3 | 53K | 1.67 | 104 | EMD-47851 | 4,2 | 25.8 | 37.4 |
| 172 | sTEC-Cas | 1 | 1 | 40 mM K+, 5 mM Mg2+ | -55 to 30 | 5 | G3/K3 | 53K | 1.67 | 94 | EMD-47851 | 4,3 | 25.9 | 38.5 |
| 173 | sTEC-Cas | 3 | 1 | 40 mM K+, 5 mM Mg2+ | -55 to 45 | 5 | G3/K3 | 53K | 1.67 | 109 | EMD-47851 | 4,4 | 26.3 | 41.2 |
| 174 | sTEC-Cas | 1 | 1 | 40 mM K+, 5 mM Mg2+ | -55 to 45 | 5 | G3/K3 | 53K | 1.67 | 109 | EMD-47851 | 4,5 | 27.2 | 37.7 |
| 175 | sTEC-Cas | 1 | 1 | 40 mM K+, 5 mM Mg2+ | -55 to 30 | 5 | G3/K3 | 53K | 1.67 | 94 | EMD-47851 | 4,6 | 27.8 | 44.4 |
| 176 | sTEC-Cas | 1 | 1 | 40 mM K+, 5 mM Mg2+ | -55 to 45 | 5 | G3/K3 | 53K | 1.67 | 109 | EMD-47851 | 4,7 | 171.4 | 34.9 |
| 177 | sTEC-Cas | 1 | 1 | 40 mM K+, 5 mM Mg2+ | -55 to 55 | 5 | G3/K3 | 53K | 1.67 | 120 | EMD-47851 | 5,1 | 26.7 | 41 |
| 178 | sTEC-Cas | 1 | 1 | 40 mM K+, 5 mM Mg2+ | -55 to 55 | 5 | G3/K3 | 53K | 1.67 | 120 | EMD-47851 | 5,2 | 26.9 | 35.7 |
| 179 | sTEC-Cas | 1 | 1 | 40 mM K+, 5 mM Mg2+ | -55 to 55 | 5 | G3/K3 | 53K | 1.67 | 120 | EMD-47851 | 5,3 | 27 | 35.9 |
| 180 | sTEC-Cas | 2 | 1 | 40 mM K+, 5 mM Mg2+ | -55 to 35 | 5 | G3/K3 | 53K | 1.67 | 99 | EMD-47851 | 5,4 | 28.1 | 44.6 |
| 181 | sTEC-Cas | 1 | 1 | 40 mM K+, 5 mM Mg2+ | -55 to 35 | 5 | G3/K3 | 53K | 1.67 | 99 | EMD-47851 | 5,5 | 26.3 | 40.6 |
| 182 | sTEC-Cas | 1 | 1 | 40 mM K+, 5 mM Mg2+ | -55 to 45 | 5 | G3/K3 | 53K | 1.67 | 109 | EMD-47851 | 5,6 | 27.9 | 43 |
| 183 | TEC-Cas | 5 | 1 | 40 mM K+, 5 mM Mg2+ | -55 to 55 | 5 | G3/K3 | 53K | 1.67 | 120 | EMD-47853 | 1,1 | 27.1 | 46.6 |
| 184 | TEC-Cas | 1 | 1 | 40 mM K+, 5 mM Mg2+ | -55 to 55 | 5 | G3/K3 | 53K | 1.67 | 120 | EMD-47853 | 1,2 | 28 | 35.1 |
| 185 | TEC-Cas | 3 | 1 | 40 mM K+, 5 mM Mg2+ | -55 to 55 | 5 | G3/K3 | 53K | 1.67 | 120 | EMD-47853 | 1,3 | 29.1 | 40.5 |
| 186 | TEC-Cas | 3 | 1 | 40 mM K+, 5 mM Mg2+ | -55 to 55 | 5 | G3/K3 | 53K | 1.67 | 120 | EMD-47853 | 1,4 | 27.6 | 45.1 |
| 187 | TEC-Cas | 1 | 1 | 40 mM K+, 5 mM Mg2+ | -55 to 55 | 5 | G3/K3 | 53K | 1.67 | 120 | EMD-47853 | 1,5 | 28.1 | 39.7 |
| 188 | TEC-Cas | 3 | 1 | 40 mM K+, 5 mM Mg2+ | -55 to 55 | 5 | G3/K3 | 53K | 1.67 | 120 | EMD-47853 | 1,6 | 27.3 | 41.3 |
| 189 | TEC-Cas | 4 | 1 | 40 mM K+, 5 mM Mg2+ | -55 to 55 | 5 | G3/K3 | 53K | 1.67 | 120 | EMD-47853 | 1,7 | 27.2 | 37.4 |
| 190 | TEC-Cas | 1 | 1 | 40 mM K+, 5 mM Mg2+ | -55 to 55 | 5 | G3/K3 | 53K | 1.67 | 120 | EMD-47853 | 1,8 | 28 | 39.6 |
| 191 | TEC-Cas | 1 | 1 | 40 mM K+, 5 mM Mg2+ | -55 to 55 | 5 | G3/K3 | 53K | 1.67 | 120 | EMD-47853 | 1,9 | 27.4 | 41.1 |
| 192 | TEC-Cas | 1 | 1 | 40 mM K+, 5 mM Mg2+ | -50 to 40 | 5 | G3/K3 | 53K | 1.67 | 99 | EMD-47853 | 2,1 | 28.2 | 37.8 |
| 193 | TEC-Cas | 1 | 0 | 40 mM K+, 5 mM Mg2+ | -50 to 40 | 5 | G3/K3 | 53K | 1.67 | 99 | EMD-47853 | 2,2 | 27.6 | 38 |
| 194 | TEC-Cas | 1 | 1 | 40 mM K+, 5 mM Mg2+ | -50 to 40 | 5 | G3/K3 | 53K | 1.67 | 99 | EMD-47853 | 2,3 | 27.1 | 41 |
| 195 | TEC-Cas | 4 | 1 | 40 mM K+, 5 mM Mg2+ | -50 to 40 | 5 | G3/K3 | 53K | 1.67 | 99 | EMD-47853 | 2,4 | 27.5 | 44.8 |
| 196 | TEC-Cas | 1 | 1 | 40 mM K+, 5 mM Mg2+ | -50 to 40 | 5 | G3/K3 | 53K | 1.67 | 99 | EMD-47853 | 2,5 | 26.9 | 43.7 |
| 197 | TEC-Cas | 2 | 1 | 40 mM K+, 5 mM Mg2+ | -50 to 40 | 5 | G3/K3 | 53K | 1.67 | 99 | EMD-47853 | 2,6 | 26.3 | 39.8 |
| 198 | TEC-Cas | 2 | 1 | 40 mM K+, 5 mM Mg2+ | -50 to 40 | 5 | G3/K3 | 53K | 1.67 | 99 | EMD-47853 | 2,7 | 27 | 41.8 |
| 199 | TEC-Cas | 3 | 1 | 40 mM K+, 5 mM Mg2+ | -50 to 40 | 5 | G3/K3 | 53K | 1.67 | 99 | EMD-47853 | 2,8 | 28.6 | 49.2 |
| 200 | TEC-Cas | 1 | 1 | 40 mM K+, 5 mM Mg2+ | -55 to 55 | 5 | G3/K3 | 53K | 1.67 | 120 | EMD-47853 | 2,9 | 27.2 | 40.3 |
| 201 | TEC-Cas | 0 | 1 | 40 mM K+, 5 mM Mg2+ | -50 to 40 | 5 | G3/K3 | 53K | 1.67 | 99 | EMD-47853 | 3,1 | 29.3 | 40.4 |
| 202 | TEC-Cas | 3 | 1 | 40 mM K+, 5 mM Mg2+ | -50 to 40 | 5 | G3/K3 | 53K | 1.67 | 99 | EMD-47853 | 3,2 | 26.5 | 44.8 |
| 203 | TEC-Cas | 4 | 1 | 40 mM K+, 5 mM Mg2+ | -55 to 55 | 5 | G3/K3 | 53K | 1.67 | 120 | EMD-47853 | 3,3 | 27.4 | 47.7 |
| 204 | TEC-Cas | 3 | 1 | 40 mM K+, 5 mM Mg2+ | -55 to 55 | 5 | G3/K3 | 53K | 1.67 | 120 | EMD-47853 | 3,4 | 27.8 | 43.1 |
| 205 | TEC-Cas | 1 | 1 | 40 mM K+, 5 mM Mg2+ | -55 to 55 | 5 | G3/K3 | 53K | 1.67 | 120 | EMD-47853 | 3,5 | 26.1 | 33.7 |
| 206 | TEC-Cas | 3 | 1 | 40 mM K+, 5 mM Mg2+ | -55 to 55 | 5 | G3/K3 | 53K | 1.67 | 120 | EMD-47853 | 3,6 | 27.3 | 42.1 |
| 207 | TEC-Cas | 1 | 1 | 40 mM K+, 5 mM Mg2+ | -55 to 50 | 5 | G3/K3 | 53K | 1.67 | 115 | EMD-47853 | 3,7 | 28 | 37.2 |
| 208 | TEC-Cas | 1 | 1 | 40 mM K+, 5 mM Mg2+ | -55 to 50 | 5 | G3/K3 | 53K | 1.67 | 115 | EMD-47853 | 3,8 | 29.3 | 43.1 |
| 209 | TEC-Cas | 1 | 1 | 40 mM K+, 5 mM Mg2+ | -55 to 55 | 5 | G3/K3 | 53K | 1.67 | 120 | EMD-47853 | 3,9 | 27.8 | 47.5 |
| 210 | TEC-Cas | 2 | 0 | 40 mM K+, 5 mM Mg2+ | -55 to 55 | 5 | G3/K3 | 53K | 1.67 | 120 | EMD-47853 | 4,1 | 27.1 | 34.9 |
| 211 | TEC-Cas | 1 | 1 | 40 mM K+, 5 mM Mg2+ | -55 to 55 | 5 | G3/K3 | 53K | 1.67 | 120 | EMD-47853 | 4,2 | 26.7 | 39.2 |
| 212 | TEC-Cas | 2 | 1 | 40 mM K+, 5 mM Mg2+ | -55 to 55 | 5 | G3/K3 | 53K | 1.67 | 120 | EMD-47853 | 4,3 | 27.3 | 45 |
| 213 | TEC-Cas | 3 | 1 | 40 mM K+, 5 mM Mg2+ | -55 to 55 | 5 | G3/K3 | 53K | 1.67 | 120 | EMD-47853 | 4,4 | 28.2 | 50.9 |
| 214 | TEC-Cas | 1 | 1 | 40 mM K+, 5 mM Mg2+ | -55 to 55 | 5 | G3/K3 | 53K | 1.67 | 120 | EMD-47853 | 4,5 | 26.7 | 37.1 |
| 215 | TEC-Cas | 2 | 1 | 40 mM K+, 5 mM Mg2+ | -55 to 55 | 5 | G3/K3 | 53K | 1.67 | 120 | EMD-47853 | 4,6 | 28.7 | 36.8 |
| 216 | TEC-Cas | 3 | 1 | 40 mM K+, 5 mM Mg2+ | -55 to 55 | 5 | G3/K3 | 53K | 1.67 | 120 | EMD-47853 | 4,7 | 29 | 48.7 |
| 217 | TEC-Cas | 2 | 2 | 40 mM K+, 5 mM Mg2+ | -55 to 55 | 5 | G3/K3 | 53K | 1.67 | 120 | EMD-47853 | 4,8 | 27.3 | 39.8 |
| 218 | TEC-Cas | 1 | 1 | 40 mM K+, 5 mM Mg2+ | -55 to 55 | 5 | G3/K3 | 53K | 1.67 | 120 | EMD-47853 | 4,9 | 24.4 | 39.8 |
| 219 | TEC-Cas | 3 | 1 | 40 mM K+, 5 mM Mg2+ | -55 to 55 | 5 | G3/K3 | 53K | 1.67 | 120 | EMD-47853 | 5,1 | 29.2 | 48.6 |
| 220 | TEC-Cas | 2 | 1 | 40 mM K+, 5 mM Mg2+ | -55 to 55 | 5 | G3/K3 | 53K | 1.67 | 120 | EMD-47853 | 5,2 | 27.6 | 43.6 |
| 221 | TEC-Cas | 1 | 1 | 40 mM K+, 5 mM Mg2+ | -55 to 55 | 5 | G3/K3 | 53K | 1.67 | 120 | EMD-47853 | 5,3 | 27.7 | 41.2 |
| 222 | TEC-Cas | 1 | 1 | 40 mM K+, 5 mM Mg2+ | -55 to 55 | 5 | G3/K3 | 53K | 1.67 | 120 | EMD-47853 | 5,4 | 27.6 | 49.9 |
| 223 | TEC-Cas | 2 | 1 | 40 mM K+, 5 mM Mg2+ | -55 to 55 | 5 | G3/K3 | 53K | 1.67 | 120 | EMD-47853 | 5,5 | 27.6 | 35.9 |
| 224 | TEC-Cas | 2 | 1 | 40 mM K+, 5 mM Mg2+ | -55 to 55 | 5 | G3/K3 | 53K | 1.67 | 120 | EMD-47853 | 5,6 | 27.7 | 46.3 |
| 225 | TEC-Cas | 3 | 1 | 40 mM K+, 5 mM Mg2+ | -55 to 55 | 5 | G3/K3 | 53K | 1.67 | 120 | EMD-47853 | 5,7 | 28.2 | 47.4 |
| 226 | TEC-Cas | 3 | 1 | 40 mM K+, 5 mM Mg2+ | -55 to 55 | 5 | G3/K3 | 53K | 1.67 | 120 | EMD-47853 | 5,8 | 26.8 | 91.1 |
| 227 | TEC-TopI | 1 | 0 | 40 mM K+, 5 mM Mg2+ | -55 to 55 | 5 | G3/K3 | 53K | 1.67 | 120 | EMD-47855 | 1,1 | 29.9 | 36.6 |
| 228 | TEC-TopI | 1 | 0 | 40 mM K+, 5 mM Mg2+ | -55 to 55 | 5 | G3/K3 | 53K | 1.67 | 120 | EMD-47855 | 1,2 | 30 | 35.7 |
| 229 | TEC-TopI | 1 | 0 | 40 mM K+, 5 mM Mg2+ | -55 to 55 | 5 | G3/K3 | 53K | 1.67 | 120 | EMD-47855 | 1,3 | 31.9 | 34.5 |
| 230 | TEC-TopI | 1 | 0 | 40 mM K+, 5 mM Mg2+ | -55 to 55 | 5 | G3/K3 | 53K | 1.67 | 120 | EMD-47855 | 1,4 | 31.3 | 35.3 |
| 231 | TEC-TopI | 2 | 0 | 40 mM K+, 5 mM Mg2+ | -55 to 55 | 5 | G3/K3 | 53K | 1.67 | 120 | EMD-47855 | 1,5 | 29.7 | 603.5 |
| 232 | TEC-TopI | 1 | 0 | 40 mM K+, 5 mM Mg2+ | -55 to 55 | 5 | G3/K3 | 53K | 1.67 | 120 | EMD-47855 | 1,6 | 31 | 34 |
| 233 | TEC-TopI | 1 | 0 | 40 mM K+, 5 mM Mg2+ | -55 to 55 | 5 | G3/K3 | 53K | 1.67 | 120 | EMD-47855 | 1,7 | 31 | 38.2 |
| 234 | TEC-TopI | 1 | 0 | 40 mM K+, 5 mM Mg2+ | -55 to 55 | 5 | G3/K3 | 53K | 1.67 | 120 | EMD-47855 | 1,8 | 32.3 | 39.4 |
| 235 | TEC-TopI | 1 | 0 | 40 mM K+, 5 mM Mg2+ | -55 to 55 | 5 | G3/K3 | 53K | 1.67 | 120 | EMD-47855 | 2,1 | 31.5 | 34.2 |
| 236 | TEC-TopI | 1 | 0 | 40 mM K+, 5 mM Mg2+ | -55 to 55 | 5 | G3/K3 | 53K | 1.67 | 120 | EMD-47855 | 2,2 | 33 | 33.7 |
| 237 | TEC-TopI | 1 | 0 | 40 mM K+, 5 mM Mg2+ | -55 to 55 | 5 | G3/K3 | 53K | 1.67 | 120 | EMD-47855 | 2,3 | 30.8 | 33.6 |
| 238 | TEC-TopI | 1 | 0 | 40 mM K+, 5 mM Mg2+ | -55 to 55 | 5 | G3/K3 | 53K | 1.67 | 120 | EMD-47855 | 2,4 | 32.5 | 33.3 |
| 239 | TEC-TopI | 1 | 0 | 40 mM K+, 5 mM Mg2+ | -55 to 55 | 5 | G3/K3 | 53K | 1.67 | 120 | EMD-47855 | 2,5 | 31.8 | 36 |
| 240 | TEC-TopI | 1 | 0 | 40 mM K+, 5 mM Mg2+ | -55 to 55 | 5 | G3/K3 | 53K | 1.67 | 120 | EMD-47855 | 2,6 | 33.2 | 32.9 |
| 241 | TEC-TopI | 1 | 0 | 40 mM K+, 5 mM Mg2+ | -55 to 55 | 5 | G3/K3 | 53K | 1.67 | 120 | EMD-47855 | 2,7 | 31.4 | 39.4 |
| 242 | TEC-TopI | 0 | 0 | 40 mM K+, 5 mM Mg2+ | -55 to 55 | 5 | G3/K3 | 53K | 1.67 | 120 | EMD-47855 | 2,8 | 28.2 | 32.3 |
| 243 | TEC-TopI | 1 | 0 | 40 mM K+, 5 mM Mg2+ | -55 to 55 | 5 | G3/K3 | 53K | 1.67 | 120 | EMD-47855 | 3,1 | 33.2 | 34.8 |
| 244 | TEC-TopI | 2 | 0 | 40 mM K+, 5 mM Mg2+ | -55 to 55 | 5 | G3/K3 | 53K | 1.67 | 120 | EMD-47855 | 3,2 | 31.5 | 37 |
| 245 | TEC-TopI | 1 | 0 | 40 mM K+, 5 mM Mg2+ | -55 to 55 | 5 | G3/K3 | 53K | 1.67 | 120 | EMD-47855 | 3,3 | 29.1 | 41.8 |
| 246 | TEC-TopI | 1 | 0 | 40 mM K+, 5 mM Mg2+ | -55 to 55 | 5 | G3/K3 | 53K | 1.67 | 120 | EMD-47855 | 3,4 | 28.1 | 32.9 |
| 247 | TEC-TopI | 0 | 0 | 40 mM K+, 5 mM Mg2+ | -55 to 55 | 5 | G3/K3 | 53K | 1.67 | 120 | EMD-47855 | 3,5 | 32.7 | 32.9 |
| 248 | TEC-TopI | 1 | 0 | 40 mM K+, 5 mM Mg2+ | -55 to 55 | 5 | G3/K3 | 53K | 1.67 | 120 | EMD-47855 | 3,6 | 27.5 | 35.6 |
| 249 | TEC-TopI | 1 | 0 | 40 mM K+, 5 mM Mg2+ | -55 to 55 | 5 | G3/K3 | 53K | 1.67 | 120 | EMD-47855 | 3,7 | 29.1 | 37.4 |

| Par. id | Name abbr. | Bound RNAP | Bound dCas9 | Salt condition | Tilt range | Tilt step | TEM /Cam. | Mag. | Apix | Total dose | EMDB map ID | Montage position | Resolution (FSC=0.143) | Resolution (FSC=0.5) |
|---|---|---|---|---|---|---|---|---|---|---|---|---|---|---|
| # | | # | # | | (°) | (°) | | | (Å²) | (e-/Å²) | # | (row, col) | Even vs. Odd (Å) | Map vs. Model (Å) |
| 250 | TEC-TopI | 1 | 0 | 40 mM K+, 5 mM Mg2+ | -55 to 55 | 5 | G3/K3 | 53K | 1.67 | 120 | EMD-47855 | 3,8 | 28.2 | 39.3 |
| 251 | TEC-TopI | 1 | 0 | 40 mM K+, 5 mM Mg2+ | -55 to 55 | 5 | G3/K3 | 53K | 1.67 | 120 | EMD-47855 | 4,1 | 28 | 38.6 |
| 252 | TEC-TopI | 1 | 0 | 40 mM K+, 5 mM Mg2+ | -55 to 55 | 5 | G3/K3 | 53K | 1.67 | 120 | EMD-47855 | 4,2 | 29.4 | 38.8 |
| 253 | TEC-TopI | 1 | 0 | 40 mM K+, 5 mM Mg2+ | -55 to 55 | 5 | G3/K3 | 53K | 1.67 | 120 | EMD-47855 | 4,3 | 28.8 | 41 |
| 254 | TEC-TopI | 1 | 0 | 40 mM K+, 5 mM Mg2+ | -55 to 55 | 5 | G3/K3 | 53K | 1.67 | 120 | EMD-47855 | 4,4 | 30.7 | 35.4 |
| 255 | TEC-TopI | 1 | 0 | 40 mM K+, 5 mM Mg2+ | -55 to 55 | 5 | G3/K3 | 53K | 1.67 | 120 | EMD-47855 | 4,5 | 28.1 | 43.8 |
| 256 | TEC-TopI | 1 | 0 | 40 mM K+, 5 mM Mg2+ | -55 to 55 | 5 | G3/K3 | 53K | 1.67 | 120 | EMD-47855 | 4,6 | 30.6 | 39.3 |
| 257 | TEC-TopI | 1 | 0 | 40 mM K+, 5 mM Mg2+ | -55 to 55 | 5 | G3/K3 | 53K | 1.67 | 120 | EMD-47855 | 4,7 | 28.4 | 37.2 |
| 258 | TEC-TopI | 1 | 0 | 40 mM K+, 5 mM Mg2+ | -55 to 55 | 5 | G3/K3 | 53K | 1.67 | 120 | EMD-47855 | 4,8 | 29.2 | 38.7 |
| 259 | TEC-TopI | 2 | 0 | 40 mM K+, 5 mM Mg2+ | -55 to 55 | 5 | G3/K3 | 53K | 1.67 | 120 | EMD-47855 | 5,1 | 29.4 | 46.8 |
| 260 | TEC-TopI | 1 | 0 | 40 mM K+, 5 mM Mg2+ | -55 to 55 | 5 | G3/K3 | 53K | 1.67 | 120 | EMD-47855 | 5,2 | 30.9 | 34.1 |
| 261 | TEC-TopI | 1 | 0 | 40 mM K+, 5 mM Mg2+ | -55 to 55 | 5 | G3/K3 | 53K | 1.67 | 120 | EMD-47855 | 5,3 | 28.4 | 30.4 |
| 262 | TEC-TopI | 1 | 0 | 40 mM K+, 5 mM Mg2+ | -55 to 55 | 5 | G3/K3 | 53K | 1.67 | 120 | EMD-47855 | 5,4 | 30.2 | 33.7 |
| 263 | TEC-TopI | 1 | 0 | 40 mM K+, 5 mM Mg2+ | -55 to 55 | 5 | G3/K3 | 53K | 1.67 | 120 | EMD-47855 | 5,5 | 29.3 | 39.4 |
| 264 | TEC-TopI | 0 | 0 | 40 mM K+, 5 mM Mg2+ | -55 to 55 | 5 | G3/K3 | 53K | 1.67 | 120 | EMD-47855 | 5,6 | 32.1 | 35.6 |
| 265 | Opp.-TEC | 1 | 0 | 40 mM K+, 5 mM Mg2+ | -51 to 51 | 3 | G3/K3 | 53K | 1.67 | 183 | EMD-71622 | 1,1 | 27.2 | 38 |
| 266 | Opp.-TEC | 1 | 0 | 40 mM K+, 5 mM Mg2+ | -51 to 51 | 3 | G3/K3 | 53K | 1.67 | 183 | EMD-71622 | 1,2 | 25.8 | 39.3 |
| 267 | Opp.-TEC | 1 | 0 | 40 mM K+, 5 mM Mg2+ | -51 to 51 | 3 | G3/K3 | 53K | 1.67 | 183 | EMD-71622 | 1,3 | 25.5 | 38 |
| 268 | Opp.-TEC | 1 | 0 | 40 mM K+, 5 mM Mg2+ | -51 to 51 | 3 | G3/K3 | 53K | 1.67 | 183 | EMD-71622 | 1,4 | 26.7 | 44.3 |
| 269 | Opp.-TEC | 1 | 0 | 40 mM K+, 5 mM Mg2+ | -51 to 51 | 3 | G3/K3 | 53K | 1.67 | 183 | EMD-71622 | 1,5 | 25.4 | 43.9 |
| 270 | Opp.-TEC | 1 | 0 | 40 mM K+, 5 mM Mg2+ | -51 to 51 | 3 | G3/K3 | 53K | 1.67 | 183 | EMD-71622 | 1,6 | 27.1 | 37 |
| 271 | Opp.-TEC | 1 | 0 | 40 mM K+, 5 mM Mg2+ | -51 to 51 | 3 | G3/K3 | 53K | 1.67 | 183 | EMD-71622 | 1,7 | 26.8 | 38.6 |
| 272 | Opp.-TEC | 1 | 0 | 40 mM K+, 5 mM Mg2+ | -51 to 51 | 3 | G3/K3 | 53K | 1.67 | 183 | EMD-71622 | 1,8 | 26.6 | 36.7 |
| 273 | Opp.-TEC | 1 | 0 | 40 mM K+, 5 mM Mg2+ | -51 to 51 | 3 | G3/K3 | 53K | 1.67 | 183 | EMD-71622 | 2,1 | 26.6 | 38.1 |
| 274 | Opp.-TEC | 1 | 0 | 40 mM K+, 5 mM Mg2+ | -51 to 51 | 3 | G3/K3 | 53K | 1.67 | 183 | EMD-71622 | 2,2 | 29.2 | 39.7 |
| 275 | Opp.-TEC | 1 | 0 | 40 mM K+, 5 mM Mg2+ | -51 to 51 | 3 | G3/K3 | 53K | 1.67 | 183 | EMD-71622 | 2,3 | 26.3 | 34.9 |
| 276 | Opp.-TEC | 1 | 0 | 40 mM K+, 5 mM Mg2+ | -51 to 51 | 3 | G3/K3 | 53K | 1.67 | 183 | EMD-71622 | 2,4 | 24.1 | 35 |
| 277 | Opp.-TEC | 2 | 0 | 40 mM K+, 5 mM Mg2+ | -51 to 51 | 3 | G3/K3 | 53K | 1.67 | 183 | EMD-71622 | 2,5 | 25.5 | 37.6 |
| 278 | Opp.-TEC | 2 | 0 | 40 mM K+, 5 mM Mg2+ | -51 to 51 | 3 | G3/K3 | 53K | 1.67 | 183 | EMD-71622 | 2,6 | 24.2 | 37.1 |
| 279 | Opp.-TEC | 2 | 0 | 40 mM K+, 5 mM Mg2+ | -51 to 51 | 3 | G3/K3 | 53K | 1.67 | 183 | EMD-71622 | 2,7 | 27.3 | 38.9 |
| 280 | Opp.-TEC | 2 | 0 | 40 mM K+, 5 mM Mg2+ | -51 to 51 | 3 | G3/K3 | 53K | 1.67 | 183 | EMD-71622 | 2,8 | 26.8 | 38.7 |
| 281 | Opp.-TEC | 2 | 0 | 40 mM K+, 5 mM Mg2+ | -51 to 51 | 3 | G3/K3 | 53K | 1.67 | 183 | EMD-71622 | 3,1 | 29.2 | 35.6 |
| 282 | Opp.-TEC | 2 | 0 | 40 mM K+, 5 mM Mg2+ | -51 to 51 | 3 | G3/K3 | 53K | 1.67 | 183 | EMD-71622 | 3,2 | 29.4 | 40.9 |
| 283 | Opp.-TEC | 2 | 0 | 40 mM K+, 5 mM Mg2+ | -51 to 51 | 3 | G3/K3 | 53K | 1.67 | 183 | EMD-71622 | 3,3 | 26.7 | 42.7 |
| 284 | Opp.-TEC | 3 | 0 | 40 mM K+, 5 mM Mg2+ | -51 to 51 | 3 | G3/K3 | 53K | 1.67 | 183 | EMD-71622 | 3,4 | 26.8 | 36.7 |
| 285 | Opp.-TEC | 3 | 0 | 40 mM K+, 5 mM Mg2+ | -51 to 51 | 3 | G3/K3 | 53K | 1.67 | 183 | EMD-71622 | 3,5 | 26.6 | 37.8 |
| 286 | Opp.-TEC | 2 | 0 | 40 mM K+, 5 mM Mg2+ | -51 to 51 | 3 | G3/K3 | 53K | 1.67 | 183 | EMD-71622 | 3,6 | 26.6 | 46.7 |
| 287 | Opp.-TEC | 2 | 0 | 40 mM K+, 5 mM Mg2+ | -51 to 51 | 3 | G3/K3 | 53K | 1.67 | 183 | EMD-71622 | 3,7 | 26.7 | 34.5 |
| 288 | Opp.-TEC | 2 | 0 | 40 mM K+, 5 mM Mg2+ | -51 to 51 | 3 | G3/K3 | 53K | 1.67 | 183 | EMD-71622 | 3,8 | 25.7 | 36.2 |
| 289 | Opp.-TEC | 2 | 0 | 40 mM K+, 5 mM Mg2+ | -51 to 51 | 3 | G3/K3 | 53K | 1.67 | 183 | EMD-71622 | 4,1 | 26.7 | 44.6 |
| 290 | Opp.-TEC | 4 | 0 | 40 mM K+, 5 mM Mg2+ | -51 to 51 | 3 | G3/K3 | 53K | 1.67 | 183 | EMD-71622 | 4,2 | 25.8 | 40.3 |
| 291 | Opp.-TEC | 4 | 0 | 40 mM K+, 5 mM Mg2+ | -51 to 51 | 3 | G3/K3 | 53K | 1.67 | 183 | EMD-71622 | 4,3 | 26.9 | 40.1 |
| 292 | Opp.-TEC | 3 | 0 | 40 mM K+, 5 mM Mg2+ | -51 to 51 | 3 | G3/K3 | 53K | 1.67 | 183 | EMD-71622 | 4,4 | 26.9 | 38 |
| 293 | Opp.-TEC | 4 | 0 | 40 mM K+, 5 mM Mg2+ | -51 to 51 | 3 | G3/K3 | 53K | 1.67 | 183 | EMD-71622 | 4,5 | 27.1 | 38.3 |
| 294 | Opp.-TEC | 3 | 0 | 40 mM K+, 5 mM Mg2+ | -51 to 51 | 3 | G3/K3 | 53K | 1.67 | 183 | EMD-71622 | 4,6 | 28.6 | 42.3 |
| 295 | Opp.-TEC | 3 | 0 | 40 mM K+, 5 mM Mg2+ | -51 to 51 | 3 | G3/K3 | 53K | 1.67 | 183 | EMD-71622 | 4,7 | 26.4 | 37.8 |
| 296 | Opp.-TEC | 3 | 0 | 40 mM K+, 5 mM Mg2+ | -51 to 51 | 3 | G3/K3 | 53K | 1.67 | 183 | EMD-71622 | 4,8 | 26.7 | 39.5 |
| 297 | Opp.-TEC | 3 | 0 | 40 mM K+, 5 mM Mg2+ | -51 to 51 | 3 | G3/K3 | 53K | 1.67 | 183 | EMD-71622 | 5,1 | 26.6 | 35.5 |
| 298 | Opp.-TEC | 3 | 0 | 40 mM K+, 5 mM Mg2+ | -51 to 51 | 3 | G3/K3 | 53K | 1.67 | 183 | EMD-71622 | 5,2 | 26.7 | 35.6 |
| 299 | Opp.-TEC | 4 | 0 | 40 mM K+, 5 mM Mg2+ | -51 to 51 | 3 | G3/K3 | 53K | 1.67 | 183 | EMD-71622 | 5,3 | 26.6 | 40.3 |
| 300 | Opp.-TEC | 4 | 0 | 40 mM K+, 5 mM Mg2+ | -51 to 51 | 3 | G3/K3 | 53K | 1.67 | 183 | EMD-71622 | 5,4 | 25.6 | 39.2 |
| 301 | Opp.-TEC | 7 | 0 | 40 mM K+, 5 mM Mg2+ | -51 to 51 | 3 | G3/K3 | 53K | 1.67 | 183 | EMD-71622 | 5,5 | 26.8 | 39.2 |
| 302 | Tan.-TEC | 1 | 0 | 40 mM K+, 5 mM Mg2+ | -51 to 51 | 3 | G3/K3 | 53K | 1.67 | 183 | EMD-71618 | 1,1 | 26.7 | 38.2 |
| 303 | Tan.-TEC | 1 | 0 | 40 mM K+, 5 mM Mg2+ | -51 to 51 | 3 | G3/K3 | 53K | 1.67 | 183 | EMD-71618 | 1,2 | 25.4 | 35.7 |
| 304 | Tan.-TEC | 1 | 0 | 40 mM K+, 5 mM Mg2+ | -51 to 51 | 3 | G3/K3 | 53K | 1.67 | 183 | EMD-71618 | 1,3 | 26.9 | 41.4 |
| 305 | Tan.-TEC | 2 | 0 | 40 mM K+, 5 mM Mg2+ | -51 to 51 | 3 | G3/K3 | 53K | 1.67 | 183 | EMD-71618 | 1,4 | 26.8 | 37 |
| 306 | Tan.-TEC | 2 | 0 | 40 mM K+, 5 mM Mg2+ | -51 to 51 | 3 | G3/K3 | 53K | 1.67 | 183 | EMD-71618 | 1,5 | 29.1 | 40.7 |
| 307 | Tan.-TEC | 2 | 0 | 40 mM K+, 5 mM Mg2+ | -51 to 51 | 3 | G3/K3 | 53K | 1.67 | 183 | EMD-71618 | 1,6 | 25 | 38.3 |
| 308 | Tan.-TEC | 2 | 0 | 40 mM K+, 5 mM Mg2+ | -51 to 51 | 3 | G3/K3 | 53K | 1.67 | 183 | EMD-71618 | 2,1 | 26.4 | 38.6 |
| 309 | Tan.-TEC | 3 | 0 | 40 mM K+, 5 mM Mg2+ | -51 to 51 | 3 | G3/K3 | 53K | 1.67 | 183 | EMD-71618 | 2,2 | 25.7 | 39.2 |
| 310 | Tan.-TEC | 5 | 0 | 40 mM K+, 5 mM Mg2+ | -51 to 51 | 3 | G3/K3 | 53K | 1.67 | 183 | EMD-71618 | 2,3 | 25.7 | 42.6 |
| 311 | Tan.-TEC | 4 | 0 | 40 mM K+, 5 mM Mg2+ | -51 to 51 | 3 | G3/K3 | 53K | 1.67 | 183 | EMD-71618 | 2,4 | 28.6 | 40.7 |
| 312 | Tan.-TEC | 3 | 0 | 40 mM K+, 5 mM Mg2+ | -51 to 51 | 3 | G3/K3 | 53K | 1.67 | 183 | EMD-71618 | 2,5 | 26.6 | 36.4 |
| 313 | Tan.-TEC | 3 | 0 | 40 mM K+, 5 mM Mg2+ | -51 to 51 | 3 | G3/K3 | 53K | 1.67 | 183 | EMD-71618 | 2,6 | 26.8 | 42.4 |
| 314 | Tan.-TEC | 7 | 0 | 40 mM K+, 5 mM Mg2+ | -51 to 51 | 3 | G3/K3 | 53K | 1.67 | 183 | EMD-71618 | 3,1 | 28.5 | 41.7 |
| 315 | Tan.-TEC | 5 | 0 | 40 mM K+, 5 mM Mg2+ | -51 to 51 | 3 | G3/K3 | 53K | 1.67 | 183 | EMD-71618 | 3,2 | 26.1 | 42.3 |
| 316 | Tan.-TEC | 6 | 0 | 40 mM K+, 5 mM Mg2+ | -51 to 51 | 3 | G3/K3 | 53K | 1.67 | 183 | EMD-71618 | 3,3 | 28.5 | 37.6 |
| 317 | Tan.-TEC | 6 | 0 | 40 mM K+, 5 mM Mg2+ | -51 to 51 | 3 | G3/K3 | 53K | 1.67 | 183 | EMD-71618 | 3,4 | 25.3 | 41.9 |
| 318 | Tan.-TEC | 5 | 0 | 40 mM K+, 5 mM Mg2+ | -51 to 51 | 3 | G3/K3 | 53K | 1.67 | 183 | EMD-71618 | 3,5 | 28.7 | 39.9 |
| 319 | Tan.-TEC | 3 | 0 | 40 mM K+, 5 mM Mg2+ | -51 to 51 | 3 | G3/K3 | 53K | 1.67 | 183 | EMD-71618 | 3,6 | 27.9 | 31.4 |
| 320 | Tan.-TEC | 5 | 0 | 40 mM K+, 5 mM Mg2+ | -51 to 51 | 3 | G3/K3 | 53K | 1.67 | 183 | EMD-71618 | 4,1 | 26.7 | 36.6 |
| 321 | Tan.-TEC | 4 | 0 | 40 mM K+, 5 mM Mg2+ | -51 to 51 | 3 | G3/K3 | 53K | 1.67 | 183 | EMD-71618 | 4,2 | 26.6 | 42.1 |
| 322 | Tan.-TEC | 4 | 0 | 40 mM K+, 5 mM Mg2+ | -51 to 51 | 3 | G3/K3 | 53K | 1.67 | 183 | EMD-71618 | 4,3 | 26.7 | 34.9 |
| 323 | Tan.-TEC | 7 | 0 | 40 mM K+, 5 mM Mg2+ | -51 to 51 | 3 | G3/K3 | 53K | 1.67 | 183 | EMD-71618 | 4,4 | 25.9 | 40.5 |
| 324 | Tan.-TEC | 7 | 0 | 40 mM K+, 5 mM Mg2+ | -51 to 51 | 3 | G3/K3 | 53K | 1.67 | 183 | EMD-71618 | 4,5 | 28.3 | 41.4 |
| 325 | Tan.-TEC | 5 | 0 | 40 mM K+, 5 mM Mg2+ | -51 to 51 | 3 | G3/K3 | 53K | 1.67 | 183 | EMD-71618 | 4,6 | 26.6 | 42.7 |
| 326 | Tan.-TEC | 3 | 0 | 40 mM K+, 5 mM Mg2+ | -51 to 51 | 3 | G3/K3 | 53K | 1.67 | 183 | EMD-71618 | 5,1 | 26.9 | 37.6 |
| 327 | Tan.-TEC | 4 | 0 | 40 mM K+, 5 mM Mg2+ | -51 to 51 | 3 | G3/K3 | 53K | 1.67 | 183 | EMD-71618 | 5,2 | 24.4 | 42.9 |
| 328 | Tan.-TEC | 9 | 0 | 40 mM K+, 5 mM Mg2+ | -51 to 51 | 3 | G3/K3 | 53K | 1.67 | 183 | EMD-71618 | 5,3 | 26.5 | 42.9 |
| 329 | Tan.-TEC | 5 | 0 | 40 mM K+, 5 mM Mg2+ | -51 to 51 | 3 | G3/K3 | 53K | 1.67 | 183 | EMD-71618 | 5,4 | 26.7 | 38.1 |
| 330 | Tan.-TEC | 5 | 0 | 40 mM K+, 5 mM Mg2+ | -51 to 51 | 3 | G3/K3 | 53K | 1.67 | 183 | EMD-71618 | 5,5 | 28.7 | 38 |
| 331 | Tan.-TEC | 4 | 0 | 40 mM K+, 5 mM Mg2+ | -51 to 51 | 3 | G3/K3 | 53K | 1.67 | 183 | EMD-71618 | 5,6 | 29.2 | 42.1 |

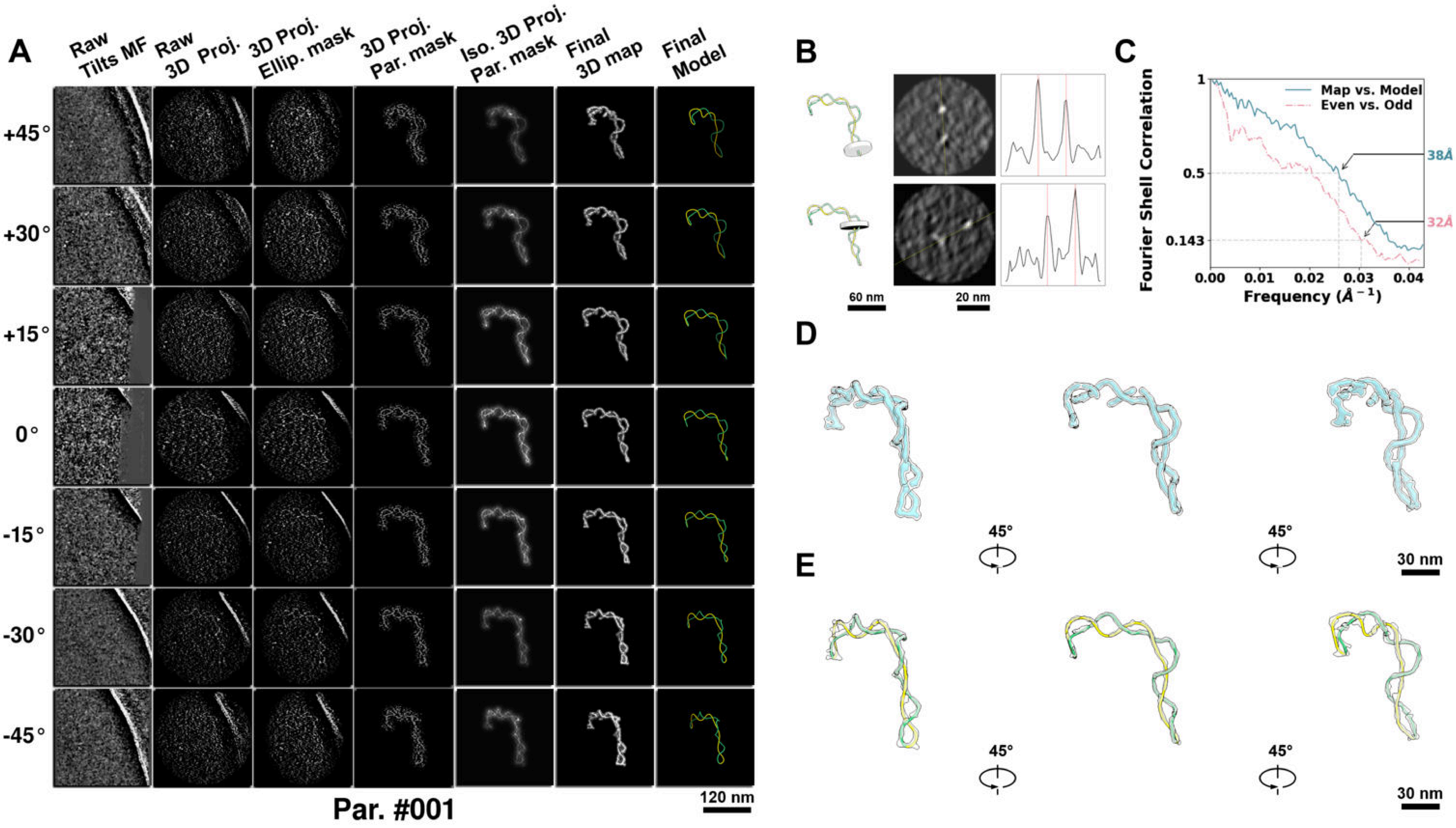

**Supplementary Particle Figure 1. Cryo-ET 3D reconstruction of an individual P.LS particle.**
(A) 3D reconstruction of the plasmid particle (index no. 1). The first column shows seven representative tilt images from +45° to -45° in step of 15°. The second, third, and fourth columns show 3D projections of the particle with spherical, ellipsoidal (thinner along the z-dimension), and particle-shaped masks, respectively. The fifth column displays the 3D projections of the enhanced and IsoNet missing-wedge-corrected particle. The sixth and seventh columns present the final 3D map and the flexibly fitted model, respectively. (B) Two cross-sectional views (12 nm thickness) of the plasmid density map along its plectoneme axis are shown in the left-middle panel. The intensity profile along the line crossing the two high-density DNA spots is displayed in the right panel. (C) Resolution assessment of the final 3D map using Fourier shell correlation (FSC). Two criteria are shown: FSC between two half-maps reconstructed from even and odd frames (evaluated at 0.143) and FSC between the final 3D map and the fitted model (evaluated at 0.5). (D) Zoomed-in views of the final 3D density map from panel A, displayed at two contour levels. (E) Superimposition of the high-contour level map from panel D onto its fitted model.

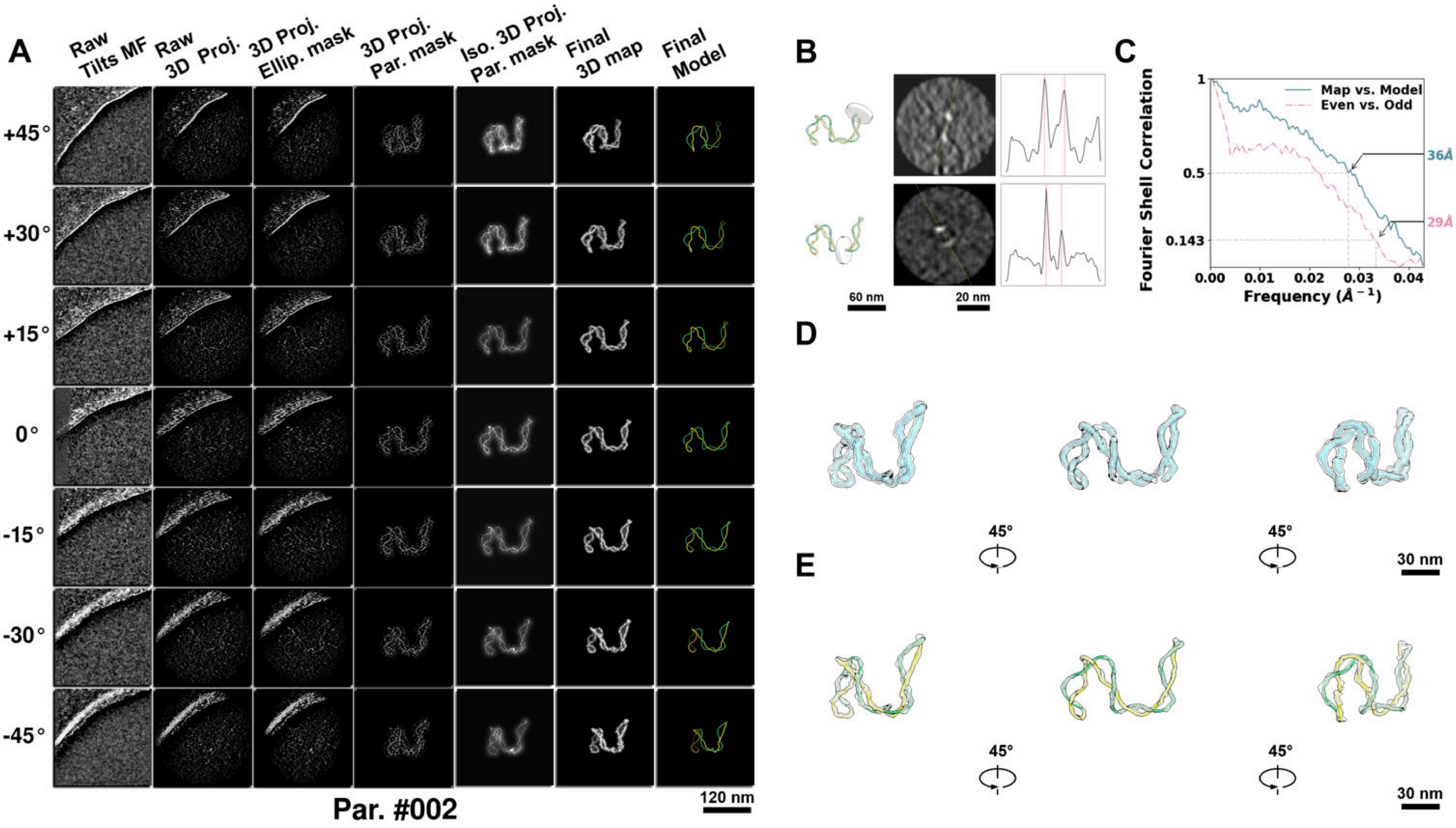

**Supplementary Particle Figure 2. Cryo-ET 3D reconstruction of an individual P.LS particle.**
(A) 3D reconstruction of the plasmid particle (index no. 2). The first column shows seven representative tilt images from +45° to -45° in step of 15°. The second, third, and fourth columns show 3D projections of the particle with spherical, ellipsoidal (thinner along the z-dimension), and particle-shaped masks, respectively. The fifth column displays the 3D projections of the enhanced and IsoNet missing-wedge-corrected particle. The sixth and seventh columns present the final 3D map and the flexibly fitted model, respectively. (B) Two cross-sectional views (12 nm thickness) of the plasmid density map along its plectoneme axis are shown in the left-middle panel. The intensity profile along the line crossing the two high-density DNA spots is displayed in the right panel. (C) Resolution assessment of the final 3D map using Fourier shell correlation (FSC). Two criteria are shown: FSC between two half-maps reconstructed from even and odd frames (evaluated at 0.143) and FSC between the final 3D map and the fitted model (evaluated at 0.5). (D) Zoomed-in views of the final 3D density map from panel A, displayed at two contour levels. (E) Superimposition of the high-contour level map from panel D onto its fitted model.

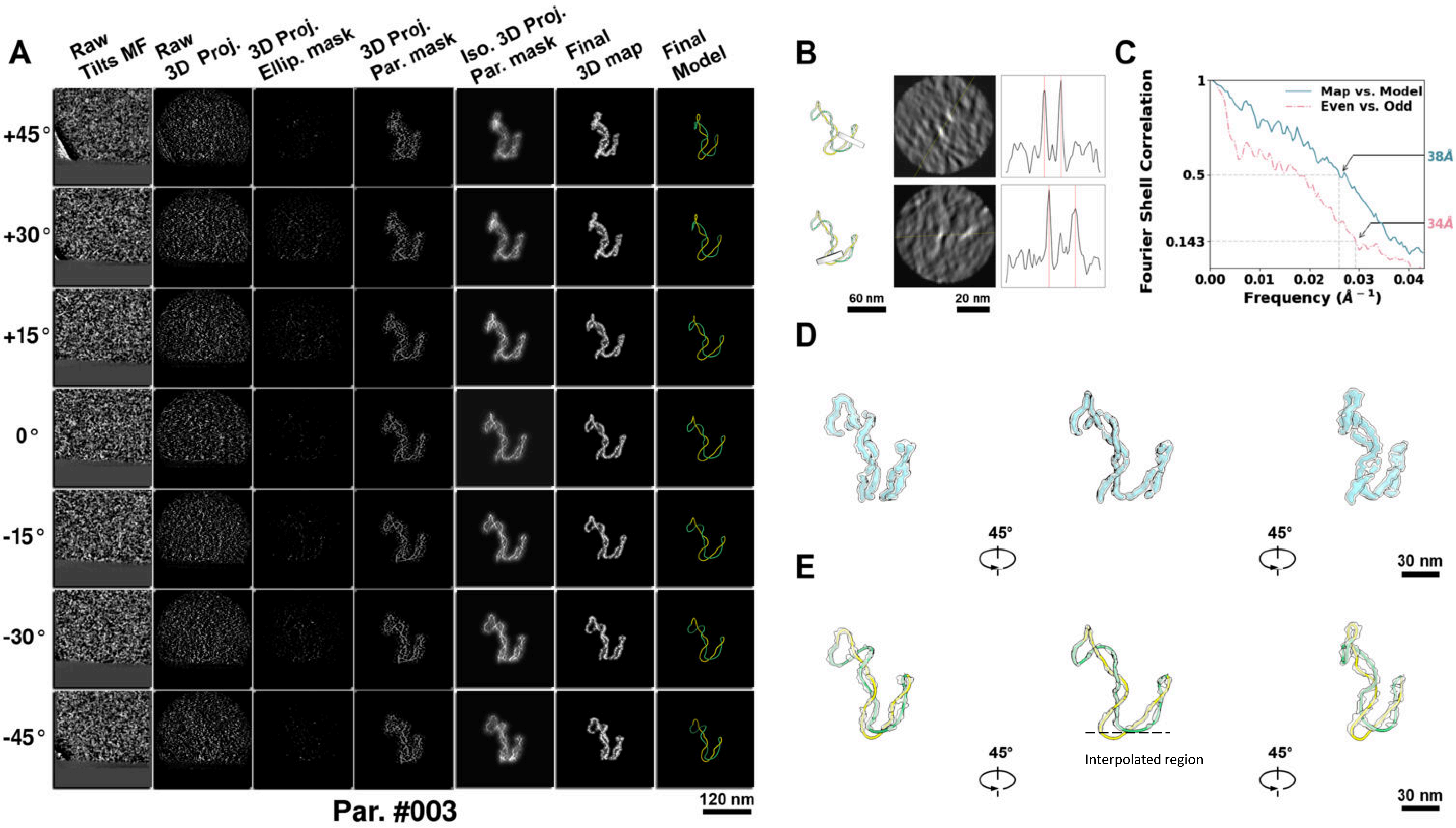

**Supplementary Particle Figure 3. Cryo-ET 3D reconstruction of an individual P.LS particle.**
(A) 3D reconstruction of the plasmid particle (index no. 3). The first column shows seven representative tilt images from +45° to -45° in step of 15°. The second, third, and fourth columns show 3D projections of the particle with spherical, ellipsoidal (thinner along the z-dimension), and particle-shaped masks, respectively. The fifth column displays the 3D projections of the enhanced and IsoNet missing-wedge-corrected particle. The sixth and seventh columns present the final 3D map and the flexibly fitted model, respectively. (B) Two cross-sectional views (12 nm thickness) of the plasmid density map along its plectoneme axis are shown in the left-middle panel. The intensity profile along the line crossing the two high-density DNA spots is displayed in the right panel. (C) Resolution assessment of the final 3D map using Fourier shell correlation (FSC). Two criteria are shown: FSC between two half-maps reconstructed from even and odd frames (evaluated at 0.143) and FSC between the final 3D map and the fitted model (evaluated at 0.5). (D) Zoomed-in views of the final 3D density map from panel A, displayed at two contour levels. (E) Superimposition of the high-contour level map from panel D onto its fitted model.

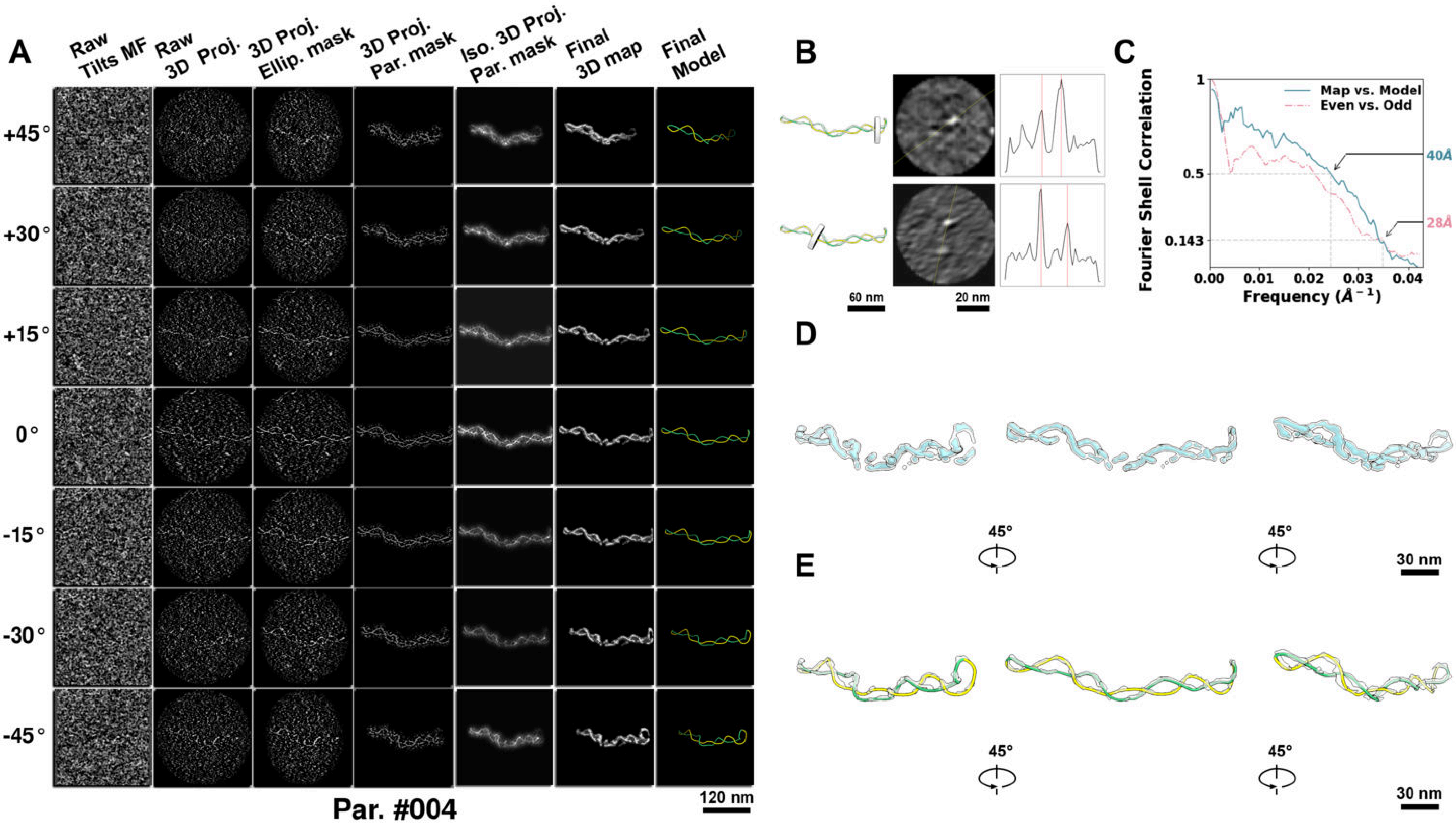

**Supplementary Particle Figure 4. Cryo-ET 3D reconstruction of an individual P.LS particle.**
(A) 3D reconstruction of the plasmid particle (index no. 4). The first column shows seven representative tilt images from +45° to -45° in step of 15°. The second, third, and fourth columns show 3D projections of the particle with spherical, ellipsoidal (thinner along the z-dimension), and particle-shaped masks, respectively. The fifth column displays the 3D projections of the enhanced and IsoNet missing-wedge-corrected particle. The sixth and seventh columns present the final 3D map and the flexibly fitted model, respectively. (B) Two cross-sectional views (12 nm thickness) of the plasmid density map along its plectoneme axis are shown in the left-middle panel. The intensity profile along the line crossing the two high-density DNA spots is displayed in the right panel. (C) Resolution assessment of the final 3D map using Fourier shell correlation (FSC). Two criteria are shown: FSC between two half-maps reconstructed from even and odd frames (evaluated at 0.143) and FSC between the final 3D map and the fitted model (evaluated at 0.5). (D) Zoomed-in views of the final 3D density map from panel A, displayed at two contour levels. (E) Superimposition of the high-contour level map from panel D onto its fitted model.

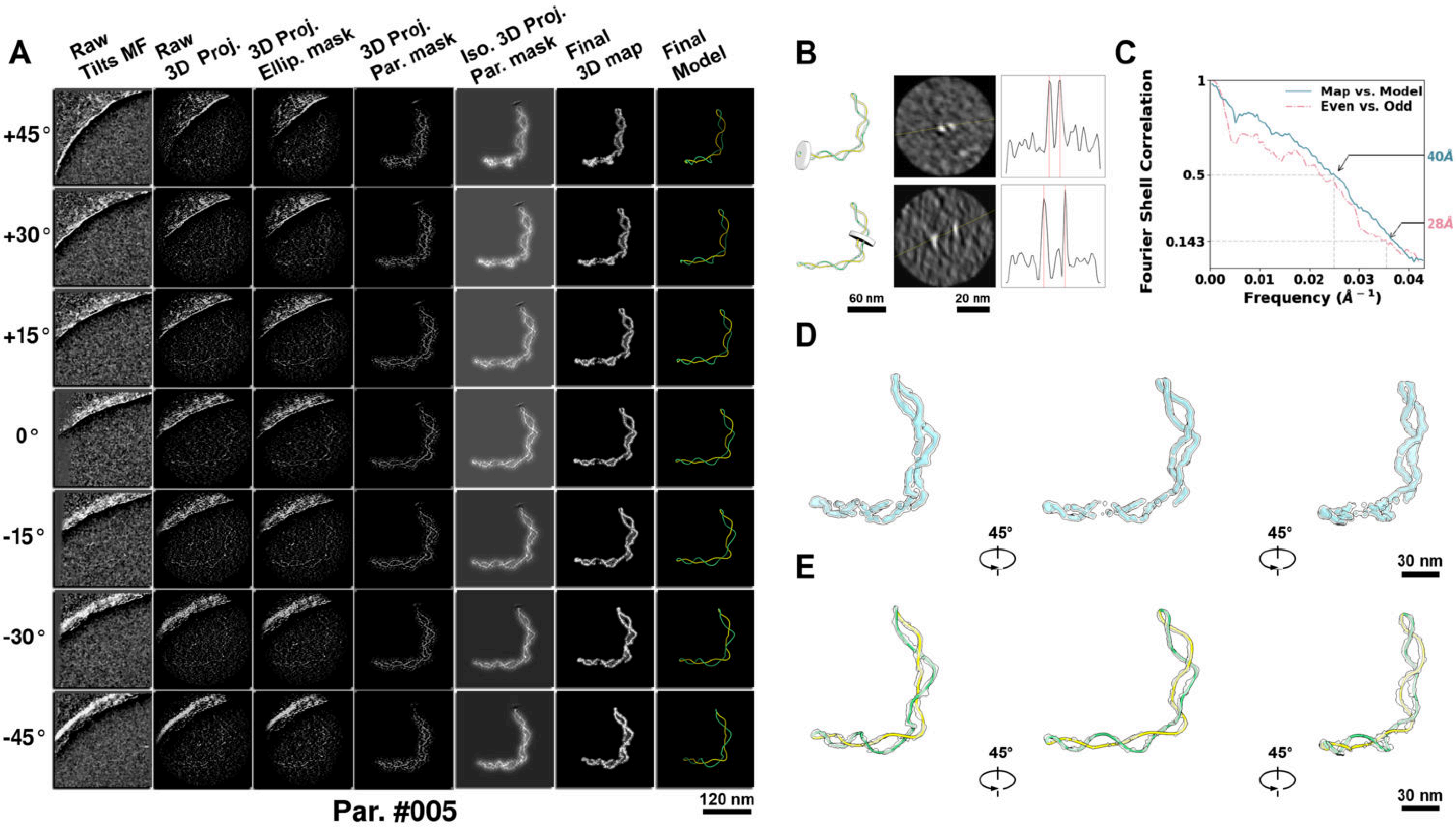

**Supplementary Particle Figure 5. Cryo-ET 3D reconstruction of an individual P.LS particle.**
(A) 3D reconstruction of the plasmid particle (index no. 5). The first column shows seven representative tilt images from +45° to -45° in step of 15°. The second, third, and fourth columns show 3D projections of the particle with spherical, ellipsoidal (thinner along the z-dimension), and particle-shaped masks, respectively. The fifth column displays the 3D projections of the enhanced and IsoNet missing-wedge-corrected particle. The sixth and seventh columns present the final 3D map and the flexibly fitted model, respectively. (B) Two cross-sectional views (12 nm thickness) of the plasmid density map along its plectoneme axis are shown in the left-middle panel. The intensity profile along the line crossing the two high-density DNA spots is displayed in the right panel. (C) Resolution assessment of the final 3D map using Fourier shell correlation (FSC). Two criteria are shown: FSC between two half-maps reconstructed from even and odd frames (evaluated at 0.143) and FSC between the final 3D map and the fitted model (evaluated at 0.5). (D) Zoomed-in views of the final 3D density map from panel A, displayed at two contour levels. (E) Superimposition of the high-contour level map from panel D onto its fitted model.

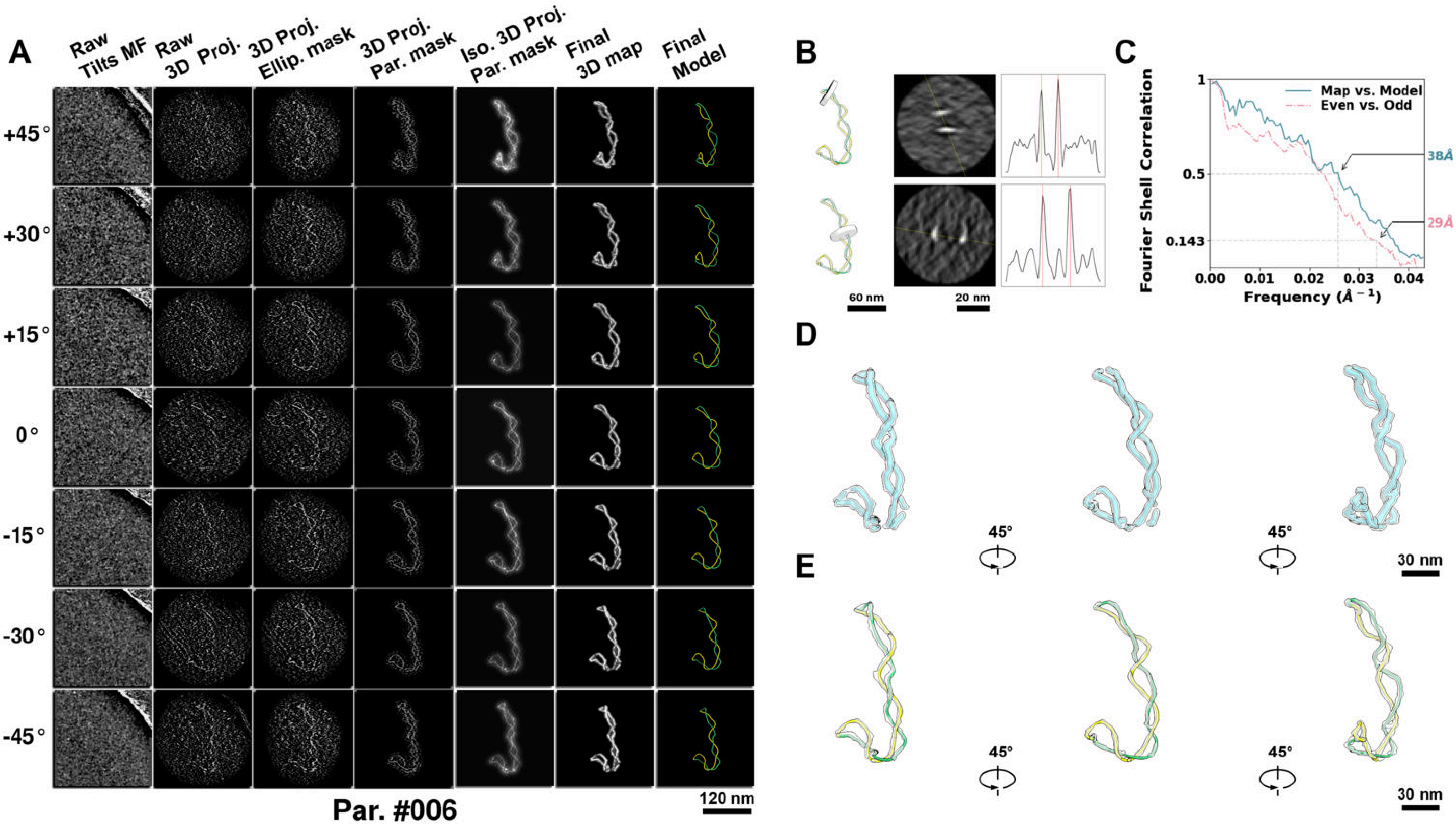

**Supplementary Particle Figure 6. Cryo-ET 3D reconstruction of an individual P.LS particle.**
(A) 3D reconstruction of the plasmid particle (index no. 6). The first column shows seven representative tilt images from +45° to -45° in step of 15°. The second, third, and fourth columns show 3D projections of the particle with spherical, ellipsoidal (thinner along the z-dimension), and particle-shaped masks, respectively. The fifth column displays the 3D projections of the enhanced and IsoNet missing-wedge-corrected particle. The sixth and seventh columns present the final 3D map and the flexibly fitted model, respectively. (B) Two cross-sectional views (12 nm thickness) of the plasmid density map along its plectoneme axis are shown in the left-middle panel. The intensity profile along the line crossing the two high-density DNA spots is displayed in the right panel. (C) Resolution assessment of the final 3D map using Fourier shell correlation (FSC). Two criteria are shown: FSC between two half-maps reconstructed from even and odd frames (evaluated at 0.143) and FSC between the final 3D map and the fitted model (evaluated at 0.5). (D) Zoomed-in views of the final 3D density map from panel A, displayed at two contour levels. (E) Superimposition of the high-contour level map from panel D onto its fitted model.

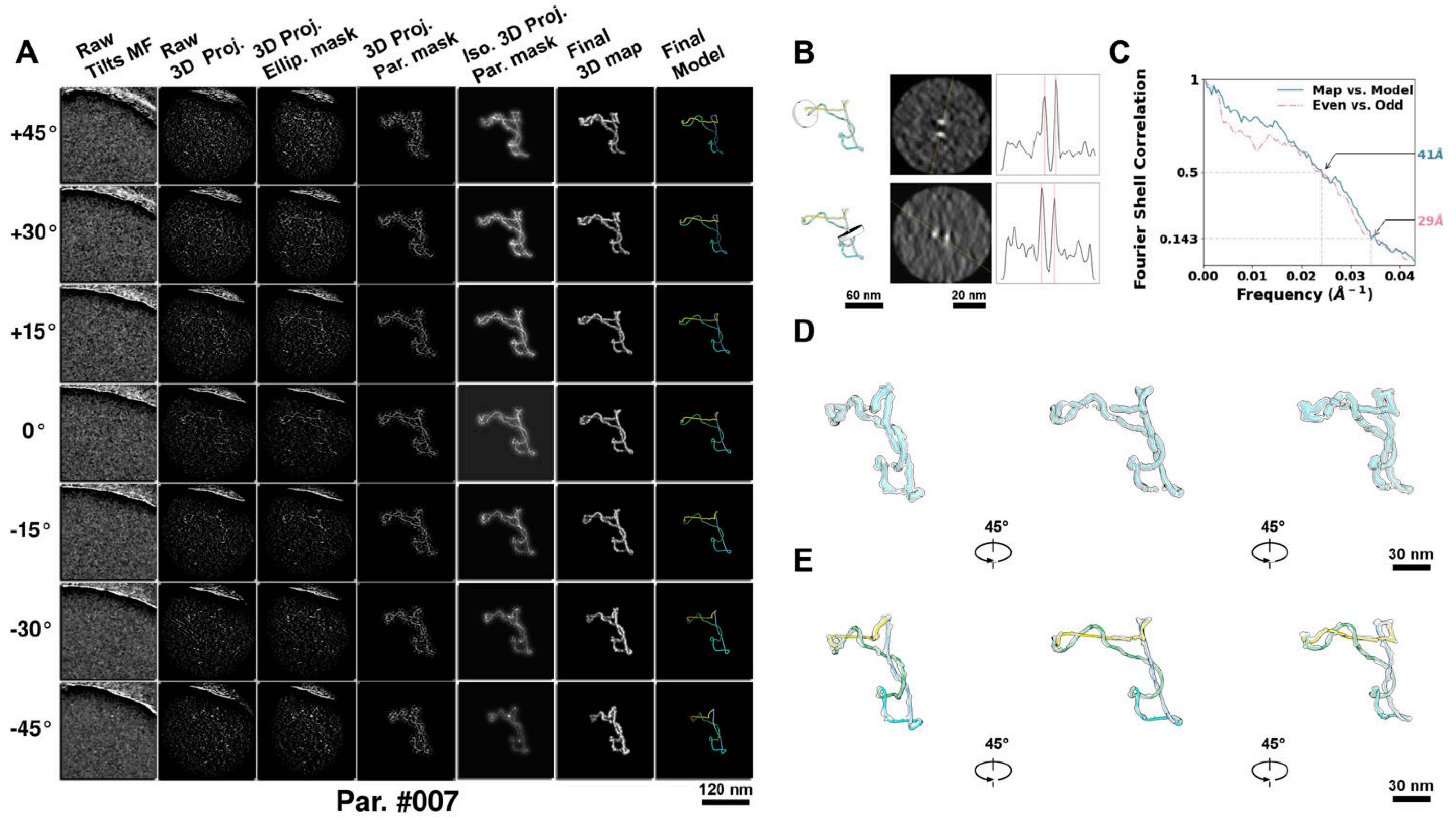

**Supplementary Particle Figure 7. Cryo-ET 3D reconstruction of an individual P.LS particle.**
(A) 3D reconstruction of the plasmid particle (index no. 7). The first column shows seven representative tilt images from +45° to -45° in step of 15°. The second, third, and fourth columns show 3D projections of the particle with spherical, ellipsoidal (thinner along the z-dimension), and particle-shaped masks, respectively. The fifth column displays the 3D projections of the enhanced and IsoNet missing-wedge-corrected particle. The sixth and seventh columns present the final 3D map and the flexibly fitted model, respectively. (B) Two cross-sectional views (12 nm thickness) of the plasmid density map along its plectoneme axis are shown in the left-middle panel. The intensity profile along the line crossing the two high-density DNA spots is displayed in the right panel. (C) Resolution assessment of the final 3D map using Fourier shell correlation (FSC). Two criteria are shown: FSC between two half-maps reconstructed from even and odd frames (evaluated at 0.143) and FSC between the final 3D map and the fitted model (evaluated at 0.5). (D) Zoomed-in views of the final 3D density map from panel A, displayed at two contour levels. (E) Superimposition of the high-contour level map from panel D onto its fitted model.

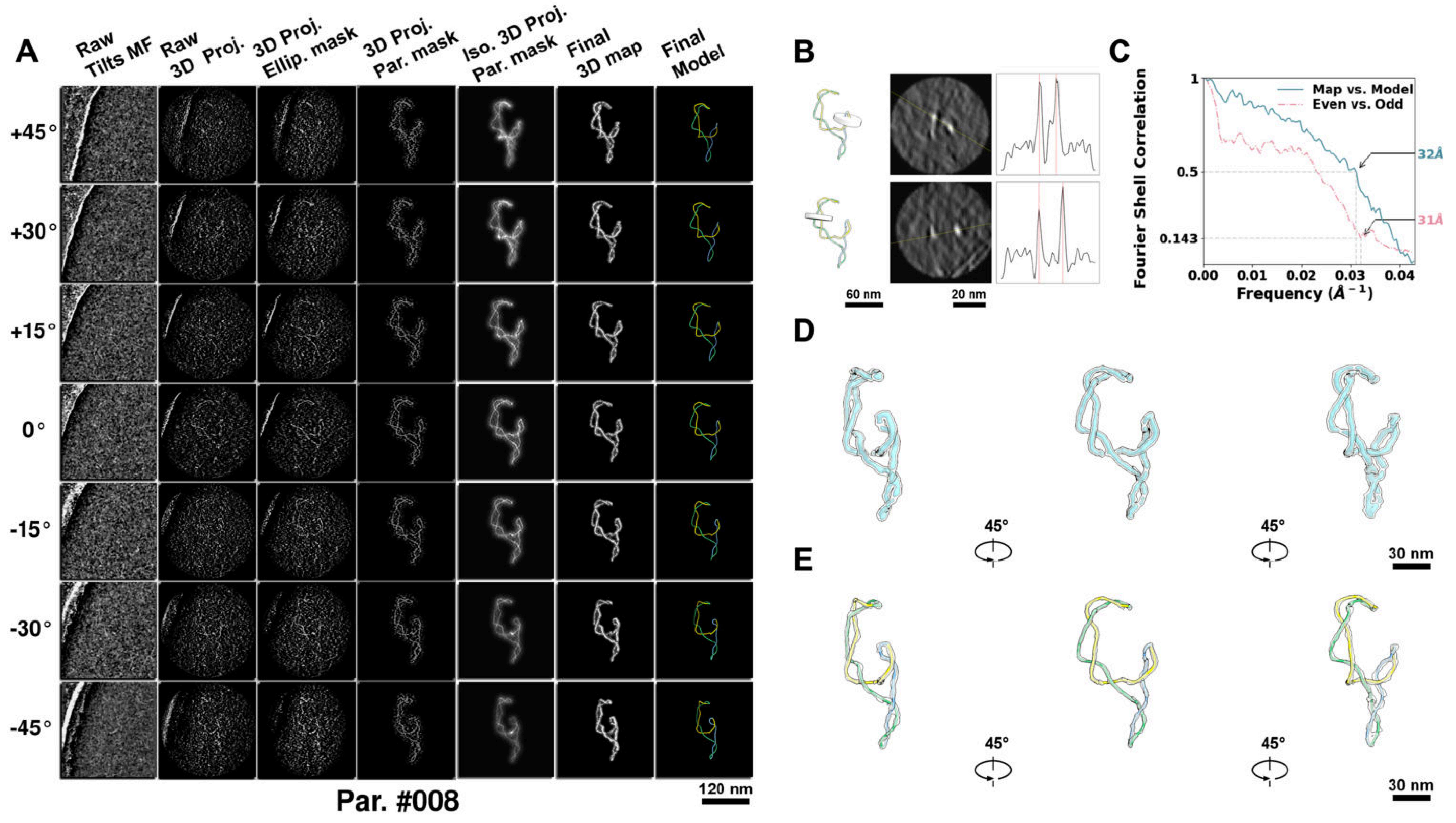

**Supplementary Particle Figure 8. Cryo-ET 3D reconstruction of an individual P.LS particle.**
(A) 3D reconstruction of the plasmid particle (index no. 8). The first column shows seven representative tilt images from +45° to -45° in step of 15°. The second, third, and fourth columns show 3D projections of the particle with spherical, ellipsoidal (thinner along the z-dimension), and particle-shaped masks, respectively. The fifth column displays the 3D projections of the enhanced and IsoNet missing-wedge-corrected particle. The sixth and seventh columns present the final 3D map and the flexibly fitted model, respectively. (B) Two cross-sectional views (12 nm thickness) of the plasmid density map along its plectoneme axis are shown in the left-middle panel. The intensity profile along the line crossing the two high-density DNA spots is displayed in the right panel. (C) Resolution assessment of the final 3D map using Fourier shell correlation (FSC). Two criteria are shown: FSC between two half-maps reconstructed from even and odd frames (evaluated at 0.143) and FSC between the final 3D map and the fitted model (evaluated at 0.5). (D) Zoomed-in views of the final 3D density map from panel A, displayed at two contour levels. (E) Superimposition of the high-contour level map from panel D onto its fitted model.

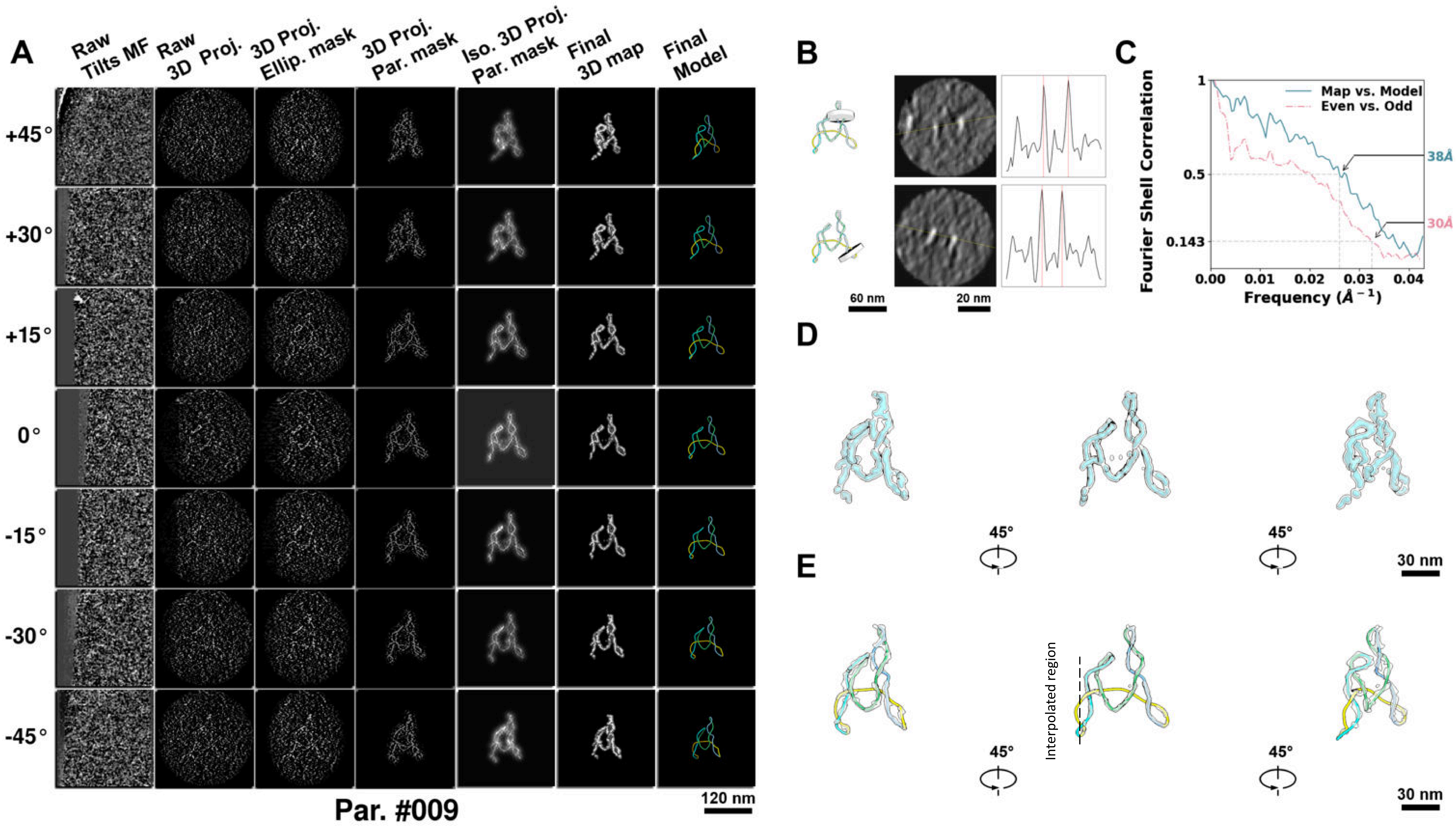

**Supplementary Particle Figure 9. Cryo-ET 3D reconstruction of an individual P.LS particle.**
(A) 3D reconstruction of the plasmid particle (index no. 9). The first column shows seven representative tilt images from +45° to -45° in step of 15°. The second, third, and fourth columns show 3D projections of the particle with spherical, ellipsoidal (thinner along the z-dimension), and particle-shaped masks, respectively. The fifth column displays the 3D projections of the enhanced and IsoNet missing-wedge-corrected particle. The sixth and seventh columns present the final 3D map and the flexibly fitted model, respectively. (B) Two cross-sectional views (12 nm thickness) of the plasmid density map along its plectoneme axis are shown in the left-middle panel. The intensity profile along the line crossing the two high-density DNA spots is displayed in the right panel. (C) Resolution assessment of the final 3D map using Fourier shell correlation (FSC). Two criteria are shown: FSC between two half-maps reconstructed from even and odd frames (evaluated at 0.143) and FSC between the final 3D map and the fitted model (evaluated at 0.5). (D) Zoomed-in views of the final 3D density map from panel A, displayed at two contour levels. (E) Superimposition of the high-contour level map from panel D onto its fitted model.

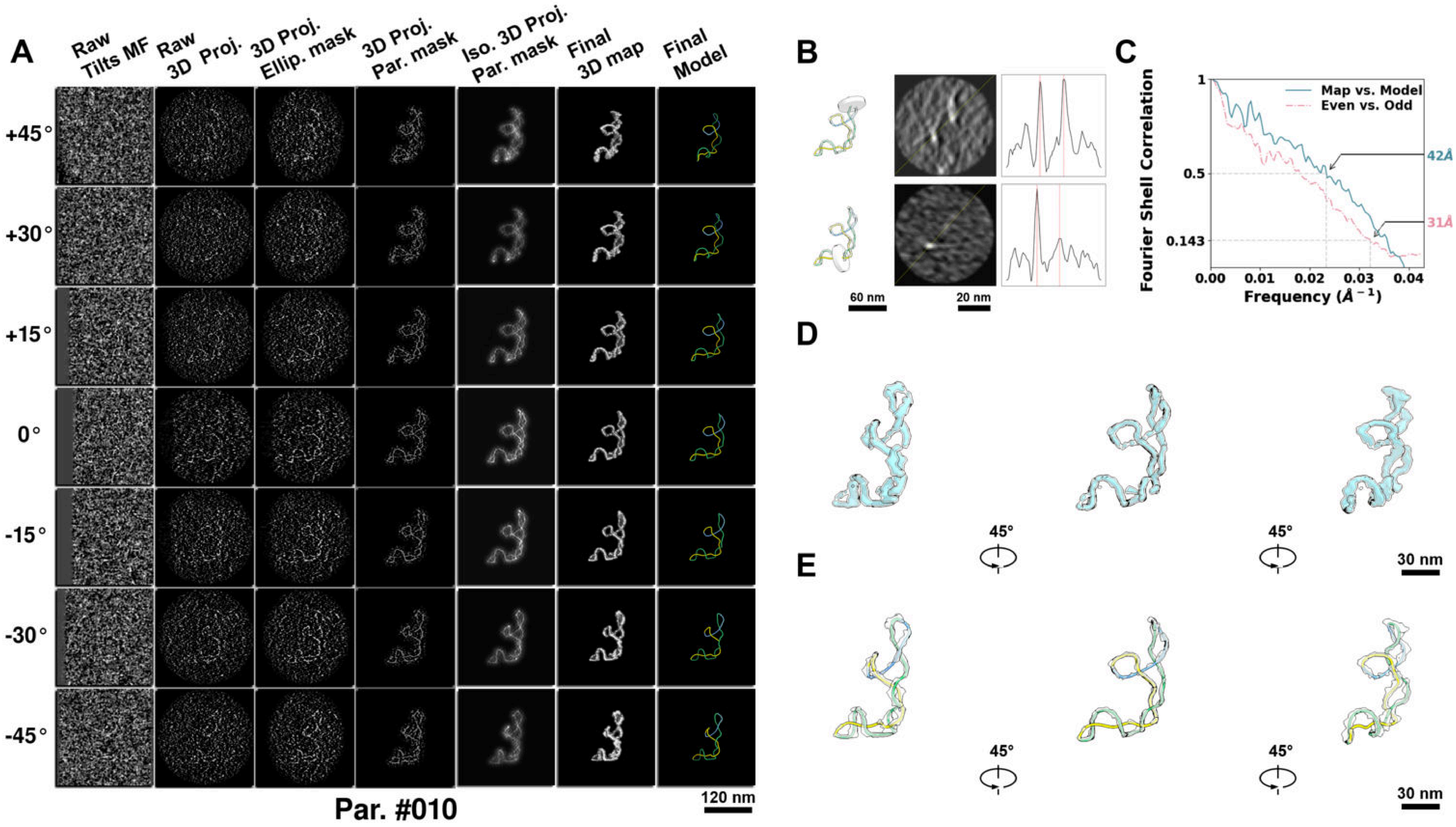

**Supplementary Particle Figure 10. Cryo-ET 3D reconstruction of an individual P.LS particle.**
(A) 3D reconstruction of the plasmid particle (index no. 10). The first column shows seven representative tilt images from +45° to -45° in step of 15°. The second, third, and fourth columns show 3D projections of the particle with spherical, ellipsoidal (thinner along the z-dimension), and particle-shaped masks, respectively. The fifth column displays the 3D projections of the enhanced and IsoNet missing-wedge-corrected particle. The sixth and seventh columns present the final 3D map and the flexibly fitted model, respectively. (B) Two cross-sectional views (12 nm thickness) of the plasmid density map along its plectoneme axis are shown in the left-middle panel. The intensity profile along the line crossing the two high-density DNA spots is displayed in the right panel. (C) Resolution assessment of the final 3D map using Fourier shell correlation (FSC). Two criteria are shown: FSC between two half-maps reconstructed from even and odd frames (evaluated at 0.143) and FSC between the final 3D map and the fitted model (evaluated at 0.5). (D) Zoomed-in views of the final 3D density map from panel A, displayed at two contour levels. (E) Superimposition of the high-contour level map from panel D onto its fitted model.

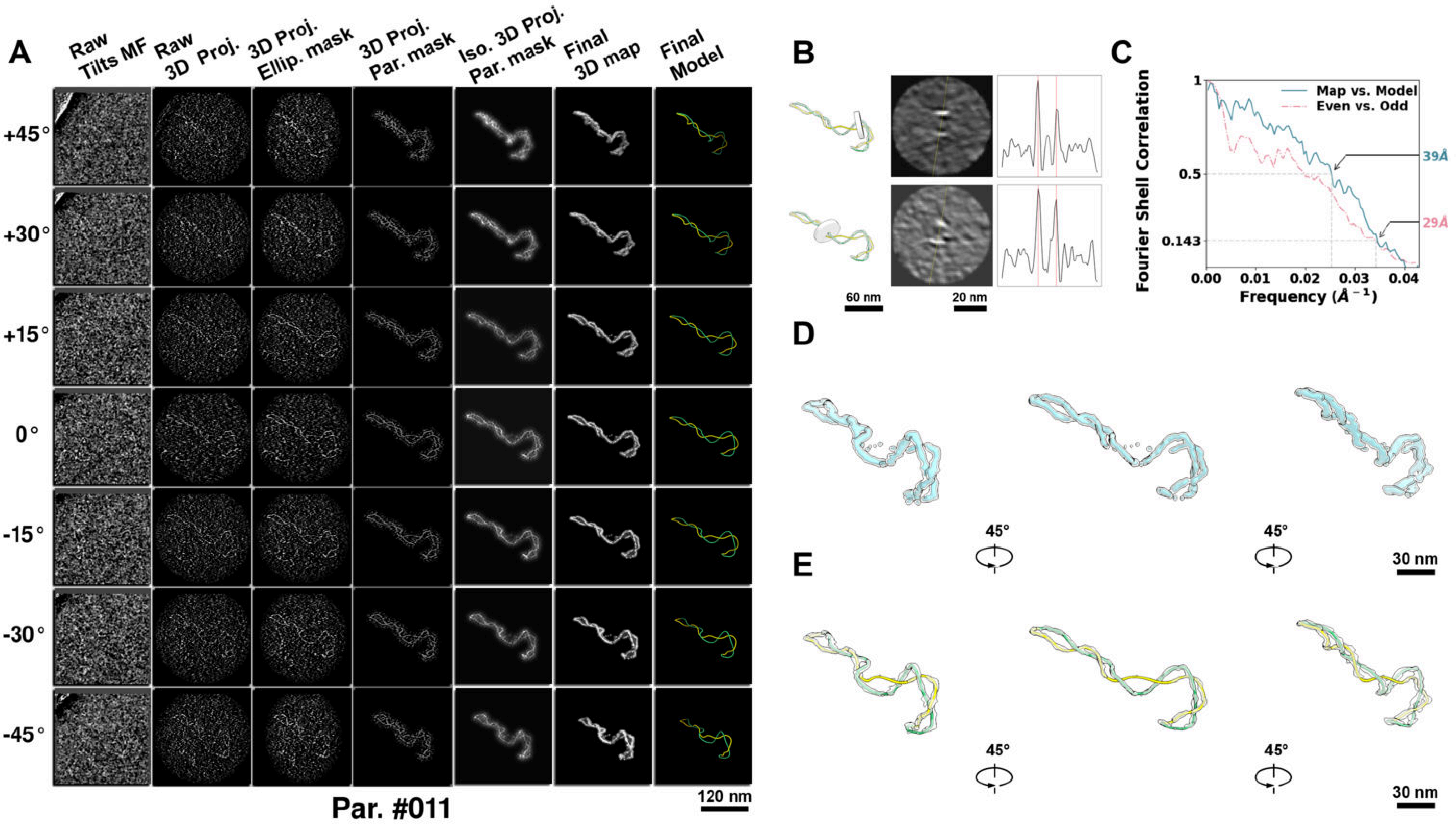

**Supplementary Particle Figure 11. Cryo-ET 3D reconstruction of an individual P.LS particle.**
(A) 3D reconstruction of the plasmid particle (index no. 11). The first column shows seven representative tilt images from +45° to -45° in step of 15°. The second, third, and fourth columns show 3D projections of the particle with spherical, ellipsoidal (thinner along the z-dimension), and particle-shaped masks, respectively. The fifth column displays the 3D projections of the enhanced and IsoNet missing-wedge-corrected particle. The sixth and seventh columns present the final 3D map and the flexibly fitted model, respectively. (B) Two cross-sectional views (12 nm thickness) of the plasmid density map along its plectoneme axis are shown in the left-middle panel. The intensity profile along the line crossing the two high-density DNA spots is displayed in the right panel. (C) Resolution assessment of the final 3D map using Fourier shell correlation (FSC). Two criteria are shown: FSC between two half-maps reconstructed from even and odd frames (evaluated at 0.143) and FSC between the final 3D map and the fitted model (evaluated at 0.5). (D) Zoomed-in views of the final 3D density map from panel A, displayed at two contour levels. (E) Superimposition of the high-contour level map from panel D onto its fitted model.

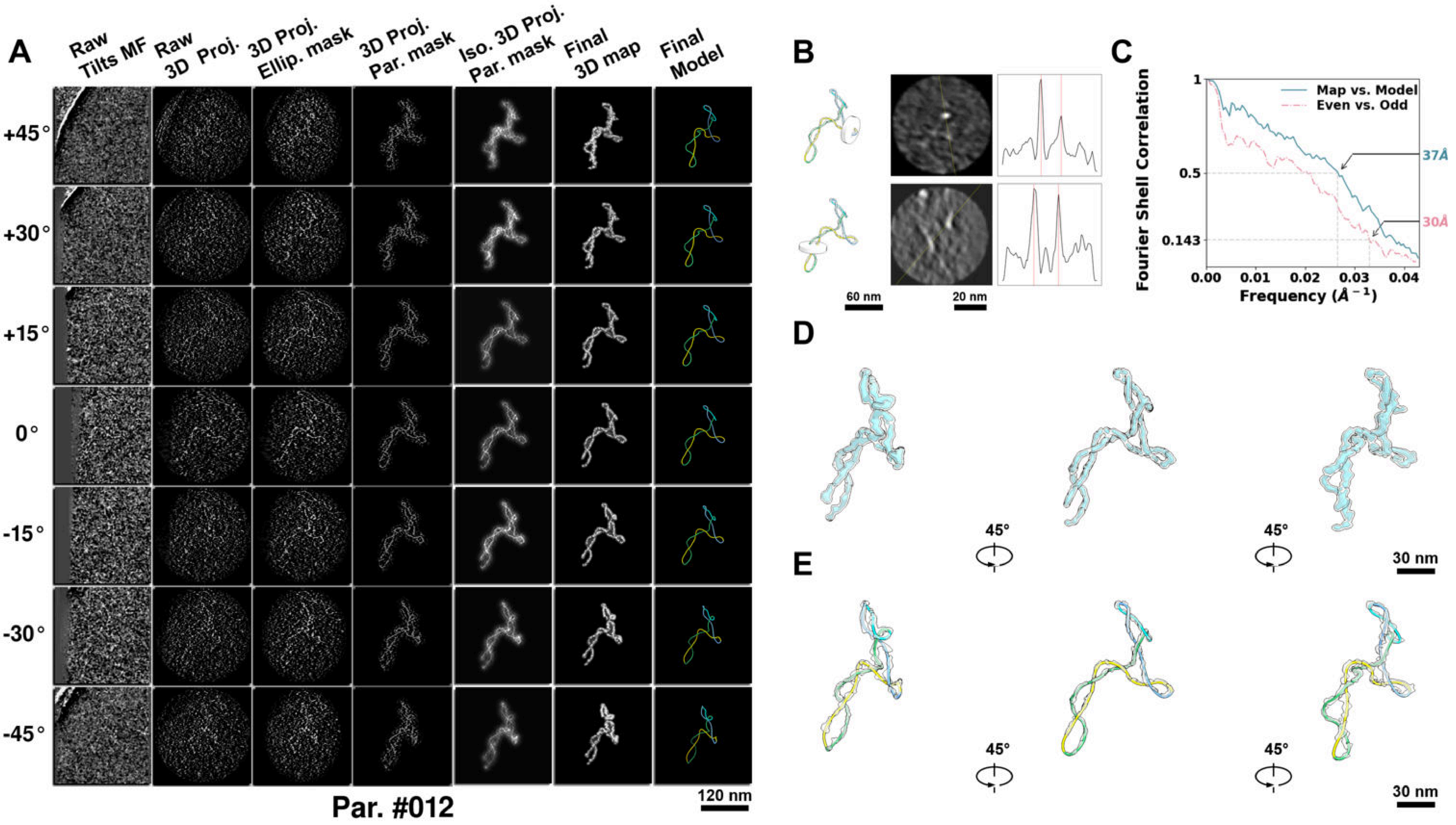

**Supplementary Particle Figure 12. Cryo-ET 3D reconstruction of an individual P.LS particle.**
(A) 3D reconstruction of the plasmid particle (index no. 12). The first column shows seven representative tilt images from +45° to -45° in step of 15°. The second, third, and fourth columns show 3D projections of the particle with spherical, ellipsoidal (thinner along the z-dimension), and particle-shaped masks, respectively. The fifth column displays the 3D projections of the enhanced and IsoNet missing-wedge-corrected particle. The sixth and seventh columns present the final 3D map and the flexibly fitted model, respectively. (B) Two cross-sectional views (12 nm thickness) of the plasmid density map along its plectoneme axis are shown in the left-middle panel. The intensity profile along the line crossing the two high-density DNA spots is displayed in the right panel. (C) Resolution assessment of the final 3D map using Fourier shell correlation (FSC). Two criteria are shown: FSC between two half-maps reconstructed from even and odd frames (evaluated at 0.143) and FSC between the final 3D map and the fitted model (evaluated at 0.5). (D) Zoomed-in views of the final 3D density map from panel A, displayed at two contour levels. (E) Superimposition of the high-contour level map from panel D onto its fitted model.

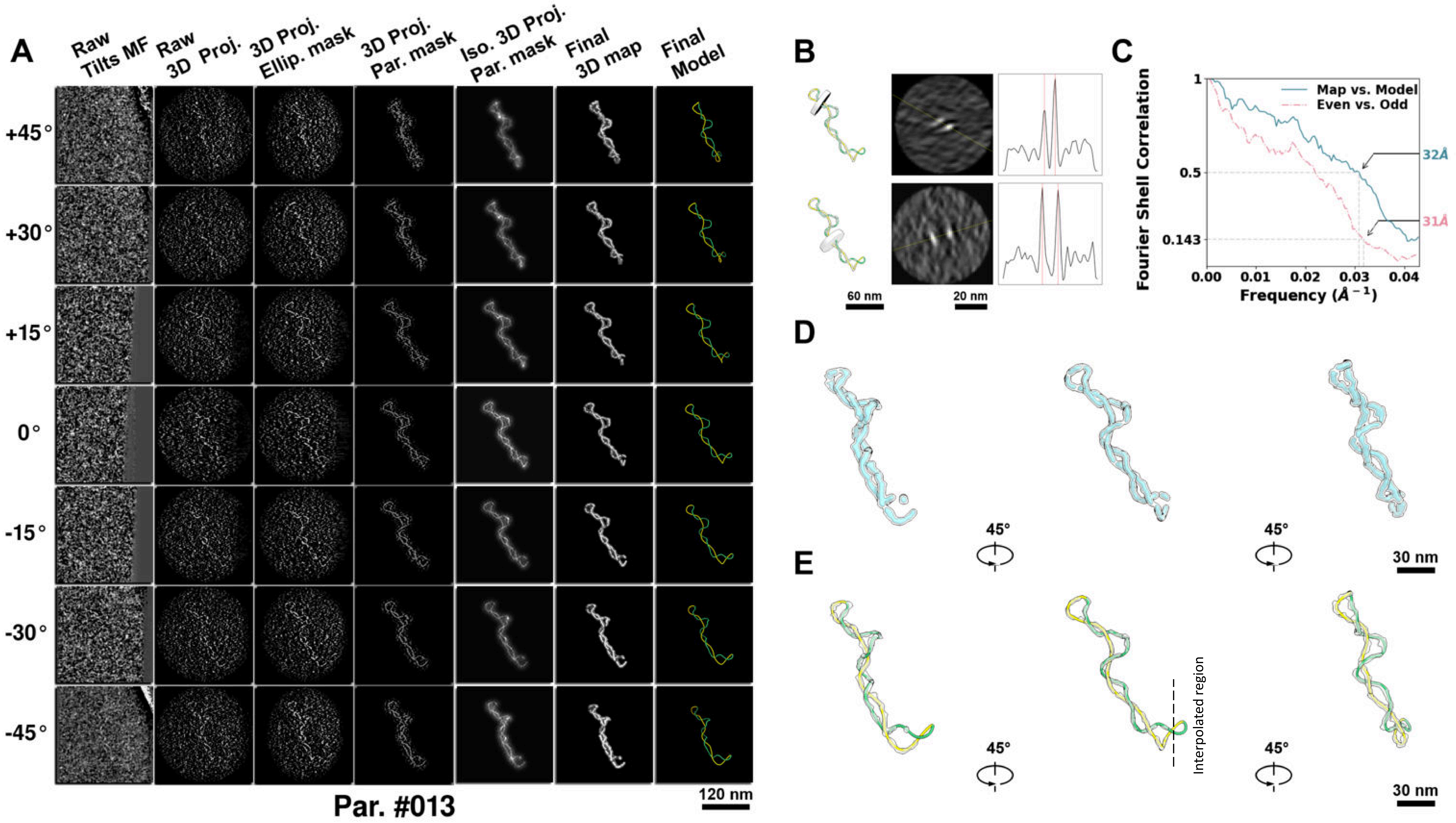

**Supplementary Particle Figure 13. Cryo-ET 3D reconstruction of an individual P.LS particle.**
(A) 3D reconstruction of the plasmid particle (index no. 13). The first column shows seven representative tilt images from +45° to -45° in step of 15°. The second, third, and fourth columns show 3D projections of the particle with spherical, ellipsoidal (thinner along the z-dimension), and particle-shaped masks, respectively. The fifth column displays the 3D projections of the enhanced and IsoNet missing-wedge-corrected particle. The sixth and seventh columns present the final 3D map and the flexibly fitted model, respectively. (B) Two cross-sectional views (12 nm thickness) of the plasmid density map along its plectoneme axis are shown in the left-middle panel. The intensity profile along the line crossing the two high-density DNA spots is displayed in the right panel. (C) Resolution assessment of the final 3D map using Fourier shell correlation (FSC). Two criteria are shown: FSC between two half-maps reconstructed from even and odd frames (evaluated at 0.143) and FSC between the final 3D map and the fitted model (evaluated at 0.5). (D) Zoomed-in views of the final 3D density map from panel A, displayed at two contour levels. (E) Superimposition of the high-contour level map from panel D onto its fitted model.

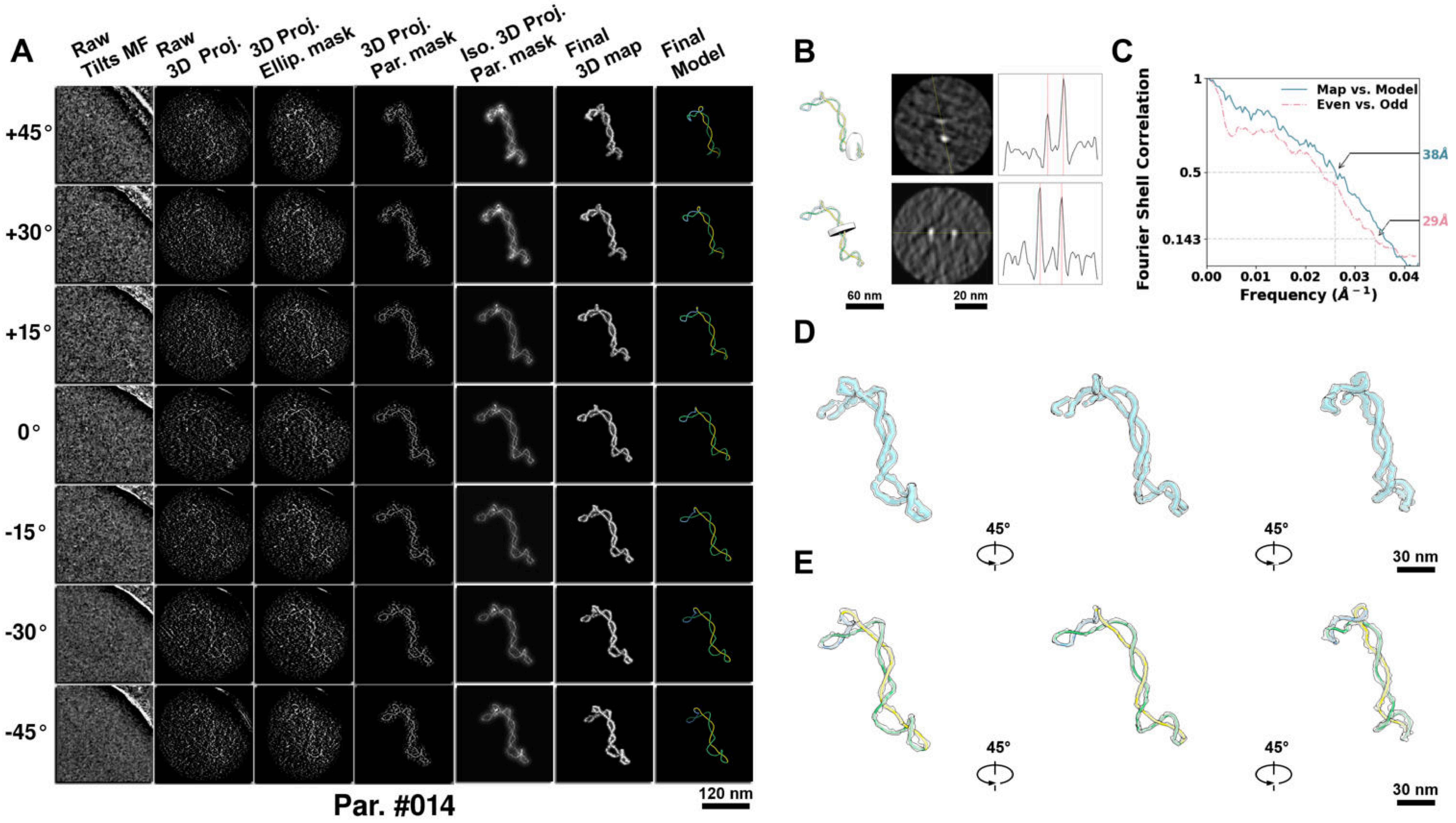

**Supplementary Particle Figure 14. Cryo-ET 3D reconstruction of an individual P.LS particle.**
(A) 3D reconstruction of the plasmid particle (index no. 14). The first column shows seven representative tilt images from +45° to -45° in step of 15°. The second, third, and fourth columns show 3D projections of the particle with spherical, ellipsoidal (thinner along the z-dimension), and particle-shaped masks, respectively. The fifth column displays the 3D projections of the enhanced and IsoNet missing-wedge-corrected particle. The sixth and seventh columns present the final 3D map and the flexibly fitted model, respectively. (B) Two cross-sectional views (12 nm thickness) of the plasmid density map along its plectoneme axis are shown in the left-middle panel. The intensity profile along the line crossing the two high-density DNA spots is displayed in the right panel. (C) Resolution assessment of the final 3D map using Fourier shell correlation (FSC). Two criteria are shown: FSC between two half-maps reconstructed from even and odd frames (evaluated at 0.143) and FSC between the final 3D map and the fitted model (evaluated at 0.5). (D) Zoomed-in views of the final 3D density map from panel A, displayed at two contour levels. (E) Superimposition of the high-contour level map from panel D onto its fitted model.

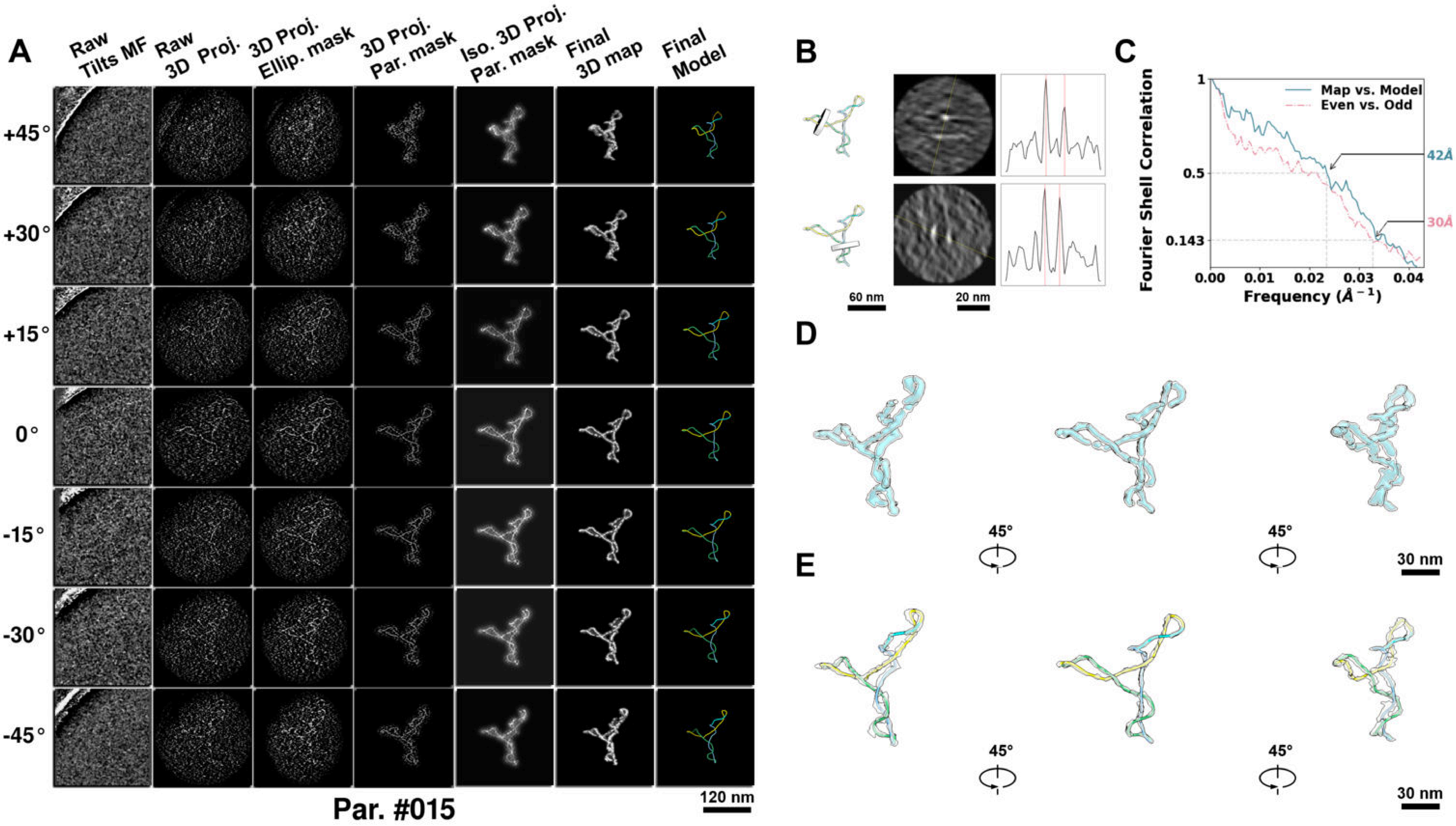

**Supplementary Particle Figure 15. Cryo-ET 3D reconstruction of an individual P.LS particle.**
(A) 3D reconstruction of the plasmid particle (index no. 15). The first column shows seven representative tilt images from +45° to -45° in step of 15°. The second, third, and fourth columns show 3D projections of the particle with spherical, ellipsoidal (thinner along the z-dimension), and particle-shaped masks, respectively. The fifth column displays the 3D projections of the enhanced and IsoNet missing-wedge-corrected particle. The sixth and seventh columns present the final 3D map and the flexibly fitted model, respectively. (B) Two cross-sectional views (12 nm thickness) of the plasmid density map along its plectoneme axis are shown in the left-middle panel. The intensity profile along the line crossing the two high-density DNA spots is displayed in the right panel. (C) Resolution assessment of the final 3D map using Fourier shell correlation (FSC). Two criteria are shown: FSC between two half-maps reconstructed from even and odd frames (evaluated at 0.143) and FSC between the final 3D map and the fitted model (evaluated at 0.5). (D) Zoomed-in views of the final 3D density map from panel A, displayed at two contour levels. (E) Superimposition of the high-contour level map from panel D onto its fitted model.

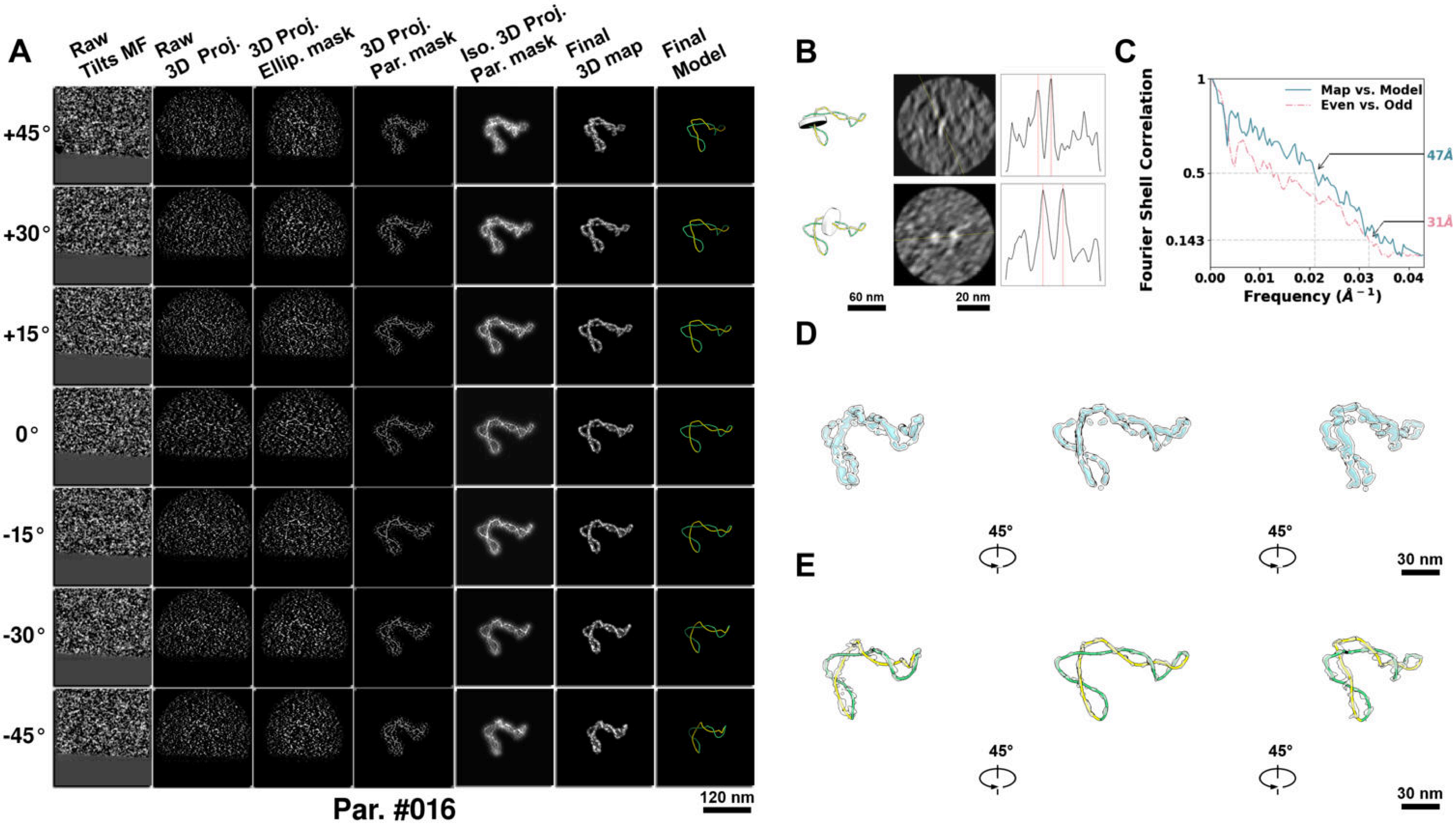

**Supplementary Particle Figure 16. Cryo-ET 3D reconstruction of an individual P.LS particle.**
(A) 3D reconstruction of the plasmid particle (index no. 16). The first column shows seven representative tilt images from +45° to -45° in step of 15°. The second, third, and fourth columns show 3D projections of the particle with spherical, ellipsoidal (thinner along the z-dimension), and particle-shaped masks, respectively. The fifth column displays the 3D projections of the enhanced and IsoNet missing-wedge-corrected particle. The sixth and seventh columns present the final 3D map and the flexibly fitted model, respectively. (B) Two cross-sectional views (12 nm thickness) of the plasmid density map along its plectoneme axis are shown in the left-middle panel. The intensity profile along the line crossing the two high-density DNA spots is displayed in the right panel. (C) Resolution assessment of the final 3D map using Fourier shell correlation (FSC). Two criteria are shown: FSC between two half-maps reconstructed from even and odd frames (evaluated at 0.143) and FSC between the final 3D map and the fitted model (evaluated at 0.5). (D) Zoomed-in views of the final 3D density map from panel A, displayed at two contour levels. (E) Superimposition of the high-contour level map from panel D onto its fitted model.

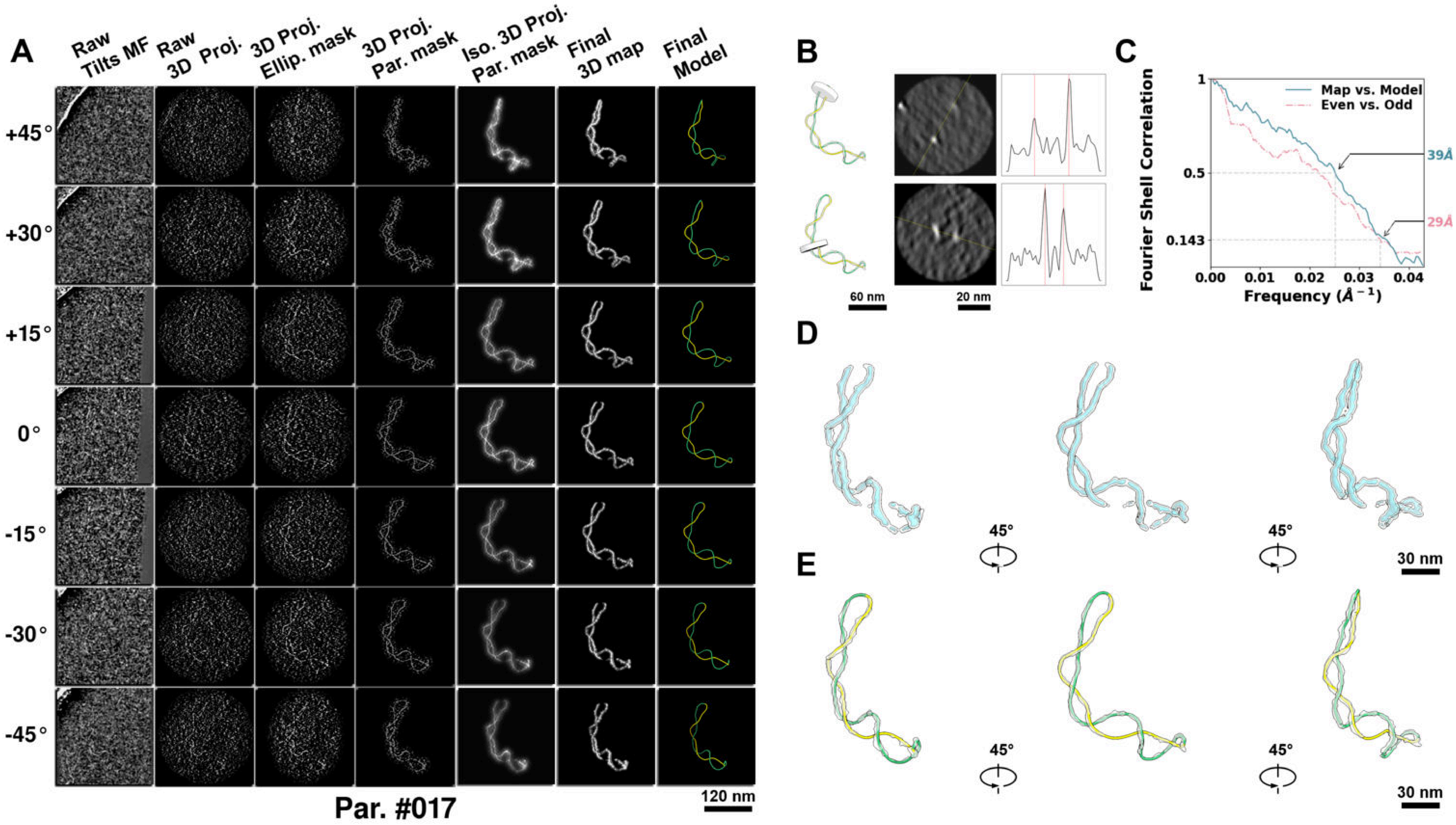

**Supplementary Particle Figure 17. Cryo-ET 3D reconstruction of an individual P.LS particle.**
(A) 3D reconstruction of the plasmid particle (index no. 17). The first column shows seven representative tilt images from +45° to -45° in step of 15°. The second, third, and fourth columns show 3D projections of the particle with spherical, ellipsoidal (thinner along the z-dimension), and particle-shaped masks, respectively. The fifth column displays the 3D projections of the enhanced and IsoNet missing-wedge-corrected particle. The sixth and seventh columns present the final 3D map and the flexibly fitted model, respectively. (B) Two cross-sectional views (12 nm thickness) of the plasmid density map along its plectoneme axis are shown in the left-middle panel. The intensity profile along the line crossing the two high-density DNA spots is displayed in the right panel. (C) Resolution assessment of the final 3D map using Fourier shell correlation (FSC). Two criteria are shown: FSC between two half-maps reconstructed from even and odd frames (evaluated at 0.143) and FSC between the final 3D map and the fitted model (evaluated at 0.5). (D) Zoomed-in views of the final 3D density map from panel A, displayed at two contour levels. (E) Superimposition of the high-contour level map from panel D onto its fitted model.

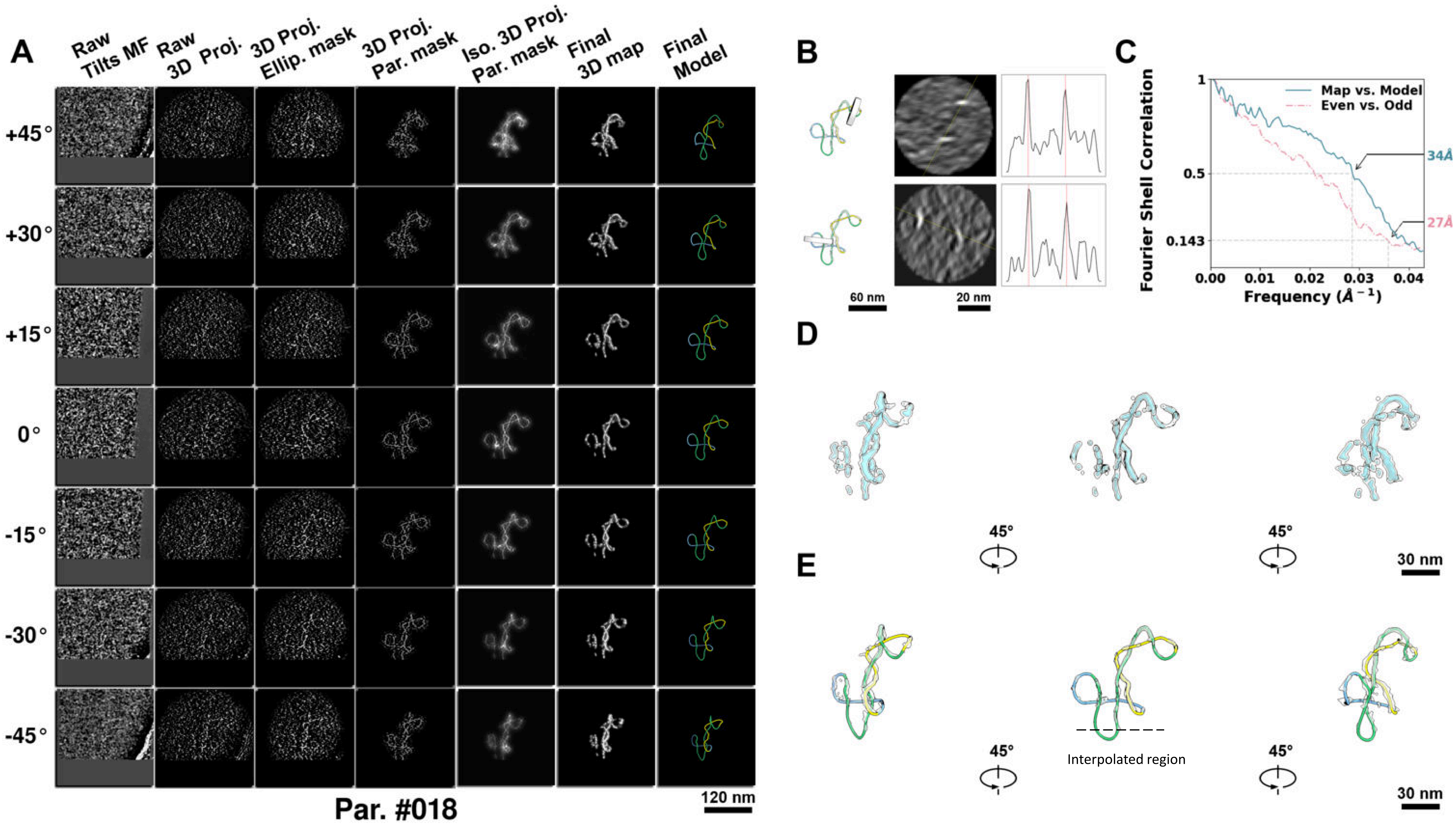

**Supplementary Particle Figure 18. Cryo-ET 3D reconstruction of an individual P.LS particle.**
(A) 3D reconstruction of the plasmid particle (index no. 18). The first column shows seven representative tilt images from +45° to -45° in step of 15°. The second, third, and fourth columns show 3D projections of the particle with spherical, ellipsoidal (thinner along the z-dimension), and particle-shaped masks, respectively. The fifth column displays the 3D projections of the enhanced and IsoNet missing-wedge-corrected particle. The sixth and seventh columns present the final 3D map and the flexibly fitted model, respectively. (B) Two cross-sectional views (12 nm thickness) of the plasmid density map along its plectoneme axis are shown in the left-middle panel. The intensity profile along the line crossing the two high-density DNA spots is displayed in the right panel. (C) Resolution assessment of the final 3D map using Fourier shell correlation (FSC). Two criteria are shown: FSC between two half-maps reconstructed from even and odd frames (evaluated at 0.143) and FSC between the final 3D map and the fitted model (evaluated at 0.5). (D) Zoomed-in views of the final 3D density map from panel A, displayed at two contour levels. (E) Superimposition of the high-contour level map from panel D onto its fitted model.

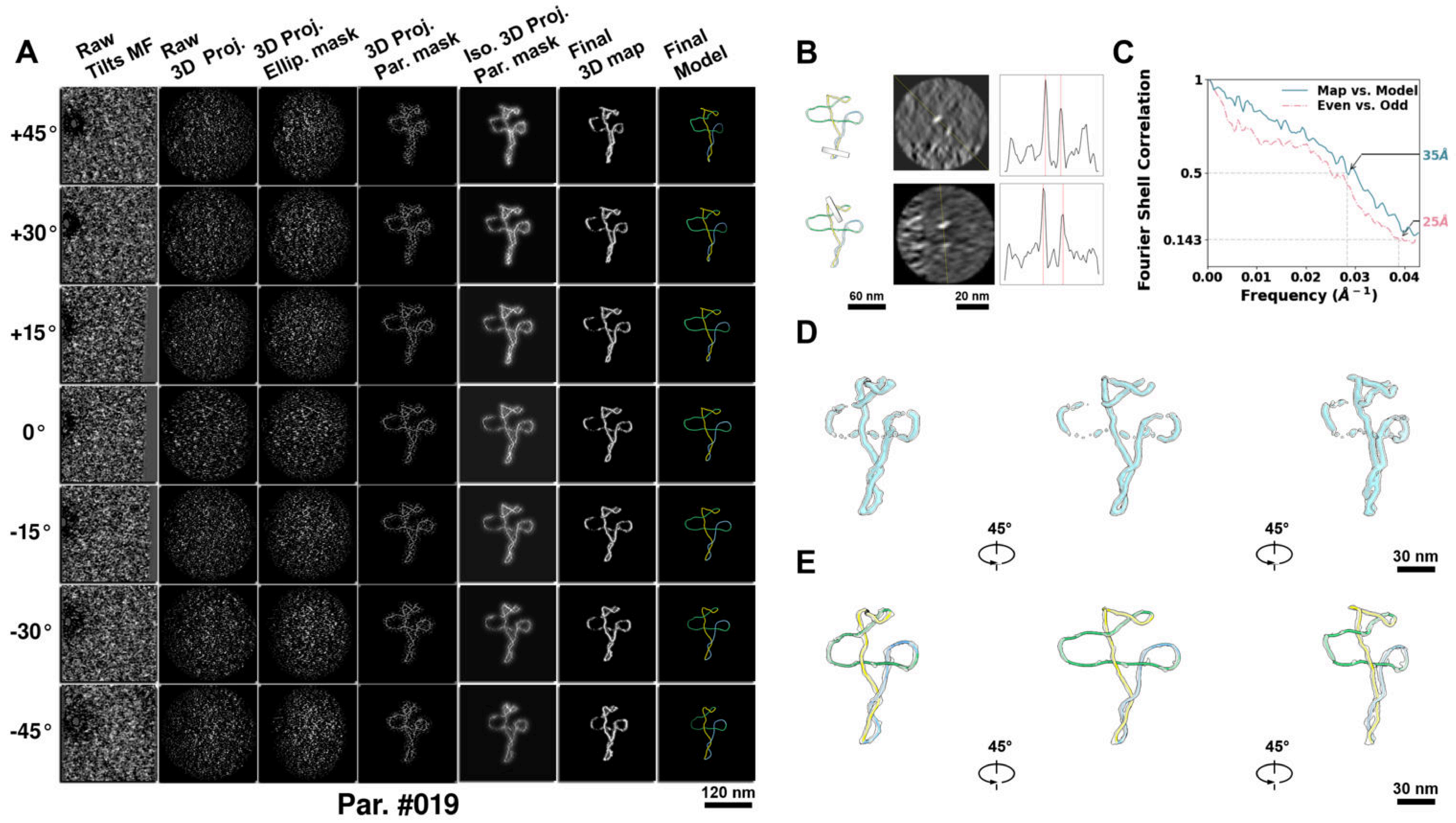

**Supplementary Particle Figure 19. Cryo-ET 3D reconstruction of an individual P.LS particle.**
(A) 3D reconstruction of the plasmid particle (index no. 19). The first column shows seven representative tilt images from +45° to -45° in step of 15°. The second, third, and fourth columns show 3D projections of the particle with spherical, ellipsoidal (thinner along the z-dimension), and particle-shaped masks, respectively. The fifth column displays the 3D projections of the enhanced and IsoNet missing-wedge-corrected particle. The sixth and seventh columns present the final 3D map and the flexibly fitted model, respectively. (B) Two cross-sectional views (12 nm thickness) of the plasmid density map along its plectoneme axis are shown in the left-middle panel. The intensity profile along the line crossing the two high-density DNA spots is displayed in the right panel. (C) Resolution assessment of the final 3D map using Fourier shell correlation (FSC). Two criteria are shown: FSC between two half-maps reconstructed from even and odd frames (evaluated at 0.143) and FSC between the final 3D map and the fitted model (evaluated at 0.5). (D) Zoomed-in views of the final 3D density map from panel A, displayed at two contour levels. (E) Superimposition of the high-contour level map from panel D onto its fitted model.

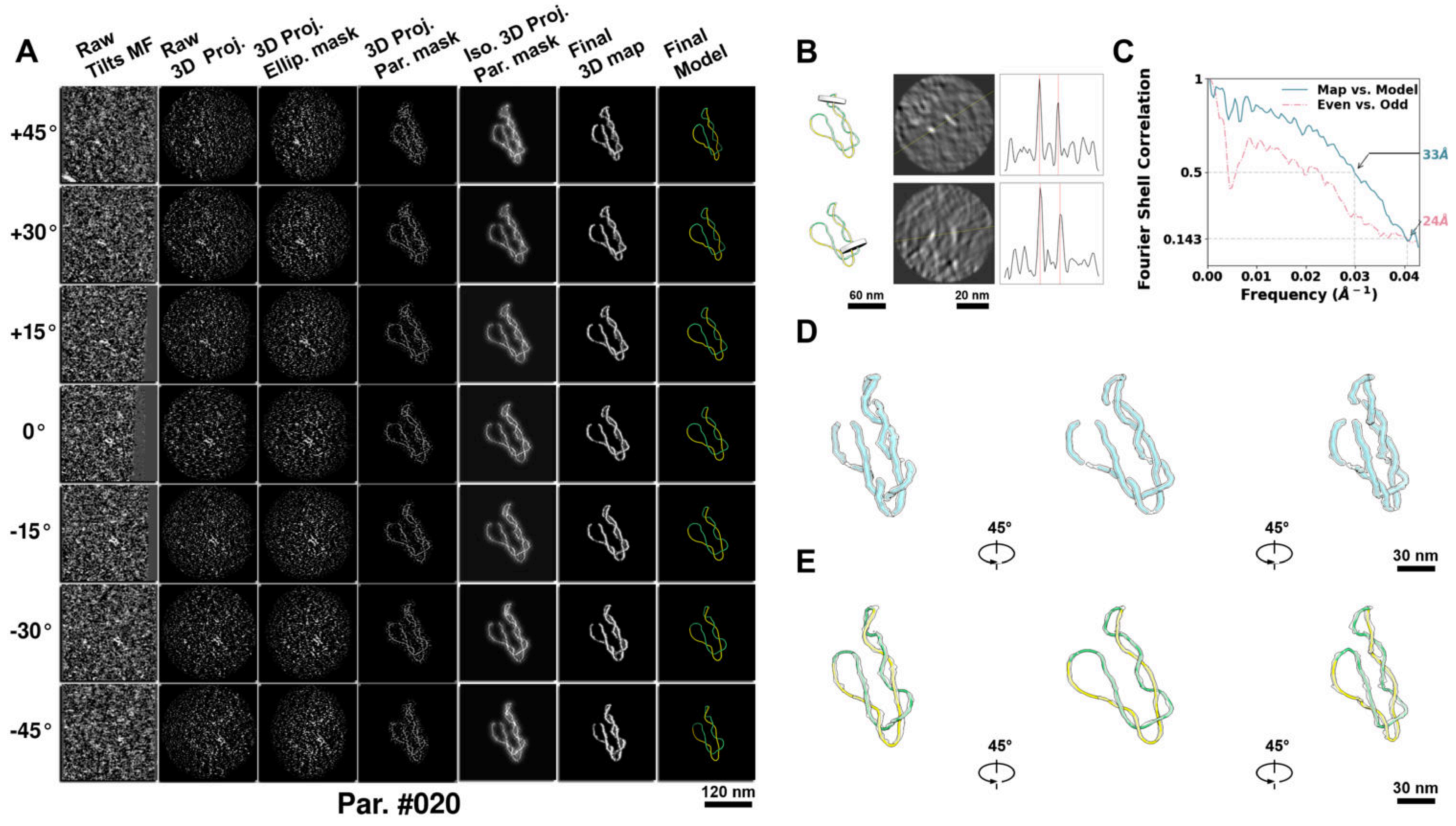

**Supplementary Particle Figure 20. Cryo-ET 3D reconstruction of an individual P.LS particle.**
(A) 3D reconstruction of the plasmid particle (index no. 20). The first column shows seven representative tilt images from +45° to -45° in step of 15°. The second, third, and fourth columns show 3D projections of the particle with spherical, ellipsoidal (thinner along the z-dimension), and particle-shaped masks, respectively. The fifth column displays the 3D projections of the enhanced and IsoNet missing-wedge-corrected particle. The sixth and seventh columns present the final 3D map and the flexibly fitted model, respectively. (B) Two cross-sectional views (12 nm thickness) of the plasmid density map along its plectoneme axis are shown in the left-middle panel. The intensity profile along the line crossing the two high-density DNA spots is displayed in the right panel. (C) Resolution assessment of the final 3D map using Fourier shell correlation (FSC). Two criteria are shown: FSC between two half-maps reconstructed from even and odd frames (evaluated at 0.143) and FSC between the final 3D map and the fitted model (evaluated at 0.5). (D) Zoomed-in views of the final 3D density map from panel A, displayed at two contour levels. (E) Superimposition of the high-contour level map from panel D onto its fitted model.

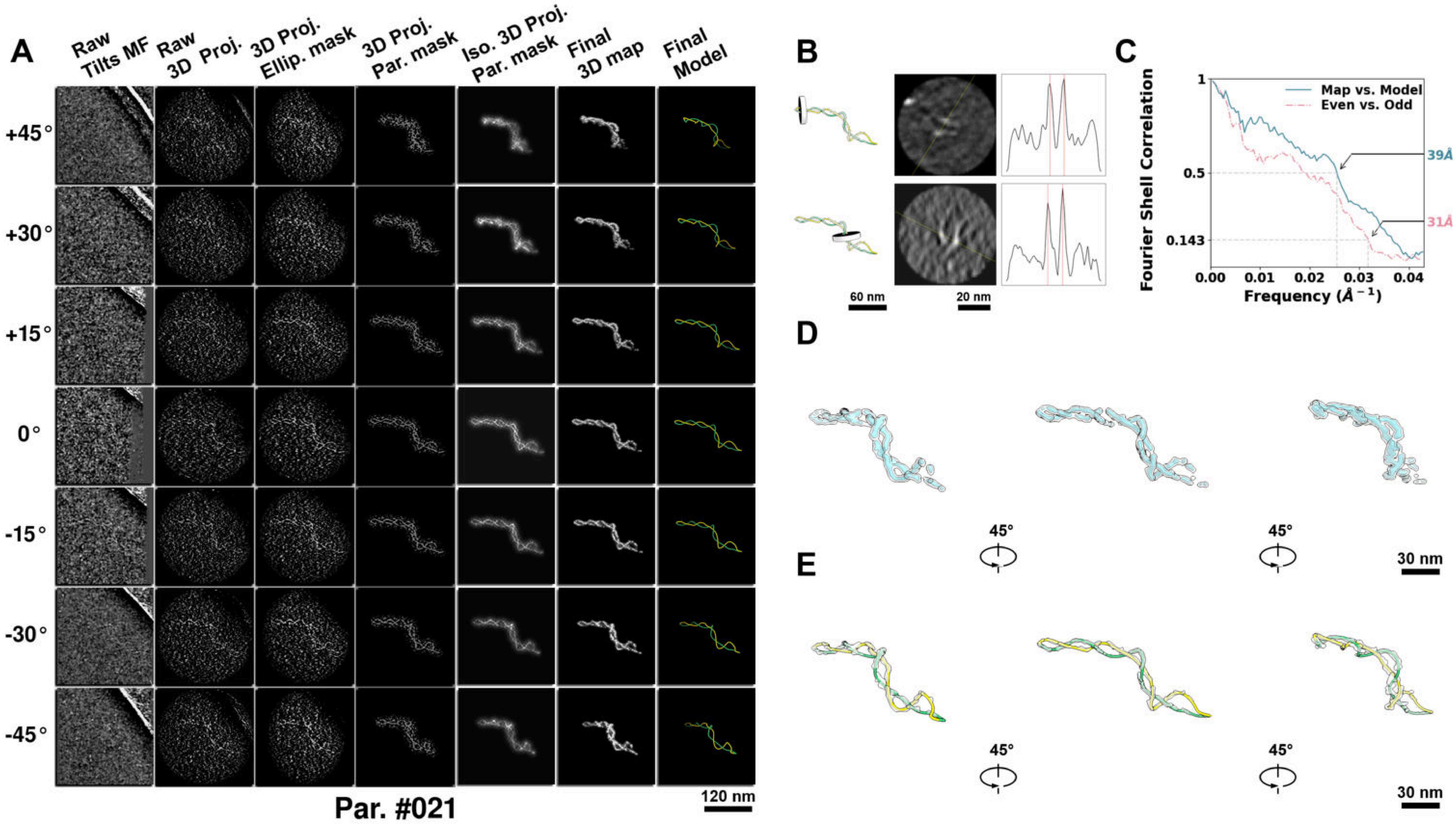

**Supplementary Particle Figure 21. Cryo-ET 3D reconstruction of an individual P.LS particle.**
(A) 3D reconstruction of the plasmid particle (index no. 21). The first column shows seven representative tilt images from +45° to -45° in step of 15°. The second, third, and fourth columns show 3D projections of the particle with spherical, ellipsoidal (thinner along the z-dimension), and particle-shaped masks, respectively. The fifth column displays the 3D projections of the enhanced and IsoNet missing-wedge-corrected particle. The sixth and seventh columns present the final 3D map and the flexibly fitted model, respectively. (B) Two cross-sectional views (12 nm thickness) of the plasmid density map along its plectoneme axis are shown in the left-middle panel. The intensity profile along the line crossing the two high-density DNA spots is displayed in the right panel. (C) Resolution assessment of the final 3D map using Fourier shell correlation (FSC). Two criteria are shown: FSC between two half-maps reconstructed from even and odd frames (evaluated at 0.143) and FSC between the final 3D map and the fitted model (evaluated at 0.5). (D) Zoomed-in views of the final 3D density map from panel A, displayed at two contour levels. (E) Superimposition of the high-contour level map from panel D onto its fitted model.

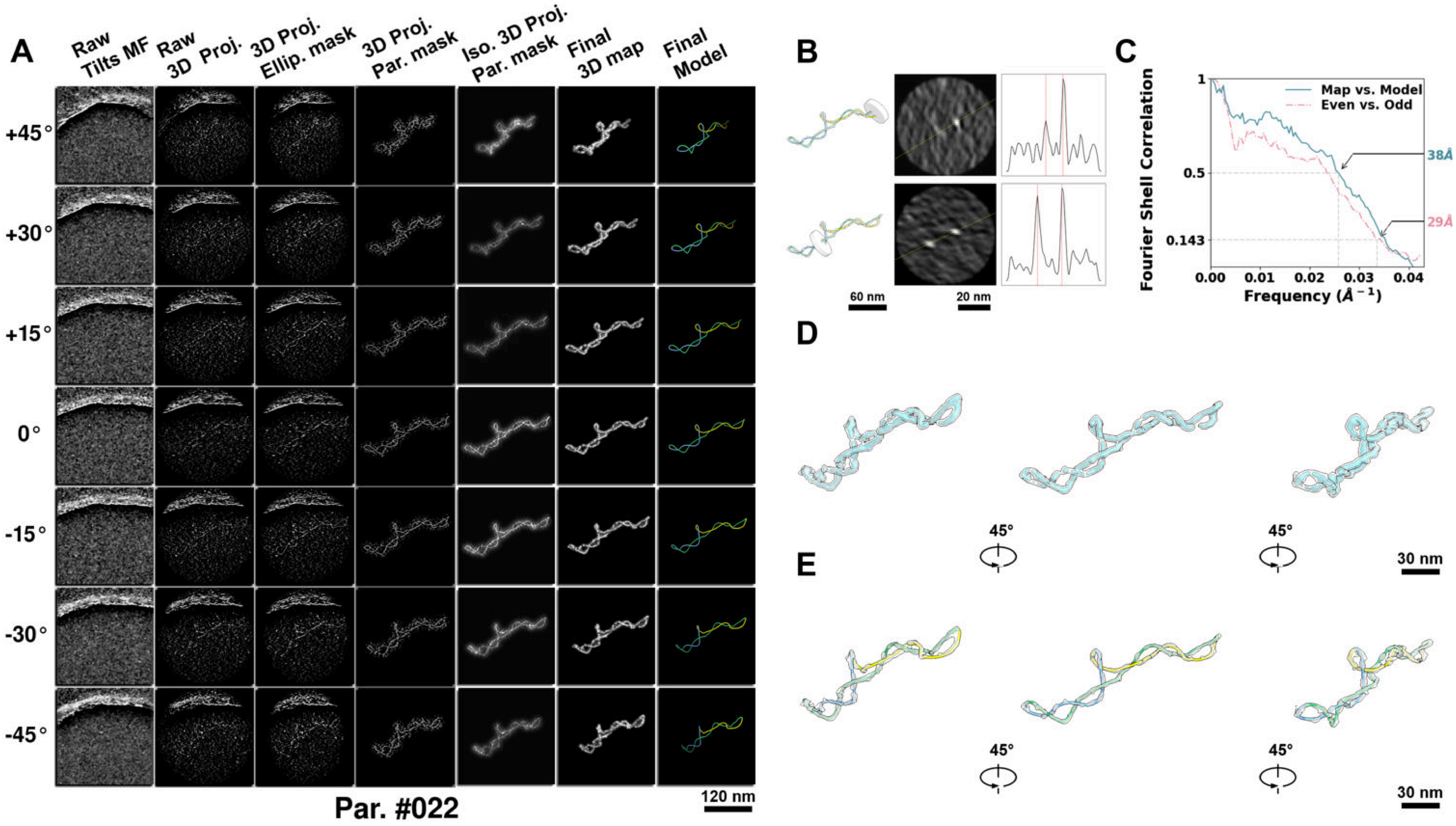

**Supplementary Particle Figure 22. Cryo-ET 3D reconstruction of an individual P.LS particle.**
(A) 3D reconstruction of the plasmid particle (index no. 22). The first column shows seven representative tilt images from +45° to -45° in step of 15°. The second, third, and fourth columns show 3D projections of the particle with spherical, ellipsoidal (thinner along the z-dimension), and particle-shaped masks, respectively. The fifth column displays the 3D projections of the enhanced and IsoNet missing-wedge-corrected particle. The sixth and seventh columns present the final 3D map and the flexibly fitted model, respectively. (B) Two cross-sectional views (12 nm thickness) of the plasmid density map along its plectoneme axis are shown in the left-middle panel. The intensity profile along the line crossing the two high-density DNA spots is displayed in the right panel. (C) Resolution assessment of the final 3D map using Fourier shell correlation (FSC). Two criteria are shown: FSC between two half-maps reconstructed from even and odd frames (evaluated at 0.143) and FSC between the final 3D map and the fitted model (evaluated at 0.5). (D) Zoomed-in views of the final 3D density map from panel A, displayed at two contour levels. (E) Superimposition of the high-contour level map from panel D onto its fitted model.

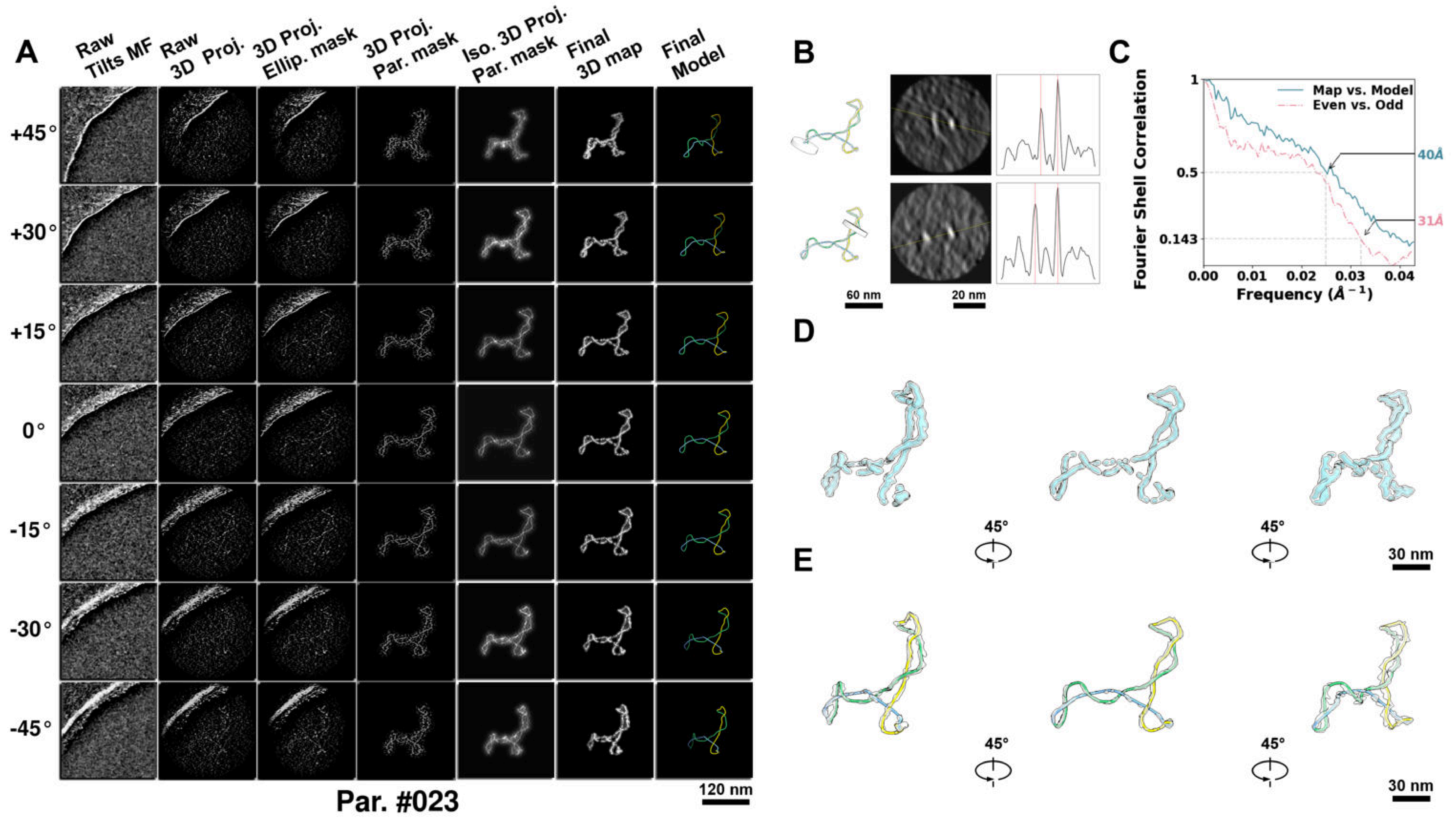

**Supplementary Particle Figure 23. Cryo-ET 3D reconstruction of an individual P.LS particle.**
(A) 3D reconstruction of the plasmid particle (index no. 23). The first column shows seven representative tilt images from +45° to -45° in step of 15°. The second, third, and fourth columns show 3D projections of the particle with spherical, ellipsoidal (thinner along the z-dimension), and particle-shaped masks, respectively. The fifth column displays the 3D projections of the enhanced and IsoNet missing-wedge-corrected particle. The sixth and seventh columns present the final 3D map and the flexibly fitted model, respectively. (B) Two cross-sectional views (12 nm thickness) of the plasmid density map along its plectoneme axis are shown in the left-middle panel. The intensity profile along the line crossing the two high-density DNA spots is displayed in the right panel. (C) Resolution assessment of the final 3D map using Fourier shell correlation (FSC). Two criteria are shown: FSC between two half-maps reconstructed from even and odd frames (evaluated at 0.143) and FSC between the final 3D map and the fitted model (evaluated at 0.5). (D) Zoomed-in views of the final 3D density map from panel A, displayed at two contour levels. (E) Superimposition of the high-contour level map from panel D onto its fitted model.

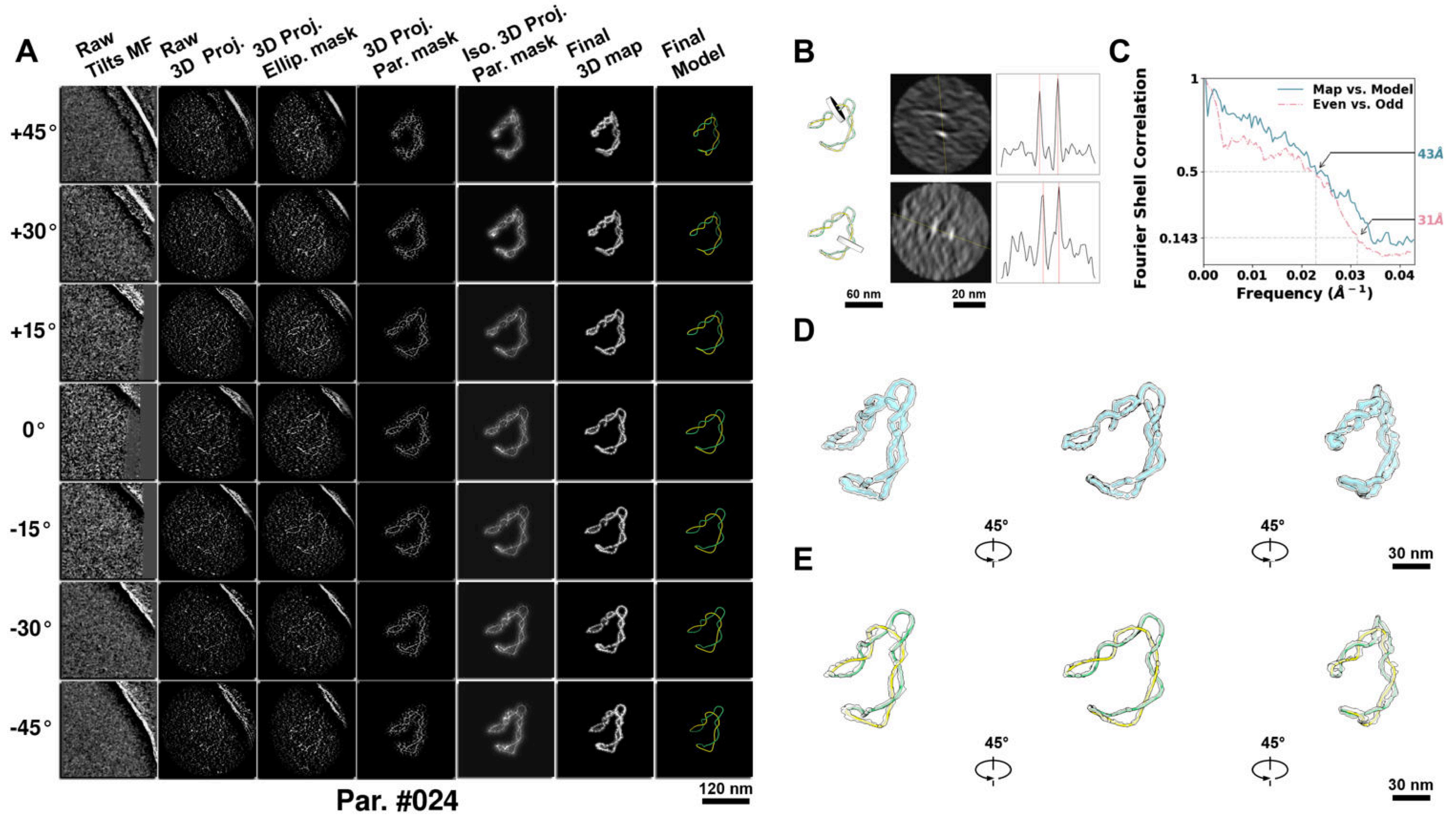

**Supplementary Particle Figure 24. Cryo-ET 3D reconstruction of an individual P.LS particle.**
(A) 3D reconstruction of the plasmid particle (index no. 24). The first column shows seven representative tilt images from +45° to -45° in step of 15°. The second, third, and fourth columns show 3D projections of the particle with spherical, ellipsoidal (thinner along the z-dimension), and particle-shaped masks, respectively. The fifth column displays the 3D projections of the enhanced and IsoNet missing-wedge-corrected particle. The sixth and seventh columns present the final 3D map and the flexibly fitted model, respectively. (B) Two cross-sectional views (12 nm thickness) of the plasmid density map along its plectoneme axis are shown in the left-middle panel. The intensity profile along the line crossing the two high-density DNA spots is displayed in the right panel. (C) Resolution assessment of the final 3D map using Fourier shell correlation (FSC). Two criteria are shown: FSC between two half-maps reconstructed from even and odd frames (evaluated at 0.143) and FSC between the final 3D map and the fitted model (evaluated at 0.5). (D) Zoomed-in views of the final 3D density map from panel A, displayed at two contour levels. (E) Superimposition of the high-contour level map from panel D onto its fitted model.

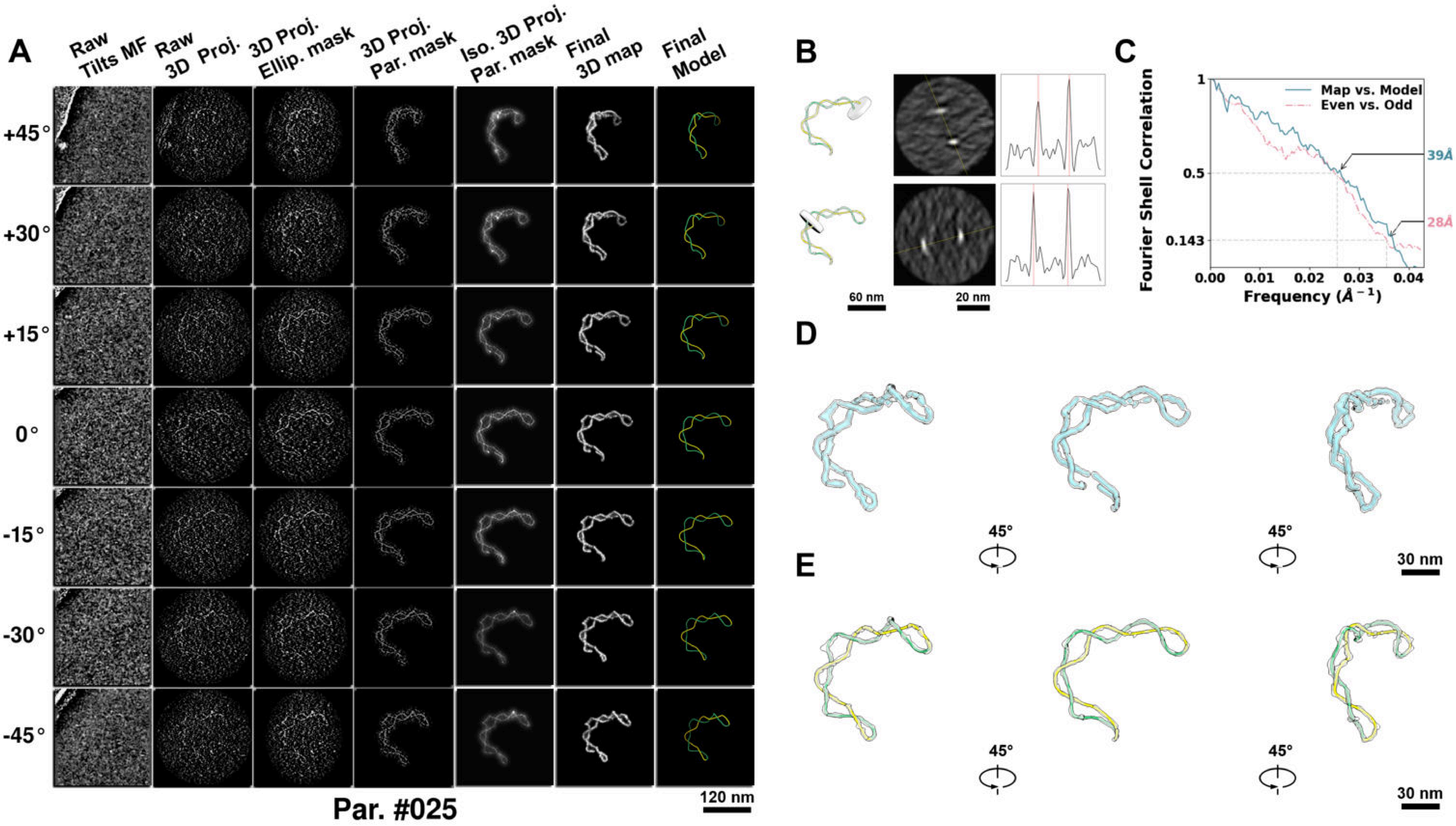

**Supplementary Particle Figure 25. Cryo-ET 3D reconstruction of an individual P.LS particle.**
(A) 3D reconstruction of the plasmid particle (index no. 25). The first column shows seven representative tilt images from +45° to -45° in step of 15°. The second, third, and fourth columns show 3D projections of the particle with spherical, ellipsoidal (thinner along the z-dimension), and particle-shaped masks, respectively. The fifth column displays the 3D projections of the enhanced and IsoNet missing-wedge-corrected particle. The sixth and seventh columns present the final 3D map and the flexibly fitted model, respectively. (B) Two cross-sectional views (12 nm thickness) of the plasmid density map along its plectoneme axis are shown in the left-middle panel. The intensity profile along the line crossing the two high-density DNA spots is displayed in the right panel. (C) Resolution assessment of the final 3D map using Fourier shell correlation (FSC). Two criteria are shown: FSC between two half-maps reconstructed from even and odd frames (evaluated at 0.143) and FSC between the final 3D map and the fitted model (evaluated at 0.5). (D) Zoomed-in views of the final 3D density map from panel A, displayed at two contour levels. (E) Superimposition of the high-contour level map from panel D onto its fitted model.

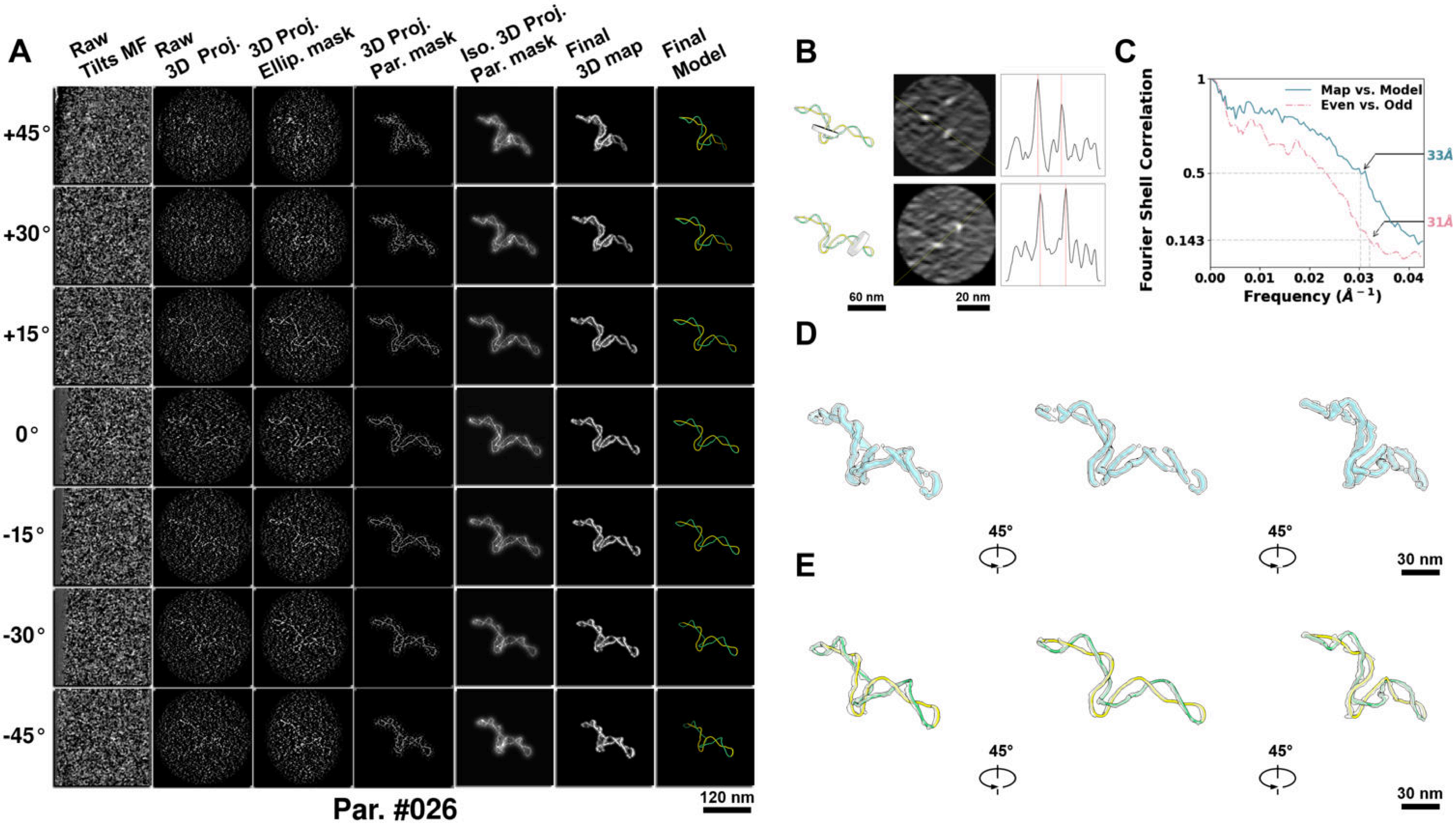

**Supplementary Particle Figure 26. Cryo-ET 3D reconstruction of an individual P.LS particle.**
(A) 3D reconstruction of the plasmid particle (index no. 26). The first column shows seven representative tilt images from +45° to -45° in step of 15°. The second, third, and fourth columns show 3D projections of the particle with spherical, ellipsoidal (thinner along the z-dimension), and particle-shaped masks, respectively. The fifth column displays the 3D projections of the enhanced and IsoNet missing-wedge-corrected particle. The sixth and seventh columns present the final 3D map and the flexibly fitted model, respectively. (B) Two cross-sectional views (12 nm thickness) of the plasmid density map along its plectoneme axis are shown in the left-middle panel. The intensity profile along the line crossing the two high-density DNA spots is displayed in the right panel. (C) Resolution assessment of the final 3D map using Fourier shell correlation (FSC). Two criteria are shown: FSC between two half-maps reconstructed from even and odd frames (evaluated at 0.143) and FSC between the final 3D map and the fitted model (evaluated at 0.5). (D) Zoomed-in views of the final 3D density map from panel A, displayed at two contour levels. (E) Superimposition of the high-contour level map from panel D onto its fitted model.

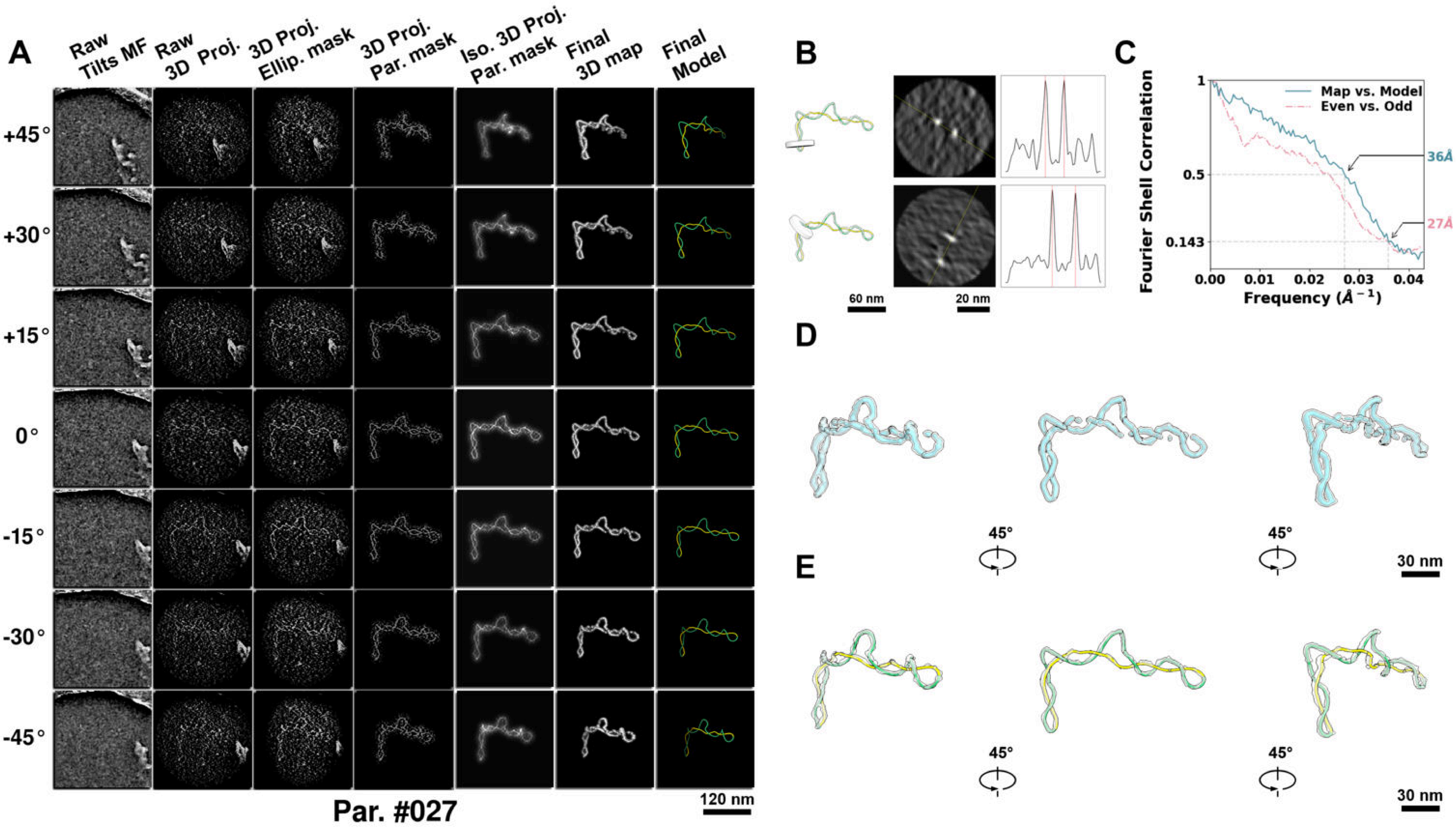

**Supplementary Particle Figure 27. Cryo-ET 3D reconstruction of an individual P.LS particle.**
(A) 3D reconstruction of the plasmid particle (index no. 27). The first column shows seven representative tilt images from +45° to -45° in step of 15°. The second, third, and fourth columns show 3D projections of the particle with spherical, ellipsoidal (thinner along the z-dimension), and particle-shaped masks, respectively. The fifth column displays the 3D projections of the enhanced and IsoNet missing-wedge-corrected particle. The sixth and seventh columns present the final 3D map and the flexibly fitted model, respectively. (B) Two cross-sectional views (12 nm thickness) of the plasmid density map along its plectoneme axis are shown in the left-middle panel. The intensity profile along the line crossing the two high-density DNA spots is displayed in the right panel. (C) Resolution assessment of the final 3D map using Fourier shell correlation (FSC). Two criteria are shown: FSC between two half-maps reconstructed from even and odd frames (evaluated at 0.143) and FSC between the final 3D map and the fitted model (evaluated at 0.5). (D) Zoomed-in views of the final 3D density map from panel A, displayed at two contour levels. (E) Superimposition of the high-contour level map from panel D onto its fitted model.

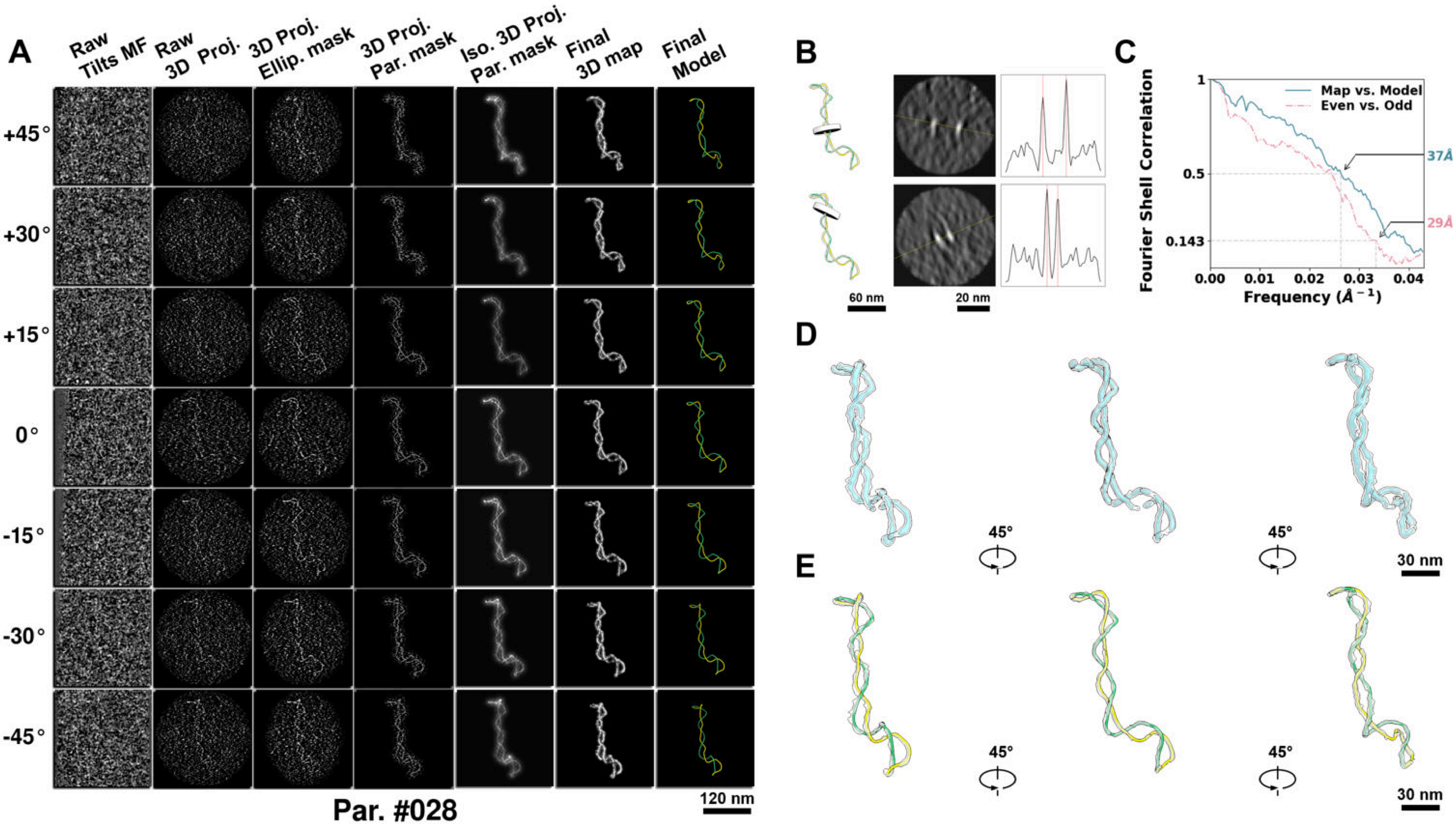

**Supplementary Particle Figure 28. Cryo-ET 3D reconstruction of an individual P.LS particle.**
(A) 3D reconstruction of the plasmid particle (index no. 28). The first column shows seven representative tilt images from +45° to -45° in step of 15°. The second, third, and fourth columns show 3D projections of the particle with spherical, ellipsoidal (thinner along the z-dimension), and particle-shaped masks, respectively. The fifth column displays the 3D projections of the enhanced and IsoNet missing-wedge-corrected particle. The sixth and seventh columns present the final 3D map and the flexibly fitted model, respectively. (B) Two cross-sectional views (12 nm thickness) of the plasmid density map along its plectoneme axis are shown in the left-middle panel. The intensity profile along the line crossing the two high-density DNA spots is displayed in the right panel. (C) Resolution assessment of the final 3D map using Fourier shell correlation (FSC). Two criteria are shown: FSC between two half-maps reconstructed from even and odd frames (evaluated at 0.143) and FSC between the final 3D map and the fitted model (evaluated at 0.5). (D) Zoomed-in views of the final 3D density map from panel A, displayed at two contour levels. (E) Superimposition of the high-contour level map from panel D onto its fitted model.

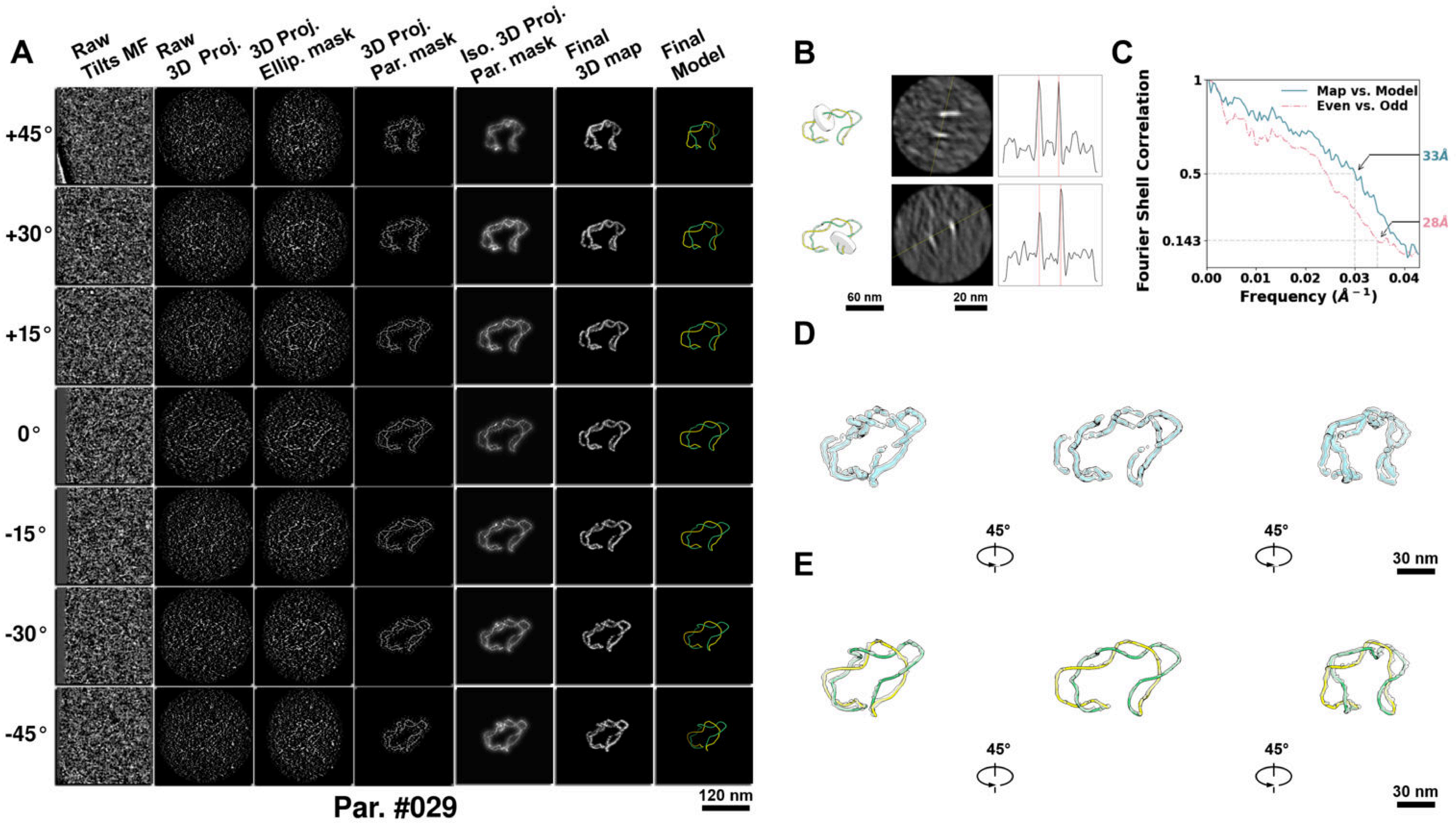

**Supplementary Particle Figure 29. Cryo-ET 3D reconstruction of an individual P.LS particle.**
(A) 3D reconstruction of the plasmid particle (index no. 29). The first column shows seven representative tilt images from +45° to -45° in step of 15°. The second, third, and fourth columns show 3D projections of the particle with spherical, ellipsoidal (thinner along the z-dimension), and particle-shaped masks, respectively. The fifth column displays the 3D projections of the enhanced and IsoNet missing-wedge-corrected particle. The sixth and seventh columns present the final 3D map and the flexibly fitted model, respectively. (B) Two cross-sectional views (12 nm thickness) of the plasmid density map along its plectoneme axis are shown in the left-middle panel. The intensity profile along the line crossing the two high-density DNA spots is displayed in the right panel. (C) Resolution assessment of the final 3D map using Fourier shell correlation (FSC). Two criteria are shown: FSC between two half-maps reconstructed from even and odd frames (evaluated at 0.143) and FSC between the final 3D map and the fitted model (evaluated at 0.5). (D) Zoomed-in views of the final 3D density map from panel A, displayed at two contour levels. (E) Superimposition of the high-contour level map from panel D onto its fitted model.

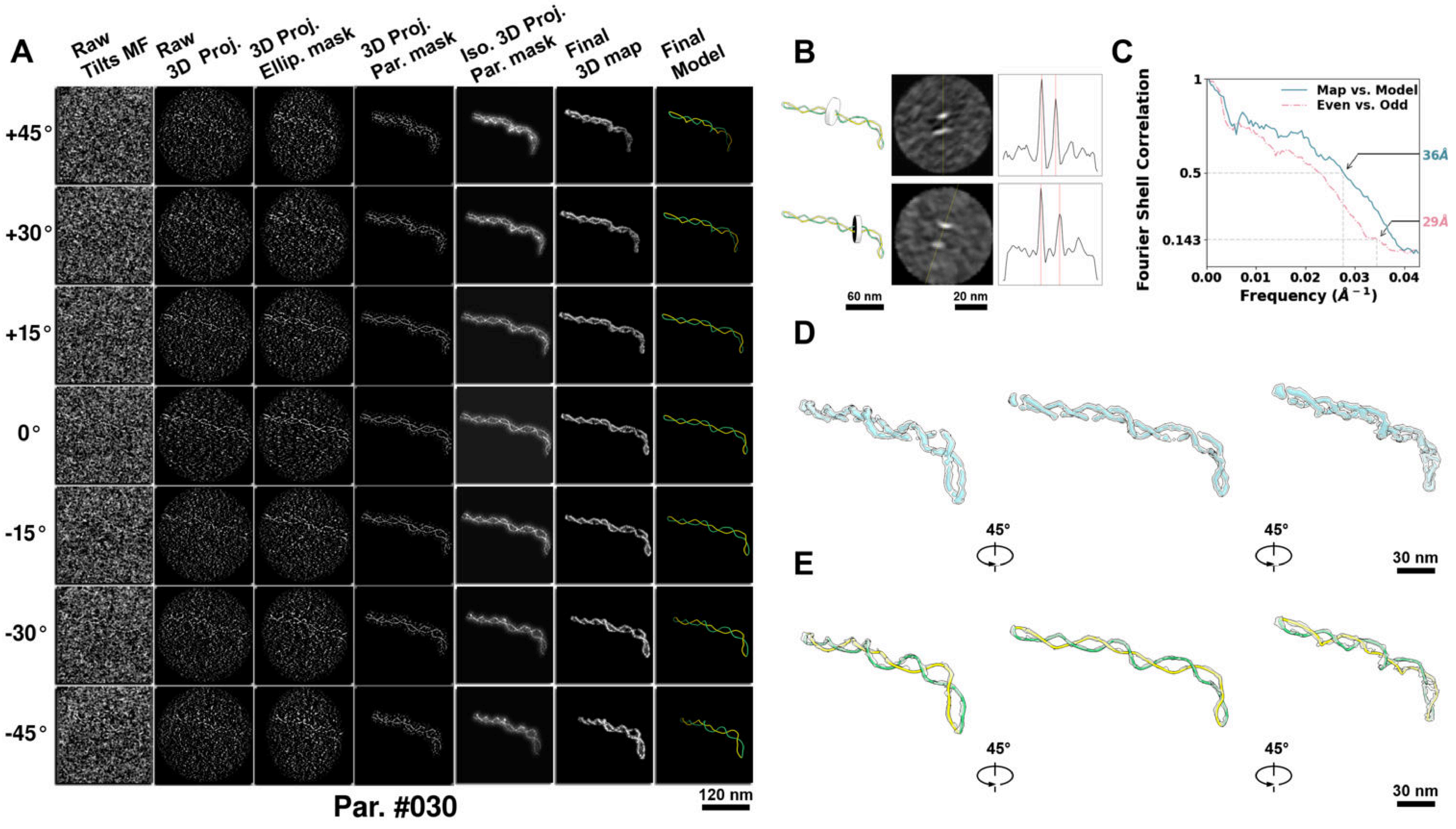

**Supplementary Particle Figure 30. Cryo-ET 3D reconstruction of an individual P.LS particle.**
(A) 3D reconstruction of the plasmid particle (index no. 30). The first column shows seven representative tilt images from +45° to -45° in step of 15°. The second, third, and fourth columns show 3D projections of the particle with spherical, ellipsoidal (thinner along the z-dimension), and particle-shaped masks, respectively. The fifth column displays the 3D projections of the enhanced and IsoNet missing-wedge-corrected particle. The sixth and seventh columns present the final 3D map and the flexibly fitted model, respectively. (B) Two cross-sectional views (12 nm thickness) of the plasmid density map along its plectoneme axis are shown in the left-middle panel. The intensity profile along the line crossing the two high-density DNA spots is displayed in the right panel. (C) Resolution assessment of the final 3D map using Fourier shell correlation (FSC). Two criteria are shown: FSC between two half-maps reconstructed from even and odd frames (evaluated at 0.143) and FSC between the final 3D map and the fitted model (evaluated at 0.5). (D) Zoomed-in views of the final 3D density map from panel A, displayed at two contour levels. (E) Superimposition of the high-contour level map from panel D onto its fitted model.

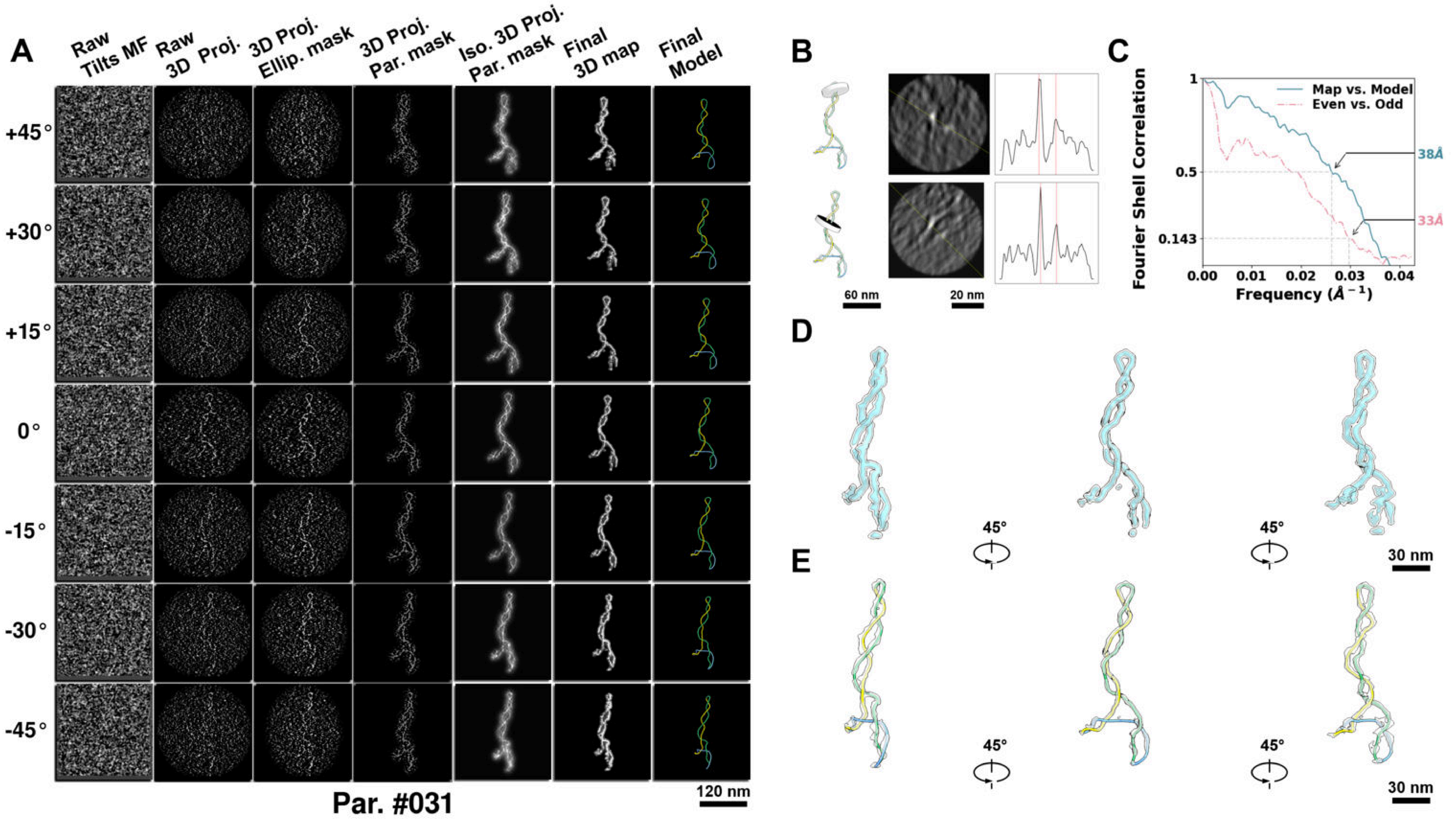

**Supplementary Particle Figure 31. Cryo-ET 3D reconstruction of an individual P.LS particle.**
(A) 3D reconstruction of the plasmid particle (index no. 31). The first column shows seven representative tilt images from +45° to -45° in step of 15°. The second, third, and fourth columns show 3D projections of the particle with spherical, ellipsoidal (thinner along the z-dimension), and particle-shaped masks, respectively. The fifth column displays the 3D projections of the enhanced and IsoNet missing-wedge-corrected particle. The sixth and seventh columns present the final 3D map and the flexibly fitted model, respectively. (B) Two cross-sectional views (12 nm thickness) of the plasmid density map along its plectoneme axis are shown in the left-middle panel. The intensity profile along the line crossing the two high-density DNA spots is displayed in the right panel. (C) Resolution assessment of the final 3D map using Fourier shell correlation (FSC). Two criteria are shown: FSC between two half-maps reconstructed from even and odd frames (evaluated at 0.143) and FSC between the final 3D map and the fitted model (evaluated at 0.5). (D) Zoomed-in views of the final 3D density map from panel A, displayed at two contour levels. (E) Superimposition of the high-contour level map from panel D onto its fitted model.

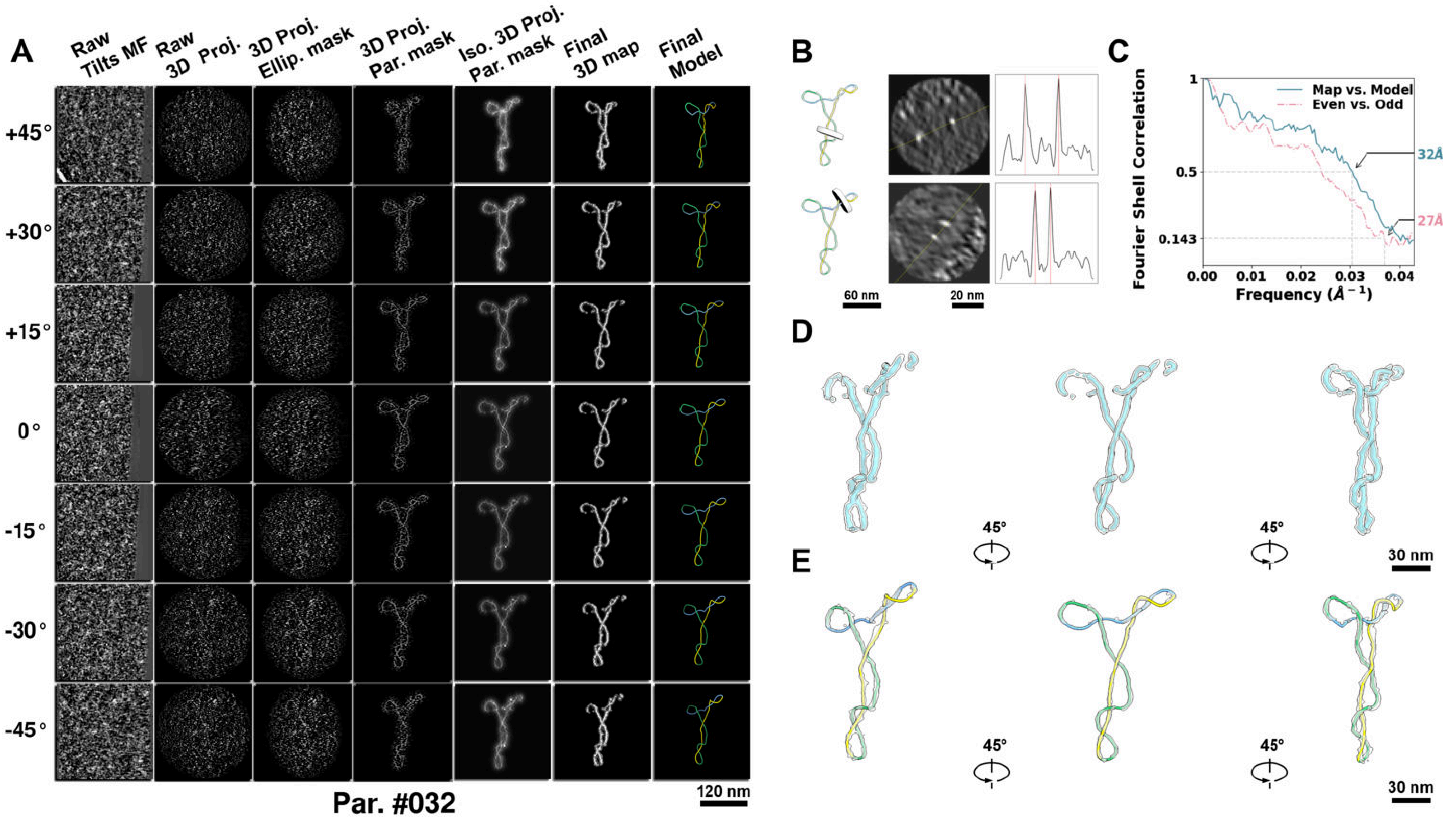

**Supplementary Particle Figure 32. Cryo-ET 3D reconstruction of an individual P.LS particle.**
(A) 3D reconstruction of the plasmid particle (index no. 32). The first column shows seven representative tilt images from +45° to -45° in step of 15°. The second, third, and fourth columns show 3D projections of the particle with spherical, ellipsoidal (thinner along the z-dimension), and particle-shaped masks, respectively. The fifth column displays the 3D projections of the enhanced and IsoNet missing-wedge-corrected particle. The sixth and seventh columns present the final 3D map and the flexibly fitted model, respectively. (B) Two cross-sectional views (12 nm thickness) of the plasmid density map along its plectoneme axis are shown in the left-middle panel. The intensity profile along the line crossing the two high-density DNA spots is displayed in the right panel. (C) Resolution assessment of the final 3D map using Fourier shell correlation (FSC). Two criteria are shown: FSC between two half-maps reconstructed from even and odd frames (evaluated at 0.143) and FSC between the final 3D map and the fitted model (evaluated at 0.5). (D) Zoomed-in views of the final 3D density map from panel A, displayed at two contour levels. (E) Superimposition of the high-contour level map from panel D onto its fitted model.

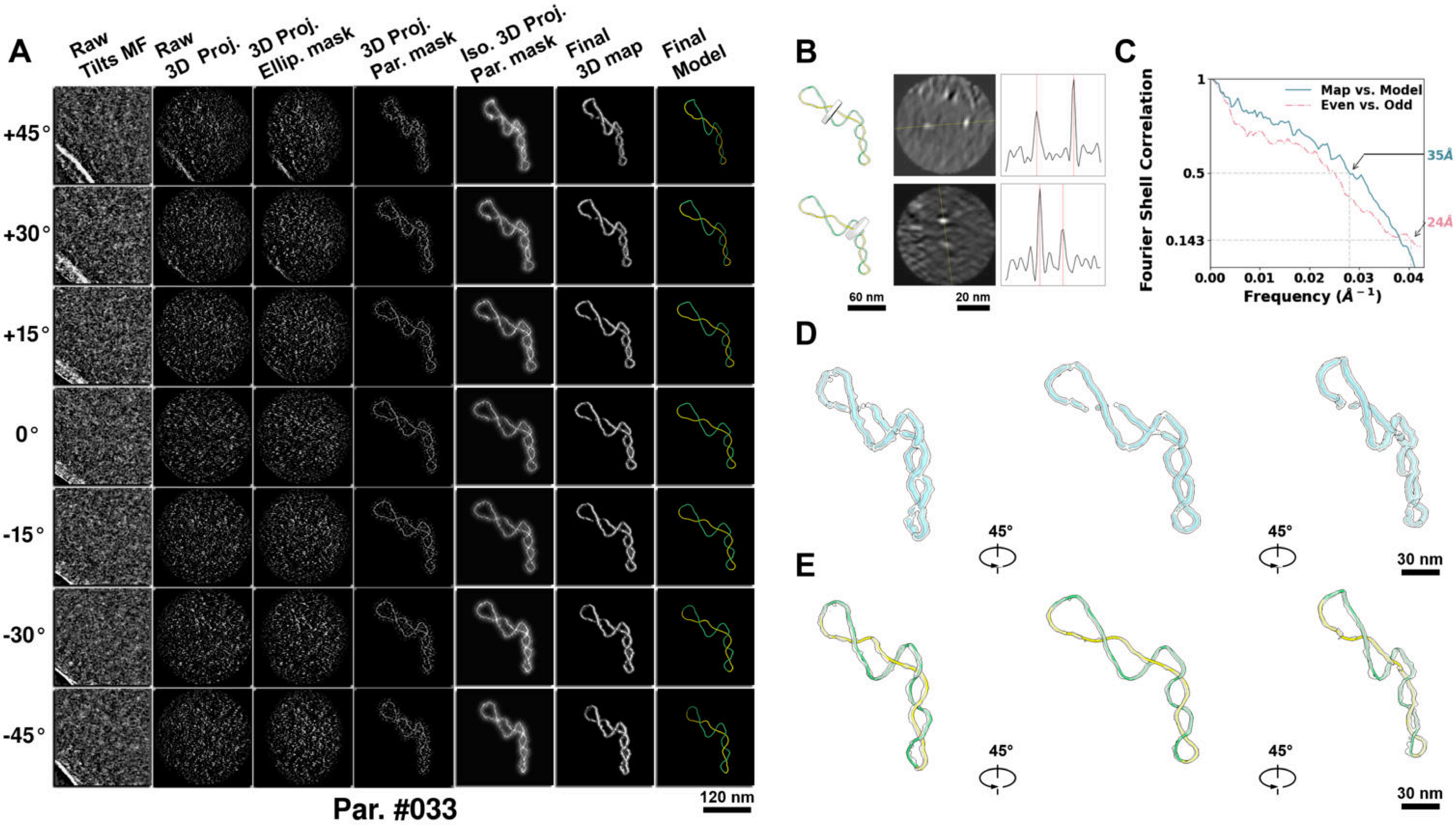

**Supplementary Particle Figure 33. Cryo-ET 3D reconstruction of an individual P.LS particle.**
(A) 3D reconstruction of the plasmid particle (index no. 33). The first column shows seven representative tilt images from +45° to -45° in step of 15°. The second, third, and fourth columns show 3D projections of the particle with spherical, ellipsoidal (thinner along the z-dimension), and particle-shaped masks, respectively. The fifth column displays the 3D projections of the enhanced and IsoNet missing-wedge-corrected particle. The sixth and seventh columns present the final 3D map and the flexibly fitted model, respectively. (B) Two cross-sectional views (12 nm thickness) of the plasmid density map along its plectoneme axis are shown in the left-middle panel. The intensity profile along the line crossing the two high-density DNA spots is displayed in the right panel. (C) Resolution assessment of the final 3D map using Fourier shell correlation (FSC). Two criteria are shown: FSC between two half-maps reconstructed from even and odd frames (evaluated at 0.143) and FSC between the final 3D map and the fitted model (evaluated at 0.5). (D) Zoomed-in views of the final 3D density map from panel A, displayed at two contour levels. (E) Superimposition of the high-contour level map from panel D onto its fitted model.

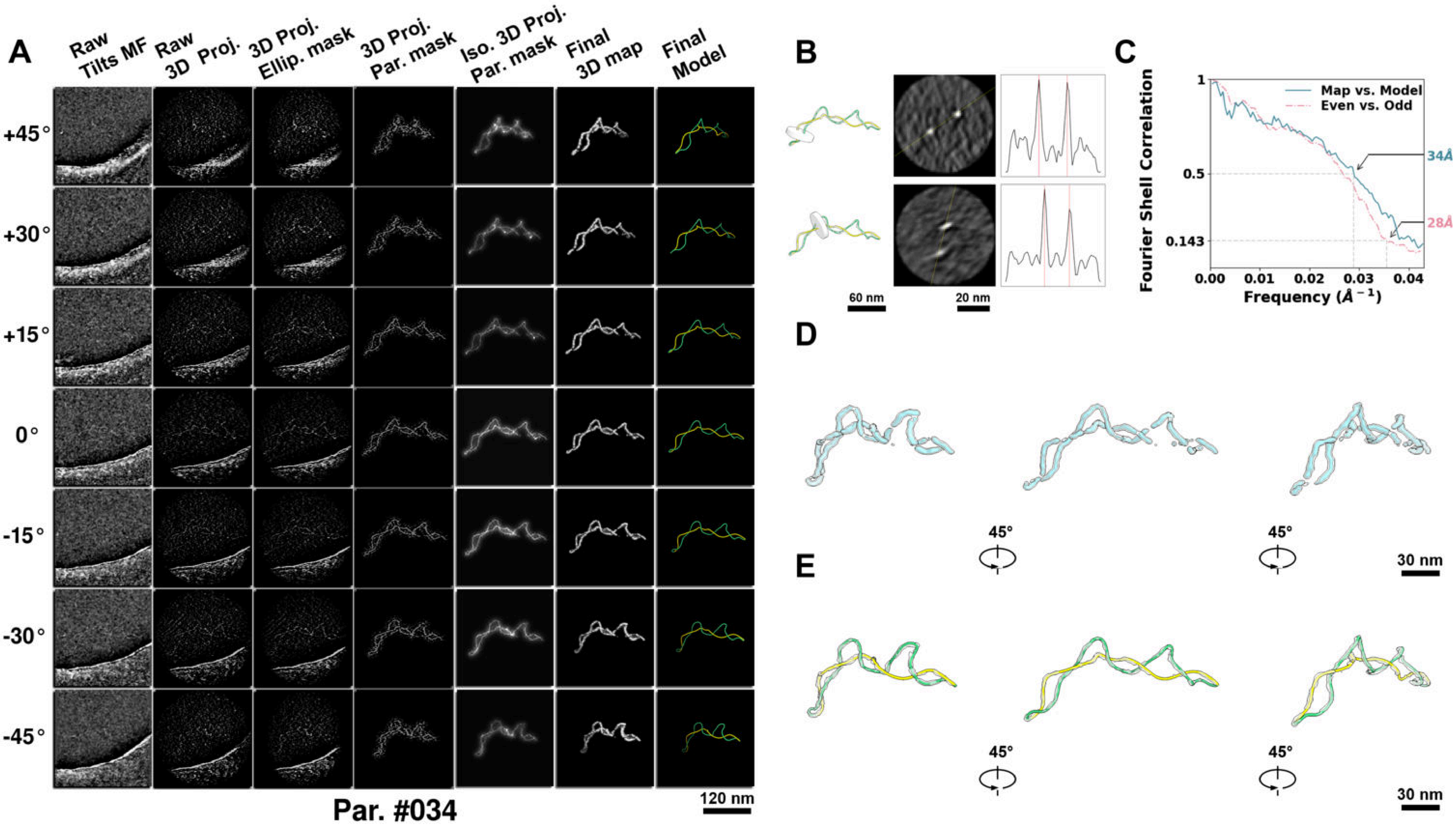

**Supplementary Particle Figure 34. Cryo-ET 3D reconstruction of an individual P.LS particle.**
(A) 3D reconstruction of the plasmid particle (index no. 34). The first column shows seven representative tilt images from +45° to -45° in step of 15°. The second, third, and fourth columns show 3D projections of the particle with spherical, ellipsoidal (thinner along the z-dimension), and particle-shaped masks, respectively. The fifth column displays the 3D projections of the enhanced and IsoNet missing-wedge-corrected particle. The sixth and seventh columns present the final 3D map and the flexibly fitted model, respectively. (B) Two cross-sectional views (12 nm thickness) of the plasmid density map along its plectoneme axis are shown in the left-middle panel. The intensity profile along the line crossing the two high-density DNA spots is displayed in the right panel. (C) Resolution assessment of the final 3D map using Fourier shell correlation (FSC). Two criteria are shown: FSC between two half-maps reconstructed from even and odd frames (evaluated at 0.143) and FSC between the final 3D map and the fitted model (evaluated at 0.5). (D) Zoomed-in views of the final 3D density map from panel A, displayed at two contour levels. (E) Superimposition of the high-contour level map from panel D onto its fitted model.

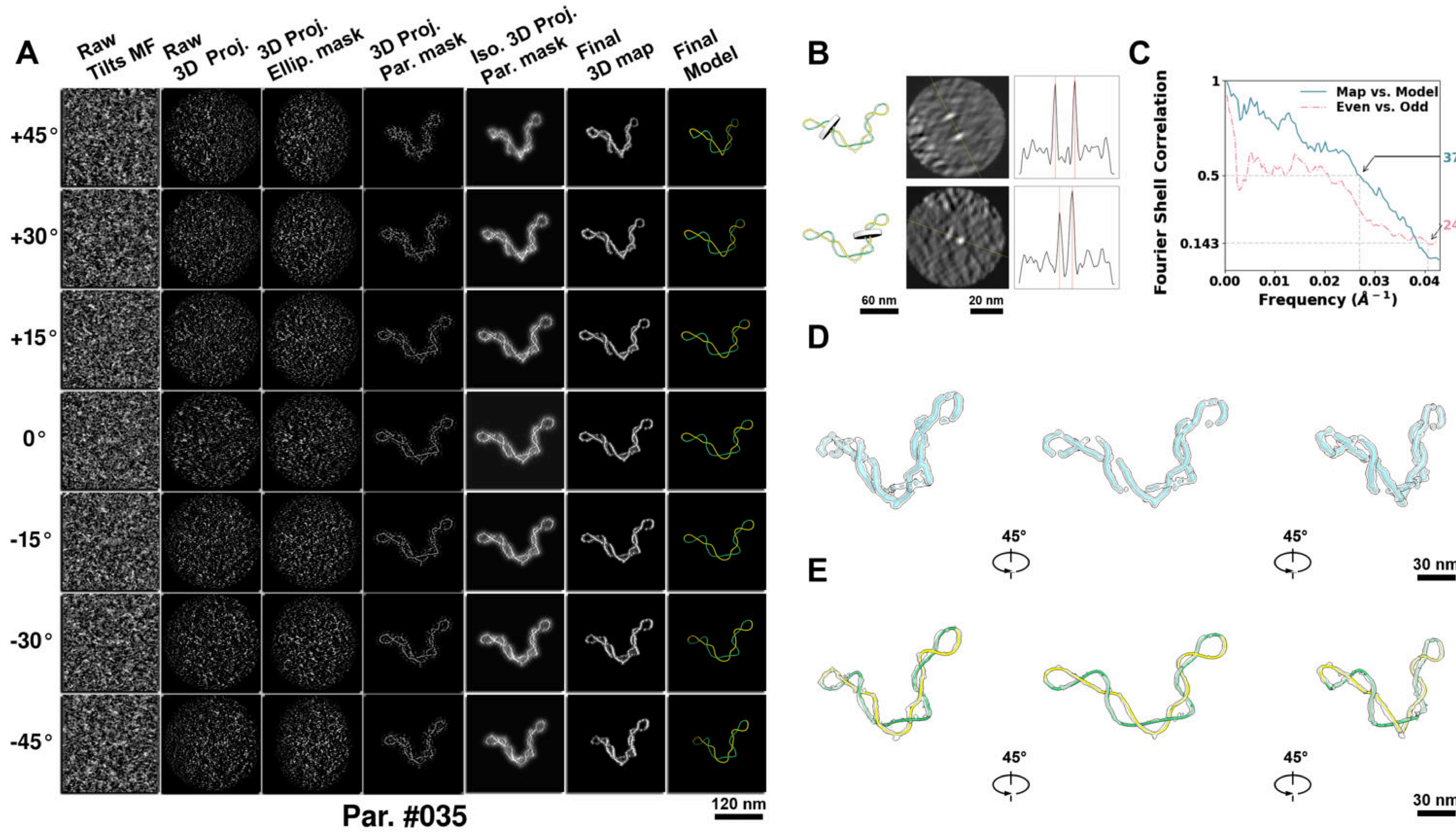

**Supplementary Particle Figure 35. Cryo-ET 3D reconstruction of an individual P.LS particle.**
(A) 3D reconstruction of the plasmid particle (index no. 35). The first column shows seven representative tilt images from +45° to -45° in step of 15°. The second, third, and fourth columns show 3D projections of the particle with spherical, ellipsoidal (thinner along the z-dimension), and particle-shaped masks, respectively. The fifth column displays the 3D projections of the enhanced and IsoNet missing-wedge-corrected particle. The sixth and seventh columns present the final 3D map and the flexibly fitted model, respectively. (B) Two cross-sectional views (12 nm thickness) of the plasmid density map along its plectoneme axis are shown in the left-middle panel. The intensity profile along the line crossing the two high-density DNA spots is displayed in the right panel. (C) Resolution assessment of the final 3D map using Fourier shell correlation (FSC). Two criteria are shown: FSC between two half-maps reconstructed from even and odd frames (evaluated at 0.143) and FSC between the final 3D map and the fitted model (evaluated at 0.5). (D) Zoomed-in views of the final 3D density map from panel A, displayed at two contour levels. (E) Superimposition of the high-contour level map from panel D onto its fitted model.

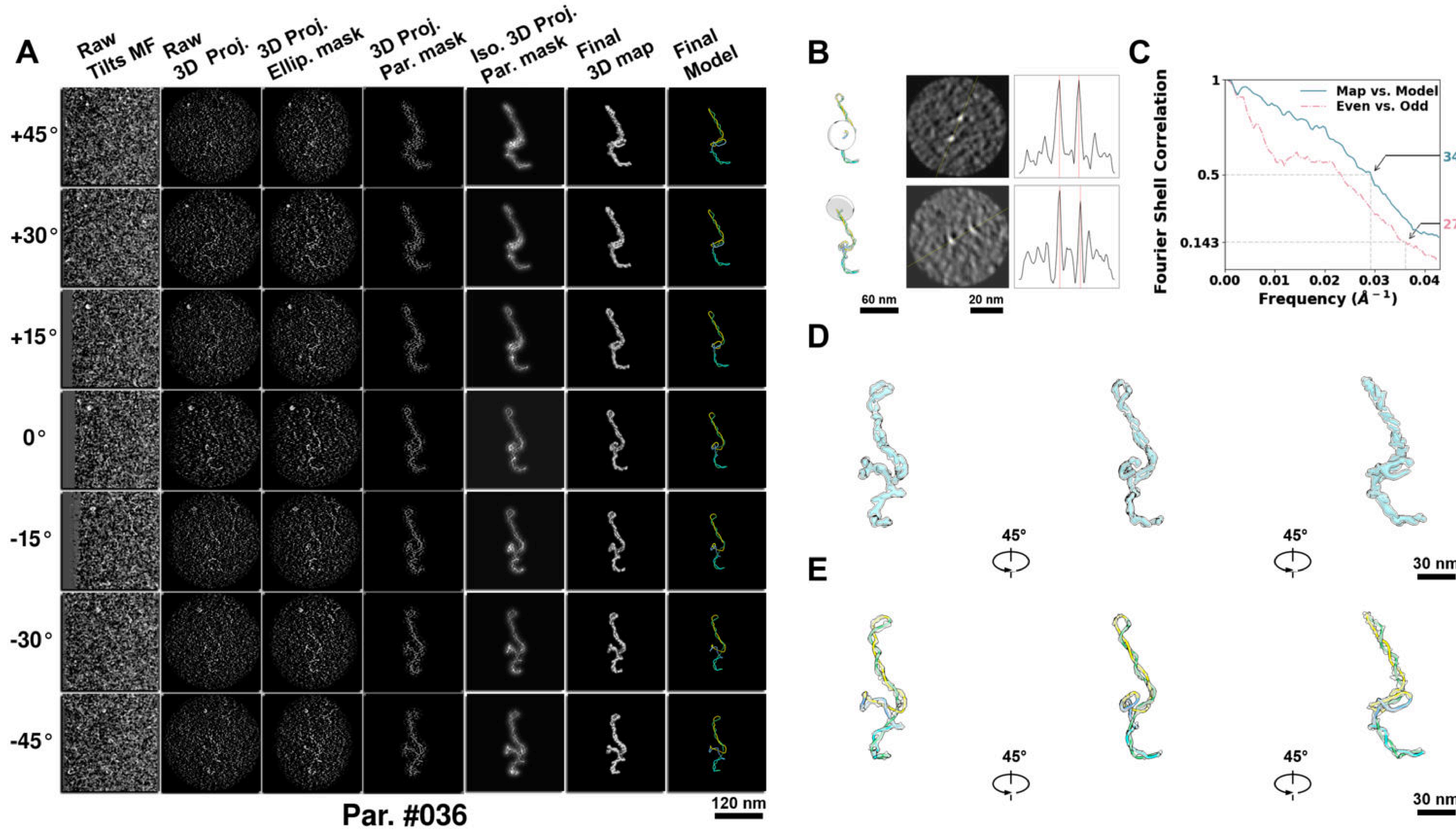

**Supplementary Particle Figure 36. Cryo-ET 3D reconstruction of an individual P.HS particle.**
(A) 3D reconstruction of the plasmid particle (index no. 36). The first column shows seven representative tilt images from +45° to -45° in step of 15°. The second, third, and fourth columns show 3D projections of the particle with spherical, ellipsoidal (thinner along the z-dimension), and particle-shaped masks, respectively. The fifth column displays the 3D projections of the enhanced and IsoNet missing-wedge-corrected particle. The sixth and seventh columns present the final 3D map and the flexibly fitted model, respectively. (B) Two cross-sectional views (12 nm thickness) of the plasmid density map along its plectoneme axis are shown in the left-middle panel. The intensity profile along the line crossing the two high-density DNA spots is displayed in the right panel. (C) Resolution assessment of the final 3D map using Fourier shell correlation (FSC). Two criteria are shown: FSC between two half-maps reconstructed from even and odd frames (evaluated at 0.143) and FSC between the final 3D map and the fitted model (evaluated at 0.5). (D) Zoomed-in views of the final 3D density map from panel A, displayed at two contour levels. (E) Superimposition of the high-contour level map from panel D onto its fitted model.

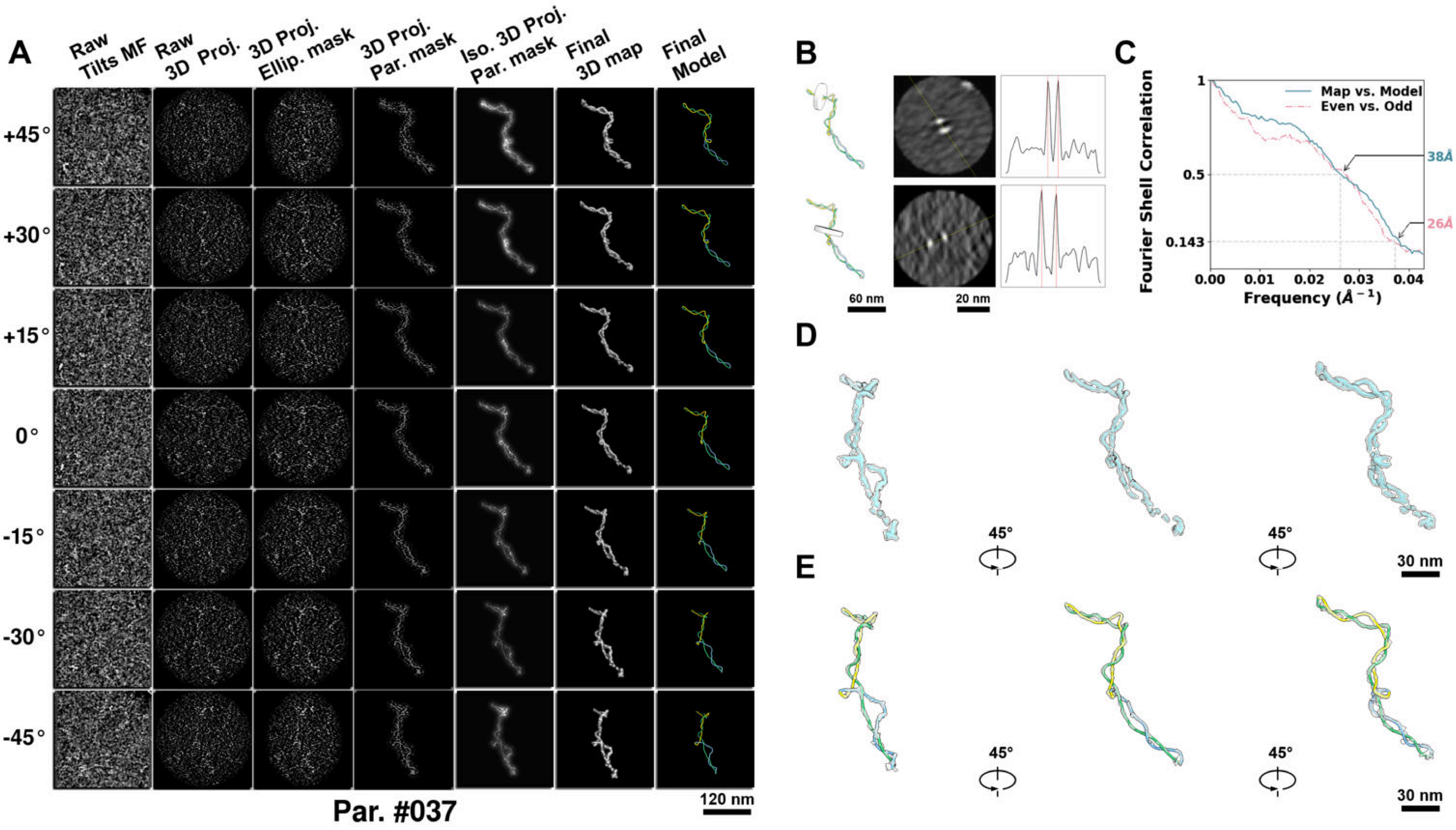

**Supplementary Particle Figure 37. Cryo-ET 3D reconstruction of an individual P.HS particle.**
(A) 3D reconstruction of the plasmid particle (index no. 37). The first column shows seven representative tilt images from +45° to -45° in step of 15°. The second, third, and fourth columns show 3D projections of the particle with spherical, ellipsoidal (thinner along the z-dimension), and particle-shaped masks, respectively. The fifth column displays the 3D projections of the enhanced and IsoNet missing-wedge-corrected particle. The sixth and seventh columns present the final 3D map and the flexibly fitted model, respectively. (B) Two cross-sectional views (12 nm thickness) of the plasmid density map along its plectoneme axis are shown in the left-middle panel. The intensity profile along the line crossing the two high-density DNA spots is displayed in the right panel. (C) Resolution assessment of the final 3D map using Fourier shell correlation (FSC). Two criteria are shown: FSC between two half-maps reconstructed from even and odd frames (evaluated at 0.143) and FSC between the final 3D map and the fitted model (evaluated at 0.5). (D) Zoomed-in views of the final 3D density map from panel A, displayed at two contour levels. (E) Superimposition of the high-contour level map from panel D onto its fitted model.

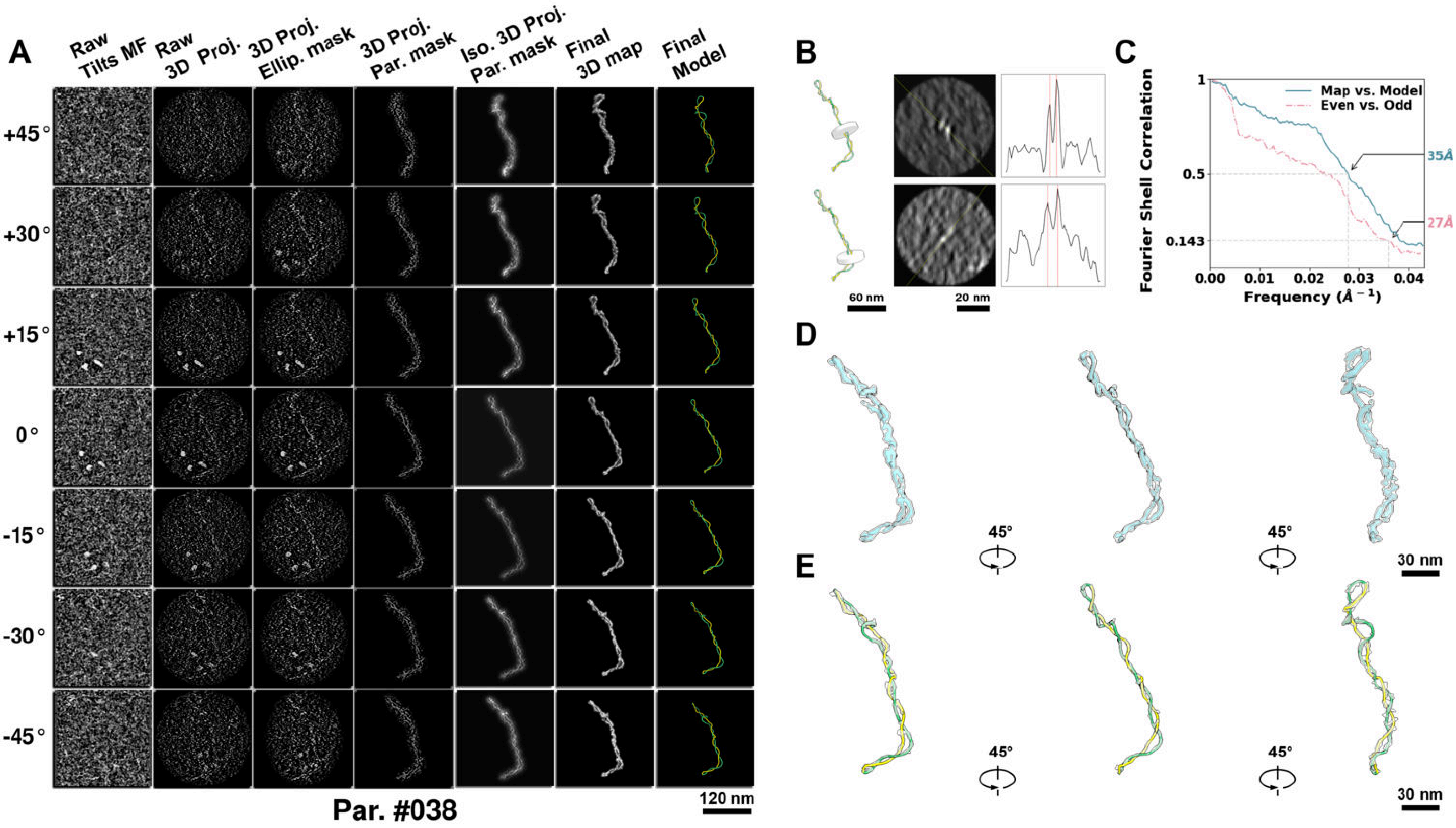

**Supplementary Particle Figure 38. Cryo-ET 3D reconstruction of an individual P.HS particle.**
(A) 3D reconstruction of the plasmid particle (index no. 38). The first column shows seven representative tilt images from +45° to -45° in step of 15°. The second, third, and fourth columns show 3D projections of the particle with spherical, ellipsoidal (thinner along the z-dimension), and particle-shaped masks, respectively. The fifth column displays the 3D projections of the enhanced and IsoNet missing-wedge-corrected particle. The sixth and seventh columns present the final 3D map and the flexibly fitted model, respectively. (B) Two cross-sectional views (12 nm thickness) of the plasmid density map along its plectoneme axis are shown in the left-middle panel. The intensity profile along the line crossing the two high-density DNA spots is displayed in the right panel. (C) Resolution assessment of the final 3D map using Fourier shell correlation (FSC). Two criteria are shown: FSC between two half-maps reconstructed from even and odd frames (evaluated at 0.143) and FSC between the final 3D map and the fitted model (evaluated at 0.5). (D) Zoomed-in views of the final 3D density map from panel A, displayed at two contour levels. (E) Superimposition of the high-contour level map from panel D onto its fitted model.

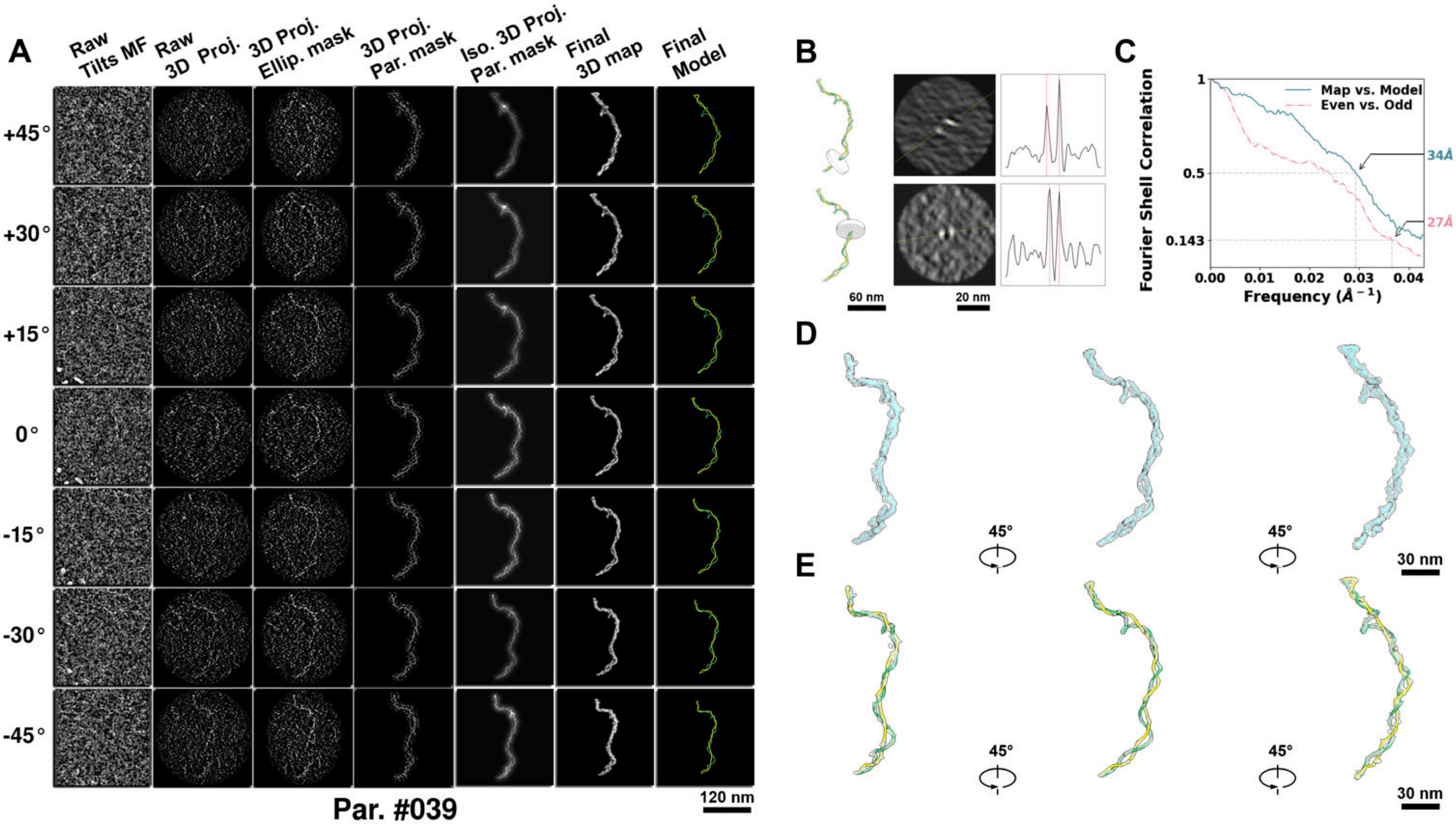

**Supplementary Particle Figure 39. Cryo-ET 3D reconstruction of an individual P.HS particle.**
(A) 3D reconstruction of the plasmid particle (index no. 39). The first column shows seven representative tilt images from +45° to -45° in step of 15°. The second, third, and fourth columns show 3D projections of the particle with spherical, ellipsoidal (thinner along the z-dimension), and particle-shaped masks, respectively. The fifth column displays the 3D projections of the enhanced and IsoNet missing-wedge-corrected particle. The sixth and seventh columns present the final 3D map and the flexibly fitted model, respectively. (B) Two cross-sectional views (12 nm thickness) of the plasmid density map along its plectoneme axis are shown in the left-middle panel. The intensity profile along the line crossing the two high-density DNA spots is displayed in the right panel. (C) Resolution assessment of the final 3D map using Fourier shell correlation (FSC). Two criteria are shown: FSC between two half-maps reconstructed from even and odd frames (evaluated at 0.143) and FSC between the final 3D map and the fitted model (evaluated at 0.5). (D) Zoomed-in views of the final 3D density map from panel A, displayed at two contour levels. (E) Superimposition of the high-contour level map from panel D onto its fitted model.

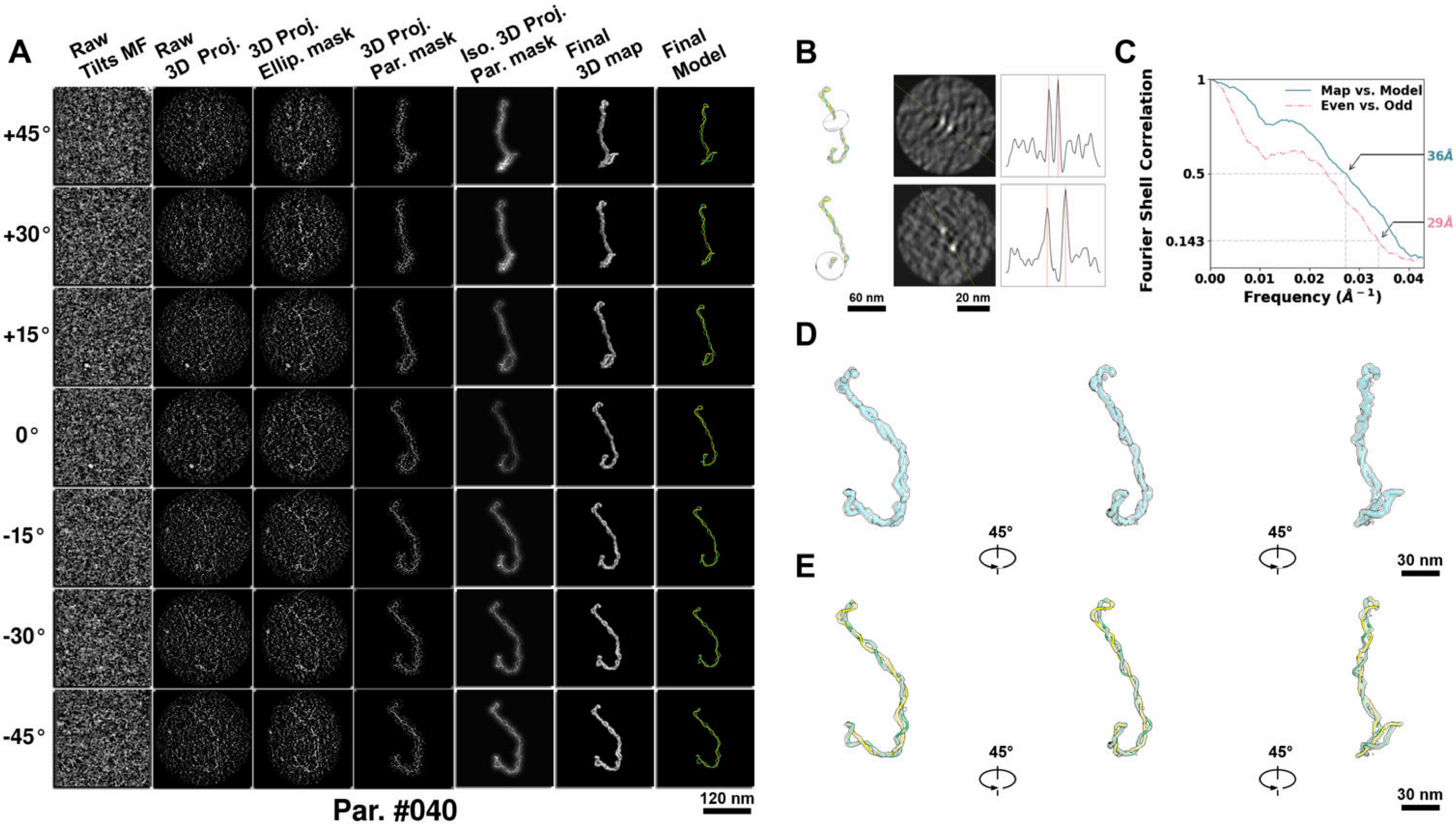

**Supplementary Particle Figure 40. Cryo-ET 3D reconstruction of an individual P.HS particle.**
(A) 3D reconstruction of the plasmid particle (index no. 40). The first column shows seven representative tilt images from +45° to -45° in step of 15°. The second, third, and fourth columns show 3D projections of the particle with spherical, ellipsoidal (thinner along the z-dimension), and particle-shaped masks, respectively. The fifth column displays the 3D projections of the enhanced and IsoNet missing-wedge-corrected particle. The sixth and seventh columns present the final 3D map and the flexibly fitted model, respectively. (B) Two cross-sectional views (12 nm thickness) of the plasmid density map along its plectoneme axis are shown in the left-middle panel. The intensity profile along the line crossing the two high-density DNA spots is displayed in the right panel. (C) Resolution assessment of the final 3D map using Fourier shell correlation (FSC). Two criteria are shown: FSC between two half-maps reconstructed from even and odd frames (evaluated at 0.143) and FSC between the final 3D map and the fitted model (evaluated at 0.5). (D) Zoomed-in views of the final 3D density map from panel A, displayed at two contour levels. (E) Superimposition of the high-contour level map from panel D onto its fitted model.

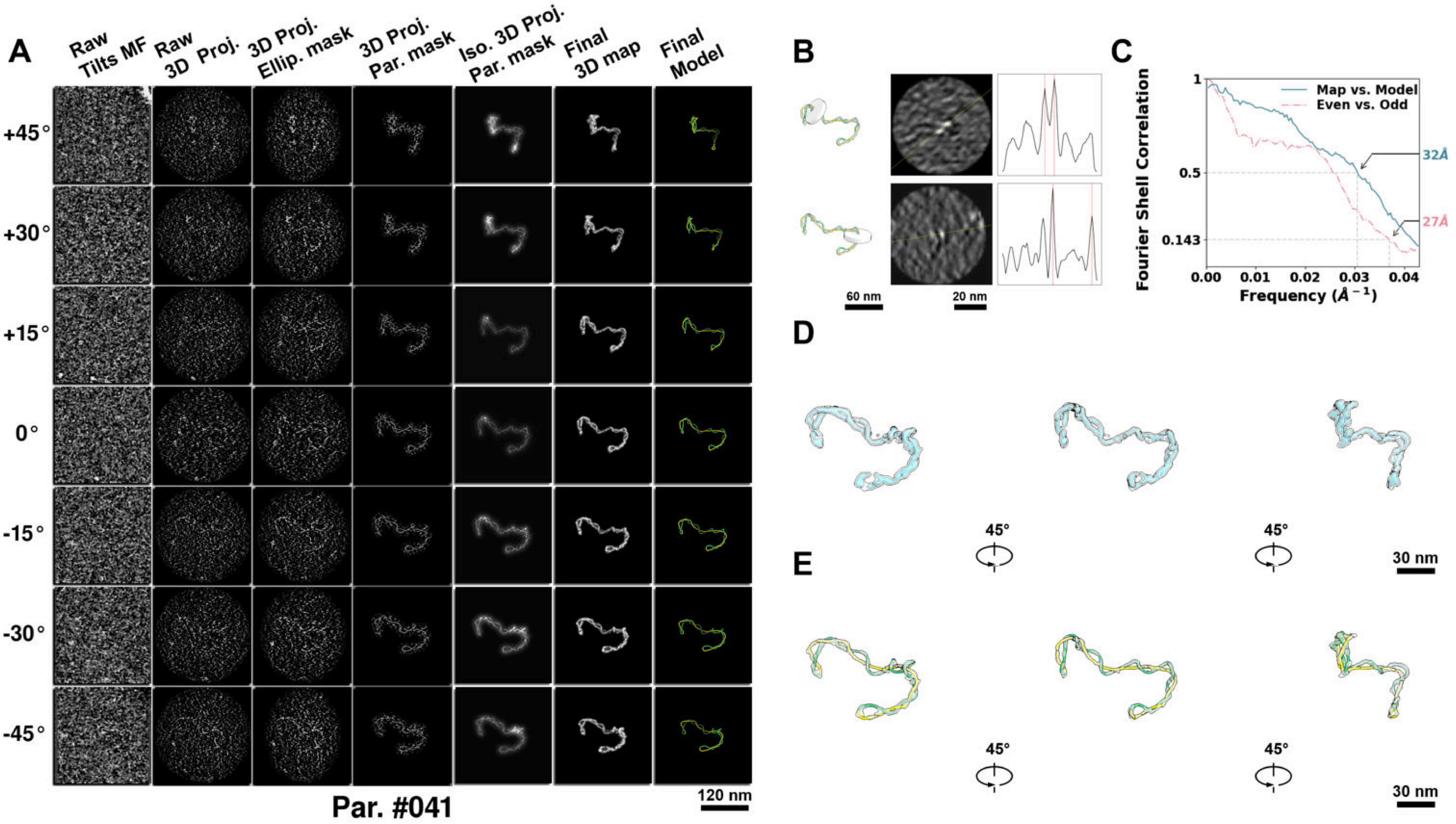

**Supplementary Particle Figure 41. Cryo-ET 3D reconstruction of an individual P.HS particle.**
(A) 3D reconstruction of the plasmid particle (index no. 41). The first column shows seven representative tilt images from +45° to -45° in step of 15°. The second, third, and fourth columns show 3D projections of the particle with spherical, ellipsoidal (thinner along the z-dimension), and particle-shaped masks, respectively. The fifth column displays the 3D projections of the enhanced and IsoNet missing-wedge-corrected particle. The sixth and seventh columns present the final 3D map and the flexibly fitted model, respectively. (B) Two cross-sectional views (12 nm thickness) of the plasmid density map along its plectoneme axis are shown in the left-middle panel. The intensity profile along the line crossing the two high-density DNA spots is displayed in the right panel. (C) Resolution assessment of the final 3D map using Fourier shell correlation (FSC). Two criteria are shown: FSC between two half-maps reconstructed from even and odd frames (evaluated at 0.143) and FSC between the final 3D map and the fitted model (evaluated at 0.5). (D) Zoomed-in views of the final 3D density map from panel A, displayed at two contour levels. (E) Superimposition of the high-contour level map from panel D onto its fitted model.

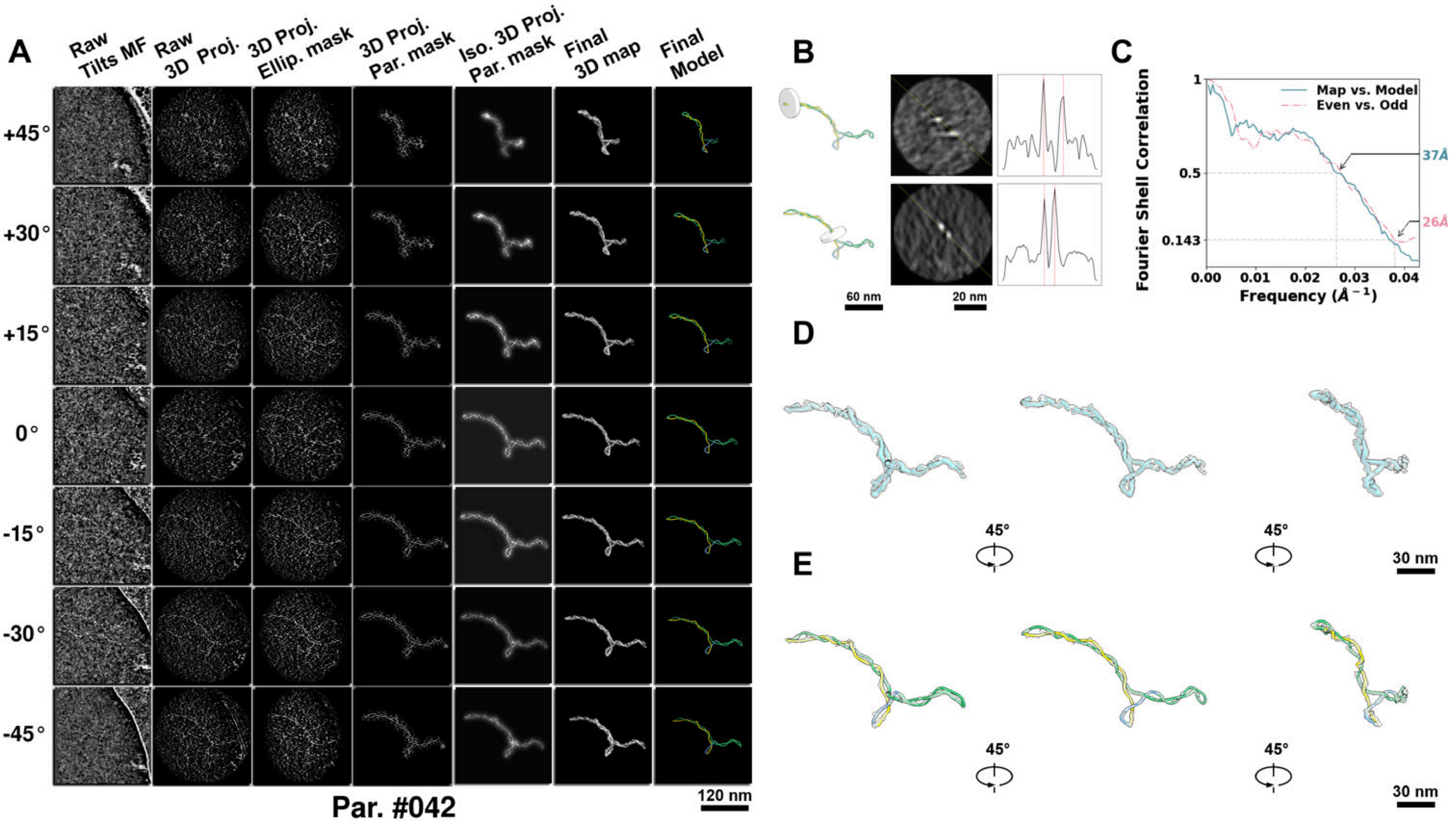

**Supplementary Particle Figure 42. Cryo-ET 3D reconstruction of an individual P.HS particle.**
(A) 3D reconstruction of the plasmid particle (index no. 42). The first column shows seven representative tilt images from +45° to -45° in step of 15°. The second, third, and fourth columns show 3D projections of the particle with spherical, ellipsoidal (thinner along the z-dimension), and particle-shaped masks, respectively. The fifth column displays the 3D projections of the enhanced and IsoNet missing-wedge-corrected particle. The sixth and seventh columns present the final 3D map and the flexibly fitted model, respectively. (B) Two cross-sectional views (12 nm thickness) of the plasmid density map along its plectoneme axis are shown in the left-middle panel. The intensity profile along the line crossing the two high-density DNA spots is displayed in the right panel. (C) Resolution assessment of the final 3D map using Fourier shell correlation (FSC). Two criteria are shown: FSC between two half-maps reconstructed from even and odd frames (evaluated at 0.143) and FSC between the final 3D map and the fitted model (evaluated at 0.5). (D) Zoomed-in views of the final 3D density map from panel A, displayed at two contour levels. (E) Superimposition of the high-contour level map from panel D onto its fitted model.

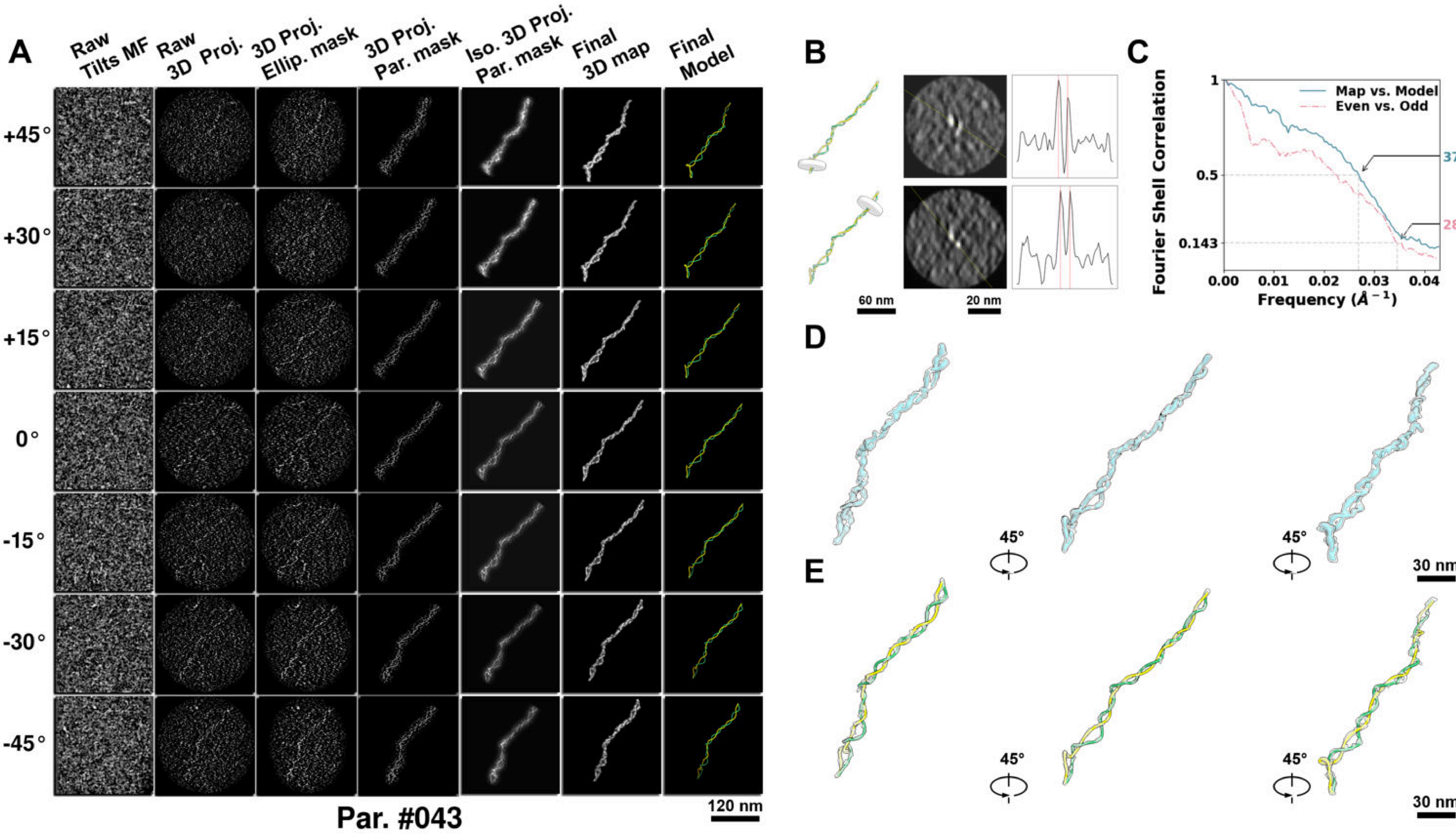

**Supplementary Particle Figure 43. Cryo-ET 3D reconstruction of an individual P.HS particle.**
(A) 3D reconstruction of the plasmid particle (index no. 43). The first column shows seven representative tilt images from +45° to -45° in step of 15°. The second, third, and fourth columns show 3D projections of the particle with spherical, ellipsoidal (thinner along the z-dimension), and particle-shaped masks, respectively. The fifth column displays the 3D projections of the enhanced and IsoNet missing-wedge-corrected particle. The sixth and seventh columns present the final 3D map and the flexibly fitted model, respectively. (B) Two cross-sectional views (12 nm thickness) of the plasmid density map along its plectoneme axis are shown in the left-middle panel. The intensity profile along the line crossing the two high-density DNA spots is displayed in the right panel. (C) Resolution assessment of the final 3D map using Fourier shell correlation (FSC). Two criteria are shown: FSC between two half-maps reconstructed from even and odd frames (evaluated at 0.143) and FSC between the final 3D map and the fitted model (evaluated at 0.5). (D) Zoomed-in views of the final 3D density map from panel A, displayed at two contour levels. (E) Superimposition of the high-contour level map from panel D onto its fitted model.

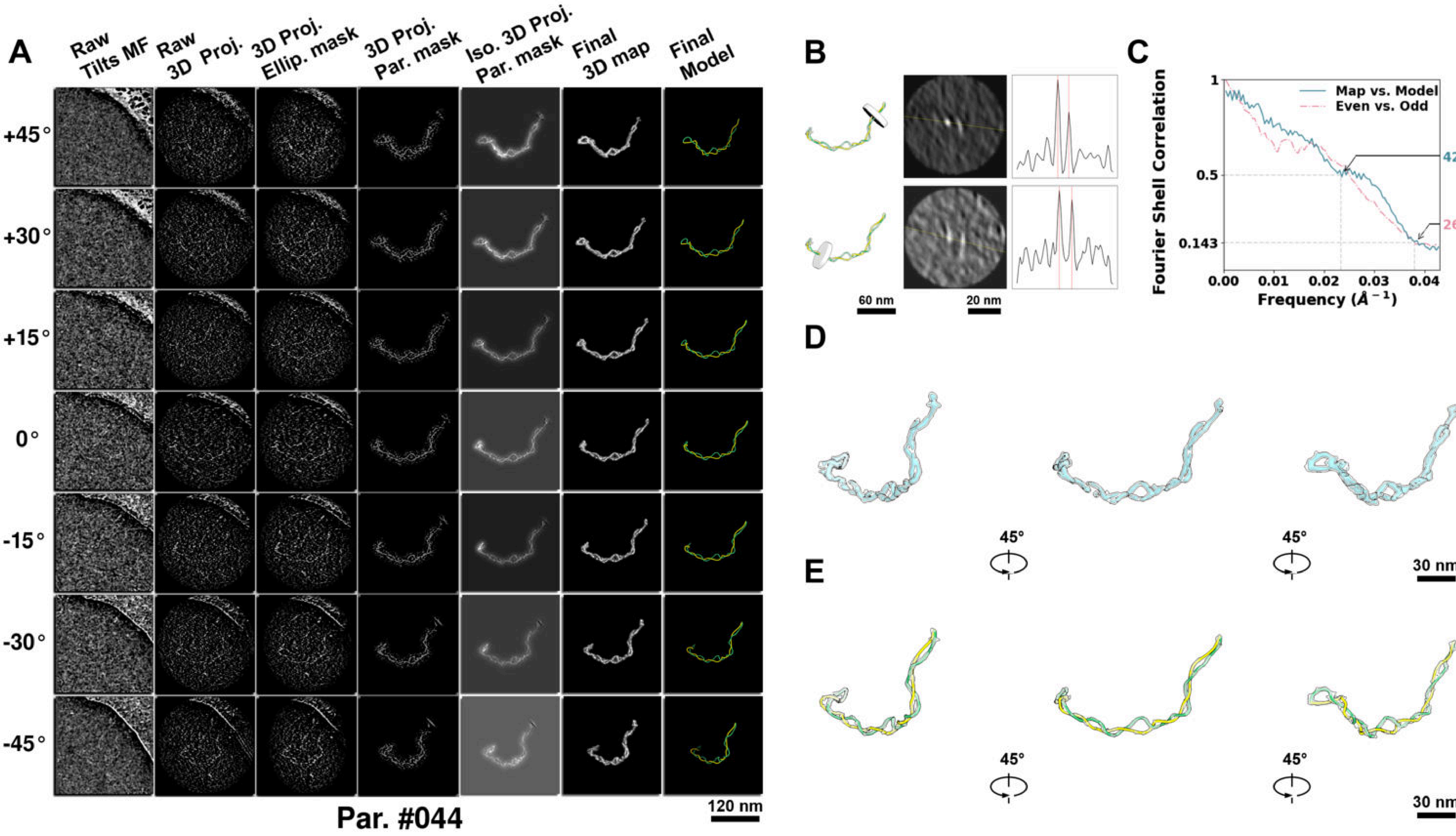

**Supplementary Particle Figure 44. Cryo-ET 3D reconstruction of an individual P.HS particle.**
(A) 3D reconstruction of the plasmid particle (index no. 44). The first column shows seven representative tilt images from +45° to -45° in step of 15°. The second, third, and fourth columns show 3D projections of the particle with spherical, ellipsoidal (thinner along the z-dimension), and particle-shaped masks, respectively. The fifth column displays the 3D projections of the enhanced and IsoNet missing-wedge-corrected particle. The sixth and seventh columns present the final 3D map and the flexibly fitted model, respectively. (B) Two cross-sectional views (12 nm thickness) of the plasmid density map along its plectoneme axis are shown in the left-middle panel. The intensity profile along the line crossing the two high-density DNA spots is displayed in the right panel. (C) Resolution assessment of the final 3D map using Fourier shell correlation (FSC). Two criteria are shown: FSC between two half-maps reconstructed from even and odd frames (evaluated at 0.143) and FSC between the final 3D map and the fitted model (evaluated at 0.5). (D) Zoomed-in views of the final 3D density map from panel A, displayed at two contour levels. (E) Superimposition of the high-contour level map from panel D onto its fitted model.

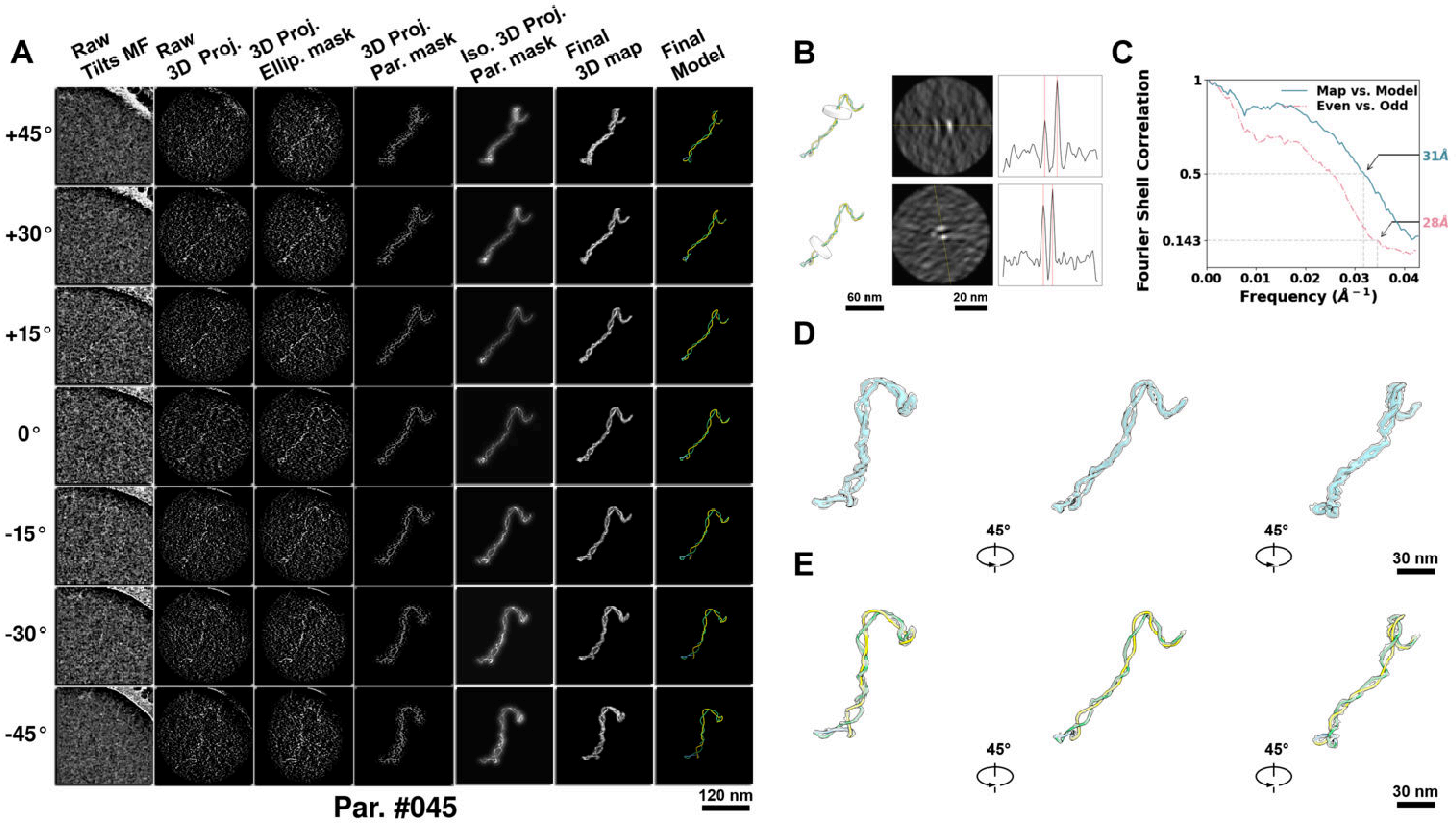

**Supplementary Particle Figure 45. Cryo-ET 3D reconstruction of an individual P.HS particle.**
(A) 3D reconstruction of the plasmid particle (index no. 45). The first column shows seven representative tilt images from +45° to -45° in step of 15°. The second, third, and fourth columns show 3D projections of the particle with spherical, ellipsoidal (thinner along the z-dimension), and particle-shaped masks, respectively. The fifth column displays the 3D projections of the enhanced and IsoNet missing-wedge-corrected particle. The sixth and seventh columns present the final 3D map and the flexibly fitted model, respectively. (B) Two cross-sectional views (12 nm thickness) of the plasmid density map along its plectoneme axis are shown in the left-middle panel. The intensity profile along the line crossing the two high-density DNA spots is displayed in the right panel. (C) Resolution assessment of the final 3D map using Fourier shell correlation (FSC). Two criteria are shown: FSC between two half-maps reconstructed from even and odd frames (evaluated at 0.143) and FSC between the final 3D map and the fitted model (evaluated at 0.5). (D) Zoomed-in views of the final 3D density map from panel A, displayed at two contour levels. (E) Superimposition of the high-contour level map from panel D onto its fitted model.

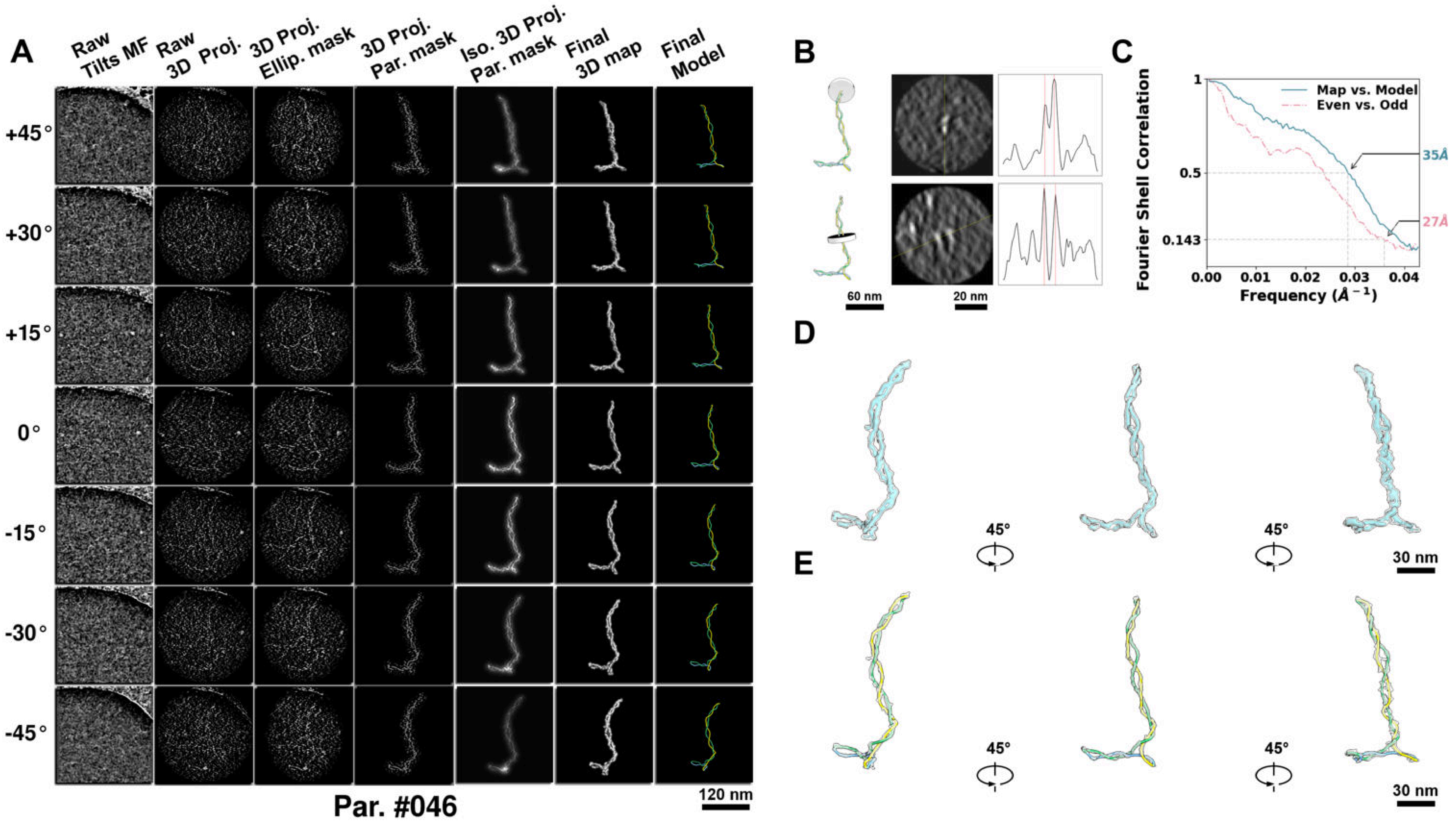

**Supplementary Particle Figure 46. Cryo-ET 3D reconstruction of an individual P.HS particle.**
(A) 3D reconstruction of the plasmid particle (index no. 46). The first column shows seven representative tilt images from +45° to -45° in step of 15°. The second, third, and fourth columns show 3D projections of the particle with spherical, ellipsoidal (thinner along the z-dimension), and particle-shaped masks, respectively. The fifth column displays the 3D projections of the enhanced and IsoNet missing-wedge-corrected particle. The sixth and seventh columns present the final 3D map and the flexibly fitted model, respectively. (B) Two cross-sectional views (12 nm thickness) of the plasmid density map along its plectoneme axis are shown in the left-middle panel. The intensity profile along the line crossing the two high-density DNA spots is displayed in the right panel. (C) Resolution assessment of the final 3D map using Fourier shell correlation (FSC). Two criteria are shown: FSC between two half-maps reconstructed from even and odd frames (evaluated at 0.143) and FSC between the final 3D map and the fitted model (evaluated at 0.5). (D) Zoomed-in views of the final 3D density map from panel A, displayed at two contour levels. (E) Superimposition of the high-contour level map from panel D onto its fitted model.

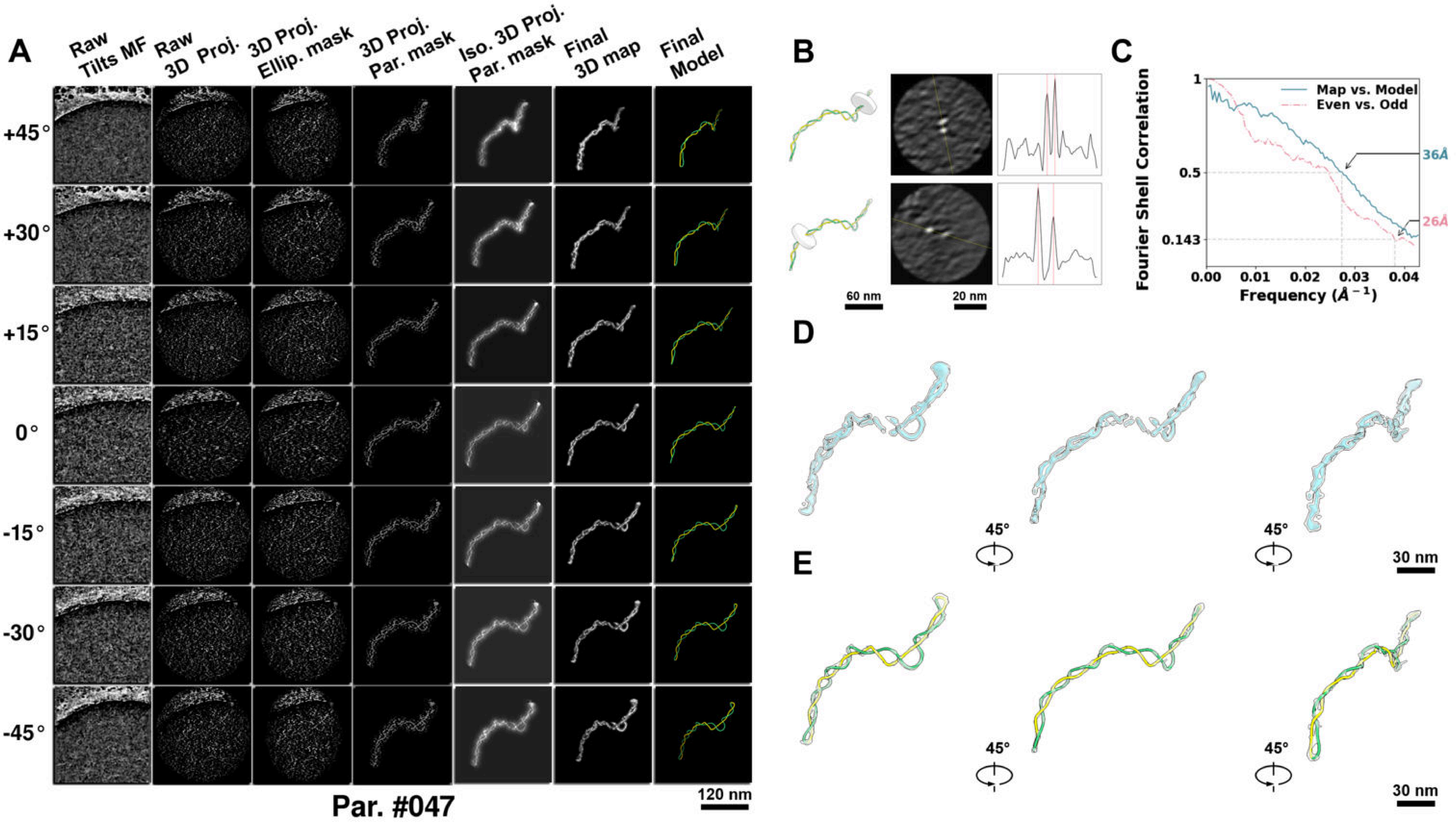

**Supplementary Particle Figure 47. Cryo-ET 3D reconstruction of an individual P.HS particle.**
(A) 3D reconstruction of the plasmid particle (index no. 47). The first column shows seven representative tilt images from +45° to -45° in step of 15°. The second, third, and fourth columns show 3D projections of the particle with spherical, ellipsoidal (thinner along the z-dimension), and particle-shaped masks, respectively. The fifth column displays the 3D projections of the enhanced and IsoNet missing-wedge-corrected particle. The sixth and seventh columns present the final 3D map and the flexibly fitted model, respectively. (B) Two cross-sectional views (12 nm thickness) of the plasmid density map along its plectoneme axis are shown in the left-middle panel. The intensity profile along the line crossing the two high-density DNA spots is displayed in the right panel. (C) Resolution assessment of the final 3D map using Fourier shell correlation (FSC). Two criteria are shown: FSC between two half-maps reconstructed from even and odd frames (evaluated at 0.143) and FSC between the final 3D map and the fitted model (evaluated at 0.5). (D) Zoomed-in views of the final 3D density map from panel A, displayed at two contour levels. (E) Superimposition of the high-contour level map from panel D onto its fitted model.

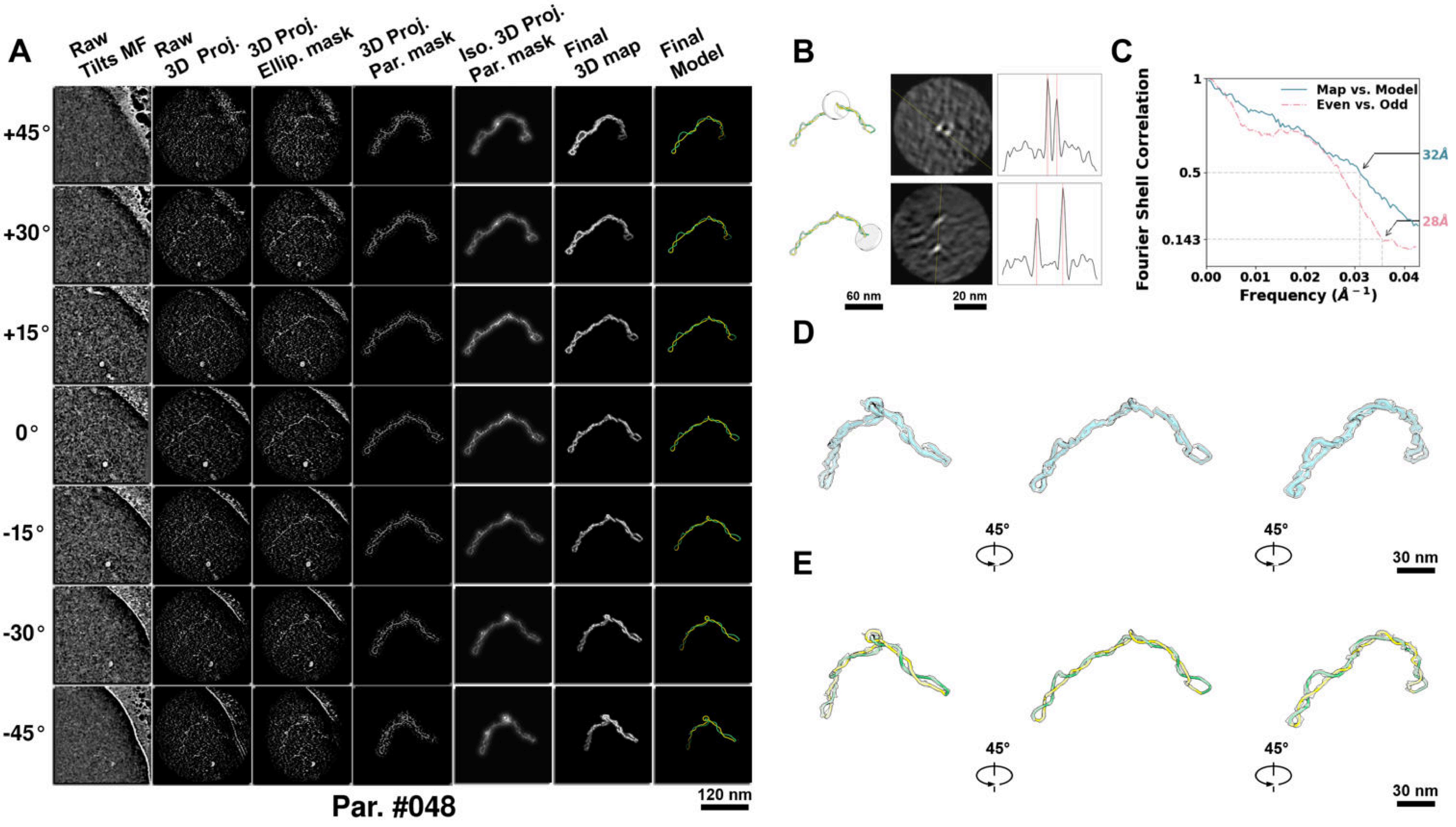

**Supplementary Particle Figure 48. Cryo-ET 3D reconstruction of an individual P.HS particle.**
(A) 3D reconstruction of the plasmid particle (index no. 48). The first column shows seven representative tilt images from +45° to -45° in step of 15°. The second, third, and fourth columns show 3D projections of the particle with spherical, ellipsoidal (thinner along the z-dimension), and particle-shaped masks, respectively. The fifth column displays the 3D projections of the enhanced and IsoNet missing-wedge-corrected particle. The sixth and seventh columns present the final 3D map and the flexibly fitted model, respectively. (B) Two cross-sectional views (12 nm thickness) of the plasmid density map along its plectoneme axis are shown in the left-middle panel. The intensity profile along the line crossing the two high-density DNA spots is displayed in the right panel. (C) Resolution assessment of the final 3D map using Fourier shell correlation (FSC). Two criteria are shown: FSC between two half-maps reconstructed from even and odd frames (evaluated at 0.143) and FSC between the final 3D map and the fitted model (evaluated at 0.5). (D) Zoomed-in views of the final 3D density map from panel A, displayed at two contour levels. (E) Superimposition of the high-contour level map from panel D onto its fitted model.

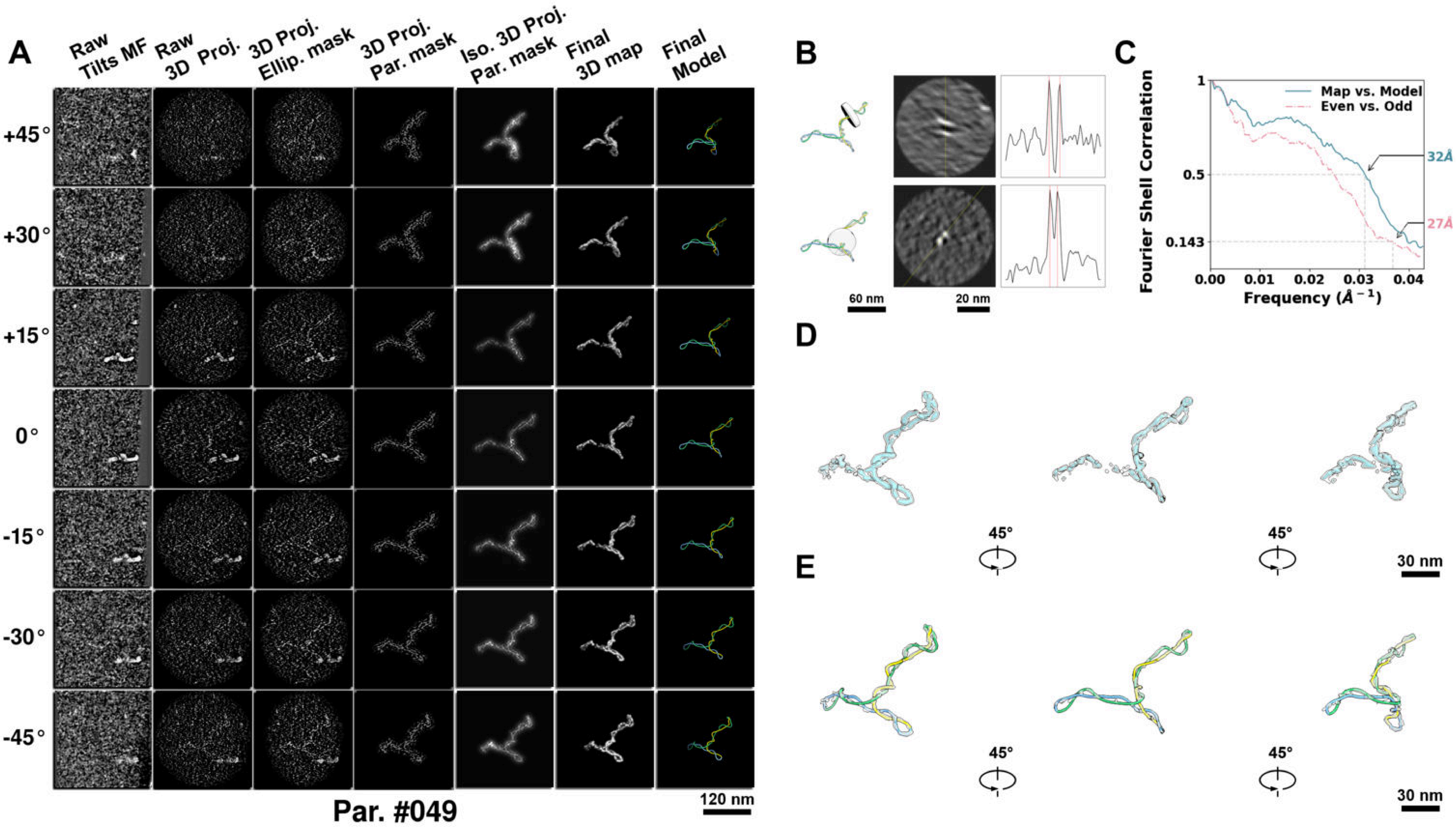

**Supplementary Particle Figure 49. Cryo-ET 3D reconstruction of an individual P.HS particle.**
(A) 3D reconstruction of the plasmid particle (index no. 49). The first column shows seven representative tilt images from +45° to -45° in step of 15°. The second, third, and fourth columns show 3D projections of the particle with spherical, ellipsoidal (thinner along the z-dimension), and particle-shaped masks, respectively. The fifth column displays the 3D projections of the enhanced and IsoNet missing-wedge-corrected particle. The sixth and seventh columns present the final 3D map and the flexibly fitted model, respectively. (B) Two cross-sectional views (12 nm thickness) of the plasmid density map along its plectoneme axis are shown in the left-middle panel. The intensity profile along the line crossing the two high-density DNA spots is displayed in the right panel. (C) Resolution assessment of the final 3D map using Fourier shell correlation (FSC). Two criteria are shown: FSC between two half-maps reconstructed from even and odd frames (evaluated at 0.143) and FSC between the final 3D map and the fitted model (evaluated at 0.5). (D) Zoomed-in views of the final 3D density map from panel A, displayed at two contour levels. (E) Superimposition of the high-contour level map from panel D onto its fitted model.

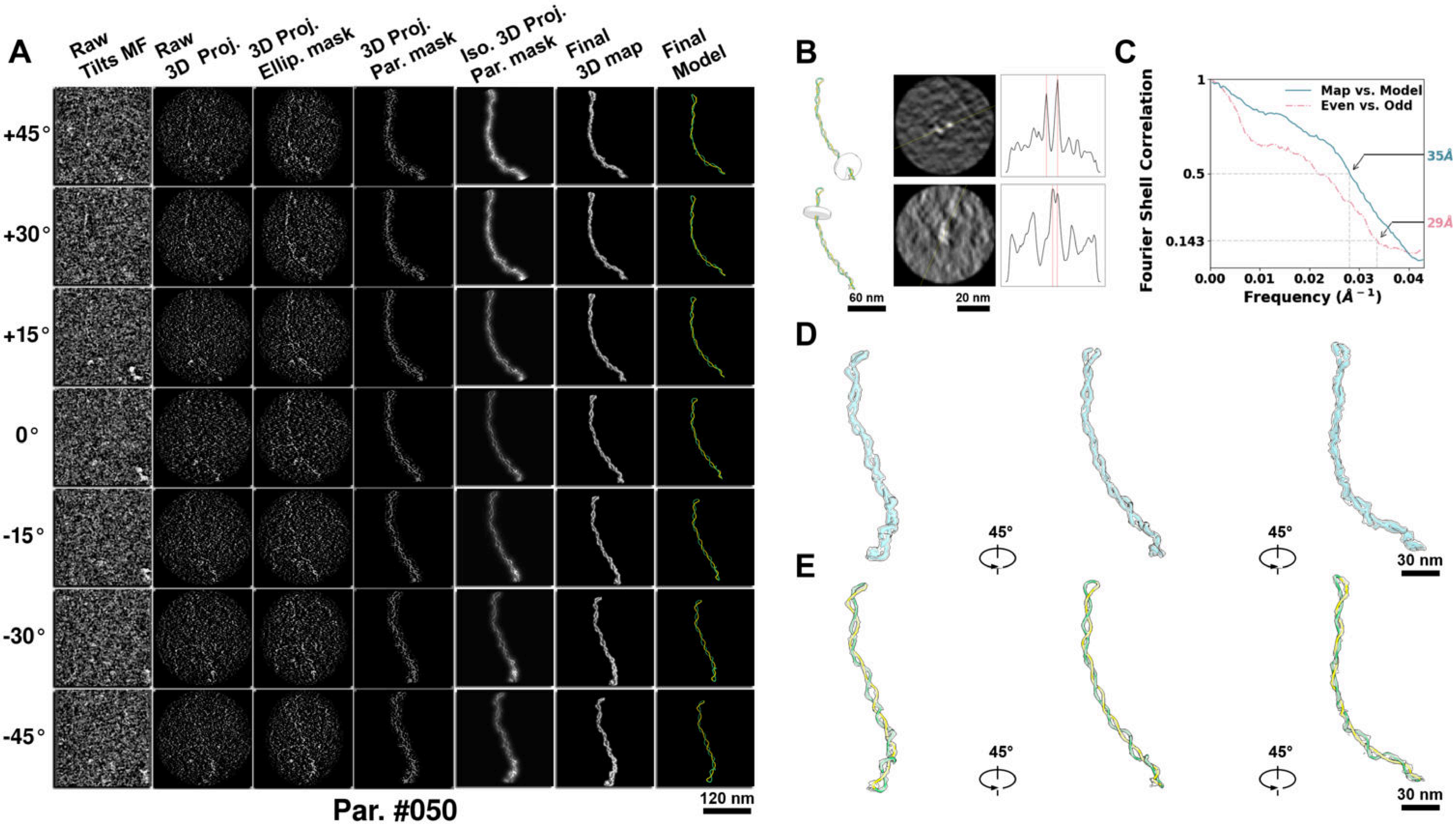

**Supplementary Particle Figure 50. Cryo-ET 3D reconstruction of an individual P.HS particle.**
(A) 3D reconstruction of the plasmid particle (index no. 50). The first column shows seven representative tilt images from +45° to -45° in step of 15°. The second, third, and fourth columns show 3D projections of the particle with spherical, ellipsoidal (thinner along the z-dimension), and particle-shaped masks, respectively. The fifth column displays the 3D projections of the enhanced and IsoNet missing-wedge-corrected particle. The sixth and seventh columns present the final 3D map and the flexibly fitted model, respectively. (B) Two cross-sectional views (12 nm thickness) of the plasmid density map along its plectoneme axis are shown in the left-middle panel. The intensity profile along the line crossing the two high-density DNA spots is displayed in the right panel. (C) Resolution assessment of the final 3D map using Fourier shell correlation (FSC). Two criteria are shown: FSC between two half-maps reconstructed from even and odd frames (evaluated at 0.143) and FSC between the final 3D map and the fitted model (evaluated at 0.5). (D) Zoomed-in views of the final 3D density map from panel A, displayed at two contour levels. (E) Superimposition of the high-contour level map from panel D onto its fitted model.

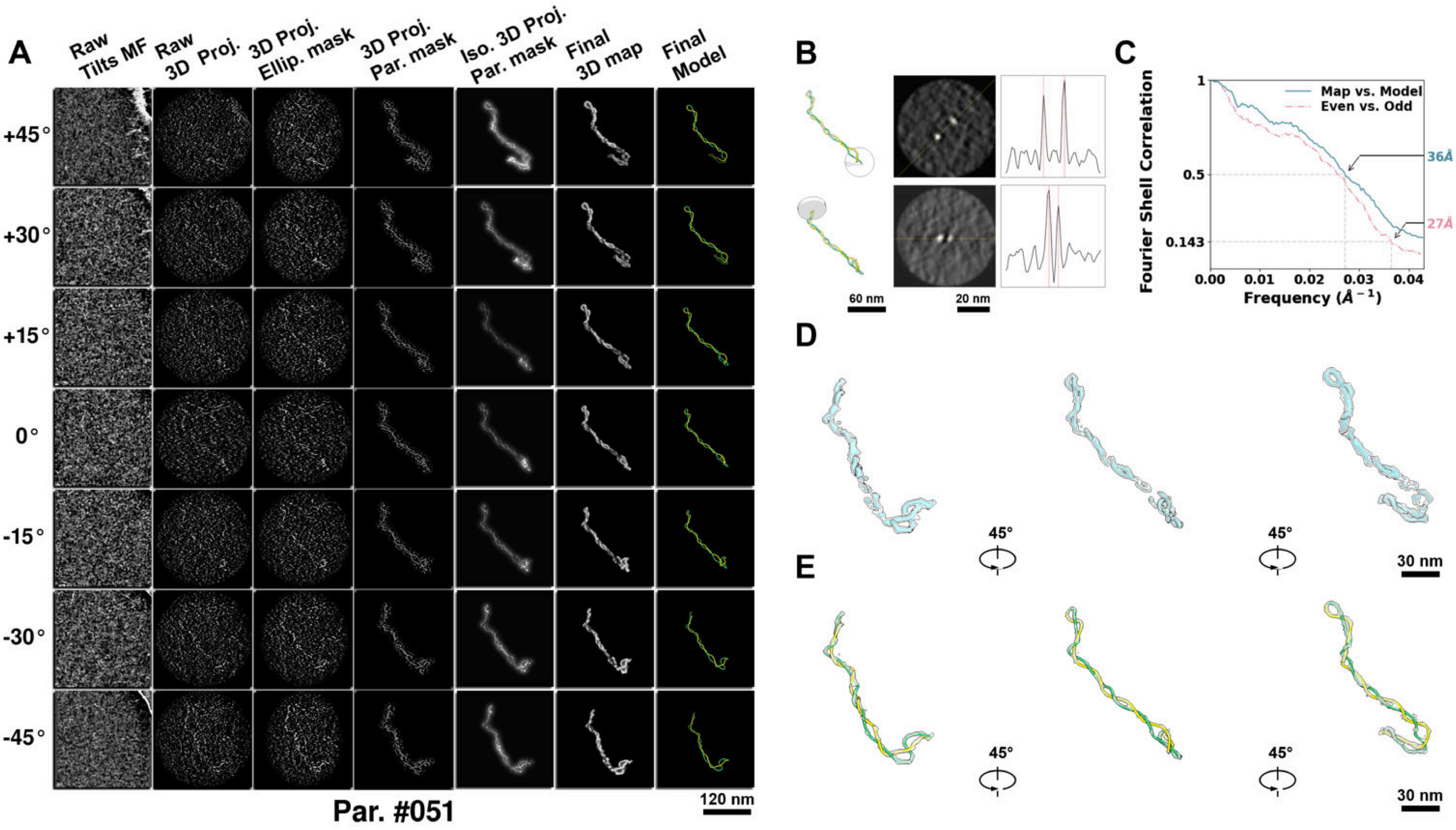

**Supplementary Particle Figure 51. Cryo-ET 3D reconstruction of an individual P.HS particle.**
(A) 3D reconstruction of the plasmid particle (index no. 51). The first column shows seven representative tilt images from +45° to -45° in step of 15°. The second, third, and fourth columns show 3D projections of the particle with spherical, ellipsoidal (thinner along the z-dimension), and particle-shaped masks, respectively. The fifth column displays the 3D projections of the enhanced and IsoNet missing-wedge-corrected particle. The sixth and seventh columns present the final 3D map and the flexibly fitted model, respectively. (B) Two cross-sectional views (12 nm thickness) of the plasmid density map along its plectoneme axis are shown in the left-middle panel. The intensity profile along the line crossing the two high-density DNA spots is displayed in the right panel. (C) Resolution assessment of the final 3D map using Fourier shell correlation (FSC). Two criteria are shown: FSC between two half-maps reconstructed from even and odd frames (evaluated at 0.143) and FSC between the final 3D map and the fitted model (evaluated at 0.5). (D) Zoomed-in views of the final 3D density map from panel A, displayed at two contour levels. (E) Superimposition of the high-contour level map from panel D onto its fitted model.

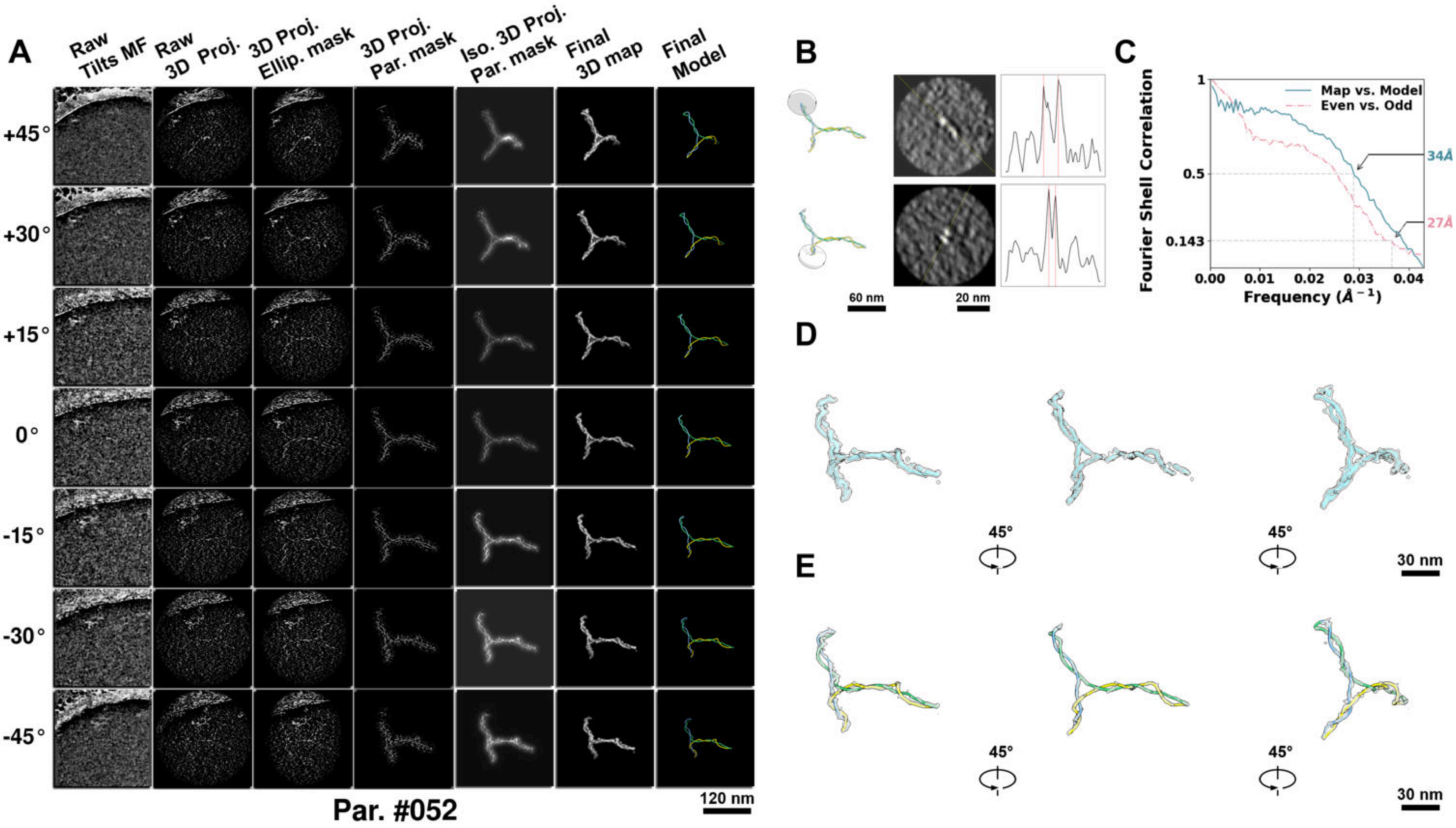

**Supplementary Particle Figure 52. Cryo-ET 3D reconstruction of an individual P.HS particle.**
(A) 3D reconstruction of the plasmid particle (index no. 52). The first column shows seven representative tilt images from +45° to -45° in step of 15°. The second, third, and fourth columns show 3D projections of the particle with spherical, ellipsoidal (thinner along the z-dimension), and particle-shaped masks, respectively. The fifth column displays the 3D projections of the enhanced and IsoNet missing-wedge-corrected particle. The sixth and seventh columns present the final 3D map and the flexibly fitted model, respectively. (B) Two cross-sectional views (12 nm thickness) of the plasmid density map along its plectoneme axis are shown in the left-middle panel. The intensity profile along the line crossing the two high-density DNA spots is displayed in the right panel. (C) Resolution assessment of the final 3D map using Fourier shell correlation (FSC). Two criteria are shown: FSC between two half-maps reconstructed from even and odd frames (evaluated at 0.143) and FSC between the final 3D map and the fitted model (evaluated at 0.5). (D) Zoomed-in views of the final 3D density map from panel A, displayed at two contour levels. (E) Superimposition of the high-contour level map from panel D onto its fitted model.

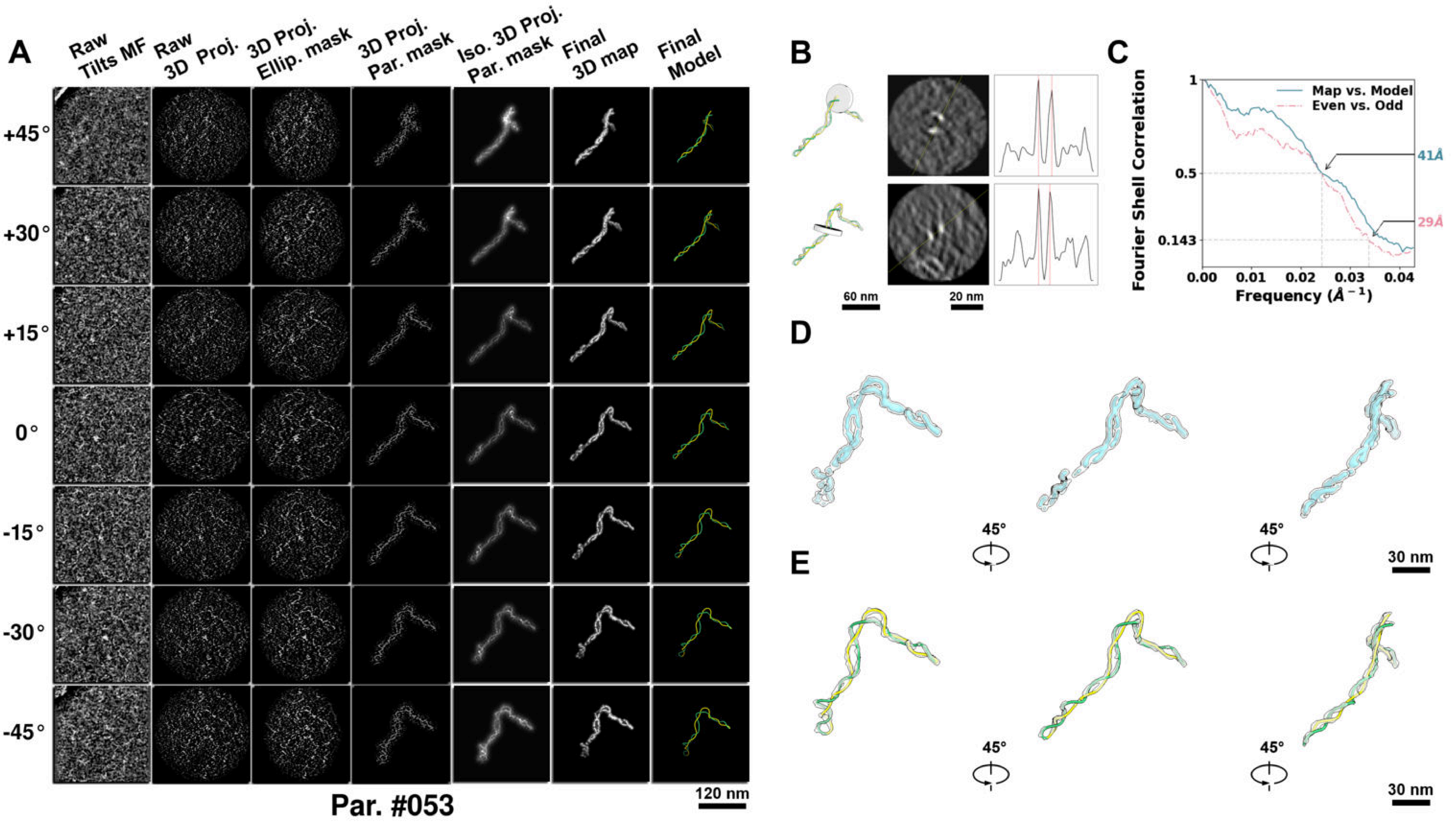

**Supplementary Particle Figure 53. Cryo-ET 3D reconstruction of an individual P.HS particle.**
(A) 3D reconstruction of the plasmid particle (index no. 53). The first column shows seven representative tilt images from +45° to -45° in step of 15°. The second, third, and fourth columns show 3D projections of the particle with spherical, ellipsoidal (thinner along the z-dimension), and particle-shaped masks, respectively. The fifth column displays the 3D projections of the enhanced and IsoNet missing-wedge-corrected particle. The sixth and seventh columns present the final 3D map and the flexibly fitted model, respectively. (B) Two cross-sectional views (12 nm thickness) of the plasmid density map along its plectoneme axis are shown in the left-middle panel. The intensity profile along the line crossing the two high-density DNA spots is displayed in the right panel. (C) Resolution assessment of the final 3D map using Fourier shell correlation (FSC). Two criteria are shown: FSC between two half-maps reconstructed from even and odd frames (evaluated at 0.143) and FSC between the final 3D map and the fitted model (evaluated at 0.5). (D) Zoomed-in views of the final 3D density map from panel A, displayed at two contour levels. (E) Superimposition of the high-contour level map from panel D onto its fitted model.

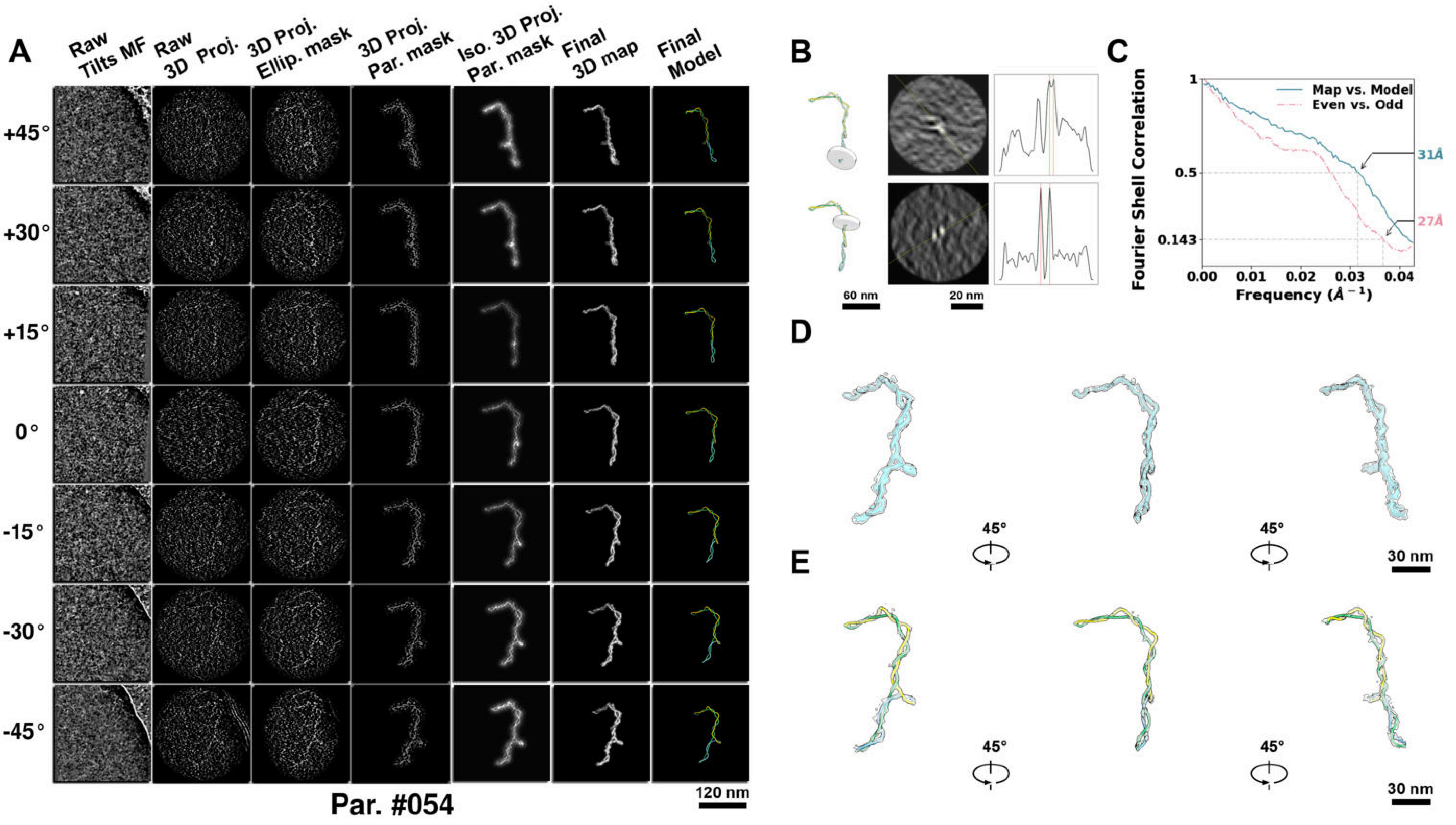

**Supplementary Particle Figure 54. Cryo-ET 3D reconstruction of an individual P.HS particle.**
(A) 3D reconstruction of the plasmid particle (index no. 54). The first column shows seven representative tilt images from +45° to -45° in step of 15°. The second, third, and fourth columns show 3D projections of the particle with spherical, ellipsoidal (thinner along the z-dimension), and particle-shaped masks, respectively. The fifth column displays the 3D projections of the enhanced and IsoNet missing-wedge-corrected particle. The sixth and seventh columns present the final 3D map and the flexibly fitted model, respectively. (B) Two cross-sectional views (12 nm thickness) of the plasmid density map along its plectoneme axis are shown in the left-middle panel. The intensity profile along the line crossing the two high-density DNA spots is displayed in the right panel. (C) Resolution assessment of the final 3D map using Fourier shell correlation (FSC). Two criteria are shown: FSC between two half-maps reconstructed from even and odd frames (evaluated at 0.143) and FSC between the final 3D map and the fitted model (evaluated at 0.5). (D) Zoomed-in views of the final 3D density map from panel A, displayed at two contour levels. (E) Superimposition of the high-contour level map from panel D onto its fitted model.

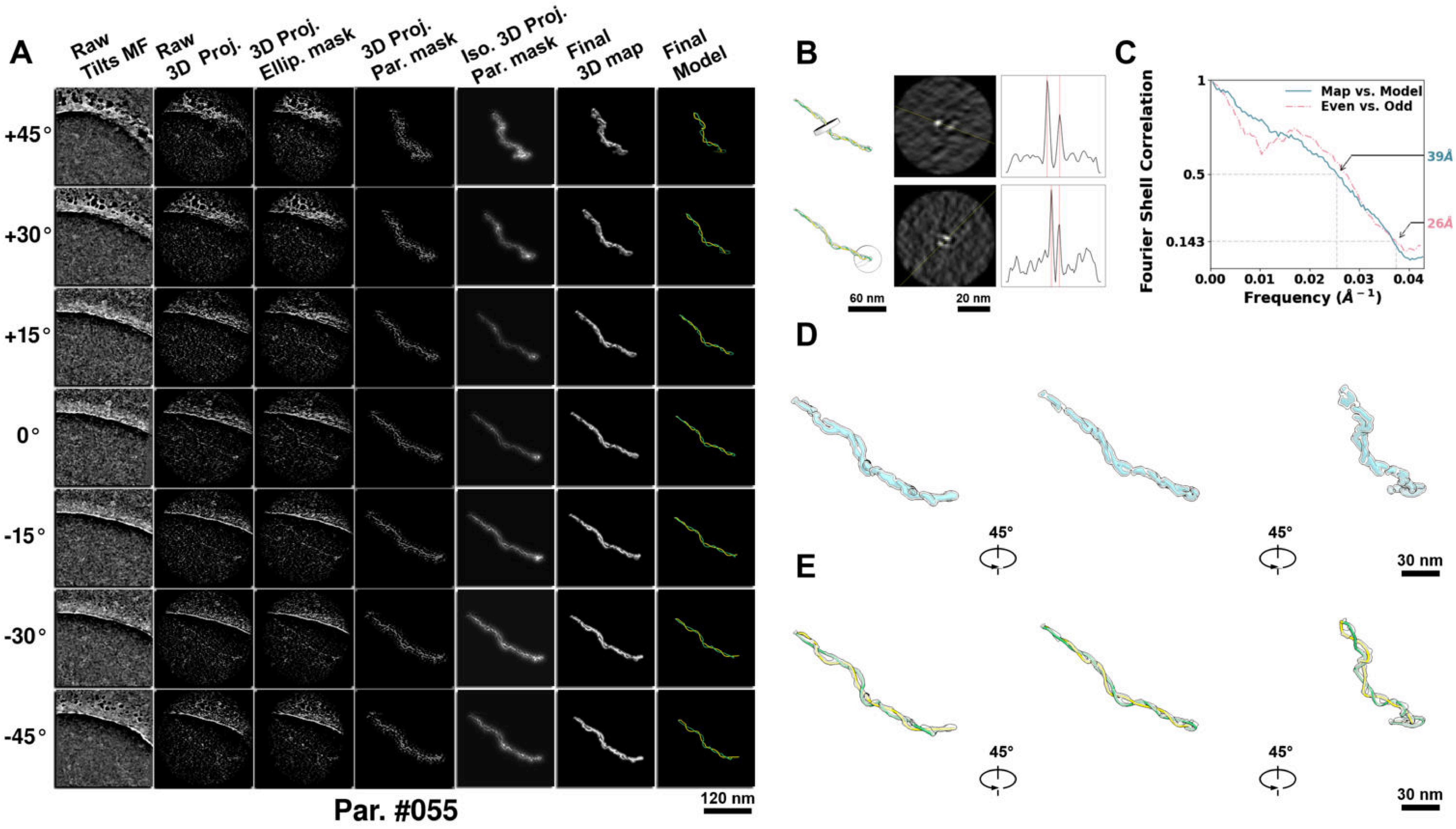

**Supplementary Particle Figure 55. Cryo-ET 3D reconstruction of an individual P.HS particle.**
(A) 3D reconstruction of the plasmid particle (index no. 55). The first column shows seven representative tilt images from +45° to -45° in step of 15°. The second, third, and fourth columns show 3D projections of the particle with spherical, ellipsoidal (thinner along the z-dimension), and particle-shaped masks, respectively. The fifth column displays the 3D projections of the enhanced and IsoNet missing-wedge-corrected particle. The sixth and seventh columns present the final 3D map and the flexibly fitted model, respectively. (B) Two cross-sectional views (12 nm thickness) of the plasmid density map along its plectoneme axis are shown in the left-middle panel. The intensity profile along the line crossing the two high-density DNA spots is displayed in the right panel. (C) Resolution assessment of the final 3D map using Fourier shell correlation (FSC). Two criteria are shown: FSC between two half-maps reconstructed from even and odd frames (evaluated at 0.143) and FSC between the final 3D map and the fitted model (evaluated at 0.5). (D) Zoomed-in views of the final 3D density map from panel A, displayed at two contour levels. (E) Superimposition of the high-contour level map from panel D onto its fitted model.

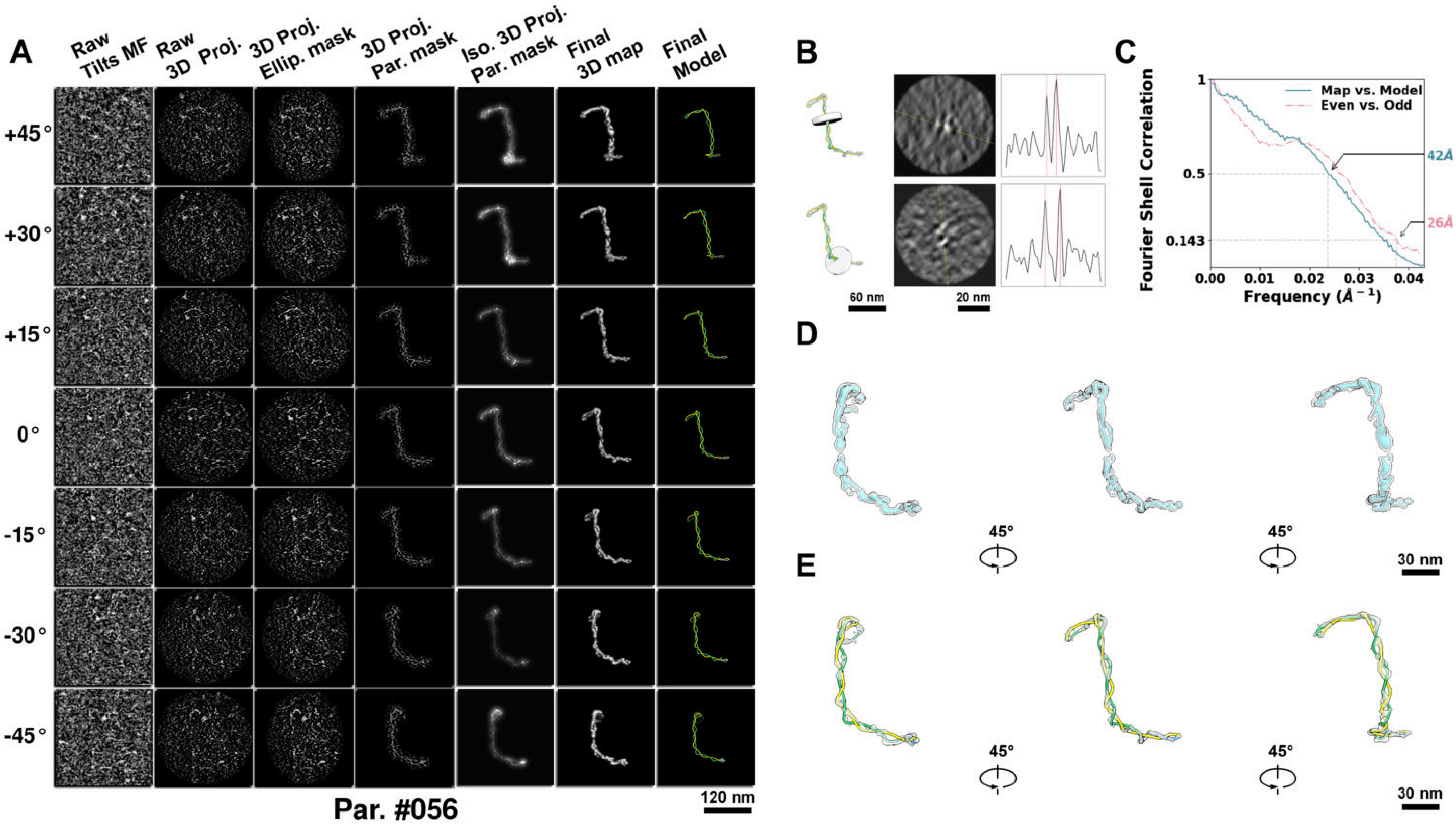

**Supplementary Particle Figure 56. Cryo-ET 3D reconstruction of an individual P.HS particle.**
(A) 3D reconstruction of the plasmid particle (index no. 56). The first column shows seven representative tilt images from +45° to -45° in step of 15°. The second, third, and fourth columns show 3D projections of the particle with spherical, ellipsoidal (thinner along the z-dimension), and particle-shaped masks, respectively. The fifth column displays the 3D projections of the enhanced and IsoNet missing-wedge-corrected particle. The sixth and seventh columns present the final 3D map and the flexibly fitted model, respectively. (B) Two cross-sectional views (12 nm thickness) of the plasmid density map along its plectoneme axis are shown in the left-middle panel. The intensity profile along the line crossing the two high-density DNA spots is displayed in the right panel. (C) Resolution assessment of the final 3D map using Fourier shell correlation (FSC). Two criteria are shown: FSC between two half-maps reconstructed from even and odd frames (evaluated at 0.143) and FSC between the final 3D map and the fitted model (evaluated at 0.5). (D) Zoomed-in views of the final 3D density map from panel A, displayed at two contour levels. (E) Superimposition of the high-contour level map from panel D onto its fitted model.

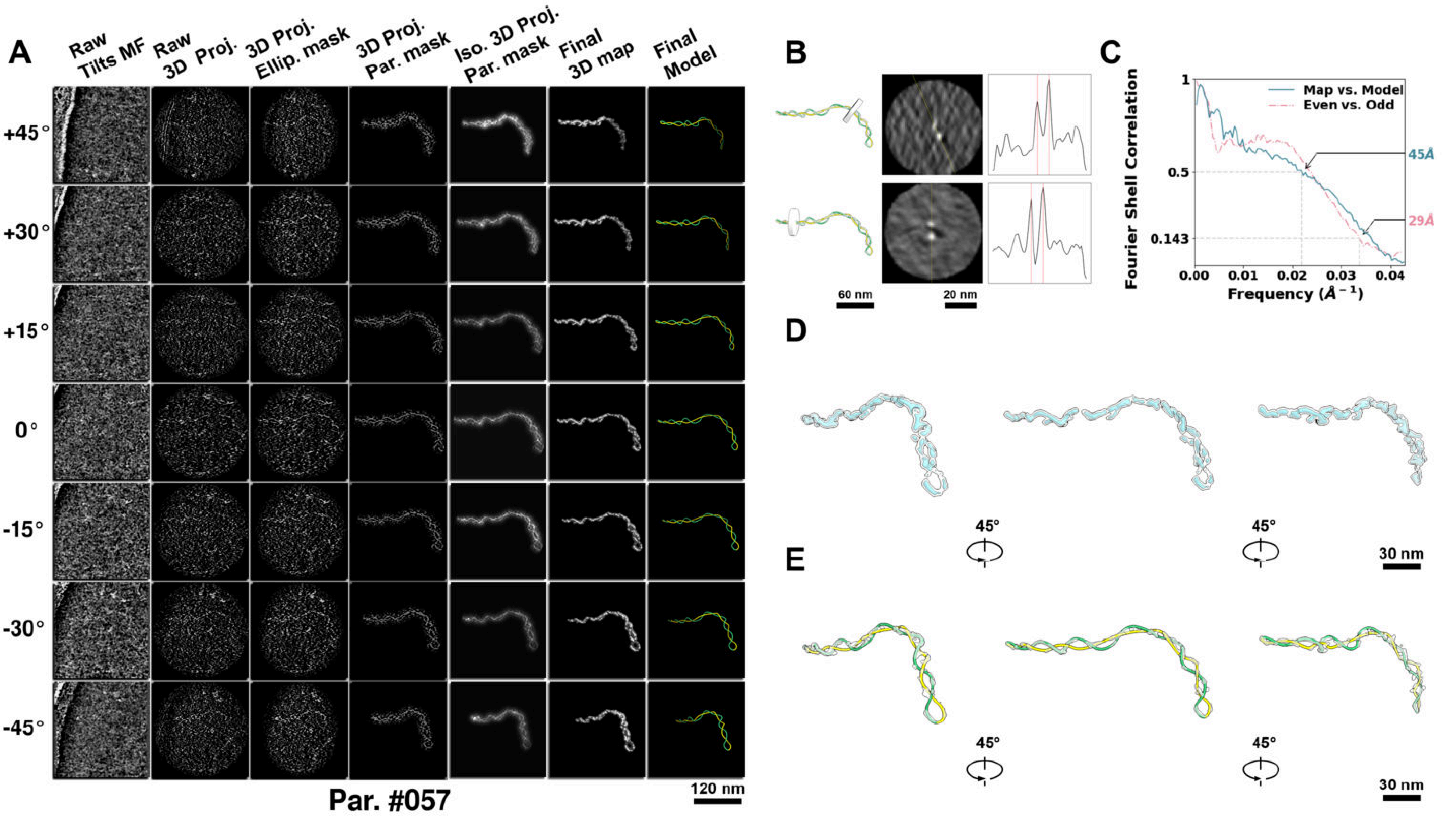

**Supplementary Particle Figure 57. Cryo-ET 3D reconstruction of an individual P.HS particle.**
(A) 3D reconstruction of the plasmid particle (index no. 57). The first column shows seven representative tilt images from +45° to -45° in step of 15°. The second, third, and fourth columns show 3D projections of the particle with spherical, ellipsoidal (thinner along the z-dimension), and particle-shaped masks, respectively. The fifth column displays the 3D projections of the enhanced and IsoNet missing-wedge-corrected particle. The sixth and seventh columns present the final 3D map and the flexibly fitted model, respectively. (B) Two cross-sectional views (12 nm thickness) of the plasmid density map along its plectoneme axis are shown in the left-middle panel. The intensity profile along the line crossing the two high-density DNA spots is displayed in the right panel. (C) Resolution assessment of the final 3D map using Fourier shell correlation (FSC). Two criteria are shown: FSC between two half-maps reconstructed from even and odd frames (evaluated at 0.143) and FSC between the final 3D map and the fitted model (evaluated at 0.5). (D) Zoomed-in views of the final 3D density map from panel A, displayed at two contour levels. (E) Superimposition of the high-contour level map from panel D onto its fitted model.

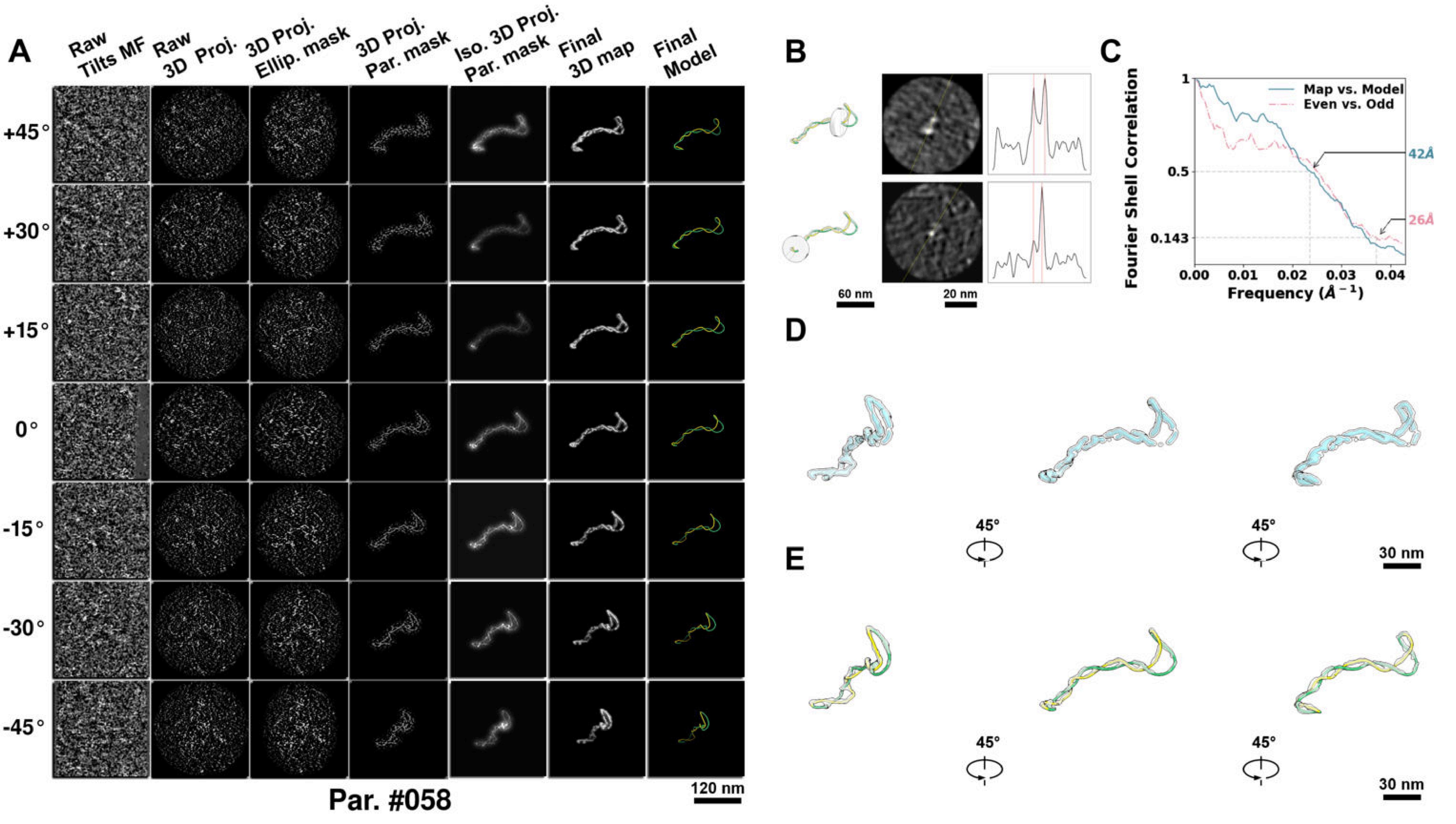

**Supplementary Particle Figure 58. Cryo-ET 3D reconstruction of an individual P.HS particle.**
(A) 3D reconstruction of the plasmid particle (index no. 58). The first column shows seven representative tilt images from +45° to -45° in step of 15°. The second, third, and fourth columns show 3D projections of the particle with spherical, ellipsoidal (thinner along the z-dimension), and particle-shaped masks, respectively. The fifth column displays the 3D projections of the enhanced and IsoNet missing-wedge-corrected particle. The sixth and seventh columns present the final 3D map and the flexibly fitted model, respectively. (B) Two cross-sectional views (12 nm thickness) of the plasmid density map along its plectoneme axis are shown in the left-middle panel. The intensity profile along the line crossing the two high-density DNA spots is displayed in the right panel. (C) Resolution assessment of the final 3D map using Fourier shell correlation (FSC). Two criteria are shown: FSC between two half-maps reconstructed from even and odd frames (evaluated at 0.143) and FSC between the final 3D map and the fitted model (evaluated at 0.5). (D) Zoomed-in views of the final 3D density map from panel A, displayed at two contour levels. (E) Superimposition of the high-contour level map from panel D onto its fitted model.

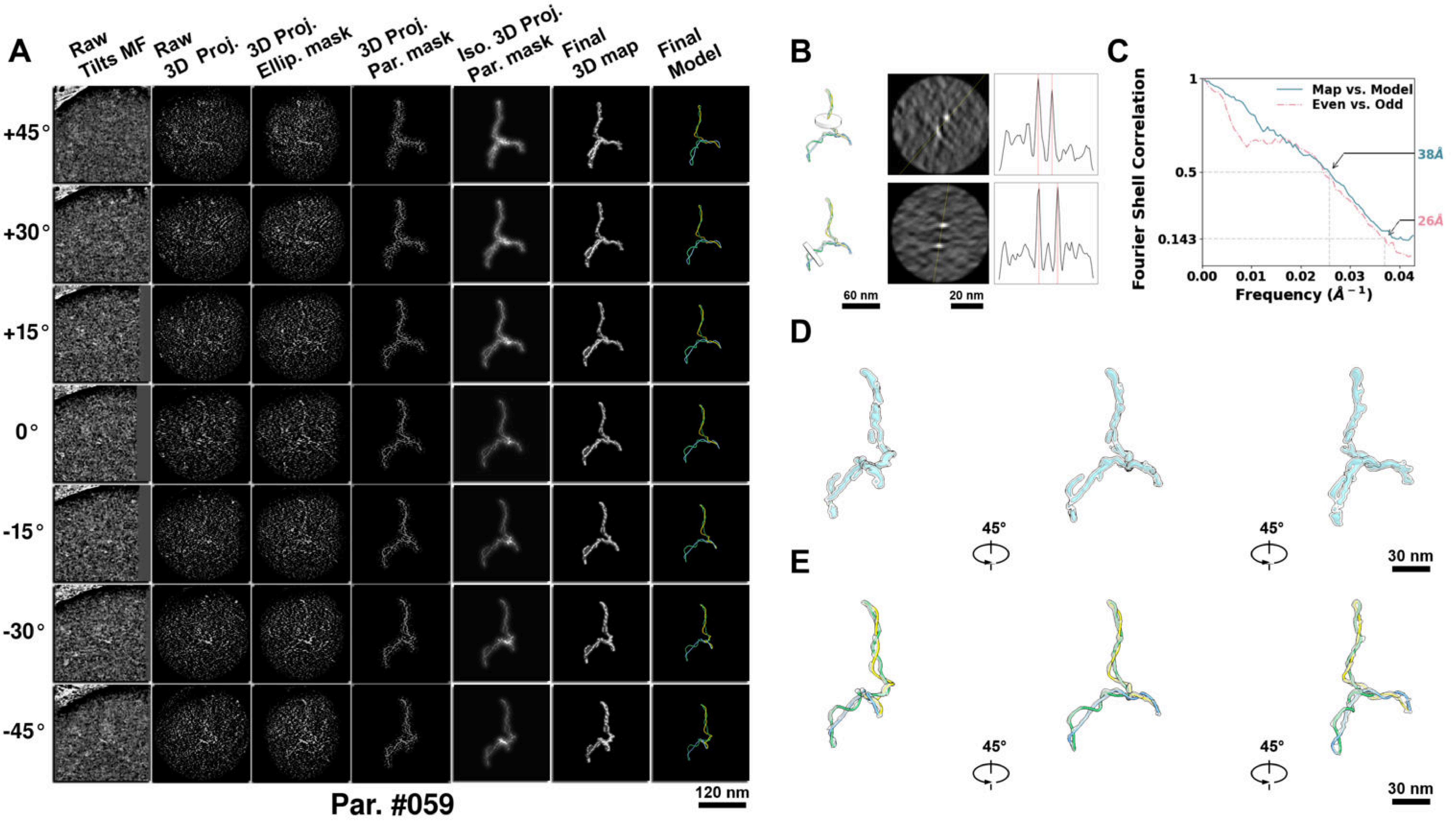

**Supplementary Particle Figure 59. Cryo-ET 3D reconstruction of an individual P.HS particle.**
(A) 3D reconstruction of the plasmid particle (index no. 59). The first column shows seven representative tilt images from +45° to -45° in step of 15°. The second, third, and fourth columns show 3D projections of the particle with spherical, ellipsoidal (thinner along the z-dimension), and particle-shaped masks, respectively. The fifth column displays the 3D projections of the enhanced and IsoNet missing-wedge-corrected particle. The sixth and seventh columns present the final 3D map and the flexibly fitted model, respectively. (B) Two cross-sectional views (12 nm thickness) of the plasmid density map along its plectoneme axis are shown in the left-middle panel. The intensity profile along the line crossing the two high-density DNA spots is displayed in the right panel. (C) Resolution assessment of the final 3D map using Fourier shell correlation (FSC). Two criteria are shown: FSC between two half-maps reconstructed from even and odd frames (evaluated at 0.143) and FSC between the final 3D map and the fitted model (evaluated at 0.5). (D) Zoomed-in views of the final 3D density map from panel A, displayed at two contour levels. (E) Superimposition of the high-contour level map from panel D onto its fitted model.

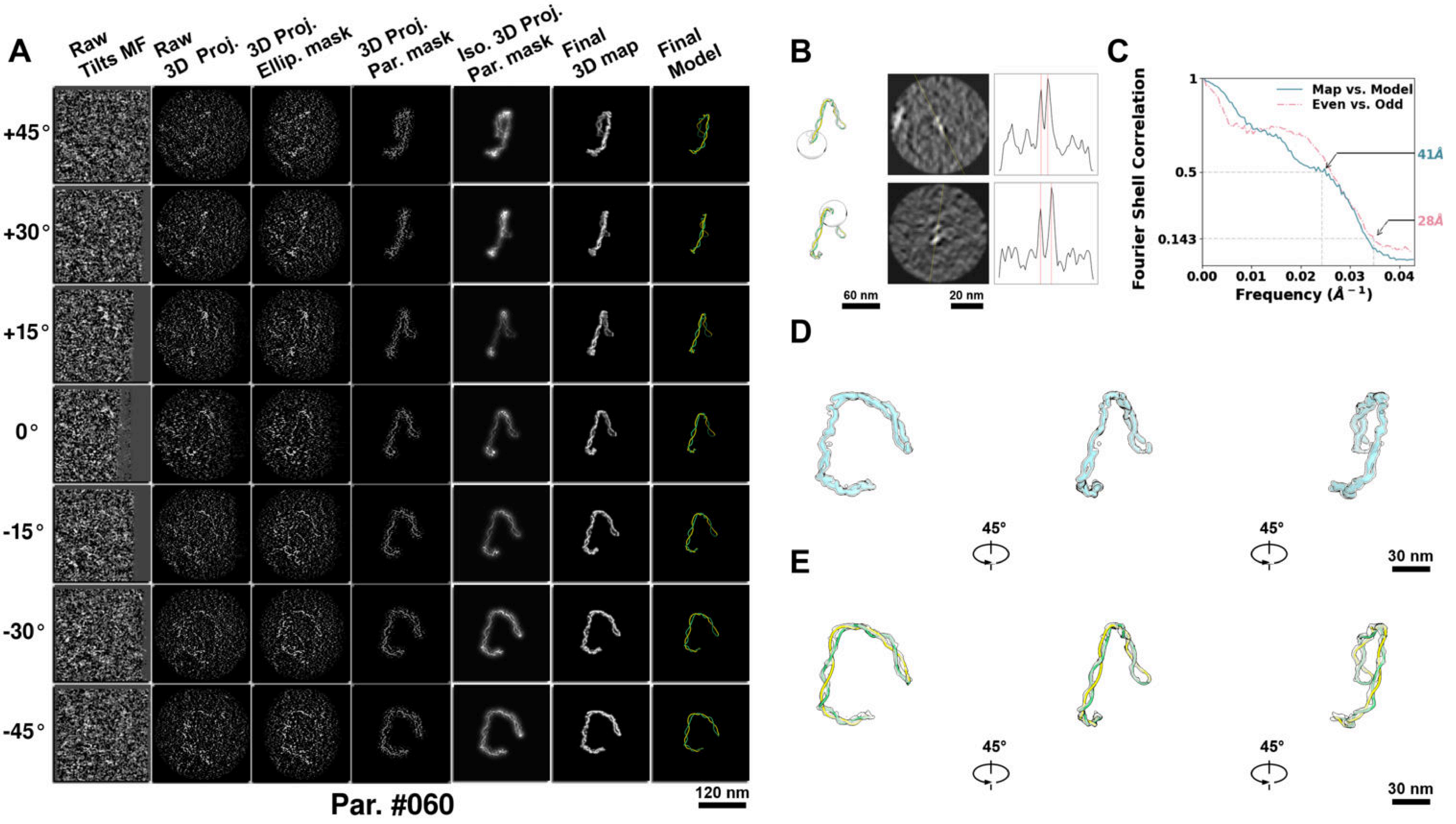

**Supplementary Particle Figure 60. Cryo-ET 3D reconstruction of an individual P.HS particle.**
(A) 3D reconstruction of the plasmid particle (index no. 60). The first column shows seven representative tilt images from +45° to -45° in step of 15°. The second, third, and fourth columns show 3D projections of the particle with spherical, ellipsoidal (thinner along the z-dimension), and particle-shaped masks, respectively. The fifth column displays the 3D projections of the enhanced and IsoNet missing-wedge-corrected particle. The sixth and seventh columns present the final 3D map and the flexibly fitted model, respectively. (B) Two cross-sectional views (12 nm thickness) of the plasmid density map along its plectoneme axis are shown in the left-middle panel. The intensity profile along the line crossing the two high-density DNA spots is displayed in the right panel. (C) Resolution assessment of the final 3D map using Fourier shell correlation (FSC). Two criteria are shown: FSC between two half-maps reconstructed from even and odd frames (evaluated at 0.143) and FSC between the final 3D map and the fitted model (evaluated at 0.5). (D) Zoomed-in views of the final 3D density map from panel A, displayed at two contour levels. (E) Superimposition of the high-contour level map from panel D onto its fitted model.

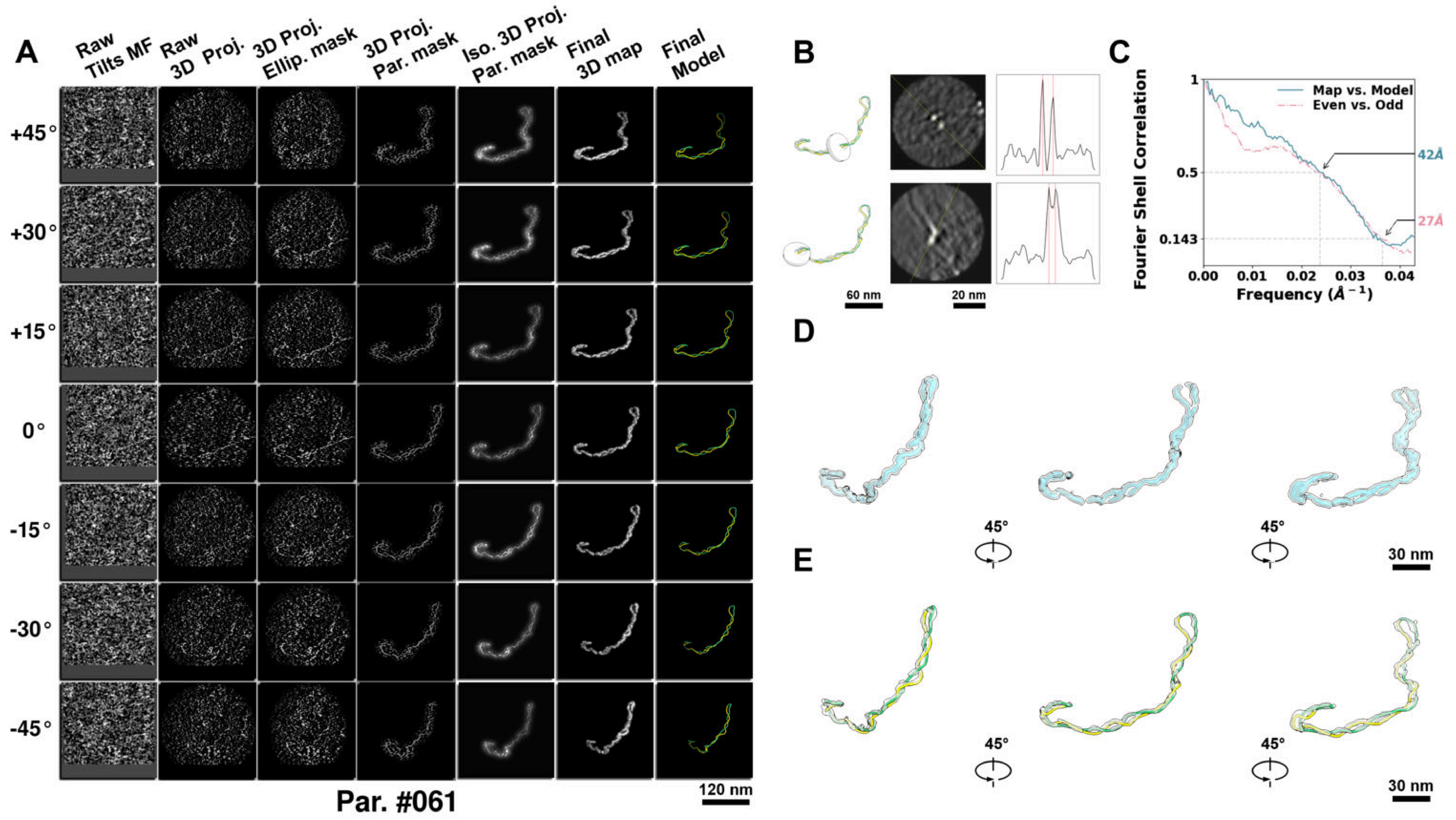

**Supplementary Particle Figure 61. Cryo-ET 3D reconstruction of an individual P.HS particle.**
(A) 3D reconstruction of the plasmid particle (index no. 61). The first column shows seven representative tilt images from +45° to -45° in step of 15°. The second, third, and fourth columns show 3D projections of the particle with spherical, ellipsoidal (thinner along the z-dimension), and particle-shaped masks, respectively. The fifth column displays the 3D projections of the enhanced and IsoNet missing-wedge-corrected particle. The sixth and seventh columns present the final 3D map and the flexibly fitted model, respectively. (B) Two cross-sectional views (12 nm thickness) of the plasmid density map along its plectoneme axis are shown in the left-middle panel. The intensity profile along the line crossing the two high-density DNA spots is displayed in the right panel. (C) Resolution assessment of the final 3D map using Fourier shell correlation (FSC). Two criteria are shown: FSC between two half-maps reconstructed from even and odd frames (evaluated at 0.143) and FSC between the final 3D map and the fitted model (evaluated at 0.5). (D) Zoomed-in views of the final 3D density map from panel A, displayed at two contour levels. (E) Superimposition of the high-contour level map from panel D onto its fitted model.

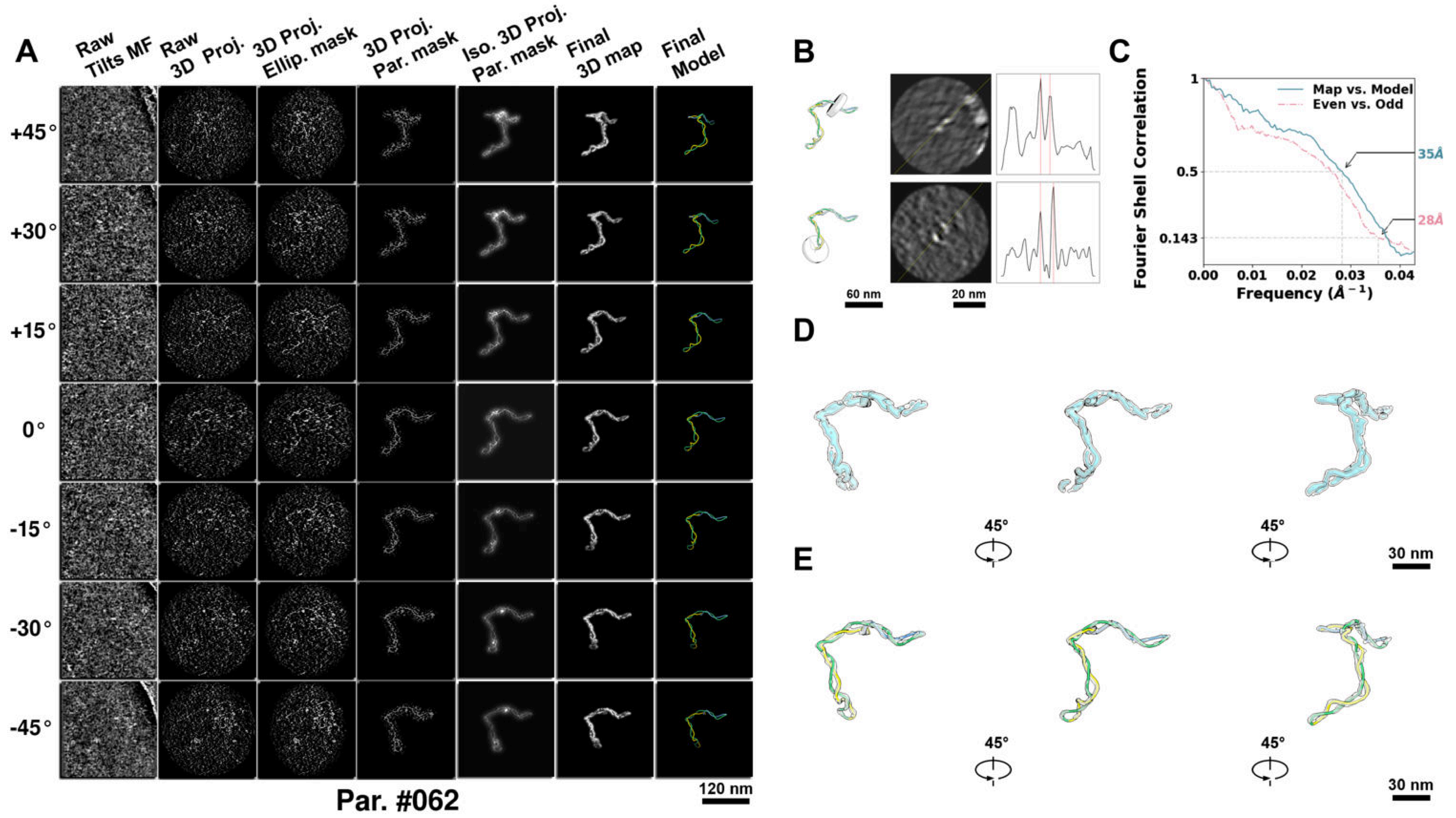

**Supplementary Particle Figure 62. Cryo-ET 3D reconstruction of an individual P.HS particle.**
(A) 3D reconstruction of the plasmid particle (index no. 62). The first column shows seven representative tilt images from +45° to -45° in step of 15°. The second, third, and fourth columns show 3D projections of the particle with spherical, ellipsoidal (thinner along the z-dimension), and particle-shaped masks, respectively. The fifth column displays the 3D projections of the enhanced and IsoNet missing-wedge-corrected particle. The sixth and seventh columns present the final 3D map and the flexibly fitted model, respectively. (B) Two cross-sectional views (12 nm thickness) of the plasmid density map along its plectoneme axis are shown in the left-middle panel. The intensity profile along the line crossing the two high-density DNA spots is displayed in the right panel. (C) Resolution assessment of the final 3D map using Fourier shell correlation (FSC). Two criteria are shown: FSC between two half-maps reconstructed from even and odd frames (evaluated at 0.143) and FSC between the final 3D map and the fitted model (evaluated at 0.5). (D) Zoomed-in views of the final 3D density map from panel A, displayed at two contour levels. (E) Superimposition of the high-contour level map from panel D onto its fitted model.

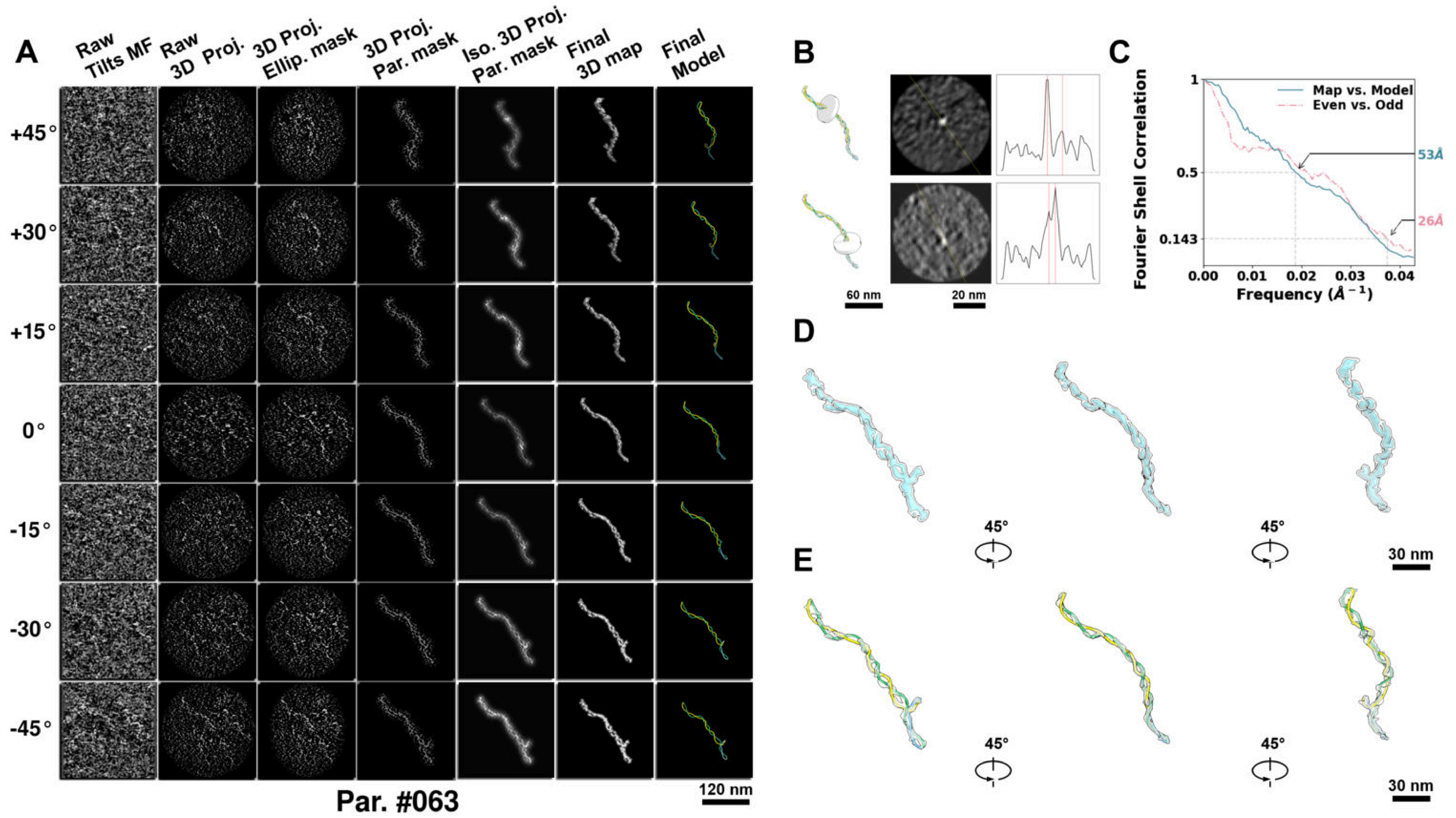

**Supplementary Particle Figure 63. Cryo-ET 3D reconstruction of an individual P.HS particle.**
(A) 3D reconstruction of the plasmid particle (index no. 63). The first column shows seven representative tilt images from +45° to -45° in step of 15°. The second, third, and fourth columns show 3D projections of the particle with spherical, ellipsoidal (thinner along the z-dimension), and particle-shaped masks, respectively. The fifth column displays the 3D projections of the enhanced and IsoNet missing-wedge-corrected particle. The sixth and seventh columns present the final 3D map and the flexibly fitted model, respectively. (B) Two cross-sectional views (12 nm thickness) of the plasmid density map along its plectoneme axis are shown in the left-middle panel. The intensity profile along the line crossing the two high-density DNA spots is displayed in the right panel. (C) Resolution assessment of the final 3D map using Fourier shell correlation (FSC). Two criteria are shown: FSC between two half-maps reconstructed from even and odd frames (evaluated at 0.143) and FSC between the final 3D map and the fitted model (evaluated at 0.5). (D) Zoomed-in views of the final 3D density map from panel A, displayed at two contour levels. (E) Superimposition of the high-contour level map from panel D onto its fitted model.

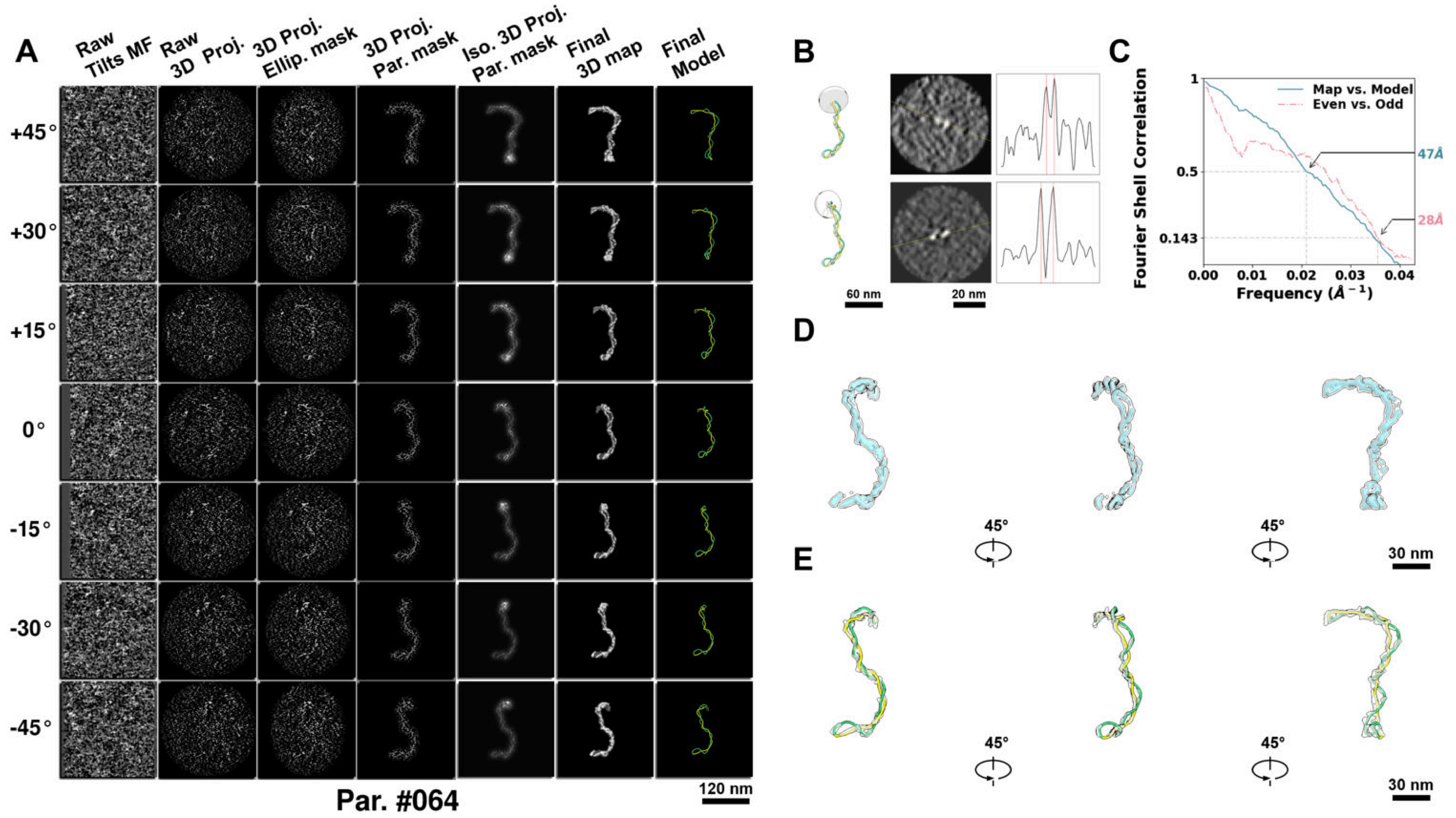

**Supplementary Particle Figure 64. Cryo-ET 3D reconstruction of an individual P.HS particle.**
(A) 3D reconstruction of the plasmid particle (index no. 64). The first column shows seven representative tilt images from +45° to -45° in step of 15°. The second, third, and fourth columns show 3D projections of the particle with spherical, ellipsoidal (thinner along the z-dimension), and particle-shaped masks, respectively. The fifth column displays the 3D projections of the enhanced and IsoNet missing-wedge-corrected particle. The sixth and seventh columns present the final 3D map and the flexibly fitted model, respectively. (B) Two cross-sectional views (12 nm thickness) of the plasmid density map along its plectoneme axis are shown in the left-middle panel. The intensity profile along the line crossing the two high-density DNA spots is displayed in the right panel. (C) Resolution assessment of the final 3D map using Fourier shell correlation (FSC). Two criteria are shown: FSC between two half-maps reconstructed from even and odd frames (evaluated at 0.143) and FSC between the final 3D map and the fitted model (evaluated at 0.5). (D) Zoomed-in views of the final 3D density map from panel A, displayed at two contour levels. (E) Superimposition of the high-contour level map from panel D onto its fitted model.

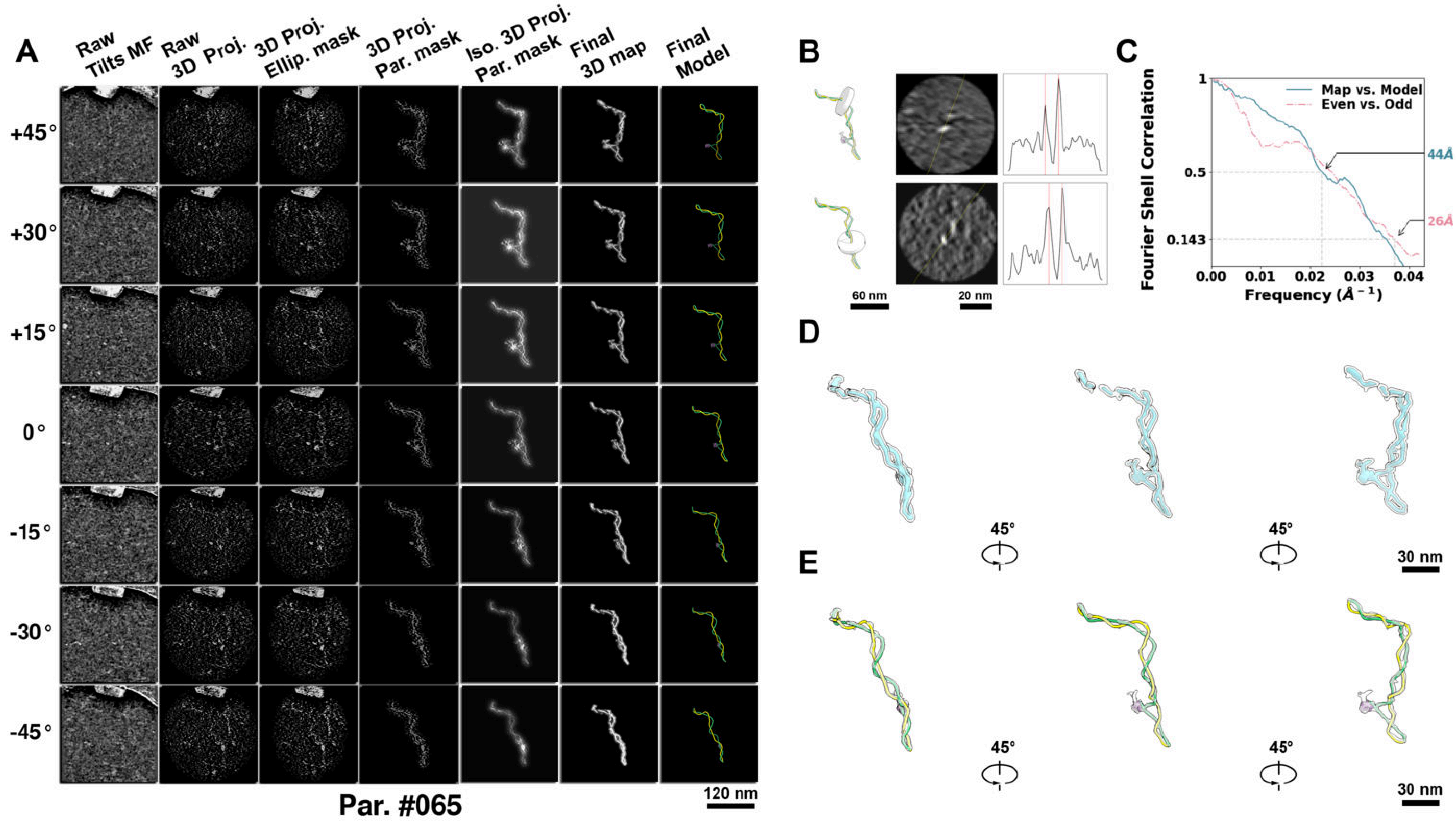

**Supplementary Particle Figure 65. Cryo-ET 3D reconstruction of an individual sTEC particle.**
(A) 3D reconstruction of the plasmid particle (index no. 65). The first column shows seven representative tilt images from +45° to -45° in step of 15°. The second, third, and fourth columns show 3D projections of the particle with spherical, ellipsoidal (thinner along the z-dimension), and particle-shaped masks, respectively. The fifth column displays the 3D projections of the enhanced and IsoNet missing-wedge-corrected particle. The sixth and seventh columns present the final 3D map and the flexibly fitted model, respectively. (B) Two cross-sectional views (12 nm thickness) of the plasmid density map along its plectoneme axis are shown in the left-middle panel. The intensity profile along the line crossing the two high-density DNA spots is displayed in the right panel. (C) Resolution assessment of the final 3D map using Fourier shell correlation (FSC). Two criteria are shown: FSC between two half-maps reconstructed from even and odd frames (evaluated at 0.143) and FSC between the final 3D map and the fitted model (evaluated at 0.5). (D) Zoomed-in views of the final 3D density map from panel A, displayed at two contour levels. (E) Superimposition of the high-contour level map from panel D onto its fitted model.

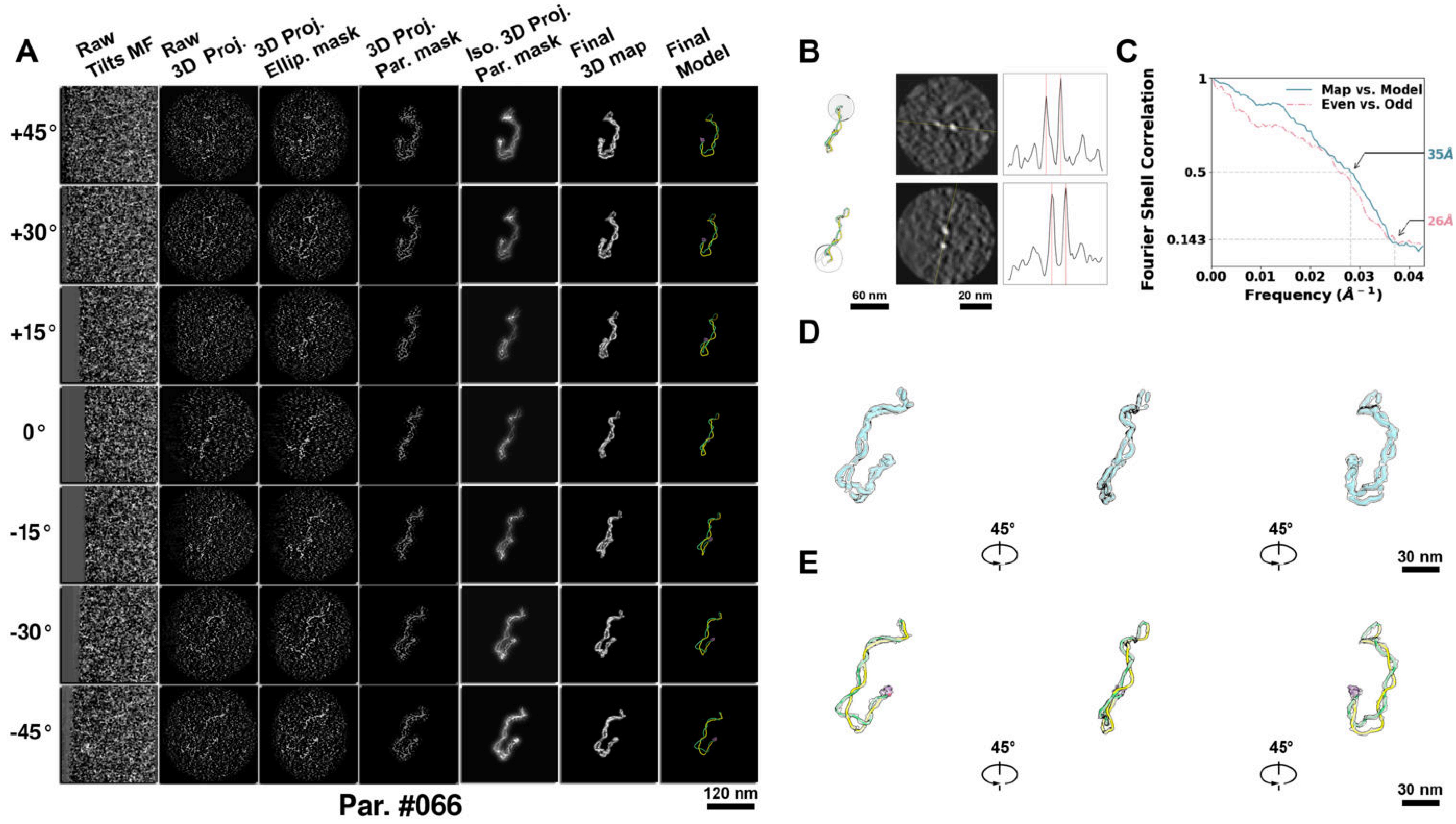

**Supplementary Particle Figure 66. Cryo-ET 3D reconstruction of an individual sTEC particle.**
(A) 3D reconstruction of the plasmid particle (index no. 66). The first column shows seven representative tilt images from +45° to -45° in step of 15°. The second, third, and fourth columns show 3D projections of the particle with spherical, ellipsoidal (thinner along the z-dimension), and particle-shaped masks, respectively. The fifth column displays the 3D projections of the enhanced and IsoNet missing-wedge-corrected particle. The sixth and seventh columns present the final 3D map and the flexibly fitted model, respectively. (B) Two cross-sectional views (12 nm thickness) of the plasmid density map along its plectoneme axis are shown in the left-middle panel. The intensity profile along the line crossing the two high-density DNA spots is displayed in the right panel. (C) Resolution assessment of the final 3D map using Fourier shell correlation (FSC). Two criteria are shown: FSC between two half-maps reconstructed from even and odd frames (evaluated at 0.143) and FSC between the final 3D map and the fitted model (evaluated at 0.5). (D) Zoomed-in views of the final 3D density map from panel A, displayed at two contour levels. (E) Superimposition of the high-contour level map from panel D onto its fitted model.

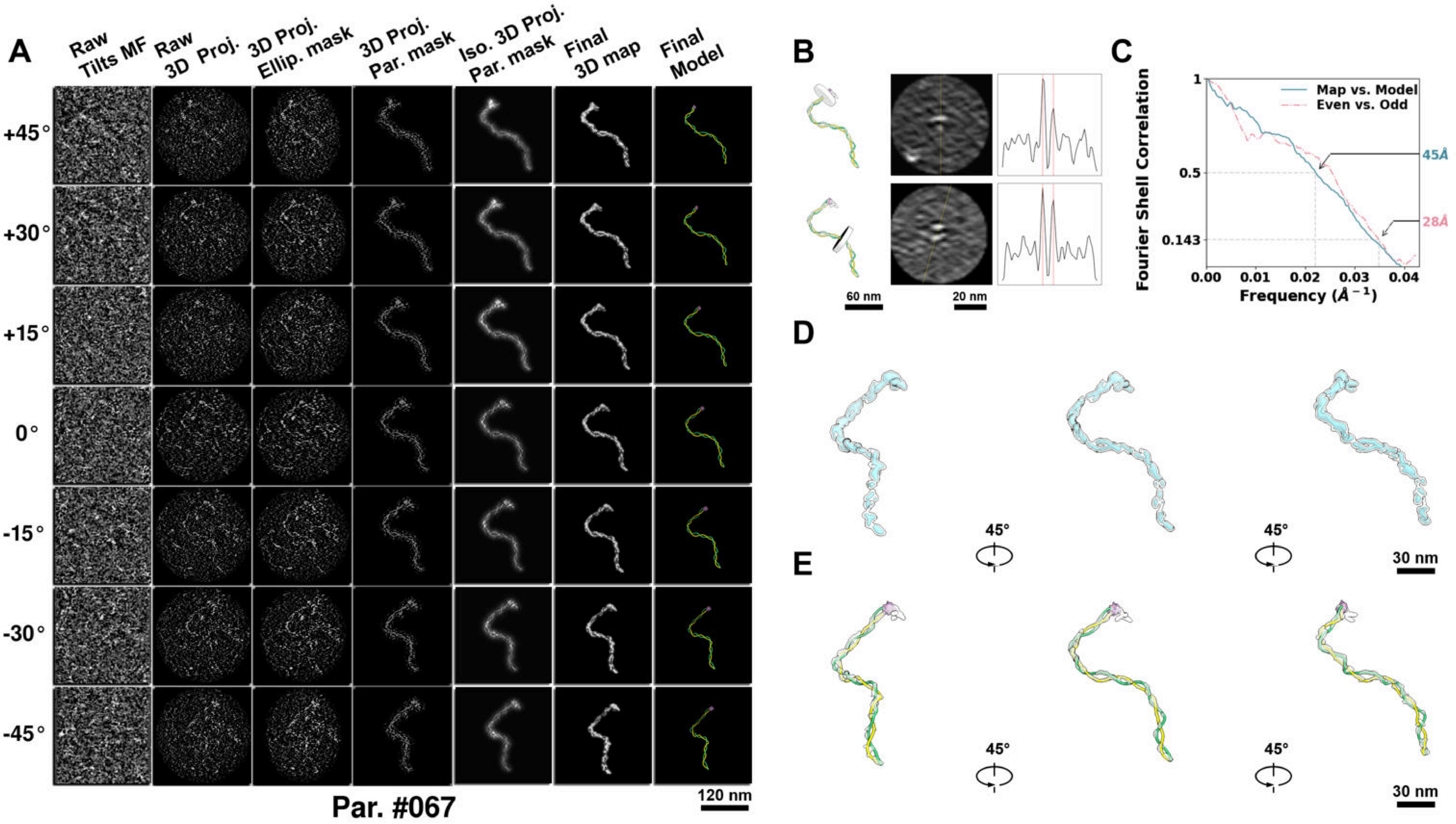

**Supplementary Particle Figure 67. Cryo-ET 3D reconstruction of an individual sTEC particle.**
(A) 3D reconstruction of the plasmid particle (index no. 67). The first column shows seven representative tilt images from +45° to -45° in step of 15°. The second, third, and fourth columns show 3D projections of the particle with spherical, ellipsoidal (thinner along the z-dimension), and particle-shaped masks, respectively. The fifth column displays the 3D projections of the enhanced and IsoNet missing-wedge-corrected particle. The sixth and seventh columns present the final 3D map and the flexibly fitted model, respectively. (B) Two cross-sectional views (12 nm thickness) of the plasmid density map along its plectoneme axis are shown in the left-middle panel. The intensity profile along the line crossing the two high-density DNA spots is displayed in the right panel. (C) Resolution assessment of the final 3D map using Fourier shell correlation (FSC). Two criteria are shown: FSC between two half-maps reconstructed from even and odd frames (evaluated at 0.143) and FSC between the final 3D map and the fitted model (evaluated at 0.5). (D) Zoomed-in views of the final 3D density map from panel A, displayed at two contour levels. (E) Superimposition of the high-contour level map from panel D onto its fitted model.

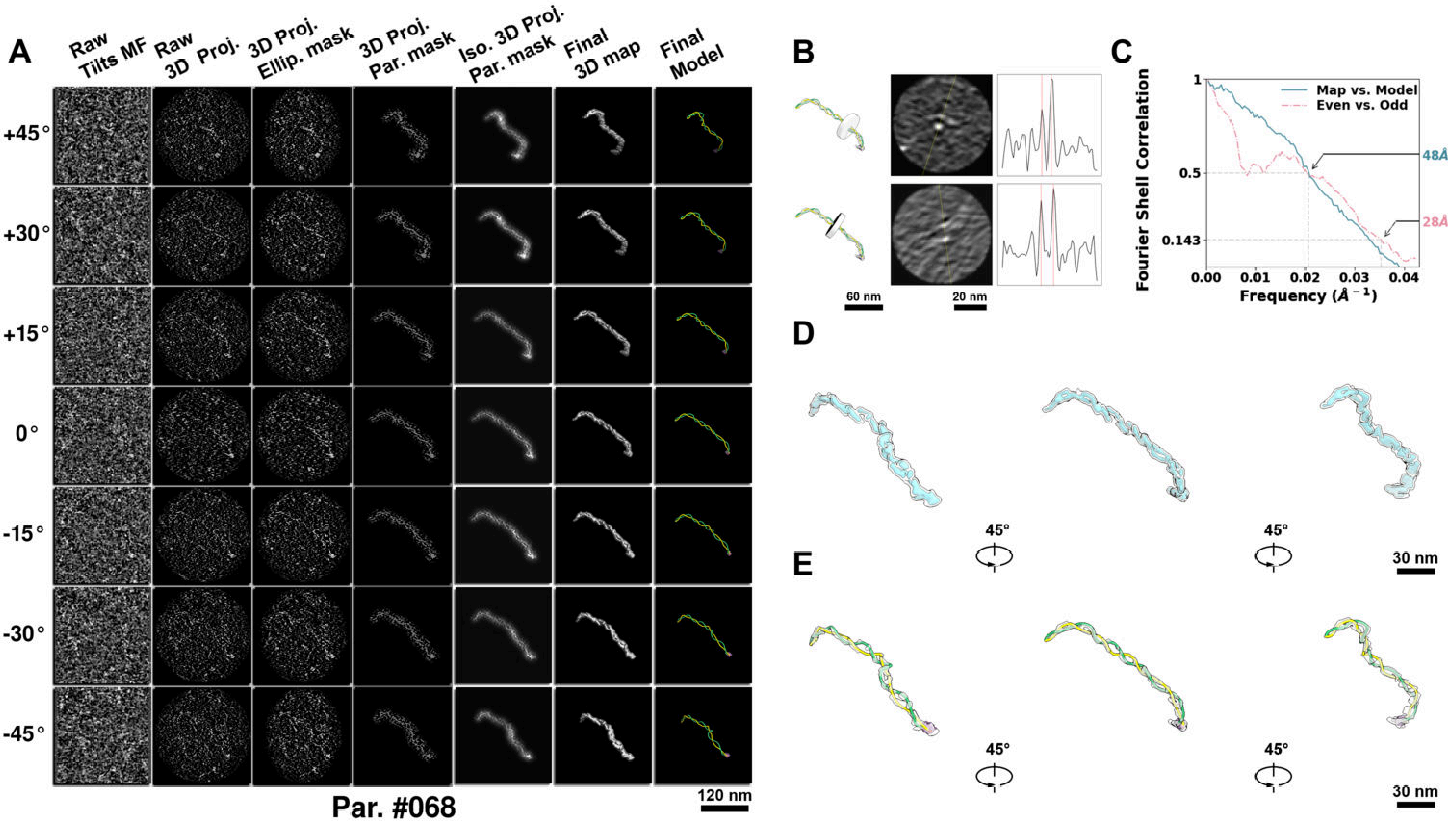

**Supplementary Particle Figure 68. Cryo-ET 3D reconstruction of an individual sTEC particle.**
(A) 3D reconstruction of the plasmid particle (index no. 68). The first column shows seven representative tilt images from +45° to -45° in step of 15°. The second, third, and fourth columns show 3D projections of the particle with spherical, ellipsoidal (thinner along the z-dimension), and particle-shaped masks, respectively. The fifth column displays the 3D projections of the enhanced and IsoNet missing-wedge-corrected particle. The sixth and seventh columns present the final 3D map and the flexibly fitted model, respectively. (B) Two cross-sectional views (12 nm thickness) of the plasmid density map along its plectoneme axis are shown in the left-middle panel. The intensity profile along the line crossing the two high-density DNA spots is displayed in the right panel. (C) Resolution assessment of the final 3D map using Fourier shell correlation (FSC). Two criteria are shown: FSC between two half-maps reconstructed from even and odd frames (evaluated at 0.143) and FSC between the final 3D map and the fitted model (evaluated at 0.5). (D) Zoomed-in views of the final 3D density map from panel A, displayed at two contour levels. (E) Superimposition of the high-contour level map from panel D onto its fitted model.

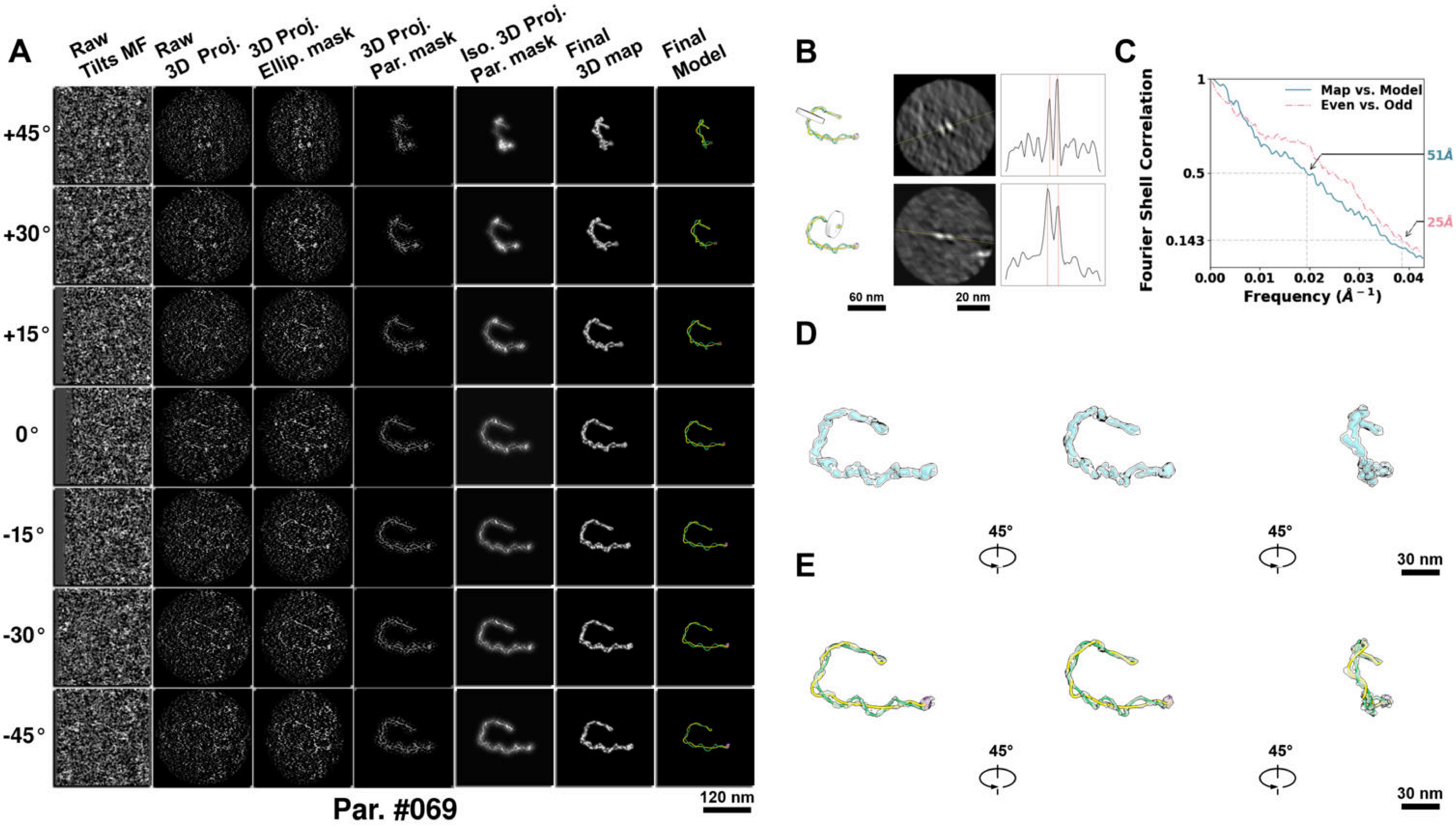

**Supplementary Particle Figure 69. Cryo-ET 3D reconstruction of an individual sTEC particle.**
(A) 3D reconstruction of the plasmid particle (index no. 69). The first column shows seven representative tilt images from +45° to -45° in step of 15°. The second, third, and fourth columns show 3D projections of the particle with spherical, ellipsoidal (thinner along the z-dimension), and particle-shaped masks, respectively. The fifth column displays the 3D projections of the enhanced and IsoNet missing-wedge-corrected particle. The sixth and seventh columns present the final 3D map and the flexibly fitted model, respectively. (B) Two cross-sectional views (12 nm thickness) of the plasmid density map along its plectoneme axis are shown in the left-middle panel. The intensity profile along the line crossing the two high-density DNA spots is displayed in the right panel. (C) Resolution assessment of the final 3D map using Fourier shell correlation (FSC). Two criteria are shown: FSC between two half-maps reconstructed from even and odd frames (evaluated at 0.143) and FSC between the final 3D map and the fitted model (evaluated at 0.5). (D) Zoomed-in views of the final 3D density map from panel A, displayed at two contour levels. (E) Superimposition of the high-contour level map from panel D onto its fitted model.

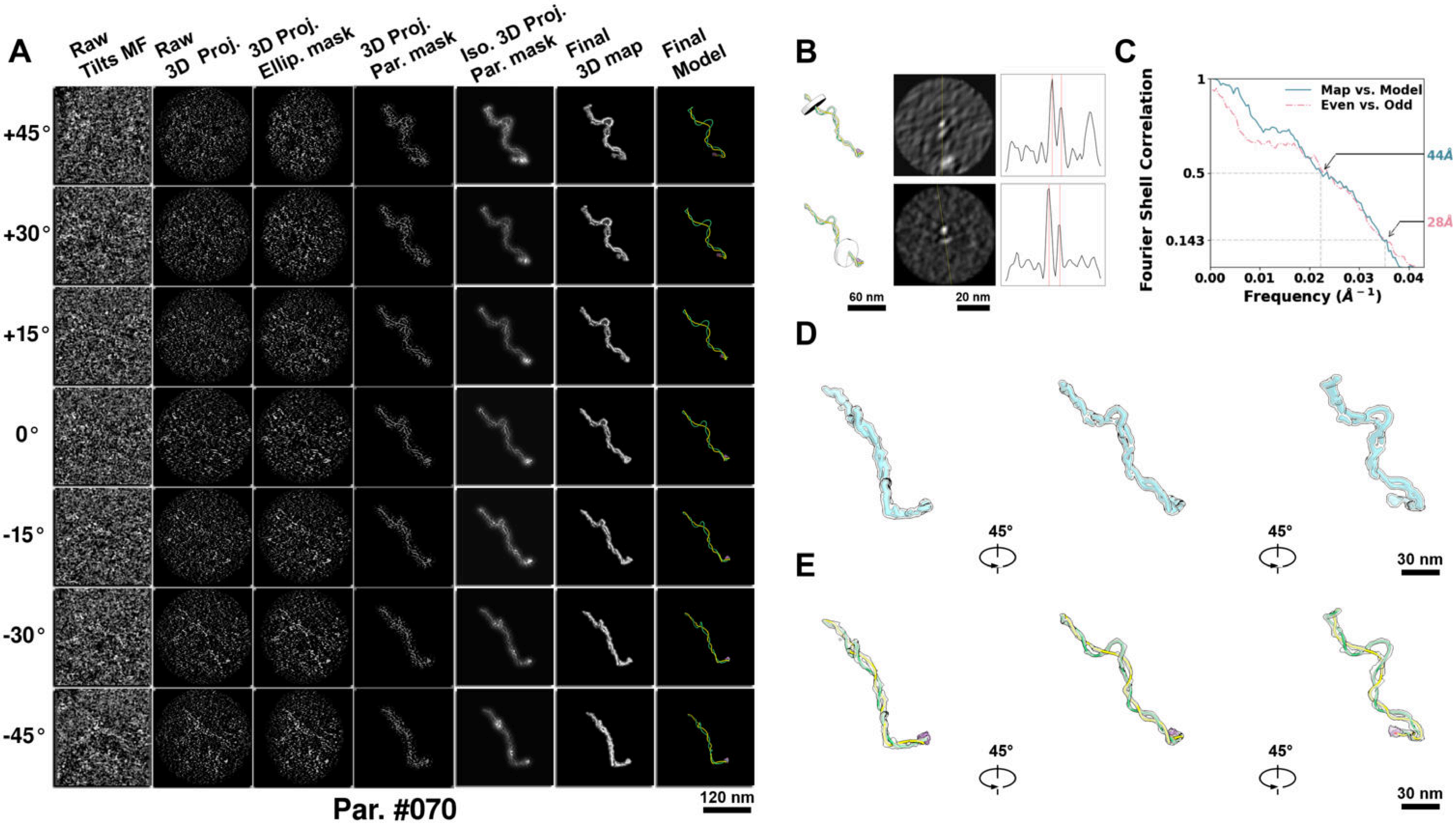

**Supplementary Particle Figure 70. Cryo-ET 3D reconstruction of an individual sTEC particle.**
(A) 3D reconstruction of the plasmid particle (index no. 70). The first column shows seven representative tilt images from +45° to -45° in step of 15°. The second, third, and fourth columns show 3D projections of the particle with spherical, ellipsoidal (thinner along the z-dimension), and particle-shaped masks, respectively. The fifth column displays the 3D projections of the enhanced and IsoNet missing-wedge-corrected particle. The sixth and seventh columns present the final 3D map and the flexibly fitted model, respectively. (B) Two cross-sectional views (12 nm thickness) of the plasmid density map along its plectoneme axis are shown in the left-middle panel. The intensity profile along the line crossing the two high-density DNA spots is displayed in the right panel. (C) Resolution assessment of the final 3D map using Fourier shell correlation (FSC). Two criteria are shown: FSC between two half-maps reconstructed from even and odd frames (evaluated at 0.143) and FSC between the final 3D map and the fitted model (evaluated at 0.5). (D) Zoomed-in views of the final 3D density map from panel A, displayed at two contour levels. (E) Superimposition of the high-contour level map from panel D onto its fitted model.

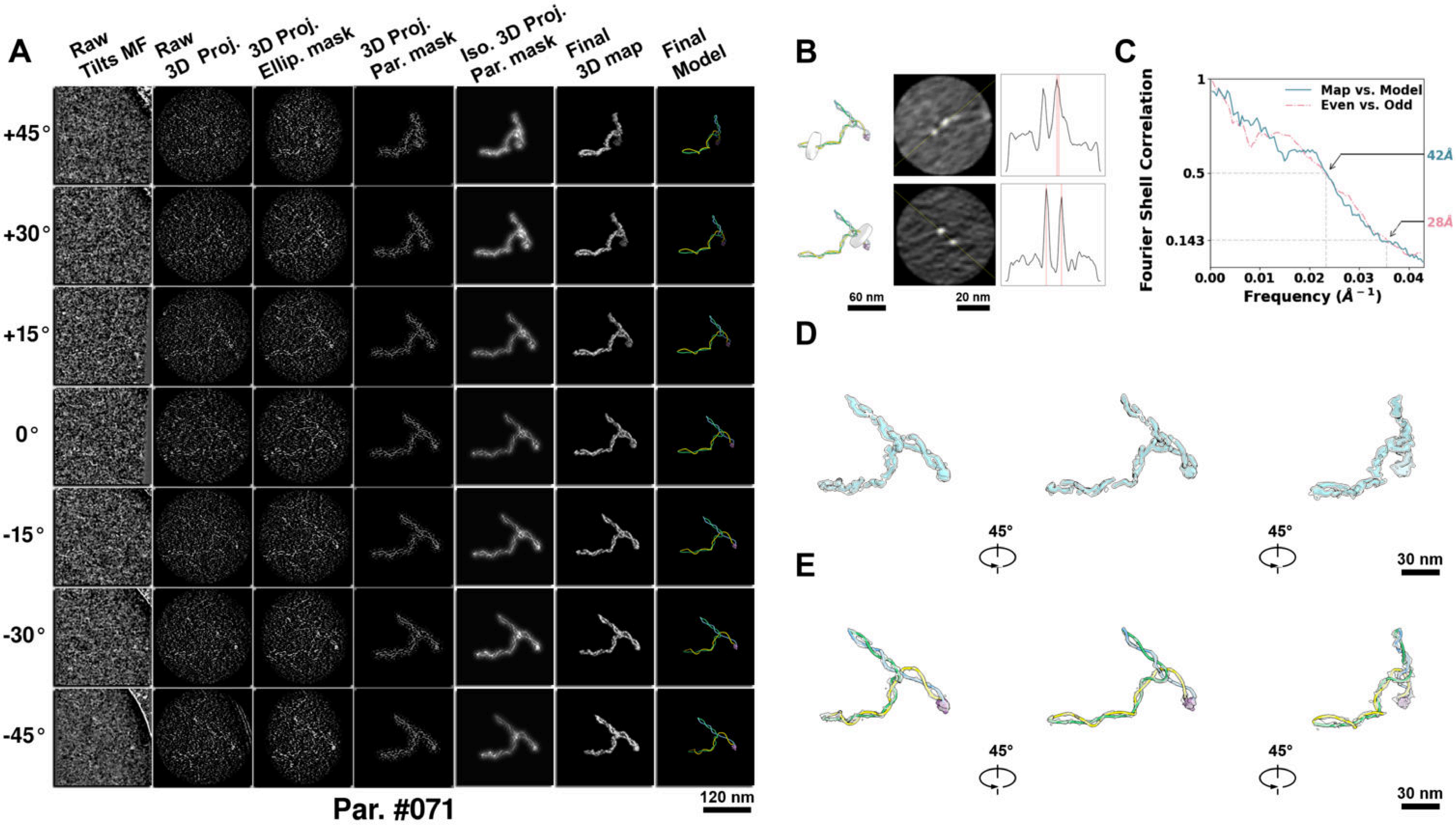

**Supplementary Particle Figure 71. Cryo-ET 3D reconstruction of an individual sTEC particle.**
(A) 3D reconstruction of the plasmid particle (index no. 71). The first column shows seven representative tilt images from +45° to -45° in step of 15°. The second, third, and fourth columns show 3D projections of the particle with spherical, ellipsoidal (thinner along the z-dimension), and particle-shaped masks, respectively. The fifth column displays the 3D projections of the enhanced and IsoNet missing-wedge-corrected particle. The sixth and seventh columns present the final 3D map and the flexibly fitted model, respectively. (B) Two cross-sectional views (12 nm thickness) of the plasmid density map along its plectoneme axis are shown in the left-middle panel. The intensity profile along the line crossing the two high-density DNA spots is displayed in the right panel. (C) Resolution assessment of the final 3D map using Fourier shell correlation (FSC). Two criteria are shown: FSC between two half-maps reconstructed from even and odd frames (evaluated at 0.143) and FSC between the final 3D map and the fitted model (evaluated at 0.5). (D) Zoomed-in views of the final 3D density map from panel A, displayed at two contour levels. (E) Superimposition of the high-contour level map from panel D onto its fitted model.

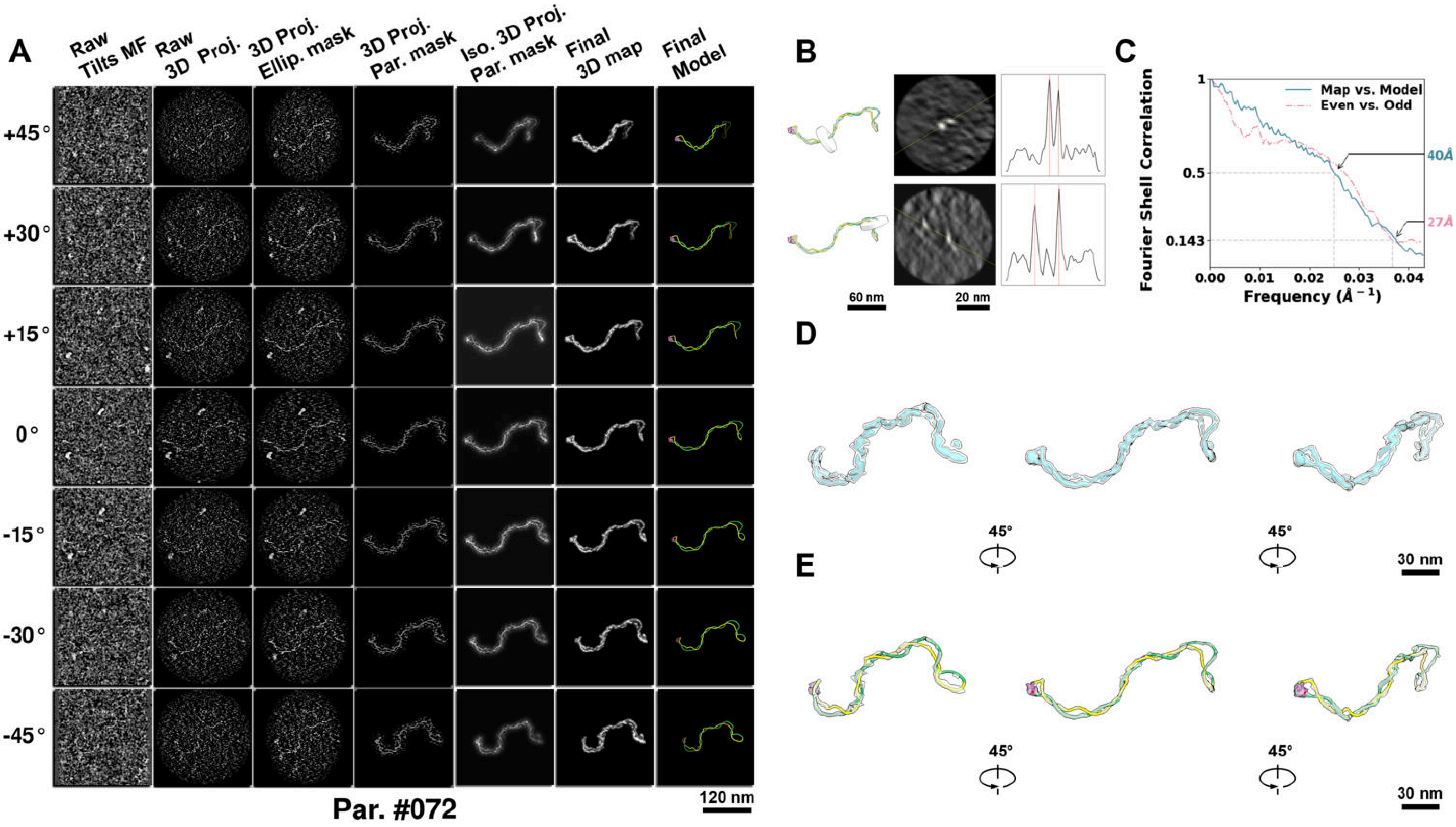

**Supplementary Particle Figure 72. Cryo-ET 3D reconstruction of an individual sTEC particle.**
(A) 3D reconstruction of the plasmid particle (index no. 72). The first column shows seven representative tilt images from +45° to -45° in step of 15°. The second, third, and fourth columns show 3D projections of the particle with spherical, ellipsoidal (thinner along the z-dimension), and particle-shaped masks, respectively. The fifth column displays the 3D projections of the enhanced and IsoNet missing-wedge-corrected particle. The sixth and seventh columns present the final 3D map and the flexibly fitted model, respectively. (B) Two cross-sectional views (12 nm thickness) of the plasmid density map along its plectoneme axis are shown in the left-middle panel. The intensity profile along the line crossing the two high-density DNA spots is displayed in the right panel. (C) Resolution assessment of the final 3D map using Fourier shell correlation (FSC). Two criteria are shown: FSC between two half-maps reconstructed from even and odd frames (evaluated at 0.143) and FSC between the final 3D map and the fitted model (evaluated at 0.5). (D) Zoomed-in views of the final 3D density map from panel A, displayed at two contour levels. (E) Superimposition of the high-contour level map from panel D onto its fitted model.

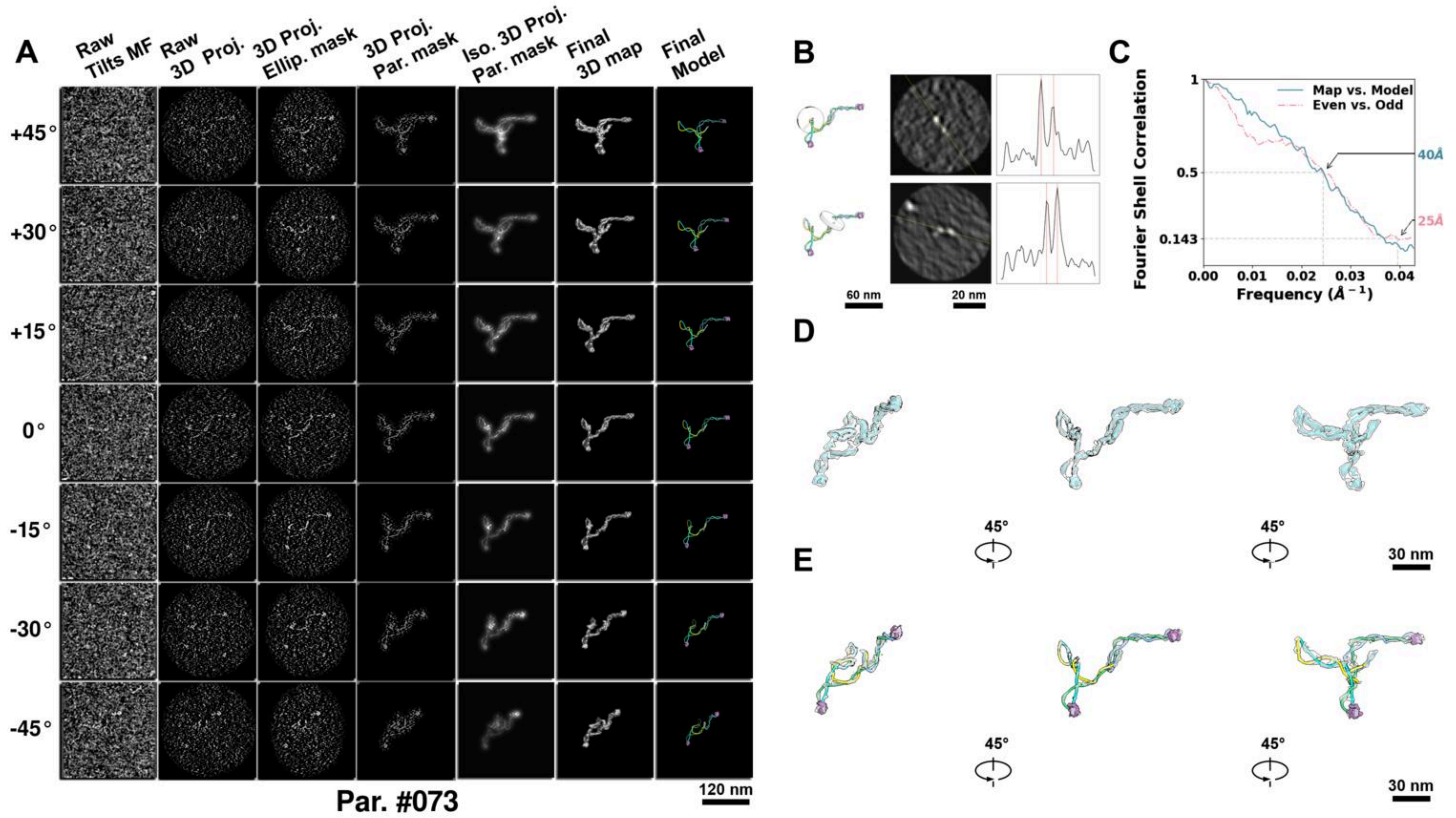

**Supplementary Particle Figure 73. Cryo-ET 3D reconstruction of an individual sTEC particle.**
(A) 3D reconstruction of the plasmid particle (index no. 73). The first column shows seven representative tilt images from +45° to -45° in step of 15°. The second, third, and fourth columns show 3D projections of the particle with spherical, ellipsoidal (thinner along the z-dimension), and particle-shaped masks, respectively. The fifth column displays the 3D projections of the enhanced and IsoNet missing-wedge-corrected particle. The sixth and seventh columns present the final 3D map and the flexibly fitted model, respectively. (B) Two cross-sectional views (12 nm thickness) of the plasmid density map along its plectoneme axis are shown in the left-middle panel. The intensity profile along the line crossing the two high-density DNA spots is displayed in the right panel. (C) Resolution assessment of the final 3D map using Fourier shell correlation (FSC). Two criteria are shown: FSC between two half-maps reconstructed from even and odd frames (evaluated at 0.143) and FSC between the final 3D map and the fitted model (evaluated at 0.5). (D) Zoomed-in views of the final 3D density map from panel A, displayed at two contour levels. (E) Superimposition of the high-contour level map from panel D onto its fitted model.

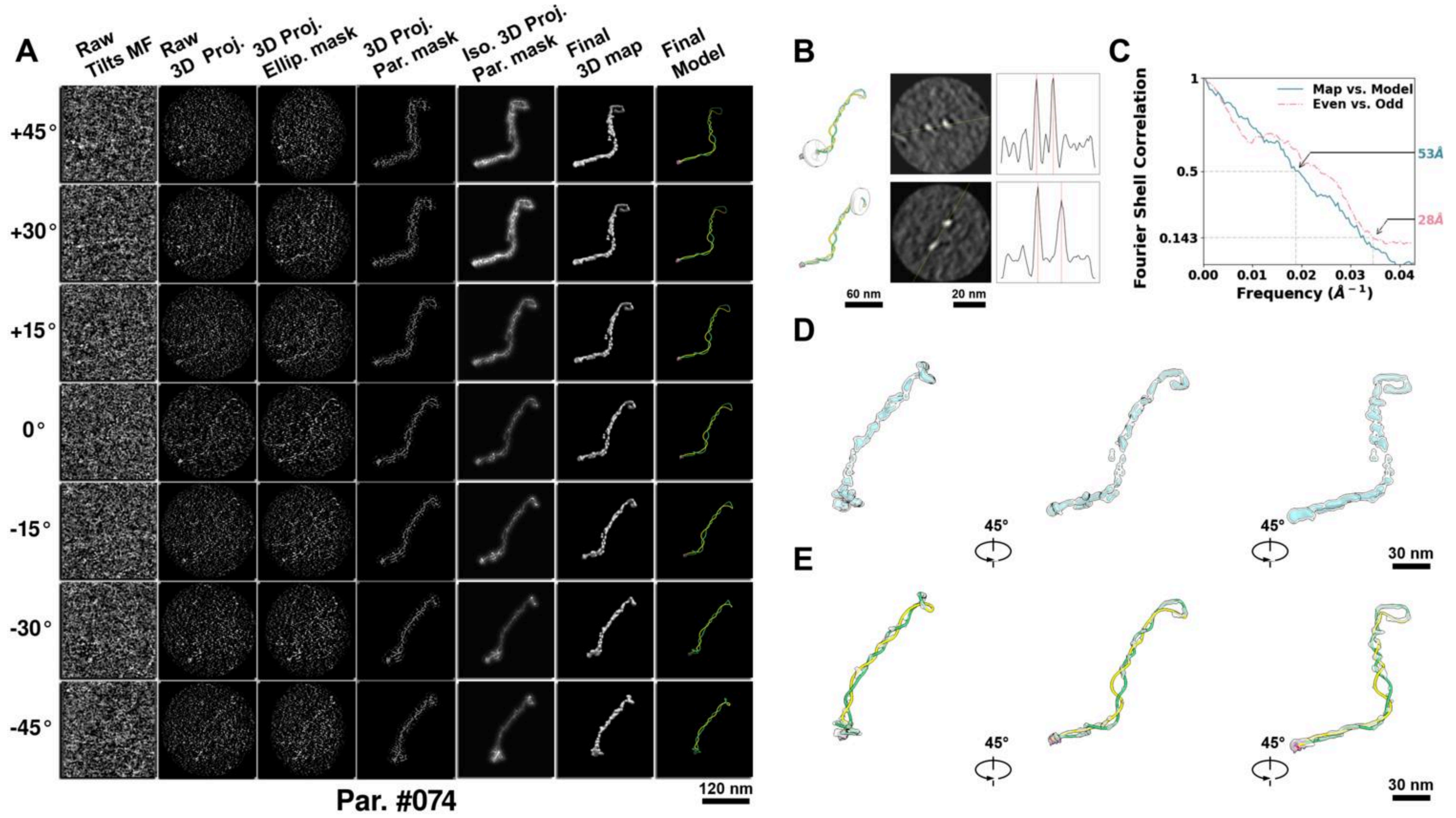

**Supplementary Particle Figure 74. Cryo-ET 3D reconstruction of an individual sTEC particle.**
(A) 3D reconstruction of the plasmid particle (index no. 74). The first column shows seven representative tilt images from +45° to -45° in step of 15°. The second, third, and fourth columns show 3D projections of the particle with spherical, ellipsoidal (thinner along the z-dimension), and particle-shaped masks, respectively. The fifth column displays the 3D projections of the enhanced and IsoNet missing-wedge-corrected particle. The sixth and seventh columns present the final 3D map and the flexibly fitted model, respectively. (B) Two cross-sectional views (12 nm thickness) of the plasmid density map along its plectoneme axis are shown in the left-middle panel. The intensity profile along the line crossing the two high-density DNA spots is displayed in the right panel. (C) Resolution assessment of the final 3D map using Fourier shell correlation (FSC). Two criteria are shown: FSC between two half-maps reconstructed from even and odd frames (evaluated at 0.143) and FSC between the final 3D map and the fitted model (evaluated at 0.5). (D) Zoomed-in views of the final 3D density map from panel A, displayed at two contour levels. (E) Superimposition of the high-contour level map from panel D onto its fitted model.

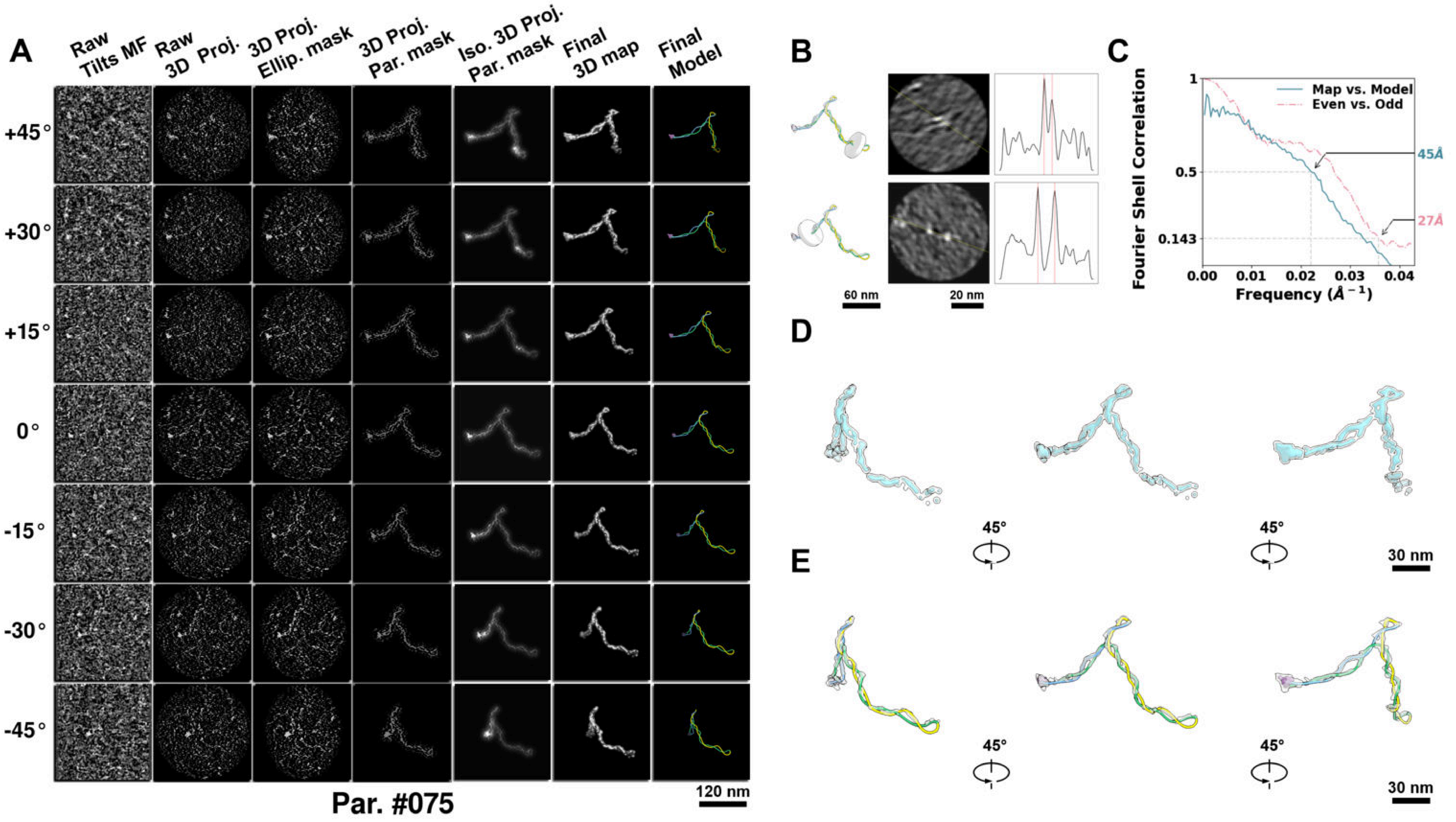

**Supplementary Particle Figure 75. Cryo-ET 3D reconstruction of an individual sTEC particle.**
(A) 3D reconstruction of the plasmid particle (index no. 75). The first column shows seven representative tilt images from +45° to -45° in step of 15°. The second, third, and fourth columns show 3D projections of the particle with spherical, ellipsoidal (thinner along the z-dimension), and particle-shaped masks, respectively. The fifth column displays the 3D projections of the enhanced and IsoNet missing-wedge-corrected particle. The sixth and seventh columns present the final 3D map and the flexibly fitted model, respectively. (B) Two cross-sectional views (12 nm thickness) of the plasmid density map along its plectoneme axis are shown in the left-middle panel. The intensity profile along the line crossing the two high-density DNA spots is displayed in the right panel. (C) Resolution assessment of the final 3D map using Fourier shell correlation (FSC). Two criteria are shown: FSC between two half-maps reconstructed from even and odd frames (evaluated at 0.143) and FSC between the final 3D map and the fitted model (evaluated at 0.5). (D) Zoomed-in views of the final 3D density map from panel A, displayed at two contour levels. (E) Superimposition of the high-contour level map from panel D onto its fitted model.

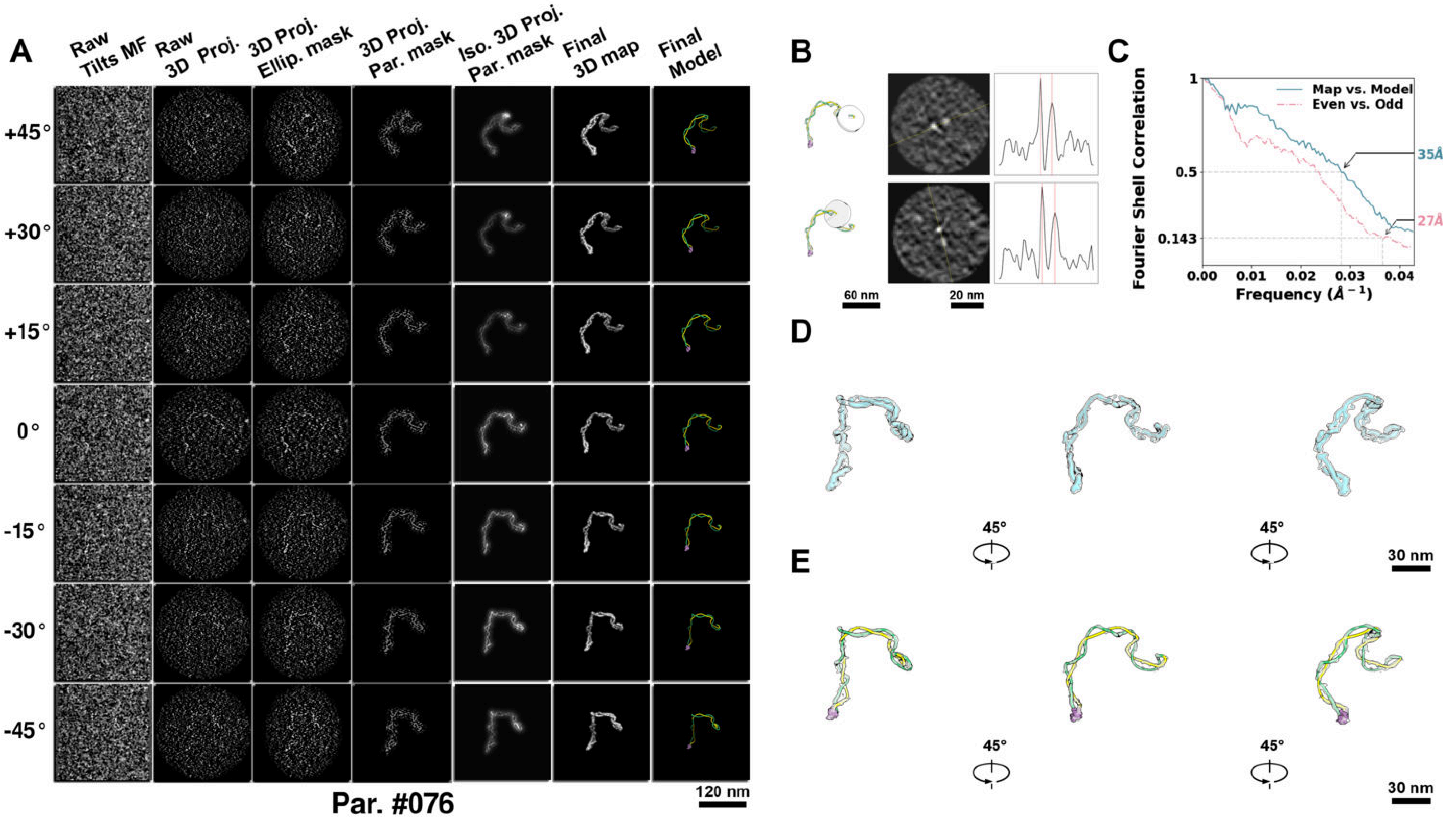

**Supplementary Particle Figure 76. Cryo-ET 3D reconstruction of an individual sTEC particle.**
(A) 3D reconstruction of the plasmid particle (index no. 76). The first column shows seven representative tilt images from +45° to -45° in step of 15°. The second, third, and fourth columns show 3D projections of the particle with spherical, ellipsoidal (thinner along the z-dimension), and particle-shaped masks, respectively. The fifth column displays the 3D projections of the enhanced and IsoNet missing-wedge-corrected particle. The sixth and seventh columns present the final 3D map and the flexibly fitted model, respectively. (B) Two cross-sectional views (12 nm thickness) of the plasmid density map along its plectoneme axis are shown in the left-middle panel. The intensity profile along the line crossing the two high-density DNA spots is displayed in the right panel. (C) Resolution assessment of the final 3D map using Fourier shell correlation (FSC). Two criteria are shown: FSC between two half-maps reconstructed from even and odd frames (evaluated at 0.143) and FSC between the final 3D map and the fitted model (evaluated at 0.5). (D) Zoomed-in views of the final 3D density map from panel A, displayed at two contour levels. (E) Superimposition of the high-contour level map from panel D onto its fitted model.

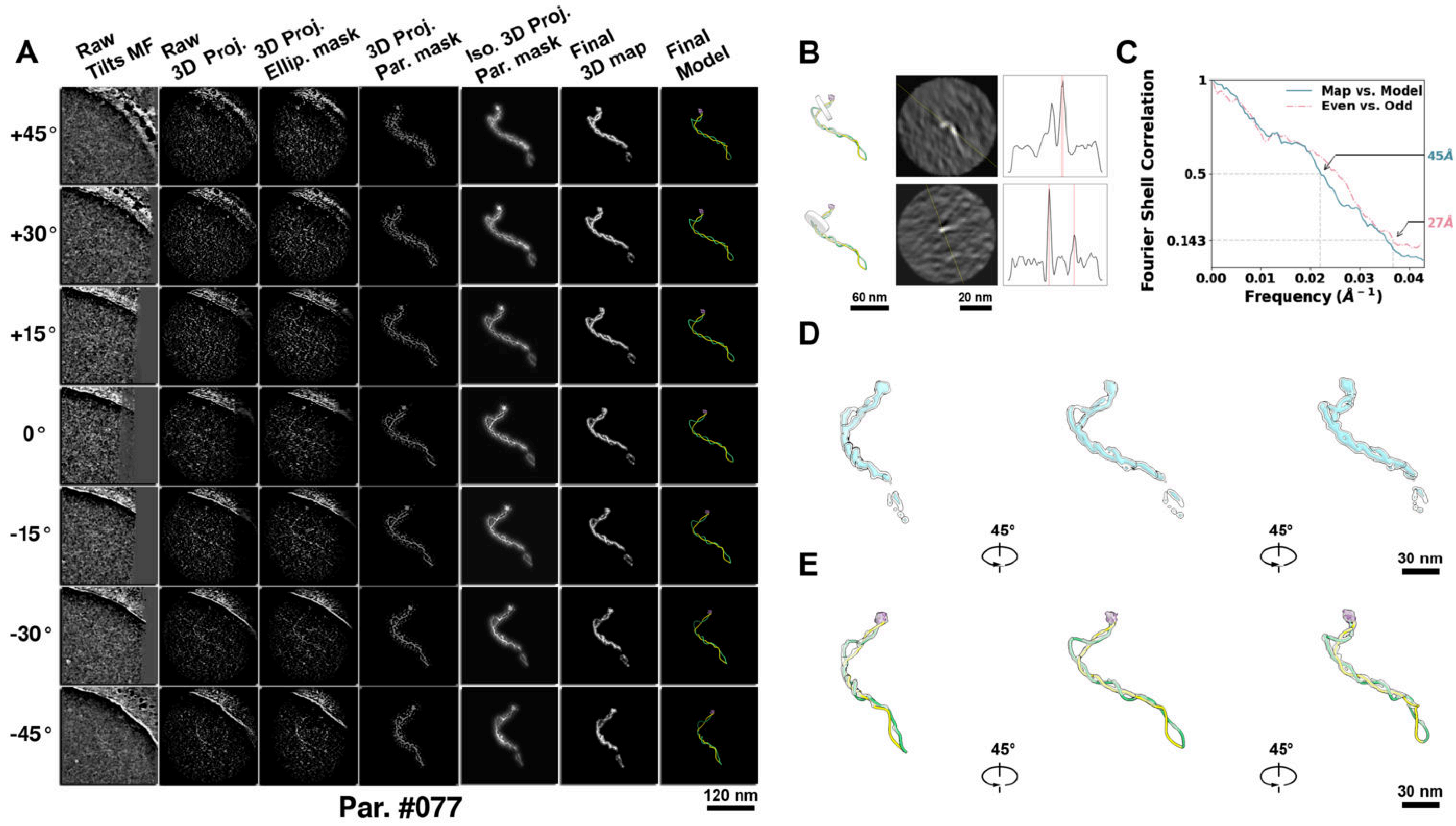

**Supplementary Particle Figure 77. Cryo-ET 3D reconstruction of an individual sTEC particle.**
(A) 3D reconstruction of the plasmid particle (index no. 77). The first column shows seven representative tilt images from +45° to -45° in step of 15°. The second, third, and fourth columns show 3D projections of the particle with spherical, ellipsoidal (thinner along the z-dimension), and particle-shaped masks, respectively. The fifth column displays the 3D projections of the enhanced and IsoNet missing-wedge-corrected particle. The sixth and seventh columns present the final 3D map and the flexibly fitted model, respectively. (B) Two cross-sectional views (12 nm thickness) of the plasmid density map along its plectoneme axis are shown in the left-middle panel. The intensity profile along the line crossing the two high-density DNA spots is displayed in the right panel. (C) Resolution assessment of the final 3D map using Fourier shell correlation (FSC). Two criteria are shown: FSC between two half-maps reconstructed from even and odd frames (evaluated at 0.143) and FSC between the final 3D map and the fitted model (evaluated at 0.5). (D) Zoomed-in views of the final 3D density map from panel A, displayed at two contour levels. (E) Superimposition of the high-contour level map from panel D onto its fitted model.

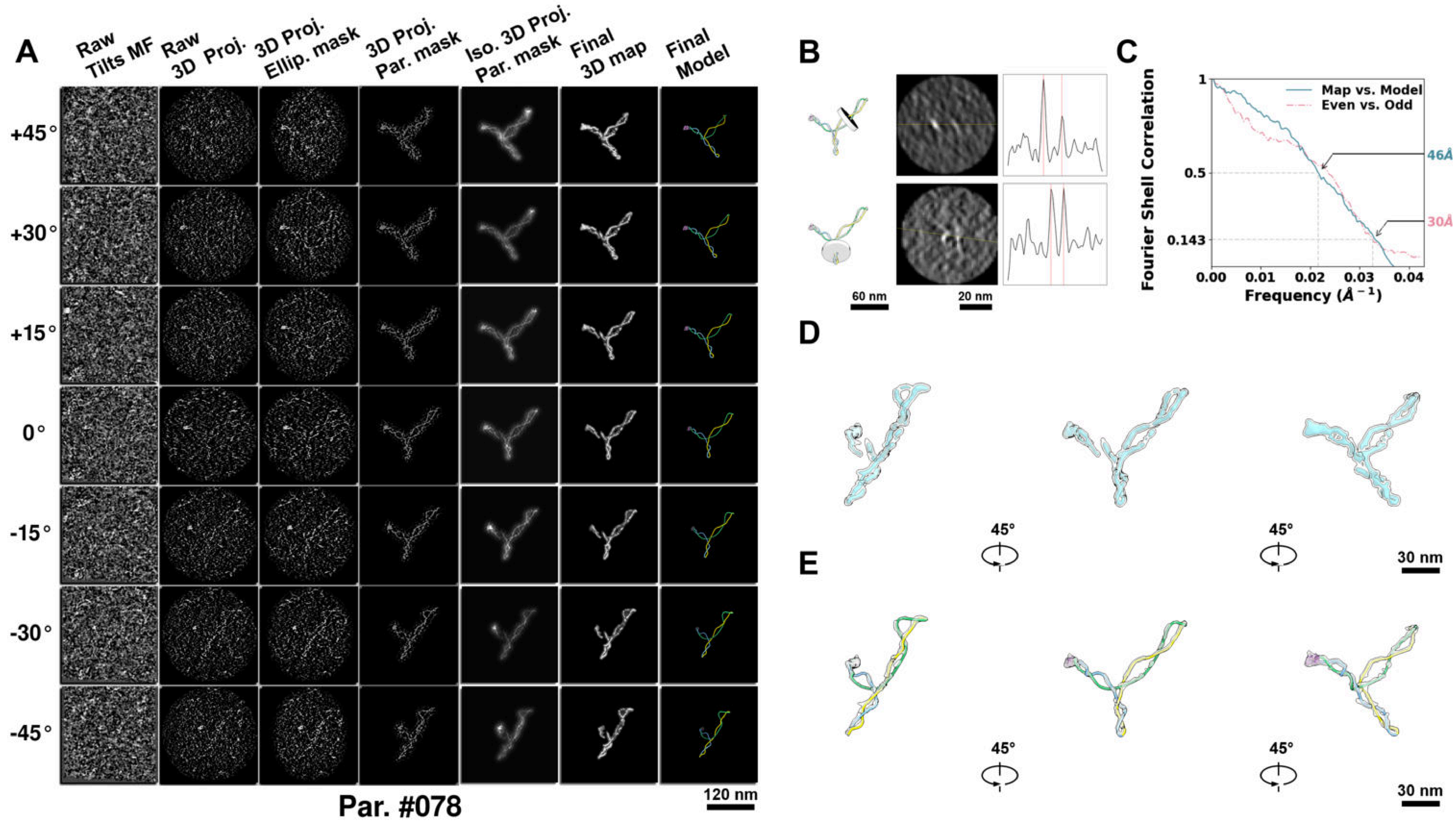

**Supplementary Particle Figure 78. Cryo-ET 3D reconstruction of an individual sTEC particle.**
(A) 3D reconstruction of the plasmid particle (index no. 78). The first column shows seven representative tilt images from +45° to -45° in step of 15°. The second, third, and fourth columns show 3D projections of the particle with spherical, ellipsoidal (thinner along the z-dimension), and particle-shaped masks, respectively. The fifth column displays the 3D projections of the enhanced and IsoNet missing-wedge-corrected particle. The sixth and seventh columns present the final 3D map and the flexibly fitted model, respectively. (B) Two cross-sectional views (12 nm thickness) of the plasmid density map along its plectoneme axis are shown in the left-middle panel. The intensity profile along the line crossing the two high-density DNA spots is displayed in the right panel. (C) Resolution assessment of the final 3D map using Fourier shell correlation (FSC). Two criteria are shown: FSC between two half-maps reconstructed from even and odd frames (evaluated at 0.143) and FSC between the final 3D map and the fitted model (evaluated at 0.5). (D) Zoomed-in views of the final 3D density map from panel A, displayed at two contour levels. (E) Superimposition of the high-contour level map from panel D onto its fitted model.

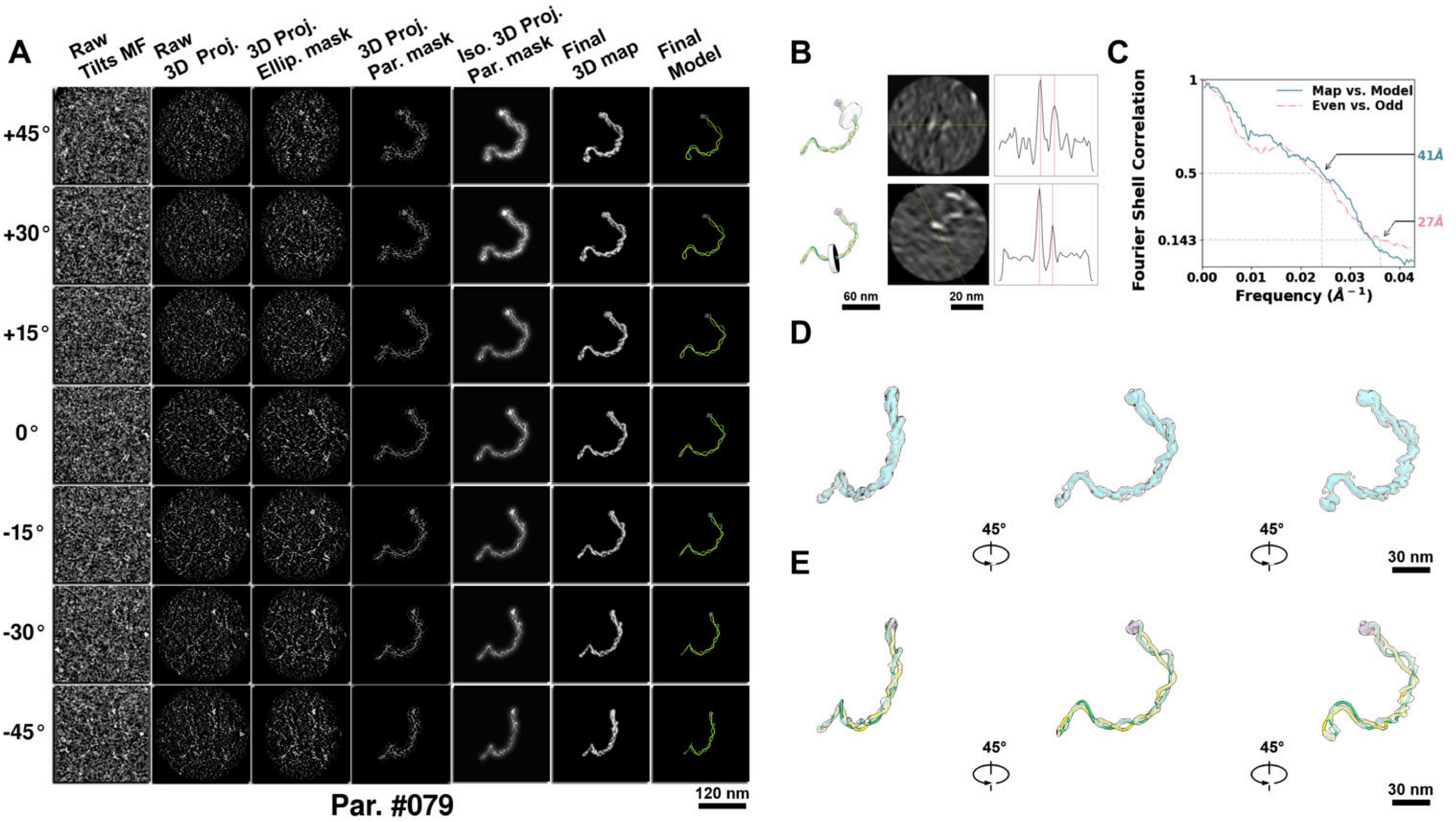

**Supplementary Particle Figure 79. Cryo-ET 3D reconstruction of an individual sTEC particle.**
(A) 3D reconstruction of the plasmid particle (index no. 79). The first column shows seven representative tilt images from +45° to -45° in step of 15°. The second, third, and fourth columns show 3D projections of the particle with spherical, ellipsoidal (thinner along the z-dimension), and particle-shaped masks, respectively. The fifth column displays the 3D projections of the enhanced and IsoNet missing-wedge-corrected particle. The sixth and seventh columns present the final 3D map and the flexibly fitted model, respectively. (B) Two cross-sectional views (12 nm thickness) of the plasmid density map along its plectoneme axis are shown in the left-middle panel. The intensity profile along the line crossing the two high-density DNA spots is displayed in the right panel. (C) Resolution assessment of the final 3D map using Fourier shell correlation (FSC). Two criteria are shown: FSC between two half-maps reconstructed from even and odd frames (evaluated at 0.143) and FSC between the final 3D map and the fitted model (evaluated at 0.5). (D) Zoomed-in views of the final 3D density map from panel A, displayed at two contour levels. (E) Superimposition of the high-contour level map from panel D onto its fitted model.

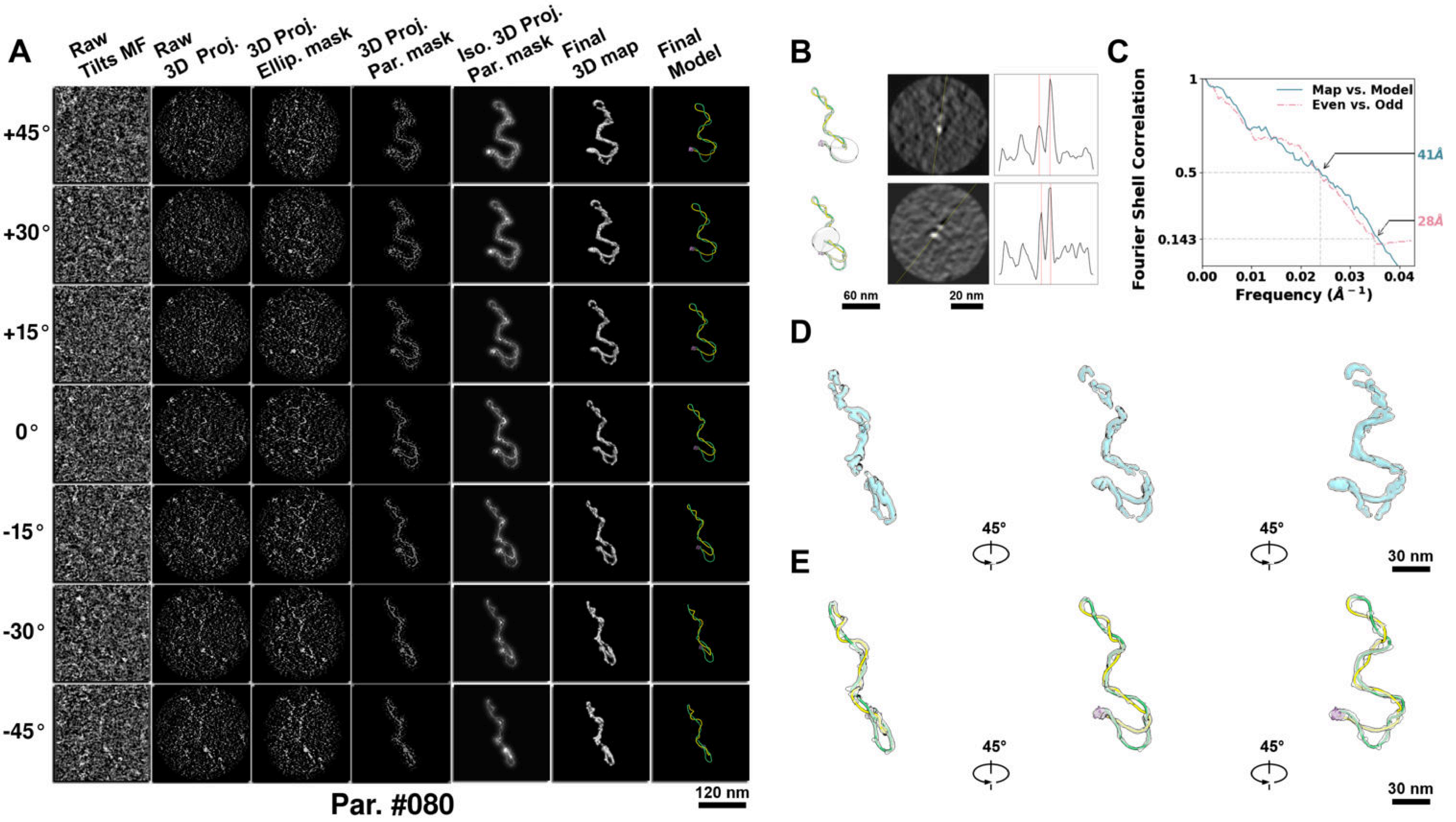

**Supplementary Particle Figure 80. Cryo-ET 3D reconstruction of an individual sTEC particle.**
(A) 3D reconstruction of the plasmid particle (index no. 80). The first column shows seven representative tilt images from +45° to -45° in step of 15°. The second, third, and fourth columns show 3D projections of the particle with spherical, ellipsoidal (thinner along the z-dimension), and particle-shaped masks, respectively. The fifth column displays the 3D projections of the enhanced and IsoNet missing-wedge-corrected particle. The sixth and seventh columns present the final 3D map and the flexibly fitted model, respectively. (B) Two cross-sectional views (12 nm thickness) of the plasmid density map along its plectoneme axis are shown in the left-middle panel. The intensity profile along the line crossing the two high-density DNA spots is displayed in the right panel. (C) Resolution assessment of the final 3D map using Fourier shell correlation (FSC). Two criteria are shown: FSC between two half-maps reconstructed from even and odd frames (evaluated at 0.143) and FSC between the final 3D map and the fitted model (evaluated at 0.5). (D) Zoomed-in views of the final 3D density map from panel A, displayed at two contour levels. (E) Superimposition of the high-contour level map from panel D onto its fitted model.

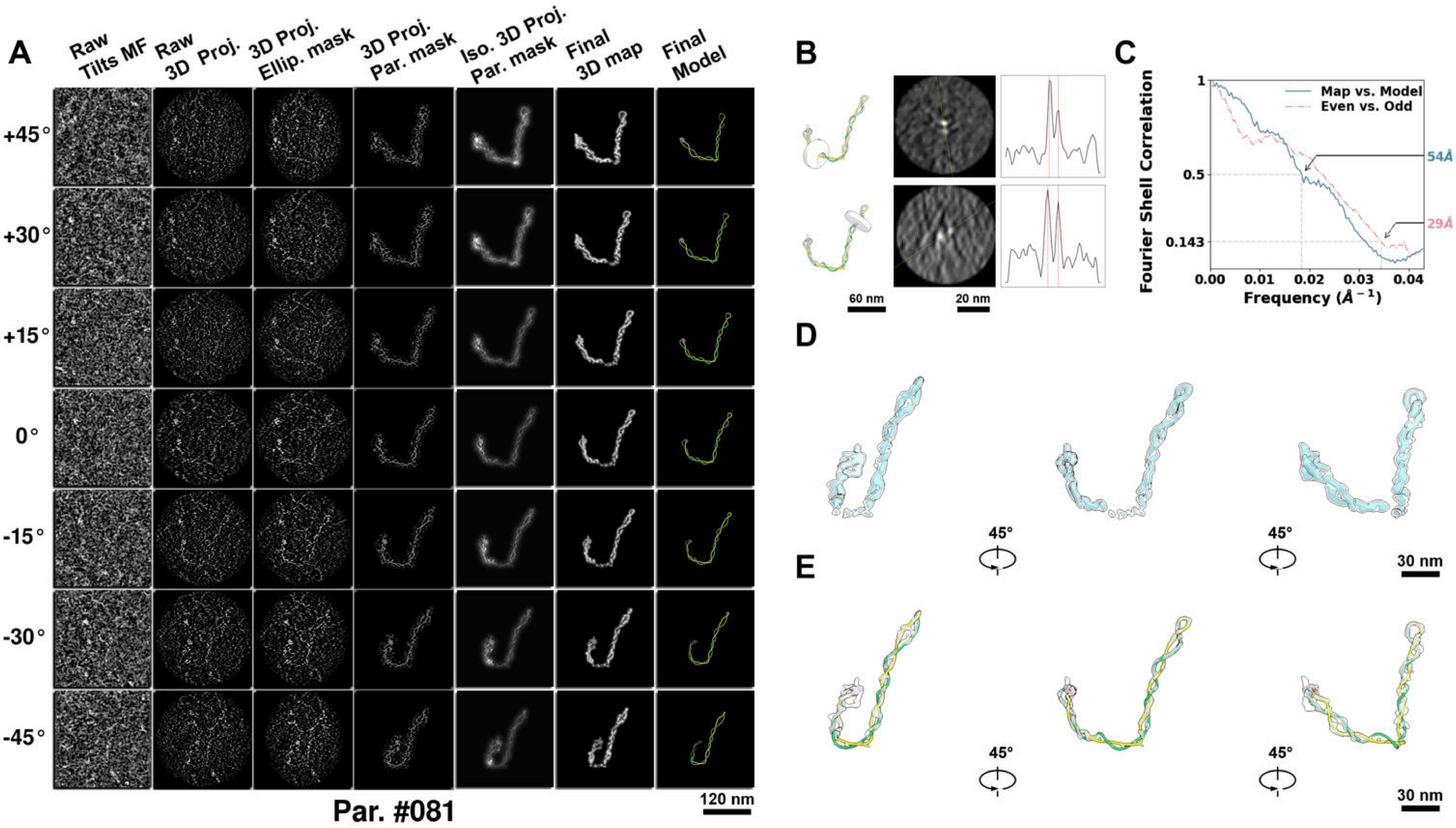

**Supplementary Particle Figure 81. Cryo-ET 3D reconstruction of an individual sTEC particle.**
(A) 3D reconstruction of the plasmid particle (index no. 81). The first column shows seven representative tilt images from +45° to -45° in step of 15°. The second, third, and fourth columns show 3D projections of the particle with spherical, ellipsoidal (thinner along the z-dimension), and particle-shaped masks, respectively. The fifth column displays the 3D projections of the enhanced and IsoNet missing-wedge-corrected particle. The sixth and seventh columns present the final 3D map and the flexibly fitted model, respectively. (B) Two cross-sectional views (12 nm thickness) of the plasmid density map along its plectoneme axis are shown in the left-middle panel. The intensity profile along the line crossing the two high-density DNA spots is displayed in the right panel. (C) Resolution assessment of the final 3D map using Fourier shell correlation (FSC). Two criteria are shown: FSC between two half-maps reconstructed from even and odd frames (evaluated at 0.143) and FSC between the final 3D map and the fitted model (evaluated at 0.5). (D) Zoomed-in views of the final 3D density map from panel A, displayed at two contour levels. (E) Superimposition of the high-contour level map from panel D onto its fitted model.

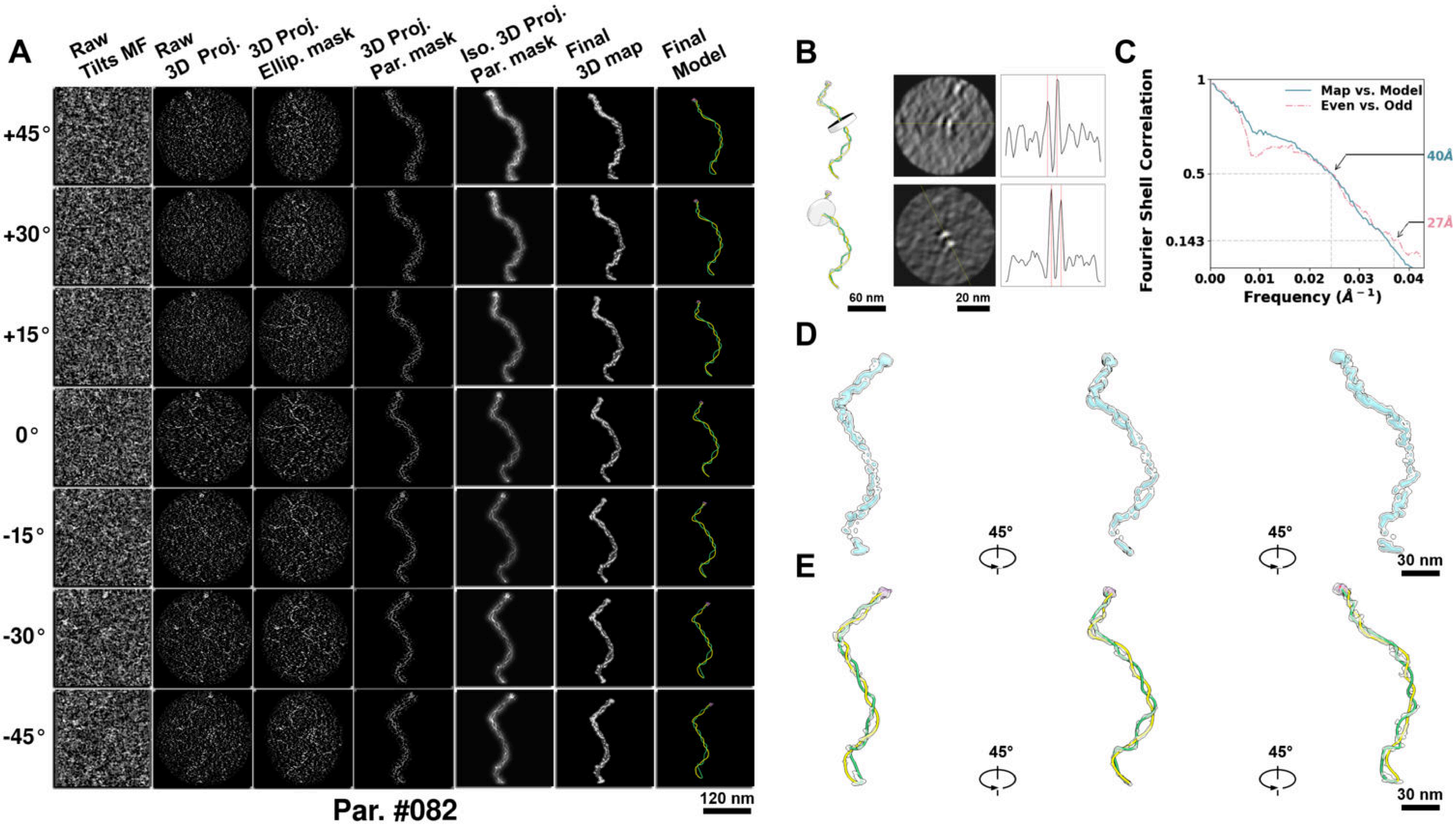

**Supplementary Particle Figure 82. Cryo-ET 3D reconstruction of an individual sTEC particle.**
(A) 3D reconstruction of the plasmid particle (index no. 82). The first column shows seven representative tilt images from +45° to -45° in step of 15°. The second, third, and fourth columns show 3D projections of the particle with spherical, ellipsoidal (thinner along the z-dimension), and particle-shaped masks, respectively. The fifth column displays the 3D projections of the enhanced and IsoNet missing-wedge-corrected particle. The sixth and seventh columns present the final 3D map and the flexibly fitted model, respectively. (B) Two cross-sectional views (12 nm thickness) of the plasmid density map along its plectoneme axis are shown in the left-middle panel. The intensity profile along the line crossing the two high-density DNA spots is displayed in the right panel. (C) Resolution assessment of the final 3D map using Fourier shell correlation (FSC). Two criteria are shown: FSC between two half-maps reconstructed from even and odd frames (evaluated at 0.143) and FSC between the final 3D map and the fitted model (evaluated at 0.5). (D) Zoomed-in views of the final 3D density map from panel A, displayed at two contour levels. (E) Superimposition of the high-contour level map from panel D onto its fitted model.

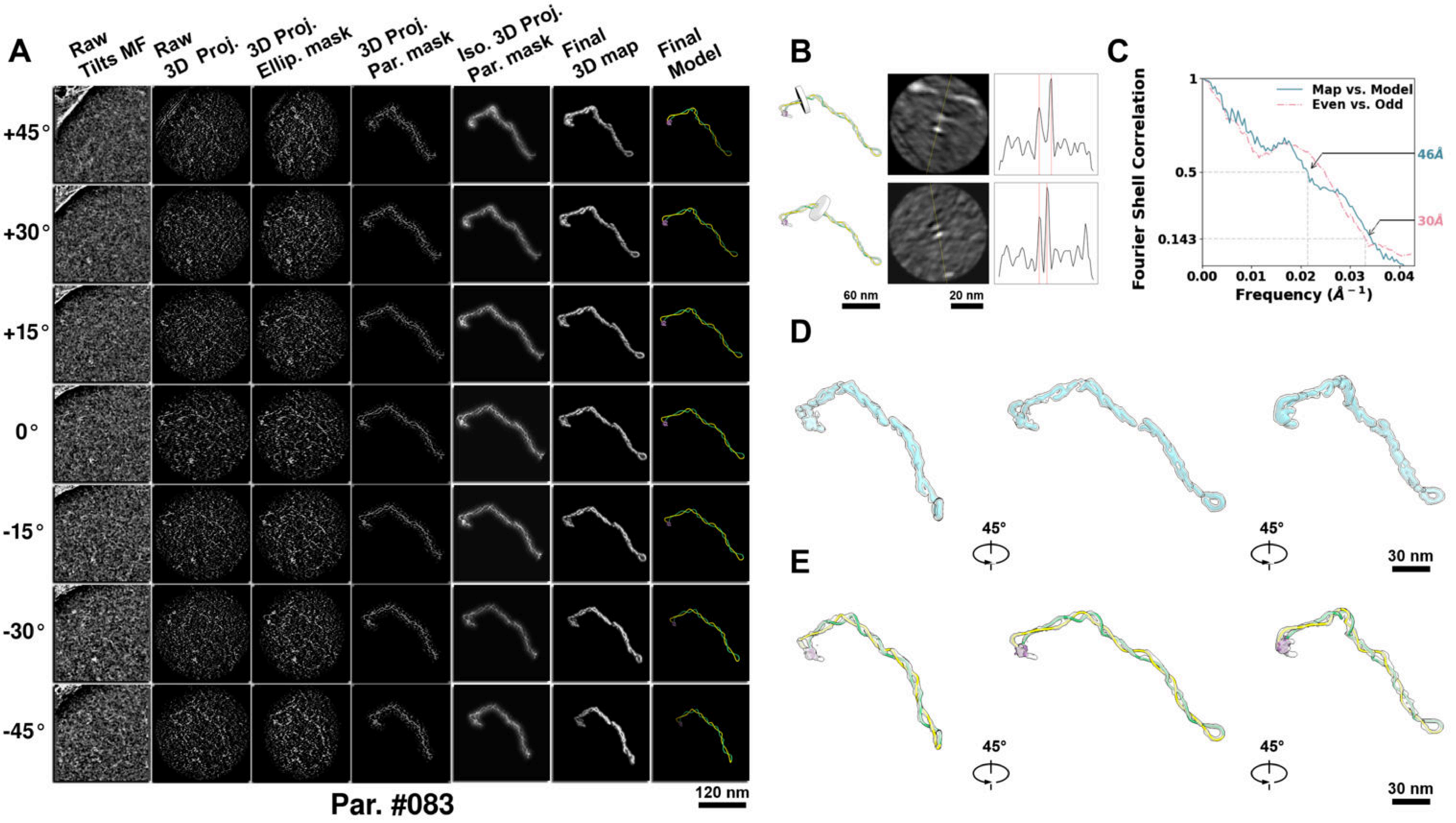

**Supplementary Particle Figure 83. Cryo-ET 3D reconstruction of an individual sTEC particle.**
(A) 3D reconstruction of the plasmid particle (index no. 83). The first column shows seven representative tilt images from +45° to -45° in step of 15°. The second, third, and fourth columns show 3D projections of the particle with spherical, ellipsoidal (thinner along the z-dimension), and particle-shaped masks, respectively. The fifth column displays the 3D projections of the enhanced and IsoNet missing-wedge-corrected particle. The sixth and seventh columns present the final 3D map and the flexibly fitted model, respectively. (B) Two cross-sectional views (12 nm thickness) of the plasmid density map along its plectoneme axis are shown in the left-middle panel. The intensity profile along the line crossing the two high-density DNA spots is displayed in the right panel. (C) Resolution assessment of the final 3D map using Fourier shell correlation (FSC). Two criteria are shown: FSC between two half-maps reconstructed from even and odd frames (evaluated at 0.143) and FSC between the final 3D map and the fitted model (evaluated at 0.5). (D) Zoomed-in views of the final 3D density map from panel A, displayed at two contour levels. (E) Superimposition of the high-contour level map from panel D onto its fitted model.

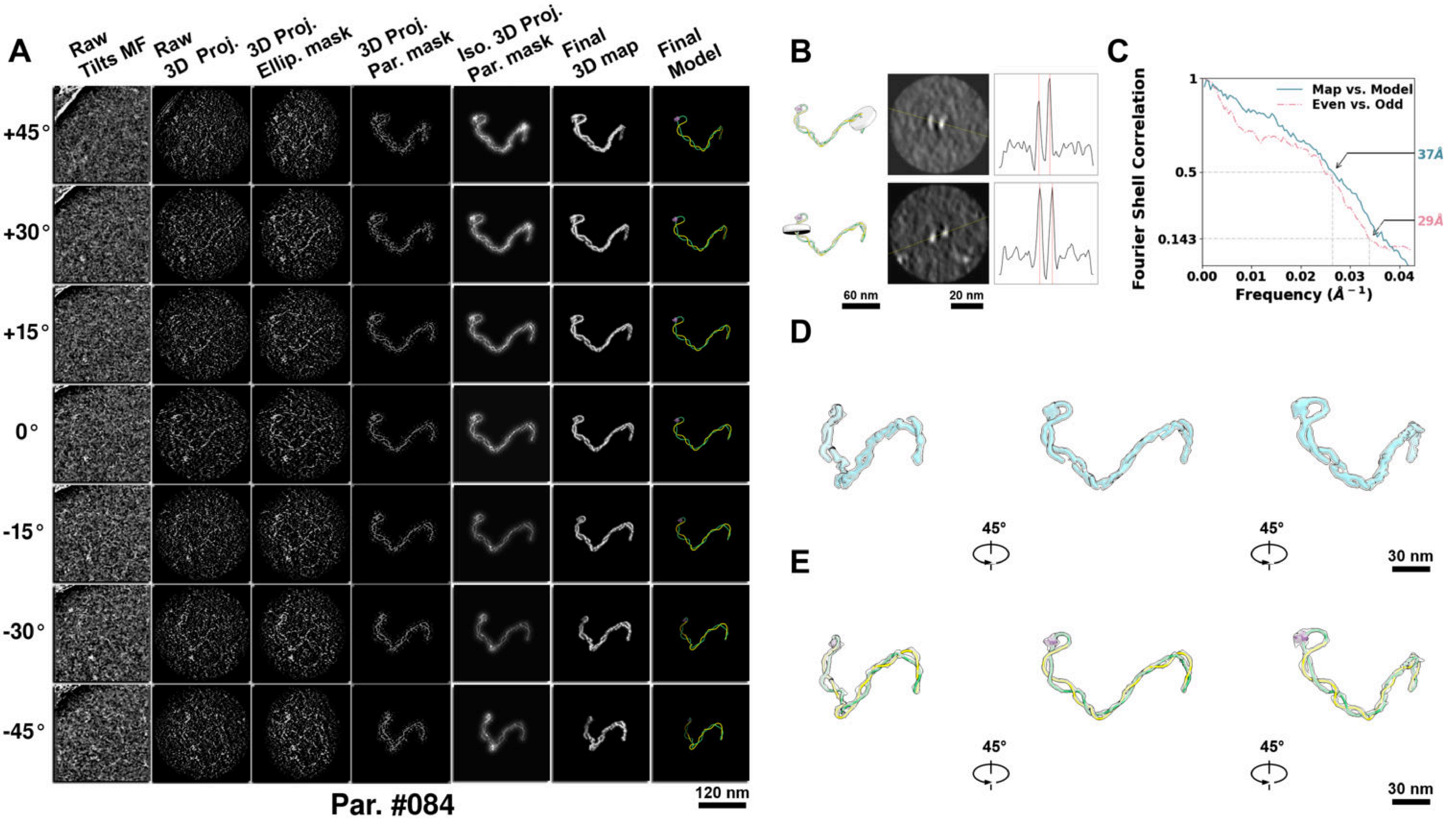

**Supplementary Particle Figure 84. Cryo-ET 3D reconstruction of an individual sTEC particle.**
(A) 3D reconstruction of the plasmid particle (index no. 84). The first column shows seven representative tilt images from +45° to -45° in step of 15°. The second, third, and fourth columns show 3D projections of the particle with spherical, ellipsoidal (thinner along the z-dimension), and particle-shaped masks, respectively. The fifth column displays the 3D projections of the enhanced and IsoNet missing-wedge-corrected particle. The sixth and seventh columns present the final 3D map and the flexibly fitted model, respectively. (B) Two cross-sectional views (12 nm thickness) of the plasmid density map along its plectoneme axis are shown in the left-middle panel. The intensity profile along the line crossing the two high-density DNA spots is displayed in the right panel. (C) Resolution assessment of the final 3D map using Fourier shell correlation (FSC). Two criteria are shown: FSC between two half-maps reconstructed from even and odd frames (evaluated at 0.143) and FSC between the final 3D map and the fitted model (evaluated at 0.5). (D) Zoomed-in views of the final 3D density map from panel A, displayed at two contour levels. (E) Superimposition of the high-contour level map from panel D onto its fitted model.

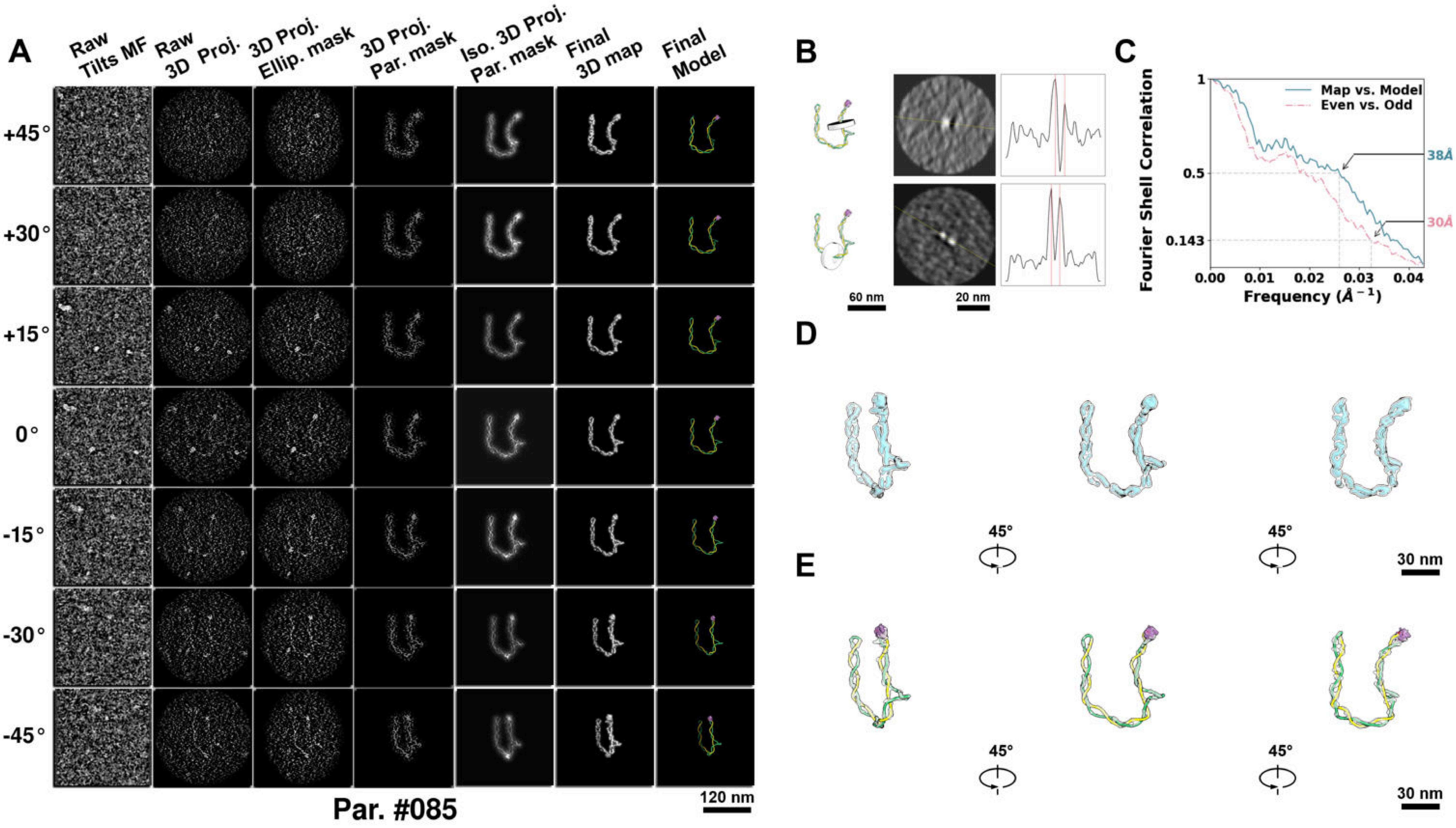

**Supplementary Particle Figure 85. Cryo-ET 3D reconstruction of an individual sTEC particle.**
(A) 3D reconstruction of the plasmid particle (index no. 85). The first column shows seven representative tilt images from +45° to -45° in step of 15°. The second, third, and fourth columns show 3D projections of the particle with spherical, ellipsoidal (thinner along the z-dimension), and particle-shaped masks, respectively. The fifth column displays the 3D projections of the enhanced and IsoNet missing-wedge-corrected particle. The sixth and seventh columns present the final 3D map and the flexibly fitted model, respectively. (B) Two cross-sectional views (12 nm thickness) of the plasmid density map along its plectoneme axis are shown in the left-middle panel. The intensity profile along the line crossing the two high-density DNA spots is displayed in the right panel. (C) Resolution assessment of the final 3D map using Fourier shell correlation (FSC). Two criteria are shown: FSC between two half-maps reconstructed from even and odd frames (evaluated at 0.143) and FSC between the final 3D map and the fitted model (evaluated at 0.5). (D) Zoomed-in views of the final 3D density map from panel A, displayed at two contour levels. (E) Superimposition of the high-contour level map from panel D onto its fitted model.

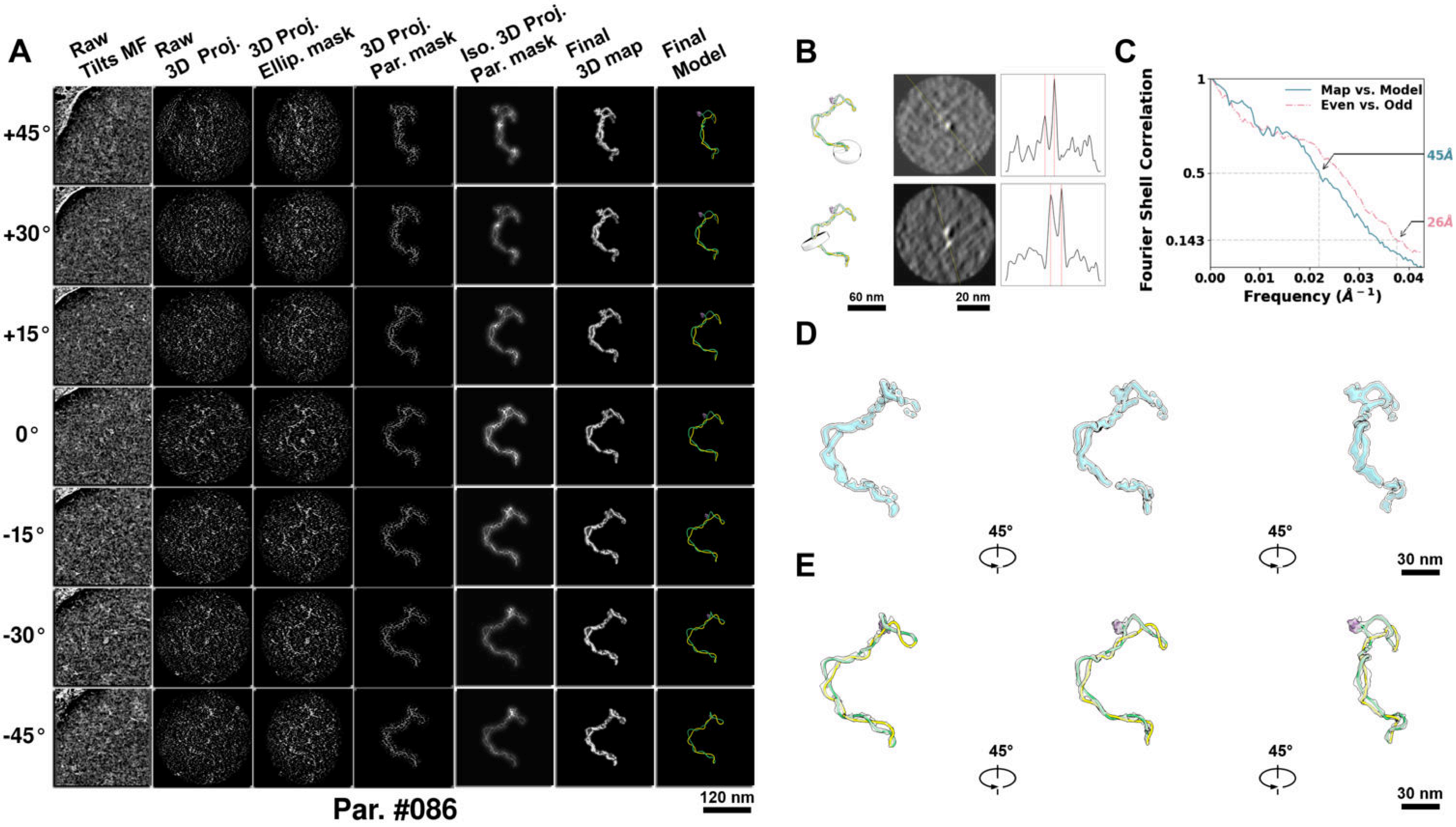

**Supplementary Particle Figure 86. Cryo-ET 3D reconstruction of an individual sTEC particle.**
(A) 3D reconstruction of the plasmid particle (index no. 86). The first column shows seven representative tilt images from +45° to -45° in step of 15°. The second, third, and fourth columns show 3D projections of the particle with spherical, ellipsoidal (thinner along the z-dimension), and particle-shaped masks, respectively. The fifth column displays the 3D projections of the enhanced and IsoNet missing-wedge-corrected particle. The sixth and seventh columns present the final 3D map and the flexibly fitted model, respectively. (B) Two cross-sectional views (12 nm thickness) of the plasmid density map along its plectoneme axis are shown in the left-middle panel. The intensity profile along the line crossing the two high-density DNA spots is displayed in the right panel. (C) Resolution assessment of the final 3D map using Fourier shell correlation (FSC). Two criteria are shown: FSC between two half-maps reconstructed from even and odd frames (evaluated at 0.143) and FSC between the final 3D map and the fitted model (evaluated at 0.5). (D) Zoomed-in views of the final 3D density map from panel A, displayed at two contour levels. (E) Superimposition of the high-contour level map from panel D onto its fitted model.

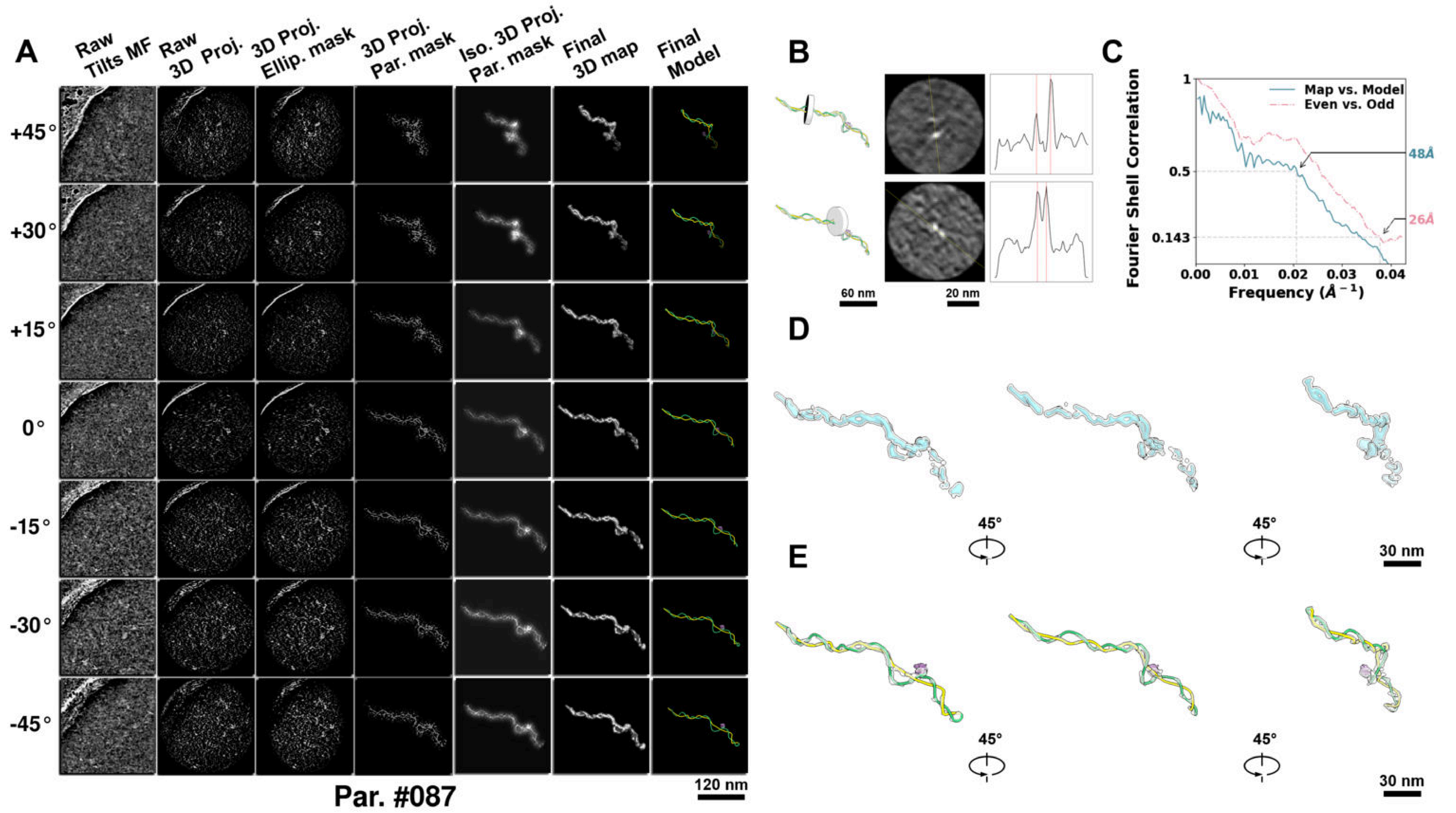

**Supplementary Particle Figure 87. Cryo-ET 3D reconstruction of an individual sTEC particle.**
(A) 3D reconstruction of the plasmid particle (index no. 87). The first column shows seven representative tilt images from +45° to -45° in step of 15°. The second, third, and fourth columns show 3D projections of the particle with spherical, ellipsoidal (thinner along the z-dimension), and particle-shaped masks, respectively. The fifth column displays the 3D projections of the enhanced and IsoNet missing-wedge-corrected particle. The sixth and seventh columns present the final 3D map and the flexibly fitted model, respectively. (B) Two cross-sectional views (12 nm thickness) of the plasmid density map along its plectoneme axis are shown in the left-middle panel. The intensity profile along the line crossing the two high-density DNA spots is displayed in the right panel. (C) Resolution assessment of the final 3D map using Fourier shell correlation (FSC). Two criteria are shown: FSC between two half-maps reconstructed from even and odd frames (evaluated at 0.143) and FSC between the final 3D map and the fitted model (evaluated at 0.5). (D) Zoomed-in views of the final 3D density map from panel A, displayed at two contour levels. (E) Superimposition of the high-contour level map from panel D onto its fitted model.

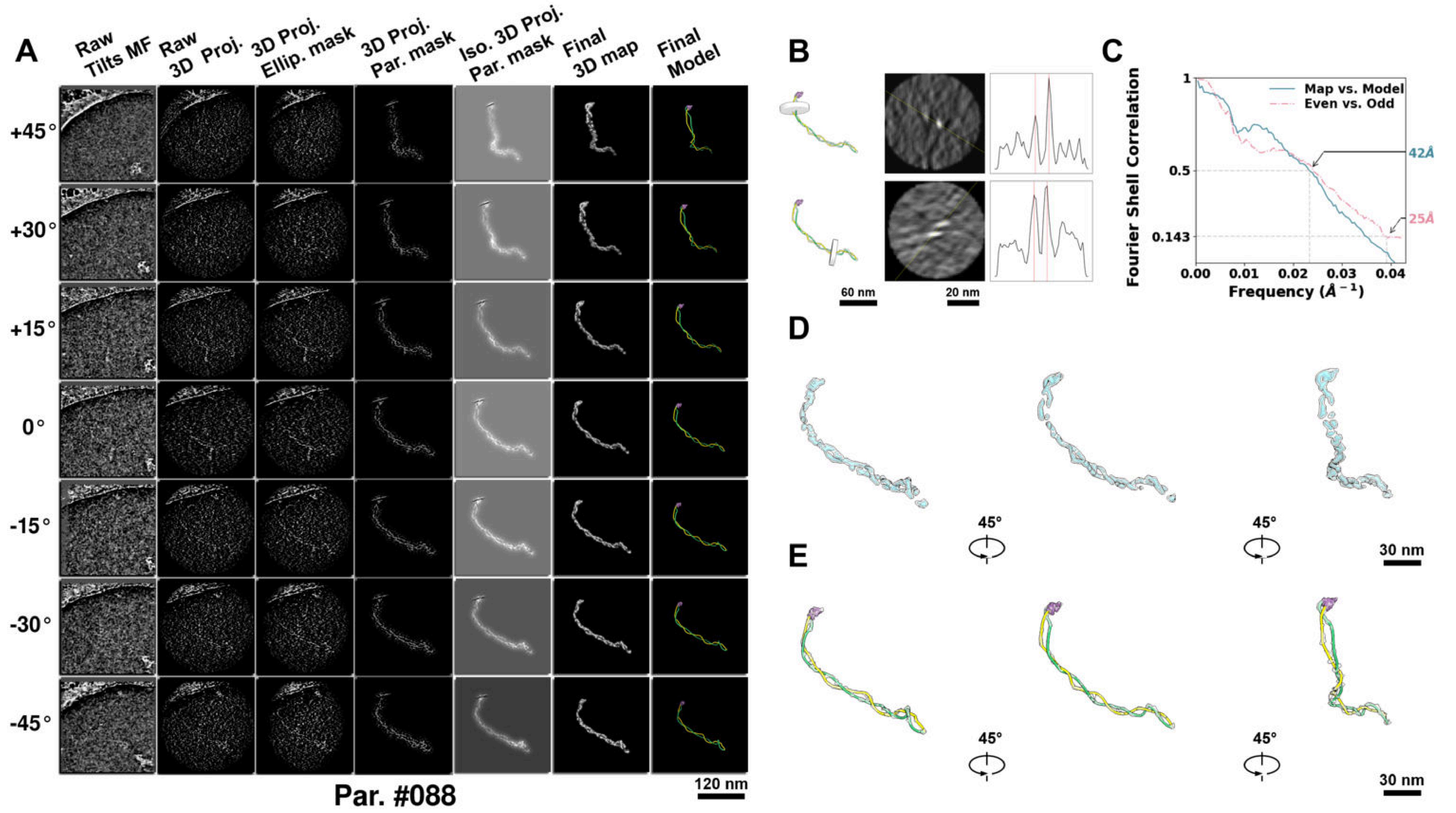

**Supplementary Particle Figure 88. Cryo-ET 3D reconstruction of an individual sTEC particle.**
(A) 3D reconstruction of the plasmid particle (index no. 88). The first column shows seven representative tilt images from +45° to -45° in step of 15°. The second, third, and fourth columns show 3D projections of the particle with spherical, ellipsoidal (thinner along the z-dimension), and particle-shaped masks, respectively. The fifth column displays the 3D projections of the enhanced and IsoNet missing-wedge-corrected particle. The sixth and seventh columns present the final 3D map and the flexibly fitted model, respectively. (B) Two cross-sectional views (12 nm thickness) of the plasmid density map along its plectoneme axis are shown in the left-middle panel. The intensity profile along the line crossing the two high-density DNA spots is displayed in the right panel. (C) Resolution assessment of the final 3D map using Fourier shell correlation (FSC). Two criteria are shown: FSC between two half-maps reconstructed from even and odd frames (evaluated at 0.143) and FSC between the final 3D map and the fitted model (evaluated at 0.5). (D) Zoomed-in views of the final 3D density map from panel A, displayed at two contour levels. (E) Superimposition of the high-contour level map from panel D onto its fitted model.

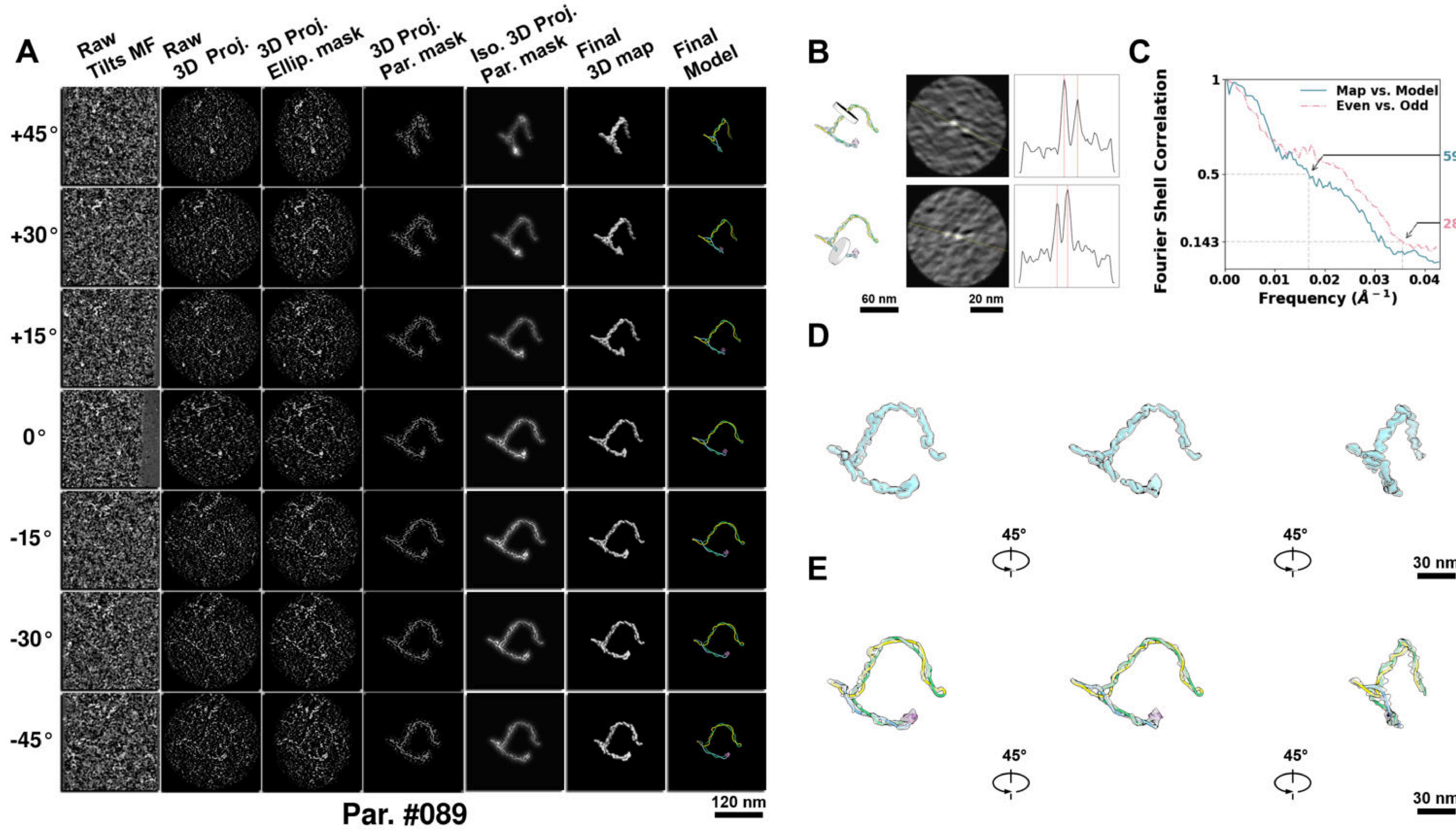

**Supplementary Particle Figure 89. Cryo-ET 3D reconstruction of an individual sTEC particle.**
(A) 3D reconstruction of the plasmid particle (index no. 89). The first column shows seven representative tilt images from +45° to -45° in step of 15°. The second, third, and fourth columns show 3D projections of the particle with spherical, ellipsoidal (thinner along the z-dimension), and particle-shaped masks, respectively. The fifth column displays the 3D projections of the enhanced and IsoNet missing-wedge-corrected particle. The sixth and seventh columns present the final 3D map and the flexibly fitted model, respectively. (B) Two cross-sectional views (12 nm thickness) of the plasmid density map along its plectoneme axis are shown in the left-middle panel. The intensity profile along the line crossing the two high-density DNA spots is displayed in the right panel. (C) Resolution assessment of the final 3D map using Fourier shell correlation (FSC). Two criteria are shown: FSC between two half-maps reconstructed from even and odd frames (evaluated at 0.143) and FSC between the final 3D map and the fitted model (evaluated at 0.5). (D) Zoomed-in views of the final 3D density map from panel A, displayed at two contour levels. (E) Superimposition of the high-contour level map from panel D onto its fitted model.

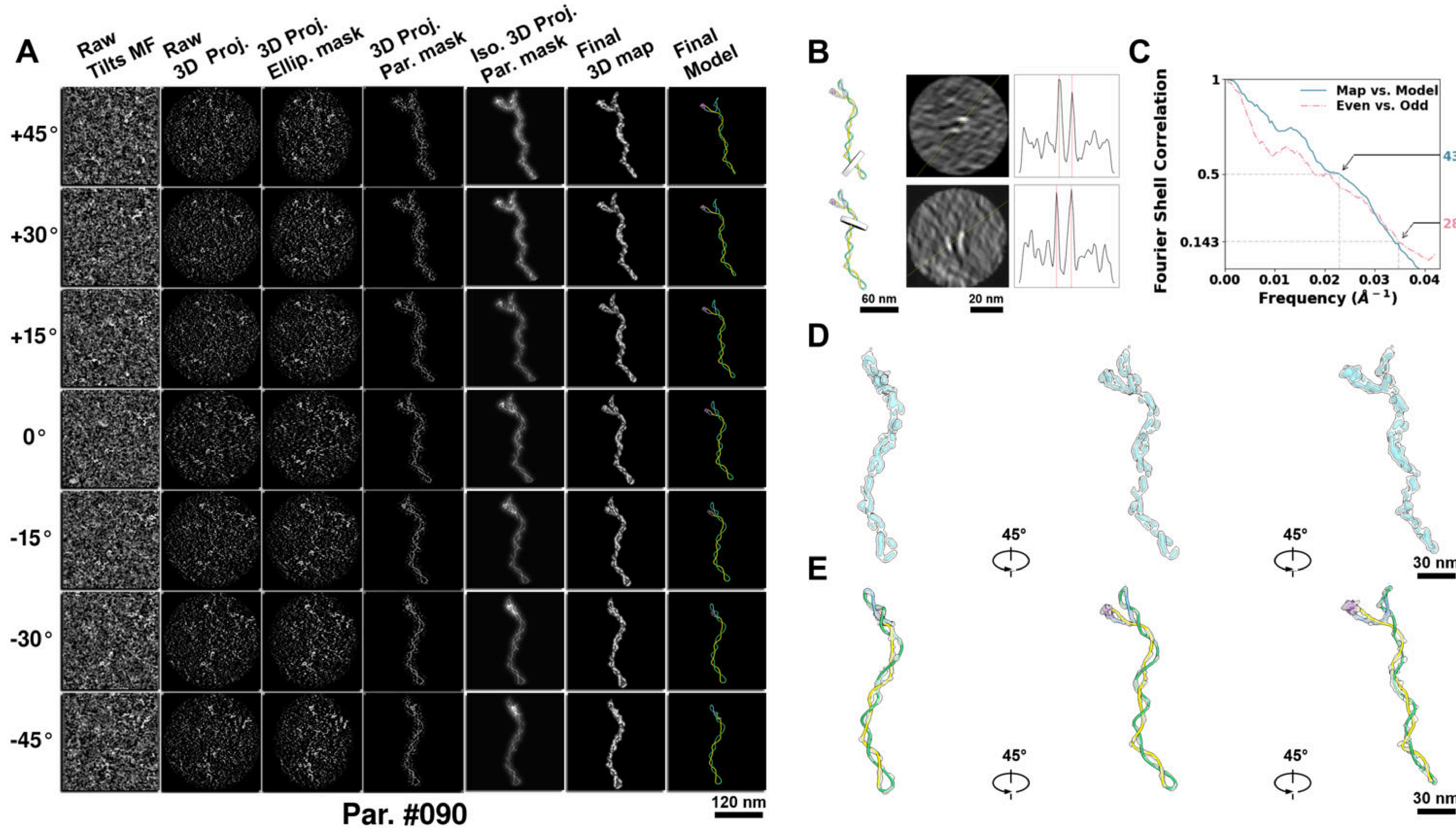

**Supplementary Particle Figure 90. Cryo-ET 3D reconstruction of an individual sTEC particle.**
(A) 3D reconstruction of the plasmid particle (index no. 90). The first column shows seven representative tilt images from +45° to -45° in step of 15°. The second, third, and fourth columns show 3D projections of the particle with spherical, ellipsoidal (thinner along the z-dimension), and particle-shaped masks, respectively. The fifth column displays the 3D projections of the enhanced and IsoNet missing-wedge-corrected particle. The sixth and seventh columns present the final 3D map and the flexibly fitted model, respectively. (B) Two cross-sectional views (12 nm thickness) of the plasmid density map along its plectoneme axis are shown in the left-middle panel. The intensity profile along the line crossing the two high-density DNA spots is displayed in the right panel. (C) Resolution assessment of the final 3D map using Fourier shell correlation (FSC). Two criteria are shown: FSC between two half-maps reconstructed from even and odd frames (evaluated at 0.143) and FSC between the final 3D map and the fitted model (evaluated at 0.5). (D) Zoomed-in views of the final 3D density map from panel A, displayed at two contour levels. (E) Superimposition of the high-contour level map from panel D onto its fitted model.

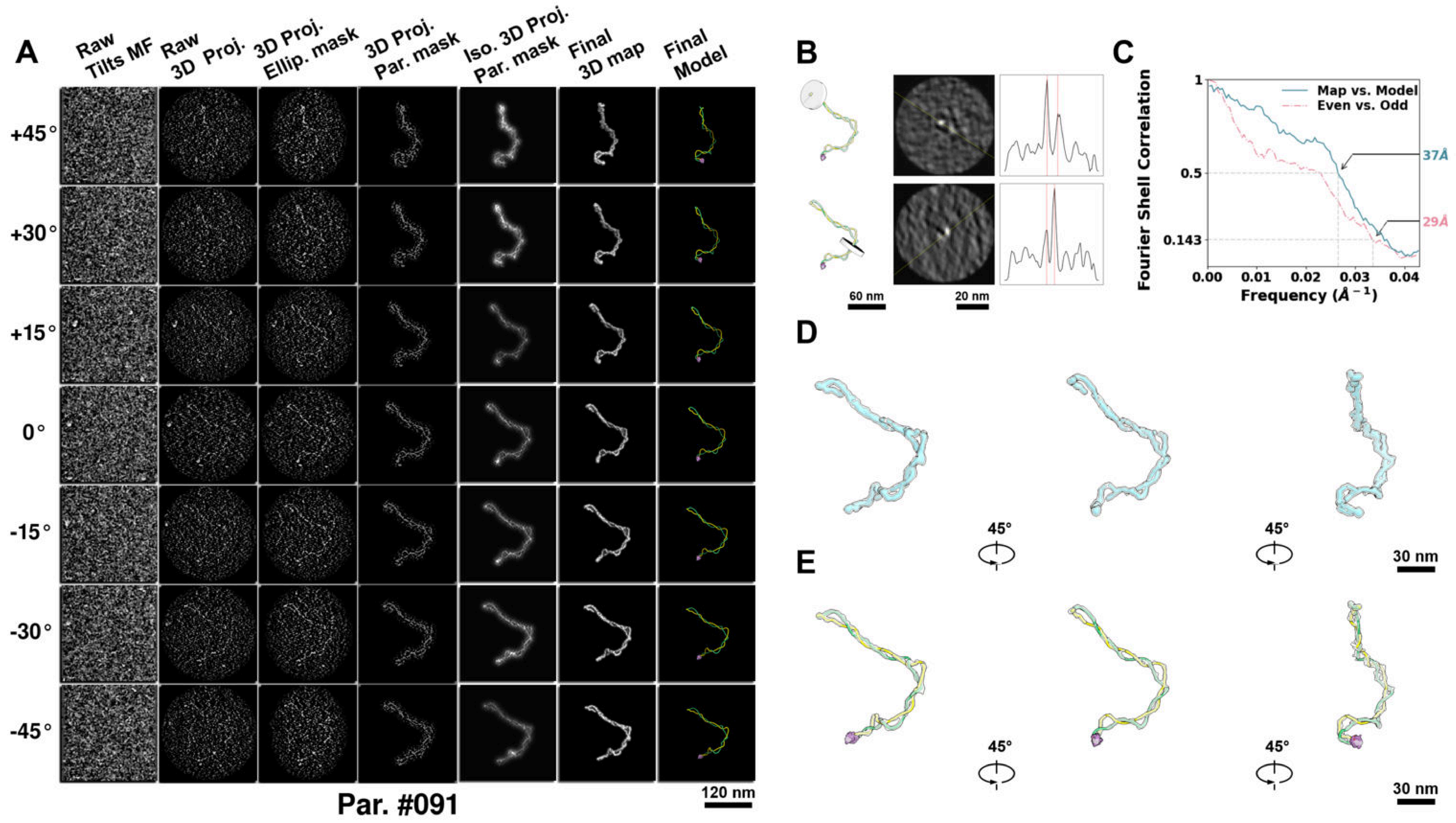

**Supplementary Particle Figure 91. Cryo-ET 3D reconstruction of an individual sTEC particle.**
(A) 3D reconstruction of the plasmid particle (index no. 91). The first column shows seven representative tilt images from +45° to -45° in step of 15°. The second, third, and fourth columns show 3D projections of the particle with spherical, ellipsoidal (thinner along the z-dimension), and particle-shaped masks, respectively. The fifth column displays the 3D projections of the enhanced and IsoNet missing-wedge-corrected particle. The sixth and seventh columns present the final 3D map and the flexibly fitted model, respectively. (B) Two cross-sectional views (12 nm thickness) of the plasmid density map along its plectoneme axis are shown in the left-middle panel. The intensity profile along the line crossing the two high-density DNA spots is displayed in the right panel. (C) Resolution assessment of the final 3D map using Fourier shell correlation (FSC). Two criteria are shown: FSC between two half-maps reconstructed from even and odd frames (evaluated at 0.143) and FSC between the final 3D map and the fitted model (evaluated at 0.5). (D) Zoomed-in views of the final 3D density map from panel A, displayed at two contour levels. (E) Superimposition of the high-contour level map from panel D onto its fitted model.

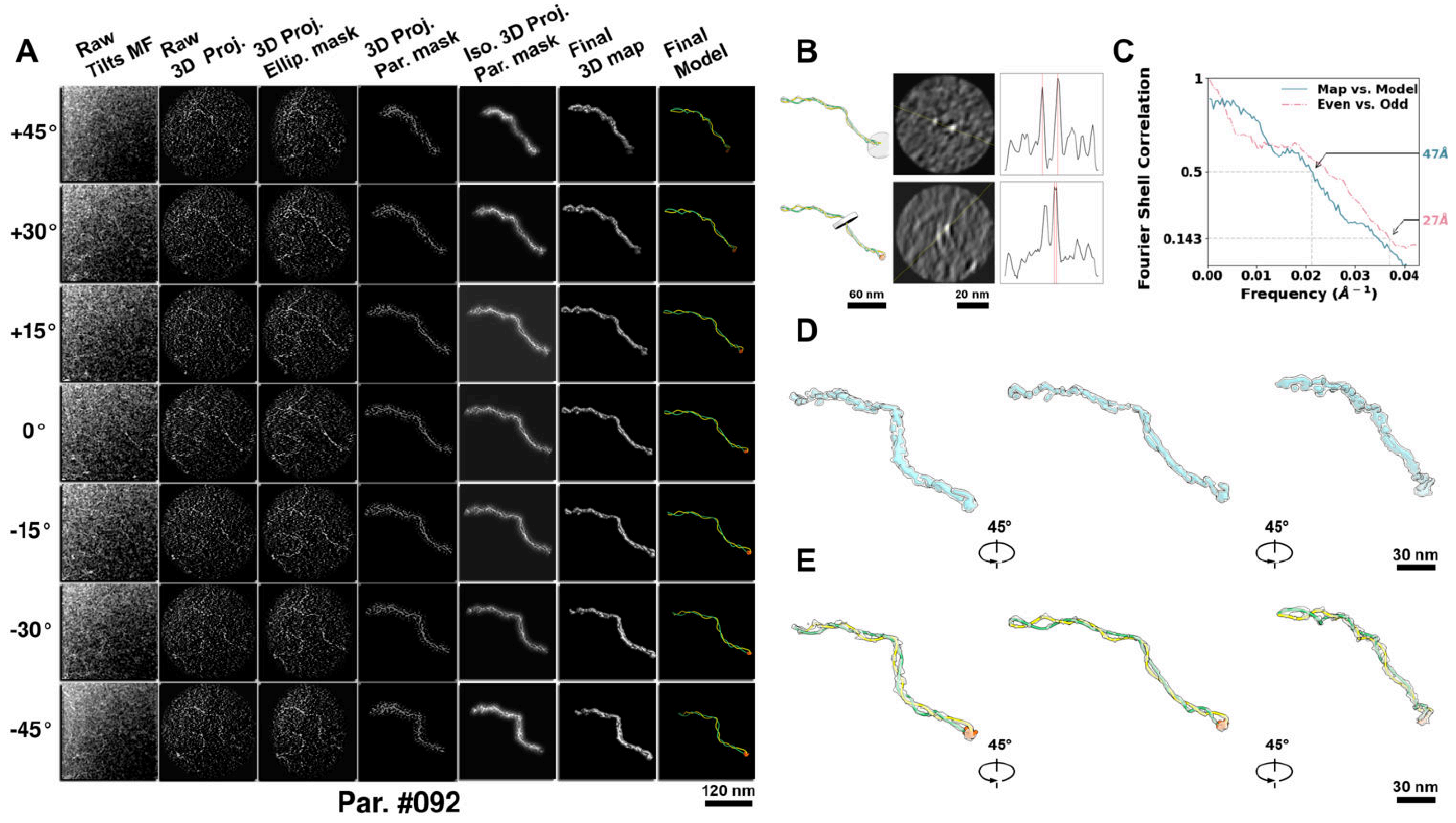

**Supplementary Particle Figure 92. Cryo-ET 3D reconstruction of an individual P.Cas particle.**
(A) 3D reconstruction of the plasmid particle (index no. 92). The first column shows seven representative tilt images from +45° to -45° in step of 15°. The second, third, and fourth columns show 3D projections of the particle with spherical, ellipsoidal (thinner along the z-dimension), and particle-shaped masks, respectively. The fifth column displays the 3D projections of the enhanced and IsoNet missing-wedge-corrected particle. The sixth and seventh columns present the final 3D map and the flexibly fitted model, respectively. (B) Two cross-sectional views (12 nm thickness) of the plasmid density map along its plectoneme axis are shown in the left-middle panel. The intensity profile along the line crossing the two high-density DNA spots is displayed in the right panel. (C) Resolution assessment of the final 3D map using Fourier shell correlation (FSC). Two criteria are shown: FSC between two half-maps reconstructed from even and odd frames (evaluated at 0.143) and FSC between the final 3D map and the fitted model (evaluated at 0.5). (D) Zoomed-in views of the final 3D density map from panel A, displayed at two contour levels. (E) Superimposition of the high-contour level map from panel D onto its fitted model.

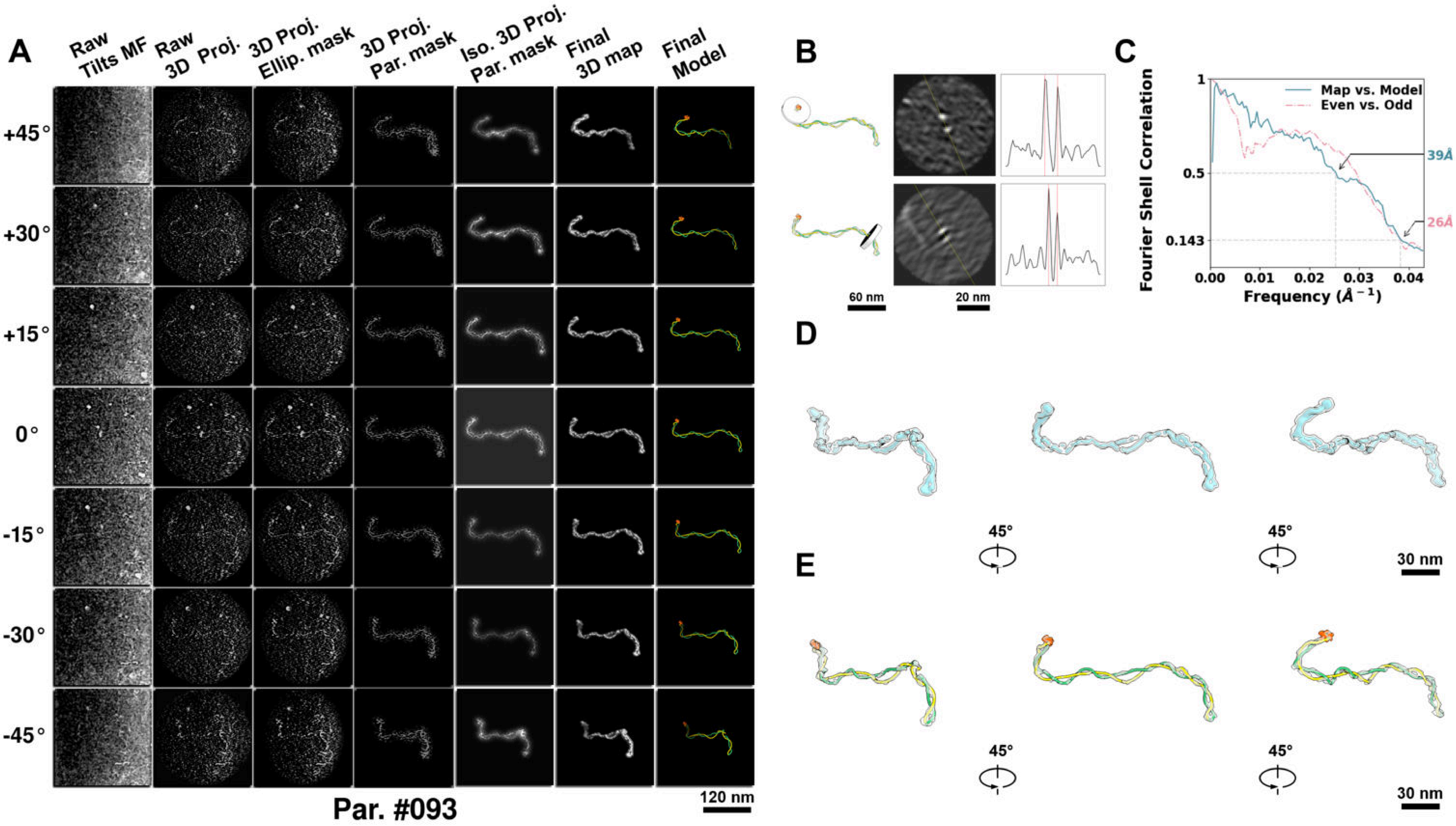

**Supplementary Particle Figure 93. Cryo-ET 3D reconstruction of an individual P.Cas particle.**
(A) 3D reconstruction of the plasmid particle (index no. 93). The first column shows seven representative tilt images from +45° to -45° in step of 15°. The second, third, and fourth columns show 3D projections of the particle with spherical, ellipsoidal (thinner along the z-dimension), and particle-shaped masks, respectively. The fifth column displays the 3D projections of the enhanced and IsoNet missing-wedge-corrected particle. The sixth and seventh columns present the final 3D map and the flexibly fitted model, respectively. (B) Two cross-sectional views (12 nm thickness) of the plasmid density map along its plectoneme axis are shown in the left-middle panel. The intensity profile along the line crossing the two high-density DNA spots is displayed in the right panel. (C) Resolution assessment of the final 3D map using Fourier shell correlation (FSC). Two criteria are shown: FSC between two half-maps reconstructed from even and odd frames (evaluated at 0.143) and FSC between the final 3D map and the fitted model (evaluated at 0.5). (D) Zoomed-in views of the final 3D density map from panel A, displayed at two contour levels. (E) Superimposition of the high-contour level map from panel D onto its fitted model.

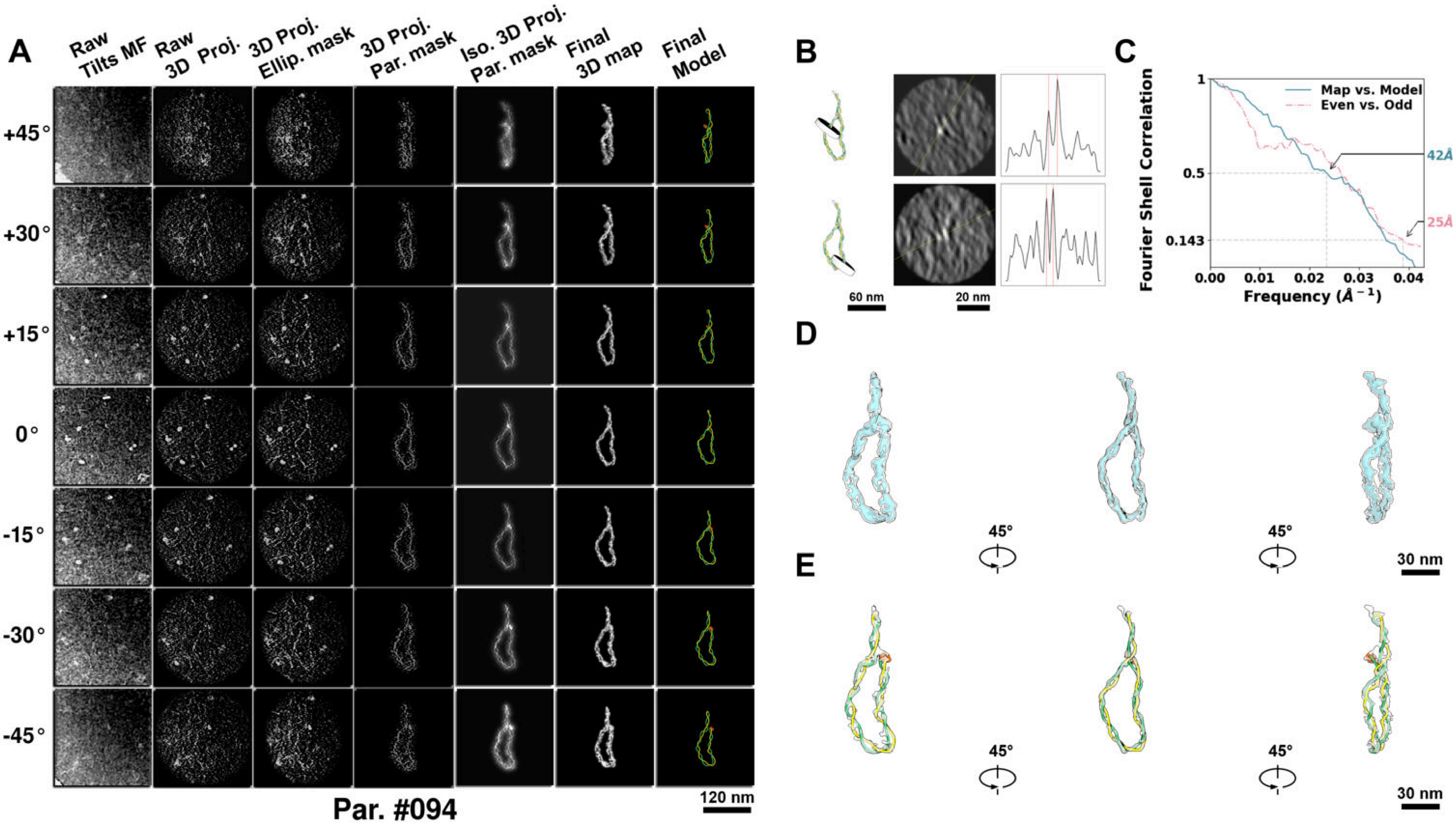

**Supplementary Particle Figure 94. Cryo-ET 3D reconstruction of an individual P.Cas particle.**
(A) 3D reconstruction of the plasmid particle (index no. 94). The first column shows seven representative tilt images from +45° to -45° in step of 15°. The second, third, and fourth columns show 3D projections of the particle with spherical, ellipsoidal (thinner along the z-dimension), and particle-shaped masks, respectively. The fifth column displays the 3D projections of the enhanced and IsoNet missing-wedge-corrected particle. The sixth and seventh columns present the final 3D map and the flexibly fitted model, respectively. (B) Two cross-sectional views (12 nm thickness) of the plasmid density map along its plectoneme axis are shown in the left-middle panel. The intensity profile along the line crossing the two high-density DNA spots is displayed in the right panel. (C) Resolution assessment of the final 3D map using Fourier shell correlation (FSC). Two criteria are shown: FSC between two half-maps reconstructed from even and odd frames (evaluated at 0.143) and FSC between the final 3D map and the fitted model (evaluated at 0.5). (D) Zoomed-in views of the final 3D density map from panel A, displayed at two contour levels. (E) Superimposition of the high-contour level map from panel D onto its fitted model.

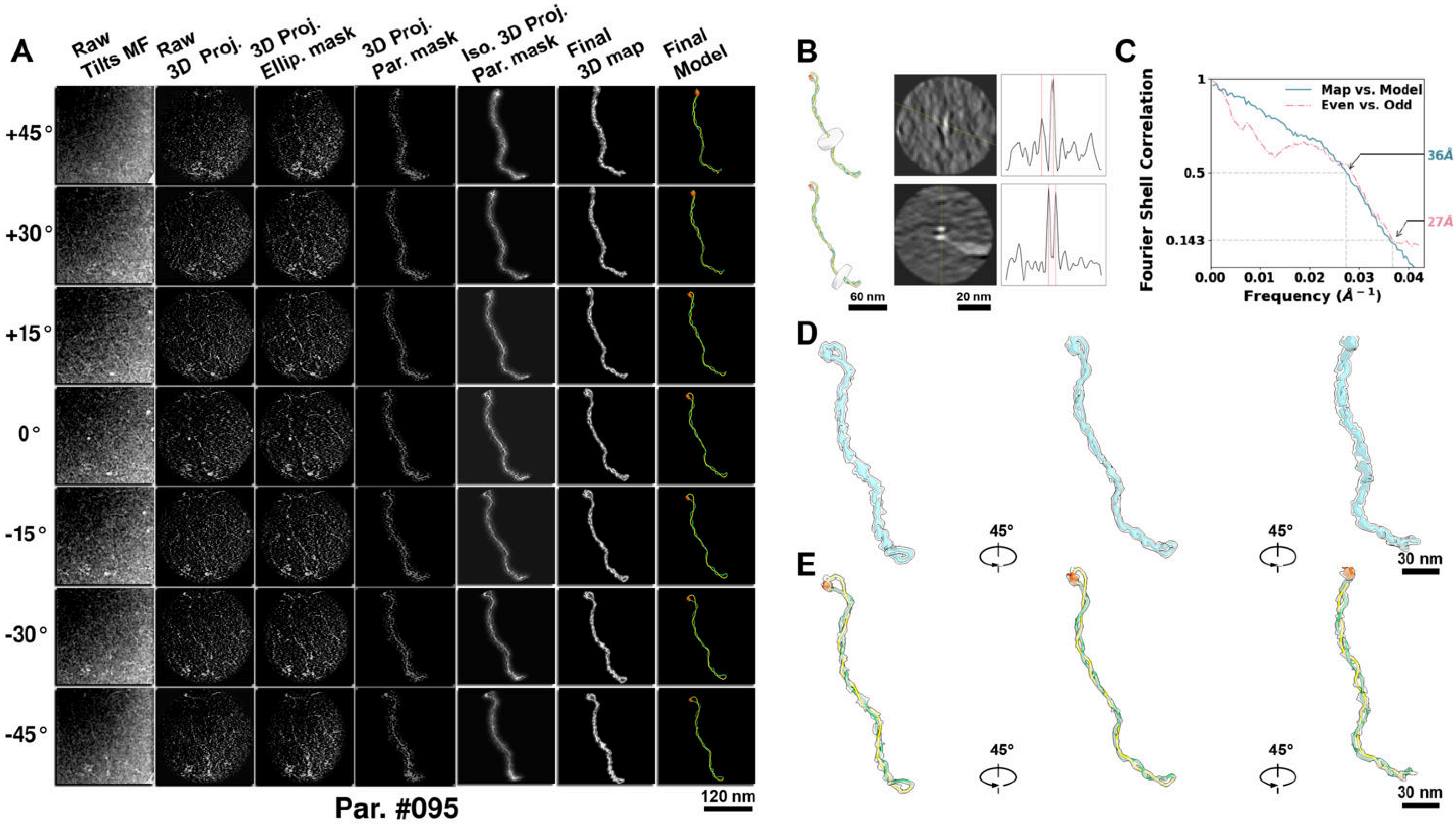

**Supplementary Particle Figure 95. Cryo-ET 3D reconstruction of an individual P.Cas particle.**
(A) 3D reconstruction of the plasmid particle (index no. 95). The first column shows seven representative tilt images from +45° to -45° in step of 15°. The second, third, and fourth columns show 3D projections of the particle with spherical, ellipsoidal (thinner along the z-dimension), and particle-shaped masks, respectively. The fifth column displays the 3D projections of the enhanced and IsoNet missing-wedge-corrected particle. The sixth and seventh columns present the final 3D map and the flexibly fitted model, respectively. (B) Two cross-sectional views (12 nm thickness) of the plasmid density map along its plectoneme axis are shown in the left-middle panel. The intensity profile along the line crossing the two high-density DNA spots is displayed in the right panel. (C) Resolution assessment of the final 3D map using Fourier shell correlation (FSC). Two criteria are shown: FSC between two half-maps reconstructed from even and odd frames (evaluated at 0.143) and FSC between the final 3D map and the fitted model (evaluated at 0.5). (D) Zoomed-in views of the final 3D density map from panel A, displayed at two contour levels. (E) Superimposition of the high-contour level map from panel D onto its fitted model.

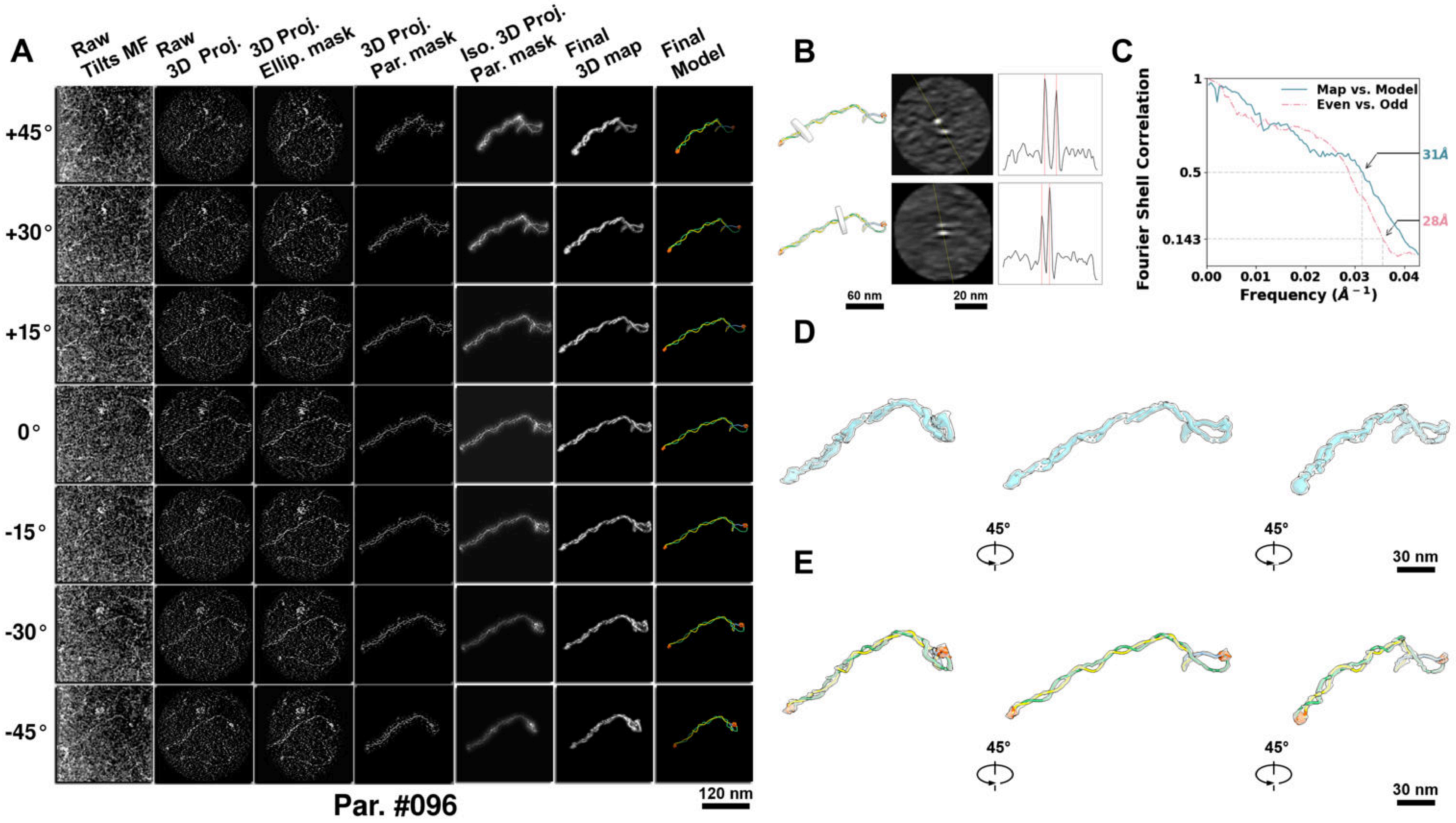

**Supplementary Particle Figure 96. Cryo-ET 3D reconstruction of an individual P.Cas particle.**
(A) 3D reconstruction of the plasmid particle (index no. 96). The first column shows seven representative tilt images from +45° to -45° in step of 15°. The second, third, and fourth columns show 3D projections of the particle with spherical, ellipsoidal (thinner along the z-dimension), and particle-shaped masks, respectively. The fifth column displays the 3D projections of the enhanced and IsoNet missing-wedge-corrected particle. The sixth and seventh columns present the final 3D map and the flexibly fitted model, respectively. (B) Two cross-sectional views (12 nm thickness) of the plasmid density map along its plectoneme axis are shown in the left-middle panel. The intensity profile along the line crossing the two high-density DNA spots is displayed in the right panel. (C) Resolution assessment of the final 3D map using Fourier shell correlation (FSC). Two criteria are shown: FSC between two half-maps reconstructed from even and odd frames (evaluated at 0.143) and FSC between the final 3D map and the fitted model (evaluated at 0.5). (D) Zoomed-in views of the final 3D density map from panel A, displayed at two contour levels. (E) Superimposition of the high-contour level map from panel D onto its fitted model.

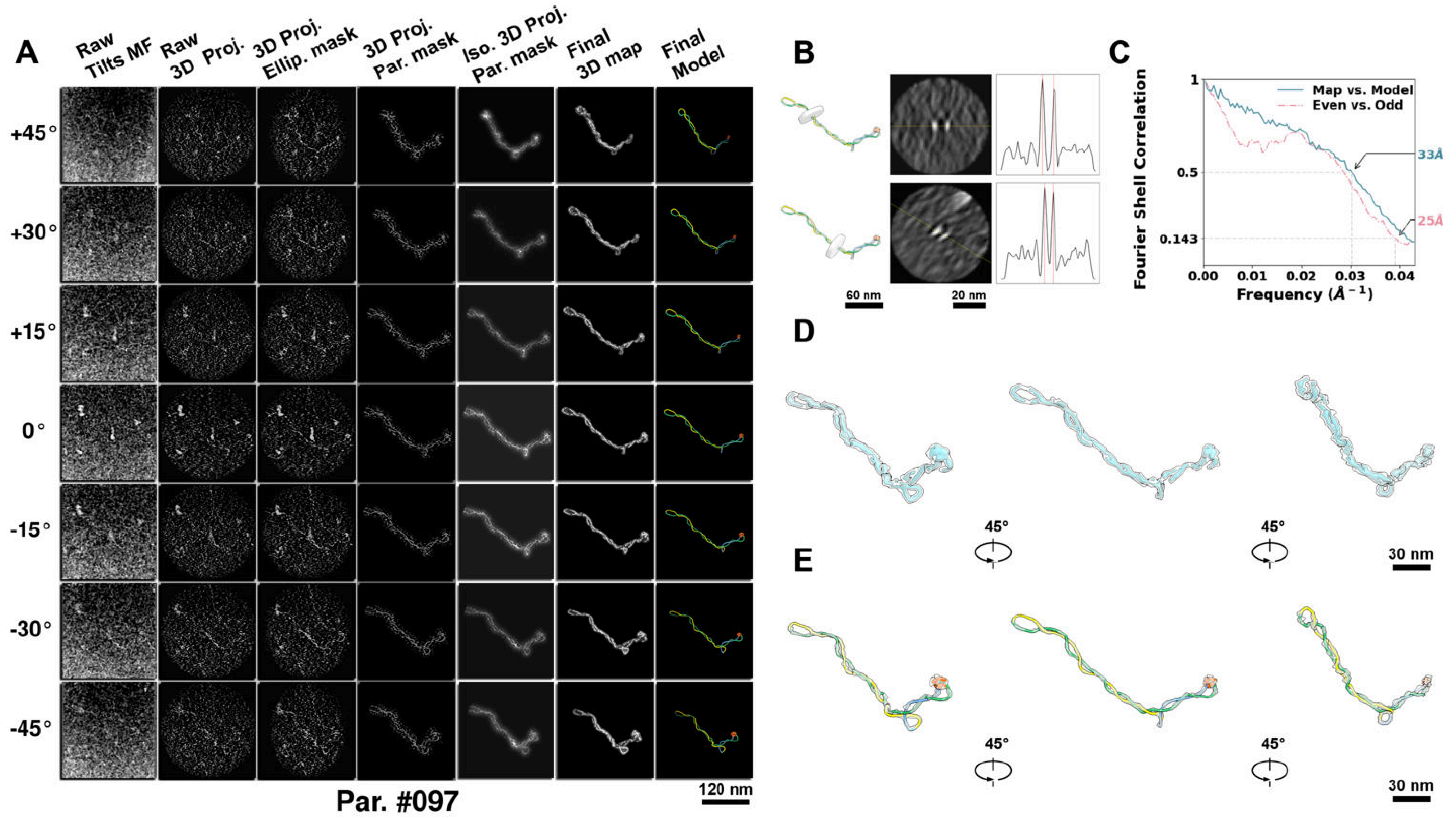

**Supplementary Particle Figure 97. Cryo-ET 3D reconstruction of an individual P.Cas particle.**
(A) 3D reconstruction of the plasmid particle (index no. 97). The first column shows seven representative tilt images from +45° to -45° in step of 15°. The second, third, and fourth columns show 3D projections of the particle with spherical, ellipsoidal (thinner along the z-dimension), and particle-shaped masks, respectively. The fifth column displays the 3D projections of the enhanced and IsoNet missing-wedge-corrected particle. The sixth and seventh columns present the final 3D map and the flexibly fitted model, respectively. (B) Two cross-sectional views (12 nm thickness) of the plasmid density map along its plectoneme axis are shown in the left-middle panel. The intensity profile along the line crossing the two high-density DNA spots is displayed in the right panel. (C) Resolution assessment of the final 3D map using Fourier shell correlation (FSC). Two criteria are shown: FSC between two half-maps reconstructed from even and odd frames (evaluated at 0.143) and FSC between the final 3D map and the fitted model (evaluated at 0.5). (D) Zoomed-in views of the final 3D density map from panel A, displayed at two contour levels. (E) Superimposition of the high-contour level map from panel D onto its fitted model.

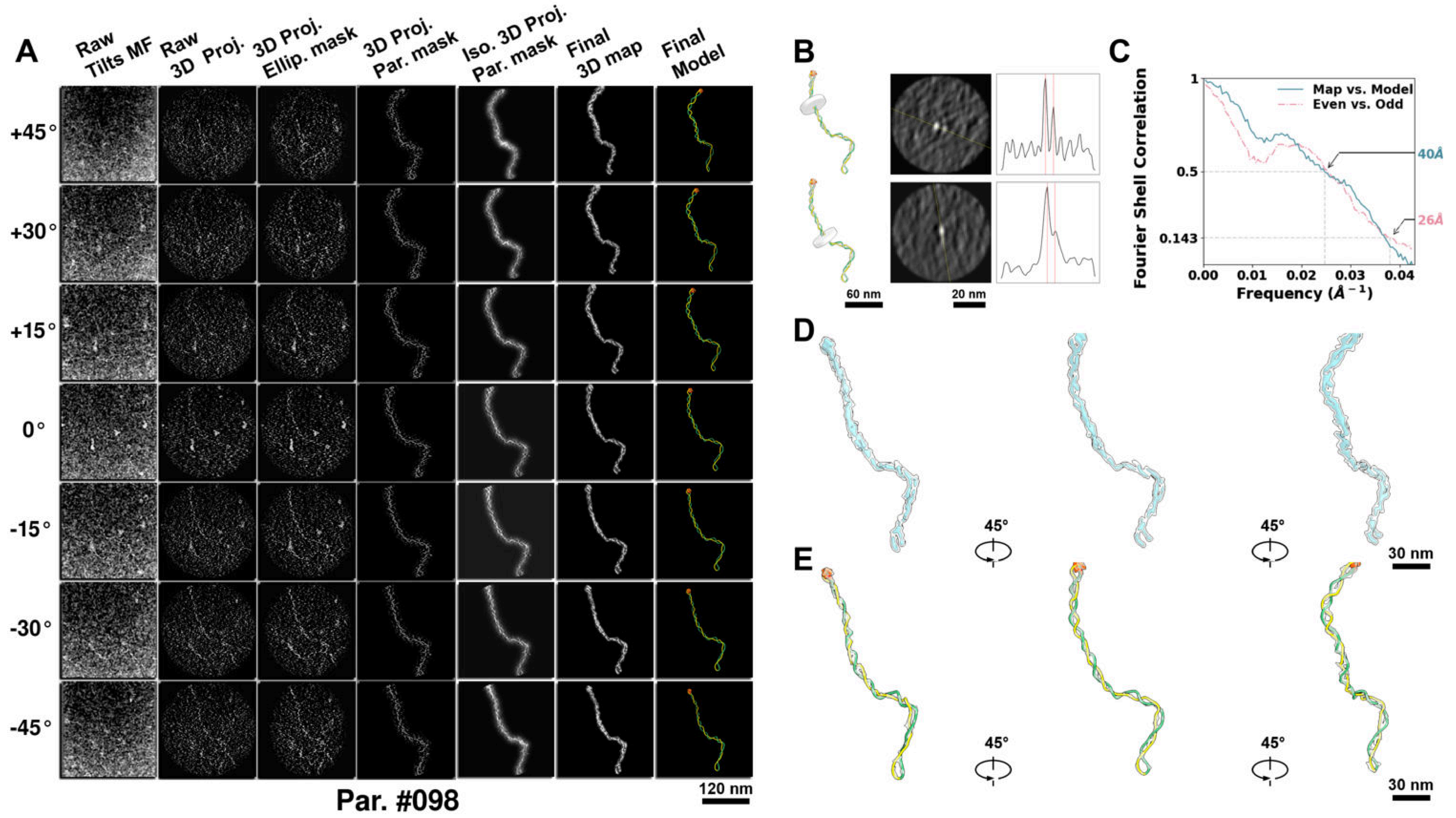

**Supplementary Particle Figure 98. Cryo-ET 3D reconstruction of an individual P.Cas particle.**
(A) 3D reconstruction of the plasmid particle (index no. 98). The first column shows seven representative tilt images from +45° to -45° in step of 15°. The second, third, and fourth columns show 3D projections of the particle with spherical, ellipsoidal (thinner along the z-dimension), and particle-shaped masks, respectively. The fifth column displays the 3D projections of the enhanced and IsoNet missing-wedge-corrected particle. The sixth and seventh columns present the final 3D map and the flexibly fitted model, respectively. (B) Two cross-sectional views (12 nm thickness) of the plasmid density map along its plectoneme axis are shown in the left-middle panel. The intensity profile along the line crossing the two high-density DNA spots is displayed in the right panel. (C) Resolution assessment of the final 3D map using Fourier shell correlation (FSC). Two criteria are shown: FSC between two half-maps reconstructed from even and odd frames (evaluated at 0.143) and FSC between the final 3D map and the fitted model (evaluated at 0.5). (D) Zoomed-in views of the final 3D density map from panel A, displayed at two contour levels. (E) Superimposition of the high-contour level map from panel D onto its fitted model.

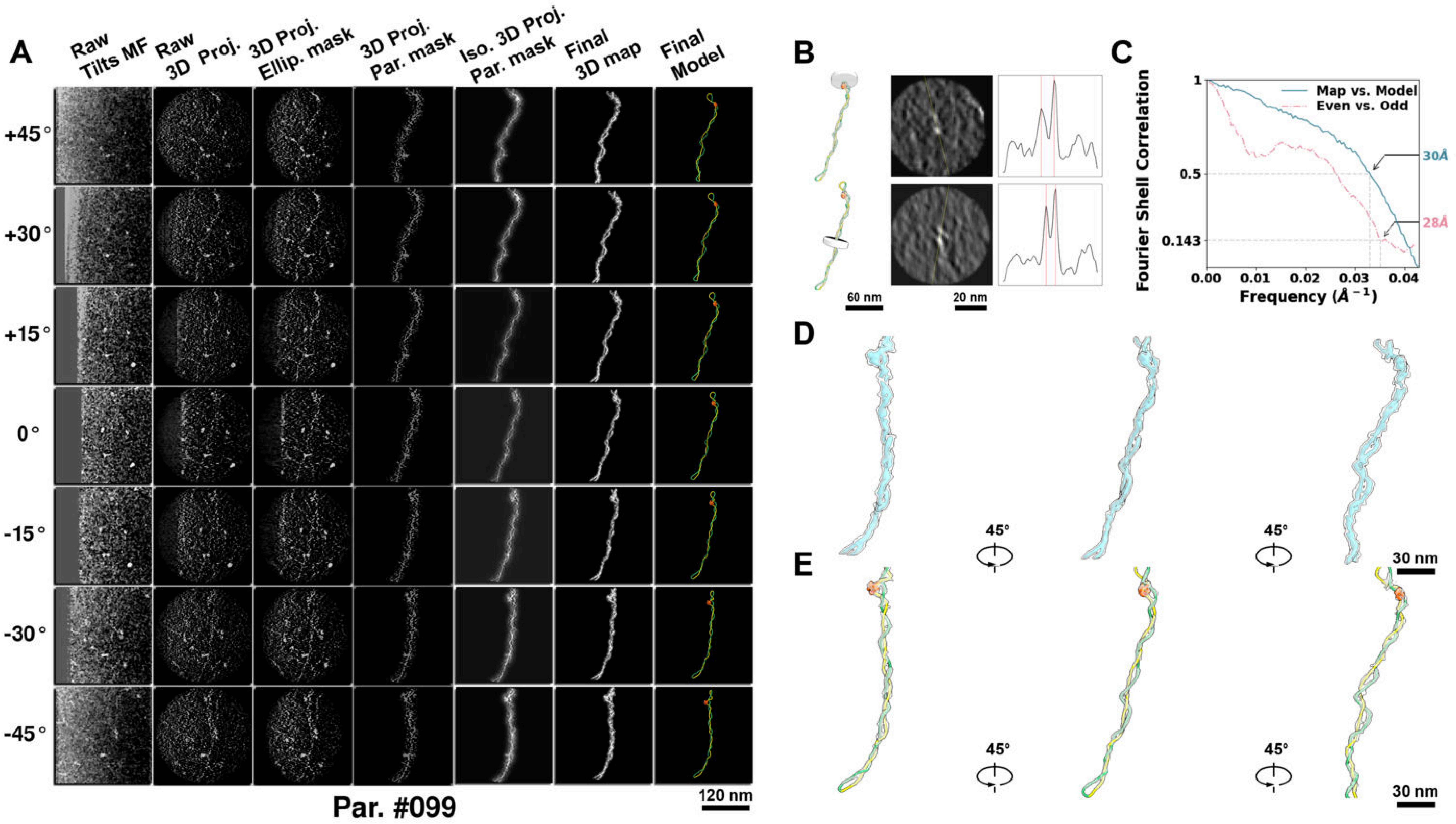

**Supplementary Particle Figure 99. Cryo-ET 3D reconstruction of an individual P.Cas particle.**
(A) 3D reconstruction of the plasmid particle (index no. 99). The first column shows seven representative tilt images from +45° to -45° in step of 15°. The second, third, and fourth columns show 3D projections of the particle with spherical, ellipsoidal (thinner along the z-dimension), and particle-shaped masks, respectively. The fifth column displays the 3D projections of the enhanced and IsoNet missing-wedge-corrected particle. The sixth and seventh columns present the final 3D map and the flexibly fitted model, respectively. (B) Two cross-sectional views (12 nm thickness) of the plasmid density map along its plectoneme axis are shown in the left-middle panel. The intensity profile along the line crossing the two high-density DNA spots is displayed in the right panel. (C) Resolution assessment of the final 3D map using Fourier shell correlation (FSC). Two criteria are shown: FSC between two half-maps reconstructed from even and odd frames (evaluated at 0.143) and FSC between the final 3D map and the fitted model (evaluated at 0.5). (D) Zoomed-in views of the final 3D density map from panel A, displayed at two contour levels. (E) Superimposition of the high-contour level map from panel D onto its fitted model.

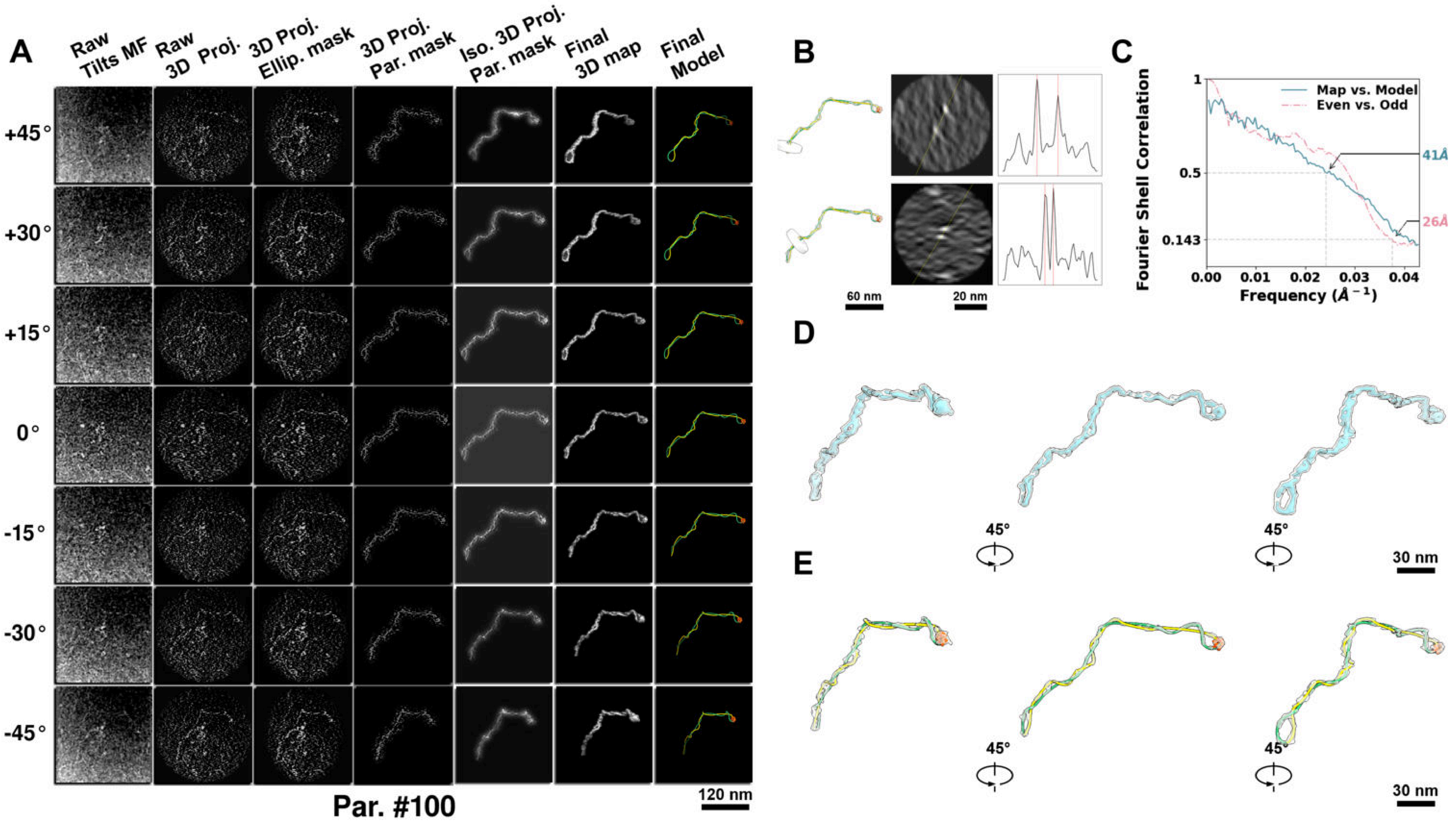

**Supplementary Particle Figure 100. Cryo-ET 3D reconstruction of an individual P.Cas particle.**
(A) 3D reconstruction of the plasmid particle (index no. 100). The first column shows seven representative tilt images from +45° to -45° in step of 15°. The second, third, and fourth columns show 3D projections of the particle with spherical, ellipsoidal (thinner along the z-dimension), and particle-shaped masks, respectively. The fifth column displays the 3D projections of the enhanced and IsoNet missing-wedge-corrected particle. The sixth and seventh columns present the final 3D map and the flexibly fitted model, respectively. (B) Two cross-sectional views (12 nm thickness) of the plasmid density map along its plectoneme axis are shown in the left-middle panel. The intensity profile along the line crossing the two high-density DNA spots is displayed in the right panel. (C) Resolution assessment of the final 3D map using Fourier shell correlation (FSC). Two criteria are shown: FSC between two half-maps reconstructed from even and odd frames (evaluated at 0.143) and FSC between the final 3D map and the fitted model (evaluated at 0.5). (D) Zoomed-in views of the final 3D density map from panel A, displayed at two contour levels. (E) Superimposition of the high-contour level map from panel D onto its fitted model.

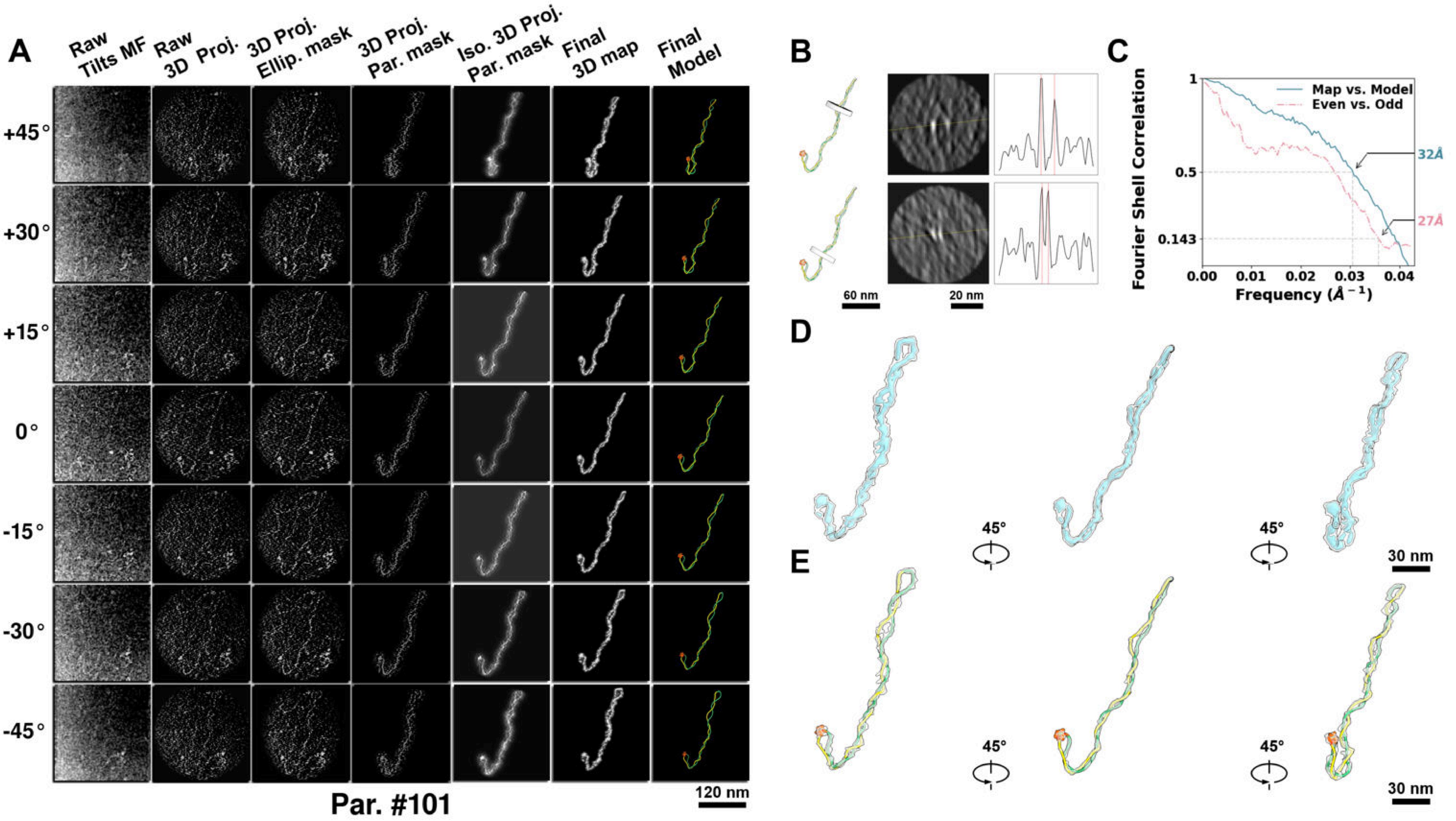

**Supplementary Particle Figure 101. Cryo-ET 3D reconstruction of an individual P.Cas particle.**
(A) 3D reconstruction of the plasmid particle (index no. 101). The first column shows seven representative tilt images from +45° to -45° in step of 15°. The second, third, and fourth columns show 3D projections of the particle with spherical, ellipsoidal (thinner along the z-dimension), and particle-shaped masks, respectively. The fifth column displays the 3D projections of the enhanced and IsoNet missing-wedge-corrected particle. The sixth and seventh columns present the final 3D map and the flexibly fitted model, respectively. (B) Two cross-sectional views (12 nm thickness) of the plasmid density map along its plectoneme axis are shown in the left-middle panel. The intensity profile along the line crossing the two high-density DNA spots is displayed in the right panel. (C) Resolution assessment of the final 3D map using Fourier shell correlation (FSC). Two criteria are shown: FSC between two half-maps reconstructed from even and odd frames (evaluated at 0.143) and FSC between the final 3D map and the fitted model (evaluated at 0.5). (D) Zoomed-in views of the final 3D density map from panel A, displayed at two contour levels. (E) Superimposition of the high-contour level map from panel D onto its fitted model.

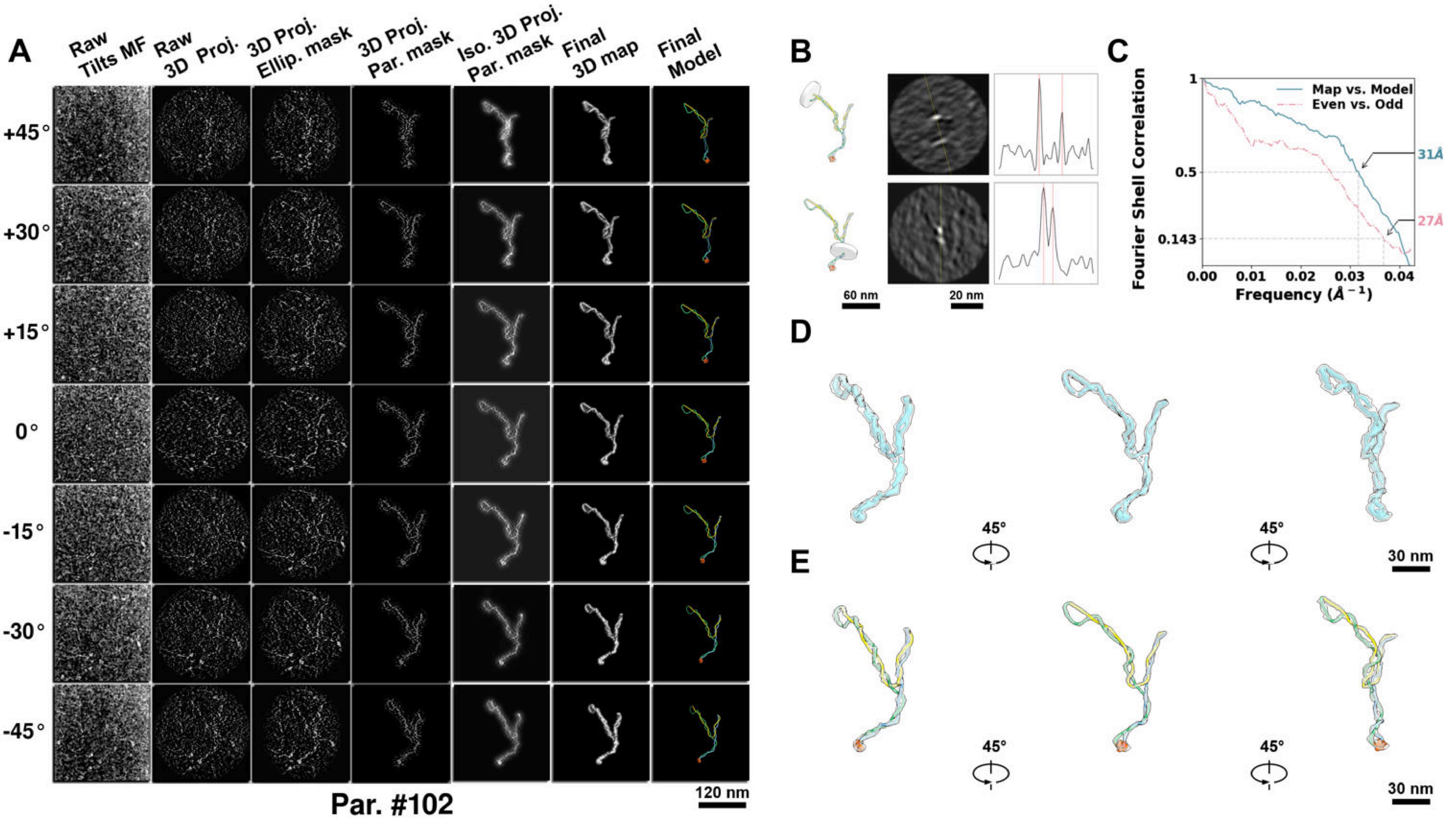

**Supplementary Particle Figure 102. Cryo-ET 3D reconstruction of an individual P.Cas particle.**
(A) 3D reconstruction of the plasmid particle (index no. 102). The first column shows seven representative tilt images from +45° to -45° in step of 15°. The second, third, and fourth columns show 3D projections of the particle with spherical, ellipsoidal (thinner along the z-dimension), and particle-shaped masks, respectively. The fifth column displays the 3D projections of the enhanced and IsoNet missing-wedge-corrected particle. The sixth and seventh columns present the final 3D map and the flexibly fitted model, respectively. (B) Two cross-sectional views (12 nm thickness) of the plasmid density map along its plectoneme axis are shown in the left-middle panel. The intensity profile along the line crossing the two high-density DNA spots is displayed in the right panel. (C) Resolution assessment of the final 3D map using Fourier shell correlation (FSC). Two criteria are shown: FSC between two half-maps reconstructed from even and odd frames (evaluated at 0.143) and FSC between the final 3D map and the fitted model (evaluated at 0.5). (D) Zoomed-in views of the final 3D density map from panel A, displayed at two contour levels. (E) Superimposition of the high-contour level map from panel D onto its fitted model.

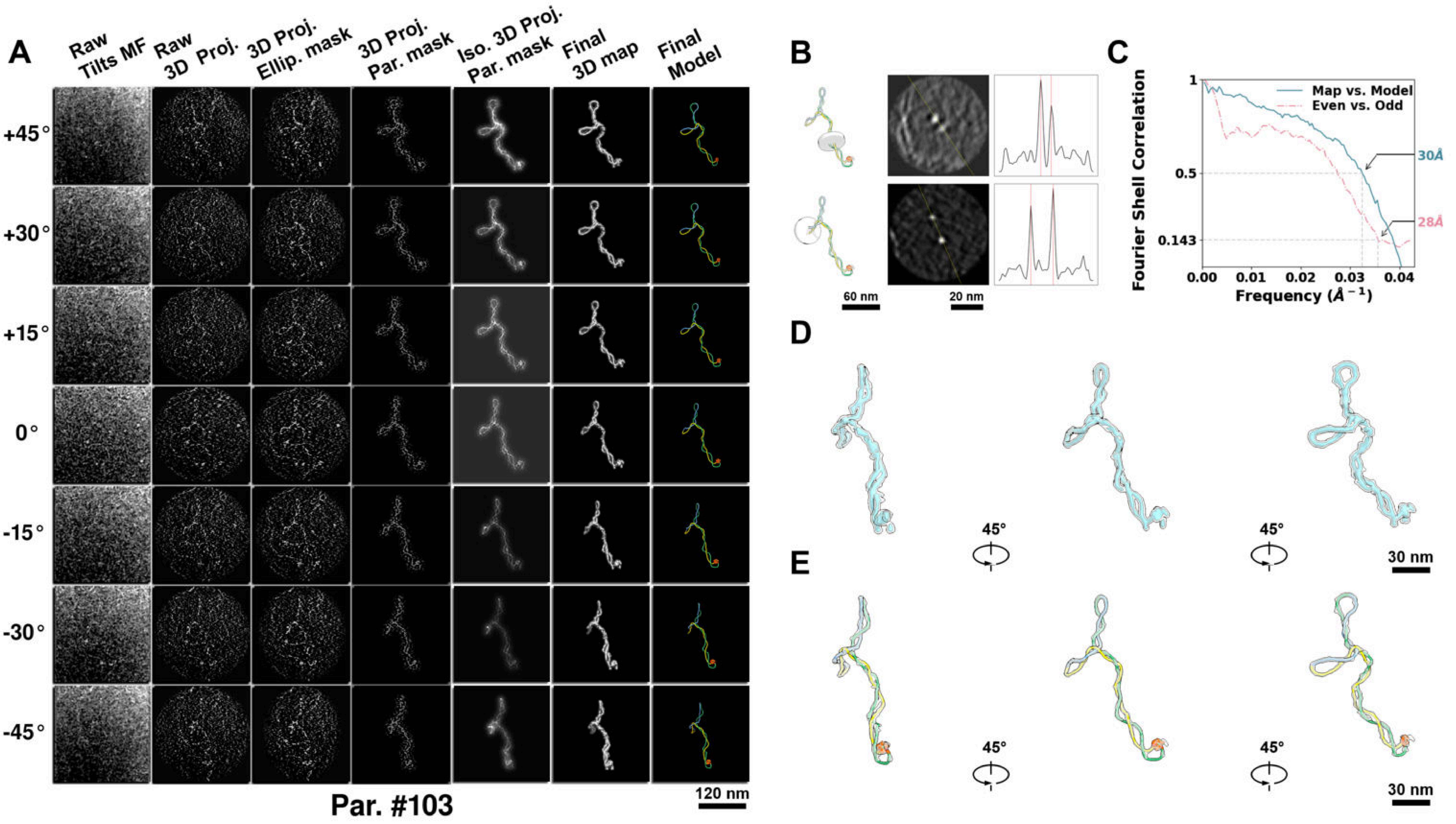

**Supplementary Particle Figure 103. Cryo-ET 3D reconstruction of an individual P.Cas particle.**
(A) 3D reconstruction of the plasmid particle (index no. 103). The first column shows seven representative tilt images from +45° to -45° in step of 15°. The second, third, and fourth columns show 3D projections of the particle with spherical, ellipsoidal (thinner along the z-dimension), and particle-shaped masks, respectively. The fifth column displays the 3D projections of the enhanced and IsoNet missing-wedge-corrected particle. The sixth and seventh columns present the final 3D map and the flexibly fitted model, respectively. (B) Two cross-sectional views (12 nm thickness) of the plasmid density map along its plectoneme axis are shown in the left-middle panel. The intensity profile along the line crossing the two high-density DNA spots is displayed in the right panel. (C) Resolution assessment of the final 3D map using Fourier shell correlation (FSC). Two criteria are shown: FSC between two half-maps reconstructed from even and odd frames (evaluated at 0.143) and FSC between the final 3D map and the fitted model (evaluated at 0.5). (D) Zoomed-in views of the final 3D density map from panel A, displayed at two contour levels. (E) Superimposition of the high-contour level map from panel D onto its fitted model.

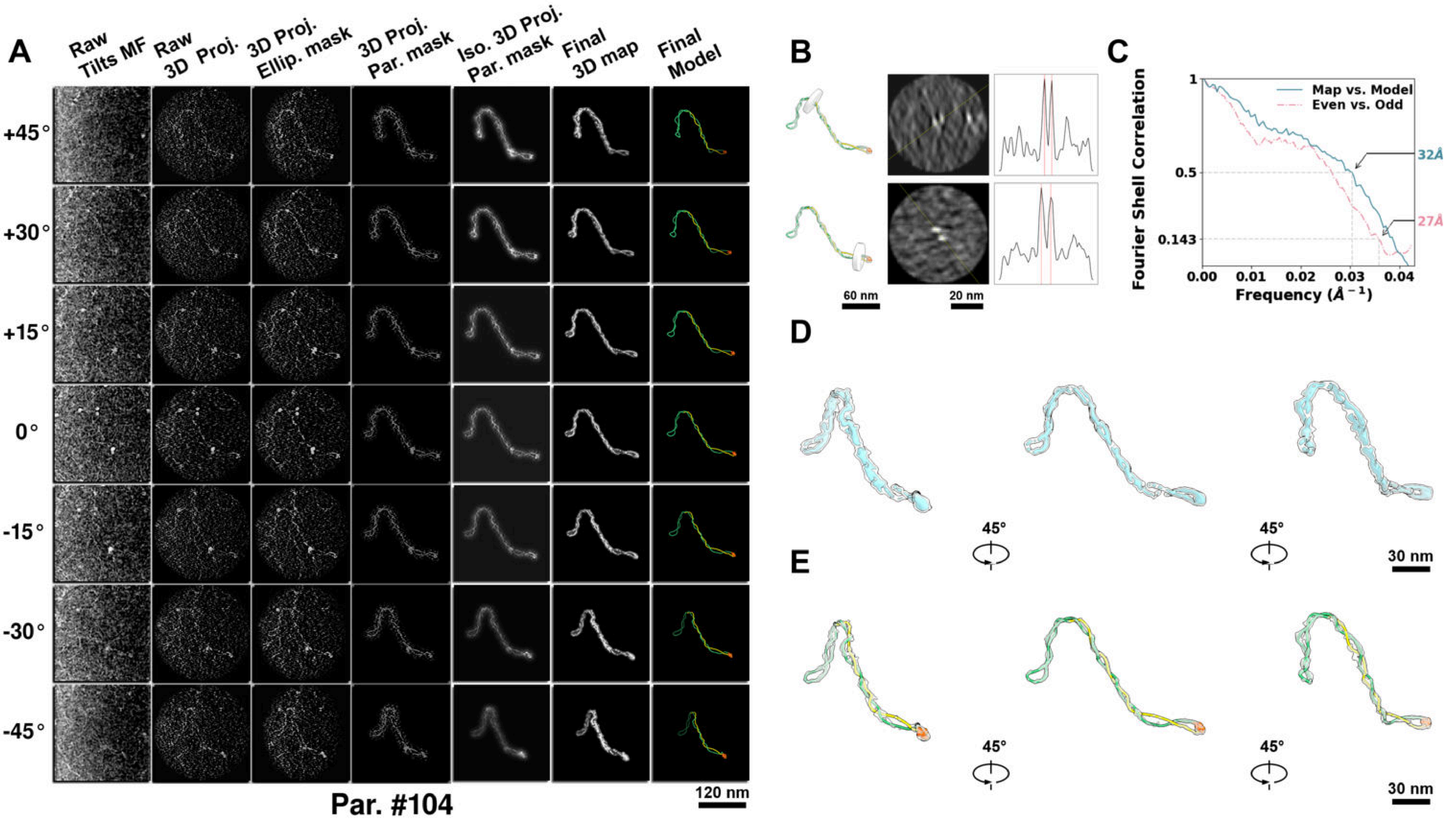

**Supplementary Particle Figure 104. Cryo-ET 3D reconstruction of an individual P.Cas particle.**
(A) 3D reconstruction of the plasmid particle (index no. 104). The first column shows seven representative tilt images from +45° to -45° in step of 15°. The second, third, and fourth columns show 3D projections of the particle with spherical, ellipsoidal (thinner along the z-dimension), and particle-shaped masks, respectively. The fifth column displays the 3D projections of the enhanced and IsoNet missing-wedge-corrected particle. The sixth and seventh columns present the final 3D map and the flexibly fitted model, respectively. (B) Two cross-sectional views (12 nm thickness) of the plasmid density map along its plectoneme axis are shown in the left-middle panel. The intensity profile along the line crossing the two high-density DNA spots is displayed in the right panel. (C) Resolution assessment of the final 3D map using Fourier shell correlation (FSC). Two criteria are shown: FSC between two half-maps reconstructed from even and odd frames (evaluated at 0.143) and FSC between the final 3D map and the fitted model (evaluated at 0.5). (D) Zoomed-in views of the final 3D density map from panel A, displayed at two contour levels. (E) Superimposition of the high-contour level map from panel D onto its fitted model.

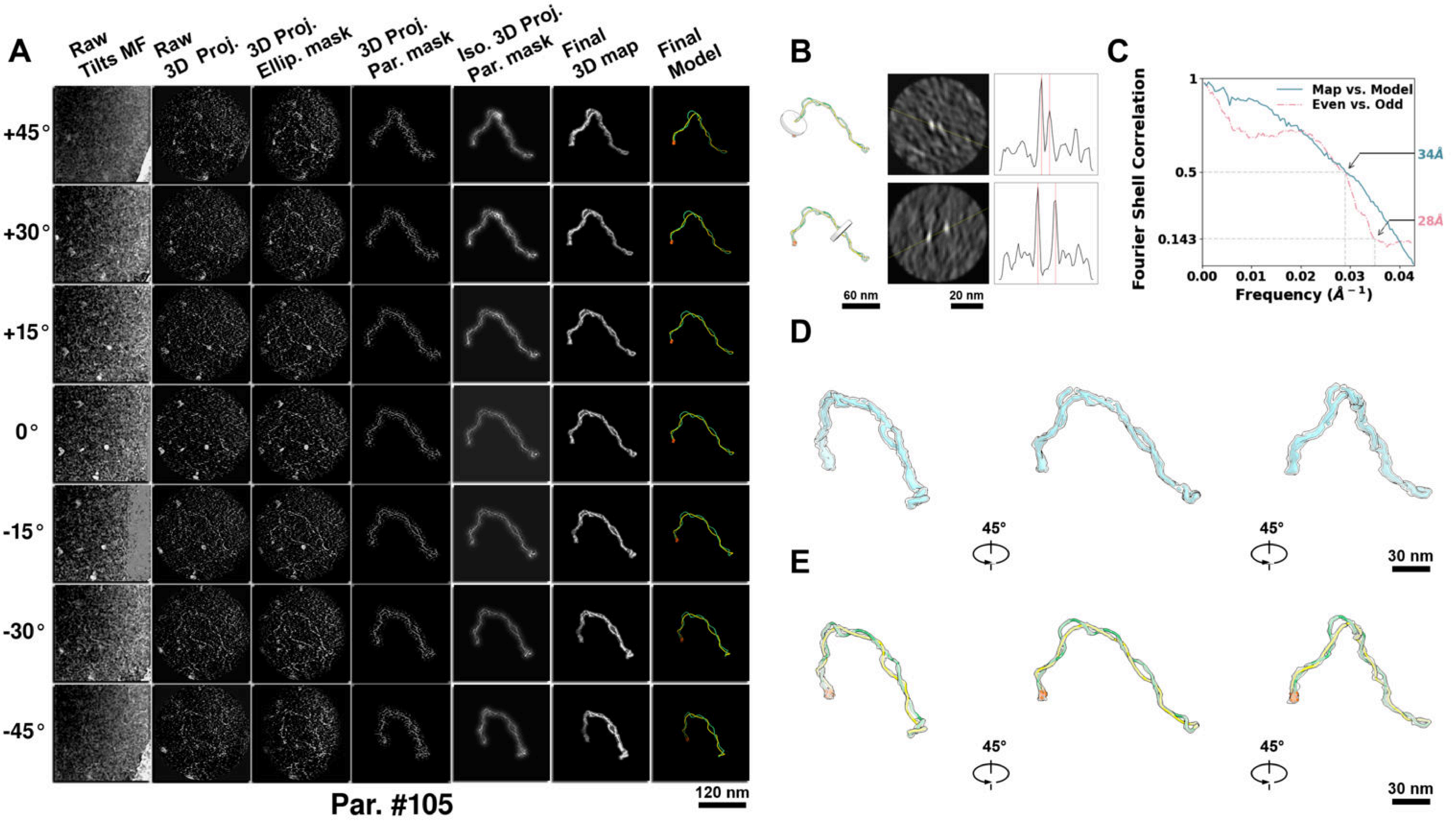

**Supplementary Particle Figure 105. Cryo-ET 3D reconstruction of an individual P.Cas particle.**
(A) 3D reconstruction of the plasmid particle (index no. 105). The first column shows seven representative tilt images from +45° to -45° in step of 15°. The second, third, and fourth columns show 3D projections of the particle with spherical, ellipsoidal (thinner along the z-dimension), and particle-shaped masks, respectively. The fifth column displays the 3D projections of the enhanced and IsoNet missing-wedge-corrected particle. The sixth and seventh columns present the final 3D map and the flexibly fitted model, respectively. (B) Two cross-sectional views (12 nm thickness) of the plasmid density map along its plectoneme axis are shown in the left-middle panel. The intensity profile along the line crossing the two high-density DNA spots is displayed in the right panel. (C) Resolution assessment of the final 3D map using Fourier shell correlation (FSC). Two criteria are shown: FSC between two half-maps reconstructed from even and odd frames (evaluated at 0.143) and FSC between the final 3D map and the fitted model (evaluated at 0.5). (D) Zoomed-in views of the final 3D density map from panel A, displayed at two contour levels. (E) Superimposition of the high-contour level map from panel D onto its fitted model.

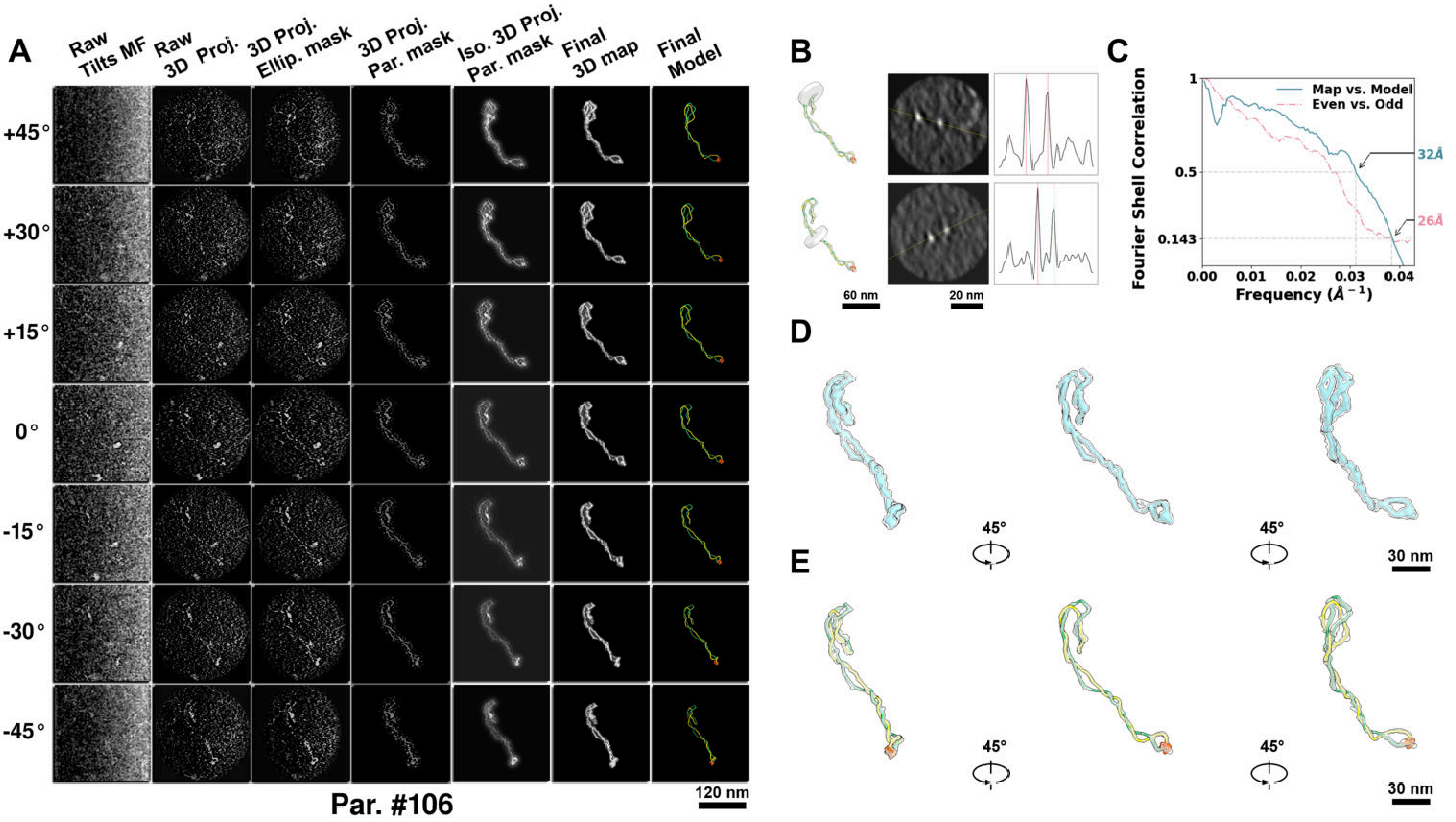

**Supplementary Particle Figure 106. Cryo-ET 3D reconstruction of an individual P.Cas particle.**
(A) 3D reconstruction of the plasmid particle (index no. 106). The first column shows seven representative tilt images from +45° to -45° in step of 15°. The second, third, and fourth columns show 3D projections of the particle with spherical, ellipsoidal (thinner along the z-dimension), and particle-shaped masks, respectively. The fifth column displays the 3D projections of the enhanced and IsoNet missing-wedge-corrected particle. The sixth and seventh columns present the final 3D map and the flexibly fitted model, respectively. (B) Two cross-sectional views (12 nm thickness) of the plasmid density map along its plectoneme axis are shown in the left-middle panel. The intensity profile along the line crossing the two high-density DNA spots is displayed in the right panel. (C) Resolution assessment of the final 3D map using Fourier shell correlation (FSC). Two criteria are shown: FSC between two half-maps reconstructed from even and odd frames (evaluated at 0.143) and FSC between the final 3D map and the fitted model (evaluated at 0.5). (D) Zoomed-in views of the final 3D density map from panel A, displayed at two contour levels. (E) Superimposition of the high-contour level map from panel D onto its fitted model.

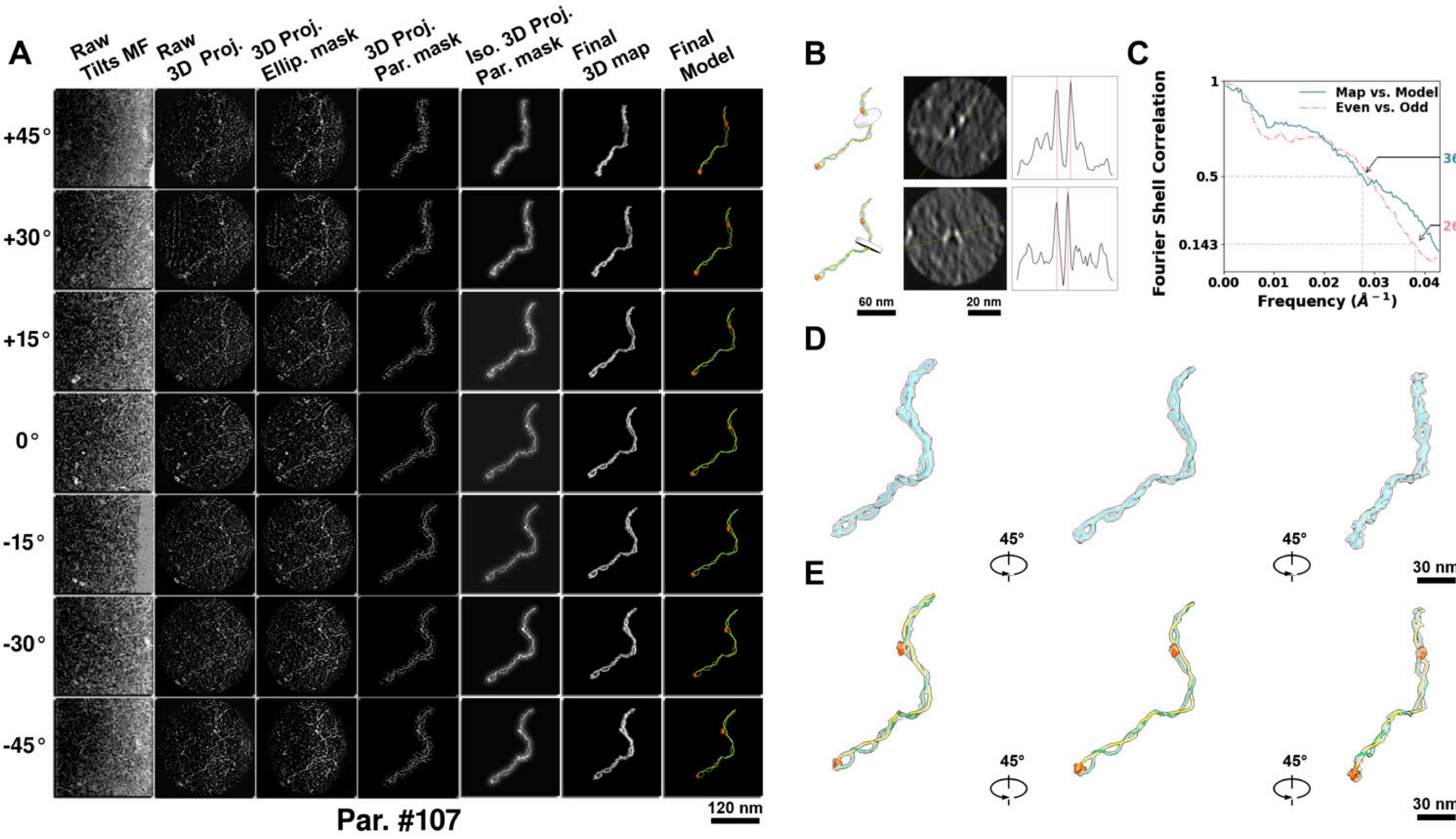

**Supplementary Particle Figure 107. Cryo-ET 3D reconstruction of an individual P.Cas particle.**
(A) 3D reconstruction of the plasmid particle (index no. 107). The first column shows seven representative tilt images from +45° to -45° in step of 15°. The second, third, and fourth columns show 3D projections of the particle with spherical, ellipsoidal (thinner along the z-dimension), and particle-shaped masks, respectively. The fifth column displays the 3D projections of the enhanced and IsoNet missing-wedge-corrected particle. The sixth and seventh columns present the final 3D map and the flexibly fitted model, respectively. (B) Two cross-sectional views (12 nm thickness) of the plasmid density map along its plectoneme axis are shown in the left-middle panel. The intensity profile along the line crossing the two high-density DNA spots is displayed in the right panel. (C) Resolution assessment of the final 3D map using Fourier shell correlation (FSC). Two criteria are shown: FSC between two half-maps reconstructed from even and odd frames (evaluated at 0.143) and FSC between the final 3D map and the fitted model (evaluated at 0.5). (D) Zoomed-in views of the final 3D density map from panel A, displayed at two contour levels. (E) Superimposition of the high-contour level map from panel D onto its fitted model.

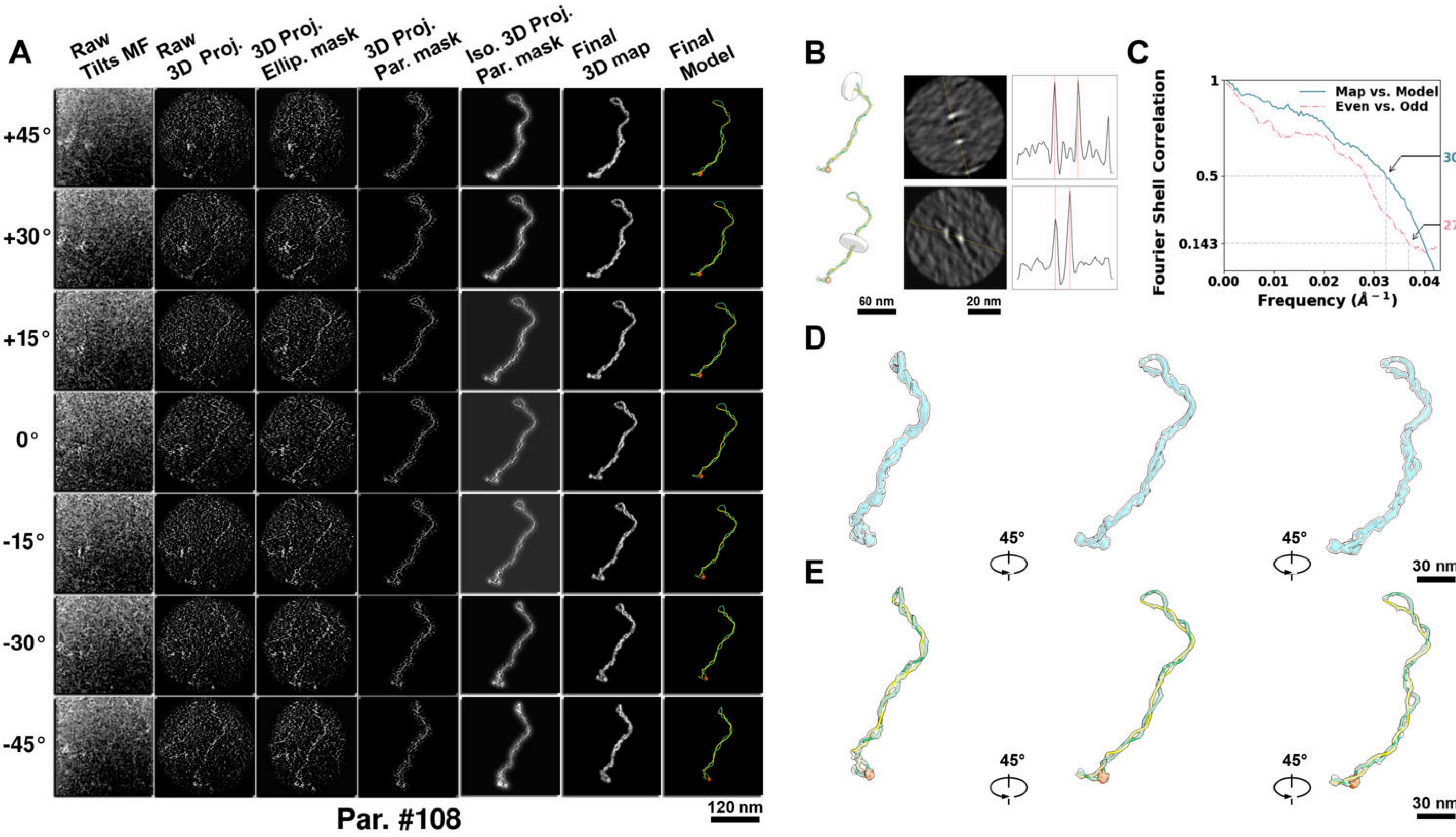

**Supplementary Particle Figure 108. Cryo-ET 3D reconstruction of an individual P.Cas particle.**
(A) 3D reconstruction of the plasmid particle (index no. 108). The first column shows seven representative tilt images from +45° to -45° in step of 15°. The second, third, and fourth columns show 3D projections of the particle with spherical, ellipsoidal (thinner along the z-dimension), and particle-shaped masks, respectively. The fifth column displays the 3D projections of the enhanced and IsoNet missing-wedge-corrected particle. The sixth and seventh columns present the final 3D map and the flexibly fitted model, respectively. (B) Two cross-sectional views (12 nm thickness) of the plasmid density map along its plectoneme axis are shown in the left-middle panel. The intensity profile along the line crossing the two high-density DNA spots is displayed in the right panel. (C) Resolution assessment of the final 3D map using Fourier shell correlation (FSC). Two criteria are shown: FSC between two half-maps reconstructed from even and odd frames (evaluated at 0.143) and FSC between the final 3D map and the fitted model (evaluated at 0.5). (D) Zoomed-in views of the final 3D density map from panel A, displayed at two contour levels. (E) Superimposition of the high-contour level map from panel D onto its fitted model.

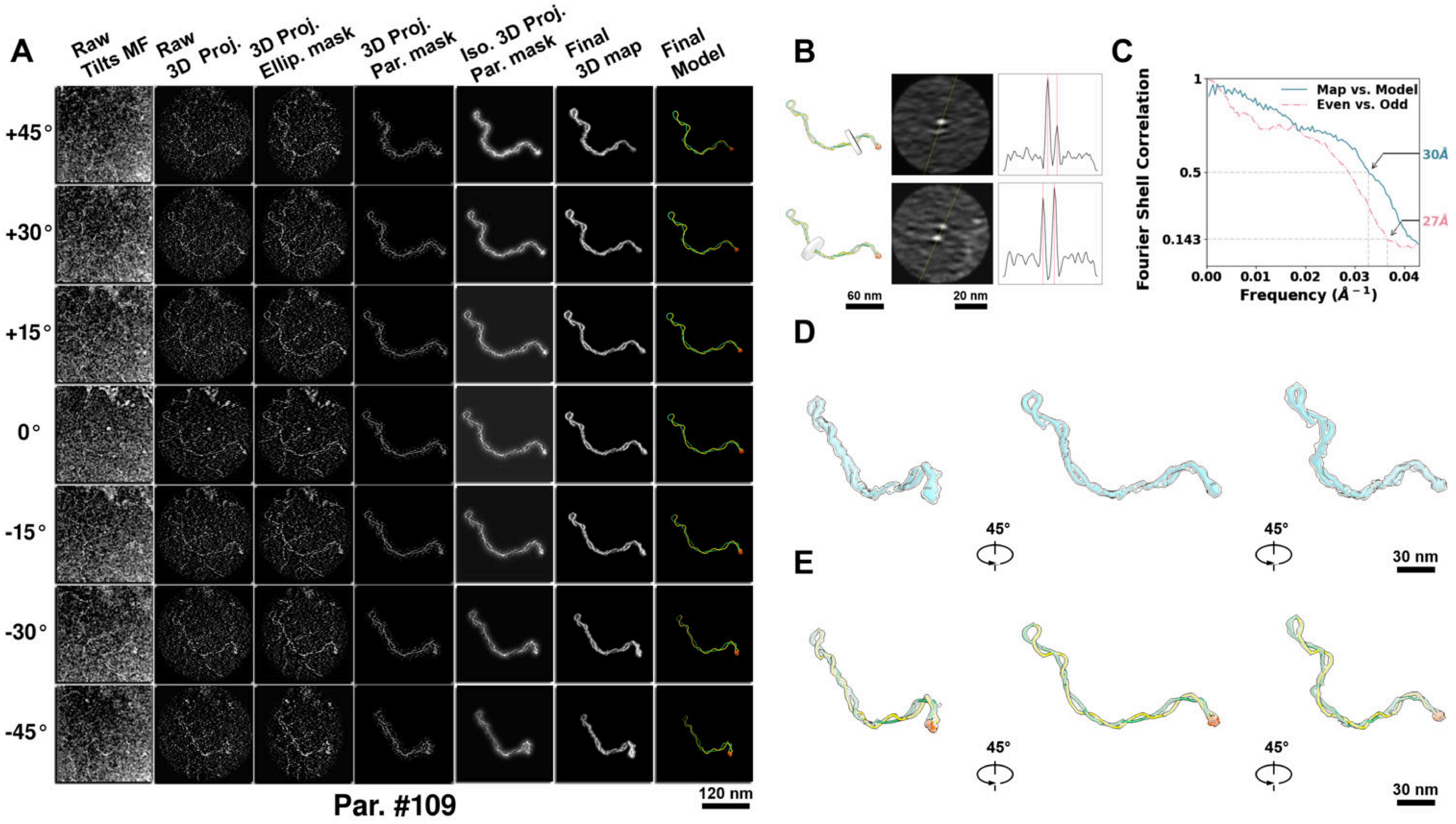

**Supplementary Particle Figure 109. Cryo-ET 3D reconstruction of an individual P.Cas particle.**
(A) 3D reconstruction of the plasmid particle (index no. 109). The first column shows seven representative tilt images from +45° to -45° in step of 15°. The second, third, and fourth columns show 3D projections of the particle with spherical, ellipsoidal (thinner along the z-dimension), and particle-shaped masks, respectively. The fifth column displays the 3D projections of the enhanced and IsoNet missing-wedge-corrected particle. The sixth and seventh columns present the final 3D map and the flexibly fitted model, respectively. (B) Two cross-sectional views (12 nm thickness) of the plasmid density map along its plectoneme axis are shown in the left-middle panel. The intensity profile along the line crossing the two high-density DNA spots is displayed in the right panel. (C) Resolution assessment of the final 3D map using Fourier shell correlation (FSC). Two criteria are shown: FSC between two half-maps reconstructed from even and odd frames (evaluated at 0.143) and FSC between the final 3D map and the fitted model (evaluated at 0.5). (D) Zoomed-in views of the final 3D density map from panel A, displayed at two contour levels. (E) Superimposition of the high-contour level map from panel D onto its fitted model.

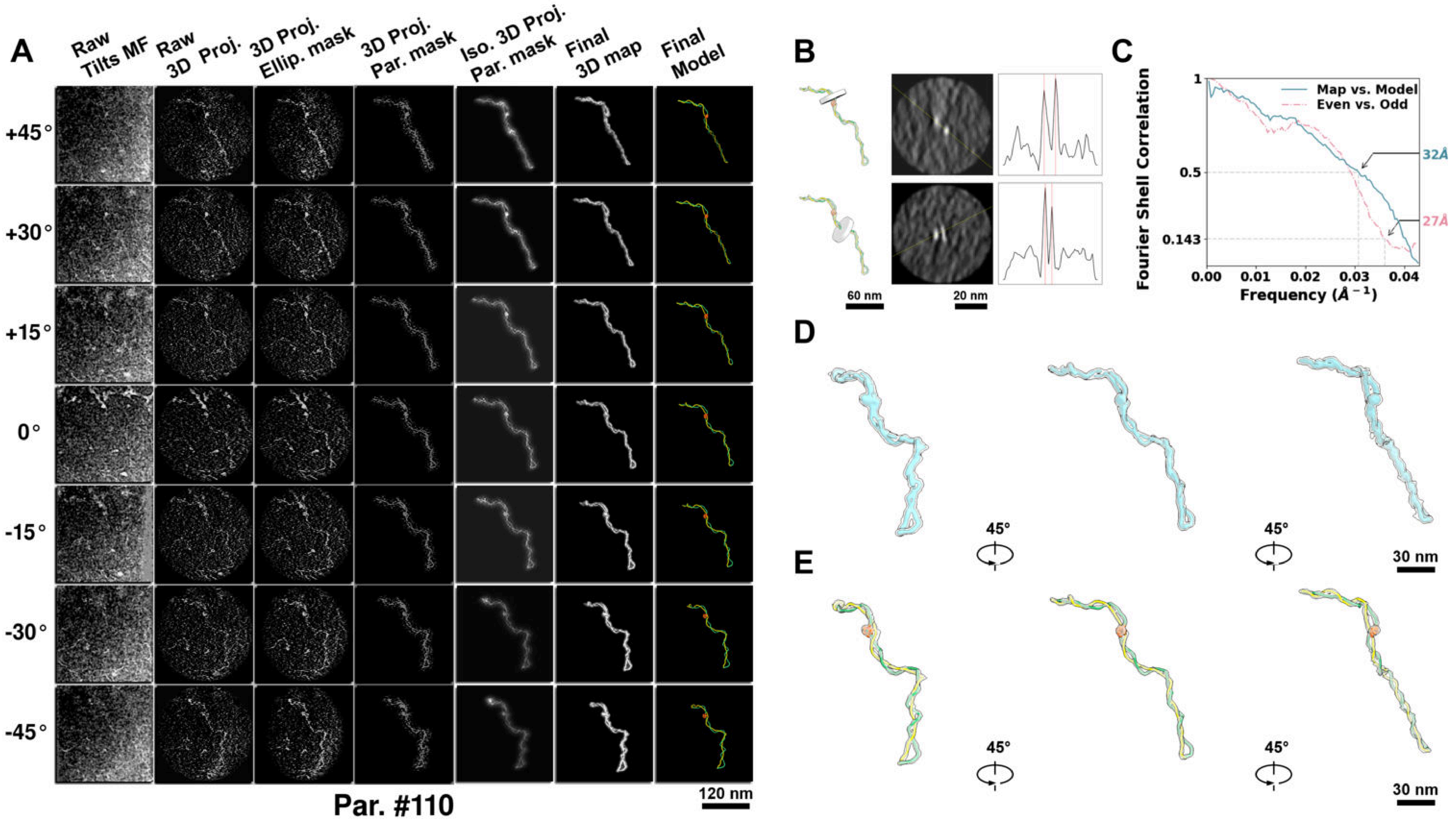

**Supplementary Particle Figure 110. Cryo-ET 3D reconstruction of an individual P.Cas particle.**
(A) 3D reconstruction of the plasmid particle (index no. 110). The first column shows seven representative tilt images from +45° to -45° in step of 15°. The second, third, and fourth columns show 3D projections of the particle with spherical, ellipsoidal (thinner along the z-dimension), and particle-shaped masks, respectively. The fifth column displays the 3D projections of the enhanced and IsoNet missing-wedge-corrected particle. The sixth and seventh columns present the final 3D map and the flexibly fitted model, respectively. (B) Two cross-sectional views (12 nm thickness) of the plasmid density map along its plectoneme axis are shown in the left-middle panel. The intensity profile along the line crossing the two high-density DNA spots is displayed in the right panel. (C) Resolution assessment of the final 3D map using Fourier shell correlation (FSC). Two criteria are shown: FSC between two half-maps reconstructed from even and odd frames (evaluated at 0.143) and FSC between the final 3D map and the fitted model (evaluated at 0.5). (D) Zoomed-in views of the final 3D density map from panel A, displayed at two contour levels. (E) Superimposition of the high-contour level map from panel D onto its fitted model.

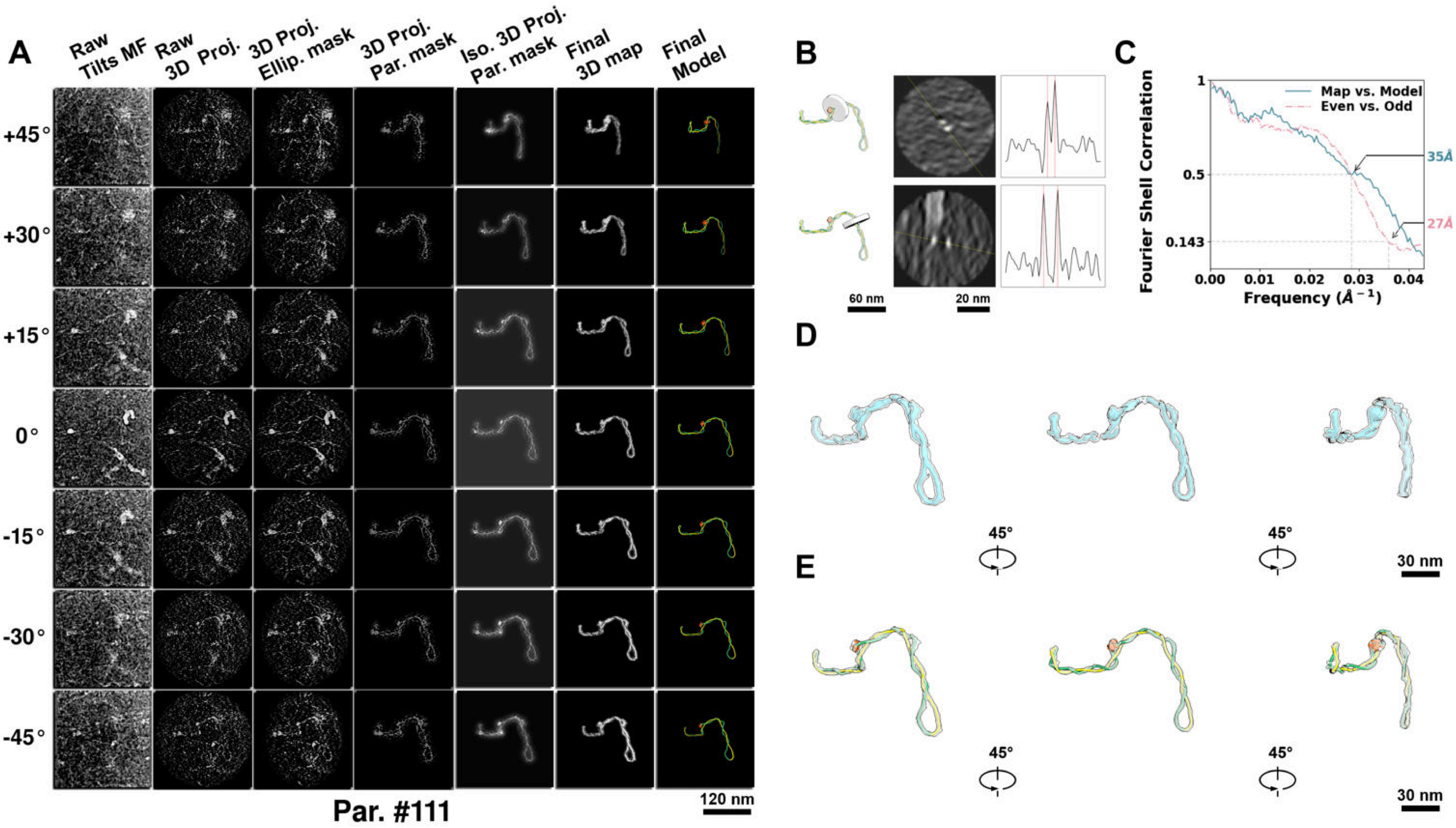

**Supplementary Particle Figure 111. Cryo-ET 3D reconstruction of an individual P.Cas particle.**
(A) 3D reconstruction of the plasmid particle (index no. 111). The first column shows seven representative tilt images from +45° to -45° in step of 15°. The second, third, and fourth columns show 3D projections of the particle with spherical, ellipsoidal (thinner along the z-dimension), and particle-shaped masks, respectively. The fifth column displays the 3D projections of the enhanced and IsoNet missing-wedge-corrected particle. The sixth and seventh columns present the final 3D map and the flexibly fitted model, respectively. (B) Two cross-sectional views (12 nm thickness) of the plasmid density map along its plectoneme axis are shown in the left-middle panel. The intensity profile along the line crossing the two high-density DNA spots is displayed in the right panel. (C) Resolution assessment of the final 3D map using Fourier shell correlation (FSC). Two criteria are shown: FSC between two half-maps reconstructed from even and odd frames (evaluated at 0.143) and FSC between the final 3D map and the fitted model (evaluated at 0.5). (D) Zoomed-in views of the final 3D density map from panel A, displayed at two contour levels. (E) Superimposition of the high-contour level map from panel D onto its fitted model.

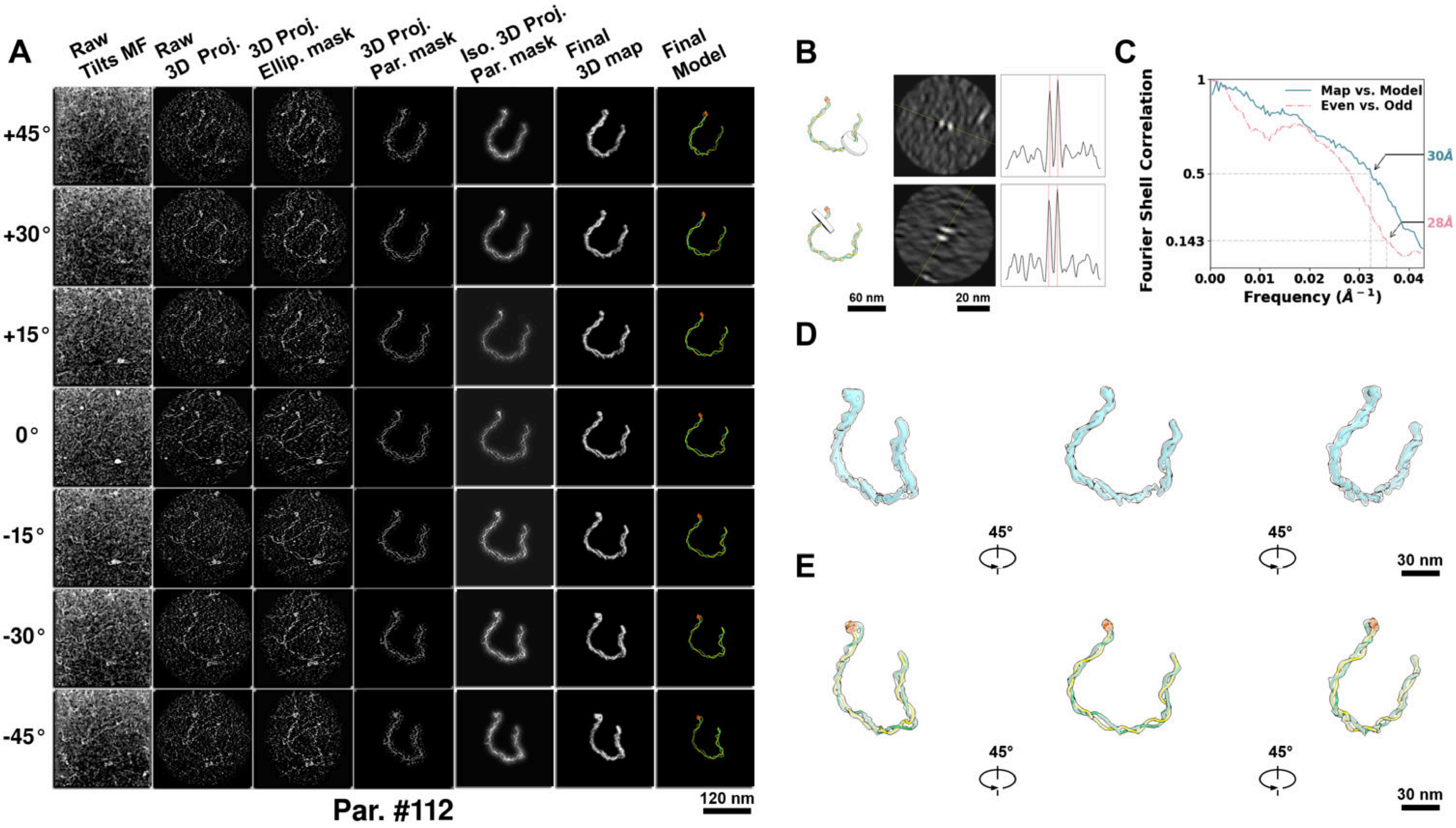

**Supplementary Particle Figure 112. Cryo-ET 3D reconstruction of an individual P.Cas particle.**
(A) 3D reconstruction of the plasmid particle (index no. 112). The first column shows seven representative tilt images from +45° to -45° in step of 15°. The second, third, and fourth columns show 3D projections of the particle with spherical, ellipsoidal (thinner along the z-dimension), and particle-shaped masks, respectively. The fifth column displays the 3D projections of the enhanced and IsoNet missing-wedge-corrected particle. The sixth and seventh columns present the final 3D map and the flexibly fitted model, respectively. (B) Two cross-sectional views (12 nm thickness) of the plasmid density map along its plectoneme axis are shown in the left-middle panel. The intensity profile along the line crossing the two high-density DNA spots is displayed in the right panel. (C) Resolution assessment of the final 3D map using Fourier shell correlation (FSC). Two criteria are shown: FSC between two half-maps reconstructed from even and odd frames (evaluated at 0.143) and FSC between the final 3D map and the fitted model (evaluated at 0.5). (D) Zoomed-in views of the final 3D density map from panel A, displayed at two contour levels. (E) Superimposition of the high-contour level map from panel D onto its fitted model.

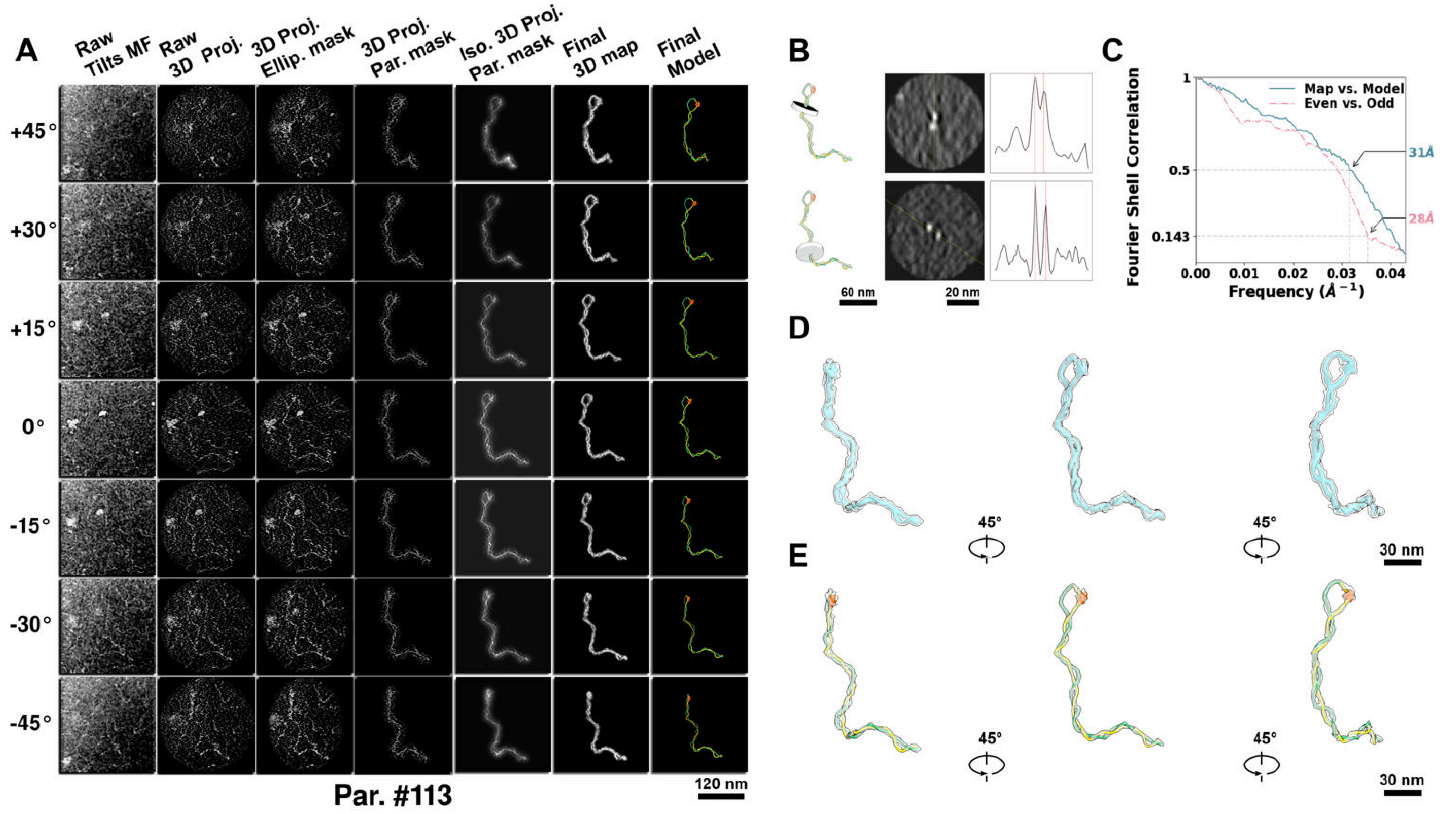

**Supplementary Particle Figure 113. Cryo-ET 3D reconstruction of an individual P.Cas particle.**
(A) 3D reconstruction of the plasmid particle (index no. 113). The first column shows seven representative tilt images from +45° to -45° in step of 15°. The second, third, and fourth columns show 3D projections of the particle with spherical, ellipsoidal (thinner along the z-dimension), and particle-shaped masks, respectively. The fifth column displays the 3D projections of the enhanced and IsoNet missing-wedge-corrected particle. The sixth and seventh columns present the final 3D map and the flexibly fitted model, respectively. (B) Two cross-sectional views (12 nm thickness) of the plasmid density map along its plectoneme axis are shown in the left-middle panel. The intensity profile along the line crossing the two high-density DNA spots is displayed in the right panel. (C) Resolution assessment of the final 3D map using Fourier shell correlation (FSC). Two criteria are shown: FSC between two half-maps reconstructed from even and odd frames (evaluated at 0.143) and FSC between the final 3D map and the fitted model (evaluated at 0.5). (D) Zoomed-in views of the final 3D density map from panel A, displayed at two contour levels. (E) Superimposition of the high-contour level map from panel D onto its fitted model.

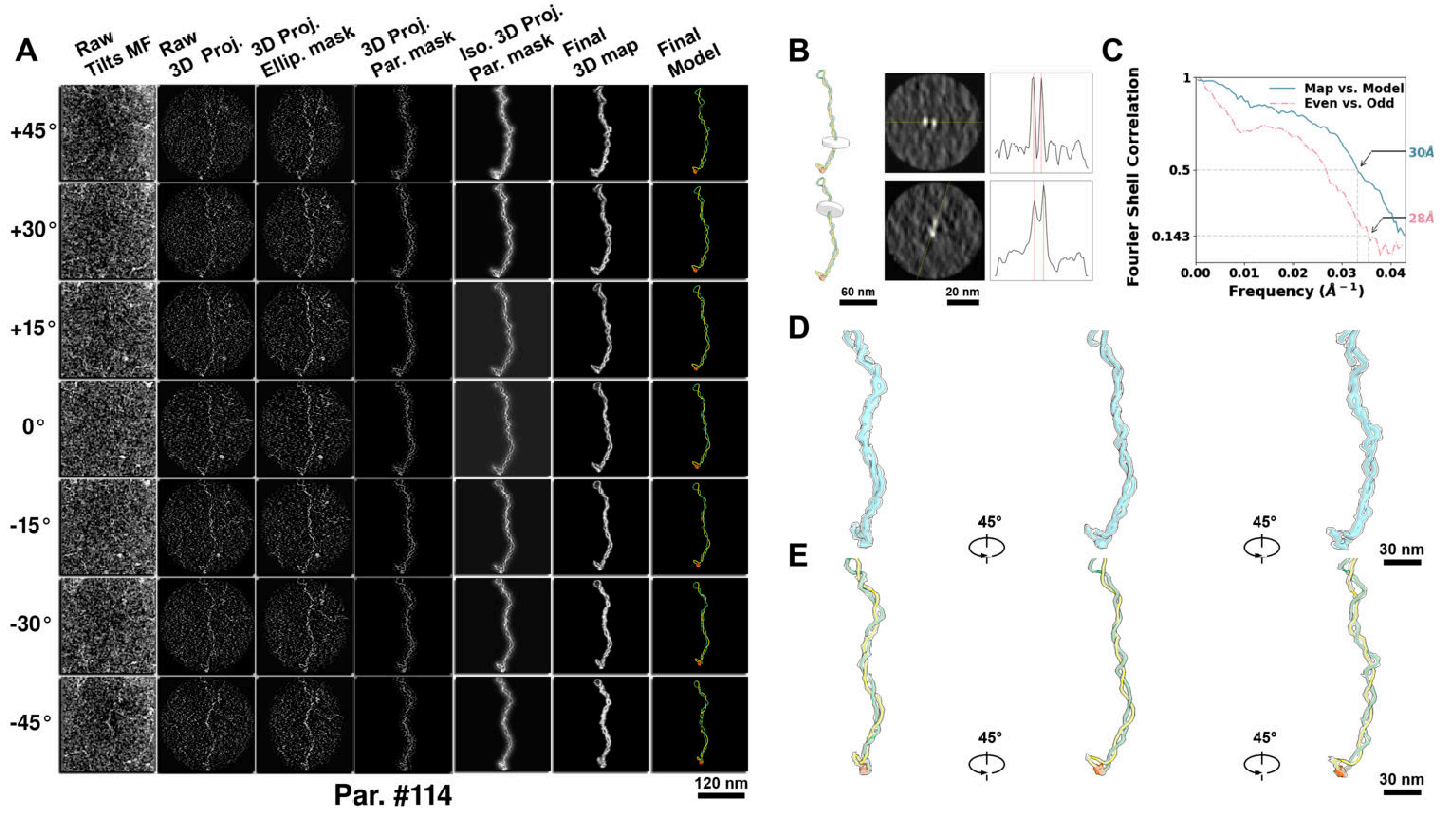

**Supplementary Particle Figure 114. Cryo-ET 3D reconstruction of an individual P.Cas particle.**
(A) 3D reconstruction of the plasmid particle (index no. 114). The first column shows seven representative tilt images from +45° to -45° in step of 15°. The second, third, and fourth columns show 3D projections of the particle with spherical, ellipsoidal (thinner along the z-dimension), and particle-shaped masks, respectively. The fifth column displays the 3D projections of the enhanced and IsoNet missing-wedge-corrected particle. The sixth and seventh columns present the final 3D map and the flexibly fitted model, respectively. (B) Two cross-sectional views (12 nm thickness) of the plasmid density map along its plectoneme axis are shown in the left-middle panel. The intensity profile along the line crossing the two high-density DNA spots is displayed in the right panel. (C) Resolution assessment of the final 3D map using Fourier shell correlation (FSC). Two criteria are shown: FSC between two half-maps reconstructed from even and odd frames (evaluated at 0.143) and FSC between the final 3D map and the fitted model (evaluated at 0.5). (D) Zoomed-in views of the final 3D density map from panel A, displayed at two contour levels. (E) Superimposition of the high-contour level map from panel D onto its fitted model.

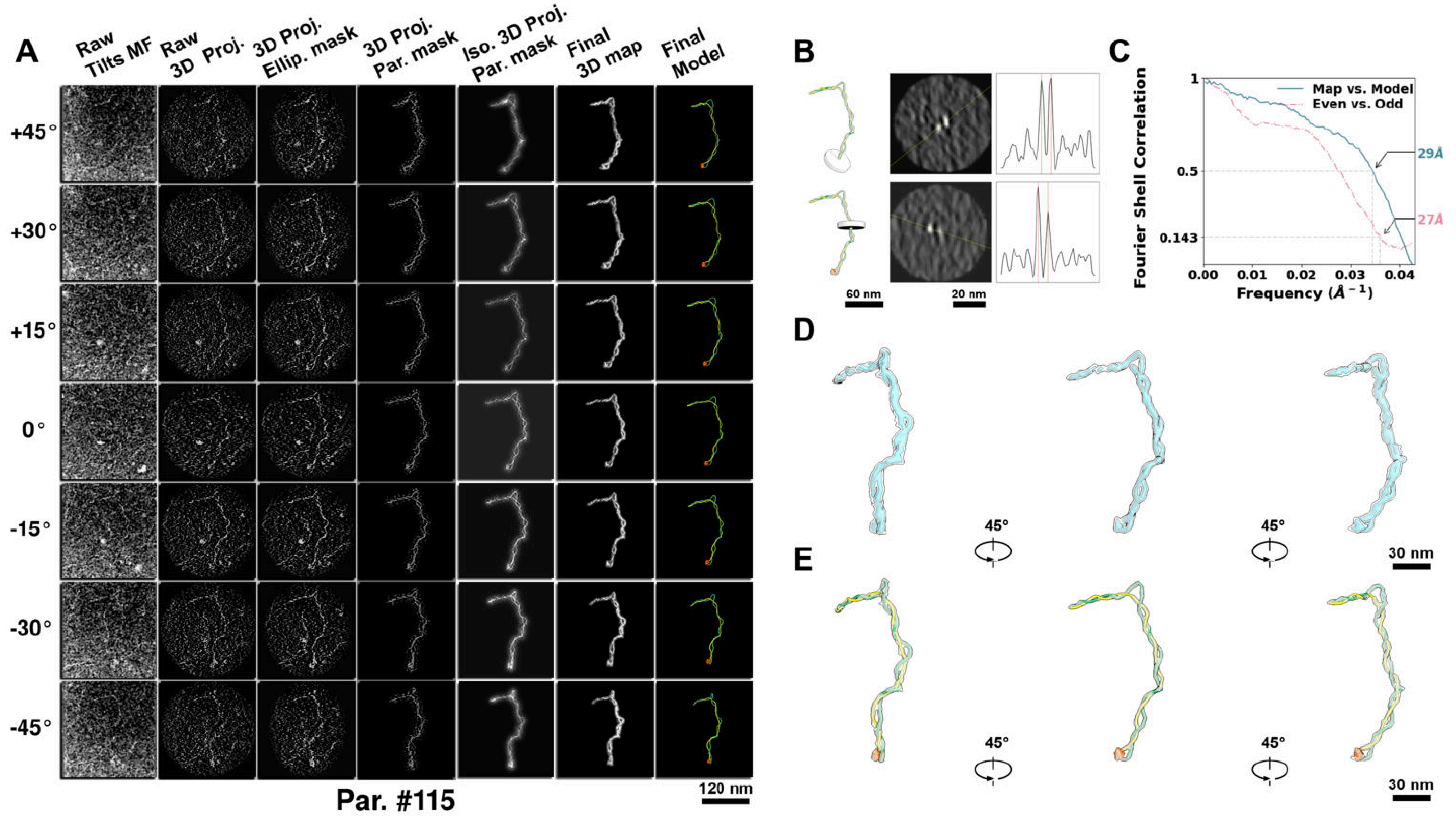

**Supplementary Particle Figure 115. Cryo-ET 3D reconstruction of an individual P.Cas particle.**
(A) 3D reconstruction of the plasmid particle (index no. 115). The first column shows seven representative tilt images from +45° to -45° in step of 15°. The second, third, and fourth columns show 3D projections of the particle with spherical, ellipsoidal (thinner along the z-dimension), and particle-shaped masks, respectively. The fifth column displays the 3D projections of the enhanced and IsoNet missing-wedge-corrected particle. The sixth and seventh columns present the final 3D map and the flexibly fitted model, respectively. (B) Two cross-sectional views (12 nm thickness) of the plasmid density map along its plectoneme axis are shown in the left-middle panel. The intensity profile along the line crossing the two high-density DNA spots is displayed in the right panel. (C) Resolution assessment of the final 3D map using Fourier shell correlation (FSC). Two criteria are shown: FSC between two half-maps reconstructed from even and odd frames (evaluated at 0.143) and FSC between the final 3D map and the fitted model (evaluated at 0.5). (D) Zoomed-in views of the final 3D density map from panel A, displayed at two contour levels. (E) Superimposition of the high-contour level map from panel D onto its fitted model.

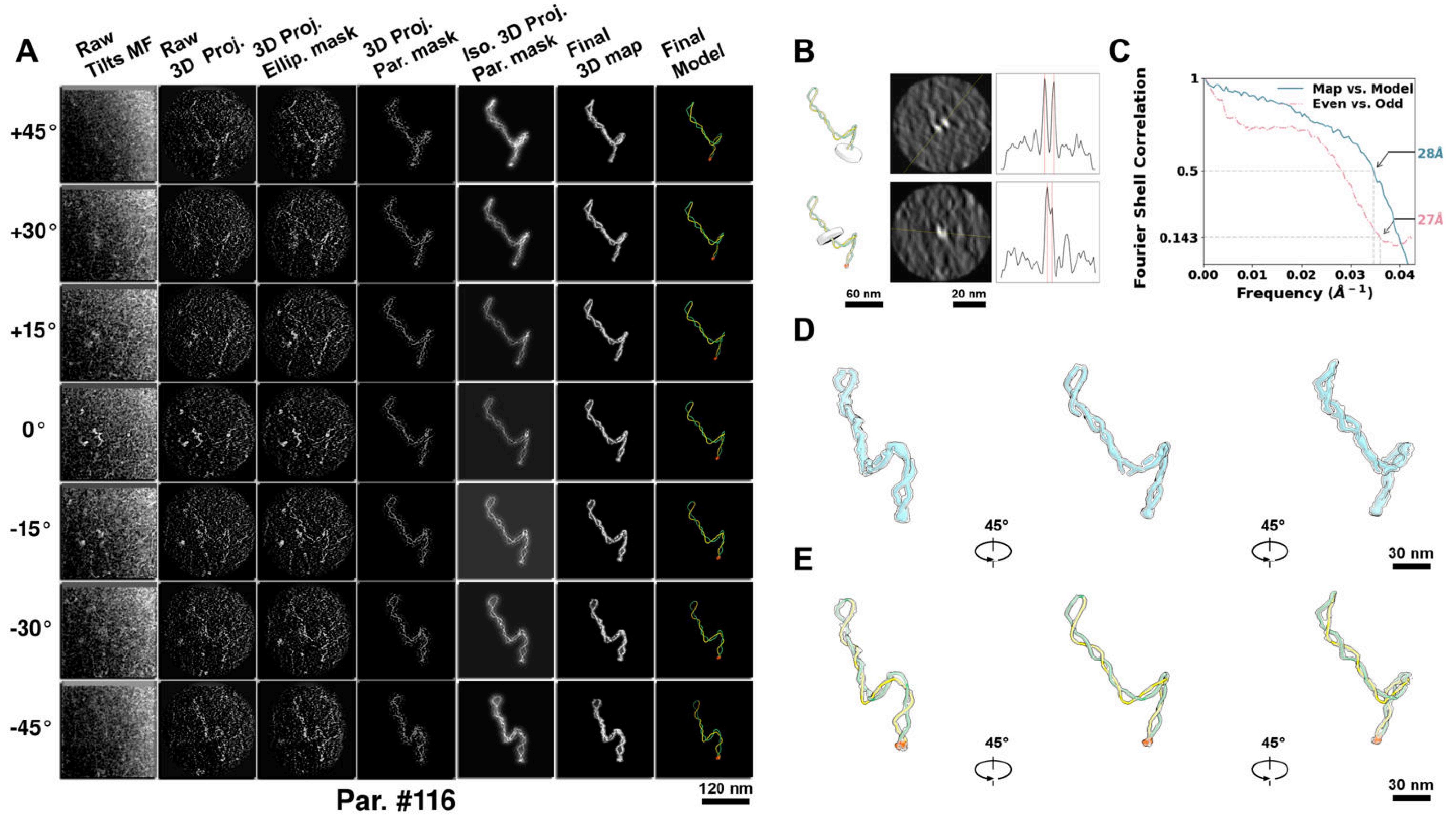

**Supplementary Particle Figure 116. Cryo-ET 3D reconstruction of an individual P.Cas particle.**
(A) 3D reconstruction of the plasmid particle (index no. 116). The first column shows seven representative tilt images from +45° to -45° in step of 15°. The second, third, and fourth columns show 3D projections of the particle with spherical, ellipsoidal (thinner along the z-dimension), and particle-shaped masks, respectively. The fifth column displays the 3D projections of the enhanced and IsoNet missing-wedge-corrected particle. The sixth and seventh columns present the final 3D map and the flexibly fitted model, respectively. (B) Two cross-sectional views (12 nm thickness) of the plasmid density map along its plectoneme axis are shown in the left-middle panel. The intensity profile along the line crossing the two high-density DNA spots is displayed in the right panel. (C) Resolution assessment of the final 3D map using Fourier shell correlation (FSC). Two criteria are shown: FSC between two half-maps reconstructed from even and odd frames (evaluated at 0.143) and FSC between the final 3D map and the fitted model (evaluated at 0.5). (D) Zoomed-in views of the final 3D density map from panel A, displayed at two contour levels. (E) Superimposition of the high-contour level map from panel D onto its fitted model.

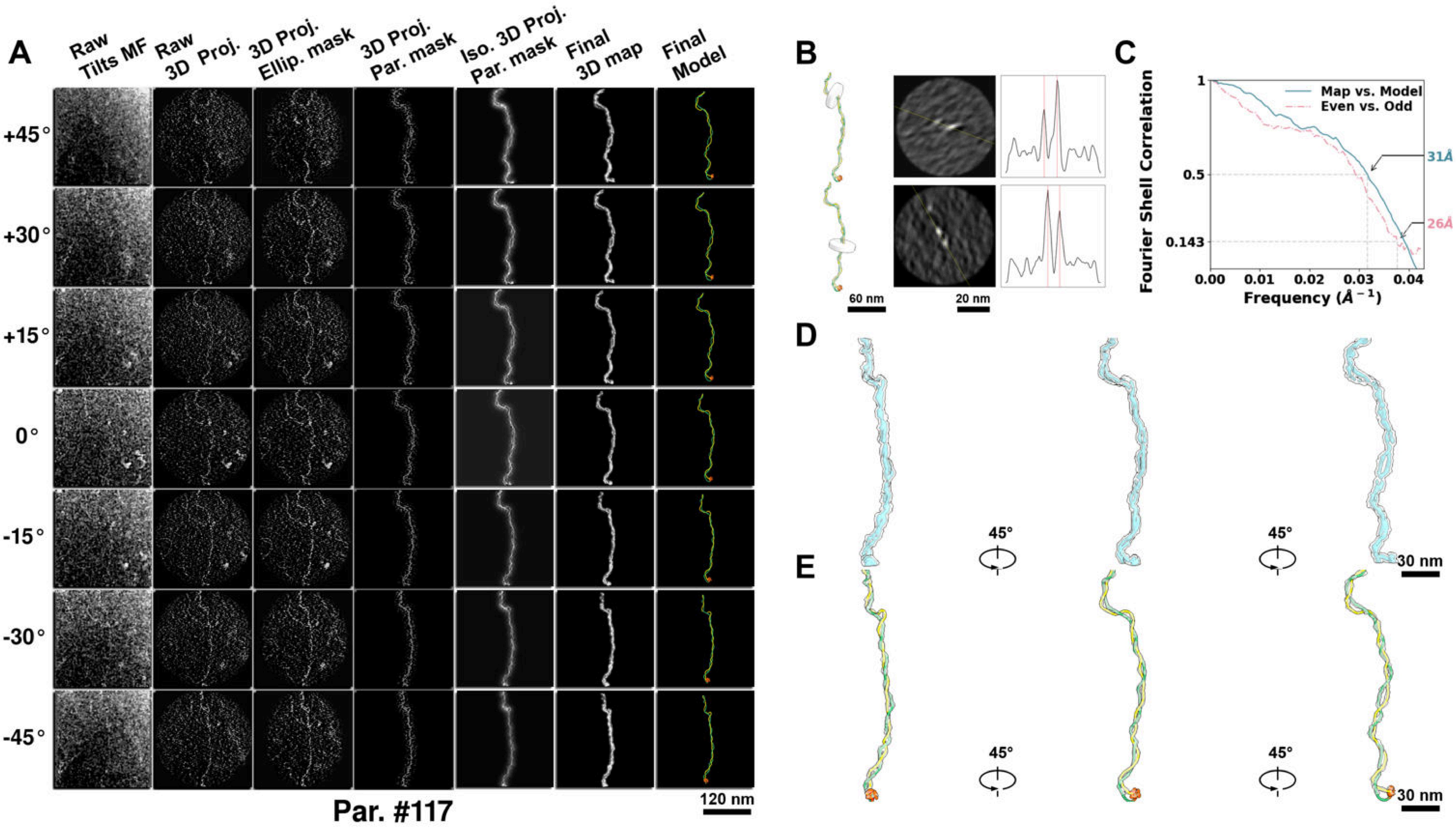

**Supplementary Particle Figure 117. Cryo-ET 3D reconstruction of an individual P.Cas particle.**
(A) 3D reconstruction of the plasmid particle (index no. 117). The first column shows seven representative tilt images from +45° to -45° in step of 15°. The second, third, and fourth columns show 3D projections of the particle with spherical, ellipsoidal (thinner along the z-dimension), and particle-shaped masks, respectively. The fifth column displays the 3D projections of the enhanced and IsoNet missing-wedge-corrected particle. The sixth and seventh columns present the final 3D map and the flexibly fitted model, respectively. (B) Two cross-sectional views (12 nm thickness) of the plasmid density map along its plectoneme axis are shown in the left-middle panel. The intensity profile along the line crossing the two high-density DNA spots is displayed in the right panel. (C) Resolution assessment of the final 3D map using Fourier shell correlation (FSC). Two criteria are shown: FSC between two half-maps reconstructed from even and odd frames (evaluated at 0.143) and FSC between the final 3D map and the fitted model (evaluated at 0.5). (D) Zoomed-in views of the final 3D density map from panel A, displayed at two contour levels. (E) Superimposition of the high-contour level map from panel D onto its fitted model.

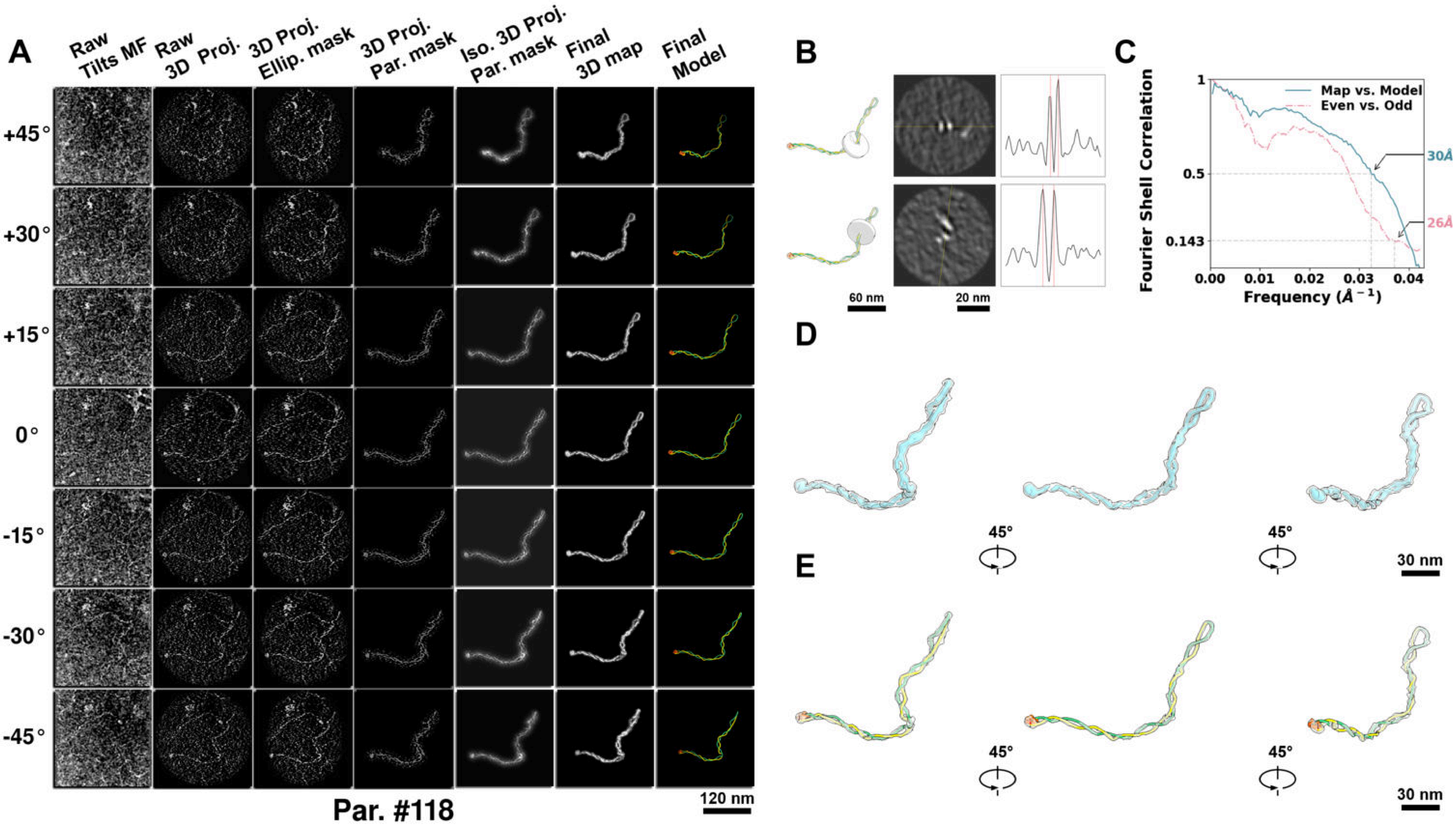

**Supplementary Particle Figure 118. Cryo-ET 3D reconstruction of an individual P.Cas particle.**
(A) 3D reconstruction of the plasmid particle (index no. 118). The first column shows seven representative tilt images from +45° to -45° in step of 15°. The second, third, and fourth columns show 3D projections of the particle with spherical, ellipsoidal (thinner along the z-dimension), and particle-shaped masks, respectively. The fifth column displays the 3D projections of the enhanced and IsoNet missing-wedge-corrected particle. The sixth and seventh columns present the final 3D map and the flexibly fitted model, respectively. (B) Two cross-sectional views (12 nm thickness) of the plasmid density map along its plectoneme axis are shown in the left-middle panel. The intensity profile along the line crossing the two high-density DNA spots is displayed in the right panel. (C) Resolution assessment of the final 3D map using Fourier shell correlation (FSC). Two criteria are shown: FSC between two half-maps reconstructed from even and odd frames (evaluated at 0.143) and FSC between the final 3D map and the fitted model (evaluated at 0.5). (D) Zoomed-in views of the final 3D density map from panel A, displayed at two contour levels. (E) Superimposition of the high-contour level map from panel D onto its fitted model.

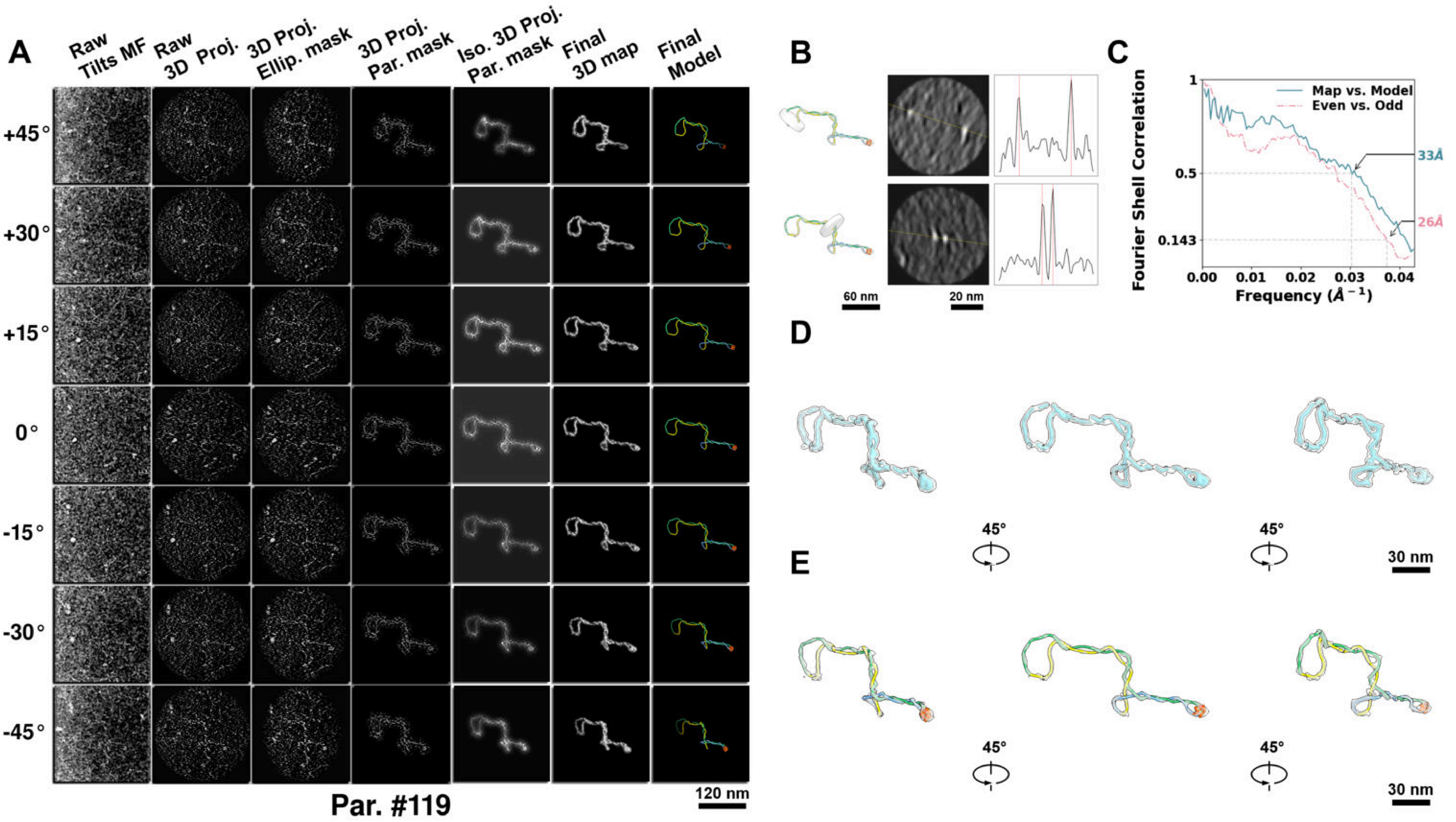

**Supplementary Particle Figure 119. Cryo-ET 3D reconstruction of an individual P.Cas particle.**
(A) 3D reconstruction of the plasmid particle (index no. 119). The first column shows seven representative tilt images from +45° to -45° in step of 15°. The second, third, and fourth columns show 3D projections of the particle with spherical, ellipsoidal (thinner along the z-dimension), and particle-shaped masks, respectively. The fifth column displays the 3D projections of the enhanced and IsoNet missing-wedge-corrected particle. The sixth and seventh columns present the final 3D map and the flexibly fitted model, respectively. (B) Two cross-sectional views (12 nm thickness) of the plasmid density map along its plectoneme axis are shown in the left-middle panel. The intensity profile along the line crossing the two high-density DNA spots is displayed in the right panel. (C) Resolution assessment of the final 3D map using Fourier shell correlation (FSC). Two criteria are shown: FSC between two half-maps reconstructed from even and odd frames (evaluated at 0.143) and FSC between the final 3D map and the fitted model (evaluated at 0.5). (D) Zoomed-in views of the final 3D density map from panel A, displayed at two contour levels. (E) Superimposition of the high-contour level map from panel D onto its fitted model.

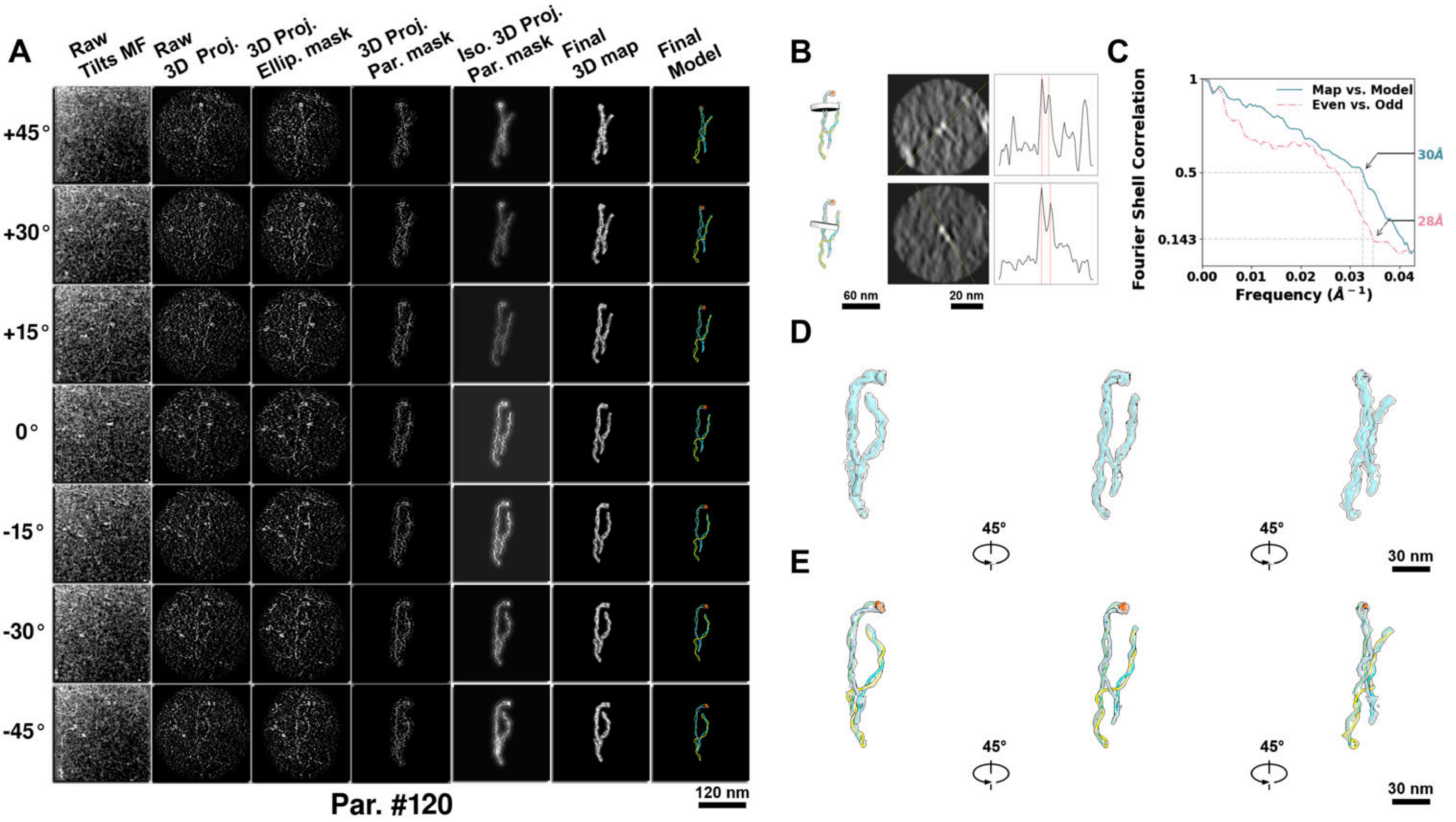

**Supplementary Particle Figure 120. Cryo-ET 3D reconstruction of an individual P.Cas particle.**
(A) 3D reconstruction of the plasmid particle (index no. 120). The first column shows seven representative tilt images from +45° to -45° in step of 15°. The second, third, and fourth columns show 3D projections of the particle with spherical, ellipsoidal (thinner along the z-dimension), and particle-shaped masks, respectively. The fifth column displays the 3D projections of the enhanced and IsoNet missing-wedge-corrected particle. The sixth and seventh columns present the final 3D map and the flexibly fitted model, respectively. (B) Two cross-sectional views (12 nm thickness) of the plasmid density map along its plectoneme axis are shown in the left-middle panel. The intensity profile along the line crossing the two high-density DNA spots is displayed in the right panel. (C) Resolution assessment of the final 3D map using Fourier shell correlation (FSC). Two criteria are shown: FSC between two half-maps reconstructed from even and odd frames (evaluated at 0.143) and FSC between the final 3D map and the fitted model (evaluated at 0.5). (D) Zoomed-in views of the final 3D density map from panel A, displayed at two contour levels. (E) Superimposition of the high-contour level map from panel D onto its fitted model.

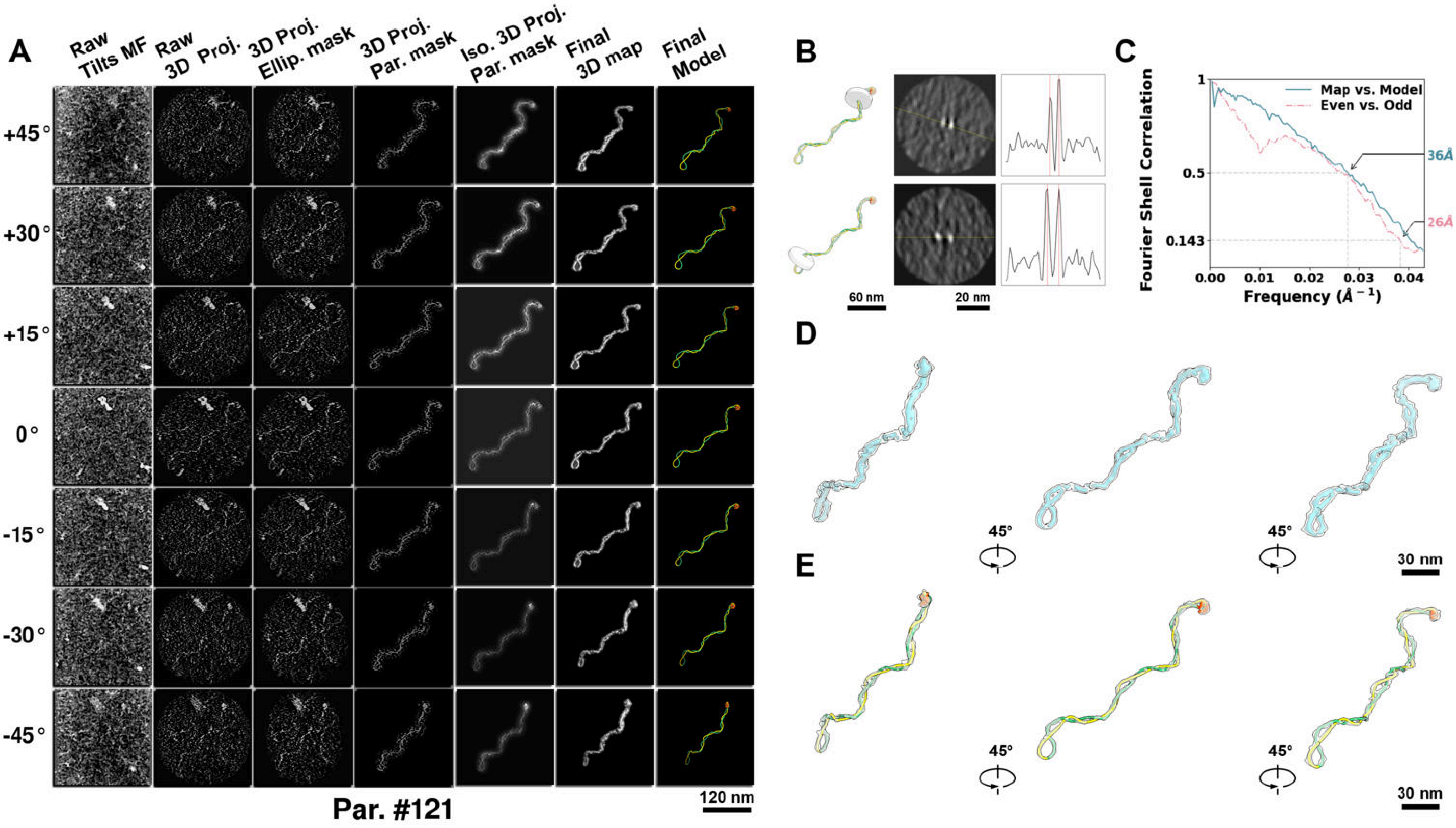

**Supplementary Particle Figure 121. Cryo-ET 3D reconstruction of an individual P.Cas particle.**
(A) 3D reconstruction of the plasmid particle (index no. 121). The first column shows seven representative tilt images from +45° to -45° in step of 15°. The second, third, and fourth columns show 3D projections of the particle with spherical, ellipsoidal (thinner along the z-dimension), and particle-shaped masks, respectively. The fifth column displays the 3D projections of the enhanced and IsoNet missing-wedge-corrected particle. The sixth and seventh columns present the final 3D map and the flexibly fitted model, respectively. (B) Two cross-sectional views (12 nm thickness) of the plasmid density map along its plectoneme axis are shown in the left-middle panel. The intensity profile along the line crossing the two high-density DNA spots is displayed in the right panel. (C) Resolution assessment of the final 3D map using Fourier shell correlation (FSC). Two criteria are shown: FSC between two half-maps reconstructed from even and odd frames (evaluated at 0.143) and FSC between the final 3D map and the fitted model (evaluated at 0.5). (D) Zoomed-in views of the final 3D density map from panel A, displayed at two contour levels. (E) Superimposition of the high-contour level map from panel D onto its fitted model.

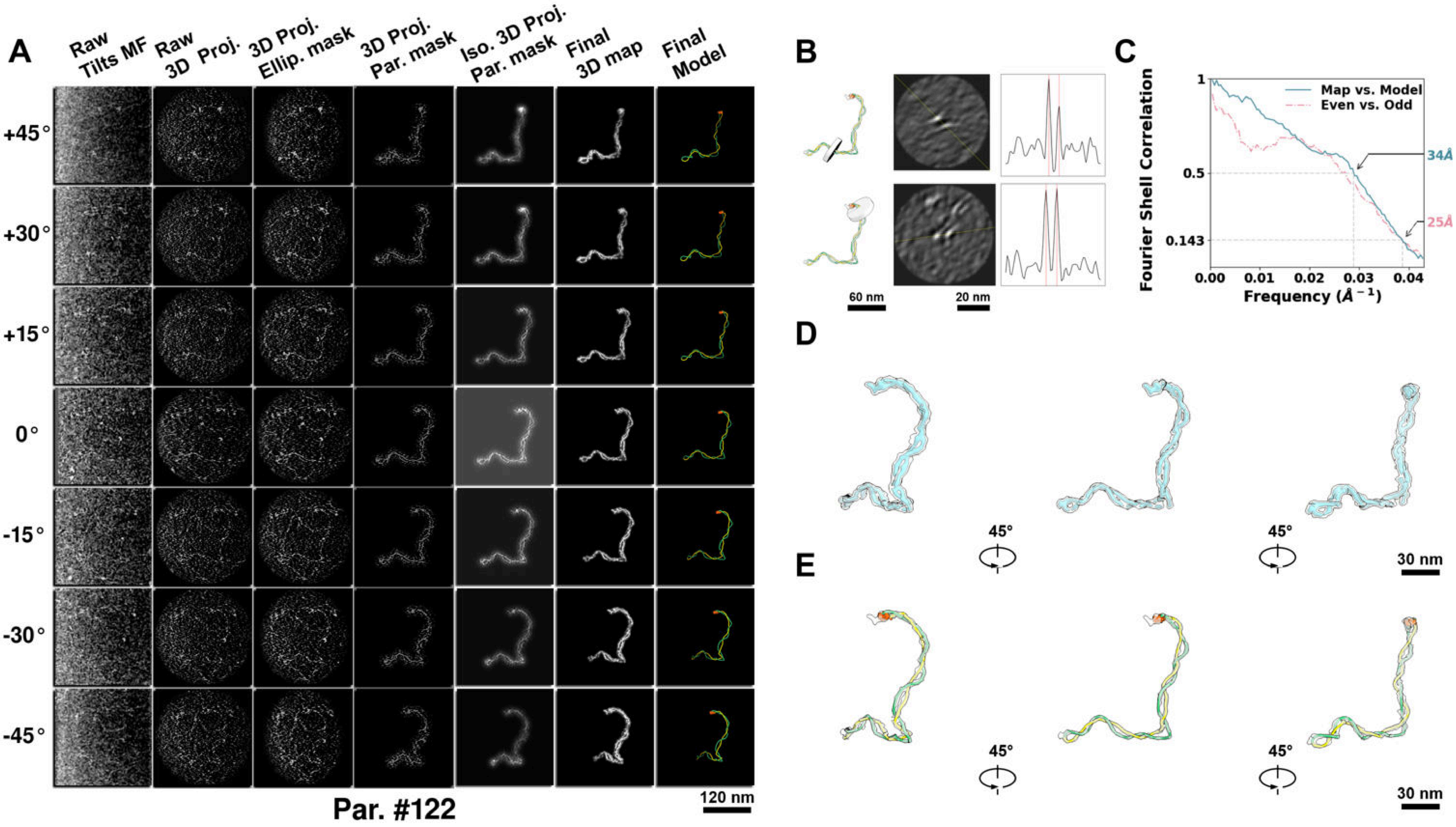

**Supplementary Particle Figure 122. Cryo-ET 3D reconstruction of an individual P.Cas particle.**
(A) 3D reconstruction of the plasmid particle (index no. 122). The first column shows seven representative tilt images from +45° to -45° in step of 15°. The second, third, and fourth columns show 3D projections of the particle with spherical, ellipsoidal (thinner along the z-dimension), and particle-shaped masks, respectively. The fifth column displays the 3D projections of the enhanced and IsoNet missing-wedge-corrected particle. The sixth and seventh columns present the final 3D map and the flexibly fitted model, respectively. (B) Two cross-sectional views (12 nm thickness) of the plasmid density map along its plectoneme axis are shown in the left-middle panel. The intensity profile along the line crossing the two high-density DNA spots is displayed in the right panel. (C) Resolution assessment of the final 3D map using Fourier shell correlation (FSC). Two criteria are shown: FSC between two half-maps reconstructed from even and odd frames (evaluated at 0.143) and FSC between the final 3D map and the fitted model (evaluated at 0.5). (D) Zoomed-in views of the final 3D density map from panel A, displayed at two contour levels. (E) Superimposition of the high-contour level map from panel D onto its fitted model.

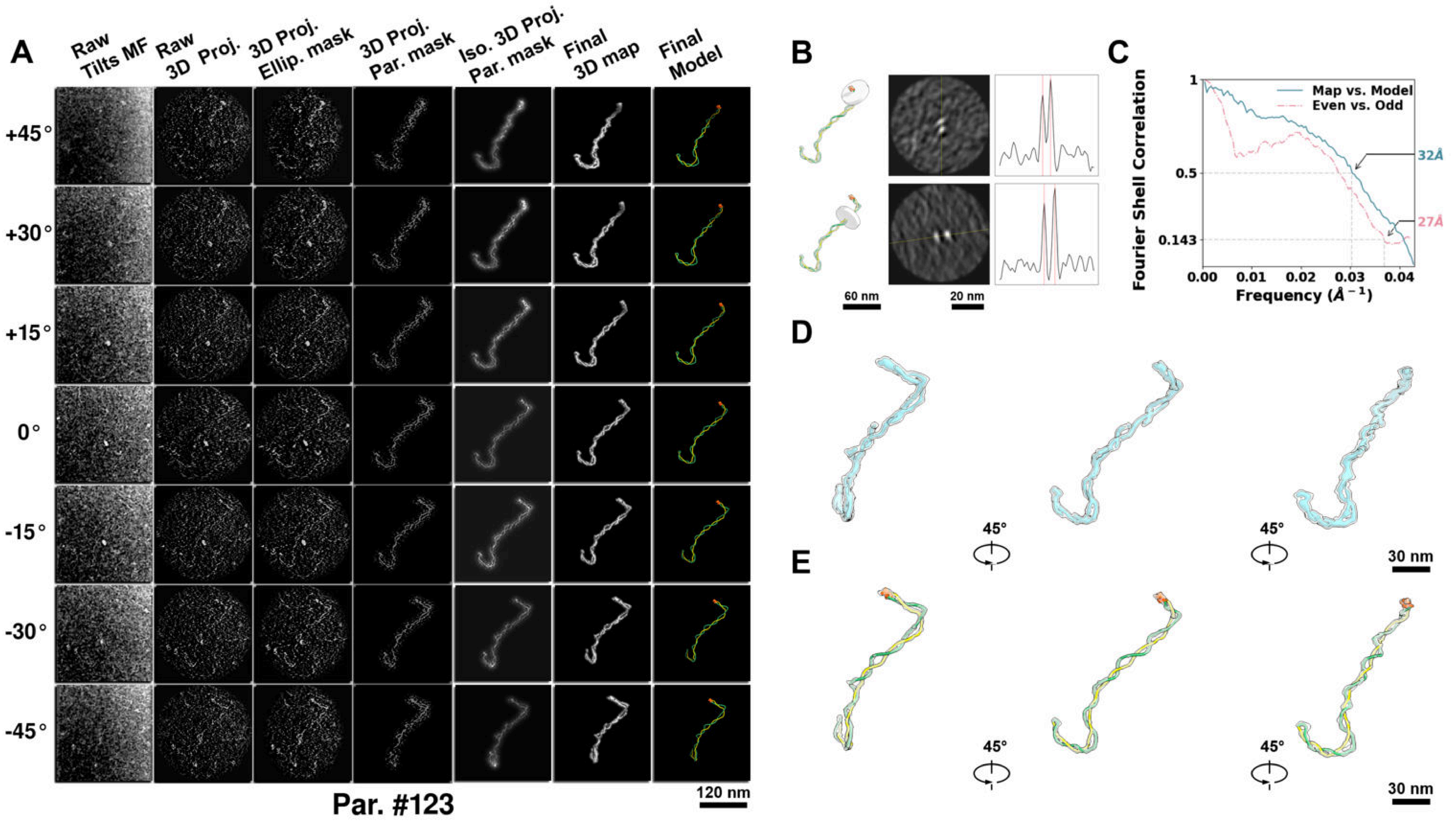

**Supplementary Particle Figure 123. Cryo-ET 3D reconstruction of an individual P.Cas particle.**
(A) 3D reconstruction of the plasmid particle (index no. 123). The first column shows seven representative tilt images from +45° to -45° in step of 15°. The second, third, and fourth columns show 3D projections of the particle with spherical, ellipsoidal (thinner along the z-dimension), and particle-shaped masks, respectively. The fifth column displays the 3D projections of the enhanced and IsoNet missing-wedge-corrected particle. The sixth and seventh columns present the final 3D map and the flexibly fitted model, respectively. (B) Two cross-sectional views (12 nm thickness) of the plasmid density map along its plectoneme axis are shown in the left-middle panel. The intensity profile along the line crossing the two high-density DNA spots is displayed in the right panel. (C) Resolution assessment of the final 3D map using Fourier shell correlation (FSC). Two criteria are shown: FSC between two half-maps reconstructed from even and odd frames (evaluated at 0.143) and FSC between the final 3D map and the fitted model (evaluated at 0.5). (D) Zoomed-in views of the final 3D density map from panel A, displayed at two contour levels. (E) Superimposition of the high-contour level map from panel D onto its fitted model.

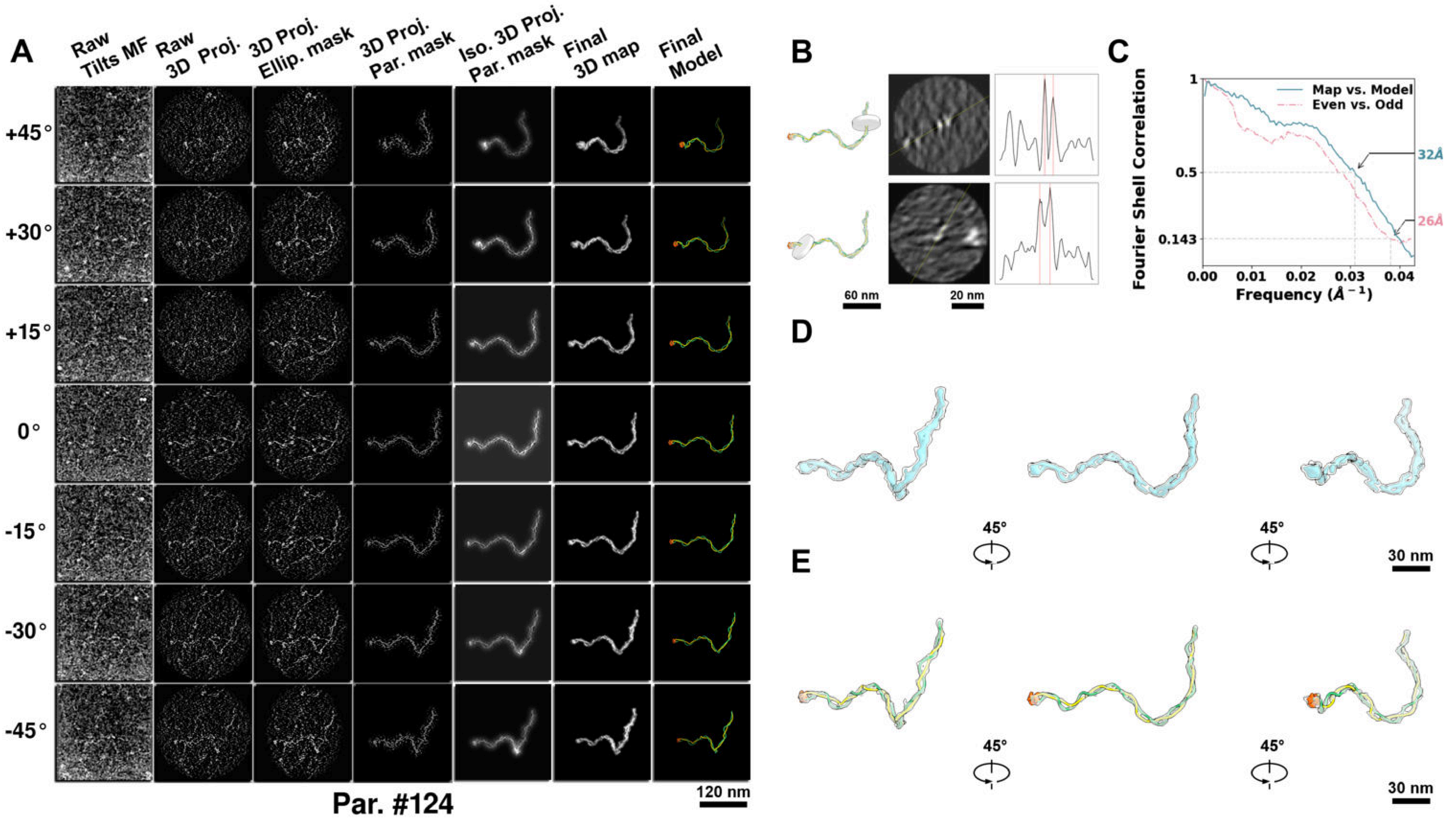

**Supplementary Particle Figure 124. Cryo-ET 3D reconstruction of an individual P.Cas particle.**
(A) 3D reconstruction of the plasmid particle (index no. 124). The first column shows seven representative tilt images from +45° to -45° in step of 15°. The second, third, and fourth columns show 3D projections of the particle with spherical, ellipsoidal (thinner along the z-dimension), and particle-shaped masks, respectively. The fifth column displays the 3D projections of the enhanced and IsoNet missing-wedge-corrected particle. The sixth and seventh columns present the final 3D map and the flexibly fitted model, respectively. (B) Two cross-sectional views (12 nm thickness) of the plasmid density map along its plectoneme axis are shown in the left-middle panel. The intensity profile along the line crossing the two high-density DNA spots is displayed in the right panel. (C) Resolution assessment of the final 3D map using Fourier shell correlation (FSC). Two criteria are shown: FSC between two half-maps reconstructed from even and odd frames (evaluated at 0.143) and FSC between the final 3D map and the fitted model (evaluated at 0.5). (D) Zoomed-in views of the final 3D density map from panel A, displayed at two contour levels. (E) Superimposition of the high-contour level map from panel D onto its fitted model.

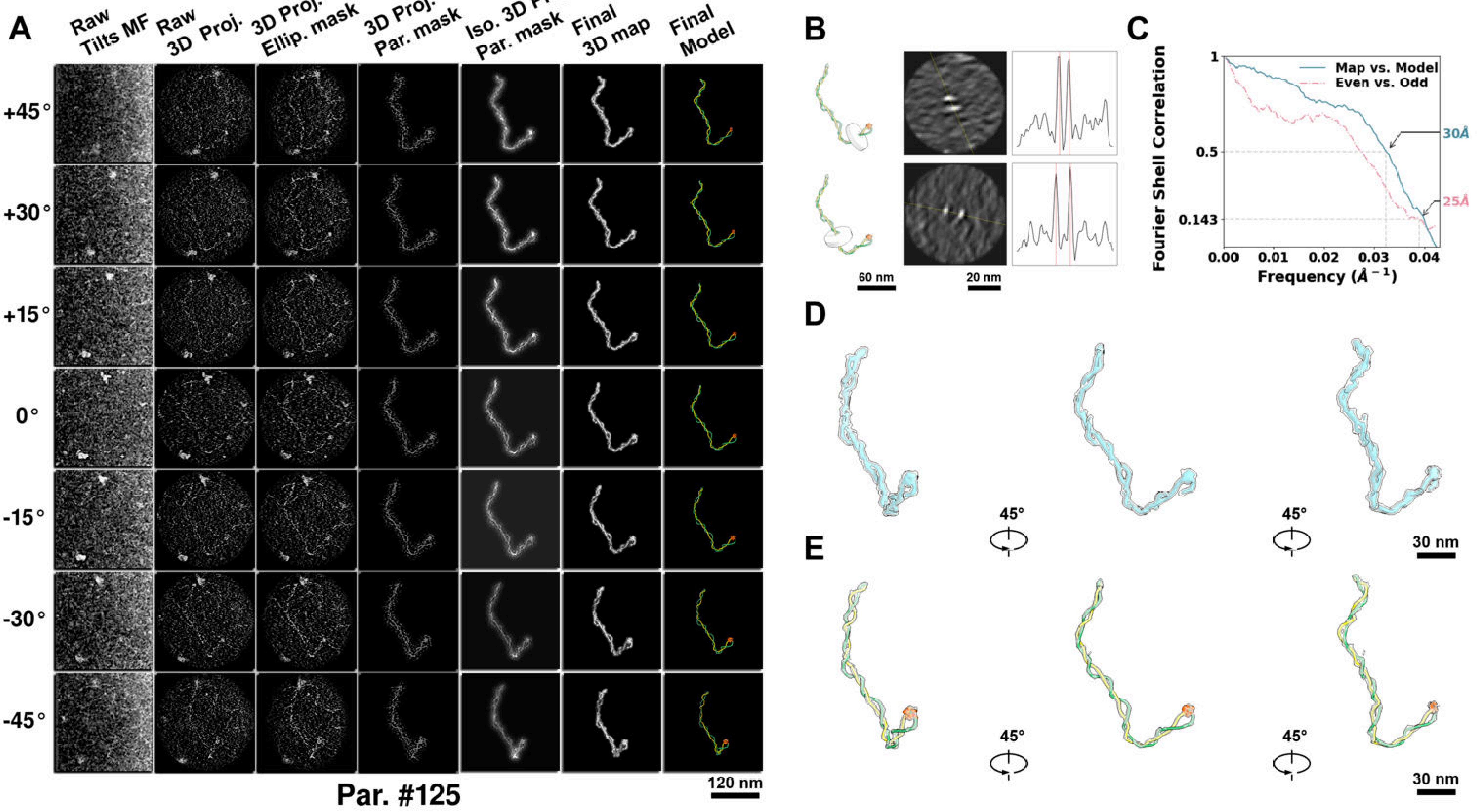

**Supplementary Particle Figure 125. Cryo-ET 3D reconstruction of an individual P.Cas particle.**
(A) 3D reconstruction of the plasmid particle (index no. 125). The first column shows seven representative tilt images from +45° to -45° in step of 15°. The second, third, and fourth columns show 3D projections of the particle with spherical, ellipsoidal (thinner along the z-dimension), and particle-shaped masks, respectively. The fifth column displays the 3D projections of the enhanced and IsoNet missing-wedge-corrected particle. The sixth and seventh columns present the final 3D map and the flexibly fitted model, respectively. (B) Two cross-sectional views (12 nm thickness) of the plasmid density map along its plectoneme axis are shown in the left-middle panel. The intensity profile along the line crossing the two high-density DNA spots is displayed in the right panel. (C) Resolution assessment of the final 3D map using Fourier shell correlation (FSC). Two criteria are shown: FSC between two half-maps reconstructed from even and odd frames (evaluated at 0.143) and FSC between the final 3D map and the fitted model (evaluated at 0.5). (D) Zoomed-in views of the final 3D density map from panel A, displayed at two contour levels. (E) Superimposition of the high-contour level map from panel D onto its fitted model.

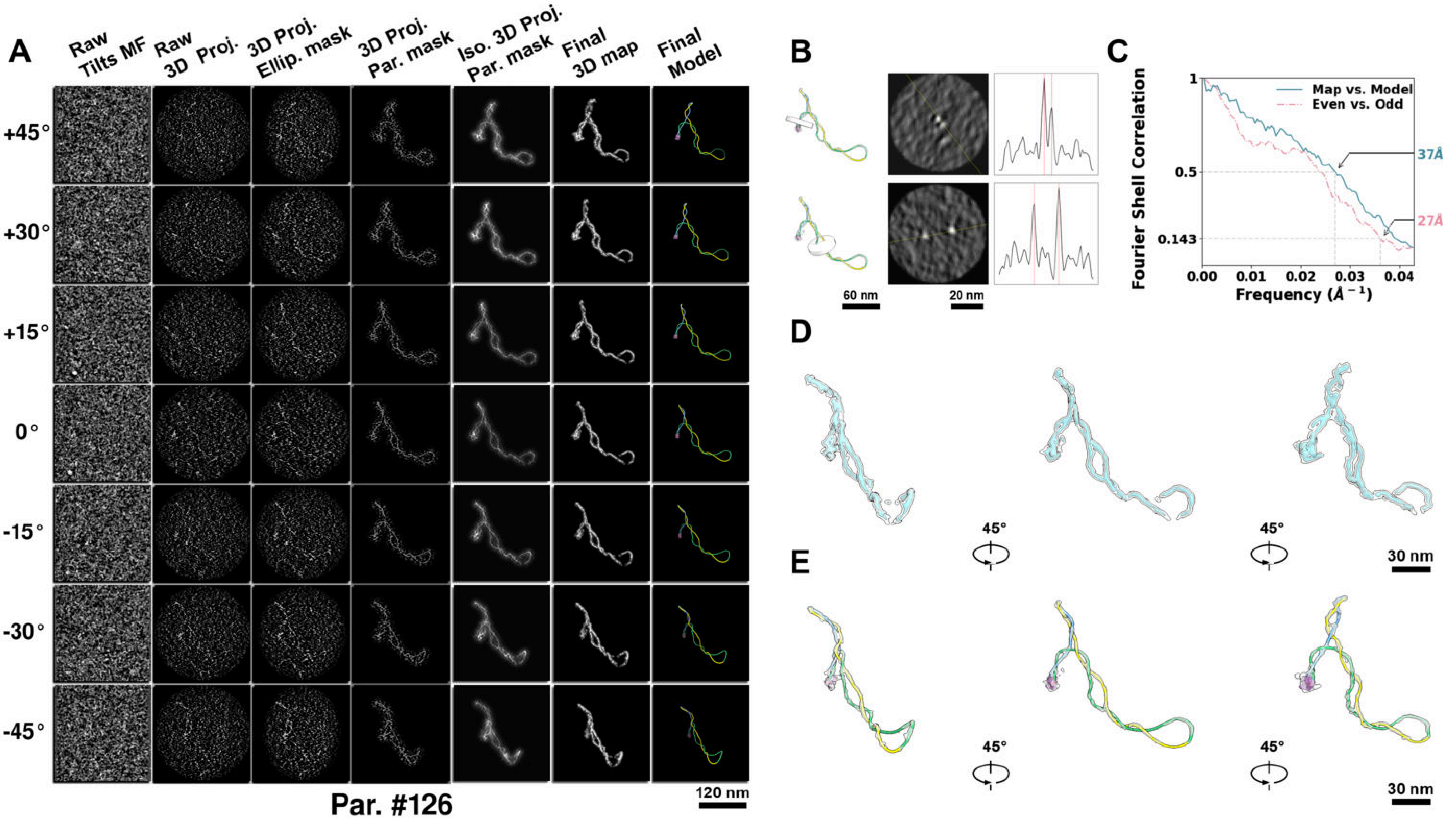

**Supplementary Particle Figure 126. Cryo-ET 3D reconstruction of an individual TEC particle.**
(A) 3D reconstruction of the plasmid particle (index no. 126). The first column shows seven representative tilt images from +45° to -45° in step of 15°. The second, third, and fourth columns show 3D projections of the particle with spherical, ellipsoidal (thinner along the z-dimension), and particle-shaped masks, respectively. The fifth column displays the 3D projections of the enhanced and IsoNet missing-wedge-corrected particle. The sixth and seventh columns present the final 3D map and the flexibly fitted model, respectively. (B) Two cross-sectional views (12 nm thickness) of the plasmid density map along its plectoneme axis are shown in the left-middle panel. The intensity profile along the line crossing the two high-density DNA spots is displayed in the right panel. (C) Resolution assessment of the final 3D map using Fourier shell correlation (FSC). Two criteria are shown: FSC between two half-maps reconstructed from even and odd frames (evaluated at 0.143) and FSC between the final 3D map and the fitted model (evaluated at 0.5). (D) Zoomed-in views of the final 3D density map from panel A, displayed at two contour levels. (E) Superimposition of the high-contour level map from panel D onto its fitted model.

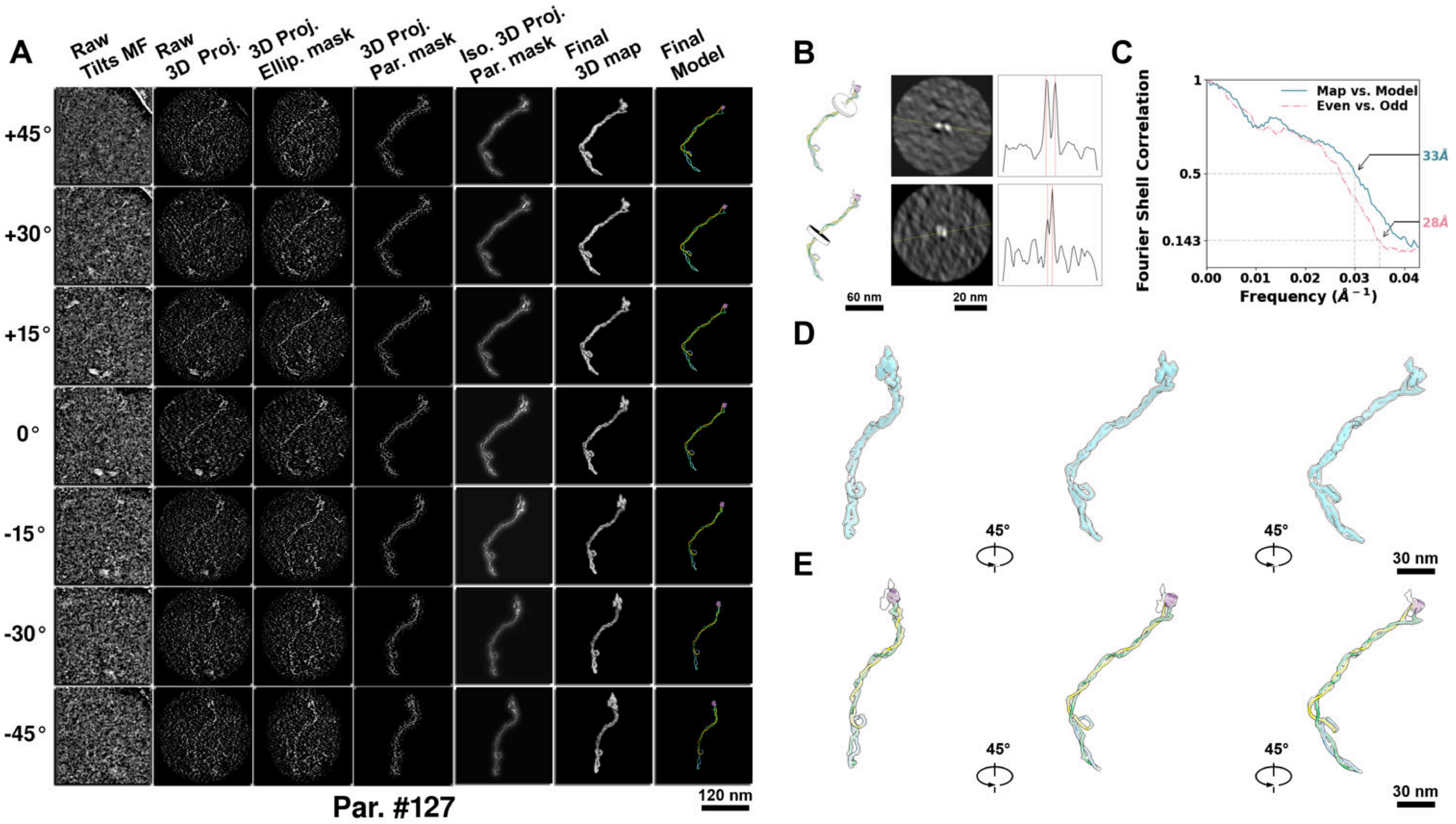

**Supplementary Particle Figure 127. Cryo-ET 3D reconstruction of an individual TEC particle.**
(A) 3D reconstruction of the plasmid particle (index no. 127). The first column shows seven representative tilt images from +45° to -45° in step of 15°. The second, third, and fourth columns show 3D projections of the particle with spherical, ellipsoidal (thinner along the z-dimension), and particle-shaped masks, respectively. The fifth column displays the 3D projections of the enhanced and IsoNet missing-wedge-corrected particle. The sixth and seventh columns present the final 3D map and the flexibly fitted model, respectively. (B) Two cross-sectional views (12 nm thickness) of the plasmid density map along its plectoneme axis are shown in the left-middle panel. The intensity profile along the line crossing the two high-density DNA spots is displayed in the right panel. (C) Resolution assessment of the final 3D map using Fourier shell correlation (FSC). Two criteria are shown: FSC between two half-maps reconstructed from even and odd frames (evaluated at 0.143) and FSC between the final 3D map and the fitted model (evaluated at 0.5). (D) Zoomed-in views of the final 3D density map from panel A, displayed at two contour levels. (E) Superimposition of the high-contour level map from panel D onto its fitted model.

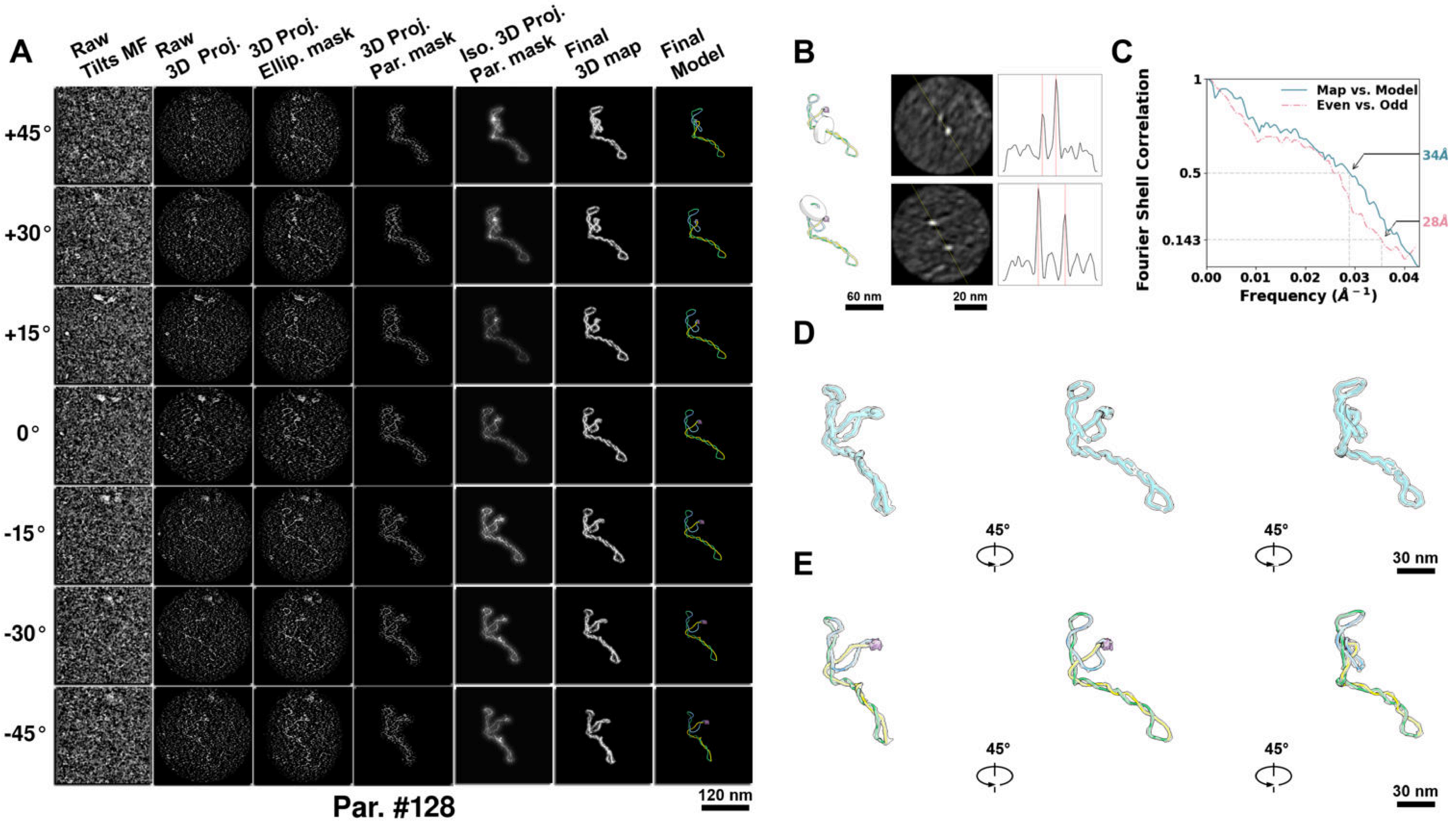

**Supplementary Particle Figure 128. Cryo-ET 3D reconstruction of an individual TEC particle.**
(A) 3D reconstruction of the plasmid particle (index no. 128). The first column shows seven representative tilt images from +45° to -45° in step of 15°. The second, third, and fourth columns show 3D projections of the particle with spherical, ellipsoidal (thinner along the z-dimension), and particle-shaped masks, respectively. The fifth column displays the 3D projections of the enhanced and IsoNet missing-wedge-corrected particle. The sixth and seventh columns present the final 3D map and the flexibly fitted model, respectively. (B) Two cross-sectional views (12 nm thickness) of the plasmid density map along its plectoneme axis are shown in the left-middle panel. The intensity profile along the line crossing the two high-density DNA spots is displayed in the right panel. (C) Resolution assessment of the final 3D map using Fourier shell correlation (FSC). Two criteria are shown: FSC between two half-maps reconstructed from even and odd frames (evaluated at 0.143) and FSC between the final 3D map and the fitted model (evaluated at 0.5). (D) Zoomed-in views of the final 3D density map from panel A, displayed at two contour levels. (E) Superimposition of the high-contour level map from panel D onto its fitted model.

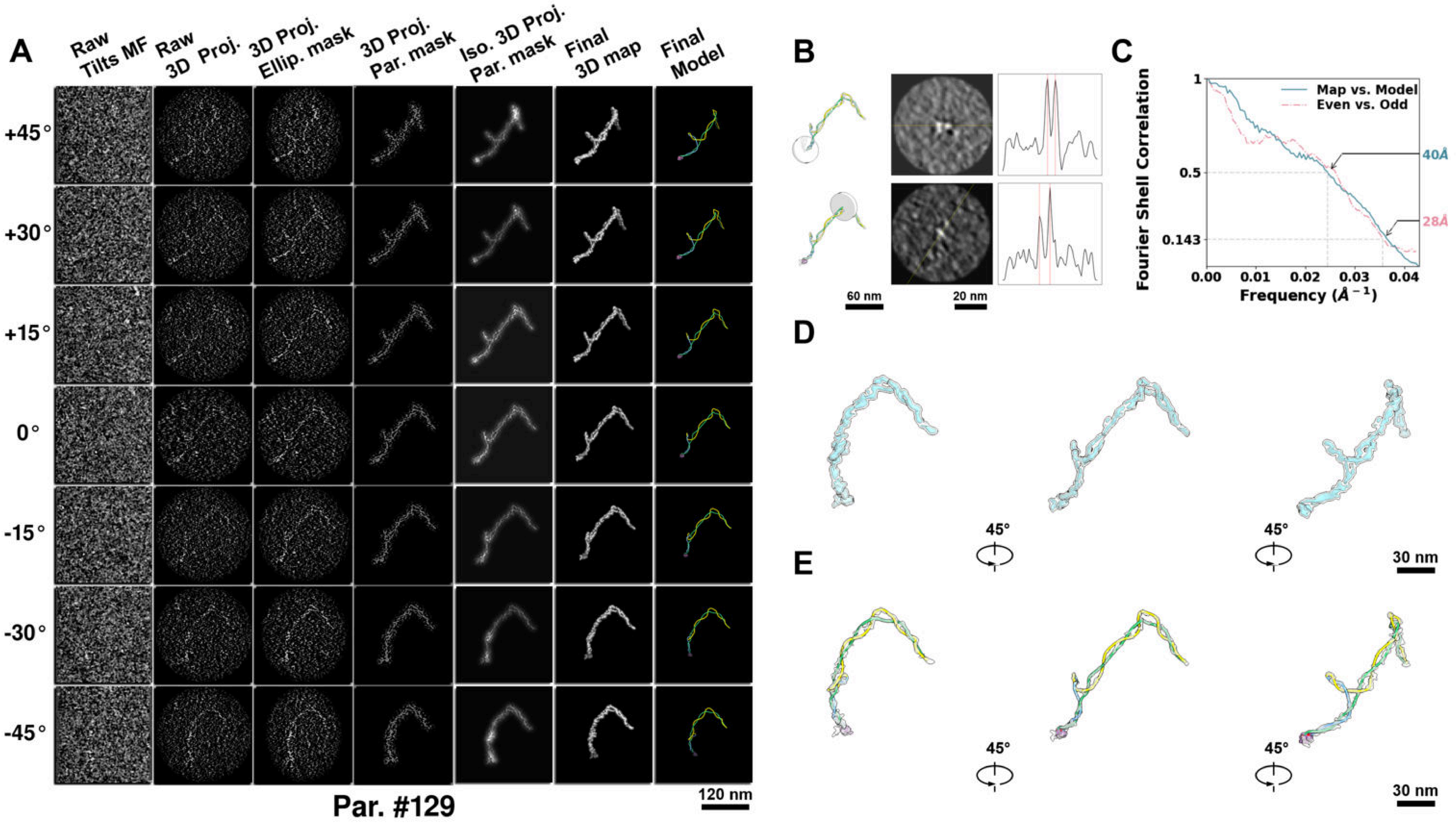

**Supplementary Particle Figure 129. Cryo-ET 3D reconstruction of an individual TEC particle.**
(A) 3D reconstruction of the plasmid particle (index no. 129). The first column shows seven representative tilt images from +45° to -45° in step of 15°. The second, third, and fourth columns show 3D projections of the particle with spherical, ellipsoidal (thinner along the z-dimension), and particle-shaped masks, respectively. The fifth column displays the 3D projections of the enhanced and IsoNet missing-wedge-corrected particle. The sixth and seventh columns present the final 3D map and the flexibly fitted model, respectively. (B) Two cross-sectional views (12 nm thickness) of the plasmid density map along its plectoneme axis are shown in the left-middle panel. The intensity profile along the line crossing the two high-density DNA spots is displayed in the right panel. (C) Resolution assessment of the final 3D map using Fourier shell correlation (FSC). Two criteria are shown: FSC between two half-maps reconstructed from even and odd frames (evaluated at 0.143) and FSC between the final 3D map and the fitted model (evaluated at 0.5). (D) Zoomed-in views of the final 3D density map from panel A, displayed at two contour levels. (E) Superimposition of the high-contour level map from panel D onto its fitted model.

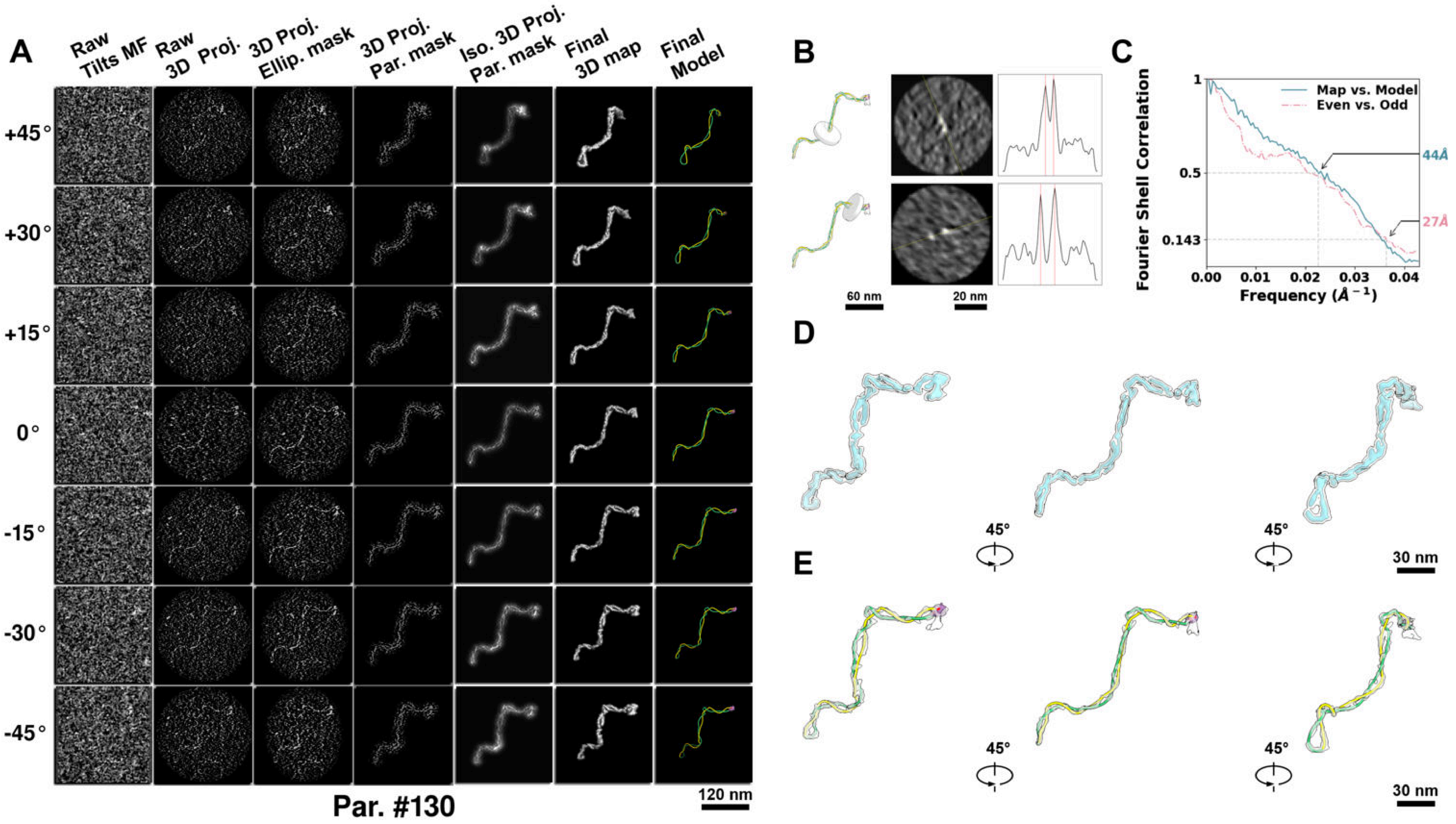

**Supplementary Particle Figure 130. Cryo-ET 3D reconstruction of an individual TEC particle.**
(A) 3D reconstruction of the plasmid particle (index no. 130). The first column shows seven representative tilt images from +45° to -45° in step of 15°. The second, third, and fourth columns show 3D projections of the particle with spherical, ellipsoidal (thinner along the z-dimension), and particle-shaped masks, respectively. The fifth column displays the 3D projections of the enhanced and IsoNet missing-wedge-corrected particle. The sixth and seventh columns present the final 3D map and the flexibly fitted model, respectively. (B) Two cross-sectional views (12 nm thickness) of the plasmid density map along its plectoneme axis are shown in the left-middle panel. The intensity profile along the line crossing the two high-density DNA spots is displayed in the right panel. (C) Resolution assessment of the final 3D map using Fourier shell correlation (FSC). Two criteria are shown: FSC between two half-maps reconstructed from even and odd frames (evaluated at 0.143) and FSC between the final 3D map and the fitted model (evaluated at 0.5). (D) Zoomed-in views of the final 3D density map from panel A, displayed at two contour levels. (E) Superimposition of the high-contour level map from panel D onto its fitted model.

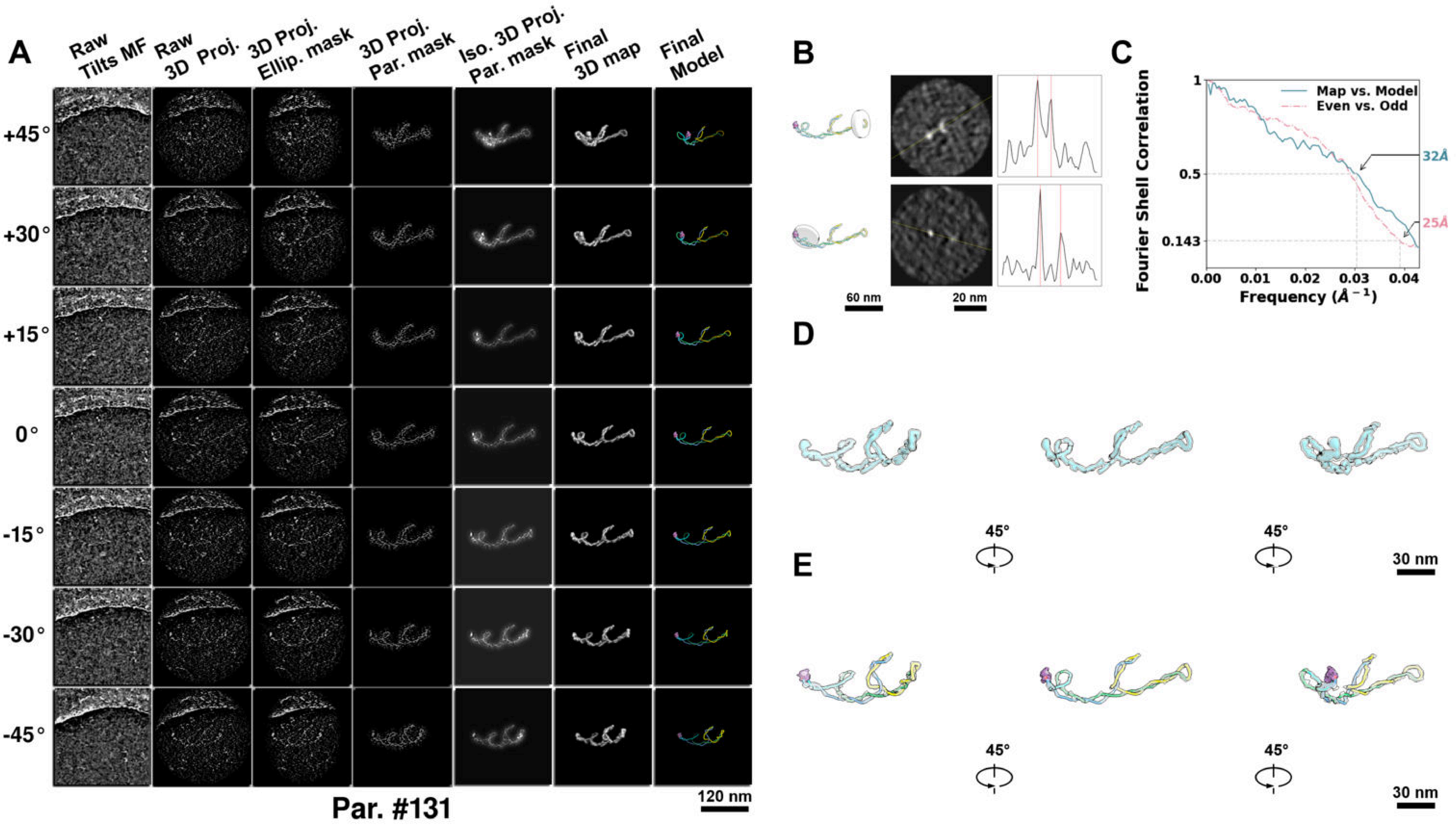

**Supplementary Particle Figure 131. Cryo-ET 3D reconstruction of an individual TEC particle.**
(A) 3D reconstruction of the plasmid particle (index no. 131). The first column shows seven representative tilt images from +45° to -45° in step of 15°. The second, third, and fourth columns show 3D projections of the particle with spherical, ellipsoidal (thinner along the z-dimension), and particle-shaped masks, respectively. The fifth column displays the 3D projections of the enhanced and IsoNet missing-wedge-corrected particle. The sixth and seventh columns present the final 3D map and the flexibly fitted model, respectively. (B) Two cross-sectional views (12 nm thickness) of the plasmid density map along its plectoneme axis are shown in the left-middle panel. The intensity profile along the line crossing the two high-density DNA spots is displayed in the right panel. (C) Resolution assessment of the final 3D map using Fourier shell correlation (FSC). Two criteria are shown: FSC between two half-maps reconstructed from even and odd frames (evaluated at 0.143) and FSC between the final 3D map and the fitted model (evaluated at 0.5). (D) Zoomed-in views of the final 3D density map from panel A, displayed at two contour levels. (E) Superimposition of the high-contour level map from panel D onto its fitted model.

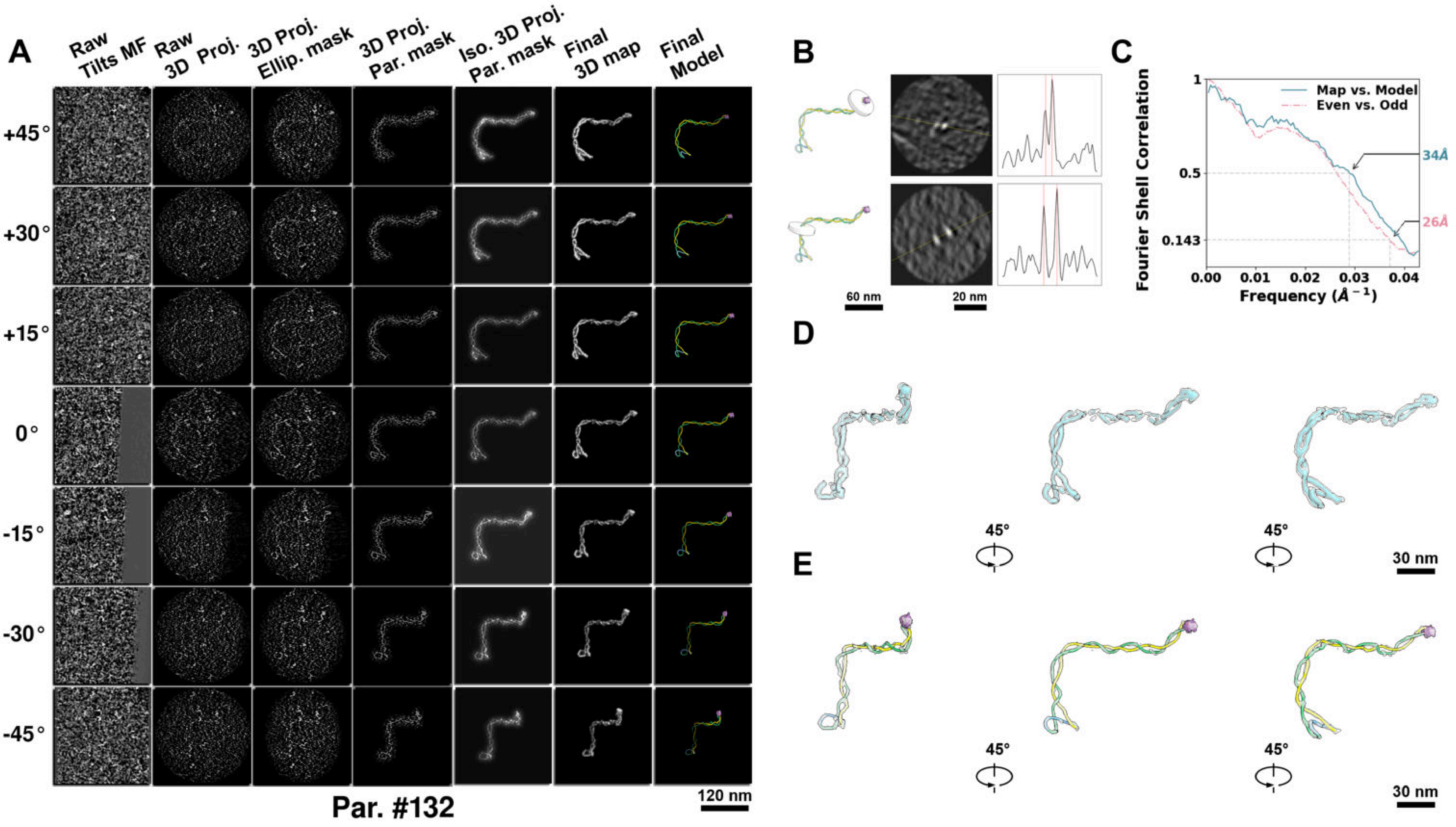

**Supplementary Particle Figure 132. Cryo-ET 3D reconstruction of an individual TEC particle.**
(A) 3D reconstruction of the plasmid particle (index no. 132). The first column shows seven representative tilt images from +45° to -45° in step of 15°. The second, third, and fourth columns show 3D projections of the particle with spherical, ellipsoidal (thinner along the z-dimension), and particle-shaped masks, respectively. The fifth column displays the 3D projections of the enhanced and IsoNet missing-wedge-corrected particle. The sixth and seventh columns present the final 3D map and the flexibly fitted model, respectively. (B) Two cross-sectional views (12 nm thickness) of the plasmid density map along its plectoneme axis are shown in the left-middle panel. The intensity profile along the line crossing the two high-density DNA spots is displayed in the right panel. (C) Resolution assessment of the final 3D map using Fourier shell correlation (FSC). Two criteria are shown: FSC between two half-maps reconstructed from even and odd frames (evaluated at 0.143) and FSC between the final 3D map and the fitted model (evaluated at 0.5). (D) Zoomed-in views of the final 3D density map from panel A, displayed at two contour levels. (E) Superimposition of the high-contour level map from panel D onto its fitted model.

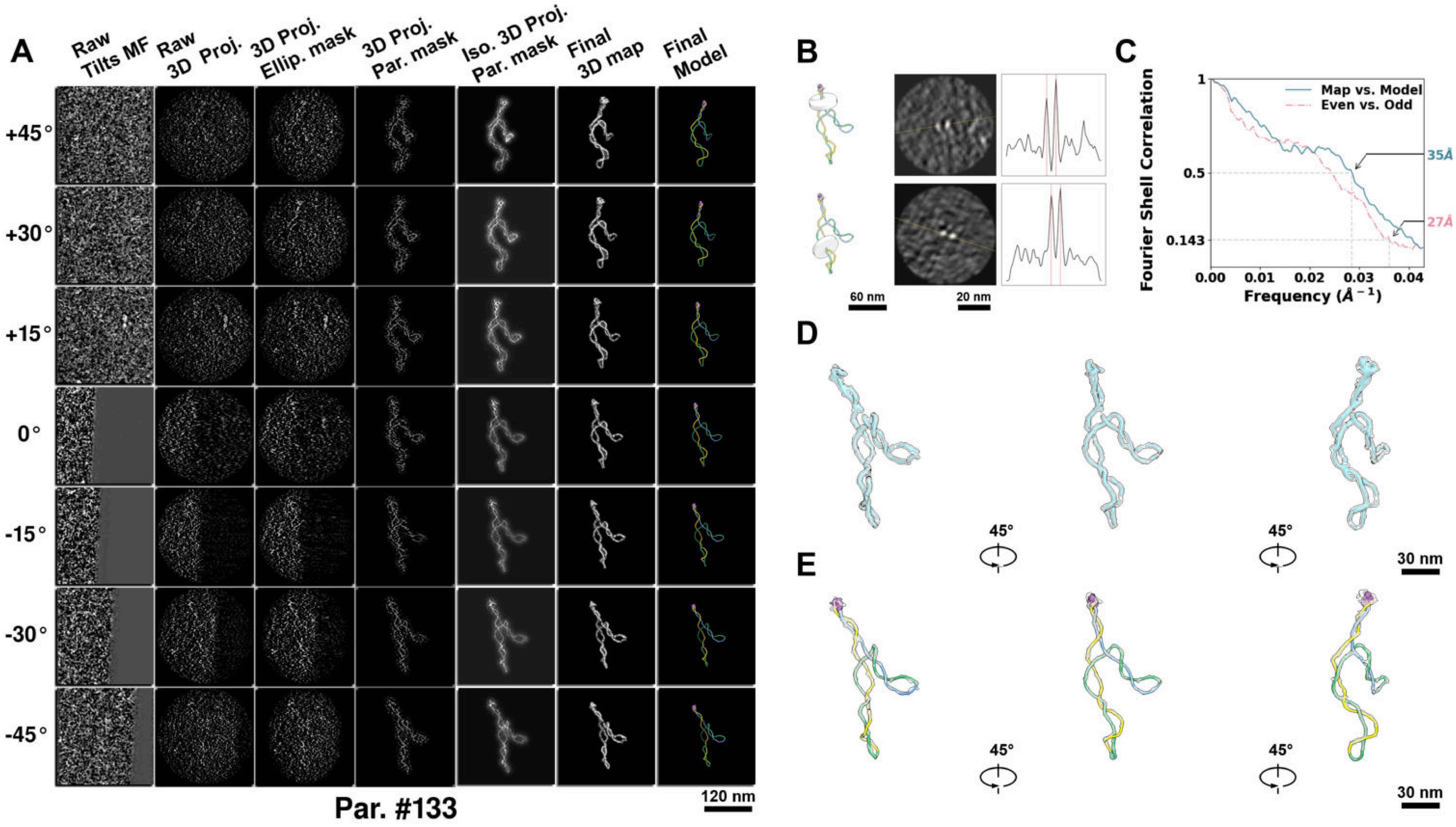

**Supplementary Particle Figure 133. Cryo-ET 3D reconstruction of an individual TEC particle.**
(A) 3D reconstruction of the plasmid particle (index no. 133). The first column shows seven representative tilt images from +45° to -45° in step of 15°. The second, third, and fourth columns show 3D projections of the particle with spherical, ellipsoidal (thinner along the z-dimension), and particle-shaped masks, respectively. The fifth column displays the 3D projections of the enhanced and IsoNet missing-wedge-corrected particle. The sixth and seventh columns present the final 3D map and the flexibly fitted model, respectively. (B) Two cross-sectional views (12 nm thickness) of the plasmid density map along its plectoneme axis are shown in the left-middle panel. The intensity profile along the line crossing the two high-density DNA spots is displayed in the right panel. (C) Resolution assessment of the final 3D map using Fourier shell correlation (FSC). Two criteria are shown: FSC between two half-maps reconstructed from even and odd frames (evaluated at 0.143) and FSC between the final 3D map and the fitted model (evaluated at 0.5). (D) Zoomed-in views of the final 3D density map from panel A, displayed at two contour levels. (E) Superimposition of the high-contour level map from panel D onto its fitted model.

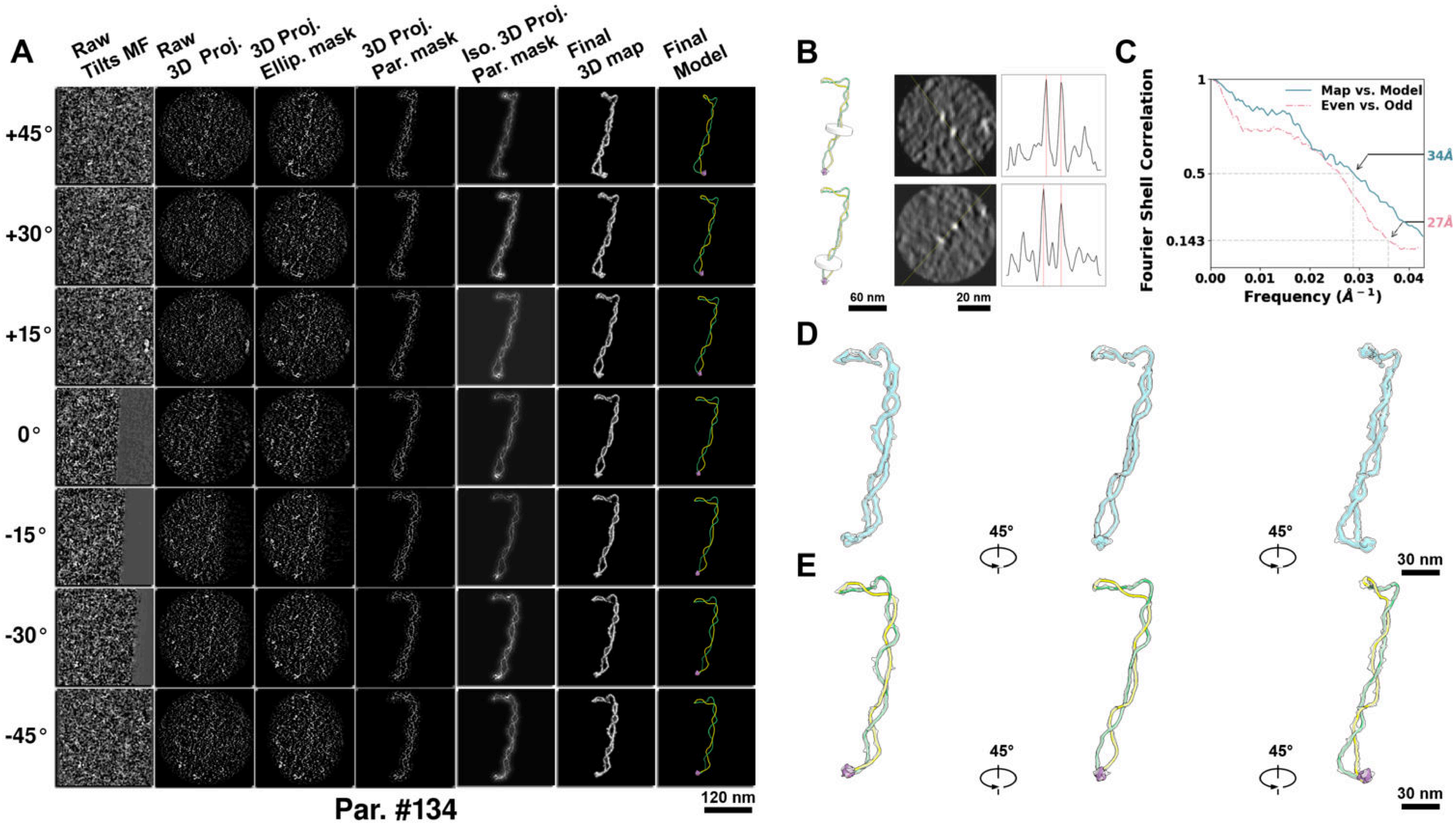

**Supplementary Particle Figure 134. Cryo-ET 3D reconstruction of an individual TEC particle.**
(A) 3D reconstruction of the plasmid particle (index no. 134). The first column shows seven representative tilt images from +45° to -45° in step of 15°. The second, third, and fourth columns show 3D projections of the particle with spherical, ellipsoidal (thinner along the z-dimension), and particle-shaped masks, respectively. The fifth column displays the 3D projections of the enhanced and IsoNet missing-wedge-corrected particle. The sixth and seventh columns present the final 3D map and the flexibly fitted model, respectively. (B) Two cross-sectional views (12 nm thickness) of the plasmid density map along its plectoneme axis are shown in the left-middle panel. The intensity profile along the line crossing the two high-density DNA spots is displayed in the right panel. (C) Resolution assessment of the final 3D map using Fourier shell correlation (FSC). Two criteria are shown: FSC between two half-maps reconstructed from even and odd frames (evaluated at 0.143) and FSC between the final 3D map and the fitted model (evaluated at 0.5). (D) Zoomed-in views of the final 3D density map from panel A, displayed at two contour levels. (E) Superimposition of the high-contour level map from panel D onto its fitted model.

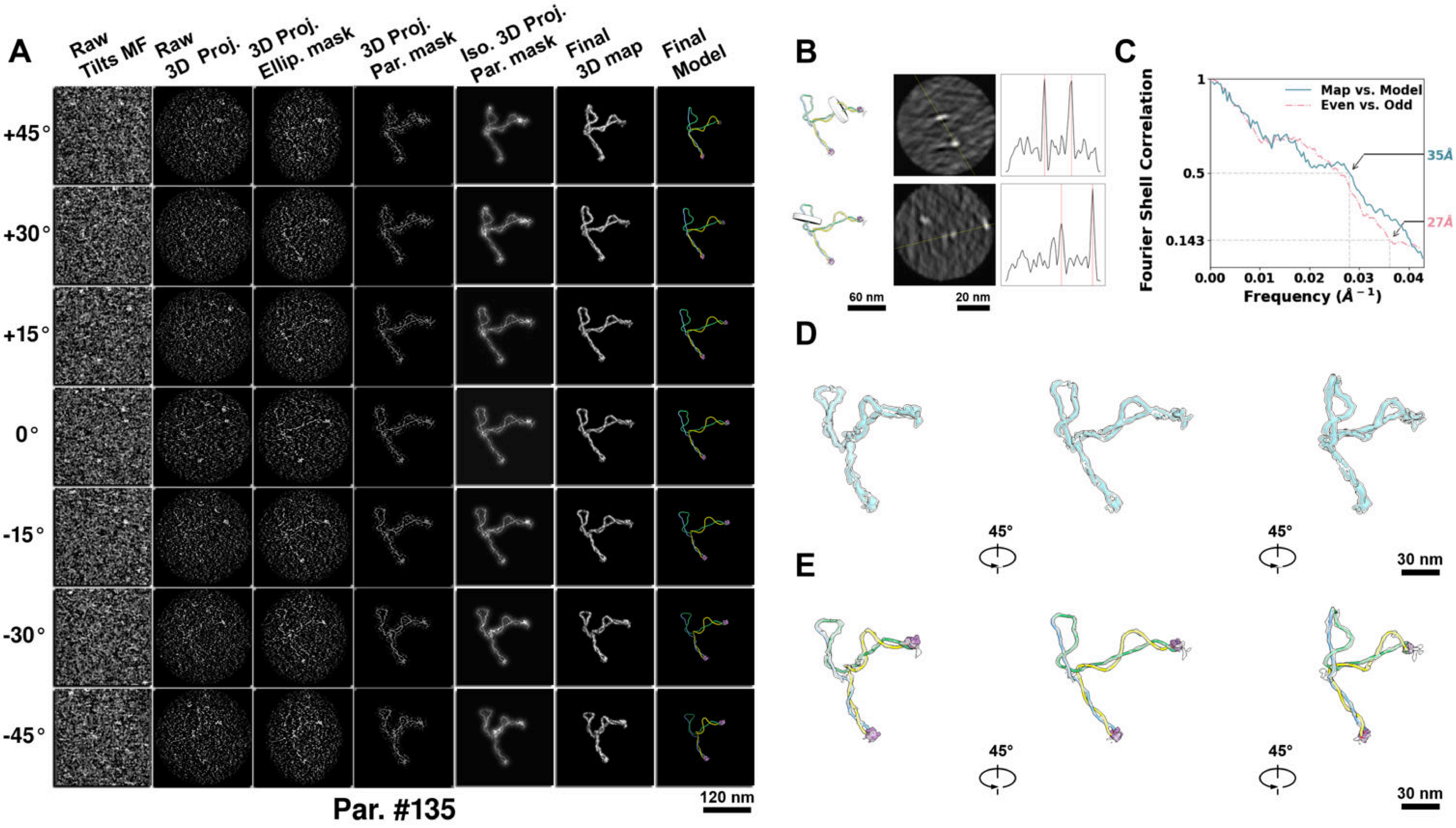

**Supplementary Particle Figure 135. Cryo-ET 3D reconstruction of an individual TEC particle.**
(A) 3D reconstruction of the plasmid particle (index no. 135). The first column shows seven representative tilt images from +45° to -45° in step of 15°. The second, third, and fourth columns show 3D projections of the particle with spherical, ellipsoidal (thinner along the z-dimension), and particle-shaped masks, respectively. The fifth column displays the 3D projections of the enhanced and IsoNet missing-wedge-corrected particle. The sixth and seventh columns present the final 3D map and the flexibly fitted model, respectively. (B) Two cross-sectional views (12 nm thickness) of the plasmid density map along its plectoneme axis are shown in the left-middle panel. The intensity profile along the line crossing the two high-density DNA spots is displayed in the right panel. (C) Resolution assessment of the final 3D map using Fourier shell correlation (FSC). Two criteria are shown: FSC between two half-maps reconstructed from even and odd frames (evaluated at 0.143) and FSC between the final 3D map and the fitted model (evaluated at 0.5). (D) Zoomed-in views of the final 3D density map from panel A, displayed at two contour levels. (E) Superimposition of the high-contour level map from panel D onto its fitted model.

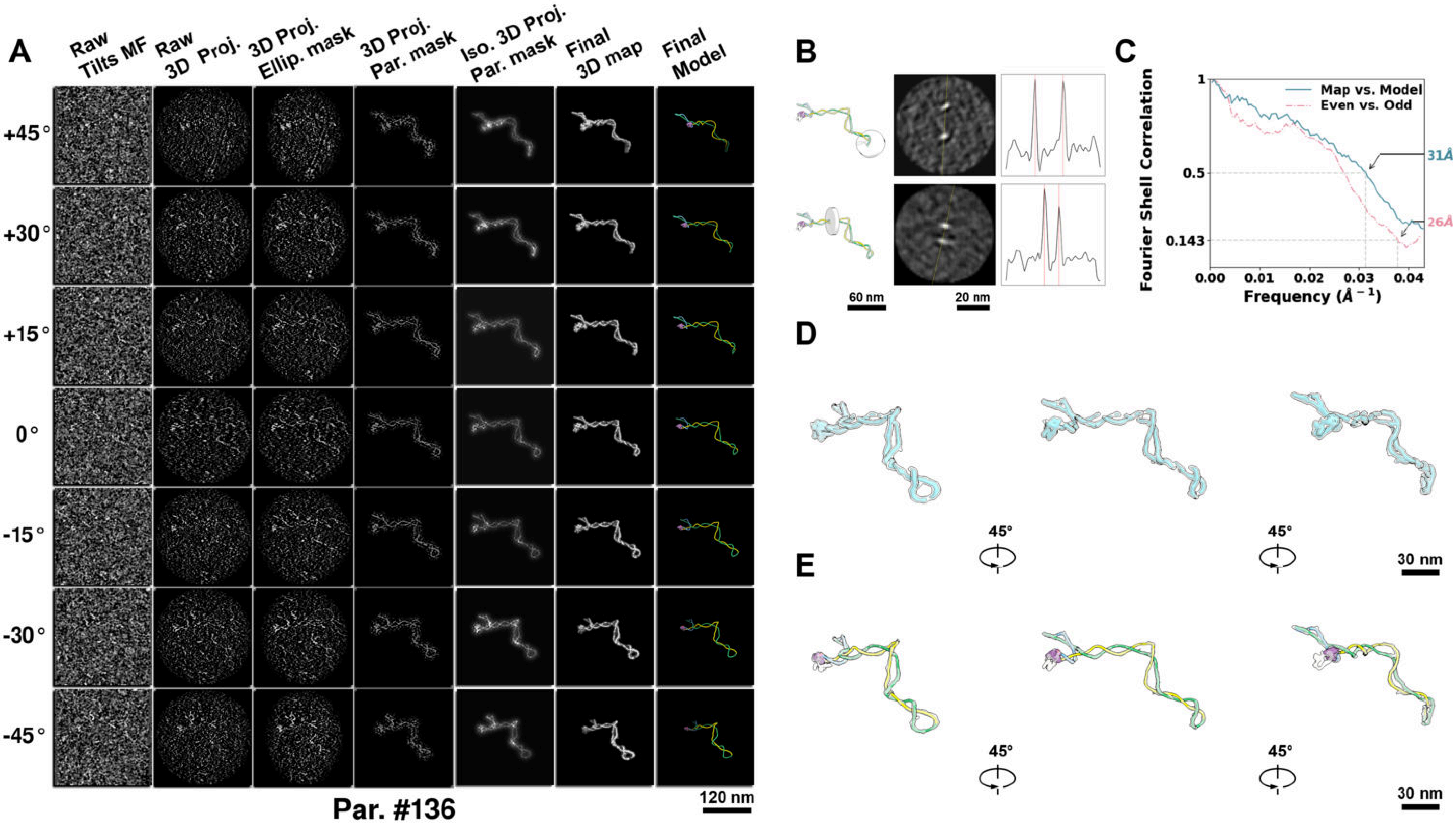

**Supplementary Particle Figure 136. Cryo-ET 3D reconstruction of an individual TEC particle.**
(A) 3D reconstruction of the plasmid particle (index no. 136). The first column shows seven representative tilt images from +45° to -45° in step of 15°. The second, third, and fourth columns show 3D projections of the particle with spherical, ellipsoidal (thinner along the z-dimension), and particle-shaped masks, respectively. The fifth column displays the 3D projections of the enhanced and IsoNet missing-wedge-corrected particle. The sixth and seventh columns present the final 3D map and the flexibly fitted model, respectively. (B) Two cross-sectional views (12 nm thickness) of the plasmid density map along its plectoneme axis are shown in the left-middle panel. The intensity profile along the line crossing the two high-density DNA spots is displayed in the right panel. (C) Resolution assessment of the final 3D map using Fourier shell correlation (FSC). Two criteria are shown: FSC between two half-maps reconstructed from even and odd frames (evaluated at 0.143) and FSC between the final 3D map and the fitted model (evaluated at 0.5). (D) Zoomed-in views of the final 3D density map from panel A, displayed at two contour levels. (E) Superimposition of the high-contour level map from panel D onto its fitted model.

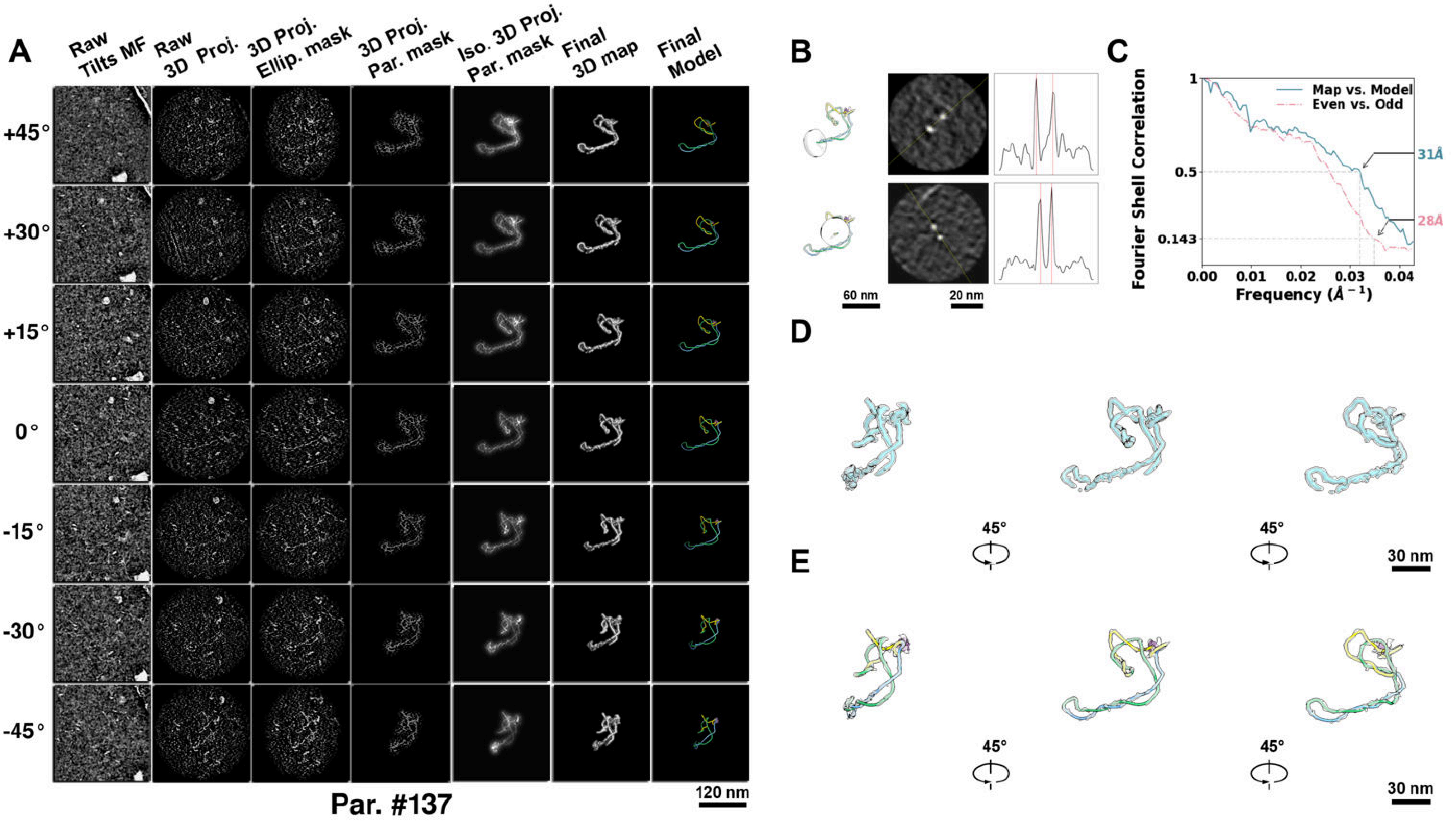

**Supplementary Particle Figure 137. Cryo-ET 3D reconstruction of an individual TEC particle.**
(A) 3D reconstruction of the plasmid particle (index no. 137). The first column shows seven representative tilt images from +45° to -45° in step of 15°. The second, third, and fourth columns show 3D projections of the particle with spherical, ellipsoidal (thinner along the z-dimension), and particle-shaped masks, respectively. The fifth column displays the 3D projections of the enhanced and IsoNet missing-wedge-corrected particle. The sixth and seventh columns present the final 3D map and the flexibly fitted model, respectively. (B) Two cross-sectional views (12 nm thickness) of the plasmid density map along its plectoneme axis are shown in the left-middle panel. The intensity profile along the line crossing the two high-density DNA spots is displayed in the right panel. (C) Resolution assessment of the final 3D map using Fourier shell correlation (FSC). Two criteria are shown: FSC between two half-maps reconstructed from even and odd frames (evaluated at 0.143) and FSC between the final 3D map and the fitted model (evaluated at 0.5). (D) Zoomed-in views of the final 3D density map from panel A, displayed at two contour levels. (E) Superimposition of the high-contour level map from panel D onto its fitted model.

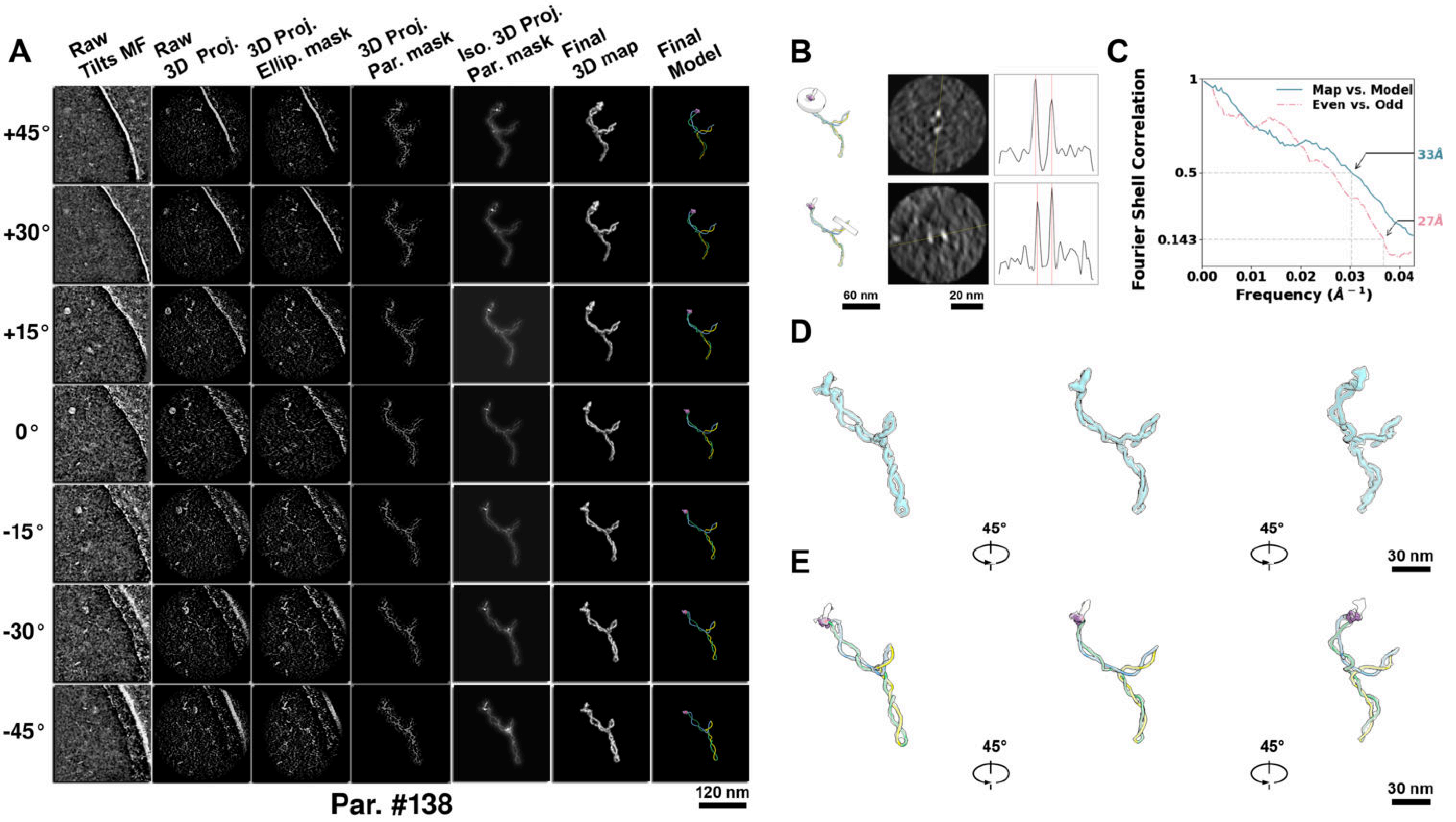

**Supplementary Particle Figure 138. Cryo-ET 3D reconstruction of an individual TEC particle.**
(A) 3D reconstruction of the plasmid particle (index no. 138). The first column shows seven representative tilt images from +45° to -45° in step of 15°. The second, third, and fourth columns show 3D projections of the particle with spherical, ellipsoidal (thinner along the z-dimension), and particle-shaped masks, respectively. The fifth column displays the 3D projections of the enhanced and IsoNet missing-wedge-corrected particle. The sixth and seventh columns present the final 3D map and the flexibly fitted model, respectively. (B) Two cross-sectional views (12 nm thickness) of the plasmid density map along its plectoneme axis are shown in the left-middle panel. The intensity profile along the line crossing the two high-density DNA spots is displayed in the right panel. (C) Resolution assessment of the final 3D map using Fourier shell correlation (FSC). Two criteria are shown: FSC between two half-maps reconstructed from even and odd frames (evaluated at 0.143) and FSC between the final 3D map and the fitted model (evaluated at 0.5). (D) Zoomed-in views of the final 3D density map from panel A, displayed at two contour levels. (E) Superimposition of the high-contour level map from panel D onto its fitted model.

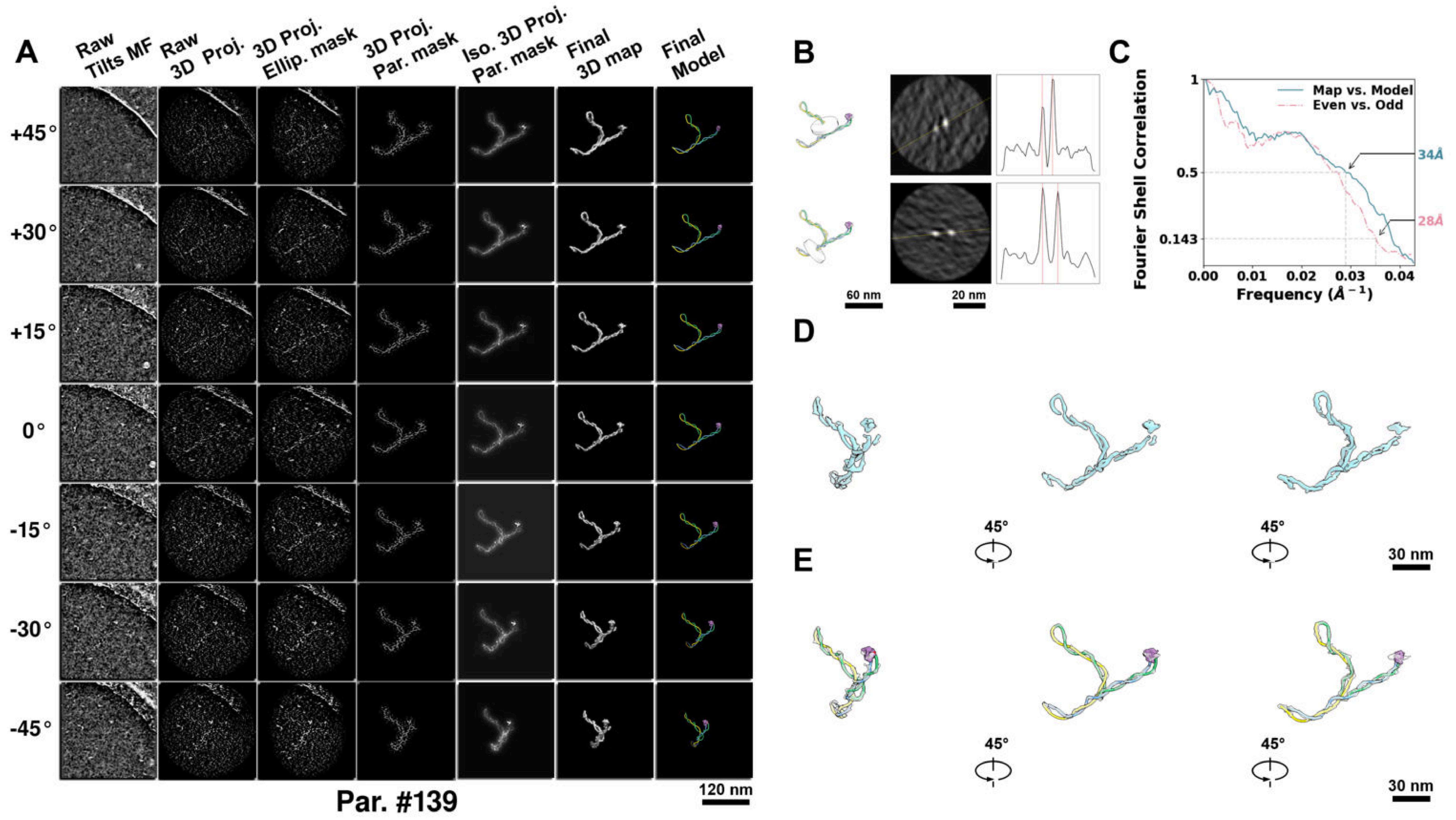

**Supplementary Particle Figure 139. Cryo-ET 3D reconstruction of an individual TEC particle.**
(A) 3D reconstruction of the plasmid particle (index no. 139). The first column shows seven representative tilt images from +45° to -45° in step of 15°. The second, third, and fourth columns show 3D projections of the particle with spherical, ellipsoidal (thinner along the z-dimension), and particle-shaped masks, respectively. The fifth column displays the 3D projections of the enhanced and IsoNet missing-wedge-corrected particle. The sixth and seventh columns present the final 3D map and the flexibly fitted model, respectively. (B) Two cross-sectional views (12 nm thickness) of the plasmid density map along its plectoneme axis are shown in the left-middle panel. The intensity profile along the line crossing the two high-density DNA spots is displayed in the right panel. (C) Resolution assessment of the final 3D map using Fourier shell correlation (FSC). Two criteria are shown: FSC between two half-maps reconstructed from even and odd frames (evaluated at 0.143) and FSC between the final 3D map and the fitted model (evaluated at 0.5). (D) Zoomed-in views of the final 3D density map from panel A, displayed at two contour levels. (E) Superimposition of the high-contour level map from panel D onto its fitted model.

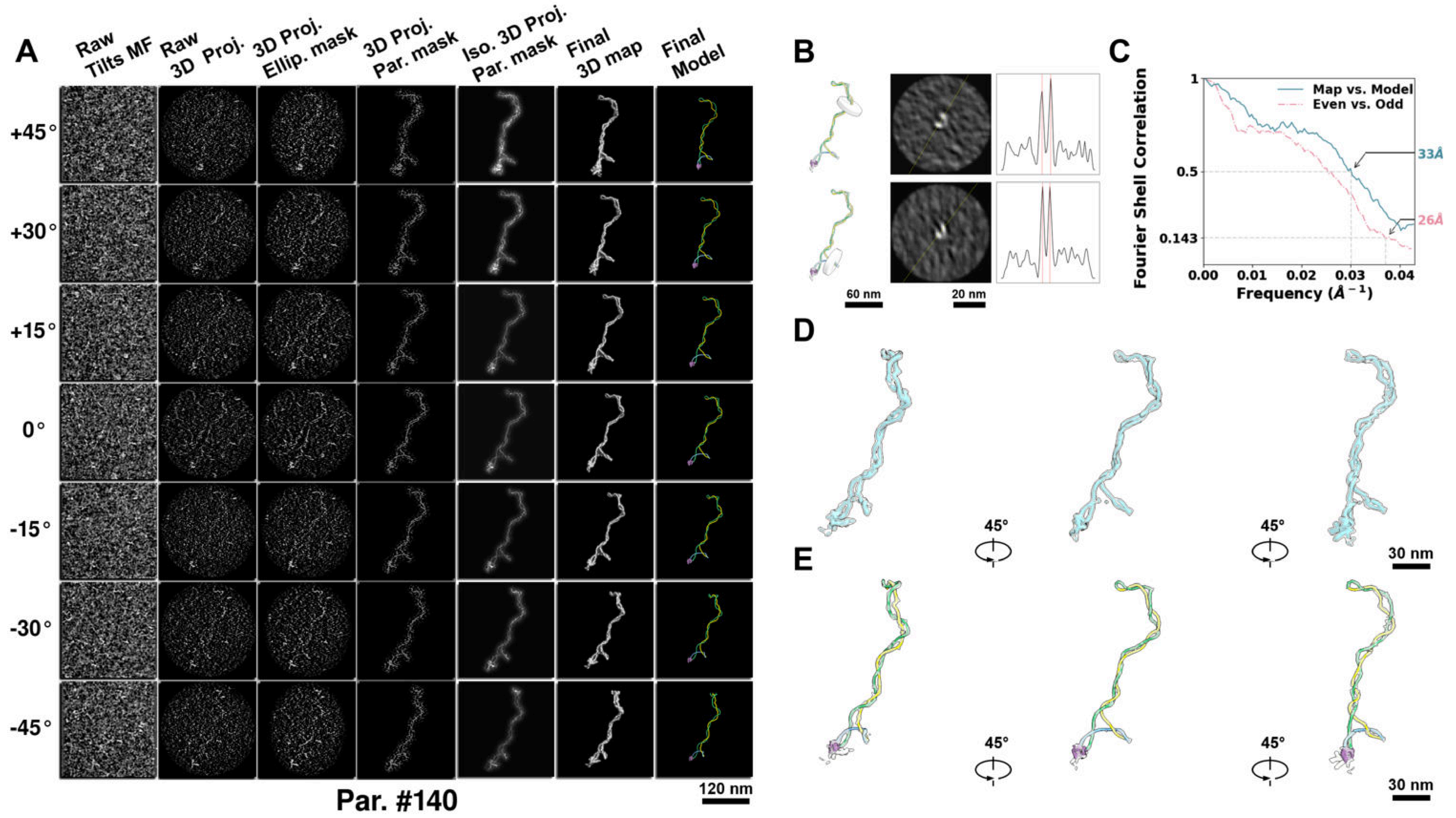

**Supplementary Particle Figure 140. Cryo-ET 3D reconstruction of an individual TEC particle.**
(A) 3D reconstruction of the plasmid particle (index no. 140). The first column shows seven representative tilt images from +45° to -45° in step of 15°. The second, third, and fourth columns show 3D projections of the particle with spherical, ellipsoidal (thinner along the z-dimension), and particle-shaped masks, respectively. The fifth column displays the 3D projections of the enhanced and IsoNet missing-wedge-corrected particle. The sixth and seventh columns present the final 3D map and the flexibly fitted model, respectively. (B) Two cross-sectional views (12 nm thickness) of the plasmid density map along its plectoneme axis are shown in the left-middle panel. The intensity profile along the line crossing the two high-density DNA spots is displayed in the right panel. (C) Resolution assessment of the final 3D map using Fourier shell correlation (FSC). Two criteria are shown: FSC between two half-maps reconstructed from even and odd frames (evaluated at 0.143) and FSC between the final 3D map and the fitted model (evaluated at 0.5). (D) Zoomed-in views of the final 3D density map from panel A, displayed at two contour levels. (E) Superimposition of the high-contour level map from panel D onto its fitted model.

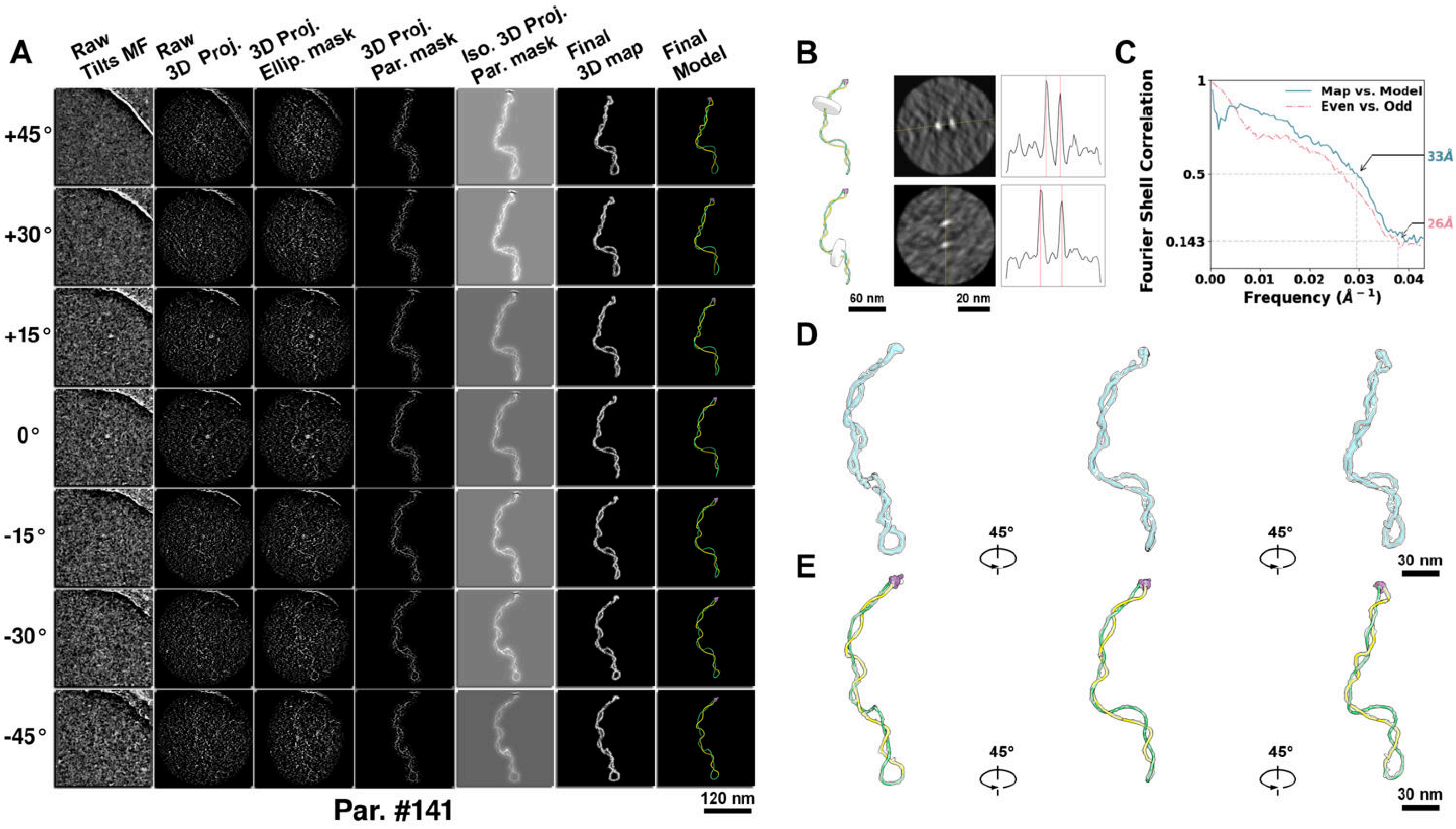

**Supplementary Particle Figure 141. Cryo-ET 3D reconstruction of an individual TEC particle.**
(A) 3D reconstruction of the plasmid particle (index no. 141). The first column shows seven representative tilt images from +45° to -45° in step of 15°. The second, third, and fourth columns show 3D projections of the particle with spherical, ellipsoidal (thinner along the z-dimension), and particle-shaped masks, respectively. The fifth column displays the 3D projections of the enhanced and IsoNet missing-wedge-corrected particle. The sixth and seventh columns present the final 3D map and the flexibly fitted model, respectively. (B) Two cross-sectional views (12 nm thickness) of the plasmid density map along its plectoneme axis are shown in the left-middle panel. The intensity profile along the line crossing the two high-density DNA spots is displayed in the right panel. (C) Resolution assessment of the final 3D map using Fourier shell correlation (FSC). Two criteria are shown: FSC between two half-maps reconstructed from even and odd frames (evaluated at 0.143) and FSC between the final 3D map and the fitted model (evaluated at 0.5). (D) Zoomed-in views of the final 3D density map from panel A, displayed at two contour levels. (E) Superimposition of the high-contour level map from panel D onto its fitted model.

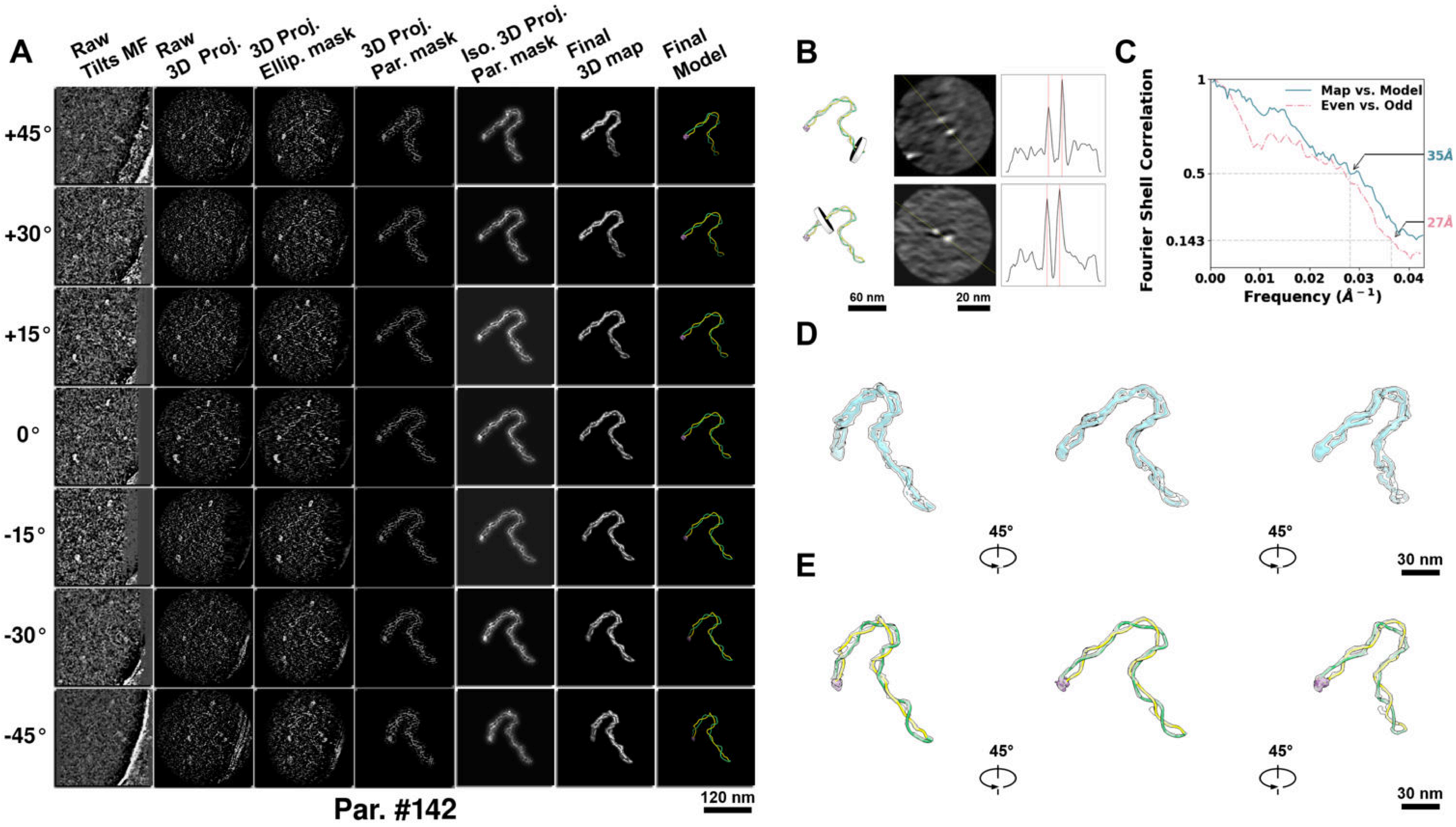

**Supplementary Particle Figure 142. Cryo-ET 3D reconstruction of an individual TEC particle.**
(A) 3D reconstruction of the plasmid particle (index no. 142). The first column shows seven representative tilt images from +45° to -45° in step of 15°. The second, third, and fourth columns show 3D projections of the particle with spherical, ellipsoidal (thinner along the z-dimension), and particle-shaped masks, respectively. The fifth column displays the 3D projections of the enhanced and IsoNet missing-wedge-corrected particle. The sixth and seventh columns present the final 3D map and the flexibly fitted model, respectively. (B) Two cross-sectional views (12 nm thickness) of the plasmid density map along its plectoneme axis are shown in the left-middle panel. The intensity profile along the line crossing the two high-density DNA spots is displayed in the right panel. (C) Resolution assessment of the final 3D map using Fourier shell correlation (FSC). Two criteria are shown: FSC between two half-maps reconstructed from even and odd frames (evaluated at 0.143) and FSC between the final 3D map and the fitted model (evaluated at 0.5). (D) Zoomed-in views of the final 3D density map from panel A, displayed at two contour levels. (E) Superimposition of the high-contour level map from panel D onto its fitted model.

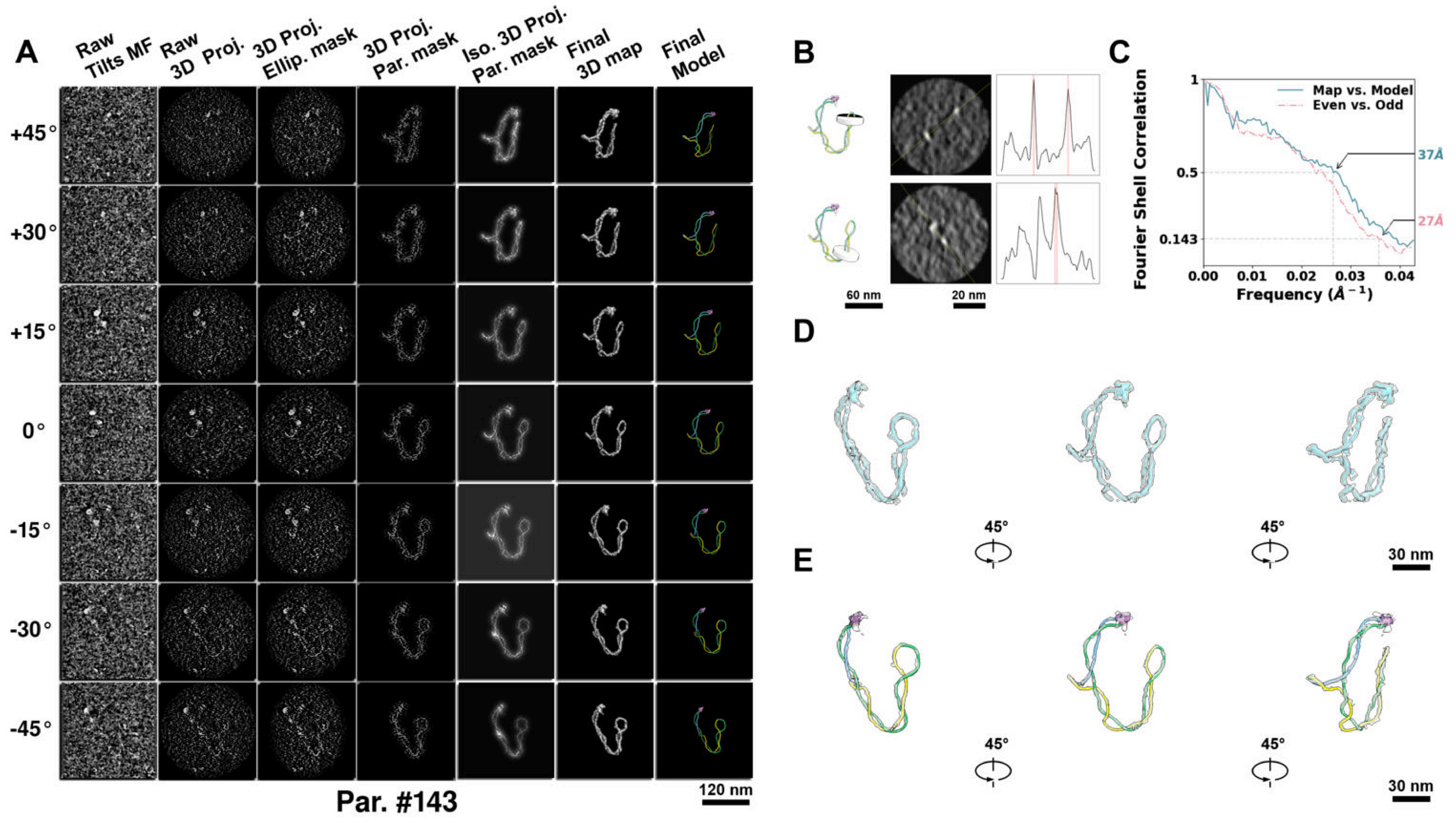

**Supplementary Particle Figure 143. Cryo-ET 3D reconstruction of an individual TEC particle.**
(A) 3D reconstruction of the plasmid particle (index no. 143). The first column shows seven representative tilt images from +45° to -45° in step of 15°. The second, third, and fourth columns show 3D projections of the particle with spherical, ellipsoidal (thinner along the z-dimension), and particle-shaped masks, respectively. The fifth column displays the 3D projections of the enhanced and IsoNet missing-wedge-corrected particle. The sixth and seventh columns present the final 3D map and the flexibly fitted model, respectively. (B) Two cross-sectional views (12 nm thickness) of the plasmid density map along its plectoneme axis are shown in the left-middle panel. The intensity profile along the line crossing the two high-density DNA spots is displayed in the right panel. (C) Resolution assessment of the final 3D map using Fourier shell correlation (FSC). Two criteria are shown: FSC between two half-maps reconstructed from even and odd frames (evaluated at 0.143) and FSC between the final 3D map and the fitted model (evaluated at 0.5). (D) Zoomed-in views of the final 3D density map from panel A, displayed at two contour levels. (E) Superimposition of the high-contour level map from panel D onto its fitted model.

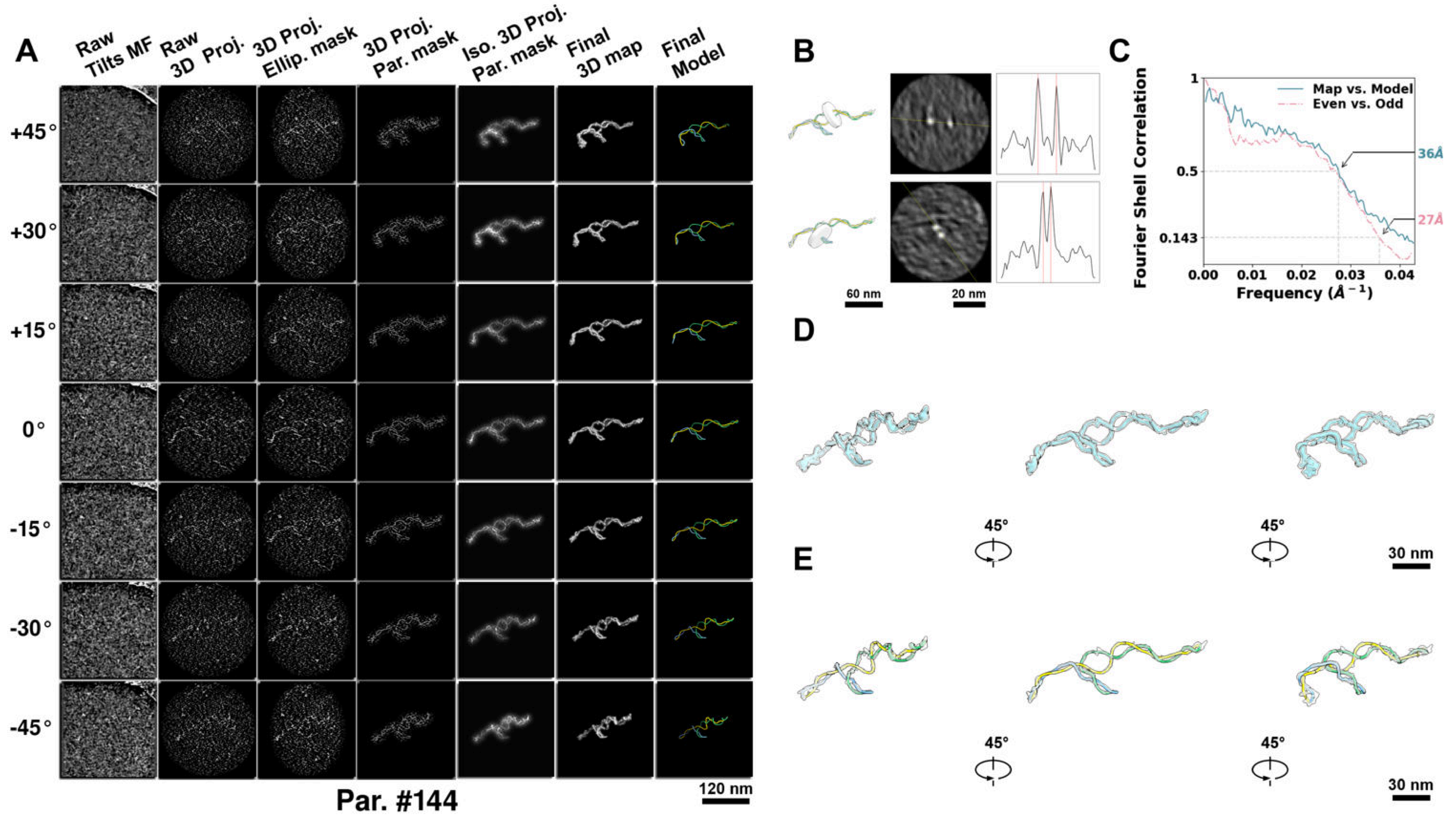

**Supplementary Particle Figure 144. Cryo-ET 3D reconstruction of an individual TEC particle.**
(A) 3D reconstruction of the plasmid particle (index no. 144). The first column shows seven representative tilt images from +45° to -45° in step of 15°. The second, third, and fourth columns show 3D projections of the particle with spherical, ellipsoidal (thinner along the z-dimension), and particle-shaped masks, respectively. The fifth column displays the 3D projections of the enhanced and IsoNet missing-wedge-corrected particle. The sixth and seventh columns present the final 3D map and the flexibly fitted model, respectively. (B) Two cross-sectional views (12 nm thickness) of the plasmid density map along its plectoneme axis are shown in the left-middle panel. The intensity profile along the line crossing the two high-density DNA spots is displayed in the right panel. (C) Resolution assessment of the final 3D map using Fourier shell correlation (FSC). Two criteria are shown: FSC between two half-maps reconstructed from even and odd frames (evaluated at 0.143) and FSC between the final 3D map and the fitted model (evaluated at 0.5). (D) Zoomed-in views of the final 3D density map from panel A, displayed at two contour levels. (E) Superimposition of the high-contour level map from panel D onto its fitted model.

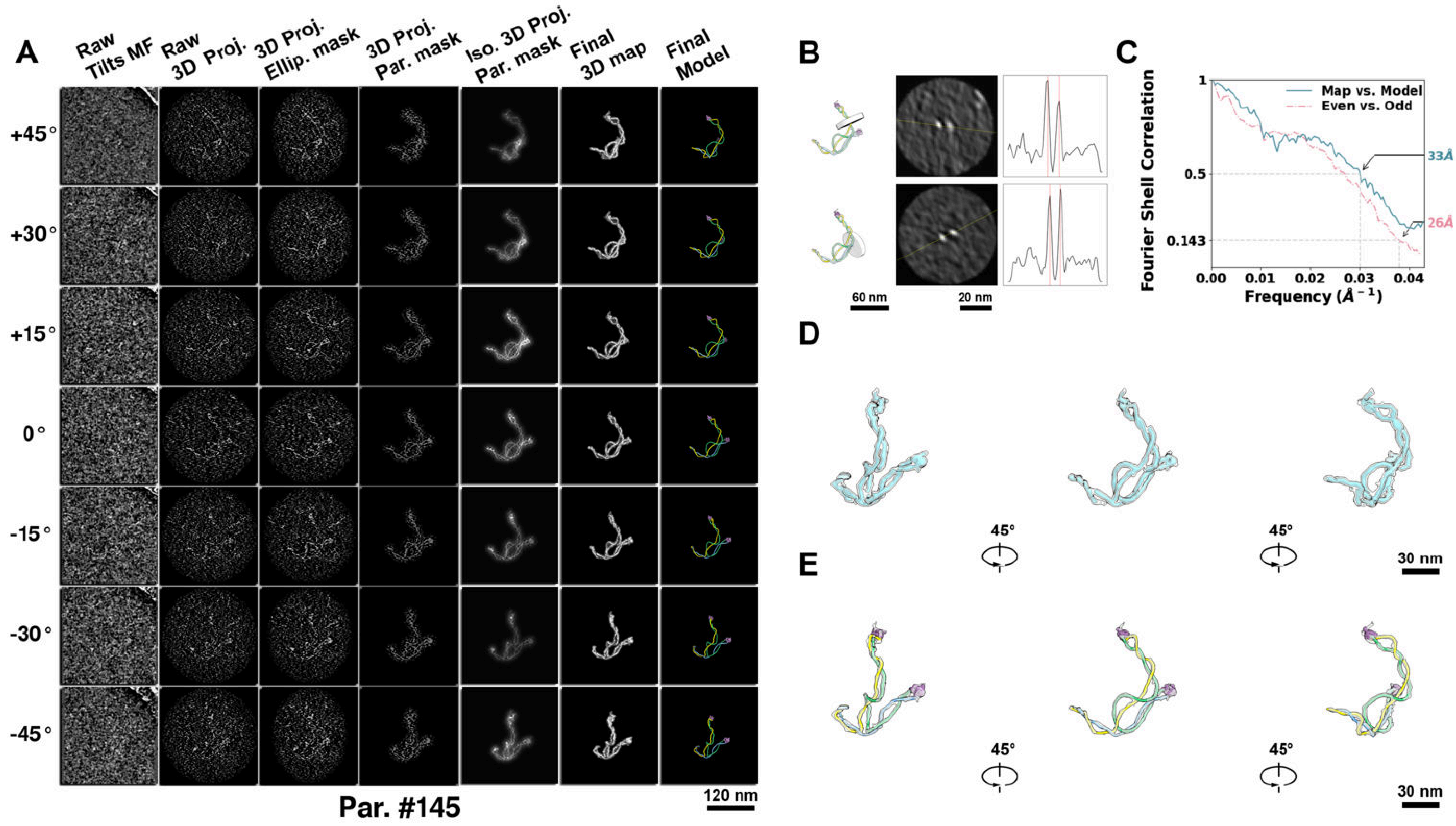

**Supplementary Particle Figure 145. Cryo-ET 3D reconstruction of an individual TEC particle.**
(A) 3D reconstruction of the plasmid particle (index no. 145). The first column shows seven representative tilt images from +45° to -45° in step of 15°. The second, third, and fourth columns show 3D projections of the particle with spherical, ellipsoidal (thinner along the z-dimension), and particle-shaped masks, respectively. The fifth column displays the 3D projections of the enhanced and IsoNet missing-wedge-corrected particle. The sixth and seventh columns present the final 3D map and the flexibly fitted model, respectively. (B) Two cross-sectional views (12 nm thickness) of the plasmid density map along its plectoneme axis are shown in the left-middle panel. The intensity profile along the line crossing the two high-density DNA spots is displayed in the right panel. (C) Resolution assessment of the final 3D map using Fourier shell correlation (FSC). Two criteria are shown: FSC between two half-maps reconstructed from even and odd frames (evaluated at 0.143) and FSC between the final 3D map and the fitted model (evaluated at 0.5). (D) Zoomed-in views of the final 3D density map from panel A, displayed at two contour levels. (E) Superimposition of the high-contour level map from panel D onto its fitted model.

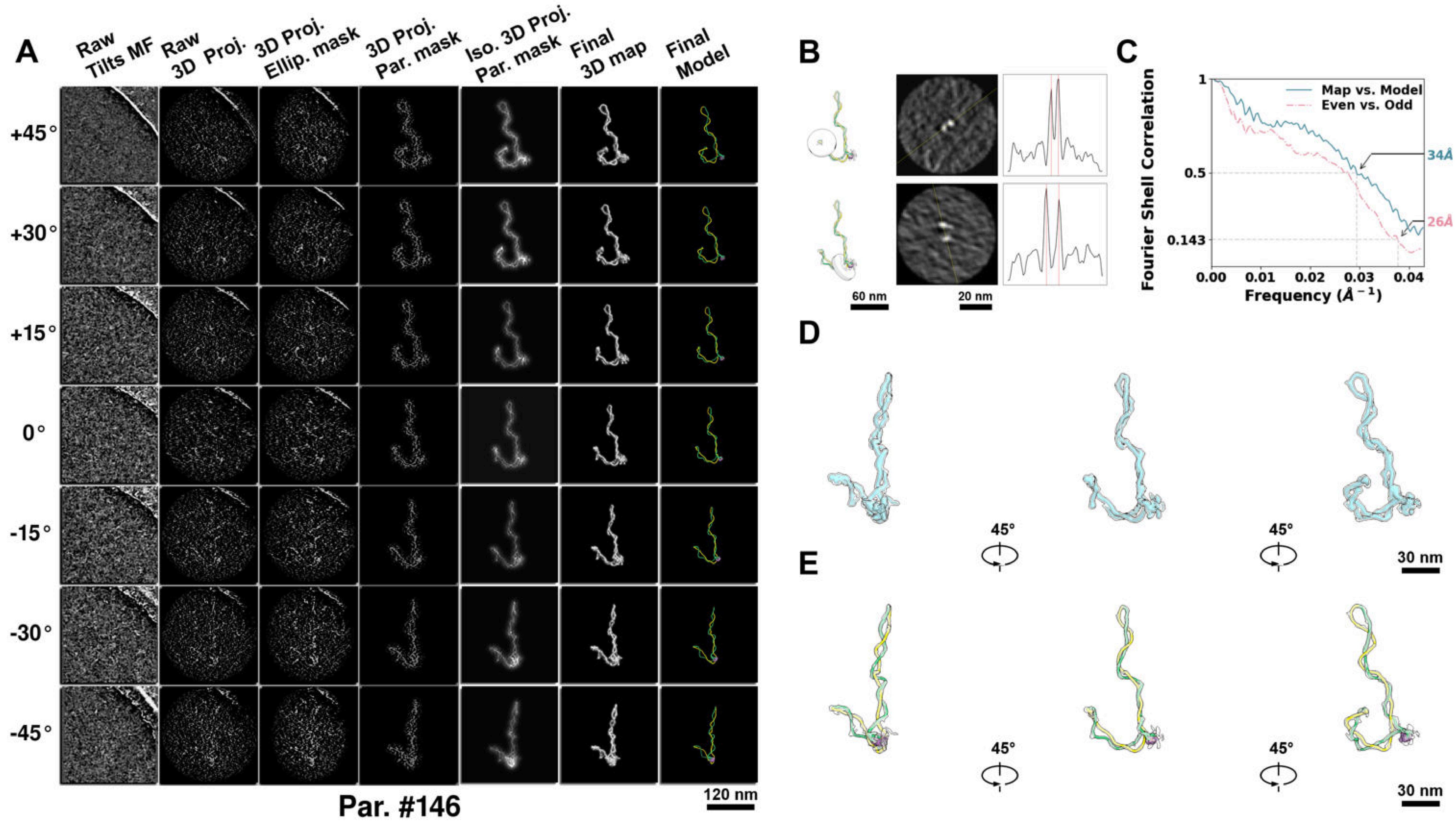

**Supplementary Particle Figure 146. Cryo-ET 3D reconstruction of an individual TEC particle.**
(A) 3D reconstruction of the plasmid particle (index no. 146). The first column shows seven representative tilt images from +45° to -45° in step of 15°. The second, third, and fourth columns show 3D projections of the particle with spherical, ellipsoidal (thinner along the z-dimension), and particle-shaped masks, respectively. The fifth column displays the 3D projections of the enhanced and IsoNet missing-wedge-corrected particle. The sixth and seventh columns present the final 3D map and the flexibly fitted model, respectively. (B) Two cross-sectional views (12 nm thickness) of the plasmid density map along its plectoneme axis are shown in the left-middle panel. The intensity profile along the line crossing the two high-density DNA spots is displayed in the right panel. (C) Resolution assessment of the final 3D map using Fourier shell correlation (FSC). Two criteria are shown: FSC between two half-maps reconstructed from even and odd frames (evaluated at 0.143) and FSC between the final 3D map and the fitted model (evaluated at 0.5). (D) Zoomed-in views of the final 3D density map from panel A, displayed at two contour levels. (E) Superimposition of the high-contour level map from panel D onto its fitted model.

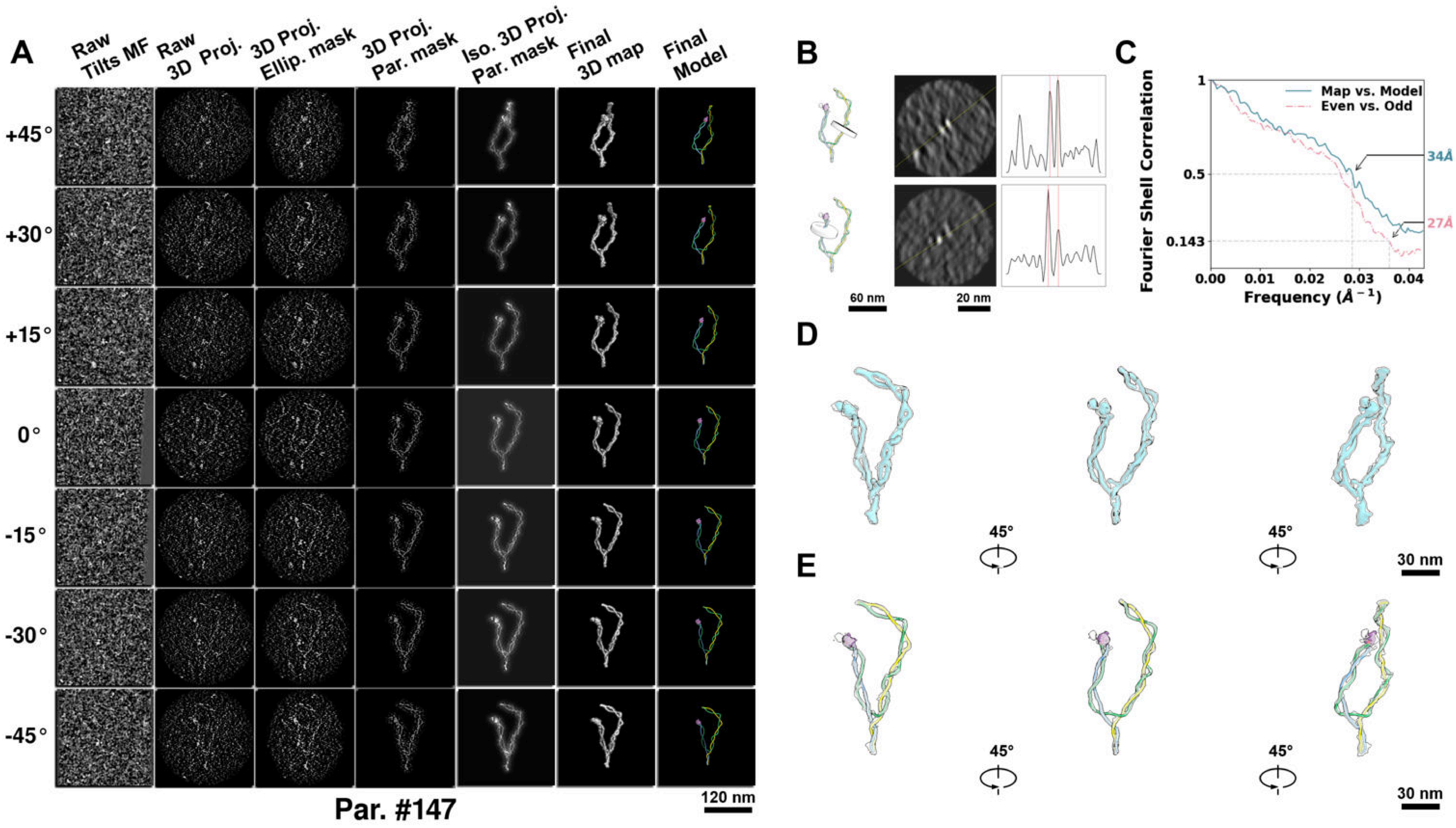

**Supplementary Particle Figure 147. Cryo-ET 3D reconstruction of an individual TEC particle.**
(A) 3D reconstruction of the plasmid particle (index no. 147). The first column shows seven representative tilt images from +45° to -45° in step of 15°. The second, third, and fourth columns show 3D projections of the particle with spherical, ellipsoidal (thinner along the z-dimension), and particle-shaped masks, respectively. The fifth column displays the 3D projections of the enhanced and IsoNet missing-wedge-corrected particle. The sixth and seventh columns present the final 3D map and the flexibly fitted model, respectively. (B) Two cross-sectional views (12 nm thickness) of the plasmid density map along its plectoneme axis are shown in the left-middle panel. The intensity profile along the line crossing the two high-density DNA spots is displayed in the right panel. (C) Resolution assessment of the final 3D map using Fourier shell correlation (FSC). Two criteria are shown: FSC between two half-maps reconstructed from even and odd frames (evaluated at 0.143) and FSC between the final 3D map and the fitted model (evaluated at 0.5). (D) Zoomed-in views of the final 3D density map from panel A, displayed at two contour levels. (E) Superimposition of the high-contour level map from panel D onto its fitted model.

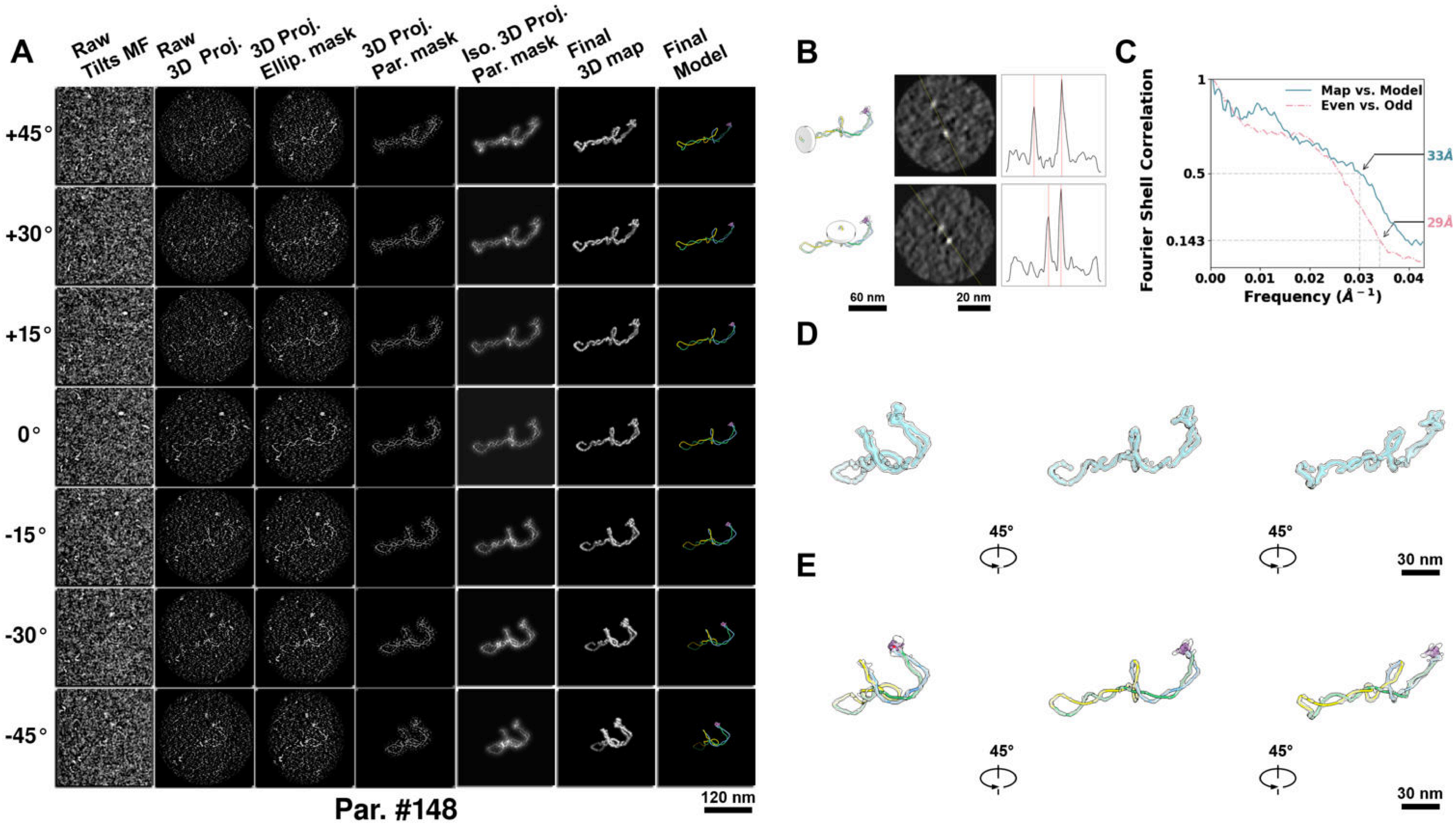

**Supplementary Particle Figure 148. Cryo-ET 3D reconstruction of an individual TEC particle.**
(A) 3D reconstruction of the plasmid particle (index no. 148). The first column shows seven representative tilt images from +45° to -45° in step of 15°. The second, third, and fourth columns show 3D projections of the particle with spherical, ellipsoidal (thinner along the z-dimension), and particle-shaped masks, respectively. The fifth column displays the 3D projections of the enhanced and IsoNet missing-wedge-corrected particle. The sixth and seventh columns present the final 3D map and the flexibly fitted model, respectively. (B) Two cross-sectional views (12 nm thickness) of the plasmid density map along its plectoneme axis are shown in the left-middle panel. The intensity profile along the line crossing the two high-density DNA spots is displayed in the right panel. (C) Resolution assessment of the final 3D map using Fourier shell correlation (FSC). Two criteria are shown: FSC between two half-maps reconstructed from even and odd frames (evaluated at 0.143) and FSC between the final 3D map and the fitted model (evaluated at 0.5). (D) Zoomed-in views of the final 3D density map from panel A, displayed at two contour levels. (E) Superimposition of the high-contour level map from panel D onto its fitted model.

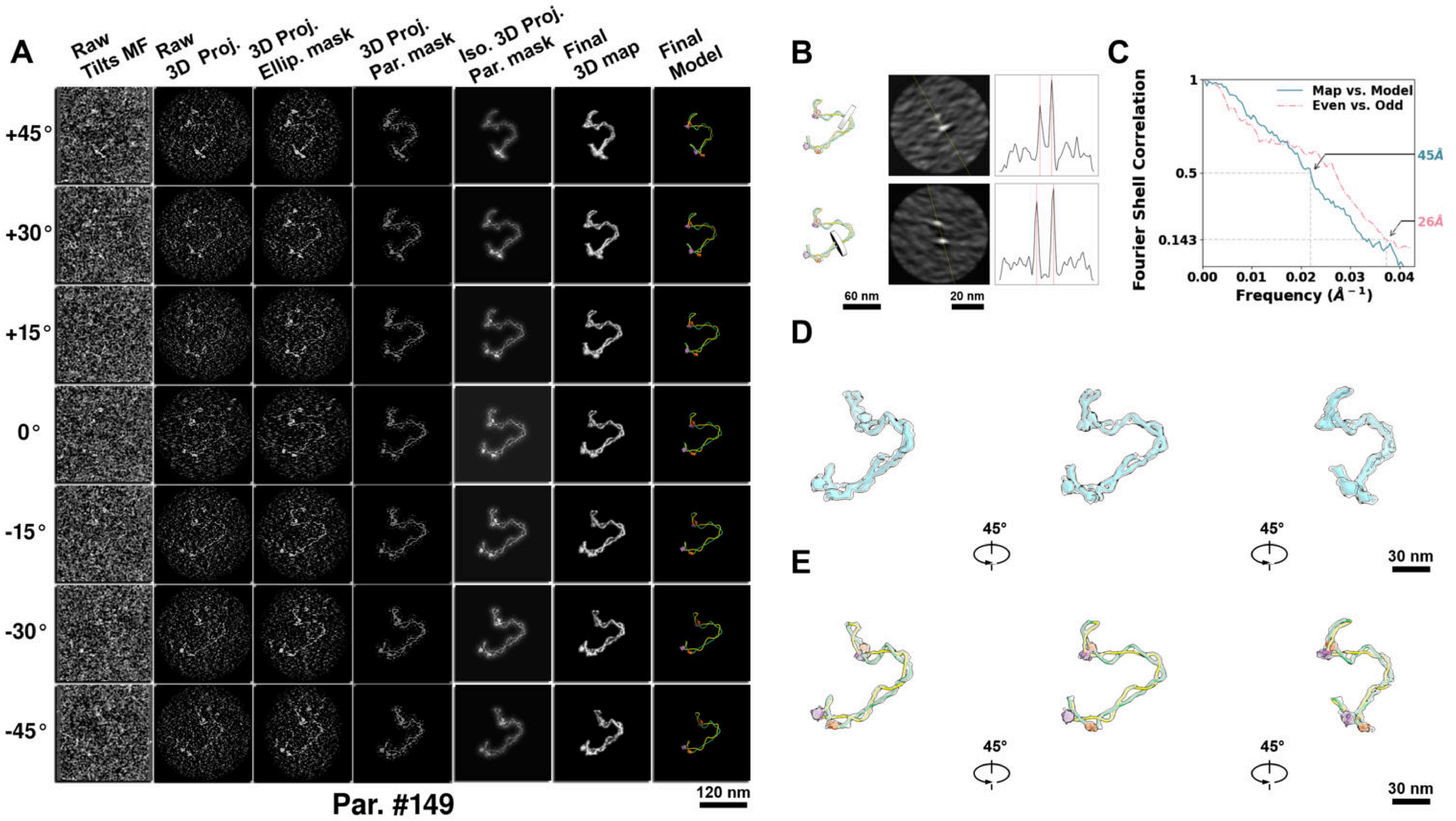

**Supplementary Particle Figure 149. Cryo-ET 3D reconstruction of an individual sTEC-Cas particle.**
(A) 3D reconstruction of the plasmid particle (index no. 149). The first column shows seven representative tilt images from +45° to -45° in step of 15°. The second, third, and fourth columns show 3D projections of the particle with spherical, ellipsoidal (thinner along the z-dimension), and particle-shaped masks, respectively. The fifth column displays the 3D projections of the enhanced and IsoNet missing-wedge-corrected particle. The sixth and seventh columns present the final 3D map and the flexibly fitted model, respectively. (B) Two cross-sectional views (12 nm thickness) of the plasmid density map along its plectoneme axis are shown in the left-middle panel. The intensity profile along the line crossing the two high-density DNA spots is displayed in the right panel. (C) Resolution assessment of the final 3D map using Fourier shell correlation (FSC). Two criteria are shown: FSC between two half-maps reconstructed from even and odd frames (evaluated at 0.143) and FSC between the final 3D map and the fitted model (evaluated at 0.5). (D) Zoomed-in views of the final 3D density map from panel A, displayed at two contour levels. (E) Superimposition of the high-contour level map from panel D onto its fitted model.

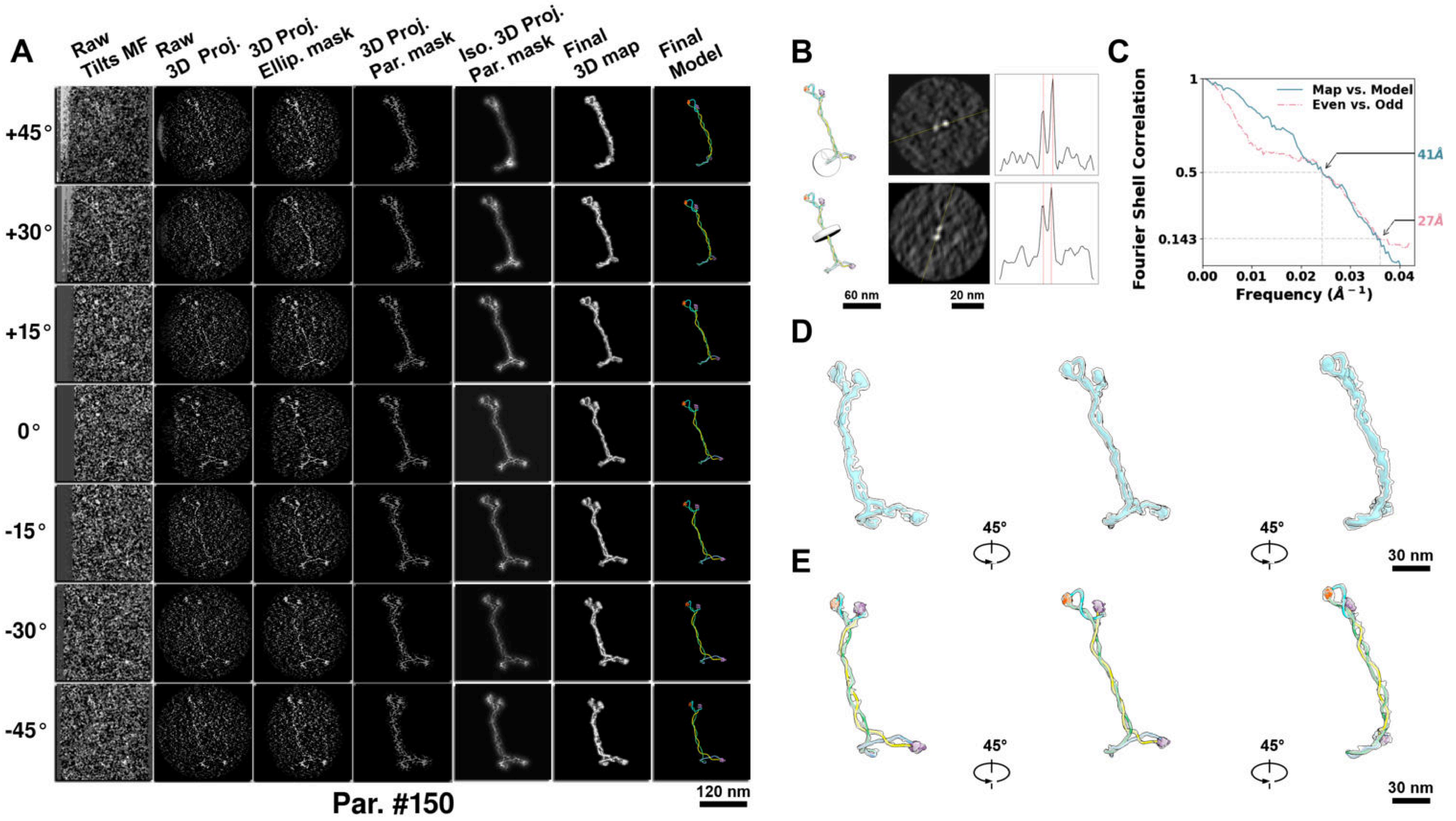

**Supplementary Particle Figure 150. Cryo-ET 3D reconstruction of an individual sTEC-Cas particle.**
(A) 3D reconstruction of the plasmid particle (index no. 150). The first column shows seven representative tilt images from +45° to -45° in step of 15°. The second, third, and fourth columns show 3D projections of the particle with spherical, ellipsoidal (thinner along the z-dimension), and particle-shaped masks, respectively. The fifth column displays the 3D projections of the enhanced and IsoNet missing-wedge-corrected particle. The sixth and seventh columns present the final 3D map and the flexibly fitted model, respectively. (B) Two cross-sectional views (12 nm thickness) of the plasmid density map along its plectoneme axis are shown in the left-middle panel. The intensity profile along the line crossing the two high-density DNA spots is displayed in the right panel. (C) Resolution assessment of the final 3D map using Fourier shell correlation (FSC). Two criteria are shown: FSC between two half-maps reconstructed from even and odd frames (evaluated at 0.143) and FSC between the final 3D map and the fitted model (evaluated at 0.5). (D) Zoomed-in views of the final 3D density map from panel A, displayed at two contour levels. (E) Superimposition of the high-contour level map from panel D onto its fitted model.

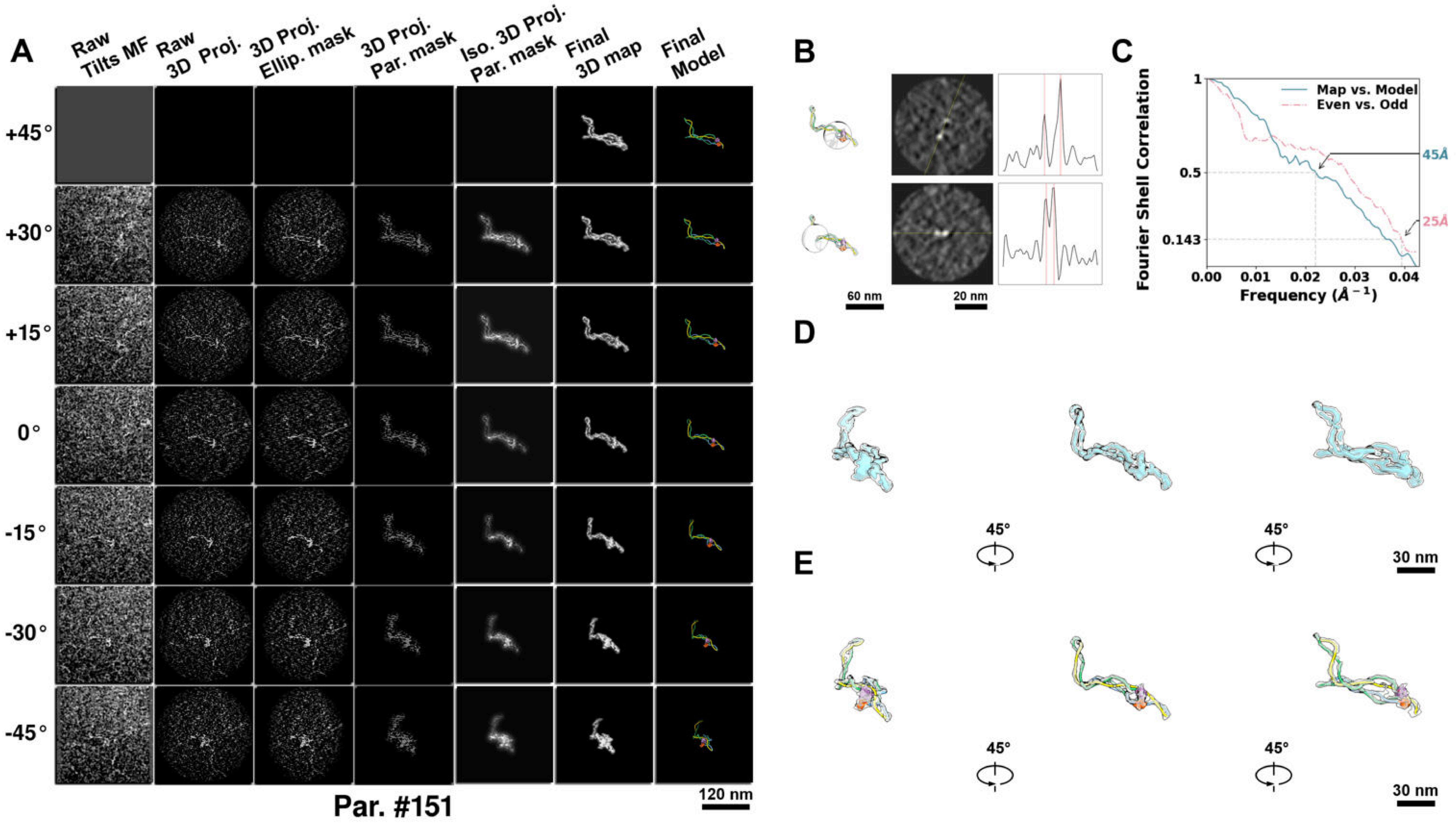

**Supplementary Particle Figure 151. Cryo-ET 3D reconstruction of an individual sTEC-Cas particle.**
(A) 3D reconstruction of the plasmid particle (index no. 151). The first column shows seven representative tilt images from +45° to -45° in step of 15°. The second, third, and fourth columns show 3D projections of the particle with spherical, ellipsoidal (thinner along the z-dimension), and particle-shaped masks, respectively. The fifth column displays the 3D projections of the enhanced and IsoNet missing-wedge-corrected particle. The sixth and seventh columns present the final 3D map and the flexibly fitted model, respectively. (B) Two cross-sectional views (12 nm thickness) of the plasmid density map along its plectoneme axis are shown in the left-middle panel. The intensity profile along the line crossing the two high-density DNA spots is displayed in the right panel. (C) Resolution assessment of the final 3D map using Fourier shell correlation (FSC). Two criteria are shown: FSC between two half-maps reconstructed from even and odd frames (evaluated at 0.143) and FSC between the final 3D map and the fitted model (evaluated at 0.5). (D) Zoomed-in views of the final 3D density map from panel A, displayed at two contour levels. (E) Superimposition of the high-contour level map from panel D onto its fitted model.

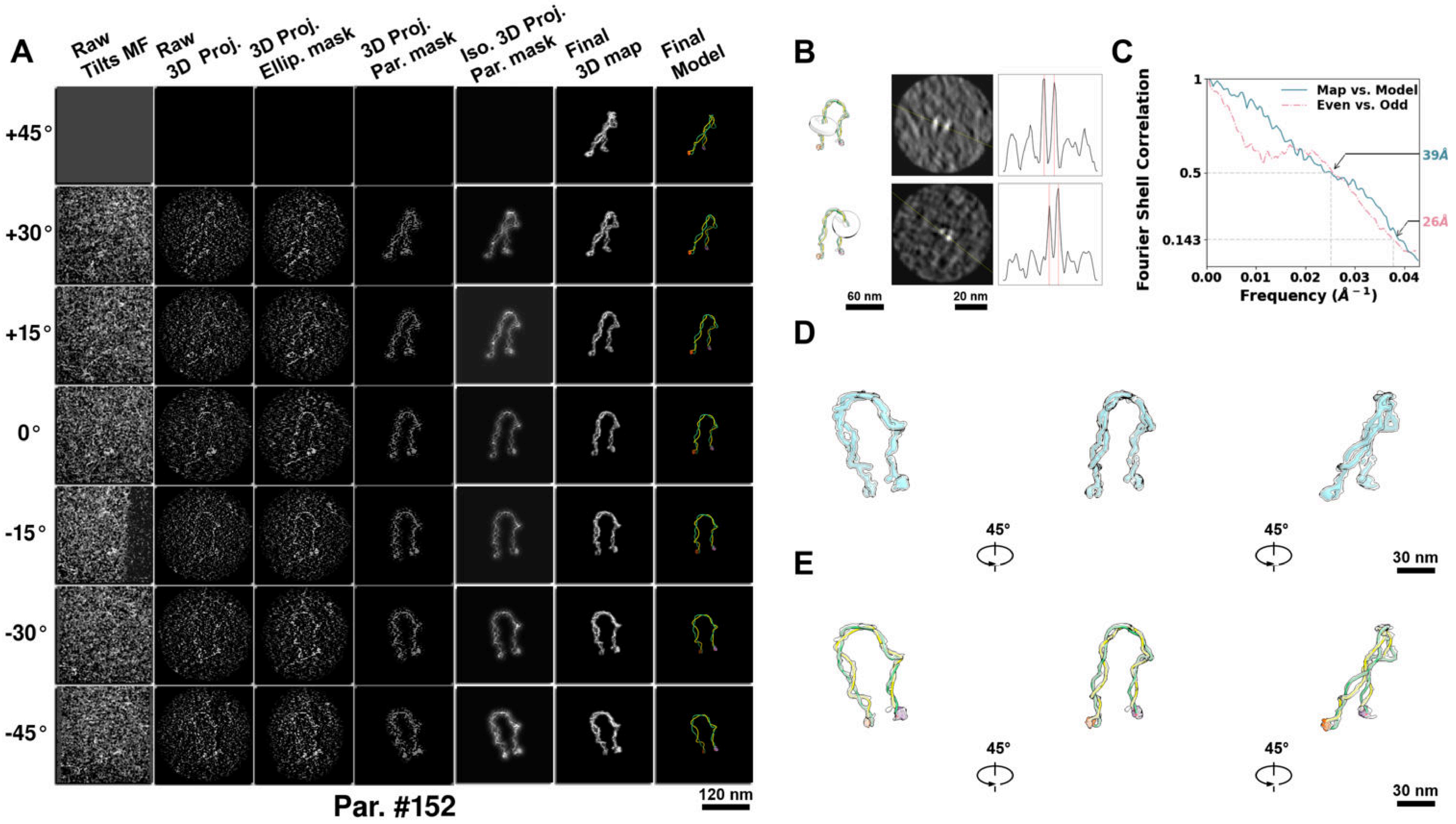

**Supplementary Particle Figure 152. Cryo-ET 3D reconstruction of an individual sTEC-Cas particle.**
(A) 3D reconstruction of the plasmid particle (index no. 152). The first column shows seven representative tilt images from +45° to -45° in step of 15°. The second, third, and fourth columns show 3D projections of the particle with spherical, ellipsoidal (thinner along the z-dimension), and particle-shaped masks, respectively. The fifth column displays the 3D projections of the enhanced and IsoNet missing-wedge-corrected particle. The sixth and seventh columns present the final 3D map and the flexibly fitted model, respectively. (B) Two cross-sectional views (12 nm thickness) of the plasmid density map along its plectoneme axis are shown in the left-middle panel. The intensity profile along the line crossing the two high-density DNA spots is displayed in the right panel. (C) Resolution assessment of the final 3D map using Fourier shell correlation (FSC). Two criteria are shown: FSC between two half-maps reconstructed from even and odd frames (evaluated at 0.143) and FSC between the final 3D map and the fitted model (evaluated at 0.5). (D) Zoomed-in views of the final 3D density map from panel A, displayed at two contour levels. (E) Superimposition of the high-contour level map from panel D onto its fitted model.

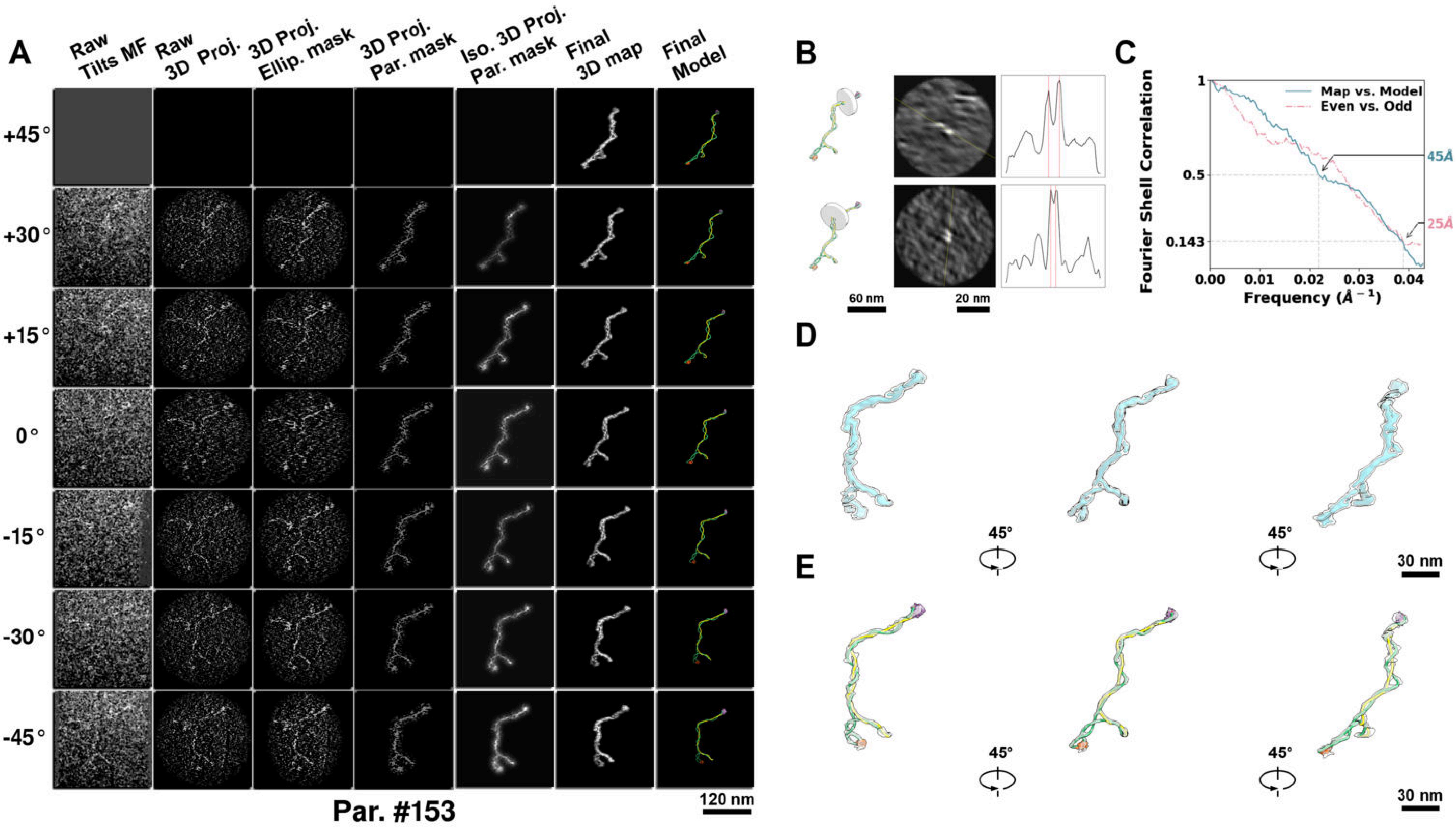

**Supplementary Particle Figure 153. Cryo-ET 3D reconstruction of an individual sTEC-Cas particle.**
(A) 3D reconstruction of the plasmid particle (index no. 153). The first column shows seven representative tilt images from +45° to -45° in step of 15°. The second, third, and fourth columns show 3D projections of the particle with spherical, ellipsoidal (thinner along the z-dimension), and particle-shaped masks, respectively. The fifth column displays the 3D projections of the enhanced and IsoNet missing-wedge-corrected particle. The sixth and seventh columns present the final 3D map and the flexibly fitted model, respectively. (B) Two cross-sectional views (12 nm thickness) of the plasmid density map along its plectoneme axis are shown in the left-middle panel. The intensity profile along the line crossing the two high-density DNA spots is displayed in the right panel. (C) Resolution assessment of the final 3D map using Fourier shell correlation (FSC). Two criteria are shown: FSC between two half-maps reconstructed from even and odd frames (evaluated at 0.143) and FSC between the final 3D map and the fitted model (evaluated at 0.5). (D) Zoomed-in views of the final 3D density map from panel A, displayed at two contour levels. (E) Superimposition of the high-contour level map from panel D onto its fitted model.

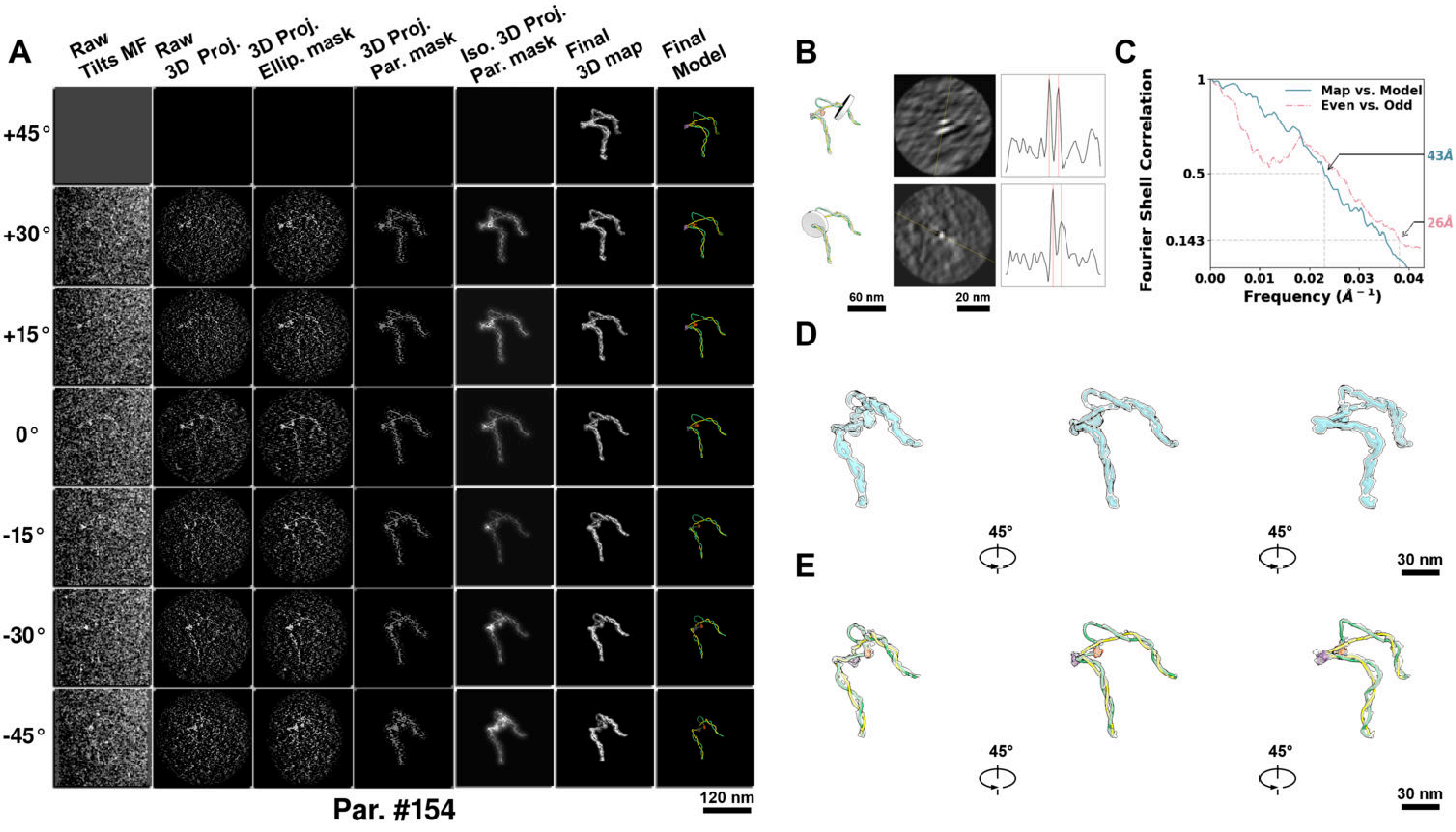

**Supplementary Particle Figure 154. Cryo-ET 3D reconstruction of an individual sTEC-Cas particle.**
(A) 3D reconstruction of the plasmid particle (index no. 154). The first column shows seven representative tilt images from +45° to -45° in step of 15°. The second, third, and fourth columns show 3D projections of the particle with spherical, ellipsoidal (thinner along the z-dimension), and particle-shaped masks, respectively. The fifth column displays the 3D projections of the enhanced and IsoNet missing-wedge-corrected particle. The sixth and seventh columns present the final 3D map and the flexibly fitted model, respectively. (B) Two cross-sectional views (12 nm thickness) of the plasmid density map along its plectoneme axis are shown in the left-middle panel. The intensity profile along the line crossing the two high-density DNA spots is displayed in the right panel. (C) Resolution assessment of the final 3D map using Fourier shell correlation (FSC). Two criteria are shown: FSC between two half-maps reconstructed from even and odd frames (evaluated at 0.143) and FSC between the final 3D map and the fitted model (evaluated at 0.5). (D) Zoomed-in views of the final 3D density map from panel A, displayed at two contour levels. (E) Superimposition of the high-contour level map from panel D onto its fitted model.

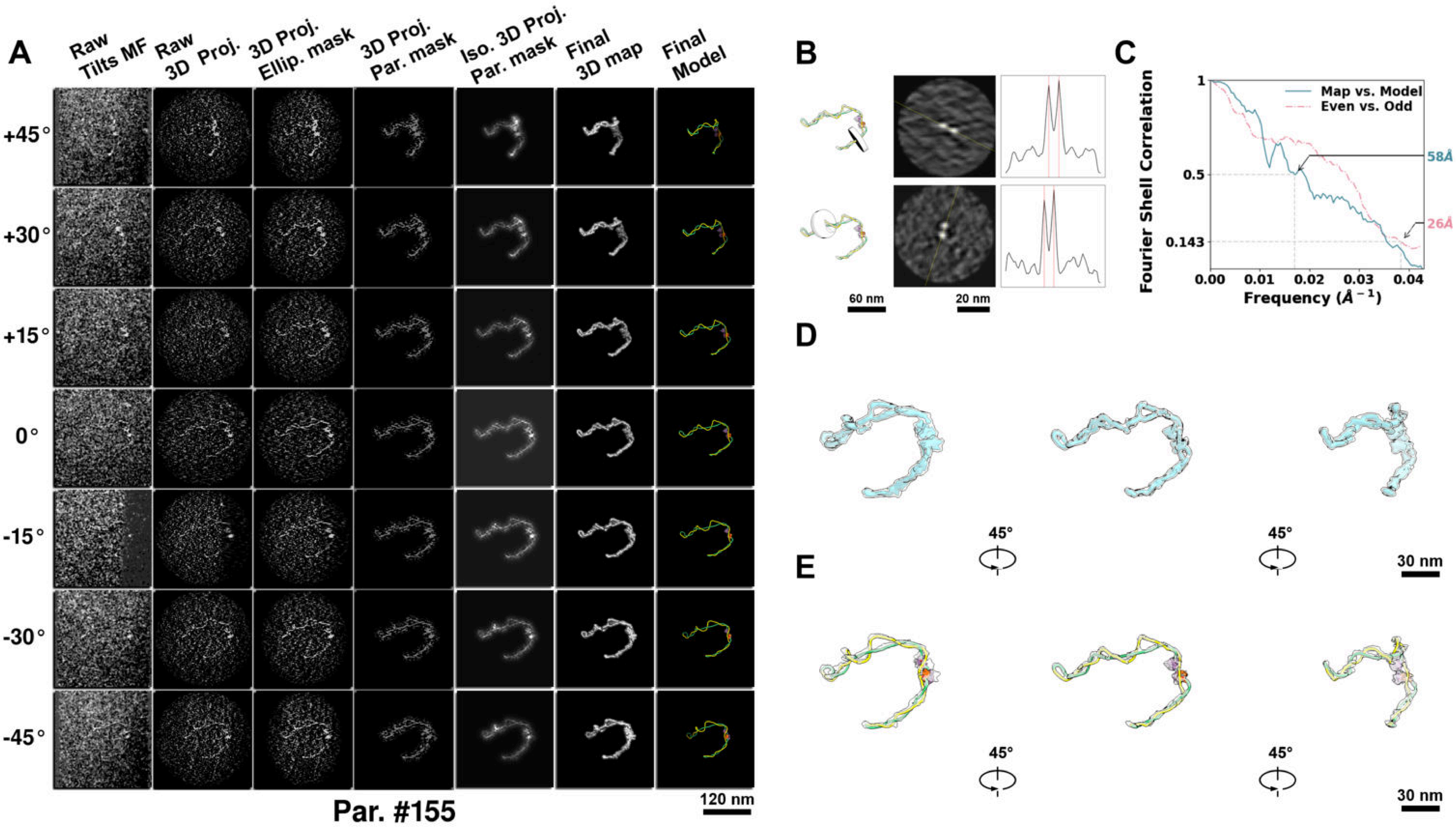

**Supplementary Particle Figure 155. Cryo-ET 3D reconstruction of an individual sTEC-Cas particle.**
(A) 3D reconstruction of the plasmid particle (index no. 155). The first column shows seven representative tilt images from +45° to -45° in step of 15°. The second, third, and fourth columns show 3D projections of the particle with spherical, ellipsoidal (thinner along the z-dimension), and particle-shaped masks, respectively. The fifth column displays the 3D projections of the enhanced and IsoNet missing-wedge-corrected particle. The sixth and seventh columns present the final 3D map and the flexibly fitted model, respectively. (B) Two cross-sectional views (12 nm thickness) of the plasmid density map along its plectoneme axis are shown in the left-middle panel. The intensity profile along the line crossing the two high-density DNA spots is displayed in the right panel. (C) Resolution assessment of the final 3D map using Fourier shell correlation (FSC). Two criteria are shown: FSC between two half-maps reconstructed from even and odd frames (evaluated at 0.143) and FSC between the final 3D map and the fitted model (evaluated at 0.5). (D) Zoomed-in views of the final 3D density map from panel A, displayed at two contour levels. (E) Superimposition of the high-contour level map from panel D onto its fitted model.

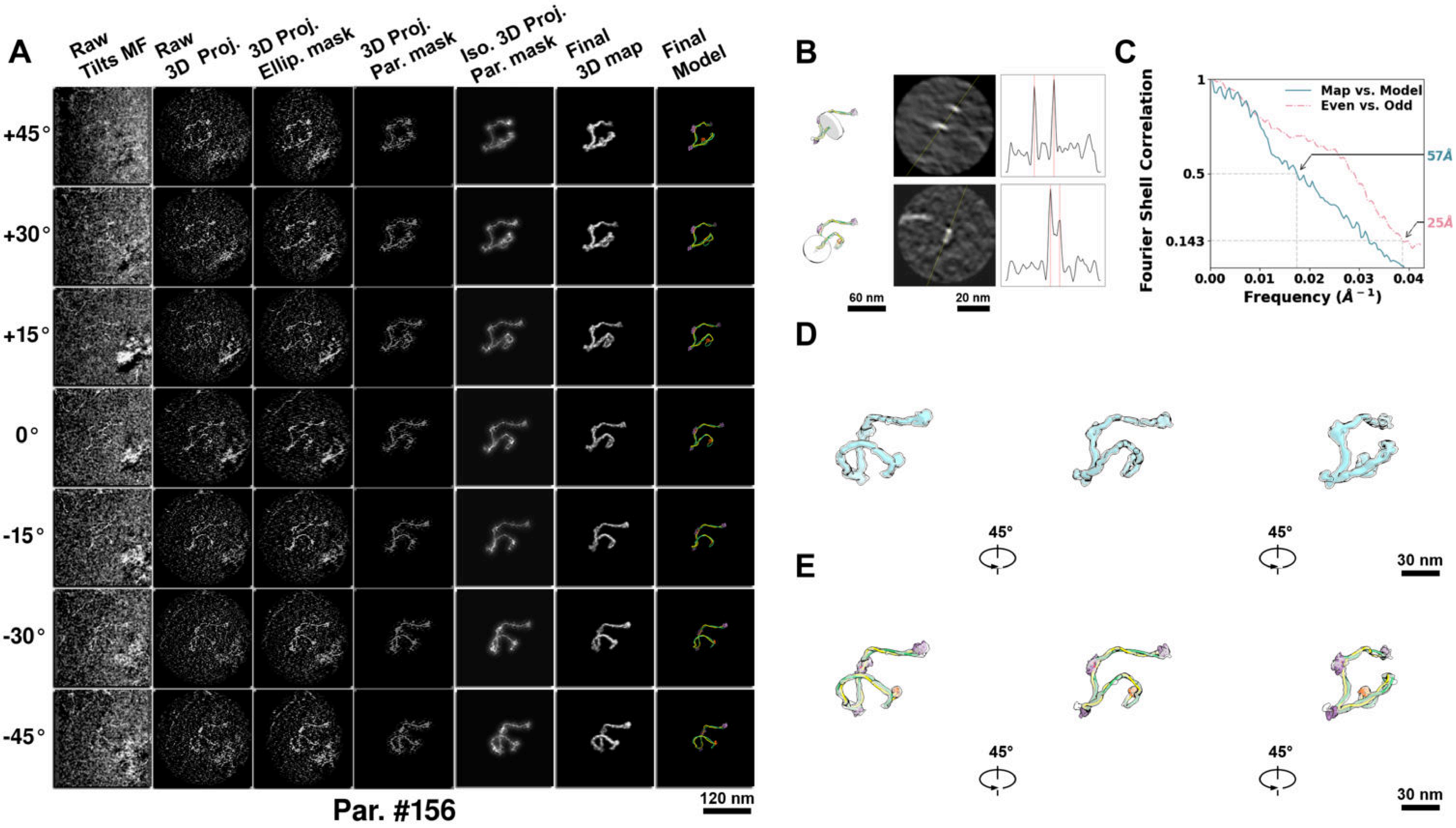

**Supplementary Particle Figure 156. Cryo-ET 3D reconstruction of an individual sTEC-Cas particle.**
(A) 3D reconstruction of the plasmid particle (index no. 156). The first column shows seven representative tilt images from +45° to -45° in step of 15°. The second, third, and fourth columns show 3D projections of the particle with spherical, ellipsoidal (thinner along the z-dimension), and particle-shaped masks, respectively. The fifth column displays the 3D projections of the enhanced and IsoNet missing-wedge-corrected particle. The sixth and seventh columns present the final 3D map and the flexibly fitted model, respectively. (B) Two cross-sectional views (12 nm thickness) of the plasmid density map along its plectoneme axis are shown in the left-middle panel. The intensity profile along the line crossing the two high-density DNA spots is displayed in the right panel. (C) Resolution assessment of the final 3D map using Fourier shell correlation (FSC). Two criteria are shown: FSC between two half-maps reconstructed from even and odd frames (evaluated at 0.143) and FSC between the final 3D map and the fitted model (evaluated at 0.5). (D) Zoomed-in views of the final 3D density map from panel A, displayed at two contour levels. (E) Superimposition of the high-contour level map from panel D onto its fitted model.

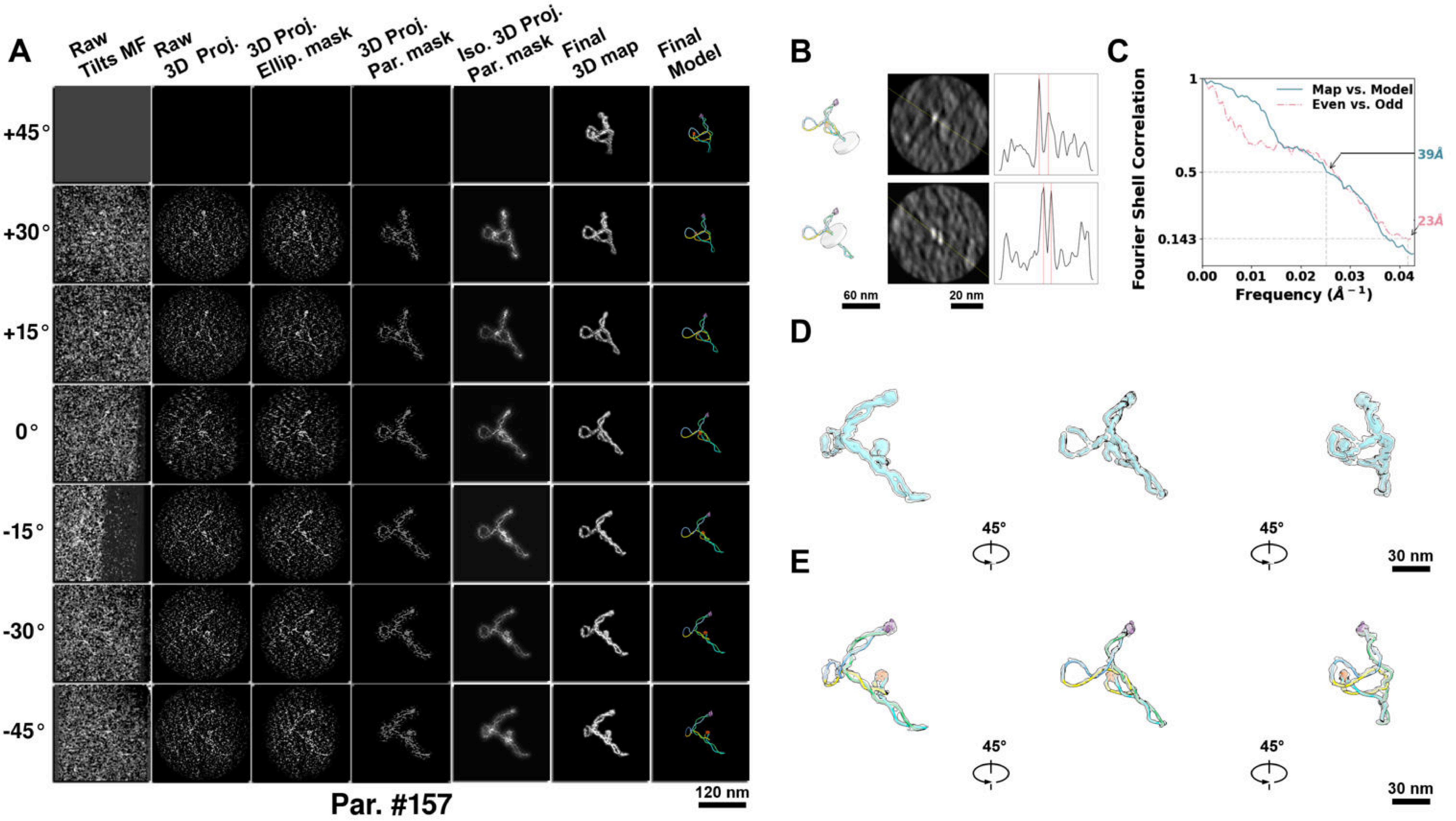

**Supplementary Particle Figure 157. Cryo-ET 3D reconstruction of an individual sTEC-Cas particle.**
(A) 3D reconstruction of the plasmid particle (index no. 157). The first column shows seven representative tilt images from +45° to -45° in step of 15°. The second, third, and fourth columns show 3D projections of the particle with spherical, ellipsoidal (thinner along the z-dimension), and particle-shaped masks, respectively. The fifth column displays the 3D projections of the enhanced and IsoNet missing-wedge-corrected particle. The sixth and seventh columns present the final 3D map and the flexibly fitted model, respectively. (B) Two cross-sectional views (12 nm thickness) of the plasmid density map along its plectoneme axis are shown in the left-middle panel. The intensity profile along the line crossing the two high-density DNA spots is displayed in the right panel. (C) Resolution assessment of the final 3D map using Fourier shell correlation (FSC). Two criteria are shown: FSC between two half-maps reconstructed from even and odd frames (evaluated at 0.143) and FSC between the final 3D map and the fitted model (evaluated at 0.5). (D) Zoomed-in views of the final 3D density map from panel A, displayed at two contour levels. (E) Superimposition of the high-contour level map from panel D onto its fitted model.

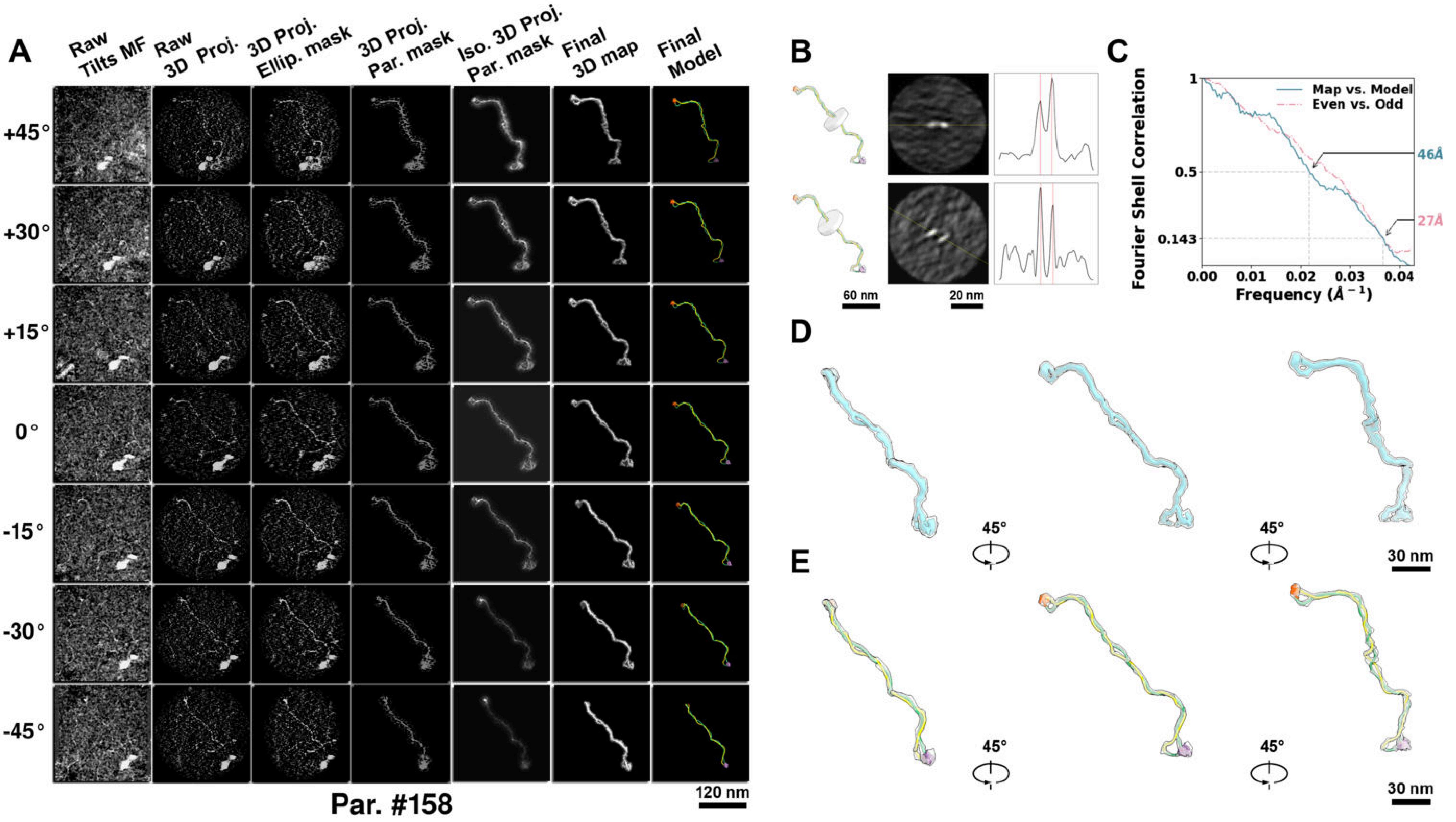

**Supplementary Particle Figure 158. Cryo-ET 3D reconstruction of an individual sTEC-Cas particle.**
(A) 3D reconstruction of the plasmid particle (index no. 158). The first column shows seven representative tilt images from +45° to -45° in step of 15°. The second, third, and fourth columns show 3D projections of the particle with spherical, ellipsoidal (thinner along the z-dimension), and particle-shaped masks, respectively. The fifth column displays the 3D projections of the enhanced and IsoNet missing-wedge-corrected particle. The sixth and seventh columns present the final 3D map and the flexibly fitted model, respectively. (B) Two cross-sectional views (12 nm thickness) of the plasmid density map along its plectoneme axis are shown in the left-middle panel. The intensity profile along the line crossing the two high-density DNA spots is displayed in the right panel. (C) Resolution assessment of the final 3D map using Fourier shell correlation (FSC). Two criteria are shown: FSC between two half-maps reconstructed from even and odd frames (evaluated at 0.143) and FSC between the final 3D map and the fitted model (evaluated at 0.5). (D) Zoomed-in views of the final 3D density map from panel A, displayed at two contour levels. (E) Superimposition of the high-contour level map from panel D onto its fitted model.

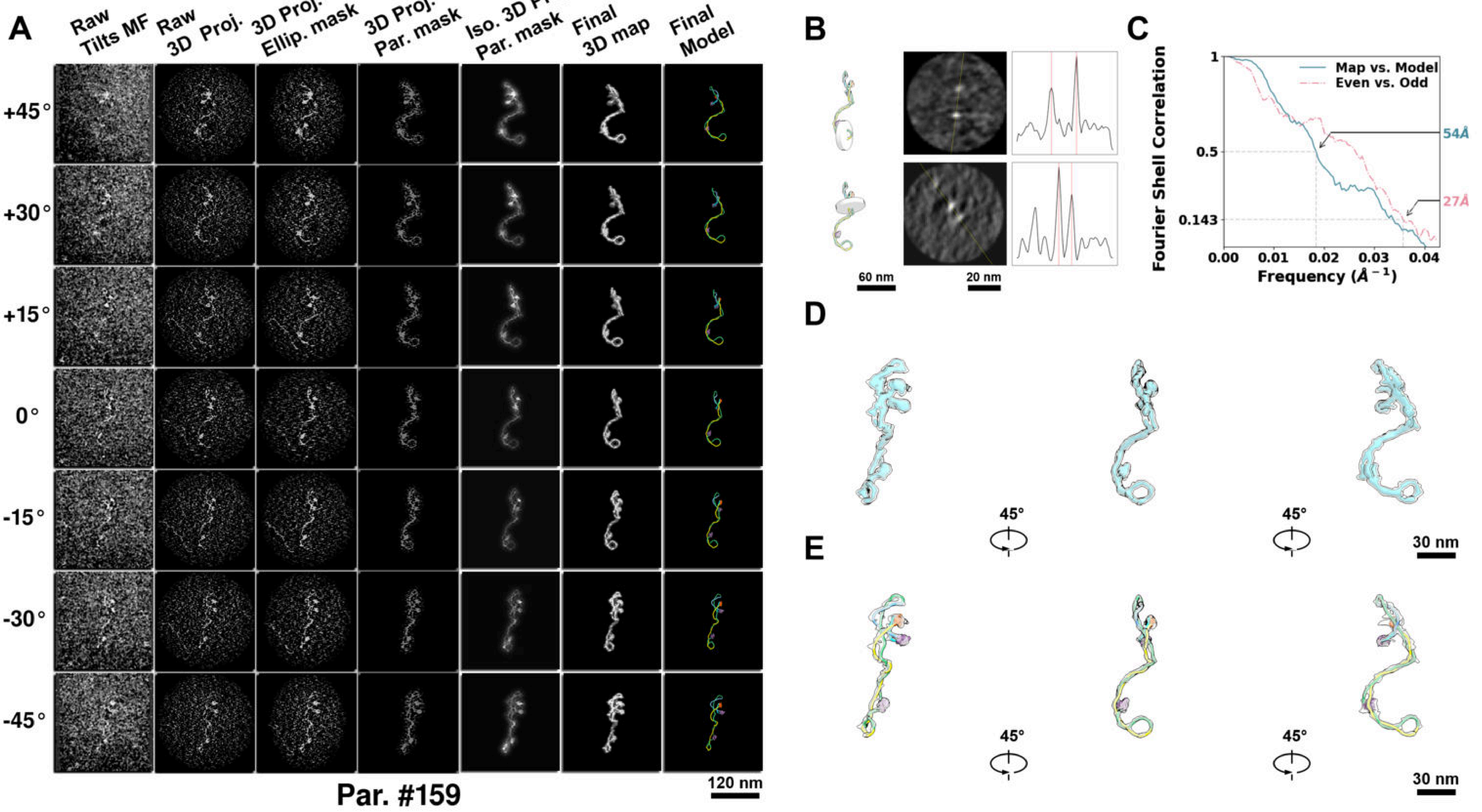

**Supplementary Particle Figure 159. Cryo-ET 3D reconstruction of an individual sTEC-Cas particle.**
(A) 3D reconstruction of the plasmid particle (index no. 159). The first column shows seven representative tilt images from +45° to -45° in step of 15°. The second, third, and fourth columns show 3D projections of the particle with spherical, ellipsoidal (thinner along the z-dimension), and particle-shaped masks, respectively. The fifth column displays the 3D projections of the enhanced and IsoNet missing-wedge-corrected particle. The sixth and seventh columns present the final 3D map and the flexibly fitted model, respectively. (B) Two cross-sectional views (12 nm thickness) of the plasmid density map along its plectoneme axis are shown in the left-middle panel. The intensity profile along the line crossing the two high-density DNA spots is displayed in the right panel. (C) Resolution assessment of the final 3D map using Fourier shell correlation (FSC). Two criteria are shown: FSC between two half-maps reconstructed from even and odd frames (evaluated at 0.143) and FSC between the final 3D map and the fitted model (evaluated at 0.5). (D) Zoomed-in views of the final 3D density map from panel A, displayed at two contour levels. (E) Superimposition of the high-contour level map from panel D onto its fitted model.

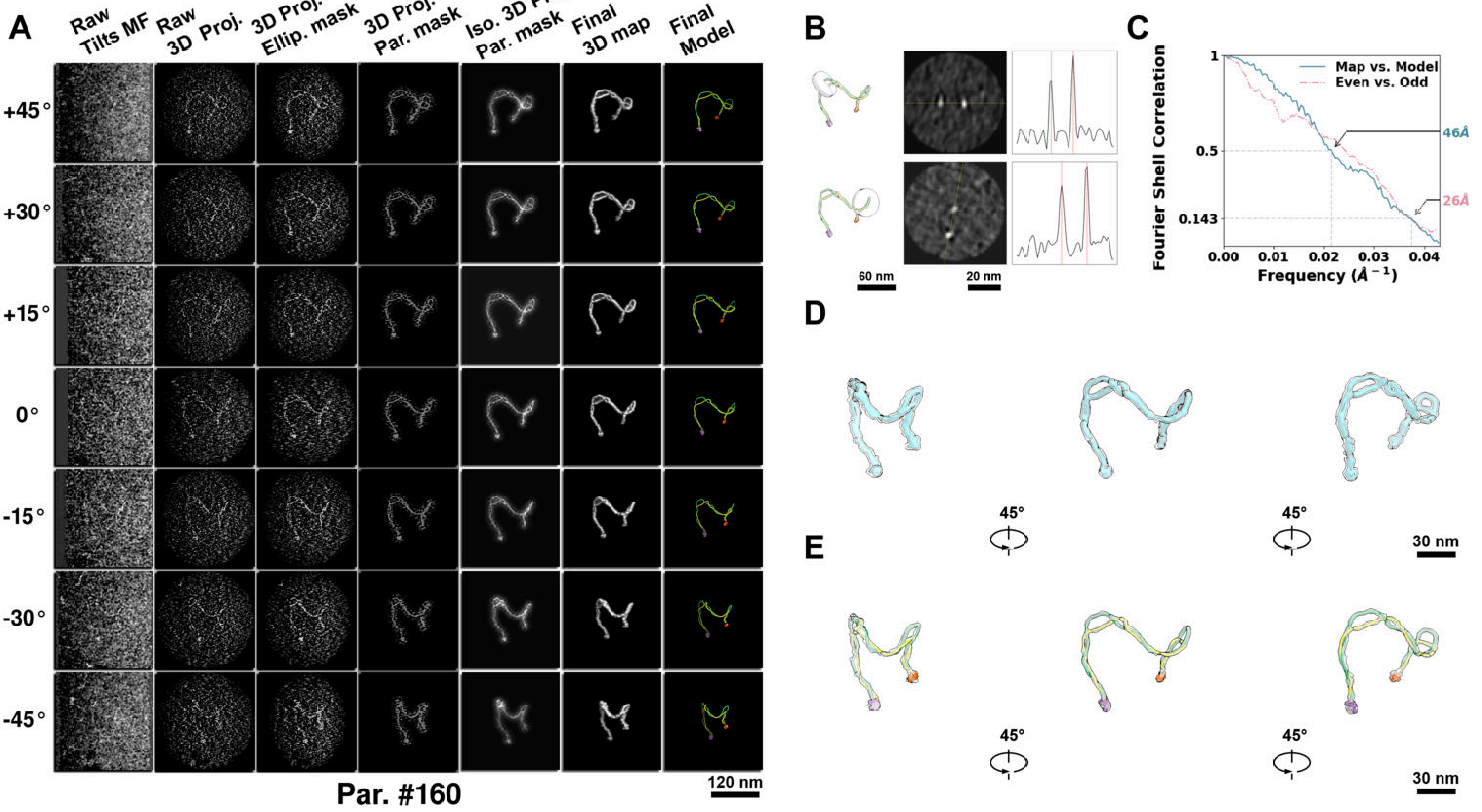

**Supplementary Particle Figure 160. Cryo-ET 3D reconstruction of an individual sTEC-Cas particle.**
(A) 3D reconstruction of the plasmid particle (index no. 160). The first column shows seven representative tilt images from +45° to -45° in step of 15°. The second, third, and fourth columns show 3D projections of the particle with spherical, ellipsoidal (thinner along the z-dimension), and particle-shaped masks, respectively. The fifth column displays the 3D projections of the enhanced and IsoNet missing-wedge-corrected particle. The sixth and seventh columns present the final 3D map and the flexibly fitted model, respectively. (B) Two cross-sectional views (12 nm thickness) of the plasmid density map along its plectoneme axis are shown in the left-middle panel. The intensity profile along the line crossing the two high-density DNA spots is displayed in the right panel. (C) Resolution assessment of the final 3D map using Fourier shell correlation (FSC). Two criteria are shown: FSC between two half-maps reconstructed from even and odd frames (evaluated at 0.143) and FSC between the final 3D map and the fitted model (evaluated at 0.5). (D) Zoomed-in views of the final 3D density map from panel A, displayed at two contour levels. (E) Superimposition of the high-contour level map from panel D onto its fitted model.

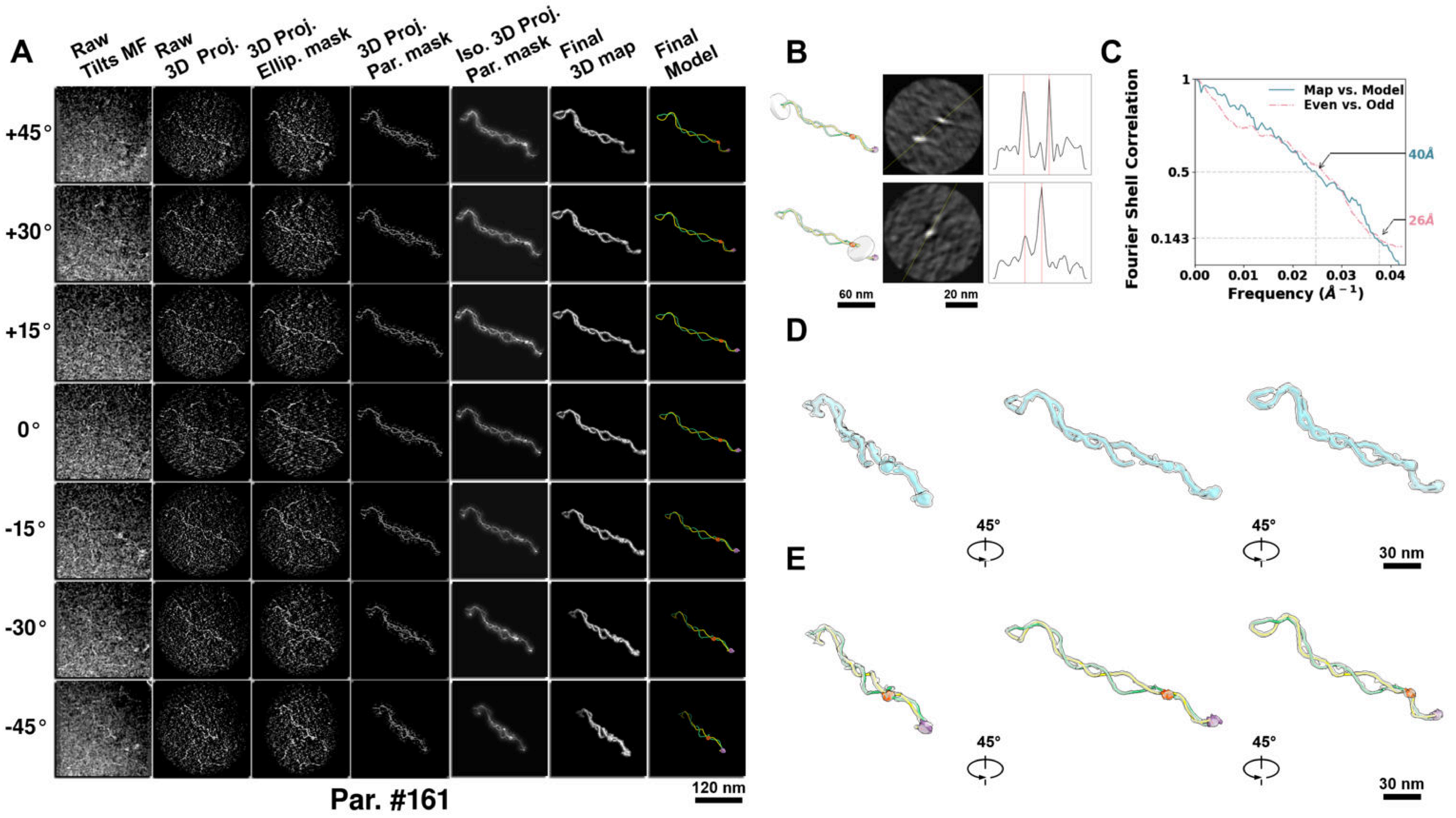

**Supplementary Particle Figure 161. Cryo-ET 3D reconstruction of an individual sTEC-Cas particle.**
(A) 3D reconstruction of the plasmid particle (index no. 161). The first column shows seven representative tilt images from +45° to -45° in step of 15°. The second, third, and fourth columns show 3D projections of the particle with spherical, ellipsoidal (thinner along the z-dimension), and particle-shaped masks, respectively. The fifth column displays the 3D projections of the enhanced and IsoNet missing-wedge-corrected particle. The sixth and seventh columns present the final 3D map and the flexibly fitted model, respectively. (B) Two cross-sectional views (12 nm thickness) of the plasmid density map along its plectoneme axis are shown in the left-middle panel. The intensity profile along the line crossing the two high-density DNA spots is displayed in the right panel. (C) Resolution assessment of the final 3D map using Fourier shell correlation (FSC). Two criteria are shown: FSC between two half-maps reconstructed from even and odd frames (evaluated at 0.143) and FSC between the final 3D map and the fitted model (evaluated at 0.5). (D) Zoomed-in views of the final 3D density map from panel A, displayed at two contour levels. (E) Superimposition of the high-contour level map from panel D onto its fitted model.

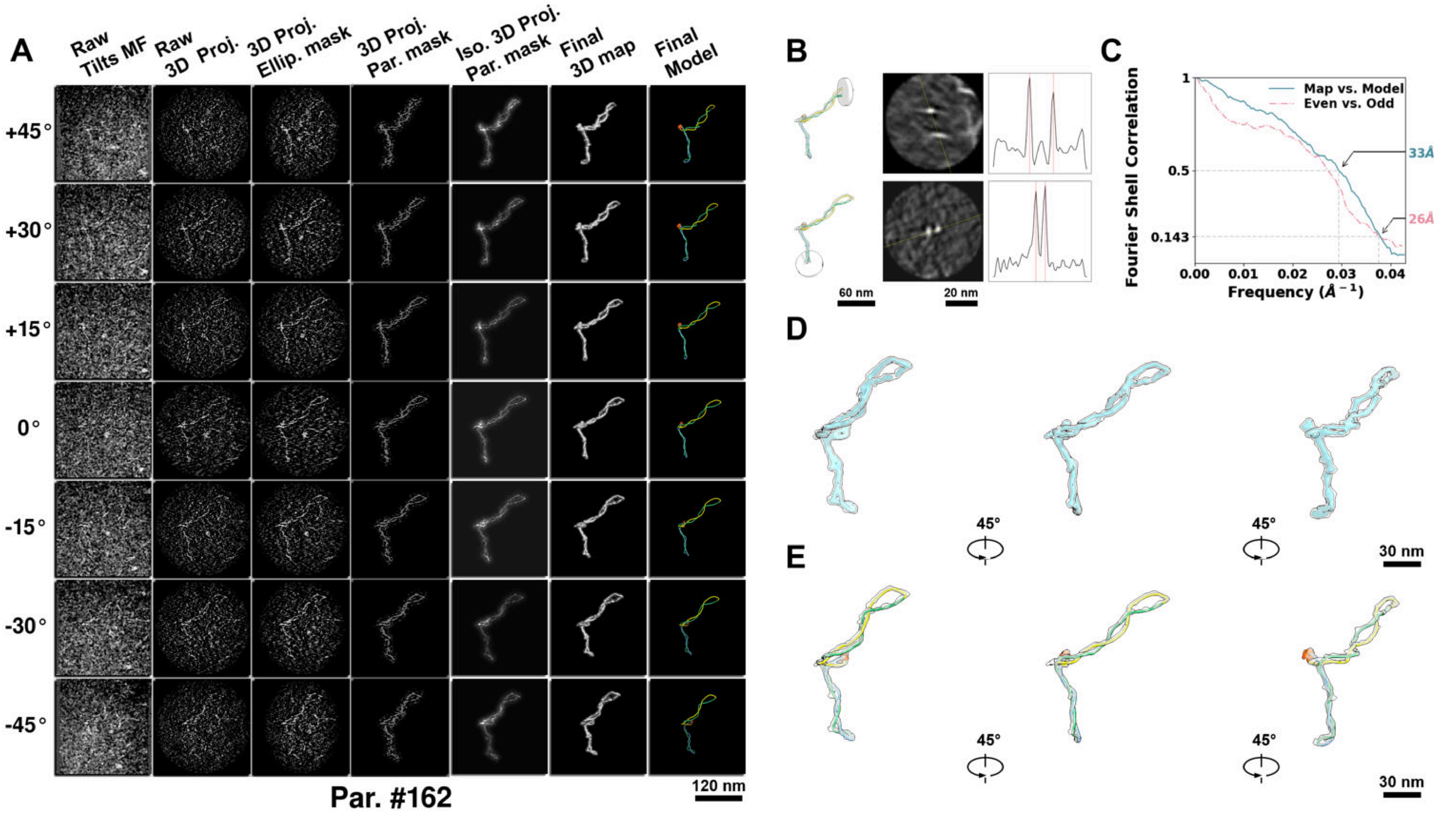

**Supplementary Particle Figure 162. Cryo-ET 3D reconstruction of an individual sTEC-Cas particle.**
(A) 3D reconstruction of the plasmid particle (index no. 162). The first column shows seven representative tilt images from +45° to -45° in step of 15°. The second, third, and fourth columns show 3D projections of the particle with spherical, ellipsoidal (thinner along the z-dimension), and particle-shaped masks, respectively. The fifth column displays the 3D projections of the enhanced and IsoNet missing-wedge-corrected particle. The sixth and seventh columns present the final 3D map and the flexibly fitted model, respectively. (B) Two cross-sectional views (12 nm thickness) of the plasmid density map along its plectoneme axis are shown in the left-middle panel. The intensity profile along the line crossing the two high-density DNA spots is displayed in the right panel. (C) Resolution assessment of the final 3D map using Fourier shell correlation (FSC). Two criteria are shown: FSC between two half-maps reconstructed from even and odd frames (evaluated at 0.143) and FSC between the final 3D map and the fitted model (evaluated at 0.5). (D) Zoomed-in views of the final 3D density map from panel A, displayed at two contour levels. (E) Superimposition of the high-contour level map from panel D onto its fitted model.

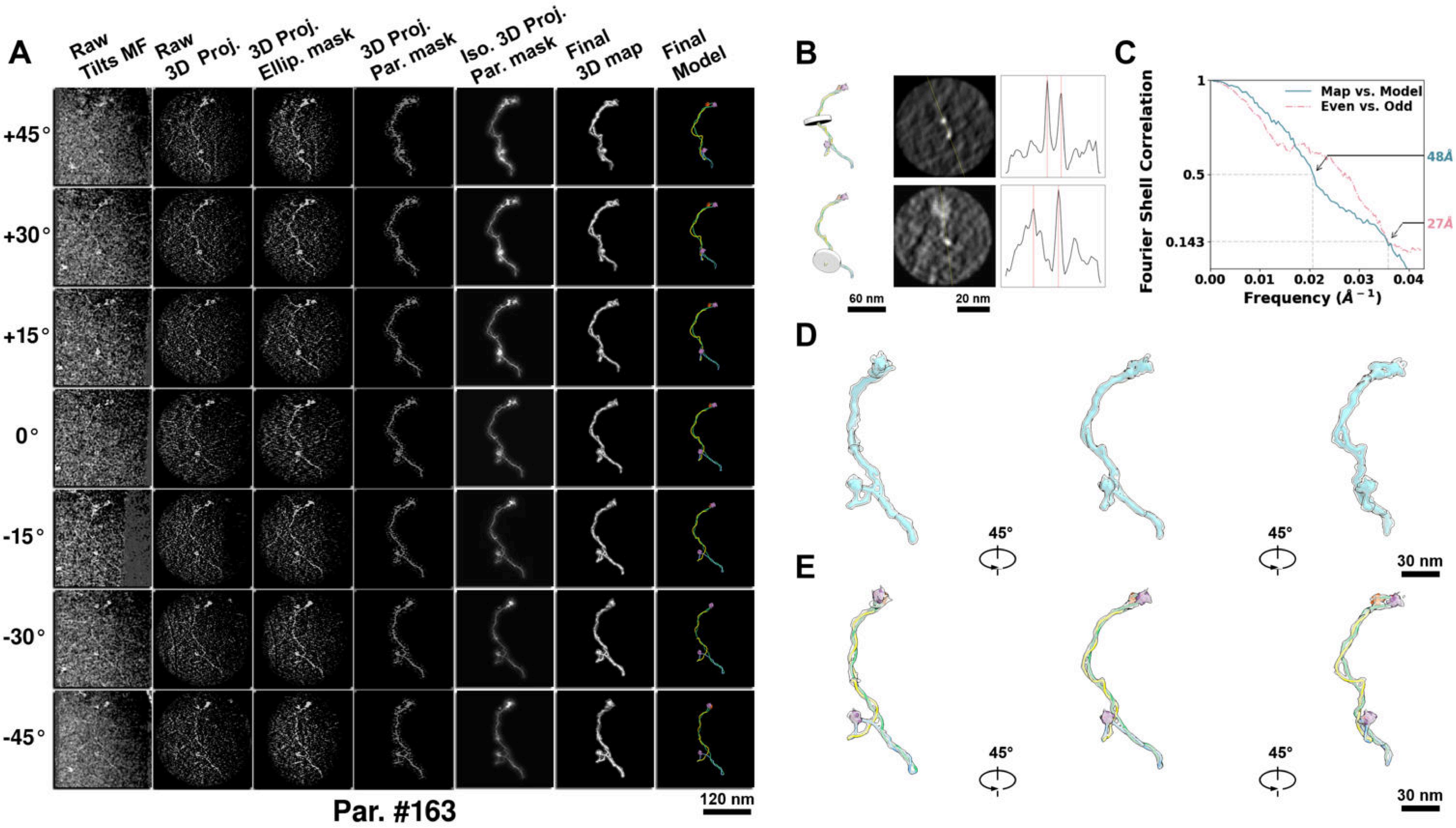

**Supplementary Particle Figure 163. Cryo-ET 3D reconstruction of an individual sTEC-Cas particle.**
(A) 3D reconstruction of the plasmid particle (index no. 163). The first column shows seven representative tilt images from +45° to -45° in step of 15°. The second, third, and fourth columns show 3D projections of the particle with spherical, ellipsoidal (thinner along the z-dimension), and particle-shaped masks, respectively. The fifth column displays the 3D projections of the enhanced and IsoNet missing-wedge-corrected particle. The sixth and seventh columns present the final 3D map and the flexibly fitted model, respectively. (B) Two cross-sectional views (12 nm thickness) of the plasmid density map along its plectoneme axis are shown in the left-middle panel. The intensity profile along the line crossing the two high-density DNA spots is displayed in the right panel. (C) Resolution assessment of the final 3D map using Fourier shell correlation (FSC). Two criteria are shown: FSC between two half-maps reconstructed from even and odd frames (evaluated at 0.143) and FSC between the final 3D map and the fitted model (evaluated at 0.5). (D) Zoomed-in views of the final 3D density map from panel A, displayed at two contour levels. (E) Superimposition of the high-contour level map from panel D onto its fitted model.

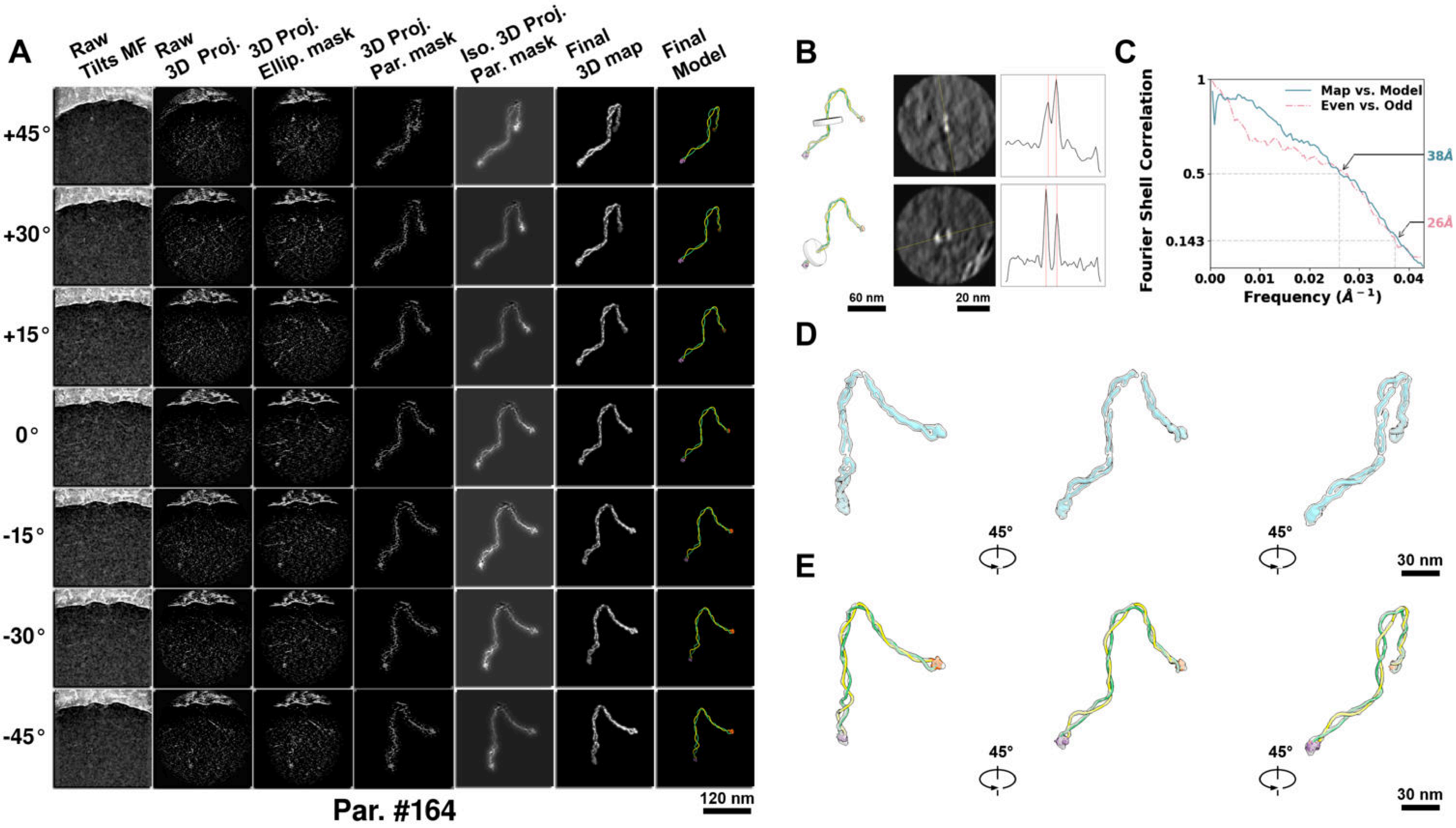

**Supplementary Particle Figure 164. Cryo-ET 3D reconstruction of an individual sTEC-Cas particle.**
(A) 3D reconstruction of the plasmid particle (index no. 164). The first column shows seven representative tilt images from +45° to -45° in step of 15°. The second, third, and fourth columns show 3D projections of the particle with spherical, ellipsoidal (thinner along the z-dimension), and particle-shaped masks, respectively. The fifth column displays the 3D projections of the enhanced and IsoNet missing-wedge-corrected particle. The sixth and seventh columns present the final 3D map and the flexibly fitted model, respectively. (B) Two cross-sectional views (12 nm thickness) of the plasmid density map along its plectoneme axis are shown in the left-middle panel. The intensity profile along the line crossing the two high-density DNA spots is displayed in the right panel. (C) Resolution assessment of the final 3D map using Fourier shell correlation (FSC). Two criteria are shown: FSC between two half-maps reconstructed from even and odd frames (evaluated at 0.143) and FSC between the final 3D map and the fitted model (evaluated at 0.5). (D) Zoomed-in views of the final 3D density map from panel A, displayed at two contour levels. (E) Superimposition of the high-contour level map from panel D onto its fitted model.

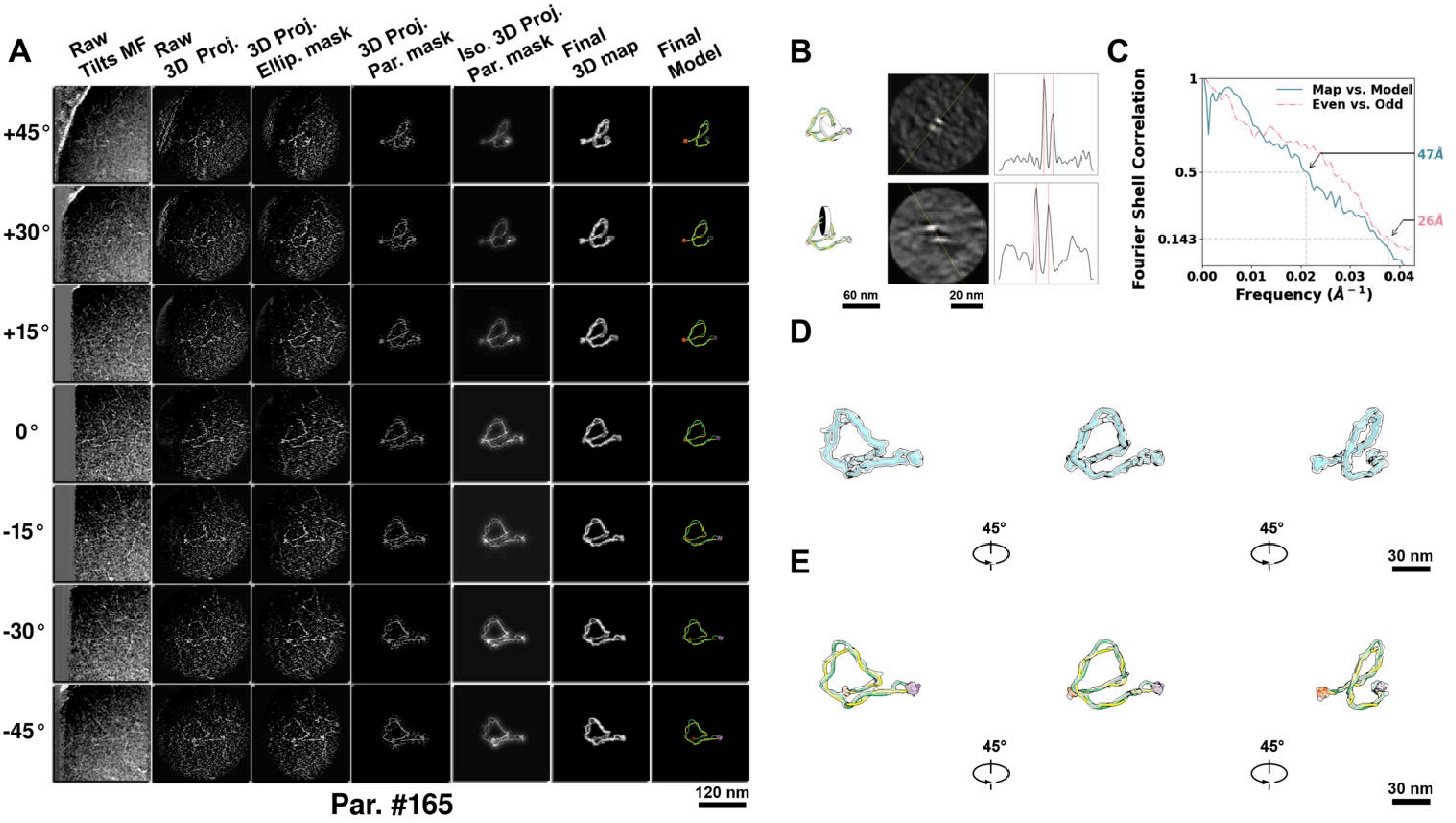

**Supplementary Particle Figure 165. Cryo-ET 3D reconstruction of an individual sTEC-Cas particle.**
(A) 3D reconstruction of the plasmid particle (index no. 165). The first column shows seven representative tilt images from +45° to -45° in step of 15°. The second, third, and fourth columns show 3D projections of the particle with spherical, ellipsoidal (thinner along the z-dimension), and particle-shaped masks, respectively. The fifth column displays the 3D projections of the enhanced and IsoNet missing-wedge-corrected particle. The sixth and seventh columns present the final 3D map and the flexibly fitted model, respectively. (B) Two cross-sectional views (12 nm thickness) of the plasmid density map along its plectoneme axis are shown in the left-middle panel. The intensity profile along the line crossing the two high-density DNA spots is displayed in the right panel. (C) Resolution assessment of the final 3D map using Fourier shell correlation (FSC). Two criteria are shown: FSC between two half-maps reconstructed from even and odd frames (evaluated at 0.143) and FSC between the final 3D map and the fitted model (evaluated at 0.5). (D) Zoomed-in views of the final 3D density map from panel A, displayed at two contour levels. (E) Superimposition of the high-contour level map from panel D onto its fitted model.

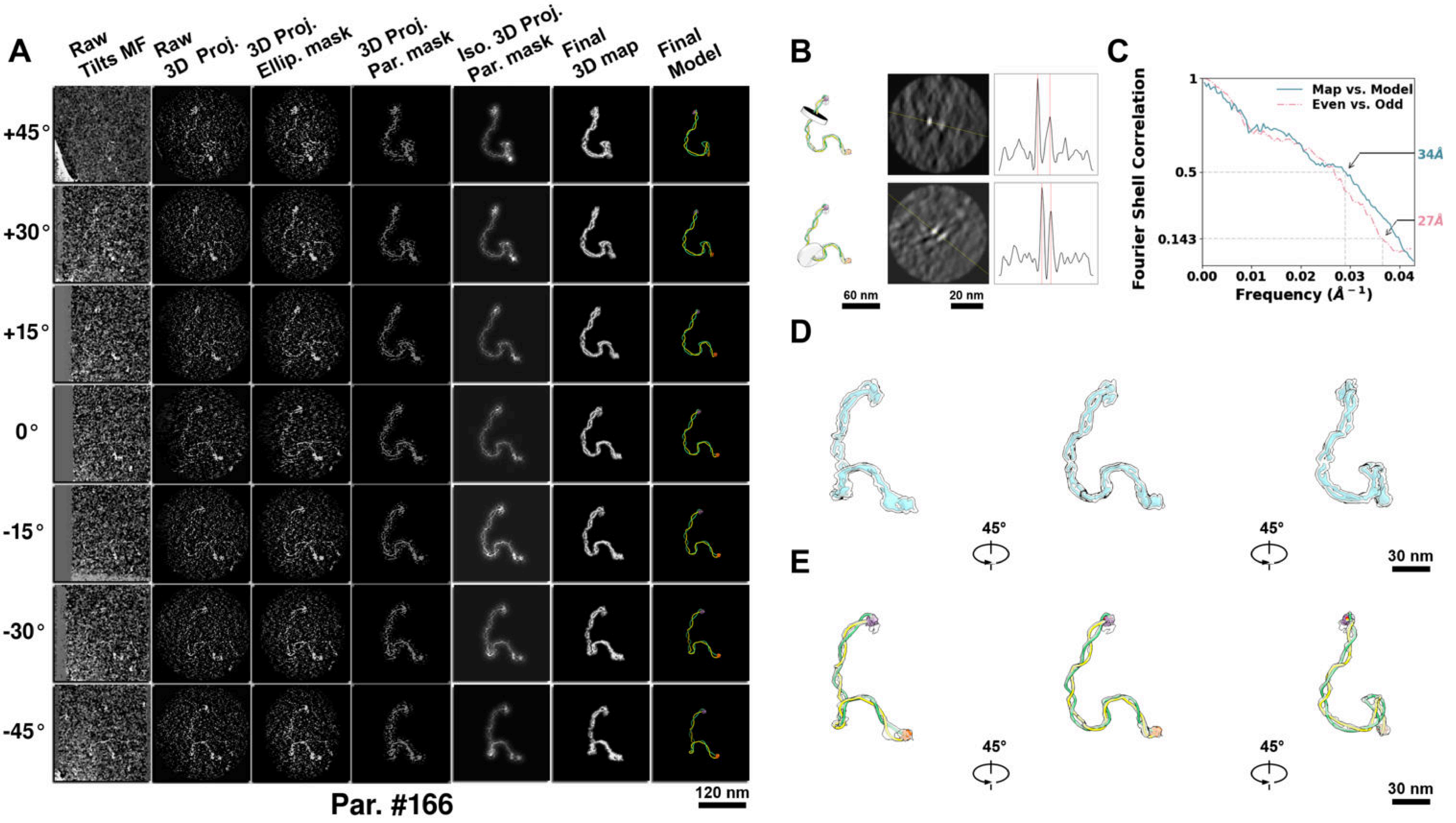

**Supplementary Particle Figure 166. Cryo-ET 3D reconstruction of an individual sTEC-Cas particle.**
(A) 3D reconstruction of the plasmid particle (index no. 166). The first column shows seven representative tilt images from +45° to -45° in step of 15°. The second, third, and fourth columns show 3D projections of the particle with spherical, ellipsoidal (thinner along the z-dimension), and particle-shaped masks, respectively. The fifth column displays the 3D projections of the enhanced and IsoNet missing-wedge-corrected particle. The sixth and seventh columns present the final 3D map and the flexibly fitted model, respectively. (B) Two cross-sectional views (12 nm thickness) of the plasmid density map along its plectoneme axis are shown in the left-middle panel. The intensity profile along the line crossing the two high-density DNA spots is displayed in the right panel. (C) Resolution assessment of the final 3D map using Fourier shell correlation (FSC). Two criteria are shown: FSC between two half-maps reconstructed from even and odd frames (evaluated at 0.143) and FSC between the final 3D map and the fitted model (evaluated at 0.5). (D) Zoomed-in views of the final 3D density map from panel A, displayed at two contour levels. (E) Superimposition of the high-contour level map from panel D onto its fitted model.

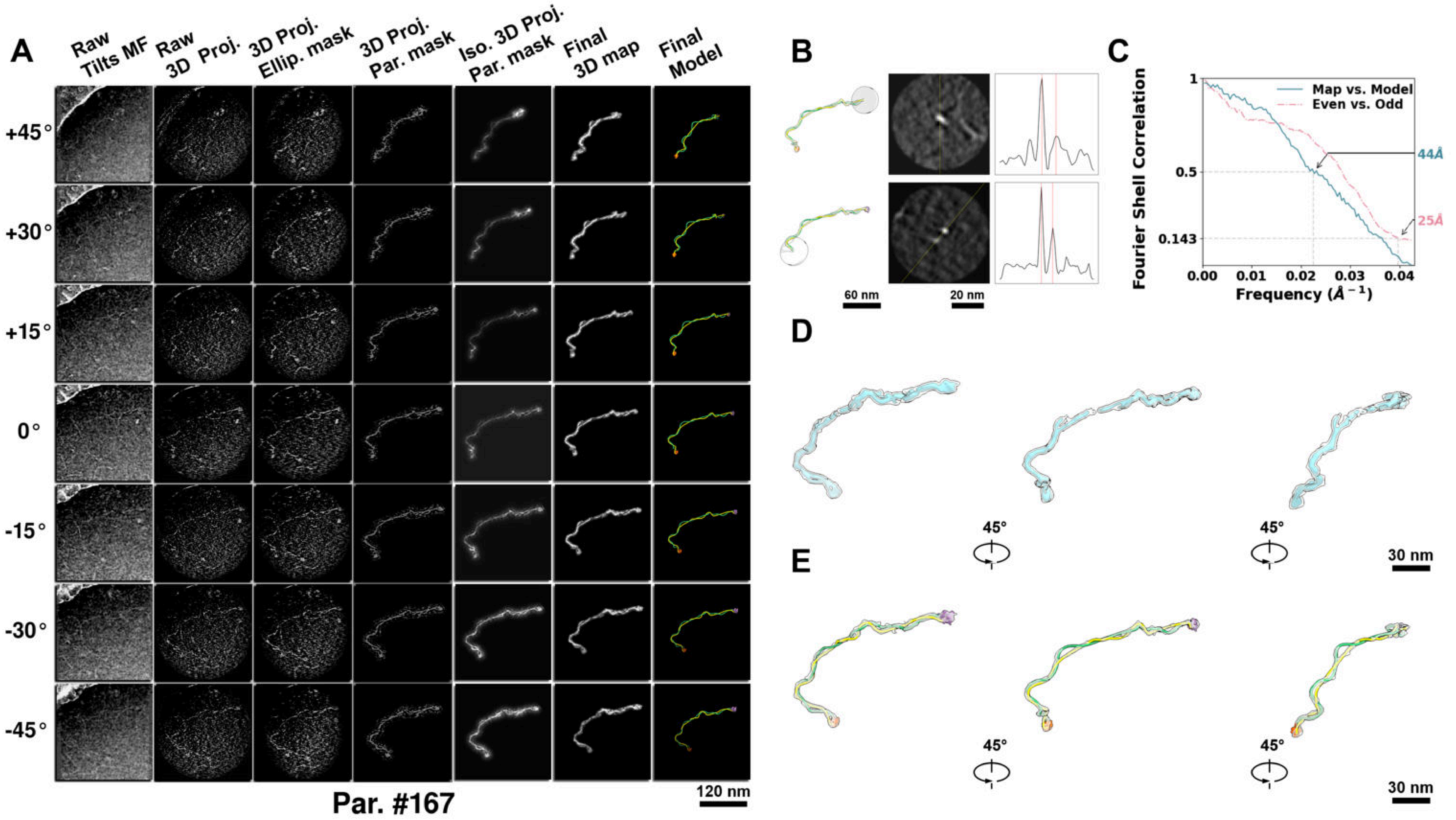

**Supplementary Particle Figure 167. Cryo-ET 3D reconstruction of an individual sTEC-Cas particle.**
(A) 3D reconstruction of the plasmid particle (index no. 167). The first column shows seven representative tilt images from +45° to -45° in step of 15°. The second, third, and fourth columns show 3D projections of the particle with spherical, ellipsoidal (thinner along the z-dimension), and particle-shaped masks, respectively. The fifth column displays the 3D projections of the enhanced and IsoNet missing-wedge-corrected particle. The sixth and seventh columns present the final 3D map and the flexibly fitted model, respectively. (B) Two cross-sectional views (12 nm thickness) of the plasmid density map along its plectoneme axis are shown in the left-middle panel. The intensity profile along the line crossing the two high-density DNA spots is displayed in the right panel. (C) Resolution assessment of the final 3D map using Fourier shell correlation (FSC). Two criteria are shown: FSC between two half-maps reconstructed from even and odd frames (evaluated at 0.143) and FSC between the final 3D map and the fitted model (evaluated at 0.5). (D) Zoomed-in views of the final 3D density map from panel A, displayed at two contour levels. (E) Superimposition of the high-contour level map from panel D onto its fitted model.

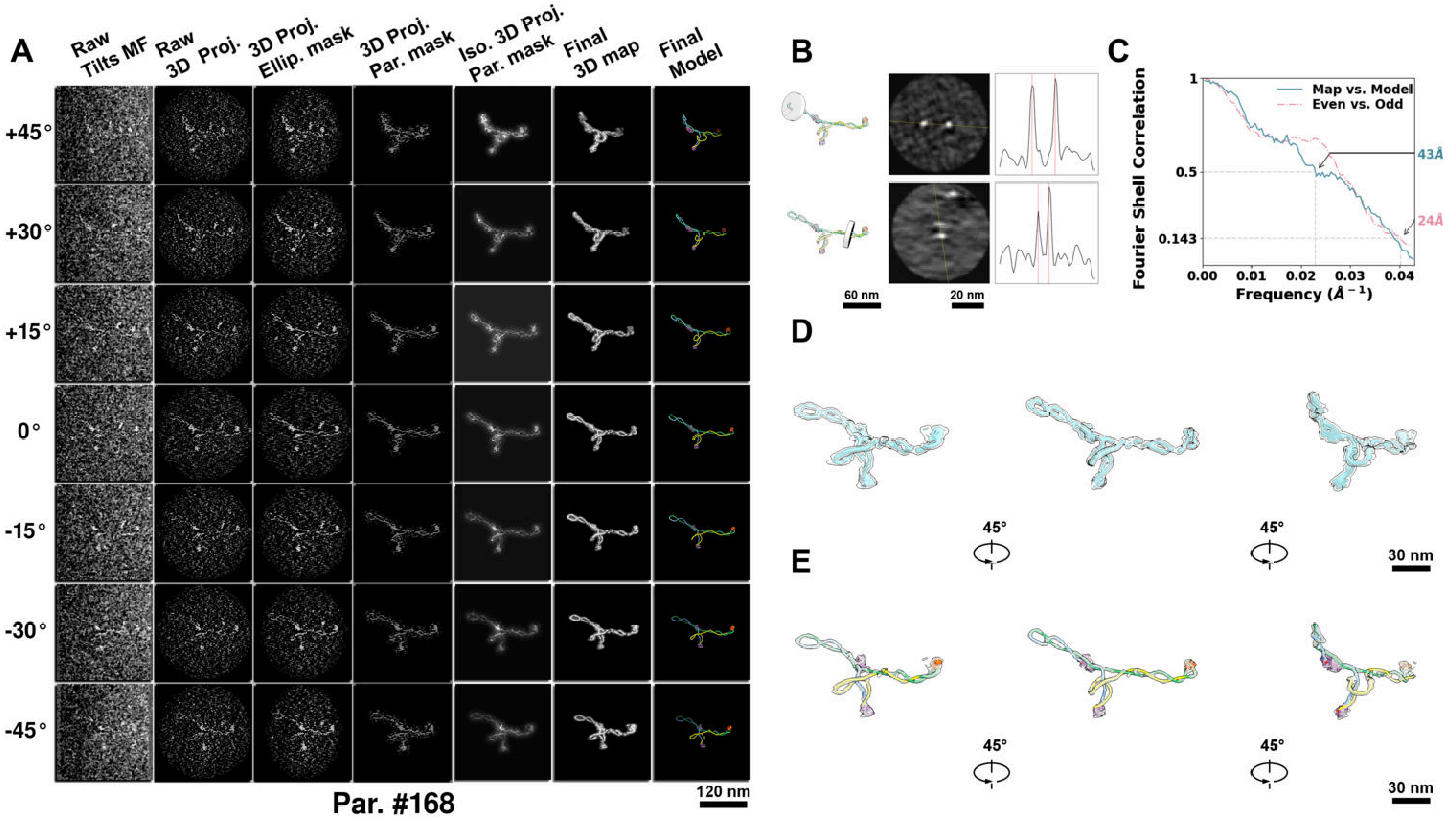

**Supplementary Particle Figure 168. Cryo-ET 3D reconstruction of an individual sTEC-Cas particle.**
(A) 3D reconstruction of the plasmid particle (index no. 168). The first column shows seven representative tilt images from +45° to -45° in step of 15°. The second, third, and fourth columns show 3D projections of the particle with spherical, ellipsoidal (thinner along the z-dimension), and particle-shaped masks, respectively. The fifth column displays the 3D projections of the enhanced and IsoNet missing-wedge-corrected particle. The sixth and seventh columns present the final 3D map and the flexibly fitted model, respectively. (B) Two cross-sectional views (12 nm thickness) of the plasmid density map along its plectoneme axis are shown in the left-middle panel. The intensity profile along the line crossing the two high-density DNA spots is displayed in the right panel. (C) Resolution assessment of the final 3D map using Fourier shell correlation (FSC). Two criteria are shown: FSC between two half-maps reconstructed from even and odd frames (evaluated at 0.143) and FSC between the final 3D map and the fitted model (evaluated at 0.5). (D) Zoomed-in views of the final 3D density map from panel A, displayed at two contour levels. (E) Superimposition of the high-contour level map from panel D onto its fitted model.

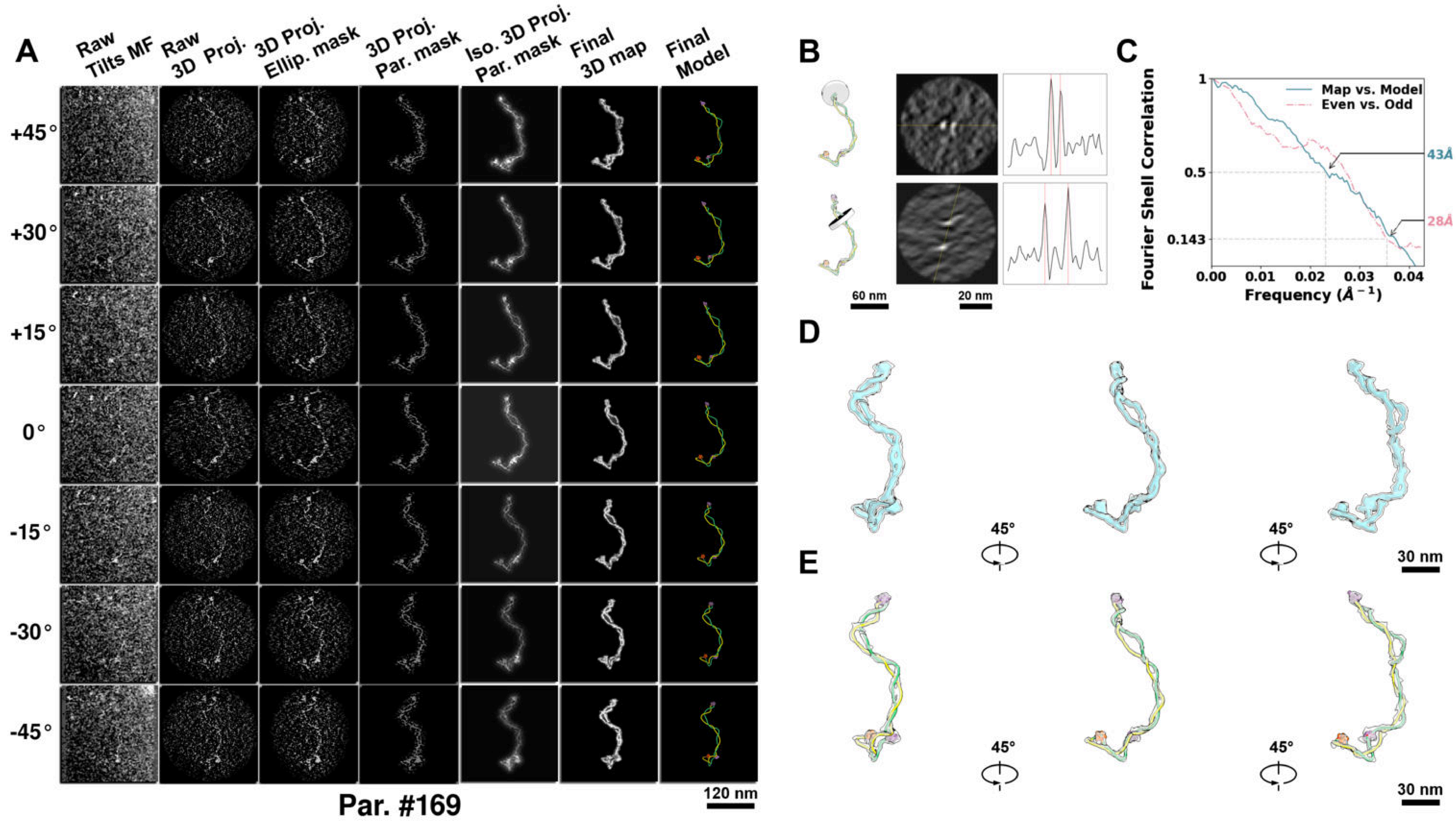

**Supplementary Particle Figure 169. Cryo-ET 3D reconstruction of an individual sTEC-Cas particle.**
(A) 3D reconstruction of the plasmid particle (index no. 169). The first column shows seven representative tilt images from +45° to -45° in step of 15°. The second, third, and fourth columns show 3D projections of the particle with spherical, ellipsoidal (thinner along the z-dimension), and particle-shaped masks, respectively. The fifth column displays the 3D projections of the enhanced and IsoNet missing-wedge-corrected particle. The sixth and seventh columns present the final 3D map and the flexibly fitted model, respectively. (B) Two cross-sectional views (12 nm thickness) of the plasmid density map along its plectoneme axis are shown in the left-middle panel. The intensity profile along the line crossing the two high-density DNA spots is displayed in the right panel. (C) Resolution assessment of the final 3D map using Fourier shell correlation (FSC). Two criteria are shown: FSC between two half-maps reconstructed from even and odd frames (evaluated at 0.143) and FSC between the final 3D map and the fitted model (evaluated at 0.5). (D) Zoomed-in views of the final 3D density map from panel A, displayed at two contour levels. (E) Superimposition of the high-contour level map from panel D onto its fitted model.

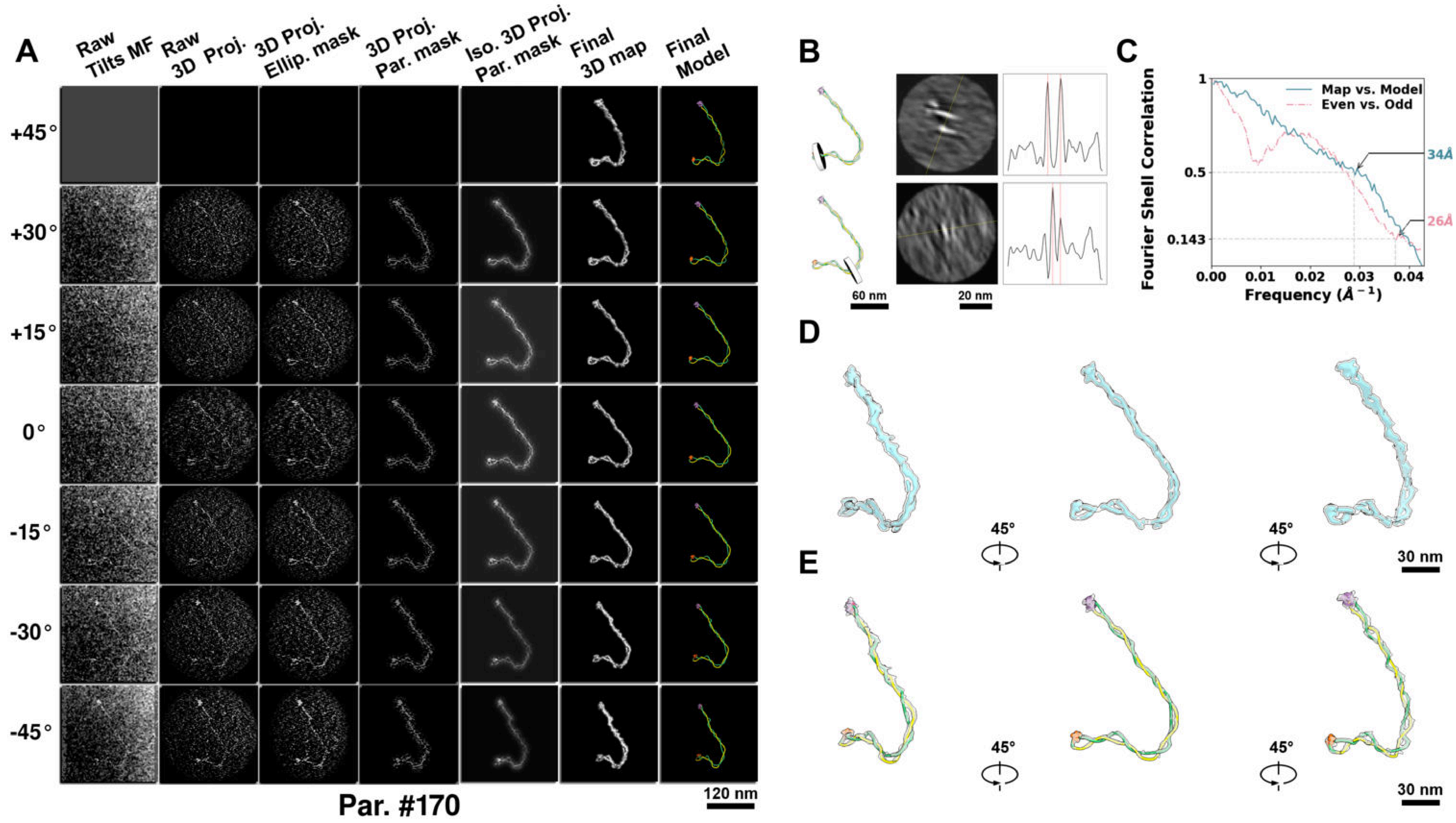

**Supplementary Particle Figure 170. Cryo-ET 3D reconstruction of an individual sTEC-Cas particle.**
(A) 3D reconstruction of the plasmid particle (index no. 170). The first column shows seven representative tilt images from +45° to -45° in step of 15°. The second, third, and fourth columns show 3D projections of the particle with spherical, ellipsoidal (thinner along the z-dimension), and particle-shaped masks, respectively. The fifth column displays the 3D projections of the enhanced and IsoNet missing-wedge-corrected particle. The sixth and seventh columns present the final 3D map and the flexibly fitted model, respectively. (B) Two cross-sectional views (12 nm thickness) of the plasmid density map along its plectoneme axis are shown in the left-middle panel. The intensity profile along the line crossing the two high-density DNA spots is displayed in the right panel. (C) Resolution assessment of the final 3D map using Fourier shell correlation (FSC). Two criteria are shown: FSC between two half-maps reconstructed from even and odd frames (evaluated at 0.143) and FSC between the final 3D map and the fitted model (evaluated at 0.5). (D) Zoomed-in views of the final 3D density map from panel A, displayed at two contour levels. (E) Superimposition of the high-contour level map from panel D onto its fitted model.

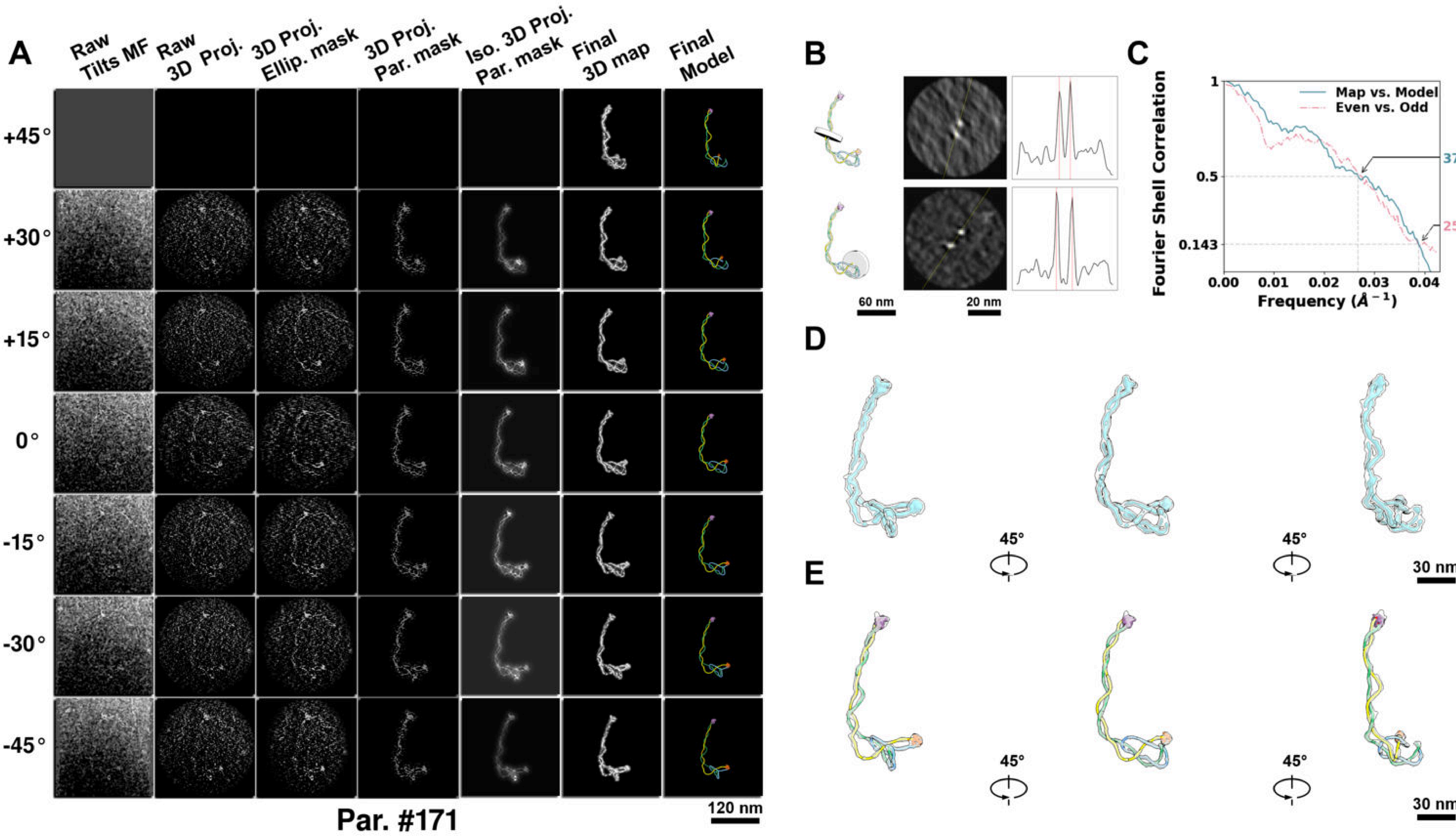

**Supplementary Particle Figure 171. Cryo-ET 3D reconstruction of an individual sTEC-Cas particle.**
(A) 3D reconstruction of the plasmid particle (index no. 171). The first column shows seven representative tilt images from +45° to -45° in step of 15°. The second, third, and fourth columns show 3D projections of the particle with spherical, ellipsoidal (thinner along the z-dimension), and particle-shaped masks, respectively. The fifth column displays the 3D projections of the enhanced and IsoNet missing-wedge-corrected particle. The sixth and seventh columns present the final 3D map and the flexibly fitted model, respectively. (B) Two cross-sectional views (12 nm thickness) of the plasmid density map along its plectoneme axis are shown in the left-middle panel. The intensity profile along the line crossing the two high-density DNA spots is displayed in the right panel. (C) Resolution assessment of the final 3D map using Fourier shell correlation (FSC). Two criteria are shown: FSC between two half-maps reconstructed from even and odd frames (evaluated at 0.143) and FSC between the final 3D map and the fitted model (evaluated at 0.5). (D) Zoomed-in views of the final 3D density map from panel A, displayed at two contour levels. (E) Superimposition of the high-contour level map from panel D onto its fitted model.

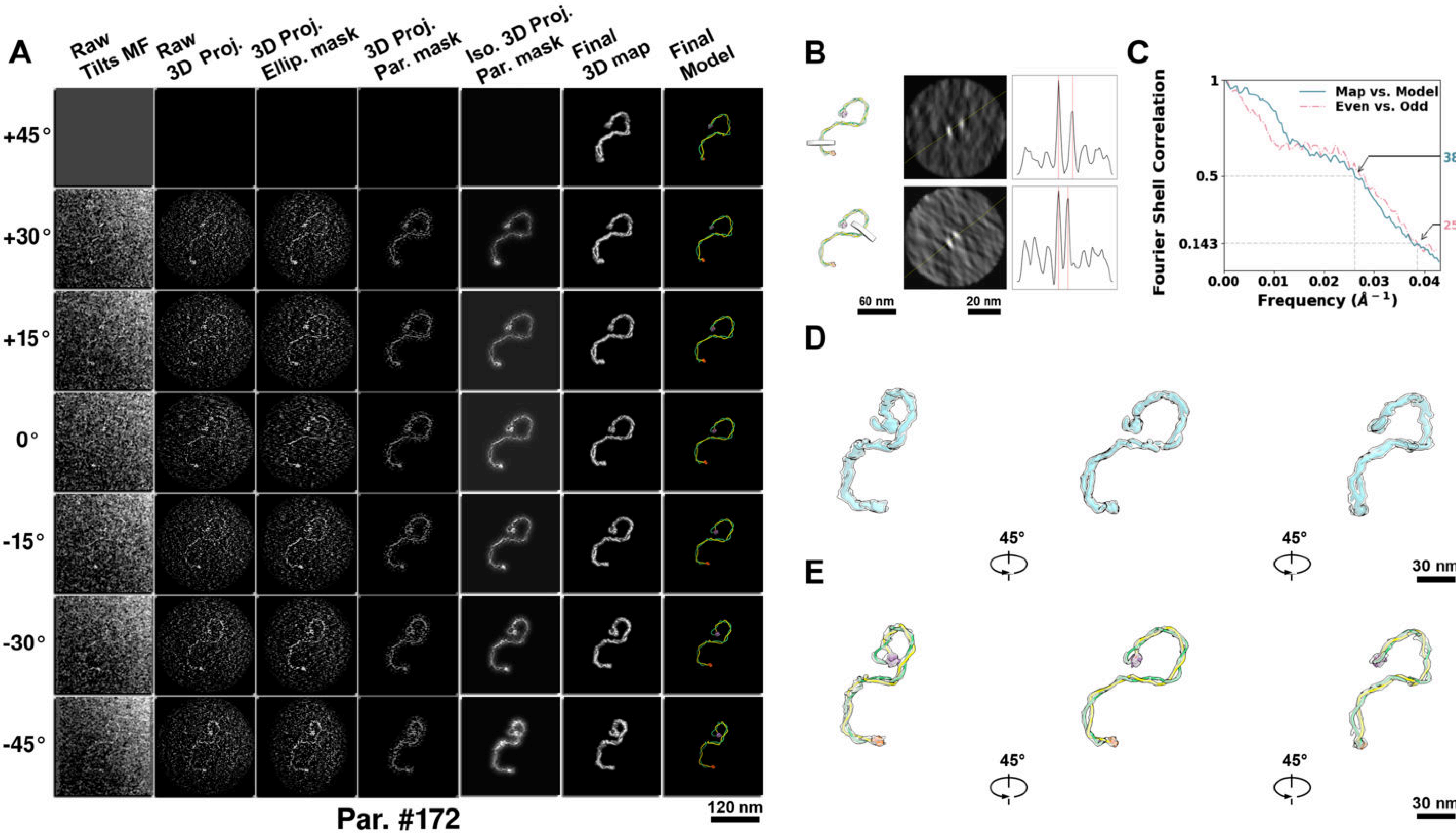

**Supplementary Particle Figure 172. Cryo-ET 3D reconstruction of an individual sTEC-Cas particle.**
(A) 3D reconstruction of the plasmid particle (index no. 172). The first column shows seven representative tilt images from +45° to -45° in step of 15°. The second, third, and fourth columns show 3D projections of the particle with spherical, ellipsoidal (thinner along the z-dimension), and particle-shaped masks, respectively. The fifth column displays the 3D projections of the enhanced and IsoNet missing-wedge-corrected particle. The sixth and seventh columns present the final 3D map and the flexibly fitted model, respectively. (B) Two cross-sectional views (12 nm thickness) of the plasmid density map along its plectoneme axis are shown in the left-middle panel. The intensity profile along the line crossing the two high-density DNA spots is displayed in the right panel. (C) Resolution assessment of the final 3D map using Fourier shell correlation (FSC). Two criteria are shown: FSC between two half-maps reconstructed from even and odd frames (evaluated at 0.143) and FSC between the final 3D map and the fitted model (evaluated at 0.5). (D) Zoomed-in views of the final 3D density map from panel A, displayed at two contour levels. (E) Superimposition of the high-contour level map from panel D onto its fitted model.

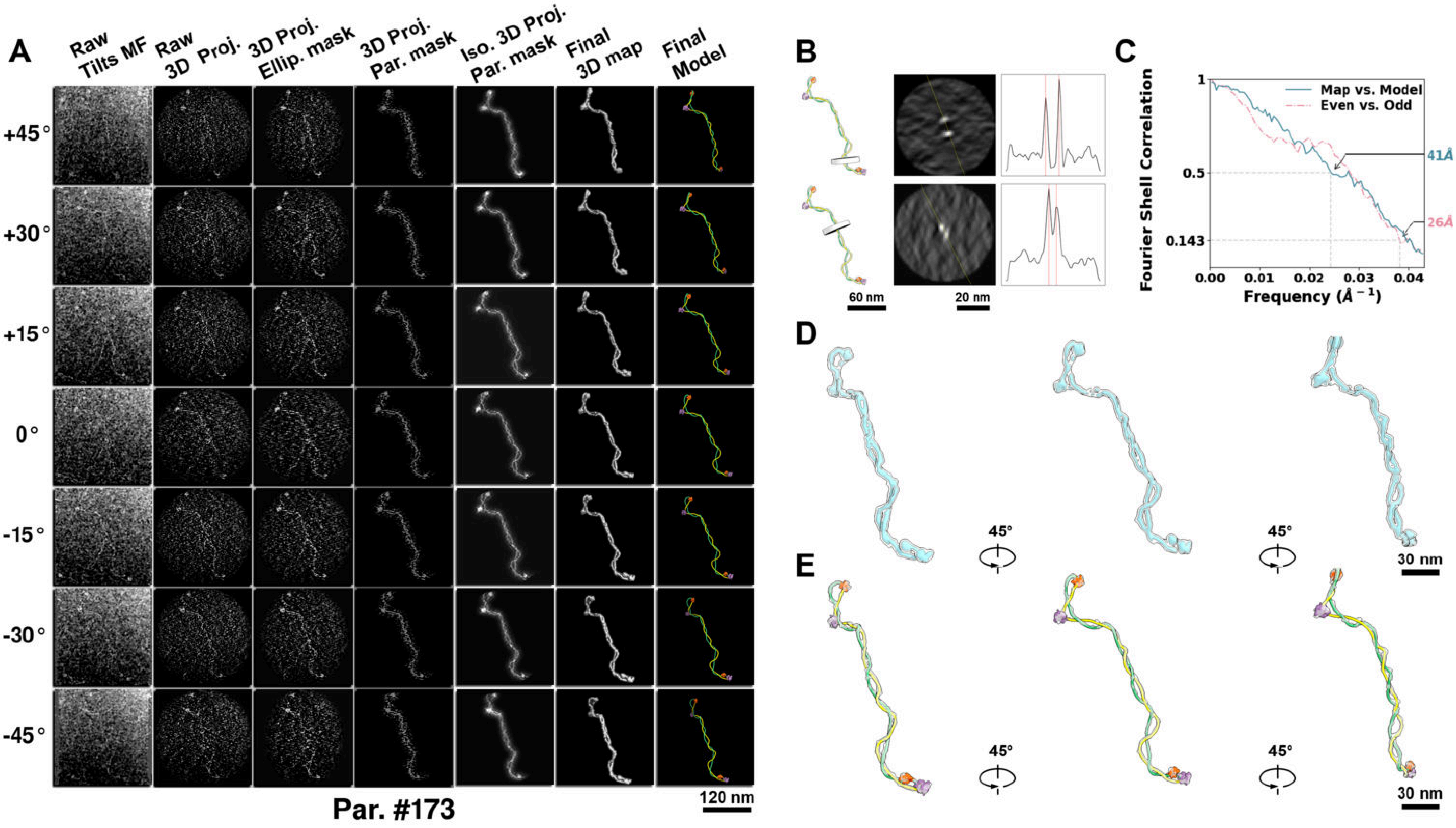

**Supplementary Particle Figure 173. Cryo-ET 3D reconstruction of an individual sTEC-Cas particle.**
(A) 3D reconstruction of the plasmid particle (index no. 173). The first column shows seven representative tilt images from +45° to -45° in step of 15°. The second, third, and fourth columns show 3D projections of the particle with spherical, ellipsoidal (thinner along the z-dimension), and particle-shaped masks, respectively. The fifth column displays the 3D projections of the enhanced and IsoNet missing-wedge-corrected particle. The sixth and seventh columns present the final 3D map and the flexibly fitted model, respectively. (B) Two cross-sectional views (12 nm thickness) of the plasmid density map along its plectoneme axis are shown in the left-middle panel. The intensity profile along the line crossing the two high-density DNA spots is displayed in the right panel. (C) Resolution assessment of the final 3D map using Fourier shell correlation (FSC). Two criteria are shown: FSC between two half-maps reconstructed from even and odd frames (evaluated at 0.143) and FSC between the final 3D map and the fitted model (evaluated at 0.5). (D) Zoomed-in views of the final 3D density map from panel A, displayed at two contour levels. (E) Superimposition of the high-contour level map from panel D onto its fitted model.

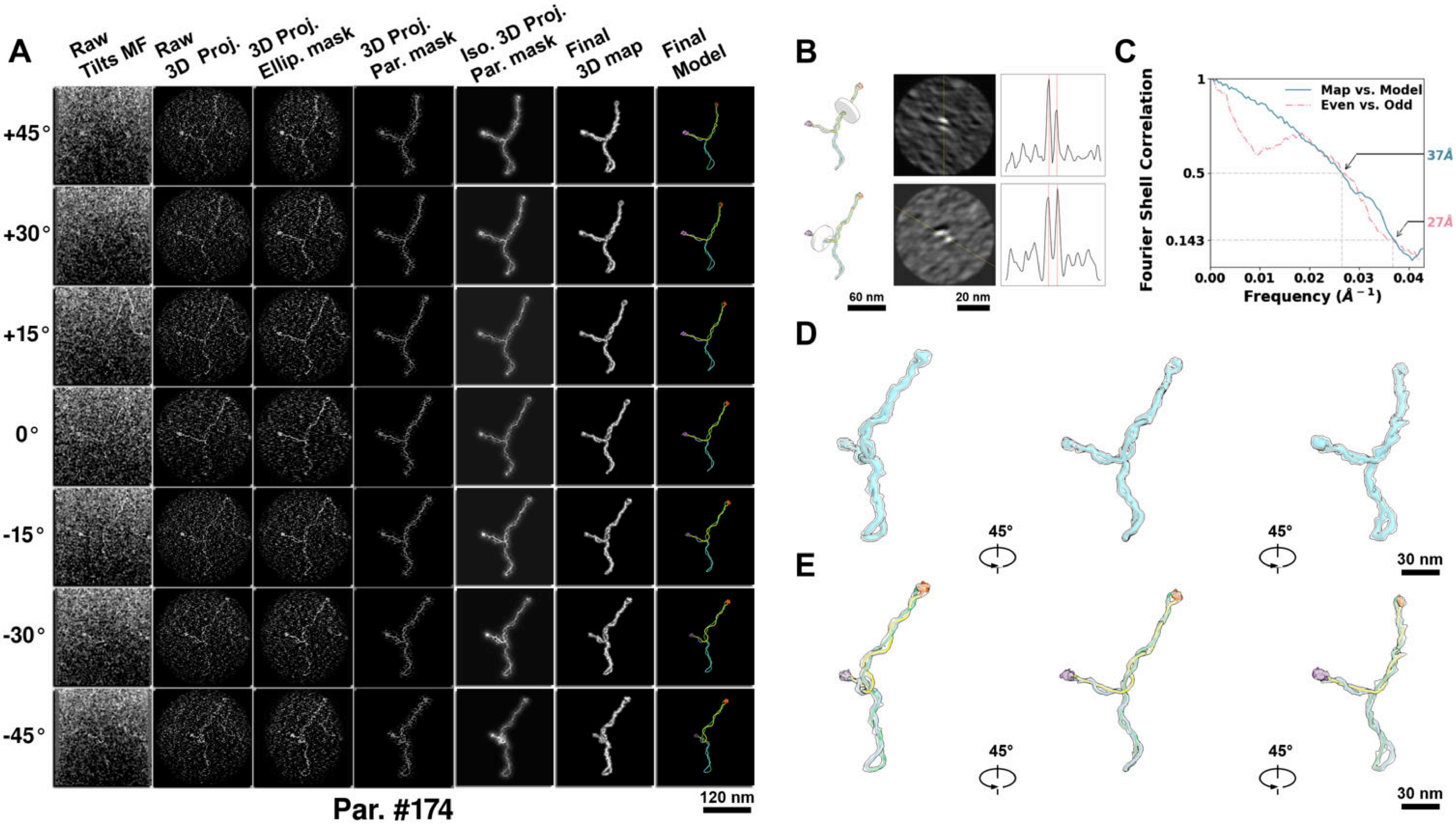

**Supplementary Particle Figure 174. Cryo-ET 3D reconstruction of an individual sTEC-Cas particle.**
(A) 3D reconstruction of the plasmid particle (index no. 174). The first column shows seven representative tilt images from +45° to -45° in step of 15°. The second, third, and fourth columns show 3D projections of the particle with spherical, ellipsoidal (thinner along the z-dimension), and particle-shaped masks, respectively. The fifth column displays the 3D projections of the enhanced and IsoNet missing-wedge-corrected particle. The sixth and seventh columns present the final 3D map and the flexibly fitted model, respectively. (B) Two cross-sectional views (12 nm thickness) of the plasmid density map along its plectoneme axis are shown in the left-middle panel. The intensity profile along the line crossing the two high-density DNA spots is displayed in the right panel. (C) Resolution assessment of the final 3D map using Fourier shell correlation (FSC). Two criteria are shown: FSC between two half-maps reconstructed from even and odd frames (evaluated at 0.143) and FSC between the final 3D map and the fitted model (evaluated at 0.5). (D) Zoomed-in views of the final 3D density map from panel A, displayed at two contour levels. (E) Superimposition of the high-contour level map from panel D onto its fitted model.

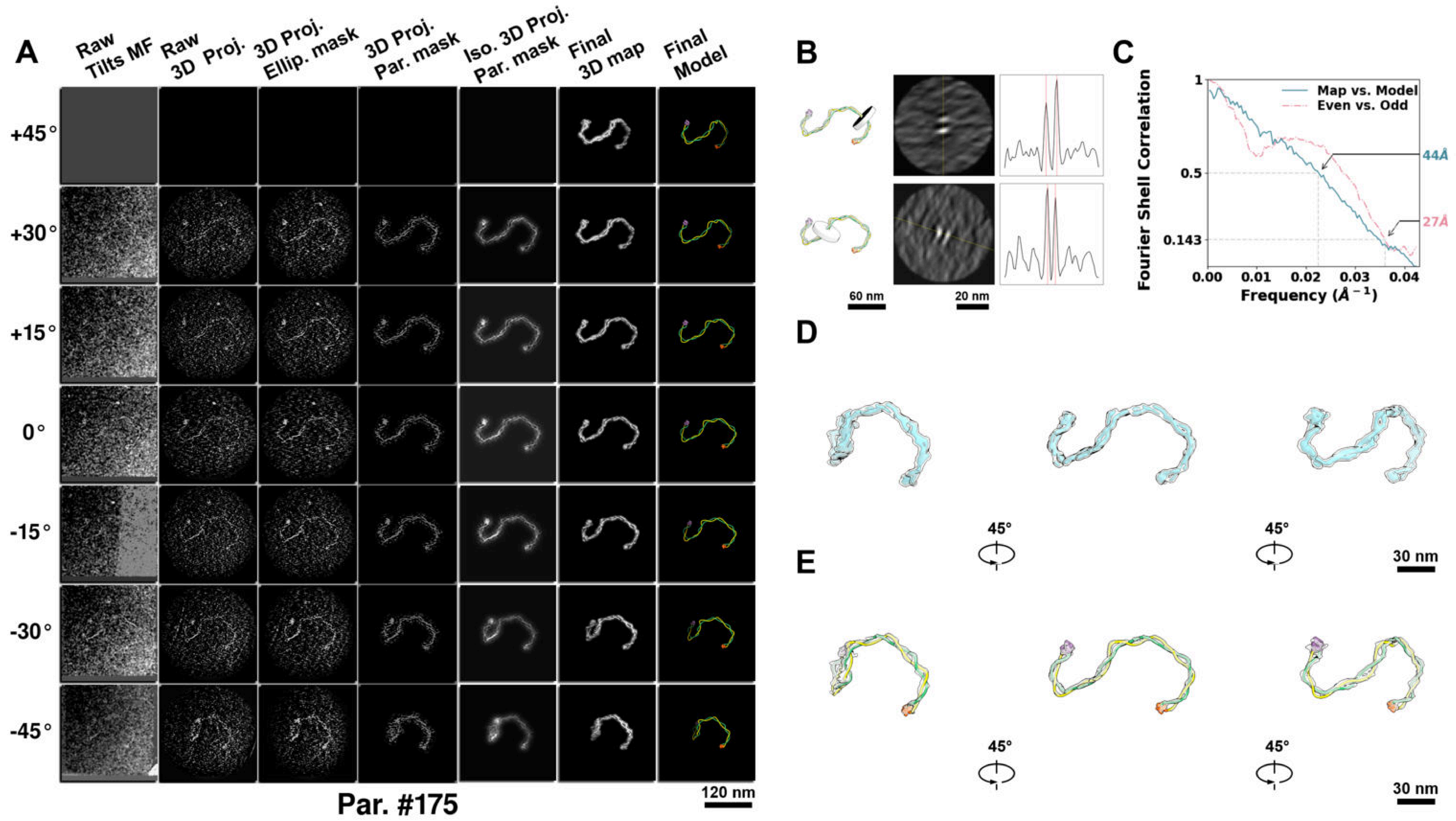

**Supplementary Particle Figure 175. Cryo-ET 3D reconstruction of an individual sTEC-Cas particle.**
(A) 3D reconstruction of the plasmid particle (index no. 175). The first column shows seven representative tilt images from +45° to -45° in step of 15°. The second, third, and fourth columns show 3D projections of the particle with spherical, ellipsoidal (thinner along the z-dimension), and particle-shaped masks, respectively. The fifth column displays the 3D projections of the enhanced and IsoNet missing-wedge-corrected particle. The sixth and seventh columns present the final 3D map and the flexibly fitted model, respectively. (B) Two cross-sectional views (12 nm thickness) of the plasmid density map along its plectoneme axis are shown in the left-middle panel. The intensity profile along the line crossing the two high-density DNA spots is displayed in the right panel. (C) Resolution assessment of the final 3D map using Fourier shell correlation (FSC). Two criteria are shown: FSC between two half-maps reconstructed from even and odd frames (evaluated at 0.143) and FSC between the final 3D map and the fitted model (evaluated at 0.5). (D) Zoomed-in views of the final 3D density map from panel A, displayed at two contour levels. (E) Superimposition of the high-contour level map from panel D onto its fitted model.

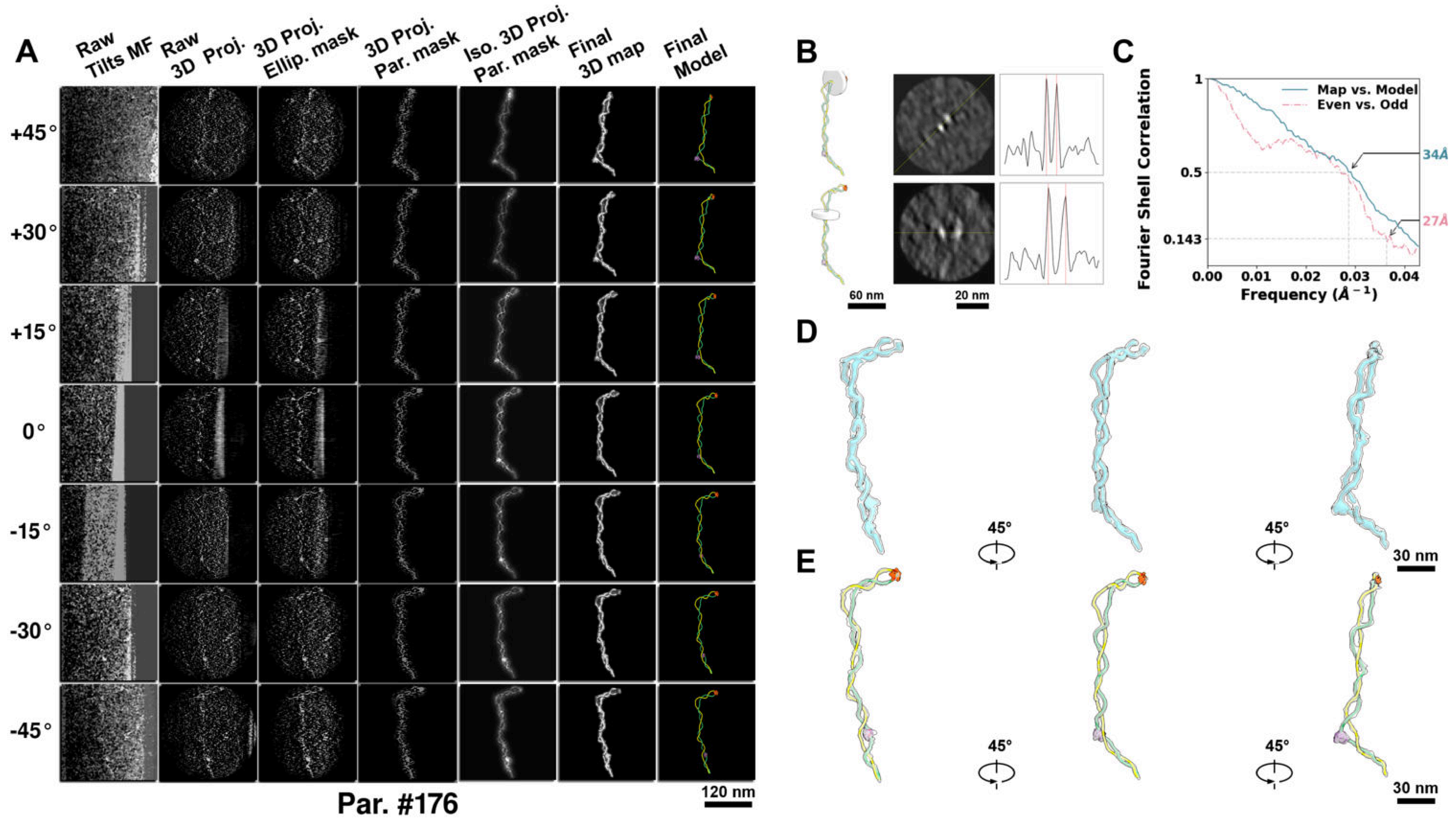

**Supplementary Particle Figure 176. Cryo-ET 3D reconstruction of an individual sTEC-Cas particle.**
(A) 3D reconstruction of the plasmid particle (index no. 176). The first column shows seven representative tilt images from +45° to -45° in step of 15°. The second, third, and fourth columns show 3D projections of the particle with spherical, ellipsoidal (thinner along the z-dimension), and particle-shaped masks, respectively. The fifth column displays the 3D projections of the enhanced and IsoNet missing-wedge-corrected particle. The sixth and seventh columns present the final 3D map and the flexibly fitted model, respectively. (B) Two cross-sectional views (12 nm thickness) of the plasmid density map along its plectoneme axis are shown in the left-middle panel. The intensity profile along the line crossing the two high-density DNA spots is displayed in the right panel. (C) Resolution assessment of the final 3D map using Fourier shell correlation (FSC). Two criteria are shown: FSC between two half-maps reconstructed from even and odd frames (evaluated at 0.143) and FSC between the final 3D map and the fitted model (evaluated at 0.5). (D) Zoomed-in views of the final 3D density map from panel A, displayed at two contour levels. (E) Superimposition of the high-contour level map from panel D onto its fitted model.

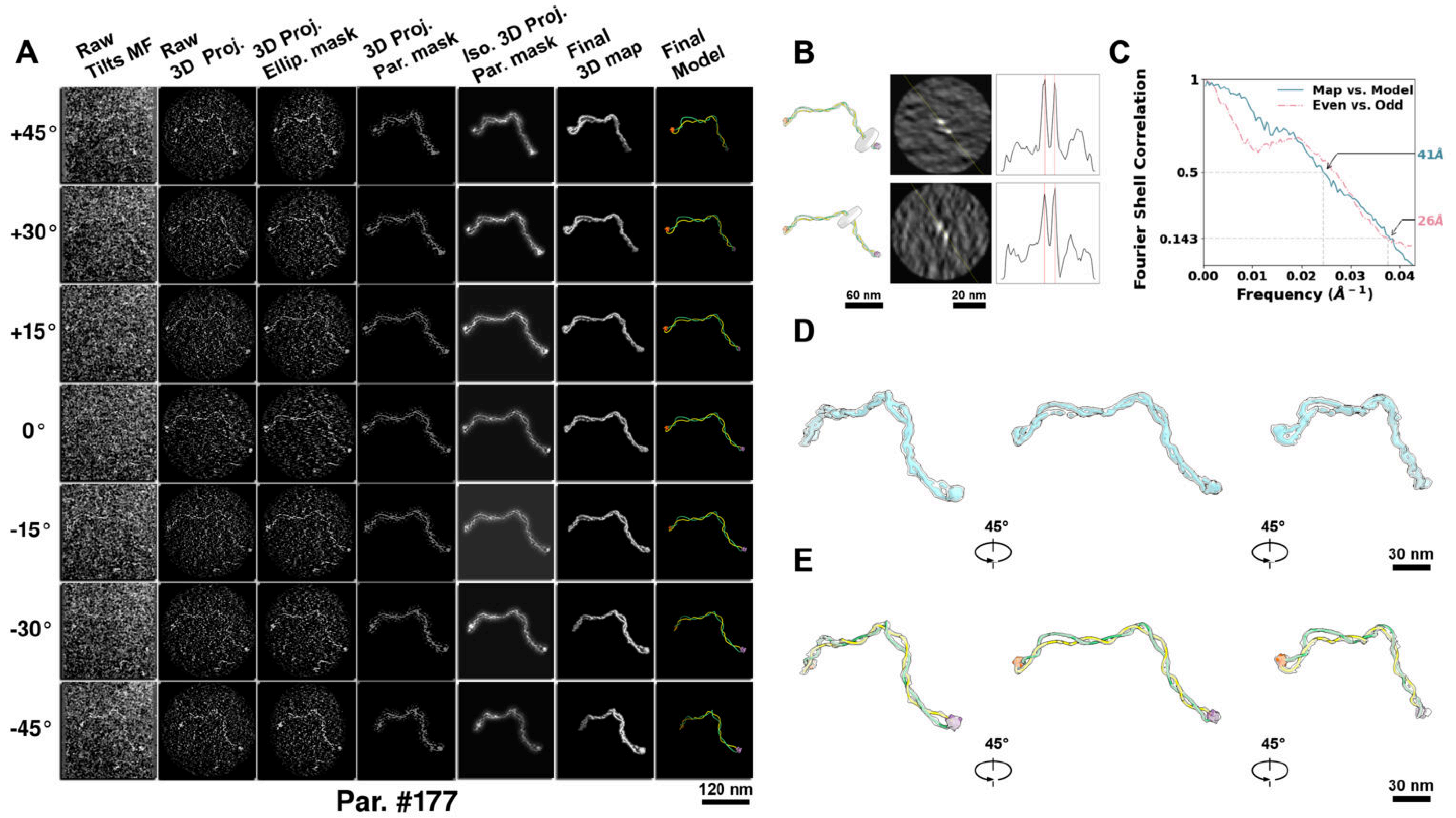

**Supplementary Particle Figure 177. Cryo-ET 3D reconstruction of an individual sTEC-Cas particle.**
(A) 3D reconstruction of the plasmid particle (index no. 177). The first column shows seven representative tilt images from +45° to -45° in step of 15°. The second, third, and fourth columns show 3D projections of the particle with spherical, ellipsoidal (thinner along the z-dimension), and particle-shaped masks, respectively. The fifth column displays the 3D projections of the enhanced and IsoNet missing-wedge-corrected particle. The sixth and seventh columns present the final 3D map and the flexibly fitted model, respectively. (B) Two cross-sectional views (12 nm thickness) of the plasmid density map along its plectoneme axis are shown in the left-middle panel. The intensity profile along the line crossing the two high-density DNA spots is displayed in the right panel. (C) Resolution assessment of the final 3D map using Fourier shell correlation (FSC). Two criteria are shown: FSC between two half-maps reconstructed from even and odd frames (evaluated at 0.143) and FSC between the final 3D map and the fitted model (evaluated at 0.5). (D) Zoomed-in views of the final 3D density map from panel A, displayed at two contour levels. (E) Superimposition of the high-contour level map from panel D onto its fitted model.

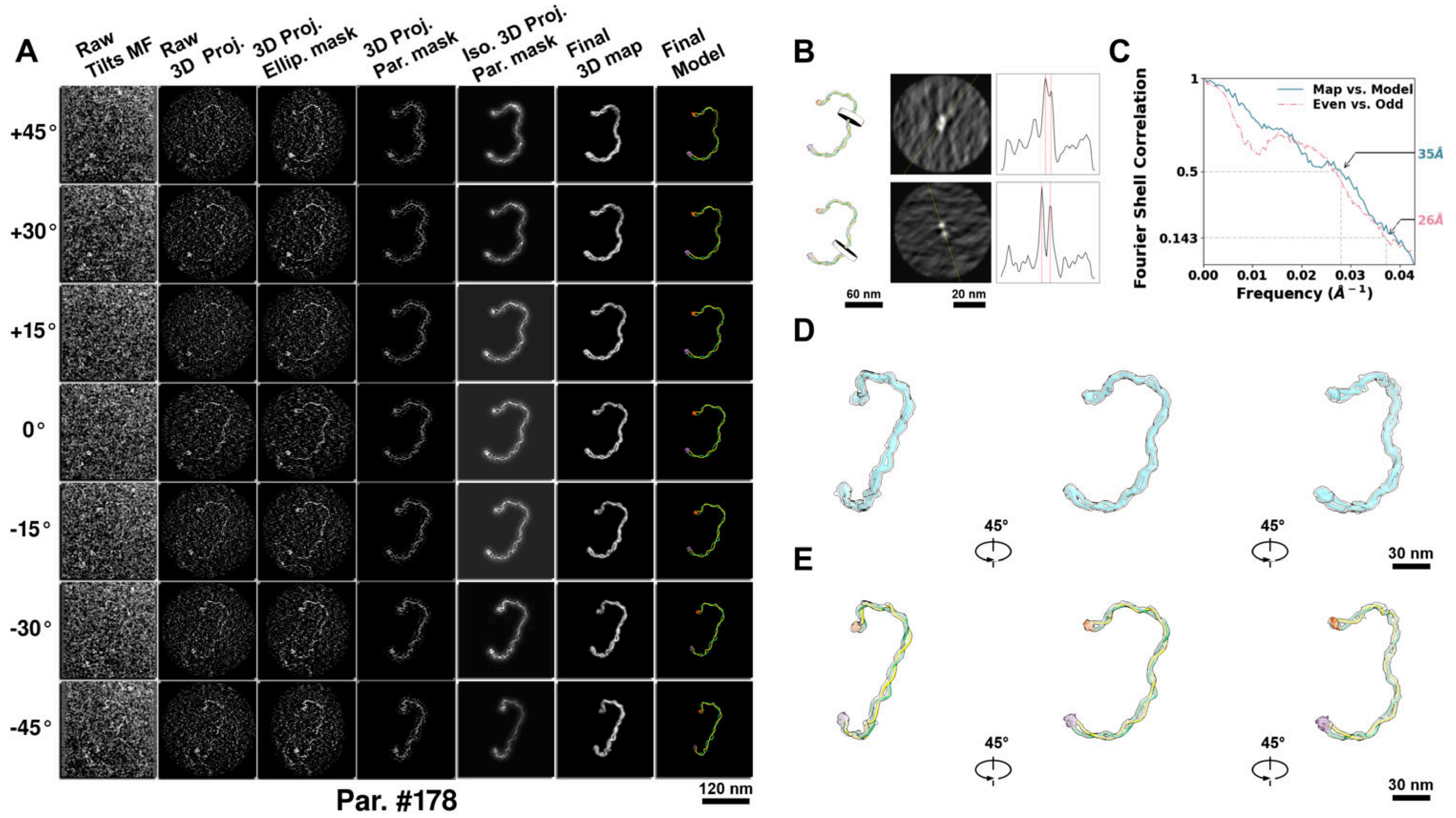

**Supplementary Particle Figure 178. Cryo-ET 3D reconstruction of an individual sTEC-Cas particle.**
(A) 3D reconstruction of the plasmid particle (index no. 178). The first column shows seven representative tilt images from +45° to -45° in step of 15°. The second, third, and fourth columns show 3D projections of the particle with spherical, ellipsoidal (thinner along the z-dimension), and particle-shaped masks, respectively. The fifth column displays the 3D projections of the enhanced and IsoNet missing-wedge-corrected particle. The sixth and seventh columns present the final 3D map and the flexibly fitted model, respectively. (B) Two cross-sectional views (12 nm thickness) of the plasmid density map along its plectoneme axis are shown in the left-middle panel. The intensity profile along the line crossing the two high-density DNA spots is displayed in the right panel. (C) Resolution assessment of the final 3D map using Fourier shell correlation (FSC). Two criteria are shown: FSC between two half-maps reconstructed from even and odd frames (evaluated at 0.143) and FSC between the final 3D map and the fitted model (evaluated at 0.5). (D) Zoomed-in views of the final 3D density map from panel A, displayed at two contour levels. (E) Superimposition of the high-contour level map from panel D onto its fitted model.

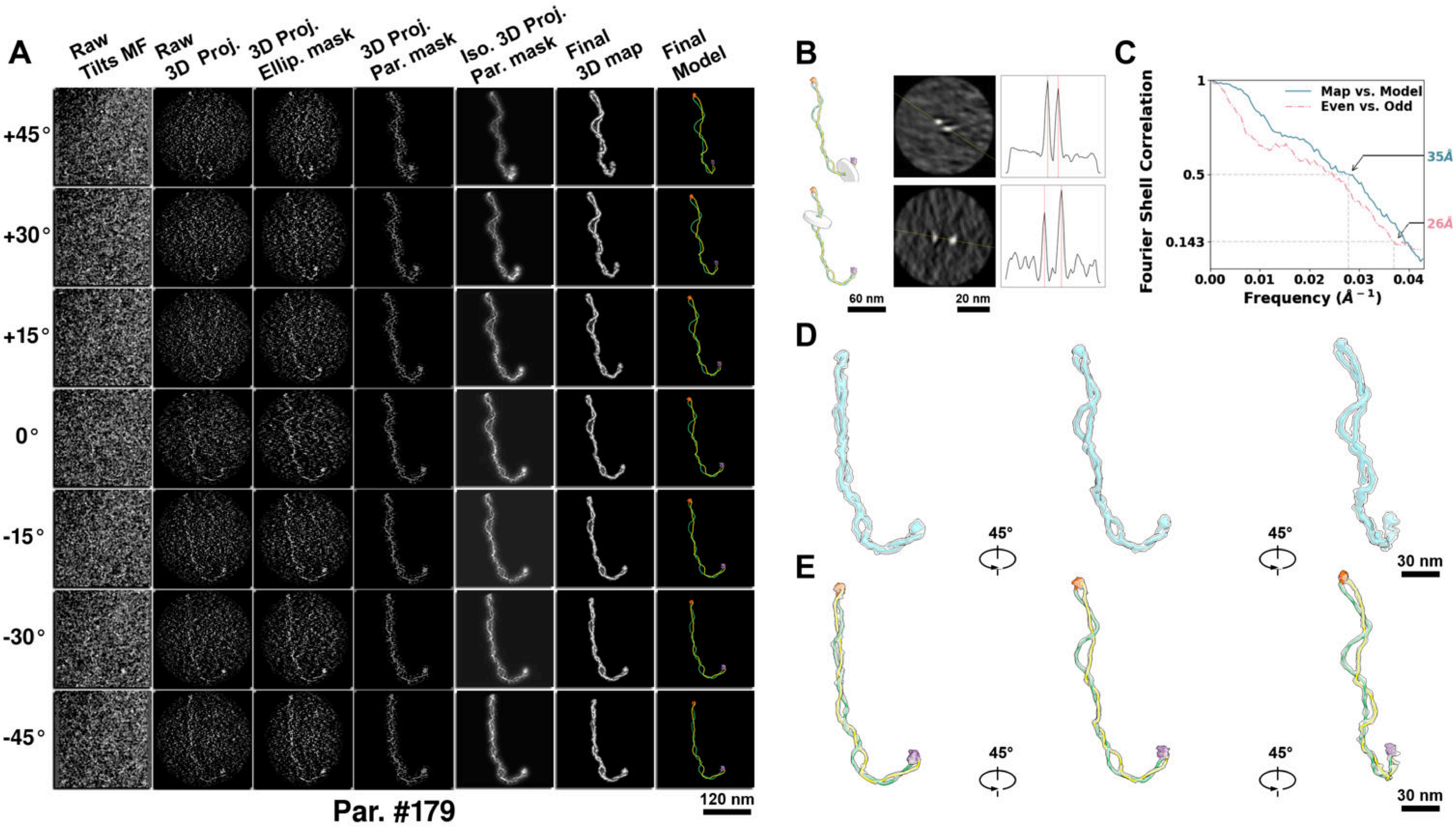

**Supplementary Particle Figure 179. Cryo-ET 3D reconstruction of an individual sTEC-Cas particle.**
(A) 3D reconstruction of the plasmid particle (index no. 179). The first column shows seven representative tilt images from +45° to -45° in step of 15°. The second, third, and fourth columns show 3D projections of the particle with spherical, ellipsoidal (thinner along the z-dimension), and particle-shaped masks, respectively. The fifth column displays the 3D projections of the enhanced and IsoNet missing-wedge-corrected particle. The sixth and seventh columns present the final 3D map and the flexibly fitted model, respectively. (B) Two cross-sectional views (12 nm thickness) of the plasmid density map along its plectoneme axis are shown in the left-middle panel. The intensity profile along the line crossing the two high-density DNA spots is displayed in the right panel. (C) Resolution assessment of the final 3D map using Fourier shell correlation (FSC). Two criteria are shown: FSC between two half-maps reconstructed from even and odd frames (evaluated at 0.143) and FSC between the final 3D map and the fitted model (evaluated at 0.5). (D) Zoomed-in views of the final 3D density map from panel A, displayed at two contour levels. (E) Superimposition of the high-contour level map from panel D onto its fitted model.

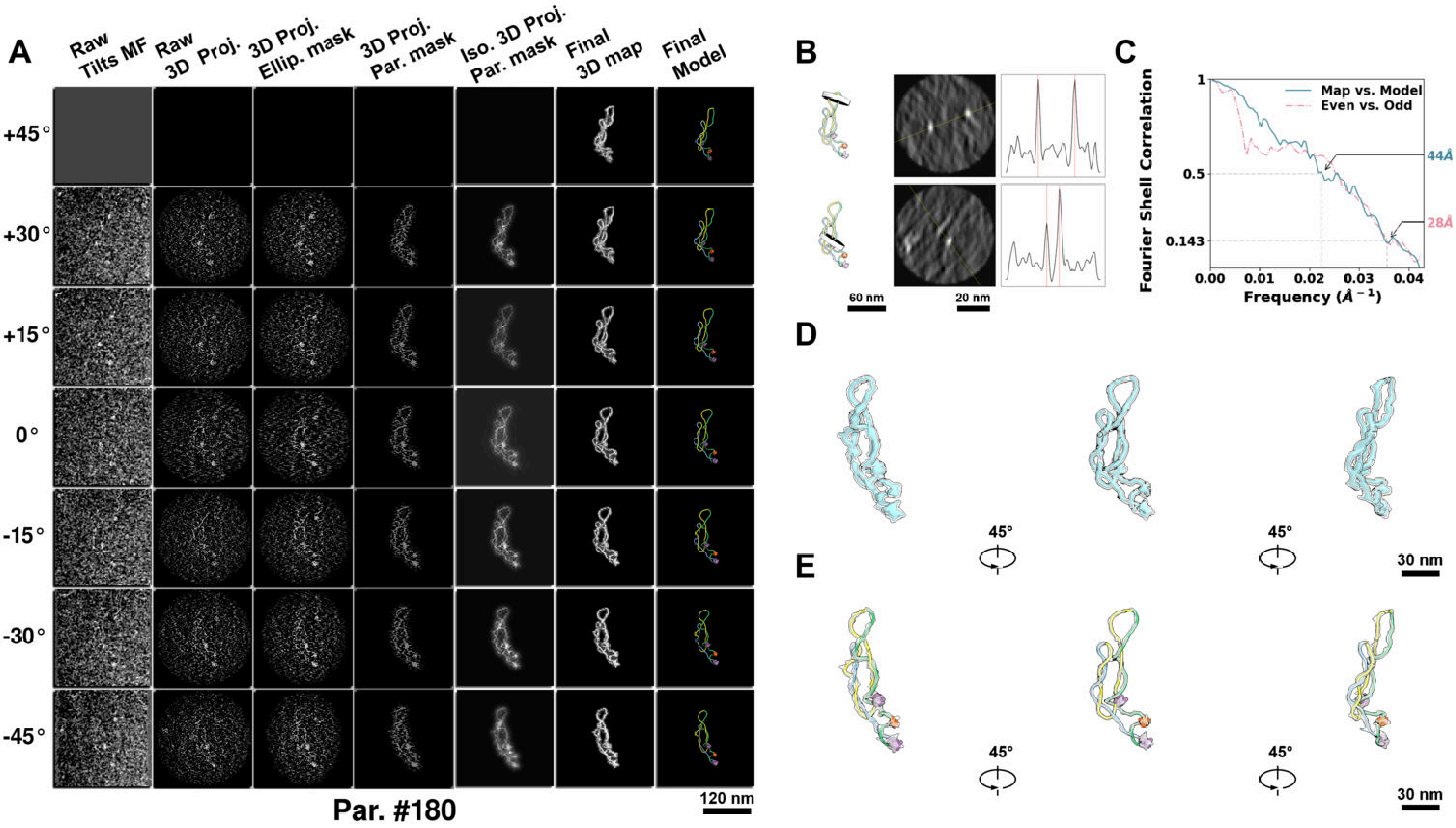

**Supplementary Particle Figure 180. Cryo-ET 3D reconstruction of an individual sTEC-Cas particle.**
(A) 3D reconstruction of the plasmid particle (index no. 180). The first column shows seven representative tilt images from +45° to -45° in step of 15°. The second, third, and fourth columns show 3D projections of the particle with spherical, ellipsoidal (thinner along the z-dimension), and particle-shaped masks, respectively. The fifth column displays the 3D projections of the enhanced and IsoNet missing-wedge-corrected particle. The sixth and seventh columns present the final 3D map and the flexibly fitted model, respectively. (B) Two cross-sectional views (12 nm thickness) of the plasmid density map along its plectoneme axis are shown in the left-middle panel. The intensity profile along the line crossing the two high-density DNA spots is displayed in the right panel. (C) Resolution assessment of the final 3D map using Fourier shell correlation (FSC). Two criteria are shown: FSC between two half-maps reconstructed from even and odd frames (evaluated at 0.143) and FSC between the final 3D map and the fitted model (evaluated at 0.5). (D) Zoomed-in views of the final 3D density map from panel A, displayed at two contour levels. (E) Superimposition of the high-contour level map from panel D onto its fitted model.

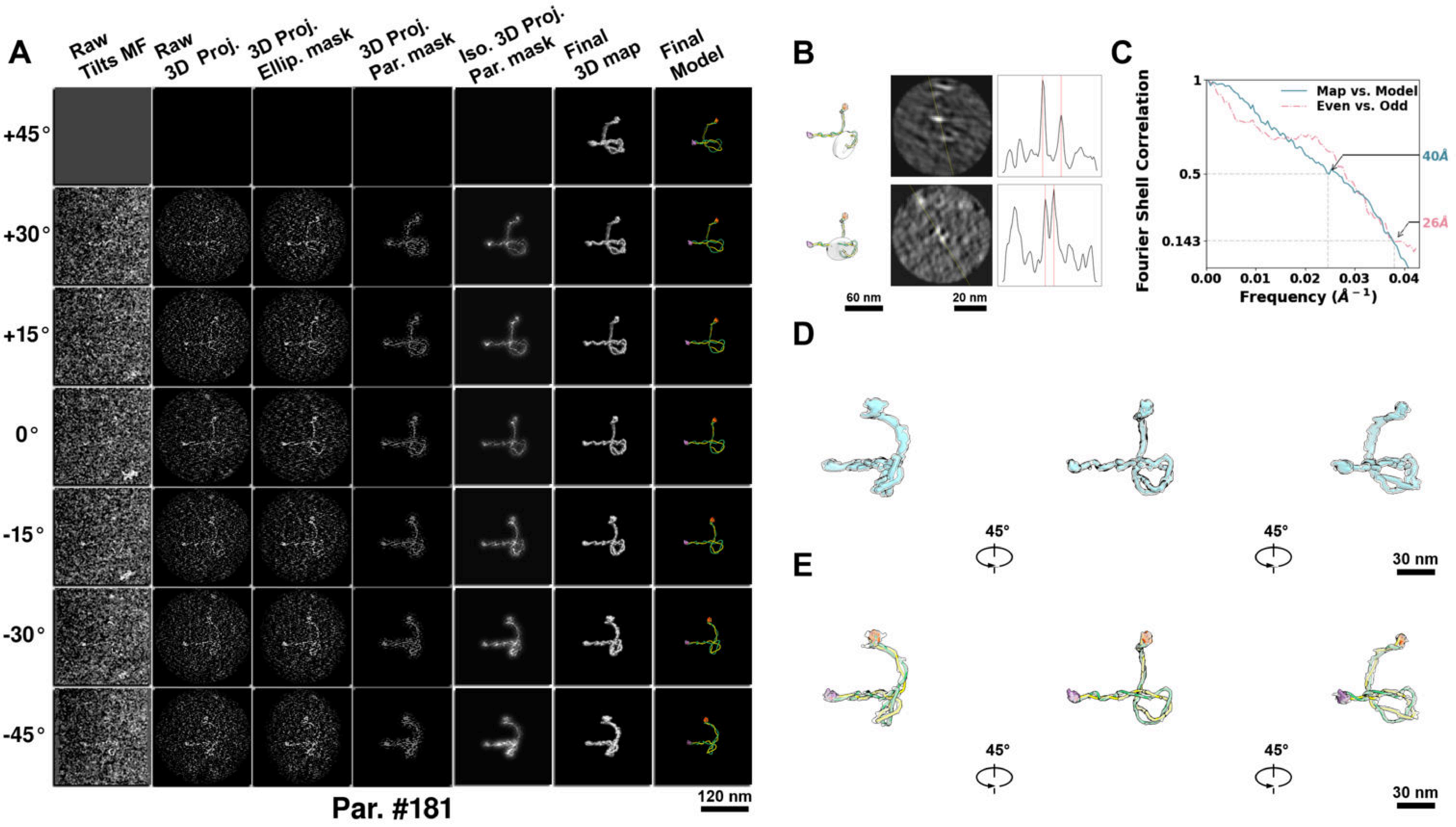

**Supplementary Particle Figure 181. Cryo-ET 3D reconstruction of an individual sTEC-Cas particle.**
(A) 3D reconstruction of the plasmid particle (index no. 181). The first column shows seven representative tilt images from +45° to -45° in step of 15°. The second, third, and fourth columns show 3D projections of the particle with spherical, ellipsoidal (thinner along the z-dimension), and particle-shaped masks, respectively. The fifth column displays the 3D projections of the enhanced and IsoNet missing-wedge-corrected particle. The sixth and seventh columns present the final 3D map and the flexibly fitted model, respectively. (B) Two cross-sectional views (12 nm thickness) of the plasmid density map along its plectoneme axis are shown in the left-middle panel. The intensity profile along the line crossing the two high-density DNA spots is displayed in the right panel. (C) Resolution assessment of the final 3D map using Fourier shell correlation (FSC). Two criteria are shown: FSC between two half-maps reconstructed from even and odd frames (evaluated at 0.143) and FSC between the final 3D map and the fitted model (evaluated at 0.5). (D) Zoomed-in views of the final 3D density map from panel A, displayed at two contour levels. (E) Superimposition of the high-contour level map from panel D onto its fitted model.

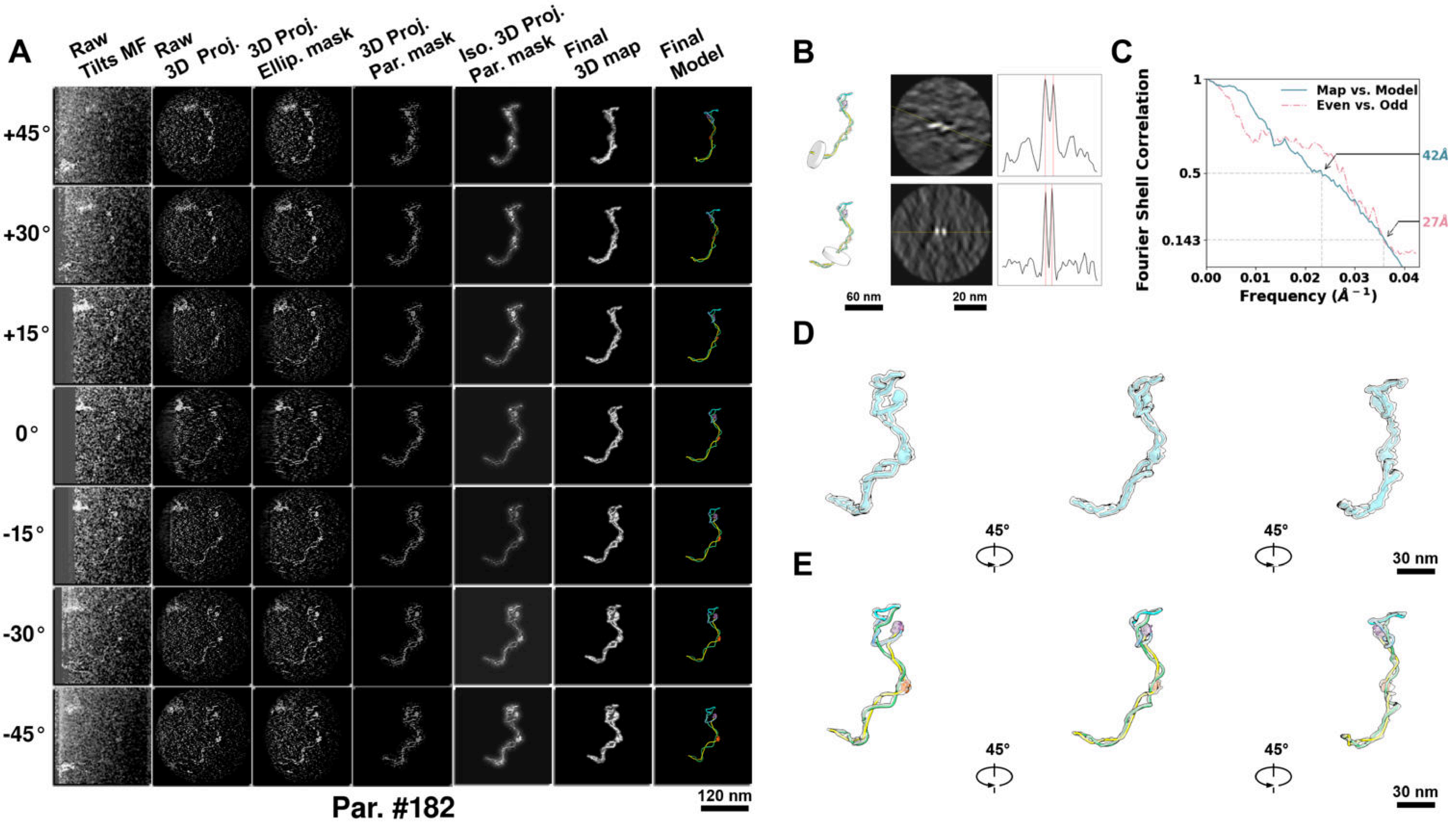

**Supplementary Particle Figure 182. Cryo-ET 3D reconstruction of an individual sTEC-Cas particle.**
(A) 3D reconstruction of the plasmid particle (index no. 182). The first column shows seven representative tilt images from +45° to -45° in step of 15°. The second, third, and fourth columns show 3D projections of the particle with spherical, ellipsoidal (thinner along the z-dimension), and particle-shaped masks, respectively. The fifth column displays the 3D projections of the enhanced and IsoNet missing-wedge-corrected particle. The sixth and seventh columns present the final 3D map and the flexibly fitted model, respectively. (B) Two cross-sectional views (12 nm thickness) of the plasmid density map along its plectoneme axis are shown in the left-middle panel. The intensity profile along the line crossing the two high-density DNA spots is displayed in the right panel. (C) Resolution assessment of the final 3D map using Fourier shell correlation (FSC). Two criteria are shown: FSC between two half-maps reconstructed from even and odd frames (evaluated at 0.143) and FSC between the final 3D map and the fitted model (evaluated at 0.5). (D) Zoomed-in views of the final 3D density map from panel A, displayed at two contour levels. (E) Superimposition of the high-contour level map from panel D onto its fitted model.

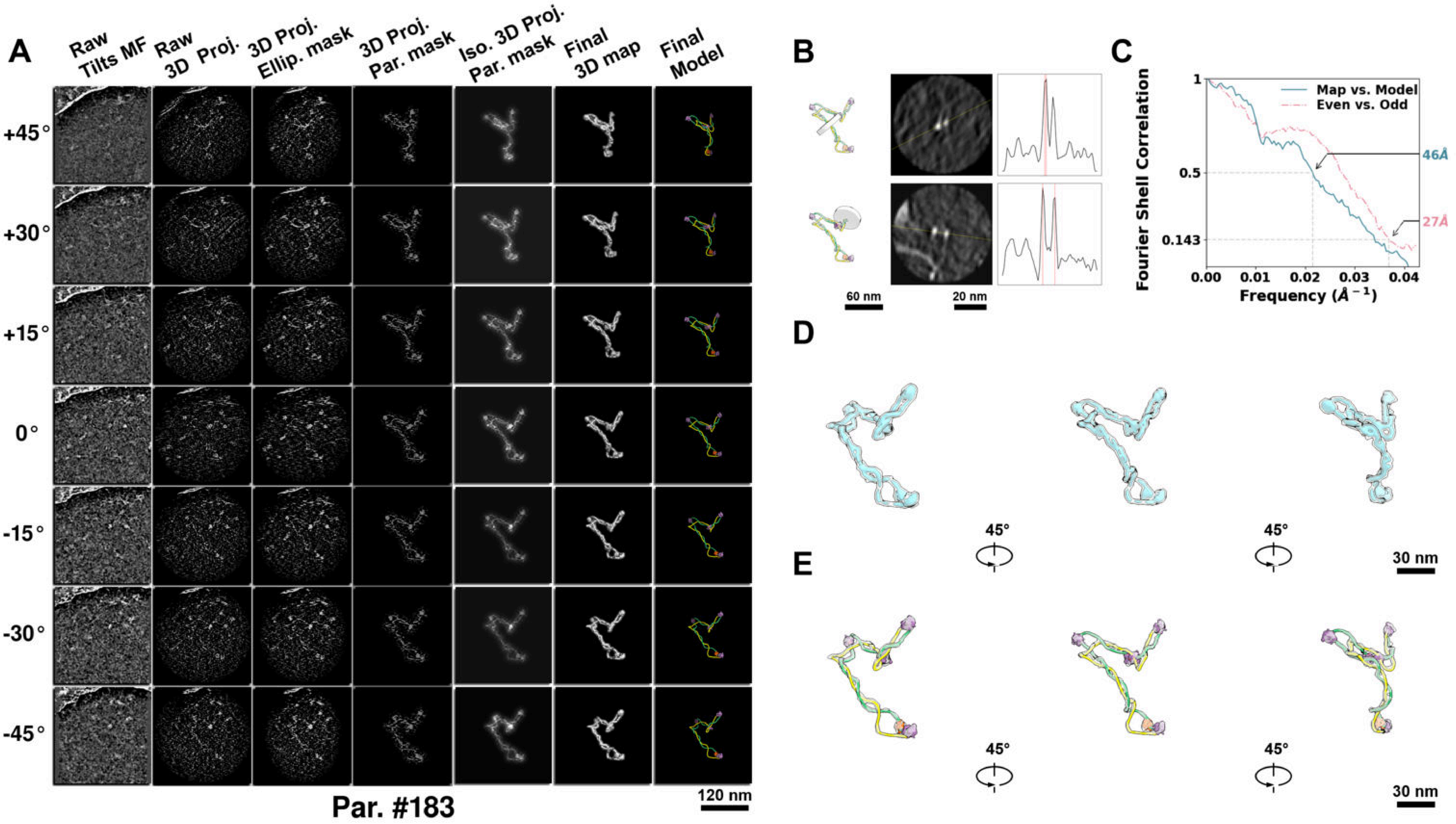

**Supplementary Particle Figure 183. Cryo-ET 3D reconstruction of an individual TEC-Cas particle.**
(A) 3D reconstruction of the plasmid particle (index no. 183). The first column shows seven representative tilt images from +45° to -45° in step of 15°. The second, third, and fourth columns show 3D projections of the particle with spherical, ellipsoidal (thinner along the z-dimension), and particle-shaped masks, respectively. The fifth column displays the 3D projections of the enhanced and IsoNet missing-wedge-corrected particle. The sixth and seventh columns present the final 3D map and the flexibly fitted model, respectively. (B) Two cross-sectional views (12 nm thickness) of the plasmid density map along its plectoneme axis are shown in the left-middle panel. The intensity profile along the line crossing the two high-density DNA spots is displayed in the right panel. (C) Resolution assessment of the final 3D map using Fourier shell correlation (FSC). Two criteria are shown: FSC between two half-maps reconstructed from even and odd frames (evaluated at 0.143) and FSC between the final 3D map and the fitted model (evaluated at 0.5). (D) Zoomed-in views of the final 3D density map from panel A, displayed at two contour levels. (E) Superimposition of the high-contour level map from panel D onto its fitted model.

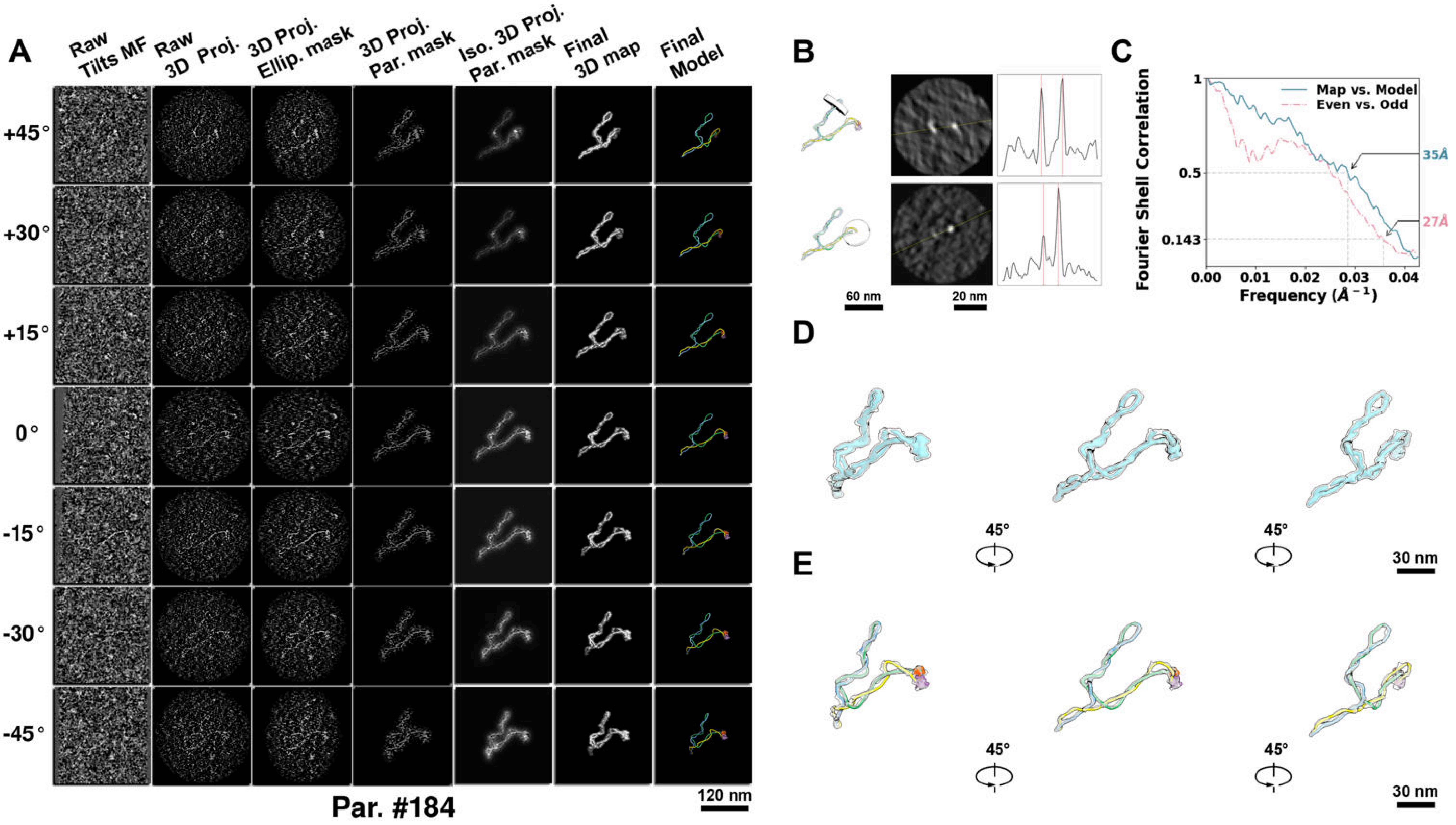

**Supplementary Particle Figure 184. Cryo-ET 3D reconstruction of an individual TEC-Cas particle.**
(A) 3D reconstruction of the plasmid particle (index no. 184). The first column shows seven representative tilt images from +45° to -45° in step of 15°. The second, third, and fourth columns show 3D projections of the particle with spherical, ellipsoidal (thinner along the z-dimension), and particle-shaped masks, respectively. The fifth column displays the 3D projections of the enhanced and IsoNet missing-wedge-corrected particle. The sixth and seventh columns present the final 3D map and the flexibly fitted model, respectively. (B) Two cross-sectional views (12 nm thickness) of the plasmid density map along its plectoneme axis are shown in the left-middle panel. The intensity profile along the line crossing the two high-density DNA spots is displayed in the right panel. (C) Resolution assessment of the final 3D map using Fourier shell correlation (FSC). Two criteria are shown: FSC between two half-maps reconstructed from even and odd frames (evaluated at 0.143) and FSC between the final 3D map and the fitted model (evaluated at 0.5). (D) Zoomed-in views of the final 3D density map from panel A, displayed at two contour levels. (E) Superimposition of the high-contour level map from panel D onto its fitted model.

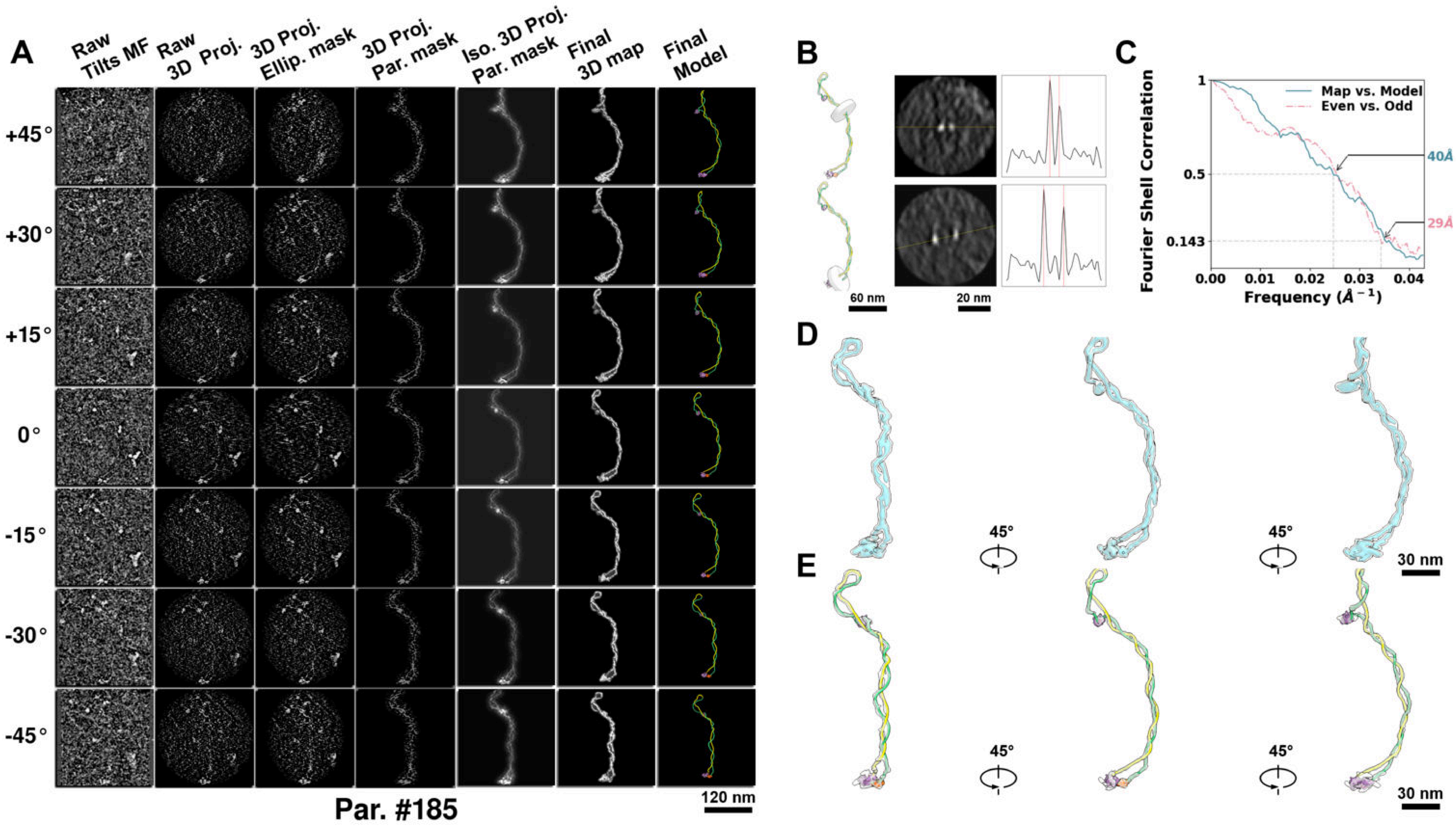

**Supplementary Particle Figure 185. Cryo-ET 3D reconstruction of an individual TEC-Cas particle.**
(A) 3D reconstruction of the plasmid particle (index no. 185). The first column shows seven representative tilt images from +45° to -45° in step of 15°. The second, third, and fourth columns show 3D projections of the particle with spherical, ellipsoidal (thinner along the z-dimension), and particle-shaped masks, respectively. The fifth column displays the 3D projections of the enhanced and IsoNet missing-wedge-corrected particle. The sixth and seventh columns present the final 3D map and the flexibly fitted model, respectively. (B) Two cross-sectional views (12 nm thickness) of the plasmid density map along its plectoneme axis are shown in the left-middle panel. The intensity profile along the line crossing the two high-density DNA spots is displayed in the right panel. (C) Resolution assessment of the final 3D map using Fourier shell correlation (FSC). Two criteria are shown: FSC between two half-maps reconstructed from even and odd frames (evaluated at 0.143) and FSC between the final 3D map and the fitted model (evaluated at 0.5). (D) Zoomed-in views of the final 3D density map from panel A, displayed at two contour levels. (E) Superimposition of the high-contour level map from panel D onto its fitted model.

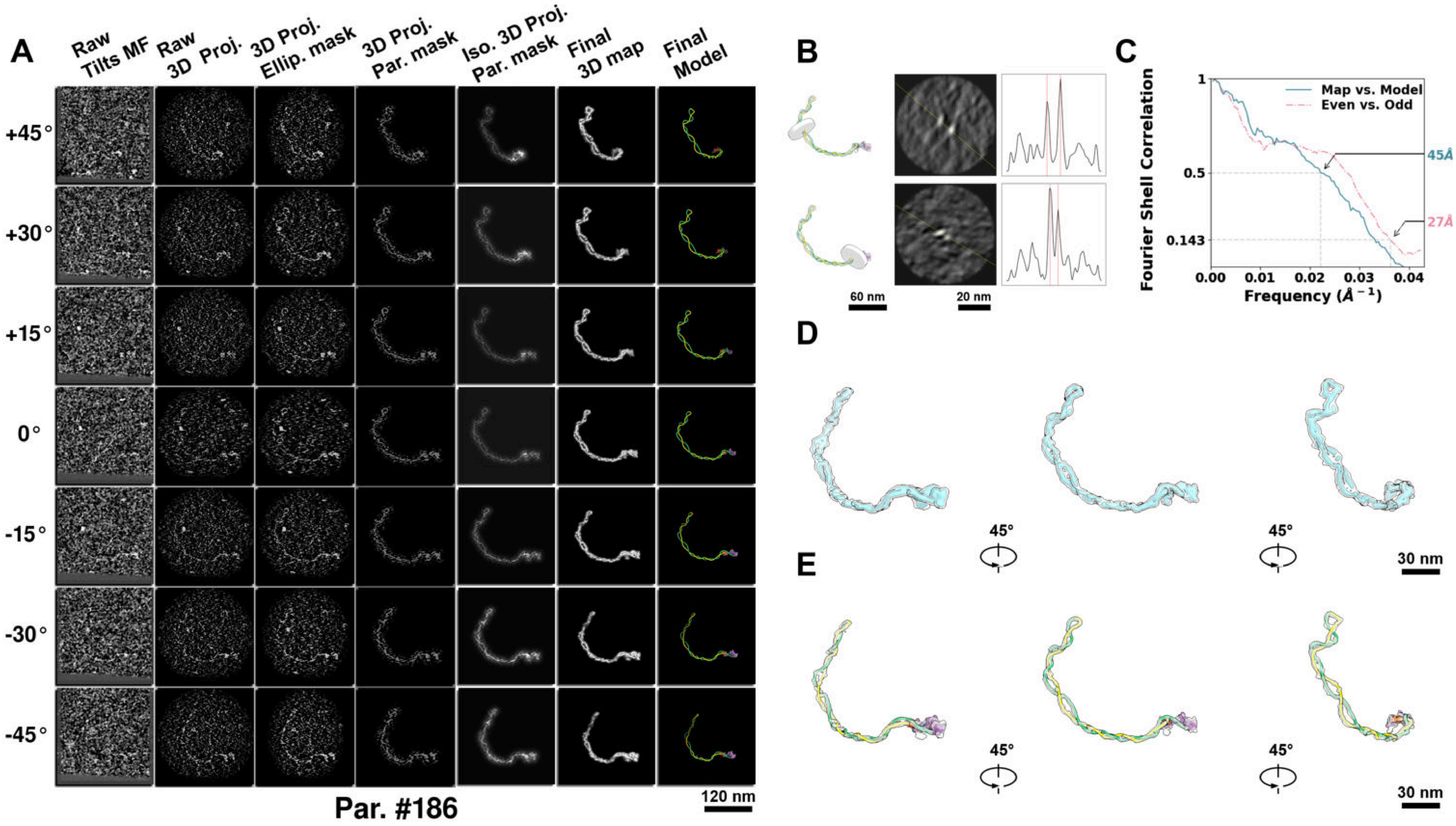

**Supplementary Particle Figure 186. Cryo-ET 3D reconstruction of an individual TEC-Cas particle.**
(A) 3D reconstruction of the plasmid particle (index no. 186). The first column shows seven representative tilt images from +45° to -45° in step of 15°. The second, third, and fourth columns show 3D projections of the particle with spherical, ellipsoidal (thinner along the z-dimension), and particle-shaped masks, respectively. The fifth column displays the 3D projections of the enhanced and IsoNet missing-wedge-corrected particle. The sixth and seventh columns present the final 3D map and the flexibly fitted model, respectively. (B) Two cross-sectional views (12 nm thickness) of the plasmid density map along its plectoneme axis are shown in the left-middle panel. The intensity profile along the line crossing the two high-density DNA spots is displayed in the right panel. (C) Resolution assessment of the final 3D map using Fourier shell correlation (FSC). Two criteria are shown: FSC between two half-maps reconstructed from even and odd frames (evaluated at 0.143) and FSC between the final 3D map and the fitted model (evaluated at 0.5). (D) Zoomed-in views of the final 3D density map from panel A, displayed at two contour levels. (E) Superimposition of the high-contour level map from panel D onto its fitted model.

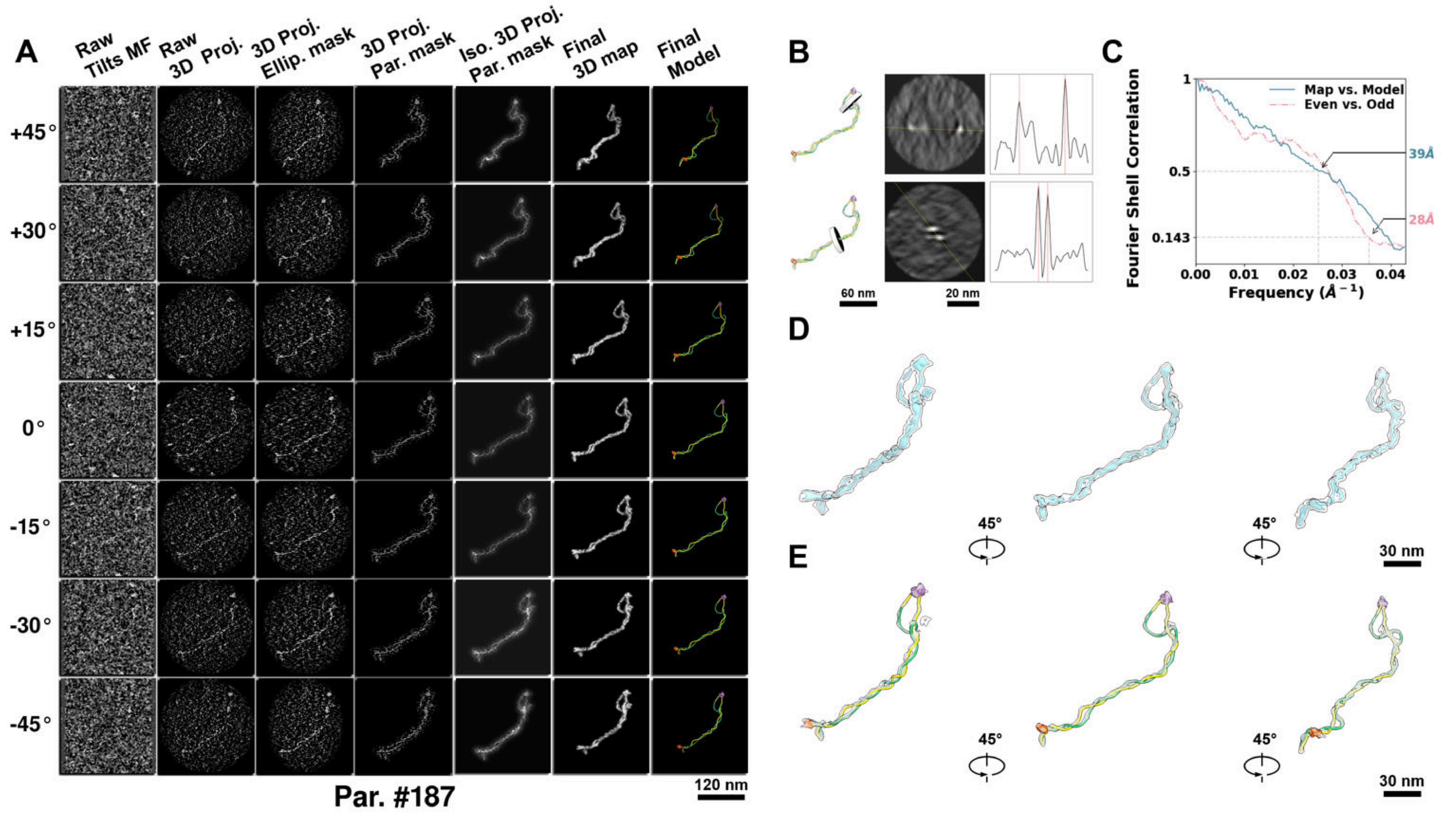

**Supplementary Particle Figure 187. Cryo-ET 3D reconstruction of an individual TEC-Cas particle.**
(A) 3D reconstruction of the plasmid particle (index no. 187). The first column shows seven representative tilt images from +45° to -45° in step of 15°. The second, third, and fourth columns show 3D projections of the particle with spherical, ellipsoidal (thinner along the z-dimension), and particle-shaped masks, respectively. The fifth column displays the 3D projections of the enhanced and IsoNet missing-wedge-corrected particle. The sixth and seventh columns present the final 3D map and the flexibly fitted model, respectively. (B) Two cross-sectional views (12 nm thickness) of the plasmid density map along its plectoneme axis are shown in the left-middle panel. The intensity profile along the line crossing the two high-density DNA spots is displayed in the right panel. (C) Resolution assessment of the final 3D map using Fourier shell correlation (FSC). Two criteria are shown: FSC between two half-maps reconstructed from even and odd frames (evaluated at 0.143) and FSC between the final 3D map and the fitted model (evaluated at 0.5). (D) Zoomed-in views of the final 3D density map from panel A, displayed at two contour levels. (E) Superimposition of the high-contour level map from panel D onto its fitted model.

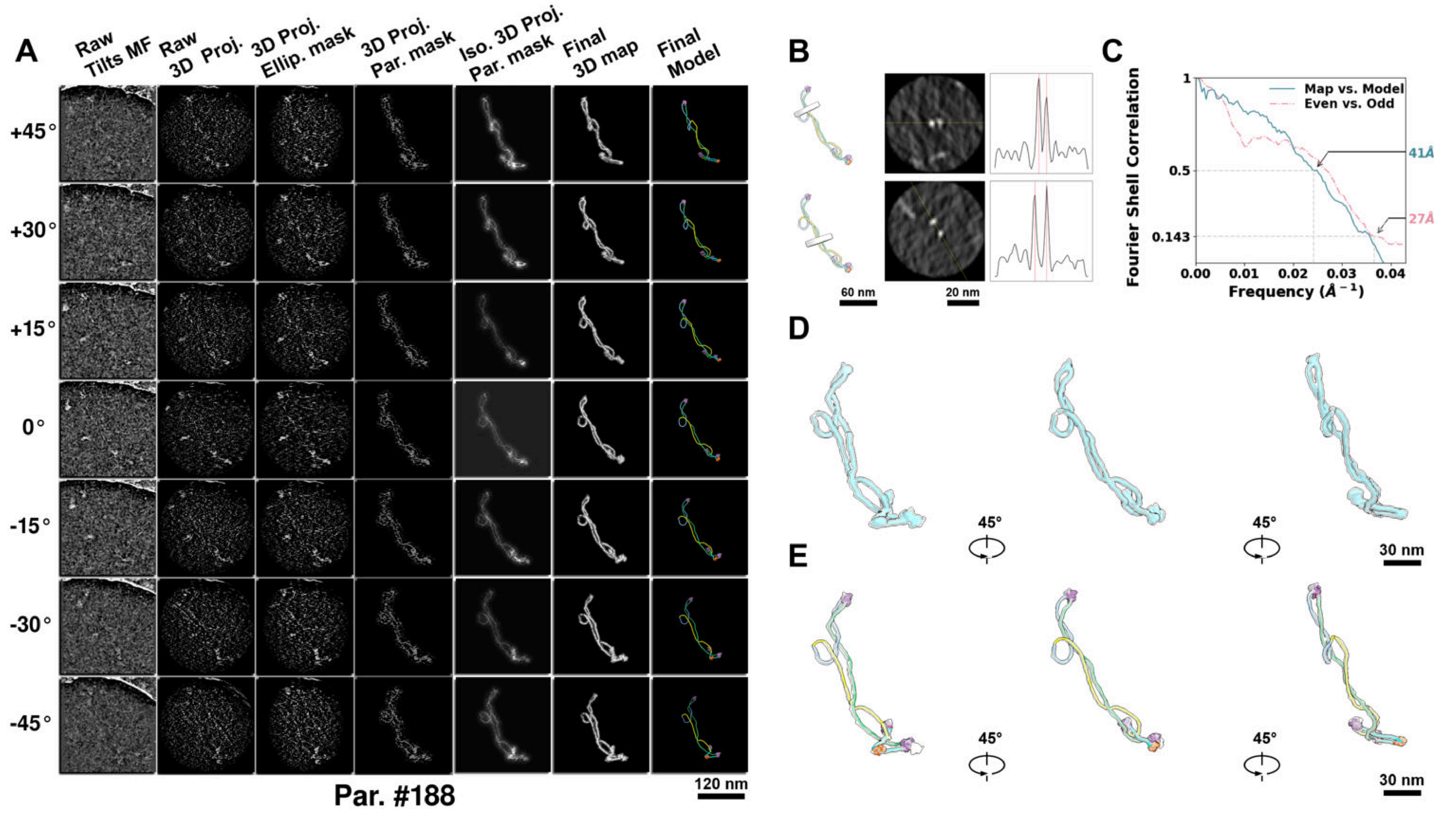

**Supplementary Particle Figure 188. Cryo-ET 3D reconstruction of an individual TEC-Cas particle.**
(A) 3D reconstruction of the plasmid particle (index no. 188). The first column shows seven representative tilt images from +45° to -45° in step of 15°. The second, third, and fourth columns show 3D projections of the particle with spherical, ellipsoidal (thinner along the z-dimension), and particle-shaped masks, respectively. The fifth column displays the 3D projections of the enhanced and IsoNet missing-wedge-corrected particle. The sixth and seventh columns present the final 3D map and the flexibly fitted model, respectively. (B) Two cross-sectional views (12 nm thickness) of the plasmid density map along its plectoneme axis are shown in the left-middle panel. The intensity profile along the line crossing the two high-density DNA spots is displayed in the right panel. (C) Resolution assessment of the final 3D map using Fourier shell correlation (FSC). Two criteria are shown: FSC between two half-maps reconstructed from even and odd frames (evaluated at 0.143) and FSC between the final 3D map and the fitted model (evaluated at 0.5). (D) Zoomed-in views of the final 3D density map from panel A, displayed at two contour levels. (E) Superimposition of the high-contour level map from panel D onto its fitted model.

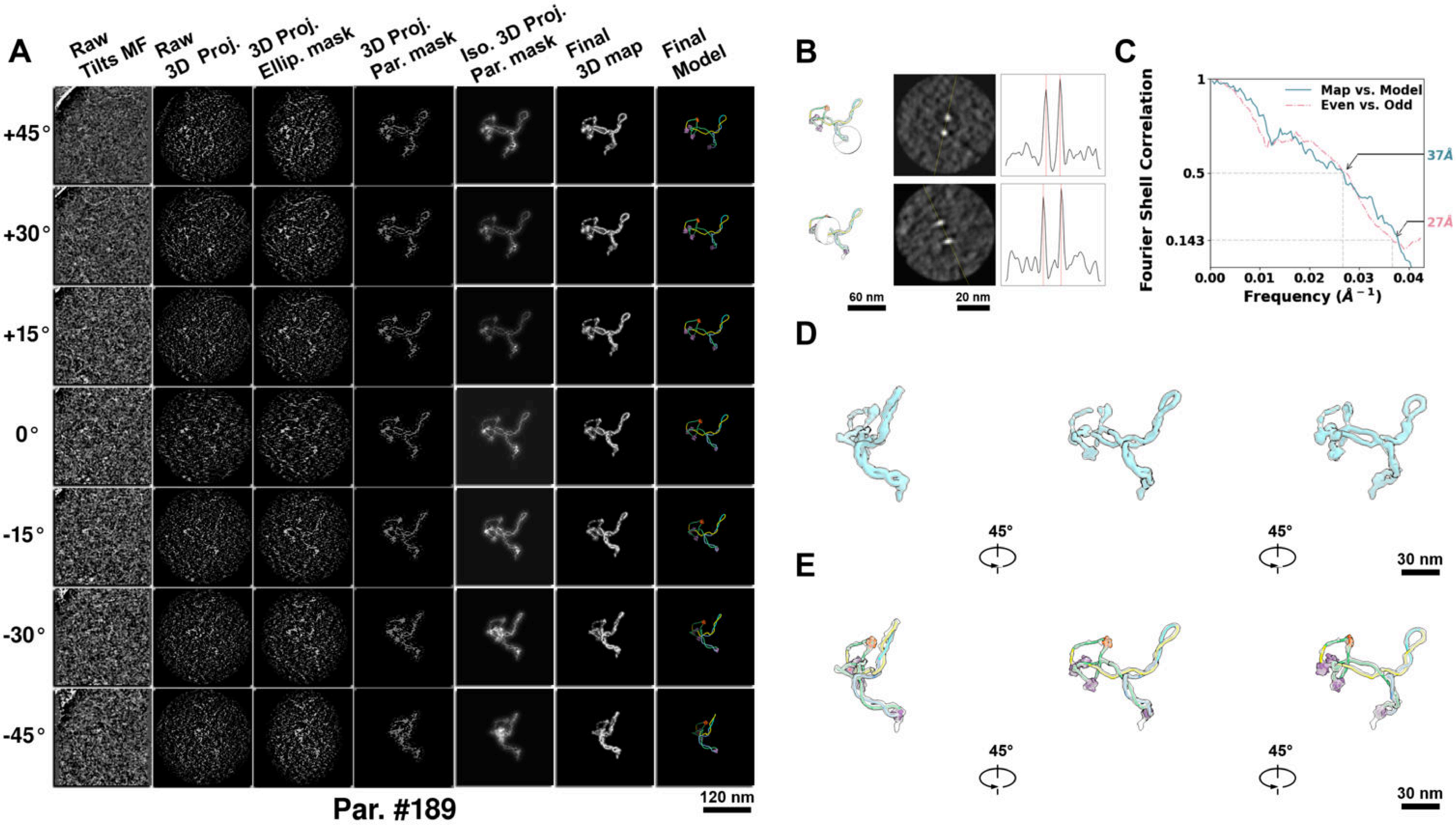

**Supplementary Particle Figure 189. Cryo-ET 3D reconstruction of an individual TEC-Cas particle.**
(A) 3D reconstruction of the plasmid particle (index no. 189). The first column shows seven representative tilt images from +45° to -45° in step of 15°. The second, third, and fourth columns show 3D projections of the particle with spherical, ellipsoidal (thinner along the z-dimension), and particle-shaped masks, respectively. The fifth column displays the 3D projections of the enhanced and IsoNet missing-wedge-corrected particle. The sixth and seventh columns present the final 3D map and the flexibly fitted model, respectively. (B) Two cross-sectional views (12 nm thickness) of the plasmid density map along its plectoneme axis are shown in the left-middle panel. The intensity profile along the line crossing the two high-density DNA spots is displayed in the right panel. (C) Resolution assessment of the final 3D map using Fourier shell correlation (FSC). Two criteria are shown: FSC between two half-maps reconstructed from even and odd frames (evaluated at 0.143) and FSC between the final 3D map and the fitted model (evaluated at 0.5). (D) Zoomed-in views of the final 3D density map from panel A, displayed at two contour levels. (E) Superimposition of the high-contour level map from panel D onto its fitted model.

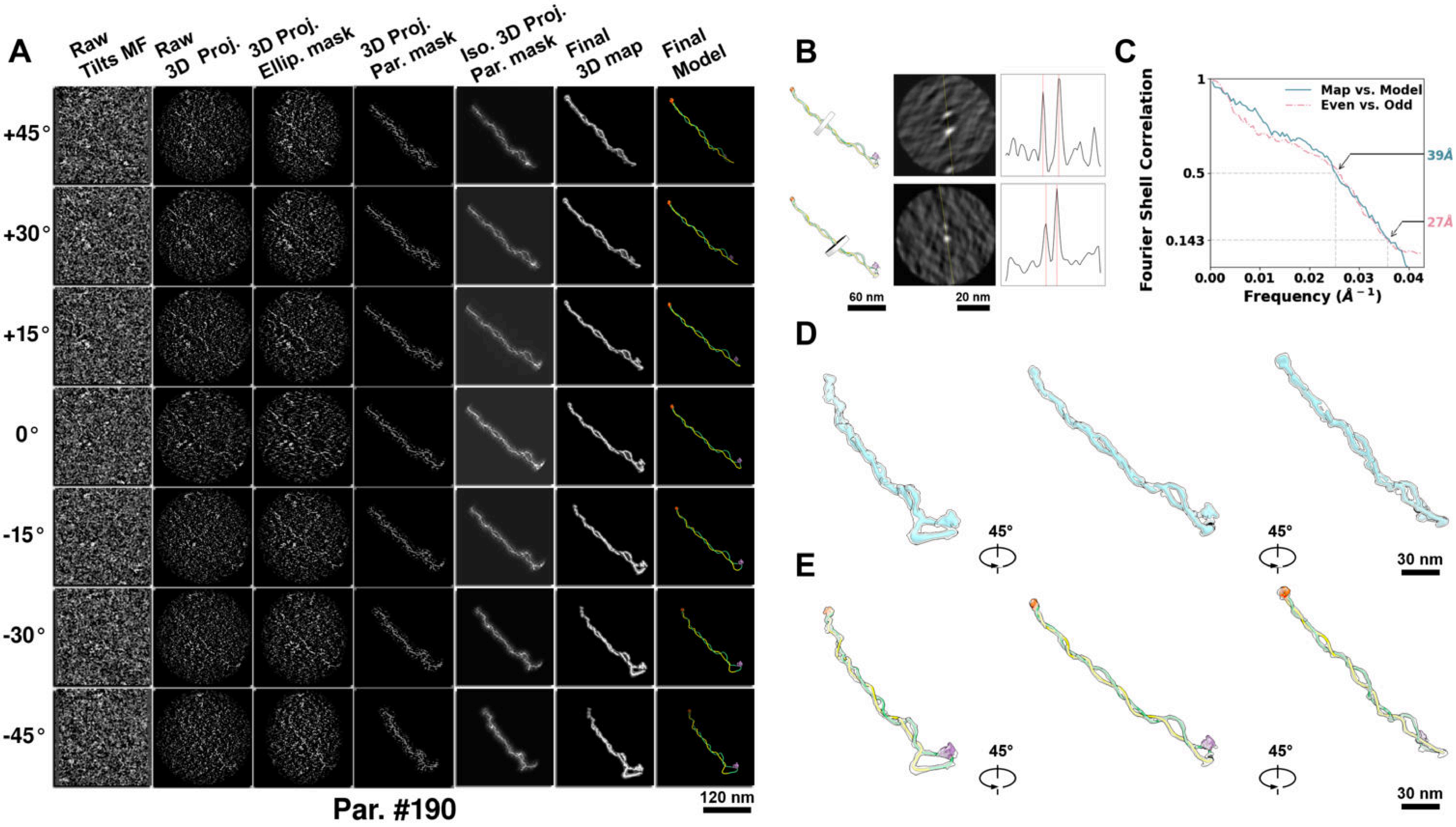

**Supplementary Particle Figure 190. Cryo-ET 3D reconstruction of an individual TEC-Cas particle.**
(A) 3D reconstruction of the plasmid particle (index no. 190). The first column shows seven representative tilt images from +45° to -45° in step of 15°. The second, third, and fourth columns show 3D projections of the particle with spherical, ellipsoidal (thinner along the z-dimension), and particle-shaped masks, respectively. The fifth column displays the 3D projections of the enhanced and IsoNet missing-wedge-corrected particle. The sixth and seventh columns present the final 3D map and the flexibly fitted model, respectively. (B) Two cross-sectional views (12 nm thickness) of the plasmid density map along its plectoneme axis are shown in the left-middle panel. The intensity profile along the line crossing the two high-density DNA spots is displayed in the right panel. (C) Resolution assessment of the final 3D map using Fourier shell correlation (FSC). Two criteria are shown: FSC between two half-maps reconstructed from even and odd frames (evaluated at 0.143) and FSC between the final 3D map and the fitted model (evaluated at 0.5). (D) Zoomed-in views of the final 3D density map from panel A, displayed at two contour levels. (E) Superimposition of the high-contour level map from panel D onto its fitted model.

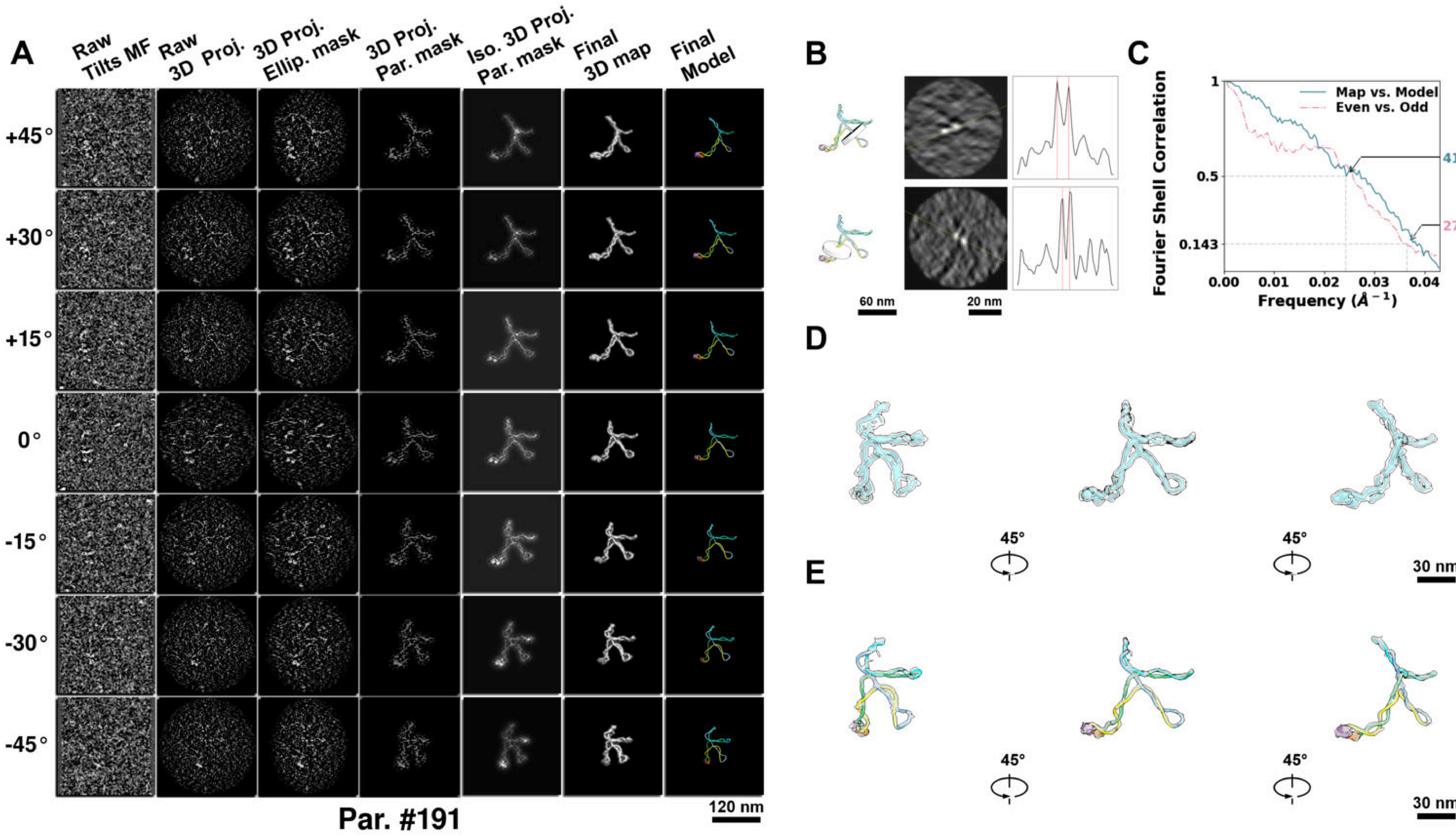

**Supplementary Particle Figure 191. Cryo-ET 3D reconstruction of an individual TEC-Cas particle.**
(A) 3D reconstruction of the plasmid particle (index no. 191). The first column shows seven representative tilt images from +45° to -45° in step of 15°. The second, third, and fourth columns show 3D projections of the particle with spherical, ellipsoidal (thinner along the z-dimension), and particle-shaped masks, respectively. The fifth column displays the 3D projections of the enhanced and IsoNet missing-wedge-corrected particle. The sixth and seventh columns present the final 3D map and the flexibly fitted model, respectively. (B) Two cross-sectional views (12 nm thickness) of the plasmid density map along its plectoneme axis are shown in the left-middle panel. The intensity profile along the line crossing the two high-density DNA spots is displayed in the right panel. (C) Resolution assessment of the final 3D map using Fourier shell correlation (FSC). Two criteria are shown: FSC between two half-maps reconstructed from even and odd frames (evaluated at 0.143) and FSC between the final 3D map and the fitted model (evaluated at 0.5). (D) Zoomed-in views of the final 3D density map from panel A, displayed at two contour levels. (E) Superimposition of the high-contour level map from panel D onto its fitted model.

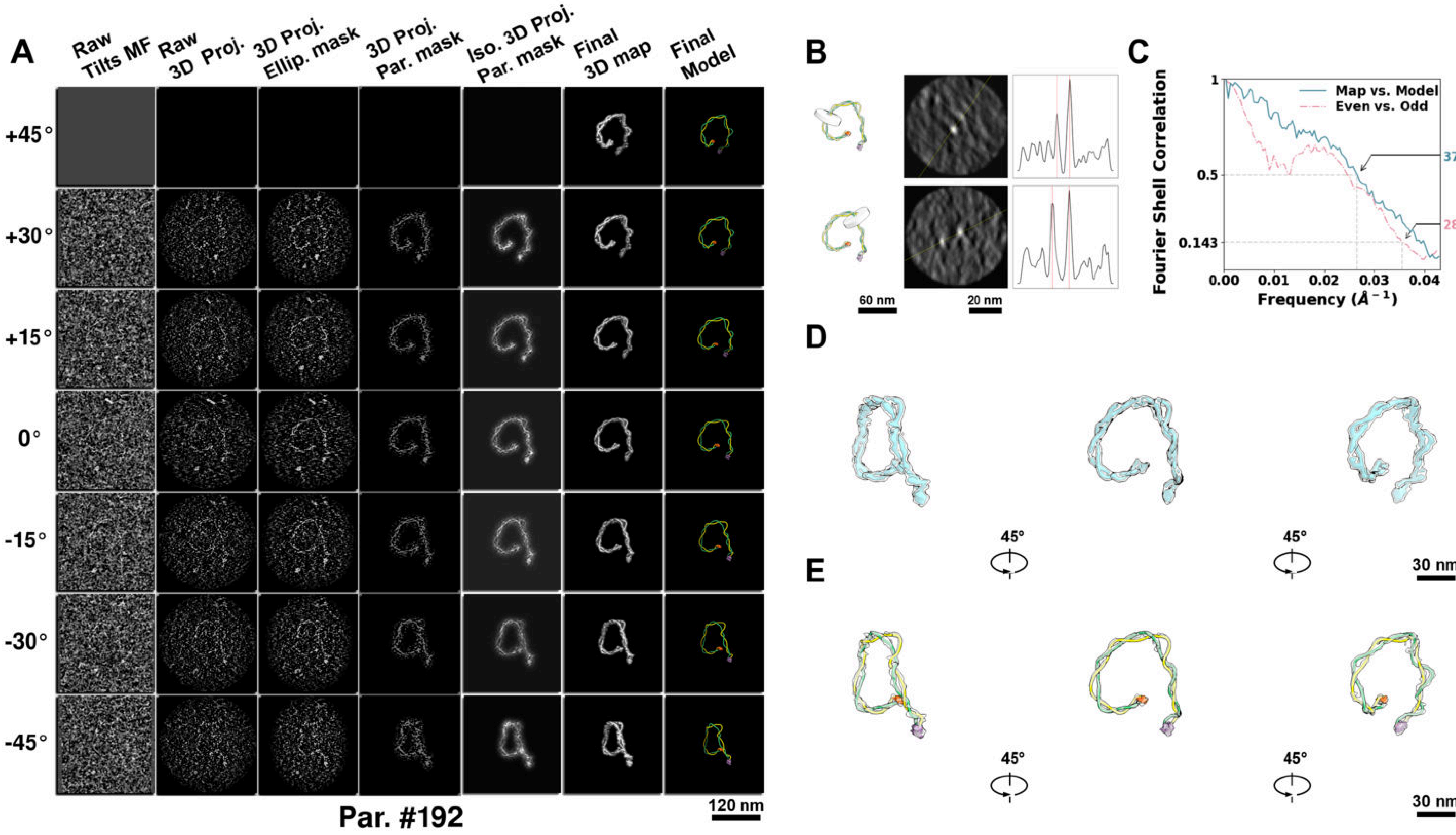

**Supplementary Particle Figure 192. Cryo-ET 3D reconstruction of an individual TEC-Cas particle.**
(A) 3D reconstruction of the plasmid particle (index no. 192). The first column shows seven representative tilt images from +45° to -45° in step of 15°. The second, third, and fourth columns show 3D projections of the particle with spherical, ellipsoidal (thinner along the z-dimension), and particle-shaped masks, respectively. The fifth column displays the 3D projections of the enhanced and IsoNet missing-wedge-corrected particle. The sixth and seventh columns present the final 3D map and the flexibly fitted model, respectively. (B) Two cross-sectional views (12 nm thickness) of the plasmid density map along its plectoneme axis are shown in the left-middle panel. The intensity profile along the line crossing the two high-density DNA spots is displayed in the right panel. (C) Resolution assessment of the final 3D map using Fourier shell correlation (FSC). Two criteria are shown: FSC between two half-maps reconstructed from even and odd frames (evaluated at 0.143) and FSC between the final 3D map and the fitted model (evaluated at 0.5). (D) Zoomed-in views of the final 3D density map from panel A, displayed at two contour levels. (E) Superimposition of the high-contour level map from panel D onto its fitted model.

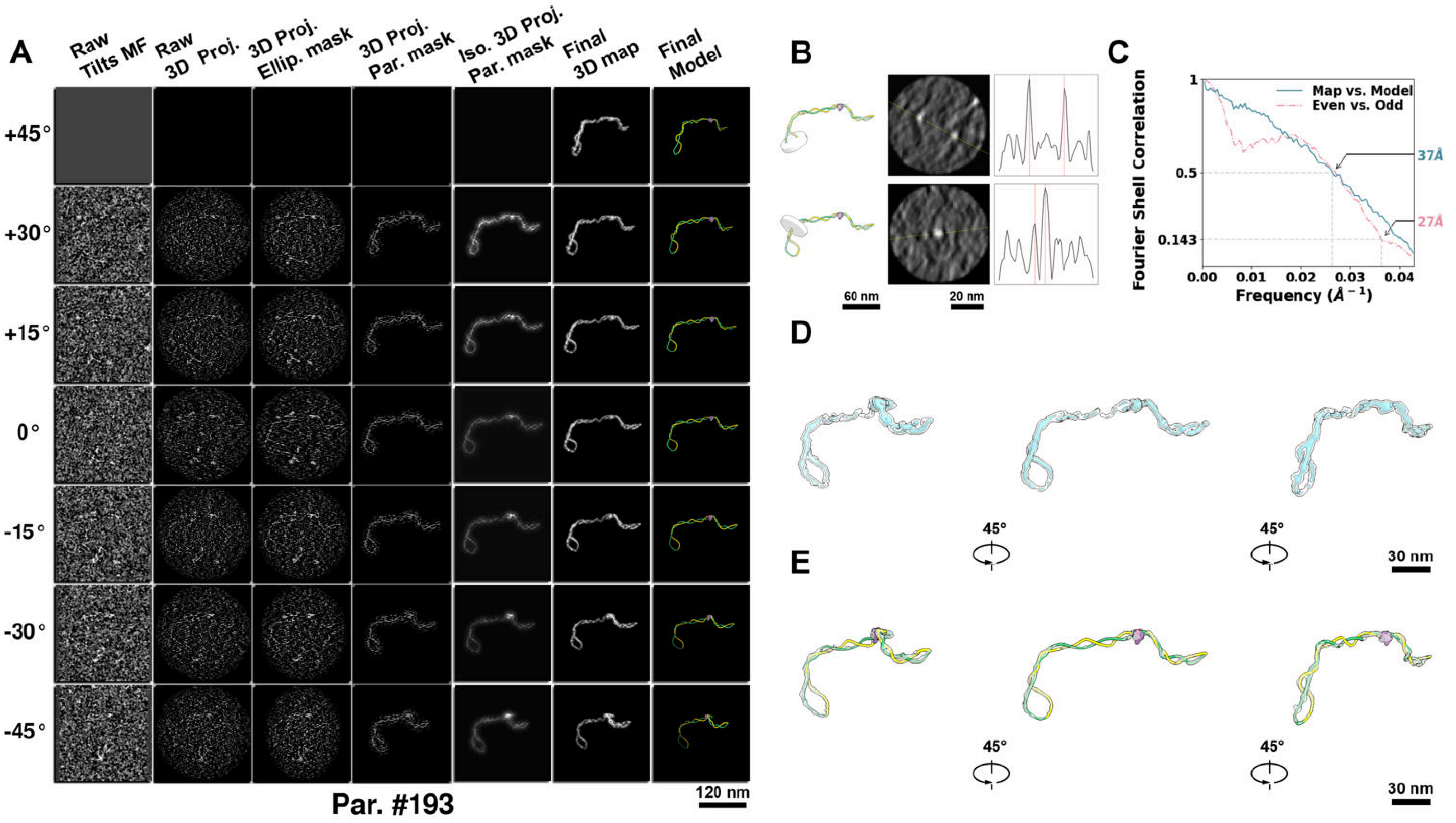

**Supplementary Particle Figure 193. Cryo-ET 3D reconstruction of an individual TEC-Cas particle.**
(A) 3D reconstruction of the plasmid particle (index no. 193). The first column shows seven representative tilt images from +45° to -45° in step of 15°. The second, third, and fourth columns show 3D projections of the particle with spherical, ellipsoidal (thinner along the z-dimension), and particle-shaped masks, respectively. The fifth column displays the 3D projections of the enhanced and IsoNet missing-wedge-corrected particle. The sixth and seventh columns present the final 3D map and the flexibly fitted model, respectively. (B) Two cross-sectional views (12 nm thickness) of the plasmid density map along its plectoneme axis are shown in the left-middle panel. The intensity profile along the line crossing the two high-density DNA spots is displayed in the right panel. (C) Resolution assessment of the final 3D map using Fourier shell correlation (FSC). Two criteria are shown: FSC between two half-maps reconstructed from even and odd frames (evaluated at 0.143) and FSC between the final 3D map and the fitted model (evaluated at 0.5). (D) Zoomed-in views of the final 3D density map from panel A, displayed at two contour levels. (E) Superimposition of the high-contour level map from panel D onto its fitted model.

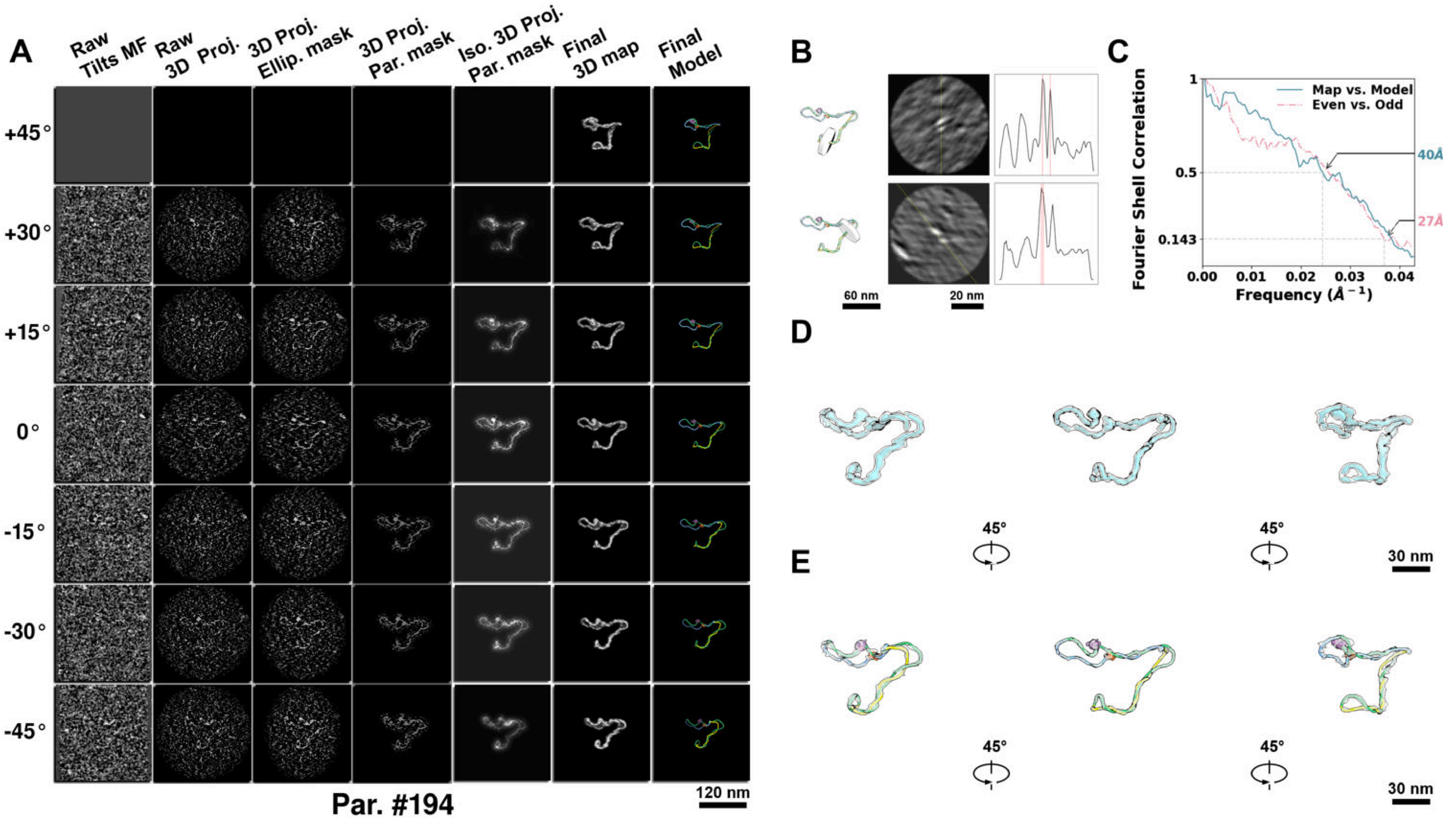

**Supplementary Particle Figure 194. Cryo-ET 3D reconstruction of an individual TEC-Cas particle.**
(A) 3D reconstruction of the plasmid particle (index no. 194). The first column shows seven representative tilt images from +45° to -45° in step of 15°. The second, third, and fourth columns show 3D projections of the particle with spherical, ellipsoidal (thinner along the z-dimension), and particle-shaped masks, respectively. The fifth column displays the 3D projections of the enhanced and IsoNet missing-wedge-corrected particle. The sixth and seventh columns present the final 3D map and the flexibly fitted model, respectively. (B) Two cross-sectional views (12 nm thickness) of the plasmid density map along its plectoneme axis are shown in the left-middle panel. The intensity profile along the line crossing the two high-density DNA spots is displayed in the right panel. (C) Resolution assessment of the final 3D map using Fourier shell correlation (FSC). Two criteria are shown: FSC between two half-maps reconstructed from even and odd frames (evaluated at 0.143) and FSC between the final 3D map and the fitted model (evaluated at 0.5). (D) Zoomed-in views of the final 3D density map from panel A, displayed at two contour levels. (E) Superimposition of the high-contour level map from panel D onto its fitted model.

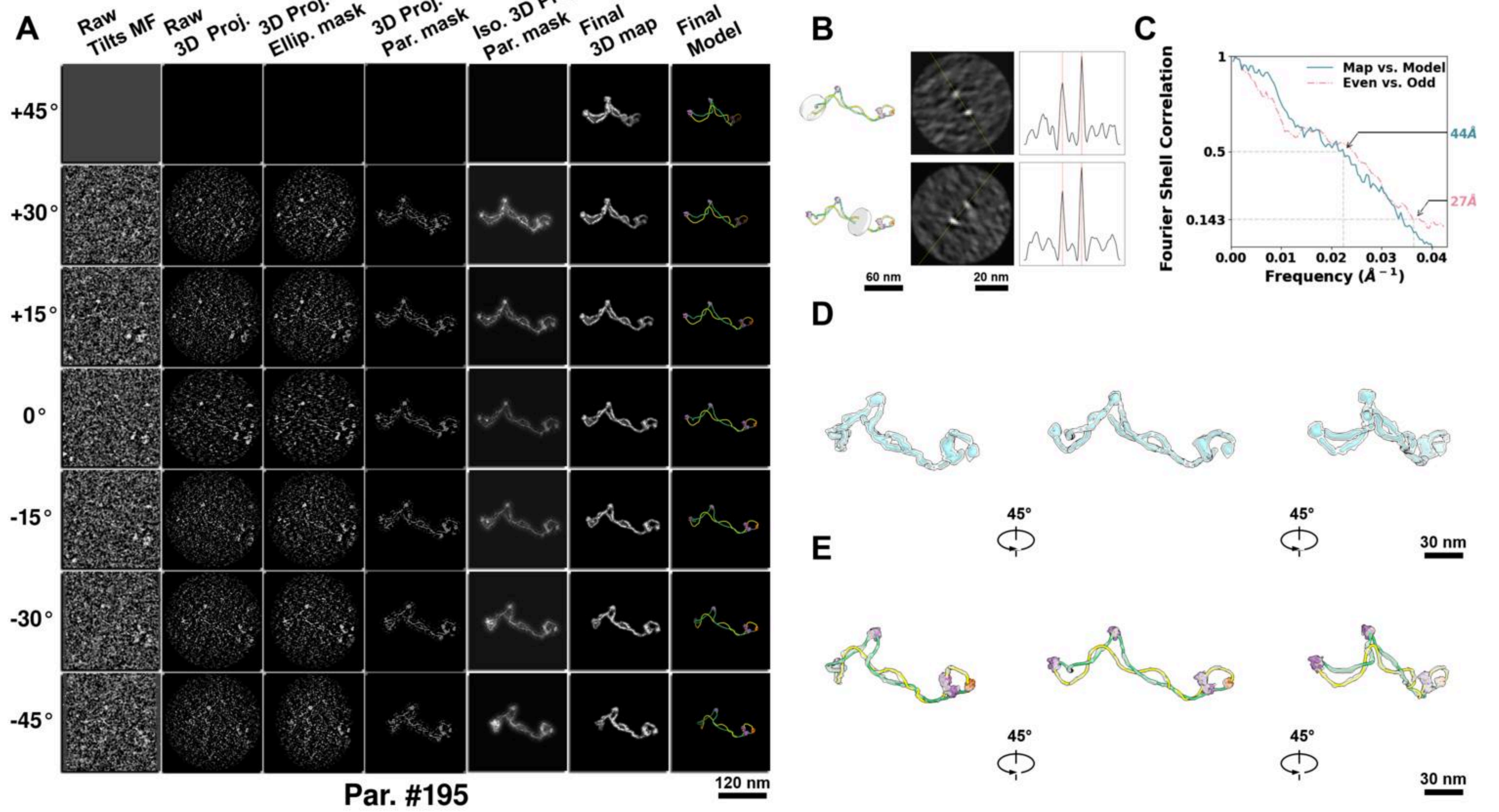

**Supplementary Particle Figure 195. Cryo-ET 3D reconstruction of an individual TEC-Cas particle.**
(A) 3D reconstruction of the plasmid particle (index no. 195). The first column shows seven representative tilt images from +45° to -45° in step of 15°. The second, third, and fourth columns show 3D projections of the particle with spherical, ellipsoidal (thinner along the z-dimension), and particle-shaped masks, respectively. The fifth column displays the 3D projections of the enhanced and IsoNet missing-wedge-corrected particle. The sixth and seventh columns present the final 3D map and the flexibly fitted model, respectively. (B) Two cross-sectional views (12 nm thickness) of the plasmid density map along its plectoneme axis are shown in the left-middle panel. The intensity profile along the line crossing the two high-density DNA spots is displayed in the right panel. (C) Resolution assessment of the final 3D map using Fourier shell correlation (FSC). Two criteria are shown: FSC between two half-maps reconstructed from even and odd frames (evaluated at 0.143) and FSC between the final 3D map and the fitted model (evaluated at 0.5). (D) Zoomed-in views of the final 3D density map from panel A, displayed at two contour levels. (E) Superimposition of the high-contour level map from panel D onto its fitted model.

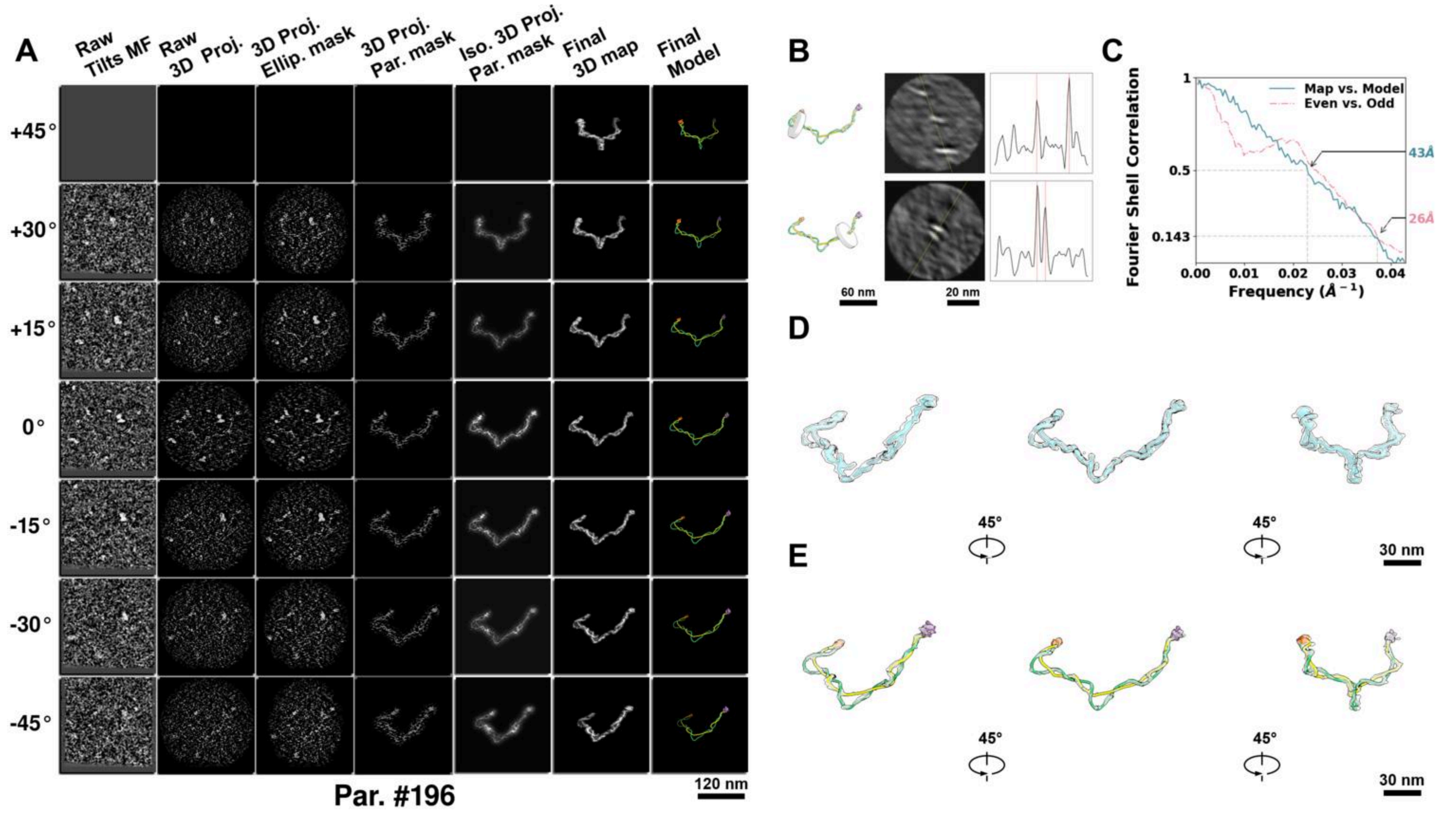

**Supplementary Particle Figure 196. Cryo-ET 3D reconstruction of an individual TEC-Cas particle.**
(A) 3D reconstruction of the plasmid particle (index no. 196). The first column shows seven representative tilt images from +45° to -45° in step of 15°. The second, third, and fourth columns show 3D projections of the particle with spherical, ellipsoidal (thinner along the z-dimension), and particle-shaped masks, respectively. The fifth column displays the 3D projections of the enhanced and IsoNet missing-wedge-corrected particle. The sixth and seventh columns present the final 3D map and the flexibly fitted model, respectively. (B) Two cross-sectional views (12 nm thickness) of the plasmid density map along its plectoneme axis are shown in the left-middle panel. The intensity profile along the line crossing the two high-density DNA spots is displayed in the right panel. (C) Resolution assessment of the final 3D map using Fourier shell correlation (FSC). Two criteria are shown: FSC between two half-maps reconstructed from even and odd frames (evaluated at 0.143) and FSC between the final 3D map and the fitted model (evaluated at 0.5). (D) Zoomed-in views of the final 3D density map from panel A, displayed at two contour levels. (E) Superimposition of the high-contour level map from panel D onto its fitted model.

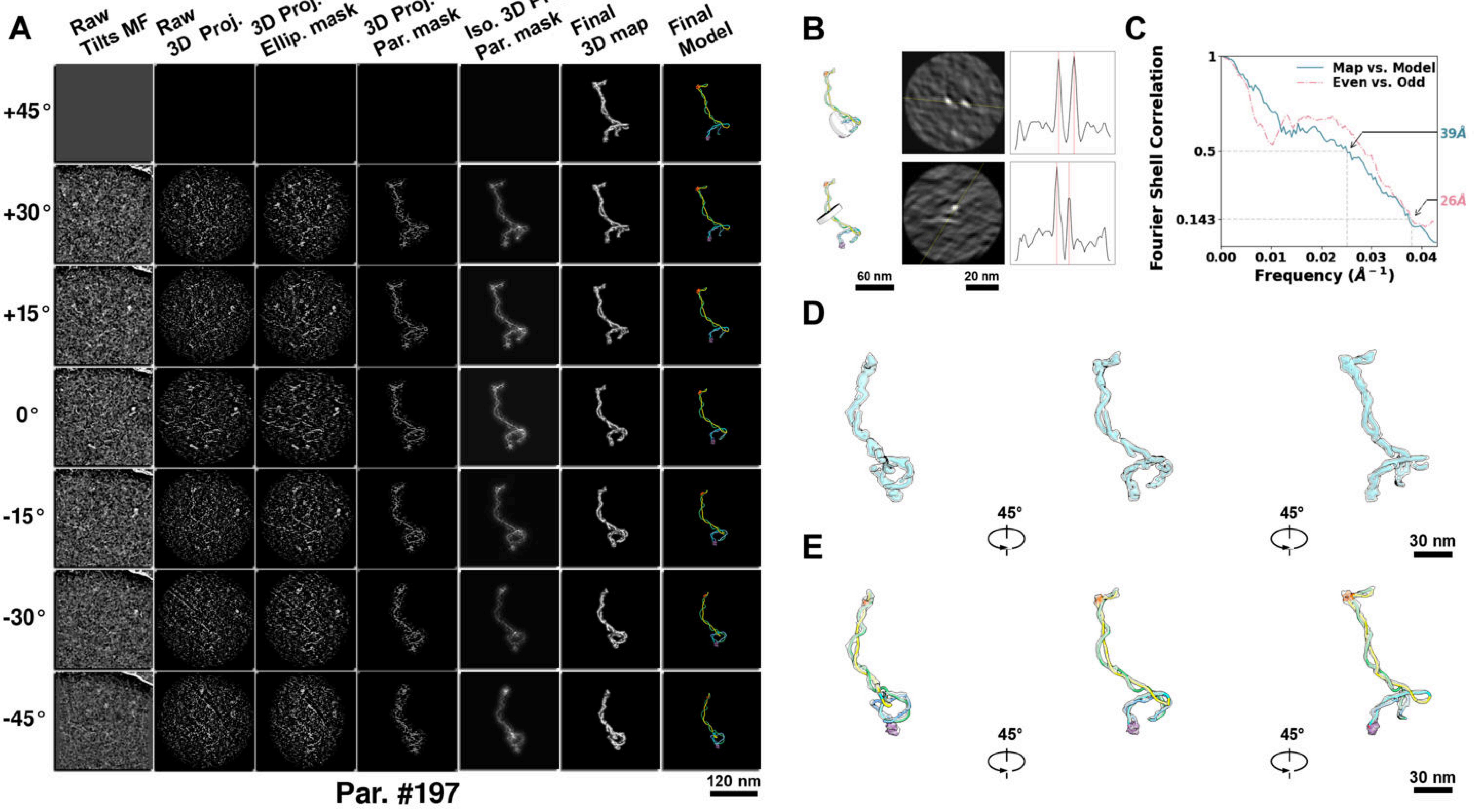

**Supplementary Particle Figure 197. Cryo-ET 3D reconstruction of an individual TEC-Cas particle.**
(A) 3D reconstruction of the plasmid particle (index no. 197). The first column shows seven representative tilt images from +45° to -45° in step of 15°. The second, third, and fourth columns show 3D projections of the particle with spherical, ellipsoidal (thinner along the z-dimension), and particle-shaped masks, respectively. The fifth column displays the 3D projections of the enhanced and IsoNet missing-wedge-corrected particle. The sixth and seventh columns present the final 3D map and the flexibly fitted model, respectively. (B) Two cross-sectional views (12 nm thickness) of the plasmid density map along its plectoneme axis are shown in the left-middle panel. The intensity profile along the line crossing the two high-density DNA spots is displayed in the right panel. (C) Resolution assessment of the final 3D map using Fourier shell correlation (FSC). Two criteria are shown: FSC between two half-maps reconstructed from even and odd frames (evaluated at 0.143) and FSC between the final 3D map and the fitted model (evaluated at 0.5). (D) Zoomed-in views of the final 3D density map from panel A, displayed at two contour levels. (E) Superimposition of the high-contour level map from panel D onto its fitted model.

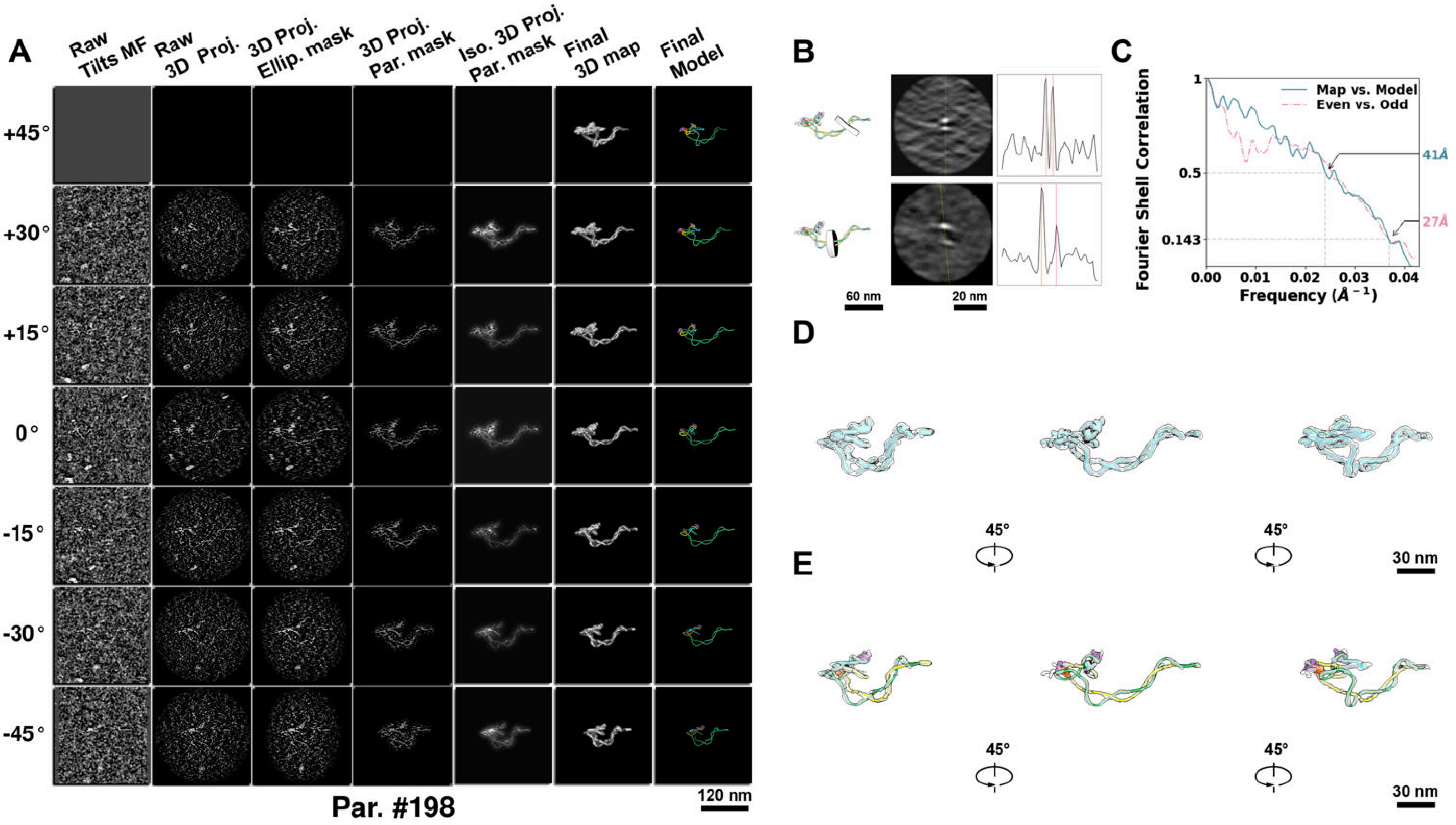

**Supplementary Particle Figure 198. Cryo-ET 3D reconstruction of an individual TEC-Cas particle.**
(A) 3D reconstruction of the plasmid particle (index no. 198). The first column shows seven representative tilt images from +45° to -45° in step of 15°. The second, third, and fourth columns show 3D projections of the particle with spherical, ellipsoidal (thinner along the z-dimension), and particle-shaped masks, respectively. The fifth column displays the 3D projections of the enhanced and IsoNet missing-wedge-corrected particle. The sixth and seventh columns present the final 3D map and the flexibly fitted model, respectively. (B) Two cross-sectional views (12 nm thickness) of the plasmid density map along its plectoneme axis are shown in the left-middle panel. The intensity profile along the line crossing the two high-density DNA spots is displayed in the right panel. (C) Resolution assessment of the final 3D map using Fourier shell correlation (FSC). Two criteria are shown: FSC between two half-maps reconstructed from even and odd frames (evaluated at 0.143) and FSC between the final 3D map and the fitted model (evaluated at 0.5). (D) Zoomed-in views of the final 3D density map from panel A, displayed at two contour levels. (E) Superimposition of the high-contour level map from panel D onto its fitted model.

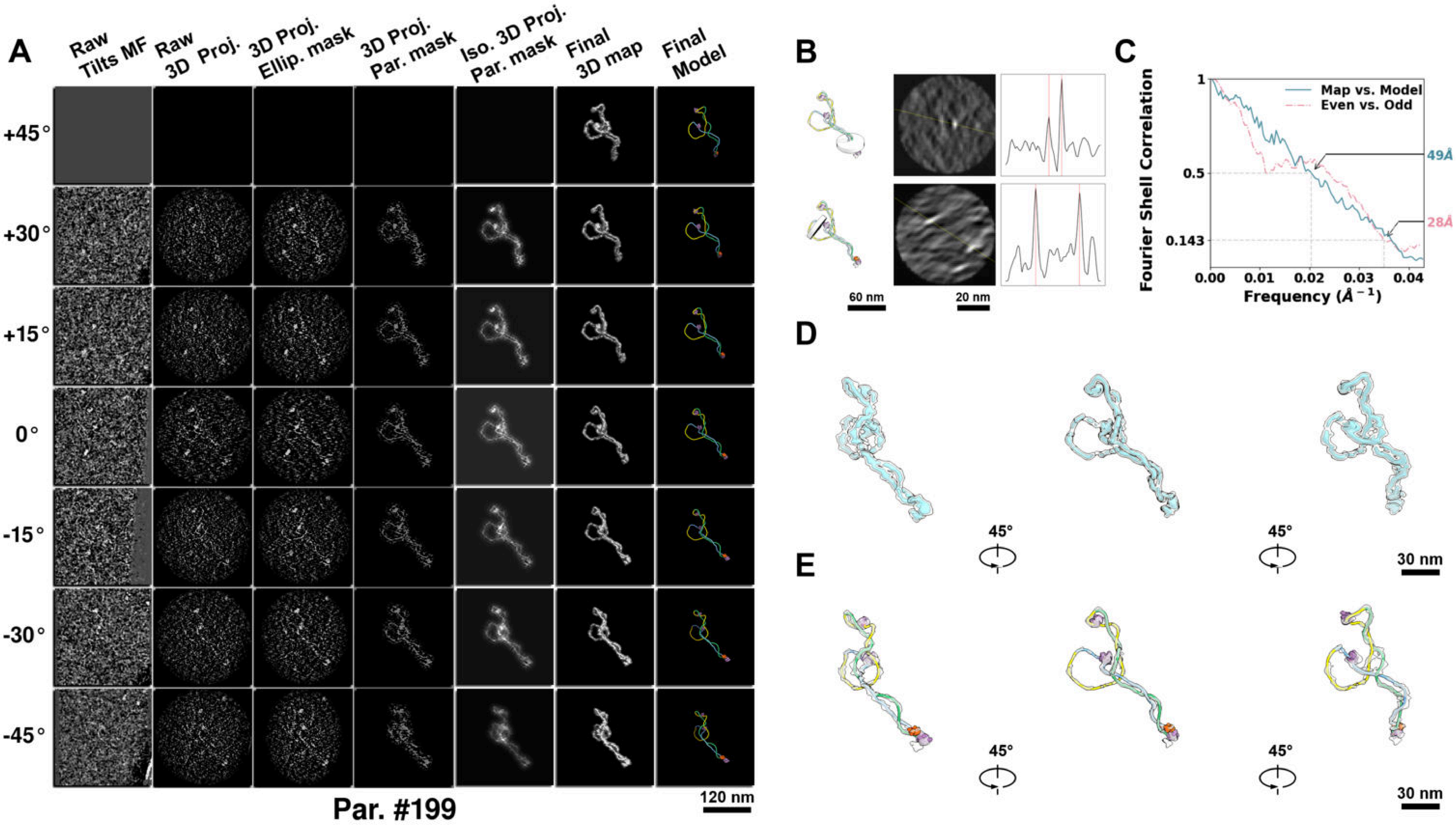

**Supplementary Particle Figure 199. Cryo-ET 3D reconstruction of an individual TEC-Cas particle.**
(A) 3D reconstruction of the plasmid particle (index no. 199). The first column shows seven representative tilt images from +45° to -45° in step of 15°. The second, third, and fourth columns show 3D projections of the particle with spherical, ellipsoidal (thinner along the z-dimension), and particle-shaped masks, respectively. The fifth column displays the 3D projections of the enhanced and IsoNet missing-wedge-corrected particle. The sixth and seventh columns present the final 3D map and the flexibly fitted model, respectively. (B) Two cross-sectional views (12 nm thickness) of the plasmid density map along its plectoneme axis are shown in the left-middle panel. The intensity profile along the line crossing the two high-density DNA spots is displayed in the right panel. (C) Resolution assessment of the final 3D map using Fourier shell correlation (FSC). Two criteria are shown: FSC between two half-maps reconstructed from even and odd frames (evaluated at 0.143) and FSC between the final 3D map and the fitted model (evaluated at 0.5). (D) Zoomed-in views of the final 3D density map from panel A, displayed at two contour levels. (E) Superimposition of the high-contour level map from panel D onto its fitted model.

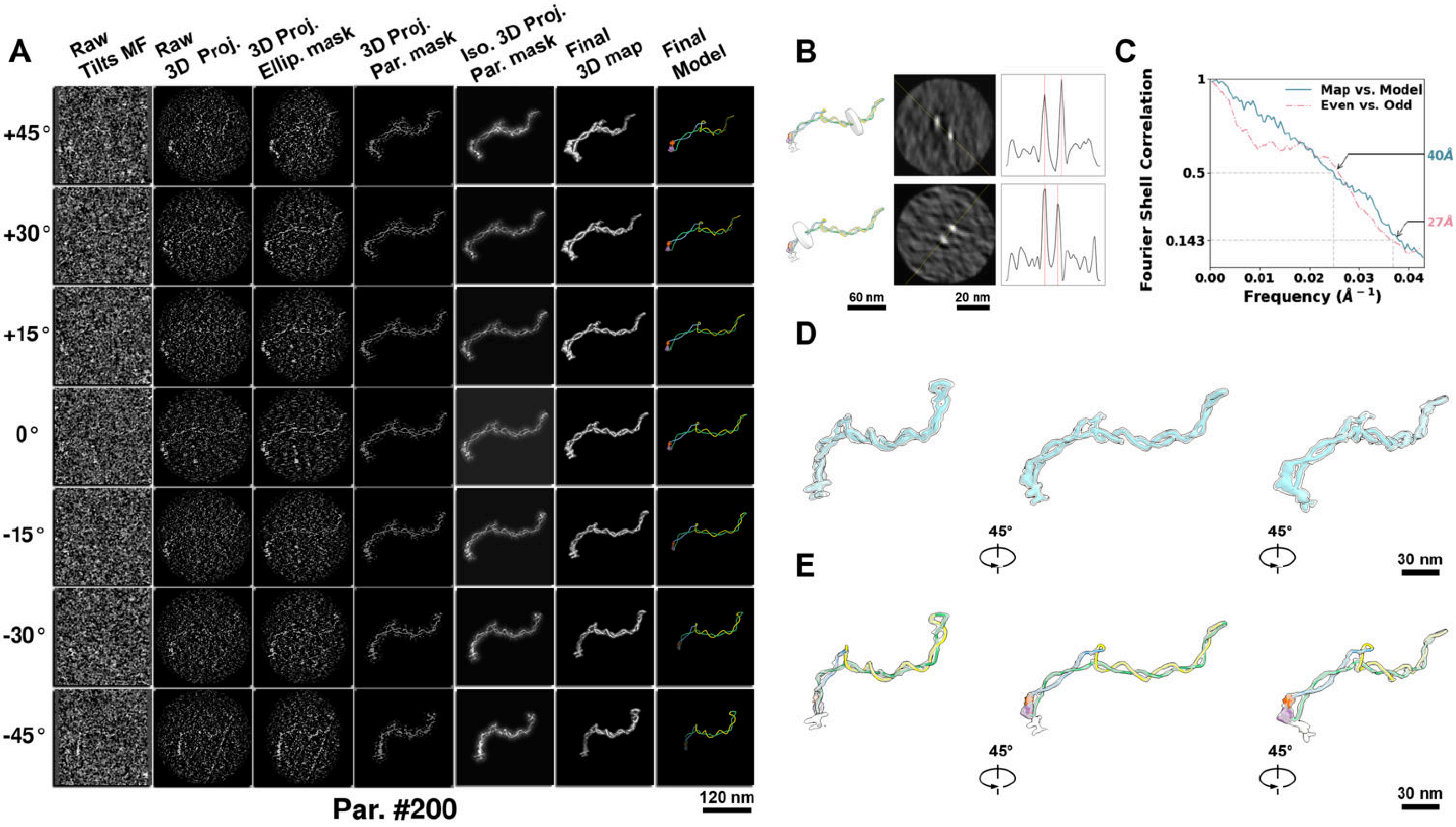

**Supplementary Particle Figure 200. Cryo-ET 3D reconstruction of an individual TEC-Cas particle.**
(A) 3D reconstruction of the plasmid particle (index no. 200). The first column shows seven representative tilt images from +45° to -45° in step of 15°. The second, third, and fourth columns show 3D projections of the particle with spherical, ellipsoidal (thinner along the z-dimension), and particle-shaped masks, respectively. The fifth column displays the 3D projections of the enhanced and IsoNet missing-wedge-corrected particle. The sixth and seventh columns present the final 3D map and the flexibly fitted model, respectively. (B) Two cross-sectional views (12 nm thickness) of the plasmid density map along its plectoneme axis are shown in the left-middle panel. The intensity profile along the line crossing the two high-density DNA spots is displayed in the right panel. (C) Resolution assessment of the final 3D map using Fourier shell correlation (FSC). Two criteria are shown: FSC between two half-maps reconstructed from even and odd frames (evaluated at 0.143) and FSC between the final 3D map and the fitted model (evaluated at 0.5). (D) Zoomed-in views of the final 3D density map from panel A, displayed at two contour levels. (E) Superimposition of the high-contour level map from panel D onto its fitted model.

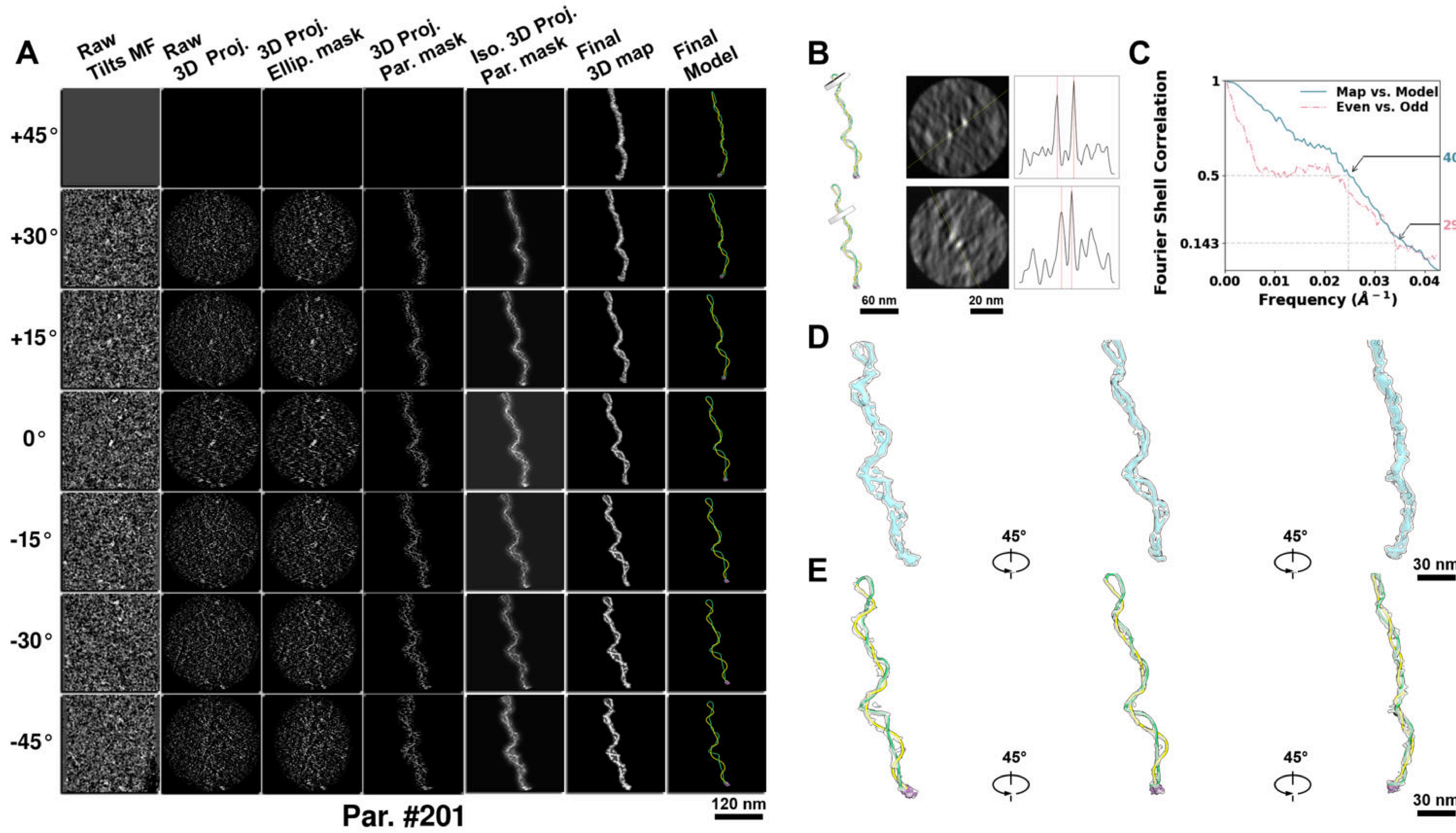

**Supplementary Particle Figure 201. Cryo-ET 3D reconstruction of an individual TEC-Cas particle.**
(A) 3D reconstruction of the plasmid particle (index no. 201). The first column shows seven representative tilt images from +45° to -45° in step of 15°. The second, third, and fourth columns show 3D projections of the particle with spherical, ellipsoidal (thinner along the z-dimension), and particle-shaped masks, respectively. The fifth column displays the 3D projections of the enhanced and IsoNet missing-wedge-corrected particle. The sixth and seventh columns present the final 3D map and the flexibly fitted model, respectively. (B) Two cross-sectional views (12 nm thickness) of the plasmid density map along its plectoneme axis are shown in the left-middle panel. The intensity profile along the line crossing the two high-density DNA spots is displayed in the right panel. (C) Resolution assessment of the final 3D map using Fourier shell correlation (FSC). Two criteria are shown: FSC between two half-maps reconstructed from even and odd frames (evaluated at 0.143) and FSC between the final 3D map and the fitted model (evaluated at 0.5). (D) Zoomed-in views of the final 3D density map from panel A, displayed at two contour levels. (E) Superimposition of the high-contour level map from panel D onto its fitted model.

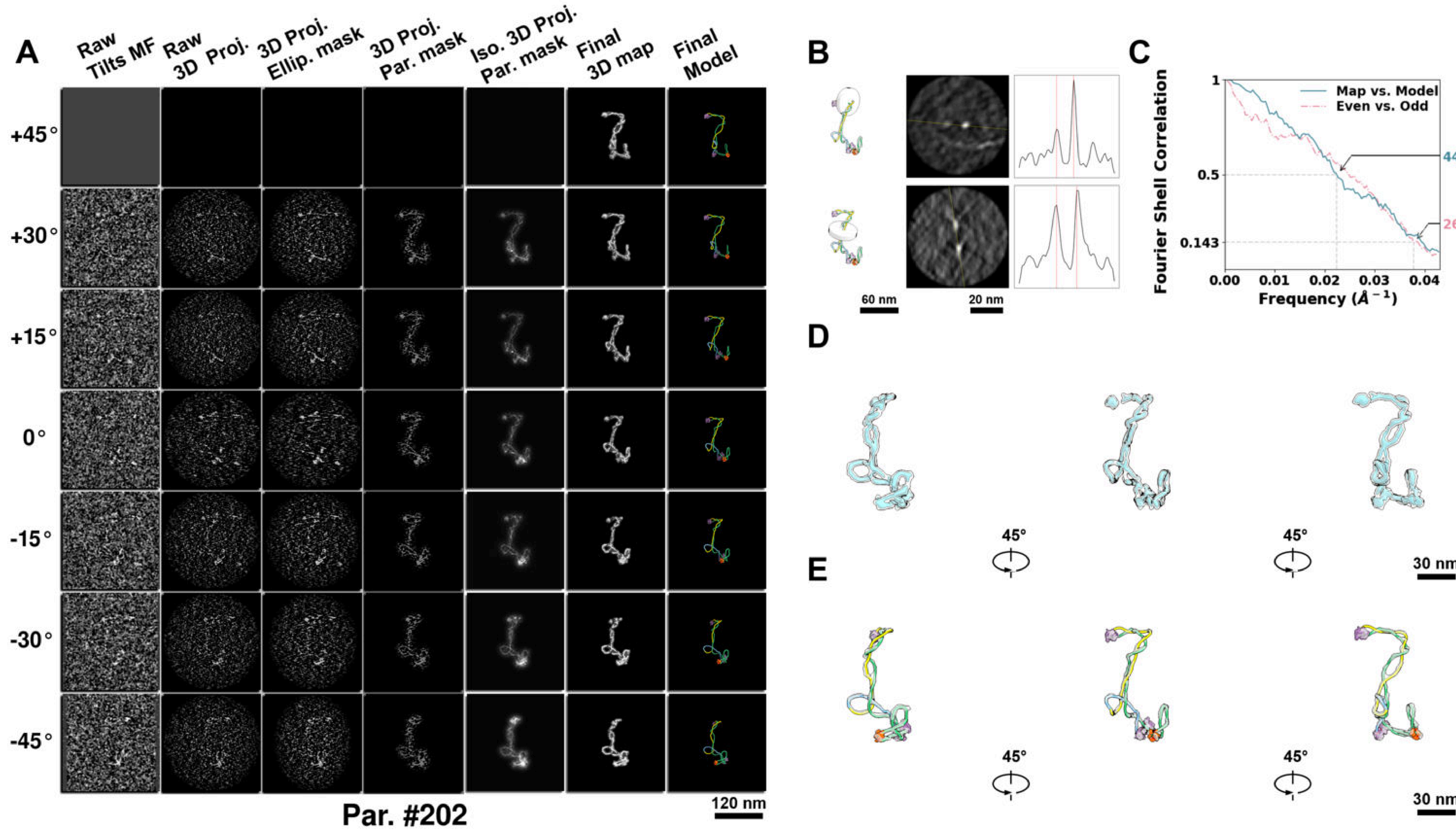

**Supplementary Particle Figure 202. Cryo-ET 3D reconstruction of an individual TEC-Cas particle.**
(A) 3D reconstruction of the plasmid particle (index no. 202). The first column shows seven representative tilt images from +45° to -45° in step of 15°. The second, third, and fourth columns show 3D projections of the particle with spherical, ellipsoidal (thinner along the z-dimension), and particle-shaped masks, respectively. The fifth column displays the 3D projections of the enhanced and IsoNet missing-wedge-corrected particle. The sixth and seventh columns present the final 3D map and the flexibly fitted model, respectively. (B) Two cross-sectional views (12 nm thickness) of the plasmid density map along its plectoneme axis are shown in the left-middle panel. The intensity profile along the line crossing the two high-density DNA spots is displayed in the right panel. (C) Resolution assessment of the final 3D map using Fourier shell correlation (FSC). Two criteria are shown: FSC between two half-maps reconstructed from even and odd frames (evaluated at 0.143) and FSC between the final 3D map and the fitted model (evaluated at 0.5). (D) Zoomed-in views of the final 3D density map from panel A, displayed at two contour levels. (E) Superimposition of the high-contour level map from panel D onto its fitted model.

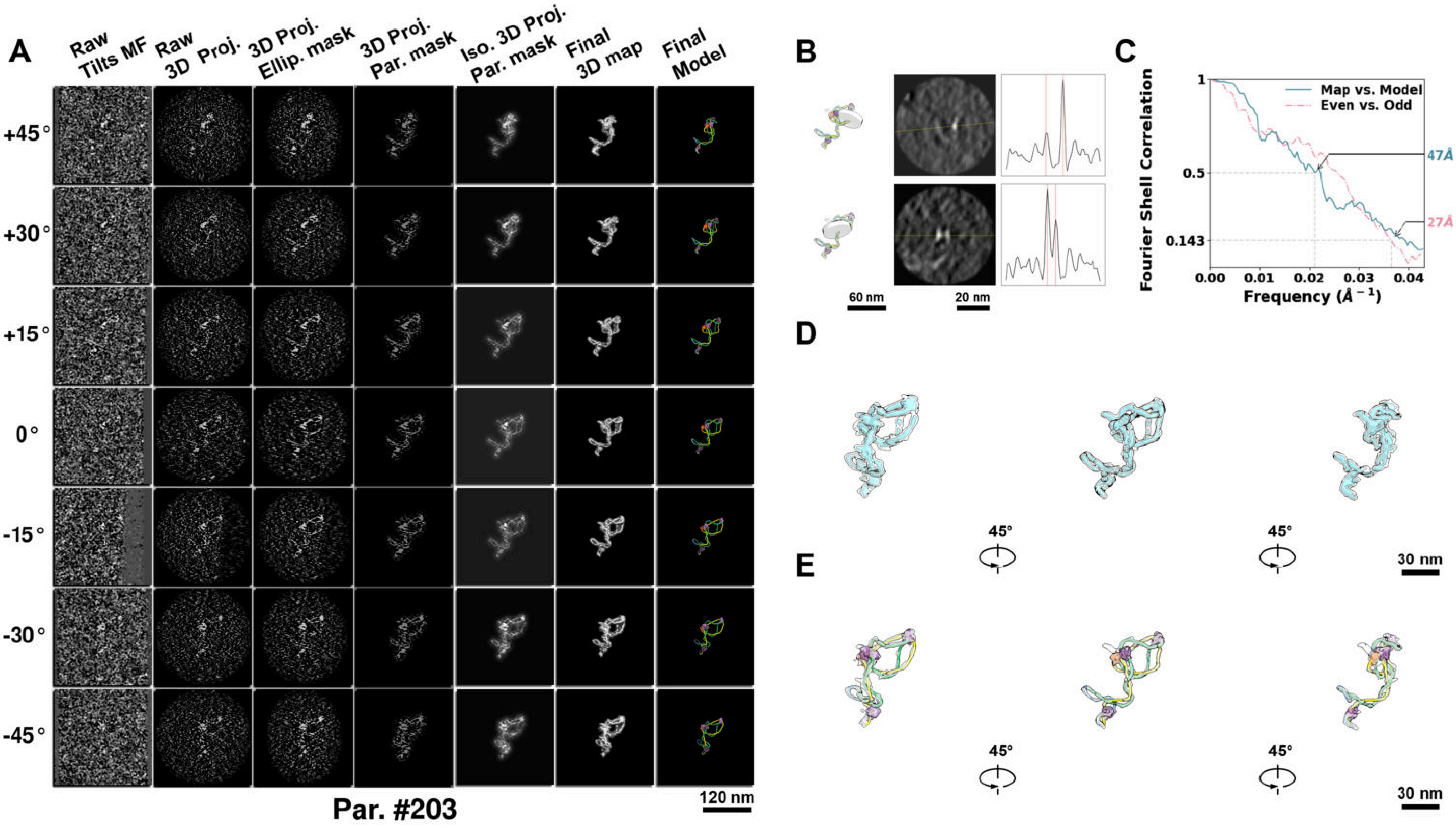

**Supplementary Particle Figure 203. Cryo-ET 3D reconstruction of an individual TEC-Cas particle.**
(A) 3D reconstruction of the plasmid particle (index no. 203). The first column shows seven representative tilt images from +45° to -45° in step of 15°. The second, third, and fourth columns show 3D projections of the particle with spherical, ellipsoidal (thinner along the z-dimension), and particle-shaped masks, respectively. The fifth column displays the 3D projections of the enhanced and IsoNet missing-wedge-corrected particle. The sixth and seventh columns present the final 3D map and the flexibly fitted model, respectively. (B) Two cross-sectional views (12 nm thickness) of the plasmid density map along its plectoneme axis are shown in the left-middle panel. The intensity profile along the line crossing the two high-density DNA spots is displayed in the right panel. (C) Resolution assessment of the final 3D map using Fourier shell correlation (FSC). Two criteria are shown: FSC between two half-maps reconstructed from even and odd frames (evaluated at 0.143) and FSC between the final 3D map and the fitted model (evaluated at 0.5). (D) Zoomed-in views of the final 3D density map from panel A, displayed at two contour levels. (E) Superimposition of the high-contour level map from panel D onto its fitted model.

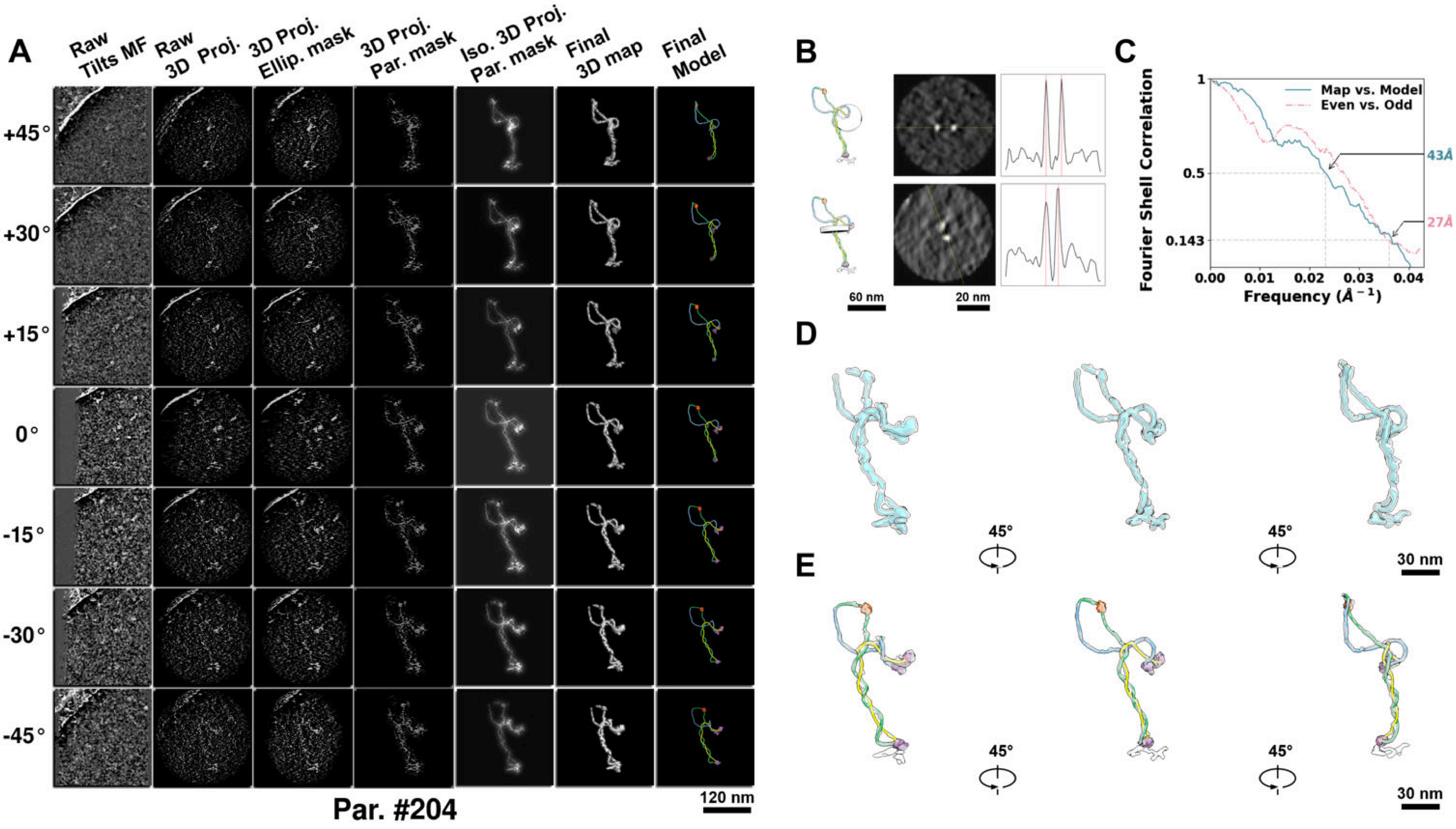

**Supplementary Particle Figure 204. Cryo-ET 3D reconstruction of an individual TEC-Cas particle.**
(A) 3D reconstruction of the plasmid particle (index no. 204). The first column shows seven representative tilt images from +45° to -45° in step of 15°. The second, third, and fourth columns show 3D projections of the particle with spherical, ellipsoidal (thinner along the z-dimension), and particle-shaped masks, respectively. The fifth column displays the 3D projections of the enhanced and IsoNet missing-wedge-corrected particle. The sixth and seventh columns present the final 3D map and the flexibly fitted model, respectively. (B) Two cross-sectional views (12 nm thickness) of the plasmid density map along its plectoneme axis are shown in the left-middle panel. The intensity profile along the line crossing the two high-density DNA spots is displayed in the right panel. (C) Resolution assessment of the final 3D map using Fourier shell correlation (FSC). Two criteria are shown: FSC between two half-maps reconstructed from even and odd frames (evaluated at 0.143) and FSC between the final 3D map and the fitted model (evaluated at 0.5). (D) Zoomed-in views of the final 3D density map from panel A, displayed at two contour levels. (E) Superimposition of the high-contour level map from panel D onto its fitted model.

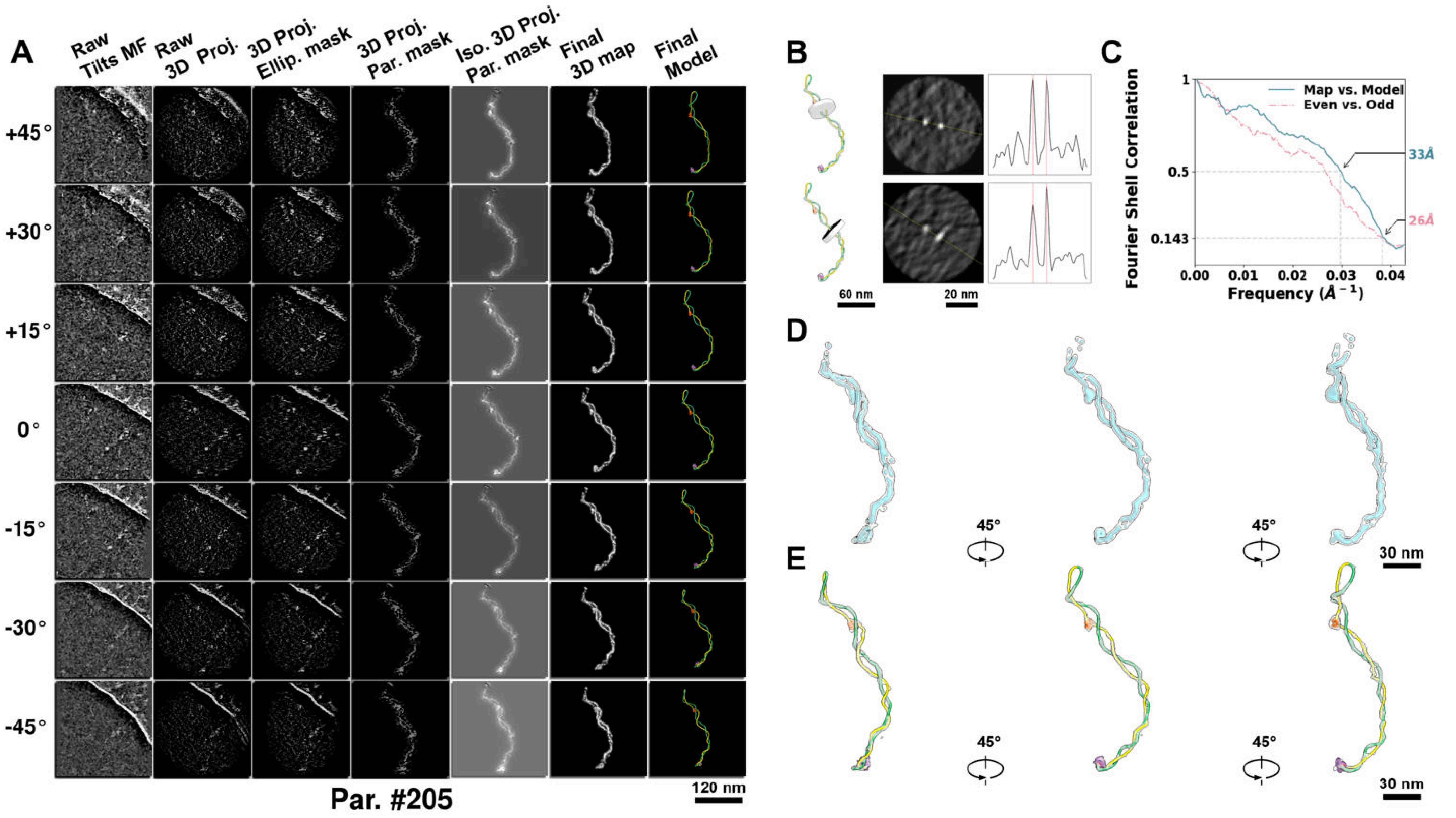

**Supplementary Particle Figure 205. Cryo-ET 3D reconstruction of an individual TEC-Cas particle.**
(A) 3D reconstruction of the plasmid particle (index no. 205). The first column shows seven representative tilt images from +45° to -45° in step of 15°. The second, third, and fourth columns show 3D projections of the particle with spherical, ellipsoidal (thinner along the z-dimension), and particle-shaped masks, respectively. The fifth column displays the 3D projections of the enhanced and IsoNet missing-wedge-corrected particle. The sixth and seventh columns present the final 3D map and the flexibly fitted model, respectively. (B) Two cross-sectional views (12 nm thickness) of the plasmid density map along its plectoneme axis are shown in the left-middle panel. The intensity profile along the line crossing the two high-density DNA spots is displayed in the right panel. (C) Resolution assessment of the final 3D map using Fourier shell correlation (FSC). Two criteria are shown: FSC between two half-maps reconstructed from even and odd frames (evaluated at 0.143) and FSC between the final 3D map and the fitted model (evaluated at 0.5). (D) Zoomed-in views of the final 3D density map from panel A, displayed at two contour levels. (E) Superimposition of the high-contour level map from panel D onto its fitted model.

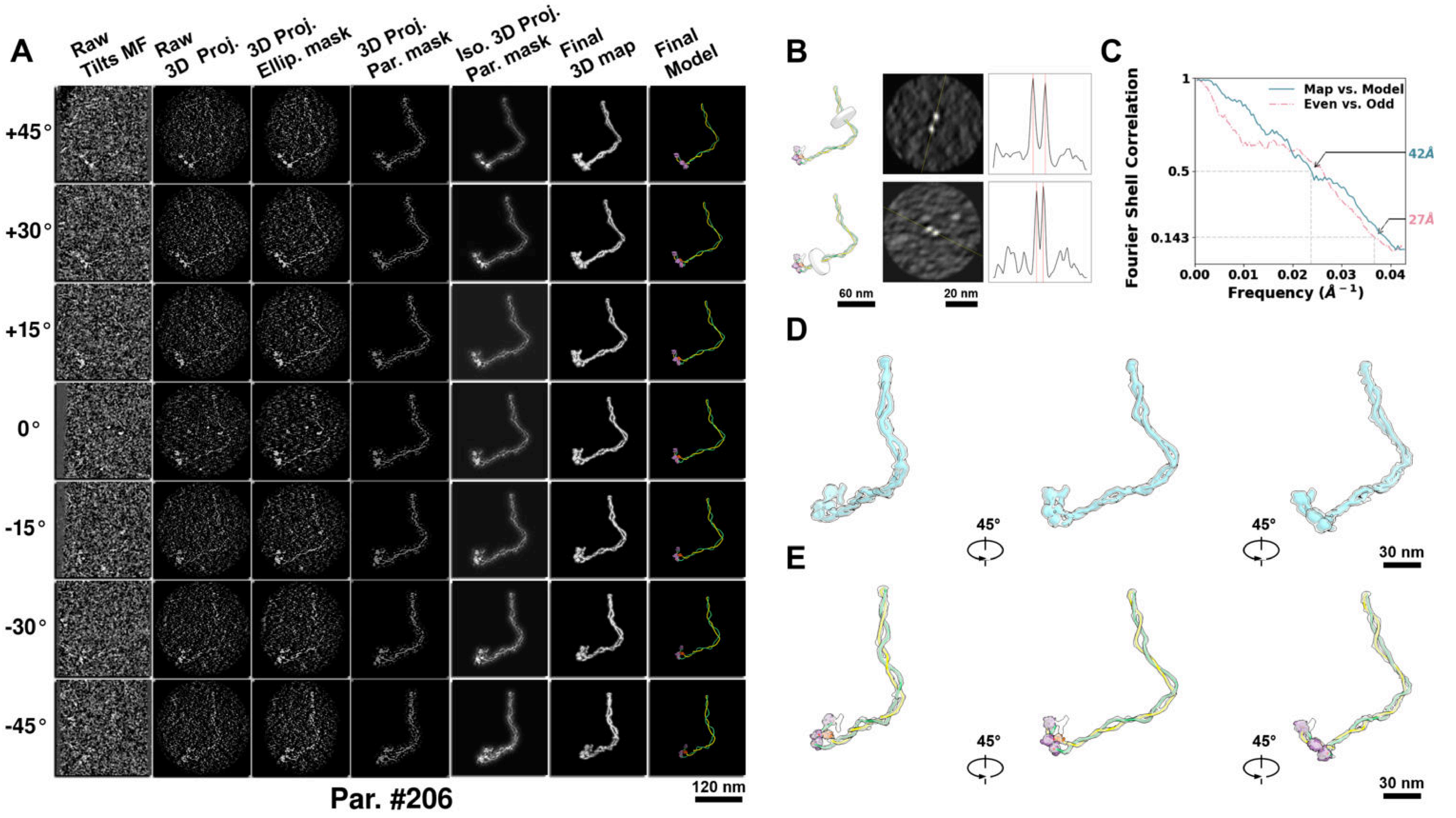

**Supplementary Particle Figure 206. Cryo-ET 3D reconstruction of an individual TEC-Cas particle.**
(A) 3D reconstruction of the plasmid particle (index no. 206). The first column shows seven representative tilt images from +45° to -45° in step of 15°. The second, third, and fourth columns show 3D projections of the particle with spherical, ellipsoidal (thinner along the z-dimension), and particle-shaped masks, respectively. The fifth column displays the 3D projections of the enhanced and IsoNet missing-wedge-corrected particle. The sixth and seventh columns present the final 3D map and the flexibly fitted model, respectively. (B) Two cross-sectional views (12 nm thickness) of the plasmid density map along its plectoneme axis are shown in the left-middle panel. The intensity profile along the line crossing the two high-density DNA spots is displayed in the right panel. (C) Resolution assessment of the final 3D map using Fourier shell correlation (FSC). Two criteria are shown: FSC between two half-maps reconstructed from even and odd frames (evaluated at 0.143) and FSC between the final 3D map and the fitted model (evaluated at 0.5). (D) Zoomed-in views of the final 3D density map from panel A, displayed at two contour levels. (E) Superimposition of the high-contour level map from panel D onto its fitted model.

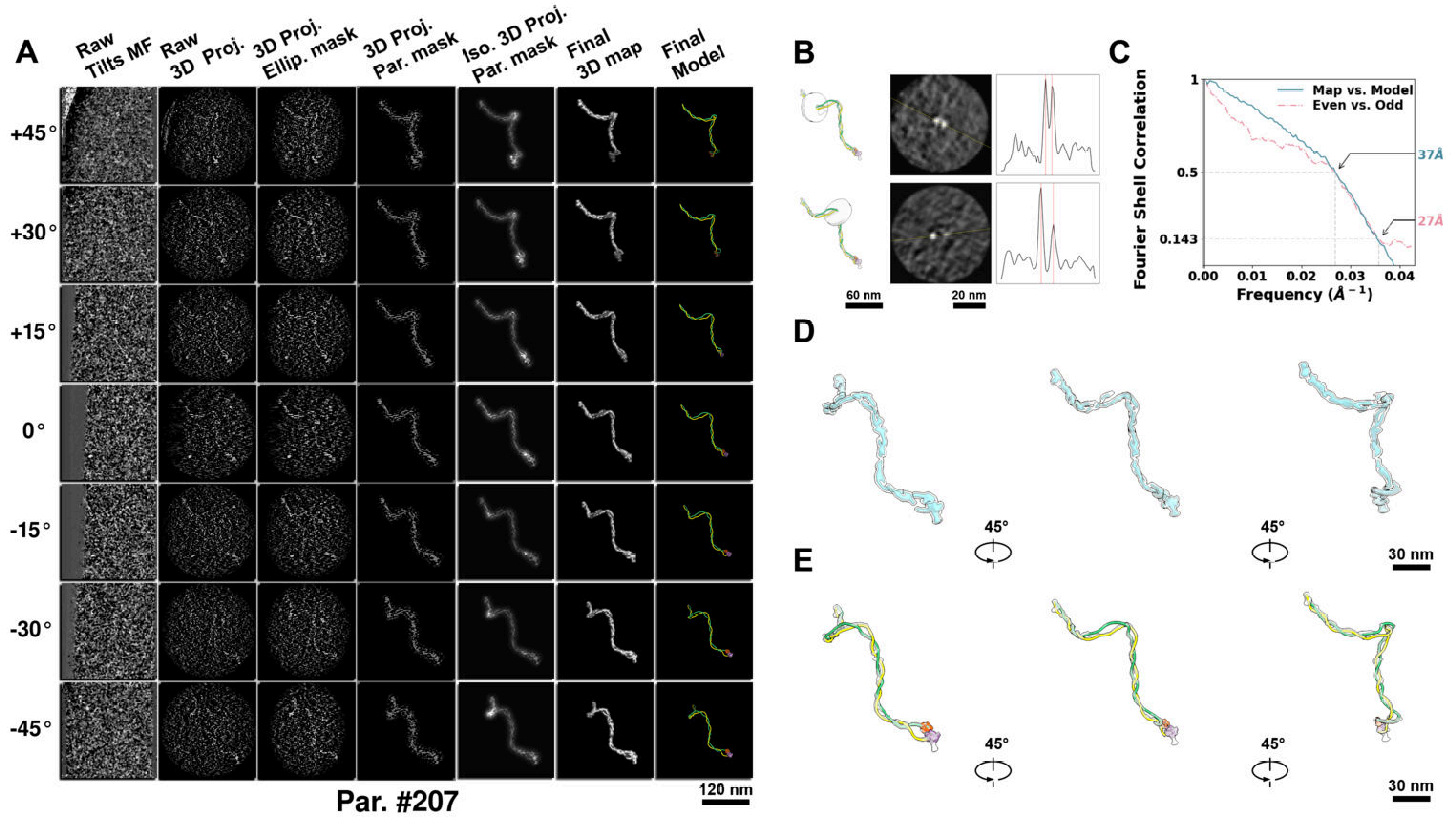

**Supplementary Particle Figure 207. Cryo-ET 3D reconstruction of an individual TEC-Cas particle.**
(A) 3D reconstruction of the plasmid particle (index no. 207). The first column shows seven representative tilt images from +45° to -45° in step of 15°. The second, third, and fourth columns show 3D projections of the particle with spherical, ellipsoidal (thinner along the z-dimension), and particle-shaped masks, respectively. The fifth column displays the 3D projections of the enhanced and IsoNet missing-wedge-corrected particle. The sixth and seventh columns present the final 3D map and the flexibly fitted model, respectively. (B) Two cross-sectional views (12 nm thickness) of the plasmid density map along its plectoneme axis are shown in the left-middle panel. The intensity profile along the line crossing the two high-density DNA spots is displayed in the right panel. (C) Resolution assessment of the final 3D map using Fourier shell correlation (FSC). Two criteria are shown: FSC between two half-maps reconstructed from even and odd frames (evaluated at 0.143) and FSC between the final 3D map and the fitted model (evaluated at 0.5). (D) Zoomed-in views of the final 3D density map from panel A, displayed at two contour levels. (E) Superimposition of the high-contour level map from panel D onto its fitted model.

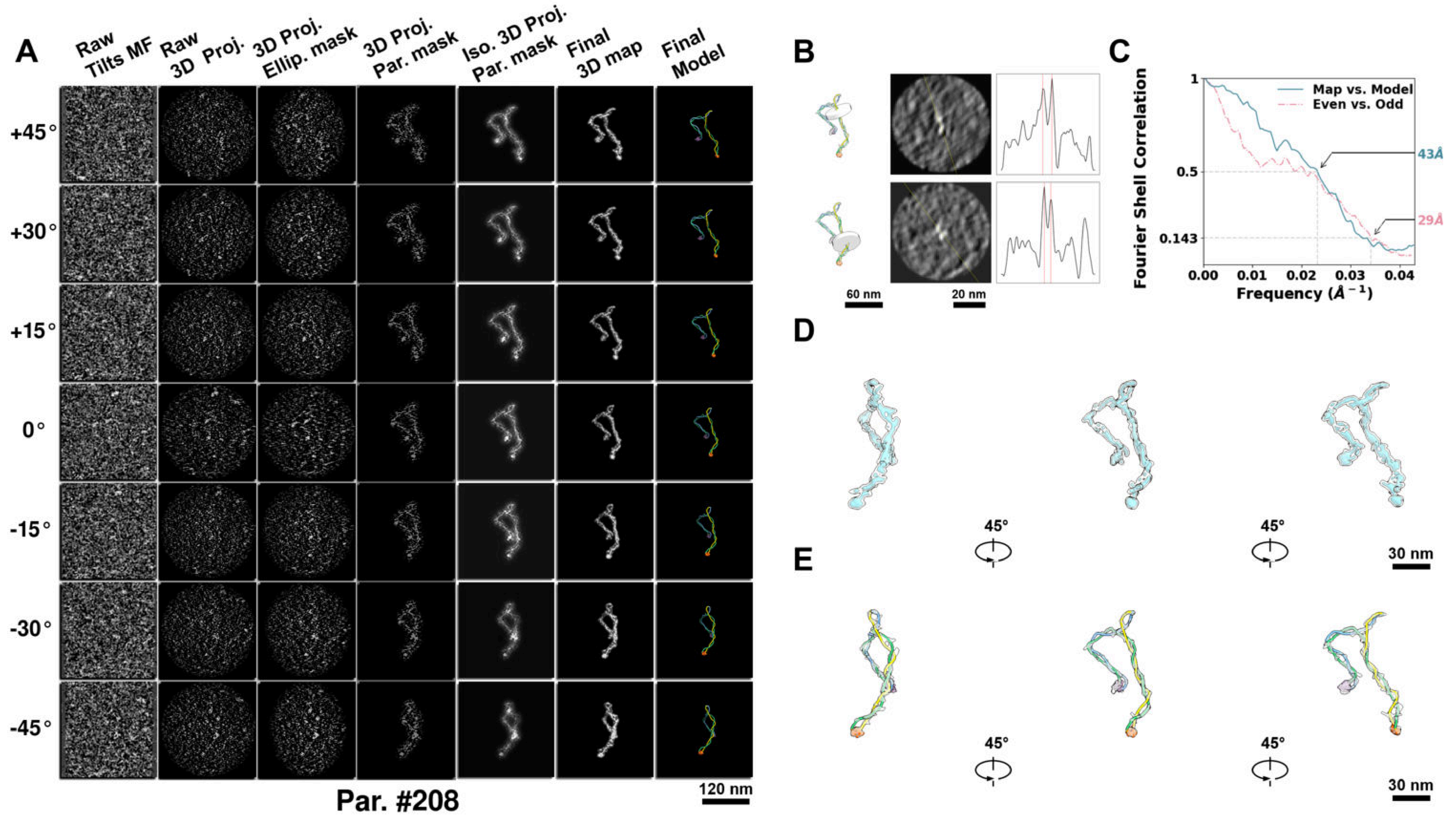

**Supplementary Particle Figure 208. Cryo-ET 3D reconstruction of an individual TEC-Cas particle.**
(A) 3D reconstruction of the plasmid particle (index no. 208). The first column shows seven representative tilt images from +45° to -45° in step of 15°. The second, third, and fourth columns show 3D projections of the particle with spherical, ellipsoidal (thinner along the z-dimension), and particle-shaped masks, respectively. The fifth column displays the 3D projections of the enhanced and IsoNet missing-wedge-corrected particle. The sixth and seventh columns present the final 3D map and the flexibly fitted model, respectively. (B) Two cross-sectional views (12 nm thickness) of the plasmid density map along its plectoneme axis are shown in the left-middle panel. The intensity profile along the line crossing the two high-density DNA spots is displayed in the right panel. (C) Resolution assessment of the final 3D map using Fourier shell correlation (FSC). Two criteria are shown: FSC between two half-maps reconstructed from even and odd frames (evaluated at 0.143) and FSC between the final 3D map and the fitted model (evaluated at 0.5). (D) Zoomed-in views of the final 3D density map from panel A, displayed at two contour levels. (E) Superimposition of the high-contour level map from panel D onto its fitted model.

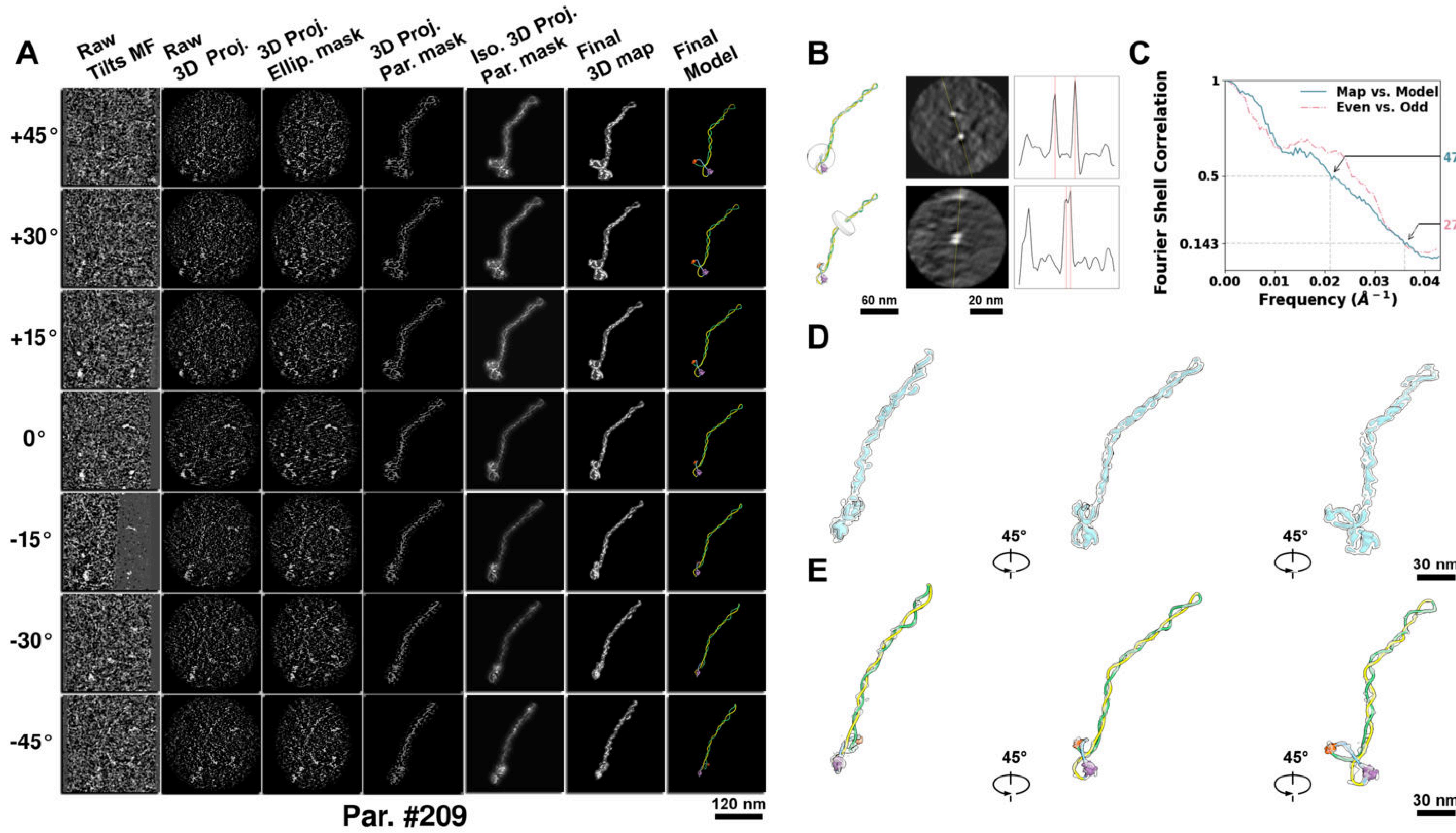

**Supplementary Particle Figure 209. Cryo-ET 3D reconstruction of an individual TEC-Cas particle.**
(A) 3D reconstruction of the plasmid particle (index no. 209). The first column shows seven representative tilt images from +45° to -45° in step of 15°. The second, third, and fourth columns show 3D projections of the particle with spherical, ellipsoidal (thinner along the z-dimension), and particle-shaped masks, respectively. The fifth column displays the 3D projections of the enhanced and IsoNet missing-wedge-corrected particle. The sixth and seventh columns present the final 3D map and the flexibly fitted model, respectively. (B) Two cross-sectional views (12 nm thickness) of the plasmid density map along its plectoneme axis are shown in the left-middle panel. The intensity profile along the line crossing the two high-density DNA spots is displayed in the right panel. (C) Resolution assessment of the final 3D map using Fourier shell correlation (FSC). Two criteria are shown: FSC between two half-maps reconstructed from even and odd frames (evaluated at 0.143) and FSC between the final 3D map and the fitted model (evaluated at 0.5). (D) Zoomed-in views of the final 3D density map from panel A, displayed at two contour levels. (E) Superimposition of the high-contour level map from panel D onto its fitted model.

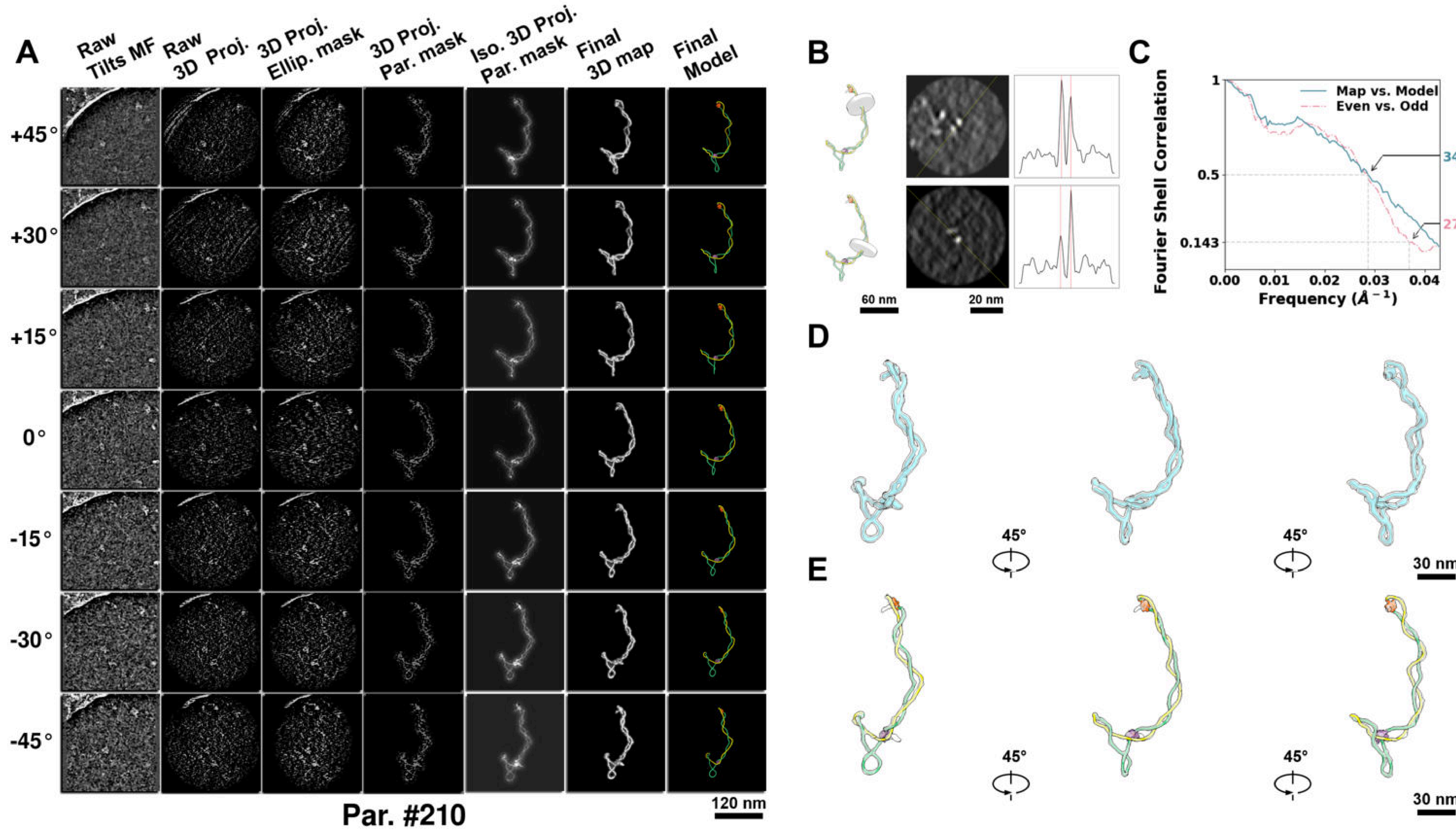

**Supplementary Particle Figure 210. Cryo-ET 3D reconstruction of an individual TEC-Cas particle.**
(A) 3D reconstruction of the plasmid particle (index no. 210). The first column shows seven representative tilt images from +45° to -45° in step of 15°. The second, third, and fourth columns show 3D projections of the particle with spherical, ellipsoidal (thinner along the z-dimension), and particle-shaped masks, respectively. The fifth column displays the 3D projections of the enhanced and IsoNet missing-wedge-corrected particle. The sixth and seventh columns present the final 3D map and the flexibly fitted model, respectively. (B) Two cross-sectional views (12 nm thickness) of the plasmid density map along its plectoneme axis are shown in the left-middle panel. The intensity profile along the line crossing the two high-density DNA spots is displayed in the right panel. (C) Resolution assessment of the final 3D map using Fourier shell correlation (FSC). Two criteria are shown: FSC between two half-maps reconstructed from even and odd frames (evaluated at 0.143) and FSC between the final 3D map and the fitted model (evaluated at 0.5). (D) Zoomed-in views of the final 3D density map from panel A, displayed at two contour levels. (E) Superimposition of the high-contour level map from panel D onto its fitted model.

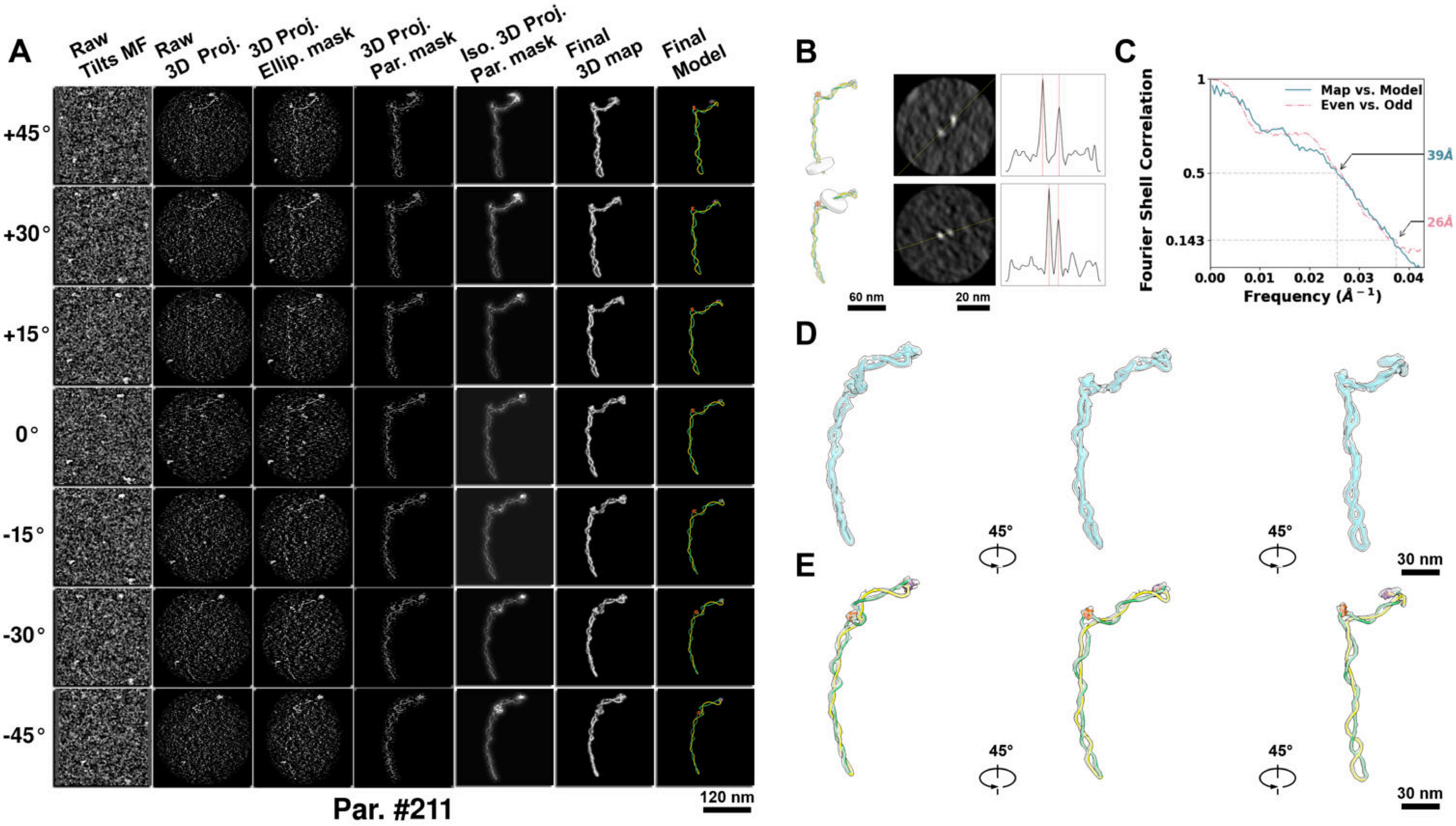

**Supplementary Particle Figure 211. Cryo-ET 3D reconstruction of an individual TEC-Cas particle.**
(A) 3D reconstruction of the plasmid particle (index no. 211). The first column shows seven representative tilt images from +45° to -45° in step of 15°. The second, third, and fourth columns show 3D projections of the particle with spherical, ellipsoidal (thinner along the z-dimension), and particle-shaped masks, respectively. The fifth column displays the 3D projections of the enhanced and IsoNet missing-wedge-corrected particle. The sixth and seventh columns present the final 3D map and the flexibly fitted model, respectively. (B) Two cross-sectional views (12 nm thickness) of the plasmid density map along its plectoneme axis are shown in the left-middle panel. The intensity profile along the line crossing the two high-density DNA spots is displayed in the right panel. (C) Resolution assessment of the final 3D map using Fourier shell correlation (FSC). Two criteria are shown: FSC between two half-maps reconstructed from even and odd frames (evaluated at 0.143) and FSC between the final 3D map and the fitted model (evaluated at 0.5). (D) Zoomed-in views of the final 3D density map from panel A, displayed at two contour levels. (E) Superimposition of the high-contour level map from panel D onto its fitted model.

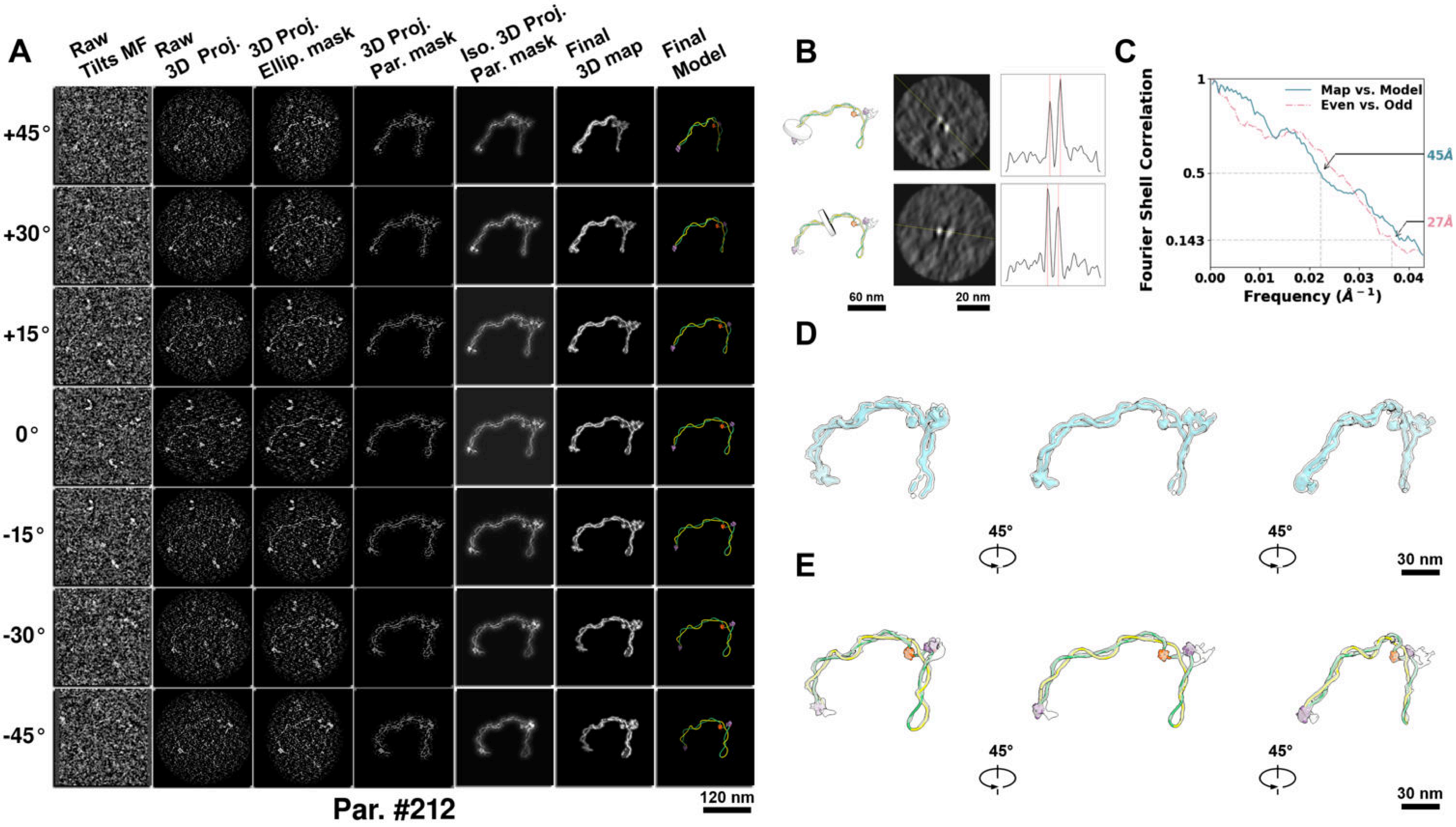

**Supplementary Particle Figure 212. Cryo-ET 3D reconstruction of an individual TEC-Cas particle.**
(A) 3D reconstruction of the plasmid particle (index no. 212). The first column shows seven representative tilt images from +45° to -45° in step of 15°. The second, third, and fourth columns show 3D projections of the particle with spherical, ellipsoidal (thinner along the z-dimension), and particle-shaped masks, respectively. The fifth column displays the 3D projections of the enhanced and IsoNet missing-wedge-corrected particle. The sixth and seventh columns present the final 3D map and the flexibly fitted model, respectively. (B) Two cross-sectional views (12 nm thickness) of the plasmid density map along its plectoneme axis are shown in the left-middle panel. The intensity profile along the line crossing the two high-density DNA spots is displayed in the right panel. (C) Resolution assessment of the final 3D map using Fourier shell correlation (FSC). Two criteria are shown: FSC between two half-maps reconstructed from even and odd frames (evaluated at 0.143) and FSC between the final 3D map and the fitted model (evaluated at 0.5). (D) Zoomed-in views of the final 3D density map from panel A, displayed at two contour levels. (E) Superimposition of the high-contour level map from panel D onto its fitted model.

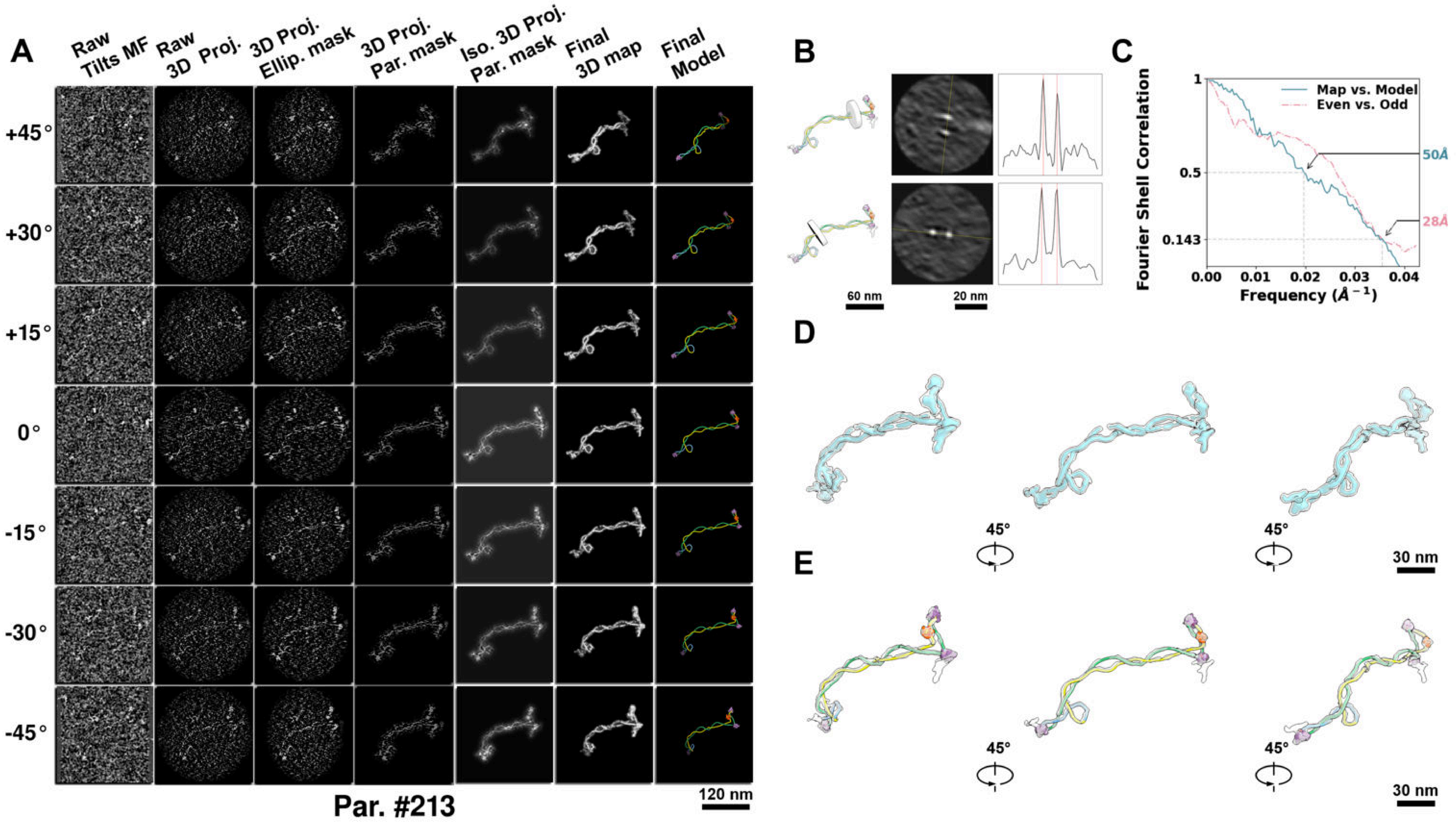

**Supplementary Particle Figure 213. Cryo-ET 3D reconstruction of an individual TEC-Cas particle.**
(A) 3D reconstruction of the plasmid particle (index no. 213). The first column shows seven representative tilt images from +45° to -45° in step of 15°. The second, third, and fourth columns show 3D projections of the particle with spherical, ellipsoidal (thinner along the z-dimension), and particle-shaped masks, respectively. The fifth column displays the 3D projections of the enhanced and IsoNet missing-wedge-corrected particle. The sixth and seventh columns present the final 3D map and the flexibly fitted model, respectively. (B) Two cross-sectional views (12 nm thickness) of the plasmid density map along its plectoneme axis are shown in the left-middle panel. The intensity profile along the line crossing the two high-density DNA spots is displayed in the right panel. (C) Resolution assessment of the final 3D map using Fourier shell correlation (FSC). Two criteria are shown: FSC between two half-maps reconstructed from even and odd frames (evaluated at 0.143) and FSC between the final 3D map and the fitted model (evaluated at 0.5). (D) Zoomed-in views of the final 3D density map from panel A, displayed at two contour levels. (E) Superimposition of the high-contour level map from panel D onto its fitted model.

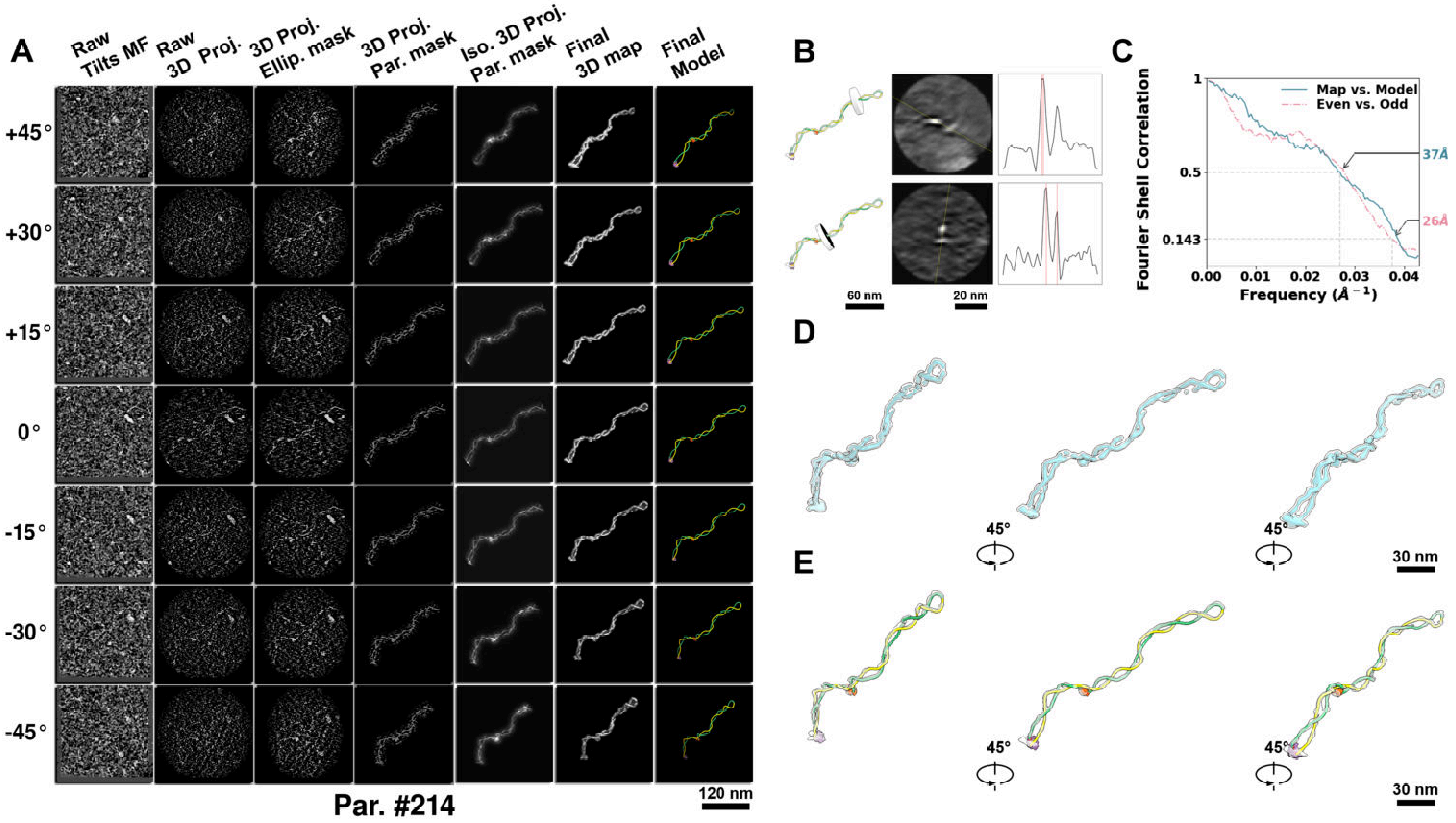

**Supplementary Particle Figure 214. Cryo-ET 3D reconstruction of an individual TEC-Cas particle.**
(A) 3D reconstruction of the plasmid particle (index no. 214). The first column shows seven representative tilt images from +45° to -45° in step of 15°. The second, third, and fourth columns show 3D projections of the particle with spherical, ellipsoidal (thinner along the z-dimension), and particle-shaped masks, respectively. The fifth column displays the 3D projections of the enhanced and IsoNet missing-wedge-corrected particle. The sixth and seventh columns present the final 3D map and the flexibly fitted model, respectively. (B) Two cross-sectional views (12 nm thickness) of the plasmid density map along its plectoneme axis are shown in the left-middle panel. The intensity profile along the line crossing the two high-density DNA spots is displayed in the right panel. (C) Resolution assessment of the final 3D map using Fourier shell correlation (FSC). Two criteria are shown: FSC between two half-maps reconstructed from even and odd frames (evaluated at 0.143) and FSC between the final 3D map and the fitted model (evaluated at 0.5). (D) Zoomed-in views of the final 3D density map from panel A, displayed at two contour levels. (E) Superimposition of the high-contour level map from panel D onto its fitted model.

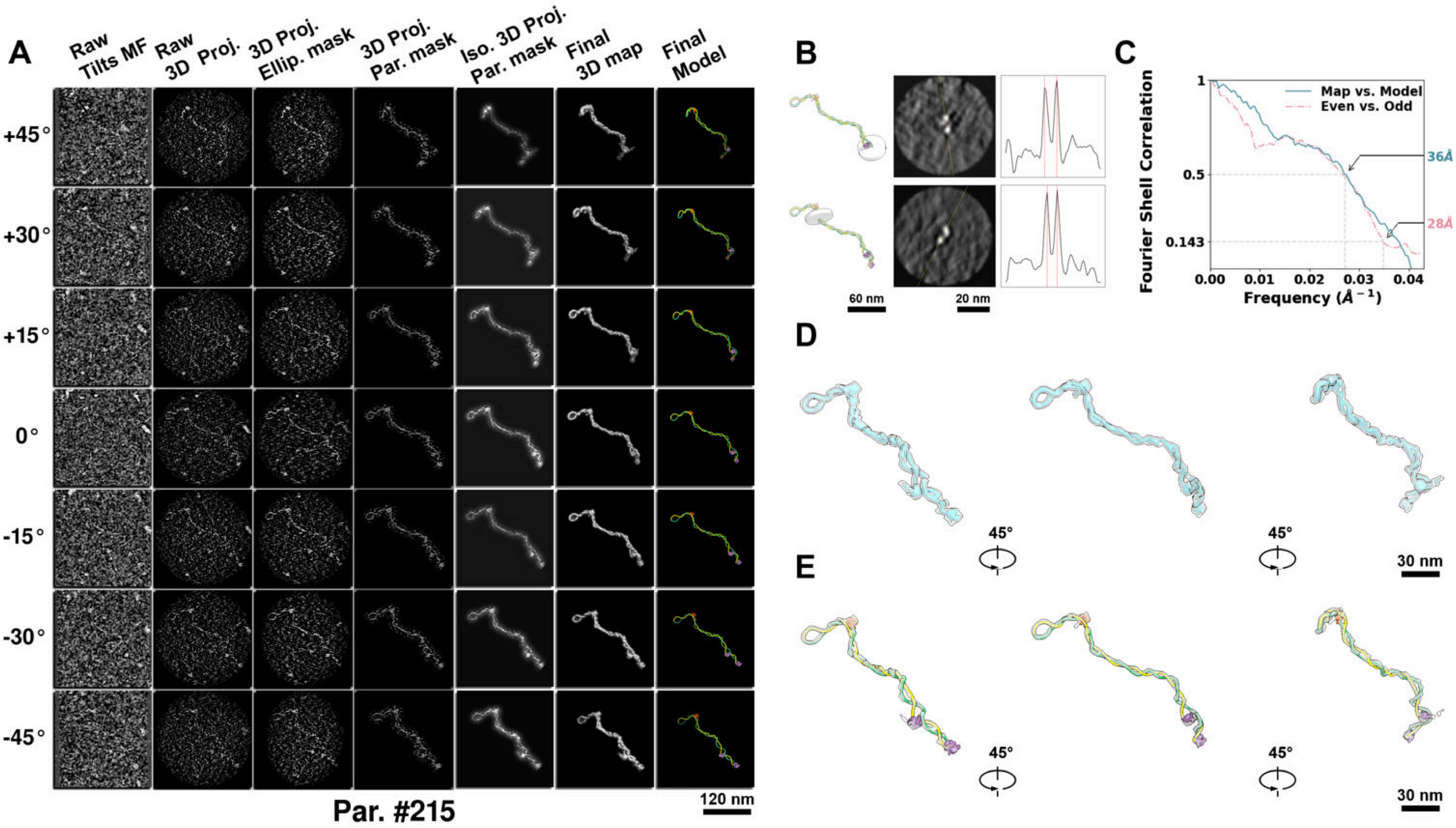

**Supplementary Particle Figure 215. Cryo-ET 3D reconstruction of an individual TEC-Cas particle.**
(A) 3D reconstruction of the plasmid particle (index no. 215). The first column shows seven representative tilt images from +45° to -45° in step of 15°. The second, third, and fourth columns show 3D projections of the particle with spherical, ellipsoidal (thinner along the z-dimension), and particle-shaped masks, respectively. The fifth column displays the 3D projections of the enhanced and IsoNet missing-wedge-corrected particle. The sixth and seventh columns present the final 3D map and the flexibly fitted model, respectively. (B) Two cross-sectional views (12 nm thickness) of the plasmid density map along its plectoneme axis are shown in the left-middle panel. The intensity profile along the line crossing the two high-density DNA spots is displayed in the right panel. (C) Resolution assessment of the final 3D map using Fourier shell correlation (FSC). Two criteria are shown: FSC between two half-maps reconstructed from even and odd frames (evaluated at 0.143) and FSC between the final 3D map and the fitted model (evaluated at 0.5). (D) Zoomed-in views of the final 3D density map from panel A, displayed at two contour levels. (E) Superimposition of the high-contour level map from panel D onto its fitted model.

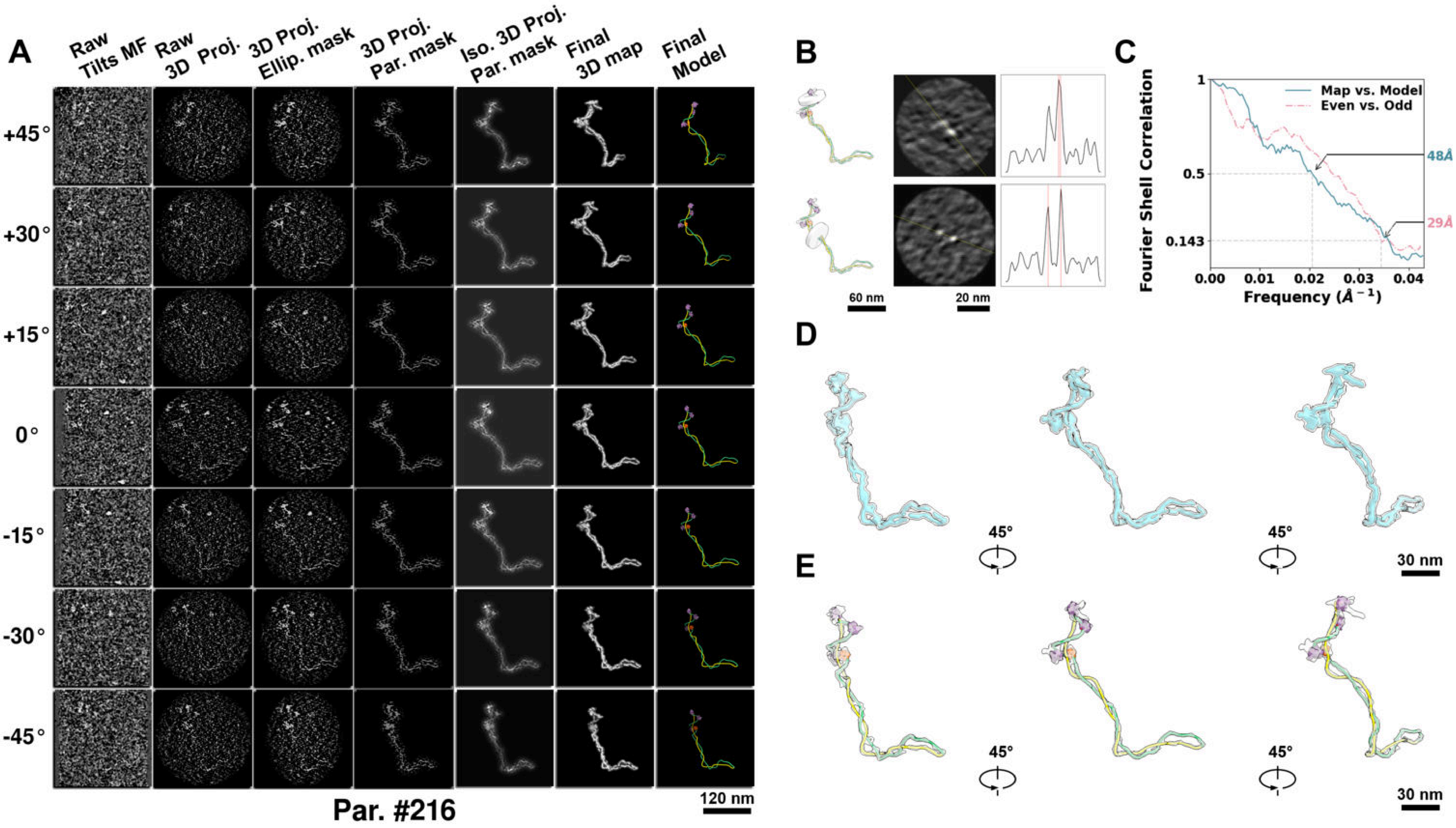

**Supplementary Particle Figure 216. Cryo-ET 3D reconstruction of an individual TEC-Cas particle.**
(A) 3D reconstruction of the plasmid particle (index no. 216). The first column shows seven representative tilt images from +45° to -45° in step of 15°. The second, third, and fourth columns show 3D projections of the particle with spherical, ellipsoidal (thinner along the z-dimension), and particle-shaped masks, respectively. The fifth column displays the 3D projections of the enhanced and IsoNet missing-wedge-corrected particle. The sixth and seventh columns present the final 3D map and the flexibly fitted model, respectively. (B) Two cross-sectional views (12 nm thickness) of the plasmid density map along its plectoneme axis are shown in the left-middle panel. The intensity profile along the line crossing the two high-density DNA spots is displayed in the right panel. (C) Resolution assessment of the final 3D map using Fourier shell correlation (FSC). Two criteria are shown: FSC between two half-maps reconstructed from even and odd frames (evaluated at 0.143) and FSC between the final 3D map and the fitted model (evaluated at 0.5). (D) Zoomed-in views of the final 3D density map from panel A, displayed at two contour levels. (E) Superimposition of the high-contour level map from panel D onto its fitted model.

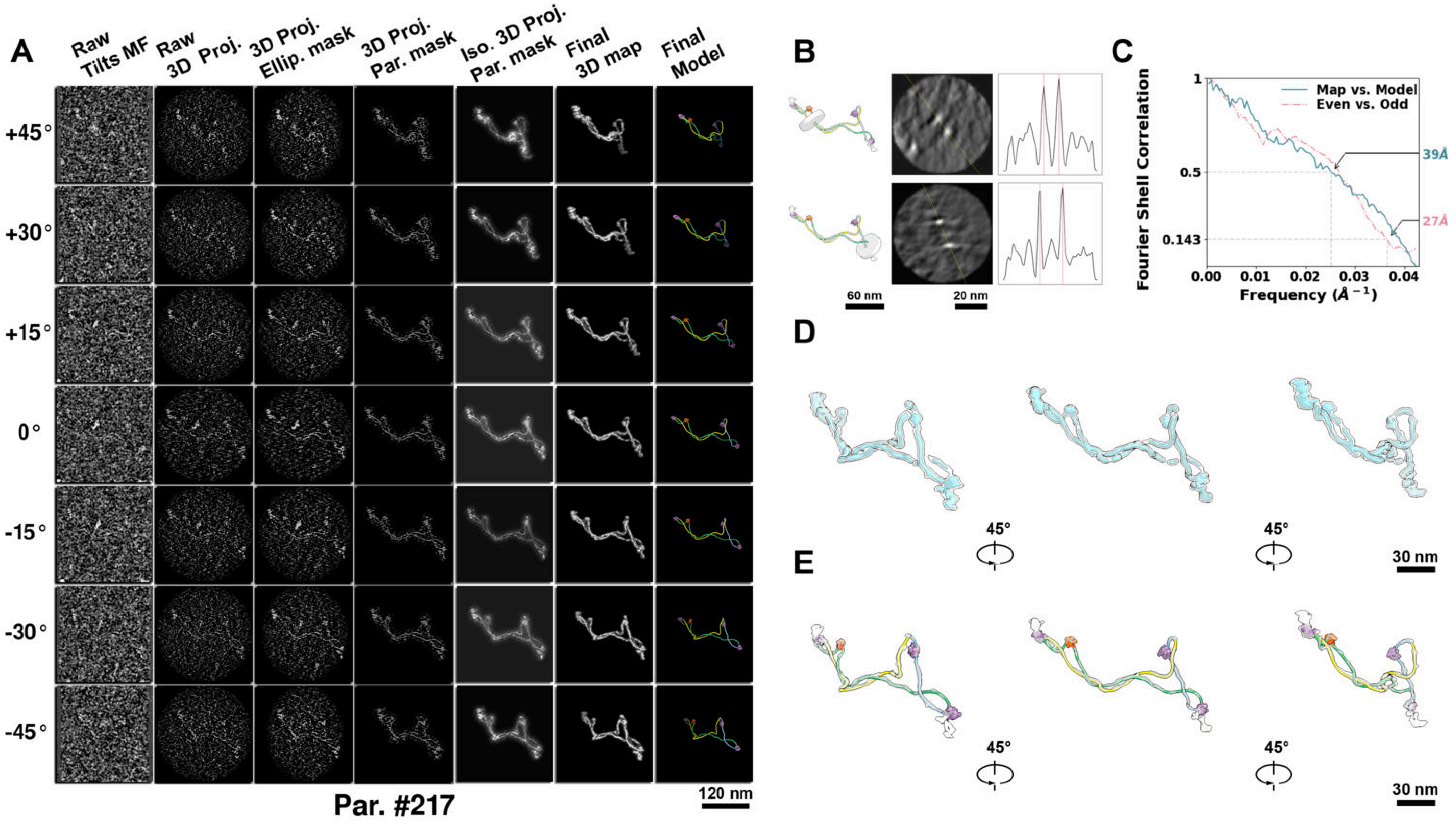

**Supplementary Particle Figure 217. Cryo-ET 3D reconstruction of an individual TEC-Cas particle.**
(A) 3D reconstruction of the plasmid particle (index no. 217). The first column shows seven representative tilt images from +45° to -45° in step of 15°. The second, third, and fourth columns show 3D projections of the particle with spherical, ellipsoidal (thinner along the z-dimension), and particle-shaped masks, respectively. The fifth column displays the 3D projections of the enhanced and IsoNet missing-wedge-corrected particle. The sixth and seventh columns present the final 3D map and the flexibly fitted model, respectively. (B) Two cross-sectional views (12 nm thickness) of the plasmid density map along its plectoneme axis are shown in the left-middle panel. The intensity profile along the line crossing the two high-density DNA spots is displayed in the right panel. (C) Resolution assessment of the final 3D map using Fourier shell correlation (FSC). Two criteria are shown: FSC between two half-maps reconstructed from even and odd frames (evaluated at 0.143) and FSC between the final 3D map and the fitted model (evaluated at 0.5). (D) Zoomed-in views of the final 3D density map from panel A, displayed at two contour levels. (E) Superimposition of the high-contour level map from panel D onto its fitted model.

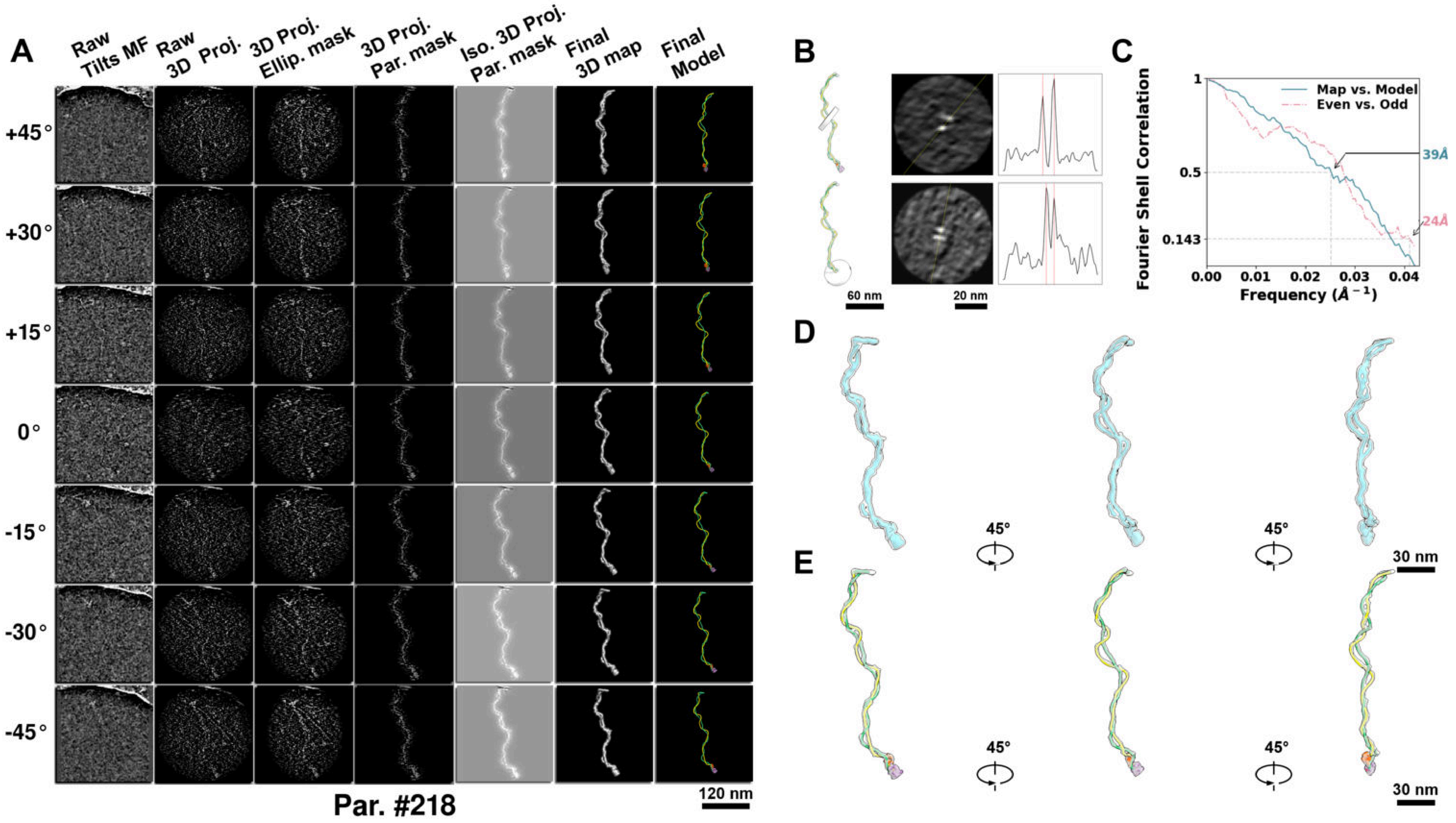

**Supplementary Particle Figure 218. Cryo-ET 3D reconstruction of an individual TEC-Cas particle.**
(A) 3D reconstruction of the plasmid particle (index no. 218). The first column shows seven representative tilt images from +45° to -45° in step of 15°. The second, third, and fourth columns show 3D projections of the particle with spherical, ellipsoidal (thinner along the z-dimension), and particle-shaped masks, respectively. The fifth column displays the 3D projections of the enhanced and IsoNet missing-wedge-corrected particle. The sixth and seventh columns present the final 3D map and the flexibly fitted model, respectively. (B) Two cross-sectional views (12 nm thickness) of the plasmid density map along its plectoneme axis are shown in the left-middle panel. The intensity profile along the line crossing the two high-density DNA spots is displayed in the right panel. (C) Resolution assessment of the final 3D map using Fourier shell correlation (FSC). Two criteria are shown: FSC between two half-maps reconstructed from even and odd frames (evaluated at 0.143) and FSC between the final 3D map and the fitted model (evaluated at 0.5). (D) Zoomed-in views of the final 3D density map from panel A, displayed at two contour levels. (E) Superimposition of the high-contour level map from panel D onto its fitted model.

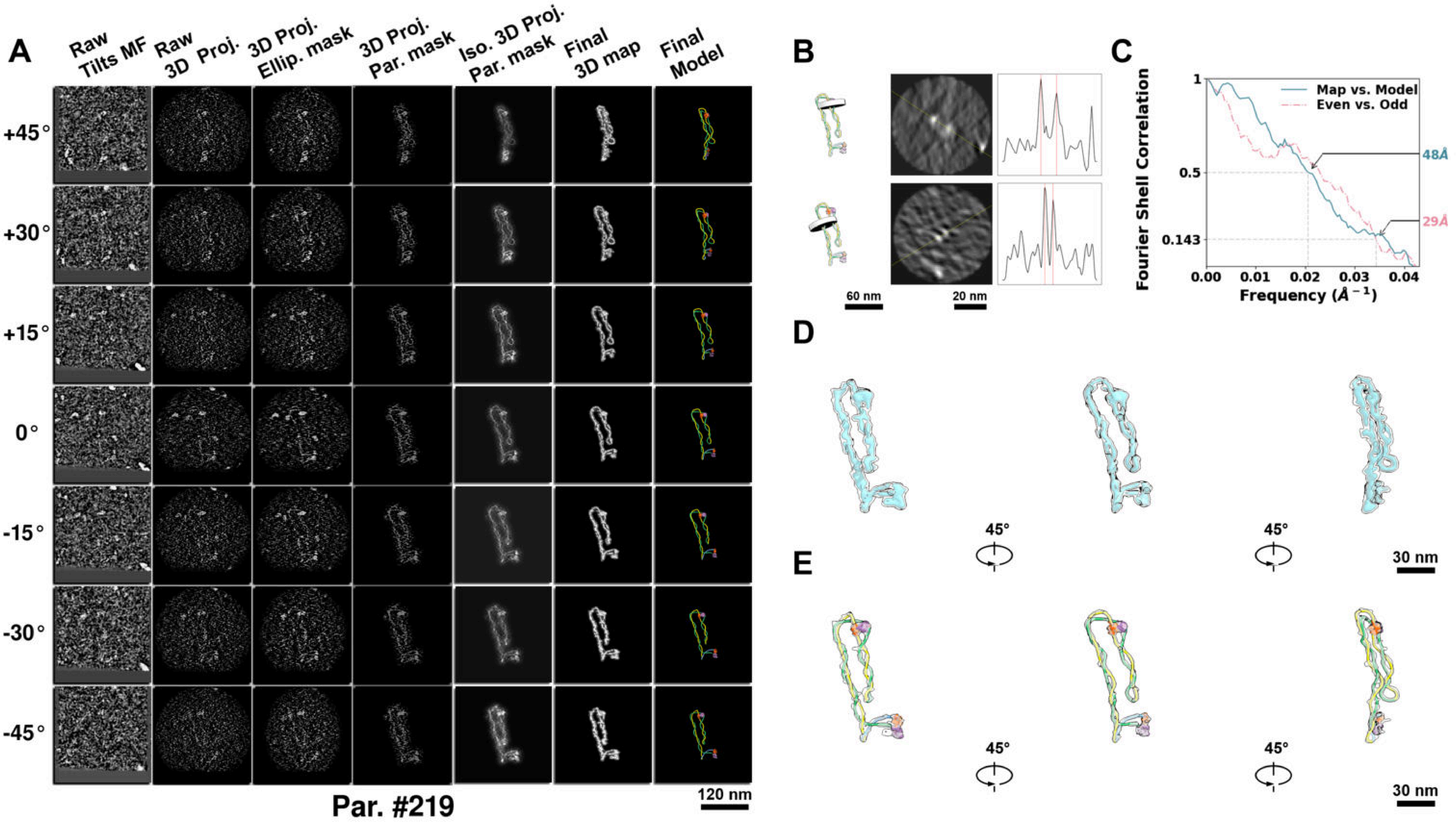

**Supplementary Particle Figure 219. Cryo-ET 3D reconstruction of an individual TEC-Cas particle.**
(A) 3D reconstruction of the plasmid particle (index no. 219). The first column shows seven representative tilt images from +45° to -45° in step of 15°. The second, third, and fourth columns show 3D projections of the particle with spherical, ellipsoidal (thinner along the z-dimension), and particle-shaped masks, respectively. The fifth column displays the 3D projections of the enhanced and IsoNet missing-wedge-corrected particle. The sixth and seventh columns present the final 3D map and the flexibly fitted model, respectively. (B) Two cross-sectional views (12 nm thickness) of the plasmid density map along its plectoneme axis are shown in the left-middle panel. The intensity profile along the line crossing the two high-density DNA spots is displayed in the right panel. (C) Resolution assessment of the final 3D map using Fourier shell correlation (FSC). Two criteria are shown: FSC between two half-maps reconstructed from even and odd frames (evaluated at 0.143) and FSC between the final 3D map and the fitted model (evaluated at 0.5). (D) Zoomed-in views of the final 3D density map from panel A, displayed at two contour levels. (E) Superimposition of the high-contour level map from panel D onto its fitted model.

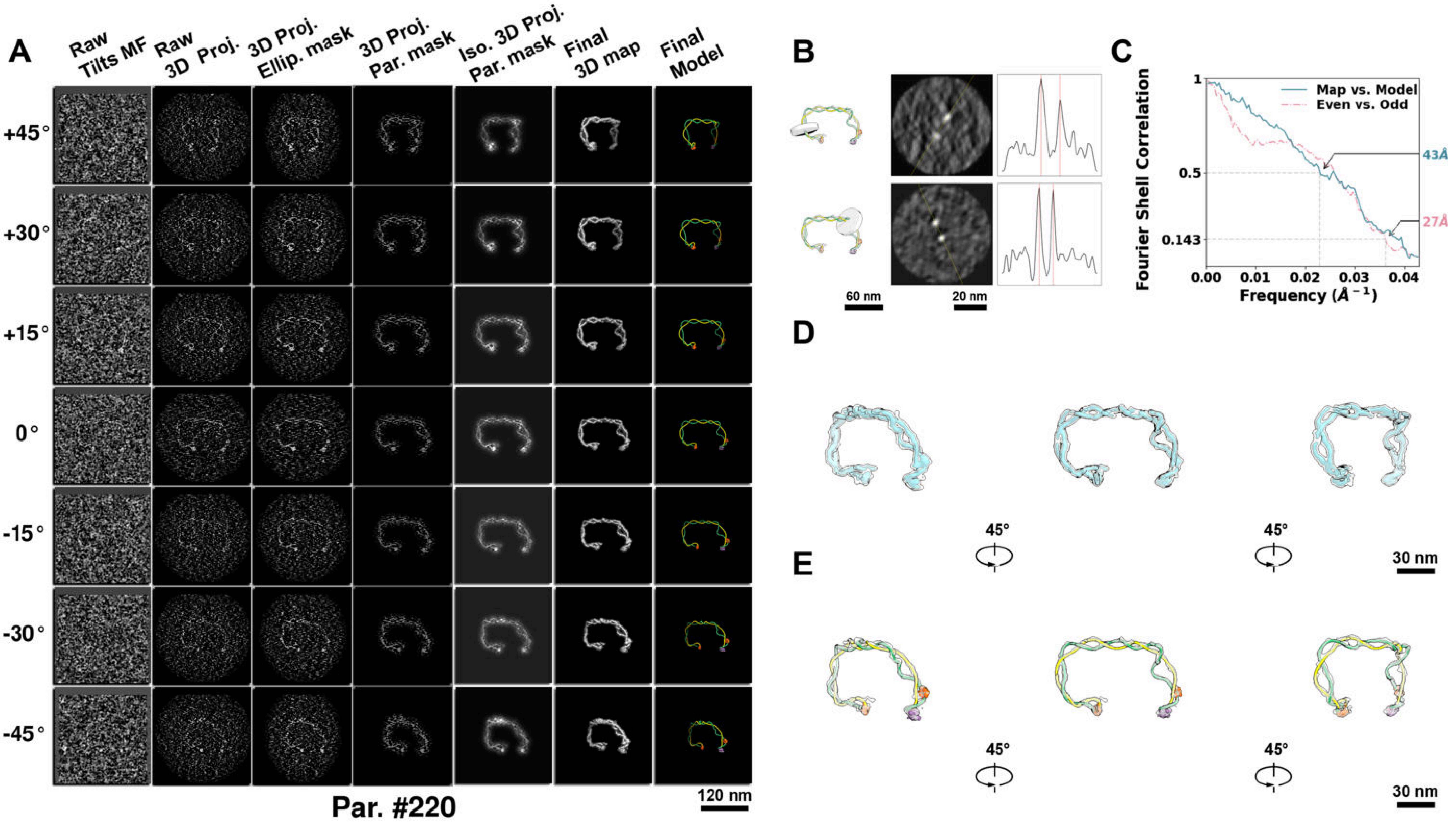

**Supplementary Particle Figure 220. Cryo-ET 3D reconstruction of an individual TEC-Cas particle.**
(A) 3D reconstruction of the plasmid particle (index no. 220). The first column shows seven representative tilt images from +45° to -45° in step of 15°. The second, third, and fourth columns show 3D projections of the particle with spherical, ellipsoidal (thinner along the z-dimension), and particle-shaped masks, respectively. The fifth column displays the 3D projections of the enhanced and IsoNet missing-wedge-corrected particle. The sixth and seventh columns present the final 3D map and the flexibly fitted model, respectively. (B) Two cross-sectional views (12 nm thickness) of the plasmid density map along its plectoneme axis are shown in the left-middle panel. The intensity profile along the line crossing the two high-density DNA spots is displayed in the right panel. (C) Resolution assessment of the final 3D map using Fourier shell correlation (FSC). Two criteria are shown: FSC between two half-maps reconstructed from even and odd frames (evaluated at 0.143) and FSC between the final 3D map and the fitted model (evaluated at 0.5). (D) Zoomed-in views of the final 3D density map from panel A, displayed at two contour levels. (E) Superimposition of the high-contour level map from panel D onto its fitted model.

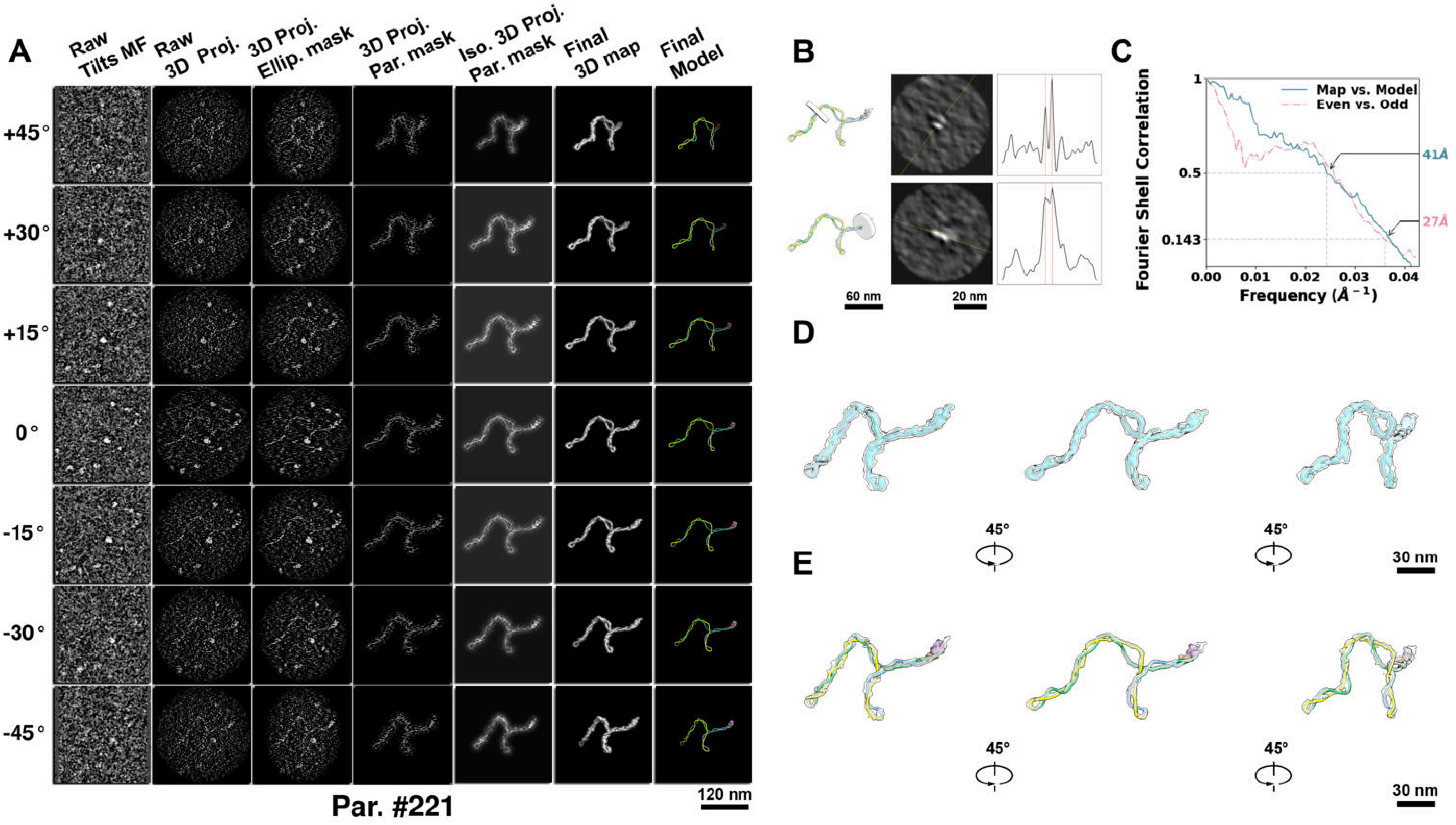

**Supplementary Particle Figure 221. Cryo-ET 3D reconstruction of an individual TEC-Cas particle.**
(A) 3D reconstruction of the plasmid particle (index no. 221). The first column shows seven representative tilt images from +45° to -45° in step of 15°. The second, third, and fourth columns show 3D projections of the particle with spherical, ellipsoidal (thinner along the z-dimension), and particle-shaped masks, respectively. The fifth column displays the 3D projections of the enhanced and IsoNet missing-wedge-corrected particle. The sixth and seventh columns present the final 3D map and the flexibly fitted model, respectively. (B) Two cross-sectional views (12 nm thickness) of the plasmid density map along its plectoneme axis are shown in the left-middle panel. The intensity profile along the line crossing the two high-density DNA spots is displayed in the right panel. (C) Resolution assessment of the final 3D map using Fourier shell correlation (FSC). Two criteria are shown: FSC between two half-maps reconstructed from even and odd frames (evaluated at 0.143) and FSC between the final 3D map and the fitted model (evaluated at 0.5). (D) Zoomed-in views of the final 3D density map from panel A, displayed at two contour levels. (E) Superimposition of the high-contour level map from panel D onto its fitted model.

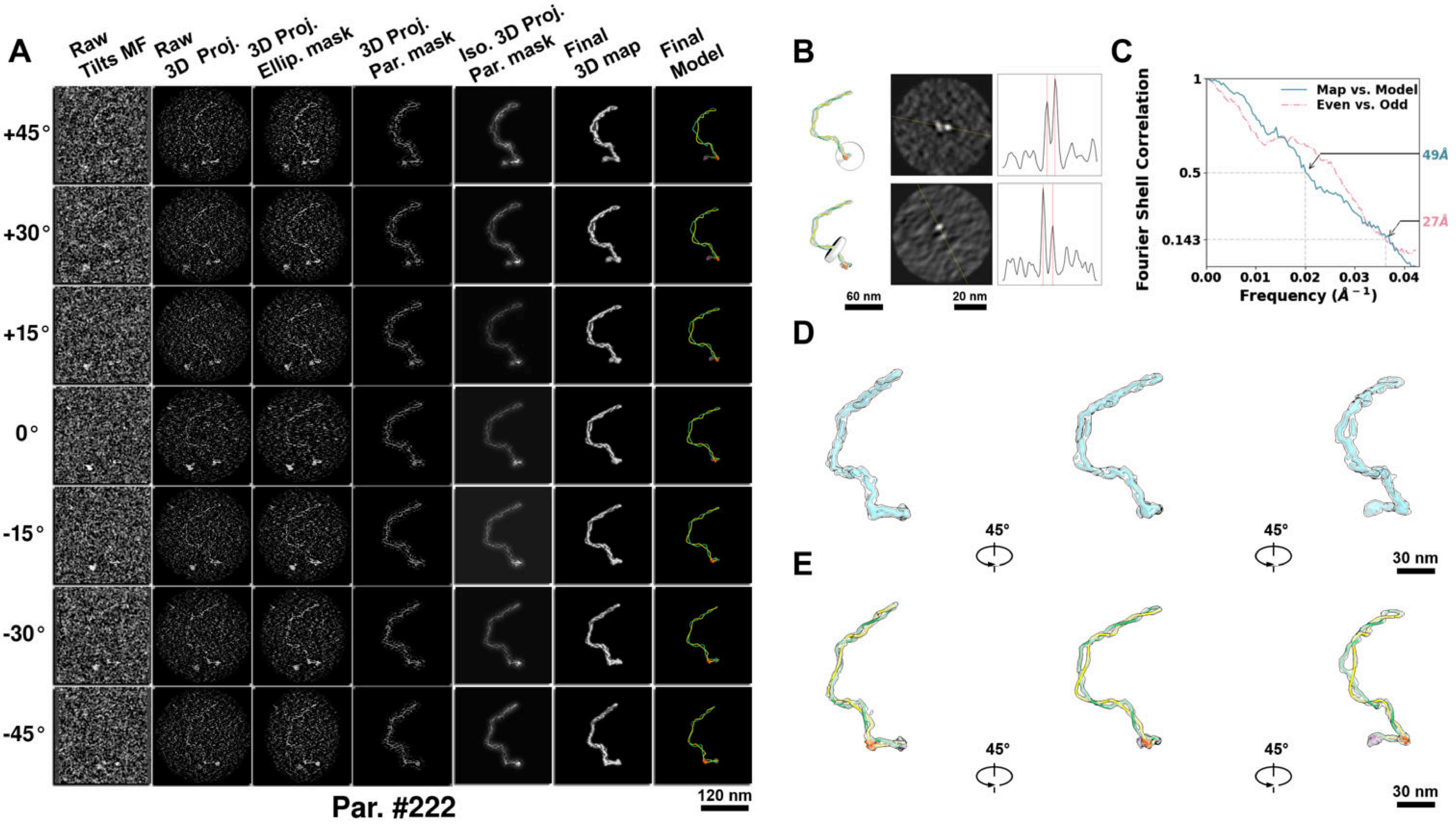

**Supplementary Particle Figure 222. Cryo-ET 3D reconstruction of an individual TEC-Cas particle.**
(A) 3D reconstruction of the plasmid particle (index no. 222). The first column shows seven representative tilt images from +45° to -45° in step of 15°. The second, third, and fourth columns show 3D projections of the particle with spherical, ellipsoidal (thinner along the z-dimension), and particle-shaped masks, respectively. The fifth column displays the 3D projections of the enhanced and IsoNet missing-wedge-corrected particle. The sixth and seventh columns present the final 3D map and the flexibly fitted model, respectively. (B) Two cross-sectional views (12 nm thickness) of the plasmid density map along its plectoneme axis are shown in the left-middle panel. The intensity profile along the line crossing the two high-density DNA spots is displayed in the right panel. (C) Resolution assessment of the final 3D map using Fourier shell correlation (FSC). Two criteria are shown: FSC between two half-maps reconstructed from even and odd frames (evaluated at 0.143) and FSC between the final 3D map and the fitted model (evaluated at 0.5). (D) Zoomed-in views of the final 3D density map from panel A, displayed at two contour levels. (E) Superimposition of the high-contour level map from panel D onto its fitted model.

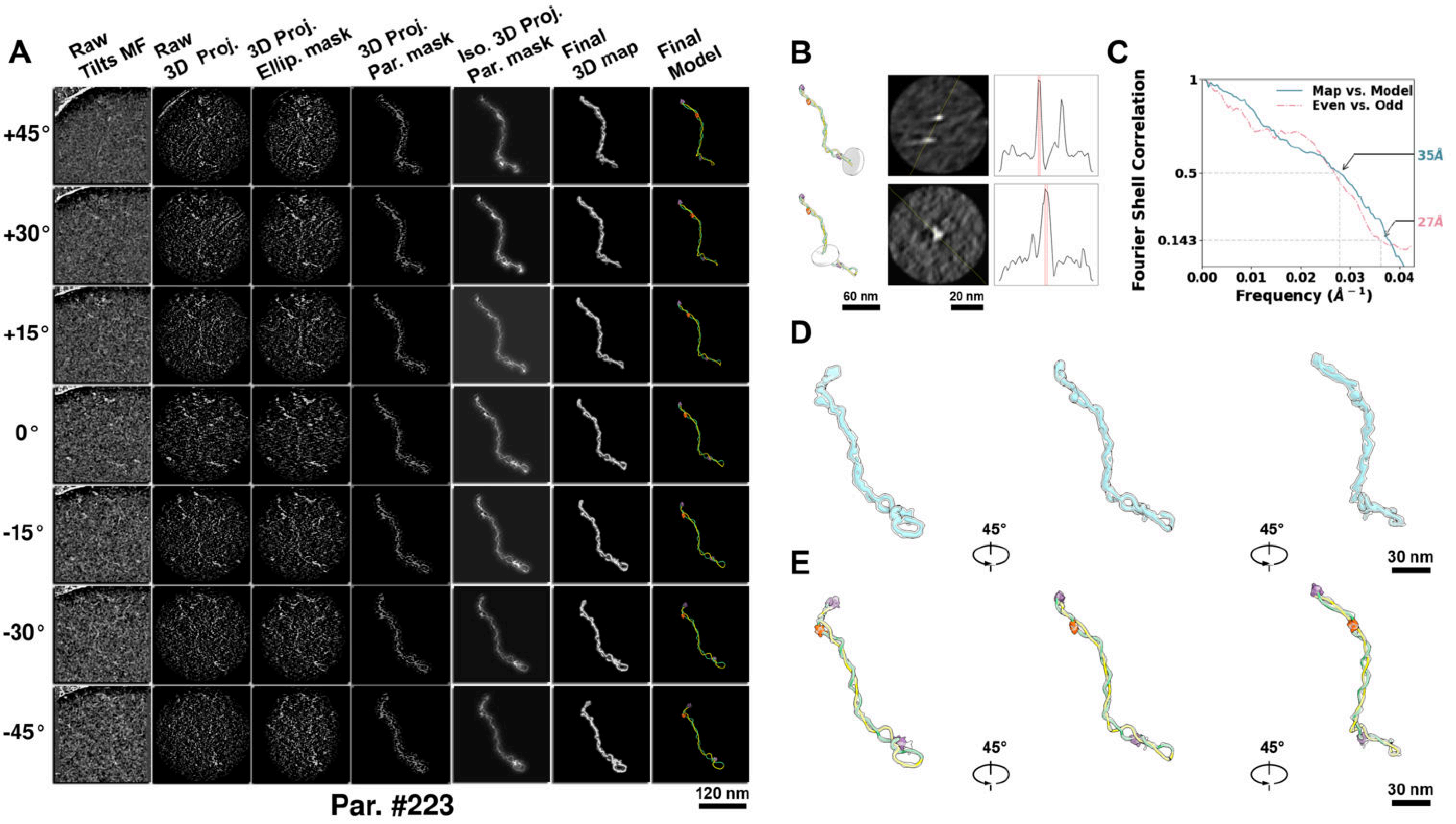

**Supplementary Particle Figure 223. Cryo-ET 3D reconstruction of an individual TEC-Cas particle.**
(A) 3D reconstruction of the plasmid particle (index no. 223). The first column shows seven representative tilt images from +45° to -45° in step of 15°. The second, third, and fourth columns show 3D projections of the particle with spherical, ellipsoidal (thinner along the z-dimension), and particle-shaped masks, respectively. The fifth column displays the 3D projections of the enhanced and IsoNet missing-wedge-corrected particle. The sixth and seventh columns present the final 3D map and the flexibly fitted model, respectively. (B) Two cross-sectional views (12 nm thickness) of the plasmid density map along its plectoneme axis are shown in the left-middle panel. The intensity profile along the line crossing the two high-density DNA spots is displayed in the right panel. (C) Resolution assessment of the final 3D map using Fourier shell correlation (FSC). Two criteria are shown: FSC between two half-maps reconstructed from even and odd frames (evaluated at 0.143) and FSC between the final 3D map and the fitted model (evaluated at 0.5). (D) Zoomed-in views of the final 3D density map from panel A, displayed at two contour levels. (E) Superimposition of the high-contour level map from panel D onto its fitted model.

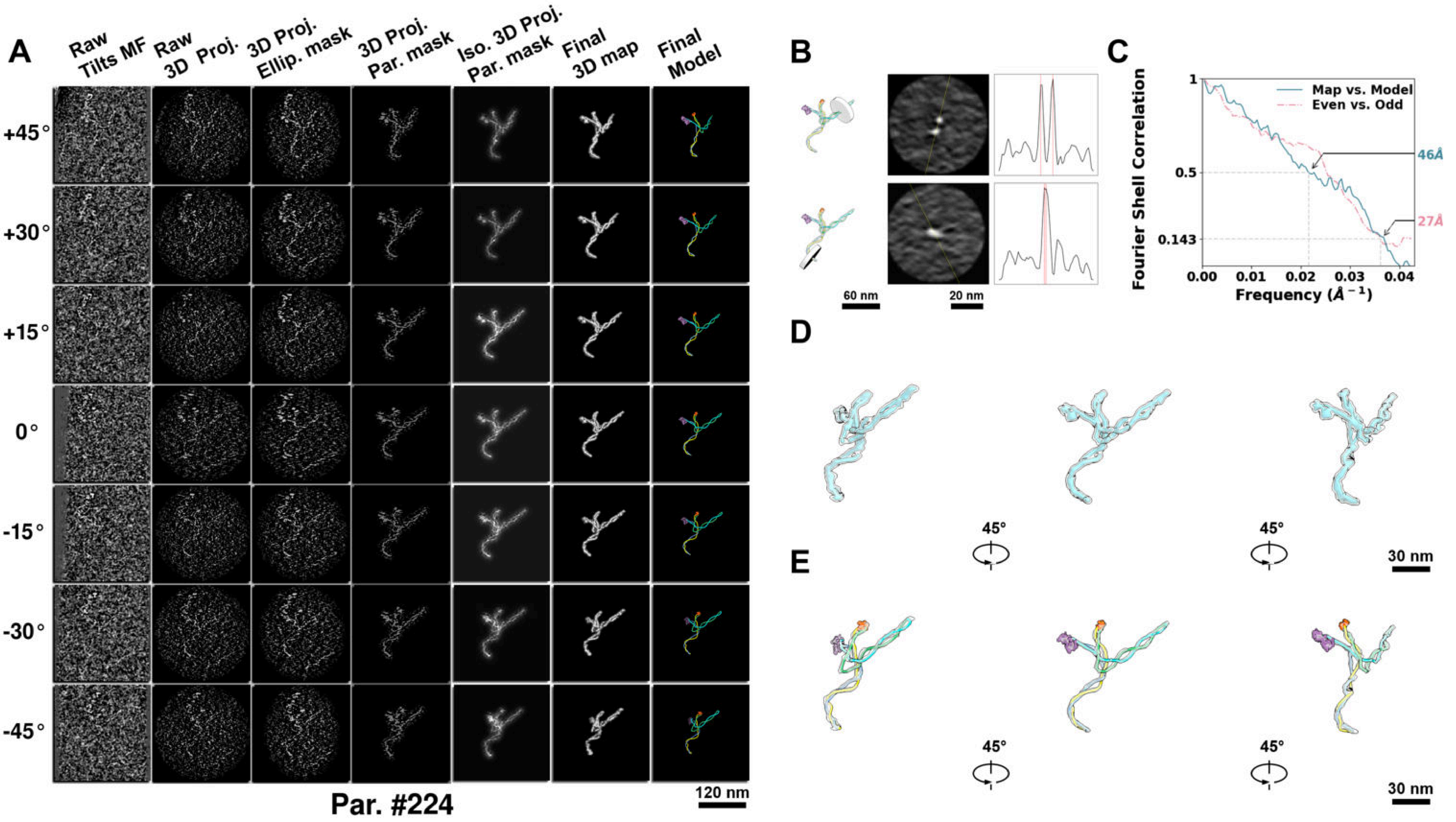

**Supplementary Particle Figure 224. Cryo-ET 3D reconstruction of an individual TEC-Cas particle.**
(A) 3D reconstruction of the plasmid particle (index no. 224). The first column shows seven representative tilt images from +45° to -45° in step of 15°. The second, third, and fourth columns show 3D projections of the particle with spherical, ellipsoidal (thinner along the z-dimension), and particle-shaped masks, respectively. The fifth column displays the 3D projections of the enhanced and IsoNet missing-wedge-corrected particle. The sixth and seventh columns present the final 3D map and the flexibly fitted model, respectively. (B) Two cross-sectional views (12 nm thickness) of the plasmid density map along its plectoneme axis are shown in the left-middle panel. The intensity profile along the line crossing the two high-density DNA spots is displayed in the right panel. (C) Resolution assessment of the final 3D map using Fourier shell correlation (FSC). Two criteria are shown: FSC between two half-maps reconstructed from even and odd frames (evaluated at 0.143) and FSC between the final 3D map and the fitted model (evaluated at 0.5). (D) Zoomed-in views of the final 3D density map from panel A, displayed at two contour levels. (E) Superimposition of the high-contour level map from panel D onto its fitted model.

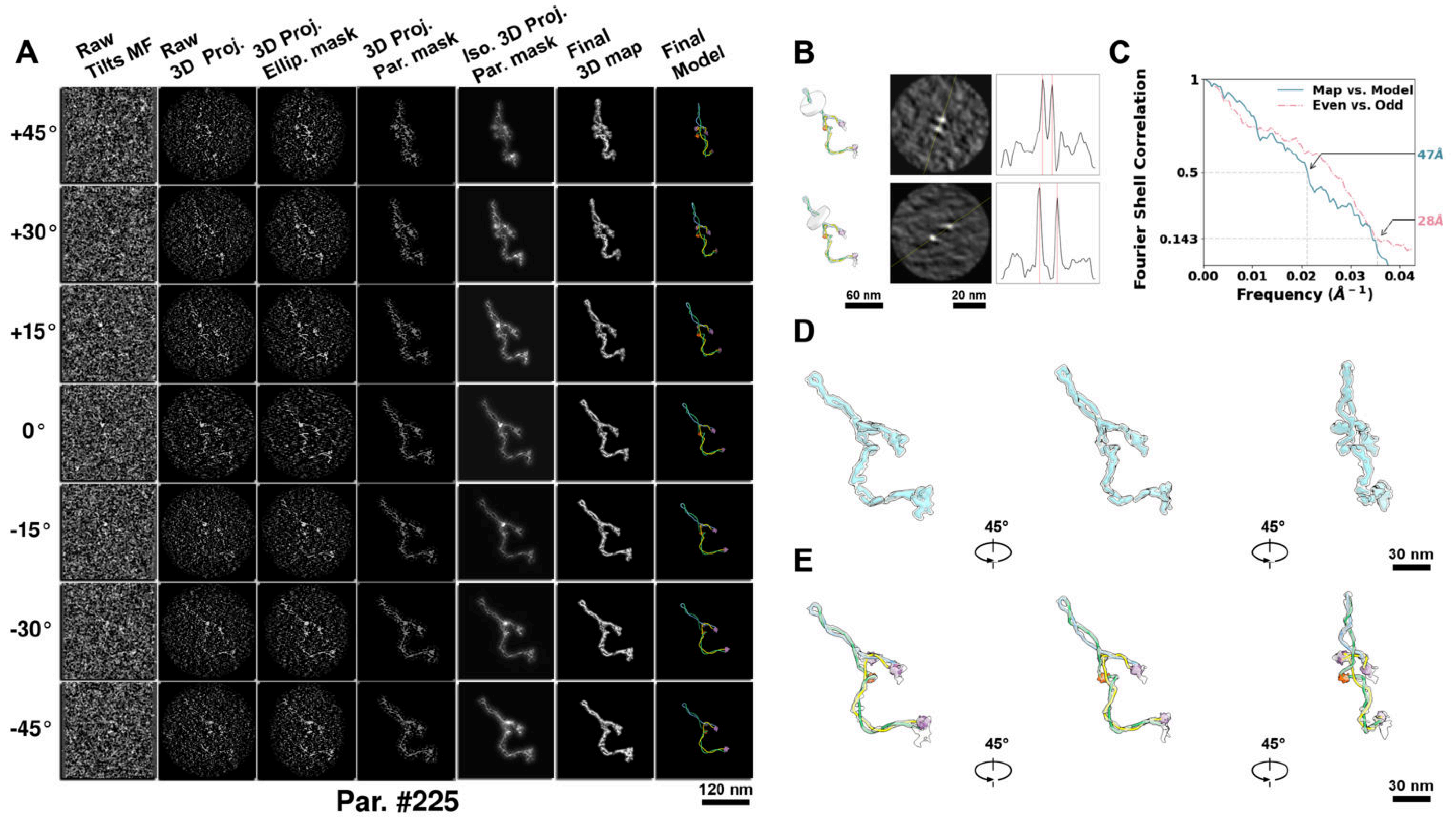

**Supplementary Particle Figure 225. Cryo-ET 3D reconstruction of an individual TEC-Cas particle.**
(A) 3D reconstruction of the plasmid particle (index no. 225). The first column shows seven representative tilt images from +45° to -45° in step of 15°. The second, third, and fourth columns show 3D projections of the particle with spherical, ellipsoidal (thinner along the z-dimension), and particle-shaped masks, respectively. The fifth column displays the 3D projections of the enhanced and IsoNet missing-wedge-corrected particle. The sixth and seventh columns present the final 3D map and the flexibly fitted model, respectively. (B) Two cross-sectional views (12 nm thickness) of the plasmid density map along its plectoneme axis are shown in the left-middle panel. The intensity profile along the line crossing the two high-density DNA spots is displayed in the right panel. (C) Resolution assessment of the final 3D map using Fourier shell correlation (FSC). Two criteria are shown: FSC between two half-maps reconstructed from even and odd frames (evaluated at 0.143) and FSC between the final 3D map and the fitted model (evaluated at 0.5). (D) Zoomed-in views of the final 3D density map from panel A, displayed at two contour levels. (E) Superimposition of the high-contour level map from panel D onto its fitted model.

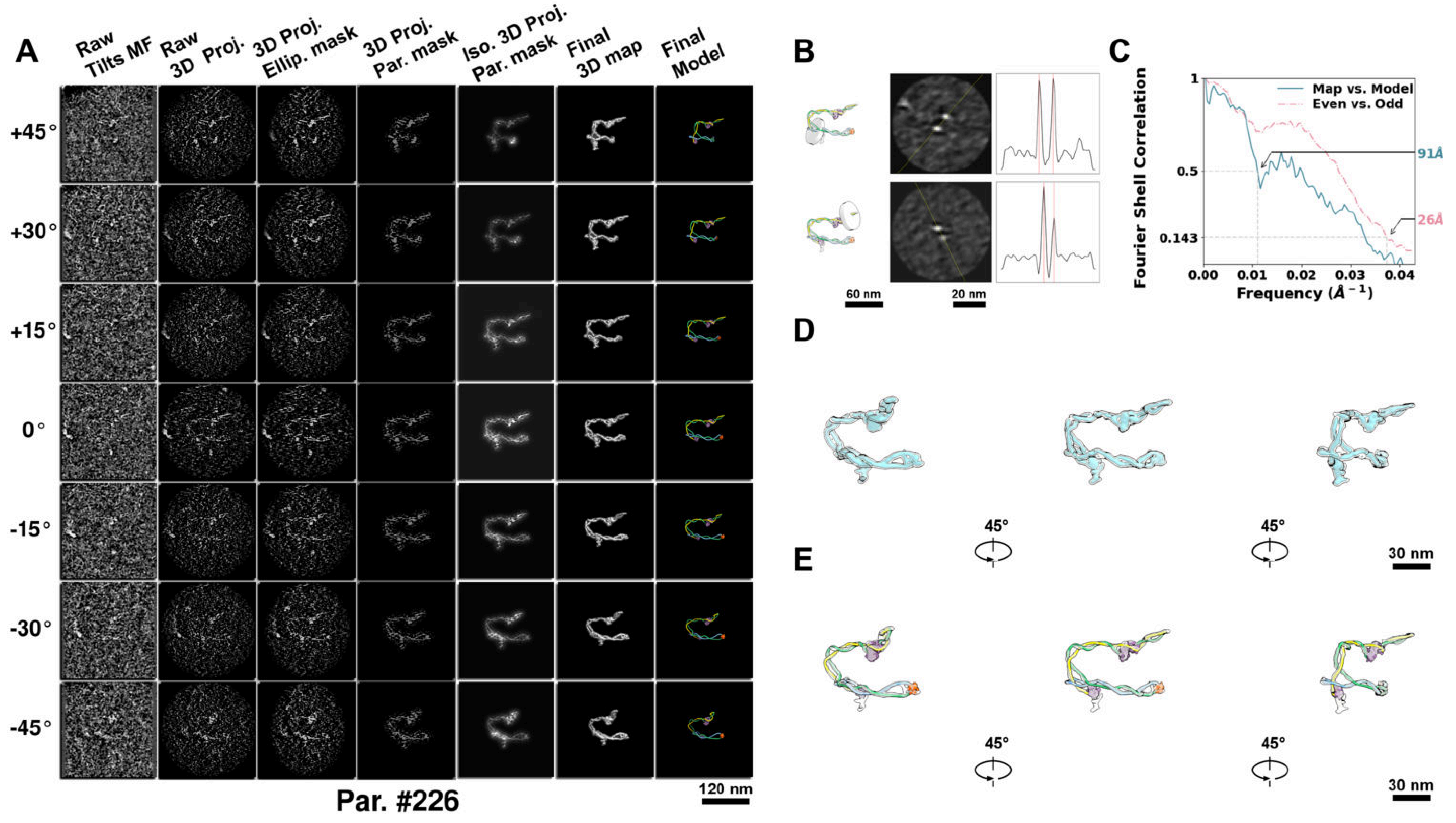

**Supplementary Particle Figure 226. Cryo-ET 3D reconstruction of an individual TEC-Cas particle.**
(A) 3D reconstruction of the plasmid particle (index no. 226). The first column shows seven representative tilt images from +45° to -45° in step of 15°. The second, third, and fourth columns show 3D projections of the particle with spherical, ellipsoidal (thinner along the z-dimension), and particle-shaped masks, respectively. The fifth column displays the 3D projections of the enhanced and IsoNet missing-wedge-corrected particle. The sixth and seventh columns present the final 3D map and the flexibly fitted model, respectively. (B) Two cross-sectional views (12 nm thickness) of the plasmid density map along its plectoneme axis are shown in the left-middle panel. The intensity profile along the line crossing the two high-density DNA spots is displayed in the right panel. (C) Resolution assessment of the final 3D map using Fourier shell correlation (FSC). Two criteria are shown: FSC between two half-maps reconstructed from even and odd frames (evaluated at 0.143) and FSC between the final 3D map and the fitted model (evaluated at 0.5). (D) Zoomed-in views of the final 3D density map from panel A, displayed at two contour levels. (E) Superimposition of the high-contour level map from panel D onto its fitted model.

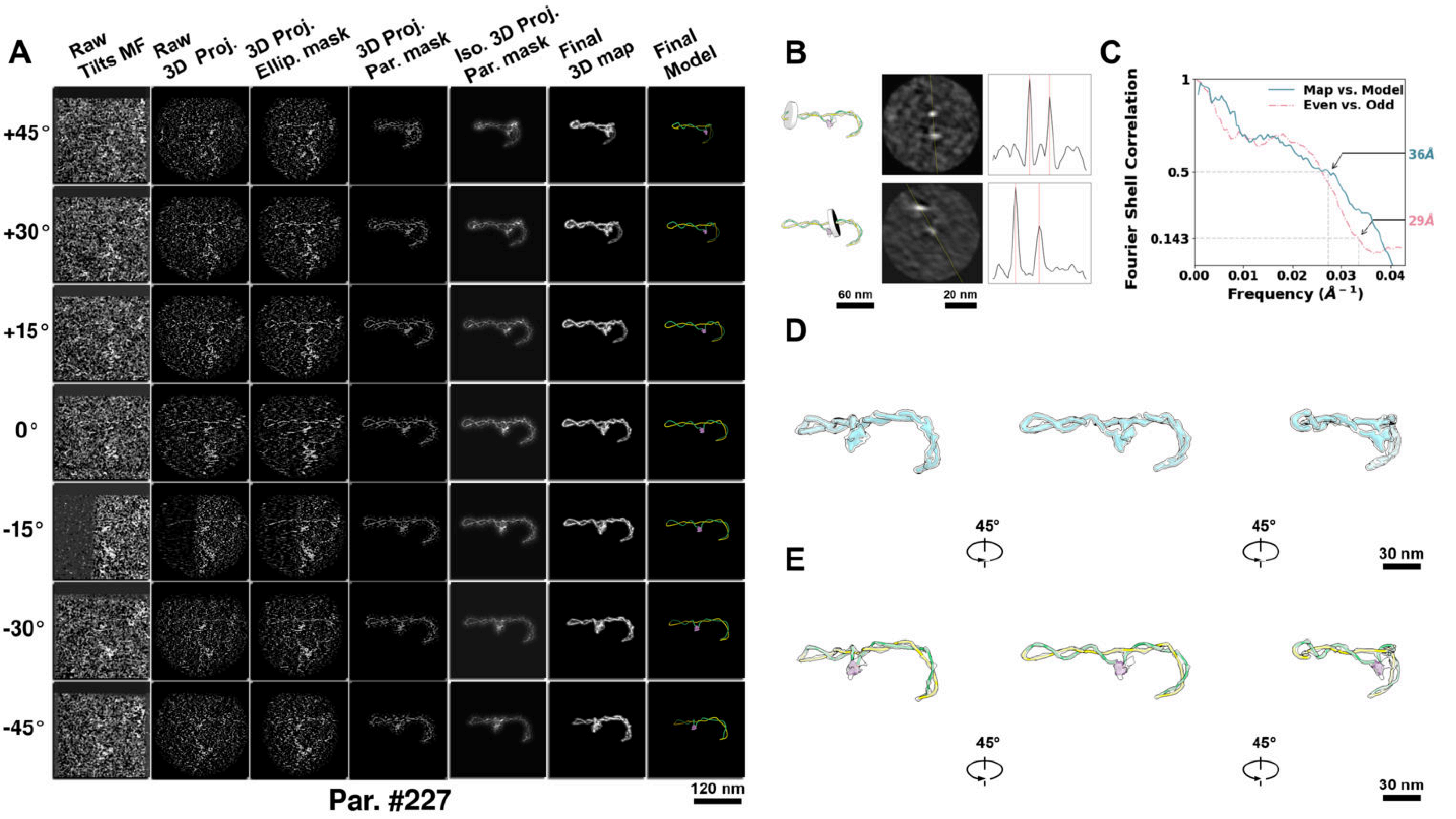

**Supplementary Particle Figure 227. Cryo-ET 3D reconstruction of an individual TEC-TopI particle.**
(A) 3D reconstruction of the plasmid particle (index no. 227). The first column shows seven representative tilt images from +45° to -45° in step of 15°. The second, third, and fourth columns show 3D projections of the particle with spherical, ellipsoidal (thinner along the z-dimension), and particle-shaped masks, respectively. The fifth column displays the 3D projections of the enhanced and IsoNet missing-wedge-corrected particle. The sixth and seventh columns present the final 3D map and the flexibly fitted model, respectively. (B) Two cross-sectional views (12 nm thickness) of the plasmid density map along its plectoneme axis are shown in the left-middle panel. The intensity profile along the line crossing the two high-density DNA spots is displayed in the right panel. (C) Resolution assessment of the final 3D map using Fourier shell correlation (FSC). Two criteria are shown: FSC between two half-maps reconstructed from even and odd frames (evaluated at 0.143) and FSC between the final 3D map and the fitted model (evaluated at 0.5). (D) Zoomed-in views of the final 3D density map from panel A, displayed at two contour levels. (E) Superimposition of the high-contour level map from panel D onto its fitted model.

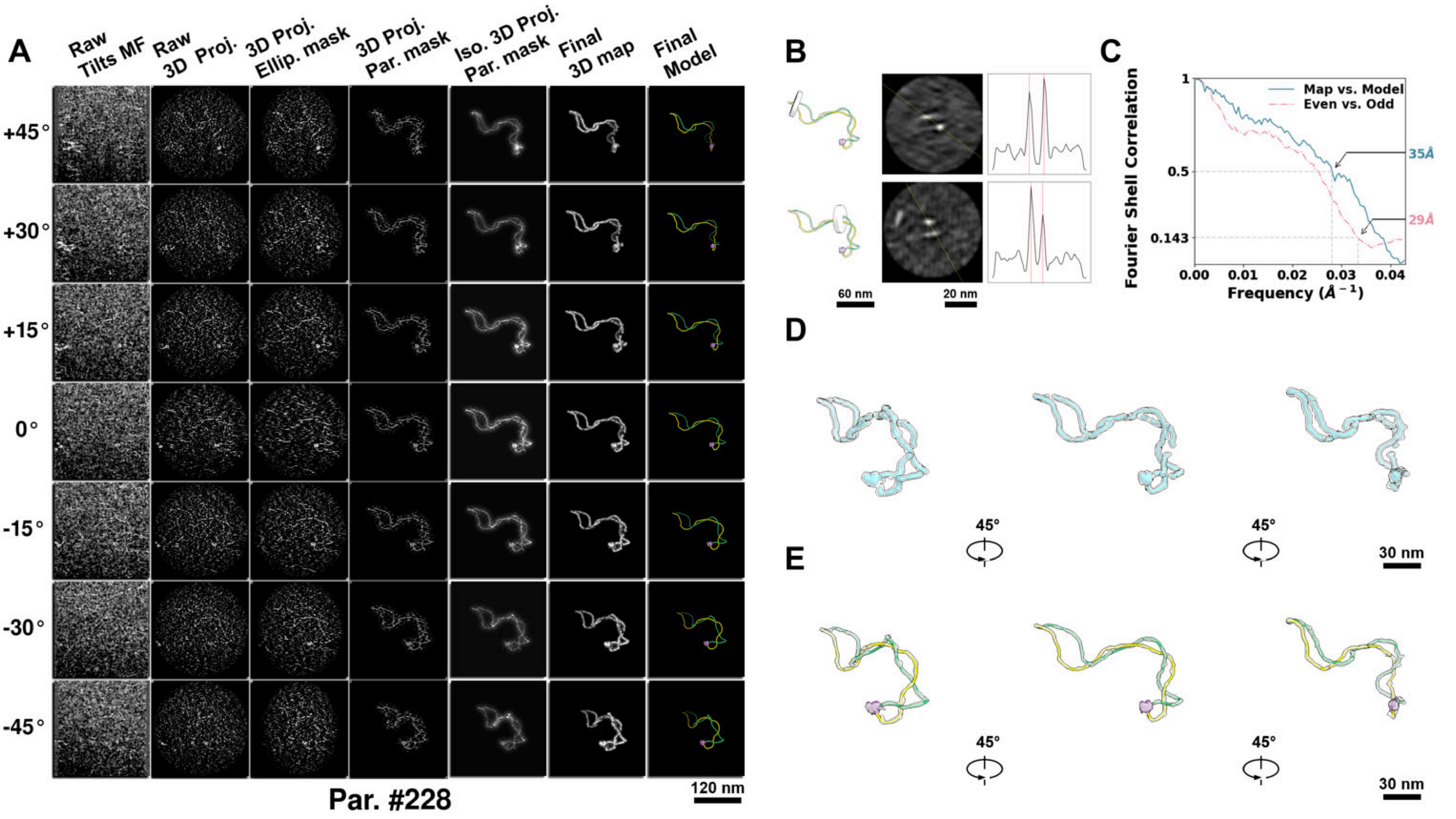

**Supplementary Particle Figure 228. Cryo-ET 3D reconstruction of an individual TEC-TopI particle.**
(A) 3D reconstruction of the plasmid particle (index no. 228). The first column shows seven representative tilt images from +45° to -45° in step of 15°. The second, third, and fourth columns show 3D projections of the particle with spherical, ellipsoidal (thinner along the z-dimension), and particle-shaped masks, respectively. The fifth column displays the 3D projections of the enhanced and IsoNet missing-wedge-corrected particle. The sixth and seventh columns present the final 3D map and the flexibly fitted model, respectively. (B) Two cross-sectional views (12 nm thickness) of the plasmid density map along its plectoneme axis are shown in the left-middle panel. The intensity profile along the line crossing the two high-density DNA spots is displayed in the right panel. (C) Resolution assessment of the final 3D map using Fourier shell correlation (FSC). Two criteria are shown: FSC between two half-maps reconstructed from even and odd frames (evaluated at 0.143) and FSC between the final 3D map and the fitted model (evaluated at 0.5). (D) Zoomed-in views of the final 3D density map from panel A, displayed at two contour levels. (E) Superimposition of the high-contour level map from panel D onto its fitted model.

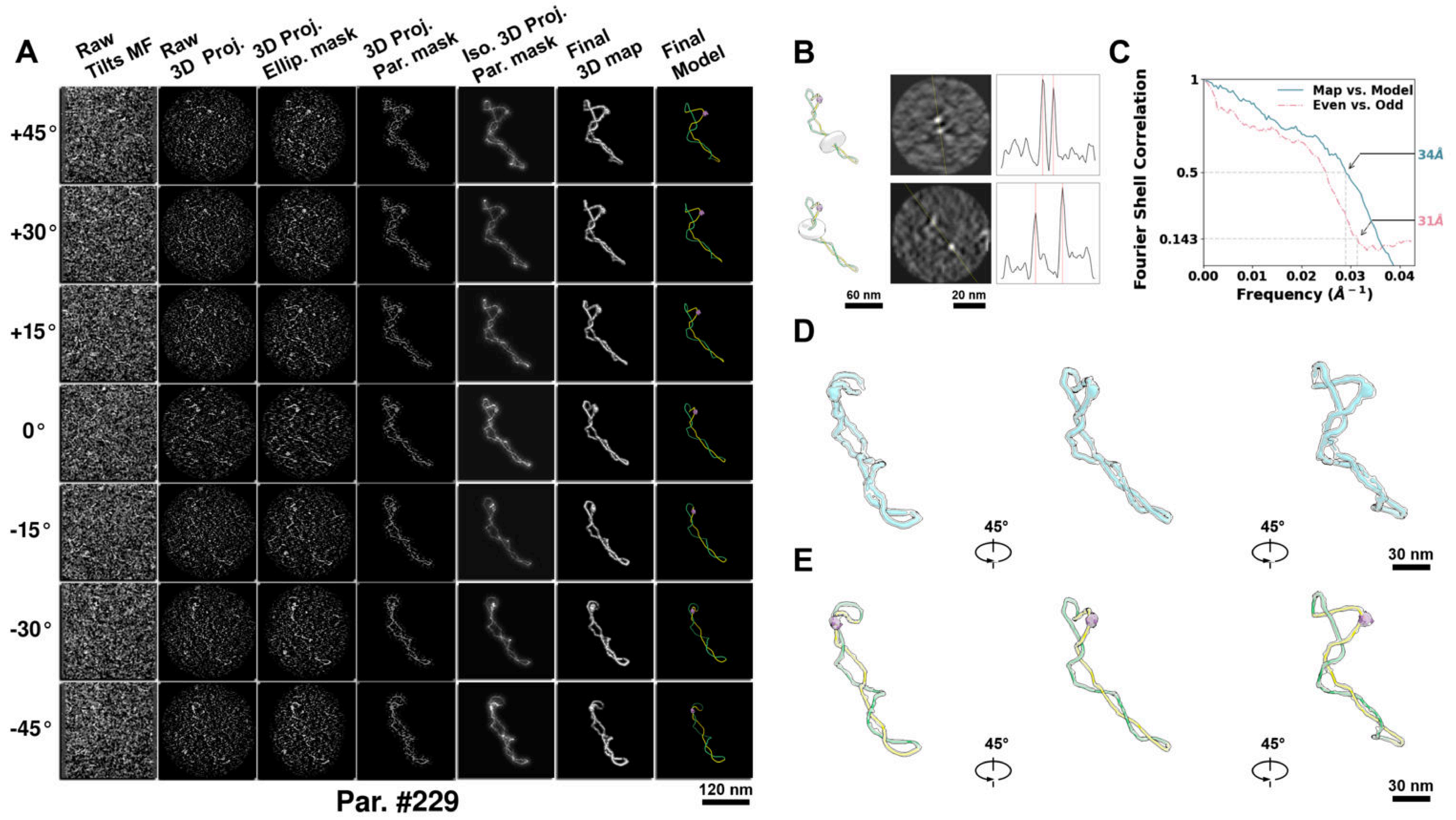

**Supplementary Particle Figure 229. Cryo-ET 3D reconstruction of an individual TEC-TopI particle.**
(A) 3D reconstruction of the plasmid particle (index no. 229). The first column shows seven representative tilt images from +45° to -45° in step of 15°. The second, third, and fourth columns show 3D projections of the particle with spherical, ellipsoidal (thinner along the z-dimension), and particle-shaped masks, respectively. The fifth column displays the 3D projections of the enhanced and IsoNet missing-wedge-corrected particle. The sixth and seventh columns present the final 3D map and the flexibly fitted model, respectively. (B) Two cross-sectional views (12 nm thickness) of the plasmid density map along its plectoneme axis are shown in the left-middle panel. The intensity profile along the line crossing the two high-density DNA spots is displayed in the right panel. (C) Resolution assessment of the final 3D map using Fourier shell correlation (FSC). Two criteria are shown: FSC between two half-maps reconstructed from even and odd frames (evaluated at 0.143) and FSC between the final 3D map and the fitted model (evaluated at 0.5). (D) Zoomed-in views of the final 3D density map from panel A, displayed at two contour levels. (E) Superimposition of the high-contour level map from panel D onto its fitted model.

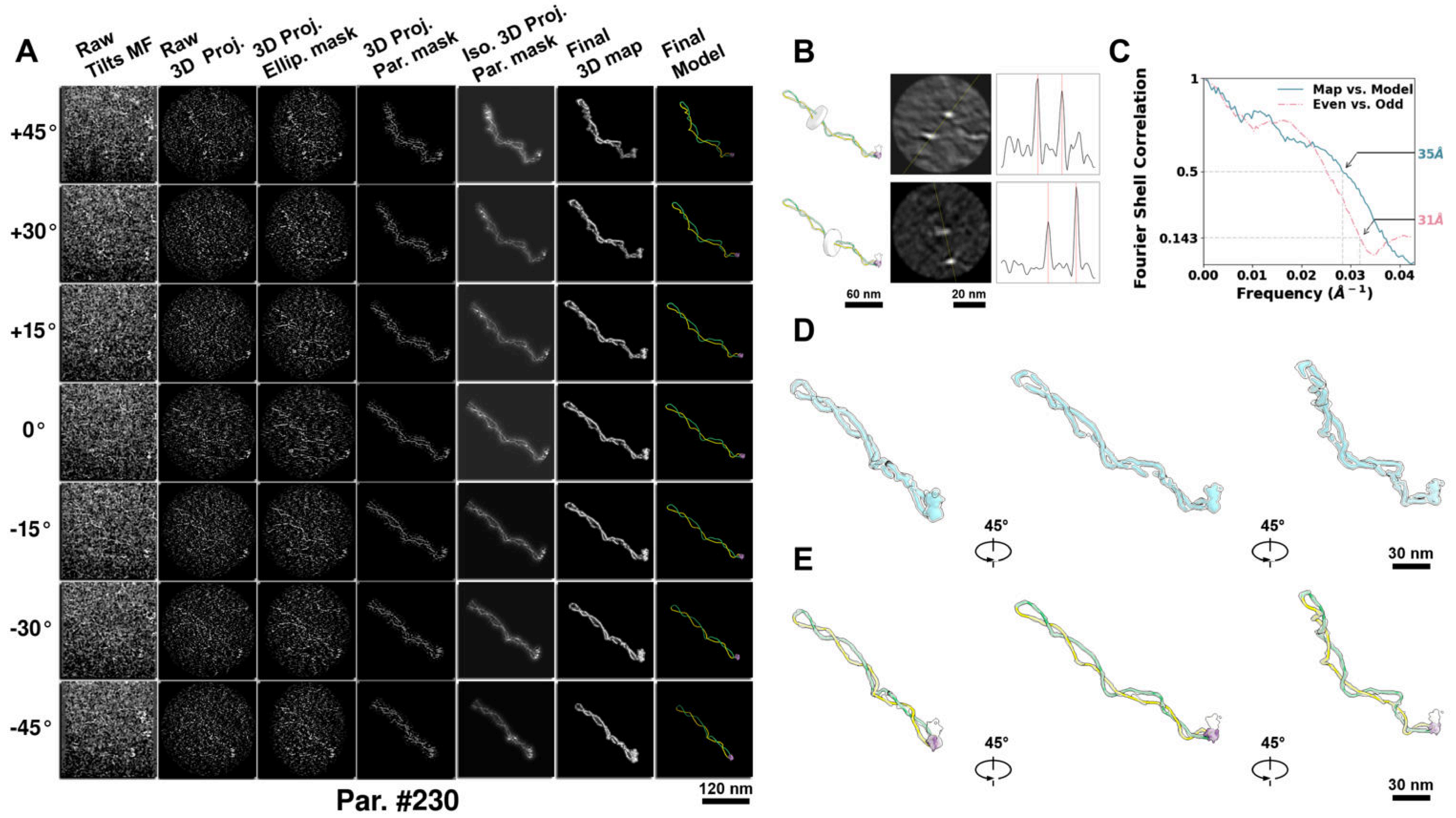

**Supplementary Particle Figure 230. Cryo-ET 3D reconstruction of an individual TEC-TopI particle.**
(A) 3D reconstruction of the plasmid particle (index no. 230). The first column shows seven representative tilt images from +45° to -45° in step of 15°. The second, third, and fourth columns show 3D projections of the particle with spherical, ellipsoidal (thinner along the z-dimension), and particle-shaped masks, respectively. The fifth column displays the 3D projections of the enhanced and IsoNet missing-wedge-corrected particle. The sixth and seventh columns present the final 3D map and the flexibly fitted model, respectively. (B) Two cross-sectional views (12 nm thickness) of the plasmid density map along its plectoneme axis are shown in the left-middle panel. The intensity profile along the line crossing the two high-density DNA spots is displayed in the right panel. (C) Resolution assessment of the final 3D map using Fourier shell correlation (FSC). Two criteria are shown: FSC between two half-maps reconstructed from even and odd frames (evaluated at 0.143) and FSC between the final 3D map and the fitted model (evaluated at 0.5). (D) Zoomed-in views of the final 3D density map from panel A, displayed at two contour levels. (E) Superimposition of the high-contour level map from panel D onto its fitted model.

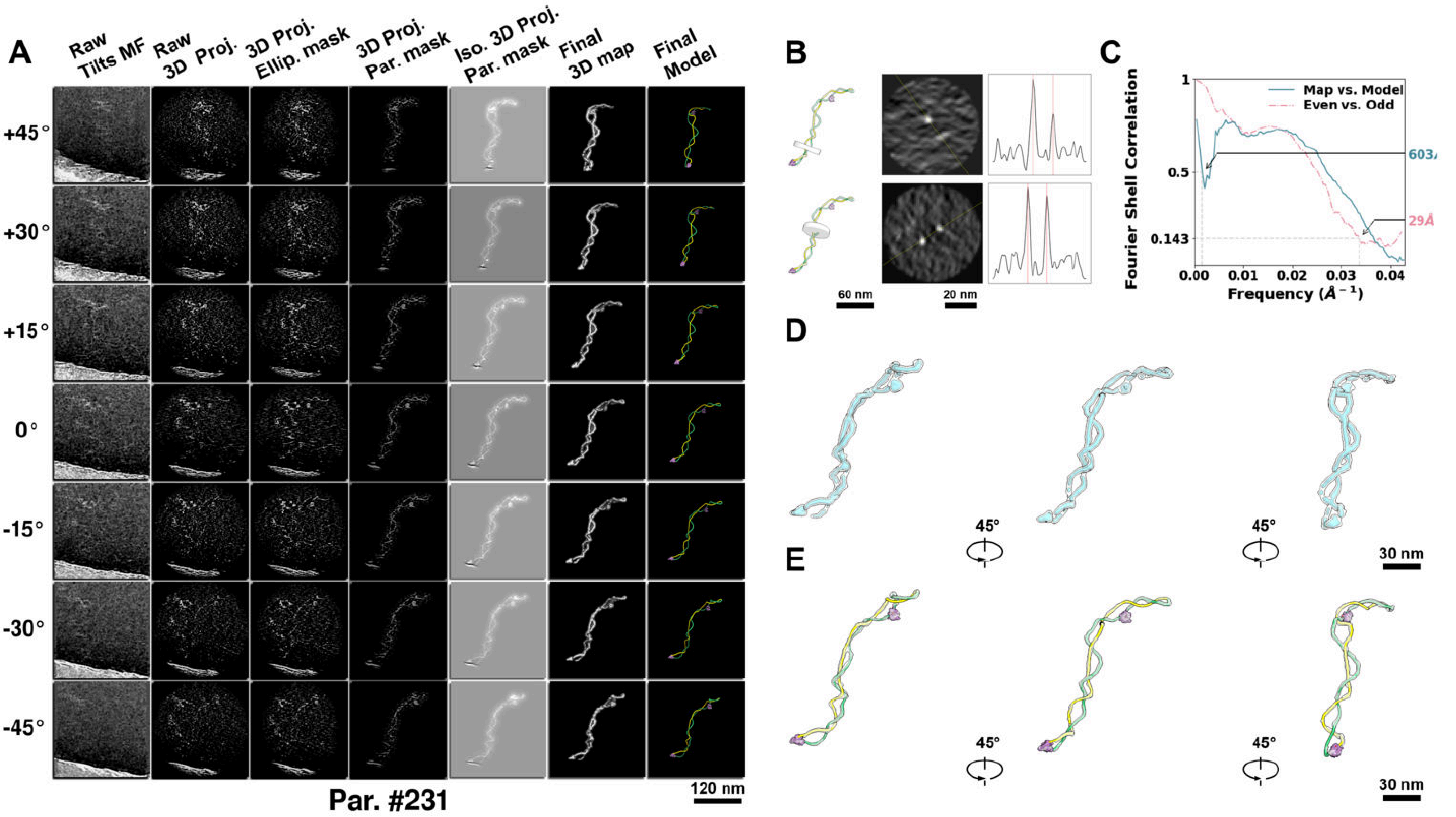

**Supplementary Particle Figure 231. Cryo-ET 3D reconstruction of an individual TEC-TopI particle.**
(A) 3D reconstruction of the plasmid particle (index no. 231). The first column shows seven representative tilt images from +45° to -45° in step of 15°. The second, third, and fourth columns show 3D projections of the particle with spherical, ellipsoidal (thinner along the z-dimension), and particle-shaped masks, respectively. The fifth column displays the 3D projections of the enhanced and IsoNet missing-wedge-corrected particle. The sixth and seventh columns present the final 3D map and the flexibly fitted model, respectively. (B) Two cross-sectional views (12 nm thickness) of the plasmid density map along its plectoneme axis are shown in the left-middle panel. The intensity profile along the line crossing the two high-density DNA spots is displayed in the right panel. (C) Resolution assessment of the final 3D map using Fourier shell correlation (FSC). Two criteria are shown: FSC between two half-maps reconstructed from even and odd frames (evaluated at 0.143) and FSC between the final 3D map and the fitted model (evaluated at 0.5). (D) Zoomed-in views of the final 3D density map from panel A, displayed at two contour levels. (E) Superimposition of the high-contour level map from panel D onto its fitted model.

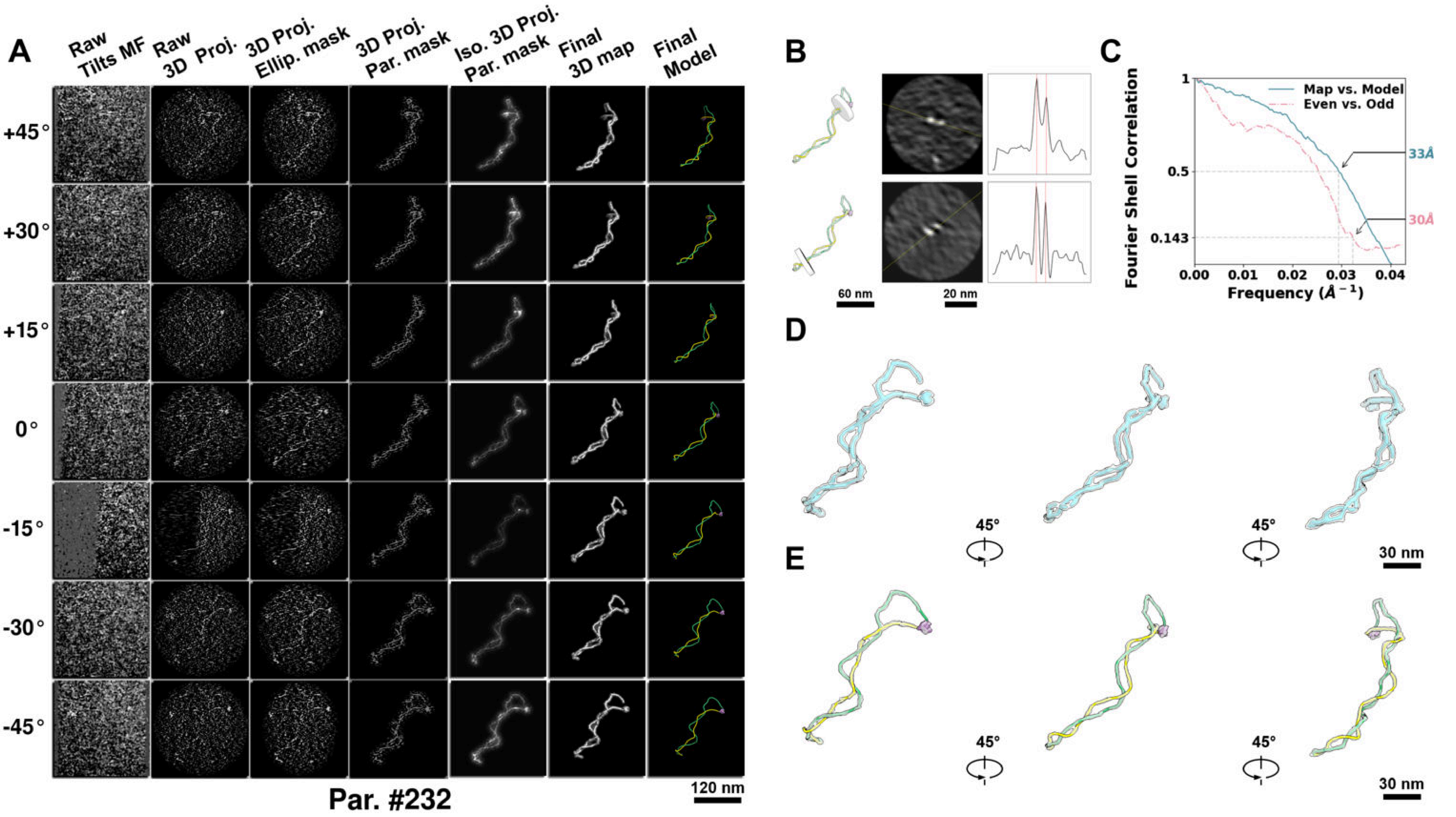

**Supplementary Particle Figure 232. Cryo-ET 3D reconstruction of an individual TEC-TopI particle.**
(A) 3D reconstruction of the plasmid particle (index no. 232). The first column shows seven representative tilt images from +45° to -45° in step of 15°. The second, third, and fourth columns show 3D projections of the particle with spherical, ellipsoidal (thinner along the z-dimension), and particle-shaped masks, respectively. The fifth column displays the 3D projections of the enhanced and IsoNet missing-wedge-corrected particle. The sixth and seventh columns present the final 3D map and the flexibly fitted model, respectively. (B) Two cross-sectional views (12 nm thickness) of the plasmid density map along its plectoneme axis are shown in the left-middle panel. The intensity profile along the line crossing the two high-density DNA spots is displayed in the right panel. (C) Resolution assessment of the final 3D map using Fourier shell correlation (FSC). Two criteria are shown: FSC between two half-maps reconstructed from even and odd frames (evaluated at 0.143) and FSC between the final 3D map and the fitted model (evaluated at 0.5). (D) Zoomed-in views of the final 3D density map from panel A, displayed at two contour levels. (E) Superimposition of the high-contour level map from panel D onto its fitted model.

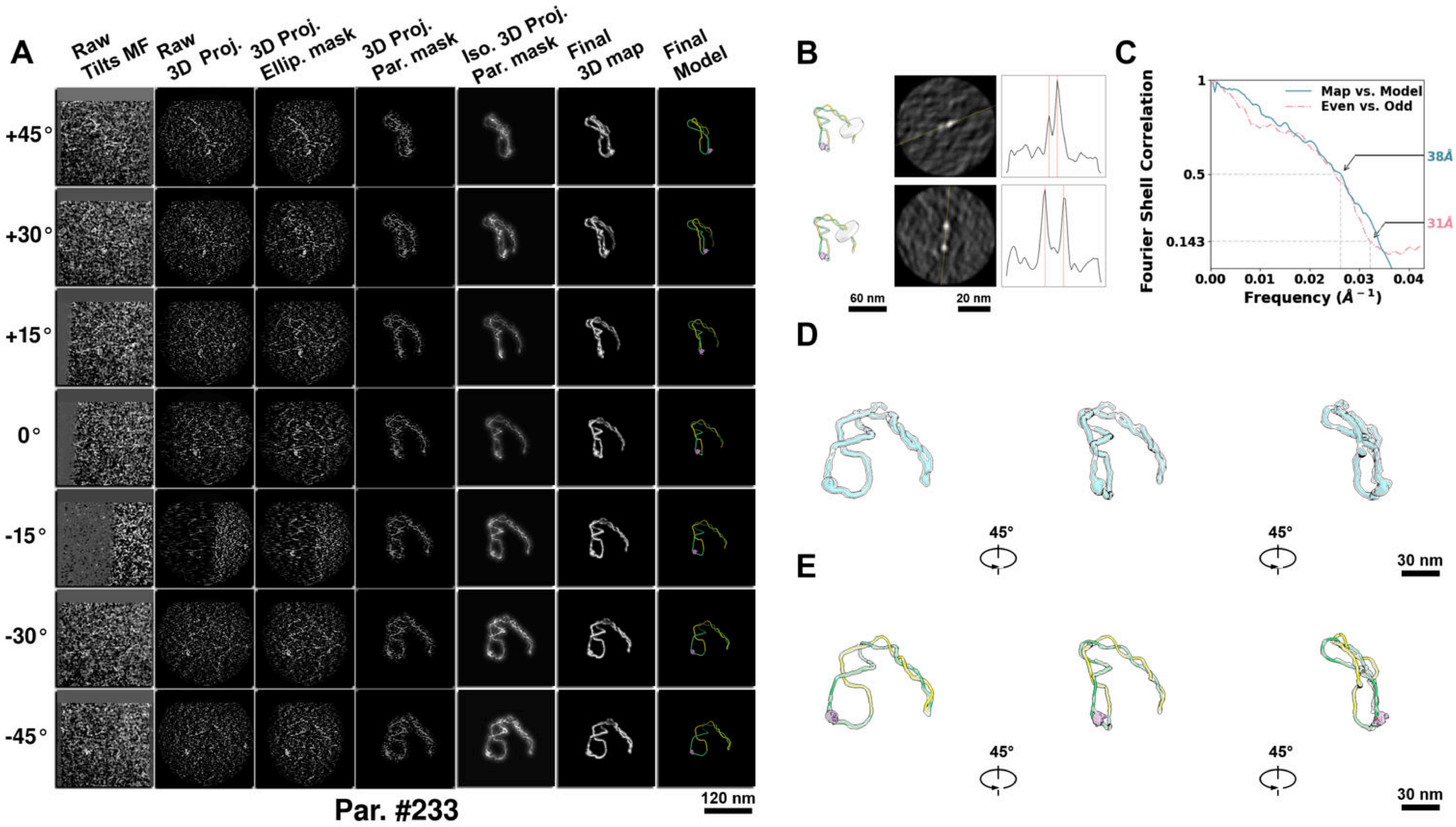

**Supplementary Particle Figure 233. Cryo-ET 3D reconstruction of an individual TEC-TopI particle.**
(A) 3D reconstruction of the plasmid particle (index no. 233). The first column shows seven representative tilt images from +45° to -45° in step of 15°. The second, third, and fourth columns show 3D projections of the particle with spherical, ellipsoidal (thinner along the z-dimension), and particle-shaped masks, respectively. The fifth column displays the 3D projections of the enhanced and IsoNet missing-wedge-corrected particle. The sixth and seventh columns present the final 3D map and the flexibly fitted model, respectively. (B) Two cross-sectional views (12 nm thickness) of the plasmid density map along its plectoneme axis are shown in the left-middle panel. The intensity profile along the line crossing the two high-density DNA spots is displayed in the right panel. (C) Resolution assessment of the final 3D map using Fourier shell correlation (FSC). Two criteria are shown: FSC between two half-maps reconstructed from even and odd frames (evaluated at 0.143) and FSC between the final 3D map and the fitted model (evaluated at 0.5). (D) Zoomed-in views of the final 3D density map from panel A, displayed at two contour levels. (E) Superimposition of the high-contour level map from panel D onto its fitted model.

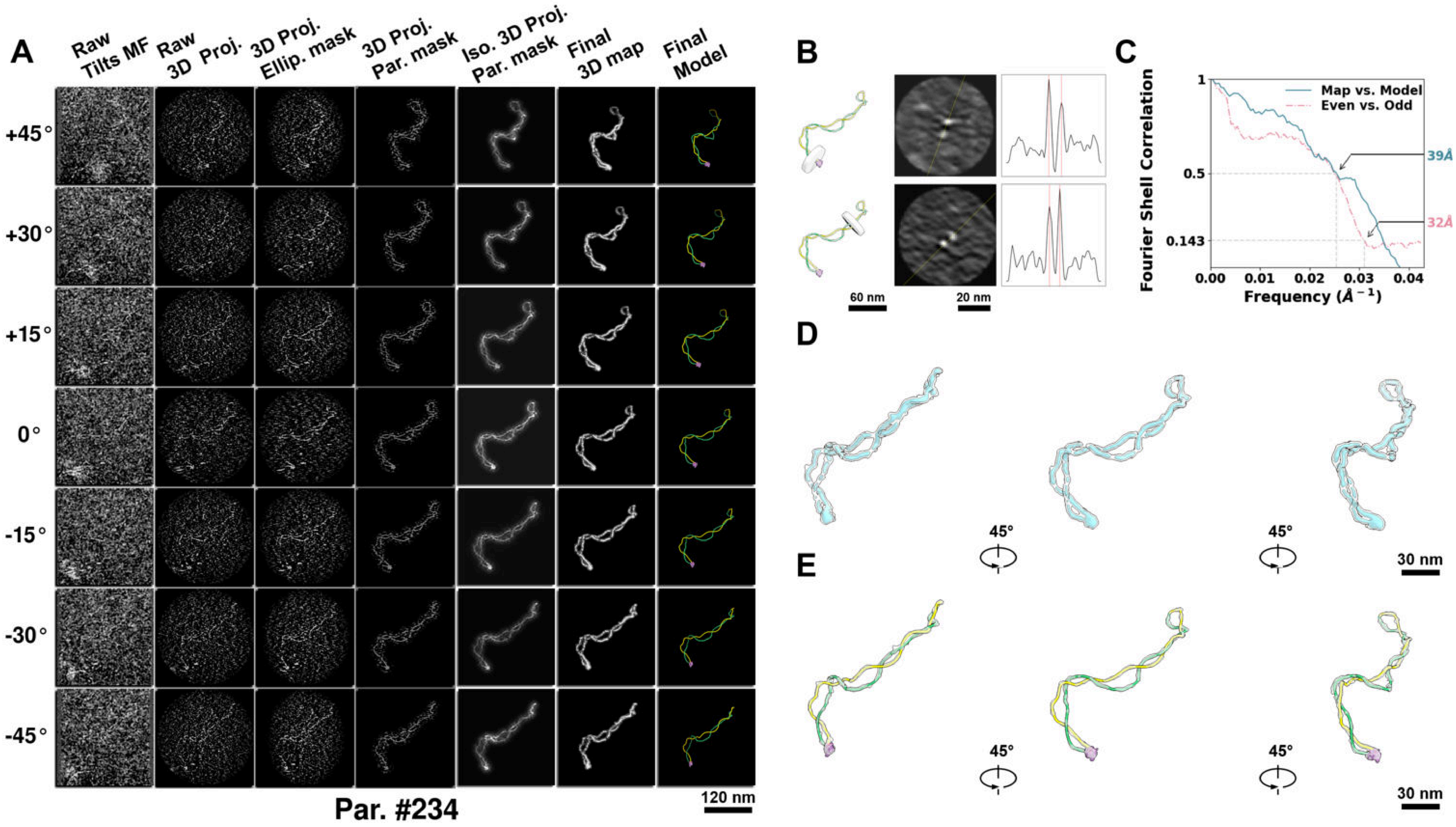

**Supplementary Particle Figure 234. Cryo-ET 3D reconstruction of an individual TEC-TopI particle.**
(A) 3D reconstruction of the plasmid particle (index no. 234). The first column shows seven representative tilt images from +45° to -45° in step of 15°. The second, third, and fourth columns show 3D projections of the particle with spherical, ellipsoidal (thinner along the z-dimension), and particle-shaped masks, respectively. The fifth column displays the 3D projections of the enhanced and IsoNet missing-wedge-corrected particle. The sixth and seventh columns present the final 3D map and the flexibly fitted model, respectively. (B) Two cross-sectional views (12 nm thickness) of the plasmid density map along its plectoneme axis are shown in the left-middle panel. The intensity profile along the line crossing the two high-density DNA spots is displayed in the right panel. (C) Resolution assessment of the final 3D map using Fourier shell correlation (FSC). Two criteria are shown: FSC between two half-maps reconstructed from even and odd frames (evaluated at 0.143) and FSC between the final 3D map and the fitted model (evaluated at 0.5). (D) Zoomed-in views of the final 3D density map from panel A, displayed at two contour levels. (E) Superimposition of the high-contour level map from panel D onto its fitted model.

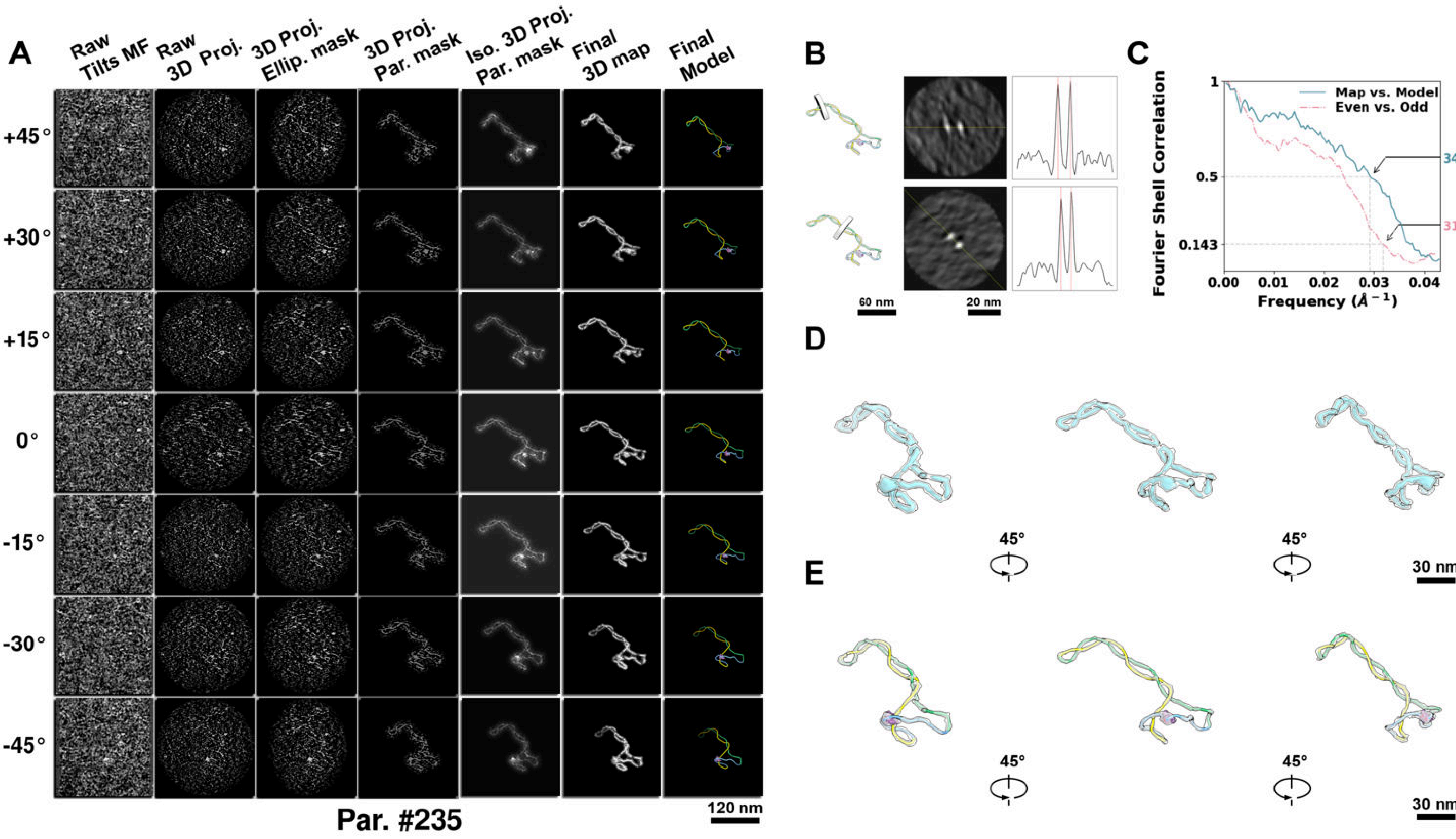

**Supplementary Particle Figure 235. Cryo-ET 3D reconstruction of an individual TEC-TopI particle.**
(A) 3D reconstruction of the plasmid particle (index no. 235). The first column shows seven representative tilt images from +45° to -45° in step of 15°. The second, third, and fourth columns show 3D projections of the particle with spherical, ellipsoidal (thinner along the z-dimension), and particle-shaped masks, respectively. The fifth column displays the 3D projections of the enhanced and IsoNet missing-wedge-corrected particle. The sixth and seventh columns present the final 3D map and the flexibly fitted model, respectively. (B) Two cross-sectional views (12 nm thickness) of the plasmid density map along its plectoneme axis are shown in the left-middle panel. The intensity profile along the line crossing the two high-density DNA spots is displayed in the right panel. (C) Resolution assessment of the final 3D map using Fourier shell correlation (FSC). Two criteria are shown: FSC between two half-maps reconstructed from even and odd frames (evaluated at 0.143) and FSC between the final 3D map and the fitted model (evaluated at 0.5). (D) Zoomed-in views of the final 3D density map from panel A, displayed at two contour levels. (E) Superimposition of the high-contour level map from panel D onto its fitted model.

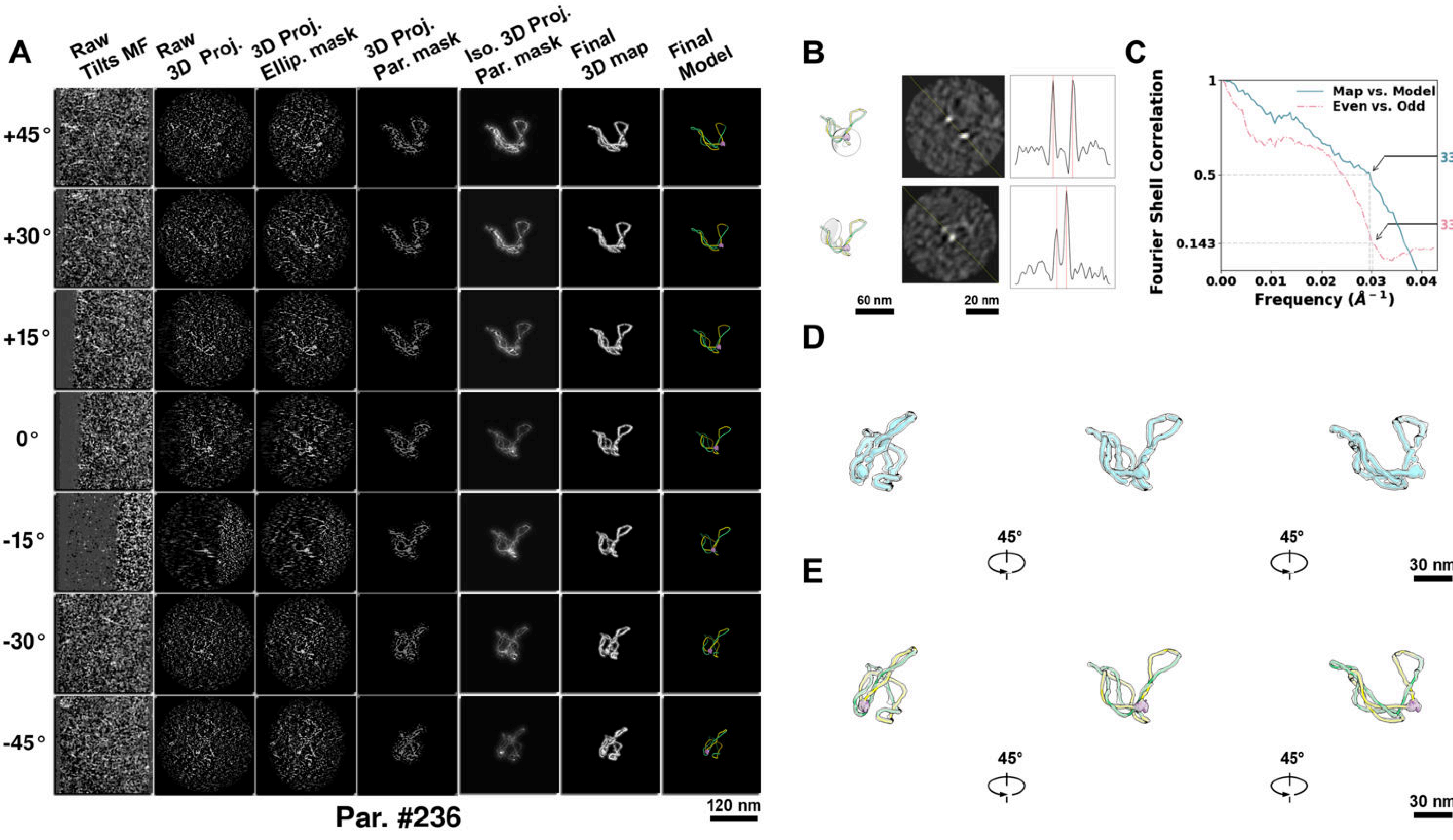

**Supplementary Particle Figure 236. Cryo-ET 3D reconstruction of an individual TEC-TopI particle.**
(A) 3D reconstruction of the plasmid particle (index no. 236). The first column shows seven representative tilt images from +45° to -45° in step of 15°. The second, third, and fourth columns show 3D projections of the particle with spherical, ellipsoidal (thinner along the z-dimension), and particle-shaped masks, respectively. The fifth column displays the 3D projections of the enhanced and IsoNet missing-wedge-corrected particle. The sixth and seventh columns present the final 3D map and the flexibly fitted model, respectively. (B) Two cross-sectional views (12 nm thickness) of the plasmid density map along its plectoneme axis are shown in the left-middle panel. The intensity profile along the line crossing the two high-density DNA spots is displayed in the right panel. (C) Resolution assessment of the final 3D map using Fourier shell correlation (FSC). Two criteria are shown: FSC between two half-maps reconstructed from even and odd frames (evaluated at 0.143) and FSC between the final 3D map and the fitted model (evaluated at 0.5). (D) Zoomed-in views of the final 3D density map from panel A, displayed at two contour levels. (E) Superimposition of the high-contour level map from panel D onto its fitted model.

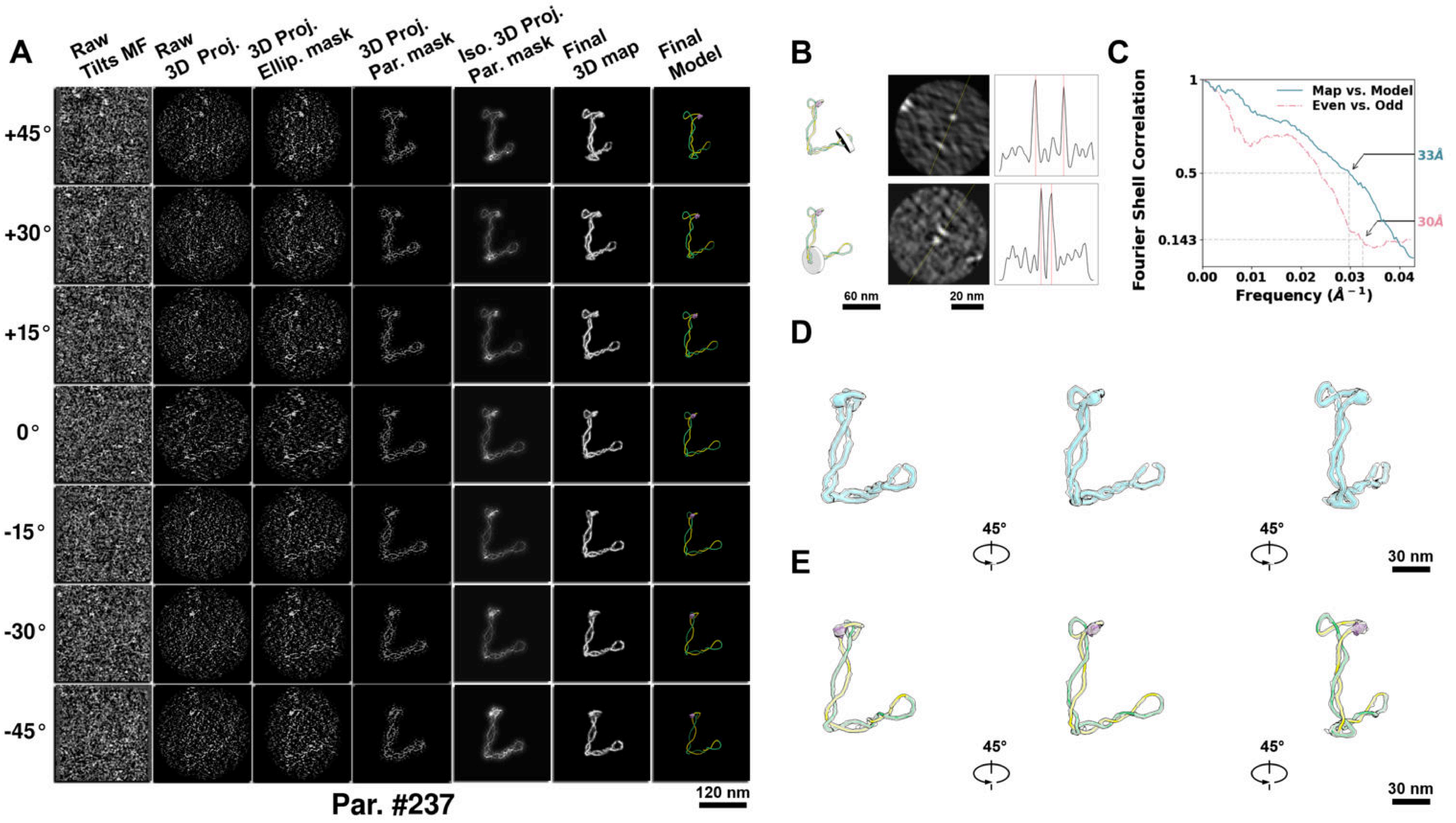

**Supplementary Particle Figure 237. Cryo-ET 3D reconstruction of an individual TEC-TopI particle.**
(A) 3D reconstruction of the plasmid particle (index no. 237). The first column shows seven representative tilt images from +45° to -45° in step of 15°. The second, third, and fourth columns show 3D projections of the particle with spherical, ellipsoidal (thinner along the z-dimension), and particle-shaped masks, respectively. The fifth column displays the 3D projections of the enhanced and IsoNet missing-wedge-corrected particle. The sixth and seventh columns present the final 3D map and the flexibly fitted model, respectively. (B) Two cross-sectional views (12 nm thickness) of the plasmid density map along its plectoneme axis are shown in the left-middle panel. The intensity profile along the line crossing the two high-density DNA spots is displayed in the right panel. (C) Resolution assessment of the final 3D map using Fourier shell correlation (FSC). Two criteria are shown: FSC between two half-maps reconstructed from even and odd frames (evaluated at 0.143) and FSC between the final 3D map and the fitted model (evaluated at 0.5). (D) Zoomed-in views of the final 3D density map from panel A, displayed at two contour levels. (E) Superimposition of the high-contour level map from panel D onto its fitted model.

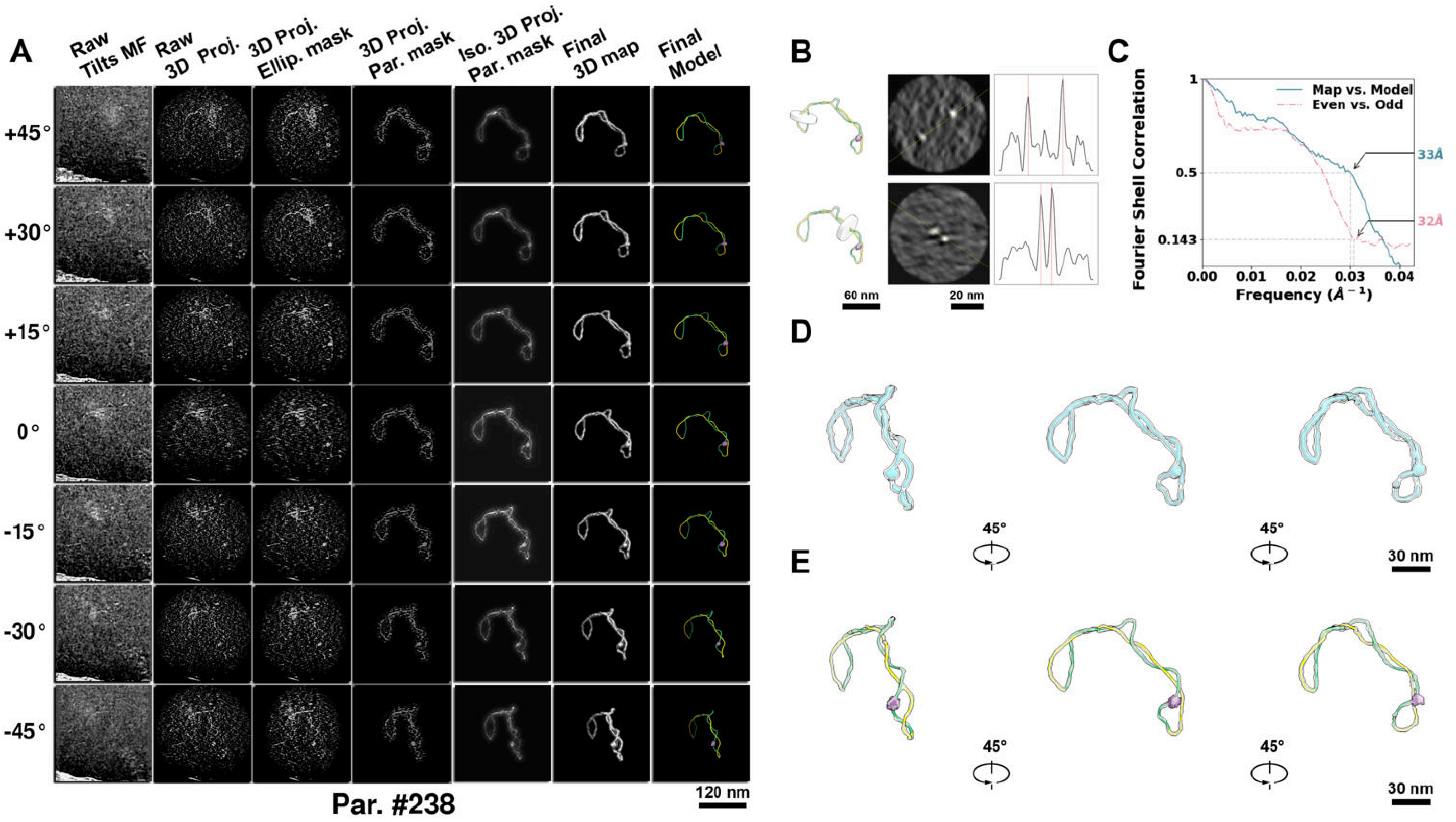

**Supplementary Particle Figure 238. Cryo-ET 3D reconstruction of an individual TEC-TopI particle.**
(A) 3D reconstruction of the plasmid particle (index no. 238). The first column shows seven representative tilt images from +45° to -45° in step of 15°. The second, third, and fourth columns show 3D projections of the particle with spherical, ellipsoidal (thinner along the z-dimension), and particle-shaped masks, respectively. The fifth column displays the 3D projections of the enhanced and IsoNet missing-wedge-corrected particle. The sixth and seventh columns present the final 3D map and the flexibly fitted model, respectively. (B) Two cross-sectional views (12 nm thickness) of the plasmid density map along its plectoneme axis are shown in the left-middle panel. The intensity profile along the line crossing the two high-density DNA spots is displayed in the right panel. (C) Resolution assessment of the final 3D map using Fourier shell correlation (FSC). Two criteria are shown: FSC between two half-maps reconstructed from even and odd frames (evaluated at 0.143) and FSC between the final 3D map and the fitted model (evaluated at 0.5). (D) Zoomed-in views of the final 3D density map from panel A, displayed at two contour levels. (E) Superimposition of the high-contour level map from panel D onto its fitted model.

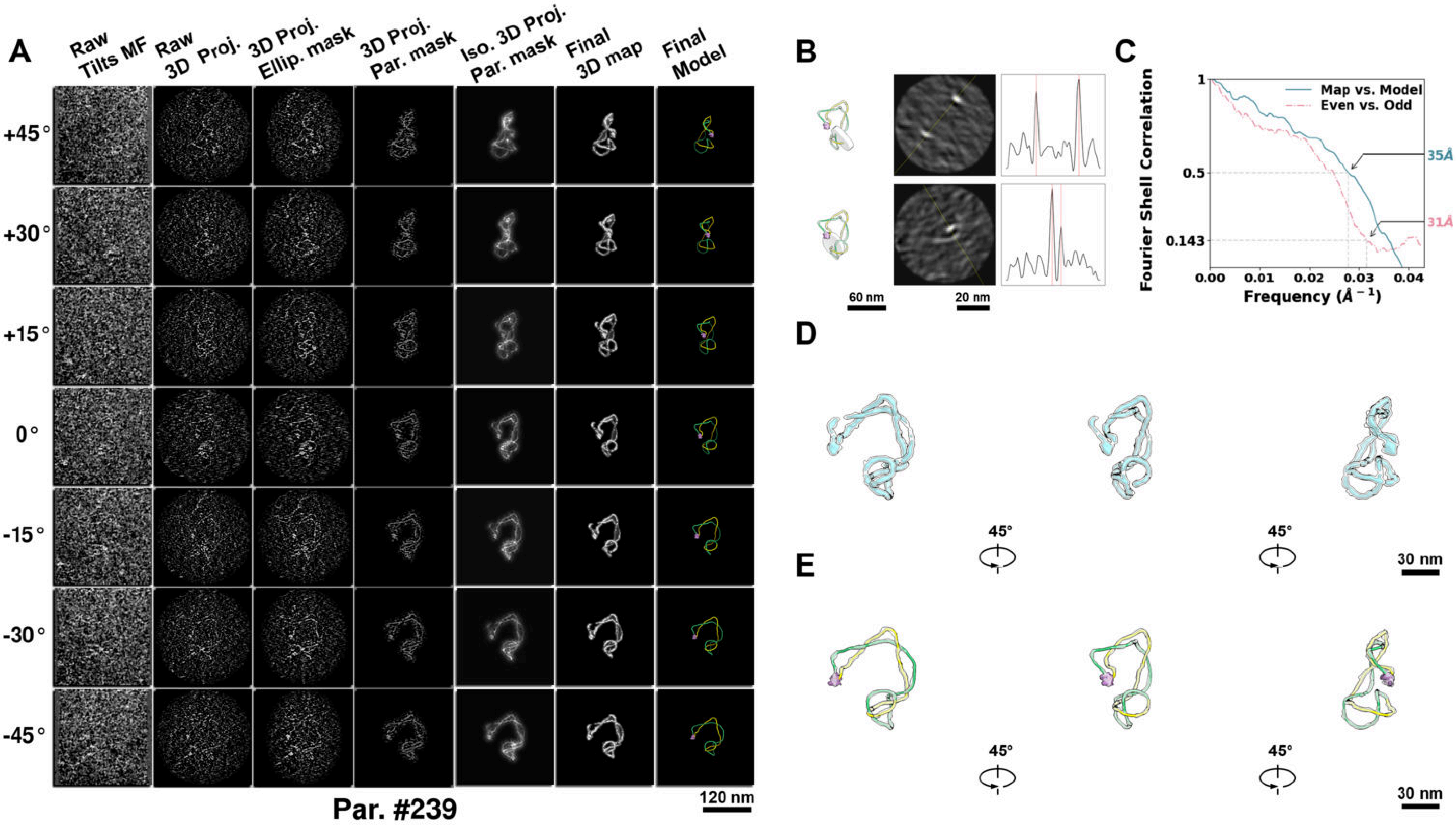

**Supplementary Particle Figure 239. Cryo-ET 3D reconstruction of an individual TEC-TopI particle.**
(A) 3D reconstruction of the plasmid particle (index no. 239). The first column shows seven representative tilt images from +45° to -45° in step of 15°. The second, third, and fourth columns show 3D projections of the particle with spherical, ellipsoidal (thinner along the z-dimension), and particle-shaped masks, respectively. The fifth column displays the 3D projections of the enhanced and IsoNet missing-wedge-corrected particle. The sixth and seventh columns present the final 3D map and the flexibly fitted model, respectively. (B) Two cross-sectional views (12 nm thickness) of the plasmid density map along its plectoneme axis are shown in the left-middle panel. The intensity profile along the line crossing the two high-density DNA spots is displayed in the right panel. (C) Resolution assessment of the final 3D map using Fourier shell correlation (FSC). Two criteria are shown: FSC between two half-maps reconstructed from even and odd frames (evaluated at 0.143) and FSC between the final 3D map and the fitted model (evaluated at 0.5). (D) Zoomed-in views of the final 3D density map from panel A, displayed at two contour levels. (E) Superimposition of the high-contour level map from panel D onto its fitted model.

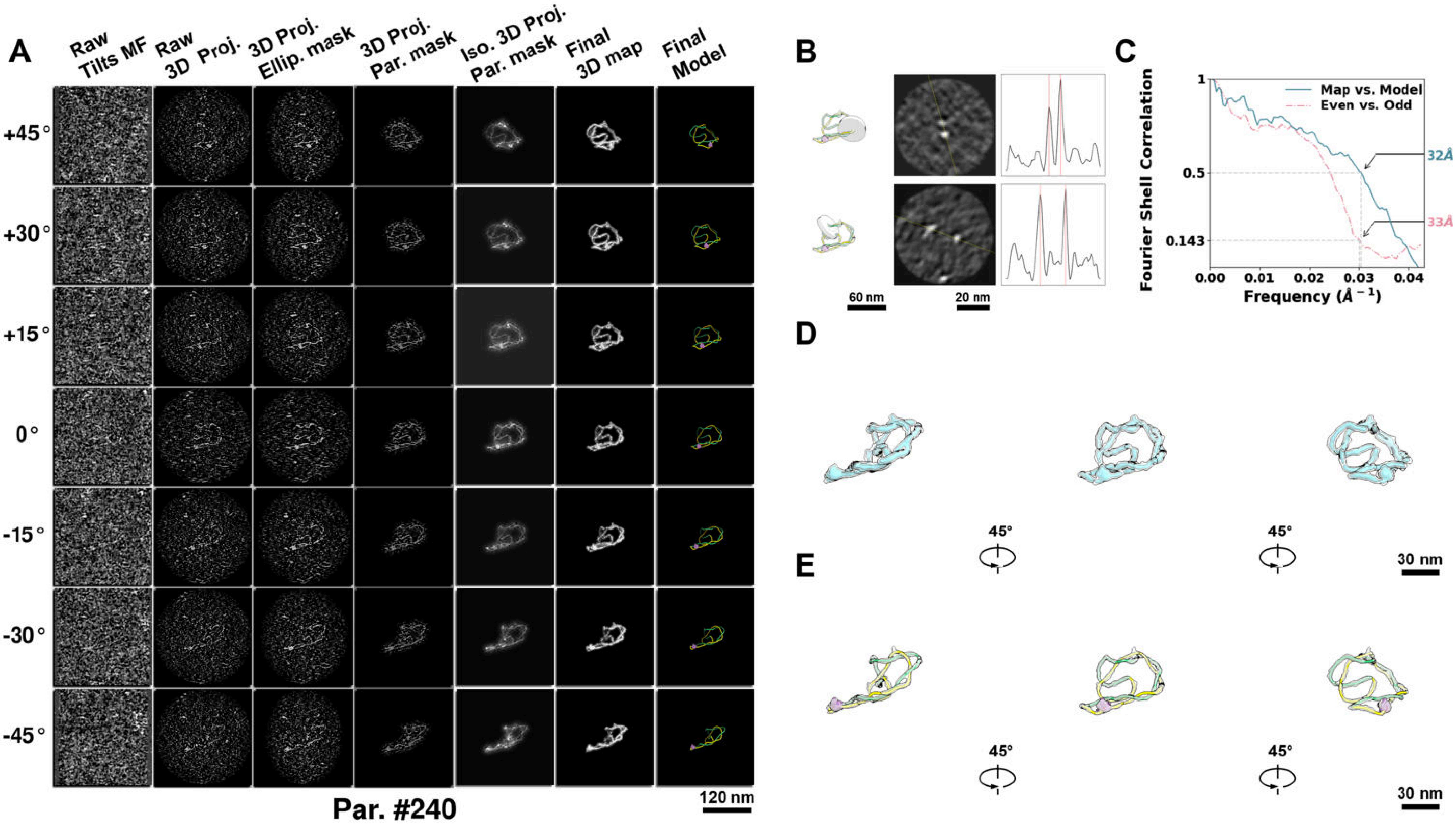

**Supplementary Particle Figure 240. Cryo-ET 3D reconstruction of an individual TEC-TopI particle.**
(A) 3D reconstruction of the plasmid particle (index no. 240). The first column shows seven representative tilt images from +45° to -45° in step of 15°. The second, third, and fourth columns show 3D projections of the particle with spherical, ellipsoidal (thinner along the z-dimension), and particle-shaped masks, respectively. The fifth column displays the 3D projections of the enhanced and IsoNet missing-wedge-corrected particle. The sixth and seventh columns present the final 3D map and the flexibly fitted model, respectively. (B) Two cross-sectional views (12 nm thickness) of the plasmid density map along its plectoneme axis are shown in the left-middle panel. The intensity profile along the line crossing the two high-density DNA spots is displayed in the right panel. (C) Resolution assessment of the final 3D map using Fourier shell correlation (FSC). Two criteria are shown: FSC between two half-maps reconstructed from even and odd frames (evaluated at 0.143) and FSC between the final 3D map and the fitted model (evaluated at 0.5). (D) Zoomed-in views of the final 3D density map from panel A, displayed at two contour levels. (E) Superimposition of the high-contour level map from panel D onto its fitted model.

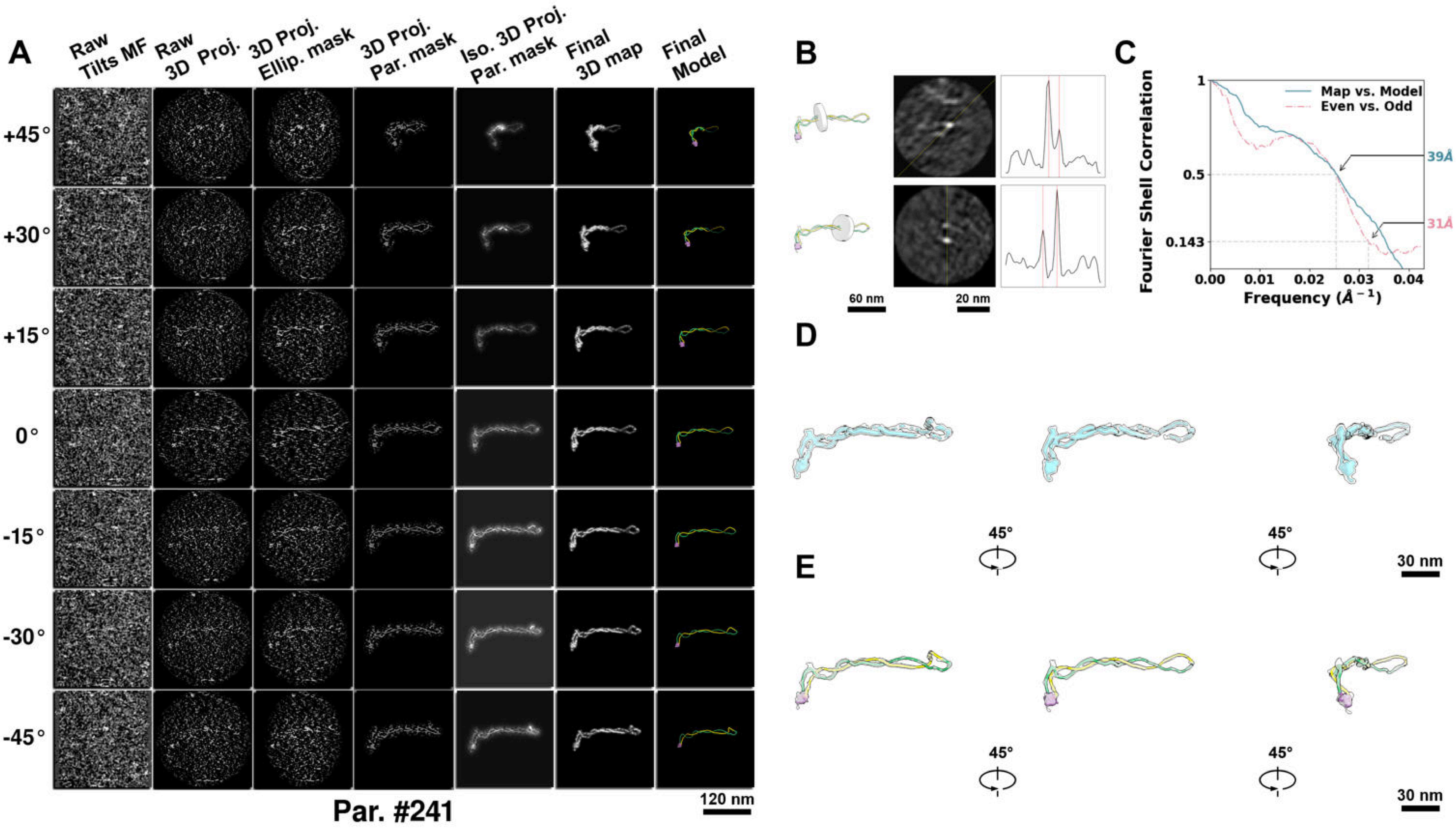

**Supplementary Particle Figure 241. Cryo-ET 3D reconstruction of an individual TEC-TopI particle.**
(A) 3D reconstruction of the plasmid particle (index no. 241). The first column shows seven representative tilt images from +45° to -45° in step of 15°. The second, third, and fourth columns show 3D projections of the particle with spherical, ellipsoidal (thinner along the z-dimension), and particle-shaped masks, respectively. The fifth column displays the 3D projections of the enhanced and IsoNet missing-wedge-corrected particle. The sixth and seventh columns present the final 3D map and the flexibly fitted model, respectively. (B) Two cross-sectional views (12 nm thickness) of the plasmid density map along its plectoneme axis are shown in the left-middle panel. The intensity profile along the line crossing the two high-density DNA spots is displayed in the right panel. (C) Resolution assessment of the final 3D map using Fourier shell correlation (FSC). Two criteria are shown: FSC between two half-maps reconstructed from even and odd frames (evaluated at 0.143) and FSC between the final 3D map and the fitted model (evaluated at 0.5). (D) Zoomed-in views of the final 3D density map from panel A, displayed at two contour levels. (E) Superimposition of the high-contour level map from panel D onto its fitted model.

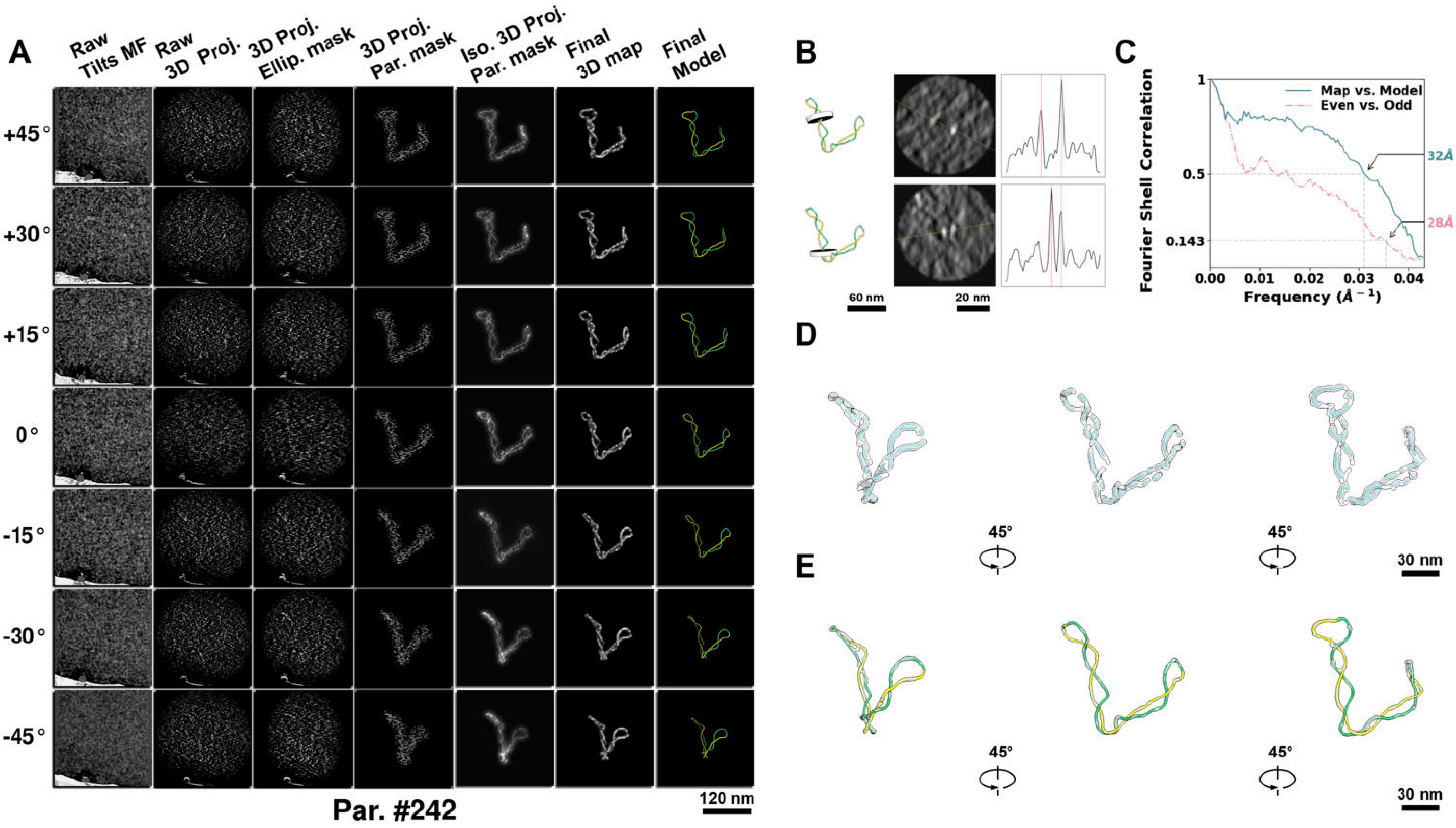

**Supplementary Particle Figure 242. Cryo-ET 3D reconstruction of an individual TEC-TopI particle.**
(A) 3D reconstruction of the plasmid particle (index no. 242). The first column shows seven representative tilt images from +45° to -45° in step of 15°. The second, third, and fourth columns show 3D projections of the particle with spherical, ellipsoidal (thinner along the z-dimension), and particle-shaped masks, respectively. The fifth column displays the 3D projections of the enhanced and IsoNet missing-wedge-corrected particle. The sixth and seventh columns present the final 3D map and the flexibly fitted model, respectively. (B) Two cross-sectional views (12 nm thickness) of the plasmid density map along its plectoneme axis are shown in the left-middle panel. The intensity profile along the line crossing the two high-density DNA spots is displayed in the right panel. (C) Resolution assessment of the final 3D map using Fourier shell correlation (FSC). Two criteria are shown: FSC between two half-maps reconstructed from even and odd frames (evaluated at 0.143) and FSC between the final 3D map and the fitted model (evaluated at 0.5). (D) Zoomed-in views of the final 3D density map from panel A, displayed at two contour levels. (E) Superimposition of the high-contour level map from panel D onto its fitted model.

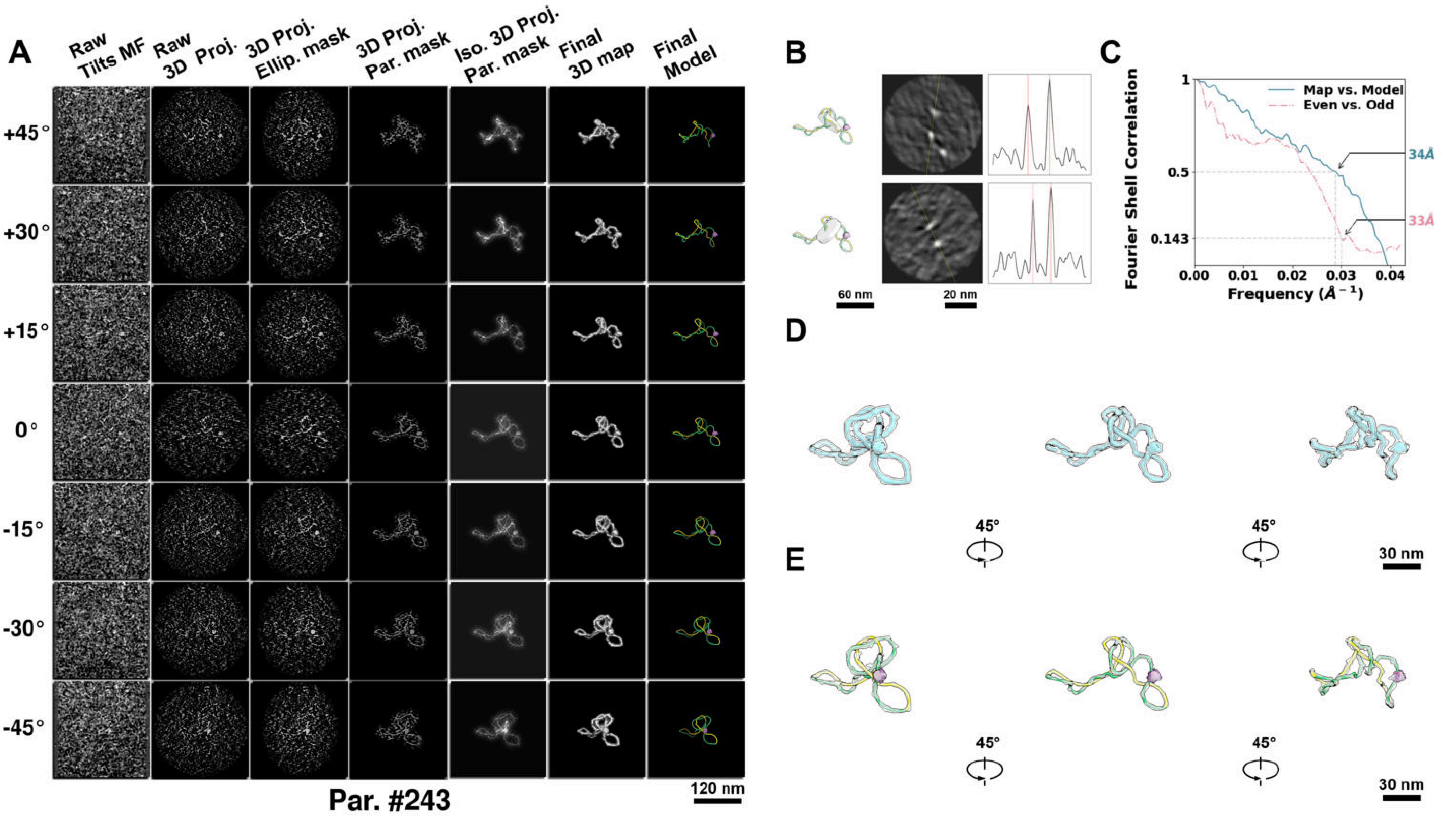

**Supplementary Particle Figure 243. Cryo-ET 3D reconstruction of an individual TEC-TopI particle.**
(A) 3D reconstruction of the plasmid particle (index no. 243). The first column shows seven representative tilt images from +45° to -45° in step of 15°. The second, third, and fourth columns show 3D projections of the particle with spherical, ellipsoidal (thinner along the z-dimension), and particle-shaped masks, respectively. The fifth column displays the 3D projections of the enhanced and IsoNet missing-wedge-corrected particle. The sixth and seventh columns present the final 3D map and the flexibly fitted model, respectively. (B) Two cross-sectional views (12 nm thickness) of the plasmid density map along its plectoneme axis are shown in the left-middle panel. The intensity profile along the line crossing the two high-density DNA spots is displayed in the right panel. (C) Resolution assessment of the final 3D map using Fourier shell correlation (FSC). Two criteria are shown: FSC between two half-maps reconstructed from even and odd frames (evaluated at 0.143) and FSC between the final 3D map and the fitted model (evaluated at 0.5). (D) Zoomed-in views of the final 3D density map from panel A, displayed at two contour levels. (E) Superimposition of the high-contour level map from panel D onto its fitted model.

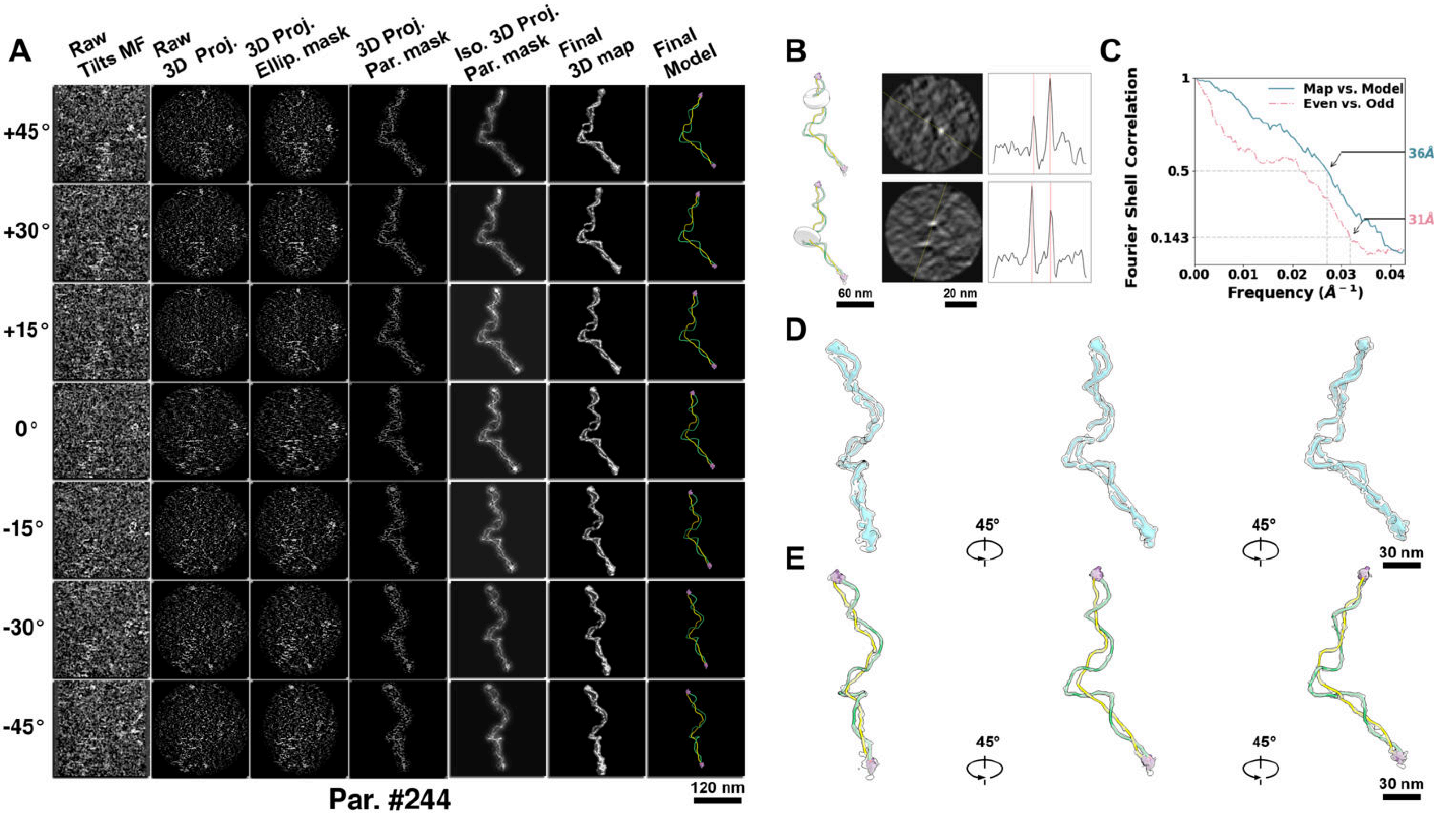

**Supplementary Particle Figure 244. Cryo-ET 3D reconstruction of an individual TEC-TopI particle.**
(A) 3D reconstruction of the plasmid particle (index no. 244). The first column shows seven representative tilt images from +45° to -45° in step of 15°. The second, third, and fourth columns show 3D projections of the particle with spherical, ellipsoidal (thinner along the z-dimension), and particle-shaped masks, respectively. The fifth column displays the 3D projections of the enhanced and IsoNet missing-wedge-corrected particle. The sixth and seventh columns present the final 3D map and the flexibly fitted model, respectively. (B) Two cross-sectional views (12 nm thickness) of the plasmid density map along its plectoneme axis are shown in the left-middle panel. The intensity profile along the line crossing the two high-density DNA spots is displayed in the right panel. (C) Resolution assessment of the final 3D map using Fourier shell correlation (FSC). Two criteria are shown: FSC between two half-maps reconstructed from even and odd frames (evaluated at 0.143) and FSC between the final 3D map and the fitted model (evaluated at 0.5). (D) Zoomed-in views of the final 3D density map from panel A, displayed at two contour levels. (E) Superimposition of the high-contour level map from panel D onto its fitted model.

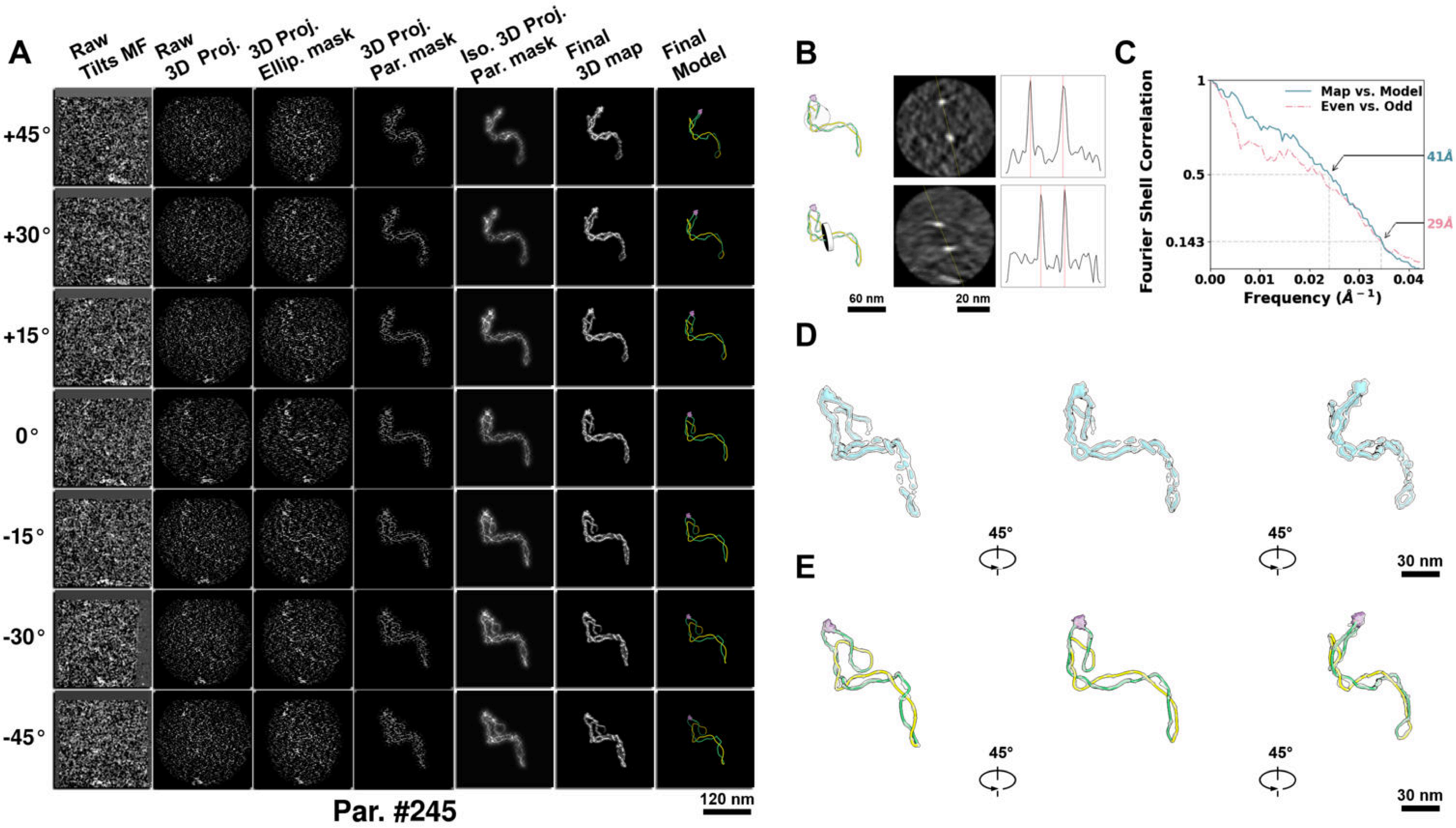

**Supplementary Particle Figure 245. Cryo-ET 3D reconstruction of an individual TEC-TopI particle.**
(A) 3D reconstruction of the plasmid particle (index no. 245). The first column shows seven representative tilt images from +45° to -45° in step of 15°. The second, third, and fourth columns show 3D projections of the particle with spherical, ellipsoidal (thinner along the z-dimension), and particle-shaped masks, respectively. The fifth column displays the 3D projections of the enhanced and IsoNet missing-wedge-corrected particle. The sixth and seventh columns present the final 3D map and the flexibly fitted model, respectively. (B) Two cross-sectional views (12 nm thickness) of the plasmid density map along its plectoneme axis are shown in the left-middle panel. The intensity profile along the line crossing the two high-density DNA spots is displayed in the right panel. (C) Resolution assessment of the final 3D map using Fourier shell correlation (FSC). Two criteria are shown: FSC between two half-maps reconstructed from even and odd frames (evaluated at 0.143) and FSC between the final 3D map and the fitted model (evaluated at 0.5). (D) Zoomed-in views of the final 3D density map from panel A, displayed at two contour levels. (E) Superimposition of the high-contour level map from panel D onto its fitted model.

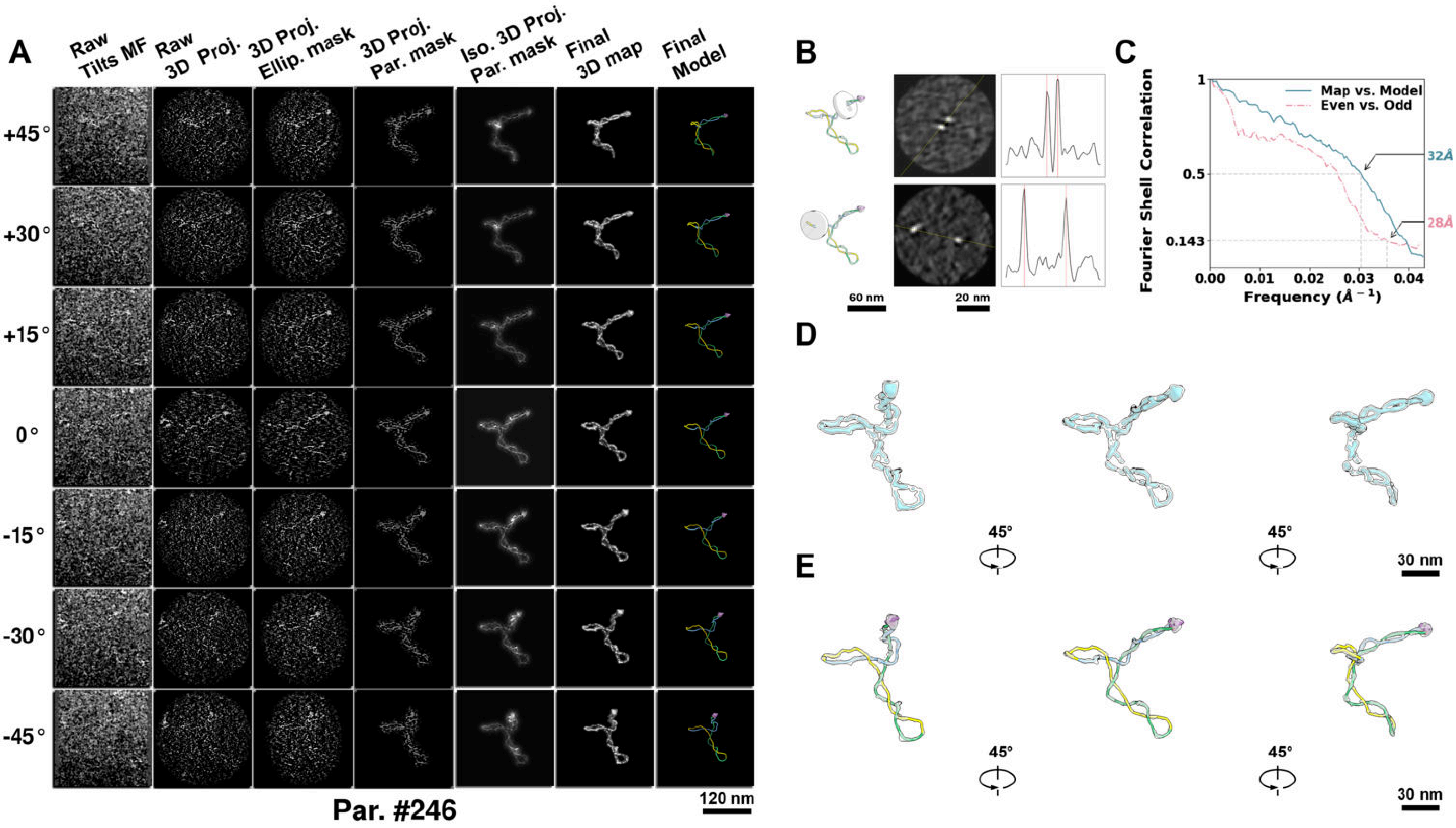

**Supplementary Particle Figure 246. Cryo-ET 3D reconstruction of an individual TEC-TopI particle.**
(A) 3D reconstruction of the plasmid particle (index no. 246). The first column shows seven representative tilt images from +45° to -45° in step of 15°. The second, third, and fourth columns show 3D projections of the particle with spherical, ellipsoidal (thinner along the z-dimension), and particle-shaped masks, respectively. The fifth column displays the 3D projections of the enhanced and IsoNet missing-wedge-corrected particle. The sixth and seventh columns present the final 3D map and the flexibly fitted model, respectively. (B) Two cross-sectional views (12 nm thickness) of the plasmid density map along its plectoneme axis are shown in the left-middle panel. The intensity profile along the line crossing the two high-density DNA spots is displayed in the right panel. (C) Resolution assessment of the final 3D map using Fourier shell correlation (FSC). Two criteria are shown: FSC between two half-maps reconstructed from even and odd frames (evaluated at 0.143) and FSC between the final 3D map and the fitted model (evaluated at 0.5). (D) Zoomed-in views of the final 3D density map from panel A, displayed at two contour levels. (E) Superimposition of the high-contour level map from panel D onto its fitted model.

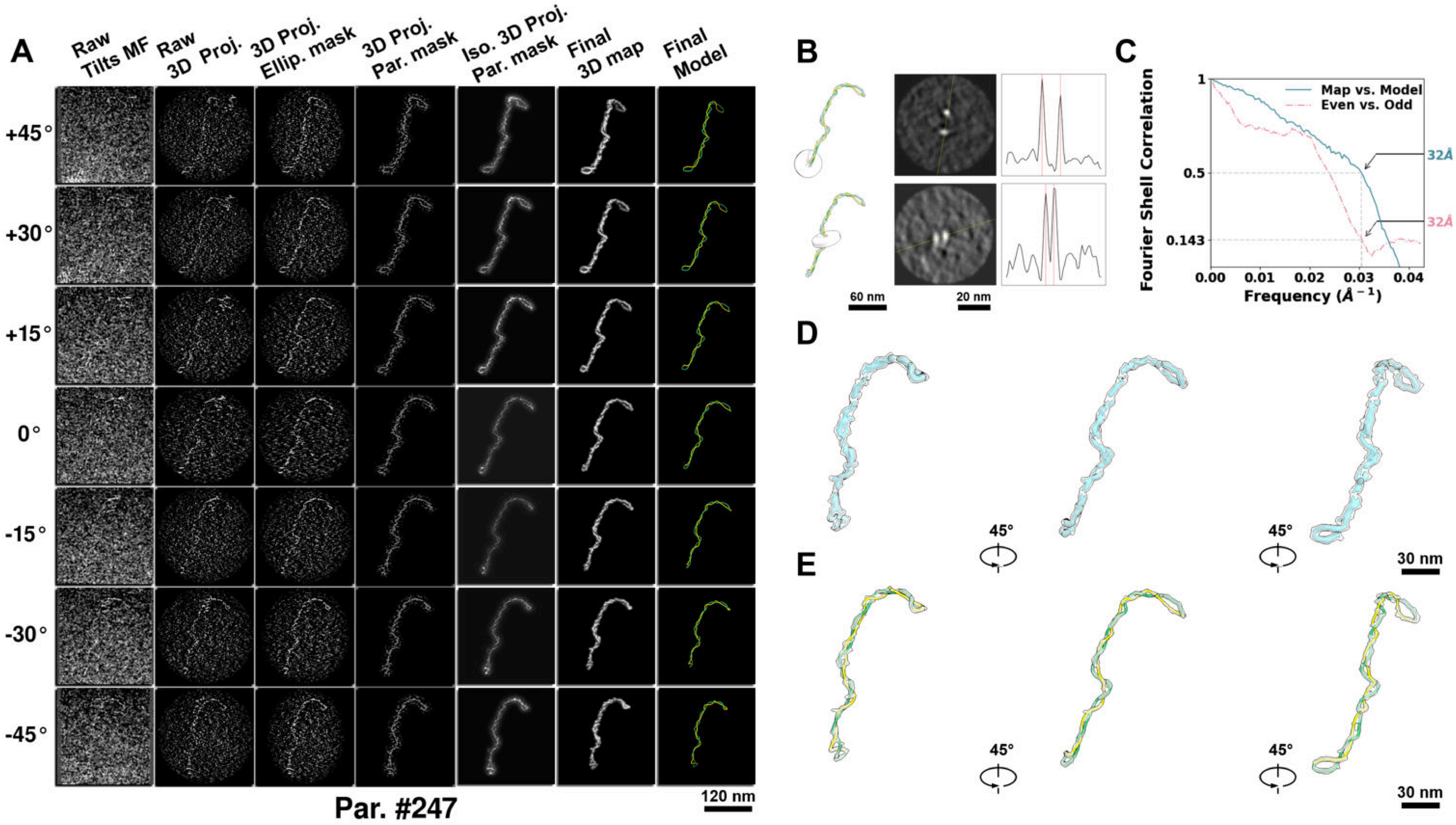

**Supplementary Particle Figure 247. Cryo-ET 3D reconstruction of an individual TEC-TopI particle.**
(A) 3D reconstruction of the plasmid particle (index no. 247). The first column shows seven representative tilt images from +45° to -45° in step of 15°. The second, third, and fourth columns show 3D projections of the particle with spherical, ellipsoidal (thinner along the z-dimension), and particle-shaped masks, respectively. The fifth column displays the 3D projections of the enhanced and IsoNet missing-wedge-corrected particle. The sixth and seventh columns present the final 3D map and the flexibly fitted model, respectively. (B) Two cross-sectional views (12 nm thickness) of the plasmid density map along its plectoneme axis are shown in the left-middle panel. The intensity profile along the line crossing the two high-density DNA spots is displayed in the right panel. (C) Resolution assessment of the final 3D map using Fourier shell correlation (FSC). Two criteria are shown: FSC between two half-maps reconstructed from even and odd frames (evaluated at 0.143) and FSC between the final 3D map and the fitted model (evaluated at 0.5). (D) Zoomed-in views of the final 3D density map from panel A, displayed at two contour levels. (E) Superimposition of the high-contour level map from panel D onto its fitted model.

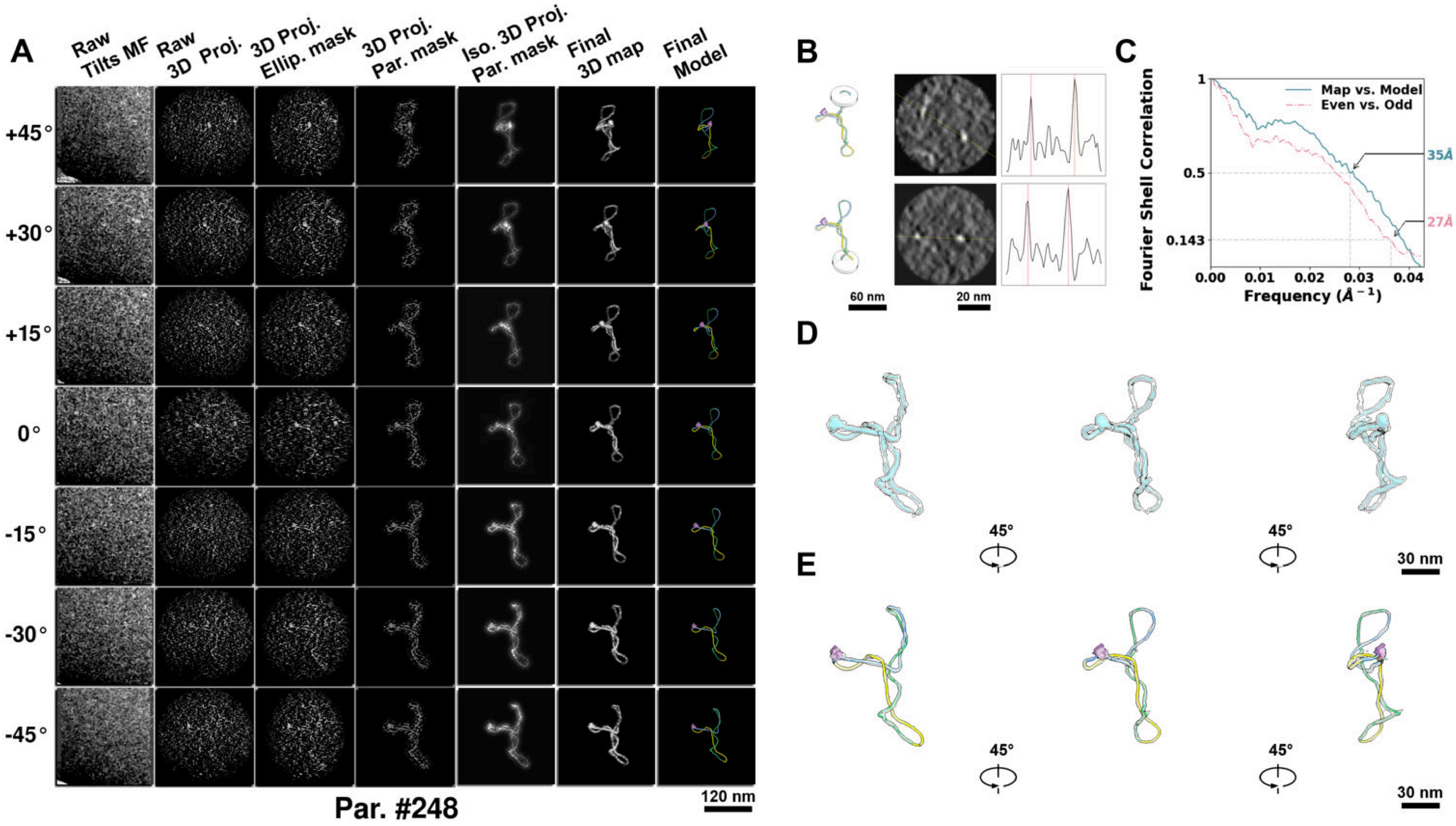

**Supplementary Particle Figure 248. Cryo-ET 3D reconstruction of an individual TEC-TopI particle.**
(A) 3D reconstruction of the plasmid particle (index no. 248). The first column shows seven representative tilt images from +45° to -45° in step of 15°. The second, third, and fourth columns show 3D projections of the particle with spherical, ellipsoidal (thinner along the z-dimension), and particle-shaped masks, respectively. The fifth column displays the 3D projections of the enhanced and IsoNet missing-wedge-corrected particle. The sixth and seventh columns present the final 3D map and the flexibly fitted model, respectively. (B) Two cross-sectional views (12 nm thickness) of the plasmid density map along its plectoneme axis are shown in the left-middle panel. The intensity profile along the line crossing the two high-density DNA spots is displayed in the right panel. (C) Resolution assessment of the final 3D map using Fourier shell correlation (FSC). Two criteria are shown: FSC between two half-maps reconstructed from even and odd frames (evaluated at 0.143) and FSC between the final 3D map and the fitted model (evaluated at 0.5). (D) Zoomed-in views of the final 3D density map from panel A, displayed at two contour levels. (E) Superimposition of the high-contour level map from panel D onto its fitted model.

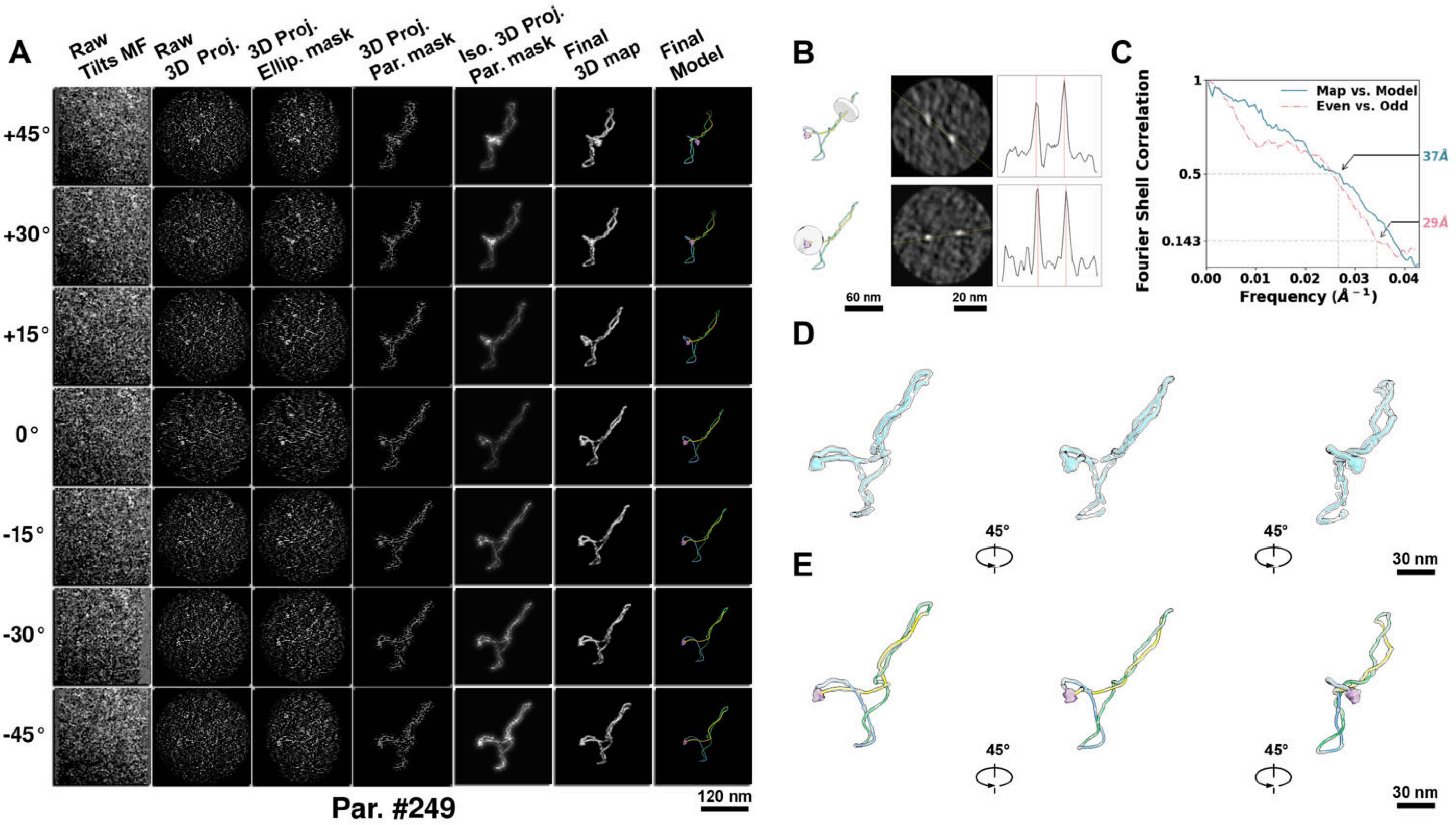

**Supplementary Particle Figure 249. Cryo-ET 3D reconstruction of an individual TEC-TopI particle.**
(A) 3D reconstruction of the plasmid particle (index no. 249). The first column shows seven representative tilt images from +45° to -45° in step of 15°. The second, third, and fourth columns show 3D projections of the particle with spherical, ellipsoidal (thinner along the z-dimension), and particle-shaped masks, respectively. The fifth column displays the 3D projections of the enhanced and IsoNet missing-wedge-corrected particle. The sixth and seventh columns present the final 3D map and the flexibly fitted model, respectively. (B) Two cross-sectional views (12 nm thickness) of the plasmid density map along its plectoneme axis are shown in the left-middle panel. The intensity profile along the line crossing the two high-density DNA spots is displayed in the right panel. (C) Resolution assessment of the final 3D map using Fourier shell correlation (FSC). Two criteria are shown: FSC between two half-maps reconstructed from even and odd frames (evaluated at 0.143) and FSC between the final 3D map and the fitted model (evaluated at 0.5). (D) Zoomed-in views of the final 3D density map from panel A, displayed at two contour levels. (E) Superimposition of the high-contour level map from panel D onto its fitted model.

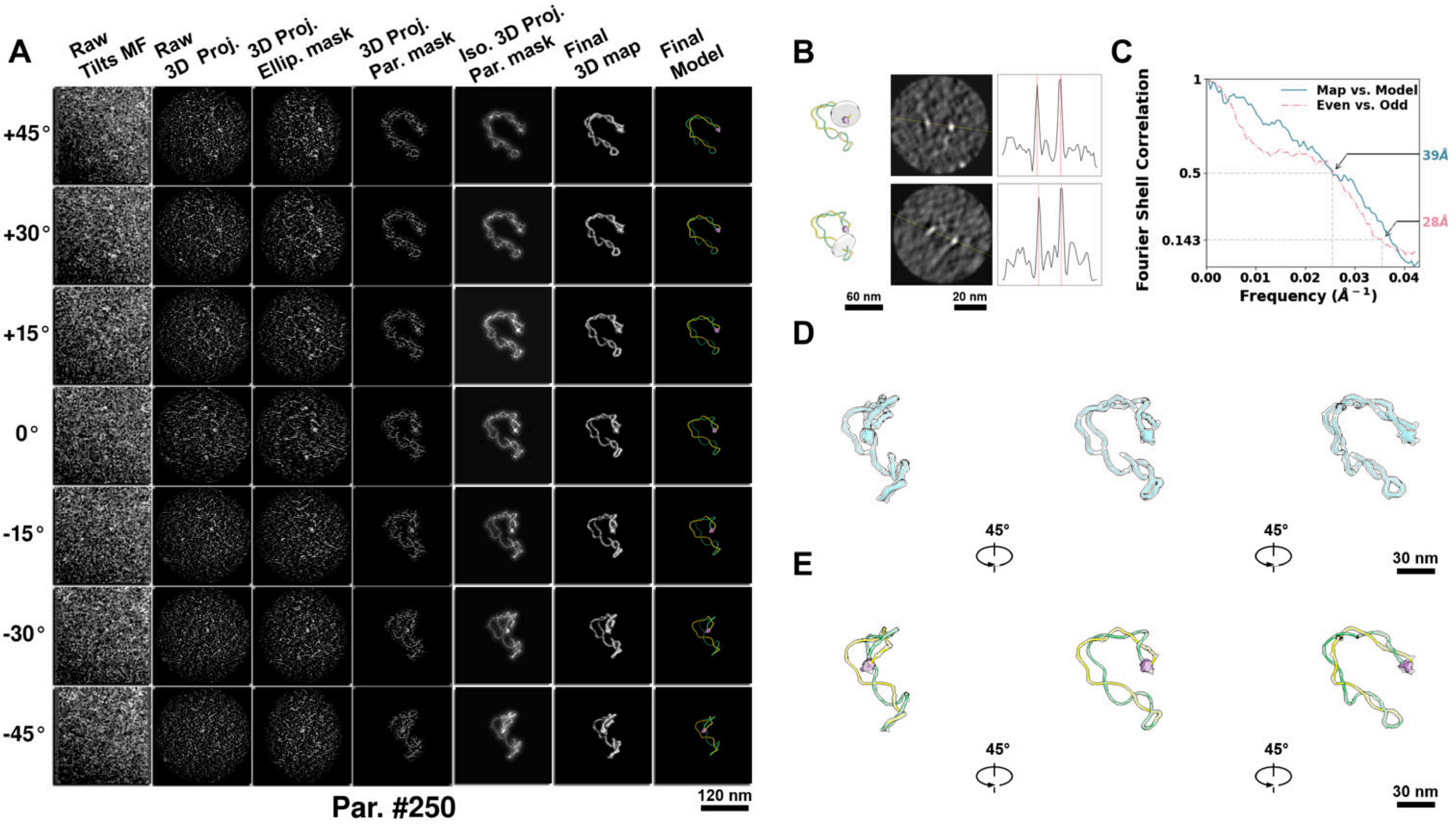

**Supplementary Particle Figure 250. Cryo-ET 3D reconstruction of an individual TEC-TopI particle.**
(A) 3D reconstruction of the plasmid particle (index no. 250). The first column shows seven representative tilt images from +45° to -45° in step of 15°. The second, third, and fourth columns show 3D projections of the particle with spherical, ellipsoidal (thinner along the z-dimension), and particle-shaped masks, respectively. The fifth column displays the 3D projections of the enhanced and IsoNet missing-wedge-corrected particle. The sixth and seventh columns present the final 3D map and the flexibly fitted model, respectively. (B) Two cross-sectional views (12 nm thickness) of the plasmid density map along its plectoneme axis are shown in the left-middle panel. The intensity profile along the line crossing the two high-density DNA spots is displayed in the right panel. (C) Resolution assessment of the final 3D map using Fourier shell correlation (FSC). Two criteria are shown: FSC between two half-maps reconstructed from even and odd frames (evaluated at 0.143) and FSC between the final 3D map and the fitted model (evaluated at 0.5). (D) Zoomed-in views of the final 3D density map from panel A, displayed at two contour levels. (E) Superimposition of the high-contour level map from panel D onto its fitted model.

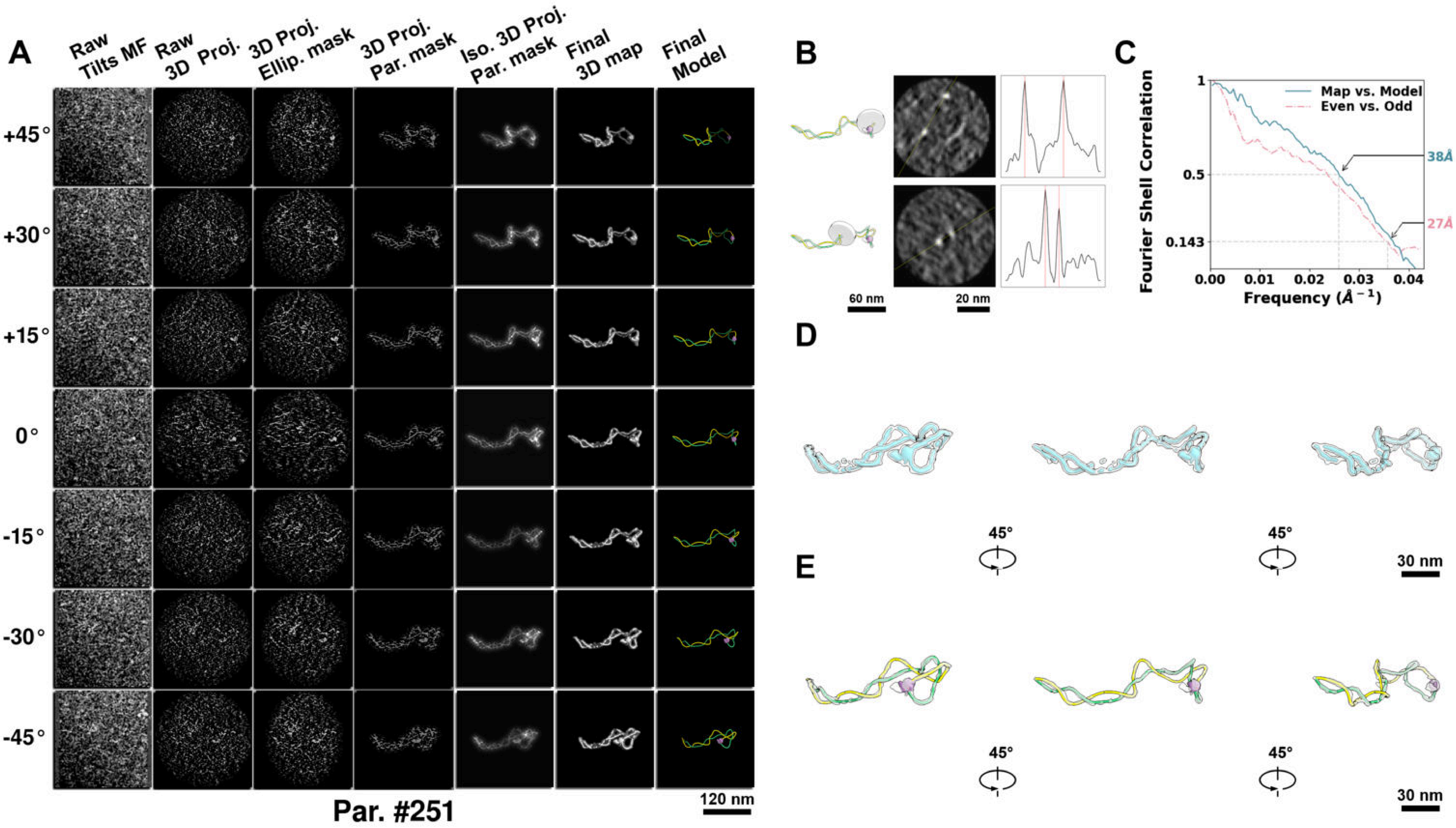

**Supplementary Particle Figure 251. Cryo-ET 3D reconstruction of an individual TEC-TopI particle.**
(A) 3D reconstruction of the plasmid particle (index no. 251). The first column shows seven representative tilt images from +45° to -45° in step of 15°. The second, third, and fourth columns show 3D projections of the particle with spherical, ellipsoidal (thinner along the z-dimension), and particle-shaped masks, respectively. The fifth column displays the 3D projections of the enhanced and IsoNet missing-wedge-corrected particle. The sixth and seventh columns present the final 3D map and the flexibly fitted model, respectively. (B) Two cross-sectional views (12 nm thickness) of the plasmid density map along its plectoneme axis are shown in the left-middle panel. The intensity profile along the line crossing the two high-density DNA spots is displayed in the right panel. (C) Resolution assessment of the final 3D map using Fourier shell correlation (FSC). Two criteria are shown: FSC between two half-maps reconstructed from even and odd frames (evaluated at 0.143) and FSC between the final 3D map and the fitted model (evaluated at 0.5). (D) Zoomed-in views of the final 3D density map from panel A, displayed at two contour levels. (E) Superimposition of the high-contour level map from panel D onto its fitted model.

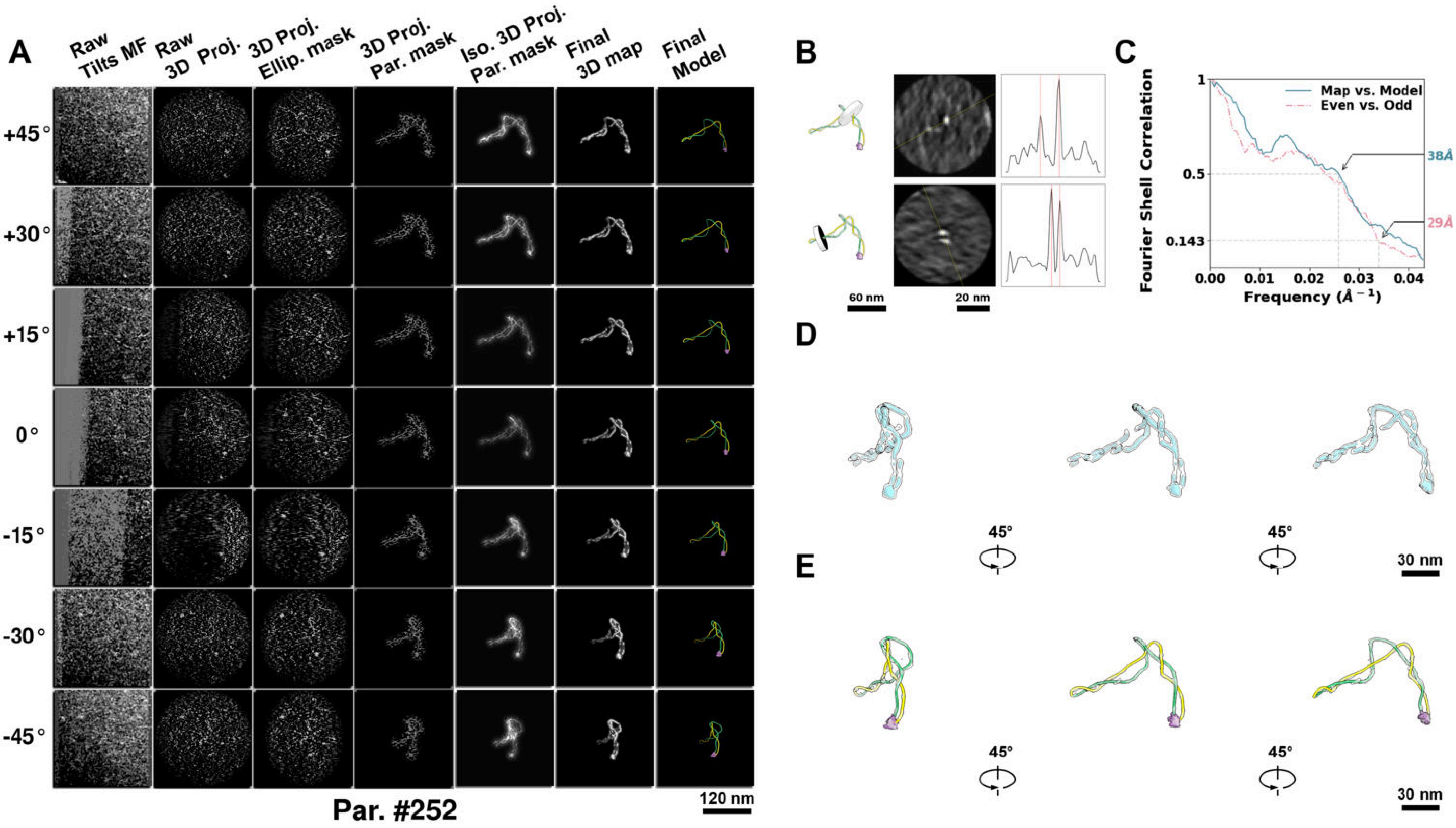

**Supplementary Particle Figure 252. Cryo-ET 3D reconstruction of an individual TEC-TopI particle.**
(A) 3D reconstruction of the plasmid particle (index no. 252). The first column shows seven representative tilt images from +45° to -45° in step of 15°. The second, third, and fourth columns show 3D projections of the particle with spherical, ellipsoidal (thinner along the z-dimension), and particle-shaped masks, respectively. The fifth column displays the 3D projections of the enhanced and IsoNet missing-wedge-corrected particle. The sixth and seventh columns present the final 3D map and the flexibly fitted model, respectively. (B) Two cross-sectional views (12 nm thickness) of the plasmid density map along its plectoneme axis are shown in the left-middle panel. The intensity profile along the line crossing the two high-density DNA spots is displayed in the right panel. (C) Resolution assessment of the final 3D map using Fourier shell correlation (FSC). Two criteria are shown: FSC between two half-maps reconstructed from even and odd frames (evaluated at 0.143) and FSC between the final 3D map and the fitted model (evaluated at 0.5). (D) Zoomed-in views of the final 3D density map from panel A, displayed at two contour levels. (E) Superimposition of the high-contour level map from panel D onto its fitted model.

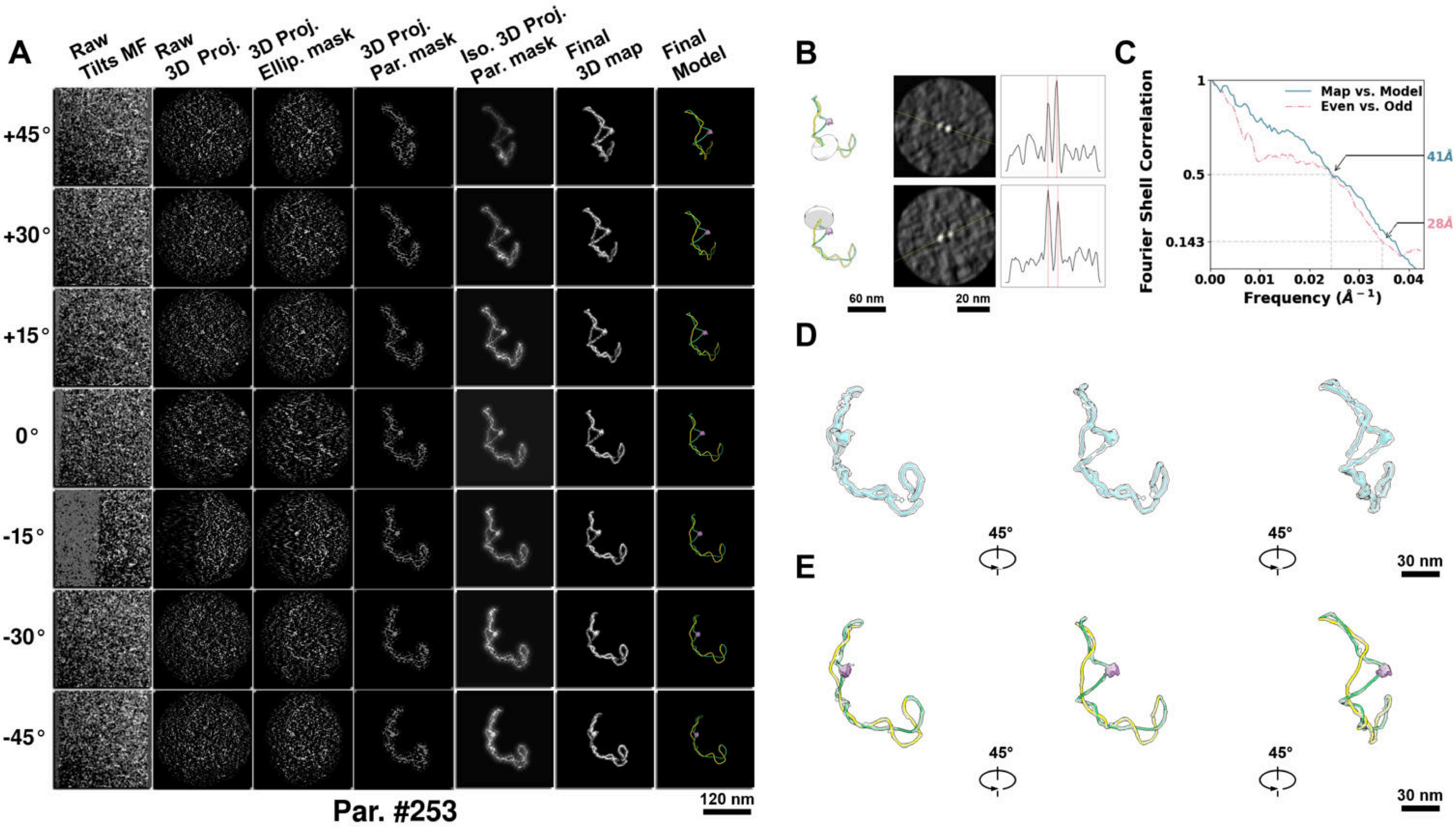

**Supplementary Particle Figure 253. Cryo-ET 3D reconstruction of an individual TEC-TopI particle.**
(A) 3D reconstruction of the plasmid particle (index no. 253). The first column shows seven representative tilt images from +45° to -45° in step of 15°. The second, third, and fourth columns show 3D projections of the particle with spherical, ellipsoidal (thinner along the z-dimension), and particle-shaped masks, respectively. The fifth column displays the 3D projections of the enhanced and IsoNet missing-wedge-corrected particle. The sixth and seventh columns present the final 3D map and the flexibly fitted model, respectively. (B) Two cross-sectional views (12 nm thickness) of the plasmid density map along its plectoneme axis are shown in the left-middle panel. The intensity profile along the line crossing the two high-density DNA spots is displayed in the right panel. (C) Resolution assessment of the final 3D map using Fourier shell correlation (FSC). Two criteria are shown: FSC between two half-maps reconstructed from even and odd frames (evaluated at 0.143) and FSC between the final 3D map and the fitted model (evaluated at 0.5). (D) Zoomed-in views of the final 3D density map from panel A, displayed at two contour levels. (E) Superimposition of the high-contour level map from panel D onto its fitted model.

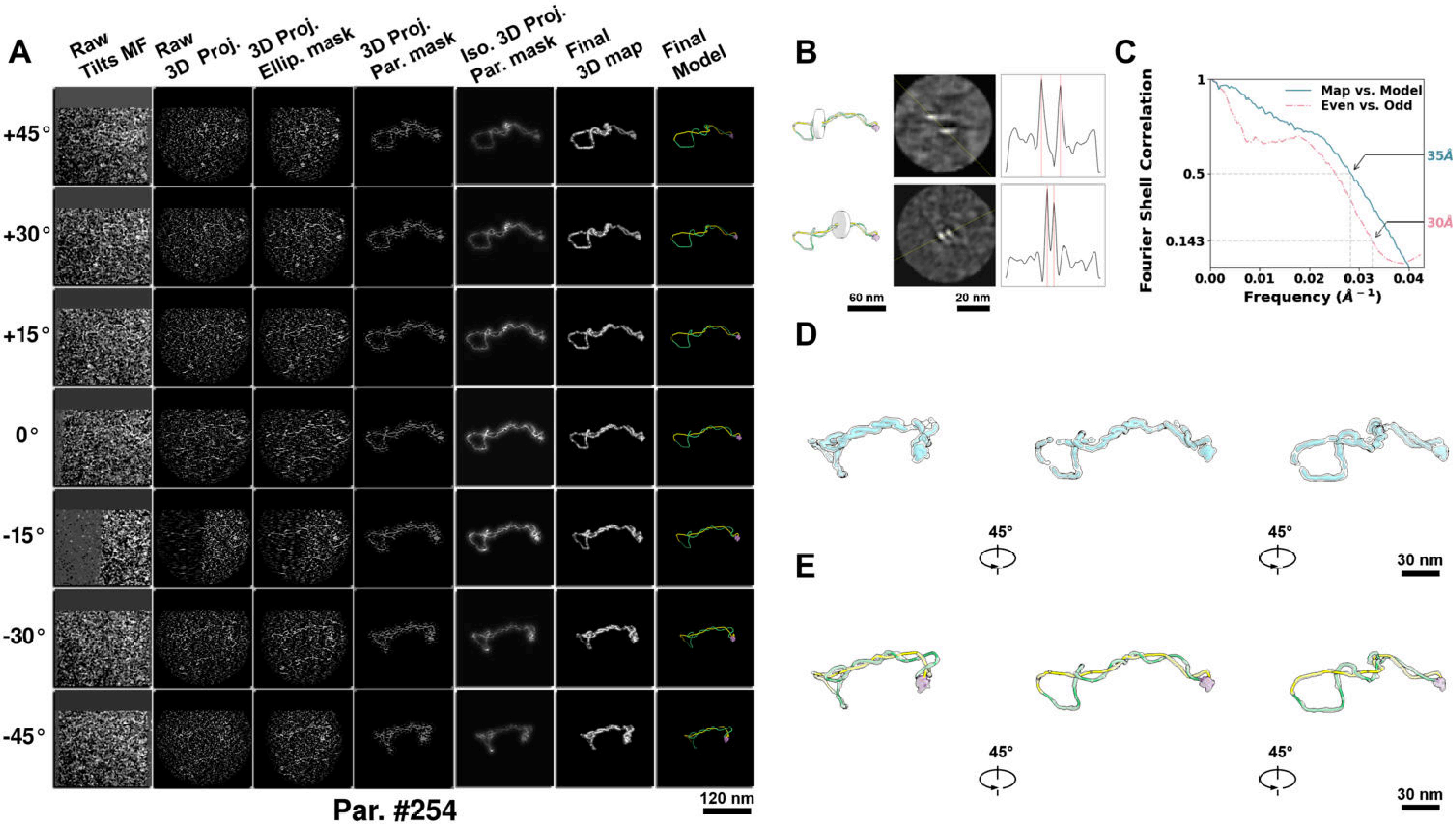

**Supplementary Particle Figure 254. Cryo-ET 3D reconstruction of an individual TEC-TopI particle.**
(A) 3D reconstruction of the plasmid particle (index no. 254). The first column shows seven representative tilt images from +45° to -45° in step of 15°. The second, third, and fourth columns show 3D projections of the particle with spherical, ellipsoidal (thinner along the z-dimension), and particle-shaped masks, respectively. The fifth column displays the 3D projections of the enhanced and IsoNet missing-wedge-corrected particle. The sixth and seventh columns present the final 3D map and the flexibly fitted model, respectively. (B) Two cross-sectional views (12 nm thickness) of the plasmid density map along its plectoneme axis are shown in the left-middle panel. The intensity profile along the line crossing the two high-density DNA spots is displayed in the right panel. (C) Resolution assessment of the final 3D map using Fourier shell correlation (FSC). Two criteria are shown: FSC between two half-maps reconstructed from even and odd frames (evaluated at 0.143) and FSC between the final 3D map and the fitted model (evaluated at 0.5). (D) Zoomed-in views of the final 3D density map from panel A, displayed at two contour levels. (E) Superimposition of the high-contour level map from panel D onto its fitted model.

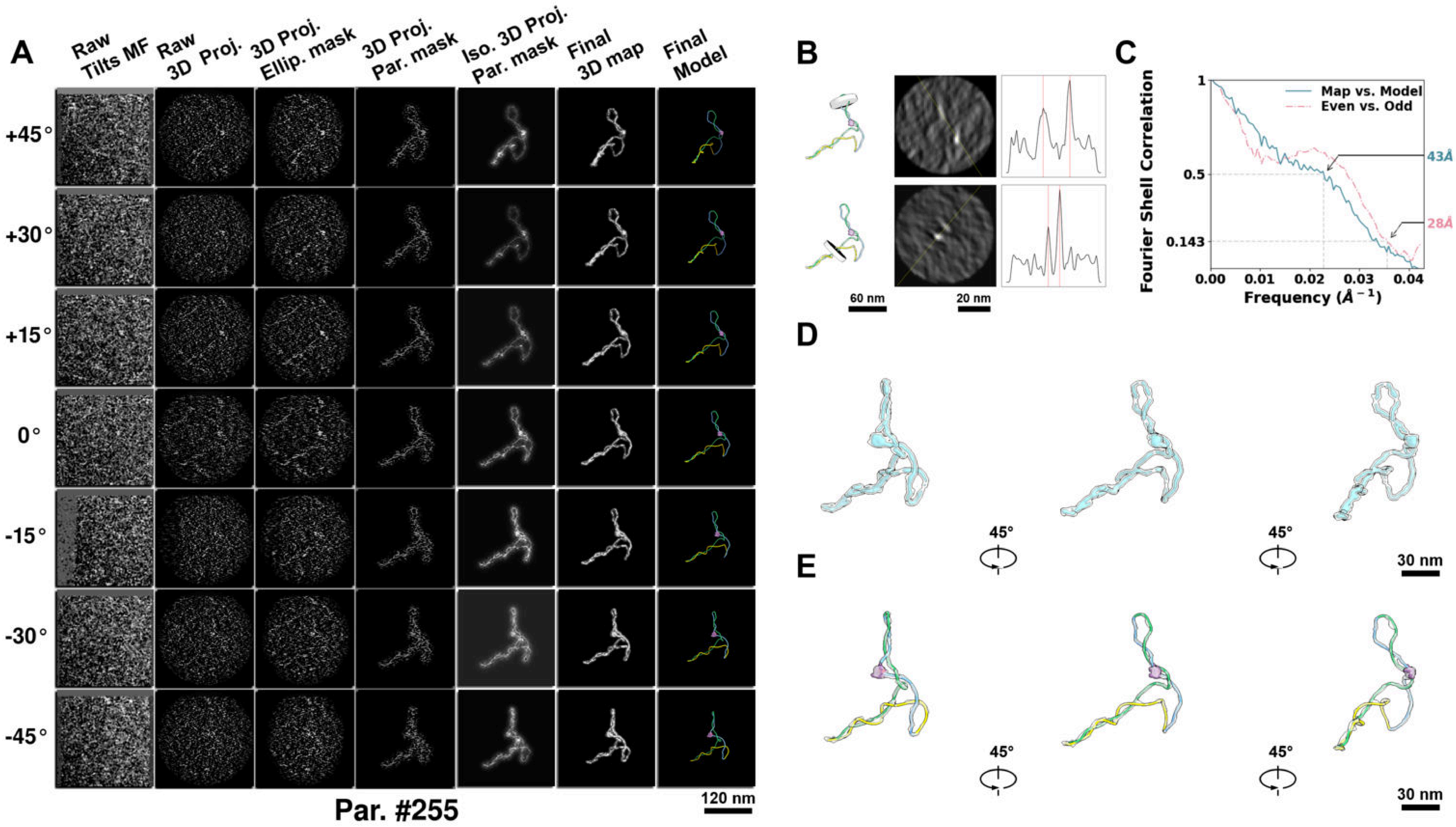

**Supplementary Particle Figure 255. Cryo-ET 3D reconstruction of an individual TEC-TopI particle.**
(A) 3D reconstruction of the plasmid particle (index no. 255). The first column shows seven representative tilt images from +45° to -45° in step of 15°. The second, third, and fourth columns show 3D projections of the particle with spherical, ellipsoidal (thinner along the z-dimension), and particle-shaped masks, respectively. The fifth column displays the 3D projections of the enhanced and IsoNet missing-wedge-corrected particle. The sixth and seventh columns present the final 3D map and the flexibly fitted model, respectively. (B) Two cross-sectional views (12 nm thickness) of the plasmid density map along its plectoneme axis are shown in the left-middle panel. The intensity profile along the line crossing the two high-density DNA spots is displayed in the right panel. (C) Resolution assessment of the final 3D map using Fourier shell correlation (FSC). Two criteria are shown: FSC between two half-maps reconstructed from even and odd frames (evaluated at 0.143) and FSC between the final 3D map and the fitted model (evaluated at 0.5). (D) Zoomed-in views of the final 3D density map from panel A, displayed at two contour levels. (E) Superimposition of the high-contour level map from panel D onto its fitted model.

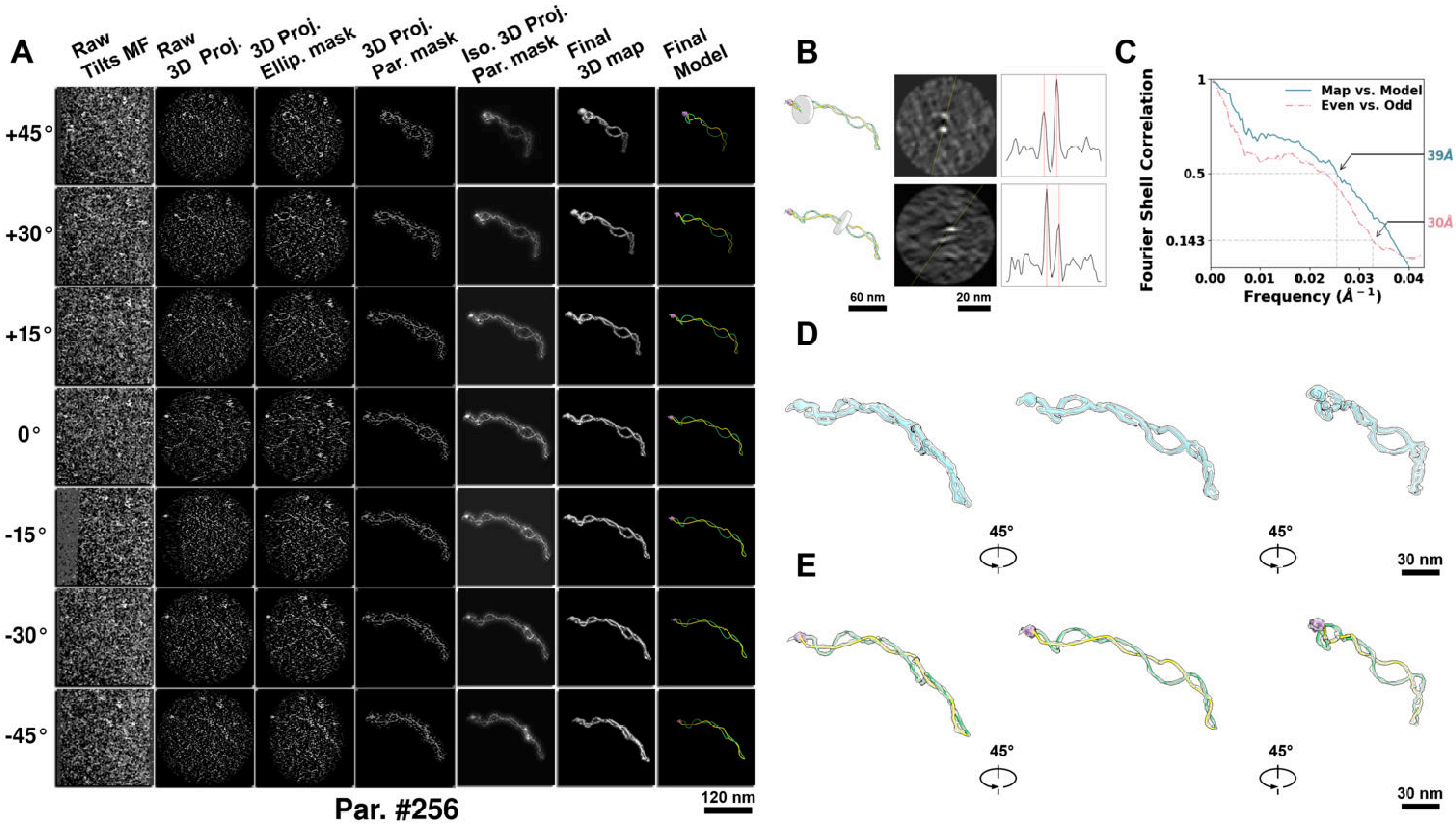

**Supplementary Particle Figure 256. Cryo-ET 3D reconstruction of an individual TEC-TopI particle.**
(A) 3D reconstruction of the plasmid particle (index no. 256). The first column shows seven representative tilt images from +45° to -45° in step of 15°. The second, third, and fourth columns show 3D projections of the particle with spherical, ellipsoidal (thinner along the z-dimension), and particle-shaped masks, respectively. The fifth column displays the 3D projections of the enhanced and IsoNet missing-wedge-corrected particle. The sixth and seventh columns present the final 3D map and the flexibly fitted model, respectively. (B) Two cross-sectional views (12 nm thickness) of the plasmid density map along its plectoneme axis are shown in the left-middle panel. The intensity profile along the line crossing the two high-density DNA spots is displayed in the right panel. (C) Resolution assessment of the final 3D map using Fourier shell correlation (FSC). Two criteria are shown: FSC between two half-maps reconstructed from even and odd frames (evaluated at 0.143) and FSC between the final 3D map and the fitted model (evaluated at 0.5). (D) Zoomed-in views of the final 3D density map from panel A, displayed at two contour levels. (E) Superimposition of the high-contour level map from panel D onto its fitted model.

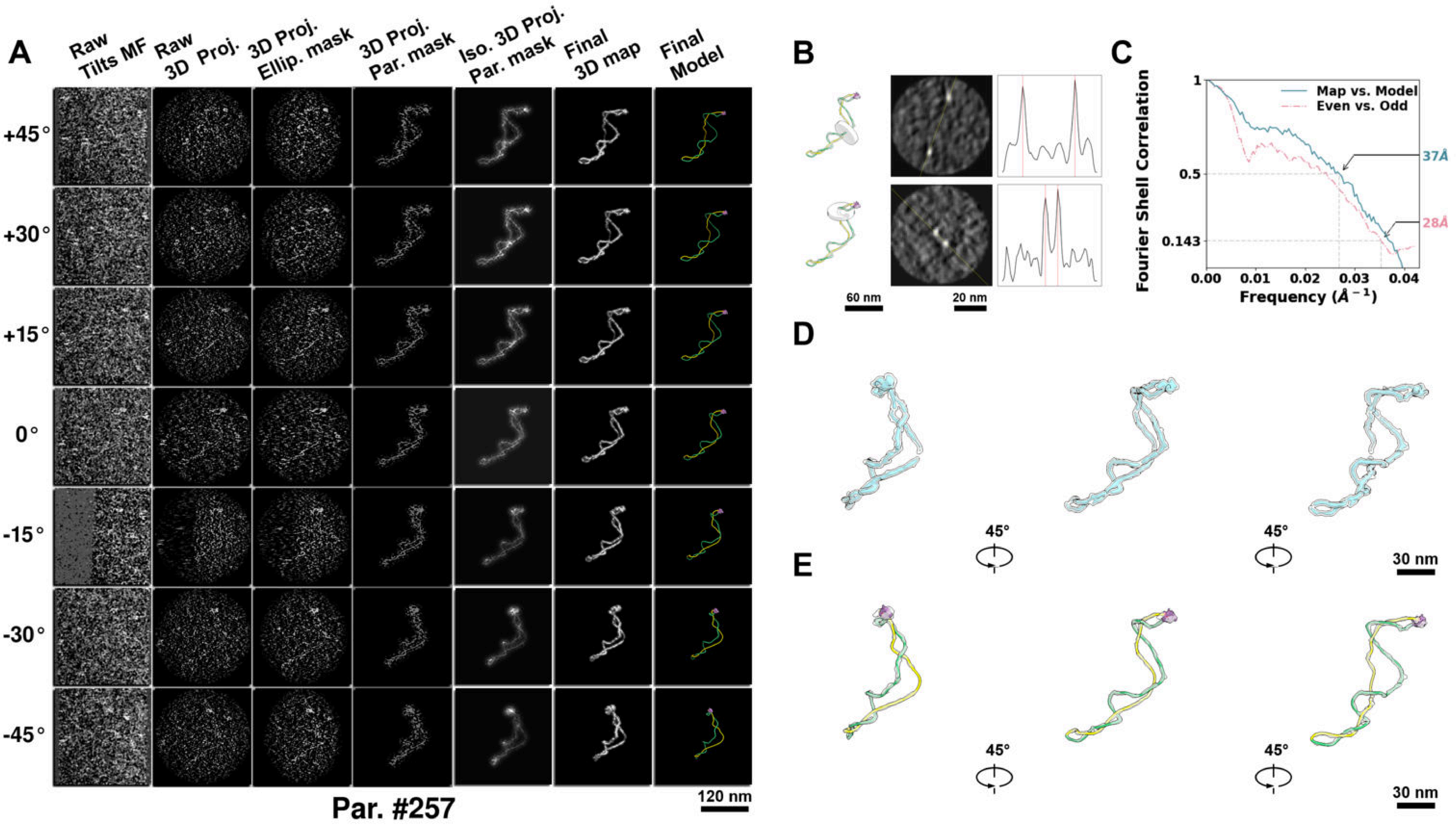

**Supplementary Particle Figure 257. Cryo-ET 3D reconstruction of an individual TEC-TopI particle.**
(A) 3D reconstruction of the plasmid particle (index no. 257). The first column shows seven representative tilt images from +45° to -45° in step of 15°. The second, third, and fourth columns show 3D projections of the particle with spherical, ellipsoidal (thinner along the z-dimension), and particle-shaped masks, respectively. The fifth column displays the 3D projections of the enhanced and IsoNet missing-wedge-corrected particle. The sixth and seventh columns present the final 3D map and the flexibly fitted model, respectively. (B) Two cross-sectional views (12 nm thickness) of the plasmid density map along its plectoneme axis are shown in the left-middle panel. The intensity profile along the line crossing the two high-density DNA spots is displayed in the right panel. (C) Resolution assessment of the final 3D map using Fourier shell correlation (FSC). Two criteria are shown: FSC between two half-maps reconstructed from even and odd frames (evaluated at 0.143) and FSC between the final 3D map and the fitted model (evaluated at 0.5). (D) Zoomed-in views of the final 3D density map from panel A, displayed at two contour levels. (E) Superimposition of the high-contour level map from panel D onto its fitted model.

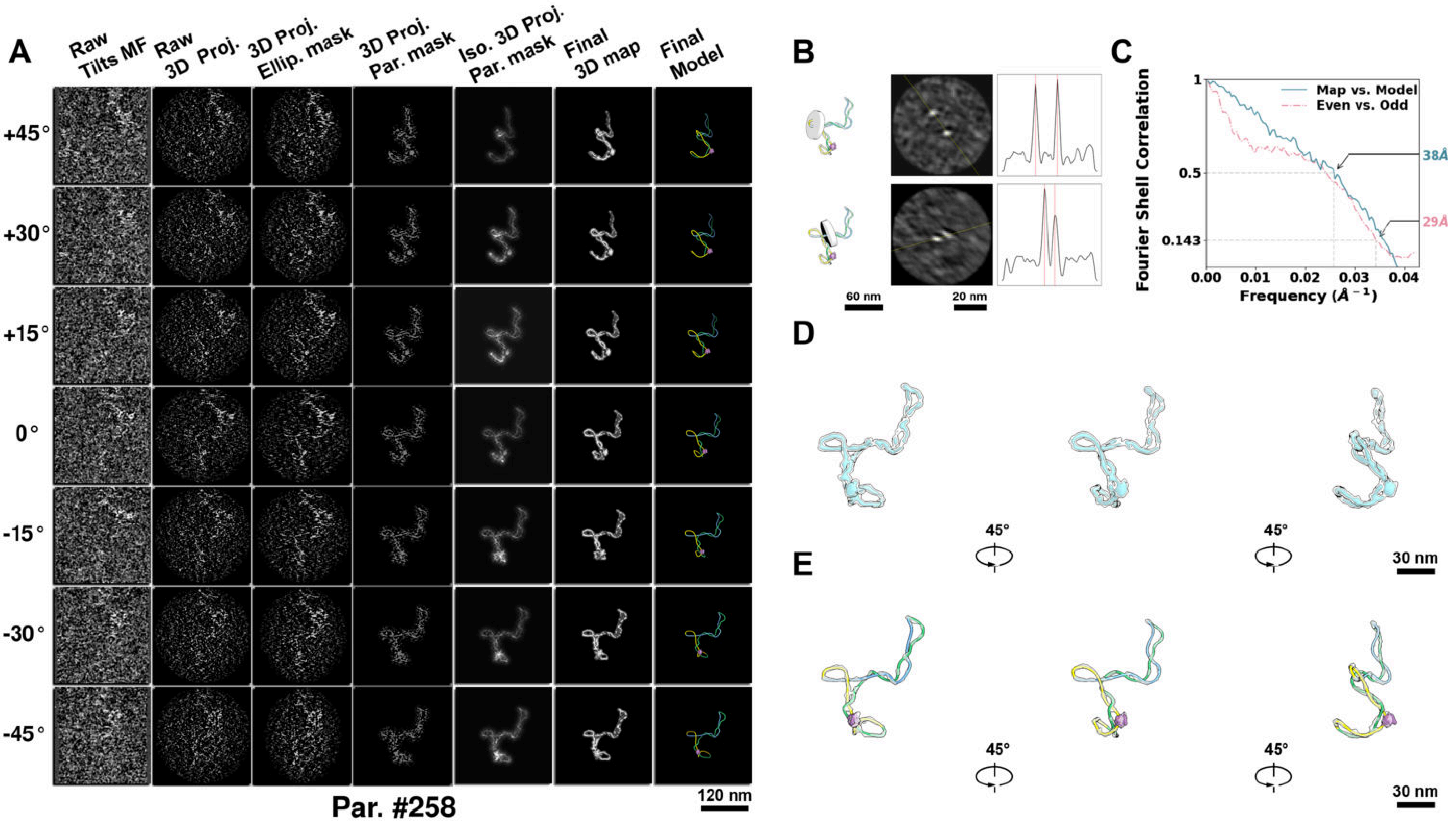

**Supplementary Particle Figure 258. Cryo-ET 3D reconstruction of an individual TEC-TopI particle.**
(A) 3D reconstruction of the plasmid particle (index no. 258). The first column shows seven representative tilt images from +45° to -45° in step of 15°. The second, third, and fourth columns show 3D projections of the particle with spherical, ellipsoidal (thinner along the z-dimension), and particle-shaped masks, respectively. The fifth column displays the 3D projections of the enhanced and IsoNet missing-wedge-corrected particle. The sixth and seventh columns present the final 3D map and the flexibly fitted model, respectively. (B) Two cross-sectional views (12 nm thickness) of the plasmid density map along its plectoneme axis are shown in the left-middle panel. The intensity profile along the line crossing the two high-density DNA spots is displayed in the right panel. (C) Resolution assessment of the final 3D map using Fourier shell correlation (FSC). Two criteria are shown: FSC between two half-maps reconstructed from even and odd frames (evaluated at 0.143) and FSC between the final 3D map and the fitted model (evaluated at 0.5). (D) Zoomed-in views of the final 3D density map from panel A, displayed at two contour levels. (E) Superimposition of the high-contour level map from panel D onto its fitted model.

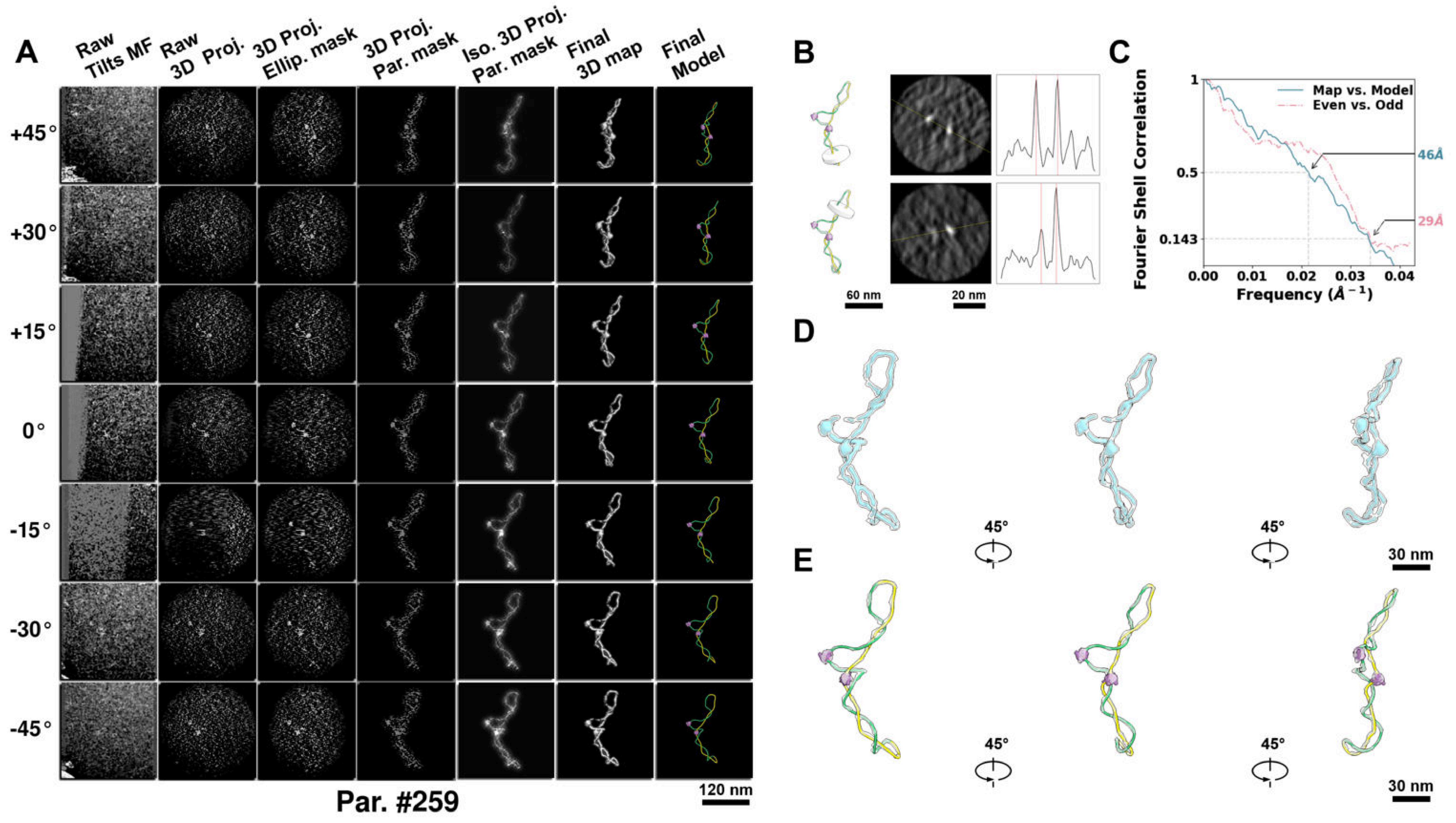

**Supplementary Particle Figure 259. Cryo-ET 3D reconstruction of an individual TEC-TopI particle.**
(A) 3D reconstruction of the plasmid particle (index no. 259). The first column shows seven representative tilt images from +45° to -45° in step of 15°. The second, third, and fourth columns show 3D projections of the particle with spherical, ellipsoidal (thinner along the z-dimension), and particle-shaped masks, respectively. The fifth column displays the 3D projections of the enhanced and IsoNet missing-wedge-corrected particle. The sixth and seventh columns present the final 3D map and the flexibly fitted model, respectively. (B) Two cross-sectional views (12 nm thickness) of the plasmid density map along its plectoneme axis are shown in the left-middle panel. The intensity profile along the line crossing the two high-density DNA spots is displayed in the right panel. (C) Resolution assessment of the final 3D map using Fourier shell correlation (FSC). Two criteria are shown: FSC between two half-maps reconstructed from even and odd frames (evaluated at 0.143) and FSC between the final 3D map and the fitted model (evaluated at 0.5). (D) Zoomed-in views of the final 3D density map from panel A, displayed at two contour levels. (E) Superimposition of the high-contour level map from panel D onto its fitted model.

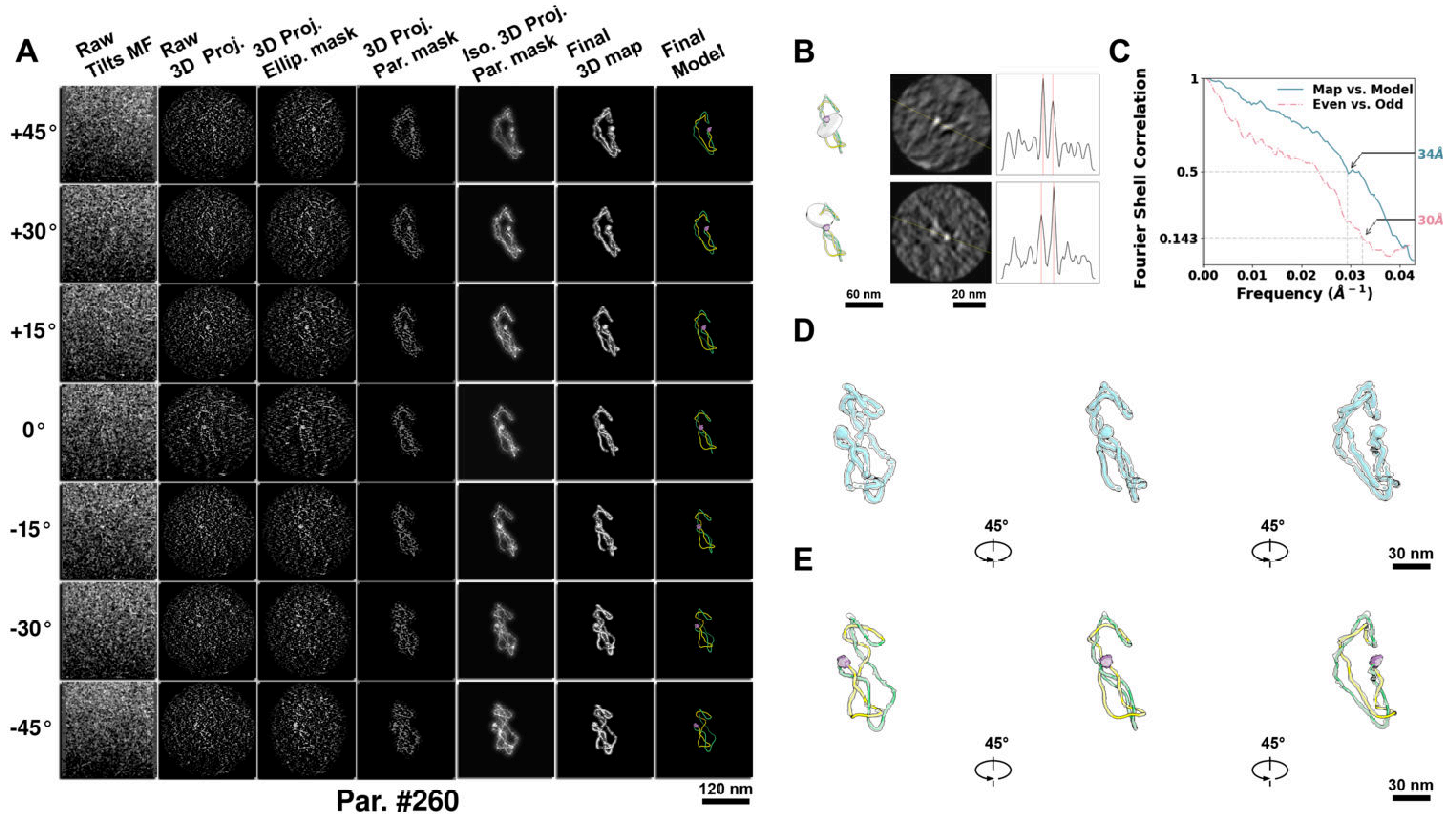

**Supplementary Particle Figure 260. Cryo-ET 3D reconstruction of an individual TEC-TopI particle.**
(A) 3D reconstruction of the plasmid particle (index no. 260). The first column shows seven representative tilt images from +45° to -45° in step of 15°. The second, third, and fourth columns show 3D projections of the particle with spherical, ellipsoidal (thinner along the z-dimension), and particle-shaped masks, respectively. The fifth column displays the 3D projections of the enhanced and IsoNet missing-wedge-corrected particle. The sixth and seventh columns present the final 3D map and the flexibly fitted model, respectively. (B) Two cross-sectional views (12 nm thickness) of the plasmid density map along its plectoneme axis are shown in the left-middle panel. The intensity profile along the line crossing the two high-density DNA spots is displayed in the right panel. (C) Resolution assessment of the final 3D map using Fourier shell correlation (FSC). Two criteria are shown: FSC between two half-maps reconstructed from even and odd frames (evaluated at 0.143) and FSC between the final 3D map and the fitted model (evaluated at 0.5). (D) Zoomed-in views of the final 3D density map from panel A, displayed at two contour levels. (E) Superimposition of the high-contour level map from panel D onto its fitted model.

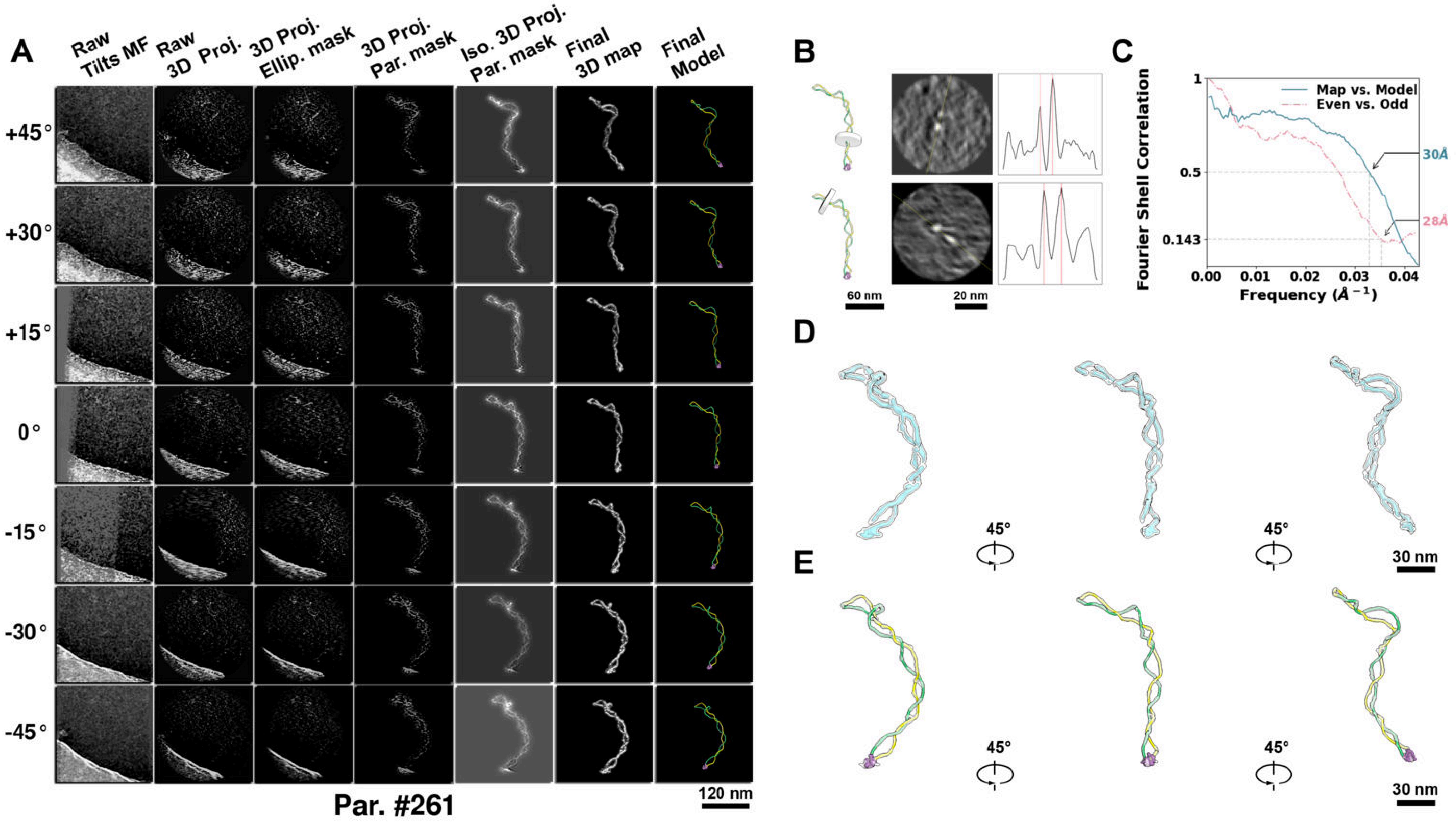

**Supplementary Particle Figure 261. Cryo-ET 3D reconstruction of an individual TEC-TopI particle.**
(A) 3D reconstruction of the plasmid particle (index no. 261). The first column shows seven representative tilt images from +45° to -45° in step of 15°. The second, third, and fourth columns show 3D projections of the particle with spherical, ellipsoidal (thinner along the z-dimension), and particle-shaped masks, respectively. The fifth column displays the 3D projections of the enhanced and IsoNet missing-wedge-corrected particle. The sixth and seventh columns present the final 3D map and the flexibly fitted model, respectively. (B) Two cross-sectional views (12 nm thickness) of the plasmid density map along its plectoneme axis are shown in the left-middle panel. The intensity profile along the line crossing the two high-density DNA spots is displayed in the right panel. (C) Resolution assessment of the final 3D map using Fourier shell correlation (FSC). Two criteria are shown: FSC between two half-maps reconstructed from even and odd frames (evaluated at 0.143) and FSC between the final 3D map and the fitted model (evaluated at 0.5). (D) Zoomed-in views of the final 3D density map from panel A, displayed at two contour levels. (E) Superimposition of the high-contour level map from panel D onto its fitted model.

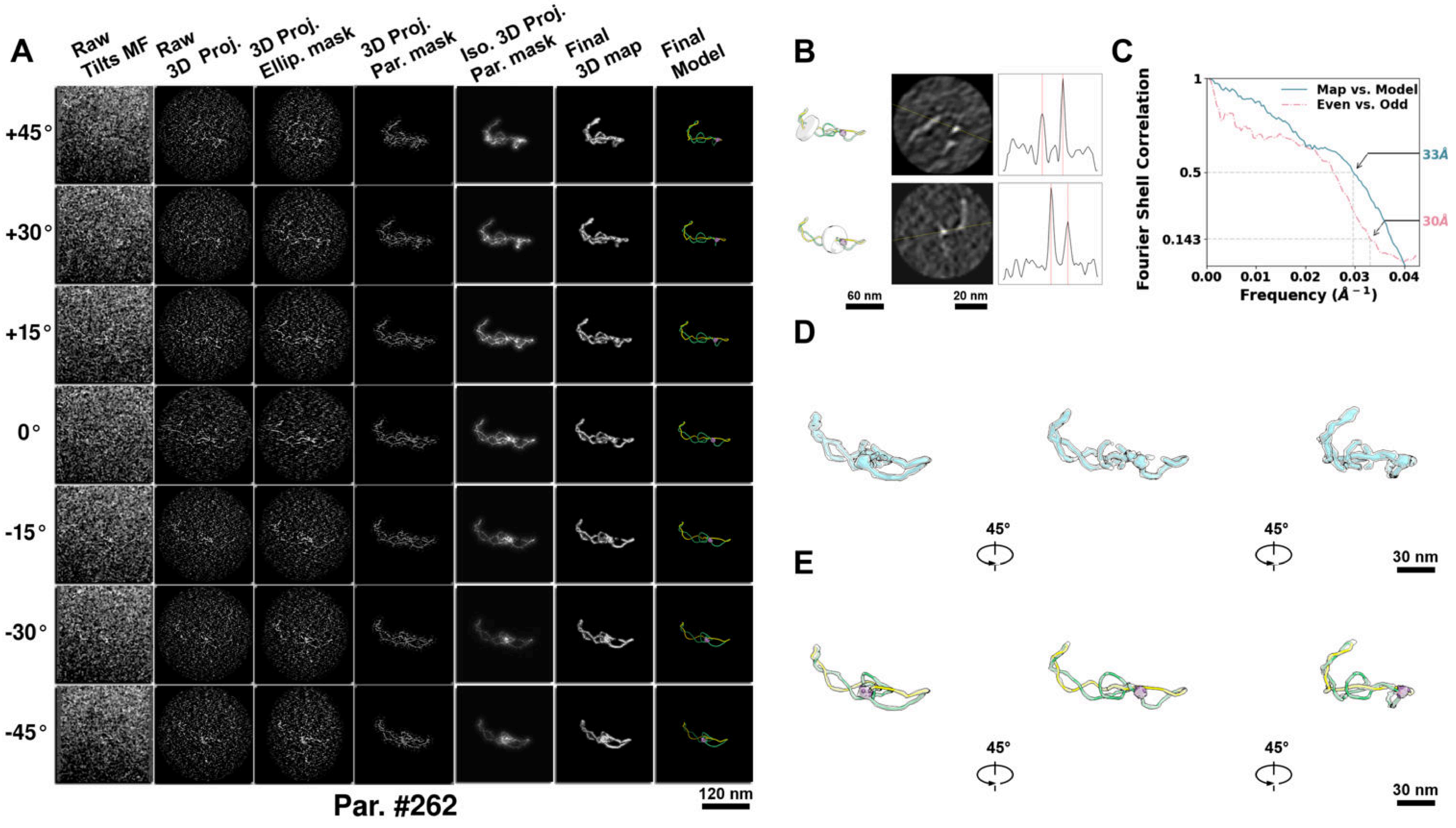

**Supplementary Particle Figure 262. Cryo-ET 3D reconstruction of an individual TEC-TopI particle.**
(A) 3D reconstruction of the plasmid particle (index no. 262). The first column shows seven representative tilt images from +45° to -45° in step of 15°. The second, third, and fourth columns show 3D projections of the particle with spherical, ellipsoidal (thinner along the z-dimension), and particle-shaped masks, respectively. The fifth column displays the 3D projections of the enhanced and IsoNet missing-wedge-corrected particle. The sixth and seventh columns present the final 3D map and the flexibly fitted model, respectively. (B) Two cross-sectional views (12 nm thickness) of the plasmid density map along its plectoneme axis are shown in the left-middle panel. The intensity profile along the line crossing the two high-density DNA spots is displayed in the right panel. (C) Resolution assessment of the final 3D map using Fourier shell correlation (FSC). Two criteria are shown: FSC between two half-maps reconstructed from even and odd frames (evaluated at 0.143) and FSC between the final 3D map and the fitted model (evaluated at 0.5). (D) Zoomed-in views of the final 3D density map from panel A, displayed at two contour levels. (E) Superimposition of the high-contour level map from panel D onto its fitted model.

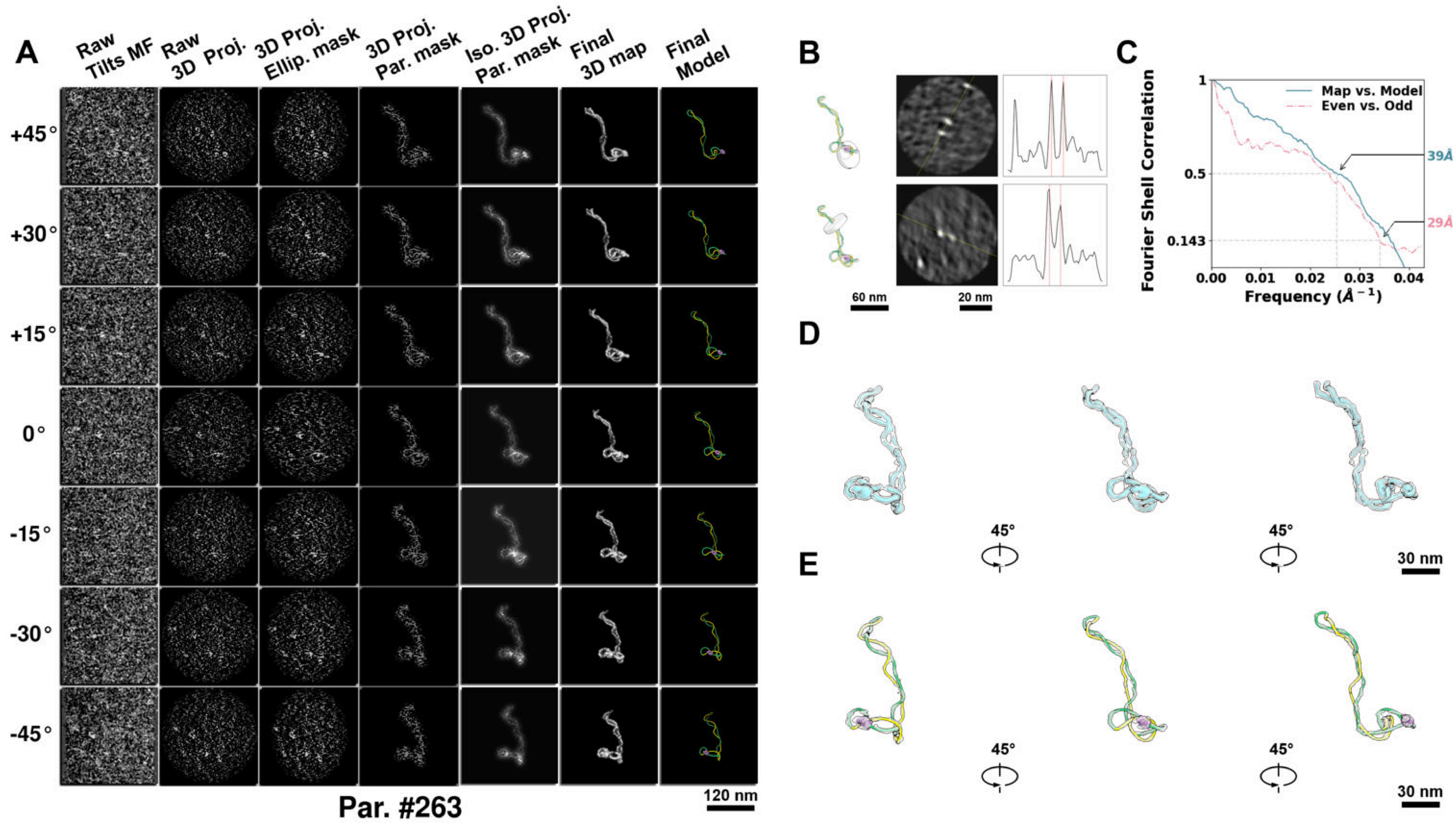

**Supplementary Particle Figure 263. Cryo-ET 3D reconstruction of an individual TEC-TopI particle.**
(A) 3D reconstruction of the plasmid particle (index no. 263). The first column shows seven representative tilt images from +45° to -45° in step of 15°. The second, third, and fourth columns show 3D projections of the particle with spherical, ellipsoidal (thinner along the z-dimension), and particle-shaped masks, respectively. The fifth column displays the 3D projections of the enhanced and IsoNet missing-wedge-corrected particle. The sixth and seventh columns present the final 3D map and the flexibly fitted model, respectively. (B) Two cross-sectional views (12 nm thickness) of the plasmid density map along its plectoneme axis are shown in the left-middle panel. The intensity profile along the line crossing the two high-density DNA spots is displayed in the right panel. (C) Resolution assessment of the final 3D map using Fourier shell correlation (FSC). Two criteria are shown: FSC between two half-maps reconstructed from even and odd frames (evaluated at 0.143) and FSC between the final 3D map and the fitted model (evaluated at 0.5). (D) Zoomed-in views of the final 3D density map from panel A, displayed at two contour levels. (E) Superimposition of the high-contour level map from panel D onto its fitted model.

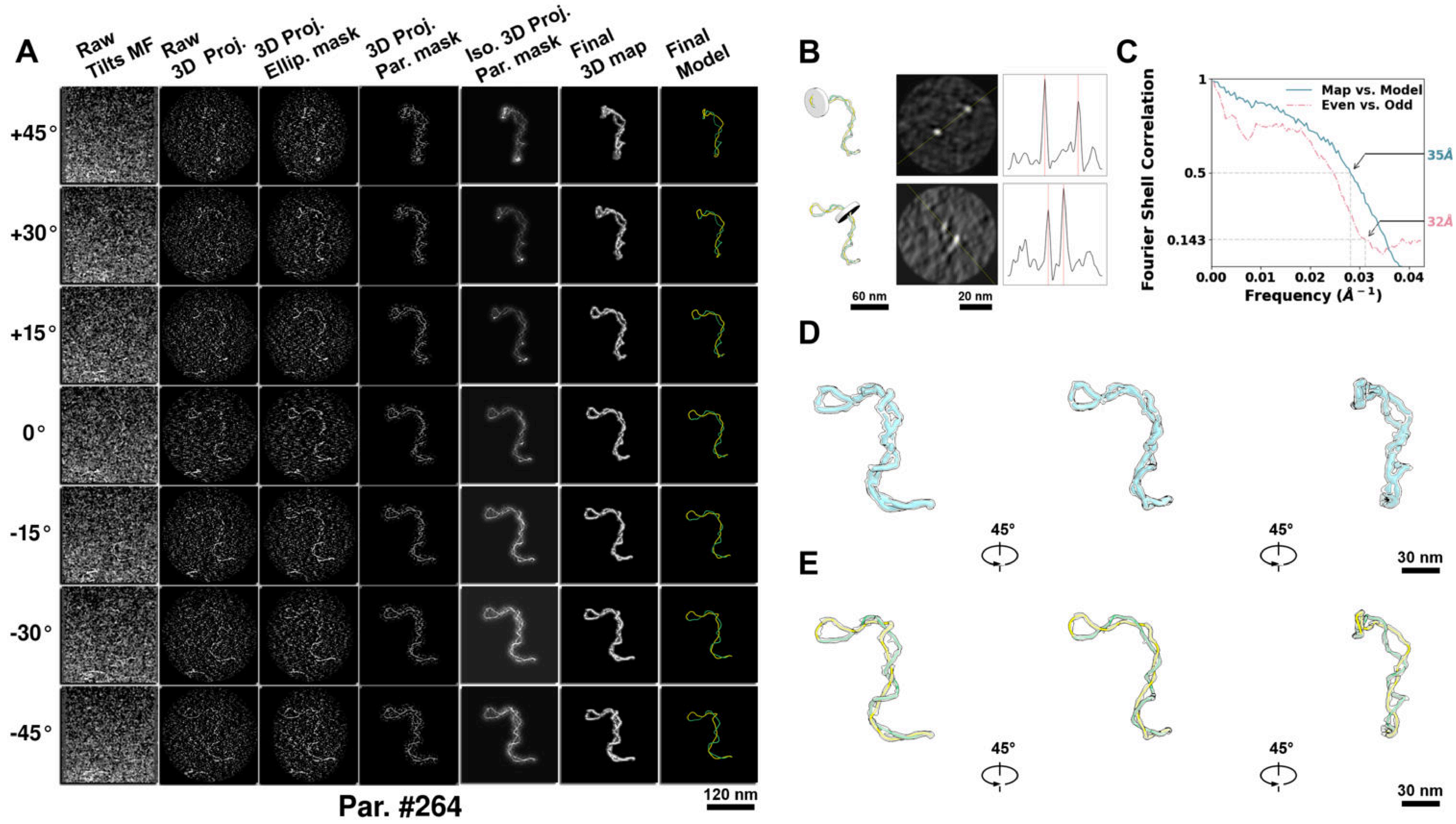

**Supplementary Particle Figure 264. Cryo-ET 3D reconstruction of an individual TEC-TopI particle.**
(A) 3D reconstruction of the plasmid particle (index no. 264). The first column shows seven representative tilt images from +45° to -45° in step of 15°. The second, third, and fourth columns show 3D projections of the particle with spherical, ellipsoidal (thinner along the z-dimension), and particle-shaped masks, respectively. The fifth column displays the 3D projections of the enhanced and IsoNet missing-wedge-corrected particle. The sixth and seventh columns present the final 3D map and the flexibly fitted model, respectively. (B) Two cross-sectional views (12 nm thickness) of the plasmid density map along its plectoneme axis are shown in the left-middle panel. The intensity profile along the line crossing the two high-density DNA spots is displayed in the right panel. (C) Resolution assessment of the final 3D map using Fourier shell correlation (FSC). Two criteria are shown: FSC between two half-maps reconstructed from even and odd frames (evaluated at 0.143) and FSC between the final 3D map and the fitted model (evaluated at 0.5). (D) Zoomed-in views of the final 3D density map from panel A, displayed at two contour levels. (E) Superimposition of the high-contour level map from panel D onto its fitted model.

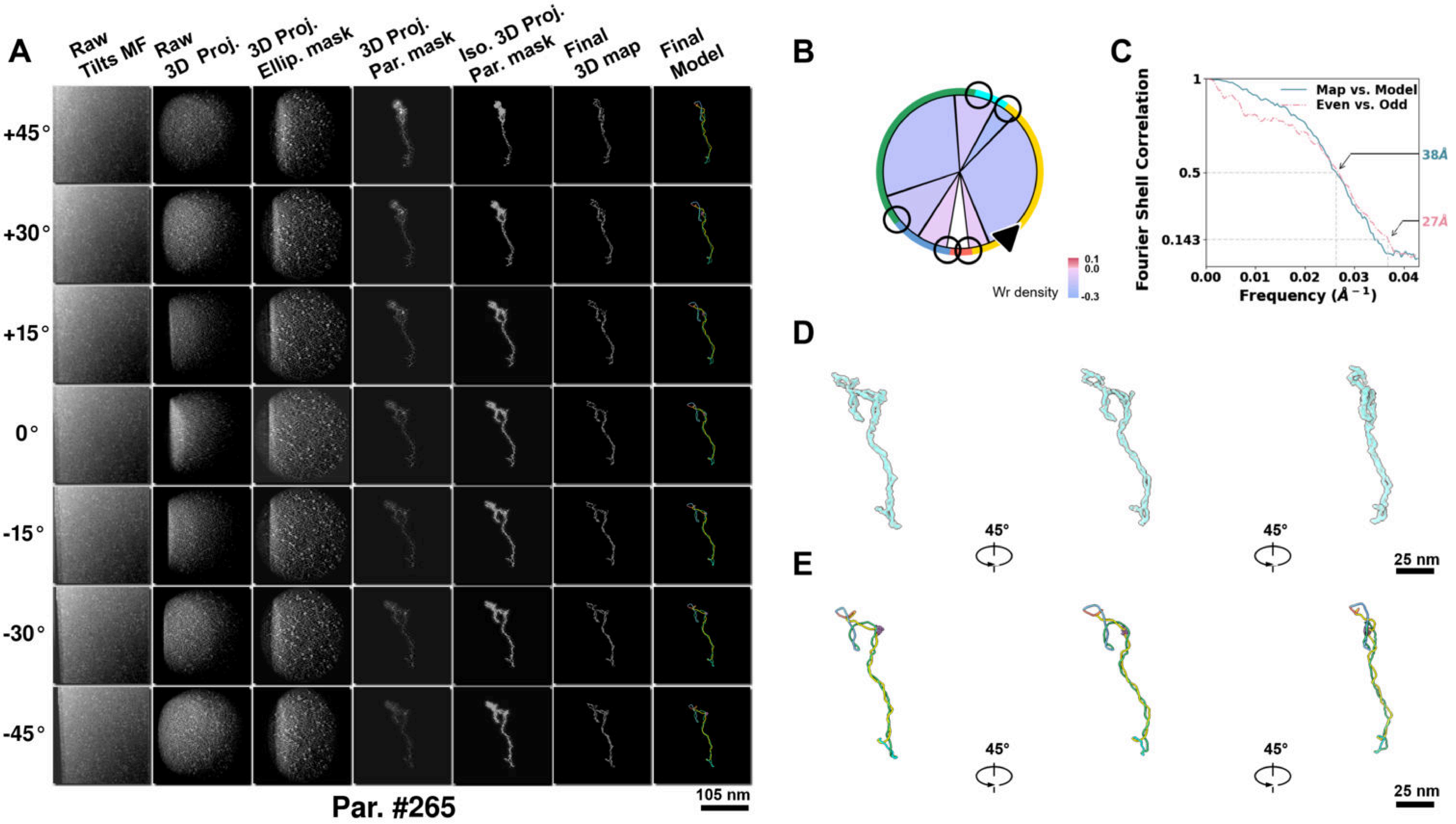

**Supplementary Particle Figure 265. Cryo-ET 3D reconstruction of an individual Opp.-TEC particle.**
(A) 3D reconstruction of the plasmid particle (index no. 265). The first column shows seven representative tilt images from +45° to -45° in step of 15°. The second, third, and fourth columns show 3D projections of the particle with spherical, ellipsoidal (thinner along the z-dimension), and particle-shaped masks, respectively. The fifth column displays the 3D projections of the enhanced and IsoNet missing-wedge-corrected particle. The sixth and seventh columns present the final 3D map and the flexibly fitted model, respectively. (B) Circular schematic representation of a plasmid particle. The outer rim is color-coded to match the corresponding 3D model. Arrowheads indicate the transcriptional direction of bound RNAPs, and circles denote apical sites. Inner circular sectors represent individual plectonemes, with colors indicating writhe density (blue to red scale, −0.3 to 0.1). (C) Resolution assessment of the final 3D map using Fourier shell correlation (FSC). Two criteria are shown: FSC between two half-maps reconstructed from even and odd frames (evaluated at 0.143) and FSC between the final 3D map and the fitted model (evaluated at 0.5). (D) Zoomed-in views of the final 3D density map from panel A, displayed at two contour levels. (E) Superimposition of the high-contour level map from panel D onto its fitted model.

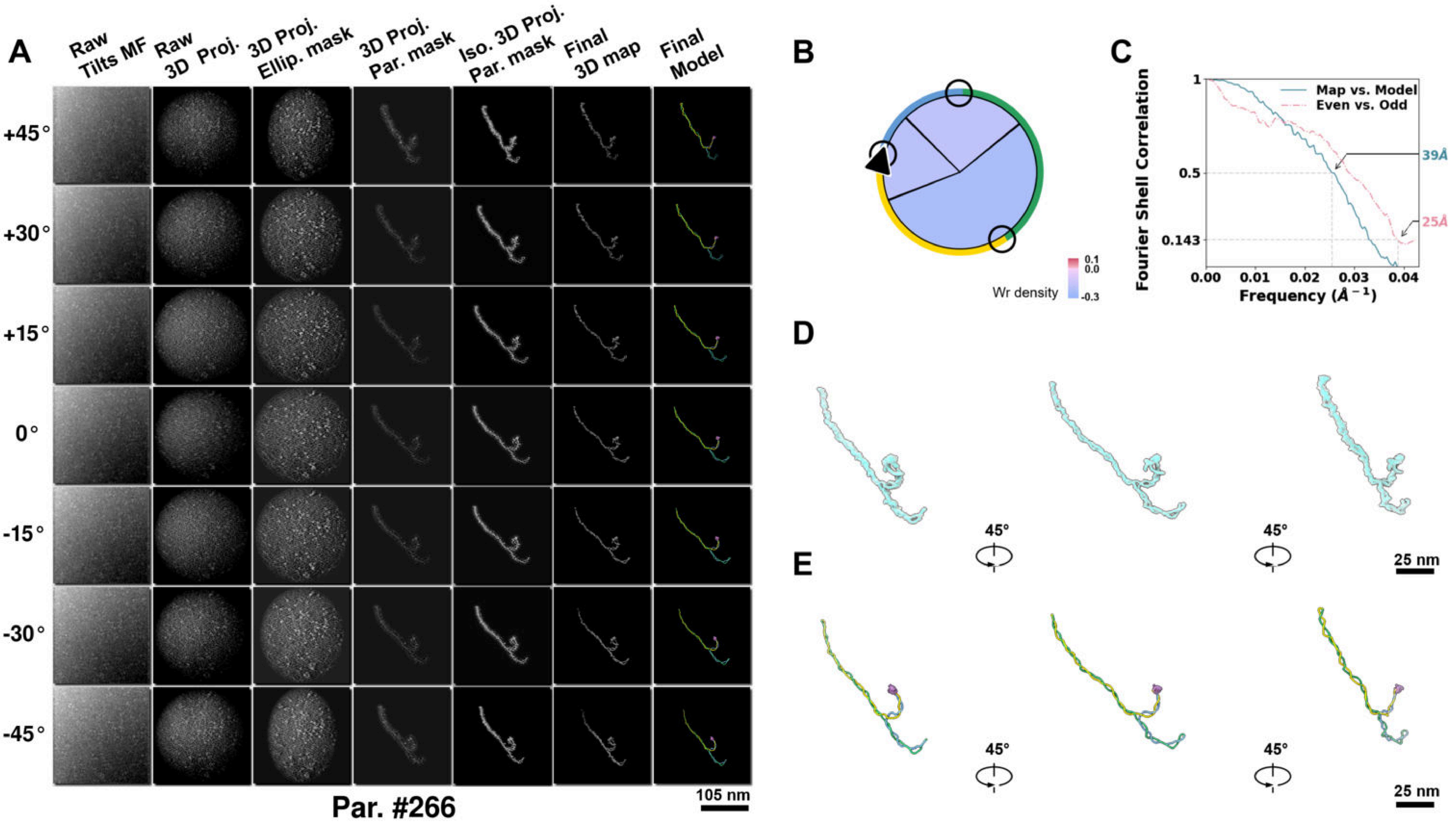

**Supplementary Particle Figure 266. Cryo-ET 3D reconstruction of an individual Opp.-TEC particle.**
(A) 3D reconstruction of the plasmid particle (index no. 266). The first column shows seven representative tilt images from +45° to -45° in step of 15°. The second, third, and fourth columns show 3D projections of the particle with spherical, ellipsoidal (thinner along the z-dimension), and particle-shaped masks, respectively. The fifth column displays the 3D projections of the enhanced and IsoNet missing-wedge-corrected particle. The sixth and seventh columns present the final 3D map and the flexibly fitted model, respectively. (B) Circular schematic representation of a plasmid particle. The outer rim is color-coded to match the corresponding 3D model. Arrowheads indicate the transcriptional direction of bound RNAPs, and circles denote apical sites. Inner circular sectors represent individual plectonemes, with colors indicating writhe density (blue to red scale, −0.3 to 0.1). (C) Resolution assessment of the final 3D map using Fourier shell correlation (FSC). Two criteria are shown: FSC between two half-maps reconstructed from even and odd frames (evaluated at 0.143) and FSC between the final 3D map and the fitted model (evaluated at 0.5). (D) Zoomed-in views of the final 3D density map from panel A, displayed at two contour levels. (E) Superimposition of the high-contour level map from panel D onto its fitted model.

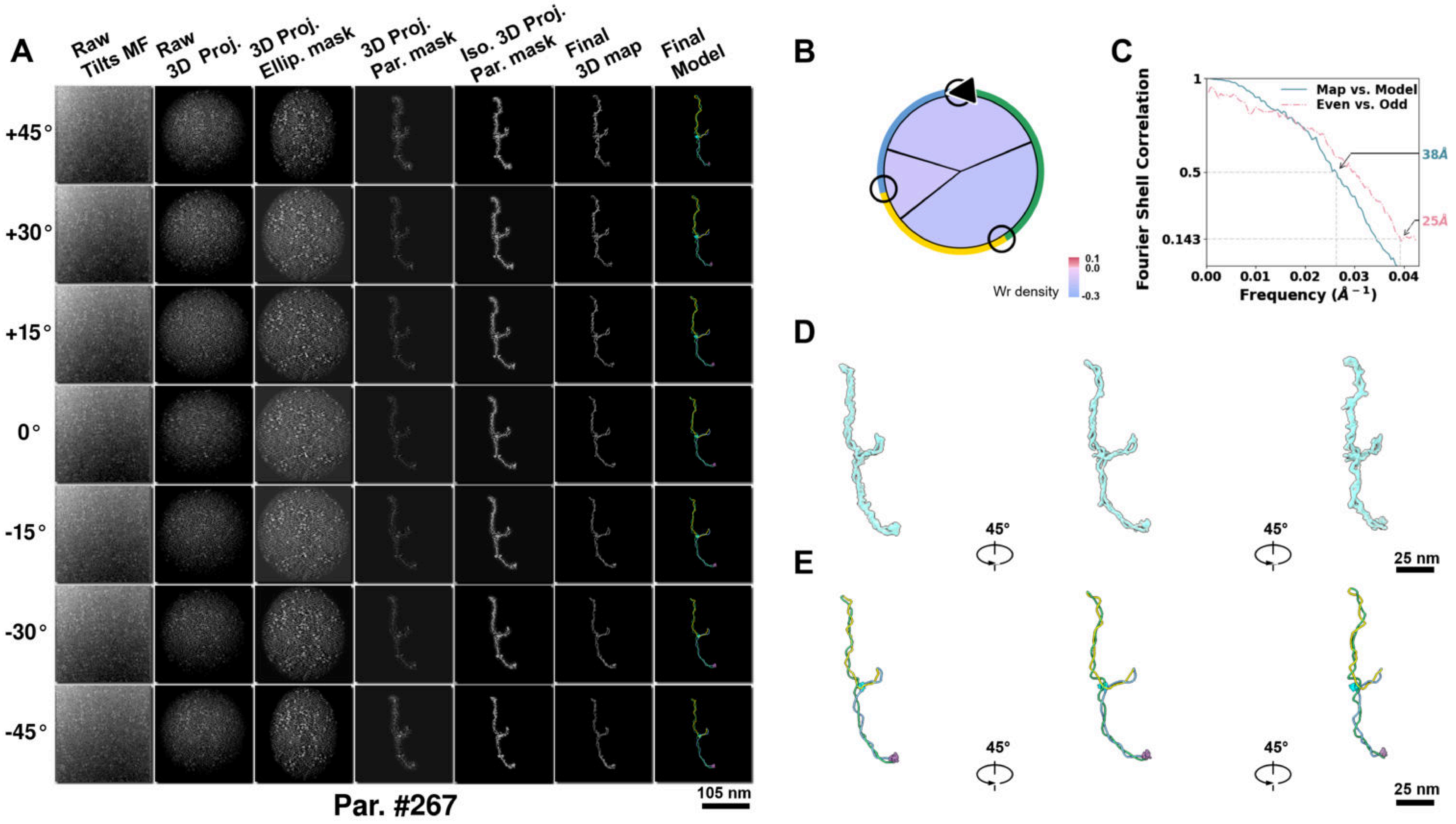

**Supplementary Particle Figure 267. Cryo-ET 3D reconstruction of an individual Opp.-TEC particle.**
(A) 3D reconstruction of the plasmid particle (index no. 267). The first column shows seven representative tilt images from +45° to -45° in step of 15°. The second, third, and fourth columns show 3D projections of the particle with spherical, ellipsoidal (thinner along the z-dimension), and particle-shaped masks, respectively. The fifth column displays the 3D projections of the enhanced and IsoNet missing-wedge-corrected particle. The sixth and seventh columns present the final 3D map and the flexibly fitted model, respectively. (B) Circular schematic representation of a plasmid particle. The outer rim is color-coded to match the corresponding 3D model. Arrowheads indicate the transcriptional direction of bound RNAPs, and circles denote apical sites. Inner circular sectors represent individual plectonemes, with colors indicating writhe density (blue to red scale, −0.3 to 0.1). (C) Resolution assessment of the final 3D map using Fourier shell correlation (FSC). Two criteria are shown: FSC between two half-maps reconstructed from even and odd frames (evaluated at 0.143) and FSC between the final 3D map and the fitted model (evaluated at 0.5). (D) Zoomed-in views of the final 3D density map from panel A, displayed at two contour levels. (E) Superimposition of the high-contour level map from panel D onto its fitted model.

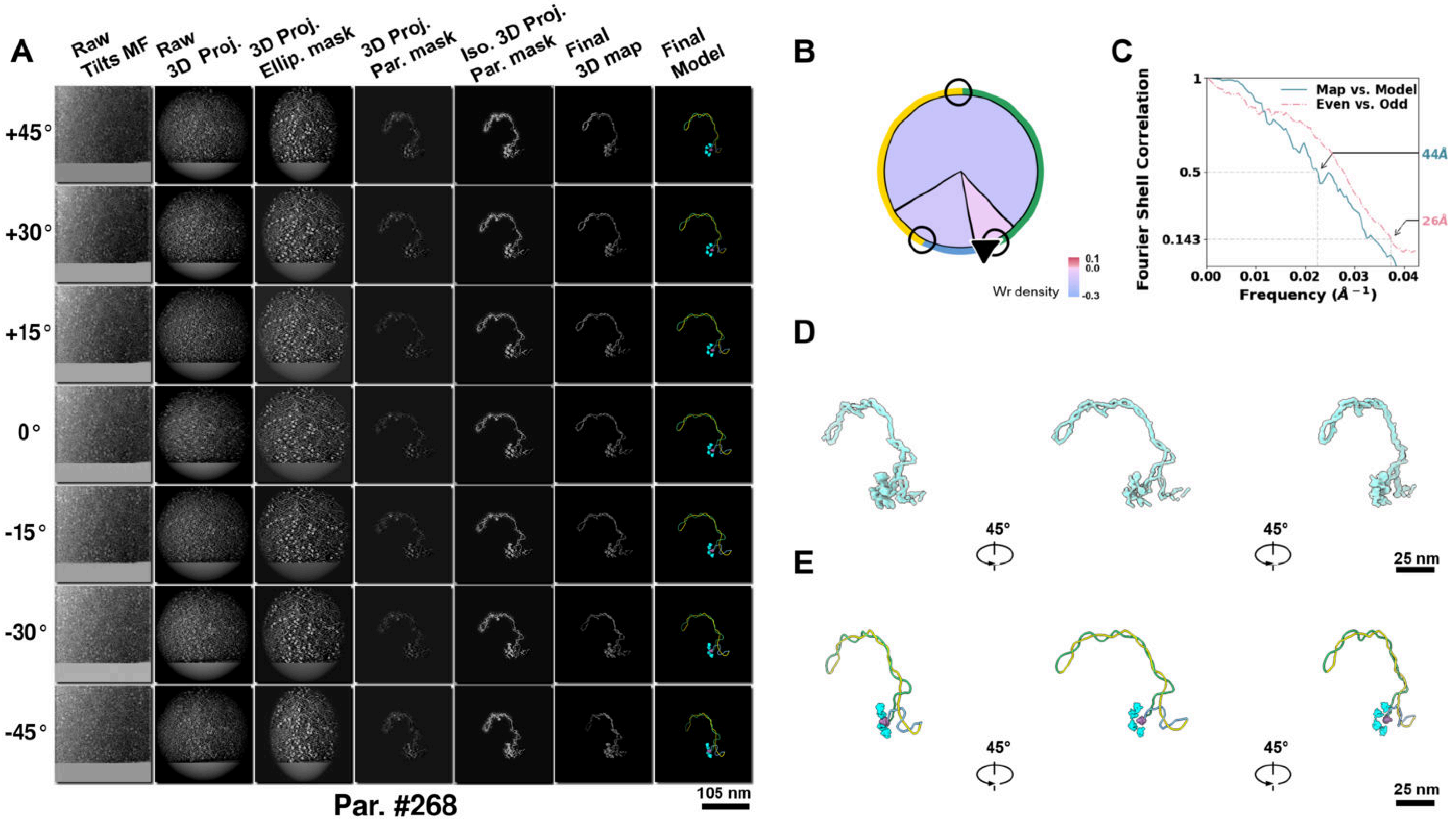

**Supplementary Particle Figure 268. Cryo-ET 3D reconstruction of an individual Opp.-TEC particle.**
(A) 3D reconstruction of the plasmid particle (index no. 268). The first column shows seven representative tilt images from +45° to -45° in step of 15°. The second, third, and fourth columns show 3D projections of the particle with spherical, ellipsoidal (thinner along the z-dimension), and particle-shaped masks, respectively. The fifth column displays the 3D projections of the enhanced and IsoNet missing-wedge-corrected particle. The sixth and seventh columns present the final 3D map and the flexibly fitted model, respectively. (B) Circular schematic representation of a plasmid particle. The outer rim is color-coded to match the corresponding 3D model. Arrowheads indicate the transcriptional direction of bound RNAPs, and circles denote apical sites. Inner circular sectors represent individual plectonemes, with colors indicating writhe density (blue to red scale, −0.3 to 0.1). (C) Resolution assessment of the final 3D map using Fourier shell correlation (FSC). Two criteria are shown: FSC between two half-maps reconstructed from even and odd frames (evaluated at 0.143) and FSC between the final 3D map and the fitted model (evaluated at 0.5). (D) Zoomed-in views of the final 3D density map from panel A, displayed at two contour levels. (E) Superimposition of the high-contour level map from panel D onto its fitted model.

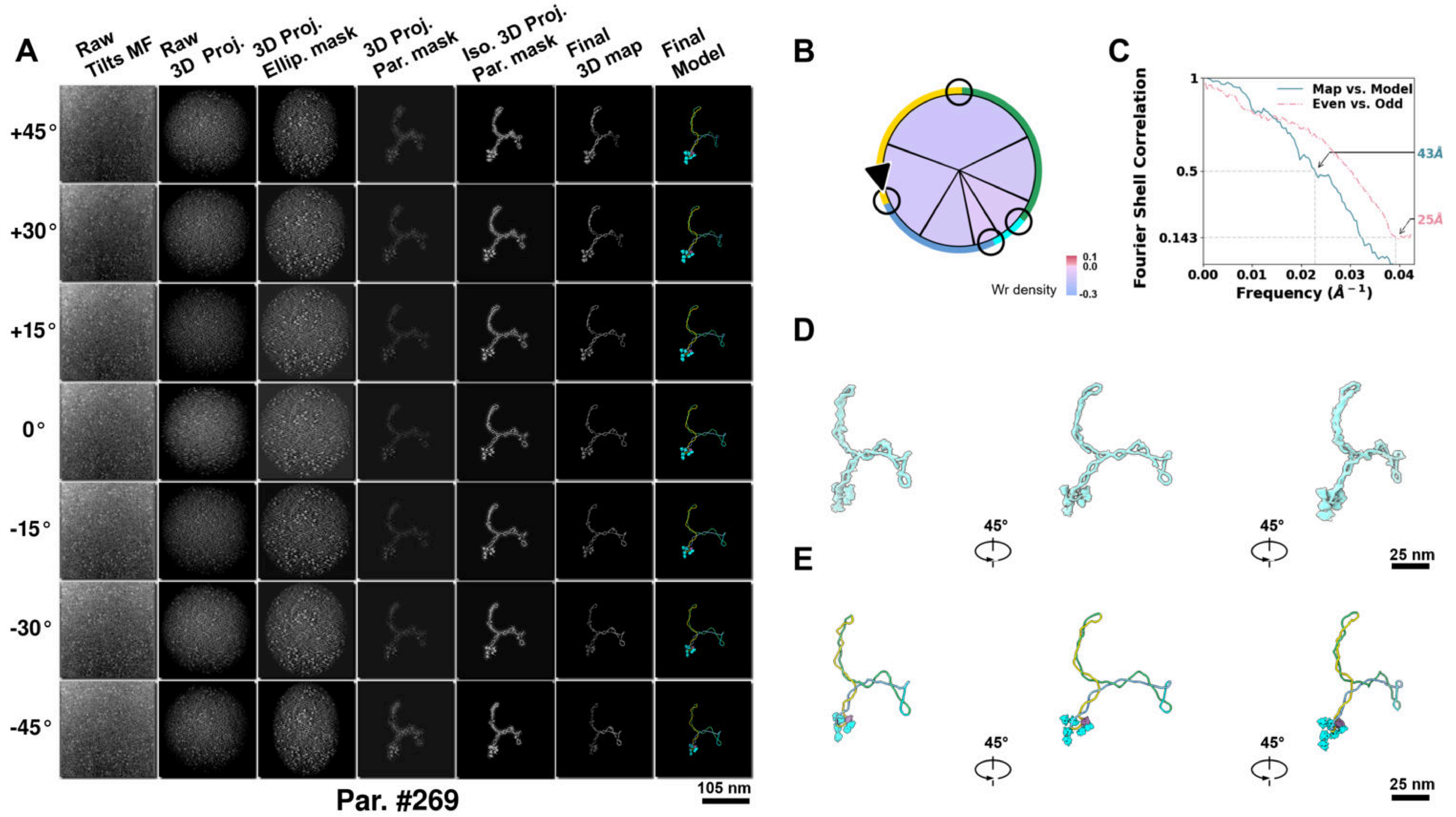

**Supplementary Particle Figure 269. Cryo-ET 3D reconstruction of an individual Opp.-TEC particle.**
(A) 3D reconstruction of the plasmid particle (index no. 269). The first column shows seven representative tilt images from +45° to -45° in step of 15°. The second, third, and fourth columns show 3D projections of the particle with spherical, ellipsoidal (thinner along the z-dimension), and particle-shaped masks, respectively. The fifth column displays the 3D projections of the enhanced and IsoNet missing-wedge-corrected particle. The sixth and seventh columns present the final 3D map and the flexibly fitted model, respectively. (B) Circular schematic representation of a plasmid particle. The outer rim is color-coded to match the corresponding 3D model. Arrowheads indicate the transcriptional direction of bound RNAPs, and circles denote apical sites. Inner circular sectors represent individual plectonemes, with colors indicating writhe density (blue to red scale, −0.3 to 0.1). (C) Resolution assessment of the final 3D map using Fourier shell correlation (FSC). Two criteria are shown: FSC between two half-maps reconstructed from even and odd frames (evaluated at 0.143) and FSC between the final 3D map and the fitted model (evaluated at 0.5). (D) Zoomed-in views of the final 3D density map from panel A, displayed at two contour levels. (E) Superimposition of the high-contour level map from panel D onto its fitted model.

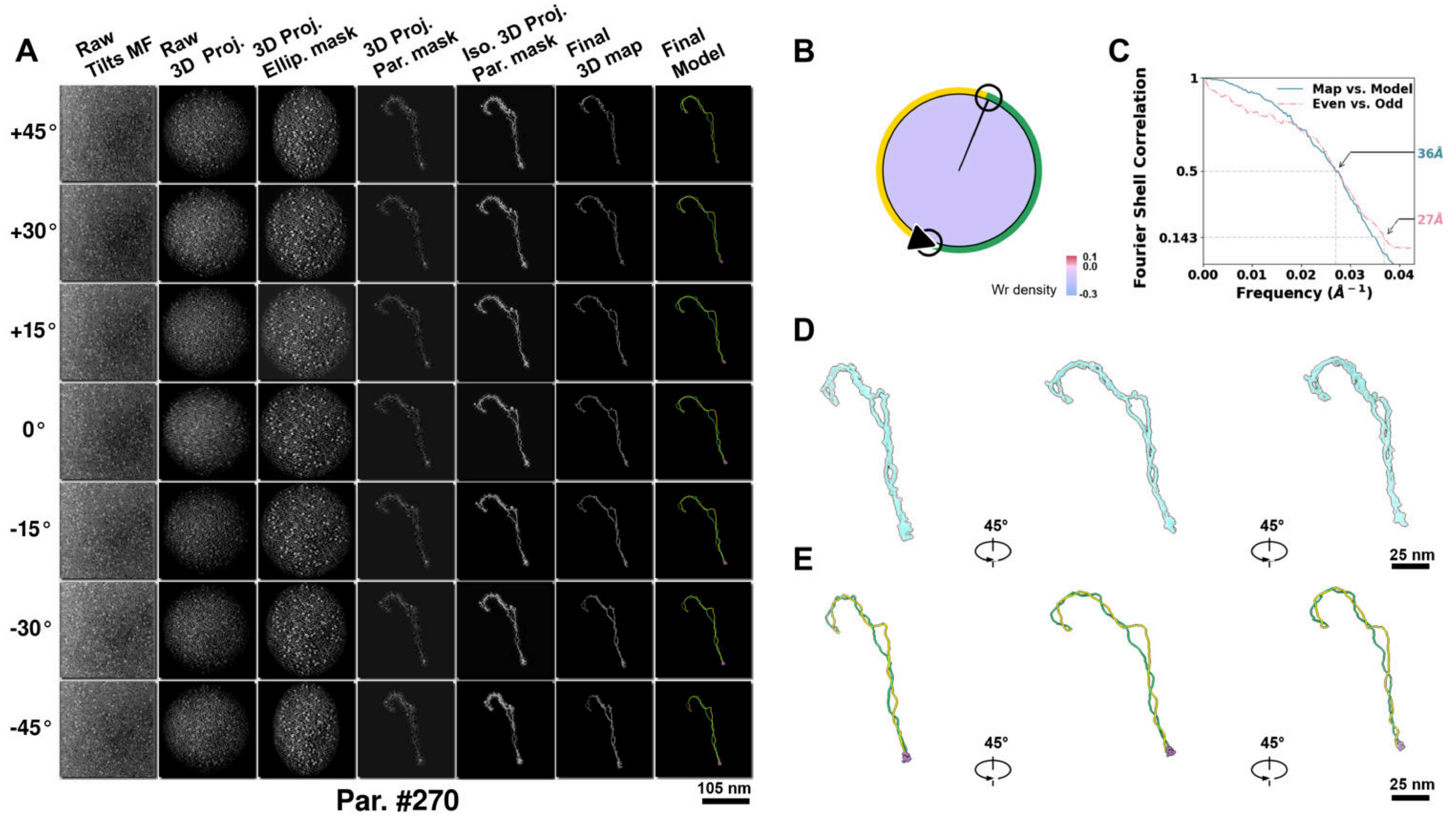

**Supplementary Particle Figure 270. Cryo-ET 3D reconstruction of an individual Opp.-TEC particle.**
(A) 3D reconstruction of the plasmid particle (index no. 270). The first column shows seven representative tilt images from +45° to -45° in step of 15°. The second, third, and fourth columns show 3D projections of the particle with spherical, ellipsoidal (thinner along the z-dimension), and particle-shaped masks, respectively. The fifth column displays the 3D projections of the enhanced and IsoNet missing-wedge-corrected particle. The sixth and seventh columns present the final 3D map and the flexibly fitted model, respectively. (B) Circular schematic representation of a plasmid particle. The outer rim is color-coded to match the corresponding 3D model. Arrowheads indicate the transcriptional direction of bound RNAPs, and circles denote apical sites. Inner circular sectors represent individual plectonemes, with colors indicating writhe density (blue to red scale, −0.3 to 0.1). (C) Resolution assessment of the final 3D map using Fourier shell correlation (FSC). Two criteria are shown: FSC between two half-maps reconstructed from even and odd frames (evaluated at 0.143) and FSC between the final 3D map and the fitted model (evaluated at 0.5). (D) Zoomed-in views of the final 3D density map from panel A, displayed at two contour levels. (E) Superimposition of the high-contour level map from panel D onto its fitted model.

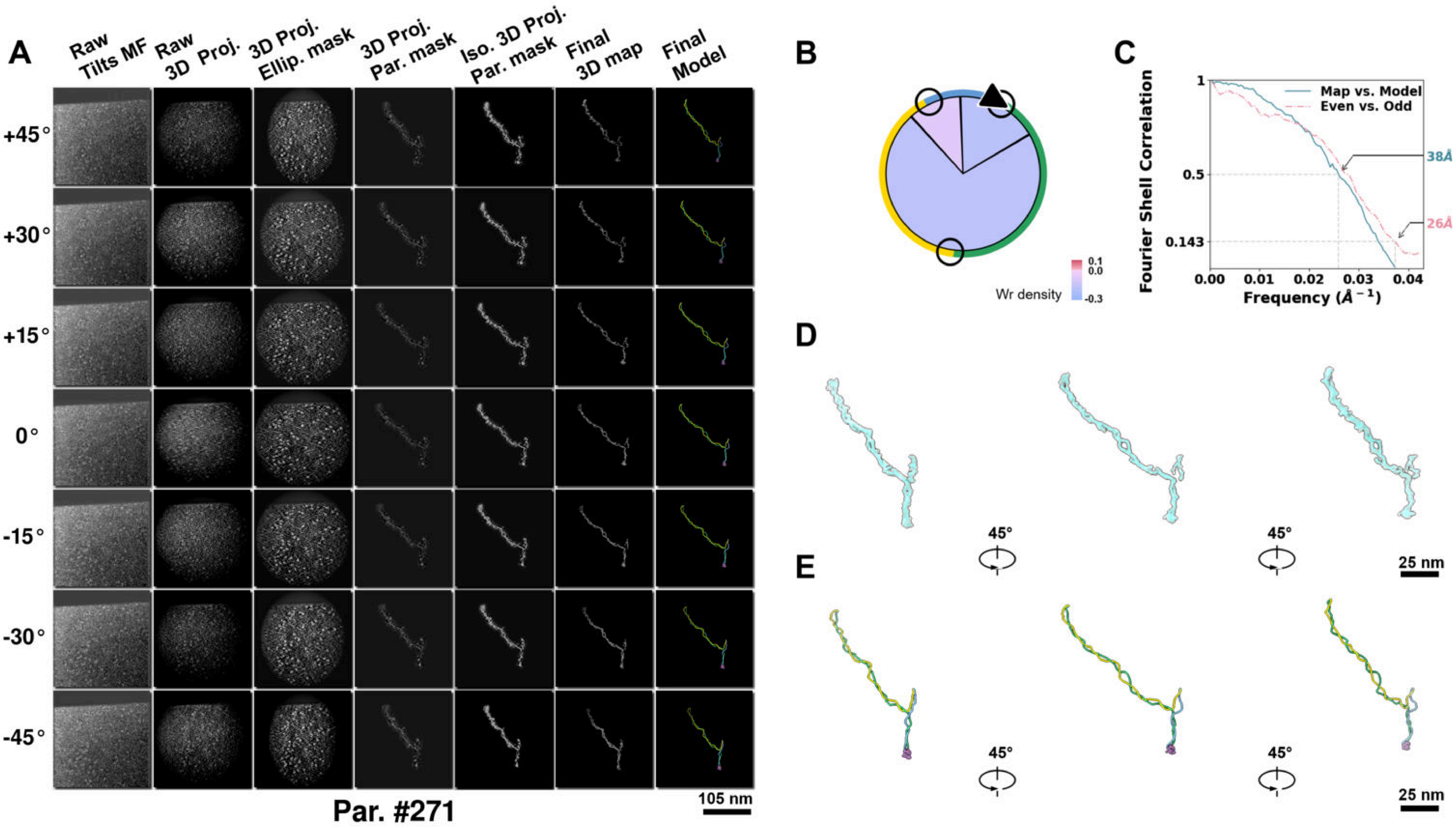

**Supplementary Particle Figure 271. Cryo-ET 3D reconstruction of an individual Opp.-TEC particle.**
(A) 3D reconstruction of the plasmid particle (index no. 271). The first column shows seven representative tilt images from +45° to -45° in step of 15°. The second, third, and fourth columns show 3D projections of the particle with spherical, ellipsoidal (thinner along the z-dimension), and particle-shaped masks, respectively. The fifth column displays the 3D projections of the enhanced and IsoNet missing-wedge-corrected particle. The sixth and seventh columns present the final 3D map and the flexibly fitted model, respectively. (B) Circular schematic representation of a plasmid particle. The outer rim is color-coded to match the corresponding 3D model. Arrowheads indicate the transcriptional direction of bound RNAPs, and circles denote apical sites. Inner circular sectors represent individual plectonemes, with colors indicating writhe density (blue to red scale, −0.3 to 0.1). (C) Resolution assessment of the final 3D map using Fourier shell correlation (FSC). Two criteria are shown: FSC between two half-maps reconstructed from even and odd frames (evaluated at 0.143) and FSC between the final 3D map and the fitted model (evaluated at 0.5). (D) Zoomed-in views of the final 3D density map from panel A, displayed at two contour levels. (E) Superimposition of the high-contour level map from panel D onto its fitted model.

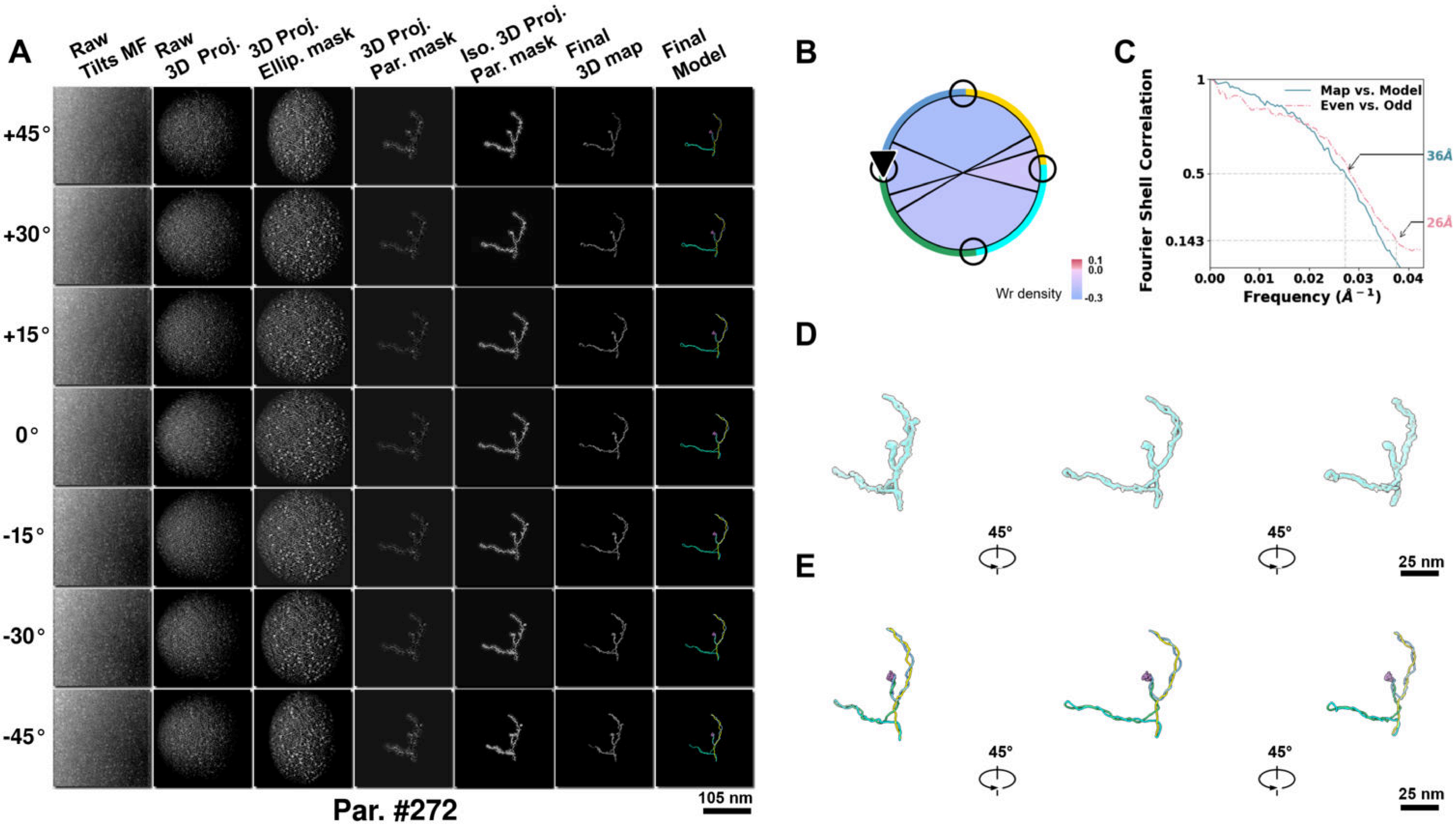

**Supplementary Particle Figure 272. Cryo-ET 3D reconstruction of an individual Opp.-TEC particle.**
(A) 3D reconstruction of the plasmid particle (index no. 272). The first column shows seven representative tilt images from +45° to -45° in step of 15°. The second, third, and fourth columns show 3D projections of the particle with spherical, ellipsoidal (thinner along the z-dimension), and particle-shaped masks, respectively. The fifth column displays the 3D projections of the enhanced and IsoNet missing-wedge-corrected particle. The sixth and seventh columns present the final 3D map and the flexibly fitted model, respectively. (B) Circular schematic representation of a plasmid particle. The outer rim is color-coded to match the corresponding 3D model. Arrowheads indicate the transcriptional direction of bound RNAPs, and circles denote apical sites. Inner circular sectors represent individual plectonemes, with colors indicating writhe density (blue to red scale, −0.3 to 0.1). (C) Resolution assessment of the final 3D map using Fourier shell correlation (FSC). Two criteria are shown: FSC between two half-maps reconstructed from even and odd frames (evaluated at 0.143) and FSC between the final 3D map and the fitted model (evaluated at 0.5). (D) Zoomed-in views of the final 3D density map from panel A, displayed at two contour levels. (E) Superimposition of the high-contour level map from panel D onto its fitted model.

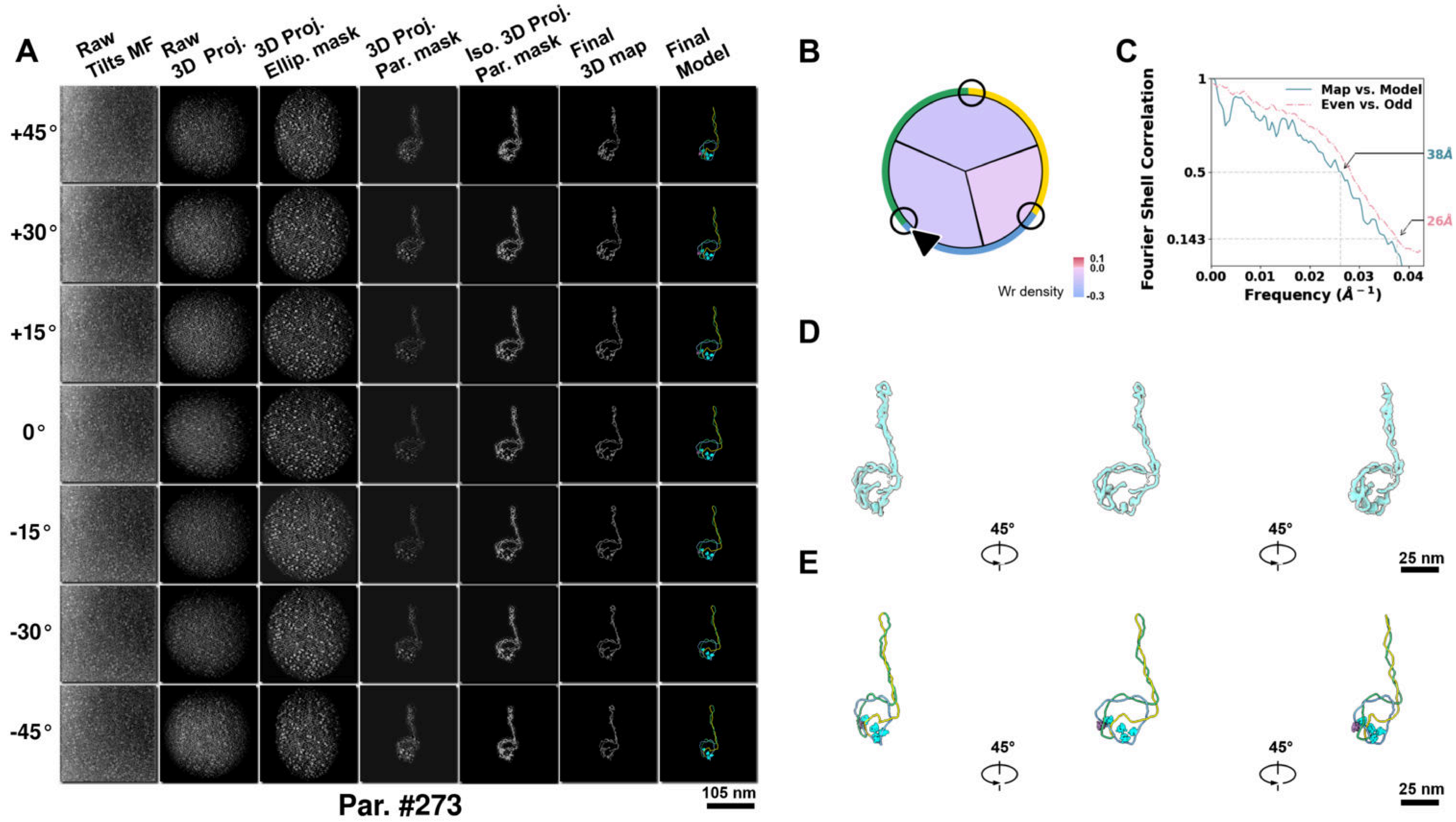

**Supplementary Particle Figure 273. Cryo-ET 3D reconstruction of an individual Opp.-TEC particle.**
(A) 3D reconstruction of the plasmid particle (index no. 273). The first column shows seven representative tilt images from +45° to -45° in step of 15°. The second, third, and fourth columns show 3D projections of the particle with spherical, ellipsoidal (thinner along the z-dimension), and particle-shaped masks, respectively. The fifth column displays the 3D projections of the enhanced and IsoNet missing-wedge-corrected particle. The sixth and seventh columns present the final 3D map and the flexibly fitted model, respectively. (B) Circular schematic representation of a plasmid particle. The outer rim is color-coded to match the corresponding 3D model. Arrowheads indicate the transcriptional direction of bound RNAPs, and circles denote apical sites. Inner circular sectors represent individual plectonemes, with colors indicating writhe density (blue to red scale, −0.3 to 0.1). (C) Resolution assessment of the final 3D map using Fourier shell correlation (FSC). Two criteria are shown: FSC between two half-maps reconstructed from even and odd frames (evaluated at 0.143) and FSC between the final 3D map and the fitted model (evaluated at 0.5). (D) Zoomed-in views of the final 3D density map from panel A, displayed at two contour levels. (E) Superimposition of the high-contour level map from panel D onto its fitted model.

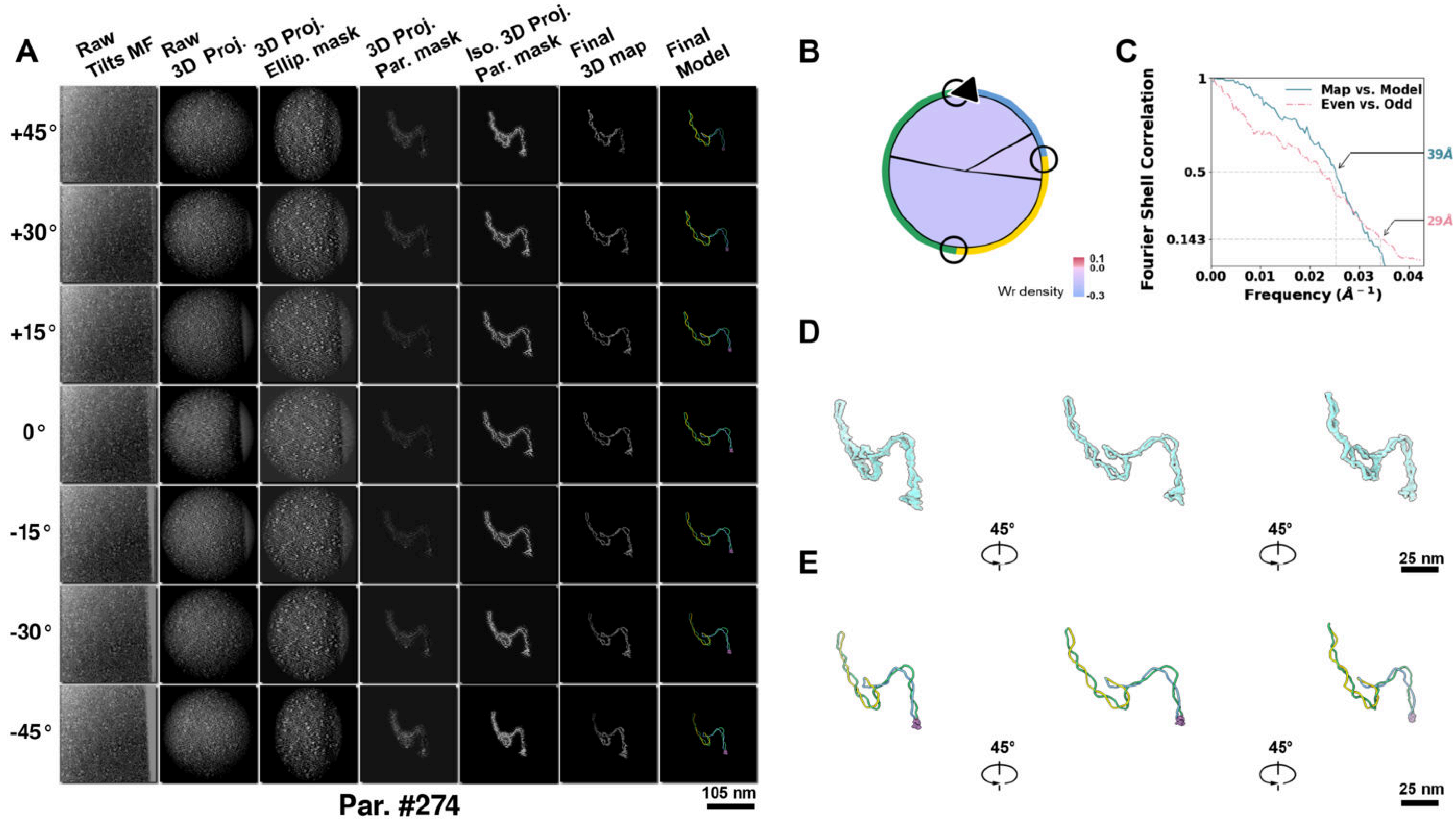

**Supplementary Particle Figure 274. Cryo-ET 3D reconstruction of an individual Opp.-TEC particle.**
(A) 3D reconstruction of the plasmid particle (index no. 274). The first column shows seven representative tilt images from +45° to -45° in step of 15°. The second, third, and fourth columns show 3D projections of the particle with spherical, ellipsoidal (thinner along the z-dimension), and particle-shaped masks, respectively. The fifth column displays the 3D projections of the enhanced and IsoNet missing-wedge-corrected particle. The sixth and seventh columns present the final 3D map and the flexibly fitted model, respectively. (B) Circular schematic representation of a plasmid particle. The outer rim is color-coded to match the corresponding 3D model. Arrowheads indicate the transcriptional direction of bound RNAPs, and circles denote apical sites. Inner circular sectors represent individual plectonemes, with colors indicating writhe density (blue to red scale, −0.3 to 0.1). (C) Resolution assessment of the final 3D map using Fourier shell correlation (FSC). Two criteria are shown: FSC between two half-maps reconstructed from even and odd frames (evaluated at 0.143) and FSC between the final 3D map and the fitted model (evaluated at 0.5). (D) Zoomed-in views of the final 3D density map from panel A, displayed at two contour levels. (E) Superimposition of the high-contour level map from panel D onto its fitted model.

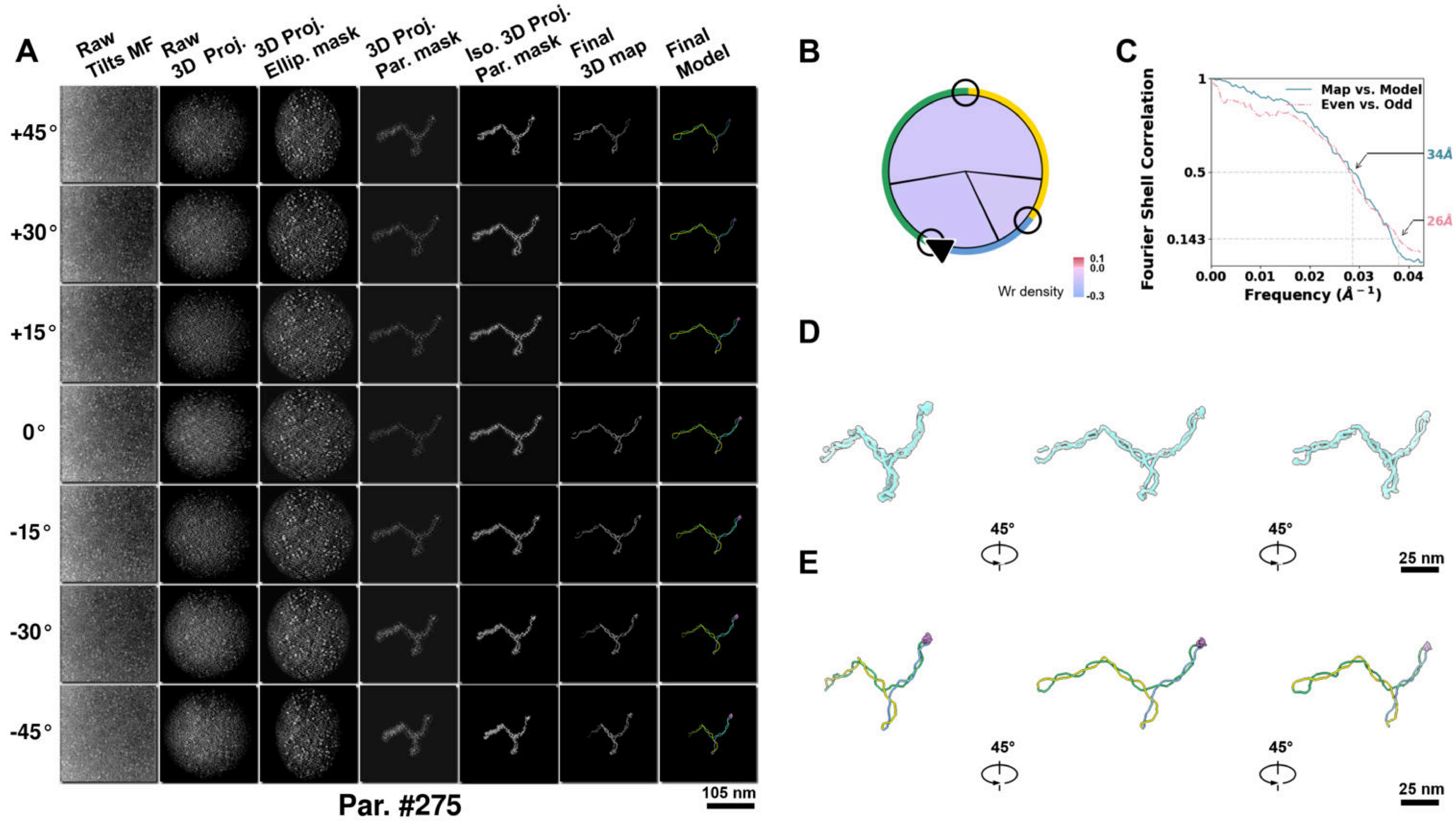

**Supplementary Particle Figure 275. Cryo-ET 3D reconstruction of an individual Opp.-TEC particle.**
(A) 3D reconstruction of the plasmid particle (index no. 275). The first column shows seven representative tilt images from +45° to -45° in step of 15°. The second, third, and fourth columns show 3D projections of the particle with spherical, ellipsoidal (thinner along the z-dimension), and particle-shaped masks, respectively. The fifth column displays the 3D projections of the enhanced and IsoNet missing-wedge-corrected particle. The sixth and seventh columns present the final 3D map and the flexibly fitted model, respectively. (B) Circular schematic representation of a plasmid particle. The outer rim is color-coded to match the corresponding 3D model. Arrowheads indicate the transcriptional direction of bound RNAPs, and circles denote apical sites. Inner circular sectors represent individual plectonemes, with colors indicating writhe density (blue to red scale, −0.3 to 0.1). (C) Resolution assessment of the final 3D map using Fourier shell correlation (FSC). Two criteria are shown: FSC between two half-maps reconstructed from even and odd frames (evaluated at 0.143) and FSC between the final 3D map and the fitted model (evaluated at 0.5). (D) Zoomed-in views of the final 3D density map from panel A, displayed at two contour levels. (E) Superimposition of the high-contour level map from panel D onto its fitted model.

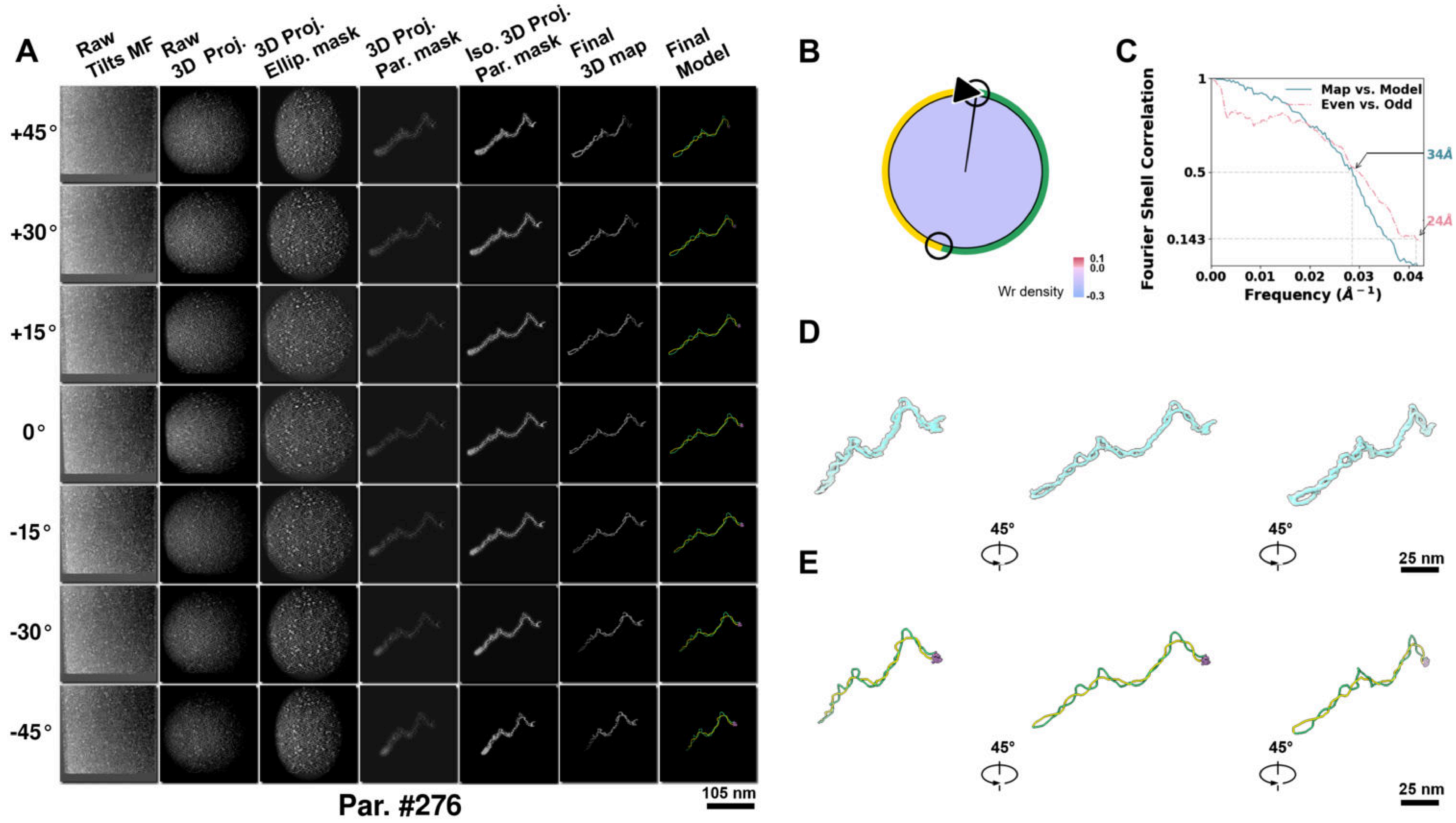

**Supplementary Particle Figure 276. Cryo-ET 3D reconstruction of an individual Opp.-TEC particle.**
(A) 3D reconstruction of the plasmid particle (index no. 276). The first column shows seven representative tilt images from +45° to -45° in step of 15°. The second, third, and fourth columns show 3D projections of the particle with spherical, ellipsoidal (thinner along the z-dimension), and particle-shaped masks, respectively. The fifth column displays the 3D projections of the enhanced and IsoNet missing-wedge-corrected particle. The sixth and seventh columns present the final 3D map and the flexibly fitted model, respectively. (B) Circular schematic representation of a plasmid particle. The outer rim is color-coded to match the corresponding 3D model. Arrowheads indicate the transcriptional direction of bound RNAPs, and circles denote apical sites. Inner circular sectors represent individual plectonemes, with colors indicating writhe density (blue to red scale, −0.3 to 0.1). (C) Resolution assessment of the final 3D map using Fourier shell correlation (FSC). Two criteria are shown: FSC between two half-maps reconstructed from even and odd frames (evaluated at 0.143) and FSC between the final 3D map and the fitted model (evaluated at 0.5). (D) Zoomed-in views of the final 3D density map from panel A, displayed at two contour levels. (E) Superimposition of the high-contour level map from panel D onto its fitted model.

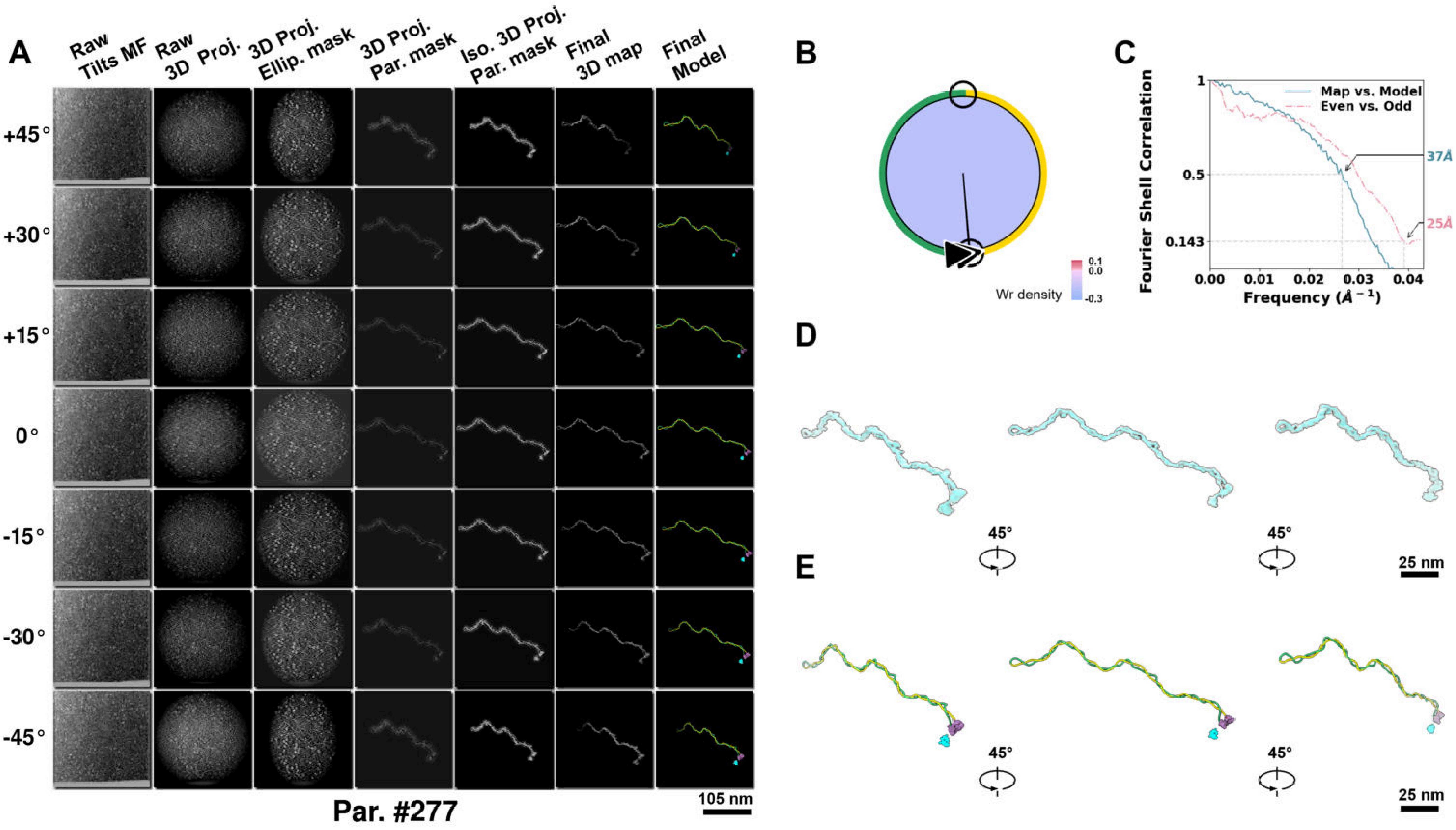

**Supplementary Particle Figure 277. Cryo-ET 3D reconstruction of an individual Opp.-TEC particle.**
(A) 3D reconstruction of the plasmid particle (index no. 277). The first column shows seven representative tilt images from +45° to -45° in step of 15°. The second, third, and fourth columns show 3D projections of the particle with spherical, ellipsoidal (thinner along the z-dimension), and particle-shaped masks, respectively. The fifth column displays the 3D projections of the enhanced and IsoNet missing-wedge-corrected particle. The sixth and seventh columns present the final 3D map and the flexibly fitted model, respectively. (B) Circular schematic representation of a plasmid particle. The outer rim is color-coded to match the corresponding 3D model. Arrowheads indicate the transcriptional direction of bound RNAPs, and circles denote apical sites. Inner circular sectors represent individual plectonemes, with colors indicating writhe density (blue to red scale, −0.3 to 0.1). (C) Resolution assessment of the final 3D map using Fourier shell correlation (FSC). Two criteria are shown: FSC between two half-maps reconstructed from even and odd frames (evaluated at 0.143) and FSC between the final 3D map and the fitted model (evaluated at 0.5). (D) Zoomed-in views of the final 3D density map from panel A, displayed at two contour levels. (E) Superimposition of the high-contour level map from panel D onto its fitted model.

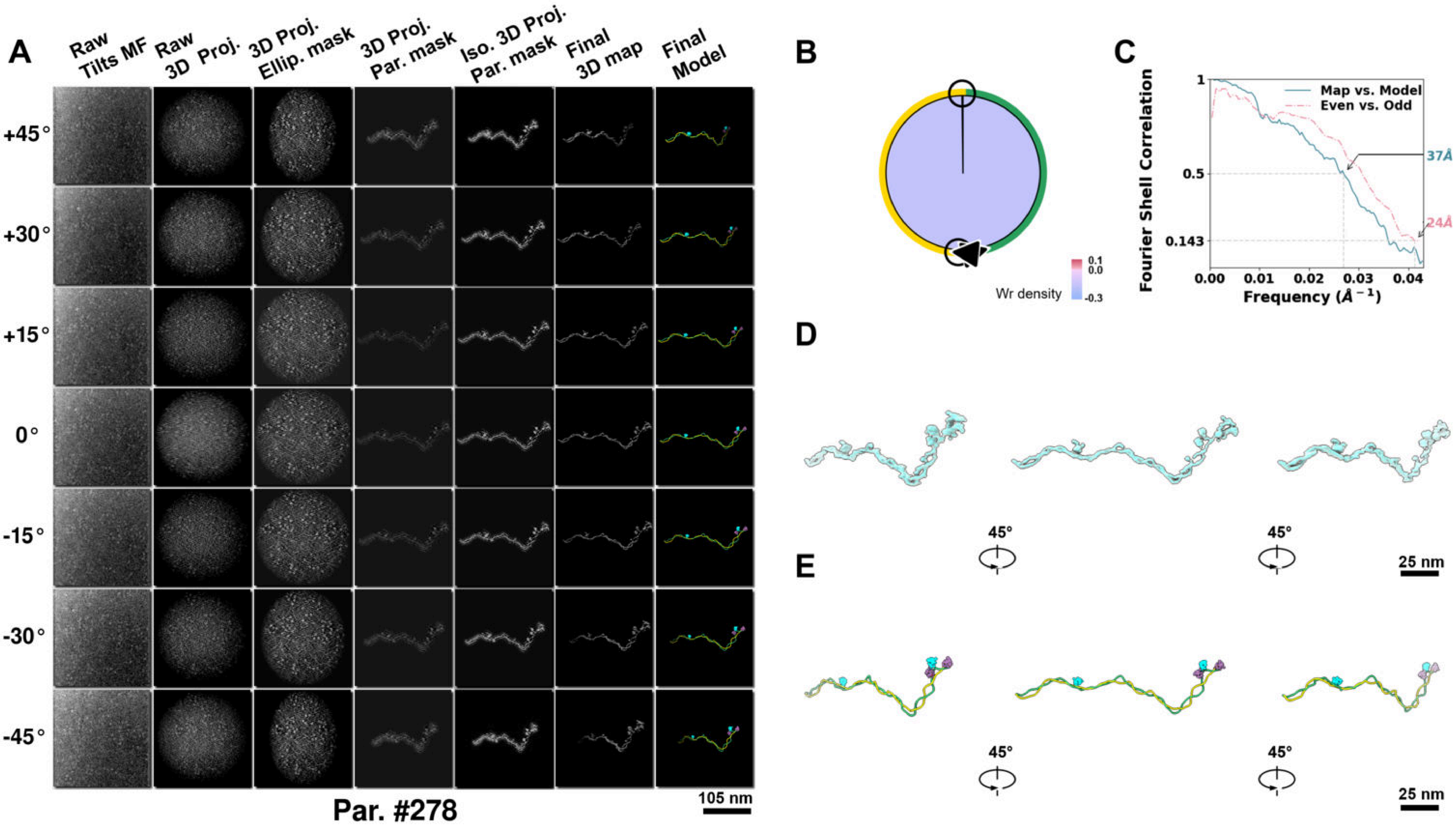

**Supplementary Particle Figure 278. Cryo-ET 3D reconstruction of an individual Opp.-TEC particle.**
(A) 3D reconstruction of the plasmid particle (index no. 278). The first column shows seven representative tilt images from +45° to -45° in step of 15°. The second, third, and fourth columns show 3D projections of the particle with spherical, ellipsoidal (thinner along the z-dimension), and particle-shaped masks, respectively. The fifth column displays the 3D projections of the enhanced and IsoNet missing-wedge-corrected particle. The sixth and seventh columns present the final 3D map and the flexibly fitted model, respectively. (B) Circular schematic representation of a plasmid particle. The outer rim is color-coded to match the corresponding 3D model. Arrowheads indicate the transcriptional direction of bound RNAPs, and circles denote apical sites. Inner circular sectors represent individual plectonemes, with colors indicating writhe density (blue to red scale, −0.3 to 0.1). (C) Resolution assessment of the final 3D map using Fourier shell correlation (FSC). Two criteria are shown: FSC between two half-maps reconstructed from even and odd frames (evaluated at 0.143) and FSC between the final 3D map and the fitted model (evaluated at 0.5). (D) Zoomed-in views of the final 3D density map from panel A, displayed at two contour levels. (E) Superimposition of the high-contour level map from panel D onto its fitted model.

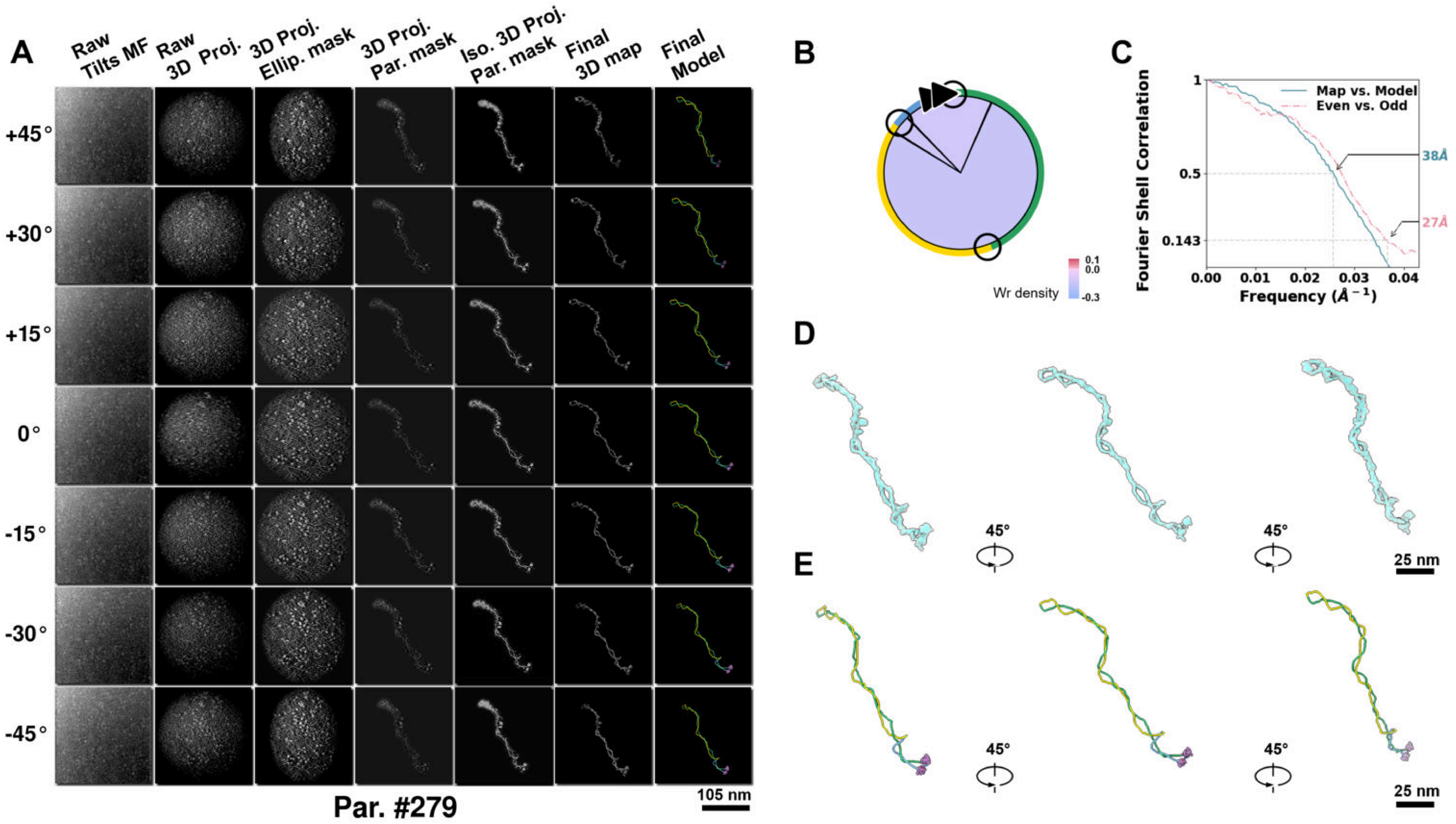

**Supplementary Particle Figure 279. Cryo-ET 3D reconstruction of an individual Opp.-TEC particle.**
(A) 3D reconstruction of the plasmid particle (index no. 279). The first column shows seven representative tilt images from +45° to -45° in step of 15°. The second, third, and fourth columns show 3D projections of the particle with spherical, ellipsoidal (thinner along the z-dimension), and particle-shaped masks, respectively. The fifth column displays the 3D projections of the enhanced and IsoNet missing-wedge-corrected particle. The sixth and seventh columns present the final 3D map and the flexibly fitted model, respectively. (B) Circular schematic representation of a plasmid particle. The outer rim is color-coded to match the corresponding 3D model. Arrowheads indicate the transcriptional direction of bound RNAPs, and circles denote apical sites. Inner circular sectors represent individual plectonemes, with colors indicating writhe density (blue to red scale, −0.3 to 0.1). (C) Resolution assessment of the final 3D map using Fourier shell correlation (FSC). Two criteria are shown: FSC between two half-maps reconstructed from even and odd frames (evaluated at 0.143) and FSC between the final 3D map and the fitted model (evaluated at 0.5). (D) Zoomed-in views of the final 3D density map from panel A, displayed at two contour levels. (E) Superimposition of the high-contour level map from panel D onto its fitted model.

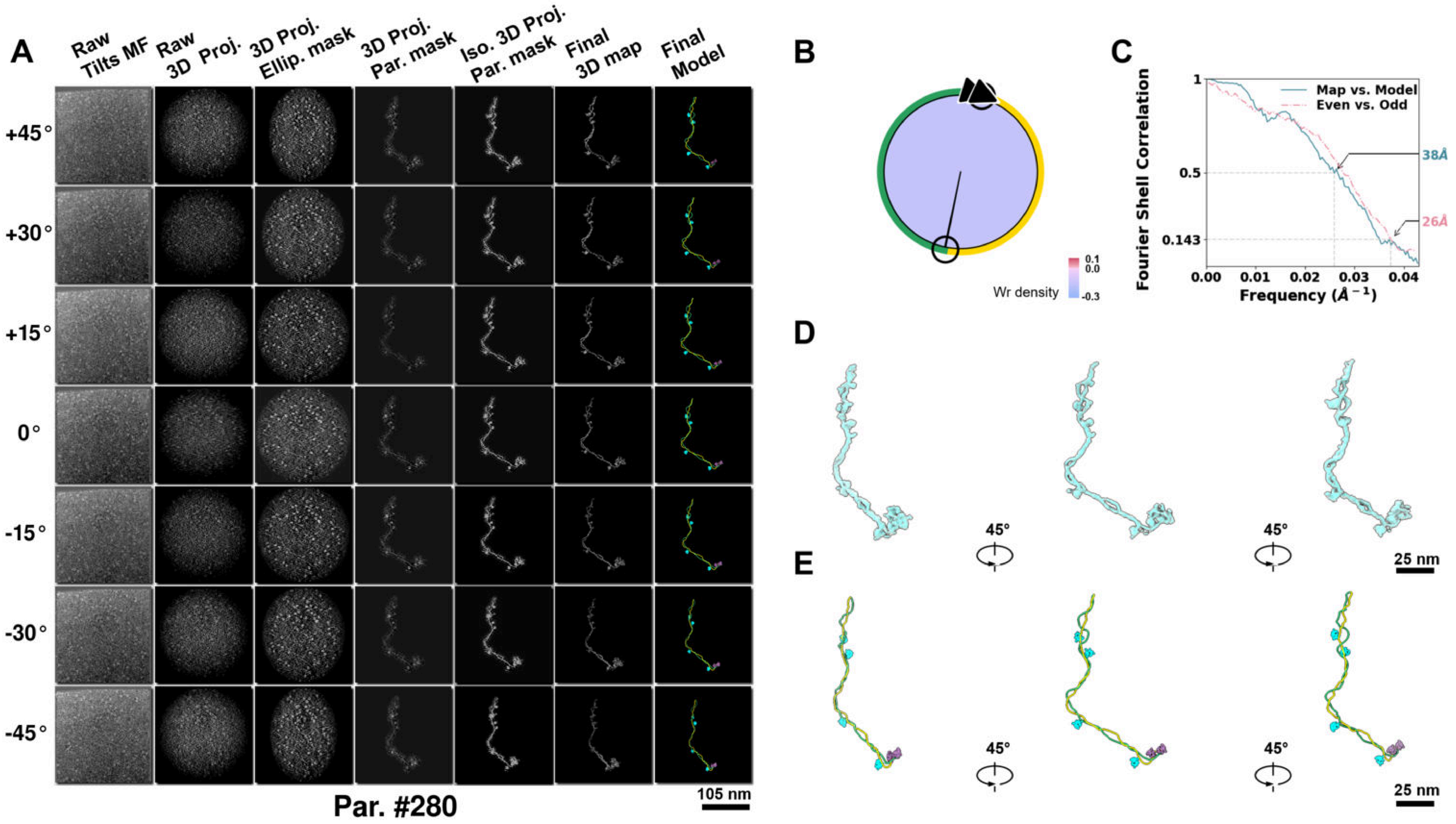

**Supplementary Particle Figure 280. Cryo-ET 3D reconstruction of an individual Opp.-TEC particle.**
(A) 3D reconstruction of the plasmid particle (index no. 280). The first column shows seven representative tilt images from +45° to -45° in step of 15°. The second, third, and fourth columns show 3D projections of the particle with spherical, ellipsoidal (thinner along the z-dimension), and particle-shaped masks, respectively. The fifth column displays the 3D projections of the enhanced and IsoNet missing-wedge-corrected particle. The sixth and seventh columns present the final 3D map and the flexibly fitted model, respectively. (B) Circular schematic representation of a plasmid particle. The outer rim is color-coded to match the corresponding 3D model. Arrowheads indicate the transcriptional direction of bound RNAPs, and circles denote apical sites. Inner circular sectors represent individual plectonemes, with colors indicating writhe density (blue to red scale, −0.3 to 0.1). (C) Resolution assessment of the final 3D map using Fourier shell correlation (FSC). Two criteria are shown: FSC between two half-maps reconstructed from even and odd frames (evaluated at 0.143) and FSC between the final 3D map and the fitted model (evaluated at 0.5). (D) Zoomed-in views of the final 3D density map from panel A, displayed at two contour levels. (E) Superimposition of the high-contour level map from panel D onto its fitted model.

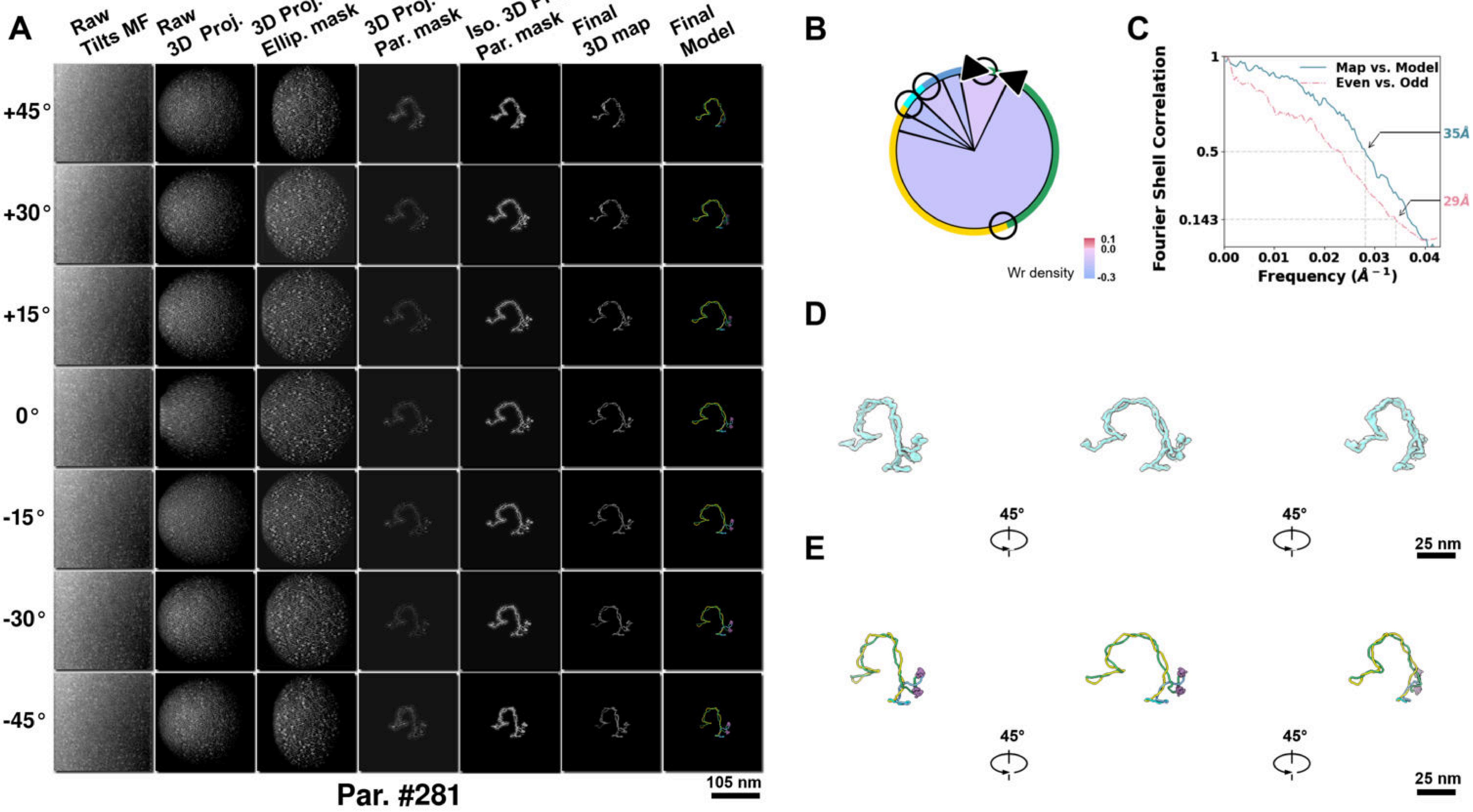

**Supplementary Particle Figure 281. Cryo-ET 3D reconstruction of an individual Opp.-TEC particle.**
(A) 3D reconstruction of the plasmid particle (index no. 281). The first column shows seven representative tilt images from +45° to -45° in step of 15°. The second, third, and fourth columns show 3D projections of the particle with spherical, ellipsoidal (thinner along the z-dimension), and particle-shaped masks, respectively. The fifth column displays the 3D projections of the enhanced and IsoNet missing-wedge-corrected particle. The sixth and seventh columns present the final 3D map and the flexibly fitted model, respectively. (B) Circular schematic representation of a plasmid particle. The outer rim is color-coded to match the corresponding 3D model. Arrowheads indicate the transcriptional direction of bound RNAPs, and circles denote apical sites. Inner circular sectors represent individual plectonemes, with colors indicating writhe density (blue to red scale, −0.3 to 0.1). (C) Resolution assessment of the final 3D map using Fourier shell correlation (FSC). Two criteria are shown: FSC between two half-maps reconstructed from even and odd frames (evaluated at 0.143) and FSC between the final 3D map and the fitted model (evaluated at 0.5). (D) Zoomed-in views of the final 3D density map from panel A, displayed at two contour levels. (E) Superimposition of the high-contour level map from panel D onto its fitted model.

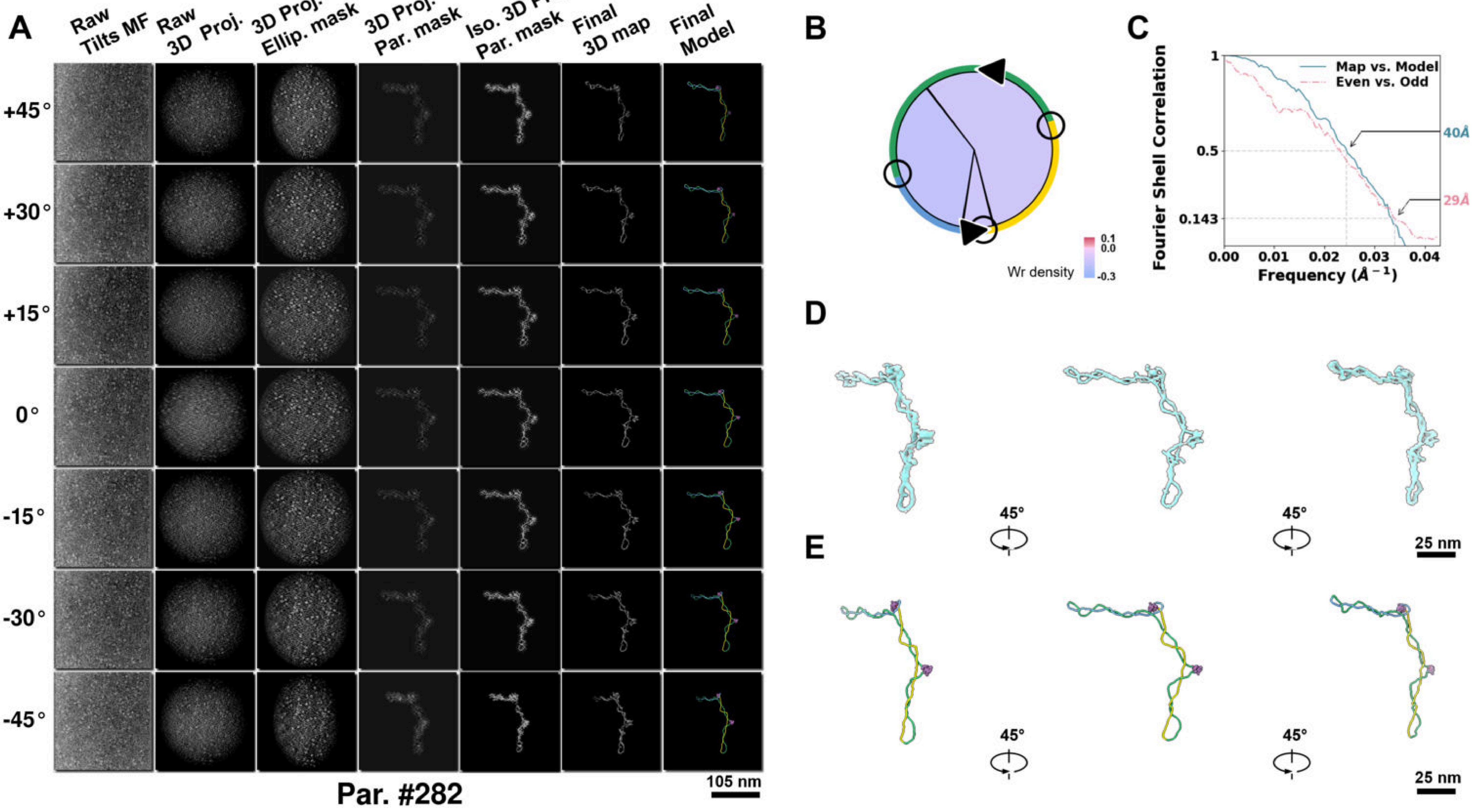

**Supplementary Particle Figure 282. Cryo-ET 3D reconstruction of an individual Opp.-TEC particle.**
(A) 3D reconstruction of the plasmid particle (index no. 282). The first column shows seven representative tilt images from +45° to -45° in step of 15°. The second, third, and fourth columns show 3D projections of the particle with spherical, ellipsoidal (thinner along the z-dimension), and particle-shaped masks, respectively. The fifth column displays the 3D projections of the enhanced and IsoNet missing-wedge-corrected particle. The sixth and seventh columns present the final 3D map and the flexibly fitted model, respectively. (B) Circular schematic representation of a plasmid particle. The outer rim is color-coded to match the corresponding 3D model. Arrowheads indicate the transcriptional direction of bound RNAPs, and circles denote apical sites. Inner circular sectors represent individual plectonemes, with colors indicating writhe density (blue to red scale, −0.3 to 0.1). (C) Resolution assessment of the final 3D map using Fourier shell correlation (FSC). Two criteria are shown: FSC between two half-maps reconstructed from even and odd frames (evaluated at 0.143) and FSC between the final 3D map and the fitted model (evaluated at 0.5). (D) Zoomed-in views of the final 3D density map from panel A, displayed at two contour levels. (E) Superimposition of the high-contour level map from panel D onto its fitted model.

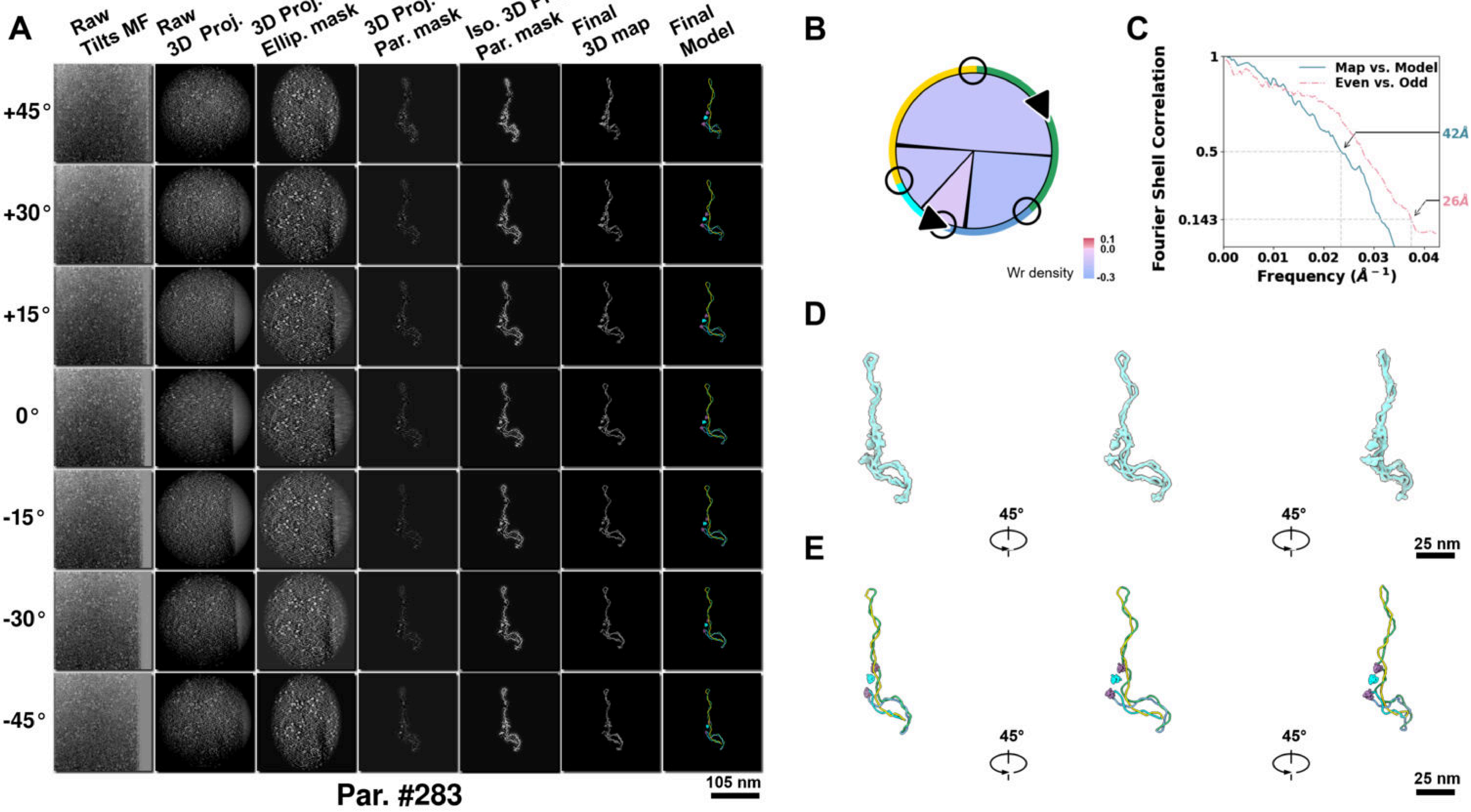

**Supplementary Particle Figure 283. Cryo-ET 3D reconstruction of an individual Opp.-TEC particle.**
(A) 3D reconstruction of the plasmid particle (index no. 283). The first column shows seven representative tilt images from +45° to -45° in step of 15°. The second, third, and fourth columns show 3D projections of the particle with spherical, ellipsoidal (thinner along the z-dimension), and particle-shaped masks, respectively. The fifth column displays the 3D projections of the enhanced and IsoNet missing-wedge-corrected particle. The sixth and seventh columns present the final 3D map and the flexibly fitted model, respectively. (B) Circular schematic representation of a plasmid particle. The outer rim is color-coded to match the corresponding 3D model. Arrowheads indicate the transcriptional direction of bound RNAPs, and circles denote apical sites. Inner circular sectors represent individual plectonemes, with colors indicating writhe density (blue to red scale, −0.3 to 0.1). (C) Resolution assessment of the final 3D map using Fourier shell correlation (FSC). Two criteria are shown: FSC between two half-maps reconstructed from even and odd frames (evaluated at 0.143) and FSC between the final 3D map and the fitted model (evaluated at 0.5). (D) Zoomed-in views of the final 3D density map from panel A, displayed at two contour levels. (E) Superimposition of the high-contour level map from panel D onto its fitted model.

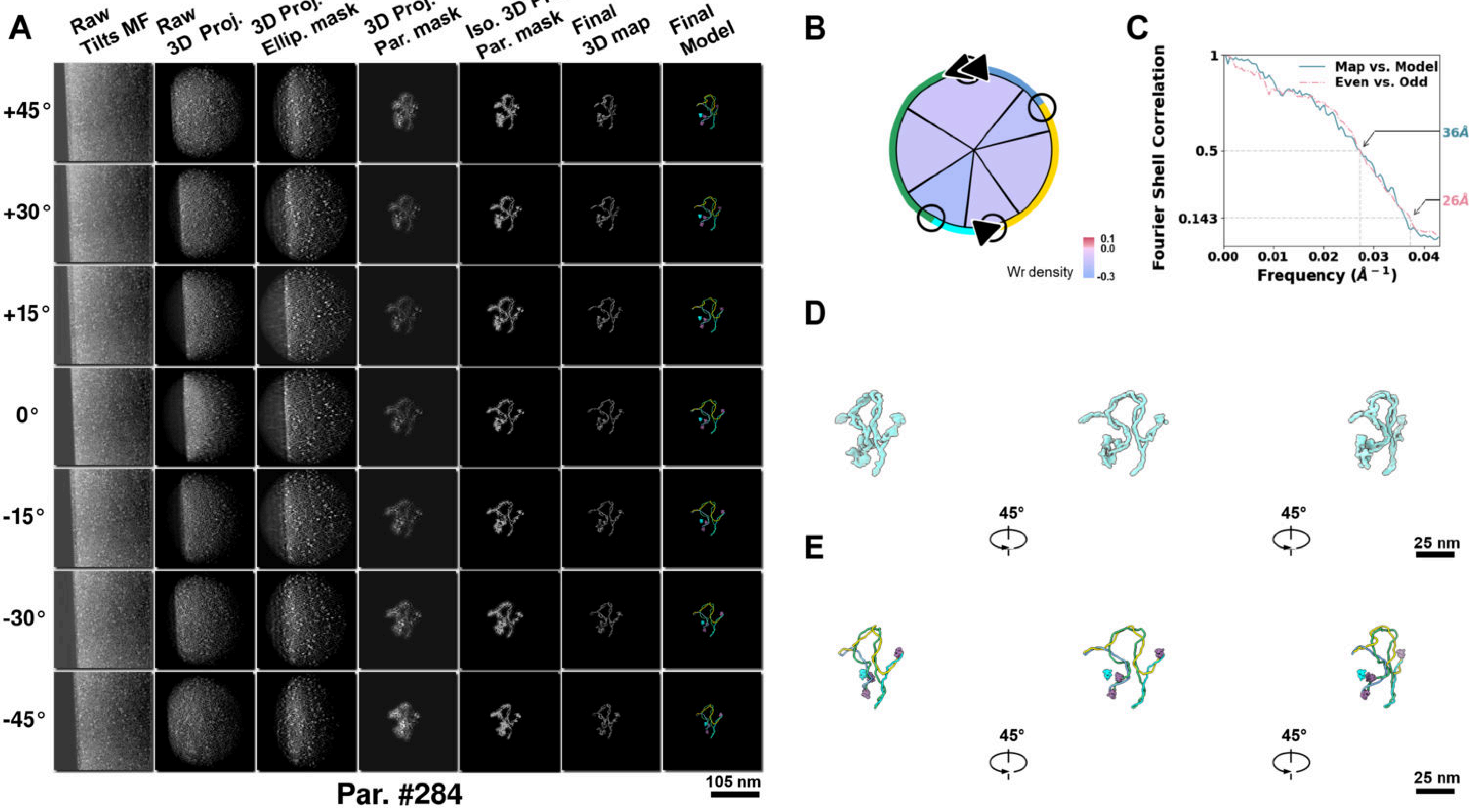

**Supplementary Particle Figure 284. Cryo-ET 3D reconstruction of an individual Opp.-TEC particle.**
(A) 3D reconstruction of the plasmid particle (index no. 284). The first column shows seven representative tilt images from +45° to -45° in step of 15°. The second, third, and fourth columns show 3D projections of the particle with spherical, ellipsoidal (thinner along the z-dimension), and particle-shaped masks, respectively. The fifth column displays the 3D projections of the enhanced and IsoNet missing-wedge-corrected particle. The sixth and seventh columns present the final 3D map and the flexibly fitted model, respectively. (B) Circular schematic representation of a plasmid particle. The outer rim is color-coded to match the corresponding 3D model. Arrowheads indicate the transcriptional direction of bound RNAPs, and circles denote apical sites. Inner circular sectors represent individual plectonemes, with colors indicating writhe density (blue to red scale, −0.3 to 0.1). (C) Resolution assessment of the final 3D map using Fourier shell correlation (FSC). Two criteria are shown: FSC between two half-maps reconstructed from even and odd frames (evaluated at 0.143) and FSC between the final 3D map and the fitted model (evaluated at 0.5). (D) Zoomed-in views of the final 3D density map from panel A, displayed at two contour levels. (E) Superimposition of the high-contour level map from panel D onto its fitted model.

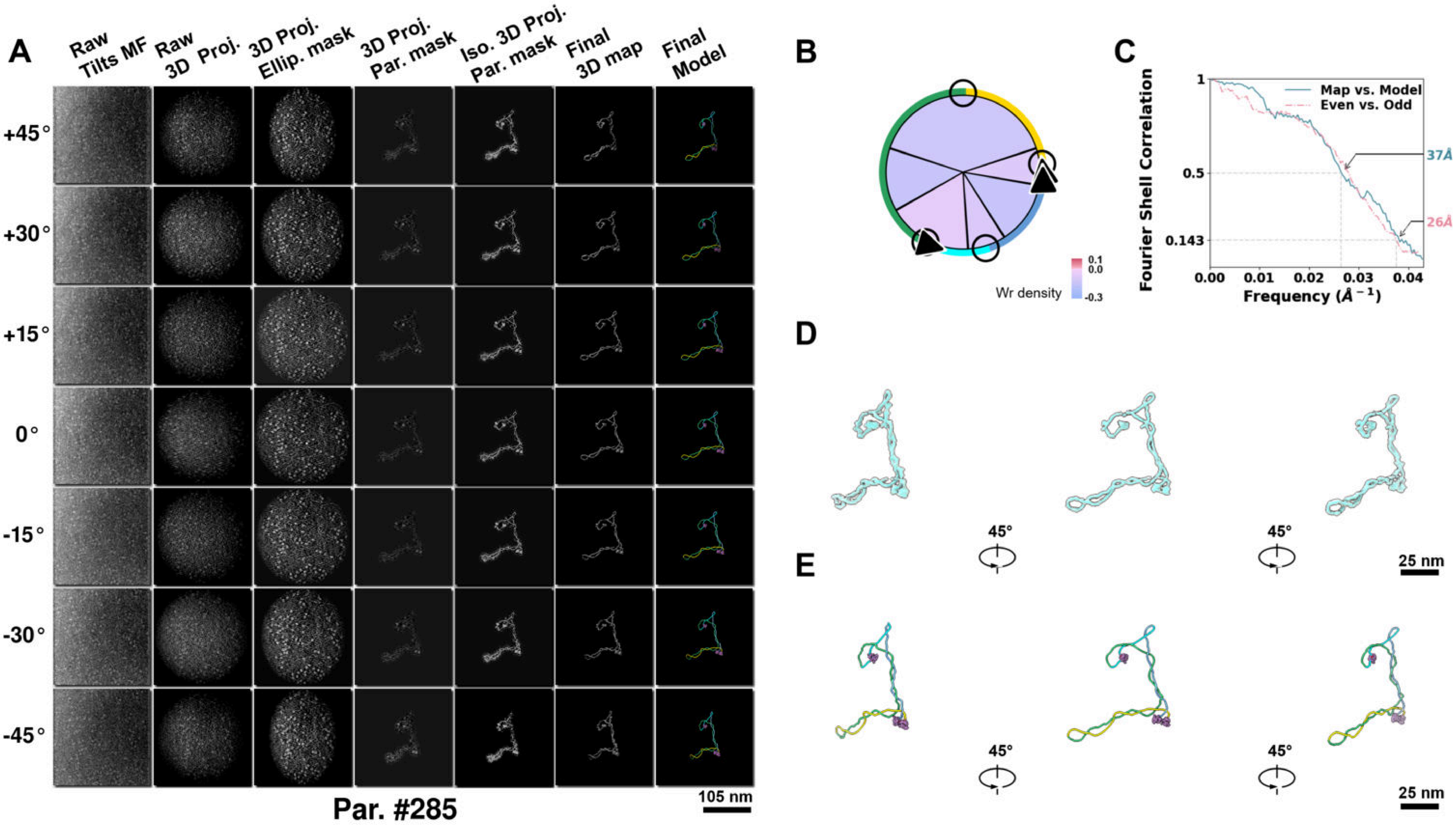

**Supplementary Particle Figure 285. Cryo-ET 3D reconstruction of an individual Opp.-TEC particle.**
(A) 3D reconstruction of the plasmid particle (index no. 285). The first column shows seven representative tilt images from +45° to -45° in step of 15°. The second, third, and fourth columns show 3D projections of the particle with spherical, ellipsoidal (thinner along the z-dimension), and particle-shaped masks, respectively. The fifth column displays the 3D projections of the enhanced and IsoNet missing-wedge-corrected particle. The sixth and seventh columns present the final 3D map and the flexibly fitted model, respectively. (B) Circular schematic representation of a plasmid particle. The outer rim is color-coded to match the corresponding 3D model. Arrowheads indicate the transcriptional direction of bound RNAPs, and circles denote apical sites. Inner circular sectors represent individual plectonemes, with colors indicating writhe density (blue to red scale, −0.3 to 0.1). (C) Resolution assessment of the final 3D map using Fourier shell correlation (FSC). Two criteria are shown: FSC between two half-maps reconstructed from even and odd frames (evaluated at 0.143) and FSC between the final 3D map and the fitted model (evaluated at 0.5). (D) Zoomed-in views of the final 3D density map from panel A, displayed at two contour levels. (E) Superimposition of the high-contour level map from panel D onto its fitted model.

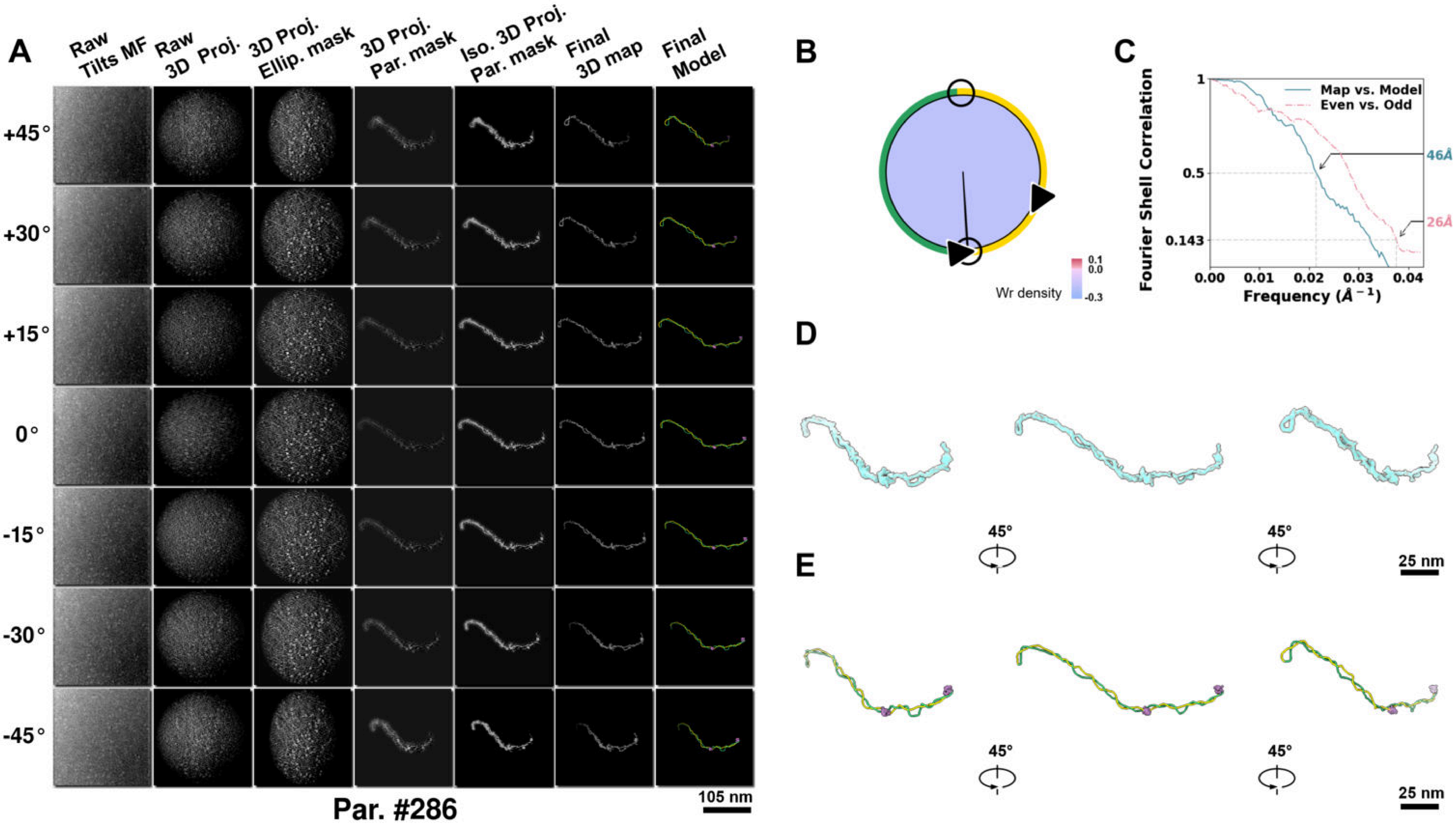

**Supplementary Particle Figure 286. Cryo-ET 3D reconstruction of an individual Opp.-TEC particle.**
(A) 3D reconstruction of the plasmid particle (index no. 286). The first column shows seven representative tilt images from +45° to -45° in step of 15°. The second, third, and fourth columns show 3D projections of the particle with spherical, ellipsoidal (thinner along the z-dimension), and particle-shaped masks, respectively. The fifth column displays the 3D projections of the enhanced and IsoNet missing-wedge-corrected particle. The sixth and seventh columns present the final 3D map and the flexibly fitted model, respectively. (B) Circular schematic representation of a plasmid particle. The outer rim is color-coded to match the corresponding 3D model. Arrowheads indicate the transcriptional direction of bound RNAPs, and circles denote apical sites. Inner circular sectors represent individual plectonemes, with colors indicating writhe density (blue to red scale, −0.3 to 0.1). (C) Resolution assessment of the final 3D map using Fourier shell correlation (FSC). Two criteria are shown: FSC between two half-maps reconstructed from even and odd frames (evaluated at 0.143) and FSC between the final 3D map and the fitted model (evaluated at 0.5). (D) Zoomed-in views of the final 3D density map from panel A, displayed at two contour levels. (E) Superimposition of the high-contour level map from panel D onto its fitted model.

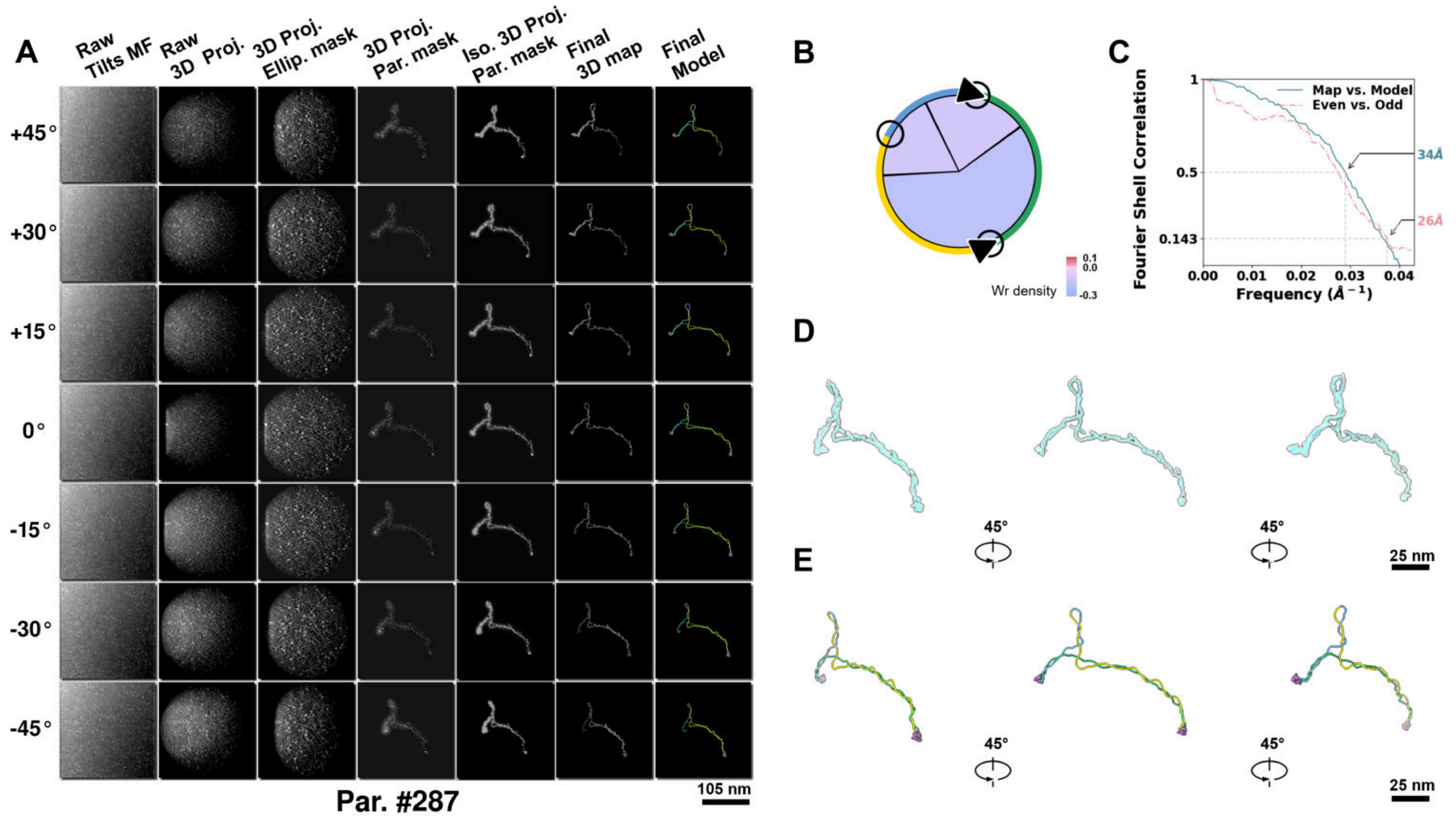

**Supplementary Particle Figure 287. Cryo-ET 3D reconstruction of an individual Opp.-TEC particle.**
(A) 3D reconstruction of the plasmid particle (index no. 287). The first column shows seven representative tilt images from +45° to -45° in step of 15°. The second, third, and fourth columns show 3D projections of the particle with spherical, ellipsoidal (thinner along the z-dimension), and particle-shaped masks, respectively. The fifth column displays the 3D projections of the enhanced and IsoNet missing-wedge-corrected particle. The sixth and seventh columns present the final 3D map and the flexibly fitted model, respectively. (B) Circular schematic representation of a plasmid particle. The outer rim is color-coded to match the corresponding 3D model. Arrowheads indicate the transcriptional direction of bound RNAPs, and circles denote apical sites. Inner circular sectors represent individual plectonemes, with colors indicating writhe density (blue to red scale, −0.3 to 0.1). (C) Resolution assessment of the final 3D map using Fourier shell correlation (FSC). Two criteria are shown: FSC between two half-maps reconstructed from even and odd frames (evaluated at 0.143) and FSC between the final 3D map and the fitted model (evaluated at 0.5). (D) Zoomed-in views of the final 3D density map from panel A, displayed at two contour levels. (E) Superimposition of the high-contour level map from panel D onto its fitted model.

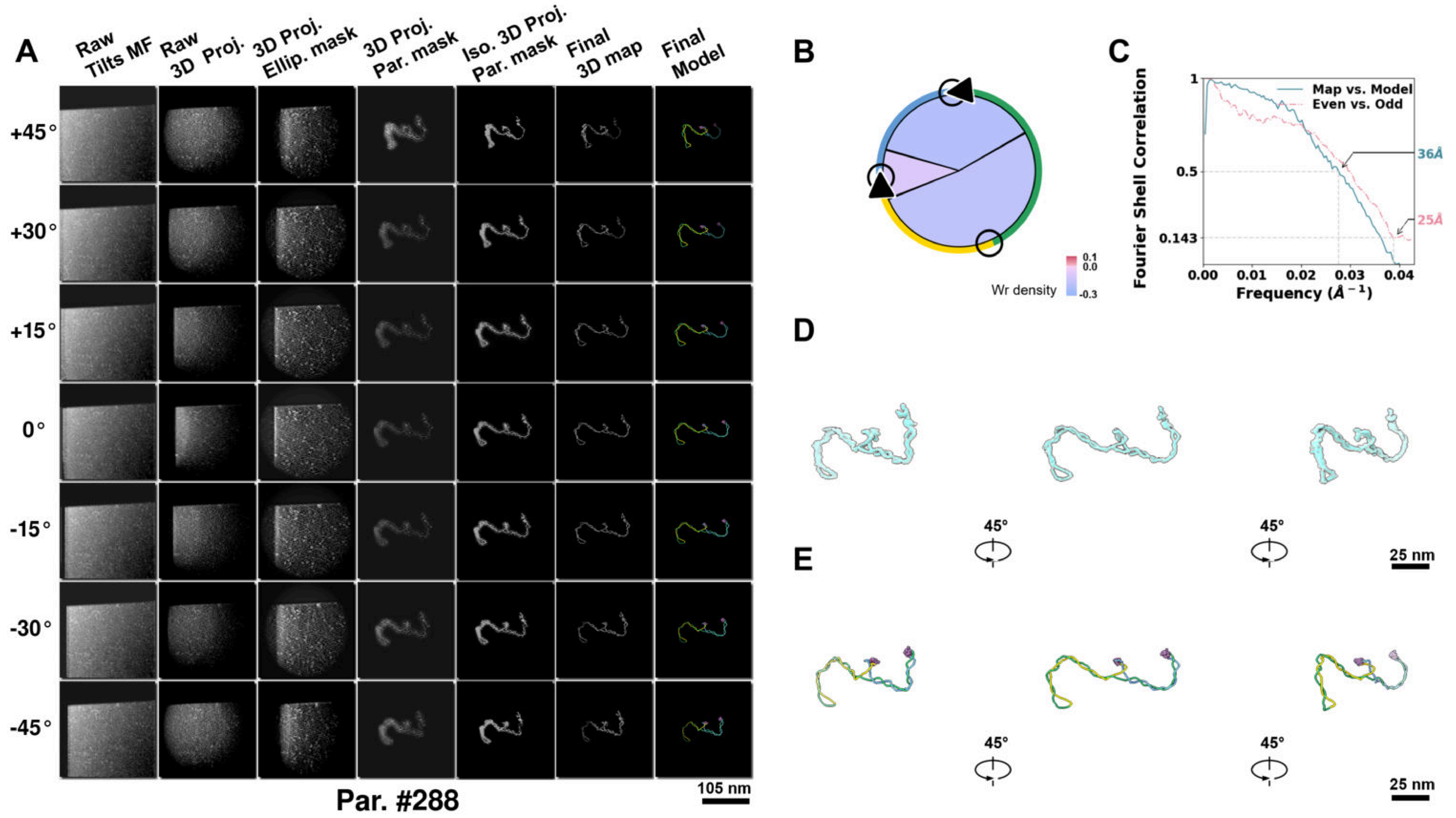

**Supplementary Particle Figure 288. Cryo-ET 3D reconstruction of an individual Opp.-TEC particle.**
(A) 3D reconstruction of the plasmid particle (index no. 288). The first column shows seven representative tilt images from +45° to -45° in step of 15°. The second, third, and fourth columns show 3D projections of the particle with spherical, ellipsoidal (thinner along the z-dimension), and particle-shaped masks, respectively. The fifth column displays the 3D projections of the enhanced and IsoNet missing-wedge-corrected particle. The sixth and seventh columns present the final 3D map and the flexibly fitted model, respectively. (B) Circular schematic representation of a plasmid particle. The outer rim is color-coded to match the corresponding 3D model. Arrowheads indicate the transcriptional direction of bound RNAPs, and circles denote apical sites. Inner circular sectors represent individual plectonemes, with colors indicating writhe density (blue to red scale, −0.3 to 0.1). (C) Resolution assessment of the final 3D map using Fourier shell correlation (FSC). Two criteria are shown: FSC between two half-maps reconstructed from even and odd frames (evaluated at 0.143) and FSC between the final 3D map and the fitted model (evaluated at 0.5). (D) Zoomed-in views of the final 3D density map from panel A, displayed at two contour levels. (E) Superimposition of the high-contour level map from panel D onto its fitted model.

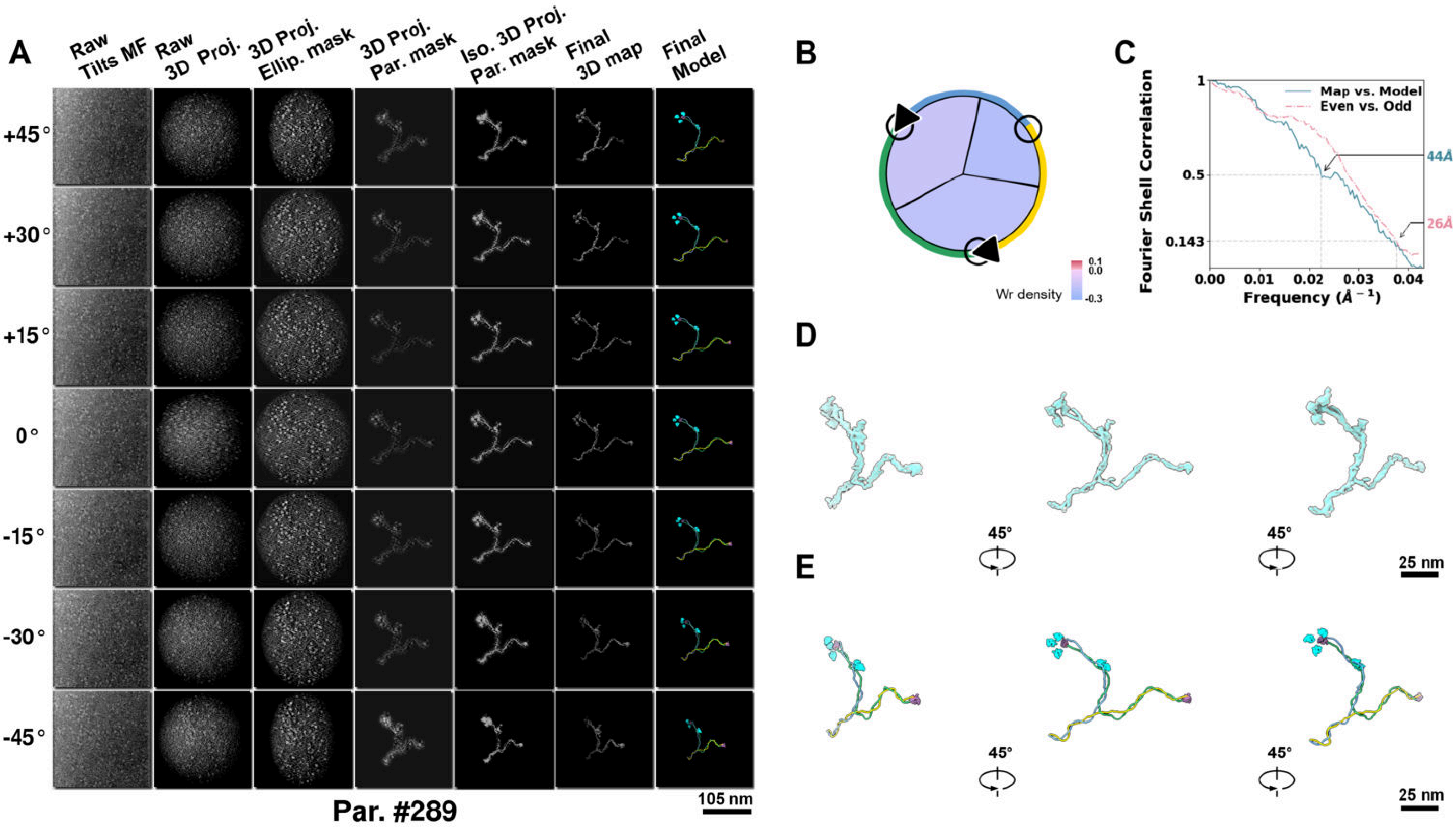

**Supplementary Particle Figure 289. Cryo-ET 3D reconstruction of an individual Opp.-TEC particle.**
(A) 3D reconstruction of the plasmid particle (index no. 289). The first column shows seven representative tilt images from +45° to -45° in step of 15°. The second, third, and fourth columns show 3D projections of the particle with spherical, ellipsoidal (thinner along the z-dimension), and particle-shaped masks, respectively. The fifth column displays the 3D projections of the enhanced and IsoNet missing-wedge-corrected particle. The sixth and seventh columns present the final 3D map and the flexibly fitted model, respectively. (B) Circular schematic representation of a plasmid particle. The outer rim is color-coded to match the corresponding 3D model. Arrowheads indicate the transcriptional direction of bound RNAPs, and circles denote apical sites. Inner circular sectors represent individual plectonemes, with colors indicating writhe density (blue to red scale, −0.3 to 0.1). (C) Resolution assessment of the final 3D map using Fourier shell correlation (FSC). Two criteria are shown: FSC between two half-maps reconstructed from even and odd frames (evaluated at 0.143) and FSC between the final 3D map and the fitted model (evaluated at 0.5). (D) Zoomed-in views of the final 3D density map from panel A, displayed at two contour levels. (E) Superimposition of the high-contour level map from panel D onto its fitted model.

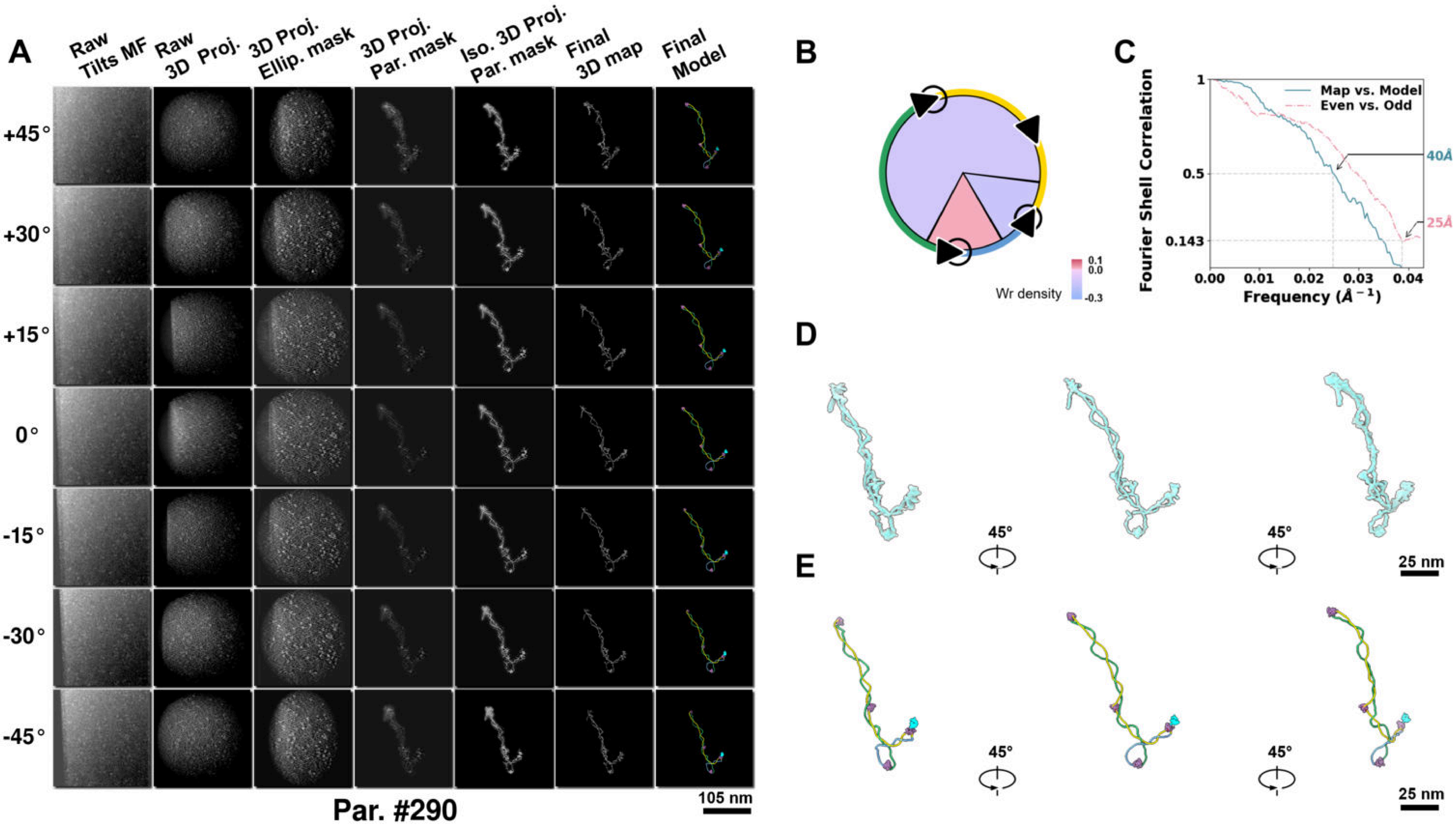

**Supplementary Particle Figure 290. Cryo-ET 3D reconstruction of an individual Opp.-TEC particle.**
(A) 3D reconstruction of the plasmid particle (index no. 290). The first column shows seven representative tilt images from +45° to -45° in step of 15°. The second, third, and fourth columns show 3D projections of the particle with spherical, ellipsoidal (thinner along the z-dimension), and particle-shaped masks, respectively. The fifth column displays the 3D projections of the enhanced and IsoNet missing-wedge-corrected particle. The sixth and seventh columns present the final 3D map and the flexibly fitted model, respectively. (B) Circular schematic representation of a plasmid particle. The outer rim is color-coded to match the corresponding 3D model. Arrowheads indicate the transcriptional direction of bound RNAPs, and circles denote apical sites. Inner circular sectors represent individual plectonemes, with colors indicating writhe density (blue to red scale, −0.3 to 0.1). (C) Resolution assessment of the final 3D map using Fourier shell correlation (FSC). Two criteria are shown: FSC between two half-maps reconstructed from even and odd frames (evaluated at 0.143) and FSC between the final 3D map and the fitted model (evaluated at 0.5). (D) Zoomed-in views of the final 3D density map from panel A, displayed at two contour levels. (E) Superimposition of the high-contour level map from panel D onto its fitted model.

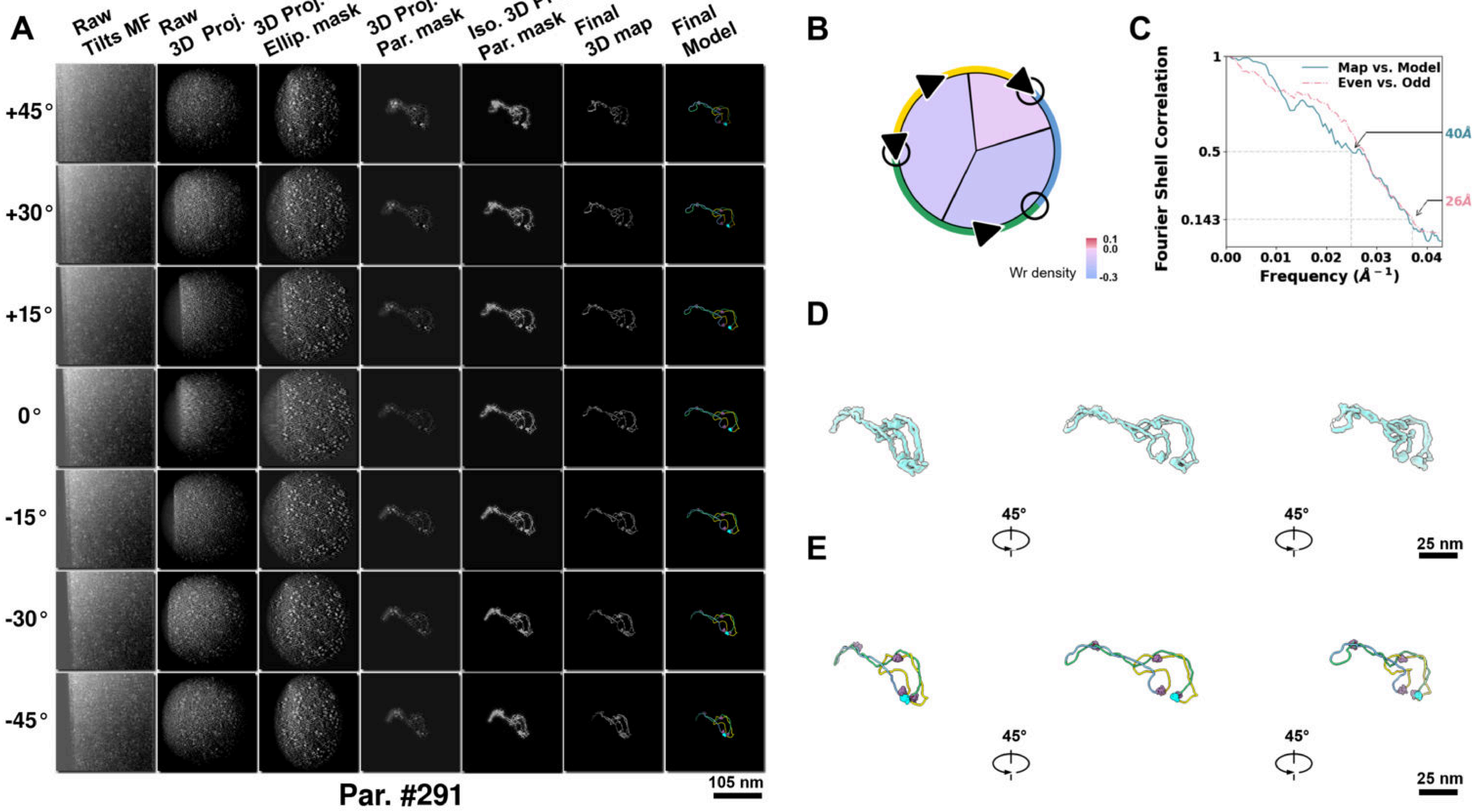

**Supplementary Particle Figure 291. Cryo-ET 3D reconstruction of an individual Opp.-TEC particle.**
(A) 3D reconstruction of the plasmid particle (index no. 291). The first column shows seven representative tilt images from +45° to -45° in step of 15°. The second, third, and fourth columns show 3D projections of the particle with spherical, ellipsoidal (thinner along the z-dimension), and particle-shaped masks, respectively. The fifth column displays the 3D projections of the enhanced and IsoNet missing-wedge-corrected particle. The sixth and seventh columns present the final 3D map and the flexibly fitted model, respectively. (B) Circular schematic representation of a plasmid particle. The outer rim is color-coded to match the corresponding 3D model. Arrowheads indicate the transcriptional direction of bound RNAPs, and circles denote apical sites. Inner circular sectors represent individual plectonemes, with colors indicating writhe density (blue to red scale, −0.3 to 0.1). (C) Resolution assessment of the final 3D map using Fourier shell correlation (FSC). Two criteria are shown: FSC between two half-maps reconstructed from even and odd frames (evaluated at 0.143) and FSC between the final 3D map and the fitted model (evaluated at 0.5). (D) Zoomed-in views of the final 3D density map from panel A, displayed at two contour levels. (E) Superimposition of the high-contour level map from panel D onto its fitted model.

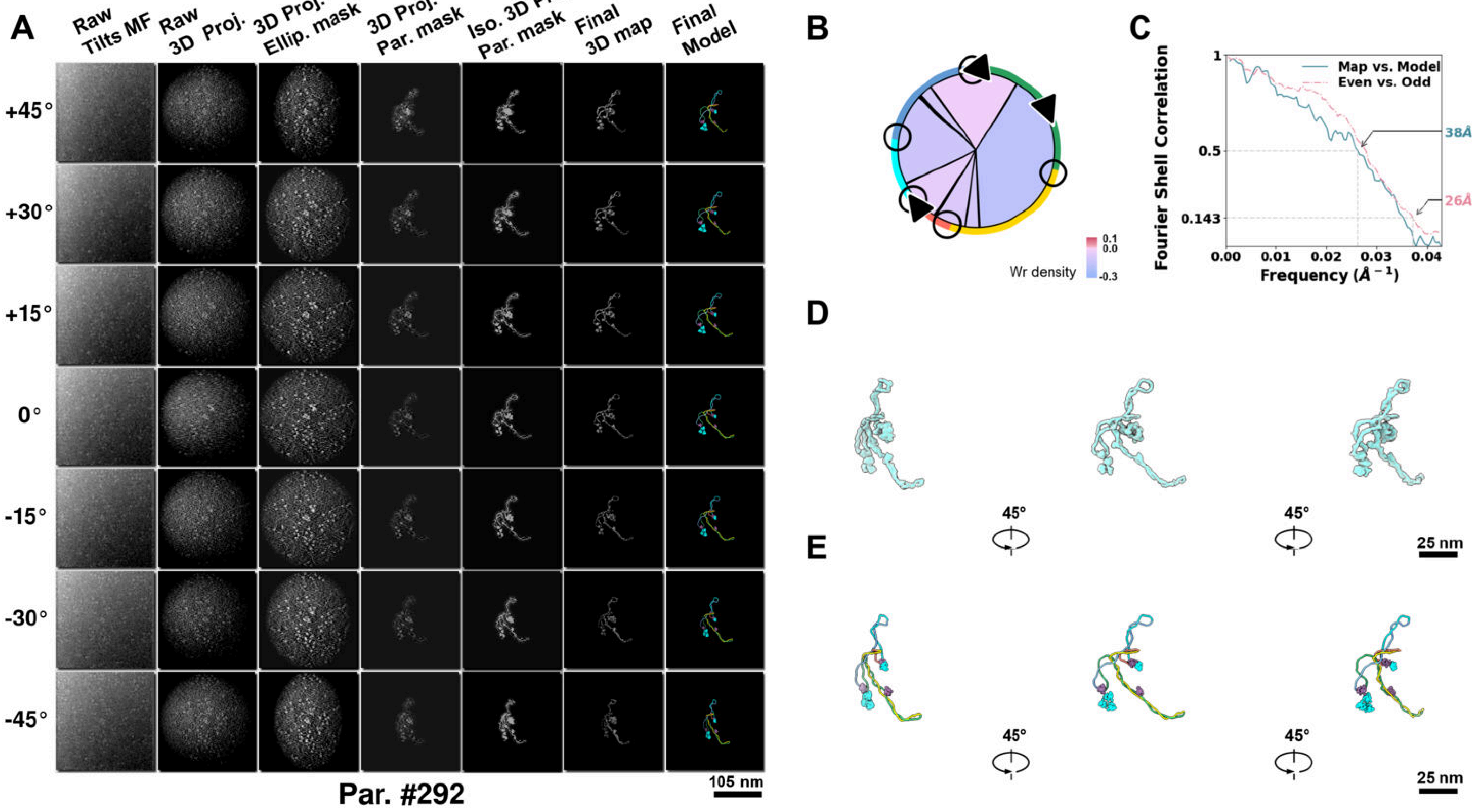

**Supplementary Particle Figure 292. Cryo-ET 3D reconstruction of an individual Opp.-TEC particle.**
(A) 3D reconstruction of the plasmid particle (index no. 292). The first column shows seven representative tilt images from +45° to -45° in step of 15°. The second, third, and fourth columns show 3D projections of the particle with spherical, ellipsoidal (thinner along the z-dimension), and particle-shaped masks, respectively. The fifth column displays the 3D projections of the enhanced and IsoNet missing-wedge-corrected particle. The sixth and seventh columns present the final 3D map and the flexibly fitted model, respectively. (B) Circular schematic representation of a plasmid particle. The outer rim is color-coded to match the corresponding 3D model. Arrowheads indicate the transcriptional direction of bound RNAPs, and circles denote apical sites. Inner circular sectors represent individual plectonemes, with colors indicating writhe density (blue to red scale, −0.3 to 0.1). (C) Resolution assessment of the final 3D map using Fourier shell correlation (FSC). Two criteria are shown: FSC between two half-maps reconstructed from even and odd frames (evaluated at 0.143) and FSC between the final 3D map and the fitted model (evaluated at 0.5). (D) Zoomed-in views of the final 3D density map from panel A, displayed at two contour levels. (E) Superimposition of the high-contour level map from panel D onto its fitted model.

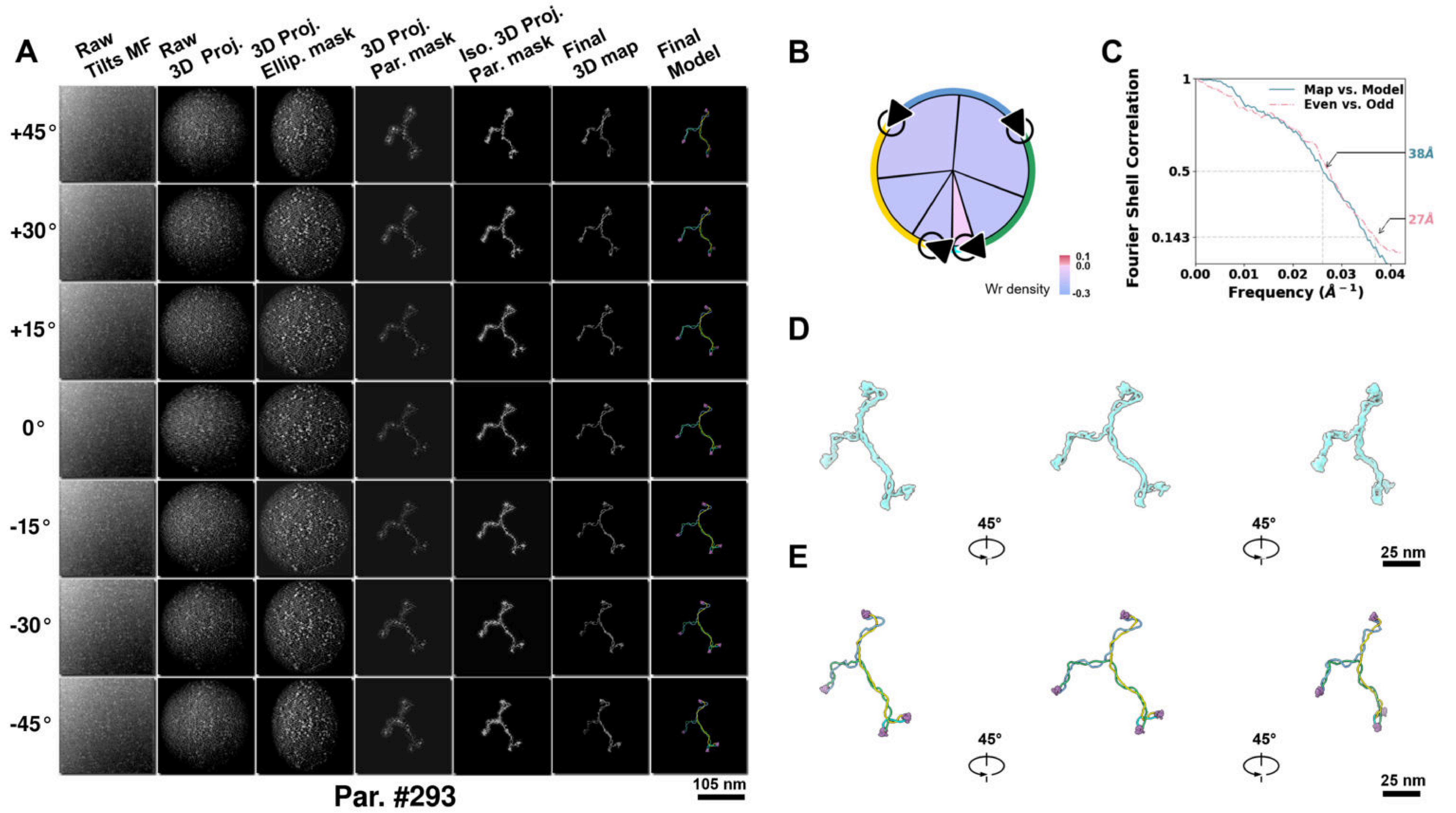

**Supplementary Particle Figure 293. Cryo-ET 3D reconstruction of an individual Opp.-TEC particle.**
(A) 3D reconstruction of the plasmid particle (index no. 293). The first column shows seven representative tilt images from +45° to -45° in step of 15°. The second, third, and fourth columns show 3D projections of the particle with spherical, ellipsoidal (thinner along the z-dimension), and particle-shaped masks, respectively. The fifth column displays the 3D projections of the enhanced and IsoNet missing-wedge-corrected particle. The sixth and seventh columns present the final 3D map and the flexibly fitted model, respectively. (B) Circular schematic representation of a plasmid particle. The outer rim is color-coded to match the corresponding 3D model. Arrowheads indicate the transcriptional direction of bound RNAPs, and circles denote apical sites. Inner circular sectors represent individual plectonemes, with colors indicating writhe density (blue to red scale, −0.3 to 0.1). (C) Resolution assessment of the final 3D map using Fourier shell correlation (FSC). Two criteria are shown: FSC between two half-maps reconstructed from even and odd frames (evaluated at 0.143) and FSC between the final 3D map and the fitted model (evaluated at 0.5). (D) Zoomed-in views of the final 3D density map from panel A, displayed at two contour levels. (E) Superimposition of the high-contour level map from panel D onto its fitted model.

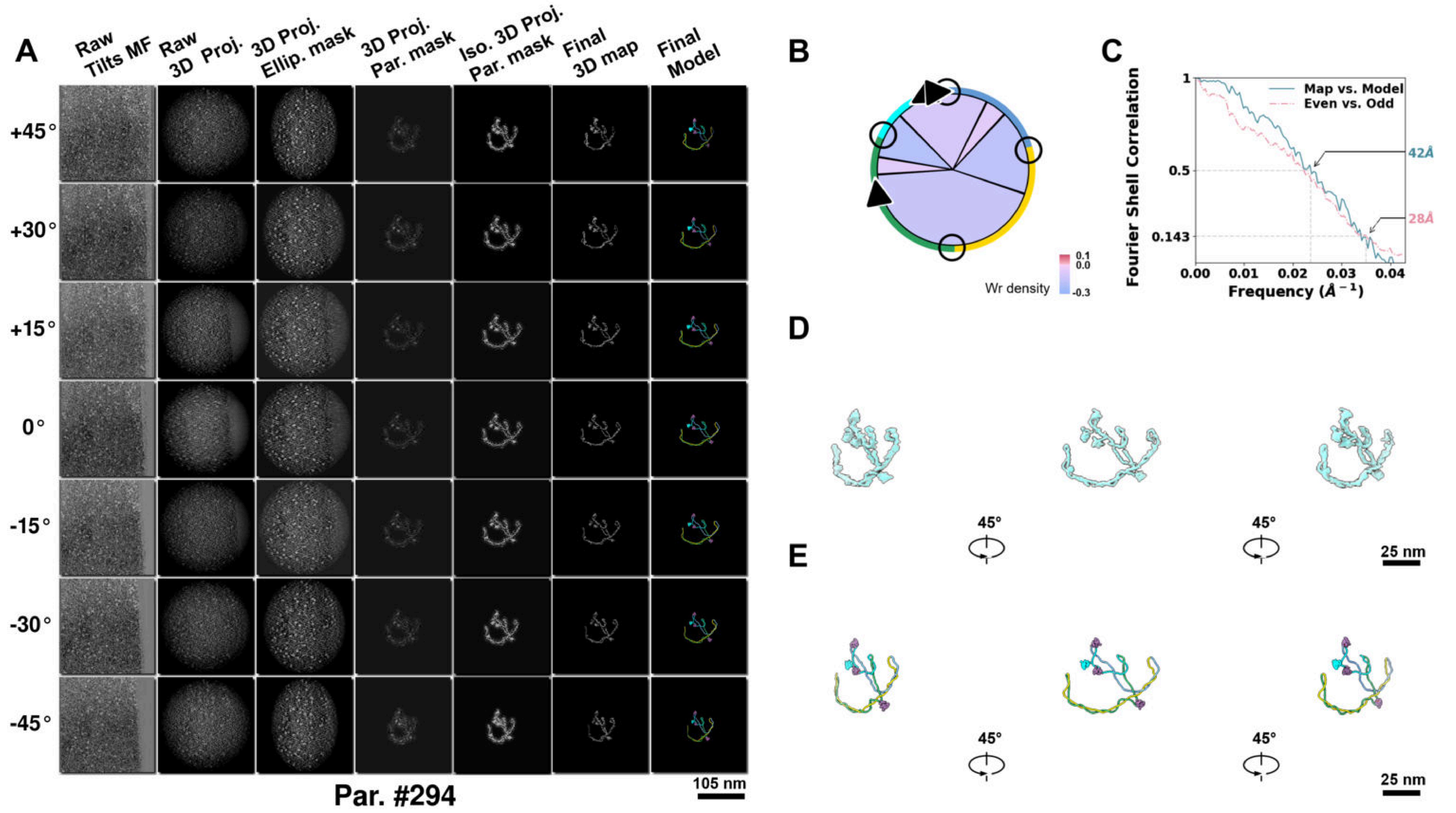

**Supplementary Particle Figure 294. Cryo-ET 3D reconstruction of an individual Opp.-TEC particle.**
(A) 3D reconstruction of the plasmid particle (index no. 294). The first column shows seven representative tilt images from +45° to -45° in step of 15°. The second, third, and fourth columns show 3D projections of the particle with spherical, ellipsoidal (thinner along the z-dimension), and particle-shaped masks, respectively. The fifth column displays the 3D projections of the enhanced and IsoNet missing-wedge-corrected particle. The sixth and seventh columns present the final 3D map and the flexibly fitted model, respectively. (B) Circular schematic representation of a plasmid particle. The outer rim is color-coded to match the corresponding 3D model. Arrowheads indicate the transcriptional direction of bound RNAPs, and circles denote apical sites. Inner circular sectors represent individual plectonemes, with colors indicating writhe density (blue to red scale, −0.3 to 0.1). (C) Resolution assessment of the final 3D map using Fourier shell correlation (FSC). Two criteria are shown: FSC between two half-maps reconstructed from even and odd frames (evaluated at 0.143) and FSC between the final 3D map and the fitted model (evaluated at 0.5). (D) Zoomed-in views of the final 3D density map from panel A, displayed at two contour levels. (E) Superimposition of the high-contour level map from panel D onto its fitted model.

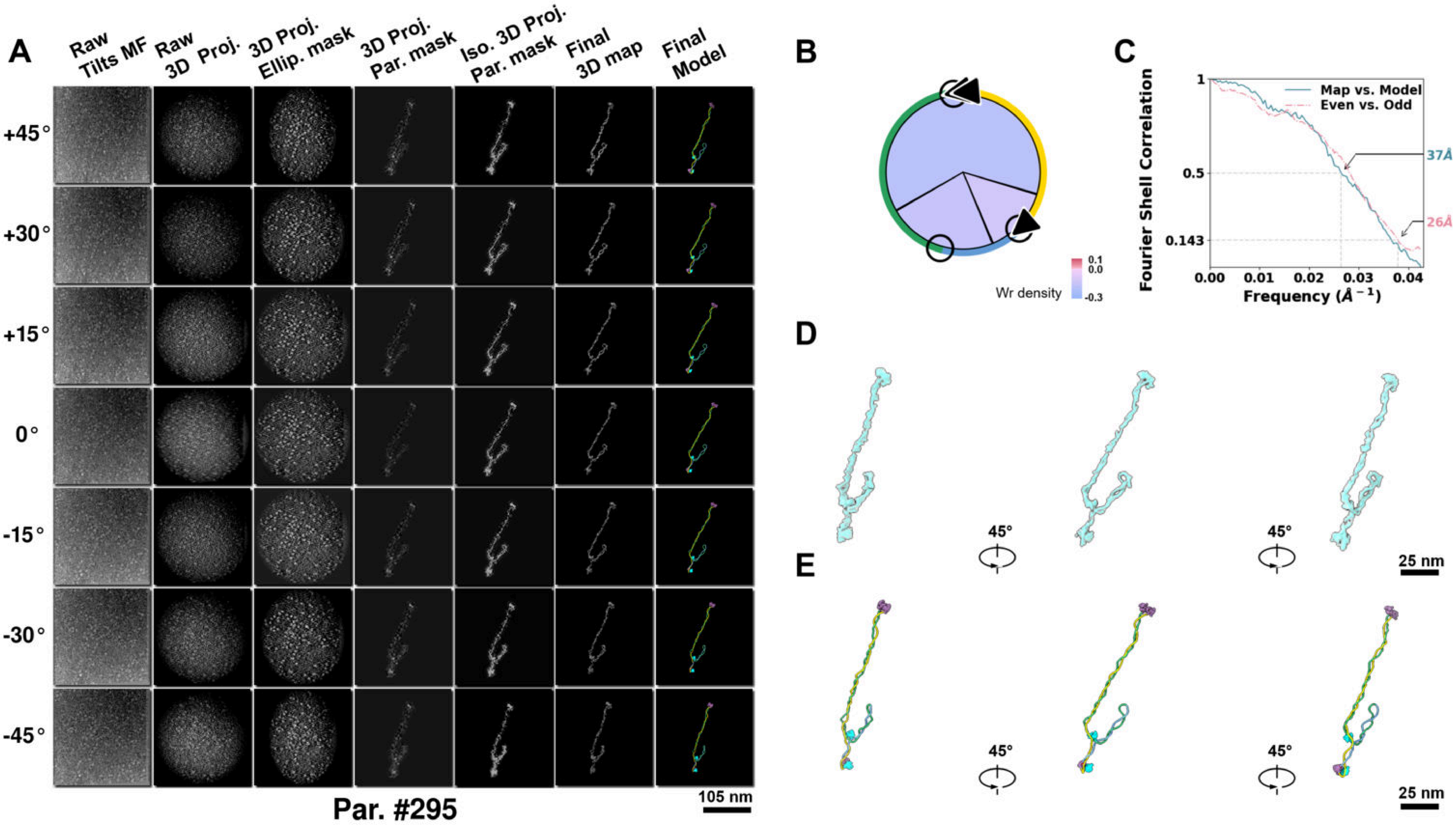

**Supplementary Particle Figure 295. Cryo-ET 3D reconstruction of an individual Opp.-TEC particle.**
(A) 3D reconstruction of the plasmid particle (index no. 295). The first column shows seven representative tilt images from +45° to -45° in step of 15°. The second, third, and fourth columns show 3D projections of the particle with spherical, ellipsoidal (thinner along the z-dimension), and particle-shaped masks, respectively. The fifth column displays the 3D projections of the enhanced and IsoNet missing-wedge-corrected particle. The sixth and seventh columns present the final 3D map and the flexibly fitted model, respectively. (B) Circular schematic representation of a plasmid particle. The outer rim is color-coded to match the corresponding 3D model. Arrowheads indicate the transcriptional direction of bound RNAPs, and circles denote apical sites. Inner circular sectors represent individual plectonemes, with colors indicating writhe density (blue to red scale, −0.3 to 0.1). (C) Resolution assessment of the final 3D map using Fourier shell correlation (FSC). Two criteria are shown: FSC between two half-maps reconstructed from even and odd frames (evaluated at 0.143) and FSC between the final 3D map and the fitted model (evaluated at 0.5). (D) Zoomed-in views of the final 3D density map from panel A, displayed at two contour levels. (E) Superimposition of the high-contour level map from panel D onto its fitted model.

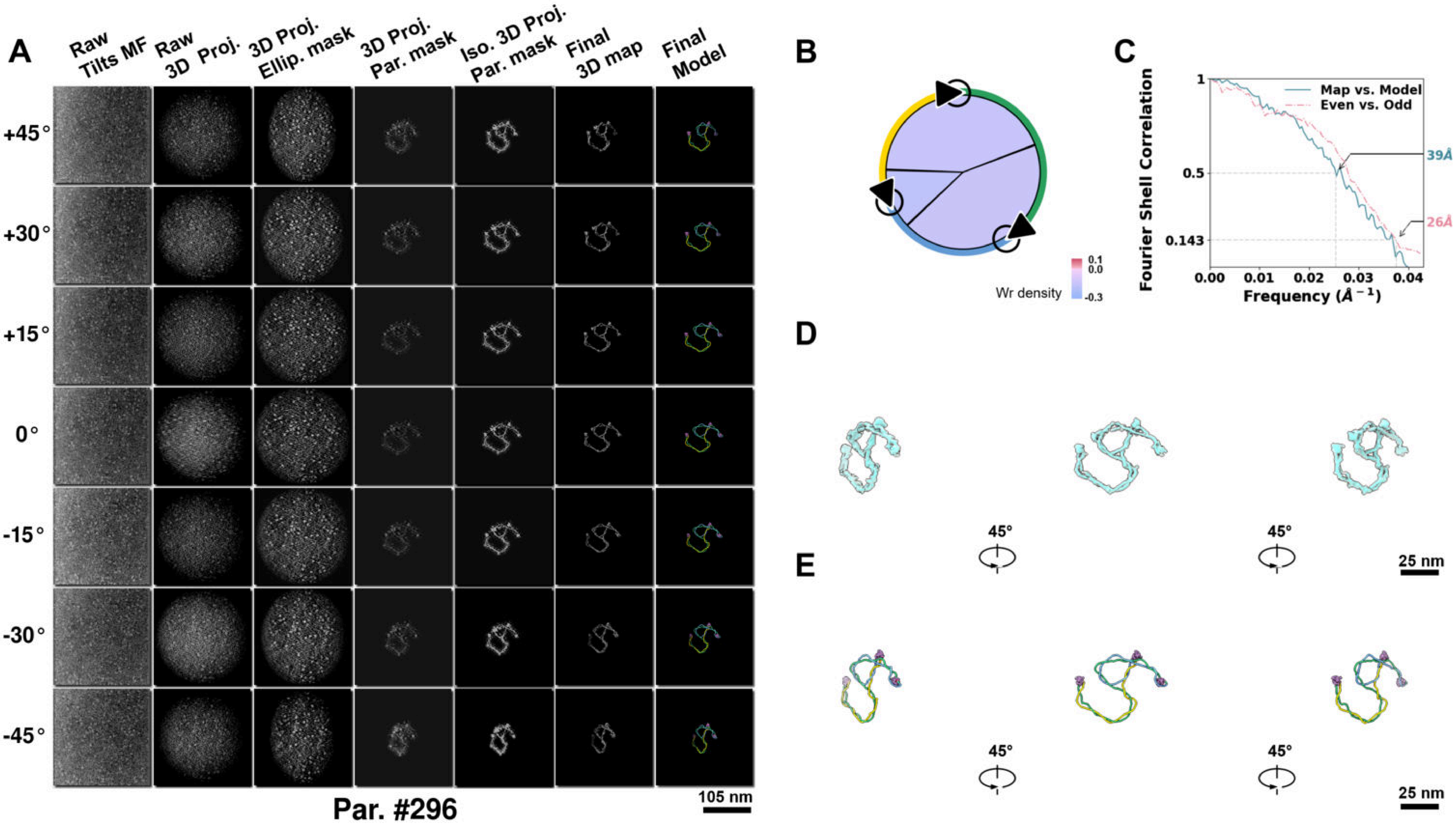

**Supplementary Particle Figure 296. Cryo-ET 3D reconstruction of an individual Opp.-TEC particle.**
(A) 3D reconstruction of the plasmid particle (index no. 296). The first column shows seven representative tilt images from +45° to -45° in step of 15°. The second, third, and fourth columns show 3D projections of the particle with spherical, ellipsoidal (thinner along the z-dimension), and particle-shaped masks, respectively. The fifth column displays the 3D projections of the enhanced and IsoNet missing-wedge-corrected particle. The sixth and seventh columns present the final 3D map and the flexibly fitted model, respectively. (B) Circular schematic representation of a plasmid particle. The outer rim is color-coded to match the corresponding 3D model. Arrowheads indicate the transcriptional direction of bound RNAPs, and circles denote apical sites. Inner circular sectors represent individual plectonemes, with colors indicating writhe density (blue to red scale, −0.3 to 0.1). (C) Resolution assessment of the final 3D map using Fourier shell correlation (FSC). Two criteria are shown: FSC between two half-maps reconstructed from even and odd frames (evaluated at 0.143) and FSC between the final 3D map and the fitted model (evaluated at 0.5). (D) Zoomed-in views of the final 3D density map from panel A, displayed at two contour levels. (E) Superimposition of the high-contour level map from panel D onto its fitted model.

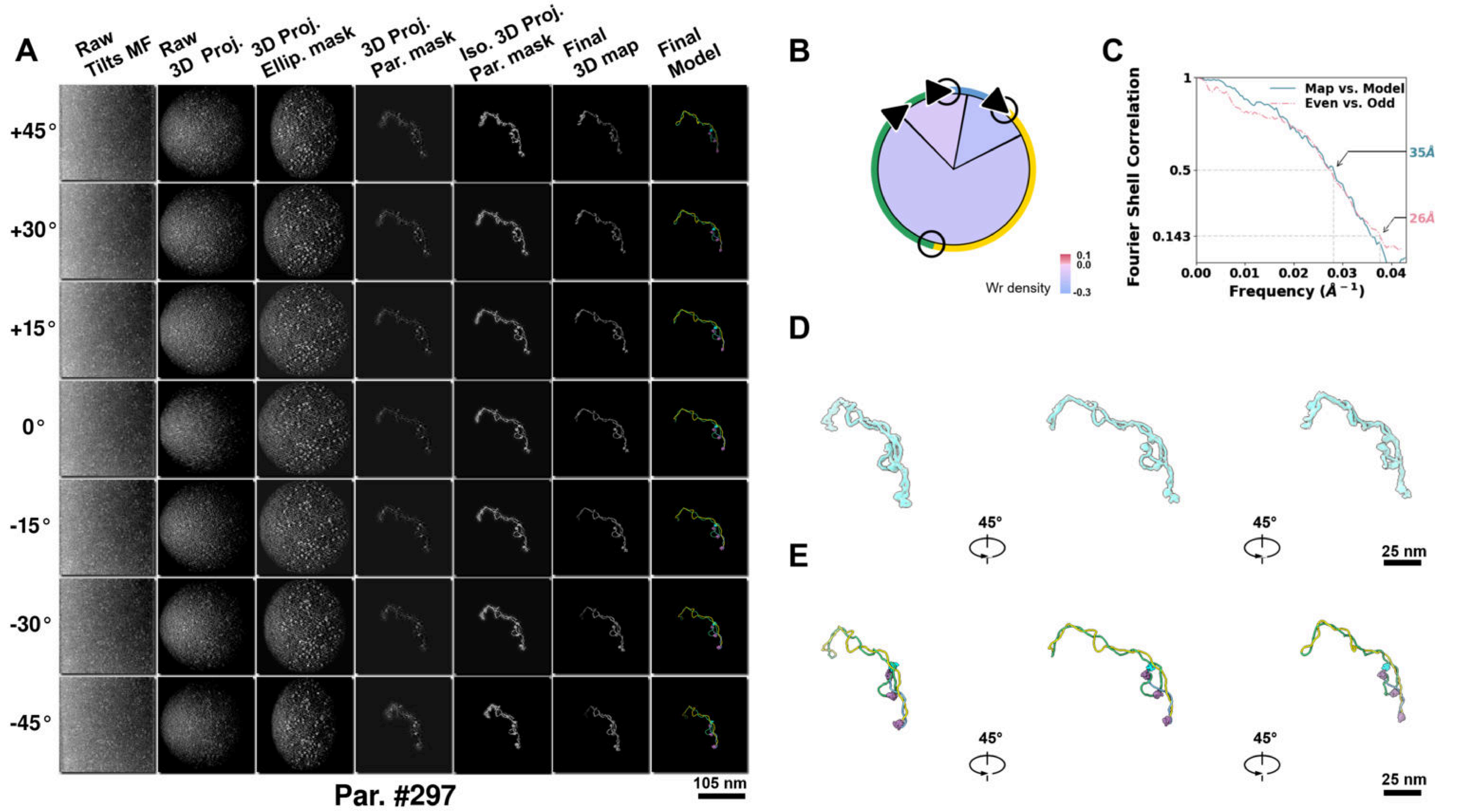

**Supplementary Particle Figure 297. Cryo-ET 3D reconstruction of an individual Opp.-TEC particle.**
(A) 3D reconstruction of the plasmid particle (index no. 297). The first column shows seven representative tilt images from +45° to -45° in step of 15°. The second, third, and fourth columns show 3D projections of the particle with spherical, ellipsoidal (thinner along the z-dimension), and particle-shaped masks, respectively. The fifth column displays the 3D projections of the enhanced and IsoNet missing-wedge-corrected particle. The sixth and seventh columns present the final 3D map and the flexibly fitted model, respectively. (B) Circular schematic representation of a plasmid particle. The outer rim is color-coded to match the corresponding 3D model. Arrowheads indicate the transcriptional direction of bound RNAPs, and circles denote apical sites. Inner circular sectors represent individual plectonemes, with colors indicating writhe density (blue to red scale, −0.3 to 0.1). (C) Resolution assessment of the final 3D map using Fourier shell correlation (FSC). Two criteria are shown: FSC between two half-maps reconstructed from even and odd frames (evaluated at 0.143) and FSC between the final 3D map and the fitted model (evaluated at 0.5). (D) Zoomed-in views of the final 3D density map from panel A, displayed at two contour levels. (E) Superimposition of the high-contour level map from panel D onto its fitted model.

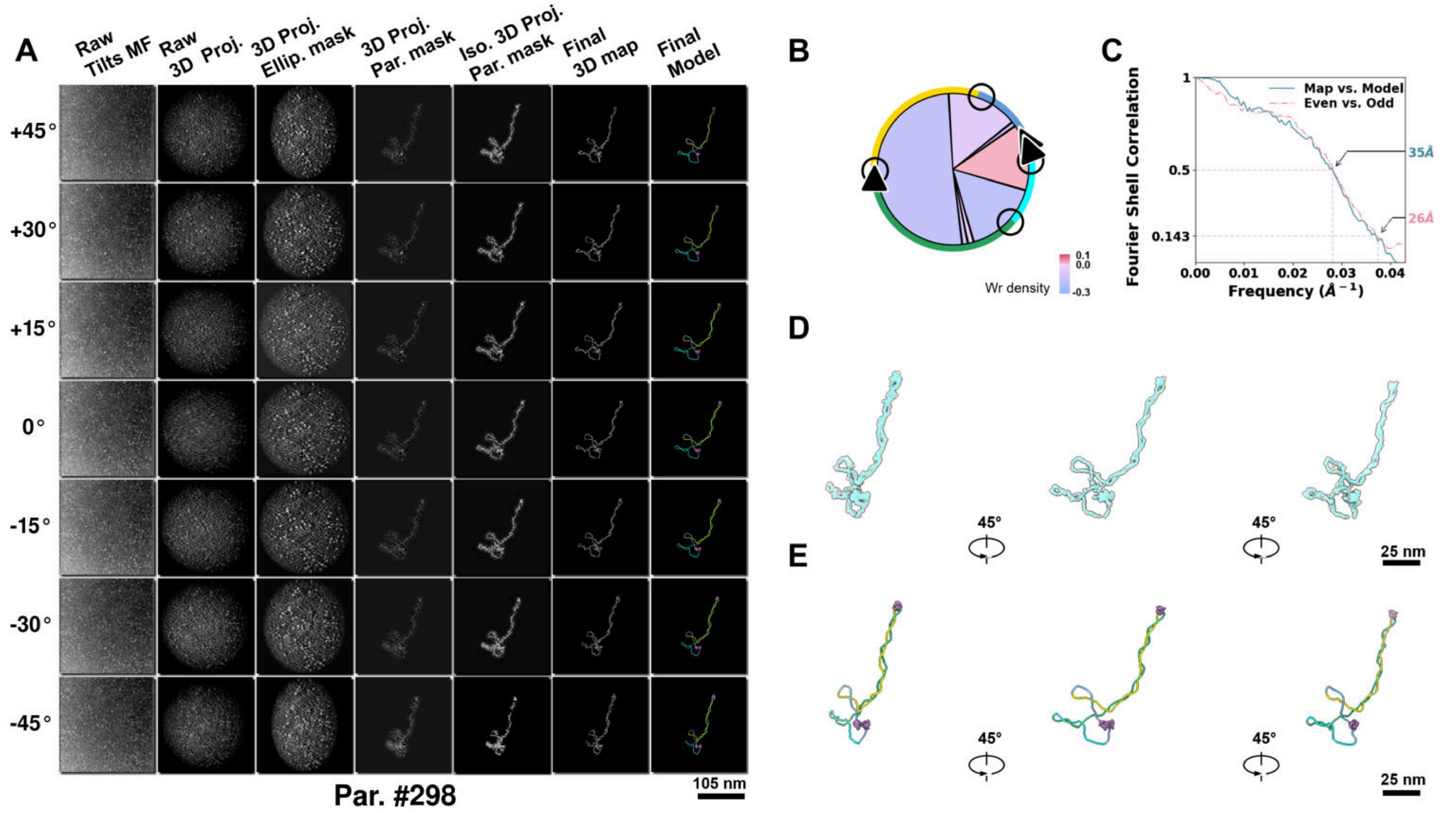

**Supplementary Particle Figure 298. Cryo-ET 3D reconstruction of an individual Opp.-TEC particle.**
(A) 3D reconstruction of the plasmid particle (index no. 298). The first column shows seven representative tilt images from +45° to -45° in step of 15°. The second, third, and fourth columns show 3D projections of the particle with spherical, ellipsoidal (thinner along the z-dimension), and particle-shaped masks, respectively. The fifth column displays the 3D projections of the enhanced and IsoNet missing-wedge-corrected particle. The sixth and seventh columns present the final 3D map and the flexibly fitted model, respectively. (B) Circular schematic representation of a plasmid particle. The outer rim is color-coded to match the corresponding 3D model. Arrowheads indicate the transcriptional direction of bound RNAPs, and circles denote apical sites. Inner circular sectors represent individual plectonemes, with colors indicating writhe density (blue to red scale, −0.3 to 0.1). (C) Resolution assessment of the final 3D map using Fourier shell correlation (FSC). Two criteria are shown: FSC between two half-maps reconstructed from even and odd frames (evaluated at 0.143) and FSC between the final 3D map and the fitted model (evaluated at 0.5). (D) Zoomed-in views of the final 3D density map from panel A, displayed at two contour levels. (E) Superimposition of the high-contour level map from panel D onto its fitted model.

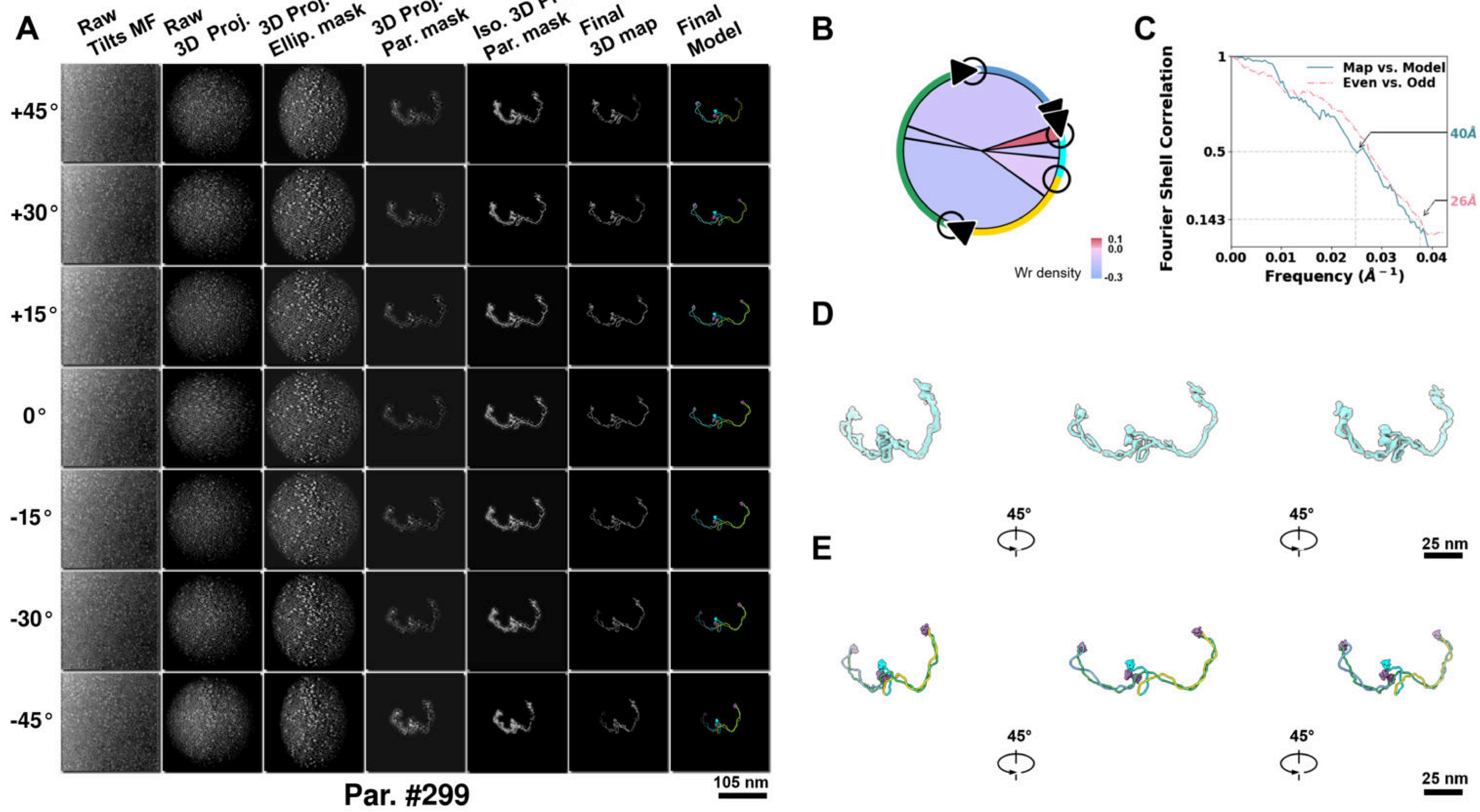

**Supplementary Particle Figure 299. Cryo-ET 3D reconstruction of an individual Opp.-TEC particle.**
(A) 3D reconstruction of the plasmid particle (index no. 299). The first column shows seven representative tilt images from +45° to -45° in step of 15°. The second, third, and fourth columns show 3D projections of the particle with spherical, ellipsoidal (thinner along the z-dimension), and particle-shaped masks, respectively. The fifth column displays the 3D projections of the enhanced and IsoNet missing-wedge-corrected particle. The sixth and seventh columns present the final 3D map and the flexibly fitted model, respectively. (B) Circular schematic representation of a plasmid particle. The outer rim is color-coded to match the corresponding 3D model. Arrowheads indicate the transcriptional direction of bound RNAPs, and circles denote apical sites. Inner circular sectors represent individual plectonemes, with colors indicating writhe density (blue to red scale, −0.3 to 0.1). (C) Resolution assessment of the final 3D map using Fourier shell correlation (FSC). Two criteria are shown: FSC between two half-maps reconstructed from even and odd frames (evaluated at 0.143) and FSC between the final 3D map and the fitted model (evaluated at 0.5). (D) Zoomed-in views of the final 3D density map from panel A, displayed at two contour levels. (E) Superimposition of the high-contour level map from panel D onto its fitted model.

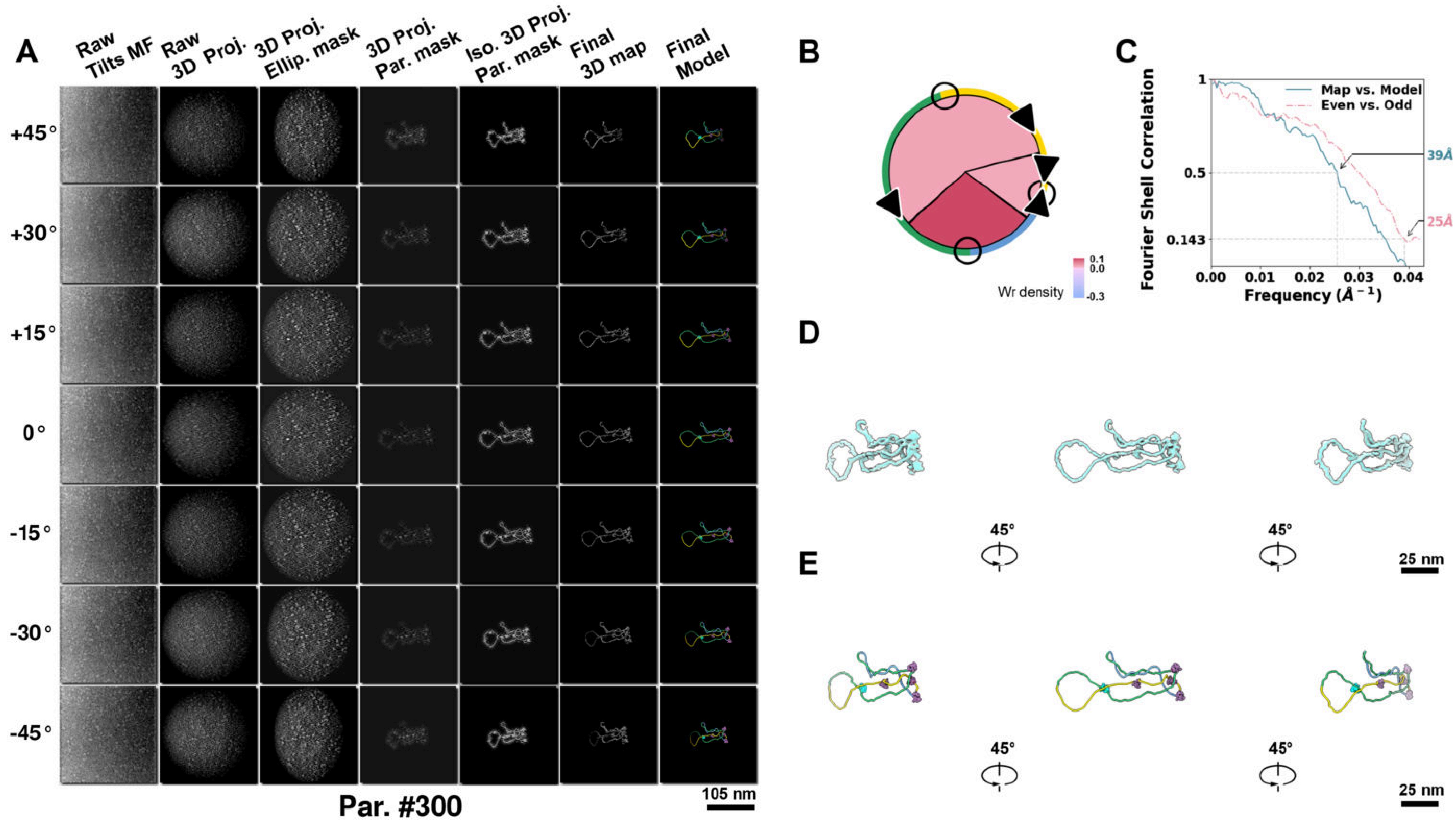

**Supplementary Particle Figure 300. Cryo-ET 3D reconstruction of an individual Opp.-TEC particle.**
(A) 3D reconstruction of the plasmid particle (index no. 300). The first column shows seven representative tilt images from +45° to -45° in step of 15°. The second, third, and fourth columns show 3D projections of the particle with spherical, ellipsoidal (thinner along the z-dimension), and particle-shaped masks, respectively. The fifth column displays the 3D projections of the enhanced and IsoNet missing-wedge-corrected particle. The sixth and seventh columns present the final 3D map and the flexibly fitted model, respectively. (B) Circular schematic representation of a plasmid particle. The outer rim is color-coded to match the corresponding 3D model. Arrowheads indicate the transcriptional direction of bound RNAPs, and circles denote apical sites. Inner circular sectors represent individual plectonemes, with colors indicating writhe density (blue to red scale, −0.3 to 0.1). (C) Resolution assessment of the final 3D map using Fourier shell correlation (FSC). Two criteria are shown: FSC between two half-maps reconstructed from even and odd frames (evaluated at 0.143) and FSC between the final 3D map and the fitted model (evaluated at 0.5). (D) Zoomed-in views of the final 3D density map from panel A, displayed at two contour levels. (E) Superimposition of the high-contour level map from panel D onto its fitted model.

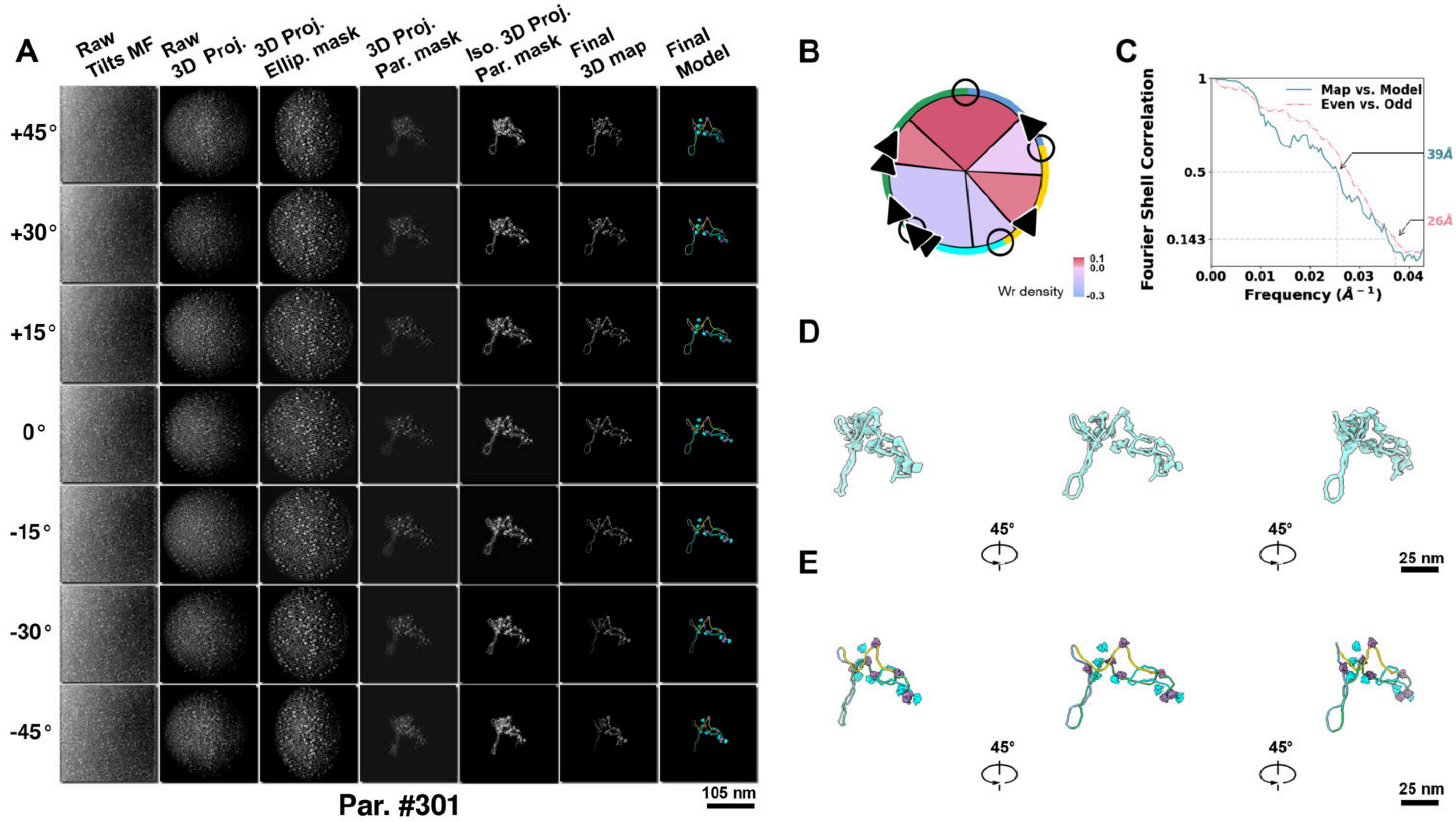

**Supplementary Particle Figure 301. Cryo-ET 3D reconstruction of an individual Opp.-TEC particle.**
(A) 3D reconstruction of the plasmid particle (index no. 301). The first column shows seven representative tilt images from +45° to -45° in step of 15°. The second, third, and fourth columns show 3D projections of the particle with spherical, ellipsoidal (thinner along the z-dimension), and particle-shaped masks, respectively. The fifth column displays the 3D projections of the enhanced and IsoNet missing-wedge-corrected particle. The sixth and seventh columns present the final 3D map and the flexibly fitted model, respectively. (B) Circular schematic representation of a plasmid particle. The outer rim is color-coded to match the corresponding 3D model. Arrowheads indicate the transcriptional direction of bound RNAPs, and circles denote apical sites. Inner circular sectors represent individual plectonemes, with colors indicating writhe density (blue to red scale, −0.3 to 0.1). (C) Resolution assessment of the final 3D map using Fourier shell correlation (FSC). Two criteria are shown: FSC between two half-maps reconstructed from even and odd frames (evaluated at 0.143) and FSC between the final 3D map and the fitted model (evaluated at 0.5). (D) Zoomed-in views of the final 3D density map from panel A, displayed at two contour levels. (E) Superimposition of the high-contour level map from panel D onto its fitted model.

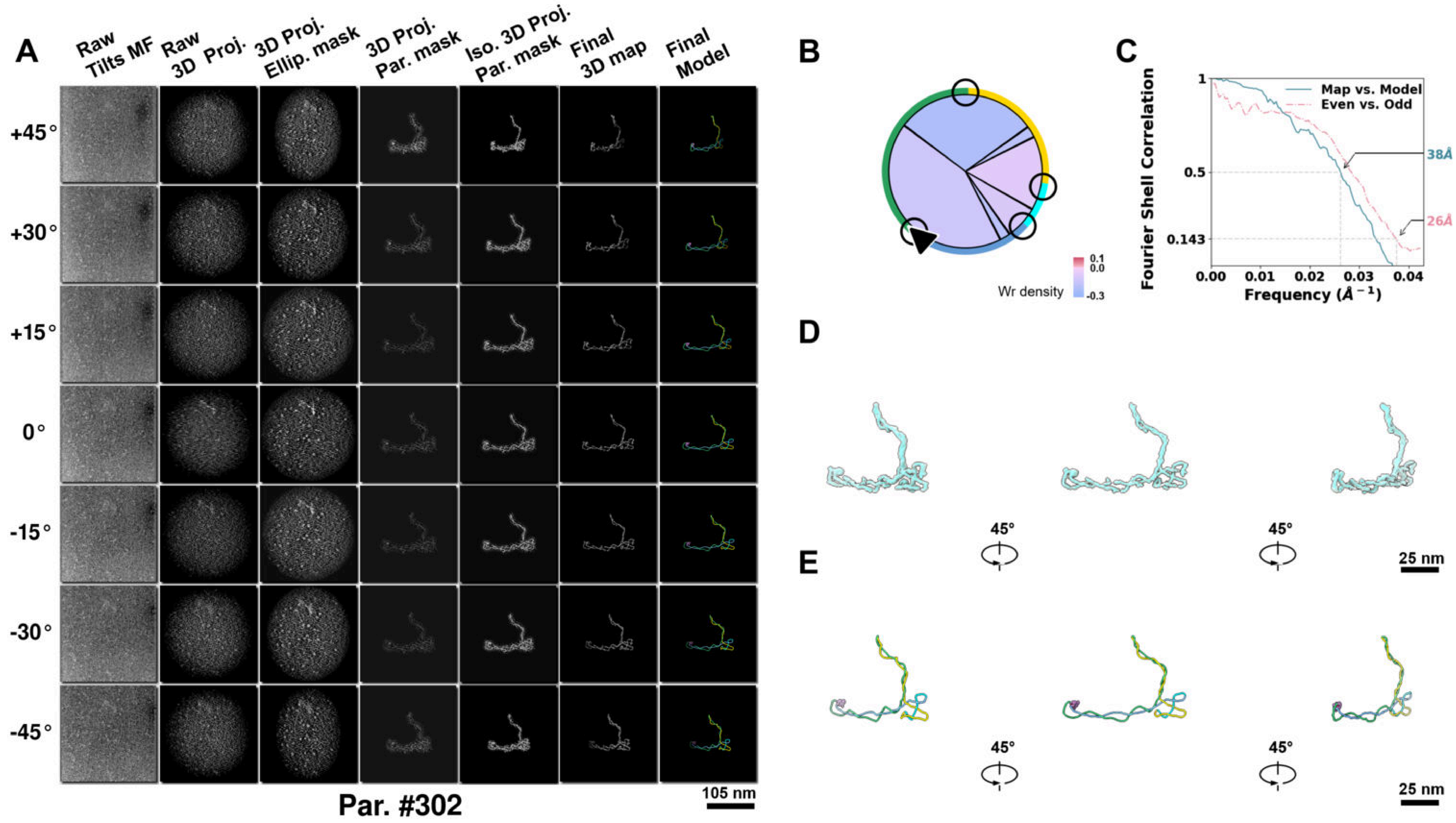

**Supplementary Particle Figure 302. Cryo-ET 3D reconstruction of an individual Tan.-TEC particle.**
(A) 3D reconstruction of the plasmid particle (index no. 302). The first column shows seven representative tilt images from +45° to -45° in step of 15°. The second, third, and fourth columns show 3D projections of the particle with spherical, ellipsoidal (thinner along the z-dimension), and particle-shaped masks, respectively. The fifth column displays the 3D projections of the enhanced and IsoNet missing-wedge-corrected particle. The sixth and seventh columns present the final 3D map and the flexibly fitted model, respectively. (B) Circular schematic representation of a plasmid particle. The outer rim is color-coded to match the corresponding 3D model. Arrowheads indicate the transcriptional direction of bound RNAPs, and circles denote apical sites. Inner circular sectors represent individual plectonemes, with colors indicating writhe density (blue to red scale, −0.3 to 0.1). (C) Resolution assessment of the final 3D map using Fourier shell correlation (FSC). Two criteria are shown: FSC between two half-maps reconstructed from even and odd frames (evaluated at 0.143) and FSC between the final 3D map and the fitted model (evaluated at 0.5). (D) Zoomed-in views of the final 3D density map from panel A, displayed at two contour levels. (E) Superimposition of the high-contour level map from panel D onto its fitted model.

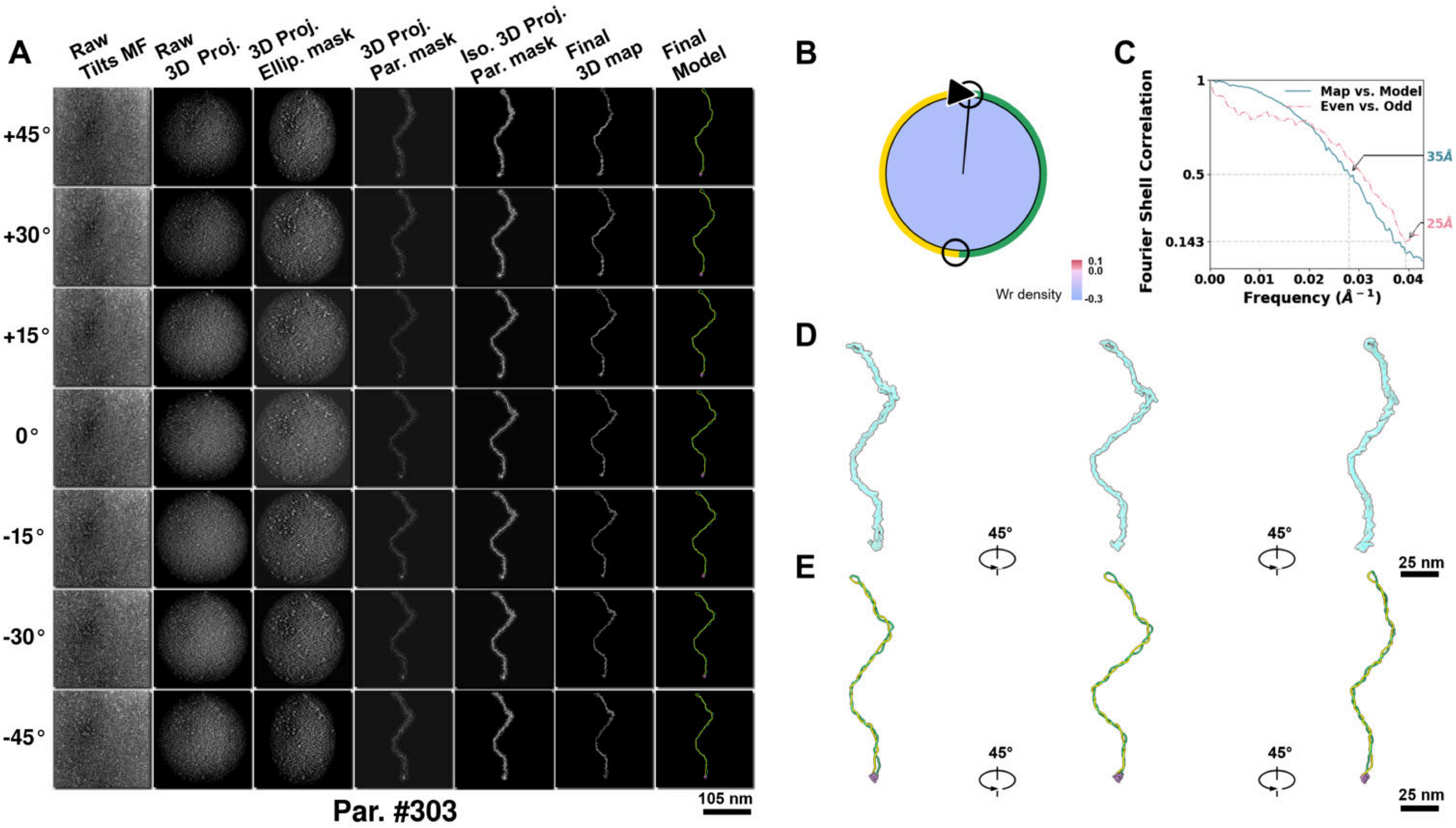

**Supplementary Particle Figure 303. Cryo-ET 3D reconstruction of an individual Tan.-TEC particle.**
(A) 3D reconstruction of the plasmid particle (index no. 303). The first column shows seven representative tilt images from +45° to -45° in step of 15°. The second, third, and fourth columns show 3D projections of the particle with spherical, ellipsoidal (thinner along the z-dimension), and particle-shaped masks, respectively. The fifth column displays the 3D projections of the enhanced and IsoNet missing-wedge-corrected particle. The sixth and seventh columns present the final 3D map and the flexibly fitted model, respectively. (B) Circular schematic representation of a plasmid particle. The outer rim is color-coded to match the corresponding 3D model. Arrowheads indicate the transcriptional direction of bound RNAPs, and circles denote apical sites. Inner circular sectors represent individual plectonemes, with colors indicating writhe density (blue to red scale, −0.3 to 0.1). (C) Resolution assessment of the final 3D map using Fourier shell correlation (FSC). Two criteria are shown: FSC between two half-maps reconstructed from even and odd frames (evaluated at 0.143) and FSC between the final 3D map and the fitted model (evaluated at 0.5). (D) Zoomed-in views of the final 3D density map from panel A, displayed at two contour levels. (E) Superimposition of the high-contour level map from panel D onto its fitted model.

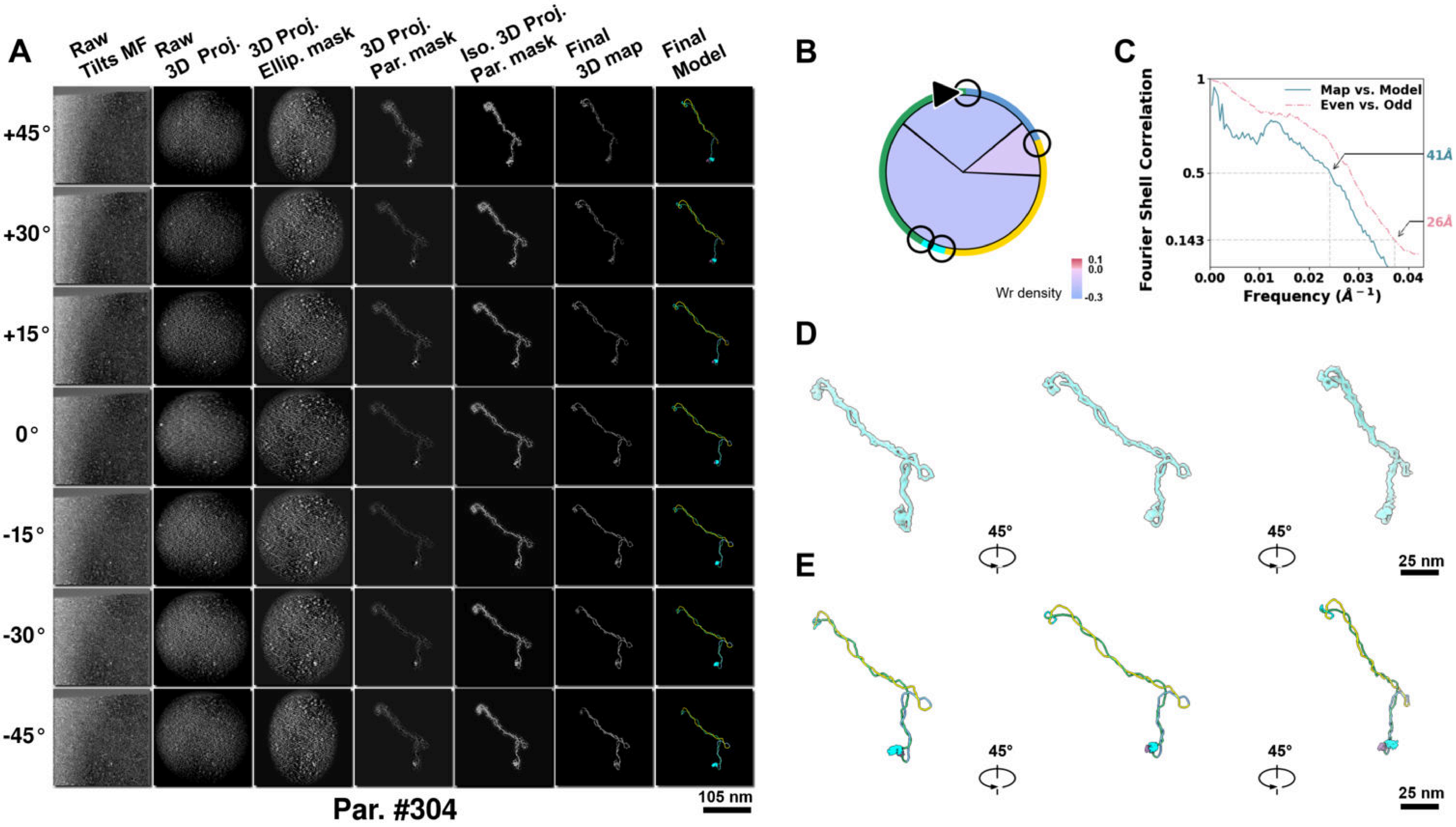

**Supplementary Particle Figure 304. Cryo-ET 3D reconstruction of an individual Tan.-TEC particle.**
(A) 3D reconstruction of the plasmid particle (index no. 304). The first column shows seven representative tilt images from +45° to -45° in step of 15°. The second, third, and fourth columns show 3D projections of the particle with spherical, ellipsoidal (thinner along the z-dimension), and particle-shaped masks, respectively. The fifth column displays the 3D projections of the enhanced and IsoNet missing-wedge-corrected particle. The sixth and seventh columns present the final 3D map and the flexibly fitted model, respectively. (B) Circular schematic representation of a plasmid particle. The outer rim is color-coded to match the corresponding 3D model. Arrowheads indicate the transcriptional direction of bound RNAPs, and circles denote apical sites. Inner circular sectors represent individual plectonemes, with colors indicating writhe density (blue to red scale, −0.3 to 0.1). (C) Resolution assessment of the final 3D map using Fourier shell correlation (FSC). Two criteria are shown: FSC between two half-maps reconstructed from even and odd frames (evaluated at 0.143) and FSC between the final 3D map and the fitted model (evaluated at 0.5). (D) Zoomed-in views of the final 3D density map from panel A, displayed at two contour levels. (E) Superimposition of the high-contour level map from panel D onto its fitted model.

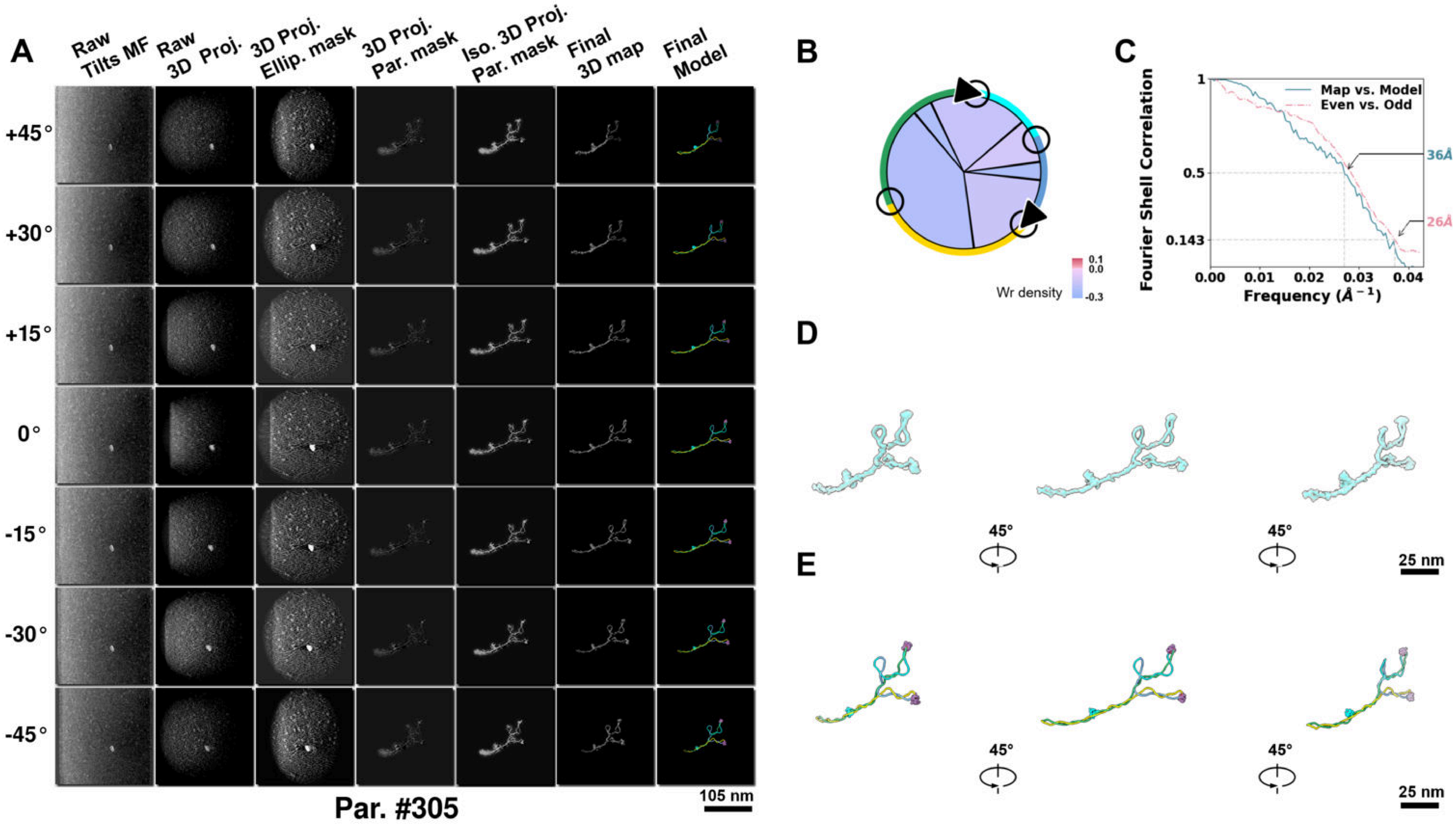

**Supplementary Particle Figure 305. Cryo-ET 3D reconstruction of an individual Tan.-TEC particle.**
(A) 3D reconstruction of the plasmid particle (index no. 305). The first column shows seven representative tilt images from +45° to -45° in step of 15°. The second, third, and fourth columns show 3D projections of the particle with spherical, ellipsoidal (thinner along the z-dimension), and particle-shaped masks, respectively. The fifth column displays the 3D projections of the enhanced and IsoNet missing-wedge-corrected particle. The sixth and seventh columns present the final 3D map and the flexibly fitted model, respectively. (B) Circular schematic representation of a plasmid particle. The outer rim is color-coded to match the corresponding 3D model. Arrowheads indicate the transcriptional direction of bound RNAPs, and circles denote apical sites. Inner circular sectors represent individual plectonemes, with colors indicating writhe density (blue to red scale, −0.3 to 0.1). (C) Resolution assessment of the final 3D map using Fourier shell correlation (FSC). Two criteria are shown: FSC between two half-maps reconstructed from even and odd frames (evaluated at 0.143) and FSC between the final 3D map and the fitted model (evaluated at 0.5). (D) Zoomed-in views of the final 3D density map from panel A, displayed at two contour levels. (E) Superimposition of the high-contour level map from panel D onto its fitted model.

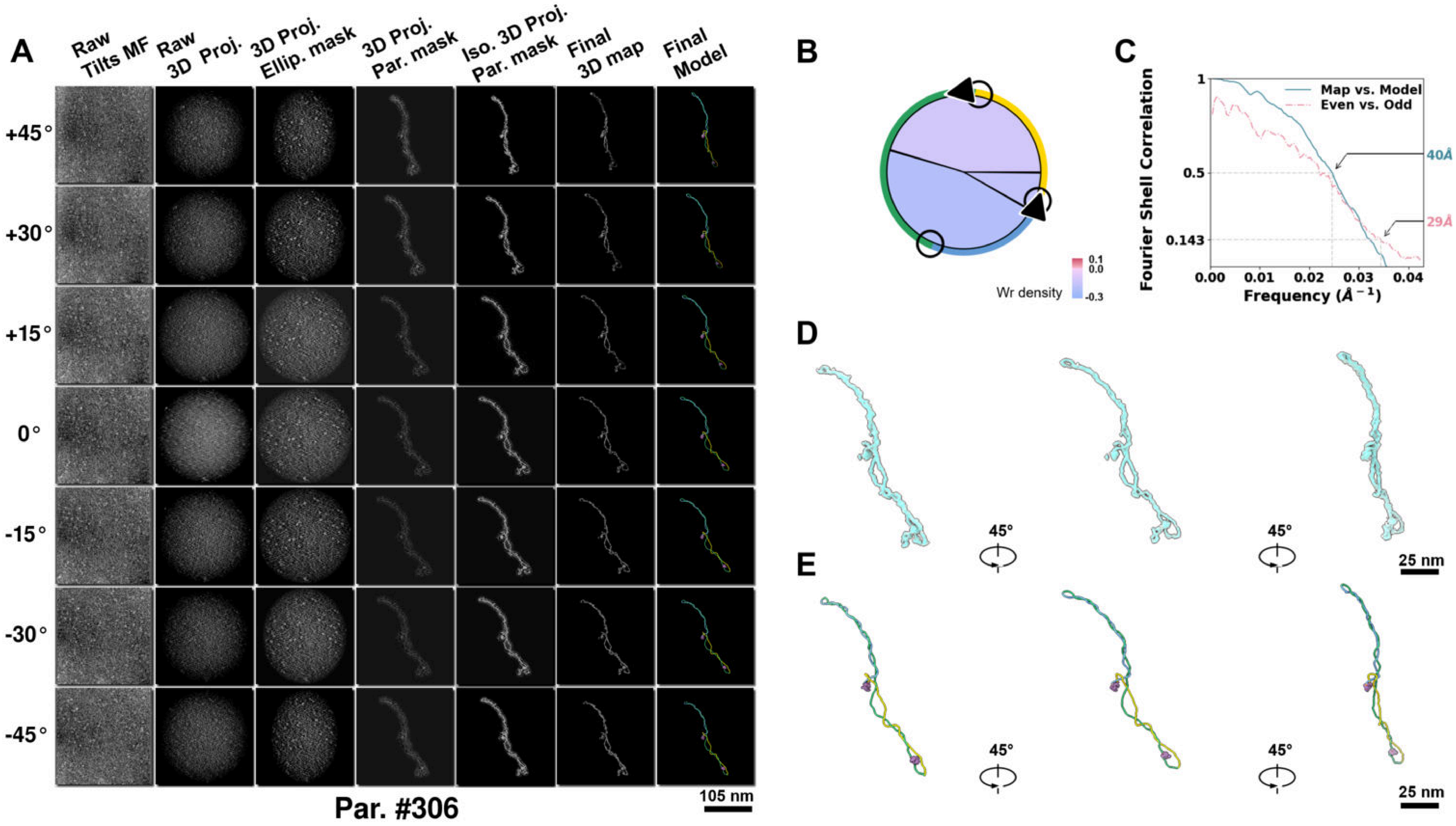

**Supplementary Particle Figure 306. Cryo-ET 3D reconstruction of an individual Tan.-TEC particle.**
(A) 3D reconstruction of the plasmid particle (index no. 306). The first column shows seven representative tilt images from +45° to -45° in step of 15°. The second, third, and fourth columns show 3D projections of the particle with spherical, ellipsoidal (thinner along the z-dimension), and particle-shaped masks, respectively. The fifth column displays the 3D projections of the enhanced and IsoNet missing-wedge-corrected particle. The sixth and seventh columns present the final 3D map and the flexibly fitted model, respectively. (B) Circular schematic representation of a plasmid particle. The outer rim is color-coded to match the corresponding 3D model. Arrowheads indicate the transcriptional direction of bound RNAPs, and circles denote apical sites. Inner circular sectors represent individual plectonemes, with colors indicating writhe density (blue to red scale, −0.3 to 0.1). (C) Resolution assessment of the final 3D map using Fourier shell correlation (FSC). Two criteria are shown: FSC between two half-maps reconstructed from even and odd frames (evaluated at 0.143) and FSC between the final 3D map and the fitted model (evaluated at 0.5). (D) Zoomed-in views of the final 3D density map from panel A, displayed at two contour levels. (E) Superimposition of the high-contour level map from panel D onto its fitted model.

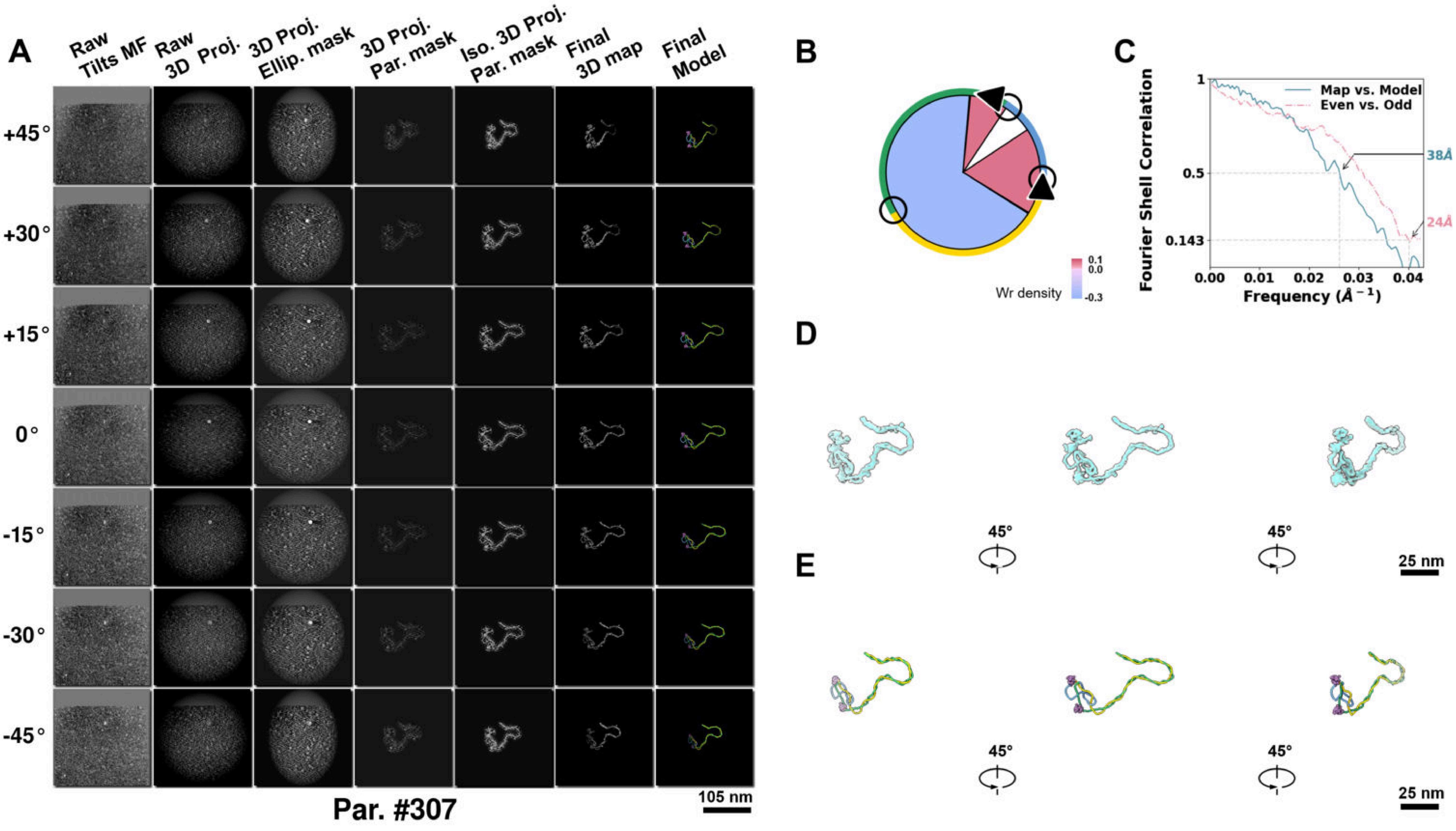

**Supplementary Particle Figure 307. Cryo-ET 3D reconstruction of an individual Tan.-TEC particle.**
(A) 3D reconstruction of the plasmid particle (index no. 307). The first column shows seven representative tilt images from +45° to -45° in step of 15°. The second, third, and fourth columns show 3D projections of the particle with spherical, ellipsoidal (thinner along the z-dimension), and particle-shaped masks, respectively. The fifth column displays the 3D projections of the enhanced and IsoNet missing-wedge-corrected particle. The sixth and seventh columns present the final 3D map and the flexibly fitted model, respectively. (B) Circular schematic representation of a plasmid particle. The outer rim is color-coded to match the corresponding 3D model. Arrowheads indicate the transcriptional direction of bound RNAPs, and circles denote apical sites. Inner circular sectors represent individual plectonemes, with colors indicating writhe density (blue to red scale, −0.3 to 0.1). (C) Resolution assessment of the final 3D map using Fourier shell correlation (FSC). Two criteria are shown: FSC between two half-maps reconstructed from even and odd frames (evaluated at 0.143) and FSC between the final 3D map and the fitted model (evaluated at 0.5). (D) Zoomed-in views of the final 3D density map from panel A, displayed at two contour levels. (E) Superimposition of the high-contour level map from panel D onto its fitted model.

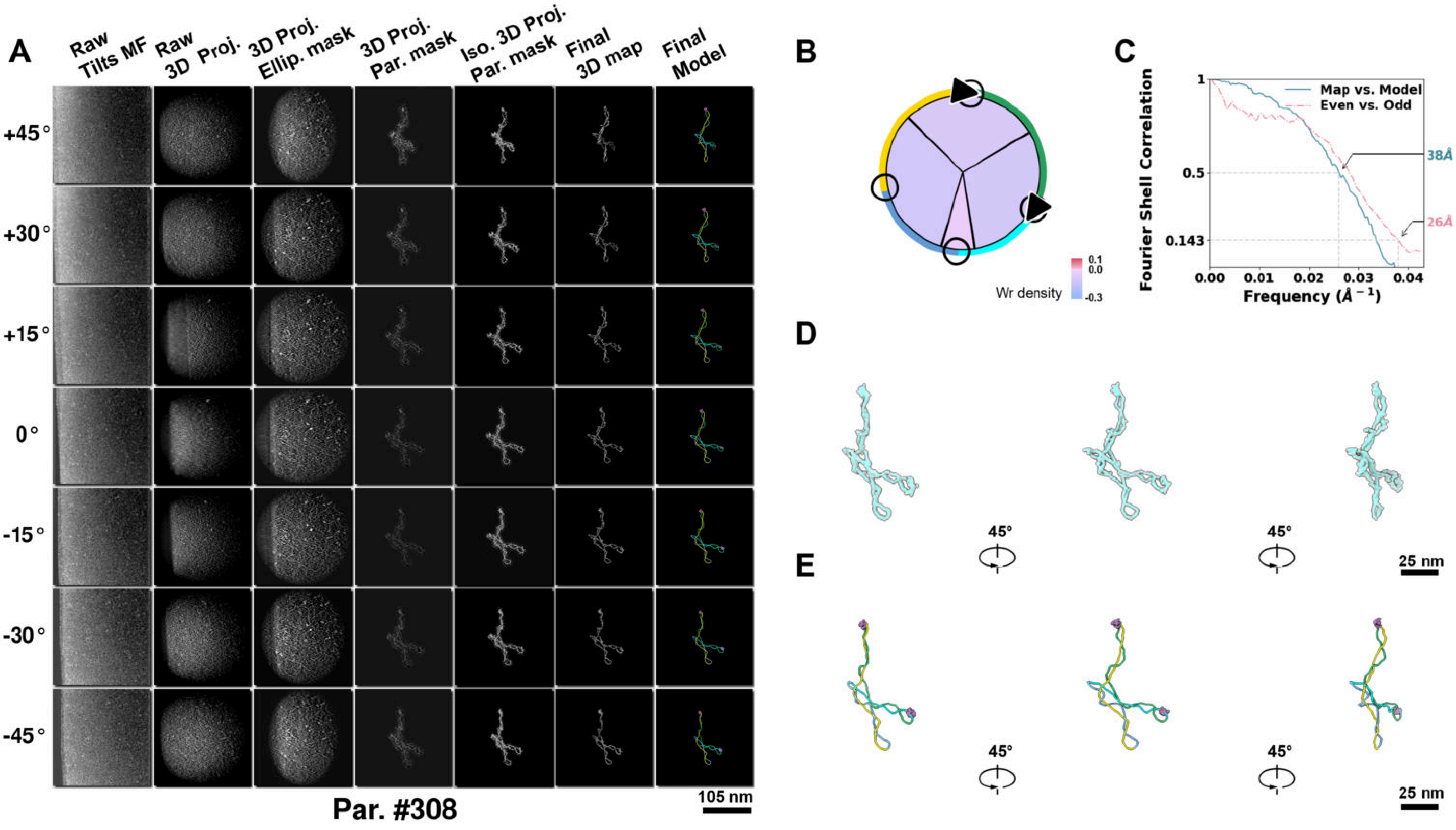

**Supplementary Particle Figure 308. Cryo-ET 3D reconstruction of an individual Tan.-TEC particle.**
(A) 3D reconstruction of the plasmid particle (index no. 308). The first column shows seven representative tilt images from +45° to -45° in step of 15°. The second, third, and fourth columns show 3D projections of the particle with spherical, ellipsoidal (thinner along the z-dimension), and particle-shaped masks, respectively. The fifth column displays the 3D projections of the enhanced and IsoNet missing-wedge-corrected particle. The sixth and seventh columns present the final 3D map and the flexibly fitted model, respectively. (B) Circular schematic representation of a plasmid particle. The outer rim is color-coded to match the corresponding 3D model. Arrowheads indicate the transcriptional direction of bound RNAPs, and circles denote apical sites. Inner circular sectors represent individual plectonemes, with colors indicating writhe density (blue to red scale, −0.3 to 0.1). (C) Resolution assessment of the final 3D map using Fourier shell correlation (FSC). Two criteria are shown: FSC between two half-maps reconstructed from even and odd frames (evaluated at 0.143) and FSC between the final 3D map and the fitted model (evaluated at 0.5). (D) Zoomed-in views of the final 3D density map from panel A, displayed at two contour levels. (E) Superimposition of the high-contour level map from panel D onto its fitted model.

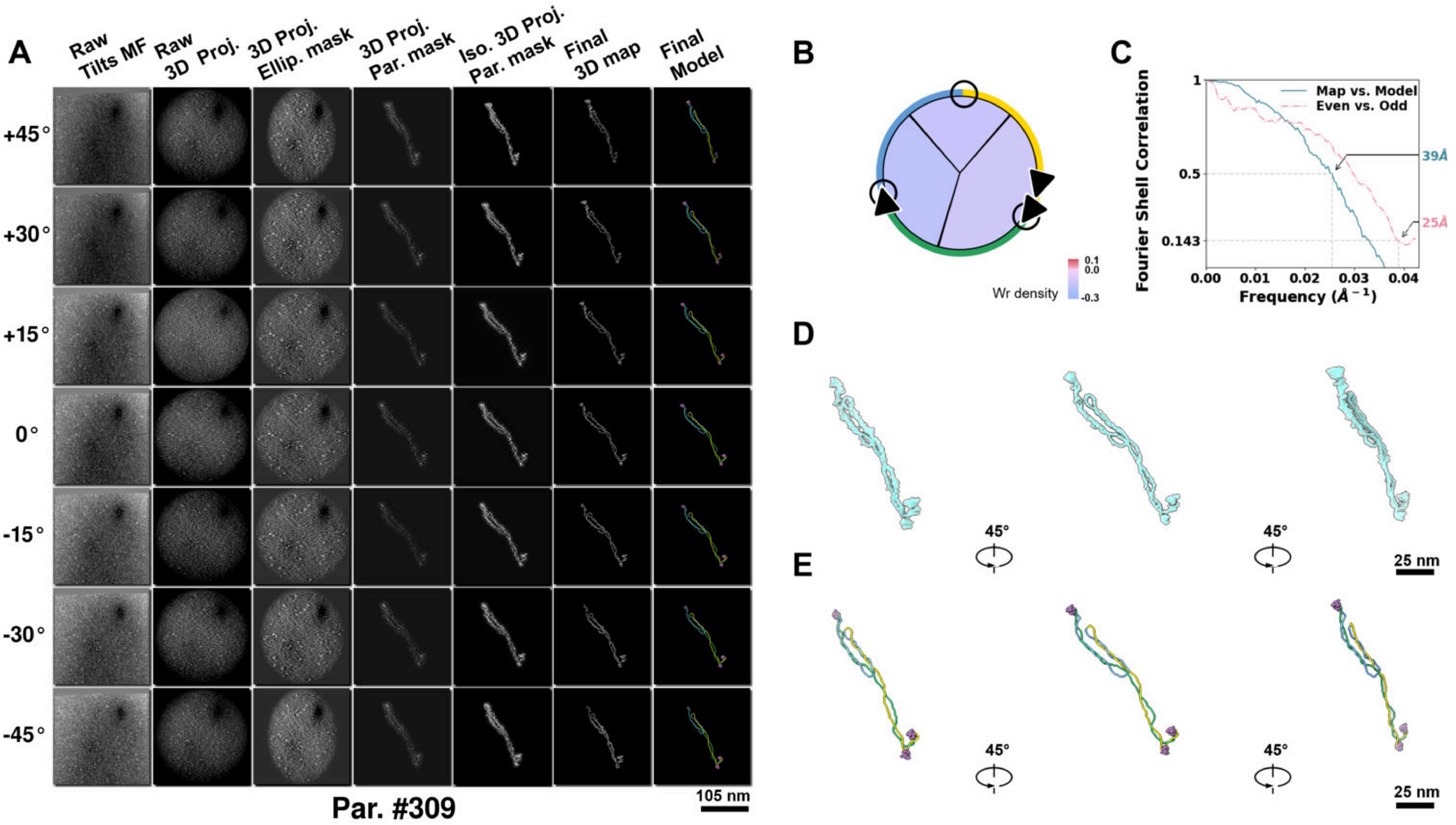

**Supplementary Particle Figure 309. Cryo-ET 3D reconstruction of an individual Tan.-TEC particle.**
(A) 3D reconstruction of the plasmid particle (index no. 309). The first column shows seven representative tilt images from +45° to -45° in step of 15°. The second, third, and fourth columns show 3D projections of the particle with spherical, ellipsoidal (thinner along the z-dimension), and particle-shaped masks, respectively. The fifth column displays the 3D projections of the enhanced and IsoNet missing-wedge-corrected particle. The sixth and seventh columns present the final 3D map and the flexibly fitted model, respectively. (B) Circular schematic representation of a plasmid particle. The outer rim is color-coded to match the corresponding 3D model. Arrowheads indicate the transcriptional direction of bound RNAPs, and circles denote apical sites. Inner circular sectors represent individual plectonemes, with colors indicating writhe density (blue to red scale, −0.3 to 0.1). (C) Resolution assessment of the final 3D map using Fourier shell correlation (FSC). Two criteria are shown: FSC between two half-maps reconstructed from even and odd frames (evaluated at 0.143) and FSC between the final 3D map and the fitted model (evaluated at 0.5). (D) Zoomed-in views of the final 3D density map from panel A, displayed at two contour levels. (E) Superimposition of the high-contour level map from panel D onto its fitted model.

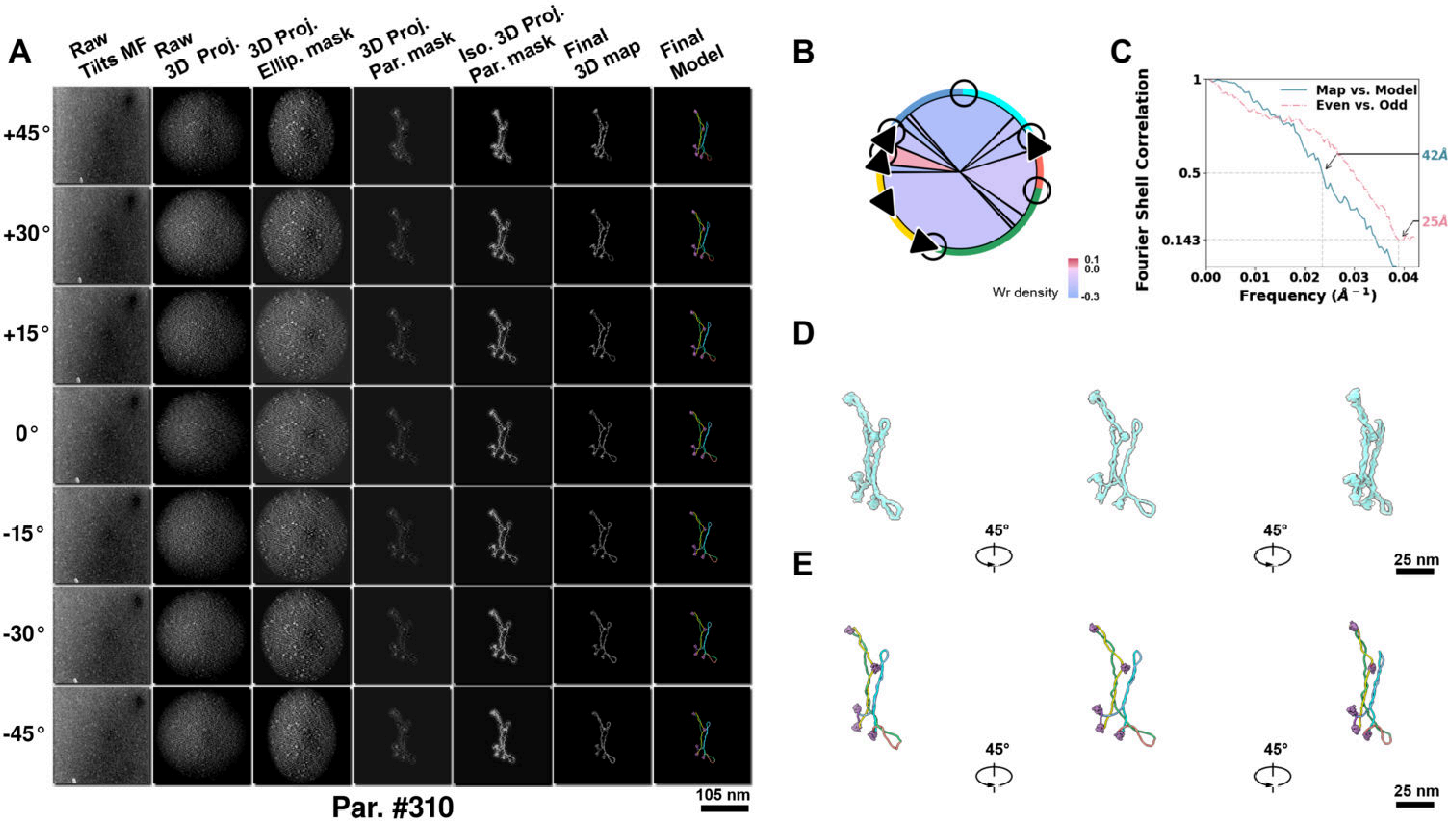

**Supplementary Particle Figure 310. Cryo-ET 3D reconstruction of an individual Tan.-TEC particle.**
(A) 3D reconstruction of the plasmid particle (index no. 310). The first column shows seven representative tilt images from +45° to -45° in step of 15°. The second, third, and fourth columns show 3D projections of the particle with spherical, ellipsoidal (thinner along the z-dimension), and particle-shaped masks, respectively. The fifth column displays the 3D projections of the enhanced and IsoNet missing-wedge-corrected particle. The sixth and seventh columns present the final 3D map and the flexibly fitted model, respectively. (B) Circular schematic representation of a plasmid particle. The outer rim is color-coded to match the corresponding 3D model. Arrowheads indicate the transcriptional direction of bound RNAPs, and circles denote apical sites. Inner circular sectors represent individual plectonemes, with colors indicating writhe density (blue to red scale, −0.3 to 0.1). (C) Resolution assessment of the final 3D map using Fourier shell correlation (FSC). Two criteria are shown: FSC between two half-maps reconstructed from even and odd frames (evaluated at 0.143) and FSC between the final 3D map and the fitted model (evaluated at 0.5). (D) Zoomed-in views of the final 3D density map from panel A, displayed at two contour levels. (E) Superimposition of the high-contour level map from panel D onto its fitted model.

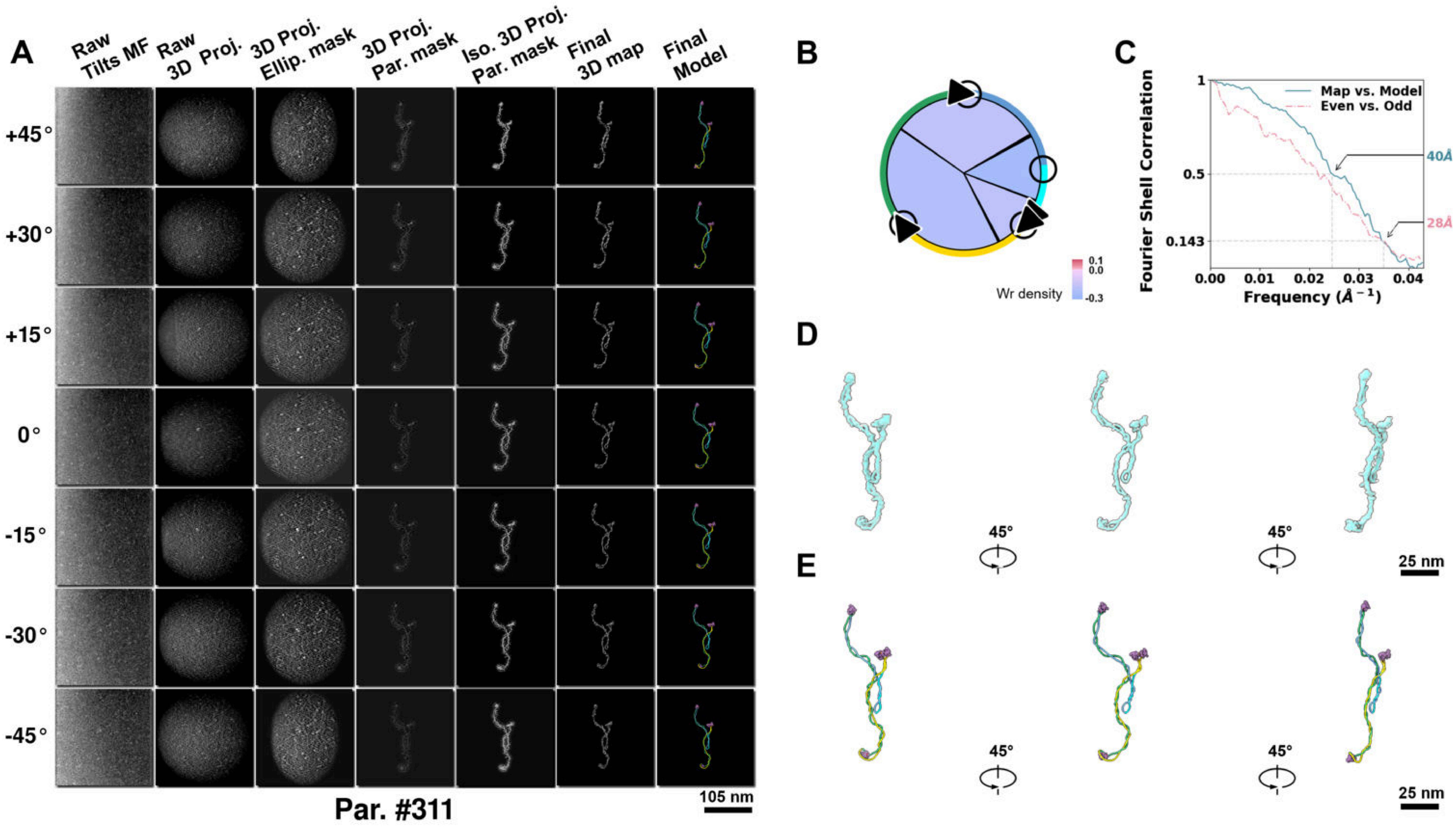

**Supplementary Particle Figure 311. Cryo-ET 3D reconstruction of an individual Tan.-TEC particle.**
(A) 3D reconstruction of the plasmid particle (index no. 311). The first column shows seven representative tilt images from +45° to -45° in step of 15°. The second, third, and fourth columns show 3D projections of the particle with spherical, ellipsoidal (thinner along the z-dimension), and particle-shaped masks, respectively. The fifth column displays the 3D projections of the enhanced and IsoNet missing-wedge-corrected particle. The sixth and seventh columns present the final 3D map and the flexibly fitted model, respectively. (B) Circular schematic representation of a plasmid particle. The outer rim is color-coded to match the corresponding 3D model. Arrowheads indicate the transcriptional direction of bound RNAPs, and circles denote apical sites. Inner circular sectors represent individual plectonemes, with colors indicating writhe density (blue to red scale, −0.3 to 0.1). (C) Resolution assessment of the final 3D map using Fourier shell correlation (FSC). Two criteria are shown: FSC between two half-maps reconstructed from even and odd frames (evaluated at 0.143) and FSC between the final 3D map and the fitted model (evaluated at 0.5). (D) Zoomed-in views of the final 3D density map from panel A, displayed at two contour levels. (E) Superimposition of the high-contour level map from panel D onto its fitted model.

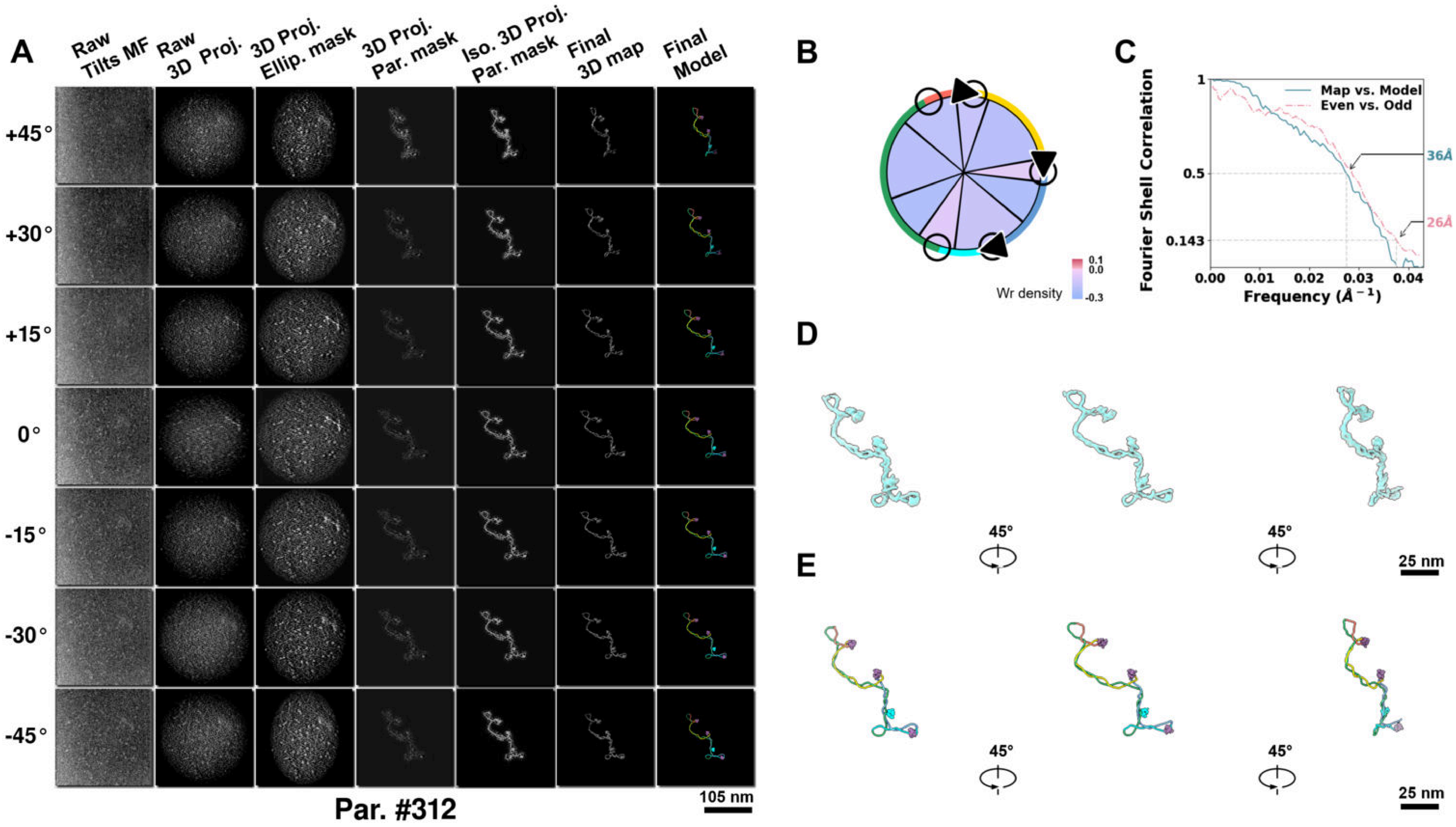

**Supplementary Particle Figure 312. Cryo-ET 3D reconstruction of an individual Tan.-TEC particle.**
(A) 3D reconstruction of the plasmid particle (index no. 312). The first column shows seven representative tilt images from +45° to -45° in step of 15°. The second, third, and fourth columns show 3D projections of the particle with spherical, ellipsoidal (thinner along the z-dimension), and particle-shaped masks, respectively. The fifth column displays the 3D projections of the enhanced and IsoNet missing-wedge-corrected particle. The sixth and seventh columns present the final 3D map and the flexibly fitted model, respectively. (B) Circular schematic representation of a plasmid particle. The outer rim is color-coded to match the corresponding 3D model. Arrowheads indicate the transcriptional direction of bound RNAPs, and circles denote apical sites. Inner circular sectors represent individual plectonemes, with colors indicating writhe density (blue to red scale, −0.3 to 0.1). (C) Resolution assessment of the final 3D map using Fourier shell correlation (FSC). Two criteria are shown: FSC between two half-maps reconstructed from even and odd frames (evaluated at 0.143) and FSC between the final 3D map and the fitted model (evaluated at 0.5). (D) Zoomed-in views of the final 3D density map from panel A, displayed at two contour levels. (E) Superimposition of the high-contour level map from panel D onto its fitted model.

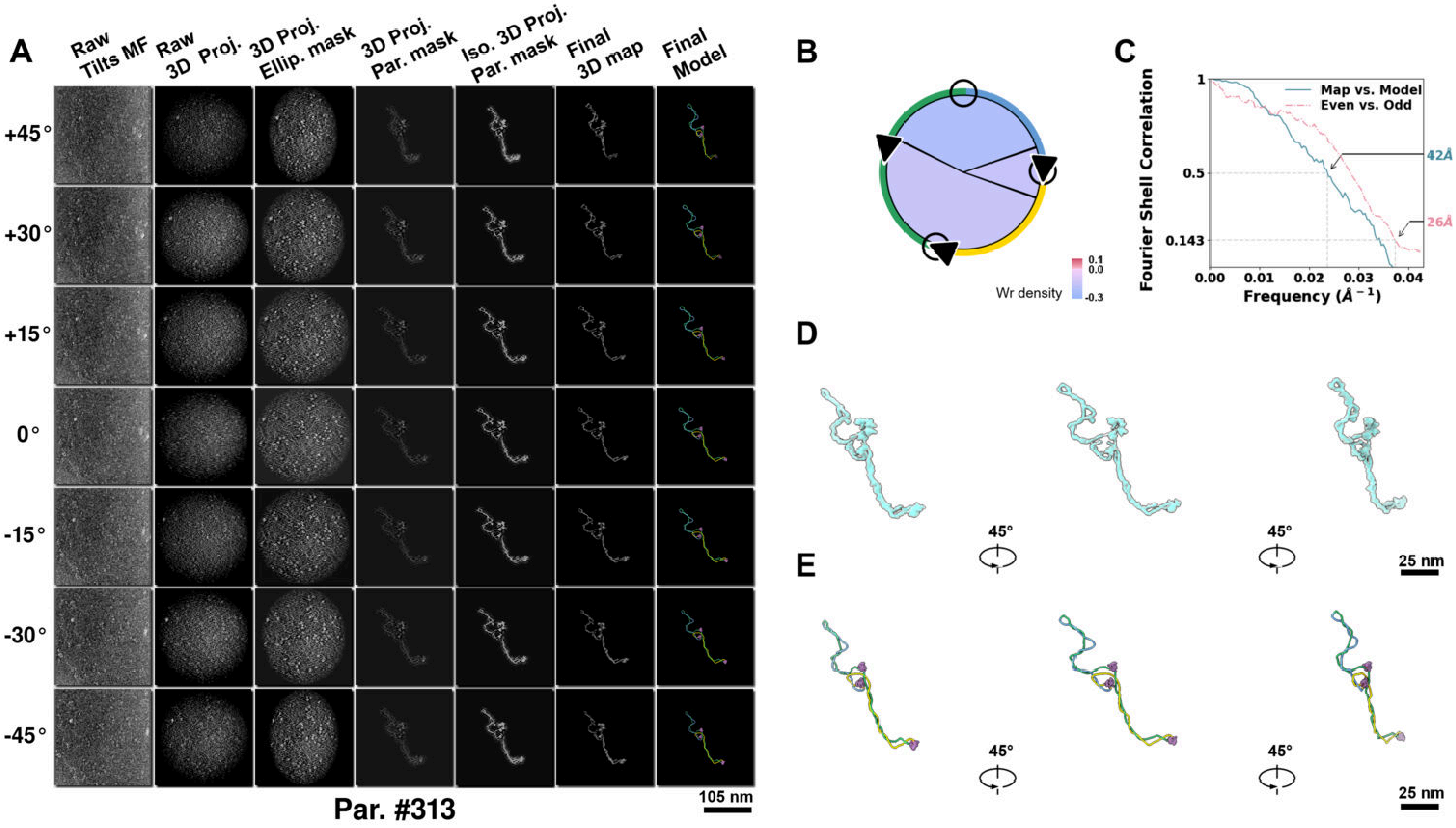

**Supplementary Particle Figure 313. Cryo-ET 3D reconstruction of an individual Tan.-TEC particle.**
(A) 3D reconstruction of the plasmid particle (index no. 313). The first column shows seven representative tilt images from +45° to -45° in step of 15°. The second, third, and fourth columns show 3D projections of the particle with spherical, ellipsoidal (thinner along the z-dimension), and particle-shaped masks, respectively. The fifth column displays the 3D projections of the enhanced and IsoNet missing-wedge-corrected particle. The sixth and seventh columns present the final 3D map and the flexibly fitted model, respectively. (B) Circular schematic representation of a plasmid particle. The outer rim is color-coded to match the corresponding 3D model. Arrowheads indicate the transcriptional direction of bound RNAPs, and circles denote apical sites. Inner circular sectors represent individual plectonemes, with colors indicating writhe density (blue to red scale, −0.3 to 0.1). (C) Resolution assessment of the final 3D map using Fourier shell correlation (FSC). Two criteria are shown: FSC between two half-maps reconstructed from even and odd frames (evaluated at 0.143) and FSC between the final 3D map and the fitted model (evaluated at 0.5). (D) Zoomed-in views of the final 3D density map from panel A, displayed at two contour levels. (E) Superimposition of the high-contour level map from panel D onto its fitted model.

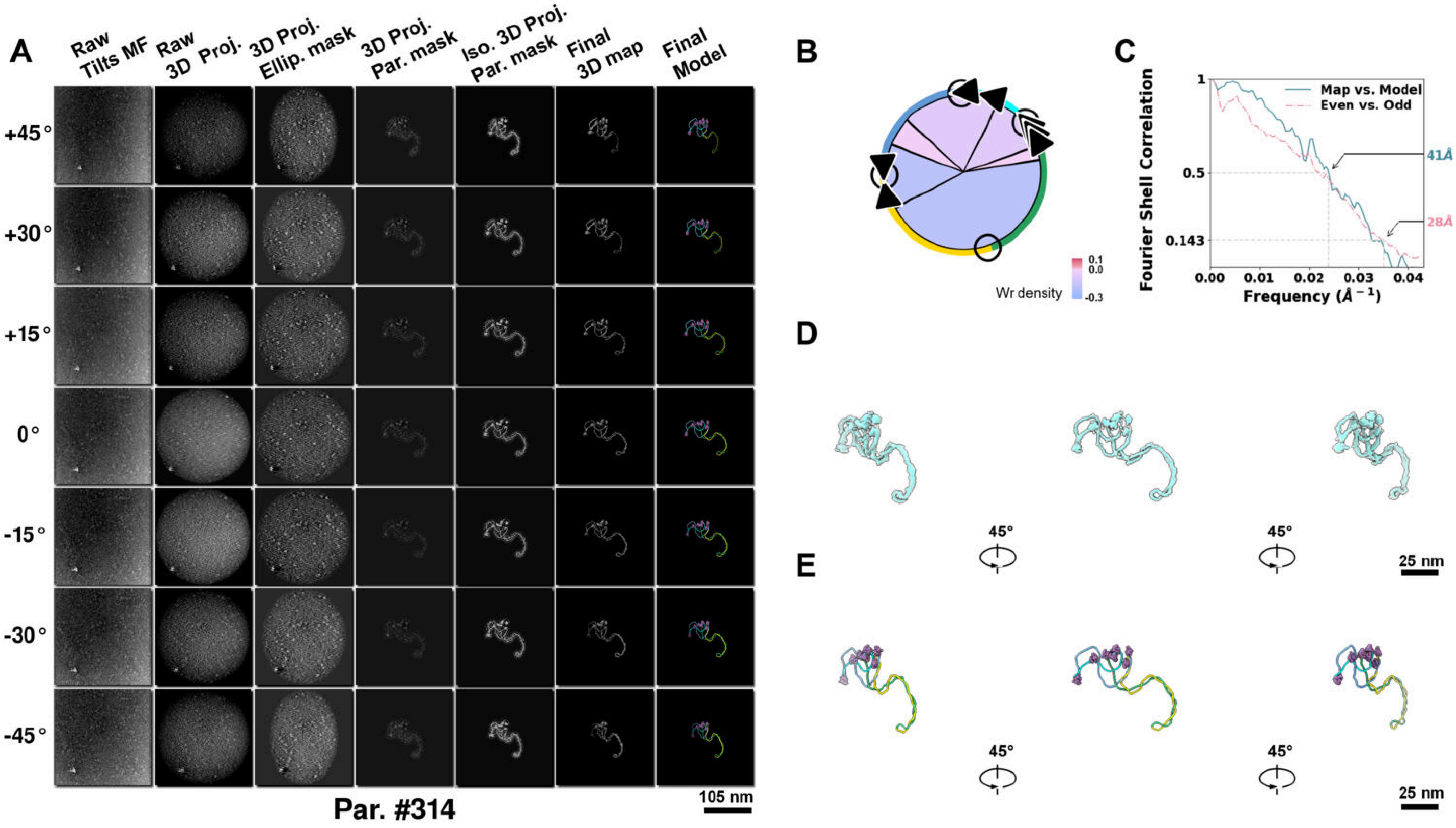

**Supplementary Particle Figure 314. Cryo-ET 3D reconstruction of an individual Tan.-TEC particle.**
(A) 3D reconstruction of the plasmid particle (index no. 314). The first column shows seven representative tilt images from +45° to -45° in step of 15°. The second, third, and fourth columns show 3D projections of the particle with spherical, ellipsoidal (thinner along the z-dimension), and particle-shaped masks, respectively. The fifth column displays the 3D projections of the enhanced and IsoNet missing-wedge-corrected particle. The sixth and seventh columns present the final 3D map and the flexibly fitted model, respectively. (B) Circular schematic representation of a plasmid particle. The outer rim is color-coded to match the corresponding 3D model. Arrowheads indicate the transcriptional direction of bound RNAPs, and circles denote apical sites. Inner circular sectors represent individual plectonemes, with colors indicating writhe density (blue to red scale, −0.3 to 0.1). (C) Resolution assessment of the final 3D map using Fourier shell correlation (FSC). Two criteria are shown: FSC between two half-maps reconstructed from even and odd frames (evaluated at 0.143) and FSC between the final 3D map and the fitted model (evaluated at 0.5). (D) Zoomed-in views of the final 3D density map from panel A, displayed at two contour levels. (E) Superimposition of the high-contour level map from panel D onto its fitted model.

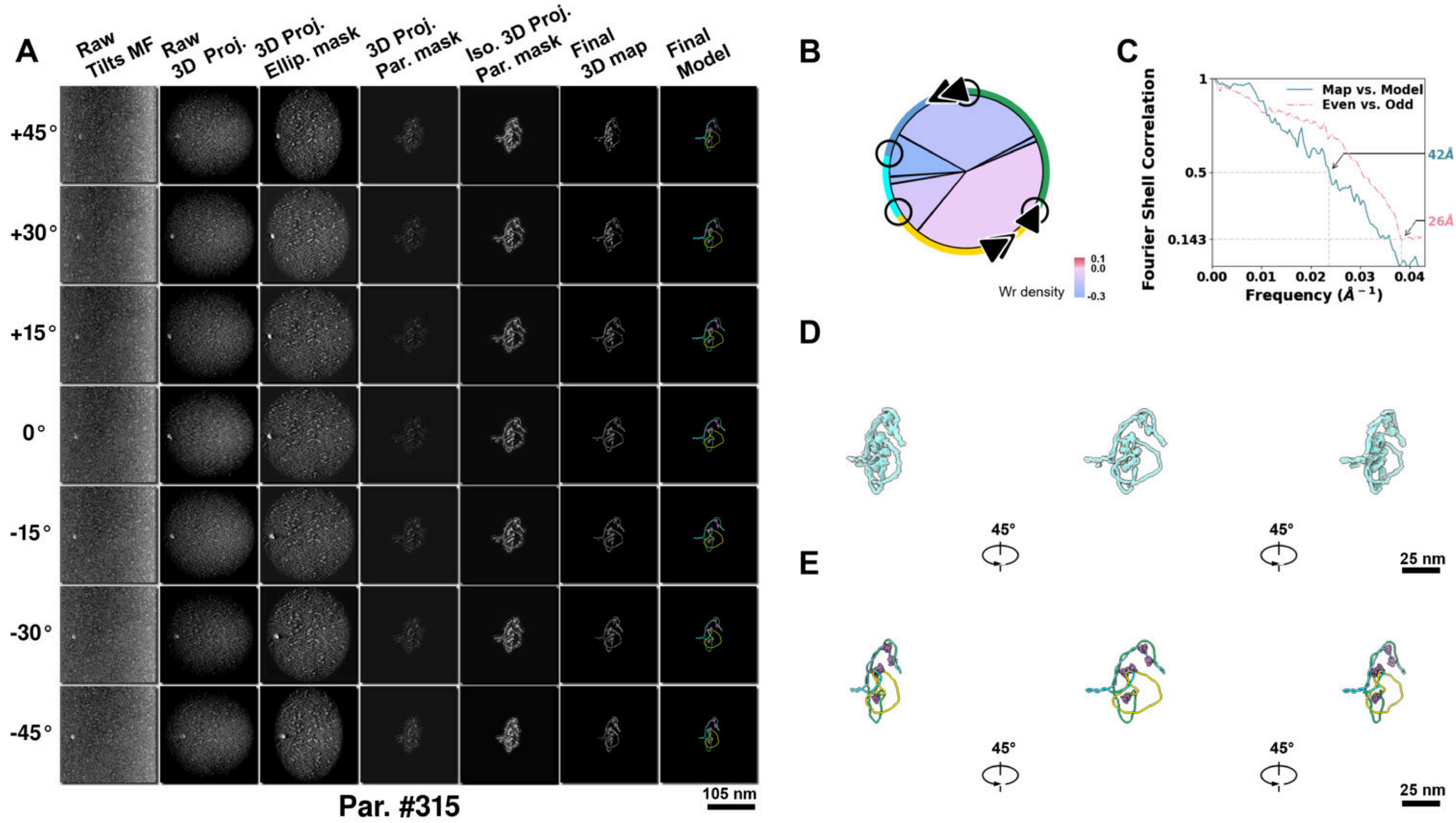

**Supplementary Particle Figure 315. Cryo-ET 3D reconstruction of an individual Tan.-TEC particle.**
(A) 3D reconstruction of the plasmid particle (index no. 315). The first column shows seven representative tilt images from +45° to -45° in step of 15°. The second, third, and fourth columns show 3D projections of the particle with spherical, ellipsoidal (thinner along the z-dimension), and particle-shaped masks, respectively. The fifth column displays the 3D projections of the enhanced and IsoNet missing-wedge-corrected particle. The sixth and seventh columns present the final 3D map and the flexibly fitted model, respectively. (B) Circular schematic representation of a plasmid particle. The outer rim is color-coded to match the corresponding 3D model. Arrowheads indicate the transcriptional direction of bound RNAPs, and circles denote apical sites. Inner circular sectors represent individual plectonemes, with colors indicating writhe density (blue to red scale, −0.3 to 0.1). (C) Resolution assessment of the final 3D map using Fourier shell correlation (FSC). Two criteria are shown: FSC between two half-maps reconstructed from even and odd frames (evaluated at 0.143) and FSC between the final 3D map and the fitted model (evaluated at 0.5). (D) Zoomed-in views of the final 3D density map from panel A, displayed at two contour levels. (E) Superimposition of the high-contour level map from panel D onto its fitted model.

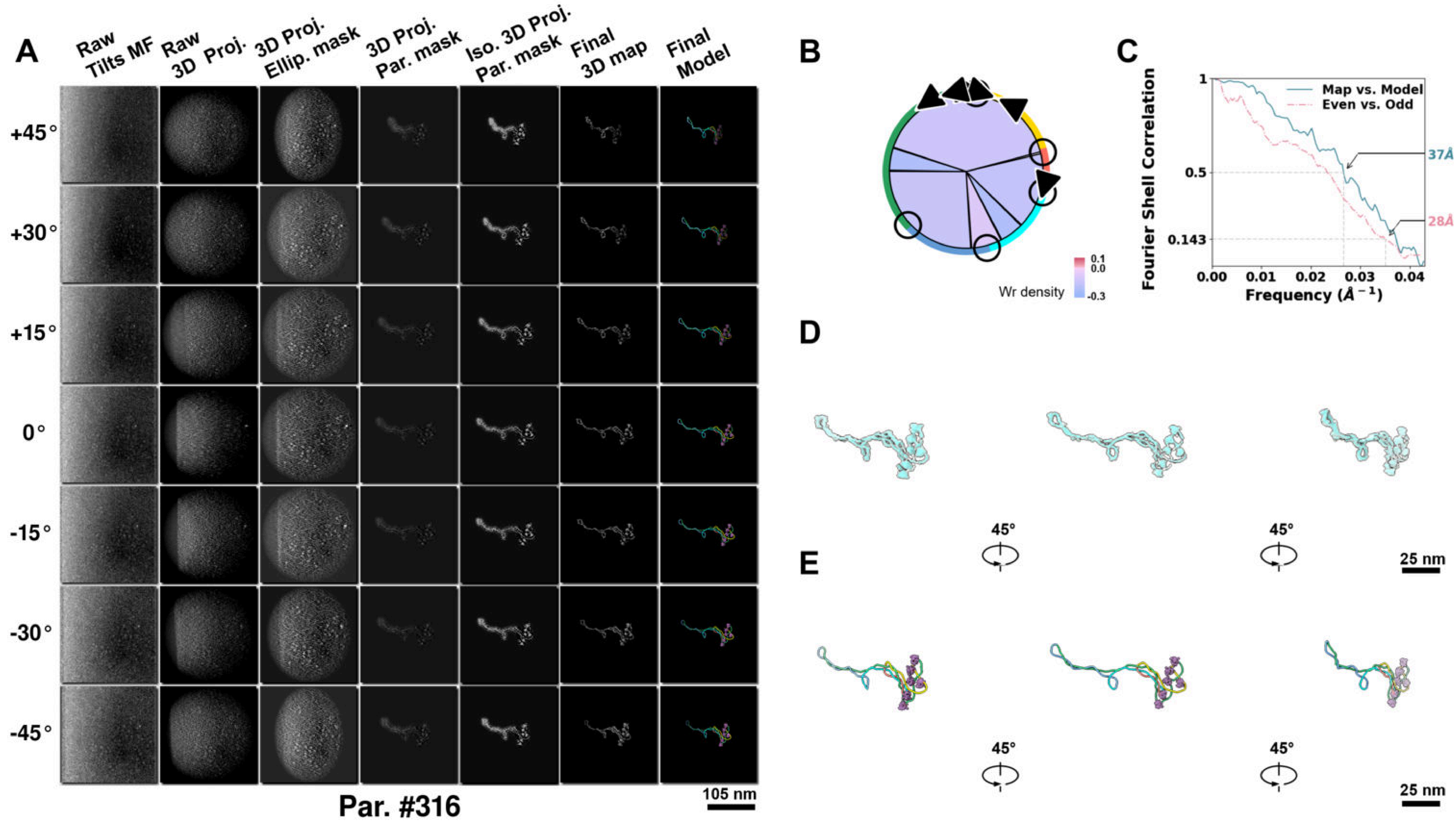

**Supplementary Particle Figure 316. Cryo-ET 3D reconstruction of an individual Tan.-TEC particle.**
(A) 3D reconstruction of the plasmid particle (index no. 316). The first column shows seven representative tilt images from +45° to -45° in step of 15°. The second, third, and fourth columns show 3D projections of the particle with spherical, ellipsoidal (thinner along the z-dimension), and particle-shaped masks, respectively. The fifth column displays the 3D projections of the enhanced and IsoNet missing-wedge-corrected particle. The sixth and seventh columns present the final 3D map and the flexibly fitted model, respectively. (B) Circular schematic representation of a plasmid particle. The outer rim is color-coded to match the corresponding 3D model. Arrowheads indicate the transcriptional direction of bound RNAPs, and circles denote apical sites. Inner circular sectors represent individual plectonemes, with colors indicating writhe density (blue to red scale, −0.3 to 0.1). (C) Resolution assessment of the final 3D map using Fourier shell correlation (FSC). Two criteria are shown: FSC between two half-maps reconstructed from even and odd frames (evaluated at 0.143) and FSC between the final 3D map and the fitted model (evaluated at 0.5). (D) Zoomed-in views of the final 3D density map from panel A, displayed at two contour levels. (E) Superimposition of the high-contour level map from panel D onto its fitted model.

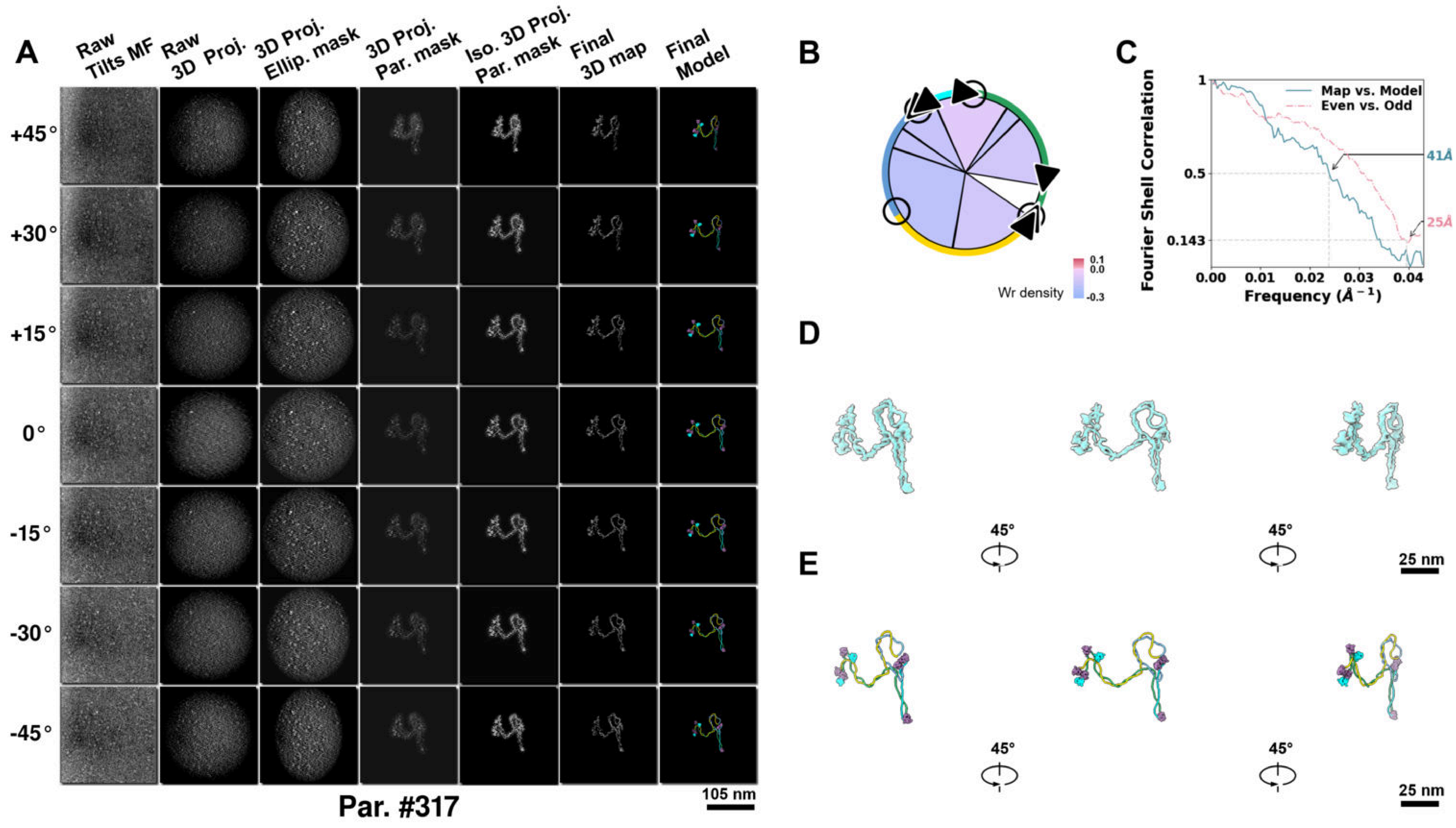

**Supplementary Particle Figure 317. Cryo-ET 3D reconstruction of an individual Tan.-TEC particle.**
(A) 3D reconstruction of the plasmid particle (index no. 317). The first column shows seven representative tilt images from +45° to -45° in step of 15°. The second, third, and fourth columns show 3D projections of the particle with spherical, ellipsoidal (thinner along the z-dimension), and particle-shaped masks, respectively. The fifth column displays the 3D projections of the enhanced and IsoNet missing-wedge-corrected particle. The sixth and seventh columns present the final 3D map and the flexibly fitted model, respectively. (B) Circular schematic representation of a plasmid particle. The outer rim is color-coded to match the corresponding 3D model. Arrowheads indicate the transcriptional direction of bound RNAPs, and circles denote apical sites. Inner circular sectors represent individual plectonemes, with colors indicating writhe density (blue to red scale, −0.3 to 0.1). (C) Resolution assessment of the final 3D map using Fourier shell correlation (FSC). Two criteria are shown: FSC between two half-maps reconstructed from even and odd frames (evaluated at 0.143) and FSC between the final 3D map and the fitted model (evaluated at 0.5). (D) Zoomed-in views of the final 3D density map from panel A, displayed at two contour levels. (E) Superimposition of the high-contour level map from panel D onto its fitted model.

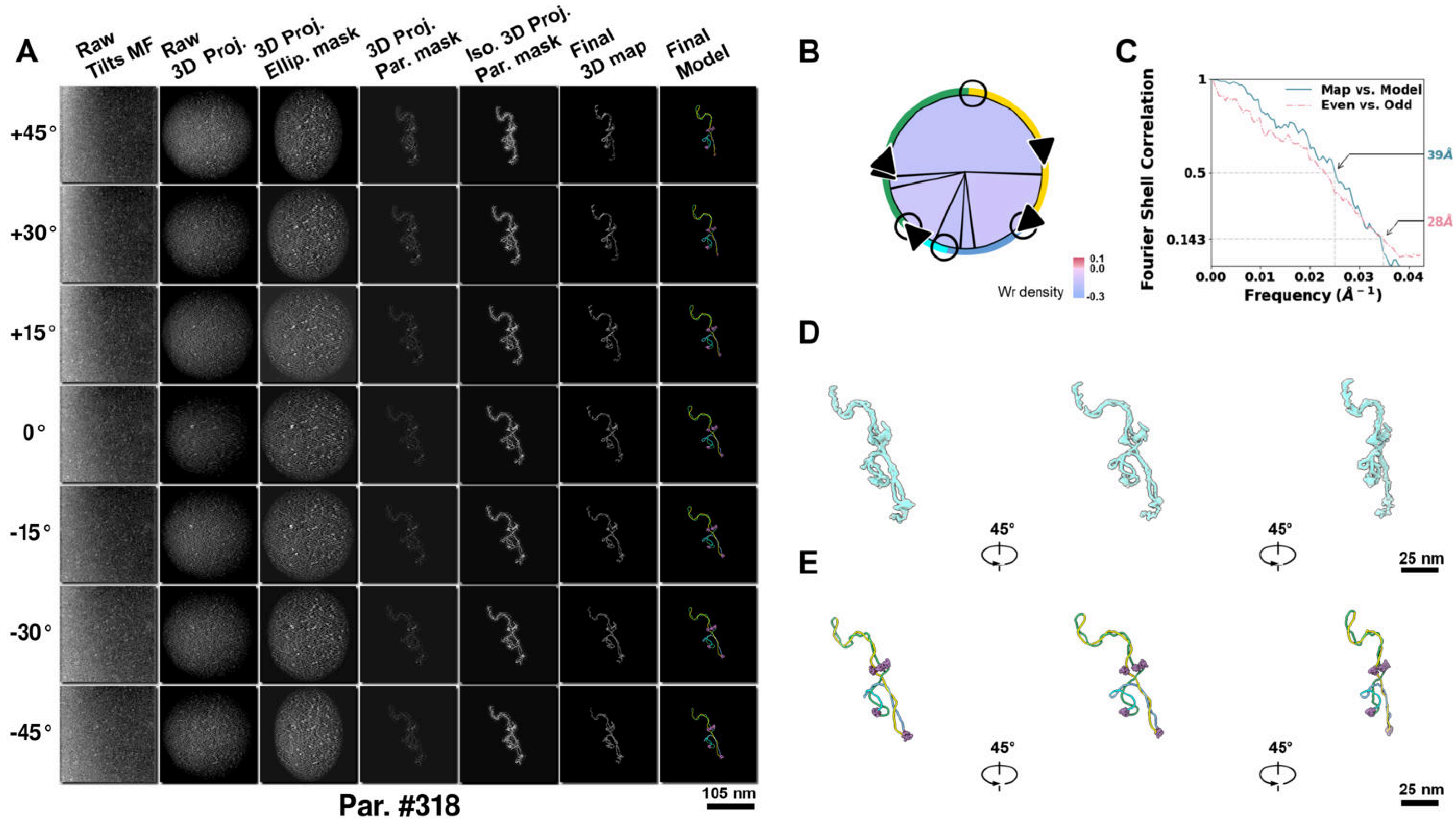

**Supplementary Particle Figure 318. Cryo-ET 3D reconstruction of an individual Tan.-TEC particle.**
(A) 3D reconstruction of the plasmid particle (index no. 318). The first column shows seven representative tilt images from +45° to -45° in step of 15°. The second, third, and fourth columns show 3D projections of the particle with spherical, ellipsoidal (thinner along the z-dimension), and particle-shaped masks, respectively. The fifth column displays the 3D projections of the enhanced and IsoNet missing-wedge-corrected particle. The sixth and seventh columns present the final 3D map and the flexibly fitted model, respectively. (B) Circular schematic representation of a plasmid particle. The outer rim is color-coded to match the corresponding 3D model. Arrowheads indicate the transcriptional direction of bound RNAPs, and circles denote apical sites. Inner circular sectors represent individual plectonemes, with colors indicating writhe density (blue to red scale, −0.3 to 0.1). (C) Resolution assessment of the final 3D map using Fourier shell correlation (FSC). Two criteria are shown: FSC between two half-maps reconstructed from even and odd frames (evaluated at 0.143) and FSC between the final 3D map and the fitted model (evaluated at 0.5). (D) Zoomed-in views of the final 3D density map from panel A, displayed at two contour levels. (E) Superimposition of the high-contour level map from panel D onto its fitted model.

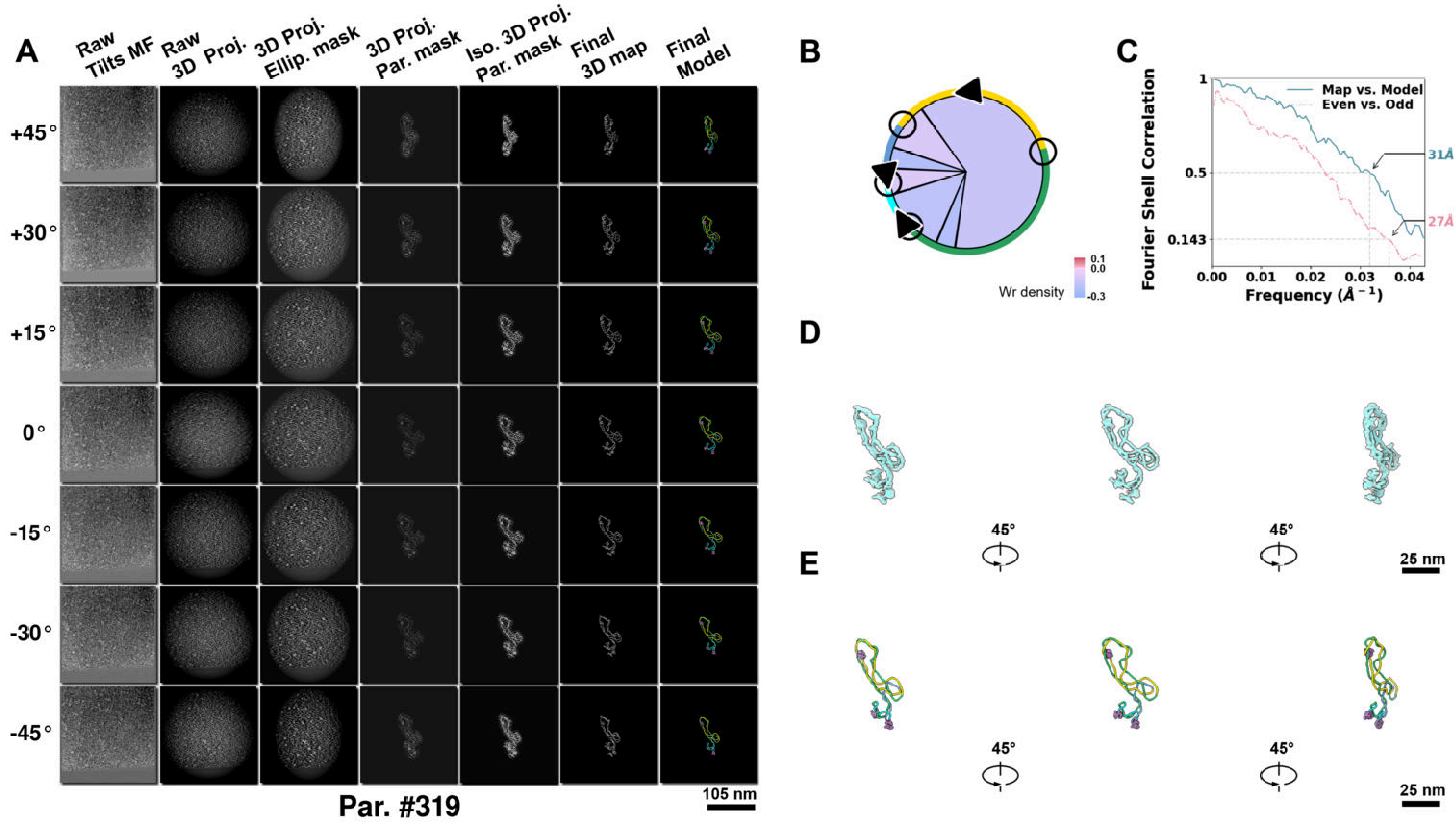

**Supplementary Particle Figure 319. Cryo-ET 3D reconstruction of an individual Tan.-TEC particle.**
(A) 3D reconstruction of the plasmid particle (index no. 319). The first column shows seven representative tilt images from +45° to -45° in step of 15°. The second, third, and fourth columns show 3D projections of the particle with spherical, ellipsoidal (thinner along the z-dimension), and particle-shaped masks, respectively. The fifth column displays the 3D projections of the enhanced and IsoNet missing-wedge-corrected particle. The sixth and seventh columns present the final 3D map and the flexibly fitted model, respectively. (B) Circular schematic representation of a plasmid particle. The outer rim is color-coded to match the corresponding 3D model. Arrowheads indicate the transcriptional direction of bound RNAPs, and circles denote apical sites. Inner circular sectors represent individual plectonemes, with colors indicating writhe density (blue to red scale, −0.3 to 0.1). (C) Resolution assessment of the final 3D map using Fourier shell correlation (FSC). Two criteria are shown: FSC between two half-maps reconstructed from even and odd frames (evaluated at 0.143) and FSC between the final 3D map and the fitted model (evaluated at 0.5). (D) Zoomed-in views of the final 3D density map from panel A, displayed at two contour levels. (E) Superimposition of the high-contour level map from panel D onto its fitted model.

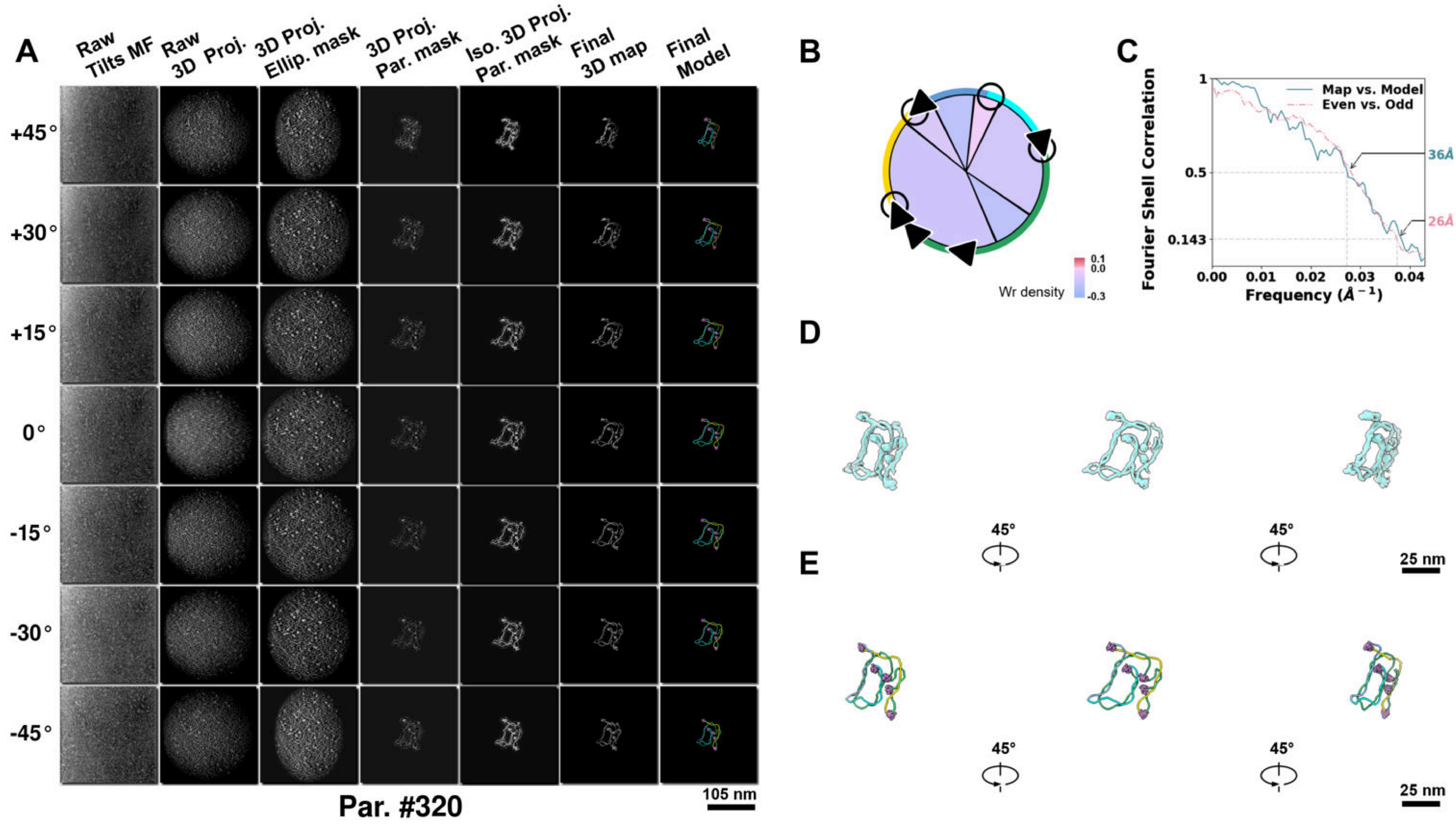

**Supplementary Particle Figure 320. Cryo-ET 3D reconstruction of an individual Tan.-TEC particle.**
(A) 3D reconstruction of the plasmid particle (index no. 320). The first column shows seven representative tilt images from +45° to -45° in step of 15°. The second, third, and fourth columns show 3D projections of the particle with spherical, ellipsoidal (thinner along the z-dimension), and particle-shaped masks, respectively. The fifth column displays the 3D projections of the enhanced and IsoNet missing-wedge-corrected particle. The sixth and seventh columns present the final 3D map and the flexibly fitted model, respectively. (B) Circular schematic representation of a plasmid particle. The outer rim is color-coded to match the corresponding 3D model. Arrowheads indicate the transcriptional direction of bound RNAPs, and circles denote apical sites. Inner circular sectors represent individual plectonemes, with colors indicating writhe density (blue to red scale, −0.3 to 0.1). (C) Resolution assessment of the final 3D map using Fourier shell correlation (FSC). Two criteria are shown: FSC between two half-maps reconstructed from even and odd frames (evaluated at 0.143) and FSC between the final 3D map and the fitted model (evaluated at 0.5). (D) Zoomed-in views of the final 3D density map from panel A, displayed at two contour levels. (E) Superimposition of the high-contour level map from panel D onto its fitted model.

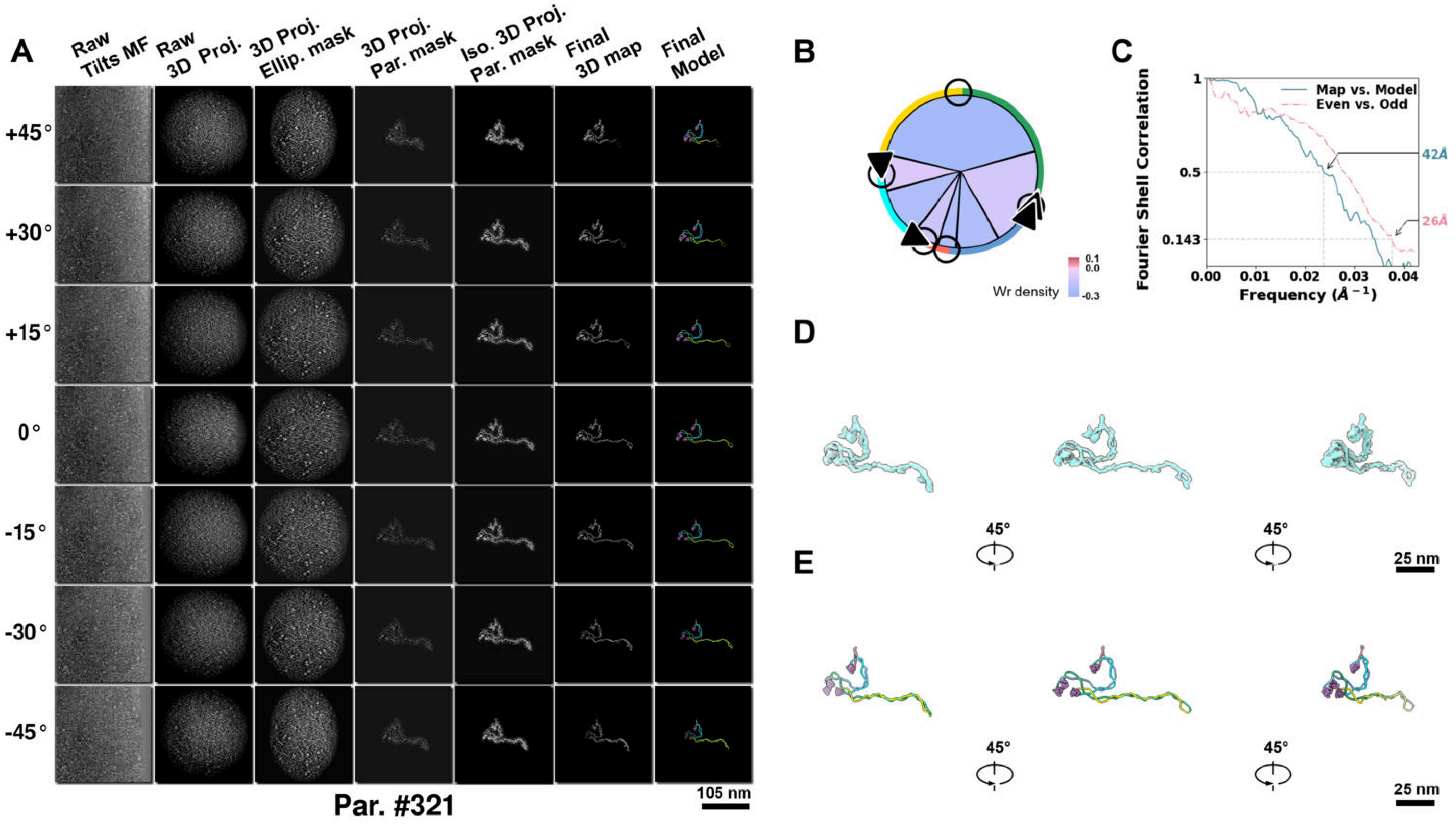

**Supplementary Particle Figure 321. Cryo-ET 3D reconstruction of an individual Tan.-TEC particle.**
(A) 3D reconstruction of the plasmid particle (index no. 321). The first column shows seven representative tilt images from +45° to -45° in step of 15°. The second, third, and fourth columns show 3D projections of the particle with spherical, ellipsoidal (thinner along the z-dimension), and particle-shaped masks, respectively. The fifth column displays the 3D projections of the enhanced and IsoNet missing-wedge-corrected particle. The sixth and seventh columns present the final 3D map and the flexibly fitted model, respectively. (B) Circular schematic representation of a plasmid particle. The outer rim is color-coded to match the corresponding 3D model. Arrowheads indicate the transcriptional direction of bound RNAPs, and circles denote apical sites. Inner circular sectors represent individual plectonemes, with colors indicating writhe density (blue to red scale, −0.3 to 0.1). (C) Resolution assessment of the final 3D map using Fourier shell correlation (FSC). Two criteria are shown: FSC between two half-maps reconstructed from even and odd frames (evaluated at 0.143) and FSC between the final 3D map and the fitted model (evaluated at 0.5). (D) Zoomed-in views of the final 3D density map from panel A, displayed at two contour levels. (E) Superimposition of the high-contour level map from panel D onto its fitted model.

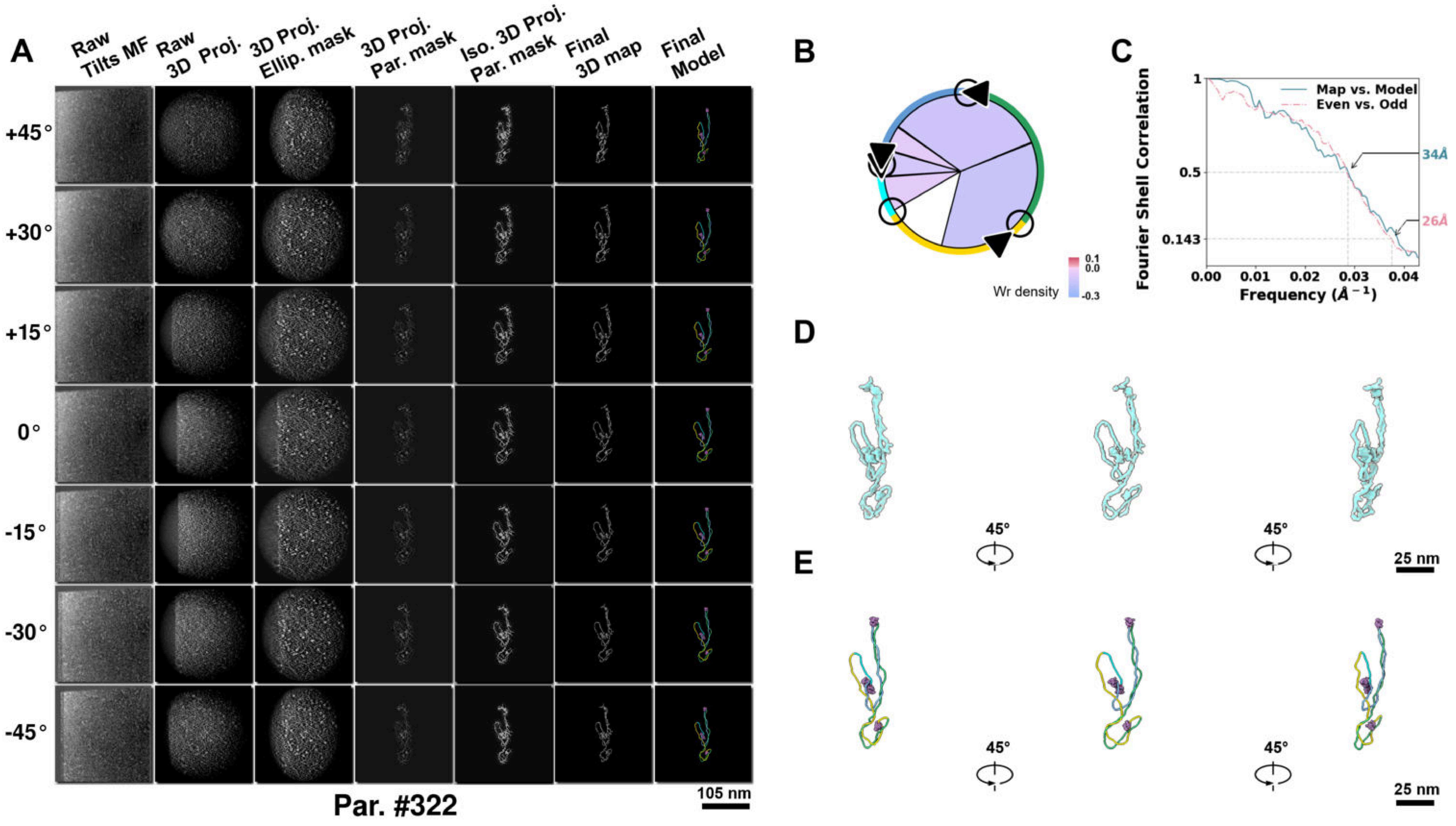

**Supplementary Particle Figure 322. Cryo-ET 3D reconstruction of an individual Tan.-TEC particle.**
(A) 3D reconstruction of the plasmid particle (index no. 322). The first column shows seven representative tilt images from +45° to -45° in step of 15°. The second, third, and fourth columns show 3D projections of the particle with spherical, ellipsoidal (thinner along the z-dimension), and particle-shaped masks, respectively. The fifth column displays the 3D projections of the enhanced and IsoNet missing-wedge-corrected particle. The sixth and seventh columns present the final 3D map and the flexibly fitted model, respectively. (B) Circular schematic representation of a plasmid particle. The outer rim is color-coded to match the corresponding 3D model. Arrowheads indicate the transcriptional direction of bound RNAPs, and circles denote apical sites. Inner circular sectors represent individual plectonemes, with colors indicating writhe density (blue to red scale, −0.3 to 0.1). (C) Resolution assessment of the final 3D map using Fourier shell correlation (FSC). Two criteria are shown: FSC between two half-maps reconstructed from even and odd frames (evaluated at 0.143) and FSC between the final 3D map and the fitted model (evaluated at 0.5). (D) Zoomed-in views of the final 3D density map from panel A, displayed at two contour levels. (E) Superimposition of the high-contour level map from panel D onto its fitted model.

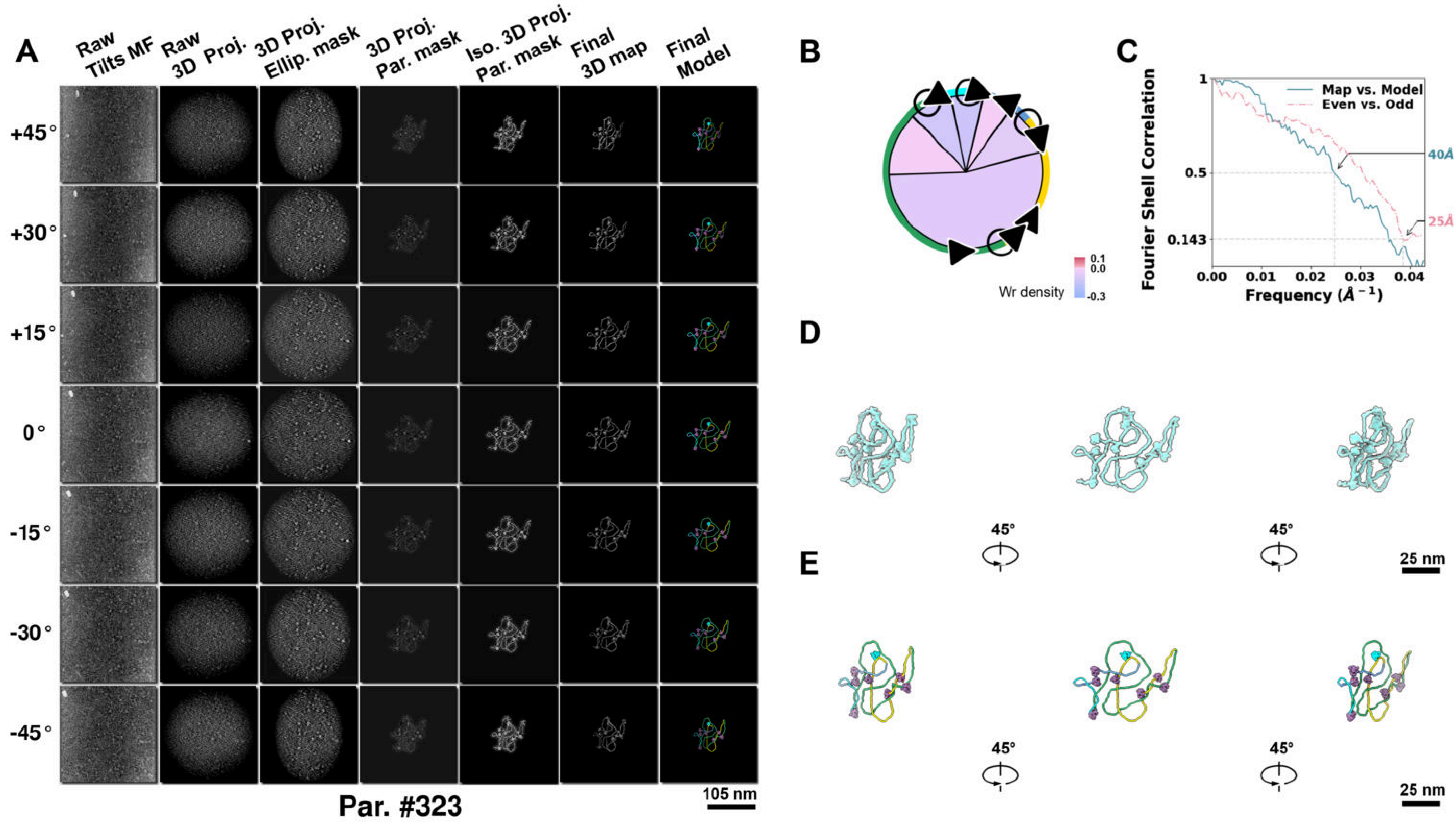

**Supplementary Particle Figure 323. Cryo-ET 3D reconstruction of an individual Tan.-TEC particle.**
(A) 3D reconstruction of the plasmid particle (index no. 323). The first column shows seven representative tilt images from +45° to -45° in step of 15°. The second, third, and fourth columns show 3D projections of the particle with spherical, ellipsoidal (thinner along the z-dimension), and particle-shaped masks, respectively. The fifth column displays the 3D projections of the enhanced and IsoNet missing-wedge-corrected particle. The sixth and seventh columns present the final 3D map and the flexibly fitted model, respectively. (B) Circular schematic representation of a plasmid particle. The outer rim is color-coded to match the corresponding 3D model. Arrowheads indicate the transcriptional direction of bound RNAPs, and circles denote apical sites. Inner circular sectors represent individual plectonemes, with colors indicating writhe density (blue to red scale, −0.3 to 0.1). (C) Resolution assessment of the final 3D map using Fourier shell correlation (FSC). Two criteria are shown: FSC between two half-maps reconstructed from even and odd frames (evaluated at 0.143) and FSC between the final 3D map and the fitted model (evaluated at 0.5). (D) Zoomed-in views of the final 3D density map from panel A, displayed at two contour levels. (E) Superimposition of the high-contour level map from panel D onto its fitted model.

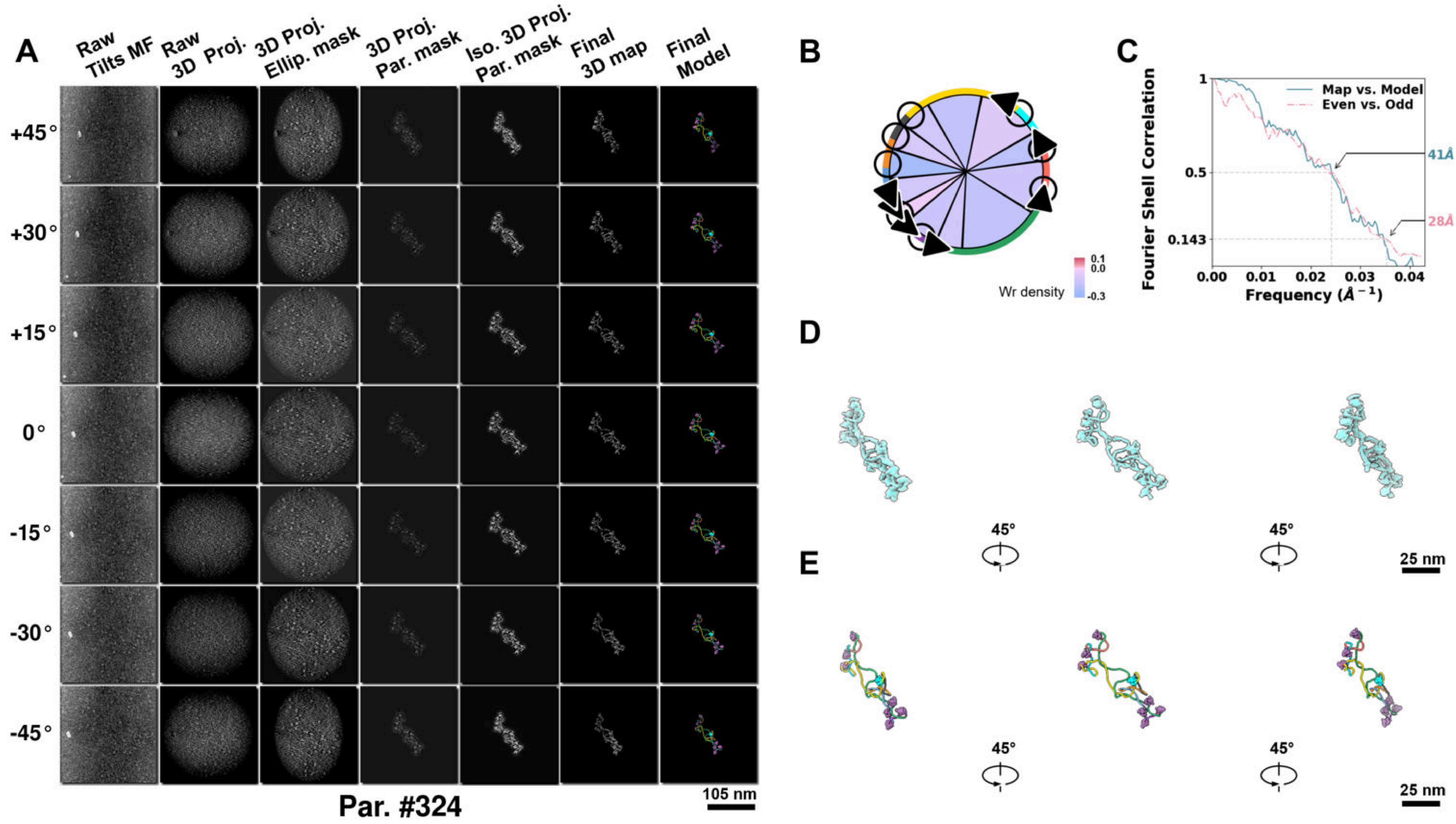

**Supplementary Particle Figure 324. Cryo-ET 3D reconstruction of an individual Tan.-TEC particle.**
(A) 3D reconstruction of the plasmid particle (index no. 324). The first column shows seven representative tilt images from +45° to -45° in step of 15°. The second, third, and fourth columns show 3D projections of the particle with spherical, ellipsoidal (thinner along the z-dimension), and particle-shaped masks, respectively. The fifth column displays the 3D projections of the enhanced and IsoNet missing-wedge-corrected particle. The sixth and seventh columns present the final 3D map and the flexibly fitted model, respectively. (B) Circular schematic representation of a plasmid particle. The outer rim is color-coded to match the corresponding 3D model. Arrowheads indicate the transcriptional direction of bound RNAPs, and circles denote apical sites. Inner circular sectors represent individual plectonemes, with colors indicating writhe density (blue to red scale, −0.3 to 0.1). (C) Resolution assessment of the final 3D map using Fourier shell correlation (FSC). Two criteria are shown: FSC between two half-maps reconstructed from even and odd frames (evaluated at 0.143) and FSC between the final 3D map and the fitted model (evaluated at 0.5). (D) Zoomed-in views of the final 3D density map from panel A, displayed at two contour levels. (E) Superimposition of the high-contour level map from panel D onto its fitted model.

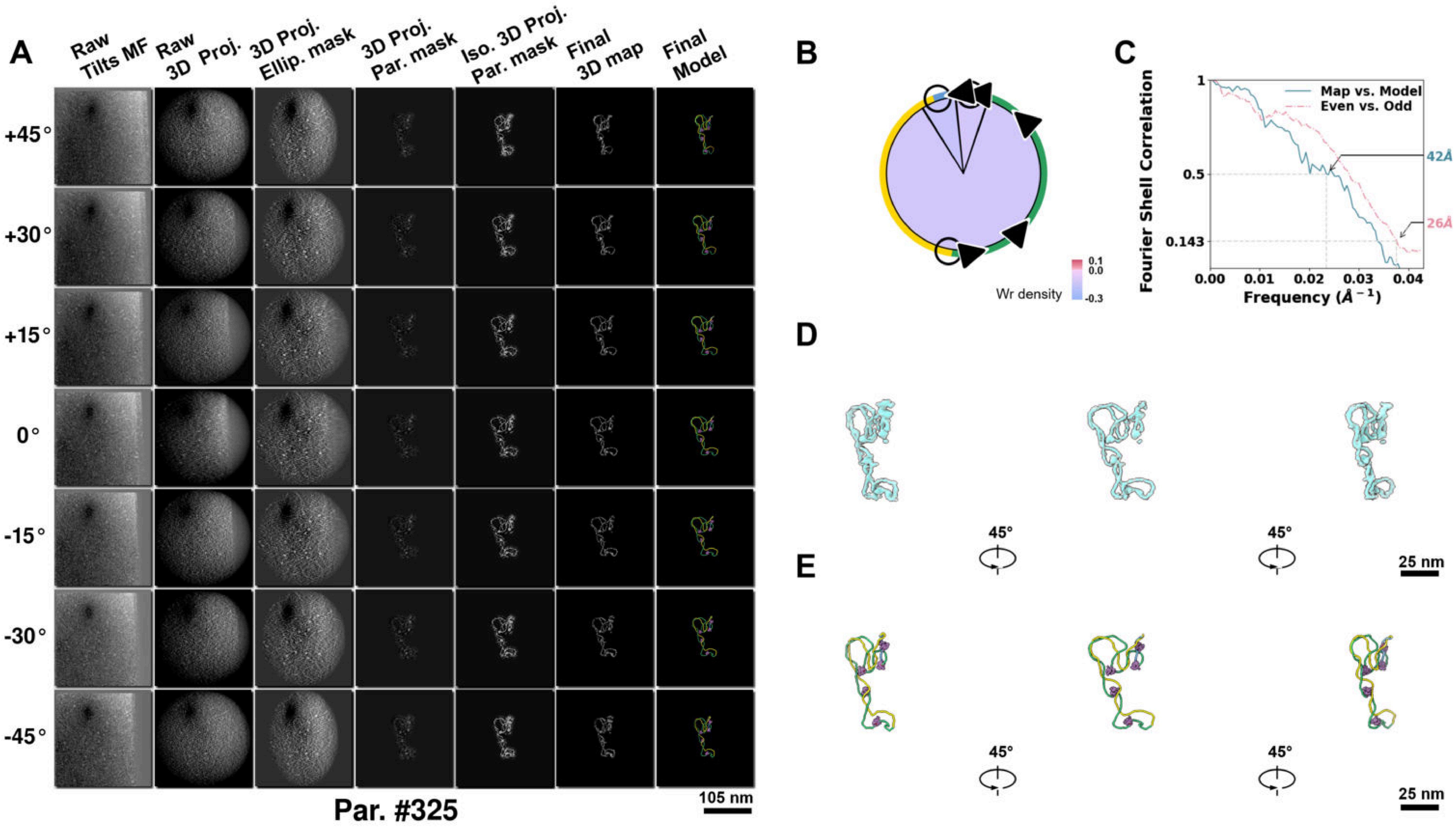

**Supplementary Particle Figure 325. Cryo-ET 3D reconstruction of an individual Tan.-TEC particle.**
(A) 3D reconstruction of the plasmid particle (index no. 325). The first column shows seven representative tilt images from +45° to -45° in step of 15°. The second, third, and fourth columns show 3D projections of the particle with spherical, ellipsoidal (thinner along the z-dimension), and particle-shaped masks, respectively. The fifth column displays the 3D projections of the enhanced and IsoNet missing-wedge-corrected particle. The sixth and seventh columns present the final 3D map and the flexibly fitted model, respectively. (B) Circular schematic representation of a plasmid particle. The outer rim is color-coded to match the corresponding 3D model. Arrowheads indicate the transcriptional direction of bound RNAPs, and circles denote apical sites. Inner circular sectors represent individual plectonemes, with colors indicating writhe density (blue to red scale, −0.3 to 0.1). (C) Resolution assessment of the final 3D map using Fourier shell correlation (FSC). Two criteria are shown: FSC between two half-maps reconstructed from even and odd frames (evaluated at 0.143) and FSC between the final 3D map and the fitted model (evaluated at 0.5). (D) Zoomed-in views of the final 3D density map from panel A, displayed at two contour levels. (E) Superimposition of the high-contour level map from panel D onto its fitted model.

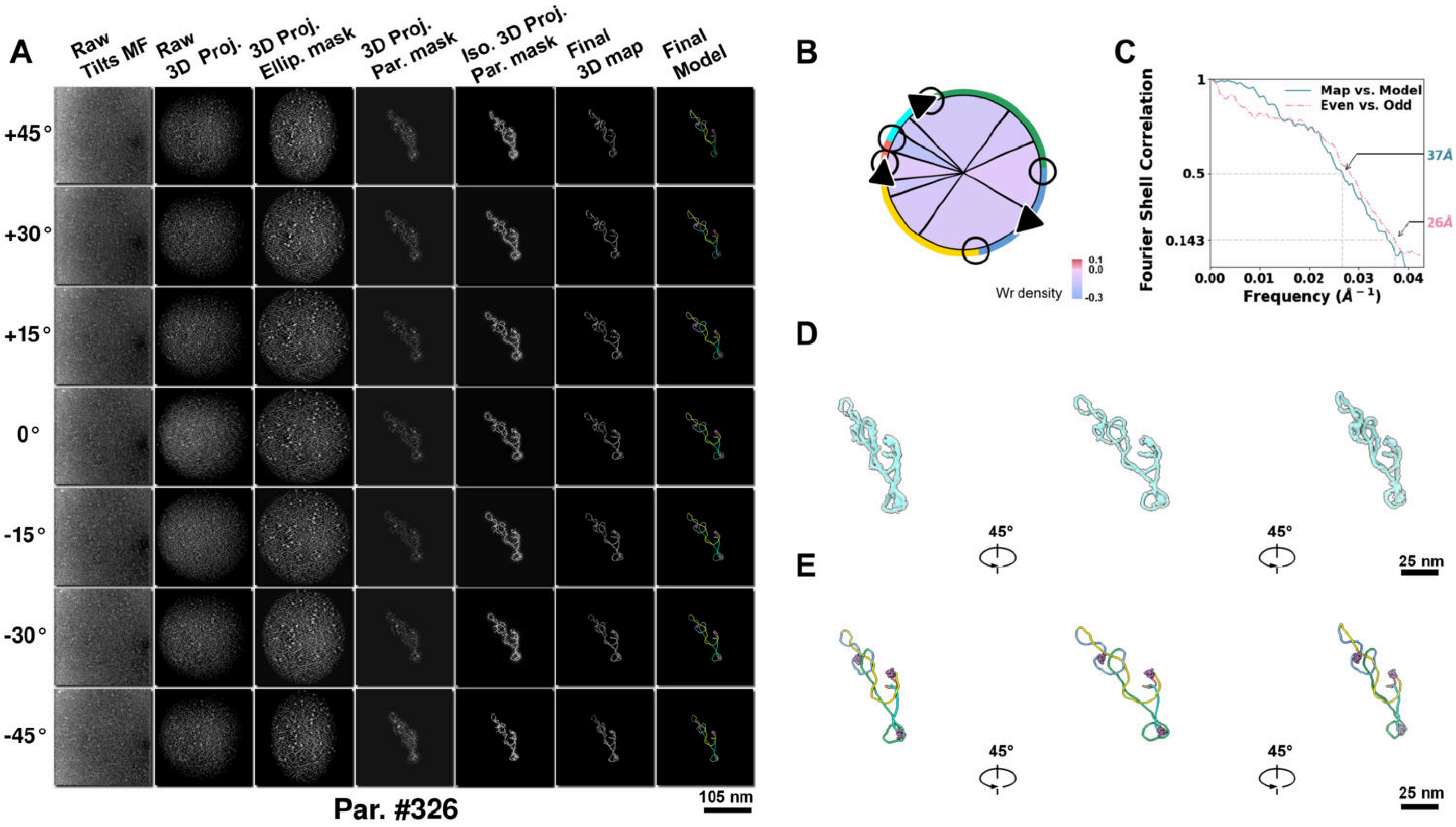

**Supplementary Particle Figure 326. Cryo-ET 3D reconstruction of an individual Tan.-TEC particle.**
(A) 3D reconstruction of the plasmid particle (index no. 326). The first column shows seven representative tilt images from +45° to -45° in step of 15°. The second, third, and fourth columns show 3D projections of the particle with spherical, ellipsoidal (thinner along the z-dimension), and particle-shaped masks, respectively. The fifth column displays the 3D projections of the enhanced and IsoNet missing-wedge-corrected particle. The sixth and seventh columns present the final 3D map and the flexibly fitted model, respectively. (B) Circular schematic representation of a plasmid particle. The outer rim is color-coded to match the corresponding 3D model. Arrowheads indicate the transcriptional direction of bound RNAPs, and circles denote apical sites. Inner circular sectors represent individual plectonemes, with colors indicating writhe density (blue to red scale, −0.3 to 0.1). (C) Resolution assessment of the final 3D map using Fourier shell correlation (FSC). Two criteria are shown: FSC between two half-maps reconstructed from even and odd frames (evaluated at 0.143) and FSC between the final 3D map and the fitted model (evaluated at 0.5). (D) Zoomed-in views of the final 3D density map from panel A, displayed at two contour levels. (E) Superimposition of the high-contour level map from panel D onto its fitted model.

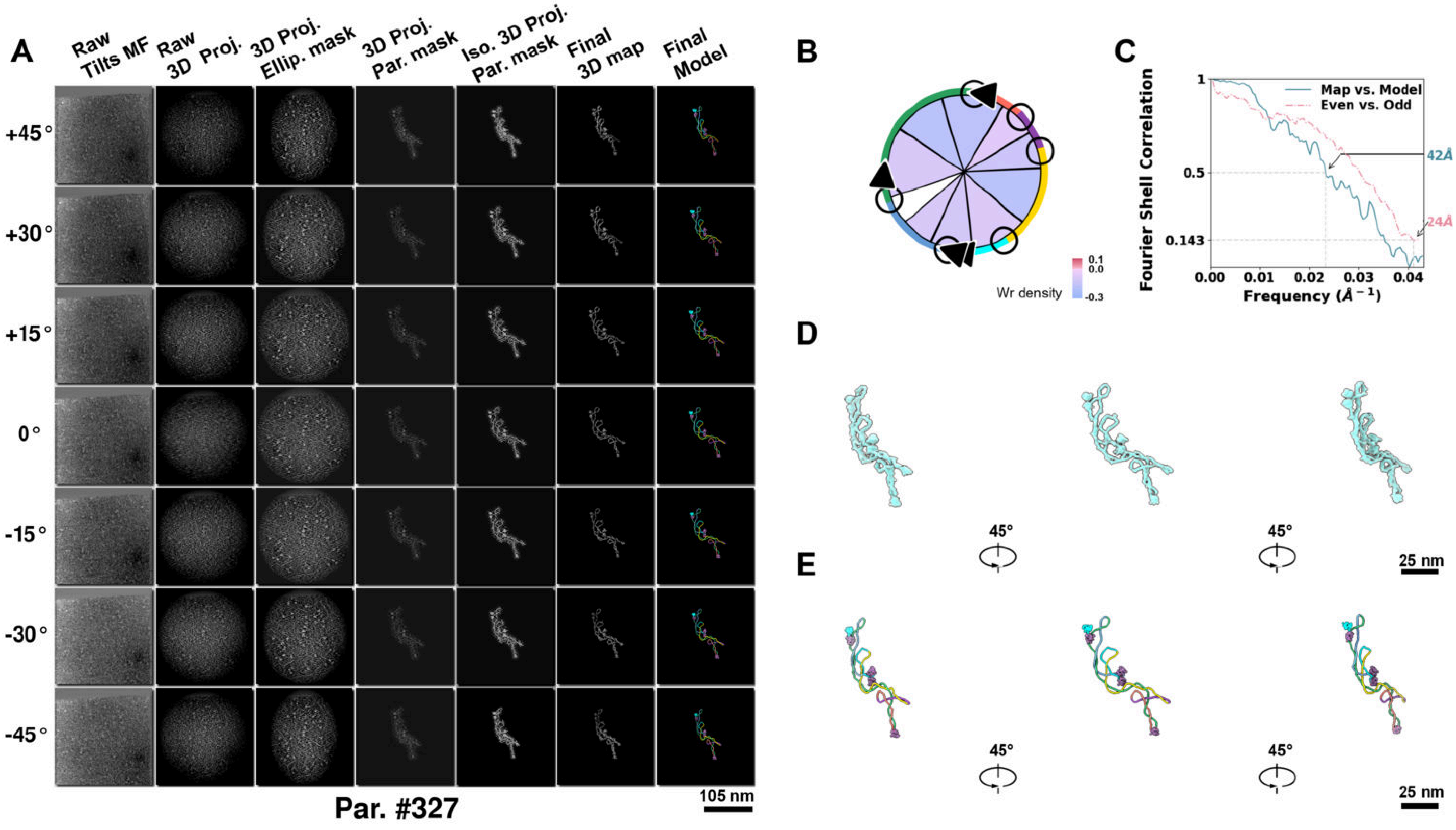

**Supplementary Particle Figure 327. Cryo-ET 3D reconstruction of an individual Tan.-TEC particle.**
(A) 3D reconstruction of the plasmid particle (index no. 327). The first column shows seven representative tilt images from +45° to -45° in step of 15°. The second, third, and fourth columns show 3D projections of the particle with spherical, ellipsoidal (thinner along the z-dimension), and particle-shaped masks, respectively. The fifth column displays the 3D projections of the enhanced and IsoNet missing-wedge-corrected particle. The sixth and seventh columns present the final 3D map and the flexibly fitted model, respectively. (B) Circular schematic representation of a plasmid particle. The outer rim is color-coded to match the corresponding 3D model. Arrowheads indicate the transcriptional direction of bound RNAPs, and circles denote apical sites. Inner circular sectors represent individual plectonemes, with colors indicating writhe density (blue to red scale, −0.3 to 0.1). (C) Resolution assessment of the final 3D map using Fourier shell correlation (FSC). Two criteria are shown: FSC between two half-maps reconstructed from even and odd frames (evaluated at 0.143) and FSC between the final 3D map and the fitted model (evaluated at 0.5). (D) Zoomed-in views of the final 3D density map from panel A, displayed at two contour levels. (E) Superimposition of the high-contour level map from panel D onto its fitted model.

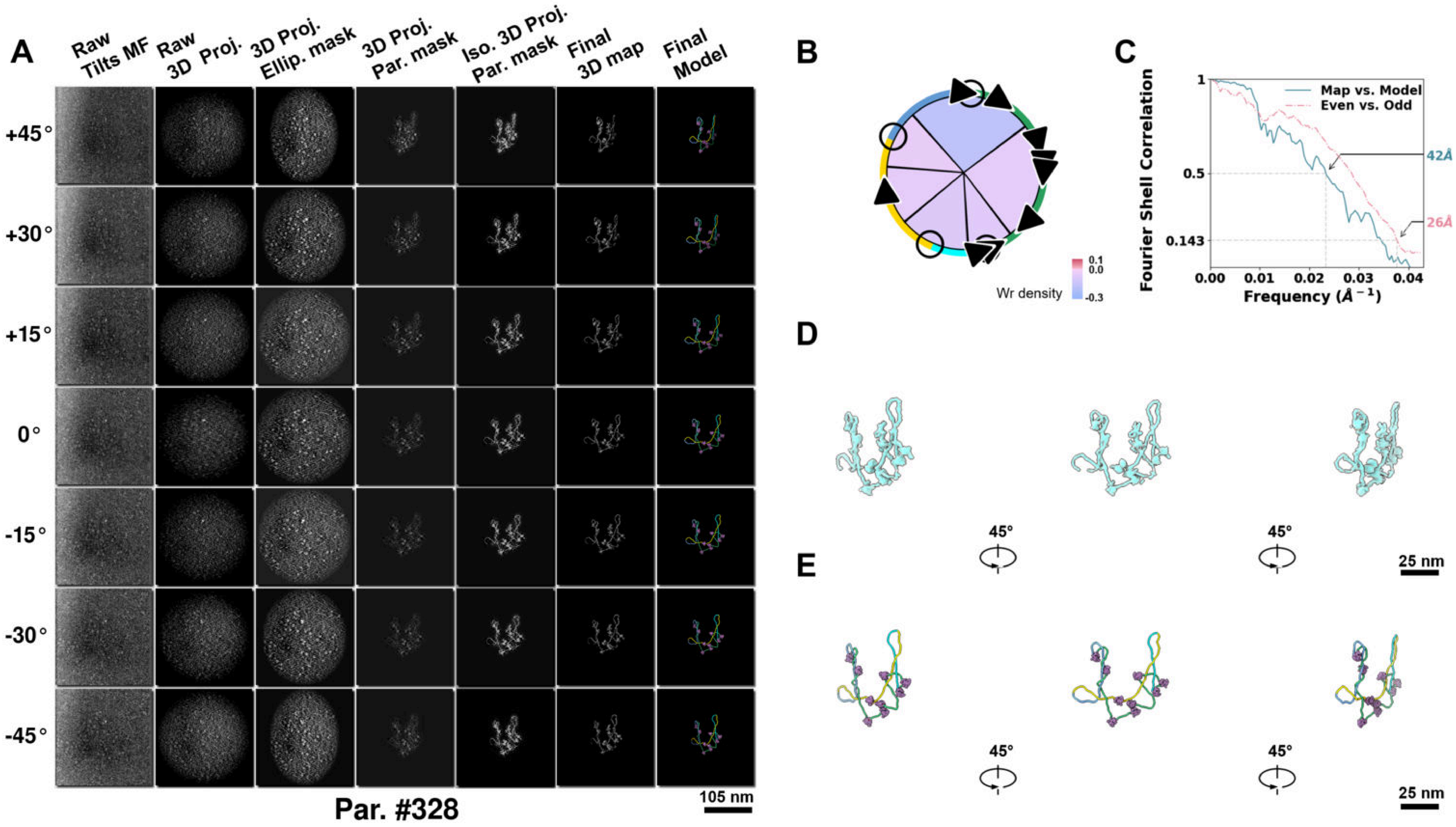

**Supplementary Particle Figure 328. Cryo-ET 3D reconstruction of an individual Tan.-TEC particle.**
(A) 3D reconstruction of the plasmid particle (index no. 328). The first column shows seven representative tilt images from +45° to -45° in step of 15°. The second, third, and fourth columns show 3D projections of the particle with spherical, ellipsoidal (thinner along the z-dimension), and particle-shaped masks, respectively. The fifth column displays the 3D projections of the enhanced and IsoNet missing-wedge-corrected particle. The sixth and seventh columns present the final 3D map and the flexibly fitted model, respectively. (B) Circular schematic representation of a plasmid particle. The outer rim is color-coded to match the corresponding 3D model. Arrowheads indicate the transcriptional direction of bound RNAPs, and circles denote apical sites. Inner circular sectors represent individual plectonemes, with colors indicating writhe density (blue to red scale, −0.3 to 0.1). (C) Resolution assessment of the final 3D map using Fourier shell correlation (FSC). Two criteria are shown: FSC between two half-maps reconstructed from even and odd frames (evaluated at 0.143) and FSC between the final 3D map and the fitted model (evaluated at 0.5). (D) Zoomed-in views of the final 3D density map from panel A, displayed at two contour levels. (E) Superimposition of the high-contour level map from panel D onto its fitted model.

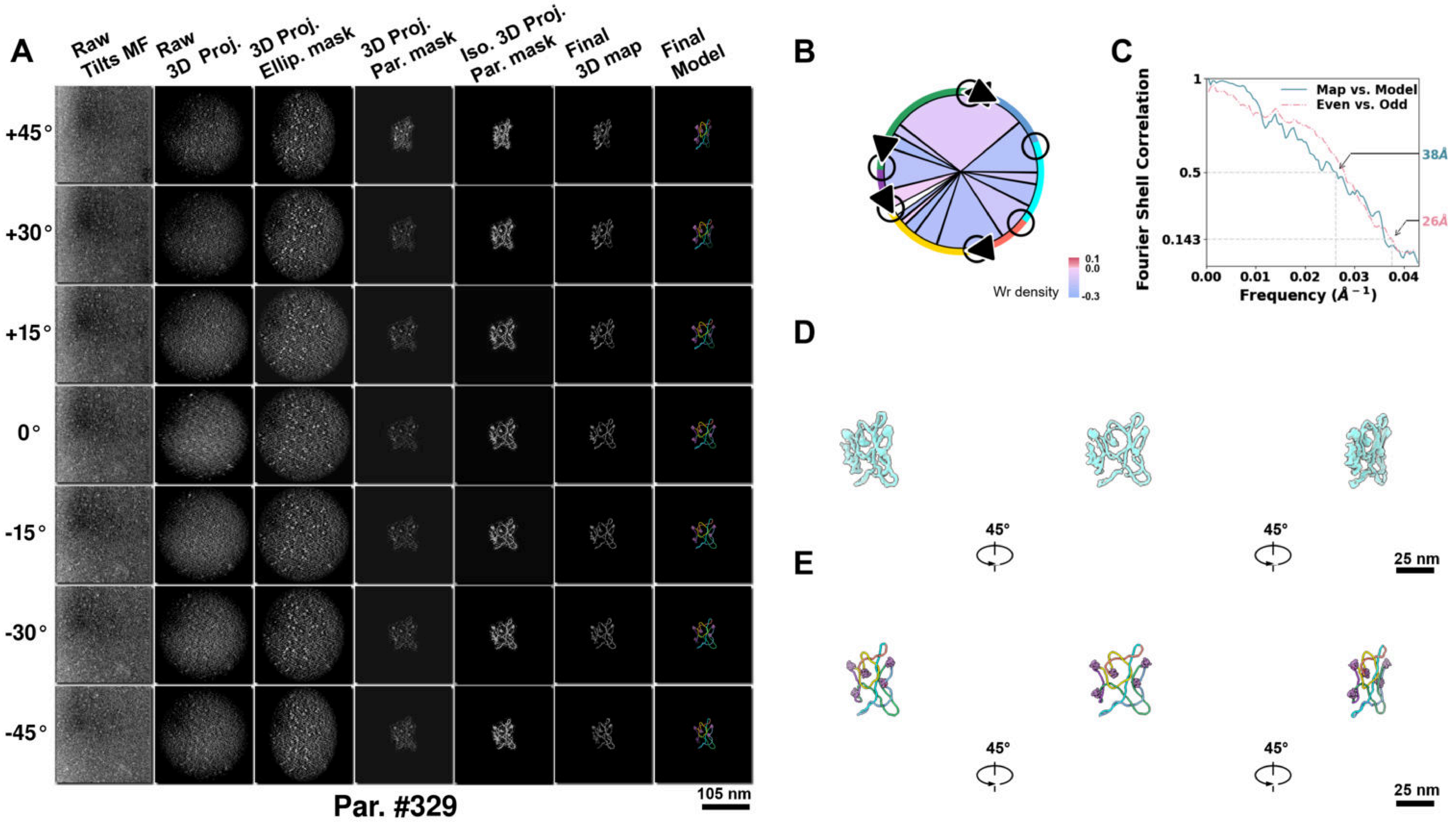

**Supplementary Particle Figure 329. Cryo-ET 3D reconstruction of an individual Tan.-TEC particle.**
(A) 3D reconstruction of the plasmid particle (index no. 329). The first column shows seven representative tilt images from +45° to -45° in step of 15°. The second, third, and fourth columns show 3D projections of the particle with spherical, ellipsoidal (thinner along the z-dimension), and particle-shaped masks, respectively. The fifth column displays the 3D projections of the enhanced and IsoNet missing-wedge-corrected particle. The sixth and seventh columns present the final 3D map and the flexibly fitted model, respectively. (B) Circular schematic representation of a plasmid particle. The outer rim is color-coded to match the corresponding 3D model. Arrowheads indicate the transcriptional direction of bound RNAPs, and circles denote apical sites. Inner circular sectors represent individual plectonemes, with colors indicating writhe density (blue to red scale, −0.3 to 0.1). (C) Resolution assessment of the final 3D map using Fourier shell correlation (FSC). Two criteria are shown: FSC between two half-maps reconstructed from even and odd frames (evaluated at 0.143) and FSC between the final 3D map and the fitted model (evaluated at 0.5). (D) Zoomed-in views of the final 3D density map from panel A, displayed at two contour levels. (E) Superimposition of the high-contour level map from panel D onto its fitted model.

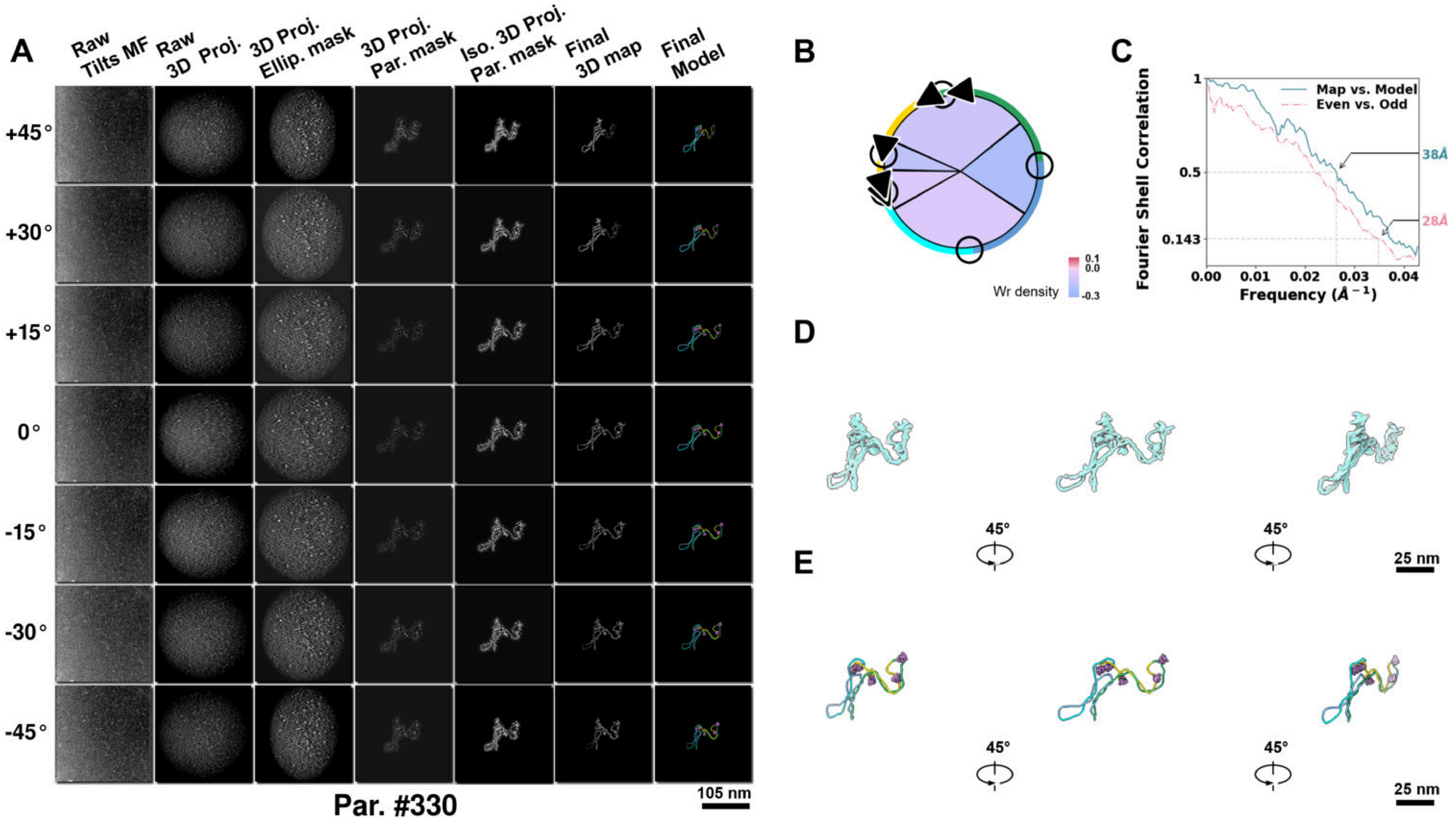

**Supplementary Particle Figure 330. Cryo-ET 3D reconstruction of an individual Tan.-TEC particle.**
(A) 3D reconstruction of the plasmid particle (index no. 330). The first column shows seven representative tilt images from +45° to -45° in step of 15°. The second, third, and fourth columns show 3D projections of the particle with spherical, ellipsoidal (thinner along the z-dimension), and particle-shaped masks, respectively. The fifth column displays the 3D projections of the enhanced and IsoNet missing-wedge-corrected particle. The sixth and seventh columns present the final 3D map and the flexibly fitted model, respectively. (B) Circular schematic representation of a plasmid particle. The outer rim is color-coded to match the corresponding 3D model. Arrowheads indicate the transcriptional direction of bound RNAPs, and circles denote apical sites. Inner circular sectors represent individual plectonemes, with colors indicating writhe density (blue to red scale, −0.3 to 0.1). (C) Resolution assessment of the final 3D map using Fourier shell correlation (FSC). Two criteria are shown: FSC between two half-maps reconstructed from even and odd frames (evaluated at 0.143) and FSC between the final 3D map and the fitted model (evaluated at 0.5). (D) Zoomed-in views of the final 3D density map from panel A, displayed at two contour levels. (E) Superimposition of the high-contour level map from panel D onto its fitted model.

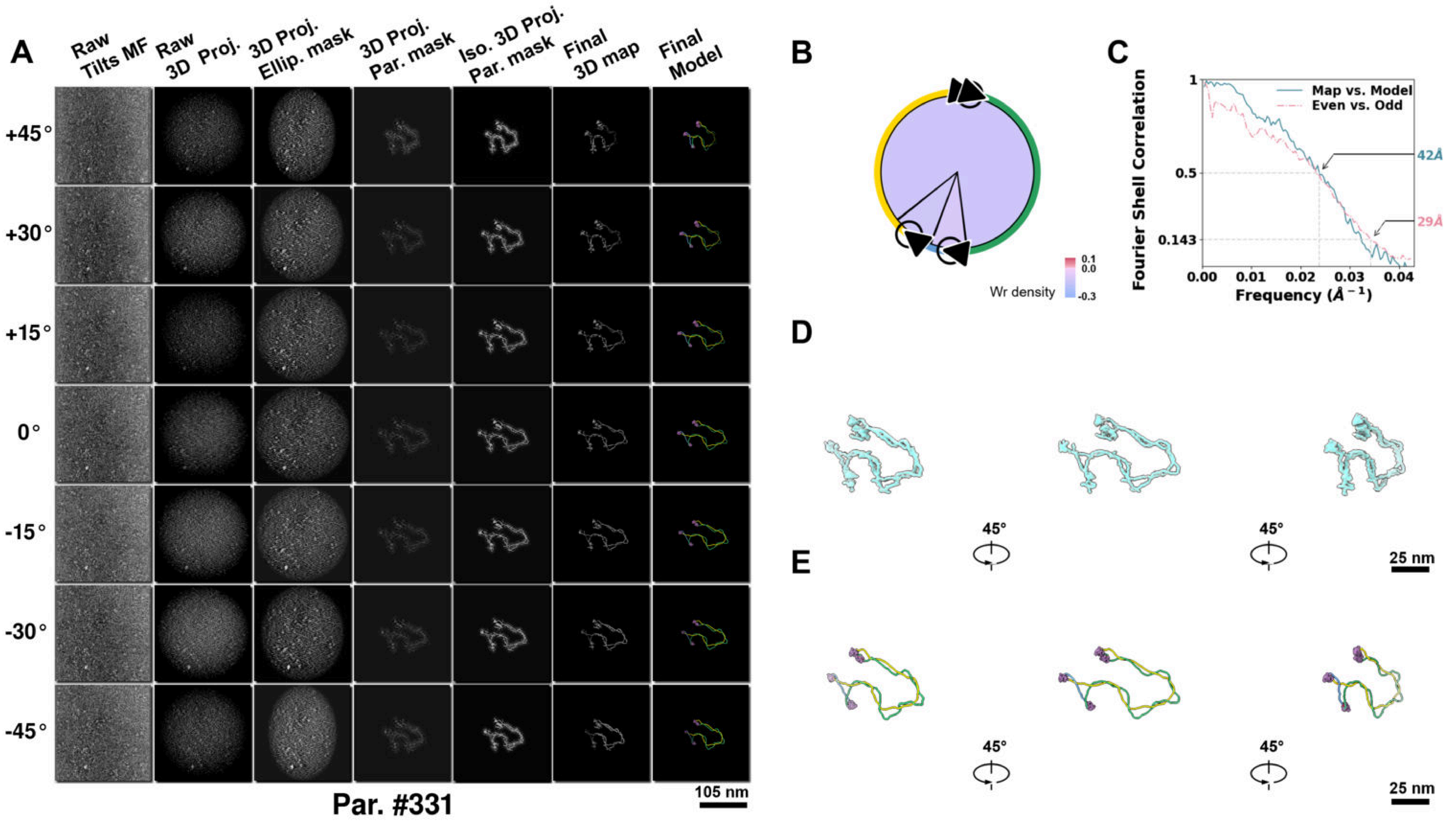

**Supplementary Particle Figure 331. Cryo-ET 3D reconstruction of an individual Tan.-TEC particle.**
(A) 3D reconstruction of the plasmid particle (index no. 331). The first column shows seven representative tilt images from +45° to -45° in step of 15°. The second, third, and fourth columns show 3D projections of the particle with spherical, ellipsoidal (thinner along the z-dimension), and particle-shaped masks, respectively. The fifth column displays the 3D projections of the enhanced and IsoNet missing-wedge-corrected particle. The sixth and seventh columns present the final 3D map and the flexibly fitted model, respectively. (B) Circular schematic representation of a plasmid particle. The outer rim is color-coded to match the corresponding 3D model. Arrowheads indicate the transcriptional direction of bound RNAPs, and circles denote apical sites. Inner circular sectors represent individual plectonemes, with colors indicating writhe density (blue to red scale, −0.3 to 0.1). (C) Resolution assessment of the final 3D map using Fourier shell correlation (FSC). Two criteria are shown: FSC between two half-maps reconstructed from even and odd frames (evaluated at 0.143) and FSC between the final 3D map and the fitted model (evaluated at 0.5). (D) Zoomed-in views of the final 3D density map from panel A, displayed at two contour levels. (E) Superimposition of the high-contour level map from panel D onto its fitted model.
